# Supersized ribosomal RNA expansion segments in Asgard archaea

**DOI:** 10.1101/2019.12.25.888164

**Authors:** Petar I. Penev, Sara Fakhretaha-Aval, Vaishnavi J. Patel, Jamie J. Cannone, Robin R. Gutell, Anton S. Petrov, Loren Dean Williams, Jennifer B. Glass

## Abstract

The ribosome’s common core, comprised of ribosomal RNA (rRNA) and universal ribosomal proteins, connects all life back to a common ancestor and serves as a window to relationships among organisms. The rRNA of the common core is most similar to rRNA of extant bacteria. In eukaryotes, the rRNA of the common core is decorated by expansion segments (ESs) that vastly increase its size. Supersized ESs have not been observed previously in Archaea, and the origin of eukaryotic ESs remains enigmatic. We discovered that the large subunit (LSU) rRNA of two Asgard phyla, *Lokiarchaeota* and *Heimdallarchaeota,* considered to be the closest modern archaeal cell lineages to Eukarya, bridge the gap in size between prokaryotic and eukaryotic LSU rRNA. The elongated LSU rRNAs in *Lokiarchaeota* and *Heimdallarchaeota* stem from the presence of two supersized ESs, ES9 and ES39. We applied chemical footprinting experiments to study the structure of *Lokiarchaeota* ES39. Furthermore, we used covariation and sequence analysis to study the evolution of Asgard ES39s and ES9s. By defining the common eukaryotic ES39 signature fold, we found that Asgard ES39s have more and longer helices than eukaryotic ES39s. While Asgard ES39s have sequences and structures distinct from eukaryotic ES39s, we found overall conservation of a three-way junction across the Asgard species that matches eukaryotic ES39 topology, a result consistent with the accretion model of ribosomal evolution.

**Significance:** Eukaryotes possess longer and more complex ribosomal RNA (rRNA) than Bacteria and Archaea, including eukaryotic-specific rRNA expansion segments (ESs). The origin and evolution of ESs has long remained a mystery. We show that two recently discovered Asgard archaeal phyla, Lokiarchaeota and Heimdallarchaeota, contain large subunit rRNA with ESs are “supersized” (>100 nt) compared to all other prokaryotes studied to date. Asgard ESs have distinct structures from eukaryotic ESs, but share a common three-way junction out of which the ES grew. Our findings raise the possibility that supersized ESs existed on the ribosomal surface before the last eukaryotic common ancestor, opening the question of whether ribosomal complexity is more deeply rooted than previously thought.

## Introduction

The ribosome connects all life on Earth back to the Last Universal Common Ancestor (LUCA) (Woese and Fox 1977). The small ribosomal subunit (SSU) decodes mRNA and the large ribosomal subunit (LSU) links amino acids together to produce coded protein. Both subunits are made of ribosomal RNA (rRNA) and ribosomal protein (rProtein). All cytoplasmic ribosomes contain a structurally conserved universal common core, comprised of 2800 nucleotides and 28 rProteins, and including the peptidyl transferase center (PTC) in the LSU and the decoding center (DCC) in the SSU (Melnikov, et al. 2012; Bernier, et al. 2018). The rRNA of the common core is a reasonable structural approximation of the rRNA in LUCA and is most similar to rRNA of extant bacteria (Melnikov, et al. 2012; Petrov, et al. 2014b; Bernier, et al. 2018).

In Eukarya, the rRNA of the common core is elaborated by expansion segments (ESs, **Fig. 1**) (Veldman, et al. 1981; Clark, et al. 1984; Hassouna, et al. 1984; Gonzalez, et al. 1985; Michot and Bachellerie 1987; Bachellerie and Michot 1989; Gutell 1992; Lapeyre, et al. 1993; Gerbi 1996; Schnare, et al. 1996). ESs emerge from a small number of conserved sites on the common core and are excluded from regions of essential ribosomal function such as the DCC, the PTC, and the subunit interface (Ben-Shem, et al. 2010; Anger, et al. 2013). Some archaea exhibit short ESs (μ-ESs) protruding from the same regions as eukaryotic ESs (Armache, et al. 2012). *Halococcus morrhuae*, a halophilic archaeon, contains a large rRNA insertion in an LSU region that lacks ESs in eukaryotes (Tirumalai, et al. 2020). Expansion segments are larger and more numerous on the LSU than on the SSU; across phylogeny, size variation of the SSU rRNA is around 10% of that of LSU rRNA (Gutell 1992; Gerbi 1996; Bernier, et al. 2018).

**Figure 1:**
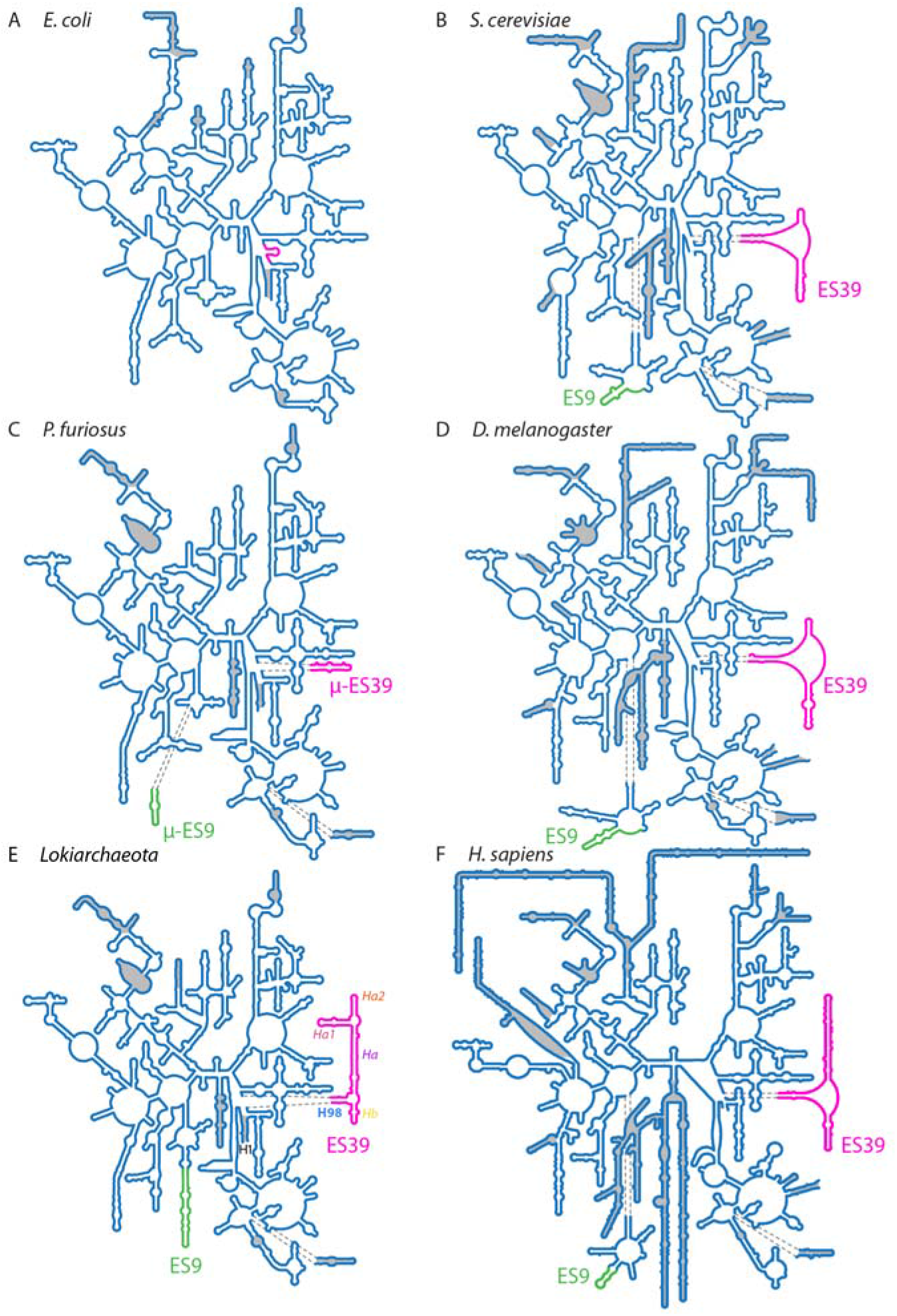
Secondary structures of the LSU rRNA from Bacteria, Archaea, and Eukarya. (A) *Escherichia coli* (Bacteria), (B) *Saccharomyces cerevisiae* (fungus, Eukarya); (C) *Pyrococcus furiosus* (Archaea); (D) *Drosophila melanogaster* (insect, Eukarya); (E) *Lokiarchaeota* B5 (Archaea); (F) *Homo sapiens* (primate, Eukarya). Secondary structures in panels A, B, C, D, and F are taken from *Petrov, et al. (2014a)*. Secondary structure in panel E is from this study. Universal rRNA common core is shown in blue lines (not shaded). ES9 is shown with a green line. ES39 is shown with a magenta line. Helices in panel E are abbreviated as Helix 1 (H1), helix 98 (H98), helix *a* (H*a*), helix *a1* (H*a1*), helix *a2* (H*a2*) and helix *b* (H*b*). ESs and helices not present in the common core are shaded in gray. Sizes of secondary structures are to scale. The numbering scheme of Noller, et al. (1981) and Leffers, et al. (1987) were used to label the helices and ESs.

ESs mutate quickly, at rates much greater than those of common core rRNA (Gonzalez, et al. 1985; Gonzalez, et al. 1988; Ajuh, et al. 1991; Gerbi 1996). For example, ES7 of *Drosophila melanogaster* (fruit fly) is AU-rich and lacks sequence homology with the GC-rich ES7 of *Aedes albopictus* (Asian tiger mosquito). The high rate of mutation is indicated by the substantial sequence differences in species of the order Diptera that diverged only 200-250 million years ago (Aris-Brosou and Yang 2002; Kumar, et al. 2017). Although sequence conservation of ESs is low, secondary and tertiary structures are excellent sources of information on ES homology and evolution. For this reason, early studies used rRNA secondary structures to establish phylogenetic relationships (Woese, et al. 1990a).

The recent discovery and characterization of the Asgard archaeal superphylum suggests that the last Asgard and eukaryotic common ancestor (LAsECA) contained key components of eukaryotic cellular systems (Spang, et al. 2015; Klinger, et al. 2016; Eme, et al. 2017; Zaremba-Niedzwiedzka, et al. 2017; Narrowe, et al. 2018; Spang, et al. 2019). Eukaryotic signature proteins (ESPs) found in Asgard archaea are involved in cytoskeleton, trafficking, ubiquitination, and translation (Spang, et al. 2015; Zaremba-Niedzwiedzka, et al. 2017; Akıl and Robinson 2018; Melnikov, et al. 2019; Stairs and Ettema 2020). Asgard archaea also contain several homologs of eukaryotic ribosomal proteins (Hartman and Fedorov 2002; Spang, et al. 2015; Zaremba-Niedzwiedzka, et al. 2017). Metazoan rRNAs contain supersized ESs of hundreds of nucleotides (nts); hereafter, we define “supersized” ESs as those with >100 nts. Prior to the current study, it was not known if Asgard rRNAs could contain features such as supersized ESs. Supersized ESs have not been observed previously in Bacteria or Archaea and were considered unique to eukaryotes (Ware, et al. 1983; Clark, et al. 1984; Hassouna, et al. 1984; Gerbi 1996; Melnikov, et al. 2012).

Here, we use computational and experimental approaches to characterize structure and evolution of Asgard rRNAs. We find that rRNAs of *Lokiarchaeota* and *Heimdallarchaeota*, both from the Asgard phyla, are elaborated on the LSU by supersized ESs. In size and complexity, *Lokiarchaeota* and *Heimdallarchaeota* LSU ESs exceed those of diverse protist rRNAs and rival those of metazoan rRNAs. No es’s were found in SSU rRNA of *Lokiarchaeota*. Based on our observations, we explore possible evolutionary pathways of ES growth in Asgard archaea and Eukarya.

## Results

### Comparative analysis reveals broad patterns of complete LSU rRNA size relationships

Previously, we developed the SEREB MSA (Sparse and Efficient Representation of Extant Biology, Multiple Sequence Alignment) for comparative analysis of sequences from the translation system (Bernier, et al. 2018). The SEREB MSA is alignments of a sparse and unbiased representation of all major phyla and is designed specifically to understand phenomena that are universal to all of biology. The MSA was manually curated and extensively cross-validated. The SEREB MSA is useful as a seed to study a variety of evolutionary phenomena. Recently, we augmented the SEREB MSA to include additional metazoan sequences, allowing us to characterize ESs and their evolution in metazoans (Mestre-Fos, et al. 2019a; Mestre-Fos, et al. 2019b). Here, we augmented the SEREB MSA to include 21 sequences from the Asgard superphylum and 12 sequences from deeply branching eukaryotes (**supplementary datasets S1,2**).

The SEREB MSA indicates that LSU rRNA size relationships are governed by the general pattern: Bacteria (2725-2960 nts, n=61 [n is number of species]) < non-Asgard Archaea (2886 to 3094 nts, n=43) < Eukarya (3300-5200 nts, n=41; **supplementary figure S1**). This pattern is broken by eukaryotic parasites (3047-4086 nts, n=2) and Asgard archaea (3038-3300 nts, n=12). The SEREB MSA confirms previous observations that archaeal rRNAs frequently contain μ-ESs at positions of attachment of eukaryotic ESs. For example, in Archaea, μ-ES39 is connected at the site of attachment of the large ES39 in eukaryotes (Armache, et al. 2012).

### *Lokiarchaeota* are intermediate between Eukarya and Archaea in LSU rRNA size

The Asgard augmentation of the SEREB MSA revealed unexpectedly large *Lokiarchaeota* LSU rRNAs. *Lokiarchaeota* LSU rRNAs range from 3100 to 3300 nts (n=7). *Lokiarchaeota* rRNAs are close to or within the observed size range of eukaryotic LSU rRNAs (**supplementary figure S1**). Other Asgard species have LSU sizes in the upper archaeal range (from 3038 to 3142 nts, n=6). Supersized archaeal ESs, which rival eukaryotic ESs, are observed in *Lokiarchaeota* and *Heimdallarchaeota* spp. These ESs connect to the universal common core rRNA at the sites of attachment of eukaryotic ES9 and ES39 and archaeal μ-ES9 and μ-ES39 (**Fig. 1**). Here we explored the Asgard augmentation of the SEREB MSA to investigate the structure, distribution, and evolution of rRNA expansions of Asgard archaea.

### ES9 and ES39 in some Asgard archaea are larger than μ-ESs of other archaea and ESs of various protists

The MSA shows that ES39 in *Lokiarchaeota* ranges in size from 95 to 200 nts and in *Heimdallarchaeota* is from 115 to 132 nts. Other Asgard species have shorter ES39s: *Thorarchaeota*, *Odinarchaeota*, and *Freyarchaeota* ES39 sequences are between 50-60 nts. These Asgard ES39 sizes are well within the range of eukaryotic ES39, which is around 80 nts in a variety of protists, 138 nts in *Saccharomyces cerevisiae,* 178 nts in *D. melanogaster,* and 231 nts in *Homo sapiens* (**Fig. 2**, **supplementary data S3)**. For *Candidatus Lokiarchaeota* archaeon 1244-F3-H4-B5 (*Lokiarchaeota* B5), the primary focus of our work here, ES39 is 191 nts **(Figs. 2**, **3**). The first cultured *Lokiarchaeota* (*Promethearchaeum. syntropicum*) and one other *Lokiarchaeota* sequence have 95 nt ES39s (**Fig. 2**, **supplementary data S2**), slightly below our definition of a supersized ES, but still more than twice as long as the largest archaeal μ-ES39, *Pyrococcus furiosus* (45 nts, **Fig. 1C**, **supplementary dataset S3)**. All Asgard have larger ES39 sequences than other archaea. Archaea from the *Euryarchaeota* phylum, except for the classes *Methanococci* and *Thermococci*, lack ES39 entirely (less than 5 nts) and μ-ES39 of species from other non-Asgard archaea varies between 0 and 45 nts (**supplementary dataset S3**).

**Figure 2:**
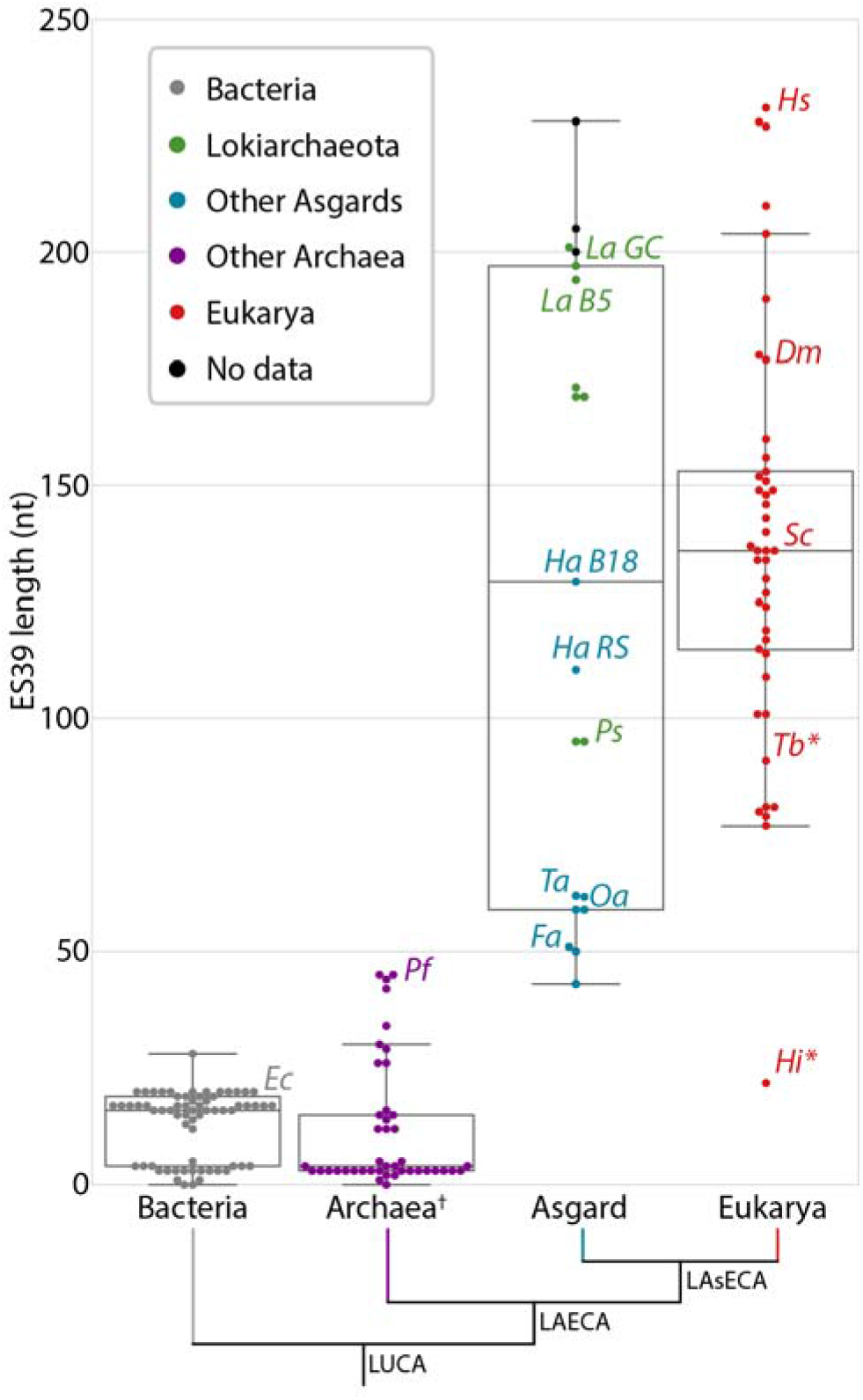
Distribution of ES39 lengths from the three domains of life, including Asgard archaea. The number of nucleotides calculated between alignment positions 4891 and 5123 (*H. sapiens* numbering) of the LSU alignment for each species (**supplementary dataset S1**). The box shows the quartiles of the dataset. Whiskers extend to show the rest of the distribution, except for points that are determined to be outliers using a function of the inter-quartile range. Bacteria sequences are gray, *Lokiarchaeota* sequences are green, other Asgard sequences are blue, other archaeal sequences are purple, eukaryotic sequences are red, and sequences from metatranscriptomic contigs (**supplementary dataset S2**) for which there is no species determination are black. Abbreviations: *Ec: Escherichia coli*; *Pf*: *Pyrococcus furiosus*; *La B5: Lokiarchaeota B5*; *La GC: Lokiarchaeota GC14_75*; *Ha B18: Heimdallarchaeota* B18G1; *Ha RS: Heimdallarchaeota* RS678; *Ps: Prometheoarchaeum syntropicum*; *Ta: Thorarchaeota*; *Oa*: *Odinarchaeota LCB4*; *Fa*: *Freyarchaeota*; *Hs*: *Homo sapiens; Dm*: *Drosophila melanogaster*; *Sc*: *Saccharomyces cerevisiae*; *Tb*: *Trypanosoma brucei*; *Hi*: *Hexamita inflata*. *: parasitic representatives of the deeply branching eukaryotic group Excavata. †: Excluding the Asgard superphylum.

**Figure 3:**
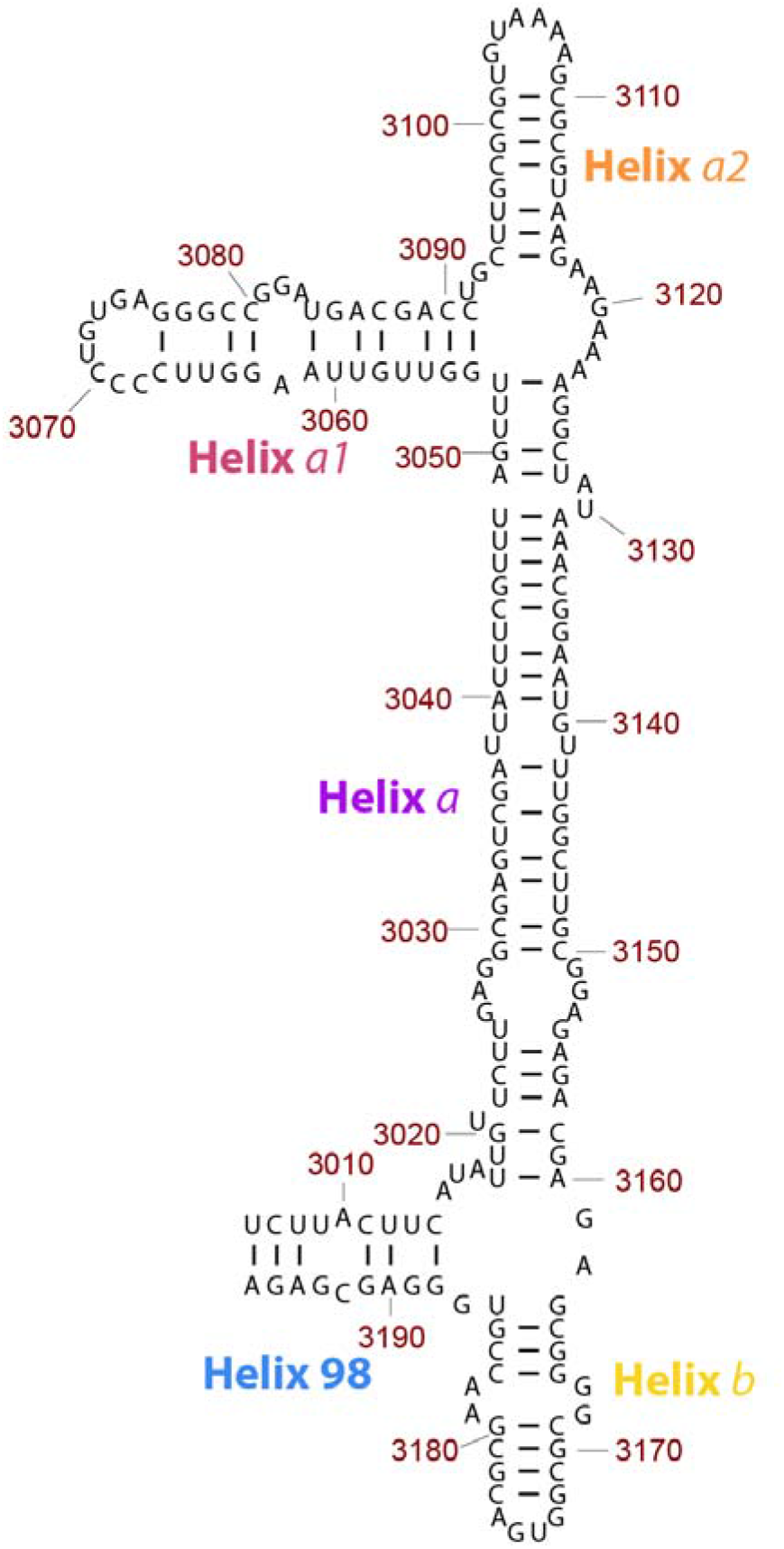
Secondary structure model of *Lokiarchaeota* B5 ES39. ES39 spans nucleotide positions 3006-3196 of the *Lokiarchaeota* B5 LSU rRNA. Canonical base-pairing positions are indicated with black lines. Helices are annotated with colored labels: blue – Helix 98, purple – Helix *a*, pink – Helix *a1*, orange – Helix *a2*, gold – Helix *b*. Figure was generated with RiboVision (Bernier, et al. 2014).

Amongst Asgard archaea, some *Lokiarchaeota* spp. exhibit large ES9, excluding *P. syntropicum*. *Lokiarchaeota* ES9 ranges from 29 to 103 nts (103 nts in *Lokiarchaeota* B5, 29 nts in *P. syntropicum*), compared to 29 nts in *S. cerevisiae,* 44 nts in *D. melanogaster* and *H. sapiens*, and 111 nts in *Guillardia theta* (**supplementary figure S2**). In other non-Asgard archaea, ES9 varies between 20-30 nts. ES9 and ES39 are the primary contributors to the large size of *Lokiarchaeota* LSU rRNAs compared to the LSU rRNAs of other archaea. Outside of the Asgard superphylum, archaea lack supersized ESs.

### Supersized ESs of *Lokiarchaeota* are transcribed *in situ*

To assess whether *Lokiarchaeota* ESs are transcribed, we assembled metatranscriptomic reads from sediment from the Gulf of Mexico, known to contain *Lokiarchaeota* sequences (Yergeau, et al. 2015; Cai, et al. 2018). We did not find metatranscriptomes that contain sequences from *Heimdallarchaeota*. Multiple transcripts from *Lokiarchaeota*-like LSU ribosomes contain ES9 and ES39 sequences, confirming that *Lokiarchaeota* ESs are indeed transcribed *in situ* **(Fig. 4D**; **supplementary dataset S3)**.

**Figure 4:**
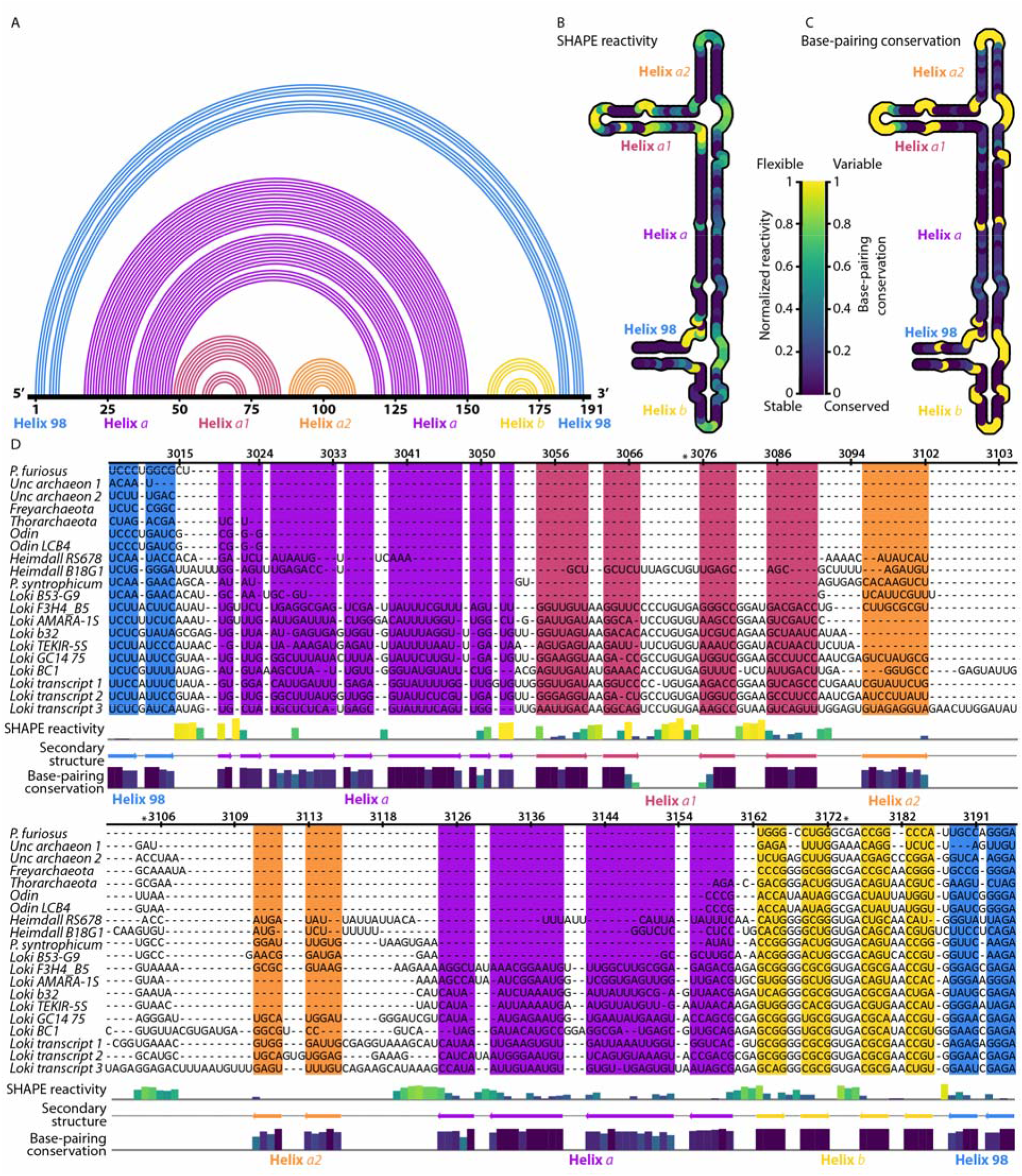
Secondary structure of *Lokiarchaeota* B5 ES39 from experiment and computation. (A) 1-D topology map of base pairs. The primary sequence of ES39 is on the horizontal line, Arcs indicate base pairs. Each helix is a distinct color. (B) SHAPE reactivity for ES39 mapped onto the secondary structure. Darker color indicates less flexible (paired) rRNA. (C) Base pairing covariation within the Asgard superphylum mapped on the secondary structure. Darker color indicates covarying (paired) rRNA across Asgard. Unpaired rRNA, for which no covariation data can be calculated, is gold. (D) SHAPE reactivity and base-pairing conservation mapped onto the ES39 MSA of Asgard sequences. *Lokiarchaeota* B5 numbering was used. The secondary structure is indicated with colored arrows bellow the alignment and as colored background. SHAPE reactivity is indicated with a bar graph above the secondary structure annotation, colors of the bars are consistent with panel B. Base-pairing conservation is indicated with a bar graph bellow the secondary structure annotation; colors of the bars are consistent with panel C. Panel D was generated with Jalview (Waterhouse, et al. 2009). Helices are labeled with colored text in each panel; blue, helix 98; violet, helix *a*; pink, helix *a1*; orange, helix *a2*; yellow, helix *b*. Full sequence names and sequencing project identifiers are available in **supplementary dataset S2**. Both SHAPE reactivity and covariation are normalized. *: positions of conserved GNRA tetraloops.

### Asgard LSU rRNA contain the common core

We determined the extent of structural similarity of *Lokiarchaeota* LSU rRNAs with those of eukaryotes. We combined computation and experiment to model the secondary structure of *Lokiarchaeota* B5 LSU rRNA (**Fig. 1E; supplementary figure S3**). Secondary and three-dimensional structures of several eukaryotic and archaeal ribosomes provided a basis for modeling by homology. Like all other LSU rRNAs, *Lokiarchaeota* LSU rRNA contains the rRNA common core, which is trivial to model because the backbone atoms of the common core are highly conserved in all cytosolic ribosomes.

### ES39 has a well-defined structural core in eukaryotes

Before comparing eukaryotic and archaeal ES39s, we characterized the conserved core of eukaryotic ES39, which we call the ES39 signature fold. We compared experimental three-dimensional structures of rRNAs in a wide range of eukaryotic ribosomes (Ben-Shem, et al. 2010; Klinge, et al. 2011; Khatter, et al. 2015; Li, et al. 2017). The ES39 signature fold, which is common to all eukaryotic ES39s, consists of helix 98 (H98; 20-30 nts), helix *b* (H*b*; 40-50 nts), and their linkage by three unpaired 15 nts long segments of rRNA (**supplementary figure S4)**. These unpaired segments are tightly associated with the ribosomal surface. In all eukaryotes, two of the unpaired segments interact with eukaryotic-specific α-helical extensions on rProteins uL13 and eL14 **(supplementary figure S6).** The third unpaired segment interacts with ES7 and rProtein aL33 **(supplementary figures S4, S7, S8)**(Khatter, et al. 2015). H*b* is terminated by a conserved GNRA tetraloop (**supplementary figures S4 and S5**). The ES39 signature fold is conserved in structure but not in sequence.

### The ES39 signature fold can be decorated by an additional helix

Many eukaryotes possess a third helix (H*a*) that projects from the ES39 signature fold **(supplementary figure S4)**. H*a* is variable in length, it is shortest in unicellular eukaryotes, such as *Tetrahymena thermophila* (no helix), *Toxoplasma gondii* (10 nts), and *S. cerevisiae* (18 nts). Metazoan eukaryotes have the longest H*a*, such as *D. melanogaster* (20 nts) and representatives of *Chordata* (106 nts for *H. sapiens*; **supplementary figure S4**).

### Initial *Lokiarchaeota* B5 ES39 secondary structure models were predicted by two methods

A preliminary secondary structural model of ES39 of *Lokiarchaeota* B5 was generated using mfold (Zuker 2003) (**Fig. 3**). The program mfold predicts a minimum free energy secondary structures using experimental nearest-neighbor parameters. We selected the mfold model with lowest free energy for further investigation. A preliminary secondary structural model assumed *Lokiarchaeota* ES39 was structurally homologous with ES39 of *H. sapiens*. The mfold model was confirmed to be correct, and the *H. sapiens* homology model was determined to be incorrect by covariation analysis and Selective 2’ Hydroxyl Acylation analyzed by Primer Extension (SHAPE) reactivity data as described below.

### Covariation supports the mfold model for the secondary structure of ES39 of *Lokiarchaeota* B5

Covariation, or coupled changes in paired nucleotides across phylogeny, can help reveal RNA secondary structure (Levitt 1969; Ninio, et al. 1969; Woese, et al. 1980; Noller, et al. 1981; Gutell, et al. 1993; Gutell, et al. 1994). Base-pairing relationships can be detected through covariation. We compared co-variation predictions of each secondary model with observed co-variation in the MSAs. The secondary structure predicted by mfold is consistent with observed covariation **(Fig. 4)** while secondary structure predicted by *H. sapiens* homology model is not. The observation of covarying nucleotides supports the model determined by mfold.

### Chemical footprinting confirms the mfold model for secondary structure of *Lokiarchaeota* B5 ES39

We further tested the secondary structural model of ES39 from *Lokiarchaeota* B5 using SHAPE. This experimental method provides data on RNA flexibility at nucleotide resolution (Merino, et al. 2005; Wilkinson, et al. 2006). SHAPE reactivity is generally high for unpaired nucleotides, which are more flexible, and low for paired nucleotides, which are less flexible. SHAPE has been widely used to probe the structure of rRNA (Leshin, et al. 2011; Lavender, et al. 2015; Gomez Ramos, et al. 2017; Lenz, et al. 2017) and other RNAs (Wilkinson, et al. 2005; Gilbert, et al. 2008; Stoddard, et al. 2008; Watts, et al. 2009; Novikova, et al. 2012; Spitale, et al. 2013; Huang, et al. 2014). The SHAPE results from *Lokiarchaeota* B5 ES39 rRNA (**supplementary figure S9**) are in agreement with the secondary structure based on mfold, which, as shown above, is consistent with observed co-variation. Reactivity is low for paired nucleotides in the mfold model and is high in loops and bulges (**Fig. 4B**). The accuracy of the SHAPE data is supported by the observation of relatively high reactivity at the vast majority of unpaired nucleotides and low reactivity for most paired nucleotides of the mfold model. The *Lokiarchaeota* SHAPE data are not consistent with models that force *Lokiarchaeota* ES39 to conform to the *H. sapiens* secondary structure.

### Asgard ES39 deviates from the eukaryotic ES39 signature fold

In eukaryotic ES39, the junction of helices H98, H*a*, and H*b* contains three unpaired segments, each 15 nts. In *Lokiarchaeota* B5, ES39 lacks unpaired segments greater than 8 nts and contains fewer unpaired nucleotides **(Fig. 3**). *Lokiarchaeota* B5 ES39 is composed of four short helices, each up to 38 nts (H98; Helix *a1:*H*a1*; Helix *a2:*H*a2*; and H*b*), and one long helix (H*a*: 72 nts). H98 and H*b* connect in a three-way junction with H*a* at the base of ES39. H*a1* and H*a2* split H*a* at the top of ES39 in a three-way junction **(Fig. 3**).

We modeled and visualized secondary structures of ES39 sequences from additional Asgard species (**Fig. 5**, **supplementary figure S10**). None of these secondary structures exhibited unpaired regions longer than 10 nts. ES39 of all Asgard archaea contain a three-way junction that connects H98, H*a,* and H*b*, as observed in ES39 of *Lokiarchaeota* B5 (**Fig. 3**). Such three-way junctions can indicate locations at which accretions in the ribosomal structure took place (Petrov, et al. 2014b). Furthermore, ES39 in *Heimdallarchaeota* B18G1 has an additional branching of H*a* into H*a1* and H*a2,* mirroring the morphology of ES39 in *Lokiarchaeota* B5 (**Fig. 5**, **supplementary figure S10**). Despite the common branching morphology, the length of the individual helices varies substantially between different species (**Figs. 4D**, **5**).

**Figure 5:**
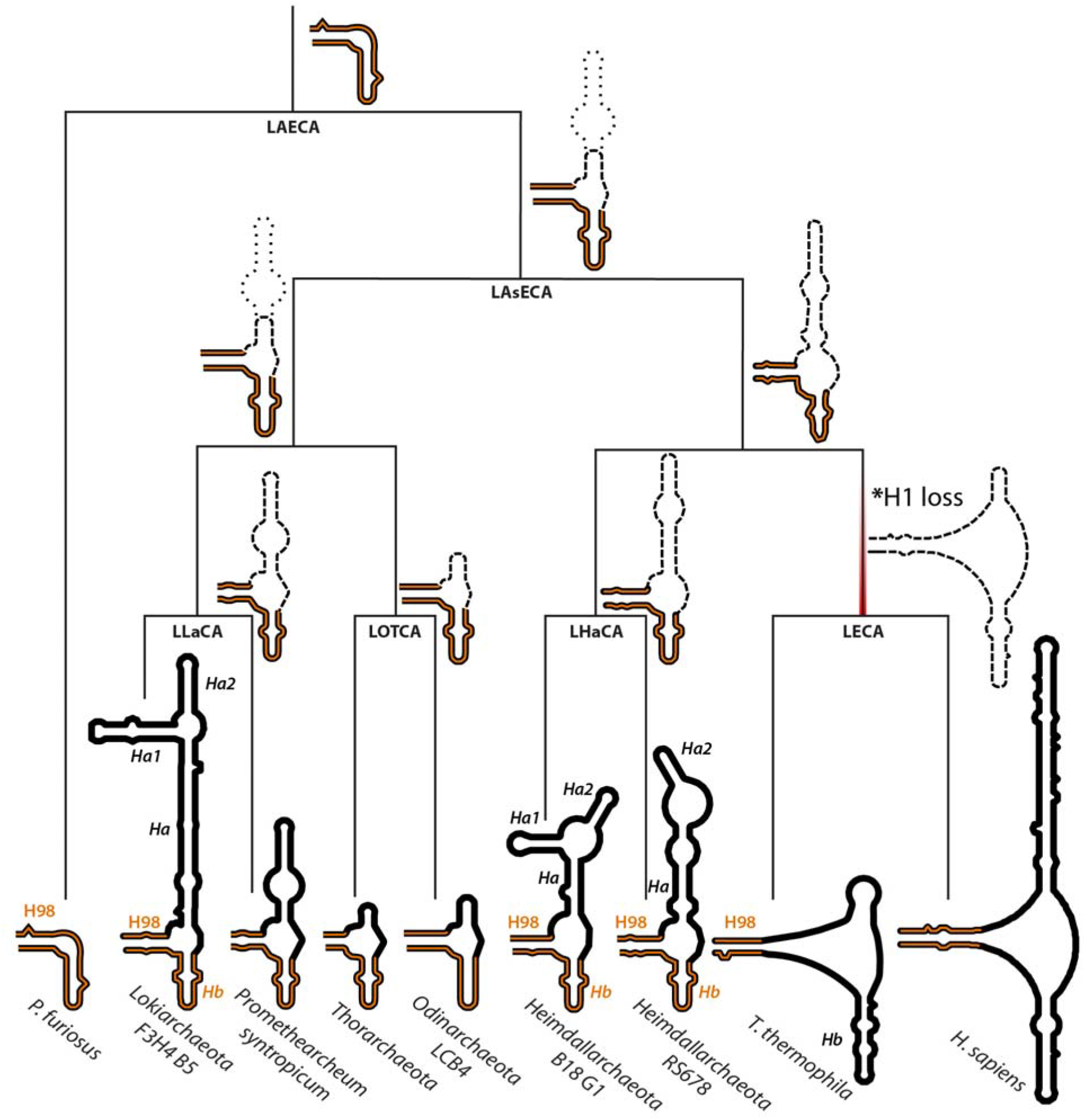
ES39 secondary structures mapped on Asgard phylogeny. Secondary structures of *Pyrococcus furiosus* (Archaea), *Lokiarchaeota* B5 (Asgard archaea), *Promethearchaeum syntropicum (*Asgard archaea), *Thorarchaeota* (Asgard archaea), *Odinarchaeota LCB4* (Asgard archaea), *Heimdallarchaeota B18 G1* and *RS678* (Asgard archaea), *Tetrahymena thermophila* (Eukarya), and *Homo sapiens* (Eukarya). Helix abbreviations are the same as in **Fig. 1E**. Helices 98 and *b* are highlighted in orange. The phylogenetic tree topology is from Williams, et al. (2020). Ancestral clades on the phylogenetic tree are labeled as LAECA: Last Archaeal and Eukaryotic Common Ancestor; LAsECA: Last Asgardian and Eukaryotic Common Ancestor; LLaCA: Last Lokiarchaeal Common Ancestor; LOTCA: Last Odin- and Thor-archaeota Common Ancestor; LHaCA: Last Heimdallarchaeota Common Ancestor; LECA: Last Eukaryotic Common Ancestor. Possible ancestral structures are indicated next to ancestral clades. Minimal common structure from related extant species is shown with bold dashes. The uncertainty in ancestral sizes of helix *a* are shown with light dashes. *: Occurrence of strand dissociation in ES39 at LECA due to loss of helix 1 is indicated with a red gradient.

### Asgard ES39 is located within an archaeal structural environment in the ribosome

ES39 in Eukarya protrudes from helices 94 and 99 of the ribosomal common core **(Fig. 6)**. In three dimensions, ES39 is close to ES7 and rProteins uL13, eL14, and aL33 (**supplementary figures S4, S6, S7**). These elements in all Asgard species are more similar to Archaea than to Eukarya. Additionally, ribosomes of both *Lokiarchaeota* and *Heimdallarchaeota*, like those of all Archaea, contain helix 1 (H1), which is in direct contact with H98 at the base of ES39, whereas eukaryotes lack H1. Combined with the large size of *Lokiarchaeota* and *Heimdallarchaeota* ES39, these characteristics predict that ribosomes from these two Asgard groups have a unique structure in this region.

**Figure 6:**
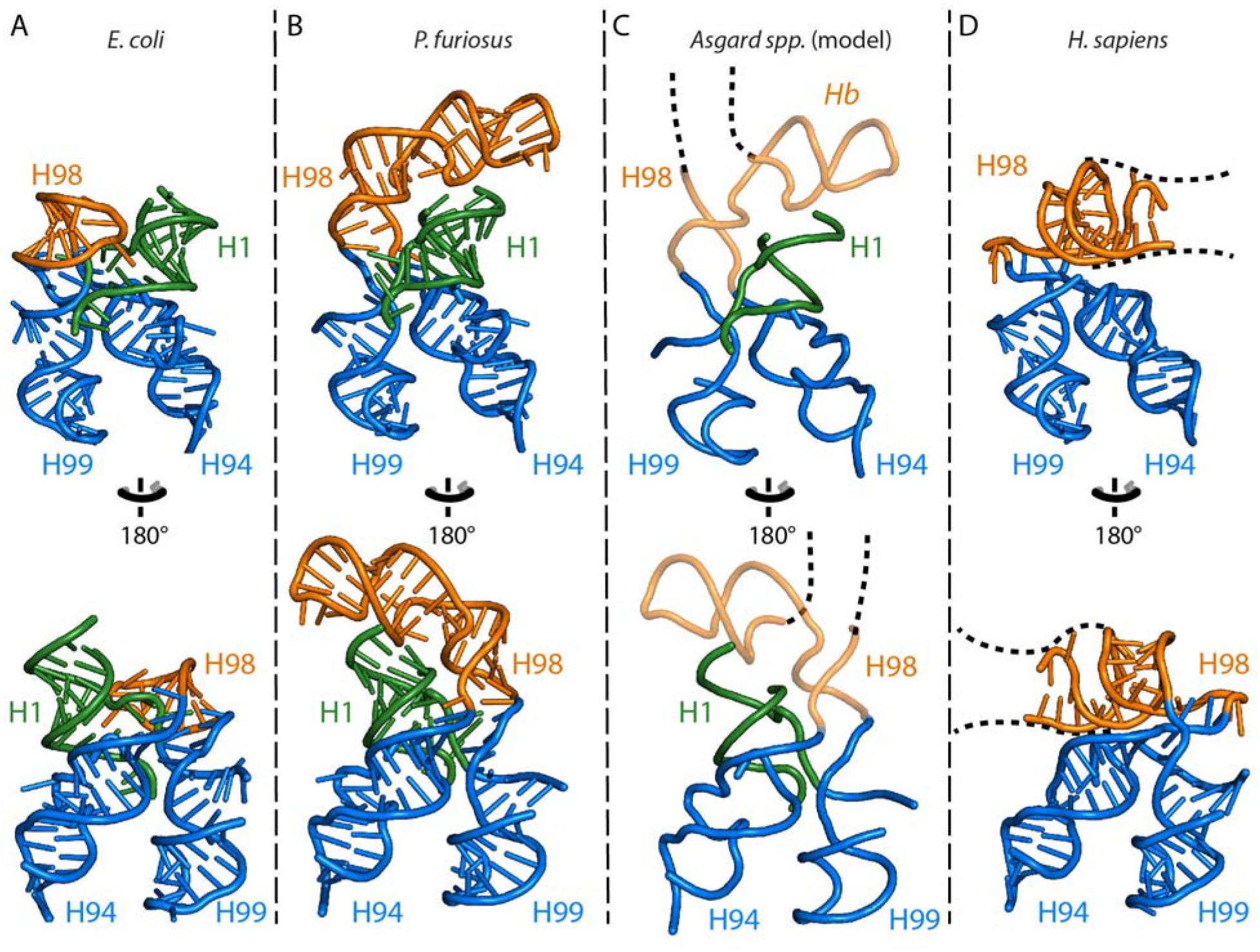
Three dimensional structures of ES39 and its neighborhood from representative species of the major life domains and Asgard archaea. (A) *Escherichia coli* (Bacteria), (B) *Pyrococcus furiosus* (Archaea), (C) Asgard archaea; (D) *Homo sapiens* (Eukarya). Helices are abbreviated as helix 1 (H1), helix 94 (H94), helix 98 (H98), helix 99 (H99) and helix *b* (H*b*). Helix 98 is orange, helix 1 is green, helices 94 and 99 are blue. The *P. furiosus* structure is used as model for the Asgard structures. Likely structure of Asgard helices 98 and *b* are shown with transparency. Likely position and direction of the Asgard ES39 continuation is indicated with a black dashed line. Direction of eukaryotic ES39 continuation is indicated with a black dashed line.

### The pathway of ES39 evolution appears to be unique

In general, ESs have increased in size over evolution via accretion. Growth by insertion of one rRNA helix into another leaves a structural mark. These growth processes are indicated by the joining of a rRNA helical trunk to an rRNA helical branch at a highly localized three- or four-way junction (an insertion fingerprint) (Petrov, et al. 2014b; Petrov, et al. 2015). Basal structure is preserved when new rRNA is acquired. For instance, ES7 shows continuous growth over phylogeny, expanding from LUCA to Archaea to diverse protist groups to metazoans to mammals (Armache, et al. 2012; Petrov, et al. 2014b; Bernier, et al. 2018). The accretion model predicts that H98, at the base of ES39, would superimpose in Bacteria, Archaea, and Eukarya, but in fact H98 does not overlap in superimposed 3D structures (**Fig. 6**). The archaeon *P. furiosus* has a slightly extended and bent H98 compared to the bacterium *E. coli* (**Fig. 6**). This spatial divergence is likely due to the difference in how *E. coli* H98 and *P. furiosus* H98 interact with H1 of the LSU. *E. coli* H98 interacts within the H1 minor groove through an A-minor interaction, while in *P. furiosus* H98 is positioned on top of H1 (**Fig. 6**). H1 is absent in eukaryotes (**Fig. 1**), allowing H98 to occupy the position of H1 (**Fig. 6**) and cause a strand dissociation in the eukaryotic ES39 structure.

### Structurally, *Lokiarchaeota* and *Heimdallarchaeota* ES39 may extend in a different direction than eukaryotic ES39

*Lokiarchaeota* and *Heimdallarchaeota* spp. have larger ES39 than other archaea (**Fig. 2**) and possess H1, unlike Eukarya (**Fig. 6**). We predict that *Lokiarchaeota* and *Heimdallarchaeota* ES39 have an archaeal-like interaction with H1 through H98 and H*b* (**Fig. 6)**. In Asgard species, the highly variable H*a* (**Figs. 4D**, **5**) likely grows out from the three-way junction between H98 and H*b*, perpendicular to the eukaryotic H*a* (**Fig. 6)**. While H*a* of eukaryotic ES39 is pointed in the direction of the sarcin-ricin loop, the large H*a* of *Lokiarchaeota* and *Heimdallarchaeota* is likely pointed in the direction of the central protuberance or the exit tunnel. The difference in directionality between Asgard and eukaryotic ES39 arises from the loss of H1 in Eukarya, where H98 and H*b* partially occupy the vacated space.

### Weak sequence homology between ES39 sequences

We searched for ES39 homology between Asgards and deeply branching protist groups. MSAs between Asgard (n=16) and eukaryotic (n=15) ES39 sequences were compared with eukaryotic-only and Asgard-only MSAs. The percent identities in the eukaryotic-only ES39 MSA varied from 6 to 90% (**supplementary figure S11**). As expected, more closely related eukaryotes had greater identities (40-90% within *Chordata*; **supplementary figure S11**). Common core rRNA, with up to 80% identity even between distantly related species, was much more conserved than ES rRNA, (**supplementary figure S12**). Within the *Lokiarchaeota*, ES39 sequences shared 40-60% identity (**supplementary figure S13**). The extent of similarity between Asgard and eukaryotic ES39 sequences (10-30% identities; **supplementary figure S14**) was similar to the extent of similarity between distantly related eukaryotes (**supplementary figure S11**).

### Conservation of a GNRA-tetraloop capping ES39 H*b* between Eukarya and Asgard

The MSA of H*b* from ES39 shows a GNRA tetraloop that is conserved in both eukaryotes and Asgard archaea. Three sequences of deeply branching eukaryotic representatives exhibit sequence similarity with H*b* sequences from Asgard archaea (**supplementary figure S5**).

## Discussion

The recent discovery of the archaeal Asgard superphylum, which contain multitudes of ESPs, has redefined our understanding of eukaryotic evolution (Spang, et al. 2015; Zaremba-Niedzwiedzka, et al. 2017). Incorporation of Asgard species into phylogenies has changed our perspective on the relationship of Archaea and Eukarya in the tree of life (Hug, et al. 2016; Fournier and Poole 2018; Doolittle 2020; Williams, et al. 2020). Here, we incorporate rRNA secondary structure and three-dimensional interactions into comparative analyses. Our work extends structure-based methods of comparative analysis to *Lokiarchaeota* and *Heimdallarchaeota* rRNAs, and mechanistic models for the evolution of common rRNA features in Eukarya and some Asgard archaea. In the following sections, we explore possible evolutionary pathways for this rRNA region.

### *Lokiarchaeota* and *Heimdallarchaeota* rRNAs adhere to most rules of ribosomal evolution

The ribosome is a window to relationships among organisms (Woese and Fox 1977; Hillis and Dixon 1991; Olsen and Woese 1993; Fournier and Gogarten 2010; Hug, et al. 2016) and was a driver of ancient evolutionary processes (Agmon, et al. 2005; Smith, et al. 2008; Bokov and Steinberg 2009; Fox 2010; Petrov, et al. 2015; Melnikov, et al. 2018; Venkataram, et al. 2020). Previous work has revealed rules of ribosomal variation over phylogeny (Hassouna, et al. 1984; Gerbi 1996; Melnikov, et al. 2012; Bernier, et al. 2018), as well as mechanisms of ribosomal change over evolutionary history (Petrov, et al. 2014b; Petrov, et al. 2015; Kovacs, et al. 2017; Melnikov, et al. 2018). Building off those studies, we assessed the extent to which *Lokiarchaeota* and *Heimdallarchaeota* ribosomes follow or deviate from previously established rules of ribosomal sequence and structure variation (Bowman, et al. 2020). We found that *Lokiarchaeota* and *Heimdallarchaeota* ribosomes:

- contain the universal common core of rRNA and rProteins (this work) (Spang, et al. 2015; Bernier, et al. 2018),
- confine rRNA variability of structure and size to ESs/μ-ESs (this work),
- restrict ESs to universally conserved sites on the common core (Ware, et al. 1983; Clark, et al. 1984; Hassouna, et al. 1984; Michot and Bachellerie 1987; Bachellerie and Michot 1989; Lapeyre, et al. 1993; Gerbi 1996),
- avoid ES attachment from the ribosomal interior or near functional centers (Ben-Shem, et al. 2010; Anger, et al. 2013), and
- concentrate variability in structure and size on LSU rRNA, not SSU rRNA (Gerbi 1996; Bernier, et al. 2018).

The ribosomes of *Lokiarchaeoacomprise the minimal form of ES39 shared t*, but not of *Heimdallarchaeota,* violate the established pattern of increasing rRNA length in the order: Bacteria < Archaea ≪ Eukarya (Melnikov, et al. 2012; Petrov, et al. 2014b; Bernier, et al. 2018). *Lokiarchaeota* LSU rRNAs are much longer than predicted for archaea; in fact, *Lokiarchaeota* rRNA eclipses the length of rRNA in many deeply branching eukaryotes. Notably, the ES9 size of the only yet cultured *Lokiarchaeota*, *P. syntropicum* (Imachi, et al. 2020) is an outlier to other *Lokiarchaeota* metagenome derived sequences. *P. syntropicum* ES9 has a similar size to non-Asgard archaea (**supplementary figure S2**). ES39 in all studied *Lokiarchaeota* are larger than in any other archaea known to date. Some *Lokiarchaeota* exhibit ES39 larger than ES39 of most eukaryotes.

### Two scenarios for the evolution of ES39

Two scenarios could explain the similarities of ES39 in Eukarya and Asgard archaea. One scenario is based on a three-domain tree of life and predicts parallel evolution of ES39 (**Fig. 7A**), while the other assumes close common ancestry of Asgard archaea and Eukarya (**Fig. 7B**). Our results do not exclude either one completely; therefore, we will discuss both.

**Figure 7:**
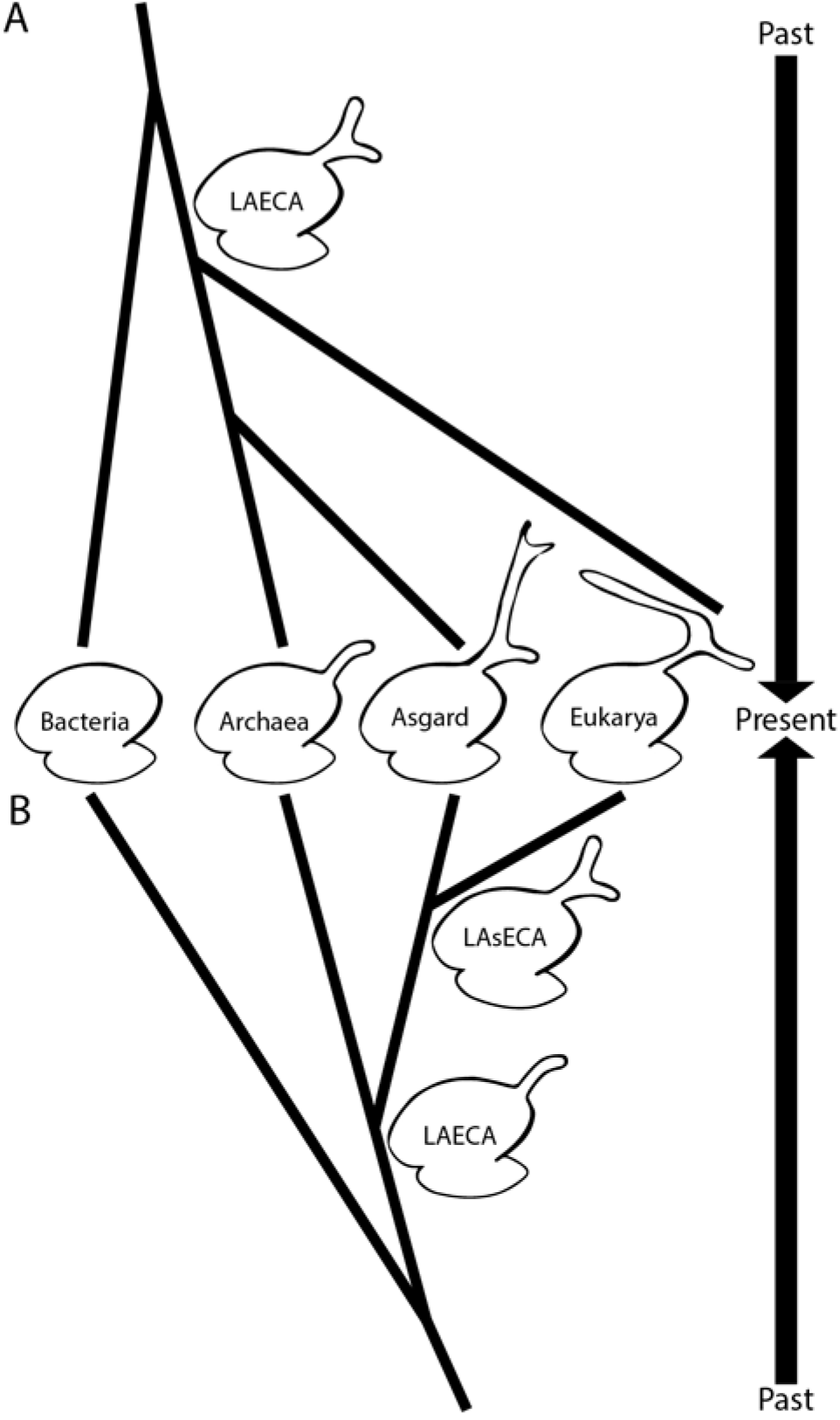
Two scenarios of ES39 evolution dependent on the tree of life topology. A) Parallel independent growth of the ES39 region between Asgard archaea and Eukarya with a three-helical ES39 in LAECA. B) Shared ancestry of three-helical ES39 between Asgard and Eukarya. Ribosomes of extant species are shown in the middle and past phylogenetic relationships extend up and down following the two scenarios. ES sizes are exaggerated for illustrative purposes.

### Parallel evolution of ES39 between Asgard archaea and Eukarya

The difference in directionality of H*a* in Asgard and Eukarya combined with the high evolutionary rates in ESs, and the lack of structural or sequence similarity, implies parallel evolution of ES39 with differential functions. Previous research has shown that archaeal ribosomes can contain μ-ESs (Armache, et al. 2012; Petrov, et al. 2014b) and large ES in a novel LSU region (Tirumalai, et al. 2020). It is possible that LAECA had a larger ES39 (**Fig. 7A**), which shrank in most archaea and was independently developed in Asgard archaea and eukaryotes. This theory is supported by the large structural diversity of ES39 in Asgard archaea. Loss of archaeal ES also matches previously documented reductions of archaeal rProteins (Lecompte, et al. 2002) and ribosome biogenesis factors (Ebersberger, et al. 2014). The parallel growth of LSU ESs in Asgard and Eukarya could also be the result of similar selection pressures. An explanation of similar ES sizes between *Lokiarchaeota*, *Heimdallarchaeota,* and higher eukaryotes (e.g. *Chordata*) may be slow growth rates and small population sizes, which are associated with characteristics that would otherwise be eliminated by purifying selection (Lynch 2007; Rajon and Masel 2011). Both *Lokiarchaeota* and *Chordata* have low growth rates and small populations (Imachi, et al. 2020), which might lead to larger size of ES rRNA.

### Possible common ancestry of ES39 in Asgard and Eukarya

Supersized ESs were thought to be unique to eukaryotic ribosomes. We now know that to be incorrect: two Asgard phyla exhibit supersized ESs. ES sequences are highly variable, even among closely related species (**supplementary figure S11**), making ESs impractical for sequence-based determination of ancient ancestry. Secondary and three dimensional structures are more conserved than sequences, allowing a more accurate detection of distant ancestry between species (Woese, et al. 1990a). By integrating the accretion model of ribosomal evolution (Petrov, et al. 2014b; Petrov, et al. 2015) and structure-informed phylogeny, we discuss the ancestral relationship of Eukarya and Archaea.

### The accretion model suggests ES39 in LAsECA contained three helices

An important structural difference between Asgard and other archaea is the three-helical topology of ES39. All Asgard have three helices in ES39, while other Archaea either lack ES39 altogether or contain a short-bent helix consisting of H98 and H*b* (**Figs. 2,5,6**). The accretion model predicts that ES39 of the Last Archaeal and Eukaryotic Common Ancestor (LAECA) contained H98 and H*b* (**Fig. 5**) because they comprise the minimal form of ES39 shared between Asgard and other archaea (**Figs. 4** and **5**). The insertion of H*a* between H98 and H*b* at LAsECA is consistent with the minimal ES39 (**Fig. 5**). Subsequently, the three-helical topology would be inherited by all Asgard species, consistent with variable H*a* sizes (**Figs. 4** and **5**). Similarly, eukaryotes have highly conserved H98 and H*b* while H*a* is variable (**supplementary figures S4 and S5**). Eukarya differ from Asgard archaea in the unpaired rRNA segments that connect H98, H*b*, and H*a* (**supplementary figure S4**). The absence of insertion fingerprints precludes a detailed model for the relative ages of eukaryotic H98, H*a*, and H*b*. It is reasonable to assume that LECA had a three-helical ES39 inherited from one of its Asgardian sister clades. This ancient three-helical ES39 underwent restructuring and strand dissociation upon the loss of H1 (**Figs. 5**, **6**). Subsequently, the unpaired structure of ES39 was inherited by all eukaryotes (**Fig. 5**).

The specific roles of μ-ESs and ESs over phylogeny are unknown but are likely complex, polymorphic, and pleotropic. The observation of μ-ESs in Archaea, ESs in Eukarya, and supersized ESs in Asgard species can be explained by either a two- or three-domain tree of life. In a two-domain model, the ES39 size and topology can be explained by a gradual growth and accretion of ribosomal complexity, consistent with a close relationship between Asgard archaea and Eukarya seen in Williams, et al. (2020). In a three-domain model, a large ES39 in LAECA could have been parallelly inherited in Asgard archaea and Eukarya without them necessarily being related. The rest of the archaeal domain experienced shrinkage of this ES region.

## Conclusions

*Lokiarchaeota* and *Heimdallarchaeota* ribosomes contain supersized ES39s with structures that are distinct from eukaryotic ES39s. *Lokiarchaeota* ES9s are larger than eukaryotic ES9s. To date, *Lokiarchaeota* and *Heimdallarchaeota* are the only prokaryotic phyla with supersized ESs, bringing the size of their LSUs close to those of Eukarya. *Lokiarchaeota* and *Heimdallarchaeota* ES39 likely grow outward from the ribosomal surface in a different direction than eukaryotic ES39s. Our findings raise the possibility that supersized ESs existed on the ribosomal surface before LECA, opening the question of whether ribosomal complexity is more deeply rooted than previously thought.

## Materials and Methods

### Genome sequencing, assembly, and binning

#### Sample collection

Sediments were cored from deep seafloor sediment at ODP site 1244 (44°35.1784°N; 125°7.1902°W; 895 m water depth) on the eastern flank of Hydrate Ridge ~3 km northeast of the southern summit on ODP Leg 204 in 2002 (Tréhu, et al. 2003) and stored at −80°C at the IODP Gulf Coast Repository.

#### DNA extraction

DNA was extracted from sediment from F3-H4 (18.10 meters below the seafloor) using a MO-BIO PowerSoil total RNA Isolation Kit with the DNA Elution Accessory Kit, following the manufacturer protocol without beads. Approximately 2 grams of sediments were used for the extraction from six extractions (12 g total) and DNA pellets from the two replicates from each depth were pooled together. DNA concentrations were measured using a Qubit 2.0 fluorometer with dsDNA High Sensitivity reagents (Invitrogen, Grand Island, NY, USA). DNA yield was 7.5 ng per gram of sediment.

#### Multiple displacement amplification, library preparation, and sequencing

Genomic DNA was amplified using a REPLI-g Single Cell Kit (Qiagen, Germantown, MD, USA) using UV-treated sterile plasticware and reverse transcription-PCR grade water (Ambion, Grand Island, NY, USA). Quantitative PCR showed that the negative control began amplifying after 5 hr of incubation at 30°C, and therefore, the 30°C incubation step was shortened to 5 hr using a Bio-Rad C1000 Touch thermal cycler (Bio-Rad, Hercules, CA, USA). DNA concentrations were measured by Qubit. Two micrograms of MDA-amplified DNA were used to generate genome libraries using a TruSeq DNA PCR-Free Kit following the manufacturer’s protocol (Illumina, San Diego, CA, USA). The resulting libraries were sequenced using a Rapid-Run on an Illumina HiSeq 2500 to obtain 100 bp paired-end reads. Metagenomic sequences were deposited into NCBI as accession numbers SAMN07256342-07256348 (BioProject PRJNA390944).

#### Metagenome assembly, binning, and annotation

Demultiplexed Illumina reads were mapped to known adapters using Bowtie2 in local mode to remove any reads with adapter contamination. Demultiplexed Illumina read pairs were quality trimmed with Trim Galore (Martin 2011) using a base Phred33 score threshold of Q25 and a minimum length cutoff of 80 bp. Paired-end reads were then assembled into contigs using SPAdes assembler (Bankevich, et al. 2012) with --meta option for assembling metagenomes, iterating over a range of k-mer values (21,27,33,37,43,47,51,55,61,65,71,75,81,85,91,95). Assemblies were assessed with reports generated with QUAST (Gurevich, et al. 2013). Features on contigs were predicted through the Prokka pipeline with RNAmmer for rRNA, Aragorn for tRNA, Infernal and Rfam for other non-coding RNA and Prodigal for protein coding genes (Laslett and Canback 2004; Lagesen, et al. 2007; Hyatt, et al. 2010; Nawrocki and Eddy 2013; Seemann 2014; Kalvari, et al. 2017). Each genomic bin was searched manually for 23S rRNA sequences. The *Lokiarchaeota* F3-H4-B5 bin (estimated 2.8% completeness and 0% contamination) was found to contain a 3300 nt 23S rRNA sequence. The *Lokiarchaeota* F3-H4-B5 bin was deposited into NCBI as BioSample SAMN13223206 and GenBank genome accession number WNEK00000000.

### Environmental 23S rRNA transcript assembly and bioinformatic analysis

#### Species selection for analysis

To efficiently sample the tree of life, we used species from the SEREB database (Bernier, et al. 2018) as a starting set. These species were selected to sparsely sample all major phyla of the tree of life where allowed by available LSU rRNA sequences. We further added LSU sequences from species of recently discovered phyla in the archaeal and eukaryotic domains (**supplementary dataset S2**). These additional species were again selected as representatives from major phyla. To gain higher level of evolutionary resolution in the Asgard superphylum we added complete LSU sequences from several available Asgard species and constructed assemblies from available meta-transcriptomes. All species used in our analysis with their respective phylogenetic groups are listed in **supplementary dataset S3**.

#### Assembly

Publicly available environmental meta-transcriptomic reads were downloaded from NCBI BioProject PRJNA288120 (Yergeau, et al. 2015). Quality evaluation of the reads was performed with FastQC (Andrews 2012) and trimming was done with TrimGalore (Martin 2011). Assembly of SRR5992925 was done using the SPADES (Bankevich, et al. 2012) assembler with --meta and --rna options, to evaluate which performs better. Basic statistic measures such as Nx, contig/transcript coverage and length were compared (**supplementary datasets S4, S5**) yielding better results for the rnaspades assembler. All subsequent meta-transcriptomic datasets were assembled with rnaspades.

#### Identifying ribosomal RNA sequences

BLAST databases were constructed (Altschul, et al. 1990) from the resulting contig files and they were queried for ribosomal regions characteristic of the Asgardian clade (ES39/ES9 sequences from GC14_75). Additionally, the program quast (Gurevich, et al. 2013) with --rna-finding option was used.

#### SEREB Multiple Sequence Alignment (MSA) augmentation

High scoring transcripts, as well as genomic sequences from added species, were included in the SEREB MSA (Bernier, et al. 2018) using the program mafft (Katoh and Standley 2013) with the --add option. Known intronic regions (Cannone, et al. 2002) were removed from new sequences. The highly variable region of ES39 was manually aligned using structural predictions from mfold (Zuker 2003).

#### Defining ES regions within Asgard archaea

To compare eukaryotic, archaeal, and Asgard ES regions we defined the basal helices of ES growth for ES9 and ES39. We calculated ES sizes by including all nucleotides within the start and end of the basal helices. In eukaryotes ES9 is a region of a cumulative expansion between helices 29, 30, and 31 (see region D3 in Hassouna, et al. (1984)). For ES9 we used helix 30 since the growth in Lokiarchaeota was contained within that helix. For ES39 we used H98, consistent with previous observations of the variability within that region (Gerbi 1996).

#### Sequence homology between Asgard and eukaryotic ES39 sequences

To uncover any sequence conservation within the ES39 region of the LSU between Asgard archaea and eukaryotes, we used the augmented SEREB MSA. The ES39 sequences for all eukaryotes and Asgard archaea were extracted from the global LSU alignment. Chordate eukaryotic sequences were removed as they were too GC rich, parasitic eukaryotic sequences were also removed. Multiple different automated alignment methods were used (Do, et al. 2005; Sievers, et al. 2011; Katoh and Standley 2013). Following the automated methods, we applied manual alignment guided by secondary structure predictions. Pairwise percent identities for the alignments where calculated with Ident and Sim from the Sequence Manipulation Suite (Stothard 2000).

#### LSU size comparison

The LSU size comparison was based on the transcribed gene for the LSU, which is comprised of a single uninterrupted rRNA sequence for bacteria and archaea (**Fig. 1A,C,E**), and is comprised of multiple concatenated rRNA sequences for the fragmented eukaryotic rRNA gene (**Fig. 1B,D,F**). The 5S rRNA, which is essentially constant, is excluded from the size calculation. The comparison takes into account whether the rRNAs are from endosymbionts and pathogens, which tend to contain reduced genomes, metabolisms, and translation systems (Peyretaillade, et al. 1998; Moran 2002; McCutcheon and Moran 2011).

#### Secondary structure models

To model the secondary structure of *Candidatus* Lokiarchaeota archaeon 1244-F3-H4-B5 LSU rRNA, we used the secondary structure of *P. furiosus* (Petrov, et al. 2014a) and a multiple sequence alignment (MSA) of archaeal LSU rRNAs broadly sampled over the phylogeny (**supplementary dataset S1**). Locations of expansion segments were unambiguously identified from the MSA. Due to the low percent identity (<50%) (Bernhart and Hofacker 2009) we applied *ab initio* modelling for ES regions. GNRA tetraloops are highly conserved structural motifs in rRNA structure (Woese, et al. 1990b; Heus and Pardi 1991; Nissen, et al. 2001). Conserved GNRA tetraloop sequences from the Asgard sequences were used to determine the limits of variable helices in ES regions. The secondary structures of the ESs were predicted by mfold (Zuker 2003).

#### Covariation

To verify the secondary structures of the highly variable ES regions base-pairing conservation was calculated with the program Jalview (Waterhouse, et al. 2009). Gaps from the MSA were ignored in the calculation to produce comparable results about available regions. The base-pairing model of secondary structures of ES9 (**supplementary figure S15**) and ES39 (**Figs. 4C,D**) was generated in the Jalview annotation format and used for the base-pairing conservation calculation.

#### Defining the eukaryotic ES39 signature old

To identify the structurally invariant part of ES39 in eukaryotes, we used superimposition based on the common core within domain VI of the ribosomal structures from 4 eukaryotes (*T. thermophila*, *T. gondii*, *S. cerevisiae*, *H. sapiens*; **supplementary figure S4**). Initially the *D. melanogaster* ribosomal structure (PDB ID: 4V6W) (Anger, et al. 2013) was used in identifying the core. However, as it has additional loops elongating the unpaired regions, we excluded it from our analysis. *D. melanogaster* is known to have AU-enriched ESs; therefore, it is not surprising that it has perturbations in its ES39.

#### Structural analysis of ES39 region in Bacteria, Archaea, and Eukarya

To identify the likely direction of Asgard ES39 we compared the ES39 region between Bacteria, Archaea, and Eukarya from available ribosomal structures. The structure used for Bacteria was from *E. coli* (PDB ID: 4V9D) (Dunkle, et al. 2011), for Archaea was from *P. furiosus* (PDB ID: 4V6U) (Armache, et al. 2012), and for Eukarya was from *H. sapiens* (PDB ID: 4V88) (Ben-Shem, et al. 2010). The model for ES39 H*b* for Asgard ES39 is based on ES39 H*b* from the *P. furiosus* ribosomal structure.

### ES39 rRNA SHAPE analysis

#### Synthesis of *Lokiarchaeota* ES39 rRNA

pUC57 constructs containing T7 promoter and the gene encoding *Lokiarchaeota* ES39 rRNA was linearized using HindIII restriction enzyme. *Lokiarchaeota ES39* rRNA was synthesized by *in vitro* transcription using HiScribe™ T7 High Yield RNA Synthesis Kit; New England Biolabs. RNA was then precipitated in ethanol/ammonium acetate and purified by G25 size exclusion chromatography (illustraTMNAPTM-10, GE Healthcare). RNA purity was assayed by denaturing gel electrophoresis.

#### SHAPE reaction

Selective 2′-hydroxyl acylation analyzed by primer extension (SHAPE; Wilkinson, et al. 2006) was performed to chemically probe local nucleotide flexibility in ES39 rRNA. *In vitro*-transcribed ES39 rRNA was added to folding buffer (180 mM NaOAc, 50mM Na-HEPES (pH 8.0) and 1 mM 1,2- diaminocyclohexanetetraacetic acid (DCTA)) to obtain 400 nM RNA in total volume of 80 μL. RNA was annealed by cooling from 75 °C to 25 °C at 1 °C/min. RNA modification reaction was performed with final concentration of 100 mM benzoyl cyanide (Sigma) prepared in dimethyl sulfoxide (DMSO). Non-modified RNA samples were incubated with DMSO. Reactions were carried out for 2 min at room temperature. Modified RNAs and control sample were purified by precipitation in ethanol and ammonium acetate at 20 °C for 2 hr. RNA samples were centrifuged at 4°C for 10 min. The RNA pellets were washed with 100 μL of 80% ethanol for two times and dried out using a SpeedVac vacuum concentrator. TE buffer [22 μL of 1 mM EDTA and 10 mM Tris-Cl (pH 8.0)] was added to each sample to resuspend the pellet.

#### Reverse transcription

Reverse transcription was conducted on 20 μL of modified RNAs and unmodified RNA sample as a control, in presence of 8 pmol 5’[6-FAM] labeled primer (5’-GAACCGGACCGAAGCCCG-3’), 2 mM DTT, 625 μM of each deoxynucleotidetriphosphate (dNTP), and 5 μL of reverse transcription (RT) 5X first-strand buffer [250 mM Tris-HCl (pH 8.3), 375 mM KCl, 15 mM MgCl_2_]. To anneal the primer, samples were heated at 95°C for 30 secs, held at 65°C for 3 min, and then 4°C for 10 min. RT mixtures were incubated at 52°C for 2 min before addition of 1 μL (200 U) of Superscript III Reverse transcriptase (Invitrogen) and reactions were incubated at 55°C for 2 hr. later, RT terminated by heating samples at 70°C for 15 min. Chain termination sequencing reaction was performed on 10 pmol unmodified RNA prepared in TE buffer, 8 pmol 5’[6-FAM] labeled primer, with a ratio of 1:1 dideoxynucleotide (ddNTP) to dNTP. A sequencing reaction was performed with the same condition without ddNTPs.

#### Capillary electrophoresis of RT reaction products and data analysis

Capillary electrophoresis of RT reactions was performed as described previously (Hsiao, et al. 2013). For each reaction 0.6 μL DNA size standard (Geneflo™ 625), 17.4 μl Hi-Di Formamide (Applied Biosystems), and 2 μL of RT reaction mixture were loaded in a 96-well plate. Samples were heated at 95°C for 5 min before electrophoresis and the RT products were resolved using applied biosystems. SHAPE data were processed using a Matlab scripts as described previously (Athavale, et al. 2012). SHAPE profile was mapped onto ES39 rRNA secondary structure with the RiboVision program (Bernier, et al. 2014).

## Acknowledgments

We thank Cecilia Kretz and Piyush Ranjan for sample preparation and genome binning. We thank Brett Baker for providing an unpublished *Heimdallarchaeota* sequence. We thank Jessica Bowman, Santi Mestre-Fos, Aaron Engelhart, Anthony Poole, and Betül Kaçar for helpful conversations.

## Author contributions

ASP, LDW, and JBG conceptualized the research. PIP curated the data and performed computational analysis. ASP, LDW, and JBG acquired the funding and administered the project. SF-A performed the SHAPE experiments. JJC and VJP performed analyses. ASP, LDW, RRG, and JBG supervised the research. RRG validated the research. PIP and VJP prepared the figures. PIP, JBG, and LDW wrote the manuscript with input from all authors.

## Data and materials availability

Metagenomic sequences were deposited into NCBI as accession numbers SAMN07256342-07256348 (BioProject PRJNA390944). The *Lokiarchaeota* F3-H4-B5 bin was deposited into NCBI as BioSample SAMN13223206 and GenBank genome accession number WNEK00000000. The *Lokiarchaeota* F3-H4-B5 23S rRNA gene is in the reverse complement of contig WNEK01000002.1, nucleotide positions 251-3579. Scripts used for generation of data figures and tables, as well as MSAs of the ES39 region are available at https://github.com/petaripenev/Asgard_LSU_project.

