## Supplementary Materials for "Supersized ribosomal RNA expansion segments in Asgard archaea"

**This PDF file includes:**

Supplementary Figures S1 – S15

Captions to Datasets S1-S5

References for SI citations

**Supplementary Figures:**

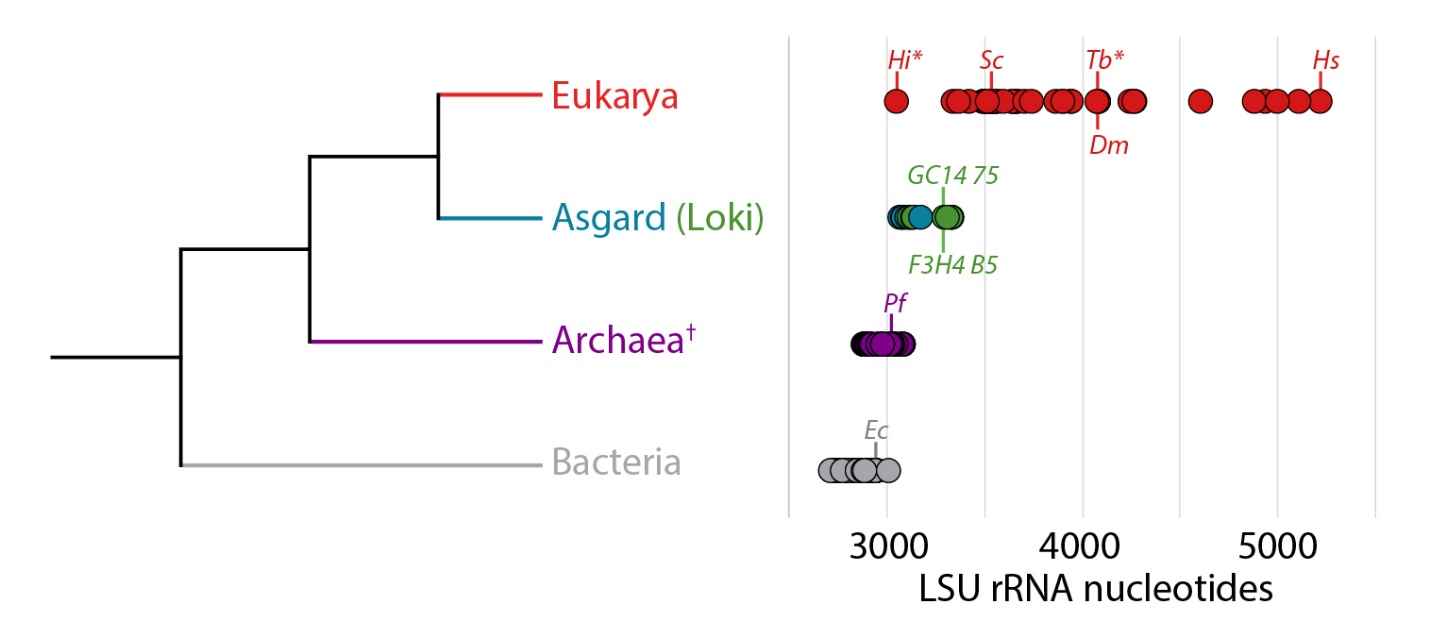
**Figure S1: Length of LSU rRNA increases from Bacteria, to the set of Archaea that excludes Asgard, to Asgard archaea, to Eukarya.** LSU rRNA lengths were obtained from the updated SEREB database. Includes all Asgard species used in the analysis. Abbreviations: *Hi, Hexamita inflata; Tb, Trypanosoma brucei* (parasite)*; Sc, Saccharomyces cerevisiae; Dm, Drosophila melanogaster; Hs, Homo sapiens; F3H4 B5, Lokiarchaeota* B5*; GC14 75, Lokiarchaeota* GC14_75*; Pf, Pyrococcus furiosus; Ec, Escherichia coli*. Eukaryotes are shown in red, Asgard representatives are shown in blue, Lokiarchaeota representatives are in green, archaeal representatives are in purple, bacterial representatives are in gray. *: parasitic representatives of the deeply branching eukaryotic group Excavata. †: excluding the Asgard group.

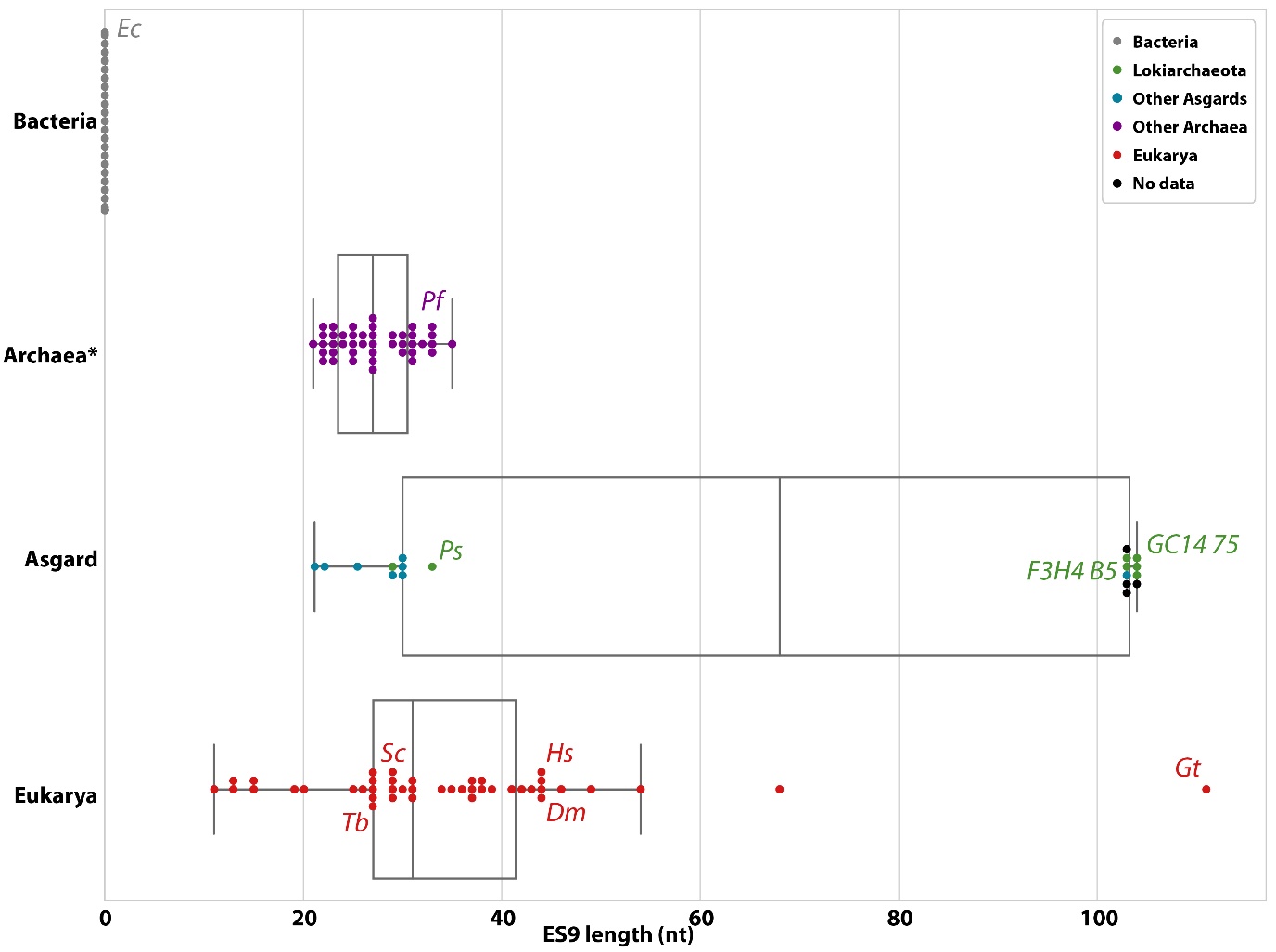

**Figure S2. Length distribution of ES9 rRNA.** The number of nucleotides calculated between alignment positions 1578 and 1621 (*H. sapiens* numbering) of the LSU alignment for each species (**supplementary dataset S1**). The box shows the quartiles of the dataset. Whiskers extend to show the rest of the distribution, except for points that are determined to be outliers using a function of the inter-quartile range. Bacteria sequences are gray, *Lokiarchaeota* sequences are green, other archaeal sequences are purple, eukaryotic sequences are red, sequences from metatranscriptomic contigs (**supplementary dataset S2**) for which there is no species determination are black. Abbreviations: *Ec: Escherichia coli; Pf: Pyrococcus furiosus; Ps: Prometheoarchaeum syntrophicum; F3H4 B5: Lokiarchaeota B5; GC14 75: Lokiarchaeota* *GC14_75; Tb: Trypanosoma brucei; Sc: Saccharomyces cerevisiae; Dm: Drosophila melanogaster; Hs: Homo sapiens*; *Gt: Guillardia theta*. *except for the Asgard superphylum.

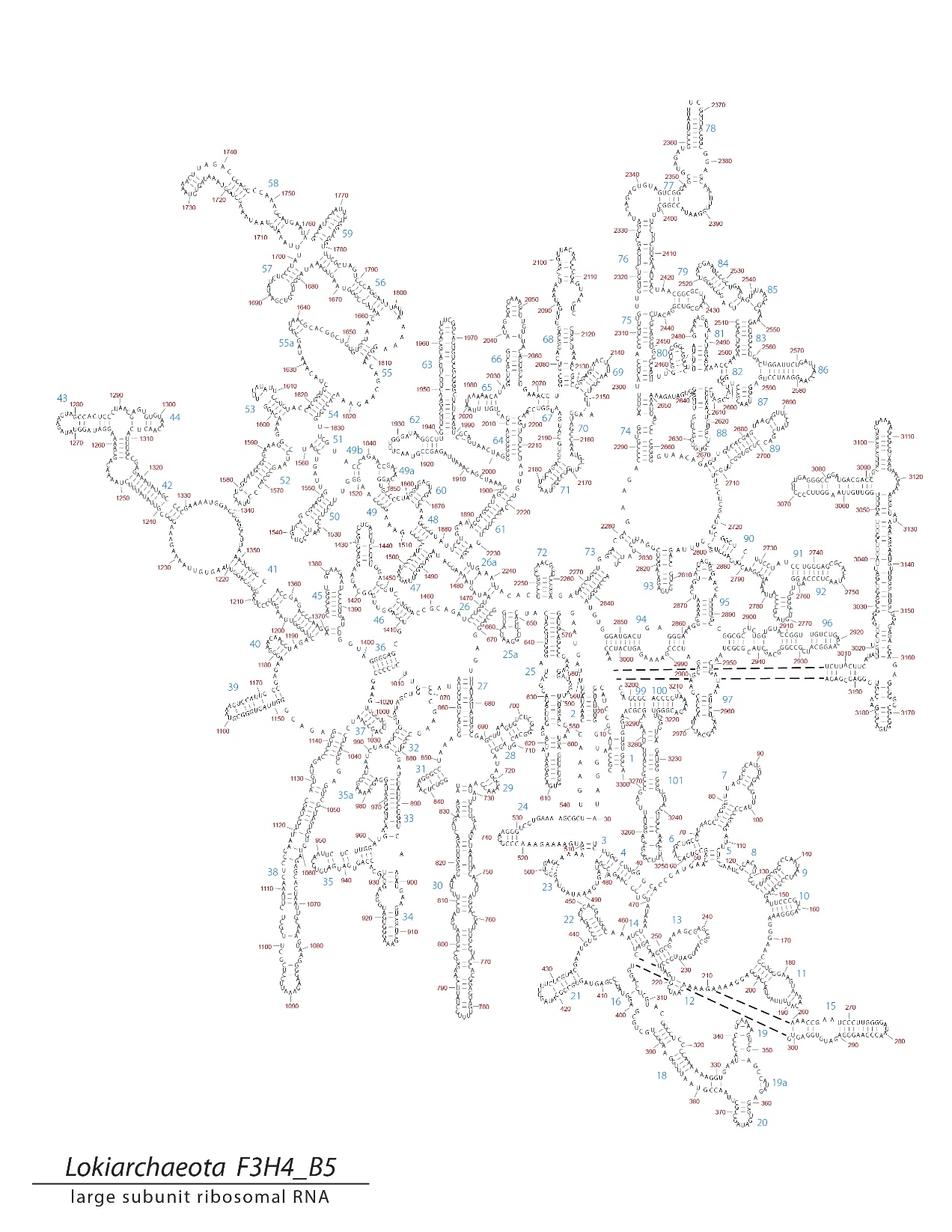

**Figure S3.** **Labeled secondary structure of *Lokiarchaeota* B5.** The numbering scheme of Noller, et al. (1981) and Leffers, et al. (1987) were used to label the helices.

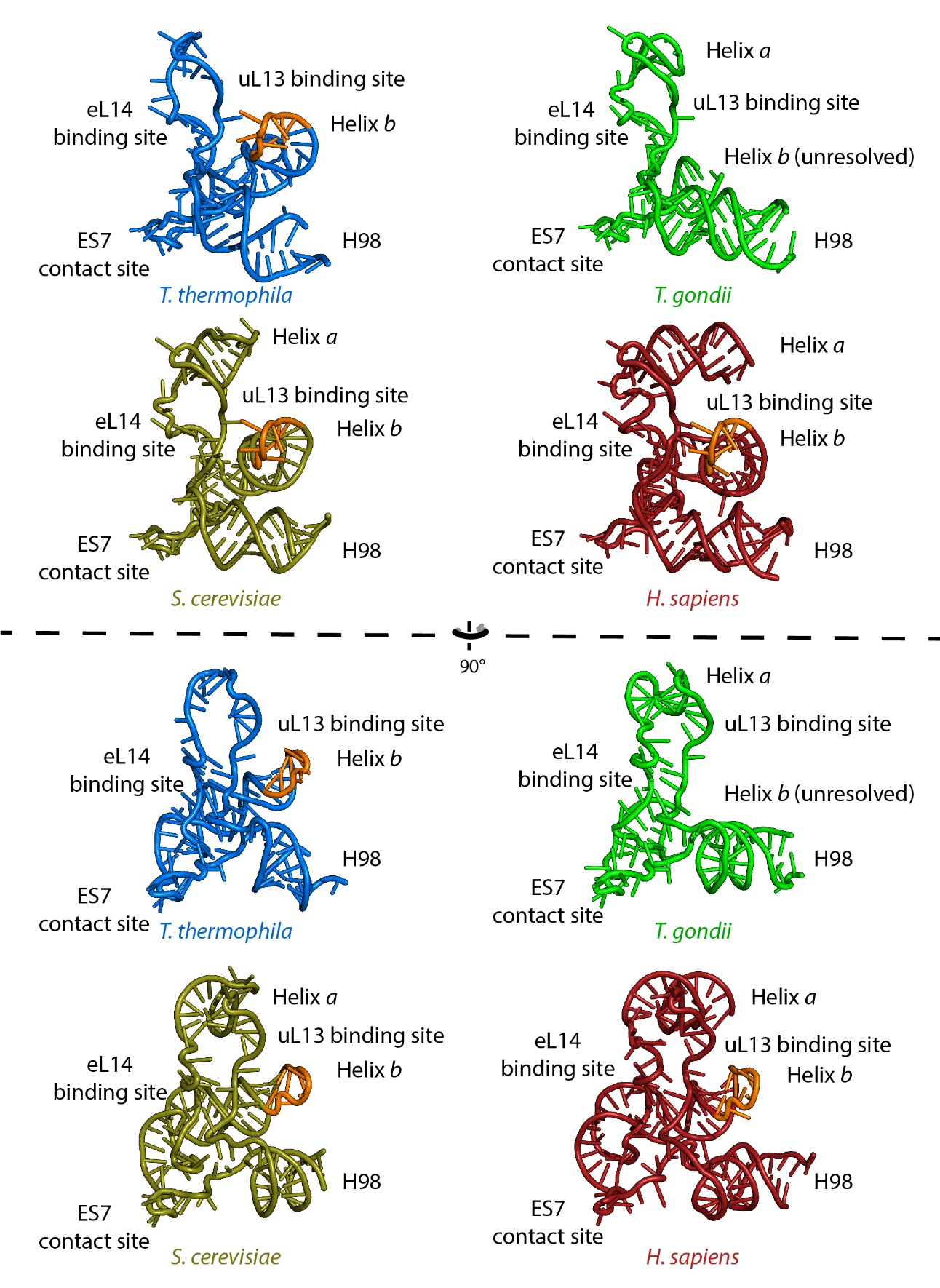

**Figure S4. Structures of ES39 from four eukaryotic species.** *Tetrahymena thermophila* (PDB ID: 4V8P), *Toxoplasma gondii* (PDB ID: 5XXB), *Saccharomyces cerevisiae* (PDB ID: 4V88), *Homo sapiens* (PDB ID: 4UG0). A conserved GNRA-tetraloop at the end of helix *b* is highlighted in orange.
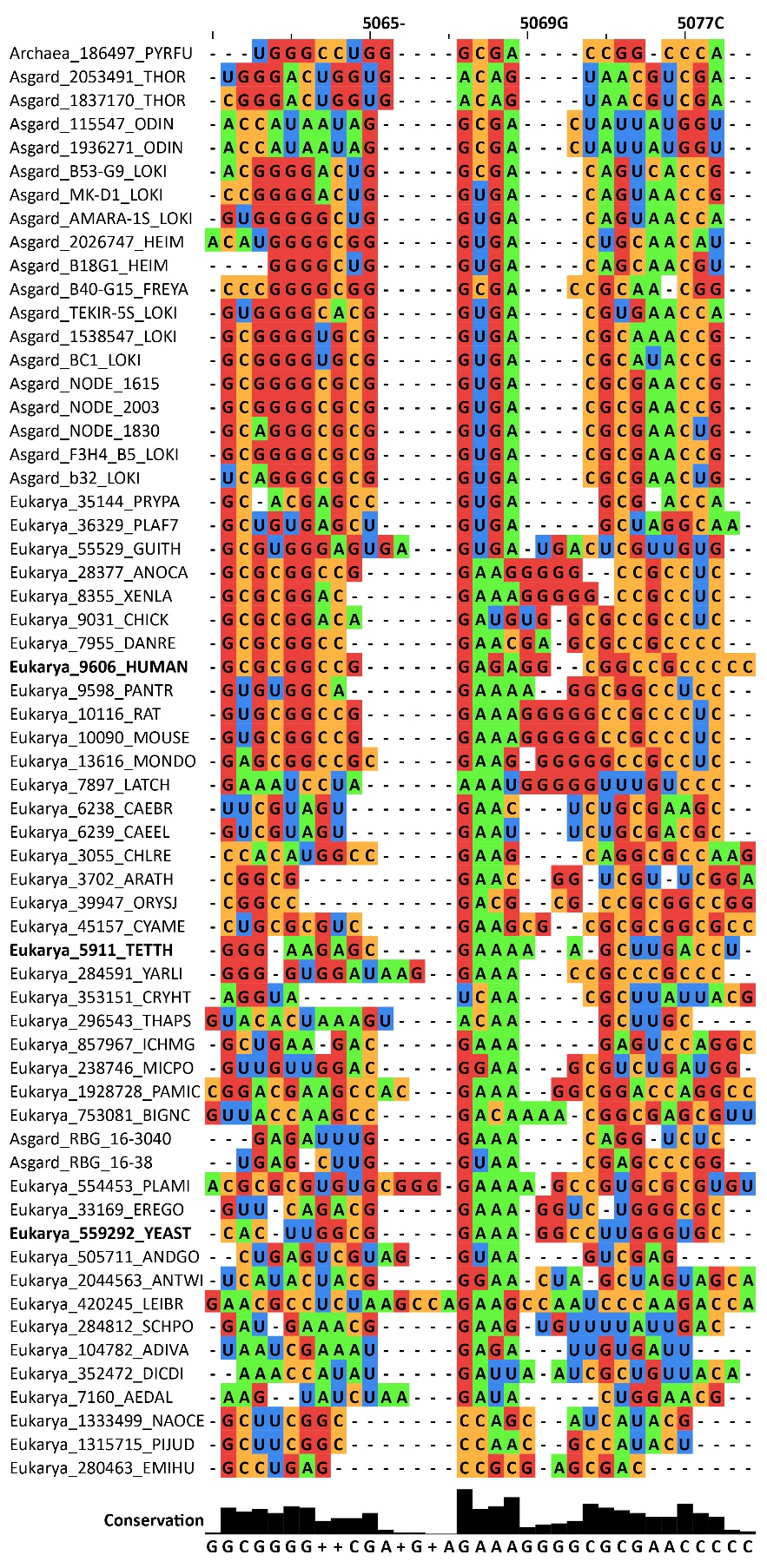

**Figure S5. MSA of ES39 helix *b* from Asgard and eukaryotic species.** Extracted MSA from **supplementary dataset S1**. Alignment index is based on the *H. sapiens* sequence. Sequences from the conserved GNRA-tetraloop shown in figure S4 are bolded. Sequence labels can be found in **supplementary dataset S3**. Figure was generated with Jalview (Waterhouse, et al. 2009).

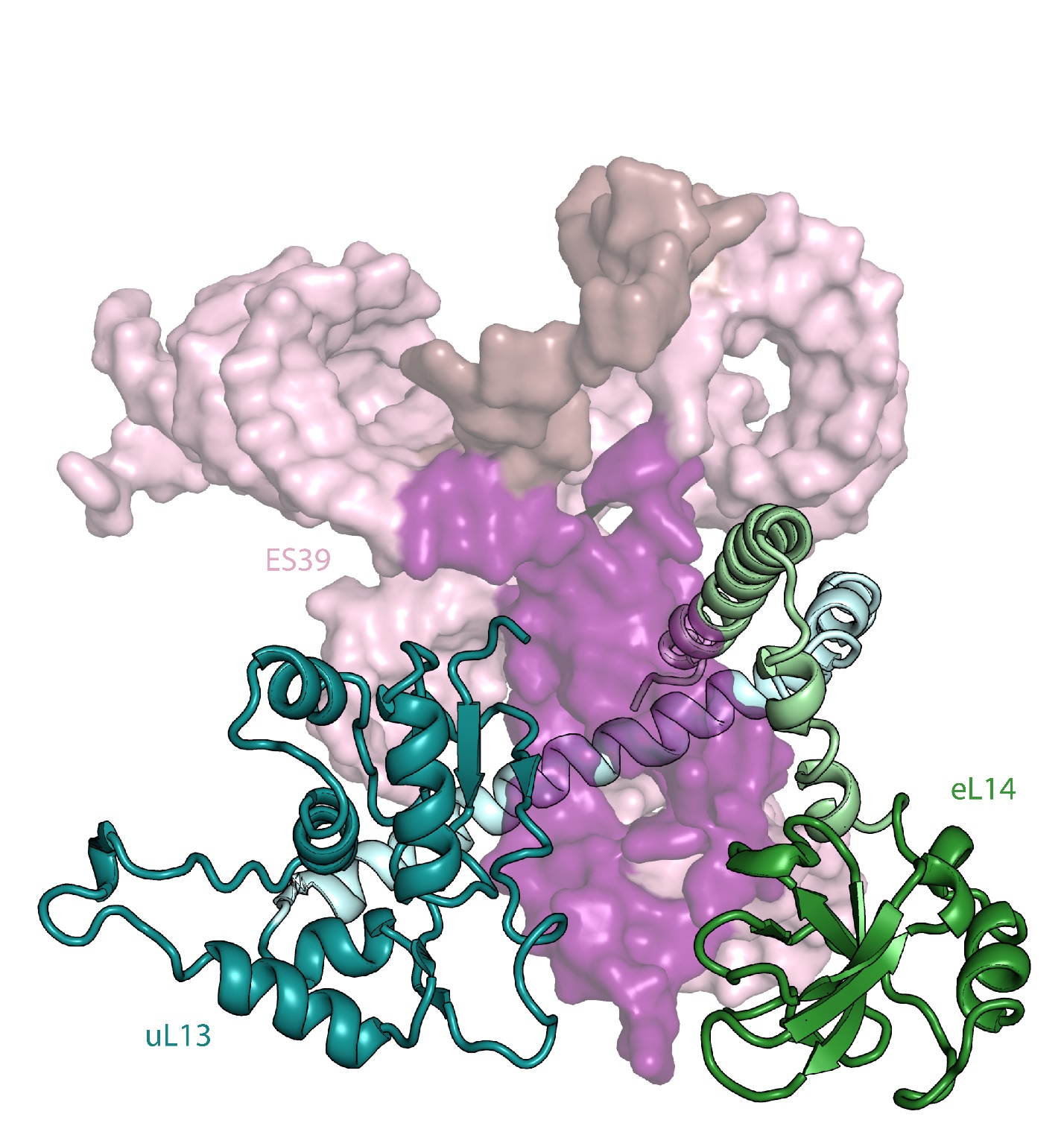

Figure S6. Structure of ES39, uL13, and eL14 from *Homo sapiens* (PDB ID: 4UG0). ES39 structure is shown in surface mode with 40% transparency, in pink are helical regions, in magenta are unpaired regions in contact with rProteins uL13 (blue) and eL14 (green), in brown is unpaired region in contact with ES7. Eukaryotic extensions in rProteins uL13 and eL14 are in light blue and green.

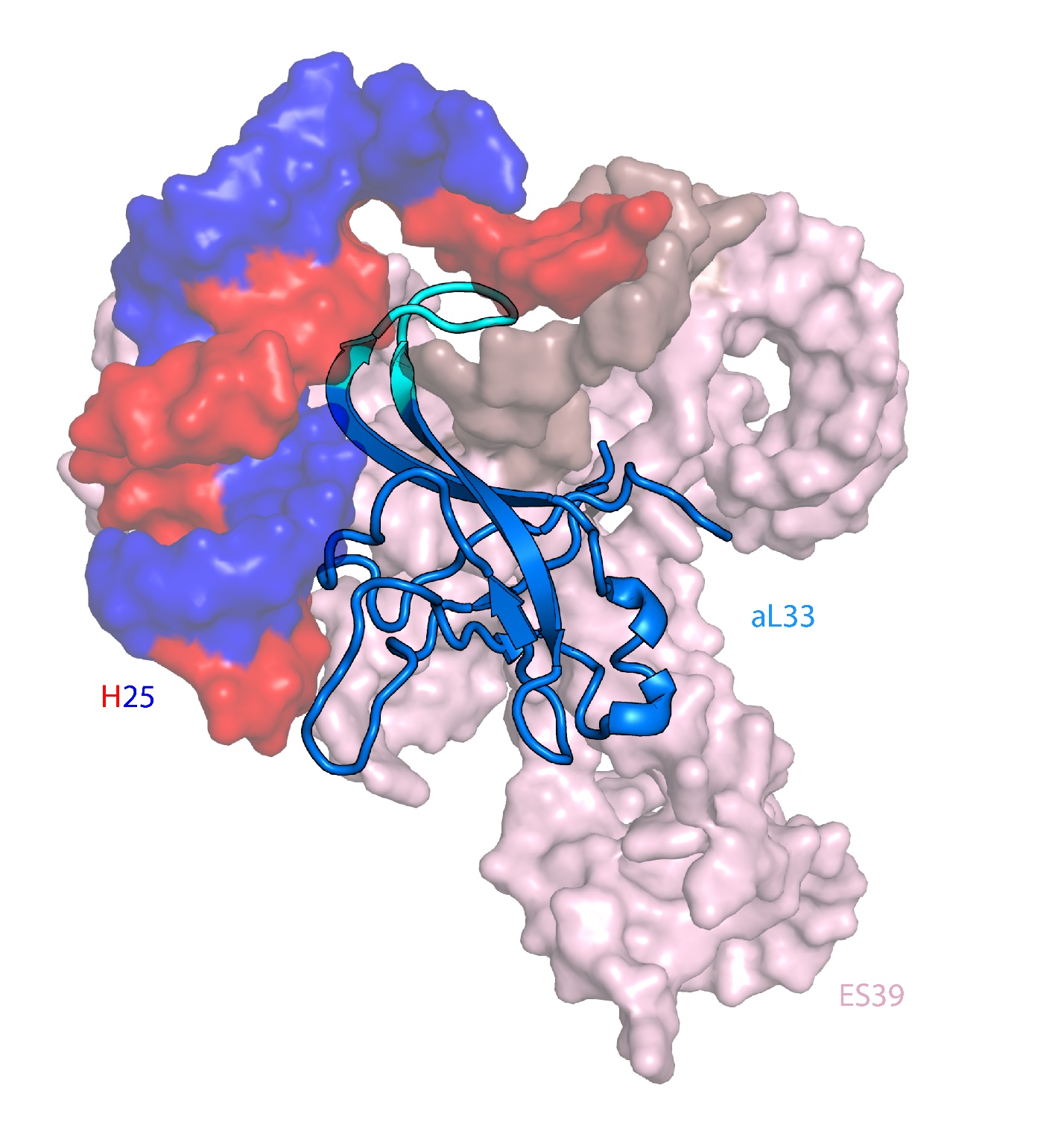

Figure S7. Structure of ES39, H25 (base of ES7), and aL33 from *Homo sapiens* (PDB ID: 4UG0). ES39 and H25 structures are shown in surface mode with 40% transparency. ES39 is in pink and H25 is in blue and red (blue – 5’, red – 3’). ES39 unpaired region in contact with ES7 is shown in brown. rProtein aL33 is shown in marine, in cyan is shown a metazoan specific extension of aL33 (nomenclature as described in Kovacs, et al. (2018), also known as eL33 according to Ban, et al. (2014)).

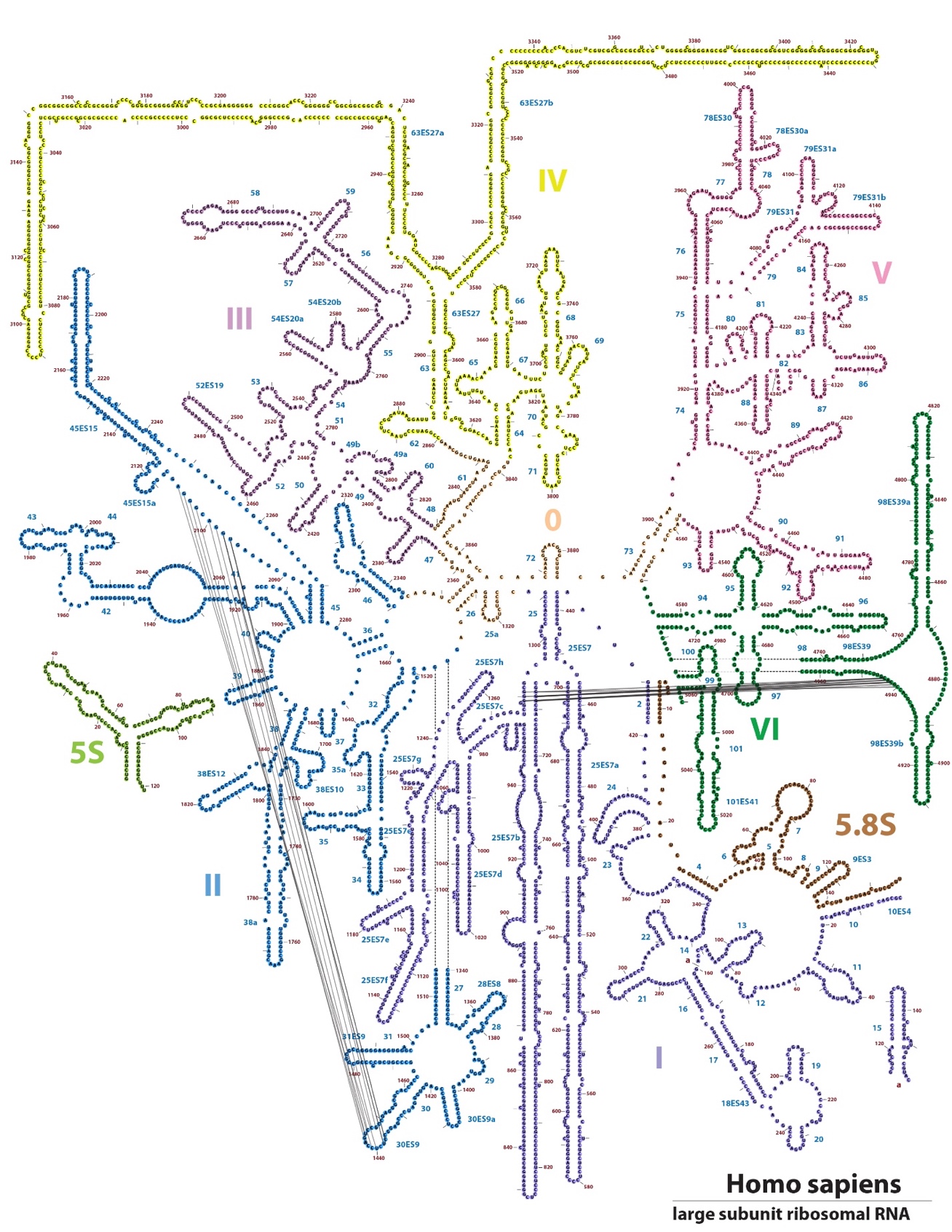

Figure S8. Secondary structure of LSU rRNA from *Homo sapiens*. Interactions between ES39 and ES7, and ES9 and ES15 are indicated with black lines. Figure was generated with RiboVision (Bernier, et al. 2014).

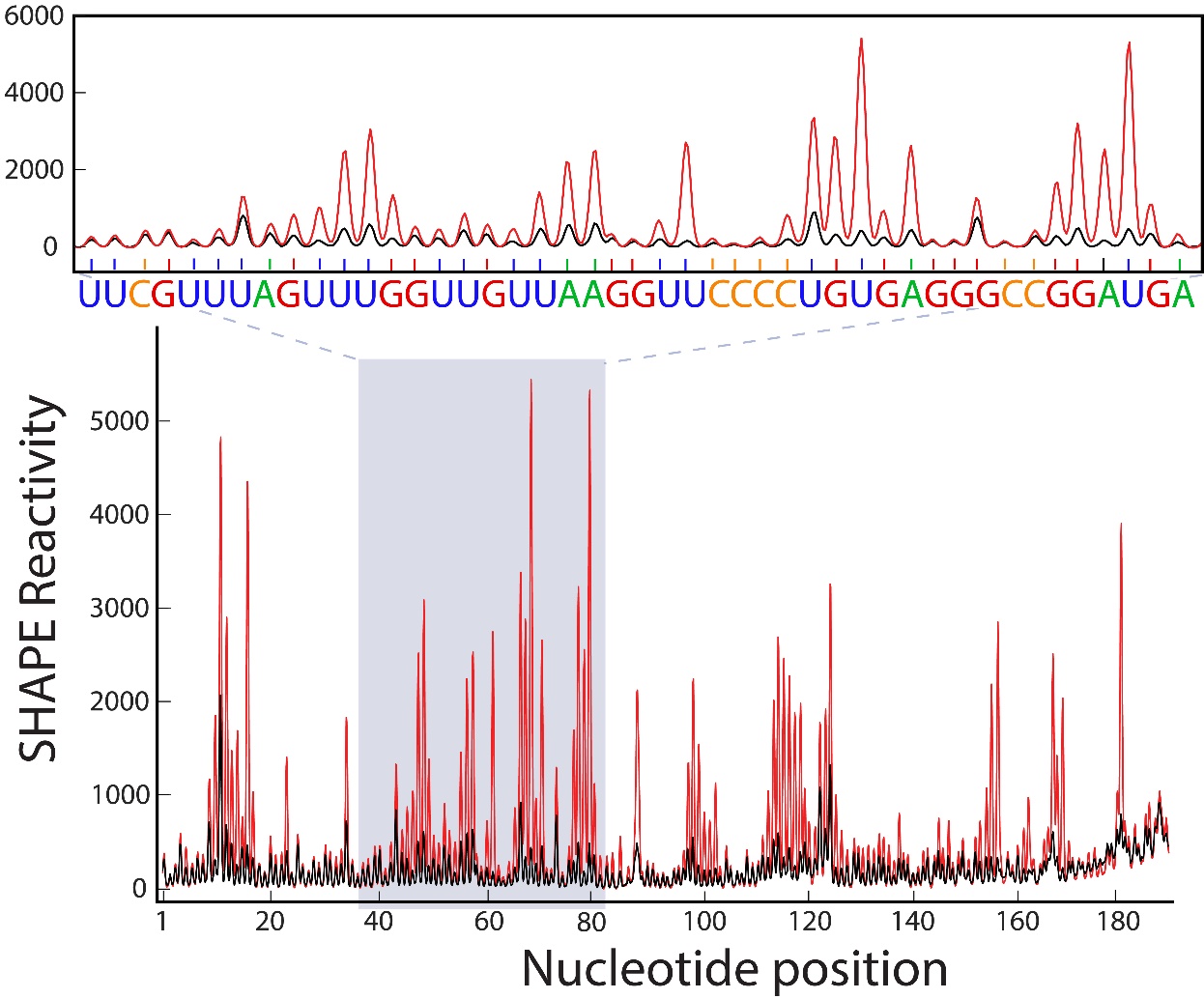

Figure S9. SHAPE reactivity profile for *Lokiarchaeota* ES39. Reactions conducted in the presence and absence of benzoyl cyanide are shown in red and black, respectively. High values show flexible nucleotides, while low values demonstrate structurally stable nucleotides.

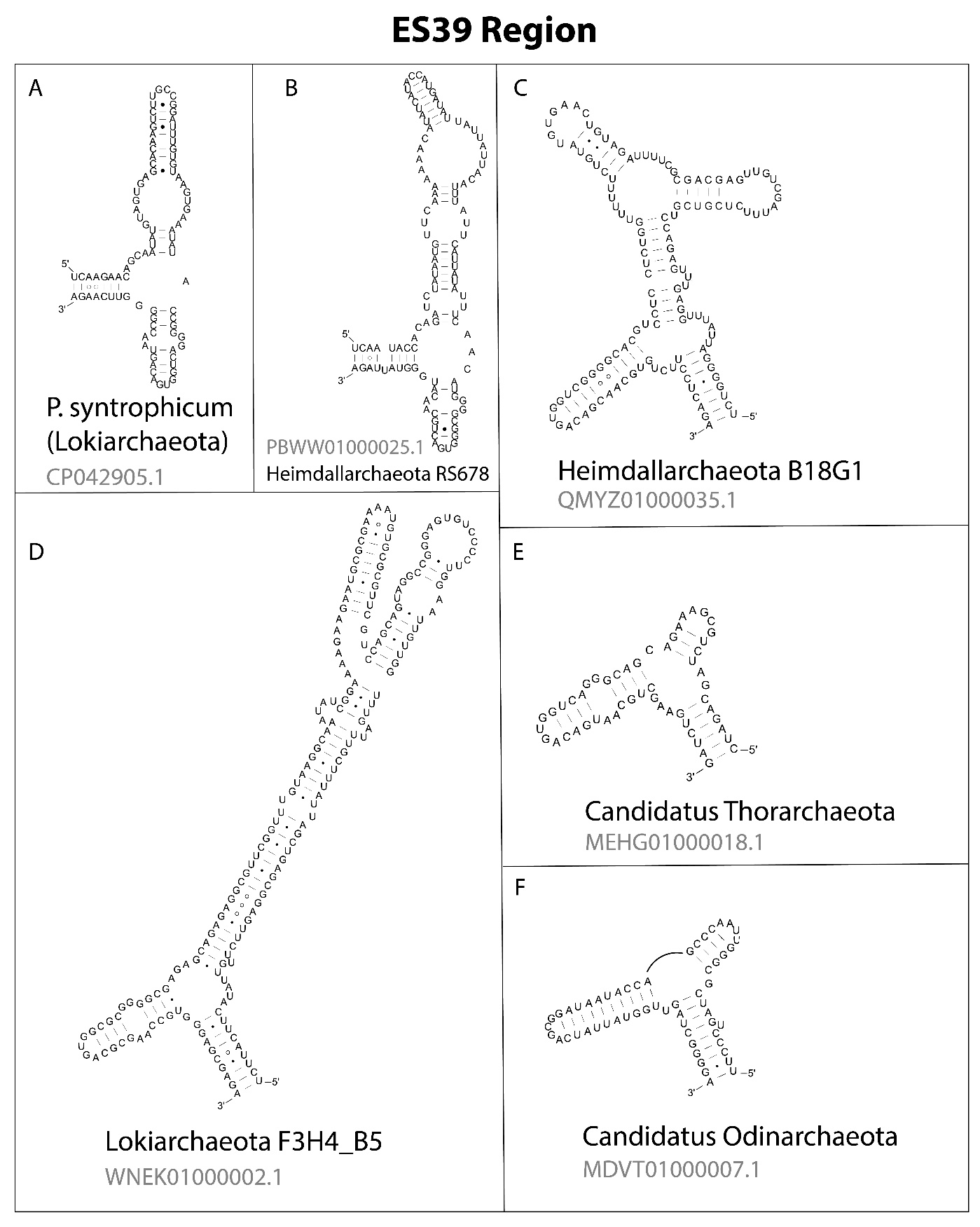

**Figure S10. Secondary structures of ES39 region sequences from Asgard representatives.** (A) and (D): *Lokiarchaeota*; (B) and (C): *Heimdallarchaeota* (complete *Heimdallarchaeota* B18G1 LSU sequence was provided by Brett Baker); (E): *Thorarchaeota*; (F): *Odinarchaeota*. Detailed lengths and genome locations are in **supplementary dataset S2**. Figure was generated with XRNA.

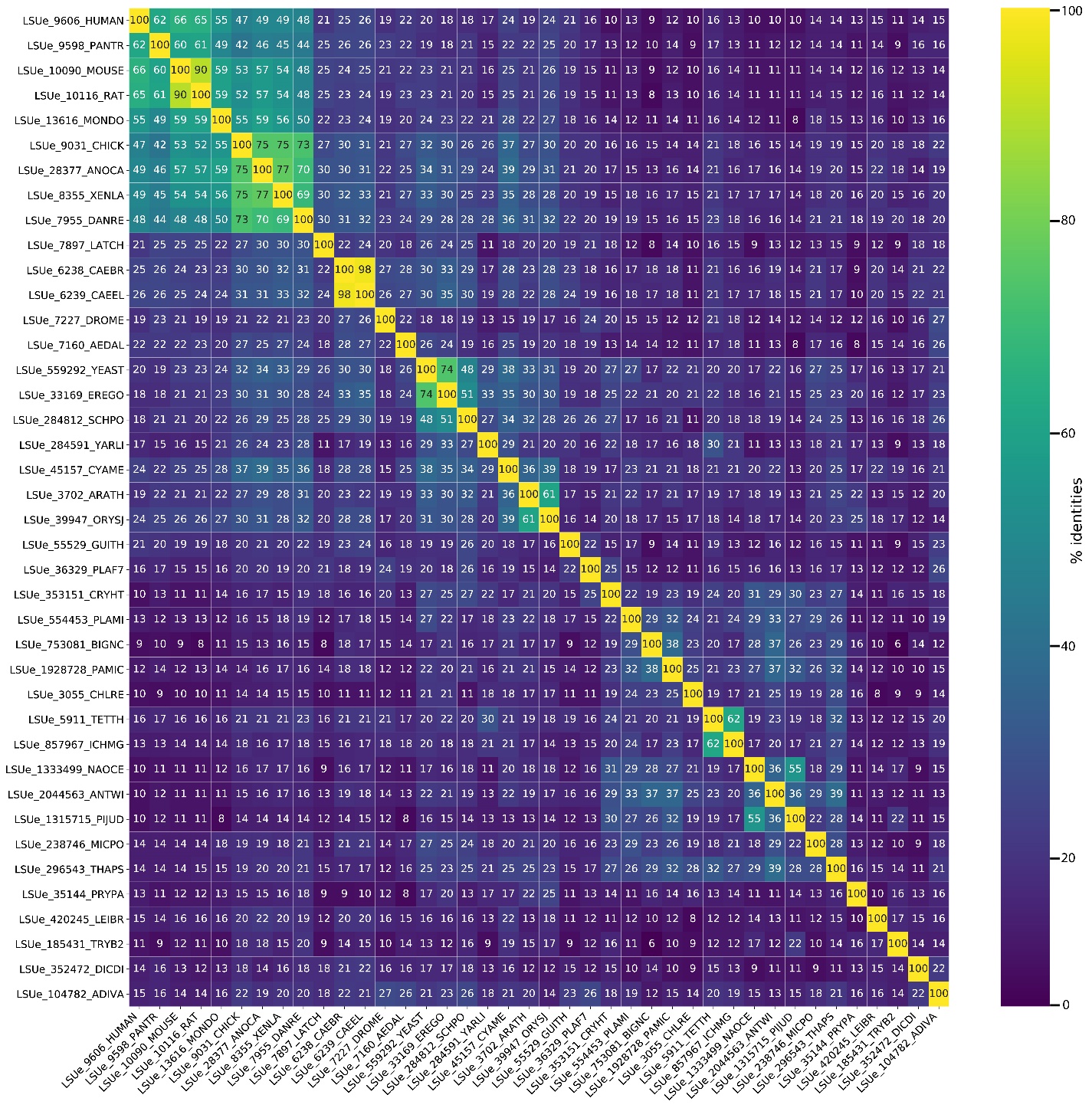

**Figure S11. Percent identity matrix from alignment of eukaryotic-only ES39 sequences.** Sequence names are indicated on both axis and full phylogeny can be found in **supplementary dataset S3**. Alignment is extracted from **supplementary dataset S1** between positions 8205-8840. The percent identities are mirrored on the two sides of the diagonal. The diagonal is set to 100 percent identity. Brighter colors indicate higher similarity between two aligned sequences and darker colors indicate lower similarity.

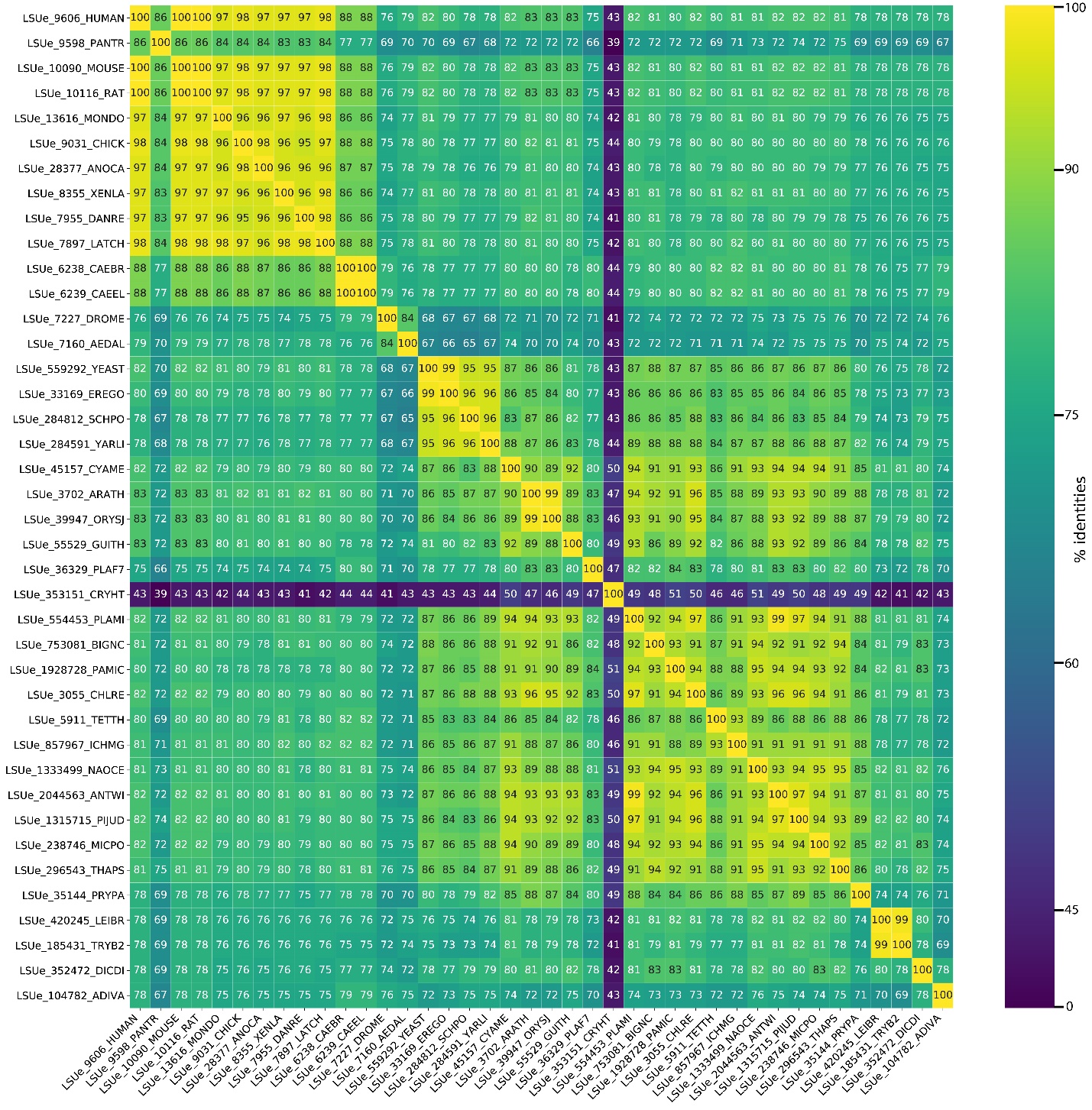

**Figure S12. Percent identity matrix from alignment of eukaryotic-only central protuberance sequences.** Sequence names are indicated on both axis and full phylogeny can be found in **supplementary dataset S3**. Alignment is extracted from **supplementary dataset S1** between positions 7512-7693. The percent identities are mirrored on the two sides of the diagonal. The diagonal is set to 100 percent identity. Brighter colors indicate higher similarity between two aligned sequences and darker colors indicate lower similarity.

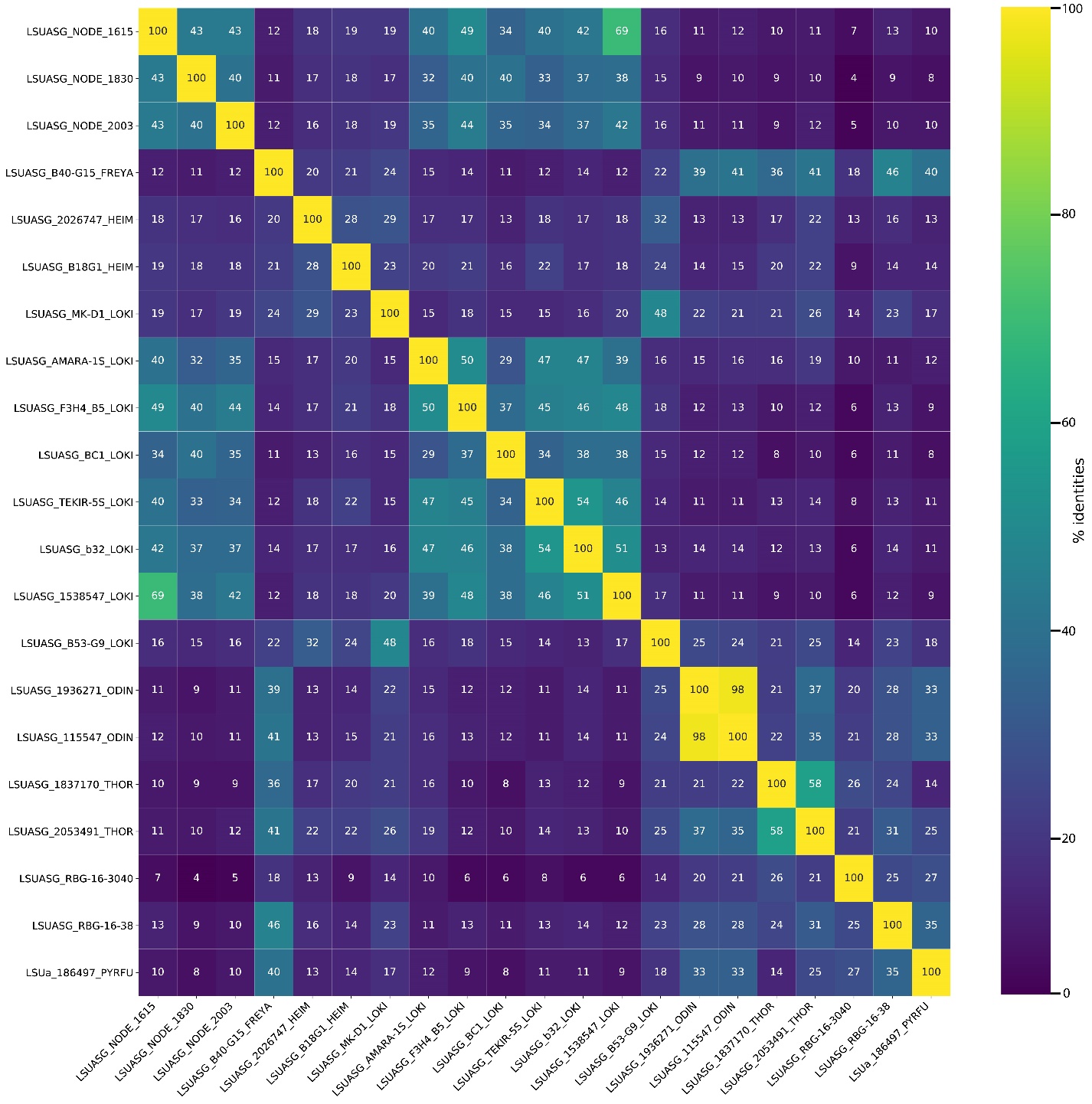

**Figure S13. Percent identity matrix from alignment of Asgard-only ES39 sequences.** Sequence names are indicated on both axis and full phylogeny can be found in **supplementary dataset S3**. Alignment is extracted from **supplementary dataset S1** between positions 8205-8840. The percent identities are mirrored on the two sides of the diagonal. The diagonal is set to 100 percent identity. Brighter colors indicate higher similarity between two aligned sequences and darker colors indicate lower similarity.

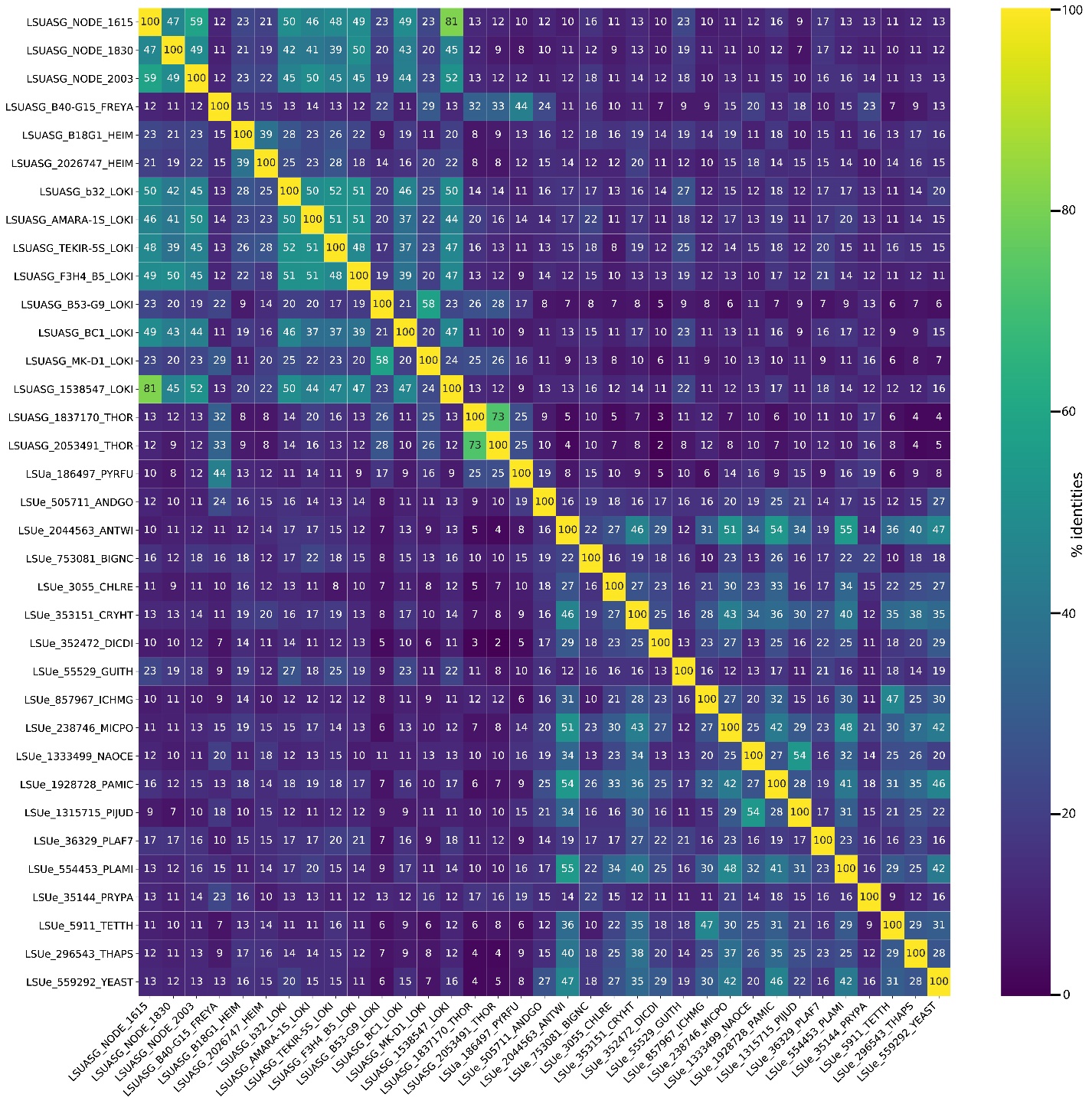

**Figure S14. Percent identity matrix from Clustal Omega alignment of Asgard and eukaryotic ES39 sequences.** Sequence names are indicated on both axis and full phylogeny can be found in **supplementary dataset S3**. Alignment is extracted from **supplementary dataset S1** between positions 8205-8840. The percent identities are mirrored on the two sides of the diagonal. The diagonal is set to 100 percent identity. Brighter colors indicate higher similarity between two aligned sequences and darker colors indicate lower similarity.

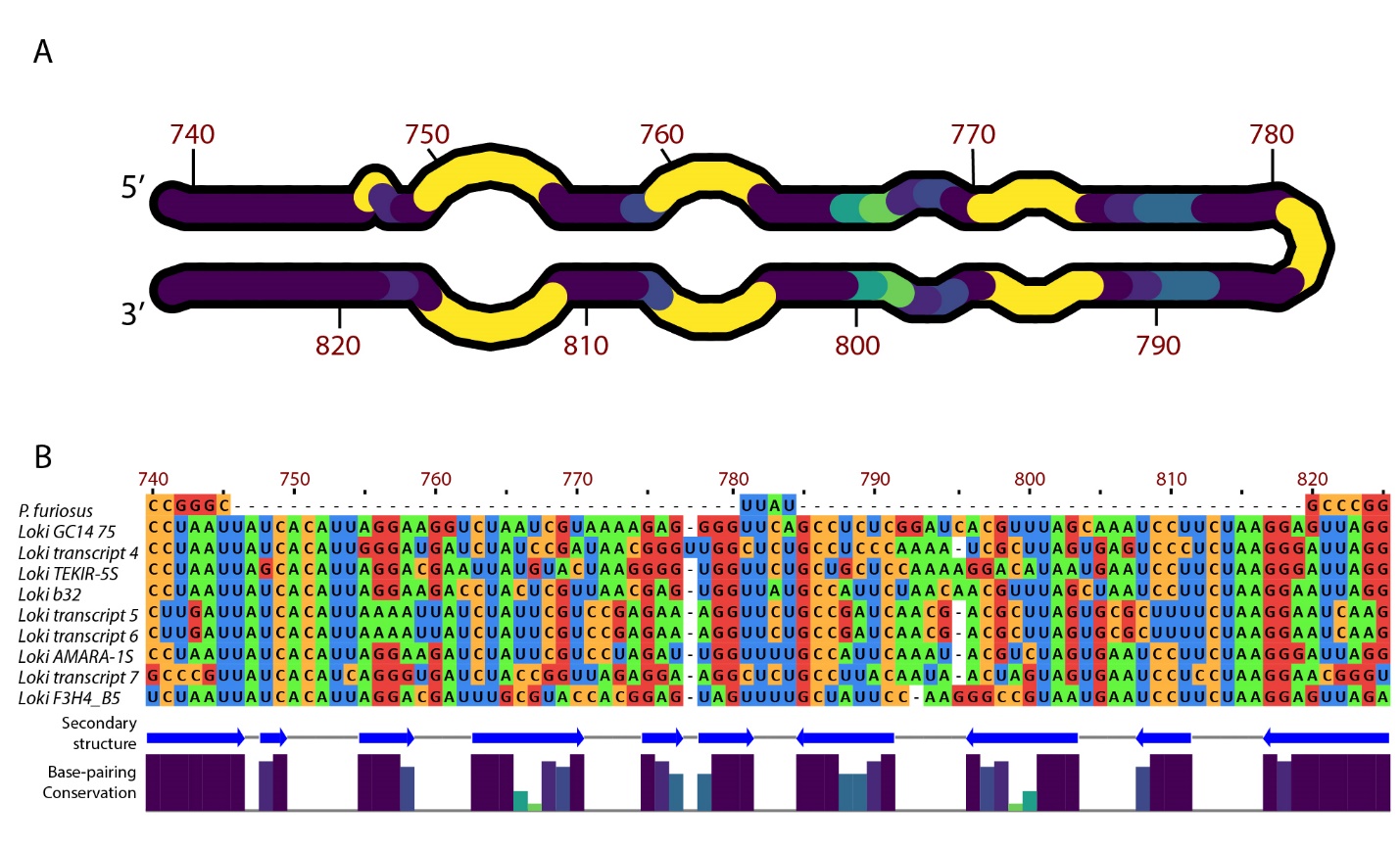

**Figure S15. Secondary structure and MSA of *Lokiarchaeota* B5 ES9.** (A) Secondary structure of ES9, nucleotide numbers for *Lokiarchaeota* B5 are labeled in red. Positions with high base-pairing conservation are shown in dark violet, while low conservation and unpaired regions are in gold. The region around positions 770 and 800 is likely unpaired in *Lokiarchaeota* B5 but paired in other *Lokiarchaeota* species due to its high base-pairing conservation*.* (B) MSA of nine *Lokiarchaeota* species and *P. furiosus*. Full sequence names and sequencing project identifiers are available in **supplementary dataset S2**. *Lokiarchaeota* B5 numbering was used. The model of secondary structure is indicated with blue arrows bellow the alignment. Base-pairing is shown with a bar graph bellow the secondary structure with the same colors as in panel A. Figure was generated with Jalview (Waterhouse, et al. 2009).

Captions to Datasets

Dataset S1. (separate file)

Multiple sequence alignment of LSU sequences from the SEREB database (Bernier, et al. 2018) expanded with sequences from Asgard representatives and metatranscriptomic assembly contigs.

Dataset S2. (separate file)

Table with sequence and gene identifiers for additional archaeal sequences, Asgard representatives, eukaryotic protists, and metatranscriptomic assembly contigs used in Figure 2 and Supp. Fig. S1.

Dataset S3. (separate file)

Table with all species used in the analysis and their alignment names in Dataset S1. The table includes taxonomic ranks of Domain, Phylum and Class, as well as LSU and ES’s nucleotide lengths.

Dataset S4. (separate file)

File containing statistic report from assembly run of SRR5992925 generated with rnaspades (Bankevich, et al. 2012).

Dataset S5. (separate file)

File containing statistic report from assembly run of SRR5992925 generated with metaspades (Bankevich, et al. 2012).

.
