## Supplemental Data S1 for "Supersized ribosomal RNA expansion segments in Asgard archaea"

```
>LSUb_575564_ACISP/1-2876
-----------------------------------------AGUGCAUGU-GGUGGAUG-C-CUUGGC-AGU
CAG---AGGCGAUGAAAGACGUGA--UAGCCUGCGAAAAGCUCCGGGGAGGCGGCAAAUAUCCU-----UUG
AUC-CGG-AGAUGUCUGAAUGGGGGAAC--------CC--------AC----CUAC-U------------UU
A-A-----------------G----G----U---AGG----U-A-UUGC-AAC-----AUGA----------
-----------A-----------UACA----------------------------------------UA-GU
G-U--------------------UG-C-AA-GGCGAACGAGGGGAAGUGAAACAUCUCAGUACCCU--UAGG
AAAAGAAAUCAA--UUGAGAUUCCCUCA--GU-AG-CGGCGAGCGAACGGGGA-U-CA-GCCCA--------
------------------------------------------------------------------------
----------UUA-------A----G-----------------------UU--AUG-----------UG---
U-------GUUUU-AGUGG------AAC-GC-UCUG------------------------------------
-------GGA-AG-UGCGA-------ACGUAGAGGGUGAUAUUCCCGUA---------CACGAAA----GGG
C------A--CACAU-----------AA----------UG-AU------GACGAGUAGGGCGAGGCACGUGA
AACCUUGUCUGAA-UAUGGGGGG-ACC--AUCCUCCAAGGCUAAAUA--CUC-CUGAC-UGACCGAUAGU-G
AAC-C-AGUACCGUGAGGGAAAG-GCGAAA-AGAAC-CCCUGUGAGGGG-A-GUGAAAUAGAUCCUGAAACC
G-CAUGCAUACAA-GCA-GU-GGGA---GCAC-C--------------------------------------
------------------------------------------------------------------------
------------------------------------------------------------------------
------------------------------------------------------------------------
------------------------------------------------------------------------
------------------------------------------------------------------------
------------------------------------------------------------------------
------------------------------------------------------------------------
------------------------------------------------------------------------
-------------------------------------------------------------UU--C-GU---
------------------------------------------------------------------------
------------------------------------------------------------------------
------------------------------------------------------------------------
------------------------------------------------------------------------
------------------------------------------------------------------------
------------------------------------------------------------------------
------------------------------------------------------------------------
------------------------------------------------------------------------
------------------------------------------------------------------------
------------------------------------------------------------------------
------------------------------------------------------------------------
------------------------------------------------------------------------
------------------------------------------------GG--U---GUGACUGCGUACCUUU
UGUAUAAUGGGUCAGCGACUUAU-A-UUCAGUAGCGA-GGUUAACC-------------GUAUA--------
---G--GGGA--GCCGUAGAGAAAU-C---------------------------------------------
------------------------------------------------------------------------
------------------------------------------------------------------------
---------------------------------------------------------------------GAG
UCU------------------------------------UAAUA-------------------------GG-
G-C-GU----------------------UUAGUUGCUGGGUAUA-GACCCGAAACC-AGGCGAUCUAUCCAU
```

```
GAGCAGGUUGAAGGUUGG-GUAACA-CUAACUGGAGGACCGA--ACCCACUG-UCGUUGAAAGCCA-GGGG
AUGACUUGUGGAUAGGGGUGAAAGGCUAAUCAAGCCUGG-UGAUAGCUGGUUCUCCCGAAA-GCUAUUUAG
GUAGCGCCUCGG-ACG--------------------------------------------------------
----------------------AA-----UA-CCAUAGGGGUAGAGC-ACUGUUU-CGG-CUA--GGGGGU
C---AUCCC--GACUUACCAAACCGAUGCAAACUCCGAAUACCUAU--------------------------
------------------------------------------------------------------------
------------------GAGU-A-CU--------AUCCGGG-AGACAGA-C-UGCG-G-GUGCUAACGUCC
GU-A-GUCAAGAG-GAAAAC---AAUCCAGACCGCCA-GCUAAGGCCCAAAAU--CA-UAG-UU-AA---G
-U-GGG-------AAACGAUGUGGGAAGGCAUAGACAGCUAGGAGGUUGGCUUAGAAGCAGCCA-CCCUUUA
AAGAAAGCGUAAUAGCUCACUAGUC-----GAGUCGGCCUGCGCGGAAGAUGUAAC-GGGGCUAA-AACUAU
G----------UGCCGAA-GCUGCGGAUUUGACA--U------------------------------------
------------------------------------------------------------------------
------------------------------------------------------------------------
---------------------------------UAGU-CAAG-UGGUAGGGGAGC-GUUCUGU---------
-----------AAG-CCGAU-GAAGGUG--U-AUUGAGAAGU--A-UGCUGGAGGU-A-UCA-GAAGUGCGA
AUGC-UGACGU-GAGUA-ACGACA---AAAC-GGGUGA--A---------------AAACCC-GUU-CGCCG
A-AAGACC-AAGGGUUCC-AGUCCAACG-UUAAUCGGGGCUGGGUGAGUCGAC-CC-CUAAGGCGAGGCCG-
A-A-AGGC-----GUA--------------------------------------------------------
----------------------------------------GUC-GA-U--GGGA-AAUUGGUU-AAUA-UUC
CAAUA----------------------------------CUUCU----GUGU--------A-AU--------
------------------------------------------------------------------------
----------------------------------------------------------GCGAU-GAGAGGAC
GGAGAA--------------GGUUAA-GUCAGCCUG-GC-G----UUGGU------------UGUCCAGGU-
GGAAGG-AUG-UA-----G-G-------------------UA-----UG------------------UA-UC
U-UAGG--------------------CAAAU-CCGGGGUACU------CU-AUACUG----AGA-UC-C-G-
AUA--GC---------------------------------------------------------------AA
----GCUGUAC-UUGUACAGCGAAG-U-GGC-U-GAUACCAUGCUUC-C-AGGAAA-AGUCU-CUAAGCUU-
-------------------------------------------------------CAGUUAC-AC-AGGAAU
CGUACCCGAAACCGAC------------------------------ACAGGUG--GUCAGGUC-GAGUAGAC
CAAGGC-G-CU-UGAG-AGAACUCU-G-CUGAAGGAACUAGGC-AAAAUGGUACCGUAACUUCGG-GAGAAG
GUACG-CUGU-U-GU--UGGUGAUG----GAA-----------CUUGC------------------------
------------------------------------------------------------------------
------------------------------------------------------------------------
------------------------------------------------------------------------
------------------------------------------------------------------------
------------------------------------------------------------------------
------------------------------------------------------------------------
------------------------------------------------------------------------
------------------------------------------------------------------------
------------------------------------------------------------------------
------------------------------------------------------------------------
------------------------------------------------------------------------
------------------------------------------------------------------------
------------------------------------------------------------------------
--------------------------------------------------------UUCC-UGA--GCUGAU
```

```
GAC--AGCCGCAGAAACCAGGCCGCUGCAA-----CUGUUUAUUAAAAACAUAGCACUCUGCAAACACG---
-AA-A-------------GUGGACGU-AUAGGGUGUGA-UGCCUGCCCGGUGCUG-G-AAGG-UUAAUUGAU
GGGGUUA-GC-----GUAA-----GCGAAGCUCUUGAUCGAAGCCC--CAGUAAAC-GGCGGCCGUAACUAU
AACGGUCCUAAGGUAGCGAAAUUCCUUGUCGGG-UAAGUUCCGACCUGCACGAAUGGCAUAAUGAUG--GCG
GCGCUGUCUCCAGC----AGA-GGCUCAGUGAAAUC--GAA---AUCGCUGUGAAGAUGCAGUGUACCCGCG
GCUAGACGGAAAGA-CCCCGUGAACC-UUUACU--GCAGCUU-GACACUGAACUUUGACCUUACUUGUGUAG
GAU-AGGUGGGAGGCUUUG----------AAGUUGG-AAC----------------------------GCUA
-------------------------------------GU-UCCAAUGGA---G--CCGU-CCUUGAAAUA-C
CACC-CUGGUAAUGUUGAGGUUCUAACUCUGU--CCCG-U-UA-U-C-------------------------
------------------------------------------------------------------------
------------------------------------------------------------------------
------------------------------------------------------------------------
------------------------------------------------------------------------
------------------------------------------------------CG----GG----------
------------------------------------------------ACGA-----GGACC-GUGUCUGGU
GGGUAGUUUGACUGGGGCGGU-CUCCUCCUA-AAGAGUAAC-GGAGGAGUACG-AAGGUGCGCUCAGCGUGG
UCGGAAAUCACGCGUA-GAGUA-UAAAGGCAA-AAGCGCGCUUAACU-GCGA--------------------
--------------GACCCA---CAAGUCGAGCAGGUACGAAA-GUAGGUCUUAGUGAUCCGGU--------
---GG-U-----UCUGU-AUG--------------------GA-AGGGCCAUCGCUCAACGG-AUAAAAGGU
ACUCUGGGG-AUAACAGGCUGAUACCGCCCAAGAGUUCAUAUCGACGGCGGUGUUUGGCACCUCGAUGUCGG
CUCAUCUCAUCCUGGGGCUGAA-GCAGGUCCCAAGGGUAUGGCUGUUCGCCAUUUAAAGAGGUACGCGAGCU
GGGUUUAGAACGUCGUGAGACAGUUCGGUCCCUAUCUACC--------------------G-UGG-------
-------------------------------GCGC--UGGAAAUUUGAGAGG-AUC-UGCUCCUAGUACGAG
AGGAC-CAGAGUGGACGAACCUC-UGGU-G-UACCGGUUGUGACGCCAGUCGCAU-C-GCCGGGUAGCUAUG
UUCGGAAGGGAUAACCGCUGAAAGCA-UC--UAAGCGGGAAGCCUACCUCAA-G-AUA--AGAUUUCC-CU-
--AGGAA-----------------------------------------------------------------
--------------------------UUU-------------------------------------------
------------------------------------------------------------------------
------------------------------------------------------------------------
------------------------------------------------------------------------
------------------------------------------------------------------------
------------------------------------------------------------------------
------------------------------------------------------------------------
------------------------------------------------AUUCCUCU--AA-AGA------GC
CGUUCGAGACUAGGACGU-UGA-UAGGUU-GGA-UGUGGAAG-CAU-A----------------GU------
----GA-----------UAUGUGAAGCUG--A-CCAA-UACUAAUU---GCUCGUGAG-GCUUGACU-----
------------------------------------------------------------------------
------------------------------------------------------------------------
------------------------------------------------------------------------
-------------------------------------------------
>LSUb_208963_PSEAB/1-2891
---------------------G-GUCAAGUG----AA--GAAGCGCAUAC-GGUGGAUG-C-CUUGGC-AGU
CAG---AGGCGAUGAAAGACGUGG--UAGCCUGCGAAAAGCUUCGGGGAGUCGGCAAACAGACU-----UUG
AUC-CGG-AGAUCUCUGAAUGGGGAAC---------CC--------AC----CUAG-G------------AU
A-A-----------------C----C----U---AGG----U-A-UCUU-GUA-----CUGA----------
```

```
-----------A-----------UCCA----------------------------------------UA-GG
U-G--------------------CA-A-GA-GGCGAACCAGGGGAACUGAAACAUCUAAGUACCCU--GAGG
AAAAGAAAUCAA--CCGAGAUUCCCUUA--GU-AG-UGGCGAGCGAACGGGGA-U-UA-GCCCU--------
------------------------------------------------------------------------
----------UAA-------G----CU--------------------------UCA-----------UU---
G-------AUUUU-AGCGG------AAC-GC-UCUG------------------------------------
-------GAA-AG-UGCGG-------CCAUAGUGGGUGAUAGCCCCGUA---------CGCGAAA----GGA
U------C--UUUGA-----------AG----------UG-AA------AUCGAGUAGGACGGAGCACGAGA
AACUUUGUCUGAA-CAUGGGGGG-ACC--AUCCUCCAAGGCUAAAUA--CUA-CUGAC-UGACCGAUAGU-G
AAC-C-AGUACCGUGAGGGAAAG-GCGAAA-AGAAC-CCCGGAGAGGGG-A-GUGAAAUAGAACCUGAAACC
G-UAUGCGUACAA-GCA-GU-GGGA---GCCU-A--------------------------------------
------------------------------------------------------------------------
------------------------------------------------------------------------
------------------------------------------------------------------------
------------------------------------------------------------------------
------------------------------------------------------------------------
------------------------------------------------------------------------
------------------------------------------------------------------------
------------------------------------------------------------------------
-------------------------------------------------------------CU--U-GU---
------------------------------------------------------------------------
------------------------------------------------------------------------
------------------------------------------------------------------------
------------------------------------------------------------------------
------------------------------------------------------------------------
------------------------------------------------------------------------
------------------------------------------------------------------------
------------------------------------------------------------------------
------------------------------------------------------------------------
------------------------------------------------------------------------
------------------------------------------------------------------------
------------------------------------------------------------------------
------------------------------------------------UA--G---GUGACUGCGUACCUUU
UGUAUAAUGGGUCAGCGACUUAU-A-UUCAGUGGCAA-GCUUAACC--G-----------UAUA--------
---G--GGUA--GGCGUAGCGAAAG-C---------------------------------------------
------------------------------------------------------------------------
------------------------------------------------------------------------
---------------------------------------------------------------------GAG
UCU------------------------------------UAAUA-------------------------GG-
G-C-GU----------------------UUAGUCGCUGGGUAUA-GACCCGAAACC-GGGCGAUCUAUCCAU
GAGCAGGUUGAAGGUUAG-GUAACA-CUGACUGGAGGACCGA--ACCCACUC-CCGUUGAAAAGGUA-GGGG
AUGACUUGUGGAUCGGAGUGAAAGGCUAAUCAAGCUCGG-AGAUAGCUGGUUCUCCUCGAAA-GCUAUUUAG
GUAGCGCCUCAU-GUA--------------------------------------------------------
-----------------------U-----CA-CUCUGGGGGGUAGAGC-ACUGUUU-CGG-CUA--GGGGGU
C---AUCCC--GACUUACCAAACCGAUGCAAACUCCGAAUACCCAG--------------------------
```

```
------------------------------------------------------------------------
------------------AAGUGCCGA--------GCAUGGG-AGACACA-C-GGCG-G-GUGCUAACGUCC
GU-C-GUGAAAAG-GGAAAC---AACCCAGACCGCCA-GCUAAGGUCCCAAAGU--UG-UGG-UU-AA---G
-U-GGU-------AAACGAUGUGGGAAGGCUUAGACAGCUAGGAGGUUGGCUUAGAAGCAGCCA-CCCUUUA
AAGAAAGCGUAAUAGCUCACUAGUC-----GAGUCGGCCUGCGCGGAAGAUGUAAC-GGGGCUCA-AACCAC
A----------CACCGAA-GCUGCGGGUGUCACG--C-----------------------------------
------------------------------------------------------------------------
------------------------------------------------------------------------
---------------------------------AAGU-GACG-CGGUAGAGGAGC-GUUCUGU---------
-----------AAG-CCUGU-GAAGGUG--A-GUUGAGAAGC--U-UGCUGGAGGU-A-UCA-GAAGUGCGA
AUGC-UGACAU-GAGUA-ACGACA---AUGG-GUGUGA--A---------------AAACAC-CCA-CGCCG
A-AAGACC-AAGGGUUCC-UGCGCAACG-UUAAUCGACGCAGGGUUAGUCGGU-UC-CUAAGGCGAGGCUG-
A-A-AAGC-----GUA--------------------------------------------------------
----------------------------------------GUC-GA-U--GGGA-AACAGGUU-AAUA-UUC
CUGUA----------------------------------CUUCU----GGUU--------A-CU--------
------------------------------------------------------------------------
----------------------------------------------------------GCGAU-GGAGGGAC
GGAGAA--------------GGCUAG-GCCAGCUUG-GC-G----UUGGU------------UGUCCAAGU-
UUAAGG-UGG-UA-----G-G-------------------CU-----GA------------------AA-UC
U-UAGG--------------------UAAAU-CCGGGGUUUC--------AAGGCCG----AGA-GC-U-G-
AUG--AC---------------------------------------------------------------GA
---GUCGUCUU-UUAGAUGACGAAG-U-GGU-U-GAUGCCAUGCUUC-C-AAGAAA-AGCUU-CUAAGCUU-
-------------------------------------------------------CAGGUAA-CC-AGGAAC
CGUACCCCAAACCGAC------------------------------ACAGGUG--GUCGGGUA-GAGAAUAC
CAAGGC-G-CU-UGAG-AGAACUCG-G-GUGAAGGAACUAGGC-AAAAUGGCACCGUAACUUCGG-GAGAAG
GUGCG-CCGG-C-UA--GGGUGAA-----GGA-----------UUUAC------------------------
------------------------------------------------------------------------
------------------------------------------------------------------------
------------------------------------------------------------------------
------------------------------------------------------------------------
------------------------------------------------------------------------
------------------------------------------------------------------------
------------------------------------------------------------------------
------------------------------------------------------------------------
------------------------------------------------------------------------
------------------------------------------------------------------------
------------------------------------------------------------------------
------------------------------------------------------------------------
------------------------------------------------------------------------
----------------------------------------------------------UCCG-UAAGCUCUG
GCU--GGUCGAAGAUACCAGGCCGCUGCGA-----CUGUUUAUUAAAAACACAGCACUCUGCAAACACG---
-AA-A-------------GUGGACGU-AUAGGGUGUGA-CGCCUGCCCGGUGCCG-G-AAGG-UUAAUUGAU
GGGGUUA-GC-----GCAA-----GCGAAGCUCUUGAUCGAAGCCC--CGGUAAAC-GGCGGCCGUAACUAU
AACGGUCCUAAGGUAGCGAAAUUCCUUGUCGGG-UAAGUUCCGACCUGCACGAAUGGCGUAACGAUG--GCG
GCGCUGUCUCCACC----CGA-GACUCAGUGAAAUU--GAA---AUCGCUGUGAAGAUGCAGUGUAUCCGCG
```

```
GCUAGACGGAAAGA-CCCCGUGAACC-UUUACU--GUAGCUU-UGCACUGGACUUUGAGCCUGCUUGUGUAG
GAU-AGGUGGGAGGCUUUG----------AAGCGUG-GAC----------------------------GCCA
--------------------------------------GU-UCGCGUGGAG-C--C-AU-CCUUGAAAUA-C
CACC-CUGGCAUGCUUGAGGUUCUAACUCUGG--UCCG-U-GA-U-C-------------------------
------------------------------------------------------------------------
------------------------------------------------------------------------
------------------------------------------------------------------------
------------------------------------------------------------------------
------------------------------------------------------CG----GA----------
------------------------------------------------UCGA-----GGACA-GUGUAUGGU
GGGCAGUUUGACUGGGGCGGU-CUCCUCCUA-AAGAGUAAC-GGAGGAGUACG-AAGGUGCGCUCAGACCGG
UCGGAAAUCGGUCGUA-GAGUA-UAAAGGCAA-AAGCGCGCUUGACU-GCGA--------------------
--------------GACAGA---CACGUCGAGCAGGUACGAAA-GUAGGUCUUAGUGAUCCGGU--------
---GG-U-----UCUGU-AUG--------------------GA-AGGGCCAUCGCUCAACGG-AUAAAAGGU
ACUCCGGGG-AUAACAGGCUGAUACCGCCCAAGAGUUCAUAUCGACGGCGGUGUUGGCACCUCGAUGUCGG
CUCAUCACAUCCUGGGGCUGAA-GCCGGUCCCAAGGGUAUGGCUGUUCGCCAUUUAAAGUGGUACGCGAGCU
GGGUUUAGAACGUCGUGAGACAGUUCGGUCCCUAUCUGCC--------------------G-UGG-------
-------------------------------ACGU--UUGAGAUUUGAGAGG-GGC-UGCUCCUAGUACGAG
AGGAC-CGGAGUGGACGAACCUC-UGGU-G-UUCCGGUUGUCACGCCAGUGGCAU-U-GCCGGGUAGCUAUG
UUCGGAAAAGAUAACCGCUGAAAGCA-UC--UAAGCGGGAAACUUGCCUCAA-G-AUG--AGAUCUCA-CU-
--GGGAA-----------------------------------------------------------------
--------------------------CU--------UGA---------------------------------
------------------------------------------------------------------------
------------------------------------------------------------------------
------------------------------------------------------------------------
------------------------------------------------------------------------
------------------------------------------------------------------------
------------------------------------------------------------------------
-------------------------------------------------UUCCCCU--GA-AGG------GC
CGUCGAAGACUACGACGU-UGA-UAGGCU-GGG-UGUGUAAG-CGU-U----------------GU------
----GA-----------GGCGUUGAGCUA--A-CCAG-UACUAAUU---GCCCGUGAG-GCUUGACCA----
------------------------------------------------------------------------
------------------------------------------------------------------------
------------------------------------------------------------------------
------------------------------------U------------
>LSUb_563178_BUCA5/1-2913
---------------------G-GUUAAGCA----AA--UAAGCGUACAU-GGUGAAUG-C-CUUGGC-AGU
CAG---AGGCGAAGAAGGACGUGC--UAAUCUGCGAAAAGCGUCGGCAAGCUGAUAUGAAGCGC-----AAU
AAC-CGU-CGAUAUCCGAAUGGGGAAAC--------CC--------GA----UGCA--------------AU
A-U----------------------U----G---CAU----C-A-UUAU-UAA-----CUGAA---------
-----------U-----------CAA----------------------------------------AUA-AG
U-U--------------------AA-U-AA-AGCAAACCAAGGGAACUGAAACAUCUAAGUACCUU--GAGG
AAAAGAAAUCAA--CCGAGAUUCCCCA--GU-AG-UGGCGAGCGAAAAGGGA-A-CA-GCCCA--------
------------------------------------------------------------------------
-------GA-AUC-------A-AA-AU----------------------CA--AUA-----------UA---
```

```
A-------AUUAU-AGUAG------AAU-GA-UUUG------------------------------------
-------GAA-AU-UUCAG-------CGAUACAAGGUGAUAGCCCGUA----------UACAAAA----UAU
U------U--UAUAU-----------UGU-----GG--AU-UC------AAAAAGUAGAGCGGGACACGAGA
AAUCCUGUUUGAA-UAUGGGGGG-ACC--AUCCUCCAAGGCUAAAUA--CUC-CUGAC-UGACCGAUAGU-G
AAC-U-AGUACCGUAAGGGAAAG-GCGAAA-AGAAC-CCCGGCGAGGGG-A-GUGAAAUAGAACCUGAAACC
G-UGUACGUACAA-GCA-GU-GGAA---GCAU-A---UU---------------------------------
------------------------------------------------------------------------
------------------------------------------------------------------------
------------------------------------------------------------------------
------------------------------------------------------------------------
------------------------------------------------------------------------
------------------------------------------------------------U-----------
------------------------------------------------------------------------
------------------------------------------------------------------------
-------------------------------------------------------------UA--A-AA---
------------------------------------------------------------------------
------------------------------------------------------------------------
------------------------------------------------------------------------
------------------------------------------------------------------------
------------------------------------------------------------------------
------------------------------------------------------------------------
------------------------------------------------------------------------
------------------------------------------------------------------------
------------------------------------------------------------------------
------------------------------------------------------------------------
------------------------------------------------------------------------
------------------------------------------------------------------------
---------------------------U------------------AAUU--U---GUGACUGCGUACCUUU
UGUAUAAUGGGUCAGCGACUUGU-A-CUCUGUAGCAA-GGUUAACU-------------UAAUA--------
---A--GGGA--GCCGAAGGGAAAC-C---------------------------------------------
------------------------------------------------------------------------
------------------------------------------------------------------------
---------------------------------------------------------------------GAG
UCC------------------------------------UAAAU-------------------------AG-
G-C-GU----------------------UAAGUUUCAGGGUACA-GACCCGAAACC-CGGUGAUCUAGCCAC
GGGCAGGUUGAAGGUUGG-GUAAAA-CCAACUGGAGGACCGA--ACCGACUG-AUGUUGAAAAAUCA-GCGG
AUGACCUAUGGCUAGGGGUGAAAGGCCAAUCAAACCGGG-AGAUAGCUGGUUCUCCCCGAAA-GCUAUUUAG
GUAGCGCCUCGU-GAA--------------------------------------------------------
----------------------UU-----CA-UCUUCGGGGGUAGAGC-ACUGUUU-CGA-CUA--GGGGGC
C---AUCCC--GGCUUACCAAACCGAUGCAAACUCCGAAUACCGUA--------------------------
------------------------------------------------------------------------
------------------GAAU-GUUA--------UCACGGG-AGACACA-C-AGCG-G-GUGCUAACGUUC
GU-U-GUGGAGAG-GGAAAC---AACCCAGACCGCCA-GCUAAGGUCCCAAAGU--CA-UAG-UU-AA---G
-U-GGG-------AAACGAUGUGGGAAGGCAUAAACAGCCAGGAUGUUGGCUUAGAAGCAGCCA-UCAUUUA
AAGAAAGCGUAAUAGCUCACUGGUC-----AAGUCGGCCUGCGCGGAAGAUGUAAC-GGGGCUCA-AACUAU
```

```
G----------CACCGAA-GCUGCGGCAGCAAAA--A-----------------------------------
------------------------------------------------------------------------
-----UU-----------------------------------------------------------------
------------------------------AUAUUUU-UGCU-GGGUAGGGGAGC-GUUCUGU---------
-----------AAG-CCAAC-GAAGAUA--U-AUUGUAAAAU--A-UAUUGGAGGU-A-UCA-GAAGUGCGA
AUGC-UGACAU-GAGUA-ACGAUA---AAGC-AGGUGA--A---------------AAACCU-GCU-CGCCG
A-AAAACC-AAGGUUUCC-UGUCCAACG-UUAAUCGGGGCAGGGUAAGCCGAC-CC-CUAAGGCGAGGCUG-
A-A-AAGC-----GUA--------------------------------------------------------
----------------------------------------GUC-GA-U--GGAC-AACAGGUU-AAUA-UUC
CUGUG----------------------------------CUUAA----UAUU--------A-CU--------
------------------------------------------------------------------------
----------------------------------------------------------GCGAA-GUGGGGAC
GAAGAA--------------GGUUAA-GUUAUCCAG-GU-G----ACGGU------------UAUCCUGGU-
UUAAAU-GUG-UA-----G-G-------------------UA-----UG------------------AU-AA
U-UAGG--------------------CAAAU-CCGAUUAUCU------UU-AAUCUG----AGG-CA-U-G-
AUG--AC---------------------------------------------------------------GA
-----AUCAU--UAAGGUGAUGAAG-U-AAC-U-GACACCACGCUUC-C-AAGAAA-AGCCU-CUAAGCAU-
-------------------------------------------------------CAGGUAA-AA-UUAAAU
CGUACCCCAAACCGAC------------------------------ACAGGUG--GUUAGGUA-GAGAAUAC
UAAGGC-G-CU-UGAG-AGAACCCA-G-GUGAAGGAACUAGGC-AAAAUAGUGCCGUAACUUCGG-GAGAAG
GCACG-CUAA-U-GU-UAAGUGAAA----GGA-----------UUUAC------------------------
------------------------------------------------------------------------
------------------------------------------------------------------------
------------------------------------------------------------------------
------------------------------------------------------------------------
------------------------------------------------------------------------
------------------------------------------------------------------------
------------------------------------------------------------------------
------------------------------------------------------------------------
------------------------------------------------------------------------
------------------------------------------------------------------------
------------------------------------------------------------------------
------------------------------------------------------------------------
------------------------------------------------------------------------
--------------------------------------------------------UCCU-AUA--GCUGAA
GUU--AGUCGAAGAUACCAGCUGGCUGCAA-----CUGUUUAUUAAAAACACAGCACUGUGCAAACACG---
-AA-A-------------GUGGAAGU-AUACGGUGUGA-CGCCUGCCCGGUGCCG-G-AAGG-UUAAUCGAA
GGAGUCA-AAG----GAAA----CUUAAAGCUCUGAACUGAAGCCC--CGGUAAAC-GGCGGCCGUAACUAU
AACGGUCCUAAGGUAGCGAAAUUCCUUGUCGGG-UAAGUUCCGACCUGCACGAAUGGCGUAAUGAUG--GCC
AGGCUGUCUCCACC----UGG-GACUCAGUGAAAUU--GAA---UUUGCUGUGAAGAUGCAGUAUAUCCGCG
GCAAGACGGAAAGA-CCCCGUGAACC-UUUACU--AUAGCUU-GACACUGAAUUCUGAAUUUUAAUGUGUAG
GAU-AGGUGGGAGGCCAUG----------AAGUUAA-AAC----------------------------GCUA
--------------------------------------GU-UUUAAUGGAG-C--C-AA-ACUUGAAAUA-C
CACC-CUUUAAAAUUCGGUAUUCUAACCUAGU--GCCG-U-AA-U-C-------------------------
------------------------------------------------------------------------
```

```
------------------------------------------------------------------------
------------------------------------------------------------------------
------------------------------------------------------------------------
------------------------------------------------------CG----GC----------
------------------------------------------------ACAG-----AGACA-GUGUCUGGU
GGGUAGUUUGACUGGGGCGGU-CUCCUCCCA-AAGAGUAAC-GGAGGAGUACG-AAGAUUGGCUAAUCACGG
UCGGACAUCGUGAGGU-UAGUG-CAAAGGCAU-AAGCCAGUUUGACU-GUGA--------------------
--------------GCGUGA---UAACGCGAGCAGGUGCGAAA-GCAGGUCUUAGUGAUCCGGU--------
---GG-U-----UCUGA-AUG--------------------GA-AGGGCCAUCGCUCAACGG-AUAAAAGGU
ACUCCGGGG-AUAACAGGCUGAUACCGCCCAAGAGUUCAUAUCGACGGCGGUGUUUGGCACCUCGAUGUCGG
CUCAUCACAUCCUGGGGCUGAA-GCAGGUCCCAAGGGUAUGGCUGUUCGCCAUUUAAAGUGGUACGCGAGCU
GGGUUUAGAACGUCGUGAGACAGUUCGGUCCCUAUCUGCC--------------------G-UGG-------
-------------------------------GCGU--UGGAAGAUUGAGGGG-GGC-UGCUUCUAGUACGAG
AGGAC-CGAAGUGGACGCAUCAC-UGGU-G-UUCGGGUUGUCAUGCCAAUGGCAU-U-GCCCGGUAGCUAAA
UGCGGGAAAGAUAAGCGCUGAACGCA-UA--UAAGCACGAAACUUGCCCCAA-G-AUG--AAUCUUCC-CU-
--GAAAC-----------------------------------------------------------------
--------------------------AGA-------AAU---------------------------------
------------------------------------------------------------------------
------------------------------------------------------------------------
------------------------------------------------------------------------
------------------------------------------------------------------------
------------------------------------------------------------------------
------------------------------------------------------------------------
-------------------------------------------------GUUUACU--GA-AGG------GA
CGUUGAAGACUACGACGU-UGA-UAGGCU-AGG-UGUGUAAG-CAU-A----------------GC------
----AA-----------UAUGUUGAGCUA--A-CUAG-UACUAAUG---ACCCAAGAG-GCUUAACCU----
------------------------------------------------------------------------
------------------------------------------------------------------------
------------------------------------------------------------------------
------------------------------------U------------
>LSUb_83333_ECOLI/1-2904
---------------------G-GUUAAGCG----AC--UAAGCGUACAC-GGUGGAUG-C-CCUGGC-AGU
CAG---AGGCGAUGAAGGACGUGC--UAAUCUGCGAUAAGCGUCGGUAAGGUGAUAUGAACCGU-----UAU
AAC-CGG-CGAUUUCCGAAUGGGGAAAC--------CC--------AG----UGUG-U------------UU
C-G-----------------A----C----A---CAC----U-A-UCAU-UAA-----CUGA----------
-----------A-----------UCCA----------------------------------------UA-GG
U-U--------------------AA-U-GA-GGCGAACCGGGGGAACUGAAACAUCUAAGUACCCC--GAGG
AAAAGAAAUCAA--CCGAGAUUCCCCCA--GU-AG-CGGCGAGCGAACGGGGA-G-CA-GCCCA--------
------------------------------------------------------------------------
----------GAG-------CCUGAAU----------------------CA--GUG-----------UG---
U-------GUGUU-AGUGG------AAG-CG-UCUG------------------------------------
-------GAA-AG-GCGCG-------CGAUACAGGGUGACAGCCCCGUA---------CACAAAA----AUG
C------A--CAUGC-----------UGU-----GA--GC-UC------GAUGAGUAGGGCGGGACACGUGG
UAUCCUGUCUGAA-UAUGGGGGG-ACC--AUCCUCCAAGGCUAAAUA--CUC-CUGAC-UGACCGAUAGU-G
AAC-C-AGUACCGUGAGGGAAAG-GCGAAA-AGAAC-CCCGGCGAGGGG-A-GUGAAAAAGAACCUGAAACC
```

```
G-UGUACGUACAA-GCA-GU-GGGA---GCAC-G---C----------------------------------
------------------------------------------------------------------------
------------------------------------------------------------------------
------------------------------------------------------------------------
------------------------------------------------------------------------
------------------------------------------------------------------------
------------------------------------------------------------------------
------------------------------------------------------------------------
------------------------------------------------------------------------
--------------------------------------------------------------U--U-AG---
------------------------------------------------------------------------
------------------------------------------------------------------------
------------------------------------------------------------------------
------------------------------------------------------------------------
------------------------------------------------------------------------
------------------------------------------------------------------------
------------------------------------------------------------------------
------------------------------------------------------------------------
------------------------------------------------------------------------
------------------------------------------------------------------------
------------------------------------------------------------------------
------------------------------------------------------------------------
-----------------------------------------------GCG--U---GUGACUGCGUACCUUU
UGUAUAAUGGGUCAGCGACUUAU-A-UUCUGUAGCAA-GGUUAACC-------------GAAUA--------
---G--GGGA--GCCGAAGGGAAAC-C---------------------------------------------
------------------------------------------------------------------------
------------------------------------------------------------------------
---------------------------------------------------------------------GAG
UCU------------------------------------UAACU-------------------------GG-
G-C-GU----------------------UAAGUUGCAGGGUAUA-GACCCGAAACC-CGGUGAUCUAGCCAU
GGGCAGGUUGAAGGUUGG-GUAACA-CUAACUGGAGGACCGA--ACCGACUA-AUGUUGAAAAAUUA-GCGG
AUGACUUGUGGCUGGGGGUGAAAGGCCAAUCAAACCGGG-AGAUAGCUGGUUCUCCCCGAAA-GCUAUUUAG
GUAGCGCCUCGU-GAA--------------------------------------------------------
----------------------UU-----CA-UCUCCGGGGGUAGAGC-ACUGUUU-CGG-CAA--GGGGGU
C---AUCCC--GACUUACCAACCCGAUGCAAACUGCGAAUACCGGA--------------------------
------------------------------------------------------------------------
------------------GAAU-GUUA--------UCACGGG-AGACACA-C-GGCG-G-GUGCUAACGUCC
GU-C-GUGAAGAG-GGAAAC---AACCCAGACCGCCA-GCUAAGGUCCCAAAGU--CA-UGG-UU-AA---G
-U-GGG-------AAACGAUGUGGGAAGGCCCAGACAGCCAGGAUGUUGGCUUAGAAGCAGCCA-UCAUUUA
AAGAAAGCGUAAUAGCUCACUGGUC-----GAGUCGGCCUGCGCGGAAGAUGUAAC-GGGGCU-A-AACCAU
G----------CACCGAA-GCUGCGGCAGCGACG--C-----------------------------------
------------------------------------------------------------------------
-----UUAU---------------------------------------------------------------
---------------------------------GCGU-UGUU-GGGUAGGGGAGC-GUUCUGU---------
-----------AAG-CCUGC-GAAGGUG--U-GCUGUGAGGC--A-UGCUGGAGGU-A-UCA-GAAGUGCGA
```

```
AUGC-UGACAU-AAGUA-ACGAUA---AAGC-GGGUGA--A---------------AAGCCC-GCU-CGCCG
G-AAGACC-AAGGGUUCC-UGUCCAACG-UUAAUCGGGGCAGGGUGAGUCGAC-CC-CUAAGGCGAGGCCG-
A-A-AGGC-----GUA--------------------------------------------------------
----------------------------------------GUC-GA-U--GGGA-AACAGGUU-AAUA-UUC
CUGUA----------------------------------CUUGG----UGUU--------A-CU--------
------------------------------------------------------------------------
----------------------------------------------------------GCGAA-GGGGGGAC
GGAGAA--------------GGCUAU-GUUGGCCGG-GC-G----ACGGU------------UGUCCCGGU-
UUAAGC-GUG-UA-----G-G-------------------CU-----GG------------------UU-UU
C-CAGG--------------------CAAAU-CCGGAAAAUC--------AAGGCUG----AGG-CG-U-G-
AUG--AC---------------------------------------------------------------GA
-----GGCAC--UACGGUGCUGAAG-C-AAC-A-AAUGCCCUGCUUC-C-AGGAAA-AGCCU-CUAAGCAU-
-------------------------------------------------------CAGGUAA-CA-UCAAAU
CGUACCCCAAACCGAC------------------------------ACAGGUG--GUCAGGUA-GAGAAUAC
CAAGGC-G-CU-UGAG-AGAACUCG-G-GUGAAGGAACUAGGC-AAAAUGGUGCCGUAACUUCGG-GAGAAG
GCACG-CUGA-U-AUGUAGGUGAGG----UCC-----------CUCGC------------------------
------------------------------------------------------------------------
------------------------------------------------------------------------
------------------------------------------------------------------------
------------------------------------------------------------------------
------------------------------------------------------------------------
------------------------------------------------------------------------
------------------------------------------------------------------------
------------------------------------------------------------------------
------------------------------------------------------------------------
------------------------------------------------------------------------
------------------------------------------------------------------------
------------------------------------------------------------------------
------------------------------------------------------------------------
--------------------------------------------------------GGAU-GGA--GCUGAA
AUC--AGUCGAAGAUACCAGCUGGCUGCAA-----CUGUUUAUUAAAAACACAGCACUGUGCAAACACG---
-AA-A-------------GUGGACGU-AUACGGUGUGA-CGCCUGCCCGGUGCCG-G-AAGG-UUAAUUGAU
GGGGUUA-GC-----GCAA-----GCGAAGCUCUUGAUCGAAGCCC--CGGUAAAC-GGCGGCCGUAACUAU
AACGGUCCUAAGGUAGCGAAAUUCCUUGUCGGG-UAAGUUCCGACCUGCACGAAUGGCGUAAUGAUG--GCC
AGGCUGUCUCCACC----CGA-GACUCAGUGAAAUU--GAA---CUCGCUGUGAAGAUGCAGUGUACCCGCG
GCAAGACGGAAAGA-CCCCGUGAACC-UUUACU--AUAGCUU-GACACUGAACAUUGAGCCUUGAUGUGUAG
GAU-AGGUGGGAGGCUUUG----------AAGUGUG-GAC----------------------------GCCA
--------------------------------------GU-CUGCAUGGAG-C--C-GA-CCUUGAAAUA-C
CACC-CUUUAAUGUUUGAUGUUCUAACGUUGA--CCCG-U-AA-U-C-------------------------
------------------------------------------------------------------------
------------------------------------------------------------------------
------------------------------------------------------------------------
------------------------------------------------------------------------
------------------------------------------------------CG----GG----------
------------------------------------------------UUGC-----GGACA-GUGUCUGGU
```

```
GGGUAGUUUGACUGGGGCGGU-CUCCUCCUA-AAGAGUAAC-GGAGGAGCACG-AAGGUUGGCUAAUCCUGG
UCGGACAUCAGGAGGU-UAGUG-CAAUGGCAU-AAGCCAGCUUGACU-GCGA--------------------
--------------GCGUGA---CGGCGCGAGCAGGUGCGAAA-GCAGGUCAUAGUGAUCCGGU--------
---GG-U-----UCUGA-AUG--------------------GA-AGGGCCAUCGCUCAACGG-AUAAAAGGU
ACUCCGGGG-AUAACAGGCUGAUACCGCCCAAGAGUUCAUAUCGACGGCGGUGUUUGGCACCUCGAUGUCGG
CUCAUCACAUCCUGGGGCUGAA-GUAGGUCCCAAGGGUAUGGCUGUUCGCCAUUUAAAGUGGUACGCGAGCU
GGGUUUAGAACGUCGUGAGACAGUUCGGUCCCUAUCUGCC--------------------G-UGG-------
-------------------------------GCGC--UGGAGAACUGAGGGG-GGC-UGCUCCUAGUACGAG
AGGAC-CGGAGUGGACGCAUCAC-UGGU-G-UUCGGGUUGUCAUGCCAAUGGCAC-U-GCCCGGUAGCUAAA
UGCGGAAGAGAUAAGUGCUGAAAGCA-UC--UAAGCACGAAACUUGCCCCGA-G-AUG--AGUUCUCC-CU-
--GACUC-----------------------------------------------------------------
--------------------------CU--------UGA---------------------------------
------------------------------------------------------------------------
------------------------------------------------------------------------
------------------------------------------------------------------------
------------------------------------------------------------------------
------------------------------------------------------------------------
------------------------------------------------------------------------
-------------------------------------------------GAGUCCU--GA-AGG------AA
CGUUGAAGACGACGACGU-UGA-UAGGCC-GGG-UGUGUAAG-CGC-A----------------GC------
----GA-----------UGCGUUGAGCUA--A-CCGG-UACUAAUG---AACCGUGAG-GCUUAACCU----
------------------------------------------------------------------------
------------------------------------------------------------------------
------------------------------------------------------------------------
------------------------------------U------------
>LSUb_214092_YERPE/1-2907
---------------------G-GUUAAGCG----AC--UAAGCGUACAC-GGUGGAUG-C-CUAGGC-AGU
CAG---AGGCGAUGAAGGGCGUGC--UAAUCUGCGAAAAGCGUCGGUAAGCUGAUAUGAAGCGU-----UAU
AAC-CGA-CGAUACCCGAAUGGGGAAAC-------CC---------AG----UGCA-A------------UA
C-G-----------------U----U----G---CAC----U-A-UCGU-UAG-----AUGA----------
-----------A-----------UACA----------------------------------------UAG-U
C-U--------------------AA-C-GA-GGCGAACCGGGGGAACUGAAACAUCUAAGUACCCC--GAGG
AAAAGAAAUCAA--CCGAGAUUCCCCCA--GU-AG-CGGCGAGCGAACGGGGA-G-GA-GCCCA--------
------------------------------------------------------------------------
----------GAG-------U----CUGAAU------------------CA--GUU-----------UG---
U-------GUGUU-AGUGG------AAG-CG-UCUG------------------------------------
-------GAA-AG-UCGCA-------CGGUACAGGGUGAUAGUCCCGUA---------CACCAAA----AUG
C------A--CAGGC-----------UGU-----GA--AC-UC------GAUGAGUAGGGCGGGACACGUGA
CAUCCUGUCUGAA-UAUGGGGGG-ACC--AUCCUCCAAGGCUAAAUA--CUC-CUGAC-UGACCGAUAGU-G
AAC-C-AGUACCGUGAGGGAAAG-GCGAAA-AGAAC-CCCGGCGAGGGG-A-GUGAAAUAGAACCUGAAACC
G-UGUACGUACAA-GCA-GU-GGGA---GCAC-C--------------------------------------
------------------------------------------------------------------------
------------------------------------------------------------------------
------------------------------------------------------------------------
------------------------------------------------------------------------
```

```
------------------------------------------------------------------------
------------------------------------------------------------------------
------------------------------------------------------------------------
------------------------------------------------------------------------
-------------------------------------------------------------UU--C-GU---
------------------------------------------------------------------------
------------------------------------------------------------------------
------------------------------------------------------------------------
------------------------------------------------------------------------
------------------------------------------------------------------------
------------------------------------------------------------------------
------------------------------------------------------------------------
------------------------------------------------------------------------
------------------------------------------------------------------------
------------------------------------------------------------------------
------------------------------------------------------------------------
------------------------------------------------------------------------
------------------------------------------------GG--U---GUGACUGCGUACCUUU
UGUAUAAUGGGUCAGCGACUUAU-A-UUUUGUAGCAA-GGUUAACC-------------GAAUA--------
---G--GGGA--GCCGUAGGGAAAC-C---------------------------------------------
------------------------------------------------------------------------
------------------------------------------------------------------------
---------------------------------------------------------------------GAG
UCU------------------------------------UAACU-------------------------GG-
G-C-GU----------------------CUAGUUGCAAGGUAUA-GACCCGAAACC-CGGUGAUCUAGCCAU
GGGCAGGUUGAAGGUUGG-GUAACA-CUAACUGGAGGACCGA--ACCGACUA-AUGUUGAAAAAUUA-GCGG
AUGACUUGUGGCUGGGGGUGAAAGGCCAAUCAAACCGGG-AGAUAGCUGGUUCUCCCCGAAA-GCUAUUUAG
GUAGCGCCUCGU-GAA--------------------------------------------------------
----------------------CU-----CA-UCUUCGGGGGUAGAGC-ACUGUUU-CGG-CUA--GGGGGC
C---AUCCC--GGCUUACCAAACCGAUGCAAACUCCGAAUACCGAA--------------------------
------------------------------------------------------------------------
------------------GAAU-GUUA--------UCACGGG-AGACACA-C-GGCG-G-GUGCUAACGUCC
GU-C-GUGAAGAG-GGAAAC---AACCCAGACCGCCA-GCUAAGGUCCCAAAGU--CA-UGG-UU-AA---G
-U-GGG-------AAACGAUGUGGGAAGGCACAGACAGCCAGGAUGUUGGCUUAGAAGCAGCCA-UCAUUUA
AAGAAAGCGUAAUAGCUCACUGGUC-----GAGUCGGCCUGCGCGGAAGAUGUAAC-GGGGCU-A-AACCAU
G----------CACCGAA-GCUGCGGCAGCGACA--C-----------------------------------
------------------------------------------------------------------------
-----UU-----------------------------------------------------------------
-------------------------------AGGUGU-UGUU-GGGUAGGGGAGC-GUUCUGU---------
-----------AAG-CCGUU-GAAGGUG--A-CCUGUGAGGG--U-UGCUGGAGGU-A-UCA-GAAGUGCGA
AUGC-UGACAU-AAGUA-ACGAUA---AUGC-GGGUGA--A---------------AAACCC-GCA-CGCCG
G-AAGACC-AAGGGUUCC-UGUCCAACG-UUAAUCGGGGCAGGGUGAGUCGAC-CC-CUAAGGCGAGGCUG-
A-A-AAGC-----GUA--------------------------------------------------------
----------------------------------------GUC-GA-U--GGGA-AACAGGUU-AAUA-UUC
CUGUA----------------------------------CUUGG----UGUU--------A-CU--------
```

```
------------------------------------------------------------------------
----------------------------------------------------------GCGAA-GGGGGGAC
GGAGAA--------------GGCUAG-GCUGGCCGG-GC-G----ACGGU------------UGUCCCGGU-
UUAA-GCGUG-UA-----G-G-------------------GG-----GA------------------GU-GA
C-CAGG--------------------UAAAU-CCGGUUGCUU-----AUCAACCCUG----AGG-CG-U-G-
AUG--AC---------------------------------------------------------------GA
-----UGCAC--CACGGUGCAGAAG-C-AGU-U-GAUGCCAUGCUUC-C-AGGAAA-AGCCU-CUAAGCAU-
-------------------------------------------------------CAGGUAA-CA-UUAAAU
CGUACCCCAAACCGAC------------------------------ACAGGUG--GUCAGGUA-GAGAAUAC
UCAGGC-G-CU-UGAG-AGAACUCG-G-GUGAAGGAACUAGGC-AAAAUGGUGCCGUAACUUCGG-GAGAAG
GCACG-CUGG-C-GU-UAGGUGAAA----GGA-----------CUAGC------------------------
------------------------------------------------------------------------
------------------------------------------------------------------------
------------------------------------------------------------------------
------------------------------------------------------------------------
------------------------------------------------------------------------
------------------------------------------------------------------------
------------------------------------------------------------------------
------------------------------------------------------------------------
------------------------------------------------------------------------
------------------------------------------------------------------------
------------------------------------------------------------------------
------------------------------------------------------------------------
------------------------------------------------------------------------
----------------------------------------------------------UCCU-GGAGCUGAA
GCC--AGUCGCAGAUACCAGCUGGCUGCAA-----CUGUUUAAUAAAAACACAGCACUGUGCAAACACG---
-AA-A-------------GUGGACGU-AUACGGUGUGA-CGCCUGCCCGGUGCUG-G-AAGG-UUAAUUGAU
GGGGUCA-GCC----GCAA----GGGGAAGCUCUUGAUCGAAGCCC--CAGUAAAC-GGCGGCCGUAACUAU
AACGGUCCUAAGGUAGCGAAAUUCCUUGUCGGG-UAAGUUCCGACCUGCACGAAUGGCGUAAUGAUG--GCC
AGGCUGUCUCCACC----CGA-GACUCAGUGAAAUU--GAA---CUCGCUGUGAAGAUGCAGUGUACCCGCG
GCAAGACGGAAAGA-CCCCGUGAACC-UUUACU--AUAGCUU-GACACUGAACAUUGAGCCUUGAUGUGUAG
GAU-AGGUGGGAGGCCUUG----------AAGUGUG-GAC----------------------------GCCA
--------------------------------------GU-CUGCAUGGAG-C--C-AA-CCUUGAAAUA-C
CACC-CUUUAAUGUUUGAUGUUCUAACUCGGC--CCCA-U-GA-U-C-------------------------
------------------------------------------------------------------------
------------------------------------------------------------------------
------------------------------------------------------------------------
------------------------------------------------------------------------
------------------------------------------------------UG----GG----------
------------------------------------------------GUGA-----GGACA-GUGUCUGGU
GGGUAGUUUGACUGGGGCGGU-CUCCUCCCA-AAGAGUAAC-GGAGGAGCACG-AAGGUUAGCUAAUCACGG
UCGGACAUCGUGAGGU-UAGUG-CAAAGGCAU-AAGCUAGCUUGACU-GCGA--------------------
--------------GAGUGA---CGGCUCGAGCAGGUACGAAA-GUAGGUCUUAGUGAUCCGGU--------
---GG-U-----UCUGA-AUG--------------------GA-AGGGCCAUCGCUCAACGG-AUAAAAGGU
ACUCCGGGG-AUAACAGGCUGAUACCGCCCAAGAGUUCAUAUCGACGGCGGUGUUUGGCACCUCGAUGUCGG
```

```
CUCAUCACAUCCUGGGGCUGAA-GUAGGUCCCAAGGGUAUGGCUGUUCGCCAUUUAAAGUGGUACGCGAGCU
GGGUUUAGAACGUCGUGAGACAGUUCGGUCCCUAUCUGCC--------------------G-UGG-------
-------------------------------GCGU--UGGAAGAUUGAGAGG-GGC-UGCUCCUAGUACGAG
AGGAC-CGGAGUGGACGAAUCAC-UGGU-G-UUCGGGUUGUCAUGCCAAUGGCAU-U-GCCCGGUAGCUAAA
UUCGGAAGAGAUAACCGCUGAAAGCA-UC--UAAGCGGGAAACUUGCCUCGA-G-AUG--AGUCUUCC-CU-
--GGGAC-----------------------------------------------------------------
--------------------------UU--------AAA---------------------------------
------------------------------------------------------------------------
------------------------------------------------------------------------
------------------------------------------------------------------------
------------------------------------------------------------------------
------------------------------------------------------------------------
------------------------------------------------------------------------
-------------------------------------------------GUCCCCU--GA-AGG------AA
CGUUAAAGACUAUGACGU-UGA-UAGGCU-GGG-UGUGUAAG-UGC-A----------------GC------
----GA-----------UGUAUUGAGCUA--A-CCAG-UACUAAUG---AUCCGUGAG-GCUUAACCU----
------------------------------------------------------------------------
------------------------------------------------------------------------
------------------------------------------------------------------------
------------------------------------U------------
>LSUb_243265_PHOLL/1-2909
---------------------G-GUUAAGCG----AC--UAAGCGUACAC-GGUGGAUG-C-CUAGGC-AGU
CAG---AGGCGAUGAAGGACGUGC--UAAUCUGCGAAAAGCGUCGGUGAGCUGAUAUGAAGCGA-----CAG
AGC-CGG-CGAUGUCCGAAUGGGGAAAC--------CC--------AG----UGCA-A------------UG
C-G-----------------U----U----G---CAC----U-A-UCAU-UAA-----CUGA----------
-----------A-----------UUCA----------------------------------------UA-GG
U-U--------------------AA-U-GA-GGCGAACCGGGGGAACUGAAACAUCUCAGUACCCC--GAGG
AAAAGAAAUCAA--CCGAGAUUCCCCCA--GU-AG-CGGCGAGCGAACGGGGA-G-GA-GCCCA--------
------------------------------------------------------------------------
-------GA-ACC-------U-GC-AU----------------------CA--GCG-----------AU---
A-------GCAUC-AGAGG------AAC-GG-UCUG------------------------------------
-------GAA-AG-GCCGG-------CGAGAGAGGGUGACAGCCCCGUA---------CUCGAAG----AUG
G------U--AUGGC-----------UGU-----GA--GU-UC------GAUGAGUAAGGCGGGACACGAGA
AAUCCUGUCUGAA-GGUGGGGGG-ACC--AUCCUCCAAGGCUAAAUA--CUC-CUGAC-UGACCGAUAGU-G
AAC-C-AGUACCGUGAGGGAAAG-GCGAAA-AGAAC-CCCGGCGAGGGG-A-GUGAAAGAGAACCUGAAACC
G-UGUACGUACAA-GCA-GU-GGGA---GCCG-U---GA---------------------------------
------------------------------------------------------------------------
------------------------------------------------------------------------
------------------------------------------------------------------------
------------------------------------------------------------------------
------------------------------------------------------------------------
------------------------------------------------------------------------
------------------------------------------------------------------------
------------------------------------------------------------------------
--------------------------------------------------------------U--U-AA---
```

```
------------------------------------------------------------------------
------------------------------------------------------------------------
------------------------------------------------------------------------
------------------------------------------------------------------------
------------------------------------------------------------------------
------------------------------------------------------------------------
------------------------------------------------------------------------
------------------------------------------------------------------------
------------------------------------------------------------------------
------------------------------------------------------------------------
------------------------------------------------------------------------
------------------------------------------------------------------------
----------------------------------------------UCAG--G---GUGACUGCGUACCUUU
UGUAUAAUGGGUCAGCGACUUAU-A-UUCUGUAGCAA-GGUUAACC-------------GGAUA--------
---G--GGGA--GCCGCAGGGAAAC-C---------------------------------------------
------------------------------------------------------------------------
------------------------------------------------------------------------
---------------------------------------------------------------------GAG
UCU------------------------------------UAACU-------------------------GG-
G-C-GA---------------------AAGAGUUGCAGGGUAUA-GACCCGAAACC-CGGUGAGCUAGCCAU
GGGCAGGUUGAAGGUUGG-GUAACA-CUAACUGGAGGACCGA--ACCGACUA-AUGUUGAAAAAUUA-GCGG
AUGACCUGUGGCUGGGGGUGAAAGGCCAAUCAAACCGGG-AGAUAGCUGGUUCUCCCCGAAA-GCUAUUUAG
GUAGCGCCUCGU-GAA--------------------------------------------------------
----------------------UU-----CA-UCUUCGGGGGUAGAGC-ACUGUUU-CGG-CUA--GGGGGC
C---AUCCC--GGCUUACCAACCCGAUGCAAACUGCGAAUACCGAA--------------------------
------------------------------------------------------------------------
------------------GAAU-GGUA--------UCACGGG-AGACACA-C-GGCG-G-GUGCUAACGUCC
GU-C-GUGAAGAG-GGAAAC---AACCCAGACCGCCA-GCUAAGGUCCCGAAGU--CA-UGG-UU-AA---G
-U-GGG-------AAACGAUGUGGGAAGGCCCAGACAGCCAGGAUGUUGGCUUAGAAGCAGCCA-UCAUUGA
AAGAAAGCGUAAUAGCUCACUGGUC-----GAGUCGGCCUGCGCGGAAGAUGUAAC-GGGGCU-A-AACCAU
G----------CACCGAA-GCUGCGGCAGUGACA--C-----------------------------------
------------------------------------------------------------------------
-----GC-----------------------------------------------------------------
-------------------------------AAGUGU-UAUU-GGGUAGGGGAGC-GUUCUGU---------
-----------AAG-CCGGA-GAAGGUG--GAGCUGUGAGGC--C-UGCUGGAGGU-A-UCA-GAAGUGCGA
AUGC-UGACAU-AAGUA-ACGAUA---AAGC-GGGUGA--A---------------AAACCC-GCU-CGCCG
G-AAGACC-AAGGGUUCC-UGUCCAACG-UUAAUCGGGGCAGGGUGAGUCGAC-CC-CUAAGGUGAGGCCG-
A-G-AGGC-----GUA--------------------------------------------------------
----------------------------------------GCU-GA-U--GGGA-AACAGGUU-AAUA-UUC
CUGUA----------------------------------CUGAA----UGUG--------A-CU--------
------------------------------------------------------------------------
----------------------------------------------------------GCGAA-GGGGGGAC
GGAGAA--------------GGCUAG-GCCAUCCGG-GC-G----ACGGU-------------UGUCCCGGU-
UUAAGC-GUG-UA-----G-G--------------------UG-----GG------------------AG-GA
G-CAGG--------------------UAAAU-CCGCUCUUCU-----GUGAACACUG----AGG-CG-U-G-
```

```
AUG--AC---------------------------------------------------------------GA
-----GCCGU--UACGGCGGUGAAG-U-GGU-U-GAUGCCCGGCUUC-C-AGGAAA-AGCCU-CUAAGCCU-
-------------------------------------------------------CAGGUCA-UA-UUGAAU
CGUACCCCAAACCGAC------------------------------ACAGGUG--GUCAGGUA-GAGAAUAC
UCAGGC-G-CU-UGAG-AGAACUCG-G-GUGAAGGAACUAGGC-AAAAUGGUGCCGUAACUUCGG-GAGAAG
GCACG-CUGG-C-GG-UAGGUGAAG----GGG-----------UUUGC------------------------
------------------------------------------------------------------------
------------------------------------------------------------------------
------------------------------------------------------------------------
------------------------------------------------------------------------
------------------------------------------------------------------------
------------------------------------------------------------------------
------------------------------------------------------------------------
------------------------------------------------------------------------
------------------------------------------------------------------------
------------------------------------------------------------------------
------------------------------------------------------------------------
------------------------------------------------------------------------
------------------------------------------------------------------------
--------------------------------------------------------CUCC-GGA--GCUGAA
GCC--AGUCGCAGAGACCAGCUGGCUGCAA-----CUGUUUAUUAAAAACACAGCACUGUGCAAACACG---
-UA-A-------------GUGGACGU-AUACGGUGUGA-CGCCUGCCCGGUGCUG-G-AAGG-UUAAUUGAU
GGGGUUA-GCC----GCAA----GGCGAGGCUCUUGACAGAAGCCC--CAGUAAAC-GGCGGCCGUAACUAU
AACGGUCCUAAGGUAGCGAAAUUCCUUGUCGGG-UAAGUUCCGACCUGCACGAAUGGCGUAAUGAUG--GCC
AGGCUGUCUCCACC----CGA-GACUCAGUGAAAUU--GAA---CUCGCUGUGAAGAUGCAGUGUCCCCGCG
GCAAGACGGAAAGA-CCCCGUGAACC-UUUACU--AUAGCUU-GACACUGAACAUGGAGCCUUGAUGUGUAG
GAU-AGGUGGGAGGCUGAG----------AAGUGCG-GAC----------------------------GCCA
--------------------------------------GU-CUGCAUGGAG-C--C-AU-CCUUGAAAUA-C
CACC-CUUGAAUGUUCGAUGUUCUCACCUGGG--CCCG-U-AA-A-C-------------------------
------------------------------------------------------------------------
------------------------------------------------------------------------
------------------------------------------------------------------------
------------------------------------------------------------------------
------------------------------------------------------CG----GG----------
------------------------------------------------CCGG-----GGACA-GUGUCUGGC
GGGUAGUUUGACUGGGGCGGU-CUCCUCCCA-AAGGGUAAC-GGAGGAGCACG-AAGGUUGGCUAAUCACGG
UCGGACAUCGUGAGGU-UAGUG-CAAAGGCAU-AAGCCAGCUUAACU-GCGA--------------------
--------------GAGUGA---CGGUUCGAGCAGGUACGAAA-GUAGGUCUUAGUGAUCCGGU--------
---GG-U-----UCUGA-AUG--------------------GA-AGGGCCAUCGCUCAACGG-AUAAAAGGU
ACUCCGGGG-AUAACAGGCUGAUACCGCCCAAGAGUUCAUAUCGACGGCGGUGUUUGGCACCUCGAUGUCGG
CUCAUCACAUCCUGGGGCUGAA-GUAGGUCCCAAGGGUAUGGCUGUUCGCCAUUUAAAGUGGUACGCGAGCU
GGGUUUAGAACGUCGUGAGACAGUUCGGUCCCUAUCUGCC--------------------G-UGG-------
-------------------------------GCGC--AGGAAGAUUGAAAGG-GGC-UGCUCCUAGUACGAG
AGGAC-CGGAGUGGACGCACCGC-UGGU-G-UACGGGUUGUCAUGCCAAUGGCAU-A-GCCCGGUAGCUAAG
UGCGGGAGAGAUAACCGCUGAAAGCA-UC--UAAGCGGGAAACUUGCCUUGA-G-AUG--AGUCUUCC-CC-
```

```
--UGAC------------------------------------------------------------------
------------------------------------UGA---------------------------------
------------------------------------------------------------------------
------------------------------------------------------------------------
------------------------------------------------------------------------
------------------------------------------------------------------------
------------------------------------------------------------------------
------------------------------------------------------------------------
-------------------------------------------------GGUCACG--GA-AGG------AA
CGUUUAAGACGAAGACGU-GGA-UAGGCU-GGG-UGUGUAAG-UGC-A----------------GC------
----GA-----------UGCAUUGAGCUG--A-CCAG-UACUAAUG---AACCGAGAG-GCUUAACCU----
------------------------------------------------------------------------
------------------------------------------------------------------------
------------------------------------------------------------------------
------------------------------------U------------
>LSUb_211586_SHEON/1-2893
---------------------G-GUUAAGCG----AC--UAAGCGUAUAC-GGUGGAUG-C-CUUGGC-AGU
CAG---AGGCGAUGAAGGACGUAG--UAACUUGCGAAAAGCGUUGGCGAGCUAGUAACAAGCAU-----UUG
AGC-UAA-CGAUGUCCGAAUGGGGGAAC--------CC--------GG----CCGC--------------AU
A-A----------------------G----C---GGU----C-A-UCAU-AAC-----GUGA----------
-----------A-----------UACA----------------------------------------UAG-C
G-U--------------------UA-U-GA-GGCAAACGAGGGGAACUGAAACAUCUAAGUACCCU--UAGG
AAAAGAAAUCAA--CCGAGAUUCCCCUA--GU-AG-CGGCGAGCGAACGGGGA-U-UA-GCCCU--------
------------------------------------------------------------------------
----------UAA-------G----UC--------------------------AGA-----------GG---
G-------GUGUU-AGUGG------AAU-GG-UCUG------------------------------------
-------GAA-AG-UCCAG-------CGGCACAGGGUGAUAGCCCCGUA---------CACGAAA----ACU
A------A--CCUUU-----------GA----------UG-AA------AACGAGUAAGGCGGGACACGUGA
UAUCCUGUUUGAA-UAUGGGGGG-ACC--AUCCUCCAAGGCUAAAUA--CUC-CUGAC-UGACCGAUAGU-G
AAC-C-AGUACCGUGAGGGAAAG-GCGAAA-AGAAC-CCCUGUGAGGGG-A-GUGAAAUAGAACCUGAAACC
G-UAUACGUACAA-GCA-GU-GGGA---GCGG-U---A----------------------------------
------------------------------------------------------------------------
------------------------------------------------------------------------
------------------------------------------------------------------------
------------------------------------------------------------------------
------------------------------------------------------------------------
------------------------------------------------------------------------
------------------------------------------------------------------------
------------------------------------------------------------------------
-------------------------------------------------------------CU--U-GA---
------------------------------------------------------------------------
------------------------------------------------------------------------
------------------------------------------------------------------------
------------------------------------------------------------------------
------------------------------------------------------------------------
```

```
------------------------------------------------------------------------
------------------------------------------------------------------------
------------------------------------------------------------------------
------------------------------------------------------------------------
------------------------------------------------------------------------
------------------------------------------------------------------------
------------------------------------------------------------------------
-----------------------------------------------GAC--C---GUGACUGCGUACCUUU
UGUAUAAUGGGUCAGCGACUUAC-G-UUUUGUAGCGA-GGUUAAGC-------------GAAUA--------
---G--CGGA--GCCGUAGGGAAAC-C---------------------------------------------
------------------------------------------------------------------------
------------------------------------------------------------------------
---------------------------------------------------------------------GAG
UGU------------------------------------UAACU-------------------------GC-
G-C-GU----------------------UUAGUUGCAAGGCGUA-GACCCGAAACC-CGGUGAUCUAGCCAU
GGGCAGGUUGAAGGUUGA-GUAACA-UCAACUGGAGGACCGA--ACCGACUA-AUGUUGAAAAAUUA-GCGG
AUGACUUGUGGCUGGGGGUGAAAGGCCAAUCAAACCGGG-AGAUAUCUGGUUCUCCUCGAAA-GCUAUUUAG
GUAGCGCCUCGA-GCG--------------------------------------------------------
----------------------AA-----UA-CCAUUGGGGGUAGAGC-ACUGUUA-AGG-CUA--GGGGGU
C---AUCCC--GACUUACCAACCCUUUGCAAACUCCGAAUACCAAU--------------------------
------------------------------------------------------------------------
------------------GAGU-A-CU--------ACUCGGG-AGACAGA-C-GGCG-G-GUGCUAACGUCC
GU-C-GUCAAAAG-GGAAAC---AACCCAGACCGUCA-GCUAAGGUCCCAAAGU--GU-AUG-UU-AA---G
-U-GGG-------AAACGAUGUGGGAAGGCUUAGACAGCUAGGAUGUUGGCUUAGAAGCAGCCA-UCAUUUA
AAGAAAGCGUAAUAGCUCACUAGUC-----GAGUCGGCCUGCGCGGAAGAUGUAAC-GGGGCU-A-AACAUA
C----------CACCGAA-GCUACGGGUGCAAUC--C------------------------------------
------------------------------------------------------------------------
-----AU-----------------------------------------------------------------
-------------------------------UAGGGU-UGCG-CGGUAGAGGAGC-GUUCUGU---------
-----------AAG-CCGUA-GAAGGUG--AAGGGGUAACCC--A-CACUGGAGGU-A-UCA-GAAGUGCGA
AUGC-UGACAU-GAGUA-ACGAUA---AAGG-GGGUGA--A----------------AAACCC-CCU-CGCCG
A-AAGACC-AAGGGUUCC-UGUCCAACG-UUAAUCGGGGCAGGGUGAGUCGAC-CC-CUAAGGUGAGGCCG-
A-A-AGGC-----GUA--------------------------------------------------------
----------------------------------------AUC-GA-U--GGGA-AACAGAUU-AAUA-UUU
CUGUA----------------------------------CUUUU----GCUA--------A-CU--------
------------------------------------------------------------------------
----------------------------------------------------------GCGAU-GGAGAGAC
GGAGAA--------------GGCUAG-GCUAGCGCG-GC-G----UUGGU------------AGUCCGCGU-
UUAAGG-UGG-UA-----G-G-------------------UG-----GG------------------UG-AC
U-UAGG--------------------CAAAU-CCGGGUCACU------AU-ACACUG----AGA-GC-U-G-
AUG--AC---------------------------------------------------------------GA
-----GUCCC--CAAGGGGAUGAAG-U-AGU-U-GAUGCCAUGCUUC-C-AGGAAA-AUCUU-CUAAGCUU-
-------------------------------------------------------CAGGUUA-GU-AGGAAU
CGUACCCCAAACCGAC------------------------------ACAGGUG--GUCGGGUA-GAGAAUAC
CAAGGC-G-CU-UGAG-AGAACUCG-G-CUGAAGGAACUAGGC-AAAAUGGUACCGUAACUUCGG-GAGAAG
```

```
GUACG-CUCC-U-GU--UGGUGAUG----AGA-----------CUUGC------------------------
------------------------------------------------------------------------
------------------------------------------------------------------------
------------------------------------------------------------------------
------------------------------------------------------------------------
------------------------------------------------------------------------
------------------------------------------------------------------------
------------------------------------------------------------------------
------------------------------------------------------------------------
------------------------------------------------------------------------
------------------------------------------------------------------------
------------------------------------------------------------------------
------------------------------------------------------------------------
------------------------------------------------------------------------
----------------------------------------------------------UCUC-UAAGCUGAC
GGG--AGUCGCAGAUACCAGGUGGCUGCAA-----CUGUUUAUCAAAAACACAGCACUGUGCAAACUCG---
-CA-A-------------GAGGAAGU-AUACGGUGUGA-CGCCUGCCCGGUGCCG-G-AAGG-UUAAUUGAU
UGGGUUA-UC-----GCAA-----GAGAAGCUCAUGAUCGAAGCCC--CGGUAAAC-GGCGGCCGUAACUAU
AACGGUCCUAAGGUAGCGAAAUUCCUUGUCGGG-UAAGUUCCGACCUGCACGAAUGGCGUAAUGAUG--GCC
ACGCUGUCUCCAGC----CGA-GACUCAGUGAAGUU--GAA---AUUGCGGUGAAGAUGCCGUAUACCCGCG
GCUAGACGGAAAGA-CCCCGUGAACC-UUUACU--AUAGCUU-GGCACUGAACAUUGACCCUACAUGUGUAG
GAU-AGGUGGGAGACUUUG----------AAGUUGG-AAC----------------------------GCUA
--------------------------------------GU-UCUGAUGGAG-U--C-GU-CCUUGAAUA-C
CACC-CUUGUAGUGUUGAUGUUCUAACUUGGC--CCCA-U-UA-U-C-------------------------
------------------------------------------------------------------------
------------------------------------------------------------------------
------------------------------------------------------------------------
------------------------------------------------------------------------
------------------------------------------------------UG----GG----------
------------------------------------------------GUAA-----GGACA-GUGCCUGGU
GGGUAGUUUGACUGGGGCGGU-CUCCUCCCA-AAGAGUAAC-GGAGGAGCACG-AAGGUUGGCUAAGUACGG
UCGGACAUCGUACGGU-UAGUG-CAAUGGCAU-AAGCCAGCUUAACU-GCGA--------------------
--------------GACAGA---CACGUCGAGCAGGUACGAAA-GUAGGUCAUAGUGAUCCGGU--------
---GG-U-----UCUGA-AUG--------------------GA-AGGGCCAUCGCUCAACGG-AUAAAAGGU
ACUCCGGGG-AUAACAGGCUGAUACCGCCCAAGAGUUCAUAUCGACGGCGGUGUUUGGCACCUCGAUGUCGG
CUCAUCACAUCCUGGGGCUGAA-GUCGGUCCCAAGGGUAUGGCUGUUCGCCAUUUAAAGUGGUACGCGAGCU
GGGUUCAGAACGUCGUGAGACAGUUCGGUCCCUAUCUGCC--------------------G-UGG-------
-------------------------------GCGU--UGGAUGAUUGAGGGG-AGU-UGCUCCUAGUACGAG
AGGAC-CGGAGUGAACGAACCGC-UGGU-G-UUCGGGUUGUCAUGCCAAUGGCAU-U-GCCCGGUAGCUAUG
UUCGGAAUCGAUAACCGCUGAAAGCA-UC--UAAGCGGGAAGCGAGCCCCAA-G-AUG--AGUCAUCC-CU-
--UGGAC-----------------------------------------------------------------
--------------------------UU--------UAA---------------------------------
------------------------------------------------------------------------
------------------------------------------------------------------------
------------------------------------------------------------------------
```

```
------------------------------------------------------------------------
------------------------------------------------------------------------
------------------------------------------------------------------------
-------------------------------------------------GUCCACU--AA-AGA------GC
CGUUCGAGACUAGGACGU-UGA-UAGGUC-AGG-UGUGUAAG-CGU-U----------------GU------
----GA-----------GGCGUUGAGCUA--A-CUGA-UACUAAUG---ACUCGAGAG-GCUUAACCA----
------------------------------------------------------------------------
------------------------------------------------------------------------
------------------------------------------------------------------------
------------------------------------U------------
>LSUb_243277_VIBCH/1-2887
---------------------G-GUUAAGUG----AC--UAAGCGUACAC-GGUGGAUG-C-CUGGGC-AGU
CAG---AGGCGAUGAAGGACGUAC--UAACUUGCGAUAAGCGCAGAUAAGGCAGUAAGAGCCGU-----UUG
AGU-CUG-CGAUUUCCGAAUGGGGAAAC--------CC--------AA----CUGC--------------AU
A-A----------------------G----C---AGU----U-A-CUGU-UAA-----CUGA----------
-----------A-----------UACA----------------------------------------UAG-G
U-U--------------------AA-C-AG-AGCAAACCGGGGGAACUGAAACAUCUAAGUACCCC--GAGG
AGAAGAAAUCAA--CCGAGAUUCCGGUA--GU-AG-CGGCGAGCGAACCUGGA-U-UA-GCCCU--------
------------------------------------------------------------------------
----------UAA-------G----CA--------------------------CUC-----------GG---
U-------GAAGU-AGGUG------AAC-AA-GCUG------------------------------------
-------GAA-AG-CUUGG-------CGAUACAGGGUGAUAGCCCCGUA---------ACCGACG----CUU
C------A--UCGAG-----------CG----------UG-AA------AUCGAGUAGGGCGGGACACGUGA
UAUCCUGUCUGAA-UAUGGGGGG-ACC--AUCCUCCAAGGCUAAAUA--CUC-CUGAC-UGACCGAUAGU-G
AAC-C-AGUACCGUGAGGGAAAG-GCGAAA-AGAAC-CCCUGUGAGGGG-A-GUGAAAUAGAACCUGAAACC
G-UGUACGUACAA-GCA-GU-AGGA---GCAC-C--------------------------------------
------------------------------------------------------------------------
------------------------------------------------------------------------
------------------------------------------------------------------------
------------------------------------------------------------------------
------------------------------------------------------------------------
------------------------------------------------------------------------
------------------------------------------------------------------------
------------------------------------------------------------------------
-------------------------------------------------------------UU--C-GU---
------------------------------------------------------------------------
------------------------------------------------------------------------
------------------------------------------------------------------------
------------------------------------------------------------------------
------------------------------------------------------------------------
------------------------------------------------------------------------
------------------------------------------------------------------------
------------------------------------------------------------------------
------------------------------------------------------------------------
------------------------------------------------------------------------
```

```
------------------------------------------------------------------------
------------------------------------------------------------------------
------------------------------------------------GG--U---GUGACUGCGUACCUUU
UGUAUAAUGGGUCAGCGACUUAU-A-UUCAGUGGCAA-GGUUAACC-------------GUAUA--------
---G--GGGA--GCCGUAGCGAAAG-C---------------------------------------------
------------------------------------------------------------------------
------------------------------------------------------------------------
---------------------------------------------------------------------GAG
UCU------------------------------------UAACU-------------------------GG-
G-C-GC----------------------UCAGUCUCUGGAUAUA-GACCCGAAACC-GGGUGAUCUAGCCAU
GGGCAGGUUGAAGGUUGA-GUAACA-UCAACUGGAGGACCGA--ACCGACUA-AUGUUGAAAAAUUA-GCGG
AUGACUUGUGGCUAGGGGUGAAAGGCCAAUCAAACUCGG-AGAUAGCUGGUUCUCCCCGAAA-GCUAUUUAG
GUAGCGCCUCGG-ACG--------------------------------------------------------
----------------------AA-----UA-CUACUGGGGGUAGAGC-ACUGUUA-AGG-CUA--GGGGGU
C---AUCCC--GACUUACCAACCCUUUGCAAACUCCGAAUACCAGU--------------------------
------------------------------------------------------------------------
------------------AAGU-A-CU--------AUCCGGG-AGACACA-C-GGCG-G-GUGCUAACGUCC
GU-C-GUGGAGAG-GGAAAC---AACCCAGACCGCCA-GCUAAGGUCCCAAAGU---A-UUG-CU-AA---G
-U-GGG-------AAACGAUGUGGGAAGGCUCAGACAGCUAGGAUGUUGGCUUAGAAGCAGCCA-UCAUUUA
AAGAAAGCGUAAUAGCUCACUAGUC-----GAGUCGGCCUGCGCGGAAGAUGUAAC-GGGGCU-A-AGCAAU
A----------CACCGAA-GCUGCGGCAAUGUCU--U-----------------------------------
------------------------------------------------------------------------
------------------------------------------------------------------------
--------------------------------UUAGA-CAUU-GGGUAGGGGAGC-GUUCUGU---------
-----------AAG-CCGUU-GAAGGUG--A-AUCGUAAGGU--U-UGCUGGAGGU-A-UCA-GAAGUGCGA
AUGC-UGACAU-GAGUA-ACGACA---AAGG-GGGUGA--A---------------AAACCU-CCU-CGCCG
G-AAGACC-AAGGGUUCC-UGUCCAACG-UUAAUCGGGGCAGGGUGAGUCGAC-CC-CUAAGGUGAGGCCG-
A-A-AGGC-----GUA--------------------------------------------------------
-----------------------------------------AUC-GA-U--GGGA-AACGGUU-AAUA-UUC
CCGUA----------------------------------CUUCU----GACU--------A-UU--------
------------------------------------------------------------------------
----------------------------------------------------------GCGAU-GGGGGGAC
GGAGAA--------------GGCUAG-GUGGGCCAG-GC-G----ACGGU------------UGUCCUGGU-
UCAAGU-GCG-UA-----G-G-------------------CU-----UG------------------AG-AG
U-UAGG--------------------UAAAU-CCGGCUCUCU------CUAAGGCUG----AGA-CA-C-G-
ACG--UC---------------------------------------------------------------GA
-----GCUAC--UACGGUAGUGAAG-U-CAU-U-GAUGCCAUGCUUC-C-AGGAAA-AGCCU-CUAAGCUU-
-------------------------------------------------------CAGAUAG-UC-AGGAAU
CGUACCCCAAACCGAC------------------------------ACAGGUG--GUCGGGUA-GAGAAUAC
CAAGGC-G-CU-UGAG-AGAACUCG-G-GUGAAGGAACUAGGC-AAAAUGGUACCGUAACUUCGG-GAGAAG
GUACG-CUCU-U-GA--UGGUGAAG----UCC-----------CUCGC------------------------
------------------------------------------------------------------------
------------------------------------------------------------------------
------------------------------------------------------------------------
------------------------------------------------------------------------
```

```
------------------------------------------------------------------------
------------------------------------------------------------------------
------------------------------------------------------------------------
------------------------------------------------------------------------
------------------------------------------------------------------------
------------------------------------------------------------------------
------------------------------------------------------------------------
------------------------------------------------------------------------
------------------------------------------------------------------------
----------------------------------------------------------GGAU-GGAGCUGAC
GAG--AGUCGCAGAUACCAGGUGGCUGCAA-----CUGUUUAUUAAAAACACAGCACUGUGCAAAAUCG---
-CA-A-------------GAUGACGU-AUACGGUGUGA-CGCCUGCCCGGUGCCG-G-AAGG-UUAAUUGAU
GGGGUUA-GC-----GCAA-----GCGAAGCUCUUGAUCGAAGCCC--CGGUAAAC-GGCGGCCGUAACUAU
AACGGUCCUAAGGUAGCGAAAUUCCUUGUCGGG-UAAGUUCCGACCUGCACGAAUGGCGUAAUGAUG--GCC
ACGCUGUCUCCACC----CGA-GACUCAGUGAAAUU--GAA---AUCGCUGUGAAGAUGCAGUGUACCCGCG
GCUAGACGGAAAGA-CCCCGUGAACC-UUUACU--ACAGCUU-GGCACUGAACAUUGAACCUACAUGUGUAG
GAU-AGGUGGGAGGCUAUG----------AAGACGU-GAC----------------------------GCCA
--------------------------------------GU-UGCGUUGGAG-C--C-GU-CCUUGAAAUA-C
CACC-CUUGUAUGUUUGAUGUUCUAACUUAGA--CCCA-U-UA-U-C-------------------------
------------------------------------------------------------------------
------------------------------------------------------------------------
------------------------------------------------------------------------
------------------------------------------------------------------------
------------------------------------------------------UG----GG----------
------------------------------------------------UUGA-----GGACA-GUGCCUGGU
GGGUAGUUUGACUGGGGCGGU-CUCCUCCCA-AAGAGUAAC-GGAGGAGCACG-AAGGUGGGCUAAUCACGG
UUGGACAUCGUGAGGU-UAGUG-CAAUGGCAU-AAGCCCGCUUAACU-GCGA--------------------
--------------GAAUGA---CGGUUCGAGCAGGUGCGAAA-GCAGGUCAUAGUGAUCCGGU--------
---GG-U-----UCUGU-AUG--------------------GA-AGGGCCAUCGCUCAACGG-AUAAAAGGU
ACUCCGGGG-AUAACAGGCUGAUACCGCCCAAGAGUUCAUAUCGACGGCGGUGUUUGGCACCUCGAUGUCGG
CUCAUCACAUCCUGGGGCUGAA-GUCGGUCCCAAGGGUAUGGCUGUUCGCCAUUUAAAGUGGUACGCGAGCU
GGGUUUAGAACGUCGUGAGACAGUUCGGUCCCUAUCUGCC--------------------G-UGG-------
-------------------------------GCGU--UGGAAGAUUGAAGGG-GGC-UGCUCCUAGUACGAG
AGGAC-CGGAGUGGACGAACCUC-UGGU-G-UUCGGGUUGUGUCGCCAGACGCAU-U-GCCCGGUAGCUAAG
UUCGGAAUUGAUAAGCGCUGAAAGCA-UC--UAAGCGCGAAGCGAGCCCUGA-G-AUG--AGUCUUCC-CU-
--GACAG-----------------------------------------------------------------
--------------------------UU--------UAA---------------------------------
------------------------------------------------------------------------
------------------------------------------------------------------------
------------------------------------------------------------------------
------------------------------------------------------------------------
------------------------------------------------------------------------
------------------------------------------------------------------------
-------------------------------------------------CUGUCCU--AA-AGG------GU
UGUUCGAGACUAGAACGU-UGA-UAGGCA-GGG-UGUAAG-CGU-U------------------GU------
```

```
----GA-----------GGCGUUGAGCUA--A-CCUG-UACUAAUU---GCCCGUGAG-GCUUAACCA----
------------------------------------------------------------------------
------------------------------------------------------------------------
------------------------------------------------------------------------
------------------------------------U------------
>LSUb_203907_BLOFL/1-2960
---------------------A-GUUAAGUG----AU-GUAAGCGUAUAC-GGUGGAUG-C-CUUGGC-AGU
CAA---AGGCGAUGAAGGACGUGC--UAAUCUGCGAAAAGUAUCGGUGAGCUGAUUUGAAGCGU-U---AUU
AAC-CGG-UGAUGUCCGAAUGGGGAAAC--------CUA-------GU----GCAA-U------------UU
A-A-----------------A----U----U---GUG----CAU-UAUU-AUUU--A-GUGA----------
-----------A-----------UUUA----------------------------------------UA-GC
U-A--------------------AA-U-GA-AGCAAACCAGAGGAACUGAAACAUCUUAGUACUCU--GAGG
AAAAGAAAUCAA--UUGAGAUUCUCGUA--GU-AG-UGGCGAGCGAACCGGGA-U-AA-GCCCA--------
------------------------------------------------------------------------
------GUA-GCA-------AAAUAUUA---------------------UA--AUU-----------AG---
U-------AUAUU-AAGAG------AAU-AA-UUUG------------------------------------
-------GAA-AA-AUUAG-------CAAUAUAGGGUGAUAGCCCGUA----------UCUGAA-----AUA
U------A--UUAGU----------UUGU-----AA--GC-UC------GAAGAGUAGGACGGGACACGUGA
UAUCUUGUCUGAA-CAUAGGGGG-ACC--AUCCUCUAAGGCUAAAUA--CUC-UUGAC-UGACCGAUAGU-G
AAC-U-AGUACCGUGAGGGAAAG-GCGAAA-AGAAC-CCCGGUUAGGGG-A-GUGAAAUAGAACCUGAAACC
G-UAUACGUACAA-GCA-GU-AGGA---GCGA-A---UA---------------------------------
------------------------------------------------------------------------
------------------------------------------------------------------------
------------------------------------------------------------------------
------------------------------------------------------------------------
------------------------------------------------------------------------
------------------------------------------------------------CA--GUGUGAU-
----U-----------AG------------------------------------------------------
------------------------------------------------------------------------
--------------------------------------------------------------U--U-GU---
------------------------------------------------------------------------
------------------------------------------------------------------------
--AUA-------------------------------------------------------------------
------------------------------------------------------------------------
------------------------------------------------------------------------
------------------------------------------------------------------------
------------------------------------------------A-----------------------
------------------------------------------------------------------------
------------------------------------------------------------------------
------------------------------------------------------------------------
------------------------------------------------------------------------
------------------------------------------------------------------------
-------------------AUGUAUAAA------------------UAUU--C---GUGACUGCGUACCUUU
UGUAUAAUGGGUCAGCGACUUAU-A-UUCUGUAGCAA-GGUUAACU-------------GAUUA--------
---A--GGGA--GCCGUAGGGAAAC-C---------------------------------------------
```

```
------------------------------------------------------------------------
------------------------------------------------------------------------
---------------------------------------------------------------------GAG
UCU------------------------------------UAAC--------------------------UG-
G-C-GU----------------------UAAGUUGCAGGAUAUA-GACCCGAAACC-CGGUGAUCUAACCAU
GAACAGGUUGAAGAUUGG-GUAAUA-CUAAUUGGAGGACCGA--ACCGACUA-AUGUUGAAAAAUUA-GCGG
AUGAUUUGUGGUUAGGGGUGAAAGACCAAUCAAACCGGG-AGAUAGCUGGUUCUCCUCGAAA-GCUAUUUAG
GUAGCGCCUUGU-AAA--------------------------------------------------------
----------------------UU-----CA-UCUUUGGGGGUAGAGC-ACUGUUU-CGA-UUA--GGGGUU
C---AAACC--GAGCUACCAACUCGAUGCAAACUACGAAUACCAAA--------------------------
------------------------------------------------------------------------
------------------GAAU-GUUA--------UUACUGG-AGACACA-C-AGCG-G-GUGCUAACAUUC
GU-U-GUGGAAAG-GGAAAC---AACCCAGAUCGCUA-GCUAAGGUCCCUAAGU--UA-UGA-UU-AA---G
-U-GGG-------AAACGAUGUGAGAAGGCAUAAACAGCCAGGAUGUUGGCUUAGAAGCAGCCA-UCAUUUA
AAGAAAGCGUAAUAGCUCACUGGUC-----GAGUCGGCUUGCGCGGAAGAUGUAAC-GGGGCU-A-AAUCAU
A----------CACCGAA-GCUGCGACAAUAGAA--CAU---------------------------------
------------------------------------------------------------------------
----UUA-----------------------------------------------------------------
-------------------------------GUGUUU-UGUU-GGGUAGAGGAGC-GUUCUGU---------
-----------AAG-CCUGA-GAAGAUA--U-AUUGUGAGAU--A-UAUUGGAGGU-A-UCA-GAAGUGCGA
AUGC-UGACAU-AAGUA-ACGAUA---AAGC-AGGUGA--A---------------AAACCU-GCU-CGCCG
A-AAAACU-AAGGUUUCC-UGUCCAACG-UUAAUCGGGGCAGGGUAAGUCGAC-CC-CUAAGGUGAGGCCA-
A-A-CGGC-----GUA--------------------------------------------------------
----------------------------------------GCU-AA-U--GGGA-AACAGGUU-AAUA-UUC
CUGUA----------------------------------CUGGA----UGUA--------A-UU--------
------------------------------------------------------------------------
----------------------------------------------------------GCGAU-GGGGUAAC
GGAGAA--------------AGCUAG-GUUCGCCGA-GG-A----UCGGU------------AGUCUCGGU-
UUAAGC-AAG-UA-----G-A-------------------UG-----AG------------------AA-GA
UAUAGG--------------------AAAAU-CCGAUCUUCU------UU-ACAUUA----AGA-UG-U-G-
AUG--AC---------------------------------------------------------------GA
-----GCUGC--UACGGUAGCGAAG-G-AAU-U-GAUGCUAUGCUUC-C-AAGAAA-AACCU-CUAAGCUU-
-------------------------------------------------------CAGAUUA-UA-UCCAAU
CGUACCGUAAACCGAC------------------------------ACAGGUA--GUUAGGUA-GAGAAUAC
UAAGGC-G-CU-UGAG-AGAACUCG-G-GUGAAGGAACUAGGC-AAAAUAGUGCCGUAACUUUGG-GAGAAG
GCACG-CUAA-C-AUGUAAGUGAAG----GGA------------GUAC------------------------
------------------------------------------------------------------------
------------------------------------------------------------------------
------------------------------------------------------------------------
------------------------------------------------------------------------
------------------------------------------------------------------------
------------------------------------------------------------------------
------------------------------------------------------------------------
------------------------------------------------------------------------
------------------------------------------------------------------------
```

```
------------------------------------------------------------------------
------------------------------------------------------------------------
------------------------------------------------------------------------
------------------------------------------------------------------------
--------------------------------------------------------UUCU-GGA--GCUGAA
GUU--AGUCGAAGAUACCAGCUGGCUGCAA-----CUGUUUAUUAAAAACACAGCACUGUGCAAACACG---
-AA-A-------------GUGGAAGU-AUACAGUGUGA-CGCCUGCCCGGUGCCG-G-AAGG-UUAAUUGAG
AUGGUUA-UCUUU--GCAA--AAAGAGAAGCUUUUGAUCGAAGCCU--CGGUAAAC-GGCGGCCGUAACUAU
AACGGUCCUAAGGUAGCGAAAUUCCUUGUCGGG-UAAGUUCCGACCUGCACGAAUGGCGUAAUGAUG--GUC
AGACUGUCUCCACC----CGA-GACUCAGUGAAAUU--GAA---UUCGCCGUGAAGAUGCGGCGUACCCGCG
GCAAGACGGAAAGA-CCCCGUGAACC-UUUACU--AUAGCUU-GAUACUGAAUGUUGGAUUUUAAUGUGUAG
GAU-AGGUGGGAGGCUAUG----------AAAUAAA-GAC----------------------------GCUA
--------------------------------------GU-UUUUAUGGAG-C--C-GU-CCUUGAAAUA-C
CACC-CUUUGAUGUUUAAUAUUCUAACUUUAG--UCCG-U-UA-U-C-------------------------
------------------------------------------------------------------------
------------------------------------------------------------------------
------------------------------------------------------------------------
------------------------------------------------------------------------
------------------------------------------------------CG----GA----------
------------------------------------------------UUAA-----GGACA-GUGUCUGGU
GGGUAGUUUGACUGGGGCGGU-CUCCUCCUA-AAGAGUAAC-GGAGGAGCACA-AAGGUUAGCUAAUUACGG
UUGGAAAUCGUAAGGU-UAGUG-CAAAGGCAU-AAGCUAGCUUAACU-GCGA--------------------
--------------GAGUGA---CGAUUCGAGCAGAUGCGAAA-GCAGGUCUUAGUGAUCCGGU--------
---GG-U-----CCUGU-AUG--------------------GA-AAGGCCAUCGCUCAACGG-AUAAAAGGU
ACUCCGGGG-AUAACAGGCUAAUACCGCCCAAGAGUUCGUAUCGACGGCGGUGUUUGGCACCUCGAUGUCGG
CUCAUCACAUCCUGGGGCUGCA-GCAGGUCCCAAGGGUAUGGCUGUUCGCCAUUUAAAGUGGUACGCGAGCU
GGGUUUAGAACGUCGUGAGACAGUUCGGUCCCUAUCUGCC--------------------G-UGG-------
-------------------------------GCGU--UGGAAGAUUGAGAGG-GGC-UGCUCCUAGUACGAG
AGGAC-CGGAGUGGACGUACCUA-UGGU-G-UUCGGGUUGUCAUGCCAAUGGCAU-U-GCCCGGUAGCUAUG
UACGGAAAAGAUAACCGCUGAAAGCA-UC--UAAGCGGGAAACUUGCCUCAA-G-AUG--AGUCUUCC-CU-
--GAGGA-----------------------------------------------------------------
--------------------------UUAUUAAUUACAA---------------------------------
------------------------------------------------------------------------
------------------------------------------------------------------------
------------------------------------------------------------------------
------------------------------------------------------------------------
------------------------------------------------------------------------
------------------------------------------------------------------------
-------------------------------------------------UCCUCCU--AA-AGG------GC
CGUUAAAGACUAUGACGU-UGA-UAGGUC-GGG-UGUGUAAG-CAU-U----------------GU------
----GA-----------GAUGUUUAGCUA--A-CCGA-UACUAAUG---ACCCGUGAGAACUUAACUU----
------------------------------------------------------------------------
------------------------------------------------------------------------
------------------------------------------------------------------------
------------------------------------C------------
```

```
>LSUb_71421_HAEIN/1-2897
------------------------UUAAGUG----AC--UAAGCGUACAA-GGUGGAUG-C-CUUGGC-AAU
CAG---AGGCGAAGAAGGACGUGC--UAAUCUGCGAAAAGCUUGGAUGAGUCGAUAAGAGGCGU------UU
AAU-CCA-AGAUAUCCGAAUGGGGAAAC--------CC--------AG----UAGA-U-G-----------A
A-G-------------A---A----U----C---UAC----U-A-UCAA-CAA-----GUGA----------
-----------A-----------UUCA----------------------------------------UA-GC
U-U--------------------GU-U-GA-GGCAAACCGGGAGAACUGAAACAUCUAAGUACCCC--GAGG
AAAAGAAAUCAA--CCGAGAUUUCGUCA--GU-AG-CGGCGAGCGAAAGCGAA-A-GA-GCCAG--------
------------------------------------------------------------------------
----------UAA-------G--U-GA----------------------UA--GCA-----------AU---
A-------UAGUG-AGGAG------AAU-GU-GUUG------------------------------------
-------GGA-AG-CACAA-------UCAAAGAGGGUGAUAAUCCCGUA---------UCUAAAA----ACC
A------U--AUUGU-----------GGUACUAAGC--UA-AC------GAGAAGUAGGGCGGGACACGUGA
UAUCCUGUUUGAA-GAAGGGGGG-CCC--AUCCUCCAAGGCUAAAUA--CUC-CUGAU-UGACCGAUAGU-G
AAC-C-AGUACUGUGAAGGAAAG-GCGAAA-AGAAC-CCCGGUGAGGGG-A-GUGAAAUAGAACCUGAAACC
U-UGUACGUACAA-GCA-GU-GGGA---GCGA-G---G----------------------------------
------------------------------------------------------------------------
------------------------------------------------------------------------
------------------------------------------------------------------------
------------------------------------------------------------------------
------------------------------------------------------------------------
------------------------------------------------------------------------
------------------------------------------------------------------------
------------------------------------------------------------------------
--------------------------------------------------------------G--C-AA---
------------------------------------------------------------------------
------------------------------------------------------------------------
------------------------------------------------------------------------
------------------------------------------------------------------------
------------------------------------------------------------------------
------------------------------------------------------------------------
------------------------------------------------------------------------
------------------------------------------------------------------------
------------------------------------------------------------------------
------------------------------------------------------------------------
------------------------------------------------------------------------
------------------------------------------------------------------------
-----------------------------------------------CCU--U---GUGACUGCGUACCUUU
UGUAUAAUGGGUCAGCGACUUAU-A-UUUUGUAGCGA-GGUUAACC-------------GAAUA--------
---G--GGGA--GCCGAAGGGAAAC-C---------------------------------------------
------------------------------------------------------------------------
------------------------------------------------------------------------
---------------------------------------------------------------------GAG
UCU------------------------------------UAACU-------------------------GG-
G-C-GA----------------------AUAGUUGCAAGGUAUA-GACCCGAAACC-CGGUGAUCUAGCCAU
```

```
GGGCAGGUUGAAGGUUGG-GUAACA-CUAACUGGAGGACCGA--ACCGACUA-AUGUUGAAAAAUUA-GCGG
AUGACUUGUGGCUGGGGGUGAAAGGCCAAUCAAACCGGG-AGAUAGCUGGUUCUCCCCGAAA-UCUAUUUAG
GUAGAGCCUUGA-GGU--------------------------------------------------------
----------------------GA-----CA-CCUUUGGGGGUAGAGC-ACUGUUU-CGG-CUA--GGGGGC
C---AUCCC--GGCUUACCAACCCGAUGCAAACUACGAAUACCAAA--------------------------
------------------------------------------------------------------------
------------------GAGU-G-AU--------ACUCAGG-AGACACA-C-GGCG-G-GUGCUAACGUCC
GU-C-GUGGAGAG-GGAAAC---AACCCAGACCGCCA-GCUAAGGUCCCCAAGU--CU-AUA-UU-AA---G
-U-GGG-------AAACGAAGUGGGAAGGCUUAGACAGCUAGGAUGUUGGCUUAGAAGCAGCCA-UCAUUUA
AAGAAAGCGUAAUAGCUCACUAGUC-----GAGUCGGCCUGCGCGGAAGAUGUAAC-GGGGCUGA-AAUAUA
G----------CACCGAA-GCUGCGGCAUCAGAA--U-----------------------------------
------------------------------------------------------------------------
------------------------------------------------------------------------
-------------------------------UUAUUC-UGUU-GGGUAGGGGAGC-GUUGUGU---------
-----------AAG-CGGAA-GAAGGUU--C-AUCGAGAGGU--G-GGCUGGACGU-A-UCA-CAAGUGCGA
AUGC-UGACAU-AAGUA-ACGAUA---AAAC-GGGUGA--A---------------AAACCC-GUU-CGCCG
G-AAGACC-AAGGGUUCC-UGUCCAACG-UUAAUCGGGGCAGGGUGAGUCGGC-UC-CUAAGGCGAGGCUG-
A-A-AAGC-----GUA--------------------------------------------------------
----------------------------------------GUC-GA-U--GGGA-AACAGGUU-AAUA-UUC
CUGUA----------------------------------CUUGG----UAAA--------G-CU--------
------------------------------------------------------------------------
----------------------------------------------------------GCGAU-GUGGGGAC
GGAGUA--------------GGUUAG-GUUAUCGCA-CU-G----UUGGA------------UAUGUGCGU-
UUAAGU-UGG-UA-----G-G-------------------UG-----GG------------------AA-GU
U-UAGG--------------------CAAAU-CCGGACUUCC------UUAACACAG----AGA-GA-U-G-
AUG--AC---------------------------------------------------------------GA
-----GGCUC--UACGGAGCUGAAG-U-AAC-U-GAUACCACACUUC-C-AGGAAA-AGCCA-CUAAGCGA-
-------------------------------------------------------AAGGCUU-UA-CUAAAC
CGUACUGAAAACCGAC------------------------------ACAGGUG--GUCAGGUA-GAGAAUAC
UCAGGC-G-CU-UGAG-AGAACUCG-G-GUGAAGGAACUAGGC-AAAAUAGCACCGUAACUUCGG-GAGAAG
GUGCG-CCGG-C-GU-AGAUUGUAA----GGG-----------CUAGC------------------------
------------------------------------------------------------------------
------------------------------------------------------------------------
------------------------------------------------------------------------
------------------------------------------------------------------------
------------------------------------------------------------------------
------------------------------------------------------------------------
------------------------------------------------------------------------
------------------------------------------------------------------------
------------------------------------------------------------------------
------------------------------------------------------------------------
------------------------------------------------------------------------
------------------------------------------------------------------------
------------------------------------------------------------------------
--------------------------------------------------------CCCU-GAA--GGUUGA
```

```
ACC--GGUCGAAGAUACCAGCUGGCUGCAA-----CUGUUUAUUAAAAACACAGCACUCUGCAAACACG---
-AA-A-------------GUGGACGU-AUAGGGUGUGA-UGCCUGCCCGGUGCUG-G-AAGG-UUAAUUGAU
GGUGUCA-UC-----GAAA-----GAGAAGCACCUGAUCGAAGCCC--CAGUAAAC-GGCGGCCGUAACUAU
AACGGUCCUAAGGUAGCGAAAUUCCUUGUCGGG-UAAGUUCCGACCUGCACGAAUGGCAUAAUGAUG--GCC
AGGCUGUCUCCACC----CGA-GACUCAGUGAAAUU--GAA---AUCGCCGUGAAGAUGCGGUGUACCCGCG
GCUAGACGGAAAGA-CCCCGUGAACC-UUUACU--AUAGCUU-GACACUGAACAUUGAAUUUUGAUGUGUAG
GAU-AGGUGGGAGCCUUUG----------AAGCAGU-GAC----------------------------GCCA
--------------------------------------GU-CAUUGUGGAG-G--C-GA-CCUUGAAAUA-C
CACC-CUUUAACGUUUGAUGUUCUAACGAAGA--UGAC-G-AA-A-C-------------------------
------------------------------------------------------------------------
------------------------------------------------------------------------
------------------------------------------------------------------------
------------------------------------------------------------------------
------------------------------------------------------GU----GG----------
------------------------------------------------UCUC-----GGACA-GUGUCUGGU
GGGUAGUUUGACUGGGGCGGU-CUCCUCCCA-AAGCGUAAC-GGAGGAGCACG-AAGGUUUGCUAAUCACGG
UCGGACAUCGUGAGGU-UAGUG-CAAUGGUAU-AAGCAAGCUUAACU-GCGA--------------------
--------------GACAGA---CAAGUCGAGCAGGUACGAAA-GUAGGUCAUAGUGAUCCGGU--------
---GG-U-----UCUGA-AUG--------------------GA-AGGGCCAUCGCUCAACGG-AUAAAAGGU
ACUCCGGGG-AUAACAGGCUGAUACCGCCCAAGAGUUCAUAUCGACGGCGGUGUUUGGCACCUCGAUGUCGG
CUCAUCACAUCCUGGGGCUGAA-GUAGGUCCCAAGGGUAUGGCUGUUCGCCAUUUAAAGUGGUACGCGAGCU
GGGUUUAGAACGUCGUGAGACAGUUCGGUCCCUAUCUGCC--------------------G-UGG-------
-------------------------------GCGU--AGGAUGAUUGAUUGG-GGC-UGCUCCUAGUACGAG
AGGAC-CGGAGUGGACGCAUCAC-UGGU-G-UUCCGGUUGUGUCGCCAGACGCAU-U-GCCGGGUAGCUAAA
UGCGGAAGAGAUAAGUGCUGAAAGCA-UC--UAAGCACGAAACUUGCCAAGA-G-AUG--AGUCAUCC-CU-
--GACUU-----------------------------------------------------------------
------------------------------------UAA---------------------------------
------------------------------------------------------------------------
------------------------------------------------------------------------
------------------------------------------------------------------------
------------------------------------------------------------------------
------------------------------------------------------------------------
------------------------------------------------------------------------
-------------------------------------------------GUCAG----UA-AGG------GU
UGUUGUAGACUACGACGU-AGA-UAGGUU-GGG-UGUGUAAG-UGA-U----------------GU------
----GA-----------GUCAUUGAGCUA--A-CCAA-UACUAAUU---GCCCGAGAG-GCUUAACU-----
------------------------------------------------------------------------
------------------------------------------------------------------------
------------------------------------------------------------------------
-------------------------------------------------
>LSUb_190486_XANAC/1-2881
---------------------G-GUCAAGCG----AA--UAAGCGCACAC-GGUGGAUG-C-CUAGGC-GGU
CAG---AGGCGAUGAAGGACGUGG--UAGCCUGCGAAAAGUGUCGGGGAGCUGGCAACAAGCUU------UG
AUC-CGG-CAAUAUCCGAAUGGGGAAAC--------CC--------AC----UGC---------------UU
C-G---------------------------G---CAG----U-A-UCCU-GCA-----GUGA----------
```

```
-----------A-----------UUCA----------------------------------------UAG-C
U-G--------------------CU-G-GA-AGCGAACCCGGUGAACUGAAAUAUCUAAGUAACCG--GAGG
AAAAGAAAUCAA--CCGAGAUUCCCUAA--GU-AG-UGACGAGCGAACGGGGA-G-CA-GCCCU--------
------------------------------------------------------------------------
----------UAA-------G----CU--------------------------GGA-----------AU---
G-------GCUUU-AGAAA------AAC-AA-UCUG------------------------------------
-------GAA-AG-AUUGG-------CCAUAGAAGGUGAUAGCCCUGUA---------UUUAAAA----GGG
C------C--ACUCC-----------AG----------UG-AA------GACGAGUAGGGCGGGGCACGUGA
AACCCUGUCUGAA-UAUGGGGGG-ACC--AUCCUCCAAGGCUAAAUA--CUC-CUGAC-CGACCGAUAGU-G
AAC-C-AGUACCGUGAGGGAAAG-GCGAAA-AGAAC-CCCGGAGAGGGG-A-GUGAAAUAGAUCCUGAAACC
G-UGUGCGUACAA-GCA-GU-AGGA---GCUC----------------------------------------
------------------------------------------------------------------------
------------------------------------------------------------------------
------------------------------------------------------------------------
------------------------------------------------------------------------
------------------------------------------------------------------------
------------------------------------------------------------------------
------------------------------------------------------------------------
------------------------------------------------------------------------
--------------------------------------------------------------G--C-AA---
------------------------------------------------------------------------
------------------------------------------------------------------------
------------------------------------------------------------------------
------------------------------------------------------------------------
------------------------------------------------------------------------
------------------------------------------------------------------------
------------------------------------------------------------------------
------------------------------------------------------------------------
------------------------------------------------------------------------
------------------------------------------------------------------------
------------------------------------------------------------------------
------------------------------------------------------------------------
-------------------------------------------------G--A---GUGACUGCGUACCUUU
UGUAUAAUGGGUCAGCGACUUAC-U-GUUCGUGGCAA-GCUUAACC-------------GUAUA--------
---G--GGGA--GGCGAAGGGAAAC-C---------------------------------------------
------------------------------------------------------------------------
------------------------------------------------------------------------
---------------------------------------------------------------------GAG
UCU------------------------------------GAUAA-------------------------GG-
G-C-GC----------------------AUAGUCGCGGCAGUA-GACCCGAAACC-GGGUGAUCUAGUCAU
GGCCAGGGUGAAGGUGCC-GUAACA-GGUACUGGAGGCCCGA--ACCCACGU-CUGUUGCAAAAGAC-GGGG
AUGAGCUGUGAUUAGGAGUGAAAAGCUAAUCGAACCCGG-AGAUAGCUGGUUCUCCUCGAAA-GCUAUUUAG
GUAGCGCCUCAU-AUG--------------------------------------------------------
----------------------UA-----UC-CUCUCGGGGGUAGAGC-ACUGUUA-UGG-CUA--GGGGGU
C---AUCGC--GACUUACCAAACCAUUGCAAACUCCGAAUACCGAG--------------------------
```

```
------------------------------------------------------------------------
------------------ACGG-A-CU--------GUAUGGG-AGACACA-C-GGCG-G-GUGCUAACGUCC
GU-C-GUGAAAAG-GGAAAC---AACCCAGACCCACA-GCUAAGGUCCCAAAUU--CA-CUG-CU-AA---G
-U-GGA-------AAACGAUGUGGAAAGGCAUAGACAGCCAGGAGGUUGGCUUAGAAGCAGCCA-CCCUUUA
AAGAAAGCGUAAUAGCUCACUGGUC-----GAGUCGGUCUGCGCGGAAGAUUUAAC-GGGGCU-A-AGCAGU
G----------AACCGAA-GCUUGGGGUGCAUAA--A-----------------------------------
------------------------------------------------------------------------
-----ACGU---------------------------------------------------------------
-------------------------------GUUUUA-UGCG-CGGUAGAGGAGC-GUUCCGU---------
-----------AAG-CCUGC-GAAGGUG--G-AUUGAGAAGU--C-UGCUGGAGGU-A-UCG-GAAGUGCGA
AUGC-UGACAU-GAGUA-ACGAUA---AUGC-GGGUGA--A---------------AAGCCC-GCA-CGCCG
A-AAGCCC-AAGGUUUCCUUGCGCAACG-UUAAUCGACGCAGGGUUAGUCGGU-CC-CUAAGGCGAGGGCG-
A-A-AGCC-----GUA--------------------------------------------------------
----------------------------------------GUC-GA-U--GGGA-AGCAGGUU-AAUA-UUC
CUGCA----------------------------------CCUCG----CGUG--------A-GU--------
------------------------------------------------------------------------
----------------------------------------------------------GCGAU-GGAGGGAC
GGAGAA--------------GGUUAG-GUGUACCGG-GC-G----UUGGU------------UGUCCCGGG-
GAAAGG-CGG-UA-----G-G-------------------UU-----UG------------------GA-UC
U-UUGG--------------------CAAAU-CCGGGAUCCU------UUAAGACCG----AGC-AC-C-G-
AG---AC---------------------------------------------------------------GA
-----GCCUU-----UUUGGCGAAG-U-CAC-U-GAUACCACGCUUC-C-AGGAAA-AGCUC-CUAAGCUU-
--------------------------------------------------------CAGCUC-AC-GAAGAC
CGUACCGUAAACCGAC------------------------------ACAGGUG--GGUAGGAU-GAGAAUUC
UCAGGC-G-CU-UGAG-AGAACUCG-G-GUGAAGGAACUAGGC-AACAUGGCACCGUAACUUCGG-GAGAAG
GUGCA-CCCU-U-UU--UGGUG--G----CUC------------AUGC------------------------
------------------------------------------------------------------------
------------------------------------------------------------------------
------------------------------------------------------------------------
------------------------------------------------------------------------
------------------------------------------------------------------------
------------------------------------------------------------------------
------------------------------------------------------------------------
------------------------------------------------------------------------
------------------------------------------------------------------------
------------------------------------------------------------------------
------------------------------------------------------------------------
------------------------------------------------------------------------
------------------------------------------------------------------------
----------------------------------------------------------GAGCUAUAGCUGAA
GAG--GGUCGCAGAAACCAGGCCGCUGCGA-----CUGUUUAUCAAAAACACAGCACUCUGCAAACACG---
-AA-A-------------GUGGACGU-AUAGGGUGUGA-CGCCUGCCCGGUGCUG-G-AAGG-UUAAUUGAU
GGGGUCA-GCC----GCAA----GGCGAAGCUCUUGAUCGAAGCCC--CAGUAAAC-GGCGGCCGUAACUAU
AACGGUCCUAAGGUAGCGAAAUUCCUUGUCGGG-UAAGUUCCGACCUGCACGAAUGGCGUAACGACA--GCG
GCGCUGUCUCCACC----CGA-GACUCAGUGAAAUU--GAA---AUCGCUGUGAAGAUGCAGCGUUCCCGUG
```

```
GCAAGACGGAAAGA-CCCCGUGAACC-UUUACU--AUAGCUU-UACACUGAACGUUGAGUUCGUCUGUGUAG
GAU-AGGUGGGAGGCUAUG----------AAACUGU-GGC----------------------------GCUA
--------------------------------------GC-UGCAGUGGAG-C--C-AU-CCUUGAAAUA-C
CACC-CUGUCGUGCUUGACGUUCUAACCUAGG--UCCG-U-UA-U-C-------------------------
------------------------------------------------------------------------
------------------------------------------------------------------------
------------------------------------------------------------------------
------------------------------------------------------------------------
------------------------------------------------------CG----GA----------
------------------------------------------------UCAG-----GGACC-GUGUAUGGU
GGGUAGUUUGACUGGGGCGGU-CUCCUCCUA-AAGAGUAAC-GGAGGAGCACG-AAGGUACGCUCAGCGCGG
UCGGACAUCGCGCACU-GUGUG-CAAAGGCAU-AAGCGUGCUUGACU-GCAA--------------------
--------------GAUCGA---CGGAUCAAGCAGGUACGAAA-GUAGGUCUUAGUGAUCCGGU--------
---GG-U-----UCUGU-AUG--------------------GA-AGGGCCAUCGCUCAACGG-AUAAAAGGU
ACUCCGGGG-AUAACAGGCUGAUACCGCCCAAGAGUUCAUAUCGACGGCGGUGUUUGGCACCUCGAUGUCGG
CUCAUCACAUCCUGGGGCUGUA-GUCGGUCCCAAGGGUAUGGCUGUUCGCCAUUUAAAGUGGUACGCGAGCU
GGGUUCAGAACGUCGUGAGACAGUUCGGUCCCUAUCUGCC--------------------A-UGG-------
-------------------------------GCGU--UGGAAGUUUGAGAGG-GGC-UGCUCCUAGUACGAG
AGGAC-CGGAGUGGACGAACCUC-UGGU-G-UUCCGGUUGUCACGCCAGUGGCAU-U-GCCGGGUAGCUAUG
UUCGGAAGCGAUAACCGCUGAAAGCA-UC--UAAGCGGGAAGCGCGCCUCAA-G-AUG--AGACUUCC-CG-
--GGGCA-----------------------------------------------------------------
------------------------------------CAA---------------------------------
------------------------------------------------------------------------
------------------------------------------------------------------------
------------------------------------------------------------------------
------------------------------------------------------------------------
------------------------------------------------------------------------
------------------------------------------------------------------------
-------------------------------------------------GCCCCCU--AA-AGG------AA
CCAUGUAGACUACGUGGU-UGA-UAGGUC-AGG-UGUGUAAG-UAC-A----------------GC------
----AA-----------UGUAUUGAGCUA--A-CUGA-UACUAAUG---AUCCGUGCG-GCUUGACCA----
------------------------------------------------------------------------
------------------------------------------------------------------------
------------------------------------------------------------------------
------------------------------------U------------
>LSUb_297245_LEGPL/1-2855
----------------------------------------AAGCGCAAAC-GGUGGAUG-C-CUUGGC-AGU
AAG---AGGCGAAGAAGGACGUGG--AAUCCUGCGAAAAGCUAUGGGGAGCUGGAAACGAGCGU------UG
AGC-CAU-AGAUGUCCGAAUGGGGGAAC--------CC--------GG----CUGC--------------CG
A-U----------------------G----C---GGU----C-A-UUUG-CAU-----UUGA----------
-----------A-----------UACA----------------------------------------UA-GG
U----------------------GC-A-AA-GGCGAACUCGGGGAACUGAAACAUCUAAGUACCCG--AAGG
AAAAGAAAUCAA--GAGAGAUUCUCCAA--GU-AG-CGGCGAGCGAACGGGGA-G-GA-GCCUG--------
------------------------------------------------------------------------
----------GCG-------U-GA-UU----------------------UA--UUA-----------UU---
```

```
G-------AACUG-AGUAG------AAC-AA-UUUG-------------------------------------
-------GGA-AA-GUUGG-------CGAUAGAGGGUGAAAGCCCCGUA---------UACGAAA----GUU
U------G--AUGAA-----------------------CU-CA------CGCGAGUAGGCCGGGACACGUGA
AAUCCUGGUUGAA-GAUGGGUGG-ACC--AUCAUCCAAGGCUAAAUA--CUA-CUUAC-UGACCGAUAGU-G
AAC-C-AGUACCGUGAGGGAAAG-GUGAAA-AGAAC-CCCGGAGAGGGG-A-GUGAAAUAGAAUCUGAAACC
G-UUUGCGUACAA-GCA-GU-GGGA---GCAU-G--------------------------------------
------------------------------------------------------------------------
------------------------------------------------------------------------
------------------------------------------------------------------------
------------------------------------------------------------------------
------------------------------------------------------------------------
------------------------------------------------------------------------
------------------------------------------------------------------------
------------------------------------------------------------------------
--------------------------------------------------------------G--U-UU---
------------------------------------------------------------------------
------------------------------------------------------------------------
------------------------------------------------------------------------
------------------------------------------------------------------------
------------------------------------------------------------------------
------------------------------------------------------------------------
------------------------------------------------------------------------
------------------------------------------------------------------------
------------------------------------------------------------------------
------------------------------------------------------------------------
------------------------------------------------------------------------
------------------------------------------------------------------------
------------------------------------------------UG--U---GUGACUGCGUACCUUU
UGUAUAAUGGGUCAGCGAGUUAC-U-UUCAGUGGCGA-GGUUAACU-------------GAAUA--------
---A--GGGA--GCCGUAGAGAAAU-C---------------------------------------------
------------------------------------------------------------------------
------------------------------------------------------------------------
---------------------------------------------------------------------GAG
UCU------------------------------------GAAUA-------------------------GG-
G-C-GA-----------------------UAGUCGCUGGGAGUA-GACCCGAAACC-GGGCGAUCUAGCCAU
GUGCAGGAUGAAGGUUGG-GUAACA-CCAACUGGAGGUCCGA--ACCGGGUA-AUGUUGAAAAAUUA-UCGG
AUGACGUGUGGCUAGGAGUGAAAGGCUAAUCAAGCCCGG-AGAUAGCUGGUUCUCCCCGAAA-GCUAUUUAG
GUAGCGCCUCGU-GAA--------------------------------------------------------
-----------------------U-----GA-UUACUGGGGGUAGAGC-ACUGUUU-CGG-CUA--GGGGGC
U---GUCAU--GGCUUACCAAACCGAUGCAAACUCCGAAUACCGGC--------------------------
------------------------------------------------------------------------
------------------UAAU-UGAA--------UCACGGG-AGACACA-C-GGCG-G-GUGCUAACGUCC
GU-C-GUGGAGAG-GGAAAC---AACCCAGACCGCCA-GCUAAGGUCCCCAAGU--AC-UAG-UU-AA---G
-U-GGG-------AAACGAUGUGGGAAGGCAUAGACAGCCAGGAGGUUGGCUUAGAAGCAGCCA-CCCUUUA
AAGAAAGCGUAAUAGCUCACUGGUC-----GAGUCGGCCUGCGCGGAAGAUGUAAC-GGGGCUAA-AACUAG
```

```
U----------CACCGAA-GCUGCGGAUGUGCGC--G-----------------------------------
------------------------------------------------------------------------
------------------------------------------------------------------------
-------------------------------AAGGCG-CACG-UGGUAGGGGAGC-GUUCUGU---------
-----------AGG-CUGAU-GAAGGUG--C-AUUGAGAAGU--G-UGCUGGAGGU-A-UCA-GAAGUGCGA
AUGC-UGACAU-GAGUA-ACGAUA---AUGU-GGGUGA--A---------------AAGCCC-ACA-CGCCG
G-AAGUCC-CAGGUUUCC-UGCACGACG-UUAAUCGGAGCAGGGUGAGUCGGC-CC-CUAAGGCGAGGCUG-
A-A-GAGC-----GUA--------------------------------------------------------
----------------------------------------GUC-GA-U--GGGA-ACCAGGUU-AAUA-UUC
CUGGA----------------------------------CUUUU----UAUA--------A-----------
------------------------------------------------------------------------
----------------------------------------------------------GUGAA-GUGGGGAC
GAAGAA--------------GGCUAG-GUGAGCCAG-GC-G----UUGGU------------UGUCCUGGU-
ACUUGC-AUG-UA-----G-G-------------------GG-----GG------------------AA-GA
C-UUGG--------------------CAAAU-CCGGUUUUCC------AUAACUCUG----AGG-UG-C-G-
AAG--UG---------------------------------------------------------------GU
-----UCUGA--CCUUUUGGAGAAG-U-CAU-U-GAUGCCCGGCUUC-C-AGGAAA-AGCUG-CUAGCCAU-
---------------------------------------------------------AACUU-AU-AGGAAC
CGUACCGCAAACCGAC------------------------------ACAGGUG--GACAGGUA-GAGAAUAC
UAAGGC-G-CU-UGAG-AGAACUCG-G-GUGAAGGAACUAGGC-AAAAUGGUACCGUAACUUCGG-GAGAAG
GUACG-CCCU-U-UC--UGGUGAUG----GGA-----------UUUAC------------------------
------------------------------------------------------------------------
------------------------------------------------------------------------
------------------------------------------------------------------------
------------------------------------------------------------------------
------------------------------------------------------------------------
------------------------------------------------------------------------
------------------------------------------------------------------------
------------------------------------------------------------------------
------------------------------------------------------------------------
------------------------------------------------------------------------
------------------------------------------------------------------------
------------------------------------------------------------------------
------------------------------------------------------------------------
--------------------------------------------------------UUUC-AGA--GCUGGA
GAG--GGCCGCAGAGACCAGGUGGCUGCGA-----CUGUUUAUUAAAAACACAGCACUCUGCAAAUUCG---
-UA-A-------------GAAGACGU-AUAGGGUGUGA-CGCCUGCCCGGUGCCG-G-AAGG-UUAAUUGAU
GGGGUUA-UC-----UUCG-----GAGAAGCUCUUGAUCGAAGCCC--CGGUAAAC-GGCGGCCGUAACUAU
AACGGUCCUAAGGUAGCGAAAUUCCUUGUCGGG-UAAGUUCCGACCUGCACGAAUGGCGUAACGAUG--GCC
ACACUGUCUCCACC----CGA-GACUCAGUGAAAUU--GAA---AUCGCUGUGAAGAUGCAGUGUACCCGCG
GCUAGACGGAAAGA-CCCCGUGAACC-UUUACU--AUAGUUU-UGCACUGGACUUUGAUGAUGACUGUGUAG
GAU-AGGUGGGAGGCUGUG----------AAGUGAG-GAC----------------------------GCUA
--------------------------------------GU-UCUCAUGGAG-C--C-GC-CCUUGAAAUA-C
CACC-CUGUUGUUAUUGAGGUUCUAACUUGGU--CCAG-U-AA-U-C-------------------------
------------------------------------------------------------------------
```

```
------------------------------------------------------------------------
------------------------------------------------------------------------
------------------------------------------------------------------------
------------------------------------------------------CU----GG----------
------------------------------------------------AUGA-----GGACA-GUGUAUGAU
GGGUAGUUUGACUGGGGCGGU-CUCCUCCCA-AAGAGUAAC-GGAGGAGCACA-AAGGUACCCUCGGUACGG
UCGGACAUCGUACCAA-GAGUG-UAAAGGCAA-AAGGGUGCUUGACU-GCGA--------------------
--------------GAGUGA---CGGCUCGAGCAGGAACGAAA-GUUGGUCUUAGUGAUCCGGU--------
---GG-U-----UCUGA-AUG--------------------GA-AGGGCCAUCGCUCAACGG-AUAAAAGGU
ACUCCGGGG-AUAACAGGCUGAUACCGCCCAAGAGUUCAUAUCGACGGCGGUGUUUGGCACCUCGAUGUCGG
CUCAUCACAUCCUGGGGCUGAA-GCAGGUCCCAAGGGUAUGGCUGUUCGCCAUUUAAAGUGGUACGCGAGCU
GGGUUUAGAACGUCGUGAGACAGUUCGGUCCCUAUCUGCC--------------------G-UGG-------
-------------------------------GCGU--AGGAAAAUUGAGAGG-AGC-UGCUCCUAGUACGAG
AGGAC-CGGAGUGGACGAACCUC-UGGU-G-CACCGGUUGUCACGCCAGUGGCAU-G-GCCGGGUAGCUAAG
UUCGGACGGGAUAACCGCUGAAAGCA-UC--UAAGCGGGAAGCCUCCCUCAA-G-AUG--AGUUUUCC-CA-
------------------------------------------------------------------------
------------------------------------------------------------------------
------------------------------------------------------------------------
------------------------------------------------------------------------
------------------------------------------------------------------------
------------------------------------------------------------------------
------------------------------------------------------------------------
------------------------------------------------------------------------
----------------------------------------------------------UG-AAG------CC
CGUUGAAGACUACGACGU-UGA-UAGGCA-AGG-UGUGGAAG-CGC-A----------------GU------
----AA-----------UGCGUGAAGCUA--A-CUUG-UACUAAUU---GGCUGAUUG-UCUUGACCA----
------------------------------------------------------------------------
------------------------------------------------------------------------
------------------------------------------------------------------------
------------------------------------U------------
>LSUb_62928_AZOSB/1-2885
---------------------G-AUCAAGCG----AC--UAAGUGCAUGU-GGUGGAUG-C-CUUGGC-GAU
CAC---AGGCGAUGAAGGACGUGC--AAGCCUGCGAAAAGCGGGGGGGAGCUGGCAAUGGAGCU-----UUG
AUC-CCC-CGAUAUCCGAAUGGGGAAAC--------CC--------GG----CCC---------------GC
A-A---------------------------G---GGU----C-A-UCCC-GUA-----CUGA----------
-----------A-----------UCCA----------------------------------------UA-GG
U-A--------------------CGUG-GA-AGCGAACGCGGCGAACUGAAACAUCUAAGUAGCCG--UAGG
AAAAGAAAUCAA--CCGAGAUUCCCCAA--GU-AG-UGGCGAGCGAACGGGGA-A-GA-GCCUG--------
------------------------------------------------------------------------
----------CAC-------G--A-CU----------------------AA--ACC-----------AU---
C-------AGCUU-AGCAA------AAC-GG-UCUG------------------------------------
-------GAA-AG-UCCGA-------CGAUACAGGGUGAUAGUCCGUA----------UGCGAAA----AGC
C------G--GUGGC--------GGGUCU---GAGC--GU-GC------GACAAGUAGGGCGGGACACGUGU
AAUCCUGUCUGAA-GAUGGGGGGG-ACC--AUCCUCCAAGGCUAAUA--CUC-GUGAU-CGACCGAUAGU-G
AAC-C-AGUACCGUGAGGGAAAG-GCGAAA-AGAAC-CCCGG-GAGGGG-A-GUGAAAUAGAUCCUGAAACC
```

```
G-CAUGCAUACAA-ACA-GU-GGGA---GCCU-G--------------------------------------
------------------------------------------------------------------------
------------------------------------------------------------------------
------------------------------------------------------------------------
------------------------------------------------------------------------
------------------------------------------------------------------------
------------------------------------------------------------------------
------------------------------------------------------------------------
------------------------------------------------------------------------
--------------------------------------------------------------G--A-AA---
------------------------------------------------------------------------
------------------------------------------------------------------------
------------------------------------------------------------------------
------------------------------------------------------------------------
------------------------------------------------------------------------
------------------------------------------------------------------------
------------------------------------------------------------------------
------------------------------------------------------------------------
------------------------------------------------------------------------
------------------------------------------------------------------------
------------------------------------------------------------------------
------------------------------------------------CG--G---GUGACUGCGUACCUUU
UGUAUAAUGGGUCAGCGACUUAC-G-UUCAGUGGCGA-GCUUAACC-------------GAAUA--------
---G--GGGA--GGCGUAGCGAAAG-C---------------------------------------------
------------------------------------------------------------------------
------------------------------------------------------------------------
---------------------------------------------------------------------GAG
UCU------------------------------------GAUAA-------------------------GG-
G-C-GA---------------------CAUAGUCGCUGGGCGUA-GACCCGAAACC-GGAUGAUCUAUCCAU
GGCCAGGAUGAAGGUGCG-GUAACA-CGCACUGGAGGUCCGA--ACCCACUA-GUGUUGAAAAACUA-GGGG
AUGAGCUGUGGAUAGGGGUGAAAGGCUAAACAAAUCCGG-AAAUAGCUGGUUCUCCCCGAAA-ACUAUUUAG
GUAGUGCGUCGU-ACG--------------------------------------------------------
----------------------GA-----CA-CUUGCGGGGGUAGAGC-ACUGUAA-UCGUUGG--GGGGGU
C---ACUGC--GAUCUACCCCGCGAUAGCAAACUCCGAAUACCGCA--------------------------
------------------------------------------------------------------------
------------------AAGU-G-AU--------AUACGGC-AGACAGA-C-AUCG-G-GUGCUAACGUCC
GG-U-GUCAAGAG-GGAAAC---AACCCAGACCGCCA-GCUAAGGUCCCAAAUG--CA-UGG-CU-AA---G
-U-GGC-------AAACGAGGUGGGAAGGCAUAGACAGCUAGGAGGUUGGCUUAGAAGCAGCCA-CCCUUUA
AAGAAAGCGUAAUAGCUCACUAGUC-----GAGUCGUCCUGCGCGGAAGAUGUAAC-GGGGCUCA-AGCCAU
G----------AACCGAA-GCUGCGGAUGUGUCU--U-----------------------------------
------------------------------------------------------------------------
------------------------------------------------------------------------
----------------------------------UGA-CACG-UGGUAGGGGAGC-GUUCCGU---------
-----------AAG-CCUGC-GAAGGUG--U-CUCGUAAGGG--A-UGCUGGAGGU-A-UCG-GAAGUGCGA
```

```
AUGC-UGACAU-GAGUA-GCGAUA---AAGG-GUGUGA--A---------------AAGCAC-CCU-CGCCG
A-AAGCCC-AAGGUUUCC-UGCGCAACG-UUCAUCGGCGCAGGGUGAGUCGGC-CC-CUAAGGCGAGGCAG-
A-A-AUGC-----GUA--------------------------------------------------------
----------------------------------------GUC-GA-U--GGGA-AACAGGUC-AAUA-UUC
CUGUA----------------------------------CCGGU----UCUA--------G-AU--------
------------------------------------------------------------------------
----------------------------------------------------------GCGAU-GGGGGGAC
GGAGAA--------------GGUUAG-GCCGGCCGG-GU-G----UUGGA------------CGUCCCGGU-
UUAAGC-GUG-UA-----G-G-------------------CG-----UG------------------CU-CU
C-UAGG--------------------CAAAU-CCGGAGAGCU---------GAGCUG----AGG-CG-U-G-
AUG--AC---------------------------------------------------------------GA
-----GCGAA--CUUGUUCGCGAAG-U-GGU-U-GAUACCCUGCUUC-C-AGGAAA-AGCCU-CUAAGCUU-
-------------------------------------------------------CAGUCUA-GA-AUCGAC
CGUACCGCAAACCGAC------------------------------ACAGGUG--GGCAGGUU-GAAAAUAC
CAAGGC-G-CU-UGAG-AGAACUCA-G-GAGAAGGAACUCGGC-AAAUUGAUACCGUAACUUCGG-GAGAAG
GUAUG-CCCC-A-UU-AGCUUGUAG----GAG-------------UAC------------------------
------------------------------------------------------------------------
------------------------------------------------------------------------
------------------------------------------------------------------------
------------------------------------------------------------------------
------------------------------------------------------------------------
------------------------------------------------------------------------
------------------------------------------------------------------------
------------------------------------------------------------------------
------------------------------------------------------------------------
------------------------------------------------------------------------
------------------------------------------------------------------------
------------------------------------------------------------------------
------------------------------------------------------------------------
--------------------------------------------------------AUCC-GAA--GGGCGA
AGG--GGCCGCAGAGAAUCGGUGGCUGCGA-----CUGUUUAUUAAAAACACAGCACUCUGCAAACACG---
-AA-A-------------GUGGACGU-AUAGGGUGUGA-CGCCUGCCCGGUGCCG-G-AAGG-UUAAGUGAU
GGGGU----------GCAA----------GCUCUUGAUCGAAGCCC--CGGUAAAC-GGCGGCCGUAACUAU
AACGGUCCUAAGGUAGCGAAAUUCCUUGUCGGG-UAAGUUCCGACCUGCACGAAUGGCGUAACGAUG--GCC
ACACUGUCUCCUCC----UGA-GACUCAGCGAAGUU--GAA---AUGUUUGUGAAGAUGCAAUCUCCCCGCG
GCUAGACGGAAAGA-CCCCAUGAACC-UUUACU--GUAGCUU-UGCAUUGGACUUUGACGGGACUUGUGUAG
GAU-AGGUGGGAGGCUUUG----------AAGCCAG-GAC----------------------------GCUA
-------------------------------------GU-UCUGGUGGA---G--CCGU-CCUUGAAAUA-C
CACC-CUGGUGUCGUUGAGGUUCUAACCCAGG--CCCG-UGAA-U-C-------------------------
------------------------------------------------------------------------
------------------------------------------------------------------------
------------------------------------------------------------------------
------------------------------------------------------------------------
------------------------------------------------------CG----GG----------
------------------------------------------------UCGG-----GAACA-GUGCAUGGU
```

```
AGGCAGUUUGACUGGGGCGGU-CUCCUCCCA-AAAGGUAAC-GGAGGAGUUCG-AAGGUCUCCUAGGUACGG
UCGGACAUCGUACUGA-UAGUG-CAAUGGCAA-AAGGAGGCUUGACU-GCGA--------------------
--------------GACCGA---CAAGUCGAGCAGGUGCGAAA-GCAGGACAUAGUGAUCCGGU--------
---GG-U-----UCUGU-AUG--------------------GA-AGGGCCAUCGCUCAACGG-AUAAAAGGU
ACUCUGGGG-AUAACAGGCUGAUUCCGCCCAAGAGUUCACAUCGACGGCGGAGUUUGGCACCUCGAUGUCGG
CUCAUCACAUCCUGGGGCUGUA-GCCGGUCCCAAGGGUAUGGCUGUUCGCCAUUUAAAGUGGUACGUGAGCU
GGGUUUAAAACGUCGUGAGACAGUUUGGUCCCUAUCUGCC--------------------G-UGG-------
-------------------------------GCGC--UGGAAGUUUGAGGGG-ACC-UGCUCCUAGUACGAG
AGGAC-CGGAGUGGACGCACCUC-UGGU-G-UACCGGUUGUGACGCCAGUCGCAU-C-GCCGGGUAGCUAAG
UGCGGAAGAGAUAACCGCUGAAAGCA-UC--UAAGCGGGAAACUCGCCUCAA-G-AUG--AGACUUCC-CU-
--GGGGA-----------------------------------------------------------------
--------------------------CU--------CGA---------------------------------
------------------------------------------------------------------------
------------------------------------------------------------------------
------------------------------------------------------------------------
------------------------------------------------------------------------
------------------------------------------------------------------------
------------------------------------------------------------------------
-------------------------------------------------UCCCCCU--GA-AGG------GU
CGUGGAAGACCACCACGU-UGA-UAGGUC-GGG-UGUGGAAG-CGC-A----------------GU------
----AA-----------UGCGUUAAGCUA--A-CCGA-UACUAAUU---GCCCGUGAG-GCUUGAUCC----
------------------------------------------------------------------------
------------------------------------------------------------------------
------------------------------------------------------------------------
------------------------------------U------------
>LSUb_159087_DECAR/1-2882
---------------------G-AUCAAGCG----AC--UAAGUGCAUGU-GGUGGAUG-C-CUUGGC-GAU
CAC---AGGCGAUGAAGGACGUGC--AAGCCUGCGAAAAGCGGGGGGGAGCUGGCAAUGAAGCU-----UUG
AUC-CCC-CGAUAUCCGAAUGGGGAAAC--------CC--------AC----UCA---------------GC
A-A---------------------------U---GAG----U-A-UCCC-UGA-----CUGA----------
-----------A-----------UACA----------------------------------------UA-GG
U-C--------------------AG-G-GA-GGCGAACCCGGUGAACUGAAACAUCUAAGUAGCCG--GAGG
AAAAGAAAUCAA--CCGAGAUUCCCCAA--GU-AG-UGGCGAGCGAACGGGGA-U-GA-GCCUG--------
------------------------------------------------------------------------
----------CUA-------G--U-GA----------------------UA--GCG-----------GA---
A-------UGGUU-AAUGG------AAC-GG-AAUG------------------------------------
-------GAA-AG-UCCGG-------CCGUAGUGGGUGAUAGCCCCGUA---------CAUGAAA----ACC
A------A--UCCGU--------GGUACU---AAGU--UA-GC------GAAAAGUAGGGCGGGGCACGUGA
AACCUUGUCUGAA-CAUGGGGGG-ACC--AUCCUCCAAGGCUAAAUA--CUC-GUGAU-CGACCGAUAGU-G
AAC-C-AGUACCGUGAGGGAAAG-GCGAAA-AGAAC-CCCGG-GAGGGG-A-GUGAAAUAGAACCUGAAACC
G-CAUGCAUACAA-ACA-GU-GGGA---GCCU-C--------------------------------------
------------------------------------------------------------------------
------------------------------------------------------------------------
------------------------------------------------------------------------
------------------------------------------------------------------------
```

```
------------------------------------------------------------------------
------------------------------------------------------------------------
------------------------------------------------------------------------
------------------------------------------------------------------------
-------------------------------------------------------------CU--U-GU---
------------------------------------------------------------------------
------------------------------------------------------------------------
------------------------------------------------------------------------
------------------------------------------------------------------------
------------------------------------------------------------------------
------------------------------------------------------------------------
------------------------------------------------------------------------
------------------------------------------------------------------------
------------------------------------------------------------------------
------------------------------------------------------------------------
------------------------------------------------------------------------
------------------------------------------------------------------------
------------------------------------------------GG--G---GUGACUGCGUACCUUU
UGUAUAAUGGGUCAGCGACUUAC-G-UUUAGUAGCAA-GCUUAACC-------------GAAUA--------
---G--GGGA--GGCGCAGCGAAAG-C---------------------------------------------
------------------------------------------------------------------------
------------------------------------------------------------------------
---------------------------------------------------------------------GAG
UCU------------------------------------GAUAA-------------------------GG-
G-C-GA---------------------UUUAGUUGCUAGGCGUA-GACCCGAAACC-GGAUGAUCUAUCCAU
GGCCAGGAUGAAGGUGCC-GUAACA-GGUACUGGAGGUCCGA--ACCCACUA-AUGUUGAAAAAUUA-GGGG
AUGAGCUGUGGAUAGGGGUGAAAGGCUAAACAAAUCCGG-AAAUAGCUGGUUCUCCCCGAAA-ACUAUUUAG
GUAGUGCGUCUU-GUA--------------------------------------------------------
-----------------------U-----CA-CUGACGGGGGUAAAGC-ACUGUUA-UGG-CUA--GGGGGU
C---AUCGC--GACUUACCAAACCAUGGCAAACUCUGAAUACCGUC--------------------------
------------------------------------------------------------------------
------------------AAGU-GCGA--------GCAAGGC-AGACAGA-C-AGUG-G-GUGCUAACGUCC
AU-U-GUCAAGAG-GGAAAC---AACCCAGACCGCCA-GCUAAGGUCCCCAAGA--CA-UGG-UU-AA---G
-U-GGG-------AAACGAAGUGGGAAGGCAUAGACAGCUAGGAGGUUGGCUUAGAAGCAGCCA-UCCUUUA
AAGAAAGCGUAAUAGCUCACUAGUC-----GAGUCGUCCUGCGCGGAAGAUGUAAC-GGGGCUCA-AACCAU
G----------CACCGAA-GCUGCGGAUAUC-----------------------------------------
------------------------------------------------------------------------
------------------------------------------------------------------------
---------------------------------GAAA-GAUA-UGGUAGGGGAGC-GUUCUGU---------
-----------AGG-UCUGU-GAAGGUG--U-CUUGUAAAGG--A-UGCUGGAGAU-A-UCA-GAAGUGCGA
AUGC-UGACAU-GAGUA-GCGAUA---AAGG-GAGUGA--A---------------AAGCUC-CCU-CGCCG
A-AAGCCC-AAGGUUUCC-UACGCAACG-UUCAUCGGCGUAGGGUGAGUCGGC-CC-CUAAGGCGAGGCAG-
A-A-AUGC-----GUA--------------------------------------------------------
----------------------------------------GUC-GA-U--GGGA-AACAGGUC-AAUA-UUC
CUGUA----------------------------------CCGAU----UCUA--------G-AU--------
```

```
------------------------------------------------------------------------
----------------------------------------------------------GCGAU-GUGGGGAC
GGAGAA--------------GGUUAG-GUCAGCCAA-CU-G----UUGGA------------AUAGUUGGU-
UUAAGC-GUG-UA-----G-G-------------------CG-----UG------------------CC-CC
U-UAGG--------------------CAAAU-CCGGGGGGCU------U---AGCUG----AGG-CG-U-G-
AUG--AC---------------------------------------------------------------GA
-----GGCAC--UACGGUGCUGAAG-U-GAU-U-GAUACCAAGCUUC-C-AGGAAA-AGCCA-CUAAGCUU-
-------------------------------------------------------CAGUCUA-GA-AUUGAC
CGUACCGCAAACCGAC------------------------------ACAGGUG--GGCAGGAU-GAAAAUUC
UAAGGC-G-CU-UGAG-AGAACUCA-G-GAGAAGGAACUCGGC-AAAUUAACACCGUAACUUCGG-GAGAAG
GUGUG-CCCC-G-GU-AGGUUGUAG----AGC-----------CUCGC------------------------
------------------------------------------------------------------------
------------------------------------------------------------------------
------------------------------------------------------------------------
------------------------------------------------------------------------
------------------------------------------------------------------------
------------------------------------------------------------------------
------------------------------------------------------------------------
------------------------------------------------------------------------
------------------------------------------------------------------------
------------------------------------------------------------------------
------------------------------------------------------------------------
------------------------------------------------------------------------
------------------------------------------------------------------------
--------------------------------------------------------GCUC-GAA--GGCCGA
UGG--GGUUGCAGUGAAAUGGUGGCUGCGA-----CUGUUUAAUAAAAACACAGCACUCUGCAAACACG---
-AA-A-------------GUGGACGU-AUAGGGUGUGA-CGCCUGCCCGGUGCCG-G-AAGG-UUAAUUGAU
GGGGU----------GCAA----------GCCCUUGAUCGAAGCCC--CGGUAAAC-GGCGGCCGUAACUAU
AACGGUCCUAAGGUAGCGAAAUUCCUUGUCGGG-UAAGUUCCGACCUGCACGAAUGGCGUAACGAUG--GCC
ACACUGUCUCCUCC----UGA-GACUCAGCGAAGUU--GAA---AUGUUUGUGAAGAUGCAAUCUCCCCGCG
GCAAGACGGAAAGA-CCCCAUGAACC-UUUACU--GUAGCUU-UGCAUUGGACUUUGAAUCGGUCUGUGUAG
GAU-AGGUGGGAGGCUGUG----------AAACCGG-GAC----------------------------GCUA
--------------------------------------GU-UUCGGUGGAG-C--C-AA-CCUUGAAAUA-C
CACC-CUGAUUUAUUUGAGGUUCUAACCUUGA--UCCG-UGAA-U-C-------------------------
------------------------------------------------------------------------
------------------------------------------------------------------------
------------------------------------------------------------------------
------------------------------------------------------------------------
------------------------------------------------------CG----GA----------
------------------------------------------------UCGG-----GGACC-GUGCAUGGU
GGGCAGUUUGACUGGGGCGGU-CUCCUCCCA-AAAGGUAAC-GGAGGAGUACG-AAGGUACGCUUAGGGCGG
UCGGACAUCGCCCAUA-AAGUG-CAAUGGCAA-AAGCGUGCUUGACU-GCGA--------------------
--------------GACCCA---CAAGUCGAGCAGGUGCGAAA-GCAGGUCAUAGUGAUCCGGU--------
---GG-U-----UCUGU-AUG--------------------GA-AGGGCCAUCGCUCAACGG-AUAAAAGGU
ACUCUGGGG-AUAACAGGCUGAUACCGCCCAAGAGUUCAUAUCGACGGCGGUGUUUGGCACCUCGAUGUCGG
```

```
CUCAUCUCAUCCUGGGGCUGUA-GCCGGUCCCAAGGGUAUGGCUGUUCGCCAUUUAAAGAGGUACGUGAGCU
GGGUUUAAAACGUCGUGAGACAGUUUGGUCCCUAUCUGCC--------------------G-UGG-------
-------------------------------GCGC--UGGAAAUUUGAAGGG-GGC-UGCUCCUAGUACGAG
AGGAC-CGGAGUGGACGAACCUC-UGGU-G-UACCGGUUAUGACGCCAGUCGUAU-C-GCCGGGUAGCUAUG
UUCGGAAGAGAUAAACGCUGAAAGCA-UC--UAAGCGUGAAACUCGCCUUAA-G-AUA--AGAUUUCC-C--
--GGAGU-----------------------------------------------------------------
--------------------------CU--------UGA---------------------------------
------------------------------------------------------------------------
------------------------------------------------------------------------
------------------------------------------------------------------------
------------------------------------------------------------------------
------------------------------------------------------------------------
------------------------------------------------------------------------
-------------------------------------------------ACUCCUU--GA-AGG------GU
CGUCGAAGACCACGACGU-UGA-UAGGUC-AGG-UGUGGAAG-CGC-A----------------GU------
----AA-----------UGCGUUAAGCUA--A-CUGA-UACUAAUU---GCCCGUACG-GCUUGAUCC----
------------------------------------------------------------------------
------------------------------------------------------------------------
------------------------------------------------------------------------
------------------------------------U------------
>LSUb_228410_NITEU/1-2886
---------------------G-AUCAAGUG----AA--UAAGUGCAUGU-GGUGGAUG-C-CUUGGC-GAU
UAC---AGGCGAUGAAGGACGUGG--AAGCCUGCGAAAAGCUUCGGGGAGCUGGCAAACAAGCU-----UUG
AUC-CGG-AGAUGUCCGAAUGGGAAAAC--------CC--------AC----CC----------------GU
A-A-------------------------------GGG----U-A-AUCA-CUC-----CUGA----------
-----------A-----------UAUA----------------------------------------UA-GG
GAG--------------------AU-G-AU-GGCUAACCUGGAGAACUGAAACAUCUAAGUAACCA--GAGG
AAAAGAAAUCAA--CAGAGAUUCCCAGA--GU-AG-UGGCGAGCGAAAUGGGA-C-CA-GCCUG--------
------------------------------------------------------------------------
----------CAG-------U----UAA---------------------UA--AUA-----------UU---
U-------AGACU-AACCG------AAU-UA-UCUG------------------------------------
-------GAA-AG-UUAAA--------CCGUAGUGGGUGAUAGUCCCGUA---------GGUGAAA----GUC
U------G--AAUGU--------GGAACU---AAGA--UU-GC------AAUAAGUAGGGCGGGGCACGUGU
AAUCCUGUCUGAA-UAUAGGGGG-ACC--AUCCUCUAAGGCUAAAUA--CUC-GUAAU-CGACCGAUAGU-G
AAC-C-AGUACCGUGAGGGAAAG-GCGAAA-AGAAC-CCCGG-GAGGGG-A-GUGAAAUAGAUCCUGAAACC
G-CAUGCAUACAA-ACA-GU-AGGA---GCCU-U--------------------------------------
------------------------------------------------------------------------
------------------------------------------------------------------------
------------------------------------------------------------------------
------------------------------------------------------------------------
------------------------------------------------------------------------
------------------------------------------------------------------------
------------------------------------------------------------------------
------------------------------------------------------------------------
--------------------------------------------------------------G--A-AA---
```

```
------------------------------------------------------------------------
------------------------------------------------------------------------
------------------------------------------------------------------------
------------------------------------------------------------------------
------------------------------------------------------------------------
------------------------------------------------------------------------
------------------------------------------------------------------------
------------------------------------------------------------------------
------------------------------------------------------------------------
------------------------------------------------------------------------
------------------------------------------------------------------------
------------------------------------------------------------------------
------------------------------------------------GA--G---GUGACUGCGUACCUUU
UGUAUAAUGGGUCAGCGACUUAC-A-UUCAGUAGCGA-GCUUAACC--------------GAUA--------
---G--GGGA--GGCGUAGUGAAAA-C---------------------------------------------
------------------------------------------------------------------------
------------------------------------------------------------------------
---------------------------------------------------------------------GAG
UCU------------------------------------UAAUA-------------------------GG-
G-C-GA----------------------UAAGUUGCUGGGUGUA-GACCCGAAACC-AGAUGAUCUACUCAU
GGCCAGGAUGAAAGGAGG-GUAAUG-CCUCGUGGAGGUCCGA--ACCCACUA-AUGUUGAAAAUUA-GGGG
AUGAGCUGUGGGUAGGGGUGAAAGGCUAAACAAAUCUGG-AAAUAGCUGGUUCUCUCCGAAA-ACUAUUUAG
GUAGUGCCUCAU-AUA--------------------------------------------------------
-----------------------U-----UA-CCUUUGGGGGUAGAGC-ACUGUUA-UGG-CUA--GGGGGU
C---GUCAA--GAUUUACCAAACCAUUGCAAACUCCGAAUACCAAA--------------------------
------------------------------------------------------------------------
------------------GAGU-GCAA--------GUAUGGG-AGACAGA-C-AUCG-G-GUGCUAACGUUC
GG-U-GUCGAAAG-GGAAAC---AACCCAGACCUUCA-GCUAAGGUCCCAAAGA--UA-CAG-UU-AA---G
-U-GGU-------AAACGAAGUGGGAAGGCAUAGACAGUCAGGAAGUUGGCUUAGAAGCAGCCA-UCCUUUA
AAGAAAGCGUAAUAGCUCACUGAUC-----GAGUCGUCCUGCGCGGAAGAUGUAAC-GGGGCU-A-AACUGU
A----------CACCGAA-GCUAAGGAUUUGCAA--U-----------------------------------
------------------------------------------------------------------------
-----U------------------------------------------------------------------
-------------------------------UAAUUG-CAAG-UGGUAGGAGAGC-GUUCCGU---------
-----------AAG-CCUGU-GAAGGUA--A-CUUGUGAAGG--U-UGCUGGAGGU-A-UCG-GAAGUGCGA
AUGC-UGACAU-GAGUA-GCGAUA---AAGG-AAGUGA--A---------------AAGCUU-CCU-CGCCG
A-AAACCC-AAGGUUUCC-UGUGCAACG-UUCAUCGGCGCAGGGUUAGUCGGU-CC-CUAAGGUGAGGUAG-
A-A-AUAC-----GUA--------------------------------------------------------
----------------------------------------GCU-GA-U--GGGA-AACUGGUU-AAUA-UUC
CAGUA----------------------------------CCUUU----AUUU--------A-AU--------
------------------------------------------------------------------------
----------------------------------------------------------GCGAU-GUGGGGAC
GAAGAA--------------GGCUAG-CUUAGCCGG-GU-G----UUGGA------------UGUCCCGGU-
UGAAGC-AGG-UA-----G-A-------------------CA-----UG------------------CU-AU
U-UAGG--------------------CAAAU-CCGGAUAGCU-------U---AGUCG----AGA-UG-U-G-
```

```
AUA--AC---------------------------------------------------------------GA
AAC--CUCUU--UCGGGAGGCCAAG-U-AAG-U-AAUGCCAUGCUUC-C-AGGAAA-AGCCA-CUAAGCUU-
-------------------------------------------------------CAGUUAA-AU-AAAGAC
CGUACCGUAAACCGAC------------------------------ACAGGUG--GGUGGGAU-GAGAAUUC
UAAGGC-G-CU-UGAG-AGAACUCA-G-GAGAAGGAACUCGGC-AAAUUGACACCGUAACUUCGG-GAGAAG
GUGUG-CCUC-U-AG-UAUGUGUAG----UGC-----------UACAC------------------------
------------------------------------------------------------------------
------------------------------------------------------------------------
------------------------------------------------------------------------
------------------------------------------------------------------------
------------------------------------------------------------------------
------------------------------------------------------------------------
------------------------------------------------------------------------
------------------------------------------------------------------------
------------------------------------------------------------------------
------------------------------------------------------------------------
------------------------------------------------------------------------
------------------------------------------------------------------------
------------------------------------------------------------------------
--------------------------------------------------------GCAU-GAA--GCAGAA
UGA--GGUUGCAAAGAAAUGGUGGCUGCGA-----CUGUUUAAUAAAAACACAGCACUCUGCAAACACG---
-AA-A-------------GUGGACGU-AUAGGGUGUGA-CGCCUGCCCGGUGCCG-G-AAGG-UUAAGUGAU
GGGGU----------GCAA----------GCUCUUGAUCGAAGCCC--CGGUAAAC-GGCGGCCGUAACUAU
AACGGUCCUAAGGUAGCGAAAUUCCUUGUCGGG-UAAGUUCCGACCUGCACGAAUGGCGUAACGAUG--GCC
ACACUGUCUCCUCC----UGA-GACUCAGCGAAGUU--GAA---AUGUUUGUGAAGAUGCAAUCUACCCGCG
GCUAGACGGAAAGA-CCCCGUGCACC-UUUACU--GUAGCUU-UACAUUGGAUUUUGAUUAUUCUUGUGUAG
GAU-AGGUGGGAGGCUGUG----------AAAUAUG-CUC----------------------------GCCA
--------------------------------------GA-GUGUAUGGAG-C--C-AA-CCUUGAAAUA-C
CACC-CUGGAAUGAUCGGAGUUCUAACCCAGG--UCCA-U-UA-U-C-------------------------
------------------------------------------------------------------------
------------------------------------------------------------------------
------------------------------------------------------------------------
------------------------------------------------------------------------
------------------------------------------------------UG----GA----------
------------------------------------------------UCGG-----GGACC-GUGUAUGGU
AGGCAGUUUGACUGGGGCGGU-CUCCUCCCA-AAGAGUAAC-GGAGGAGUGCG-AAGGUACGCUAGGUACGG
UCGGAAAUCGUACUGA-UAGUG-CAAUGGCAU-AAGCGUGCUUAACU-GCGA--------------------
--------------GACUGA---CAAGUCGAGCAGAUGCGAAA-GCAGGUCAUAGUGAUCCGGU--------
---GG-U-----UCUGU-AUG--------------------GA-AGGGCCAUCGCUCAACGG-AUAAAAGGU
ACGCCGGGG-AUAACAGGCUGAUUCCUCCCAAGAGUUCAUAUCGACGGGGGAGUUUGGCACCUCGAUGUCGG
CUCAUCACAUCCUGGGGCUGUA-GUCGGUCCCAAGGGUAUGGCUGUUCGCCAUUUAAAGUGGUACGUGAGCU
GGGUUUAAAACGUCGUGAGACAGUUUGGUCCCUAUCUGCC--------------------G-UGG-------
-------------------------------GCGU--UGGAAAUUUGAGUGG-GGC-UGCUCCUAGUACGAG
AGGAC-CGGAGUGGACGCACCUC-UGGU-G-UACCGGUUAUGACGCCAGUCGUAU-C-GCCGGGUAGCUAAG
UGCGGAAGAGAUAACCGCUGAAAGCA-UC--UAAGCGGGAAACUUGCCAUAA-G-AUU--AGAUUUCC-C--
```

```
--GGGAA-----------------------------------------------------------------
--------------------------UU--------UAA---------------------------------
------------------------------------------------------------------------
------------------------------------------------------------------------
------------------------------------------------------------------------
------------------------------------------------------------------------
------------------------------------------------------------------------
------------------------------------------------------------------------
-------------------------------------------------UUCCCCU--AA-AGG------UU
CGUUGAAGACUACAACGU-UGA-UAGGUC-GAA-UGUGUAAG-CGU-A----------------GU------
----AA-----------UGCGUUAAGCUA--A-UCGA-UACUAAUU---GACCGUGAG-GCUUGAUCC----
------------------------------------------------------------------------
------------------------------------------------------------------------
------------------------------------------------------------------------
------------------------------------U------------
>LSUb_482957_BURL3/1-2882
---------------------G-GUCAAGCG----AA--CAAGUGCAUGU-GGUGGAUG-C-CUUGGC-GAU
CAC---AGGCGAUGAAGGACGCGG--UAGCCUGCGAAAAGCUACGGGGAGCUGGCAAACAAGCU-----UUG
AUC-CGU-AGAUGUCCGAAUGGGGAAAC--------CC--------AC----UCC---------------UU
U-U---------------------------G---GAG----U-A-UCCA-UGG-----CUGA----------
-----------A-----------UACA----------------------------------------UA-GG
C-C--------------------AU-GUGA-AGCGAACGCGGUGAACUGAAACAUCUAAGUAACCG--CAGG
AAAAGAAAUCAA--CCGAGAUUCCCAAA--GU-AG-UGGCGAGCGAAAUGGGA-U-GA-GCCUU--------
------------------------------------------------------------------------
-------GC-ACU-------C-UU-UA--------------------------UUU-----------GU---
A-------UUGUU-AGCCG------AAC-GC-UCUG------------------------------------
-------GAA-AG-UGCGG-------CCAUAGCGGGUGAUAGCCCCGUA---------GGCGAAA----ACA
G------U--AUGAA--------GGAACU---AGGU--GU-GC------GACAAGUAGGGCGGGACACGUGA
AAUCCUGUCUGAA-GAUGGGGGG-ACC--AUCCUCCAAGGCUAAAUA--CUC-GUGAU-CGACCGAUAGU-G
AAC-C-AGUACCGUGAGGGAAAG-GCGAAA-AGAAC-CCCGG-GAGGGG-A-GUGAAAUAGAUCCUGAAACC
G-CAUGCAUACAA-ACA-GU-CGGA---GCCU-C--------------------------------------
------------------------------------------------------------------------
------------------------------------------------------------------------
------------------------------------------------------------------------
------------------------------------------------------------------------
------------------------------------------------------------------------
------------------------------------------------------------------------
------------------------------------------------------------------------
------------------------------------------------------------------------
--------------------------------------------------------------G--U-AA---
------------------------------------------------------------------------
------------------------------------------------------------------------
------------------------------------------------------------------------
------------------------------------------------------------------------
------------------------------------------------------------------------
```

```
------------------------------------------------------------------------
------------------------------------------------------------------------
------------------------------------------------------------------------
------------------------------------------------------------------------
------------------------------------------------------------------------
------------------------------------------------------------------------
------------------------------------------------------------------------
------------------------------------------------GG--G---GUGACGGCGUACCUUU
UGUAUAAUGGGUCAGCGACUUAC-G-UUCAGUAGCAA-GCUUAACC-------------GUAUA--------
---G--GGCA--GGCGUAGCGAAAG-C---------------------------------------------
------------------------------------------------------------------------
------------------------------------------------------------------------
---------------------------------------------------------------------GAG
UCC------------------------------------GAAUA-------------------------GG-
G-C-GU----------------------UCAGUUGCUGGGCGUA-GACCCGAAACC-AAGUGAUCUAUCCAU
GGCCAGGAUGAAGGUGCG-GUAACA-CGUACUGGAGGUCCGA--ACCCACUA-ACGUUGAAAAGUUA-GGGG
AUGAGCUGUGGAUAGGGGUGAAAGGCUAAACAAACUUGG-AAAUAGCUGGUUCUCUCCGAAA-ACUAUUUAG
GUAGUGCCUCGU-GUC--------------------------------------------------------
-----------------------U-----CA-CCUUCGGGGGUAGAGC-ACUGUCA-UGG-UUG--GGGGGU
C-U-AU-UG-CAGAUUACCCCGCCAUAGCAAACUCCGAAUACCGAA--------------------------
------------------------------------------------------------------------
------------------GAGU-GCAA--------UCACGGG-AGACAGA-C-AUCG-G-GUGCUAACGUCC
GG-U-GUCAAGAG-GGAAAC---AACCCAGACCGCCA-GCUAAGGUCCCCAAAU---A-UAG-CU-AA---G
-U-GGG-------AAACGAAGUGGGAAGGCUAAAACAGUCAGGAGGUUGGCUUAGAAGCAGCCA-CCCUUUA
AAGAAAGCGUAAUAGCUCACUGAUC-----GAGUCGUCCUGCGCGGAAGAUGUAAC-GGGGCU-A-AGCUAU
A----------UACCGAA-GCUGCGGAUGCGUGC--U-----------------------------------
------------------------------------------------------------------------
------------------------------------------------------------------------
--------------------------------UUGCA-CGCA-UGGUAGGAGAGC-GUUCUGU---------
-----------AAG-CCUGC-GAAGGUG--U-CUCGGAAGGG--A-UGCUGGAGGU-A-UCA-GAAGUGCGA
AUGC-UGACAU-GAGUA-GCGAUA---AAGG-GGGUGA--A---------------AGGCCC-CCU-CGCCG
U-AAGCCC-AAGGUUUCC-UACGCAACG-UUCAUCGGCGUAGGGUGAGUCGGC-CC-CUAAGGCGAGGCAG-
A-A-AUGC-----GUA--------------------------------------------------------
----------------------------------------GCU-GA-U--GGGA-AGCAGGUC-AAUA-UUC
CUGCA----------------------------------CCAUU----GUUA--------G-AU--------
------------------------------------------------------------------------
----------------------------------------------------------GCGAU-GGGGGGAC
GGAUCG--------------CGGAAA-GUUGUCCGG-GU-G----UUGGA------------AGUCCCGGU-
CGCUGC-AU--UG-----G-A-------------------GA-----AG------------------GC-GC
U-UAGG--------------------CAAAU-CCGGGCGCGG--------AAUUCAA----GGG-UG-U-G-
GC---GC---------------------------------------------------------------GA
-----GCUCC----UUAGGGAGCGA-A-GCA-A-UUGGAAGUGGUUC-C-AAGAAA-AGCCU-CUAAGCUU-
-------------------------------------------------------CAGUCUA-AU-GAUGAC
CGUACCGCAAACCGAC------------------------------ACAGGUG--GGCGAGAU-GAGUAUUC
UAAGGC-G-CU-UGAG-AGAACUCG-G-GAGAAGGAACUCGGC-AAAUUGGUACCGUAACUUCGG-GAUAAG
```

```
GUACG-CCCU-U-GU-AGCUUGAUG----CCC------------CUGC------------------------
------------------------------------------------------------------------
------------------------------------------------------------------------
------------------------------------------------------------------------
------------------------------------------------------------------------
------------------------------------------------------------------------
------------------------------------------------------------------------
------------------------------------------------------------------------
------------------------------------------------------------------------
------------------------------------------------------------------------
------------------------------------------------------------------------
------------------------------------------------------------------------
------------------------------------------------------------------------
------------------------------------------------------------------------
--------------------------------------------------------GGGC-AAA--GGGUGA
AGG--GGUUGCAAUAAACUGGUGGCUGCGA-----CUGUUUAAUAAAAACACAGCACUCUGCAAACACG---
-AA-A-------------GUGGACGU-AUAGGGUGUGA-CGCCUGCCCGGUGCCG-G-AAGA-UUAAAUGAU
GGGGU----------GCAA----------GCUCUUGAUUGAAGUCC--CGGUAAAC-GGCGGCCGUAACUAU
AACGGUCCUAAGGUAGCGAAAUUCCUUGUCGGG-UAAGUUCCGACCUGCACGAAUGGCGUAACGAUG--GCC
ACACUGUCUCCUCC----CGA-GACUCAGCGAAGUU--GAA---GUGUUUGUGAUGAUGCAAUCUACCCGCG
GCUAGACGGAAAGA-CCCCAUGAACC-UUUACU--GUAGCUU-UGCAUUGGACUUUGAACCGAUCUGUGUAG
GAU-AGGUGGGAGGCUAUG----------AAACCGG-AAC----------------------------GCUA
--------------------------------------GU-UUCGGUGGAG-C--C-GU-CCUUGAAAUA-C
CACC-CUGGUUUGUUUGAGGUUCUAACCUUGG--CCCG-U-GA-U-C-------------------------
------------------------------------------------------------------------
------------------------------------------------------------------------
------------------------------------------------------------------------
------------------------------------------------------------------------
------------------------------------------------------CG----GG----------
------------------------------------------------UCGG-----GGACA-GUGCAUGGU
AGGCAGUUUGACUGGGGCGGU-CUCCUCCCA-AAGCGUAAC-GGAGGAGUACG-AAGGUACGCUAGGUACGG
UCGGAAAUCGUGCUGA-UAGUG-CAAUGGCAU-AAGCGUGCUUAACU-GCGA--------------------
--------------GACCGA---CAAGUCGAGCAGGUGCGAAA-GCAGGUCAUAGUGAUCCGGU--------
---GG-U-----UCUGU-AUG--------------------GA-AGGGCCAUCGCUCAACGG-AUAAAAGGU
ACUCUGGGG-AUAACAGGCUGAUACCGCCCAAGAGUUCAUAUCGACGGCGGUGUUUGGCACCUCGAUGUCGG
CUCAUCUCAUCCUGGGGCUGUA-GCCGGUCCCAAGGGUAUGGCUGUUCGCCAUUUAAAGAGGUACGUGAGCU
GGGUUUAAAACGUCGUGAGACAGUUUGGUCCCUAUCUGCC--------------------G-UGG-------
-------------------------------GCGU--UGGAUAUUUGAAGGG-GGC-UGCUCCUAGUACGAG
AGGAC-CGGAGUGGACGAACCUC-UGGU-G-UACCGGUUGUCACGCCAGUGGCAU-C-GCCGGGUAGCUAUG
UUCGGAAGAGAUAACCGCUGAAAGCA-UC--UAAGCGGGAAACUCGCCUUAA-G-AUG--AGAUAUCC-CU-
--GGGGA-----------------------------------------------------------------
--------------------------CU--------AGA---------------------------------
------------------------------------------------------------------------
------------------------------------------------------------------------
------------------------------------------------------------------------
```

```
------------------------------------------------------------------------
------------------------------------------------------------------------
------------------------------------------------------------------------
-------------------------------------------------UCCCCUU--GA-AGG------GU
CGUUCGAGACCAGGACGU-UGA-UAGGUC-AGG-UGUGUAAG-CGC-A----------------GU------
----AA-----------UGCGUUCAGCUA--A-CUGA-UACUAAUU---GCCCGUAAG-GCUUGAUCC----
------------------------------------------------------------------------
------------------------------------------------------------------------
------------------------------------------------------------------------
------------------------------------U------------
>LSUb_264198_CUPNJ/1-2881
---------------------G-GUCAAGCG----AA--CAAGUGCAUGU-GGUGGAUG-C-CUUGGC-GAU
CAC---AGGCGAUGAAGGACGCGG--UAGCCUGCGAAAAGCUUCGGGGAGCUGGCAAACGAGCU-----UUG
AUC-CGG-AGAUGUCCGAAUGGGGAAAC--------CC--------GG----CCC---------------GA
A-U---------------------------G---GGU----C-A-UCCC-UGA-----CUGA----------
-----------A-----------UACA----------------------------------------UA-GG
U-C--------------------AG-G-GA-AGCGAACGCGGCGAACUGAAACAUCUAAGUAGCUG--CAGG
AACAGAAAUCAA--CCGAGAUUCCCAAA--GU-AG-UGGCGAACGAAAUGGGA-A-GA-GCCUU--------
------------------------------------------------------------------------
----------GUA-------C-UC-UU----------------------UA--GCA-----------GU---
G-------UUGUU-AGCAG------AAC-GG-GAUG------------------------------------
-------GAA-AG-CCCGG-------CCGUAGCAGGUGAUAGCCCUGUA---------UGCGAAA----ACA
G------C--GUUGU--------GGAACU---AGGU--GU-AC------GACAAGUAGGGCGGGACACGUGA
AAUCCUGUCUGAA-GAUGGGGGG-ACC--AUCCUCCAAGGCUAAAUA--CUC-GUGAU-CGACCGAUAGU-G
AAC-C-AGUACCGUGAGGGAAAG-GCGAAA-AGAAC-CCCGG-GAGGGG-A-GUGAAAUAGAUCCUGAAACC
G-CAUGCAUACAA-ACA-GU-CGGA---GCCU-C--------------------------------------
------------------------------------------------------------------------
------------------------------------------------------------------------
------------------------------------------------------------------------
------------------------------------------------------------------------
------------------------------------------------------------------------
------------------------------------------------------------------------
------------------------------------------------------------------------
------------------------------------------------------------------------
--------------------------------------------------------------G--U-AA---
------------------------------------------------------------------------
------------------------------------------------------------------------
------------------------------------------------------------------------
------------------------------------------------------------------------
------------------------------------------------------------------------
------------------------------------------------------------------------
------------------------------------------------------------------------
------------------------------------------------------------------------
------------------------------------------------------------------------
------------------------------------------------------------------------
```

------------------------------------------------------------------------
------------------------------------------------------------------------
------------------------------------------------GG--G---GUGACGGCGUACCUUU
UGUAUAAUGGGUCAGCGACUUAC-A-UUCAGUGGCGA-GCUUAACC--G-----------AUUA--------
---G--GGCA--GGCGUAGCGAAAG-C---------------------------------------------
------------------------------------------------------------------------
------------------------------------------------------------------------
---------------------------------------------------------------------GAG
UCC------------------------------------GAACA-------------------------GG-
G-C-GA----------------------UUAGUCGCUGGGUGUA-GACCCGAAACC-AGAUGAUCUAUCCAU
GGCCAGGUUGAAGGUGCG-GUAACA-CGUACUGGAGGACCGA--ACCCACUA-ACGUUGAAAGUUA-GGGG
AUGAGCUGUGGAUAGGGGUGAAAGGCUAAACAAAUCUGG-AAAUAGCUGGUUCUCUCCGAAA-ACUAUUUAG
GUAGUGCCUCGU-GUG--------------------------------------------------------
-----------------------U-----CA-CCUUCGGGGGUAGAGC-ACUGUCA-UGG-UUG--GGGGGU
C-U-AU-UG-CUGAUUACCCCGCCAUAGCAAACUCCGAAUACCGAA--------------------------
------------------------------------------------------------------------
------------------GAGU-GCAA--------UCACGGG-AGACAGA-C-AUCG-G-GUGCUAACGUCC
GG-U-GUCAAGAG-GGAAAC---AACCCAGACCGCCA-GCUAAGGUCCCCAAAU---A-UGG-CU-AA---G
-U-GGG-------AAACGAAGUGGGAAGGCUAAAACAGUCAGGAGGUUGGCUUAGAAGCAGCCA-CCCUUUA
AAGAAAGCGUAAUAGCUCACUGAUC-----GAGUCGUCCUGCGCGGAAGAUGUAAC-GGGGCU-A-AGCCAU
A----------UACCGAA-GCUGCGGACGCACAG--C-----------------------------------
------------------------------------------------------------------------
------------------------------------------------------------------------
---------------------------------AAUG-UGCG-UGGUAGGAGAGC-GUUCCGU---------
-----------AAG-CCUGU-GAAGGUG--U-CUUGUAAAGG--A-UGCUGGAGGU-A-UCG-GAAGUGCGA
AUGC-UGACAU-GAGUA-GCGAUA---AAGG-GGGUGA--A---------------AGGCCC-CCU-CGCCG
U-AAGCCC-AAGGUUUCC-UACGCAACG-UUCAUCGGCGUAGGGUGAGUCGGC-CC-CUAAGGCGAGGCAG-
A-G-AUGC-----GUA--------------------------------------------------------
----------------------------------------GCU-GA-U--GGGA-AGCAGGUU-AAUA-UUC
CUGCA----------------------------------CCGUC----GUAU--------G-AU--------
------------------------------------------------------------------------
----------------------------------------------------------GCGAU-GGGGGGAC
GGAUCG--------------CGGAAG-GUUGUCCGG-GU-G----UUGGA------------AGUCCCGGU-
CCCUGC-AG--UG-----G-A-------------------GA-----AG------------------GC-GC
U-UAGG--------------------CAAAU-CCGGGCGCGU--------AAUUCAA----GGC-UG-U-G-
GG---GC---------------------------------------------------------------GA
-----GUGAGC---UAGGCUCGCGA-A-GCA-A-UUGGAAGUGGUUC-C-AAGAAA-AGCCU-CUAAGCUU-
--------------------------------------------------------CAGUCA-UA-CGAGAC
CGUACCGCAAACCGAC------------------------------ACAGGUG--GGCGAGAU-GAGUAUUC
UAAGGC-G-CU-UGAG-AGAACUCG-G-GAGAAGGAACUCGGC-AAAUUGGUACCGUAACUUCGG-GAUAAG
GUACG-CCCU-G-GU-AGCUUGACU----GGC------------CUGC------------------------
------------------------------------------------------------------------
------------------------------------------------------------------------
------------------------------------------------------------------------
------------------------------------------------------------------------

```
------------------------------------------------------------------------
------------------------------------------------------------------------
------------------------------------------------------------------------
------------------------------------------------------------------------
------------------------------------------------------------------------
------------------------------------------------------------------------
------------------------------------------------------------------------
------------------------------------------------------------------------
------------------------------------------------------------------------
----------------------------------------------------------GCCA-GAAGGGUGA
AGG--GGUUGCAAUAAAAUGGUGGCUGCGA-----CUGUUUAAUAAAAACACAGCACUCUGCAAACACG---
-AA-A-------------GUGGACGU-AUAGGGUGUGA-CGCCUGCCCGGUGCCG-G-AAGA-UUAAAUGAU
GGGGU----------GCAA----------GCUCUUGAUUGAAGUCC--CGGUAAAC-GGCGGCCGUAACUAU
AACGGUCCUAAGGUAGCGAAAUUCCUUGUCGGG-UAAGUUCCGACCUGCACGAAUGGCGUAACGAUG--GCC
ACACUGUCUCCUCC----CGA-GACUCAGCGAAGUU--GAA---GUGUUUGUGAUGAUGCAAUCUCCCCGCG
GCUAGACGGAAAGA-CCCCAUGAACC-UUUACU--GUAGCUU-UGCAUUGGACUUUGAACCGAUCUGUGUAG
GAU-AGGUGGGAGGCUUUG----------AAGCGUG-GAC----------------------------GCUA
--------------------------------------GU-UCACGUGGAG-C--C-GU-CCUUGAAAUA-C
CACC-CUGGUUUGUUUGAGGUUCUAACCUUGG--CCCG-UGAA-U-C-------------------------
------------------------------------------------------------------------
------------------------------------------------------------------------
------------------------------------------------------------------------
------------------------------------------------------------------------
------------------------------------------------------CG----GG----------
------------------------------------------------UCGG-----GGACA-GUGCAUGGU
AGGCAGUUUGACUGGGGCGGU-CUCCUCCCA-AAGUGUAAC-GGAGGAGUUCG-AAGGUACGCUUGGUACGG
UCGGACAUCGUACCUA-AAGUG-CAAUGGCAA-AAGCGUGCUUAACU-GCGA--------------------
--------------GACCGA---CAAGUCGAGCAGGUGCGAAA-GCAGGACAUAGUGAUCCGGU--------
---GG-U-----UCUGA-AUG--------------------GA-AGGGCCAUCGCUCAACGG-AUAAAAGGU
ACUCUGGGG-AUAACAGGCUGAUACCGCCCAAGAGUUCAUAUCGACGGCGGUGUUUGGCACCUCGAUGUCGG
CUCAUCUCAUCCUGGGGCUGUA-GCCGGUCCCAAGGGUAUGGCUGUUCGCCAUUUAAAGAGGUACGUGAGCU
GGGUUUAAAACGUCGUGAGACAGUUUGGUCCCUAUCUGCC--------------------G-UGG-------
-------------------------------GCGU--UGGAAUCUUGACGGG-GGC-UGCUCCUAGUACGAG
AGGAC-CGGAGUGGACGUACCGC-UGGU-G-UACCUGUUGUCUCGCCAGAGGCAU-C-GCAGGUAGCUAUG
UACGGAAGAGAUAACCGCUGAAAGCA-UC--UAAGCGGGAAACUCGCCUGAA-G-AUG--AGGAUUCC-CU-
--GGAGG-----------------------------------------------------------------
--------------------------CU--------UGA---------------------------------
------------------------------------------------------------------------
------------------------------------------------------------------------
------------------------------------------------------------------------
------------------------------------------------------------------------
------------------------------------------------------------------------
------------------------------------------------------------------------
-------------------------------------------------CCUCCUU--GA-AGG------GU
CGUUCGAGACCAGGACGU-UGA-UAGGCU-GGG-UGUGGAAG-CGC-A----------------GU------
```

```
----AA-----------UGCGUUAAGCUA--A-CCAG-UACUAAUU---GCCCGUAAG-GCUUGAUCC----
------------------------------------------------------------------------
------------------------------------------------------------------------
------------------------------------------------------------------------
------------------------------------U------------
>LSUb_242231_NEIG1/1-2879
-----------------------------------------AGUGCAUCA-GGCGGAUG-C-CUUGGC-GAU
GAU---AGGCGACGAAGGACGUGU--AAGCCUGCGAAAAGCGCGGGGGAGCUGGCAAUAAAGCA-----AUG
AUC-CCG-CGGUGUCCGAAUGGGGAAAC--------CC--------AC----UGCA-U-------------U
C-U-----------------G----U----G---CAG----U-A-UCCU-AAG-----UUGA----------
-----------A-----------UACA----------------------------------------UA-GG
C-U--------------------UA-GAGA-AGCGAACCCGGAGAACUGAACCAUCUAAGUACCCG--GAGG
AAAAGAAAUCAA--CCGAGAUUCCGCAA--GU-AG-UGGCGAGCGAACGCGGA-G-GA-GCCUG--------
------------------------------------------------------------------------
----------UAC-------G----UAA---------------------UA--ACU-----------GU---
C-------GAGGU-AGAAG------AAC-AA-GCUG------------------------------------
-------GGA-AG-CUUGA-------CCAUAGCGGGUGACAGUCCCGUA---------UUCGAAA----UCU
C------A--ACAGC--------GGUACU---AAGC--GU-AC------GAAAAGUAGGGCGGGACACGUGA
AAUCCUGUCUGAA-UAUGGGGGG-ACC--AUCCUCCAAGGCUAAAUA--CUC-AUCAU-CGACCGAUAGU-G
AAC-C-AGUACCGUGAGGGAAAG-GCGAAA-AGAAC-CCCGG-GAGGGG-A-GUGAAACAGAACCUGAAACC
U-GAUGCAUACAA-ACA-GU-GGGA---GCGC-C--------------------------------------
------------------------------------------------------------------------
------------------------------------------------------------------------
------------------------------------------------------------------------
------------------------------------------------------------------------
------------------------------------------------------------------------
------------------------------------------------------------------------
------------------------------------------------------------------------
------------------------------------------------------------------------
-------------------------------------------------------------CU--A-GU---
------------------------------------------------------------------------
------------------------------------------------------------------------
------------------------------------------------------------------------
------------------------------------------------------------------------
------------------------------------------------------------------------
------------------------------------------------------------------------
------------------------------------------------------------------------
------------------------------------------------------------------------
------------------------------------------------------------------------
------------------------------------------------------------------------
------------------------------------------------------------------------
------------------------------------------------------------------------
------------------------------------------------------------------------
------------------------------------------------GG--U---GUGACUGCGUACCUUU
UGUAUAAUGGGUCAACGACUUAC-A-UUCAGUAGCGA-GCUUAACC-------------GGAUA--------
---G--GGGA--GGCGUAGGGAAAC-C---------------------------------------------
```

```
------------------------------------------------------------------------
------------------------------------------------------------------------
---------------------------------------------------------------------GAG
UCU------------------------------------UAAUA-------------------------GG-
G-C-GA----------------------UGAGUUGCUGGGUGUA-GACCCGAAACC-GAGUGAUCUAUCCAU
GGCCAGGUUGAAGGUGCC-GUAACA-GGUACUGGAGGACCGA--ACCCACGC-AUGUUGCAAAAUGC-GGGG
AUGAGCUGUGGGUAGGGGUGAAAGGCUAAACAAACUCGG-AGAUAGCUGGUUCUCCCCGAAA-ACUAUUUAG
GUAGUGCCUCGA-GCAA-------------------------------------------------------
----------------------GA-----CA-CUGAUGGGGGUAAAGC-ACUGUUA-UGG-CUA--GGGGGU
U---AUUGC--AACUUACCAACCCAUGGCAAACUCAGAAUACCAUC--------------------------
------------------------------------------------------------------------
------------------AAGU-GGUU--------CCUCGGG-AGACAGA-C-AGCG-G-GUGCUAACGUCC
GU-U-GUCAAGAG-GGAAAC---AACCCAGACCGCCG-GCUAAGGUCCCAAAUG--AU-AGA-UU-AA---G
-U-GGU-------AAACGAAGUGGGAAGGCACAGACAGCCAGGAUGUUGGCUUAGAAGCAGCCA-UCAUUUA
AAGAAAGCGUAAUAGCUCACUGGUC-----GAGUCGUCCUGCGCGGAAGAUGUAAC-GGGGCUCA-AAUCUA
U----------AACCGAA-GCUGCGGAUGCCGGU--U-----------------------------------
------------------------------------------------------------------------
------------------------------------------------------------------------
---------------------------------UACC-GGCA-UGGUAGGGGAGC-GUUCUGU---------
-----------AGG-CUGAU-GAAGGUG--C-AUUGUAAAGU--G-UGCUGGAGGU-A-UCA-GAAGUGCGA
AUGU-UGACAU-GAGUA-GCGAUA---AAGC-GGGUGA--A---------------AAGCCC-GCU-CGCCG
A-AAGCCC-AAGGUUUCC-UACGCAACG-UUCAUCGGCGUAGGGUGAGUCGGC-CC-CUAAGGCGAGGCAG-
A-A-AUGC-----GUA--------------------------------------------------------
----------------------------------------GUC-GA-U--GGGA-AACAGGUU-AAUA-UUC
CUGUA----------------------------------CUUGA----UUCA--------A-AU--------
------------------------------------------------------------------------
----------------------------------------------------------GCGAU-GUGGGGAC
GGAGAA--------------GGUUAG-GUUGGCAAG-CU-G----UUGGA------------AUAGCUUGU-
UUAAGC-CGG-UA-----G-G-------------------UG-----GA------------------AG-AC
U-UAGG--------------------CAAAU-CCGGGUUUC-------UUAACACCG----AGA-AG-U-G-
AUG--AC---------------------------------------------------------------GA
-----GUGUC--UACGGACACGAAG-C-AAC-C-GAUACCACGCUUC-C-AGGAAA-AGCCA-CUAAGCUU-
-------------------------------------------------------CAGUUUG-AA-UCGAAC
CGUACCGCAAACCGAC------------------------------ACAGGUG--GGCAGGAU-GAGAAUUC
UAAGGC-G-CU-UGAG-AGAACUCG-G-GAGAAGGAACUCGGC-AAAUUGAUACCGUAACUUCGG-GAGAAG
GUAUG-CCCU-C-UA--AGGUUAA-----GGA-----------CUUGC------------------------
------------------------------------------------------------------------
------------------------------------------------------------------------
------------------------------------------------------------------------
------------------------------------------------------------------------
------------------------------------------------------------------------
------------------------------------------------------------------------
------------------------------------------------------------------------
------------------------------------------------------------------------
------------------------------------------------------------------------
```

```
------------------------------------------------------------------------
------------------------------------------------------------------------
------------------------------------------------------------------------
------------------------------------------------------------------------
--------------------------------------------------------UCCG-UAA--GCCCCG
GAG--GGUCGCAGAGAAUAGGUGGCUGCGA-----CUGUUUAUUAAAAACACAGCACUCUGCCAACACG---
-AA-A-------------GUGGACGU-AUAGGGUGUGA-CGCCUGCCCGGUGCCG-G-AAGG-UUAAUUGAA
GAUGU----------GCAA----------GCAUCGGAUCGAAGCCC--CGGUAAAC-GGCGGCCGUAACUAU
AACGGUCCUAAGGUAGCGAAAUUCCUUGUCGGG-UAAGUUCCGACCCGCACGAAUGGCGUAACGAUG--GCC
ACACUGUCUCCUCC----CGA-GACUCAGCGAAGUU--GAA---GUGGUUGUGAAGAUGCAAUCUACCCGCU
GCUAGACGGAAAGA-CCCCGUGAACC-UUUACU--GUAGCUU-UGCAUUGGACUUUGAAGUCACUUGUGUAG
GAU-AGGUGGGAGGCUUGG----------AAGCAGA-GAC----------------------------GCCA
--------------------------------------GU-CUCUGUGGAG-U--C-GU-CCUUGAAAUA-C
CACC-CUGGUGUCUUUGAGGUUCUAACCCAGA--CCCG-U-CA-U-C-------------------------
------------------------------------------------------------------------
------------------------------------------------------------------------
------------------------------------------------------------------------
------------------------------------------------------------------------
------------------------------------------------------CG----GG----------
------------------------------------------------UCGG-----GGACC-GUGCAUGGU
AGGCAGUUUGACUGGGGCGGU-CUCCUCCCA-AAGCGUAAC-GGAGGAGUUCG-AAGGUUACCUAGGUCCGG
UCGGAAAUCGGACUGA-UAGUG-CAAUGGCAA-AAGGUAGCUUAACU-GCGA--------------------
--------------GACCGA---CAAGUCGGGCAGGUGCGAAA-GCAGGACAUAGUGAUCCGGU--------
---GG-U-----UCUGU-AUG--------------------GA-AGGGCCAUCGCUCAACGG-AUAAAAGGU
ACUCCGGGG-AUAACAGGCUGAUUCCGCCCAAGAGUUCAUAUCGACGGCGGAGUUUGGCACCUCGAUGUCGG
CUCAUCACAUCCUGGGGCUGUA-GUCGGUCCCAAGGGUAUGGCUGUUCGCCAUUUAAAGUGGUACGUGAGCU
GGGUUUAAAACGUCGUGAGACAGUUUGGUCCCUAUCUGCA--------------------G-UGG-------
-------------------------------GCGU--UGGAAGUUUGACGGG-GGC-UGCUCCUAGUACGAG
AGGAC-CGGAGUGGACGAACCUC-UGGU-G-UACCGGUUGUAACGCCAGUUGCAU-A-GCCGGGUAGCUAAG
UUCGGAAGAGAUAAGCGCUGAAAGCA-UC--UAAGCGCGAAACUCGCCUGAA-G-AUG--AGACUUCC-CU-
--UGCGG-----------------------------------------------------------------
--------------------------UU--------UAA---------------------------------
------------------------------------------------------------------------
------------------------------------------------------------------------
------------------------------------------------------------------------
------------------------------------------------------------------------
------------------------------------------------------------------------
------------------------------------------------------------------------
-------------------------------------------------CCGCACU--AA-AGG------GU
CGUUCGAGACCAGGACGU-UGA-UAGGUG-GGG-UGUGGAAG-CGC-G----------------GU------
----AA-----------CGCGUGAAGCUA--A-CCCA-UACUAAUU---GCCCGUGAG-GCUUGACUC----
------------------------------------------------------------------------
------------------------------------------------------------------------
------------------------------------------------------------------------
------------------------------------U------------
```

```
>LSUb_296591_POLSJ/1-2879
---------------------G-GUCAAGUG----AA--UAAGAGCACAU-GGUGGAUG-C-CUUGGC-AAU
GAU---AGGCGACGAAAGACGUGA--UAGCCUGCGAUAAGCUUCGGGGAGCUGGCAAAUUAGCU-----UUG
AUC-CGG-AGAUUUCUGAAUGGGGAAAC--------CC--------AC----CC----------------UU
C-G-------------------------------GGG----U-A-UCGC-AUA-----CUGA----------
-----------A-----------UACA----------------------------------------UA-GG
U-A--------------------UG-C-GA-GGCGAACCGGGUGAACUGAAACAUCUCAGUAGCUC--GAGG
AAAAUACAUCAA--CCGAGAUUCCGAAA--GU-AG-UGGCGAGCGAAAUCGGA-G-AA-GCCUG--------
------------------------------------------------------------------------
----------UUA-------G----UGA---------------------UA--GCA-----------AG---
A-------CUCUU-AGCAG------AAC-AG-CCUG-------------------------------------
-------GAA-AG-GCUGG-------CCAUAGCGGGUGAUAGCCCCGUA---------UGCAAAA----AGA
G------A--CUUGU--------GGUACU---GAGC--UA-AC------GAAAAGUAGGGCGGGACACGAGA
AAUCCUGUCUGAA-UAUGGGGGG-ACC--AUCCUCCAAGGCUAAAUA--CUC-AUCAU-UGACCGAUAGU-G
AAC-U-AGUACCGUGAGGGAAAG-GCGAAA-AGAAC-CCCGG-GAGGGG-A-GUGAAAUAGAUCCUGAAACC
G-UGUGCUUACAA-AAA-GU-AGGA---GCCU-C--------------------------------------
------------------------------------------------------------------------
------------------------------------------------------------------------
------------------------------------------------------------------------
------------------------------------------------------------------------
------------------------------------------------------------------------
------------------------------------------------------------------------
------------------------------------------------------------------------
------------------------------------------------------------------------
--------------------------------------------------------------G--C-AA---
------------------------------------------------------------------------
------------------------------------------------------------------------
------------------------------------------------------------------------
------------------------------------------------------------------------
------------------------------------------------------------------------
------------------------------------------------------------------------
------------------------------------------------------------------------
------------------------------------------------------------------------
------------------------------------------------------------------------
------------------------------------------------------------------------
------------------------------------------------------------------------
------------------------------------------------------------------------
------------------------------------------------GG--G---GUGACUGCGUACCUUU
UGUAUAAUGGGUCAGCGACUUAC-A-UUCAGUGGCAA-GGUUAACC-------------GAAUA--------
---G--GGUA--GCCGUAGAGAAAU-C---------------------------------------------
------------------------------------------------------------------------
------------------------------------------------------------------------
---------------------------------------------------------------------GAG
UCC------------------------------------GAAUA-------------------------GG-
G-C-GA----------------------UCAGUCGCUGGGUGUA-GACCCGAAACC-AAGUGAGCUAUCCAU
```

```
GGCCAGGAUGAAGGUGCG-GUAACA-CGCACUGGAGGUCCGA--ACCGACUA-GUGUUGCAAAACUA-GCGG
AUGAGCUGUGGAUAGGGGUGAAAGGCUAAACAAACUUGG-AAAUAGCUGGUUCUCUCCGAAA-ACUAUUUAG
GUAGUGCCUCAA-GUA--------------------------------------------------------
-----------------------U-----UA-CCGUCGGGGGUAGAGC-ACUGUUU-UGG-CUA--GGGGGU
C---AUGGC--GACUUACCAAACCAAUGCAAACUCCGAAUACCGAC--------------------------
------------------------------------------------------------------------
------------------GAGU-A-CA--------GCUUGGG-AGACAGAGC-ACCG-G-GUGCUAACGUCC
GG-A-CUCAAGAG-GGAAAC---AACCCAGACCGCCA-GCUAAGGUCCCUAAAA--U--UGG-CU-AA---G
-U-GGG-------AAACGAAGUGGGAAGGCUAUAACAGUCAGGAUGUUGGCUUAGAAGCAGCCA-UCAUUUA
AAGAAAGCGUAAUAGCUCACUGAUC-----GAGUCGUCCUGCGCGGAAGAUGUAAC-GGGGCU-A-AGCCAG
U----------UACCGAA-GCUGCGGAUUUGCAA--U-----------------------------------
------------------------------------------------------------------------
------------------------------------------------------------------------
-------------------------------UUAUUG-CAAG-UGGUAGGAGAGC-GUUCUGU---------
-----------AAG-CCUGU-GAAGGUG--U-GGUGUAAACC--C-UGCUGGAGGU-A-UCA-GAAGUGCGA
AUGC-UGACAU-GAGUA-GCGUUA---AAGG-GGGUGA--A---------------AAGCCC-CCU-CGCCG
U-AAGCGC-AAGGUUUUC-UACGCAACG-UUCAUCGGCGUAGAGUGAGUCGGC-CC-CUAAGGCGAGGCAG-
A-G-AUGC-----GUA--------------------------------------------------------
----------------------------------------GCU-GA-U--GGGA-AACAGGUC-AAUA-UUC
CUGUA----------------------------------CCGAU----GUGU--------A-GU--------
------------------------------------------------------------------------
----------------------------------------------------------GCGAU-GUGGGGAC
GGAGAA--------------GGUUAA-CUCAGCCAA-CU-G----UUGGA------------UAUGUUGGU-
UCAAGC-CUG-UA-----G-U-------------------CG-----UG------------------CC-UG
G-UAGG--------------------UAAAU-CCGCCGGGCU------U---AGAUG----AGG-GG-U-G-
AUA--AC---------------------------------------------------------------GA
-----GUCUG--CUUGCAGACGAAG-U-GAG-U-GAUACCCUGCUUC-C-AGGAAA-AGCCA-CUAAGCUU-
--------------------------------------------------------CAGCUA-CA-CACGAC
CGUACCGCAAACCGAC------------------------------ACUGGUG--CGCGAGAU-GAGUAUUC
UAAGGC-G-CU-UGAG-AGAACUCA-G-GAGAAGGAACUCGGC-AAAUUGACACCGUAACUUCGG-AAGAAG
GUGUG-CCUU-UAGU--AGGUGAAC----CA------------UUUAC------------------------
------------------------------------------------------------------------
------------------------------------------------------------------------
------------------------------------------------------------------------
------------------------------------------------------------------------
------------------------------------------------------------------------
------------------------------------------------------------------------
------------------------------------------------------------------------
------------------------------------------------------------------------
------------------------------------------------------------------------
------------------------------------------------------------------------
------------------------------------------------------------------------
------------------------------------------------------------------------
------------------------------------------------------------------------
----------------------------------------------------------UUGG-GGAGCCCAA
```

```
UGA--GGCUGCAAAAAAUCGGUGGCUGCAA-----CUGUUUAUUAAAAACACAGCACUCUGCUAAGACG---
-AA-A-------------GUCGACGU-AUAGGGUGUGA-CGCCUGCCCGGUGCUG-G-AAGA-UUAAAUGAU
GGGGU----------GCAA----------GCUCUUGAUUGAAGUCC--CAGUAAAC-GGCGGCCGUAACUAU
AACGGUCCUAAGGUAGCGAAAUUCCUUGUCGGG-UAAGUUCCGACCUGCACGAAUGGCGUAAUGAUG--GCC
ACACUGUCUCCUCC----UGA-GACUCAGCGAAGUU--GAA---AUGUUUGUGAUGAUGCAAUCUCCCCGCG
GAAAGACGGAAAGA-CCCCAUGAACC-UUUACU--GUAGCUU-UGUAUUGGACUUUGAACAGAUCUGUGUAG
GAU-AGGUGGGAGGCUUUG----------AAGCCCG-GUC----------------------------GCUA
--------------------------------------GA-UCGGGUGGAG-C--C-AA-CGUUGAAAUA-C
CACC-CUGGUGUGUUUGAGGUUCUAACCUAGG--UCCA-U-UA-U-C-------------------------
------------------------------------------------------------------------
------------------------------------------------------------------------
------------------------------------------------------------------------
------------------------------------------------------------------------
------------------------------------------------------UG----GA----------
------------------------------------------------UCGG-----GGACA-GUGCAUGGU
AGGCAGUUUGACUGGGGCGGU-CUCCUCCCA-AAGUGUAAC-GGAGGAGUUCG-AAGGUACGCUAGGUACGG
UCGGACAUCGUGCUAA-UAGUG-CAAUGGCAU-AAGCGUGCUUAACU-GCGA--------------------
--------------GACUGA---CAAGUCGAGCAGAUGCGAAA-GCAGGACAUAGUGAUCCGGU--------
---GG-U-----UCUGU-AUG--------------------GA-AGGGCCAUCGCUCAACGG-AUAAAAGGU
ACUCUGGGG-AUAACAGGCUGAUACCGCCCAAGAGUUCAUAUCGACGGCGGUGUUUGGCACCUCGAUGUCGG
CUCAUCUCAUCCUGGGGCUGUA-GCCGGUCCCAAGGGUAUGGCUGUUCGCCAUUUAAAGAGGUACGUGAGCU
GGGUUUAAAACGUCGUGAGACAGUUUGGUCCCUAUCUUCC--------------------G-UGG-------
-------------------------------GCGC--UGCAGAUUUGAGGAA-GCC-UGCUCCUAGUACGAG
AGGAC-CGGAGUGGACGAACCUC-UGGU-G-UACCGGUUGUCACGCCAGUGGCAU-U-GCCGGGUAGCUAAG
UUCGGAAGAGAUAACCGCUGAAAGCA-UC--UAAGCGGGAAACUCGUUUCAA-G-AUU--AGAUCUGC-C--
--GGGGC-----------------------------------------------------------------
--------------------------CU--------UGA---------------------------------
------------------------------------------------------------------------
------------------------------------------------------------------------
------------------------------------------------------------------------
------------------------------------------------------------------------
------------------------------------------------------------------------
------------------------------------------------------------------------
-------------------------------------------------GCCCCCU--GA-AGA------GU
CGUUCAAGACCAGGACGU-UGA-UAGGUC-AGG-UGUGGAAG-CGC-A----------------GU------
----AA-----------UGCGUUAAGCUA--A-CUGA-UACUAAU---GCUCGUGCG-GCUUGACCC-----
------------------------------------------------------------------------
------------------------------------------------------------------------
------------------------------------------------------------------------
------------------------------------U------------
>LSUb_663917_GEOSK/1-2958
---------------------G-GUCAAGCU----AC--UAAGGGCGUAC-GGUGGAUG-C-CUAGGC-AGA
GAG---AGGCGAUGAAGGACGUGG--UAAGCUGCGAAAAGCCUCGGGGAGCCGCUAAACAGGCU-----UUG
AUC-CGG-GGGUGUCCGAAUGGGGAAAC--------CC--------GG----CGGA-GGU-----------C
A-U-------------G---C----U----C---CGU----C-A-UCGU-GCA-----CUGA----------
```

```
-----------A-----------UCCA----------------------------------------UA-GG
U-G--------------------CA-U-GA-AGCGAACGGGGGGAACUGAAACAUCUAAGUACCCC--CAGG
AAAAGAAAACAA--UAGUGAUUCCGUCA--GU-AG-UGGCGAGCGAAAGCGGA-A-CA-GCCCAAACCGAGA
-GGA-CUACG--------------------------------G-UCCUUUCGGGG--UUG------------
-----UGGG-ACC-------CCGA-CG----------------------UG--GGA---------UUGA-UG
A-------UGGGU-AGCUA------AAC-GG-UCUG------------------------------------
-------GAA-AG-UCCGG-------CCAUAGUGGGUGAUAGCCCGUA----------AGCGAAA----CCU
U------G--AUUCA----------CCCU---AGGG--GU-AU------CCCGAGUACCGCGGGACACGUGG
AAUCCCGUGGGAA-GCUGGGAGG-ACC--AUCUCCCAAGGCUAAAUA--CUA-CUCUC-UGACCGAUAGU-G
CAC-A-AGUACCGUGAGGGAAAG-GUGAAA-AGUAC-UCCGAUGAGGAG-G-GUGAAAUAGAACCUGAAACC
G-UAUGCCUACAA-GCA-GU-GGGA---GCAC-U---AU---------------------------------
------------------------------------------------------------------------
------------------------------------------------------------------------
------------------------------------------------------------------------
------------------------------------------------------------------------
------------------------------------------------------------------------
------------------------------------------------------------G-----------
------------------------------------------------------------------------
------------------------------------------------------------------------
-------------------------------------------------------------UA--U-AU---
------------------------------------------------------------------------
------------------------------------------------------------------------
------------------------------------------------------------------------
------------------------------------------------------------------------
------------------------------------------------------------------------
------------------------------------------------------------------------
------------------------------------------------------------------------
------------------------------------------------------------------------
------------------------------------------------------------------------
------------------------------------------------------------------------
------------------------------------------------------------------------
------------------------------------------------------------------------
---------------------------C-------------------AAG--U---GUGACCGCGUGCCUUU
UGCAUAAUGAGUCAGCGAGUUAC-U-CUCGGCAGCGA-GGUUAAGC--U----------CAUAG--------
--AG--UGGA--GCCGCAGCGAAAG-C--------------------------------------------
------------------------------------------------------------------------
------------------------------------------------------------------------
---------------------------------------------------------------------GAG
UCU------------------------------------GAAUA-------------------------GG-
G-C-GU---------------------UUGAGUUGCCGGGAGUA-GACCCGAAACC-GGGUGAUCUAUCCAU
GUCCAGGGUGAAAGGAAG-GUAACA-CUUCGUGGAGGCCCGA--ACCCACUG-GCGUUGAAAAGCCA-GGGG
AUGAGGUGUGGAUAGGAGUGAAAGGCUAAUCAAACUCGG-AGAUAGCUGGUUCUCCCCGAAA-UAUAUUUAG
GUAUAGCCUCGU-GAA--------------------------------------------------------
----------------------GU-----AA-GUAGUGGGGGUAGAGC-ACUGAAU-GGG-CUA--GGGGUC
C---UA-CC--AGAUUACCAAACCCAAUCAAACUCCGAAUACCACU--------------------------
```

```
------------------------------------------------------------------------
------------------AACU-GUUA--------UCACGGG-AGUCAGA-C-UGCG-G-GUGAUAAGAUCC
GU-A-GUCAAAAG-GGAAAG---AGCCCAGAUCGUCG-GCUAAGGUCCCAAAAU--CC-AUG-CU-AA---G
-U-GGA-------AAACGAUGUGGAAAUGCCCAGACAACCAGGAGGUUGGCUUAGAAGCAGCCA-CCCUUUA
AAGAAAGCGUAAUAGCUCACUGGUC-----GAGUGGGUCUGCGCGGAAAAUGUAAC-GGGGCUCA-AGCAUG
G----------UACCGAA-GCCACGGGUUGAUAC--C-----------------------------------
------------------------------------------------------------------------
-----UU-----------------------------------------------------------------
-------------------------------AAGGUA-UCAG-CGGUAGGGGAGC-AUUGUGU---------
-----------AAG-CCUGC-GAAGGUC--G-ACCGCGAGGA--C-GGCUGGAGGU-A-UCA-CAAGAGAUU
AUGC-UGACAU-GAGUA-GCGAAA---AUGC-GGGUGA--G---------------AAACCC-GCA-CACCG
A-AAACCC-AAGGGUUCC-UCAGUAAAG-GUAAUCUGCUGAGGGUUAGUCGGU-CC-CUAAGGCGAGGCCG-
A-A-AGGC-----GUA--------------------------------------------------------
----------------------------------------GUU-GA-U--GGGA-AACAGGUU-AAUA-UUC
CUGUA----------------------------------CCACU----UGUC--------G-CU--------
------------------------------------------------------------------------
----------------------------------------------------------GCGAU-GGGGGGAC
GGAGAA--------------GGGUAG-GCGAUCCGG-GU-G----CUGGA------------CGUCCCGGU-
UCAAGC-GUG-UA-----G-G-------------------CG-----GG------------------AA-GA
G-GAGG--------------------CAAAU-CCCUUCUUCCA----UACAACGCCA----AGG-CG-U-G-
AUG--AC---------------------------------------------------------------GA
-----GGGGG--UAUCCCCACAAAG-U-CGU-U-GAUCCCAUGCUUC-C-AAGAAA-AGCCU-CUAAGCUU-
--------------------------------------------------------CAGGCG-CA-GGUGAC
CGUACCGUAAACCGAC------------------------------UCAGGUG--GGUGAGGAUUAAUGUCC
UAAGGU-G-CU-UGAG-AGAACACU-G-GUUAAGGAACUCGGC-AAAUUGACACCGUAACUUCGG-GAGAAG
GUGUG-CCCU-C-AC-UAGGUGUAG----CGA-----------UUCGC------------------------
------------------------------------------------------------------------
------------------------------------------------------------------------
------------------------------------------------------------------------
------------------------------------------------------------------------
------------------------------------------------------------------------
------------------------------------------------------------------------
------------------------------------------------------------------------
------------------------------------------------------------------------
------------------------------------------------------------------------
------------------------------------------------------------------------
------------------------------------------------------------------------
------------------------------------------------------------------------
------------------------------------------------------------------------
--------------------------------------------------------ACGU-GAA--GCCGAA
GAG--GGUCGCAGAGAAAUGGCGGUAGCGA-----CUGUUUACUAAAAACACAGGACUCUGCUAAGUCG---
-CA-A-------------GACGAUGU-AUAGGGUCUGA-CGCCUGCCCGGUGCCG-G-AAGG-UUAAGGGGA
UUUGUUA-GC-----GCAA-----GCGAAGCUUUGAACCGAAGCCC--CGGUAAAC-GGCGGCCGUAACUAU
AACGGUCCUAAGGUAGCGAAAUUCCUUGUCGGG-UAAGUUCCGACCUGCACGAAUGGCGUAACGAUU--UCC
GCGCUGUCUCAACC----AGU-GGCUCAGCGAAAUU--GAA---UUCUCGGUGAAGAUGCCGAGUACCCGCG
```

```
GCAAGACGGAAAGA-CCCCGUGAACC-UUUACU--ACAGCUU-GACAGUGACAUUCGGAAUAGCUUGUGUAG
GAU-AGGUGGGAGGCUGUG----------AAACCGG-GAC----------------------------GCCA
--------------------------------------GU-UUCGGUGGAG-C--C-AA-CCUUGAAAUA-C
CACC-CUGGUUGUUUUGGAUGUCUAACCCAGG--CCCG-U-CA-U-C-------------------------
------------------------------------------------------------------------
------------------------------------------------------------------------
------------------------------------------------------------------------
------------------------------------------------------------------------
------------------------------------------------------CG----GG----------
------------------------------------------------UCGG-----GGACA-CUGUCUGGU
GGGUAGUUUGACUGGGGCGGU-CGCCUCCCA-AAGCGUAAC-GGAGGCGCGCG-AAGGUUCCCUCAGGCUGA
UUGGAAACCAGCCGUA-GAGUG-CAAAGGCAU-AAGGGAGCUUGACU-GCGA--------------------
--------------GACAGA---CACGUCGAGCAGGUACGAAA-GUAGGCCUUAGUGAUCCGGC--------
---GG-U-----UCUGU-AUG--------------------GA-AGGGCCGUCGCUCAACGG-AUAAAAGGU
ACUCCGGGG-AUAACAGGCUGAUCUCCCCCAAGAGUUCACAUCGACGGGGAGGUUUGGCACCUCGAUGUCGG
CUCAUCGCAUCCUGGGGCUGAA-GUCGGUCCCAAGGGUUUGGCUGUUCGCCAAUUAAAGCGGUACGCGAGCU
GGGUUUAAAACGUCGUGAGACAGUUUGGUCCCUAUCUGCC--------------------G-UGG-------
-------------------------------GCGC--AGGAUACUUGAGAAG-AGC-UGUCCUUAGUACGAG
AGGAC-CGGGAUGGACGUACCUC-UAGU-G-UUCCAGUUGUUCCGCCAGGAGCAU-UCGCUGGGUAGCUAUG
UACGGAAAGGAUAACCGCUGAAAGCA-UC--UAAGCGGGAAGCCUCCUUCAA-G-AUU--AGGUAUCC-CC-
--GGGAC-----------------------------------------------------------------
--------------------------GC---------GA---------------------------------
------------------------------------------------------------------------
------------------------------------------------------------------------
------------------------------------------------------------------------
------------------------------------------------------------------------
------------------------------------------------------------------------
------------------------------------------------------------------------
-------------------------------------------------GUCCCCU--GA-AGG------CC
CGUUGUAGACCACAACGU-UGA-UAGGCC-GGG-UGUGUAAG-CAG-G----------------GU------
----AA-----------CCUGCUCAGCUG--A-CCGG-UACUAAUC---GGCCGUGAG-GCUUGACCA----
------------------------------------------------------------------------
------------------------------------------------------------------------
------------------------------------------------------------------------
------------------------------------U------------
>LSUb_176299_AGRFC/1-2771
---------------------A--AGAAGUG----AA-UUAAGGGCAUUU-GGUGGAUG-C-CUUGGC-AUG
CAC---AGGCGAAGAAGGACGUGA--UACGCUGCGAAAAGCCGUGGGGAGCUGCGAAUAAGCUU------UG
AUC-CAU-GGAUCUCCGAAUGGGGCAAC--------CC--------AC----CUUA-A-AUGCUUGGGAA-A
U-CU---UUCCAAGCAU---UGUGAU----A---AGG----U-A-UCUU-ACUU----UCGA----------
-----------A-----------UACA----------------------------------------UA-GG
G-G--------------------UA-A-GA-AGCGAACGCAGGGAACUGAAACAUCUAAGUACCUG--CAGG
AAAGGACAUCAA--CCGAGACUCCGCAA--GU-AG-UGGCGAGCGAACGCGGA-C-CA-GGCCA--------
------------------------------------------------------------------------
----------GUG-------G----------------------------CA--AUG-----------AU---
```

```
G-------AAUAA-AGCGG------AAC-GA-UUUG------------------------------------
-------GAA-AA-GUCGG-------CCAUAGAGGGUGAUAGCCCCUUA---------CGCGUAG----AAC
A------G--UCAU------------------------UG-UC-------CUUGAGUAGGGCGGACACGUGA
AAUCCUGUCUGAA-CAUGGGGAG-ACC--ACUCUCCAAGCCUAAGUA--CUC-GUGCA-UGACCGAUAGC-G
AAC-A-AGUACCGUGAGGGAAAG-GUGAAA-AGCAC-CCCGACGAGGGG-A-GUGAAAUAGAACCUGAAACC
G-GAUGCCUACAA-ACA-GU-CGGA---GCCC----------------------------------------
------------------------------------------------------------------------
------------------------------------------------------------------------
------------------------------------------------------------------------
------------------------------------------------------------------------
------------------------------------------------------------------------
------------------------------------------------------------------------
------------------------------------------------------------------------
------------------------------------------------------------------------
--------------------------------------------------------------G--C-AA---
------------------------------------------------------------------------
------------------------------------------------------------------------
------------------------------------------------------------------------
------------------------------------------------------------------------
------------------------------------------------------------------------
------------------------------------------------------------------------
------------------------------------------------------------------------
------------------------------------------------------------------------
------------------------------------------------------------------------
------------------------------------------------------------------------
------------------------------------------------------------------------
------------------------------------------------------------------------
-------------------------------------------------G--G---GUGACGGCGUACCUUU
UGUAUAAUGGGUCAACGACUUAG-U-GUAACUAGCAA-GCUUAAGC--C-----------GGUA--------
--GG--UGUA--GGCGCAGCGAAAG-C---------------------------------------------
------------------------------------------------------------------------
------------------------------------------------------------------------
---------------------------------------------------------------------GAG
UCU------------------------------------GAAUA-------------------------GG-
G-C-GA---------------------UUGAGUUAGUUGCAUUA-GACCCGAAACC-GAGUGAUCUAGCCAU
GAGCAGGCUGAAGGUUGG-GUAACA-CCAACUGGAGGGCCGA--ACCCGUAU-CUGUUGCAAUAGAU-UGGG
AUGACUUGUGGCUAGGGGUGAAAGGCCAAUCAAACUCGG-AAAUAGCUGGUUCUCCGCGAAA-UCUAUUUAG
GUAGAGCGUCGA-CCG--------------------------------------------------------
----------------------AA-----UA-CCCUCGGGGGUAGAGC-ACUGGAU-GGG-CUA--UGGGGA
C---UCACC--GUCUUACUGAUCCUAACCAAACUCCGAAUACCGAG--------------------------
------------------------------------------------------------------------
------------------GAGU-A-CU--------AGUCGGC-AGACACA-C-GGCG-G-GUGCUAACGUCC
GU-C-GUGAAAAG-GGCAAC---AACCCUGACCUCCA-GCUAAGGUCCCCAAGU--CA-UGG-CU-AA---G
-U-GGG-------AAAGGAUGUGAGGAUCCCAAAACAACCAGGAUGUUGGCUUAGAAGCAGCCA-UCAUUUA
AAGAAAGCGUAACAGCUCACUGGUCUAAAUAAGGG-UCUUGCGCCGAAAAUGUAAC-GGGGCUAA-AGCCAU
```

```
G----------CACCGAA-GCUGAGGAUUUGCAG--U-----------------------------------
------------------------------------------------------------------------
------------------------------------------------------------------------
---------------------------------AAUG-CAAG-UGGUAGCGGAGC-GUUCCGU---------
-----------AAG-UCUGU-GAAGGCG--GACCCGUGAGGG--C-UGCUGGAGAU-A-UCG-GAAGUGCGA
AUGU-UGACAU-GAGUA-ACGAUA---AAGG-GAGUGA--G---------------AGACUC-CCU-CGCCG
A-AAGACC-AAGGGUUCC-UGCUUAAAG-UUAAUCUGAGCAGGGUUAGCCGGC-CC-CUAAGACGAGGCGG-
A-C-ACGC-----GUA--------------------------------------------------------
----------------------------------------GUC-GA-U--GGGA-ACCACGUU-AAUA-UUC
GUGGG----------------------------------CCUGG----UG----------------------
------------------------------------------------------------------------
----------------------------------------------------------------GUAGUGAC
GGAUCU--------------CGUG-U-GUUGUACAA-CC-U----UAUUG-------------GAUUGGUUG-
------------------------------------------------------------------------
------------------------------------------------------------------------
------------------------------------------------------------------------
-----------------------UG-C-GGC-G----AAG-AGGUUC-C-AGGAAAUAGCUC-CACC-----
---------------------------------------------------------------GU-AUAGAC
CGUACCCGAAACCGAC------------------------------ACAGGUG--GUCAGGUA-GAGUAUAC
CAAGGC-G-CU-UGAG-AGAACUAU-G-UUGAAGGAACUCGGC-AAAUUGCACGCGUAACUUCGG-AAGAAG
CGUGA-CCCC-A-UU--UUAGG--------------------------------------------------
------------------------------------------------------------------------
------------------------------------------------------------------------
------------------------------------------------------------------------
------------------------------------------------------------------------
------------------------------------------------------------------------
------------------------------------------------------------------------
------------------------------------------------------------------------
------------------------------------------------------------------------
------------------------------------------------------------------------
------------------------------------------------------------------------
------------------------------------------------------------------------
------------------------------------------------------------------------
------------------------------------------------------------------------
-------------------------------------------------------------CAA--CUAUGA
UGG--GGUGGCACAGACCAGGGGGUAGCGA-----CUGUUUAUCAAAAACACAGGGCUCUGCGAAGUAG---
-CA-A-------------UACGACGU-AUAGGGUCUGA-CGCCUGCCCGGUGCUG-G-AAGG-UUAAAGGGA
GGGGU----------GCAA----------GCUCUGAACUGAAGCCC--CAGUAAAC-GGCGGCCGUAACUAU
AACGGUCCUAAGGUAGCGAAAUUCCUUGUCGGG-UAAGUUCCGACCUGCACGAAUGGCGUAACGACU--UCC
CCGCUGUCUCCAAC----AUA-GACUCAGUGAAAUU--GAA---UUCCCCGUGAAGAUGCGGGGUUCCUGCG
GUCAGACGGAAAGA-CCCCGUGCACC-UUUACU--AUAGCUU-UACACUGGCAUUCGCCAAGGCAUGUGUAG
GAU-AGGUGGUAGGCUUUG----------AAGCAGG-GAC----------------------------GCCA
-------------------------------------GU-UUCUGUGGA---G--CCAU-CCUUGAAAUA-C
CACC-CUUAUCUUCGUGGAUGUCUAACCGCGG--UCCG-U-UA-U-C-------------------------
------------------------------------------------------------------------
```

------------------------------------------------------------------------
------------------------------------------------------------------------
------------------------------------------------------------------------
------------------------------------------------------CG----GA----------
------------------------------------------------UCCG-----GGACA-GUGUAUGGU
GGGUAGUUUGACUGGGGCGGU-CGCCUCCGA-AAGAGUAAC-GGAGGCGCGCG-AUGGUUAGCUCAGACCGG
UCGGAAAUCGGUCGUC-GAGUG-CAAUGGCAU-AAGCUAGCCUGACU-GCGA--------------------
--------------GACUGA---CAAGUCGAGCAGAGACGAAA-GUCGGUCAUAGUGAUCCGGU--------
---GG-U-----CCCGU-GUG--------------------GA-AGGGCCAUCGCUCAACGG-AUAAAAGGU
ACGCCGGGG-AUAACAGGCUGAUGACCCCCAAGAGUCCAUAUCGACGGGGUUGUUGGCACCUCGAUGUCGA
CUCAUCGCAUCCUGGGGCUGGA-GCAGGUCCCAAGGGUAUGGCUGUUCGCCAUUUAAAGCGGUACGUGAGUU
GGGUUCAGAACGUCGUGAGACAGUUCGGUCCCUAUCUGCC--------------------G-UGG-------
-------------------------------GUGU--AGGAAUAUUGACAGG-AUC-UGUCCCUAGUACGAG
AGGAC-CGGGAUGGACGUAUCUC-UGGU-G-GAUCUGUUGUCCUGCCAAGGGCAU-A-GCAGAGUAGCUAUA
UACGGAAUGGAUAACCGCUGAAGGCA-UC--UAAGCGGGAAACCAACCUGAA-A-ACG--AGUGUUCC-CU-
--A---------------------------------------------------------------------
------------------------------------------------------------------------
------------------------------------------------------------------------
------------------------------------------------------------------------
------------------------------------------------------------------------
------------------------------------------------------------------------
------------------------------------------------------------------------
------------------------------------------------------------------------
----------------------------------------------------------UC-AGA------GC
CGUGGAAGACGACCACGU-UGA-UAGGAC-GGG-UGUGGAAG-CAC-A----------------GC------
----AA-----------UGUGUGAAGCUU--A-CCGU-UACUAAUA---GCUCGAUCG-ACUUCUUCG----
------------------------------------------------------------------------
------------------------------------------------------------------------
------------------------------------------------------------------------
------------------------------------U------------
>LSUb_266834_RHIME/1-2769
---------------------G-AUCAAGUG----AA-AGAAGGGCAUUU-GGUGGAUG-C-CUUGGC-AUG
CAC---AGGCGAUGAAGGACGUGA--UACGCUGCGAAAAGCCGUGGGGAGCUGCGAAUGAGCUU------UG
AUC-CAU-GGAUCUCCGAAUGGGGCAAC--------CC--------AC----CUUA-G-AUGCUUGGAA--A
A-U----UUCCAAGCAU---U----GUUGAA---AGG----U-A-UCUU-AUU-----CUGA----------
-----------A-----------UACA----------------------------------------UAG-G
G-A--------------------UA-A-GA-AGCGAACGCAGGGAACUGAAACAUCUAAGUACCUG--CAGG
AAAGGACAUCAA--CCGAGACUCCGCAA--GU-AG-UGGCGAGCGAACGCGGA-C-CA-GGCCA--------
------------------------------------------------------------------------
----------GUG-------G----CA--------------------------AUG-----------AG---
G-------AAUAA-AGUGG------AAC-GA-UCUG------------------------------------
-------GAA-AG-UUCGG-------CCGUAGCGGGUGAUAGCCCGUA----------CACGUAG----AAC
A------C--UCAU------------------------UG-UC------CUUGAGUAAGGCGGGACACGUGA
AAUCCUGUCUGAA-CAUGGGGAG-ACC--ACUCUCCAAGCCUAAGUA--CUC-GUGCA-UGACCGAUAGC-G
AAC-A-AGUACCGUGAGGGAAAG-GUGAAA-AGCAC-CCCGACAAGGGG-A-GUGAAAUAGAACCUGAAACC

```
G-GAUGCCUACAA-ACA-GU-CGGA---GCCC----------------------------------------
------------------------------------------------------------------------
------------------------------------------------------------------------
------------------------------------------------------------------------
------------------------------------------------------------------------
------------------------------------------------------------------------
------------------------------------------------------------------------
------------------------------------------------------------------------
------------------------------------------------------------------------
--------------------------------------------------------------G--C-AA---
------------------------------------------------------------------------
------------------------------------------------------------------------
------------------------------------------------------------------------
------------------------------------------------------------------------
------------------------------------------------------------------------
------------------------------------------------------------------------
------------------------------------------------------------------------
------------------------------------------------------------------------
------------------------------------------------------------------------
------------------------------------------------------------------------
------------------------------------------------------------------------
------------------------------------------------------------------------
-------------------------------------------------G--G---GUGACGGCGUACCUUU
UGUAUAAUGGGUCAACGACUUAG-U-GUGGCAUGCAA-GCUUAAGC--C-----------GGUA--------
--GG--UGGA--GGCGUAGCGAAAG-C---------------------------------------------
------------------------------------------------------------------------
------------------------------------------------------------------------
---------------------------------------------------------------------GAG
UCU------------------------------------GAACA-------------------------GG-
G-C-GC---------------------UUUAGUAUGUCGCAUUA-GACCCGAAACC-GAGUGAUCUAGCCAU
GAGCAGGUUGAAGGUUGG-GUAACA-CCAACUGGAGGACCGA--ACCCGCAU-CUGUUGCAAUAGAU-UGGG
AUGACUUGUGGCUAGGGGUGAAAGGCCAAUCAAACUCGG-AAAUAGCUGGUUCUCCGCGAAA-ACUAUUUAG
GUAGUGCGUCGA-CCG--------------------------------------------------------
----------------------AA-----UA-CCCUCGGGGGUAGAGC-ACUGGAU-GGG-CUA--UGGGGA
C---UCACC--GUCUUACUGAUCCUAACCAAACUCCGAAUACCGAG--------------------------
------------------------------------------------------------------------
------------------GAGU-A-CU---------AGUCGGC-AGACACA-C-GGCG-G-GUGCUAACGUCC
GU-C-GUGAAAAG-GGCAAC---AACCCUGACCUCCA-GCUAAGGUCCCCAAGU--CA-UGG-CU-AA---G
-U-GGG-------AAAGGAUGUGAGGAUCCCAAAACAACCAGGAUGUUGGCUUAGAAGCAGCCA-UCAUUUA
AAGAAAGCGUAACAGCUCACUGGUCUAAAUAAGGGUCUUUGCGCCGAAAAUGUAAC-GGGGCUGA-AGCCAU
G----------CACCGAA-GCUGAGGAUACGUCG--U-----------------------------------
------------------------------------------------------------------------
------------------------------------------------------------------------
---------------------------------AAGA-CGUG-UGGUAGCGGAGC-GUUCCGU---------
-----------AAG-CCAGU-GAAGGGA--UACCCGUGAGGG--G-UCCUGGAGGU-A-UCG-GAAGUGCGA
```

```
AUGU-UGACAU-GAGUA-ACGAUA---AAGA-GGGUGA--G---------------AGACCC-UCU-CGCCG
A-AAGACC-AAGGGUUCC-UGCUUAAAG-UUAAUCUGAGCAGGGUUAGCCGGC-CC-CUAAGACGAGGCGG-
A-C-ACGC-----GUA--------------------------------------------------------
----------------------------------------GUC-GA-U--GGGA-ACCACGUU-AAUA-UUC
GUGGG----------------------------------CCUGG----UG----------------------
------------------------------------------------------------------------
----------------------------------------------------------------GUAGUGAC
GGAUUG--------------CUCA-A-CUUGUCCGG-AC-U----UAUUG------------GAUUGUCCG-
GG----------------------------------------------------------------------
------------------------------------------------------------------------
------------------------------------------------------------------------
--------------------------C-GGG-G----ACG-CGGUUC-C-AGGAAAUAGCUC-CACC-----
---------------------------------------------------------------GU-AUAGAC
CGUACCCGAAACCGAC------------------------------ACAGGUG--GUCAGGUA-GAGUAUAC
CAAGGC-G-CU-UGAG-AGAACUAU-G-UUGAAGGAACUCGGC-AAAUUGCACGCGUAACUUCGG-AAGAAG
CGUGA-CCCU-U-UU--CUACG--------------------------------------------------
------------------------------------------------------------------------
------------------------------------------------------------------------
------------------------------------------------------------------------
------------------------------------------------------------------------
------------------------------------------------------------------------
------------------------------------------------------------------------
------------------------------------------------------------------------
------------------------------------------------------------------------
------------------------------------------------------------------------
------------------------------------------------------------------------
------------------------------------------------------------------------
------------------------------------------------------------------------
------------------------------------------------------------------------
---------------------------------------------------------------CAAGUAGGA
UGG--GGUGGCACAGACCAGGGGGUAGCGA-----CUGUUUAUCAAAAACACAGGGCUCUGCGAAGUCG---
-CA-A-------------GACGACGU-AUAGGGUCUGA-CGCCUGCCCGGUGCUG-G-AAGG-UUAAGAGGA
GGGGU----------GCAA----------GCUCUGAAUCGAAGCCC--CAGUAAAC-GGCGGCCGUAACUAU
AACGGUCCUAAGGUAGCGAAAUUCCUUGUCGGG-UAAGUUCCGACCUGCACGAAUGGCGUAACGACU--UCC
CCGCUGUCUCCAAC----AUA-GACUCAGUGAAAUU--GAA---UUCCCCGUGAAGAUGCGGGGUUCCUGCG
GUCAGACGGAAAGA-CCCCGUGCACC-UUUACU--AUAGCUU-UACACUGGCAUUCGUGUCGGCAUGUGUAG
GAU-AGGUGGUAGGCUUUG----------AAGCGG-GAC-----------------------------GCCA
--------------------------------------GU-UCUCGUGGAG-C--C-AU-CCUUGAAAUA-C
CACC-CUUAUCGUCAUGGAUGUCUAACCGCGG--UCCG-U-CA-U-C-------------------------
------------------------------------------------------------------------
------------------------------------------------------------------------
------------------------------------------------------------------------
------------------------------------------------------------------------
------------------------------------------------------CG----GA----------
------------------------------------------------UCCG-----GGACA-GUGUAUGGU
```

```
GGGUAGUUUGACUGGGGCGGU-CGCCUCCGA-AAGAGUAAC-GGAGGCGCGCG-AUGGUGGGCUCAGAGCGG
UCGGAAAUCGCUCGUC-GAGUG-CAAUGGCAU-AAGCCCGCCUGACU-GCGA--------------------
--------------GACUGA---CAAGUCGAGCAGAGACGAAA-GUCGGUCAUAGUGAUCCGGU--------
---GG-U-----CCCGC-GUG--------------------GA-AGGGCCAUCGCUCAACGG-AUAAAAGGU
ACGCCGGGG-AUAACAGGCUGAUGACCCCCAAGAGUCCAUAUCGACGGGGUUGUUUGGCACCUCGAUGUCGG
CUCAUCGCAUCCUGGGGCUGGA-GCAGGUCCCAAGGGUUUGGCUGUUCGCCAAUUAAAGCGGUACGUGAGCU
GGGUUCAGAACGUCGUGAGACAGUUCGGUCCCUAUCUGCC--------------------G-UGG-------
-------------------------------GUGU--AGGAAUAUUGACAGG-AUC-UGUCCCUAGUACGAG
AGGAC-CGGGAUGGACAUAUCUC-UGGU-G-GACCUGUUGUCCUGCCAAGGGCAU-A-GCAGGGUAGCUAUA
UAUGGAAGGGAUAACCGCUGAAGGCA-UC--UAAGCGGGAAACCCACCUGAA-A-ACG--AGUAUUCC-CU-
------------------------------------------------------------------------
------------------------------------------------------------------------
------------------------------------------------------------------------
------------------------------------------------------------------------
------------------------------------------------------------------------
------------------------------------------------------------------------
------------------------------------------------------------------------
------------------------------------------------------------------------
----------------------------------------------------------UG-AGA------AC
CGUGGAAGACGACCACGU-UGA-UAGGCC-GGG-UGUGGAAG-UGC-G----------------GC------
----AA-----------CGCAUGAAGCUU--A-CCGG-UACUAAUC---GUUCGAUUG-GCUUGAUCG----
------------------------------------------------------------------------
------------------------------------------------------------------------
------------------------------------------------------------------------
------------------------------------U------------
>LSUb_283166_BARHE/1-2785
---------------------A--UCAAGUG----UC-UUAAGGGCAUUU-GGUGGAUG-C-CUUGGC-AUG
CAC---AGGCGAUGAAGGACGUGA--UACGCUGCGAUAAGCUACGGGGAGGUGCGAAUACCCUU------UG
AUC-CGU-AGAUCUCCGAAUGGGGCAAC--------CC--------AC----CUUU-G-AUGGCUAGAAAAA
U-UU---UUCUAGUCGU---C----AGAUAA---AGG----U-A-UCUA-CAC-----CUGAA---------
-----------U-----------AAA----------------------------------------AUA-GG
GUG--------------------UA-A-GA-AGCAAACGCAGGGAACUGAAACAUCUAAGUACCUG--UAGG
AAAGGACAUCAA--ACGAGACUCCGUUA--GU-AG-UGGCGAGCGAAAGCGGA-C-CA-GGCCA--------
------------------------------------------------------------------------
----------GUG-------G----CU--------------------------UGA-----------AG--C
A-------AGAAA-AGUAG------AAU-CG-ACUG------------------------------------
-------GAA-AG-UCGAA-------CCAAAGAGGGUGAUAGUCCCGUA---------UACGUAA----CUC
U------U--CUUUA-----------------------AG-UC------CUAGAGUAGGGCGGGACACGUGA
AAUCCUGUCUGAA-UAUGGGUCG-ACC--ACGAUCCAAGCCUAAGUA--CUC-GUGCA-UGACCGAUAGC-G
AAC-C-AGUACCGUGAGGGAAAG-GUGAAA-AGUAC-CCCGACAAGGGG-A-GUGAAAAAGUACCUGAAACC
G-AAUGCCUACAA-ACA-GU-CGGA---GCCC-A---AG---------------------------------
------------------------------------------------------------------------
------------------------------------------------------------------------
------------------------------------------------------------------------
------------------------------------------------------------------------
```

------------------------------------------------------------------------
------------------------------------------------------------A-----------
------------------------------------------------------------------------
------------------------------------------------------------------------
--------------------------------------------------------------U--U-UG---
------------------------------------------------------------------------
------------------------------------------------------------------------
------------------------------------------------------------------------
------------------------------------------------------------------------
------------------------------------------------------------------------
------------------------------------------------------------------------
------------------------------------------------------------------------
------------------------------------------------------------------------
------------------------------------------------------------------------
------------------------------------------------------------------------
------------------------------------------------------------------------
------------------------------------------------------------------------
---------------------------U------------------UCUG--G---GUGACGGCGUACCUUU
UGUAUAAUGGGUCAGCGACUUAG-U-CUAACGAGCAA-GCUUAAGC--C-----------GGUA--------
--GG--UGUA--GGCGUAGCGAAAG-C---------------------------------------------
------------------------------------------------------------------------
------------------------------------------------------------------------
---------------------------------------------------------------------GAG
UCU------------------------------------GAACA-------------------------GG-
G-C-GU----------------------UCAGUUCGUUGGAUUA-GACCCGAAACC-GAGUGAUCUAGCCAU
GAGCAGGCUGAAGGUAAG-GUAACA-CUUACUGAAGGGCCGA--ACCCGUAU-CUGUUGCAAUAGAU-UGGG
AUGACUUGUGGCUAGGGGUGAAAGGCCAAUCAAACUCGG-AAAUAGCUGGUUCUCCGCGAAA-UCUAUUUAG
GUAGAGCGUUAG-UCG--------------------------------------------------------
----------------------AA-----UU-CUCCAGGGGGUAGAGC-ACUGGAU-GGG-CUA--GGGGUC
C---UCACC--GGAUUACCAAACCUAACCAAACUCCGAAUACCUGG--------------------------
------------------------------------------------------------------------
------------------AAGA-A-CU--------AACUAGC-AGACACA-C-GGCG-G-GUGCUAACGUCC
GU-C-GUGGAGAG-GGAAAC---AACCCUGACCACCA-UCUAAGGUCCCCAAGU--UA-UGG-CU-AA---G
-U-GGG-------AAAGGAUGUGAGGAUCCCAAAACAACCAGGAUGUUGGCUUAGAAGCAGCCA-CCAUUUA
AAGAAAGCGUAACAGCUCACUGGUCUAAAUAAGGGUCCCUGCGCCGAAAAUGUAAC-GGGGCUAA-AGCCAU
A----------CACCGAA-GCUGUGGAUUUACUC--C-----------------------------------
------------------------------------------------------------------------
---UUAA-----------------------------------------------------------------
---------------------------------GGAG-UAAG-UGGUAGCGGAGC-GUUCCGU---------
-----------AAG-CCUGU-GAAGGGA--GACUCGUGAGAG--C-UCCUGGAGGU-A-UCG-GAAGUGAGA
AUGC-UGACAU-GAGUA-ACGAUA---AAGG-GAGUGA--G----------------AGACUC-CCU-CGCCG
A-AAGUCC-AAGGGUUCC-UGCUUAAAG-UUAAUCUGAGCAGGGUGAGUCGGC-CC-CUAAGGCGAGGCCG-
A-A-AGGC-----GUA--------------------------------------------------------
----------------------------------------GUC-GA-U--GGGA-ACCACGUU-AAUA-UUC
GUGGA----------------------------------CCUGU----GG----------------------

```
------------------------------------------------------------------------
----------------------------------------------------------------GUAGUGAC
GGAUUG--------------CGUG-U-AUUGUAAGG-UC-U----UAUUG------------GAUUGAUCU-
UG----------------------------------------------------------------------
------------------------------------------------------------------------
------------------------------------------------------------------------
--------------------------C-AGU-G----AAG-CAGUUC-C-AGGAAAUAGCUC-CCAC-----
---------------------------------------------------------------AU-AGAGAC
CGUACCCCAAACCGAC------------------------------ACAGGUG--GACUGGUA-GAGAAUAC
UAAGGC-G-CU-UGAG-AGAACUAC-G-UUGAAGGAACUCGGC-AAAUUGCACGCGUAACUUCGG-AAGAAG
CGUGA-CCCU-U-UA--GCGGG--------------------------------------------------
------------------------------------------------------------------------
------------------------------------------------------------------------
------------------------------------------------------------------------
------------------------------------------------------------------------
------------------------------------------------------------------------
------------------------------------------------------------------------
------------------------------------------------------------------------
------------------------------------------------------------------------
------------------------------------------------------------------------
------------------------------------------------------------------------
------------------------------------------------------------------------
------------------------------------------------------------------------
------------------------------------------------------------------------
-------------------------------------------------------------CAA--CCGUUA
GAG--GGUGGCACAGACCAGGGGGUAGCGA-----CUGUUUACCAAAAACACAGGGCUCUGCGAAGUCG---
-CA-A-------------GACGACGU-AUAGGGUCUGA-CGCCUGCCCGGUGCUG-G-AAGG-UUAAGAGGA
GAUGU----------GCAA----------GCAUUGAAUUGAAGCCC--CAGUAAAC-GGCGGCCGUAACUAU
AACGGUCCUAAGGUAGCGAAAUUCCUUGUCGGG-UAAGUUCCGACCUGCACGAAUGGCGUAACGACU--UCC
CCGCUGUCUCCAAC----GUA-GACUCAGUGAAAUU--GAA---UUCCCCGUGAAGAUGCGGGGUUCCCGCG
GUUAGACGGAAAGA-CCCCGUGCACC-UUUACU--AUAACUU-UACACUGGCAUUUGUGUCUGUAUGUGUAG
GAU-AGGUGGUAGACUUCG----------AAGCAGG-GGC----------------------------GCCA
--------------------------------------GC-CCUUGUGGAG-U--C-AU-CCUUGAAAUA-C
CACC-CUUACCGAUAUGGAUGUCUAACCGCAG--UCCG-U-UA-U-C-------------------------
------------------------------------------------------------------------
------------------------------------------------------------------------
------------------------------------------------------------------------
------------------------------------------------------------------------
------------------------------------------------------CG----GA----------
------------------------------------------------UUCG-----GGACC-GUGUAUGGU
GGGUAGUUUGACUGGGGCGGU-CGCCUCCUA-AAGAGUAAC-GGAGGCGCGCG-AUGGUAGGCUCAGAACGG
UCGGAAAUCGUUUGUU-GAGUG-CAAUGGCAU-AAGCCUGCCUGACU-GUGA--------------------
--------------GACUGA---CAAGUCGAGCAGAGUCGAAA-GACGGUCAUAGUGAUCCGGU--------
---GG-U-----CCCGU-GUG--------------------GA-AGGGCCAUCGCUCAACGG-AUAAAAGGU
ACGCCGGGG-AUAACAGGCUGAUGACCCCCAAGAGUCCAUAUCGACGGGGUUGUUUGGCACCUCGAUGUCGA
```

```
CUCAUCGCAUCCUGGGGCUGGA-GCAGGUCCCAAGGGUAUGGCUGUUCGCCAUUUAAAGCGGUACGUGAGUU
GGGUUCAGAACGUCGUGAGACAGUUCGGUCCCUAUCUGCC--------------------G-UGG-------
-------------------------------GUGU--UGGAAUAUUGACAGG-ACC-UGUCCCUAGUACGAG
AGGAC-CGGGAUGGACGUAUCUC-UGGU-G-GACCUGUUGUGGCGCCAGCCGCAU-A-GCAGGGUAGCUAUA
UACGGACGAGAUAACCGCUGAAGGCA-UC--UAAGCGGGAAACUCACCUGAA-A-ACA--AGUAUUCC-C--
------------------------------------------------------------------------
------------------------------------------------------------------------
------------------------------------------------------------------------
------------------------------------------------------------------------
------------------------------------------------------------------------
------------------------------------------------------------------------
------------------------------------------------------------------------
------------------------------------------------------------------------
----------------------------------------------------------UG-AGA------AC
CGUGGUAGACUACCACGU-UGA-UAGGUC-AGG-UGUGGAAG-UGU-G----------------GU------
----AA-----------CACAUGAAGCUU--A-CUGA-UACUAAUC---GUUCAAUCG-ACUUGAUCG----
------------------------------------------------------------------------
------------------------------------------------------------------------
------------------------------------------------------------------------
------------------------------------U------------
>LSUb_224911_BRADU/1-2783
---------------------A-AUCAAGUG----CC-UUAAGGGUGUUC-GGUGGAUG-C-CUUGGC-GCU
GAG---AGGCGAUGAAGGACGUGC--UACGCUGCGAUAAGCCGUGGGGAGCUGCGAAGAAGCUU------UG
AUC-CGC-GGAUUUCCGAAUGGGGAAAC--------CC--------AC----CUUC-G-AUAGCCGGAA--C
U-U----UUCCGGUUAU---CAAGAG----A---AGG----U-A-UGAG-ACUU----CUGA----------
-----------A-----------UACA----------------------------------------UA-GG
AGG--------------------UU-U-CA-AGCAAACCCAGGGAACUGAAACAUCUAAGUACCUG--GAGG
AAAGGACAUCAA--CAGAGACUCCGUUA--GU-AG-UGGCGAGCGAACGCGGA-C-CA-GGCCA--------
------------------------------------------------------------------------
----------GUG-------A----UA--------------------------CAU-----------CA---
A-------AGACA-AUCGG------AACCAG-UCAG------------------------------------
-------GAA-AG-CUGGG-------CCUCAGAGGGUGAUAGCCCGUA----------CGAGUAA----UGC
G------A--UGAUG-----------------------UA-UC------CACGAGUAAGGCGGGACACGUGC
AAUCCUGUCUGAA-CGCGGGGGG-ACC--ACCCUCCAAGCCUAAGUA--CUC-CUCAG-CGACCGAUAGU-G
AAC-C-AGUACCGUGAGGGAAAG-GUGAAA-AGCAC-CCCGACGAGGGG-A-GUGAAAUAG-ACCUGAAACC
G-GACACCUACAA-ACA-GA-UGGA---GCCC-A---AG---------------------------------
------------------------------------------------------------------------
------------------------------------------------------------------------
------------------------------------------------------------------------
------------------------------------------------------------------------
------------------------------------------------------------------------
------------------------------------------------------------A-----------
------------------------------------------------------------------------
------------------------------------------------------------------------
-------------------------------------------------------------UU--C-GU---
```

```
------------------------------------------------------------------------
------------------------------------------------------------------------
------------------------------------------------------------------------
------------------------------------------------------------------------
------------------------------------------------------------------------
------------------------------------------------------------------------
------------------------------------------------------------------------
------------------------------------------------------------------------
------------------------------------------------------------------------
------------------------------------------------------------------------
------------------------------------------------------------------------
------------------------------------------------------------------------
---------------------------U-------------------CUG--G---GUGACAUCGUACCUUU
UGUAUUAUGGGCCAGCGACUUAA-U-UUAACGAGCAA-GCUUAAGC--C-----------GAUA--------
--GG--CGAA--GGCGUAGCGAAAG-C---------------------------------------------
------------------------------------------------------------------------
------------------------------------------------------------------------
---------------------------------------------------------------------GAG
UCU------------------------------------GAAUA-------------------------GG-
G-C-GU----------------------CAAGUUCGUUGUAUUA-GACCCGAAACC-UAGUGAUCUAGCCAU
GAGCAGGUUGAAGGUGAG-GUAACA-CUCACUGGAGGACCGA--ACGGGUGC-CUGUUGAAAAAGGC-UCCG
AUGACUUGUGGUUAGGGGUGAAAGGCCAAUCAAACUGGG-AAAUAGCUGGUUCUCCGCGAAA-GAUAUUUAG
GUAUCGCCUCGG-AUG--------------------------------------------------------
----------------------AA-----UA-CCUCAGGGGGUAGAGC-ACUGGAU-GGG-CUA--GGGGGA
C---UUACC--GUCUUACCAAACCCAACCAAACUCCGAAUACCUGA--------------------------
------------------------------------------------------------------------
------------------GAGU-A-CU--------AUCCGGG-AGUCACA-C-GGCG-G-GUGCUAACGUCC
GU-C-GUGGAGAG-GGAAAC---AACCCGGACCUACA-GCUAAGGCCCCUAAUU--CG-UGG-CU-AA---G
-U-GGG-------AAAGGAUGUGGAAAUCCCAAAACAACCAGGAGGUUGGCUUAGAAGCAGCCA-UCCUUUA
AAGAAAGCGUAACAGCUCACUGGUCUAAAUAAGGGUUUCUGCGCCGAAGAUGUAAC-GGGGCUCA-AGCCAC
G----------AGCCGAA-GCUUAGGGUGUGAUC--C-----------------------------------
------------------------------------------------------------------------
-----GC-----------------------------------------------------------------
-------------------------------AAGGGU-CACG-CGGUAGCGGAGC-GUUCUGU---------
-----------AAG-CCUGC-GAAGGGC--GACUCGUGAGAG--C-GCCUGGAGGU-A-UCA-GAAGUGCGA
AUGC-UGGCAU-GAGUA-ACGACA---AACA-CUGUGA--A---------------AGACAG-UGU-CGCCG
A-AAGUCC-AAGGGUUCC-UGCGUAAAG-UUAAUCUUCGCAGGGUUAGCCGGU-CC-CUAAGGCGAGGCCG-
A-A-AGGC-----GUA--------------------------------------------------------
----------------------------------------GUC-GA-U--GGGA-AUGCAGUG-AAUA-UUC
UGCAG----------------------------------CCAGU----GG----------------------
------------------------------------------------------------------------
----------------------------------------------------------------AUGGUGAC
GAAUCC--------------CGUG-U-GUUGUCCGA-CC-U----UAAUG------------GAUUGGUUG-
GG----------------------------------------------------------------------
------------------------------------------------------------------------
```

```
------------------------------------------------------------------------
--------------------------C-UUC-G----AAG-GGGUUC-C-AGGAAAUAGCCU-CCA------
---------------------------------------------------------------CA-UUAGAC
CGUACCCGAAACCGAC------------------------------ACAGGUG--GACUGGUA-GAGUAUAC
CAAGGC-G-CU-UGAG-AGAACUAU-G-UUGAAGGAACUCGGC-AAUUUACCUCCGUAACUUCGG-GAUAAG
GAGGC-CCAU-U-GC--UCGCG--------------------------------------------------
------------------------------------------------------------------------
------------------------------------------------------------------------
------------------------------------------------------------------------
------------------------------------------------------------------------
------------------------------------------------------------------------
------------------------------------------------------------------------
------------------------------------------------------------------------
------------------------------------------------------------------------
------------------------------------------------------------------------
------------------------------------------------------------------------
------------------------------------------------------------------------
------------------------------------------------------------------------
------------------------------------------------------------------------
-------------------------------------------------------------CAA--GCGGGC
AGU--GGGGGCACAGACCAGGGGGUGGCAA-----CUGUUUAACAAAAACACAGGGCUCUGCGAAAUCG---
-CA-A-------------GAUGACGU-AUAGGGUCUGA-CGCCUGCCCGGUGCCG-G-AAGG-UUAAGAGGA
GAGGU----------GCAA----------GCCUUGAAUCGAAGCCC--CGGUAAAC-GGCGGCCGUAACUAU
AACGGUCCUAAGGUAGCGAAAUUCCUUGUCGGG-UAAGUUCCGACCUGCACGAAUGGCGUAAUGACU--UCC
CCGCUGUCUCCAAC----AUA-GACUCAGUGAAAUU--GAA---UUCCCCGUGAAGAUGCGGGGUUCCUGCG
GUCAGACGGAAAGA-CCCCGUGCACC-UUUACU--GUAGCUU-UGCGCUGGUAUUCGUGACUGUUUGUGUAG
AAU-AGGUGGUAGGCUUUG----------AAGCCGU-GGC----------------------------GCCA
--------------------------------------GC-CAUGGUGGAG-C--C-GAAAUGUGAAAUA-C
CACC-CUAAUGGUUAUGGAUAUCUAACCGCGU--UCCC-U-CA-G-C-------------------------
------------------------------------------------------------------------
------------------------------------------------------------------------
------------------------------------------------------------------------
------------------------------------------------------------------------
------------------------------------------------------GG----GA----------
------------------------------------------------ACCG-----GGACA-GCGCAUGGU
GGGCAGUUUGACUGGGGCGGU-CGCCUCCCA-AAGAGUAAC-GGAGGCGUGCG-AAGGUAGGCUCAGAACGG
UCGGAAAUCGUUCGUC-GAGUA-UAAUGGCAU-AAGCCUGCCUGACU-GCGA--------------------
--------------GAUCUA---CGAAUCGAGCAGAGACGAAA-GUCGGUCAUAGUGAUCCGGU--------
---GG-U-----CCCGC-GUG--------------------GA-UGGGCCAUCGCUCAACGG-AUAAAAGGU
ACGCCGGGG-AUAACAGGCUGAUGACGCCCAAGAGUCCAUAUCGACGGCGUCGUUUGGCACCUCGAUGUCGG
CUCAUCACAUCCUGGGGCUGGA-GAAGGUCCCAAGGGUUCGGCUGUUCGCCGAUUAAAGUGGUACGUGAGCU
GGGUUCAGAACGUCGUGAGACAGUUCGGUCCCUAUCUGCC--------------------G-UGG-------
-------------------------------GUGU--UGGAAUGUUGAGAGG-AUU-UGCCCCUAGUACGAG
AGGAC-CGGGGUGAACGUACCUC-UGGU-G-GAGCUGUUGUCGCGCCAGCGGCAG-U-GCAGCAUAGCUAUG
UACGGACGGGAUAACCGCUGAAAGCA-UC--UAAGCGGGAAACCCACCUCAA-A-ACG--AGCAUUCC-CU-
```

```
------------------------------------------------------------------------
------------------------------------------------------------------------
------------------------------------------------------------------------
------------------------------------------------------------------------
------------------------------------------------------------------------
------------------------------------------------------------------------
------------------------------------------------------------------------
------------------------------------------------------------------------
----------------------------------------------------------UG-AGA------AC
CGUGGAAGACCACCACGU-UGA-UAGGCC-GGA-UGUGGAAG-UGC-A----------------GU------
----AA-----------UGCAUGCAGCUU--A-CCGG-UACUAAUC---GUUCGAUUG-GCUUGAUUG----
------------------------------------------------------------------------
------------------------------------------------------------------------
------------------------------------------------------------------------
------------------------------------C------------
>LSUb_290633_GLUOX/1-2725
---------------------G----------------AGAAGGGCGUUC-GGUGGAUG-C-CUUGGC-ACU
AAG---AGGCGAUGAAGGACGUGG--UACGCUGCGAAAAGCCAUGGGGAGCCGCGAACAGGCUU------UG
AUC-CGU-GGAUGUCCGAAUGGGGCAAC--------CC--------CC----UC----------------GC
A-A-------------------------------GAG----G-A-UCAU-GCA-----CUGA----------
-----------A-----------UACA----------------------------------------UA-GG
U-G--------------------UA-U-GA-GGCAAACCCGGGGAACUGAAACAUCUCAGUACCUG--GAGG
AAAAGACAUCAA--CAGAGAUUCCGCUA--GU-AG-UGGCGAGCGAACGCGGA-G-CA-GGCCA--------
------------------------------------------------------------------------
----------AUG-------C----CU--------------------------GAU-----------CA---
G-------GAAGA-AGCUG------AAC-GG-UCUG------------------------------------
-------GAA-AG-UCCGG-------CAAGAAUGGGUGAUAGCCCGUA----------AGCGUAG----UGU
U------C--UGAUG-----------------------AG-GA------UUCGAGUAGGGCGGGGCACGUGA
AACCCUGUCUGAA-CAUGGGGGG-ACC--ACCCUCCAAGCCUAAAUA--CUC-CUUAG-UGACCGAUAGC-G
AAC-A-AGUACCGUGAGGGAAAG-GUGAAA-AGCAC-CCCGAUGAGGGG-A-GUGAAAGAG-ACCUGAAACC
G-GACGCCUACAA-GCA-GU-CGGA---GCCU-C--------------------------------------
------------------------------------------------------------------------
------------------------------------------------------------------------
------------------------------------------------------------------------
------------------------------------------------------------------------
------------------------------------------------------------------------
------------------------------------------------------------------------
------------------------------------------------------------------------
------------------------------------------------------------------------
--------------------------------------------------------------U--U-AU---
------------------------------------------------------------------------
------------------------------------------------------------------------
------------------------------------------------------------------------
------------------------------------------------------------------------
------------------------------------------------------------------------
```

```
------------------------------------------------------------------------
------------------------------------------------------------------------
------------------------------------------------------------------------
------------------------------------------------------------------------
------------------------------------------------------------------------
------------------------------------------------------------------------
------------------------------------------------------------------------
------------------------------------------------GG--G---GUGACGGCGUACCUUU
UGUAUAAUGGGUCAGCGAGUUUC-U-GUUUGCAGCGA-GCUUAAGC--C-----------GUUA--------
--GG--UGUA--GGCGUAGCGAAAG-C---------------------------------------------
------------------------------------------------------------------------
------------------------------------------------------------------------
---------------------------------------------------------------------GAG
UCU------------------------------------GAAUA-------------------------GG-
G-C-GA---------------------AUGAGUUGCUGGCAGAA-GACCCGAAACC-GAGUGAUCUAGCCAU
GGCCAGGCUGAAGGUGCG-GUAACA-CGCACUGGAGGGCCGA--ACCCACGC-CUGUUGAAAAAGUC-GGGG
AUGAGCUGUGGCUAGGGGUGAAAGGCCAAUCAAACUCGG-AGAUAGCUGGUUCUCCGCGAAA-UCUAUUGAG
GUAGAUCGUCGG-GUG--------------------------------------------------------
----------------------UU-----UA-CCCUGGGGGGUAGAGC-ACUGGAU-GGG-CUA--GGGGGG
C---CCAAA--GCCUUACCAAACCUAACCAAACUCCGAAUACCCAG--------------------------
------------------------------------------------------------------------
------------------GAGU-A-UA---------GCCCGGC-AGACAGA-C-AGUG-G-GUGCUAAGGUCC
AU-U-GUCGAGAG-GGAAAC---AGCCCAGACCACCA-GCUAAGGCCCCUAAAU--CG-UGG-CU-AA---G
-U-GGG-------AAAGGAUGUGGGGAUUCCAAAACAACCAGGAGGUUGGCUUAGAAGCAGCCA-UCCUUUA
AAGAAAGCGUAAUAGCUCACUGGUCU--AUUAGAAACCCUGCGCCGAAAAUGUAAC-GGGGCUCA-AGCCAC
G----------UGCCGAA-GCUGUGGGUGCAUAC--U-----------------------------------
------------------------------------------------------------------------
------------------------------------------------------------------------
--------------------------------AUGUA-UGCG-CGGUAGCGGAGC-GUUCCGU---------
-----------AAG-UCUGC-GAAGGAG--ACGGGGUGACCC--U-CUCUGGAGAU-A-UCG-GAAGUGCGA
AUGC-UGACAU-GAGUA-GCGACA---AACA-GUGCGA--G---------------AAACAC-UGU-CGCCG
A-AAGUCC-AAGGGUUCC-UGCGCAAGG-UUAAUCCACGCAGGGUGAGCCGGC-CC-CUAAGGCGAGGGCG-
A-G-AGCC-----GUA--------------------------------------------------------
----------------------------------------GUC-GA-U--GGGA-ACCAGUUC-AAUA-UUA
CUGGG----------------------------------CCUGC----CA----------------------
------------------------------------------------------------------------
----------------------------------------------------------------GAAGUGAC
GAAUGA--------------GAGA-U-GUUGUCUGU-CC-U----UAACG------------GAUUGAACA-
GG----------------------------------------------------------------------
------------------------------------------------------------------------
------------------------------------------------------------------------
--------------------------C-UUU-U----CAA-UCAUUC-C-AGGAAAUAGCUC-UGGC-----
---------------------------------------------------------------GU-AUAGAC
CGUACCCGAAACCGAC------------------------------ACAGGUG--GACUGGUA-GAGAAUAC
CAAGGC-G-CU-UGAG-AGAACGAU-G-CUGAAGGAACUAGGC-AAAUUGCUCGUGUAACUUCGG-GAUAAA
```

CGAGA-CCCG-C-UC--GUGGG--------------------------------------------------
------------------------------------------------------------------------
------------------------------------------------------------------------
------------------------------------------------------------------------
------------------------------------------------------------------------
------------------------------------------------------------------------
------------------------------------------------------------------------
------------------------------------------------------------------------
------------------------------------------------------------------------
------------------------------------------------------------------------
------------------------------------------------------------------------
------------------------------------------------------------------------
------------------------------------------------------------------------
------------------------------------------------------------------------
-------------------------------------------------------------CAA--CCAUGG
ACG--GGUGGCACAGACCAGGGGGUAGCGA-----CUGUUUAGUAAAAACACAGGGCUCUGCGAAAUCG---
-UG-A-------------GAUGACGU-AUAGGGCCUGA-CGCCUGCCCGGUGCCG-G-AAGG-UUAAGAGGA
GGUGU----------GCAA----------GCACUGAAUUGAAGCCC--CGGUAAAC-GGCGGCCGUAACUAU
AACGGUCCUAAGGUAGCGAAAUUCCUUGUCGGG-UAAGUUCCGACCUGCACGAAUGGCGUAACGACU--UCC
CCACUGUCUCCAGC----AUC-GACUCAGCGAAAUU--GAA---UUCCCCGUGAAGAUGCGGGGUACCCGCG
GUCAGACGGAAAGA-CCCUAUGAACC-UUUACU--GCAGCUU-UGCAGUGGCAUCAGAGACAUUCUGUGUAG
GAU-AGGUCGGAGGCUUUG----------AAACCCA-GGC----------------------------GCCA
--------------------------------------GC-UUGGGUGGAG-C--C-AU-CCUUGAAAUA-C
GACC-CUGAAUUUUUCUGAUGUCUAACCGAGA--CCAG-U-AA-G-C-------------------------
------------------------------------------------------------------------
------------------------------------------------------------------------
------------------------------------------------------------------------
------------------------------------------------------------------------
------------------------------------------------------CU----GG----------
------------------------------------------------UCCG-----GGACC-CUGCAUGGU
GGGCAGUUUGACUGGGGCGGU-CGCCUCCCA-AAGUGUAAC-GGAGGCGCGCG-AUGGUGGGCUCAGGCCGG
UCGGAAACCGGCUGUC-GAGUG-CAAUGGCAU-AAGCCCGCCUGACU-GUGA--------------------
--------------GAGUGA---CAGCUCGAUCAGAGACGAAA-GUCGGCCAUAGUGAUCCGGU--------
---GG-U-----CCCAU-GUG--------------------GA-CGGGCCAUCGCUCAACGG-AUAAAAGGU
ACUCUAGGG-AUAACAGGCUGAUCUCCCCCAAGAGUCCACAUCGACGGGGAGGUUUGGCACCUCGAUGUCGG
CUCAUCACAUCCUGGGGCUGGA-GCAGGUCCCAAGGGUUCGGCUGUUCGCCGAUUAAAGUGGUACGUGAGCU
GGGUUUAGAACGUCGUGAGACAGUUCGGUCCCUAUCUGCC--------------------G-UGG-------
-------------------------------GUGU--AAGAGACUUGAGAGG-AUU-UGUCCCUAGUACGAG
AGGAC-CGGGAUGAACAUACCUC-UGGU-G-CACCGGUUGUCACGCCAGUGGCAC-A-GCCGGGUAGCUAAG
UAUGGACGGGAUAACCGCUGAAAGCA-UC--UAAGCGGGAAACCCACCUCAA-A-ACG--AGGUCUCA-U--
------------------------------------------------------------------------
------------------------------------------------------------------------
------------------------------------------------------------------------
------------------------------------------------------------------------
------------------------------------------------------------------------

------------------------------------------------------------------------
------------------------------------------------------------------------
------------------------------------------------------------------------
-------------------------------------------------------------AGA------GC
CGUGACAGACCAUCACGU-UGA-UAGGCC-GGG-UGUGAAAG-UGC-A----------------GU------
----AA-----------UGCAUGCAGCUA--A-CCGG-UCCUAAUC---GCUCAUA-A-UCCUCACU-----
------------------------------------------------------------------------
------------------------------------------------------------------------
------------------------------------------------------------------------
-------------------------------------------------
>LSUb_272627_MAGMG/1-2739
------------------------UCAAGCG----UC-AAAAGGGCAUUU-CGUGGAUG-C-CUUGGC-ACU
GAG---AGGCGAUGAAGGACGUGG--CACUCUGCGAAAAGCCACGGGGAGCCGAGAGCAGGCUU------UG
AUC-CGU-GGAUAUCCGAAUGGGGAAAC--------CC--------CA----CC----------------GU
A-A-------------------------------GGU----G-A-UCCC-ACA-----CUGA----------
-----------A-----------UACA----------------------------------------UA-GG
U-G--------------------UG-G-GA-GGCGAACCGGGCGAACUGAAACAUCUAAGUAGCCC--GAGG
AAAGGACAUCAA--CCGAGACUCCGCUA--GU-AG-UGGCGAGCGAACGCGGA-C-CA-GCCCA--------
------------------------------------------------------------------------
----------GUG-------A----UG--------------------------UUG-----------UG---
U-------ACAUA-AUCUG------AAC-CG-UCUG------------------------------------
-------GAA-AG-UCGGG-------CCAUAGAGCGUGAUAGCCGCUUA---------AGAGUAA----ACC
G------C--ACAG------------------------CA-UC------CUCGAGUAAGGCGGGACACGUGA
AAUCCUGUCUGAA-CAUGGGGGG-ACC--ACCCUCCAAGGCUAAGUA--CUC-CUCAG-UGACCGAUAGC-G
AAC-U-AGUACCGUGAGGGAAAG-GUGAAA-AGCAC-CCCGACAAGGGG-A-GUGAAAUAGUACCUGAAACG
G-AAUGCCUACAA-GCA-GU-CGGA---GCCC-C--------------------------------------
------------------------------------------------------------------------
------------------------------------------------------------------------
------------------------------------------------------------------------
------------------------------------------------------------------------
------------------------------------------------------------------------
------------------------------------------------------------------------
------------------------------------------------------------------------
------------------------------------------------------------------------
--------------------------------------------------------------C--U-UG---
------------------------------------------------------------------------
------------------------------------------------------------------------
------------------------------------------------------------------------
------------------------------------------------------------------------
------------------------------------------------------------------------
------------------------------------------------------------------------
------------------------------------------------------------------------
------------------------------------------------------------------------
------------------------------------------------------------------------
------------------------------------------------------------------------

```
------------------------------------------------------------------------
------------------------------------------------------------------------
------------------------------------------------GG--G---GUGACGGCGUACCUUU
UGUAUAAUGGGUCAGCGAGUUAG-U-GUAUGCAGCAA-GCUUAAGC--C-----------GGUA--------
--GG--CGUA--GGCGCAGCGAAAG-C---------------------------------------------
------------------------------------------------------------------------
------------------------------------------------------------------------
---------------------------------------------------------------------GAG
UCU------------------------------------GAAUA-------------------------GG-
G-C-GA----------------------CGAGUUGCAUGCAUUA-GACCCGAAACC-GGGUGAUCUAGCCAU
GGGCAGGUUGAAGGUGGG-GUAACA-CCCACUGGAGGACCGA--ACUCACGU-CUGUUGAAAAAGAC-GGAG
AUGACCUGUGGUUAGGGGUGAAAGGCCAAUCAAACUCGG-AAAUAGCUGGUUCUCCGCGAAA-GCUAUUUAG
GUAGCGCCUCGG-ACG--------------------------------------------------------
----------------------AA-----UA-CCUACGGGGGUAGAGC-ACUGGAU-GGG-CUA--GGGGGU
C---CCAAA--GACCUACCAAACCUAACCAAACUCCGAAUACCGUA--------------------------
------------------------------------------------------------------------
------------------GAGU-A-CU--------AUCCGGG-AGACAGA-C-GGUG-G-GUGCUAAGGUCC
AU-C-GUCAAAAG-GGAAAC---AGCCCAGACCGCCA-GCUAAGGUCCCCAAGU--CA-UGG-UU-AA---G
-U-GGG-------AAAGGAUGUAAGACGGCCAAAACAACCAGGAGGUUGGCUUAGAAGCAGCCA-UCCUUUA
AAGAAAGCGUAAUAGCUCACUGGUCUAA--UAGCUGUCUCGCGCCGAAAAUGUACC-GGGGCUCA-AACCAU
G----------CACCGAA-GCUGCGGGUGUGCAG--C-----------------------------------
------------------------------------------------------------------------
------------------------------------------------------------------------
---------------------------------AAUG-CACG-CGGUAGCGGAGC-GUUCCGU---------
-----------AAG-CCUGC-GAAGGCG--UCAUCGUGAGAG--G-CGCUGGAGGU-A-UCG-GAAGUGAGA
AUGC-UGACAU-GAGUA-GCGACA---AACA-GUGUGA--G---------------AAACAC-UGU-CGCCG
A-AAGUCC-AAGGGUUCC-UGCGCAAGG-CUAAUCCGCGCAGGGUAAGCCGGC-CC-CUAAGGCGAGGGCG-
A-A-AGCC-----GUA--------------------------------------------------------
----------------------------------------GUC-GA-U--GGGA-ACCACGUU-AAUA-UUC
GUGGG----------------------------------CCUGC----UG----------------------
------------------------------------------------------------------------
----------------------------------------------------------------GAAGUGAC
GACCUC--------------CGUG-U-AUCGUAUCC-UC-U----UAUCG------------GAUUGAGGG-
UG----------------------------------------------------------------------
------------------------------------------------------------------------
------------------------------------------------------------------------
--------------------------C-GGU-G----AA--GGGGGU-C-CAGGAA-AUAAC-UCCAGC---
---------------------------------------------------------------GU-AUAGAC
CGUACCCUAAACCGAC------------------------------ACAGGUG--GACUGGUA-GAGUAUAC
CAAGGC-G-CU-UGAG-AGAACGAU-G-UUGAAGGAACUAGGC-AAAUUGACCCCGUAACUUCGG-GAUAAG
GGGUA-CCUC-U-CU--UUGCG--------------------------------------------------
------------------------------------------------------------------------
------------------------------------------------------------------------
------------------------------------------------------------------------
------------------------------------------------------------------------
```

```
------------------------------------------------------------------------
------------------------------------------------------------------------
------------------------------------------------------------------------
------------------------------------------------------------------------
------------------------------------------------------------------------
------------------------------------------------------------------------
------------------------------------------------------------------------
------------------------------------------------------------------------
------------------------------------------------------------------------
-------------------------------------------------------------CAA--GCAGAG
GGG--GGUGGCACAGACCAGGGGGUGGCGA-----CUGUUUACUAAAAACACAGGGCUCUGCGAAGUCG---
-AA-A-------------GACGACGU-AUAGGGUCUGA-CGCCUGCCCGGUGCCG-G-AAGG-UUAAAAGGA
GGGGU----------GCAA----------GCUCUGAAUUGAAGCCC--CGGUAAAC-GGCGGCCGUAACUAU
AACGGUCCUAAGGUAGCGAAAUUCCUUGUCGGG-UAAGUUCCGACCUGCACGAAUGGCGUAACGACU--UCC
CCGCUGUCUCCAAC----AUC-GACUCAGCGAAAUU--GAA---UUCUCCGUGAAGAUGCGGAGUACCCGUG
GCUAGACGGAAAGA-CCCCGUGCACC-UUUACU--ACAGCUU-UGCAGUGGUACUAGGAUGUGGAUGUGUAG
GAU-AGGUGGGAGCCUAUG----------AAGCACC-GGC----------------------------GCCA
--------------------------------------GC-UGGUGUGGAG-G--C-AA-CCUUGAAAUA-C
CACC-CUUCUGCUUCCUGGUAUCUAACCGCGC--CCCG-UGAA-C-C-------------------------
------------------------------------------------------------------------
------------------------------------------------------------------------
------------------------------------------------------------------------
------------------------------------------------------------------------
------------------------------------------------------CG----GG----------
------------------------------------------------GCCG-----GGACC-CUGCAUGGC
GGGUAGUUUGACUGGGGCGGU-CGCCUCCCA-AAGAGUAAC-GGAGGCGCGCG-AUGGUGGGCUCAGGCUGG
UCGGAAAUCAGCUGUC-GAGUG-CAAUGGCAU-AAGCCCGCCUGACU-GCGA--------------------
--------------GACAAA---CAAGUCGAGCAGAGACGAAA-GUCGGUCAUAGUGAUCCGGU--------
---GG-U-----CCCGC-GUG--------------------GA-AGGGCCAUCGCUCAACGG-AUAAAAGGU
ACGCCGGGG-AUAACAGGCUGAUCUCCCCCAAGAGUCCACAUCGACGGGGAGGUUUGGCACCUCGAUGUCGG
CUCAUCACAUCCUGGGGCUGGA-GCAGGUCCCAAGGGUUCGGCUGUUCGCCGAUUAAAGUGGUACGUGAGCU
GGGUUUAGAACGUCGUGAGACAGUUCGGUCCCUAUCUACC--------------------A-UGG-------
-------------------------------GUGU--UGGAAACUUGAGAGG-ACC-UGUCCCUAGUACGAG
AGGAC-CGGGAUGGACAUACCUC-UAGU-G-UACCGGUUGUCGCGCCAGCGGCAU-C-GCCGGGUAGCCAUG
UAUGGACGAGAUAACCGCUGAAUGCA-UC--UAAGCGGGAAGCUCCCCUCAA-A-ACC--AGGUUUCC-CU-
--A---------------------------------------------------------------------
------------------------------------------------------------------------
------------------------------------------------------------------------
------------------------------------------------------------------------
------------------------------------------------------------------------
------------------------------------------------------------------------
------------------------------------------------------------------------
------------------------------------------------------------------------
----------------------------------------------------------UC-AGA------GC
CGUGGAAGACCACCACGU-UGA-UAGGCA-GCA-UGUGGAAG-CGU-G----------------GU------
```

```
----AA-----------CACGUGAAGCUA--A-GCUG-UACUAAUC---GCUCGAUCG-GCUUGAUCG----
------------------------------------------------------------------------
------------------------------------------------------------------------
------------------------------------------------------------------------
------------------------------------C------------
>LSUb_565050_CAUCN/1-2763
---------------------G-AUCAAGCG----CA--UAAGGGCUUCU-GACGGAUG-C-CUUGGC-AUU
GAG---AGGCGAUGAAGGACGUGG--CACGCUGCGAUAAGAGCCGGGGAGGCGCGAGCACCCUU------UG
AUC-CGG-CUAUCUCCGAAUGGGGAAAC--------CC--------AC----CUUU-A-UGGUCACCCGA-C
U-AU---CGGUUGAUCA---G----A---CA---AGG----U-A-UAAU-GAC-----CUGA----------
-----------A-----------UACA----------------------------------------UA-GG
GUU--------------------CA-U-UA-AGCAAACCCGGGGAACUGAAACAUCUCAGUACCCG--GAGG
AAAGGACAUCAA--CCGAGACUCCCGUA--GU-AG-UGGCGAGCGAACCGGGA-C-CA-GGCCA--------
------------------------------------------------------------------------
----------GUG-------C----UG--------------------------UCG-----------UG---
A-------CAUAA-AGCUG------AAU-GU-GUUG------------------------------------
-------GGA-AA-CACAG-------CCAUAGUGGGUGAUAGCCCGUA----------AGCGUCA----AAU
A------G--CGA-------------------------CA-GA------CUCGAGUAGGGCGGGACACGUGA
AAUCCUGUCUGAA-CAUGGGGGG-ACC--ACCCUCCAAGCCUAAGUA--CUC-CUCAA-UGACCGAUAGC-G
AAC-A-AGUACCGUGAGGGAAAG-GUGAAA-AGCAC-CCCGACAAGGGG-A-GUGAAACAGAACCUGAAAUC
G-GAAGCCUACAA-GCA-GU-CGGA---GCCA-C--------------------------------------
------------------------------------------------------------------------
------------------------------------------------------------------------
------------------------------------------------------------------------
------------------------------------------------------------------------
------------------------------------------------------------------------
------------------------------------------------------------------------
------------------------------------------------------------------------
------------------------------------------------------------------------
-------------------------------------------------------------CG--C-GC---
------------------------------------------------------------------------
------------------------------------------------------------------------
------------------------------------------------------------------------
------------------------------------------------------------------------
------------------------------------------------------------------------
------------------------------------------------------------------------
------------------------------------------------------------------------
------------------------------------------------------------------------
------------------------------------------------------------------------
------------------------------------------------------------------------
------------------------------------------------------------------------
------------------------------------------------------------------------
------------------------------------------------GU--G---GUGACGGCGUACCUUU
UGUAUAAUGGGUCAGCGACUUCA-U-GUGCCGUGCAA-GCUUAAGC--C-----------GUUA--------
--GG--UGUA--GGCGCAGCGAAAG-C---------------------------------------------
```

```
------------------------------------------------------------------------
------------------------------------------------------------------------
---------------------------------------------------------------------GAG
UCU------------------------------------GAAUA-------------------------GG-
G-C-GA---------------------AUAAGUACGUCGCAUGACGACCCGAAACC-AGGUGAUCUAUCCAU
GAGCAGGUUGAAGGUAAG-GUAACA-CUUACUGGAGGACCGA--ACCGGUGA-AUGUUGAAAAAUUC-UCGG
AUGACUUGUGGAUAGGGGUGAAAGGCCAAUCAAACCUGG-ACAUAGCUGGUUCUCCGCGAAA-ACUAUUUAG
GUAGUGCCUCGG-ACG--------------------------------------------------------
----------------------AA-----UU-CCUUGGGGGGUAGAGC-ACUGAAU-GGA-UGC--GGGCGG
C---GC-GA--GCUGUACCAAUUCUAAUCAAACUCCGAAUACCCAA--------------------------
------------------------------------------------------------------------
------------------GAGA-A-CU--------AUCCGGG-AGACACA-C-GGCG-G-GUGCUAACGUCC
GU-C-GUGAAAAG-GGAAAC---AACCCUAACCAUCA-UCUAAGGCCCCCAAGU--AC-UGG-CU-AA---G
-U-GGG-------AAACGAUGUGGGUUUGCUUUGACAACCAGGAUGUUGGCUUAGAAGCAGCCA-UCAUUUA
AAGAAAGCGUAACAGCUCACUGGUC-----AAGCGAACCUGCGCGGAAAAUGUAAC-GGGGCUAA-AGCCAG
U----------CGCCGAA-GGUAUGGGUGUGCAC--U-------------------------------------
------------------------------------------------------------------------
------------------------------------------------------------------------
--------------------------------CUGUG-CACG-CGGUAGCGGAGC-GUUCCGU---------
-----------AAG-CCGAU-GAAGGUG--A-GGCGUGAGCC--U-UGCUGGAGGU-A-UCG-GAAGUGAGA
AUGC-UGACAU-GAGUA-GCGAUA---AAGA-GGGUGA--G---------------AGACCC-UCU-CGCCG
A-AAGCCC-AAGGGUUCC-UGCGUAAAG-CUAAUCUGCGCAGGGUUAGUCGGC-CC-CUAAGGCGAGGCCG-
A-A-AGGC-----GUA--------------------------------------------------------
----------------------------------------GUC-GA-U--GGGA-AUCAGGUG-AAUA-UUC
CUGAA----------------------------------CCAGU----UG----------------------
------------------------------------------------------------------------
----------------------------------------------------------------GAAGUGAC
GGAUCU--------------GGUA-A-AUUGUCAGG-GC-U----UAUUG------------GAUUGCUCC-
UG----------------------------------------------------------------------
------------------------------------------------------------------------
------------------------------------------------------------------------
-------------------------GC-AGU-G----AAC-AGGUCC-C-UGGAAAUAACUC-CA-------
---------------------------------------------------------------AC-GGAGAC
CGUACCCGAAACCGAC------------------------------ACAGGUG--GGCAGGUA-GAGUAUAC
CAAGGC-G-CU-UGAG-AGAACUAU-G-CUGAAGGAACUCGGC-AAAUUGCACGCGUAACUUCGG-GAUAAG
CGUGA-CUCU-C-UU--UUGGG--------------------------------------------------
------------------------------------------------------------------------
------------------------------------------------------------------------
------------------------------------------------------------------------
------------------------------------------------------------------------
------------------------------------------------------------------------
------------------------------------------------------------------------
------------------------------------------------------------------------
------------------------------------------------------------------------
------------------------------------------------------------------------
```

```
------------------------------------------------------------------------
------------------------------------------------------------------------
------------------------------------------------------------------------
------------------------------------------------------------------------
-------------------------------------------------------------CAA--CCAGAU
GAG--AGUGGCACAAGCCAGGGGGUAGCGA-----CUGUUUAUCAAAAACACAGGGCUCUGCGAAGCCG---
-CA-A-------------GGCGACGU-AUAGGGUCUGA-CGCCUGCCCGGUGCCU-G-AAGG-UUAAAAGGA
GGGGU----------GCAA----------GCUCUGAAUUGAAGCCC--AGGUAAAC-GGCGGCCGUAACUAU
AACGGUCCUAAGGUAGCGAAAUUCCUUGUCGGG-UAAGUUCCGACCUGCACGAAUGGCGUAACGACU--UCC
CCACUGUCUCCAGC----AUA-GGCUCAGCGAAAUU--GAA---UUCCCCGUGAAGAUGCGGGGUUCCCGCG
GUCAGACGGAAAGA-CCCUAUGAACC-UUUACU--GCAGCUU-CGCCUUGGCGUUAGCAGCAACAUGUGUAG
GAU-AGGUGGGAGGCUAUG----------AAACUCG-GGC----------------------------GCCA
--------------------------------------GU-UCGAGUGGAG-C--C-AU-CCUUGAAAUA-C
CACC-CUUAUUGUUGCUGACGUCUAACCGCGG--CCCG-U-UA-U-C-------------------------
------------------------------------------------------------------------
------------------------------------------------------------------------
------------------------------------------------------------------------
------------------------------------------------------------------------
------------------------------------------------------CG----GG----------
------------------------------------------------UCCG-----GGACA-UGGCGUGGC
GGGCAGUUUGACUGGGGCGGU-CGCCUCCCA-AAGUGUAAC-GGAGGCGCGCG-AUGGUGGGCUCAGACCGG
UCGGAAAUCGGUCGUC-GAGUG-CAAUGGCAU-AAGCCCGCCUGACU-GCGA--------------------
--------------GACUGA---CAAGUCGAGCAGAGACGAAA-GUCGGCCAUAGUGAUCCGGU--------
---GG-U-----CCUGC-GUG--------------------GA-AGGGCCAUCGCUCAACGA-AUAAAAGGU
ACUCUAGGG-AUAACAGGCUGAUUUUGCCCAAGAGUCCAUAUCGACGGCAAAGUUUGGCACCUCGAUGUCGG
CUCAUCACAUCCUGGGGCUGGA-GCAGGUCCCAAGGGUUCGGCUGUUCGCCGAUUAAAGUGGUACGUGAGCU
GGGUUCAGAACGUCGUGAGACAGUUUGGUCCCUAUCUGCC--------------------G-UGG-------
-------------------------------GUGU--ACGAGACUUGAGAGG-AUC-UGUCCCUAGUACGAG
AGGAC-CGGGAUGGACACACCUC-UGGU-G-GACCUGUCAUGGCGCCAGCUGUGC-A-GCAGGGUAGCUAAG
UGUGGAAUAGAUAACCGCUGAAAGCA-UC--UAAGCGGGAAACUAACCUCAA-A-ACA--AGGUCUCG-C--
------------------------------------------------------------------------
------------------------------------------------------------------------
------------------------------------------------------------------------
------------------------------------------------------------------------
------------------------------------------------------------------------
------------------------------------------------------------------------
------------------------------------------------------------------------
------------------------------------------------------------------------
----------------------------------------------------------UG-AGA------GC
CGUGGAAGACCACCACGU-UGA-UAGGCC-GGG-UGUGGAAG-UGC-G----------------GU------
----GA-----------CGCAUGGAGCUU--A-CCGG-UACUAAUA---GCUCGAUAG-GCUUGAUCG----
------------------------------------------------------------------------
------------------------------------------------------------------------
------------------------------------------------------------------------
------------------------------------U------------
```

```
>LSUb_349102_RHOS5/1-2765
------------------------UCAAGCG----CG-AAAAGGGCGUUU-GGUGGAUG-C-CUAGGC-AGC
AAG---AGGCGAUGAAGGACGUGA--UACCCUGCGUUAAGCCAUGGGGAGCCGGGAAUAGGCUU------UG
AUC-CAU-GGAUGUCCGAAUGGGGAAAC--------CC--------AC----CUGA-U-UGUUCGUU---GU
U-C------AAUGGGCA---AA---G---AC---AGG----U-A-UCAU-UAC-----CCUGA---------
-----------A-----------UACA----------------------------------------UAG-G
G-G--------------------UU-U-UGAAGCGAACCCGGGGAACUGAAACAUCUAAGUACCCG--GAGG
AAAGGAAAUCAA--CAGAGACUCCGUUA--GU-AG-UGGCGAGCGAACGCGGA-C-CA-GCCGA--------
------------------------------------------------------------------------
----------UCC-------U----CG-----------------------------------------A---
A-------GAGUG-ACUGG------AAU-GC-GCUG------------------------------------
-------GAA-AG-CGCAG-------CGAUACAGGGUGACAGCCCGUA----------CAGGAAG----CUC
C------A--CAGG------------------------AC-AU------AUCAAGUAGGGCGGGACACGUGA
AAUCCUGUCUGAA-GAUCGGGGG-ACC--ACCCCCGAAGGCUAAGUA--CUC-CUUGC-UGACCGAUAGC-G
AAC-C-AGUACCGUGAGGGAAAG-GUGAAA-AGCAC-CCCGACGAGGGG-A-GUGAAACAGUACCUGAAACC
G-GACGCCUACAA-GCA-GU-CGGA---GGCC-C---AU---------------------------------
------------------------------------------------------------------------
------------------------------------------------------------------------
------------------------------------------------------------------------
------------------------------------------------------------------------
------------------------------------------------------------------------
------------------------------------------------------------------------
------------------------------------------------------------------------
------------------------------------------------------------------------
-------------------------------------------------------------GU--U-CU---
------------------------------------------------------------------------
------------------------------------------------------------------------
------------------------------------------------------------------------
------------------------------------------------------------------------
------------------------------------------------------------------------
------------------------------------------------------------------------
------------------------------------------------------------------------
------------------------------------------------------------------------
------------------------------------------------------------------------
------------------------------------------------------------------------
------------------------------------------------------------------------
------------------------------------------------------------------------
----------------------------------------------UCCC--G---CUGACGGCGUACCUUU
UGUAUAAUGGGUCAACGACUUGG-U-CUCACGAGCAA-GCUUAAGC--C-----------GGUA--------
--GG--UGGA--GGCGCAGCGAAAG-C---------------------------------------------
------------------------------------------------------------------------
------------------------------------------------------------------------
---------------------------------------------------------------------GAG
UCU------------------------------------UAAAA-------------------------GG-
G-C-GU----------------------CGAGUUCGUGGGAUCA-GACCCGAAACC-AGGUGAUCUAGCCAU
```

```
GAGCAGGAUGAAGUCAGG-GUAACA-CCUGAUGGAGGUCCGA--ACCAACAC-CCGUUGAAAAGGGU-CUGG
AUGACUUGUGGCUAGGGGUGAAAGGCCAAUCAAACCUGG-AGAUAGCUGGUUCUCCGCGAAA-GCUAUUUAG
GUAGCGCCUCGG-ACG--------------------------------------------------------
----------------------AA-----UA-CCUCGGGGGGUAGAGC-ACUGCAU-GGA-UGA--UGGGGG
C---CCACA--GCCUUACUGAGUCUAAGCAAACUCCGAAUACCCGA--------------------------
------------------------------------------------------------------------
------------------GAGU-A-CU--------AUCCGGG-AGACACA-C-GGCG-G-GUGCUAACGUCC
GU-C-GUGAAGAG-GGAAAC---AACCCUGACCUGCA-GCUAAGGCCCCCAAUU--CG-UGG-CU-AA---G
-U-GGG-------AAAGCAUGUGGGACGGCCAAAACAACCAGGAGGUUGGCUUAGAAGCAGCCA-UCCUUUA
AAGAUAGCGUAACAGCUCACUGGUCUAGAUAAGCUGUCCUGCGGCGAAGAUGUAAC-GGGGCUCA-AGCCAC
G----------AGCCGAA-GCUCAGGAUGCACAG--C-----------------------------------
------------------------------------------------------------------------
------------------------------------------------------------------------
---------------------------------AAUG-UGCG-UGGUAGCGGAGC-GUUCCGU---------
-----------GAU-UCUGU-GAAGCCG--G-GCCGUAAGGCAUC-CGGUGGAGAG-A-UCG-GAAGCGAGA
AUGU-UGACAU-GAGUA-GCGACA---AACA-GGGUGA--G---------------AGACCC-UGU-CGCCG
A-AAGUCC-AAGGGUUCC-UGCUUAAAG-CUAAUCUGAGCAGGGUGAGCCGGC-CC-CUAAGGCGAGGCCG-
A-A-AGGC-----GUA--------------------------------------------------------
----------------------------------------GUC-GA-U--GGGA-ACCAGGUU-AAUA-UUC
CUGGG----------------------------------CCAGG----AG----------------------
------------------------------------------------------------------------
----------------------------------------------------------------GAUGUGAC
GGAUCG--------------CAGG-U-GUAGUUCGG-CC-U----UAUCG------------GAUUGGCCG-
GG----------------------------------------------------------------------
------------------------------------------------------------------------
------------------------------------------------------------------------
--------------------------C-UGC-U----GAG-CGGUCC-C-UGGAAAUAGCCC-UC-------
---------------------------------------------------------------CA-UCAGAC
CGUACCCUAAACCGAC------------------------------ACAGGUG--GACUGGUA-GAGAAUAC
CAAGGC-G-CU-UGAG-AGAACCAC-A-UCAAAGGAACUCGGC-AAAAUGCCUCCGUAAGUUCGC-GAGAAG
GAGGC-CCCG-U-CU--UUGGG--------------------------------------------------
------------------------------------------------------------------------
------------------------------------------------------------------------
------------------------------------------------------------------------
------------------------------------------------------------------------
------------------------------------------------------------------------
------------------------------------------------------------------------
------------------------------------------------------------------------
------------------------------------------------------------------------
------------------------------------------------------------------------
------------------------------------------------------------------------
------------------------------------------------------------------------
------------------------------------------------------------------------
------------------------------------------------------------------------
---------------------------------------------------------------CAACCAGAG
```

GCG--GGGGGCACAAACCAGGGGGUGGCGA-----CUGUUUACUUAAAACACAGGGCUCUGCGAAGCCG---
-CA-A-------------GGCGACGU-AUAGGGUCUGA-CGCCUGCCCGGUGCUG-G-AAGG-UUAAAAGGA
GAGGU----------GCAA----------GCCUUGAAUUGAAGCCC--CAGUAAAC-GGCGGCCGUAACUAU
AACGGUCCUAAGGUAGCGAAAUUCCUUGUCGGG-UAAGUUCCGACCUGCACGAAUGGCGUAACGAUC--UCC
CCGCUGUCUCUGAU----GUG-GACUCAGCGAAAUU--GAA---CUGUGUGUCAAGAUGCACACUUCCCGCG
GUUAGACGGAAAGA-CCCCAUGAACC-UUUACU--AUAGCUU-CGCACUGGCAUCAGGAAUGUGAUGUGCAG
GAU-AGGUGGUAGGCAUCG----------AAACGGG-GAC----------------------------GCCA
--------------------------------------GU-UCCCGUGGAG-C--C-AA-CCUUGAGAUA-C
CACC-CUUCGCAUUCUUGAUGUCUAACCGCGG--UCCA-U-UA-C-C-------------------------
------------------------------------------------------------------------
------------------------------------------------------------------------
------------------------------------------------------------------------
------------------------------------------------------------------------
------------------------------------------------------UG----GA----------
------------------------------------------------UCCG-----GGACC-CUGCGUGGU
GGGUAGUUUGACUGGGGCGGU-CGCCUCCCA-AACAGUAAC-GGAGGCGCGCG-AUGGUGGGCUCAGACCGG
UCGGAAAUCGGUCGUC-GAGUG-CAAUGGCAG-AAGCCCGCCUGACU-GCAA--------------------
--------------GACUGA---CAAGUCGAGCAGAGACGAAA-GUCGGCCAUAGUGAUCCGGU--------
---GG-U-----CCCGA-GUG--------------------GA-AGGGCCAUCGCUCAACGG-AUAAAAGGU
ACUCUGGGG-AUAACAGGCUGAUGAUGCCCAAGAGUCCAUAUCGACGGCAUCGUUUGGCACCUCGAUGUCGG
CUCAUCUCAUCCUGGGGCUGGA-GCAGGUCCCAAGGGUAUGGCUGUUCGCCAUUUAAAGAGGUACGUGAGCU
GGGUUUAGAACGUCGUGAGACAGUUCGGUCCCUAUCUGCC--------------------G-UGG-------
-------------------------------GUGU--AGGAGACUUGAGAAG-AGU-UGCCCCUAGUACGAG
AGGAC-CGGGGUGAACGAACCAC-UGGU-G-GACCAGUUGUCGUGCCAACGGCAG-U-GCUGGGUAGCUAUG
UUCGGACAGGAUAACCGCUGAAGGCA-UC--UAAGCGGGAAGCCCCCUUCAA-A-ACA--AGGUCUCC-CU-
------------------------------------------------------------------------
------------------------------------------------------------------------
------------------------------------------------------------------------
------------------------------------------------------------------------
------------------------------------------------------------------------
------------------------------------------------------------------------
------------------------------------------------------------------------
------------------------------------------------------------------------
----------------------------------------------------------UG-AGG------GC
CGUGGAAGACCACCACGU-CGA-UAGGCC-AGA-GGUGUAAG-CGC-A----------------GC------
----AA-----------UGCGUUCAGCUG--A-CUGG-UACUAAUU---GCCCGAUUG-GCUUGAUCU----
------------------------------------------------------------------------
------------------------------------------------------------------------
------------------------------------------------------------------------
------------------------------------G------------
>LSUb_279238_NOVAD/1-2754
------------------------UGAGG----------UAAGAGCAUUU-GGUGGAUG-C-CUUGGC-AUG
UAC---AGGCGAUGAAGGACGUGG--CACGCUGCGAUAAGCGUCGGGGAGUUGUGAGCAAACUU------UG
AUC-CGG-CGAUUUCCGAAUGGGGAAAC--------CC--------AC----CUUC-A-CCAU-------UU
C-U----------GAGG---U----G----GAUAAGG----U-A-UCCC-AGA-----UUGA----------

```
-----------A-----------UACA----------------------------------------UA-GA
U-C--------------------UG-GUGA-AGCGAACCCGGGGAACUGAAACAUCUCAGUACCUG--GAGG
AAAAGACAUCAA--CCGAGAUUCCGUUA--GU-AG-UGGCGAGCGAACGCGGA-C-CA-GGCCA--------
------------------------------------------------------------------------
----------GUG-------C----CU--------------------------CAU-----------CU---
U-------CAACU-AGCAG------AAC-AC-UUUG------------------------------------
-------GAA-AG-AGUGG-------CCAUAGCGGGUGACAGCCCGUA----------UGCGAAA----GUG
A------U--GGAUG-----------------------AG-GA------CUCGAGUAGGGCGGGACACGUGA
AAUCCUGUCUGAA-CAUGGGGGG-ACC--ACCCUCCAAGCCUAAAUA--CUC-GUACA-UGACCGAUAGC-G
AACAC-AGUACCGUGAGGGAAAG-GUGAAA-AGCAC-CCCGAUGAGGGG-A-GUGAAACAGUACCUGAAACC
G-AAUGCUUACAA-GCA-GU-UGGA---GCCU-C--------------------------------------
------------------------------------------------------------------------
------------------------------------------------------------------------
------------------------------------------------------------------------
------------------------------------------------------------------------
------------------------------------------------------------------------
------------------------------------------------------------------------
------------------------------------------------------------------------
------------------------------------------------------------------------
-------------------------------------------------------------UU--U-AG---
------------------------------------------------------------------------
------------------------------------------------------------------------
------------------------------------------------------------------------
------------------------------------------------------------------------
------------------------------------------------------------------------
------------------------------------------------------------------------
------------------------------------------------------------------------
------------------------------------------------------------------------
------------------------------------------------------------------------
------------------------------------------------------------------------
------------------------------------------------------------------------
------------------------------------------------------------------------
------------------------------------------------GG--G---GUGACAGCGUACCUCU
UGCAUAAUGGGUCAGUGACUUAG-U-UUAUCGAGCAA-GCUUAAGC--C-----------GUUA--------
--GG--UGUA--GGCGUAGCGAAAG-C---------------------------------------------
------------------------------------------------------------------------
------------------------------------------------------------------------
---------------------------------------------------------------------GAG
UCU------------------------------------GAAUA-------------------------GG-
G-C-GC---------------------UGAAGUUCGAUGAAUUA-GACCCGAAACC-CGGUGAUCUAGGCAU
GACCAGGCUGAAGGUGCG-GUAACA-CGCACUGGAGGGCCGA--ACCGUUGA-AUGUUGAAAAAUUC-UCGG
AUGAGUUGUGUUUAGGGGUGAAAGGCCAAUCAAACCGGG-AAAUAGCUGGUUCUCCGCGAAA-UCUAUUGAG
GUAGAGCGUCGA-AUG--------------------------------------------------------
----------------------UA-----UG-CCUCUGGGGGUAGAGC-ACUGGAU-GGA-UGC--GGGGGU
C---GC-GA--GAUCUACCAAUUCUAACCAAACUCCGAAUACCAGA--------------------------
```

```
------------------------------------------------------------------------
------------------GAGA-C-UU--------GUUCGGC-AGACAGA-C-GGCG-G-GUGCUAAGGUCC
GU-C-GUCAAAAG-GGAAAC---AGCCCUAACCUACA-GCUAAGGUCCCCAAGU--CA-UCA-CU-AA---G
-U-GGG-------AAAGCAUGUGGGAAUCCCAAAACAACCAGGAGGUUGGCUUAGAAGCAGCCA-UCCUUUA
AAGAAAGCGUAACAGCUCACUGGUCUAAACAAGGGUUCCUGCGGCGAAGAUGUAAC-GGGGCUAA-AGUGAU
G----------CACCGAA-GCUUAGGGUGUGAUC--U-----------------------------------
------------------------------------------------------------------------
------------------------------------------------------------------------
-------------------------------UCGGAU-CACG-CGGUAGCGGAGC-GUUCCGU----------
-----------AAG-CCUGU-GAAGCGG--U-CUGGUAAUGG--G-CCGUGGAGGU-A-UCG-GAAGUGCGA
AUGC-UGACAU-GAGUA-GCGACA---AAGA-GGGUGA--G---------------AUGCCC-UCU-CGCCG
A-AAGACC-AAGGGUUCC-UGCUUAAAG-CUAAUCUGAGCAGGGUGAGCCGGC-CC-CUAAGACGAGCCCG-
A-A-GGGG-----GUA--------------------------------------------------------
----------------------------------------GUC-GA-U--GGGA-ACCACGUU-AAUA-UUC
GUGGG----------------------------------CCUGG----UGG---------------------
------------------------------------------------------------------------
----------------------------------------------------------------UGUGUGAC
GGAUCA--------------CGUG-U-GUUGUCUGG-CC-U----UAUUG------------GAUUGGUCA-
GG----------------------------------------------------------------------
------------------------------------------------------------------------
------------------------------------------------------------------------
--------------------------C-UUC-G----AA--GUGGUU-C-CAGGAA-AUAGC-CCCACC---
---------------------------------------------------------------GU-AUAGAC
CGUACCCGAAACCGAC------------------------------ACAGGUG--GUCAGGUA-GAGUAUAC
CAAGGC-G-CU-UGAG-AGAAGUAU-C-CUGAAGGAACUCGGC-AAAUUGCCUCCGUACCUUCGG-AAGAAG
GAGGC-CCAA-C-CU--GUGCG--------------------------------------------------
------------------------------------------------------------------------
------------------------------------------------------------------------
------------------------------------------------------------------------
------------------------------------------------------------------------
------------------------------------------------------------------------
------------------------------------------------------------------------
------------------------------------------------------------------------
------------------------------------------------------------------------
------------------------------------------------------------------------
------------------------------------------------------------------------
------------------------------------------------------------------------
------------------------------------------------------------------------
------------------------------------------------------------------------
-------------------------------------------------------------CAA--GCACUU
GUU--GGGGGCACAGGCCAGGGGGUAGCGA-----CUGUUUAGCAAAAACACAGGACUCUGCUAAGUCGGCU
UCA-A-------------GACGACGU-AUAGGGUCUGA-CGCCUGCCCGGUGCUC-G-AAGG-UUAAGAGGA
GGAGU----------GCAA----------GCUCCGAAUUGAAGCCC--GAGUAAAC-GGCGGCCGUAACUAU
AACGGUCCUAAGGUAGCGAAAUUCCUUGUCGGG-UAAGUUCCGACCUGCACGAAUGGCGUAACGACU--UCC
CCACUGUCUCCAGG----AUA-UGCUCAGCGAAAUU--GAA---UUCUCCGUGAAGAUGCGGAGUACCCGCG
```

```
GUUAGACGGAAAGA-CCCCGUGCACC-UUUACU--GCAGCUU-CAGAGUGGCAUUAGGAAAGAAUUGUGUAG
CAU-AGGUGGGAGGCUUUG----------AAGCACC-GGC----------------------------GCCA
--------------------------------------GC-UGGUGUGGAG-C--C-AU-AGGUGAAAUA-C
CACC-CUGUUGUUUUCUGAUGUCUAACCCAGG--ACCG-U-UA-G-C-------------------------
------------------------------------------------------------------------
------------------------------------------------------------------------
------------------------------------------------------------------------
------------------------------------------------------------------------
------------------------------------------------------CG----GU----------
------------------------------------------------UCGG-----GGACC-CUCUGUGGC
GGGUAGUUUGACUGGGGCGGU-CGCCUCCUA-AAGAGUAAC-GGAGGCGCGCG-AUGGUAGGCUCAGGCCGG
UUGGAAACCGGCUGUUAGAGUG-CAAUGGCAU-AAGCCUGCCUGACU-GCGA--------------------
--------------GACUGA---CGAGUCGAGCAGAGACGAAA-GUCGGUCAUAGUGAUCCGGU--------
---GG-U-----CCCUC-GUG--------------------GA-AGGGCCAUCGCUCAACGG-AUAAAAGGU
ACGCCGGGG-AUAACAGGCUGAUGAUUCCCAAGAGCUCAUAUCGACGGAAUCGUUUGGCACCUCGAUGUCGG
CUCAUCACAUCCUGGGGCUGGA-GCAGGUCCCAAGGGUUUGGCUGUUCGCCAAUUAAAGUGGUACGUGAGCU
GGGUUCAGAACGUCGCGAGACAGUUUGGUCCCUAUCUGCC--------------------G-UGG-------
-------------------------------GCGU--C-GAUACUUGAGAGG-AGU-UGCCCCUAGUACGAG
AGGAC-CGGGGUGAACAUGCCUC-UGGU-G-UACCUGUCGUUCCGCCAGGAGCGC-A-GCAGGGUAGCUAUG
CAUGGACGGGAUAACCGCUGAAAGCA-UC--UAAGCGGGAAGCCUCCCUCAA-G-AUA--AGGUAUCA-UC-
------------------------------------------------------------------------
------------------------------------------------------------------------
------------------------------------------------------------------------
------------------------------------------------------------------------
------------------------------------------------------------------------
------------------------------------------------------------------------
------------------------------------------------------------------------
------------------------------------------------------------------------
--------------------------------------------------------------GA------GU
CGUGGUAGACCACCACGU-UGA-UAGGCC-GGG-UGUGGAAG-UGC-G----------------GU------
----AA-----------CGCAUGAAGCUA--A-CCGG-UCCUAAUA---ACUCU-----GUUCAUGCU----
------------------------------------------------------------------------
------------------------------------------------------------------------
------------------------------------------------------------------------
------------------------------------U------------
&gt;LSUb_555217_ZYMMA/1-2758
---------------------A-------------GG--UAAGAGCAUUU-GGUGGAUG-C-CUAGGC-AUA
CAC---AGGCGAUGAAGGACGUGG--CACGCUGCGAUAAGCUACGGCGAGAUGUGAGCAAUCUU------UG
ACC-CGU-AGAUUUCCGAAUGGGGAAAC--------CC--------AU----CCUC-A-CCAUUUA-----A
G-U-------UAGGUGG---G----A----A---GGA----U-A-UUAC-CAA-----AGUGA---------
-----------A-----------UACA---------------------------------------UAGC-U
U-U--------------------GG-U-GA-AGCGAACCCGGCGAACUGAAACAUCUAAGUACCCG--GAGG
AAAAGACAUCAA--CCGAGAUUCCGUUA--GU-AG-UGGCGAGCGAACGCGGA-G-UA-GGCCA--------
------------------------------------------------------------------------
----------GUA-------C----CU--------------------------UUG-----------AU---
```

```
U-------GCGUU-AGCAG------AAG-UC-UUUG------------------------------------
-------GAA-AG-AGACA-------CCAUAGUGGGUGACAGUCCCGUA---------UGUUAAA----ACA
A------U--AUUGA-----------------------AG-GU------CUUGAGUAGGGCGGGGCACGUGA
AACCUUGUCUGAA-CAUGGGGGG-ACC--ACCCUCCAAGCCUAAAUA--CUC-GUGUA-UGACCGAUAGU-G
AAC-A-AGUACCGUGAGGGAAAG-GUGAAA-AGAAC-CCCGAUUAGGGG-A-GUGAAACAGUUCCUGAAACC
G-AAUGCUUACAA-GCA-GU-AGGA---GGGU-C--------------------------------------
------------------------------------------------------------------------
------------------------------------------------------------------------
------------------------------------------------------------------------
------------------------------------------------------------------------
------------------------------------------------------------------------
------------------------------------------------------------------------
------------------------------------------------------------------------
------------------------------------------------------------------------
-------------------------------------------------------------UU--U-AA---
------------------------------------------------------------------------
------------------------------------------------------------------------
------------------------------------------------------------------------
------------------------------------------------------------------------
------------------------------------------------------------------------
------------------------------------------------------------------------
------------------------------------------------------------------------
------------------------------------------------------------------------
------------------------------------------------------------------------
------------------------------------------------------------------------
------------------------------------------------------------------------
------------------------------------------------------------------------
------------------------------------------------GG--C---CUGACUGCGUACCUCU
UGCAUAAUGGGUCUGUGACUUAA-U-GUAUCAAGCAA-GCUUAAGC--C-----------GAUA--------
--GG--UGUA--GGCGCAGCGAAAG-C---------------------------------------------
------------------------------------------------------------------------
------------------------------------------------------------------------
---------------------------------------------------------------------GAG
UCU------------------------------------GAAUA-------------------------GG-
G-C-GA---------------------UUUAGUUUGAUGCAUUA-GACCCGAAACC-CGGCGAUCUAGGCAU
GGUCAGGAUGAAGGUAAG-GUAACA-CUUACUGGAGGUCCGA--ACCGAUUA-ACGUUGAAAGUUA-UCGG
AUGAACUGUGUUUAGGGGUGAAAGGCCAAUCAAGCCGGG-AAAUAGCUGGUUCUCCGCGAAA-ACUAUUGAG
GUAGUGCCUCAU-GUG--------------------------------------------------------
----------------------AU-----GA-CCGUUGGGGGUAGAGC-ACUGGAU-GGA-UGC--GGGGGU
U---UC-GC--GACCUACCAAAUCUAACCAAACUCCGAAUACCAAC--------------------------
------------------------------------------------------------------------
------------------GAGU-U-UA--------ACAUGGG-AGACAGA-C-GGCG-G-GUGCUAAGGUCC
GU-C-GUCGAGAG-GGAAAC---AGCCCUGACCUACA-GCUAAGGUCCCCAAGU--CA-UGU-CU-AA---G
-U-GGG-------AAAGCUUGUGAAGACCCCAAAACAACCAGGAGGUUGGCUUAGAAGCAGCCA-UCCUUUA
AAGAAAGCGUAACAGCUCACUGGUCUAAAUAAGGGUCUUUGCGGCGAAGAUGUAAC-GGGGCUCA-AGACAU
```

```
G----------CACCGAA-GCUUAGGAUUGUUAC--A-----------------------------------
------------------------------------------------------------------------
------------------------------------------------------------------------
-------------------------------UUAGUA-ACAG-UGGUAGCGGAGC-GUUCCGU---------
-----------AAG-CCUGU-GAAGCAG--U-CUGGUAAUGG--A-CUGUGGAGGU-A-UCG-GAAGUGCGA
AUGC-AGACAU-GAGUA-GCGAUA---AAUA-GGGUGA--G---------------AUGCCC-UAU-CGCCG
A-AAGUCC-AAGGGUUCC-UGCGCAAGG-CUAAUCCGCGCAGGGUGAGUCGGC-CC-CUAAGACGAGCCCG-
A-A-GGGG-----GUA--------------------------------------------------------
----------------------------------------GUC-GA-U--GGGA-AACAGGUU-AAUA-UUC
CUGUA----------------------------------CCUGG----AGA---------------------
------------------------------------------------------------------------
----------------------------------------------------------------GAUGUGAC
GGAUCG--------------UGUA-A-AUUGUAUGU-CC-U----UAUCG------------GAUUGGACA-
UG----------------------------------------------------------------------
------------------------------------------------------------------------
------------------------------------------------------------------------
--------------------------C-UCA-U----AAG-CGGUUC-C-AGGAAAUAGCCU-CUCC-----
---------------------------------------------------------------AU-AUAGAC
CGUACCCUAAACCGAC------------------------------ACAGGUG--GACAGGUA-GAGUAUAC
CAAGGC-G-CU-UGAG-AGAAGGGU-G-UUGAAGGAACUCGGC-AAAUUGCCUCCGUACCUUCGG-AAGAAG
GAGGC-CCUU-A-UU--AUAGG--------------------------------------------------
------------------------------------------------------------------------
------------------------------------------------------------------------
------------------------------------------------------------------------
------------------------------------------------------------------------
------------------------------------------------------------------------
------------------------------------------------------------------------
------------------------------------------------------------------------
------------------------------------------------------------------------
------------------------------------------------------------------------
------------------------------------------------------------------------
------------------------------------------------------------------------
------------------------------------------------------------------------
------------------------------------------------------------------------
---------------------------------------------------------------CAACUAUUC
UAA--GGGGGCACAGGCCAGGGGGUAGCGA-----CUGUUUAGCAAAAACACAGGGCUCUGCAAAGUCGGCU
UCA-A-------------GACGACGU-AUAGGGCCUGA-CGCCUGCCCGGUGCUG-G-AAGG-UUAAGUGGA
GGAGU----------GCAA----------GCUCUGAAAUGAAGCCC--CAGUAAAC-GGCGGCCGUAACUAU
AACGGUCCUAAGGUAGCGAAAUUCCUUGUCGGG-UAAGUUCCGACCUGCACGAAUGGCGUAACGACU--UCC
CCACUGUCUCCAAC----ACC-UGCUCAGCGAAAUU--GAA---UUCUCCGUGAAGAUGCGGAGUACCCGCG
GUUAGACGGAAAGA-CCCCGUGCACC-UUUACU--GCAGCUU-CAGAGUGGCAUUAGGAAAGAACUGUGUAG
CAU-AGGUGGGAGGCUUUG----------AAACUUG-AGC----------------------------GCCA
--------------------------------------GC-UUGAGUGGAG-C--C-AU-AGGUGAAAUA-C
CACC-CUGUUGUUUCUGAUGUCUAACCUAGA--ACCA-UGAA-A-C-------------------------
------------------------------------------------------------------------
```

```
------------------------------------------------------------------------
------------------------------------------------------------------------
------------------------------------------------------------------------
------------------------------------------------------UG----GU----------
------------------------------------------------UCAG-----GGACC-CUCUGUGGC
GGGUAGUUUGACUGGGGCGGU-CGCCUCCUA-AAGAGUAAC-GGAGGCGCGCG-AUGGUUGGCUCAGGCCGG
UUGGAAACCGGCUGCAAGAGUG-CAAUGGCAU-AAGCCAGCCUGACU-GUGA--------------------
--------------GACUGA---CAAGUCGAACAGAGACGAAA-GUCGGUCAUAGUGAUCCGGU--------
---GG-U-----CCCUC-GUG--------------------GA-AGGGCCAUCGCUCAACGG-AUAAAAGGU
ACGCCGGGG-AUAACAGGCUGAUAACCCCCAAGAGCUCAUAUCGACGGGGUUGUUUGGCACCUCGAUGUCGG
CUCAUCACAUCCUGGGGCUGGA-GCAGGUCCCAAGGGUUUGGCUGUUCGCCAAUUAAAGUGGUACGUGAGCU
GGGUUCAGAACGUCGCGAGACAGUUUGGUCCCUAUCUGCC--------------------G-UGG-------
-------------------------------GCGU--C-GAUAUUUGAGAGG-AGU-UGCCCUUAGUACGAA
AGGAC-CGGGGUGAACAUGCCUC-UGGU-G-GACCUGUCGUGGCGCCAGCCGCGC-A-GCAGGGUAGCUAUG
CAUGGACGGGAUAACCGCUGAAAGCA-UC--UAAGCGGGAAGCCUCCCUCAA-G-AUA--AGAUAUCU-C--
------------------------------------------------------------------------
------------------------------------------------------------------------
------------------------------------------------------------------------
------------------------------------------------------------------------
------------------------------------------------------------------------
------------------------------------------------------------------------
------------------------------------------------------------------------
------------------------------------------------------------------------
-------------------------------------------------------------AGA------GC
CGUCGAAGACUACGACGU-UGA-UAGGCC-AGA-UGUGGAAG-UGU-G----------------GU------
----AA-----------CACAUGUAGCUA--A-CUGG-UCCUAAUU---GCUCUAUUC-ACGCUUUUG----
------------------------------------------------------------------------
------------------------------------------------------------------------
------------------------------------------------------------------------
-------------------------------------------------
>LSUb_156889_MAGMM/1-2846
---------------------A-AGUAAAC---------GAAGGGCAUAC-GGUGGAUG-C-CUUGGU-GCC
AAG---AGGCGAUGAAGGACGUAG--CACGUUGCGAUAAGCUUCGGAGAGCCGCGAGCAGGCUU------UG
ACC-CGG-AGAUUUCCGAAUGGGGAAAC--------CC--------GG----CGGG-A-U-----------G
A-A-------------A---U----C----C---CGU----C-A-CUAC-UAG-----CUGA----------
-----------A-----------UAAA----------------------------------------UA-GG
C-U--------------------AU-U-AG-AGCGAACGCGGGGAACUGAAACAUCUAAGUACCCG--UAGG
AACAGAAAUCAA--UUGAGAUUCCCUAA--GU-AG-CGGCGAGCGAACGGGGA-C-CA-GGCCA--------
------------------------------------------------------------------------
----------GUA-------U----CA---------------------------UA-----------AA---
U-------CAUUU-AGCCG------AAC-CA-ACUG------------------------------------
-------GAA-AG-UUGGA-------CCAUAGUGGGUGAAAGUCCUUUA---------GGUGAAA----GGG
C------G--UGUAU-----------GA-----GGG--AU-AC------GAAGAGUAGGGCGGGACACGUGA
AAUCCUGUCUGAA-CAUGGGGGG-CCC--UCCCUCCAAGCCUAAAUA--CUA-CUUGG-CAACCGAUAGU-G
AAC-C-AGUACCGUGAGGGAAAG-GUGAAA-AGAAC-CCCGAUAAGGGG-A-GUGAAAUAGAACCUGAAACC
```

```
G-UAUGCUUACAA-GCA-GU-CGGA---GGCU----------------------------------------
------------------------------------------------------------------------
------------------------------------------------------------------------
------------------------------------------------------------------------
------------------------------------------------------------------------
------------------------------------------------------------------------
------------------------------------------------------------------------
------------------------------------------------------------------------
------------------------------------------------------------------------
-------------------------------------------------------------GA--U-AA---
------------------------------------------------------------------------
------------------------------------------------------------------------
------------------------------------------------------------------------
------------------------------------------------------------------------
------------------------------------------------------------------------
------------------------------------------------------------------------
------------------------------------------------------------------------
------------------------------------------------------------------------
------------------------------------------------------------------------
------------------------------------------------------------------------
------------------------------------------------------------------------
------------------------------------------------------------------------
-------------------------------------------------G--C---CUGACGGCGUACCUUU
UGUAUAAUGGGUCAGCGAGUUAA-U-CUGCGUAGCAA-GCUUAAGC--------------GAAA--------
---G--UGUA--GGCGAAGCGAAAG-C---------------------------------------------
------------------------------------------------------------------------
------------------------------------------------------------------------
---------------------------------------------------------------------GAG
UCU------------------------------------GAAUA-------------------------GG-
G-C-GU----------------------UAAGUUUCGUAGAUUA-GACCCGAAACC-GAGUGAUCUACCCAU
GGUCAGGUUGAAGGUGCG-GUAACA-CGCACUGGAGGACCGA--ACCCACCU-AUGUUGAAAAAUGG-GGGG
AUGAACUGUGGGUAGGAGUGAAAGGCUAAACAAACUCGG-AAAUAGCUGGUUCUCUCCGAAA-ACUAUUUAG
GUAGUGCCUCGA-AUU--------------------------------------------------------
-----------------------G-----GC-AUGCCGGGGGUAGAGC-ACUGGAU-GGG-CUA--GGGGGG
C---CCAAA--GCCUUACCAAACCUAACCAAACUCCGAAUACCGGU--------------------------
------------------------------------------------------------------------
------------------CAGUUA-UA--------GUUCGGG-AGACAGA-C-UGCG-G-GUGCUAAGGUCC
GU-A-GUCGAGAG-GGAAAA---AGCCCAGACCAUCA-UCUAAGGUCCCCAAGU--CA-UAC-UU-AA---G
-U-GGG-------AAAGGAUGUGGGAAUGCCCUGACAGCCAGGAGGUUGGCUUAGAAGCAGCCA-CCCUUUA
AAGAAAGCGUAAUAGCUCACUGGUC-----GAGUGGUCCUGCGCCGAAAAUAUAAC-GGGGCUCA-AAGUAU
G----------CACCGAA-GAUAUGGAUGCCGCG--------------------------------------
------------------------------------------------------------------------
------------------------------------------------------------------------
------------------------------------A-GGCA-UGGUAGGAGAGC-GUUCCGU---------
-----------AAG-CCGUU-GAAGCCG--U-ACCGAGAGGA--G-CGGUGGAGGU-A-UCG-GAAGUGAGA
```

```
AUGC-UGACAU-GAGUA-ACGAUA---AUCA-AGGUGA--G---------------AAUCCU-UGA-CGCCG
A-AAACCC-AAGGUUUCC-UACGCAAGG-CUAUUCCGCGUAGGGUUAGUCGGC-CC-CUAAGGUGAGGCAU-
A-C-AUGC-----GUA--------------------------------------------------------
----------------------------------------ACC-GA-U--GGGA-AACAGGUU-AAUA-UUC
CUGUA----------------------------------CCUCU----UAUA--------A-CU--------
------------------------------------------------------------------------
----------------------------------------------------------GCGAU-GGGGGGAC
GGAGAA--------------GGAUAG-AUCUACCGG-GC-G----UUGGU------------AGUCCCGGG-
ACAAGC-GUG-UA-----G-G-------------------U------UG------------------ACAGA
C-UAGG--------------------UAAAU-CCGGACUGUU--------AAGACUG----AGG-CG-U-G-
AUG--UG-----------------------------------------------------------------
-----GAGCCA----CUAGGCGAAC-U-GAU-U-GAUUCCAAGCUUC-C-AAGAAA-AGCCU-CUAAGCUU-
-------------------------------------------------------CAGGUUA-UG-AGAGAC
CGUACCCCAAACCGAC------------------------------ACAGGUG--GGUGGGAU-GAGAAUUC
UAAGGC-G-UG-UGAG-AGAACUCU-G-GUAAAGGAACUCGGC-AAAU--GCCCCGUAACUUCGG-GAGAAG
GGGUG-CCUC-U-UG-UAUGUGAAG----GCC-----------UUUAC------------------------
------------------------------------------------------------------------
------------------------------------------------------------------------
------------------------------------------------------------------------
------------------------------------------------------------------------
------------------------------------------------------------------------
------------------------------------------------------------------------
------------------------------------------------------------------------
------------------------------------------------------------------------
------------------------------------------------------------------------
------------------------------------------------------------------------
------------------------------------------------------------------------
------------------------------------------------------------------------
------------------------------------------------------------------------
--------------------------------------------------------GGCU-GGA--GCAGAA
GGG--GGCCGCAGUGACUAGGGGGGAGCGA-----CUGUUUACCAAAAACAUAGGACUCUGCUAAAUCG---
-CA-A-------------GAUGAAGU-AUAGGGUCUGA-CGCCUGCCCGGUGCUG-G-AAGG-UUAAGAGGA
GGGGU----------GCAA----------GCUCUGAAUCGAAGCCC--CAGUAAAC-GGCGGCCGUAACUAU
AACGGUCCUAAGGUAGCGAAAUUCCUUGUCGGG-UAAGUUCCGACCUGCACGAAUGGCGUAACGACU--UCC
CCGCUGUCUCUAUC----AGA-GGCUCAGUGAAAUU--GAA---UUUGCUGUGAAGAUGCAGCAUACCCGCG
GCAAGACGGAAAGA-CCCCGUGAACC-UUUACU--AUAGCUU-UGCAUUGGUACUAGAGACAGUUUGUGUAG
GAU-AGGUGGGAGGCUUUG----------AAGCGUG-AGC----------------------------GCCA
--------------------------------------GC-UCAUGUGGAG-C--C-AC-CCUUGAAAUA-C
CACC-CUGGCUCUCUUUGGUAUCUAACUGCGC--GCCG-U-UA-U-C-------------------------
------------------------------------------------------------------------
------------------------------------------------------------------------
------------------------------------------------------------------------
------------------------------------------------------------------------
------------------------------------------------------CG----GC----------
------------------------------------------------GUCA-----GGACA-GUGCAUGGU
```

```
GGGUAGUUUGACUGGGGCGGU-CGCCUCCCA-AACAGUAAC-GGAGGCGCACG-AUGGUUCGCUCAGGUUGG
UCGGAAAUCAACCGUA-GAGUG-UAAUGGCAU-AAGCGAGCCUGACU-GUGA--------------------
--------------GACUGA---CAAGUCGAGCAGGUACGAAA-GUAGGUCAUAGUGAUCCGGC--------
---GA-U-----UCCGA-GAG--------------------GA-AGGGUCGUCGCUCAACGG-AUAAAAGGU
ACUCCGGGG-AUAACAGGCUGAUCUCCCCCAAGAGUUCACAUCGACGGGGAGGUUUGGCACCUCGAUGUCGG
CUCAUCGCAUCCUGGGGCUGUA-GUUGGUCCCAAGGGUUUGGCUGUUCGCCAAUUAAAGCGGUACGCGAGCU
GGGUUCAGAACGUCGUGAGACAGUUCGGUCCCUAUCUGCC--------------------G-UGG-------
-------------------------------GCGU---GGAAAAUUGAGAGG-AUC-UGCUCCUAGUACGAG
AGGAC-CGGAGUGGACAAGCCUC-UGGU-G-UUCGGGUUGUCGUGCCAACGGCAU-C-GCCCGGUAGCUACG
CUUGGAAGGGAUAACCGCUGAAAGCA-UC--UAAGCGGGAAACCCACCUCAA-G-AUG--AAUUUUCC-CU-
--A---------------------------------------------------------------------
------------------------------------------------------------------------
------------------------------------------------------------------------
------------------------------------------------------------------------
------------------------------------------------------------------------
------------------------------------------------------------------------
------------------------------------------------------------------------
------------------------------------------------------------------------
----------------------------------------------------------UG-AGA------UC
CGUGGAAGACCACCACGU-UGA-UAGGCA-GGA-UGUGGAAG-UGG-G----------------GU------
----AA-----------CCCAUGCAGCUA--A-CCUG-UACUAAUC---GAUCCACUG-ACUUCUA------
------------------------------------------------------------------------
------------------------------------------------------------------------
------------------------------------------------------------------------
-------------------------------------------------
>LSUb_292805_WOLTR/1-2789
---------------------UAAAUAAAUU----AU--UAAGAGCACUU-GAUGGAUG-C-CUUGGC-GUU
AAG---AGGCGAUGAAGGACGUGG--CAGGCUGCGAUAAGCUGCGGGAAUUGUCAACAGGUUU------UG
AUC-CGC-AGAUUUCCGAAUGGGAAAC--------CC--------AA----CUAG-U-------------U
G-A-----------------G----C----U---AGU----U-A-UUGC-AUG-----UU------------
-----------A-----------GAUU------------------------------------------A-A
U-A--------------------UG-U-AA-AGCGAACUUGGUGAACUGAAAUAUCUAAGUAGCCA--AAGG
AAAAGAAAUCAA--CCGAGAUUCUGUUA--GU-AG-UGACGAGCGAAAGCGGG-A-AA-GGCUA--------
------------------------------------------------------------------------
----------GUG-------G----UU--------------------------UAA-----------GA---
A-------UAAGA-AUUAG------AAU-AC-UCUG------------------------------------
-------GAA-AG-AGUAA-------CCAUAGAAGGUGAUAGUCCUGUA---------UAGGUAG----AAA
G------U--UUUUA-----------------------AA-UC------CUCGAGUAGAGCGGGGCACGUGA
AAUCCUGUUUGAA-CAUGGGGGG-ACC--AUCCUCCAAGCCUAAAUA--CUC-CUUGA-CGACCGAUAGU-G
AAC-A-AGUACCGUGAGGGAAAG-GUGAAA-AGAAC-CCCGG-GAGGGG-A-GUGAAAUAGAAUCUGAAAUC
G-AGUGCUUACAA-ACA-GU-UGGA---GCUU-U---AU---------------------------------
------------------------------------------------------------------------
------------------------------------------------------------------------
------------------------------------------------------------------------
------------------------------------------------------------------------
```

```
------------------------------------------------------------------------
------------------------------------------------------------GU-UUUUGCUA-
----U-----------GG-G-------------------A--------------------------------
------------------------------------------------------------------------
--------------------------------------------------------------G--C-GA---
------------------------------------------------------------------------
------------------------------------------------------------------------
UCUUG-------------------------------------------------------------------
------------------------------------------------------------------------
------------------------------------------------------------------------
------------------------------------------------------------------------
------------------------------------------------U-----------------------
------------------------------------------------------------------------
------------------------------------------------------------------------
------------------------------------------------------------------------
------------------------------------------------------------------------
------------------------------------------------------------------------
-------------------AGCUAAUUU------------------AUAG--A---GUGACAGCGUACCUUU
UGCAUAAUGGGUCAGCGAGUUAA-U-CUAUAAAGCAA-GCUUAAGC--C-----------GUUA--------
--GG--UAUA--GGCAUAGCGAAAG-C---------------------------------------------
------------------------------------------------------------------------
------------------------------------------------------------------------
---------------------------------------------------------------------GAG
UCU------------------------------------GAAUA-------------------------UG-
G-C-GU---------------------UUUAGUUUUAUGGAUUA-GACCCGAAACC-AAGUGAUCUAGUCAU
GACCAGAUUGAAGGUGUG-GUAAAA-CACACUGGAGGAUCGA--ACCAGUUA-AUGUUGCAACAUUA-UUGG
AUGAGUUGUGAUUAGGGGUGAAAGGCCAAUCAAACUUGG-AAAUAGCUGGUUCUCCGCGAAA-UCUAUUUAG
GUAGAGCGUUGU-AUG--------------------------------------------------------
----------------------UA-----UG-UUGUUGGGGGUAGAGC-ACUUGAU-AGA-CUA--GGGGGG
C---UCACU--GUCUUACCAAAUCUAACUAAACUCCGAAUACCAAU--------------------------
------------------------------------------------------------------------
------------------AAUUGA-UU--------AUACAGC-AGACACA-C-UGCG-G-GUGCUAA-GUCC
GU-G-GUGGAAAG-GGAAAC---AGCCCAGAUUACUA-UCUAAGGUCCCAAAAU--UA-CAG-CU-AA---G
-U-GGG-------GAAGGAAGUAGAAAAACCAUUACAGCUAGGAGGUUGGCUUGGAAGCAGCUA-UCCUUUA
AAGAAAGCGUAACAGCUCACUUGUCUAAAUAAGUUUUUCUGCGCUGAAAAUGUACC-GGGGCUAA-AGCUGU
A----------UACCGAA-GAUGUAGGUACUUAU--U-----------------------------------
------------------------------------------------------------------------
----GAUUG---------------------------------------------------------------
----------------------------CAGUUAAUA-AGUG-CGGUAGCGGAGC-GUUCUGU---------
-----------AAA-UCUGU-GAAGGUA--G-UUUGUAAAAA--C-UGCUGGAGAU-A-UCA-GAAGUGAGA
AUGC-UGACAU-AAGUA-GCGUAA---AAGA-GUGUGA--A---------------AAACAC-UCU-CACCA
A-AAAUCU-AAGGGUUCC-UACGUUAAG-UUAAUCUGCGUAGGGUUAGUCGGU-UC-CUAAGGCAAGUCCGU
A-A-AGGA-----GUA--------------------------------------------------------
----------------------------------------GUU-GA-U--GGCA-AUCAGGUU-AAUA-UUC
CUGAA----------------------------------CCUCU----UA----------------------
```

```
------------------------------------------------------------------------
----------------------------------------------------------------AGUGUGAC
GGGUUU--------------UGUA-U-UUGUAUAGG-UC-U----UAUUG------------GAUUGACCU-
AUG---------------------------------------------------------------------
------------------------------------------------------------------------
------------------------------------------------------------------------
--------------------------C-UUA-A----AAA-AAGCUC-C-AGGAAAUAGCAC-UUAUAUU--
---------------------------------------------------------------UA-UGAGGC
CGUACCGUAAACCGAC------------------------------ACUGGUG--GAUGAGUA-GAGUAUAC
UAAGGU-G--U-UGAA-AGAAUGAU-G-UUGAAGGAACUCGGC-AAAUUGUACCUGUAACUUCGG-AAGAAG
GGUAA-CCUG-C-UU--UUAGG--------------------------------------------------
------------------------------------------------------------------------
------------------------------------------------------------------------
------------------------------------------------------------------------
------------------------------------------------------------------------
------------------------------------------------------------------------
------------------------------------------------------------------------
------------------------------------------------------------------------
------------------------------------------------------------------------
------------------------------------------------------------------------
------------------------------------------------------------------------
------------------------------------------------------------------------
------------------------------------------------------------------------
------------------------------------------------------------------------
---------------------------------------------------------------CAACUAUAA
GUA--GGUGGCACAAAAUAGGGAGUAGUGA-----CUGUUUACUAAAAACACAGGACUCUGCAAACACG---
-UA-A-------------GUGGAAGU-AUAGGGUCUGA-CGCCUGCCCGGUGCUG-G-AAGG-UUAAUAAGA
GGGGU----------GUAA----------GCUCUAAAUUGAAGCCC--CAGUAAAC-GGCGGCCGUAACACU
GACGGUCCUAAGGUAGCGAAAUUCCUUGUCGGG-UAAGUUCCGACCCGCACGAAUGGCGUAACAAUU--UCU
CCACUGUCUCCAAC----AUC-AAUUCAGCGAAAUU--GAA---UUCCCCGUGCAGAUGCGGGGUACCCGCG
GUUAGACGAAGAGA-CCCCGUGCACC-UUUACU--AUAGCUU-UACAUUGGUAUUAAAAAUGUGAUGUGCAG
AAU-AGGUGGGAGACUUUG----------AAGUUAU-GGC----------------------------GCUA
--------------------------------------GC-UAUAAUGGAG-U--C-AA-CCUUGAGAUA-C
CACC-CUUUACACUUUUGAUACCUAACUAUGU--UUCA-U-UA-U-C-------------------------
------------------------------------------------------------------------
------------------------------------------------------------------------
------------------------------------------------------------------------
------------------------------------------------------------------------
------------------------------------------------------UG----AA----------
------------------------------------------------ACUA-----GGACA-UUGUAUGGU
GGGUAGUUUGACUGGGGCGGU-CGCCUCCUA-AAGAGUAAC-GGAGGCGUGCG-AAGGUAAGCUAGAGCUGG
UCGGAAAUCAGCUUGA-UAGUA-UAAUGGCAU-AAGCUUGCCUGACU-GCGA--------------------
--------------GGCUGA---CAAGCCAAGCAGAGACGAAA-GUCGGUCAUAGUGAUCCGGU--------
---GA-U-----UCUGU-AUG--------------------GA-AGGGUCAUCGCUCAACGG-AUAAAAGGU
ACGCCGGGG-AUAACAGGCUGAUGGUGUUCAAGAGUUCAUAUCGACGACACCGUUUGGCACCUCGAUGUCGA
```

```
CUCAUCACAUCCUGGGGCUGAA-GAAGGUCCCAAGGGUUCGGCUGUUCGCCGAUUAAAGUGAUACGUGAGUU
GGGUUUAGAACGUCGUGAGACAGUUCGGUUUCUAUCUGCC--------------------G-UGG-------
-------------------------------GUGA--AGGAAAUUUGAGAAG-AUC-UGACUCUAGUACGAG
AGGAC-CGAGAUGGAUAUACCUC-UGGU-G-UACCAGCUGUUAUGCCAAUAGCAU-U-GCUGGGUAGCUAUG
UAUAGGUGGGAUAAUUGCUGAAAGCA-UA--UAAGCAAGAACCCUCUUCAA-A-A-----AGAUUUCC-CA-
--A---------------------------------------------------------------------
------------------------------------------------------------------------
------------------------------------------------------------------------
------------------------------------------------------------------------
------------------------------------------------------------------------
------------------------------------------------------------------------
------------------------------------------------------------------------
------------------------------------------------------------------------
----------------------------------------------------------UU-AAG------GC
CGUGGAAGACUACCACGU-UGA-UAGGCU-AGG-UGUGGAAG-CAU-G----------------GU------
----AA-----------CGUGUGGAGCUA--A-CUAG-UACUAAUA---GCCUGAUUG-AUUUAUUUG----
------------------------------------------------------------------------
------------------------------------------------------------------------
------------------------------------------------------------------------
------------------------------------U------------
>LSUb_227377_COXBU/1-2897
---------------------G-GUCAAGUG----AA--UAAGCGCAUAU-GGUGGAUG-C-CUUGGC-AGU
AAG---AGGCGAUGAAGGACGUAG--UAGCCUGCGAUAACUUCGGGGAGCUGGCAAACAAGCGA------UG
AUC-CGG-ACAUUUCCGAAUGGGGAAAC--------CC--------AA----CC----------------UU
U-C---------------------------G---AGG----U-UAUGUA-UAC-----UGA-----------
-----------A-----------UUCA-----------------------------------------U-AG
G-U--------------------AU-A-CGAGGCGAACCUGGGGAACUGAAACAUCUAAGUACCCA--GAGG
AAAAGAAAUCAA--CCGAGAUUCCGUCA--GU-AG-CGGCGAGCGAAAG---A-A-CA-GCCCA--------
------------------------------------------------------------------------
-------GU-UAC-------UAAA-UC-----------------------A--UUA-----------UU---
U-------GUUCU-AGCAG------AAU-GU-UCUG------------------------------------
-------GAA-AG-UCAG--------CCAUAGCGGGUGAUAGCCCGGUA---------CCCGAAA----GAG
U------A--AAUAA-----------UGU----GGG--UA-AC------GAUGAGUAGGUCGGGACACGUGG
UAUCUUGACUGAA-CAUGGGGGG-ACC--AUCCUCCAAGGCUAAAUA--CUC-CUUAC-UGACCGAUAGC-G
AAC-C-AGUACCGUGAGGAAAGG-UGAAAA-AGAAC-CCCGG-CGAGGG-A-GUGAAAUAGAACCUGAAACC
G-UAUGCGUACAA-GCU-GU-AGGA---GCAU-U---UC---------------------------------
------------------------------------------------------------------------
------------------------------------------------------------------------
------------------------------------------------------------------------
------------------------------------------------------------------------
------------------------------------------------------------------------
------------------------------------------------------------------------
------------------------------------------------------------------------
------------------------------------------------------------------------
--------------------------------------------------------------U--U-CG---
```

```
------------------------------------------------------------------------
------------------------------------------------------------------------
------------------------------------------------------------------------
------------------------------------------------------------------------
------------------------------------------------------------------------
------------------------------------------------------------------------
------------------------------------------------------------------------
------------------------------------------------------------------------
------------------------------------------------------------------------
------------------------------------------------------------------------
------------------------------------------------------------------------
------------------------------------------------------------------------
----------------------------------------------GAAA--U---GUGACUGCGUACCUUU
UGUAUAAUGGGUCAGCGACUUAC-U-UGUUGUAGCGA-GCUUAACC-------------GUCUA--------
---G--GGGA--GGCGUAGGGAAAC-C---------------------------------------------
------------------------------------------------------------------------
------------------------------------------------------------------------
---------------------------------------------------------------------GAG
UCC------------------------------------GAAAU-------------------------GG-
G-C-GU----------------------UUAGUUGCAACGAGUA-GACCGAACCCG-AAGCGAUCUAUCUAU
GGCCAGGGUGAAGGUCAG-GUAACA-CUGACUGGAGGCC--G--AACCACUA-AUGUGAAAAAGUUA-GGGG
AUGAGCUGUGGAUAGGAGUGAAAGGCUAAUCAAGCUCGG-AGAUAGCUGGUUCUCCUCGAAA-GCUAUUUAG
GUAGCGCCUCGU-GUA--------------------------------------------------------
-----------------------U-----GA-CUCUUGGGGGUAGAGC-ACUGUUU-CGG-CUA--GGGGGC
C---AUCCC--GGCCUACCAAACCGAGGCAAACUCCGAAUACCAAG--------------------------
------------------------------------------------------------------------
------------------AAGU-UUUA--------GCACGGG-AGACACA-C-UGCU-GGGUGAUAAGGUCC
GU-G-GUGGAAAG-GAACGCCAGAUCGCCAGAUCGCA-GCUAAGGUCCCAAAUA--CA-CGU-----A---G
-U-GG--------AAACGAUGUGGGAAGGCUCAGACAGCCAGGACCUUGGCUUAGAAGCAGCCA-UCCUUUA
AAGAAAGCCUAAUAGCUCACUGGUC-----GAGUCGUCCUGCGCGGAAGAUUUAAC-GGGGCUCA-AACUGU
G----------UACCGAA-GCUGCGG----------------------------------------------
------------------------------------------------------------------------
------------------------------------------------------------------------
--------------------------------------------GGUAGAGGAGC-GUUCUGU---------
-----------A-A-CCUGU-GAAGGUG----AUCGAGAGGU--U-UGCUGGAGGU-A-UCA-GAAGUGCGA
AUGC-UGACAUAAAGUA-ACGAUA---AUGU-GGGUGA--A---------------AAACCC-ACA-CGCCG
A-AAGUCU-AAGGUUUCC-UGCGC--CG-UUAAUC--CGCAGGGUGAGUCGGC-CC-UAAGGGAGGA---G-
A-A-A-UC-----GUA--------------------------------------------------------
----------------------------------------GUC-GA-U--GGGA-AACGGGUU-AAUA-UUC
CCGUA----------------------------------CUUUA----UAAU--------A-CU--------
------------------------------------------------------------------------
--------------------------------------------------------GCGAUGG-GGGGGGAC
GGAGAA--------------GGCUAG-GUCAGCCCC-CC-G----AUGGU------------UGUGGUGGG-
CUAGUC-AGC-CACCGAUGUG--------------------UG-----UGUGUUAAGUGAUGUGUAGAAA-GG
A-UCUU--------------------CAAAU-CCGGAACUCA---------AUUCCG----AGA-CA-U-G-
```

AUG--AC---------------------------------------------------------------GA
-GUACGAACUU-GUUCGUGCAAAGU-G-AUU-G-AUGCCACGACUUC-C-AGGAAA-AGUCC-CUAAGCUU-
-------------------------------------------------------CAGGUAU-UG-UAAAAC
CGUACUAUAAACCGAC------------------------------ACAGGUG--GACAGGUA-GAGAAUAC
CAAGGC-G-CU-UGAG-AGAACUUG-G-GUGAAGGAACUAGGC-AAAAUGGCACCGUAACUUCGG-GAGAAG
GUGCGCCCUU-G-GU--AAGUGAAG----GUC-----------CUUGC------------------------
------------------------------------------------------------------------
------------------------------------------------------------------------
------------------------------------------------------------------------
------------------------------------------------------------------------
------------------------------------------------------------------------
------------------------------------------------------------------------
------------------------------------------------------------------------
------------------------------------------------------------------------
------------------------------------------------------------------------
------------------------------------------------------------------------
------------------------------------------------------------------------
------------------------------------------------------------------------
------------------------------------------------------------------------
--------------------------------------------------------GACU--GA--GCU--G
AAA--GGUUGCAGAUACCAGGUGGCUGCGAAAGACUAGUUUACUAAAAACACAGCACUCUGCAAACUCG---
-UA-A-------------GAGAGCGUAAUAGGGUGUGAGCGCCUGCCCGGUGCCG-GAAAGG-UUAAGUGAU
GGGGUUA-GCC----CUCG----GGCGAAGCUCUUGAUCGAAGCCC--CGGUAAACGGGCGGCCGUAACUAU
AACGGUCCUAAGGUAGCGAAAUUCCUUGUCAGU-GAAGUUCCGACCUGCACGAAUGGCGUAACGAUA--GCC
ACGCUGUCUCCACC----CAA-GACUCAGUGAAAUU--GAA---AUCGCUGUGAAGAUGCAGCGUACCCGCG
GCUAGACGGAAAGA-CCCCGUGAACC-UUUACU--ACAGCUU-UACACUGAACUUUGAAUAUGCCUGUGUAG
GAU-AGGUGGGAGGCUUUG----------AAGCUGG-AGC----------------------------GCUA
---------------------------------------GC-UCCAGUGGAG-C--C-AA-CCUUGAAAUA-C
CACC-CUGGCGUGUUUGAGGUUCUAACCCUGU--CCCG-U-AA-U-C-------------------------
------------------------------------------------------------------------
------------------------------------------------------------------------
------------------------------------------------------------------------
------------------------------------------------------------------------
------------------------------------------------------CG----GG----------
------------------------------------------------UCGG-----GAACA-GUGUAUGGU
GGGUAGUUUGACUGGGGCGGU-CUCCUCCUA-AAGAGUAAC-GGAGGAGUACA-AAGGUACCCUCAGCACGG
UCGGAAAUCGUGCAUU-GUGUG-CAAAGGCAU-AAG-GUGCUUGACU-GCGA--------------------
--------------GACGUA---CAAGUCGAGCAGGUACGAAA-GUAGGUCUUAGUGAUCCGGU--------
---GG-U-----CCUUU-AUG--------------------GA-AGGGC--ACGCUCAACGG-AUAAAAGGU
ACUCCGGGG-AUAACAGGCUGAUACCACCCAAGAGUUCAUAUCGACGGUGGUGUUUGGCACCUCGAUGUCGG
CUCAUCACAUCCUGGGGCUGUA-GCCGGUCCCAAGGGUAUGGCUGUUCGCCAUUUAAAGUGGUACGUGAGCU
GGGUUUAGAACGUCGUGAGACAGUUCGGUCCCUAUCUGCC--------------------GUUGG-------
-------------------------------GCGU--UGGAGAUUUGAGAGG-AGC-UGCUCCUAGUACGAG
AGGAC-CGGAGUGGACGUACCUC-UGGU-G-UUCCGGUUGUCACGCCAGUGGCAU-UUGCCGGGUAGCUAAG
UACGGACGGGAUAACCGCUGAAAGCA-UC--UAAGCGGGAAACCC-CCUCAA-G-AUG--AGAUCUCC-GG-

```
--ACCUU-----------------------------------------------------------------
--------------------------U-----------A---------------------------------
------------------------------------------------------------------------
------------------------------------------------------------------------
------------------------------------------------------------------------
------------------------------------------------------------------------
------------------------------------------------------------------------
------------------------------------------------------------------------
-------------------------------------------------AGGUCCU--AA-AGA------UU
CGUUGAAGACGACAACGU-UGA-UAGGCA-GGG-UGUGGAAGUCUC-A----------------GU------
----AA-----------UGAGUGAAGCUA--A-CCUG-UACUAAUU---AAUCGUGCG-ACUUGACUA----
------------------------------------------------------------------------
------------------------------------------------------------------------
------------------------------------------------------------------------
------------------------------------U------------
>LSUb_198094_BACAN/1-2908
------------------------------------------GGGCGCAC-GGUGGAUG-C-CUUGAC-ACU
AGG---AGUCGAUGAAGGACGGGA--CUAACGCCGAUAUGCUUCGGGGAGCUGUAAGUAAGCUU------UG
AUC-CGA-AGAUUUCCGAAUGGGGAAAC--------CC--------AC----CAUA-C-G----------UA
A-U-------------G---G----U----A---UGG----U-A-UCCU-UAU-----CUGA----------
-----------A-----------UACA----------------------------------------UA-GG
G-U--------------------AA-G-GAAGACAGACCCAGGGAACUGAAACAUCUAAGUACCUG--GAGG
AAGAGAAAGCAA--AUGCGAUUUCCUGA--GU-AG-CGGCGAGCGAAACGGAACA-UA-GCCCAAACCAAG-
--AG-GCUUG----------------------------------CCU-CUUGGGG--UUG------------
-----UAGG-ACA-------UUCUAUA----------------------CG--GAGUUACAAA--GGAA---
U-------GAGGU-AGACG------AAGCGA-CCUG------------------------------------
-------GAA-AG-GUCCG-------UCGUAGAGGGUAACAACCCCGUA---------GUCGAAA----CUU
C------G--UUCUC----------UCUU---GAAU--GU-AU------CCUGAGUACGGCGGAACACGUGA
AAUUCCGUCGGAA-UCUGGGAGG-ACC--AUCUCCCAAGGCUAAAUA--CUC-CCUAG-UGAUCGAUAGU-G
AAC-C-AGUACCGUGAGGGAAAG-GUGAAA-AGCAC-CCCGG-AAGGGG-A-GUGAAAGAGAUCCUGAAACC
G-UGUGCCUACAA-AUA-GU-CAGA---GCCC-G--------------------------------------
------------------------------------------------------------------------
------------------------------------------------------------------------
------------------------------------------------------------------------
------------------------------------------------------------------------
------------------------------------------------------------------------
------------------------------------------------------------------------
------------------------------------------------------------------------
------------------------------------------------------------------------
--------------------------------------------------------------U--U-AA---
------------------------------------------------------------------------
------------------------------------------------------------------------
------------------------------------------------------------------------
------------------------------------------------------------------------
------------------------------------------------------------------------
```

```
------------------------------------------------------------------------
------------------------------------------------------------------------
------------------------------------------------------------------------
------------------------------------------------------------------------
------------------------------------------------------------------------
------------------------------------------------------------------------
------------------------------------------------------------------------
------------------------------------------------CG--G---GUGAUGGCGUGCCUUU
UGUAGAAUGAACCGGCGAGUUAC-G-AUCCCGUGCGA-GGUUAAGC--U----------GAAGA--------
--GG--CGGA--GCCGCAGCGAAAG-C---------------------------------------------
------------------------------------------------------------------------
------------------------------------------------------------------------
---------------------------------------------------------------------GAG
UCU------------------------------------GAAUA-------------------------GG-
G-C-GU----------------------UUAGUACGUGGUCGUA-GACCCGAAACC-AGGUGAUCUACCCAU
GUCCAGGGUGAAGUUCAG-GUAACA-CUGAAUGGAGGCCCGA--ACCCACGC-AUGUUGAAAAGUGC-GGGG
AUGAGGUGUGGGUAGCGGAGAAAUUCCAAUCGAACCUGG-AGAUAGCUGGUUCUCCCCGAAA-UAGCUUUAG
GGCUAGCCUUAA-GUGU-------------------------------------------------------
----------------------AA-----GA-GUCUUGGAGGUAGAGC-ACUGAUU-GGA-CUA--GGGGUC
C---UCAUC--GGAUUACCGAAUUCAGUCAAACUCCGAAUGCCAAU--------------------------
------------------------------------------------------------------------
------------------GACU-U-AU---------CCUUAGG-AGUCAGA-C-UGCG-A-GUGAUAAGAUCC
GU-A-GUCAAAAG-GGAAAC---AGCCCAGACCGCCA-GCUAAGGUCCCAAAGU--GU-GUA-UU-AA---G
-U-GGA-------AAAGGAUGUGGAGUUGCUUAGACAACUAGGAUGUUGGCUUAGAAGCAGCCA-CCAUUUA
AAGAGUGCGUAAUAGCUCACUAGUC-----GAGUGACUCUGCGCCGAAAAUGUACC-GGGGCU-A-AAUACA
C----------CACCGAA-GCUGCGGAUUGAUAC--CA----------------------------------
------------------------------------------------------------------------
------A-----------------------------------------------------------------
--------------------------------UGGUA-UCAG-UGGUAGGGGAGC-GUUCUAA---------
-----------G-G-ACAGU-GAAGUCA--G-ACCGGAAGGA--C-UGGUGGAGUG-C-UUA-GAAGUGAGA
AUGC-CGGUAU-GAGUA-GCGAAA----GAC-GGGUGA--G---------------AAUCCC-GUC-CACCG
A-AUGCCU-AAGGUUCC-UGAGGAAGG-CUCGUCCGCUCAGGGUUAGUCAGG-AC-CUAAGCCGAGGCCG-
A-C-AGGC-----GUA--------------------------------------------------------
----------------------------------------GGC-GA-U--GGAC-AACAGGUU-GAUA-UUC
CUGUA----------------------------------CCACC----UCUU--------U-AUC-------
------------------GUUUGA------------------------------------------------
----------------------------------------------------------GCAAU-GGAGGGCC
GCAGAA--------------GGAUAG-AAGAAGCGU-GCGA----UUGGU------------UGUGCACGU-
CCAAGC-AGU-UA-----G-G-------------------C------UG------------------AU-AA
G-UAGG--------------------CAAAU-CCGCUUAUCG------UGAAGGCUG----AGC-UG-U-G-
AUG--GG---------------------------------------------------------------GA
-----AGCUCC-UUAUGGAGCGAAG-U-CUU-U-GAUUCCCCGCUGC-C-AAGAAA-AGCUU-CUAGCG---
--------------------------------------------------------AGAUAA-AA-GGUGCC
UGUACCGCAAACCGAC------------------------------ACAGGUA--GGCGAGGA-GAGAAUCC
UAAGGU-G-UG-CGAG-AGAACUCU-G-GUUAAGGAACUCGGC-AAAAUGACCCCGUAACUUCGG-GAGAAG
```

```
GGGUG-CUUU-C-UU---------------------------------------------------------
------------------------------------------------------------------------
------------------------------------------------------------------------
------------------------------------------------------------------------
------------------------------------------------------------------------
------------------------------------------------------------------------
------------------------------------------------------------------------
------------------------------------------------------------------------
------------------------------------------------------------------------
------------------------------------------------------------------------
------------------------------------------------------------------------
------------------------------------------------------------------------
------------------------------------------------------------------------
------------------------------------------------------------------------
--------------------------------------------------------------------AACG
GAA--AGCCGCAGUGAAUAGGCCCAAGCGA-----CUGUUUAGCAAAAACACAGCUCUCUGCGAAGCCG---
-UA-A-------------GGCGAAGU-AUAGGGGGUGA-CACCUGCCCGGUGCUG-G-AAGG-UUAAGGAGA
GGGGUUA-GC-----GUAA-----GCGAAGCUCUGAACUGAAGCCC--CAGUAAAC-GGCGGCCGUAACUAU
AACGGUCCUAAGGUAGCGAAAUUCCUUGUCGGG-UAAGUUCCGACCCGCACGAAAGGUGUAACGAUU--UGG
GCACUGUCUCAACC----AGA-GACUCGGUGAAAUU--AUA---GUACCUGUGAAGAUGCAGGUUACCCGCG
ACAGGACGGAAAGA-CCCCGUGGAGC-UUUACU--GUAGCCU-GAUAUUGAAUUUUGGUACAGUUUGUACAG
GAU-AGGCGGGAGCCUUUG----------AAACCGG-AGC----------------------------GCUA
--------------------------------------GC-UUCGGUGGAG-G--C-GC-UGGUGGGAUA-C
CGCC-CUGACUGUAUUGAAAUUCUAACCUACG--GGUC-U-UA-U-C-------------------------
------------------------------------------------------------------------
------------------------------------------------------------------------
------------------------------------------------------------------------
------------------------------------------------------------------------
------------------------------------------------------GA----CC----------
------------------------------------------------CGGG-----AGACA-GUGUCAGGU
GGGCAGUUUGACUGGGGCGGU-CGCCUCCUA-AAGUGUAAC-GGAGGCGCCCA-AAGGUUCCCUCAGAAUGG
UUGGAAAUCAUUCGUA-GAGUG-CAAAGGCAU-AAGGGAGCUUGACU-GCGA--------------------
--------------GAACUA---CAAGUCGAGCAGGGACGAAA-GUCGGGCUUAGUGAUCCGGU--------
---GG-U-----UCCGC-AUG--------------------GA-AGGGCCAUCGCUCAACGG-AUAAAAGCU
ACCCCGGGG-AUAACAGGCUUAUCUCCCCCAAGAGUCCACAUCGACGGGGAGGUUUGGCACCUCGAUGUCGG
CUCAUCGCAUCCUGGGGCUGUA-GUCGGUCCCAAGGGUUGGGCUGUUCGCCCAUUAAAGCGGUACGCGAGCU
GGGUUCAGAACGUCGUGAGACAGUUCGGUCCCUAUCCGUC--------------------G-UGG-------
-------------------------------GCGU--AGGAAAUUUGAGAGG-AGC-UGUCCUUAGUACGAG
AGGAC-CGGGAUGGACGCACCGC-UGGU-G-UACCAGUUGUUCUGCCAAGGGCAU-A-GCUGGGUAGCUAUG
UGCGGAAGGGAUAAGUGCUGAAAGCA-UC--UAAGCAUGAAGCCCCCCUCAA-G-AUG--AGAUUUCC-CA-
--UAGCG-----------------------------------------------------------------
--------------------------U---------------------------------------------
------------------------------------------------------------------------
------------------------------------------------------------------------
------------------------------------------------------------------------
```

```
------------------------------------------------------------------------
------------------------------------------------------------------------
------------------------------------------------------------------------
-------------------------------------------------AAGCUAG--UA-AGA------UC
CCUGAAAGAUGAUCAGGU-UGA-UAGGUU-CGA-GGUGGAAG-CAU-G----------------GU------
----GA-----------CAUGUGGAGCUG--A-CGAA-UACUAAUA---GAUCGAG-G-ACUUAACCA----
------------------------------------------------------------------------
------------------------------------------------------------------------
------------------------------------------------------------------------
------------------------------------U------------
>LSUb_235909_GEOKA/1-2930
---------------------G-GUUAAGCU----GG--AAAGGGCGCAC-GGUGGAUG-C-CUUGGC-ACU
AGG---AGCCGAUGAAGGACGGGG--CAAACGCCGAAACGCUUCGGGGAGCUGUAAGCAAGCGU------UG
AUC-CGG-AGAUGUCCGAAUGGGGAAC---------CC--------AC----UGUC-CGU-----------A
A-U-------------G---G----G----G---CAG----U-A-UCCA-UGC-----CUGA----------
-----------A-----------UCCA----------------------------------------UA-GG
G-C--------------------AU-G-GAGGGCACACCCGGGGAACUGAAACAUCUUAGUACCCG--GAGG
AGAAGAAAGCAA--CCGCGAUUCCCUGA--GU-AG-CGGCGAGCGAAACGGGA-A-CA-GCCCAAACCAAG-
--AG-GCAUG--------------------------------U-CCU-CUUGGGG--UUG------------
-----UAGG-ACC-------GCUCACGA--------------------UGG--GAGUGAGAAA--GGGA---
C-------GGGGU-AGACG------AACCGG-UCUG------------------------------------
-------GAA-CG-GCCGG-------CCAGAGAAGGUGAGAGCCCUGUA---------GUCGAAA----CUU
C-----G--UUCCC-----------UCCC---GAGC--GG-AU------CCUGAGUACGGCGGGACACGAGG
AAUCCCGUCGGAA-GCAGGGAGG-ACC--AUCUCCCAAGGCUAAAUA--CUC-CCUAG-UGACCGAUAGU-G
CAC-C-AGUACCGUGAGGGAAAG-GUGAAA-AGCAC-CCCGG-GAGGGG-A-GUGAAAGAGAACCUGAAACC
G-UGUGCCUACAA-GUA-GU-CAGA---GCCC-G--------------------------------------
------------------------------------------------------------------------
------------------------------------------------------------------------
------------------------------------------------------------------------
------------------------------------------------------------------------
------------------------------------------------------------------------
------------------------------------------------------------------------
------------------------------------------------------------------------
------------------------------------------------------------------------
--------------------------------------------------------------U--U-GA---
------------------------------------------------------------------------
------------------------------------------------------------------------
------------------------------------------------------------------------
------------------------------------------------------------------------
------------------------------------------------------------------------
------------------------------------------------------------------------
------------------------------------------------------------------------
------------------------------------------------------------------------
------------------------------------------------------------------------
------------------------------------------------------------------------
```

```
------------------------------------------------------------------------
------------------------------------------------------------------------
------------------------------------------------UG--G---GUGAUGGCGUGCCUUU
UGUAGAAUGAACCGGCGAGUGAC-G-AUGGCGUGCGA-GGUUAAGC--C----------GAAGA--------
--GG--CGGA--GCCGCAGCGAAAG-C---------------------------------------------
------------------------------------------------------------------------
------------------------------------------------------------------------
---------------------------------------------------------------------GAG
UCU------------------------------------GAACA-------------------------GG-
G-C-GU---------------------GUGAGUACGUCGUCGUC-GACCCGAAACC-AGGUGAUCUACCCAU
GUCCAGGGUGAAGGCCGG-GUAACA-CCGGCUGGAGGCCCGA--ACCCACGC-ACGUUGAAAAGUGC-GGGG
AUGAGGUGUGGGUAGGGGUGAAAUGCCAAUCGAACUUGG-AGAUAGCUGGUUCUCCCCGAAA-UAGCUUUAG
GGCUAGCCUCGG-GUUU-------------------------------------------------------
----------------------GA-----GA-GUCUUGGAGGUAGAGC-ACUGAUU-GGG-CUA--GGGGCC
C---CAAAC--GGGUUACCGAACCCAGUCAAACUCCGAAUGCCAAU--------------------------
------------------------------------------------------------------------
------------------GACU-U-AU--------GCCCGGG-AGUCAGA-C-CGCG-A-GUGAUAAGAUCC
GU-G-GUCGAGAG-GGGAAC---AGCCCAGACCGCCA-GCUAAGGCCCCGAAGU--GC-ACG-UU-CA---G
-U-GGA-------AAAGGAUGUGGAGUUGCCGAGACAACCAGGAUGUUGGCUUAGAAGCAGCCA-CCAUUUA
AAGAGUGCGUAAUAGCUCACUGGUC-----GAGUGACUCUGCGCCGAAAAUGUACC-GGGGCU-A-AACGUG
C----------CGCCGAA-GCUGCGGGAUGACCG--U-----------------------------------
------------------------------------------------------------------------
------------------------------------------------------------------------
----------------------------------UGG-UCAU-CGGUAGGGGAGC-GUUCUAA---------
-----------G-G-GCAGA-GAAGCCA--G-ACCGGAAGGA--C-UGGUGGAGCG-C-UUA-GAAGUGAGA
AUGC-CGGUAU-GAGUA-GCGAAA---ACAG-AGGUGA--G---------------AAUCCU-CUG-CGCCG
A-AAGCCU-AAGGGUUCC-UGAGGAAGG-UUCGUCCGCUCAGGGUUAGUCGGG-AC-CUAAGCCGAGGCCG-
A-A-AGGC-----GUA--------------------------------------------------------
----------------------------------------GGU-GA-U--GGAC-AACAGGUU-GAGA-UUC
CUGUA----------------------------------CCACC----UUCU--------U-CCC-------
------------------GUUUGA------------------------------------------------
----------------------------------------------------------GCGAU-GGGGGGAC
GCAGGA--------------GGAUAG-GGCGAGCAG-GCGG----CUGGA------------AGAGCCUGU-
CCAAGC-CGU-GA------G-G-------------------C------UG------------------AU-CC
G-CAGG--------------------CAAAU-CCGCGGAUCA------U-AAGGCCA----AGC-GG-U-G-
ACG--GC---------------------------------------------------------------GA
-----CGGA----UUGAUCCGGAAG-U-CCC-C-GAUUUCACACUGC-C-AAGAAA-AGCCU-CUAGCG---
--------------------------------------------------------AGGGAA-GA-GGUGCC
CGUACCGCAAACCGAC------------------------------ACAGGUA--GGCGAGGA-GAGAAUCC
UAAGGC-G-CG-CGGG-AGAACUCU-C-GUUAAGGAACUCGGC-AAAAUGACCCCGUAACUUCGG-GAGAAG
GGGUG-CUCU-U-UG--GGGUGA-------------------------------------------------
------------------------------------------------------------------------
------------------------------------------------------------------------
------------------------------------------------------------------------
------------------------------------------------------------------------
```

```
------------------------------------------------------------------------
------------------------------------------------------------------------
------------------------------------------------------------------------
------------------------------------------------------------------------
------------------------------------------------------------------------
------------------------------------------------------------------------
------------------------------------------------------------------------
------------------------------------------------------------------------
------------------------------------------------------------------------
-------------------------------------------------------------AGA--GCCCUG
AAG--AGCCGCAGUGAAAAGGCCCAAGCGA-----CUGUUUAUCAAAAACACAGGUCUCUGCGAAGCCG---
-AA-A-------------GGCGACGU-AUAGGGGCUGA-CACCUGCCCGGUGCUG-G-AAGG-UUAAGGGGA
GCGCUUA-GC-----GGAA-----GCGAAGGUGCGAACCGAAGCCC--CAGUAAAC-GGCGGCCGUAACUAU
AACGGUCCUAAGGUAGCGAAAUUCCUUGUCGGG-UAAGUUCCGACCCGCACGAAAGGUGUAACGACU--UGG
GCGCUGUCUCAACG----AGA-GACCCGGUGAAAUU--AUA---CUACCUGUGAAGAUGCAGGUUACCCGCG
ACAGGACGGAAAGA-CCCCGUGGAGC-UUUACU--GCAGCCU-GAUAUGGAAUUUUGGUAUCGCUUGUACAG
GAU-AGGUGGGAGCCUGGG----------AAGCCGG-AGC----------------------------GCCA
--------------------------------------GC-UUCGGUGGAG-G--C-GA-CGGUGGGAUA-C
CACC-CUGGCGGUAUUGAAAUUCUAACCCGCA--CCCC-U-UA-G-C-------------------------
------------------------------------------------------------------------
------------------------------------------------------------------------
------------------------------------------------------------------------
------------------------------------------------------------------------
------------------------------------------------------GG----GG----------
------------------------------------------------UGGG-----AGACA-GUGUCAGGU
GGGCAGUUUGACUGGGGCGGU-CGCCUCCCA-AAAGGUAAC-GGAGGCGCCCA-AAGGUUCCCUCAGAAUGG
UUGGAAAUCAUUCGGA-GAGUG-CAAAGGCAC-AAGGGAGCUUGACU-GCGA--------------------
--------------GACGGA---CAGGUCGAGCAGGGACGAAA-GUCGGCUUAGUGAUCCGGU---------
---GG-U-----UCCGC-AUG--------------------GA-AGGGCCAUCGCUCAACGG-AUAAAAGCU
ACCCCGGGG-AUAACAGGCUGAUCUCCCCCAAGAGUCCACAUCGACGGGGAGGUUUGGCACCUCGAUGUCGG
CUCAUCGCAUCCUGGGGCUGUA-GUCGGUCCCAAGGGUUGGGCUGUUCGCCCAUUAAAGCGGUACGCGAGCU
GGGUUCAGAACGUCGUGAGACAGUUCGGUCCCUAUCCGUC--------------------G-CGG-------
-------------------------------GCGG--AGGAAAUUUGAGAGG-AGC-UGUCCUUAGUACGAG
AGGAC-CGGGAUGGACGCACCGC-UGGU-G-UACCAGUUGUCCCGCCAGGGGCAC-C-GCUGGGUAGCUAUG
UGCGGACGGGAUAAGCGCUGAAAGCA-UC--UAAGCGUGAAGCCCCCCUCAA-G-AUG--AGAUUUCC-CA-
--CCGCG-----------------------------------------------------------------
--------------------------CA--------------------------------------------
------------------------------------------------------------------------
------------------------------------------------------------------------
------------------------------------------------------------------------
------------------------------------------------------------------------
------------------------------------------------------------------------
------------------------------------------------------------------------
-------------------------------------------------AAGCGGG--UA-AGA------UC
CCUCGAAGAUGACGAGGU-CGA-UAGGUC-CGA-GGUGGAAG-CGU-G----------------GC------
```

```
----GA-----------CACGUGGAGCUG--A-CGGA-UACUAAUC---GAUCGAG-G-GCUUGACCU----
------------------------------------------------------------------------
------------------------------------------------------------------------
------------------------------------------------------------------------
------------------------------------A------------
>LSUb_221109_OCEIH/1-2922
---------------------A-GUUAAGUG----AA--UAAGGGCGCAC-GGUGGAUG-C-CUUGGC-ACU
AGG---AGCUGACGAAGGACGGGA--CUAACACCGAUAUGCCUCGGGGAGUUGUAAGUAAACUU------UG
AUC-CGG-GGAUUUCCGAAUGGGGAAAC--------CC--------AC----UGUU-CGU-----------A
A-U-------------G---G----A----G---CAG----U-A-CCUA-UAC-----CUCA----------
-----------A-----------UACA----------------------------------------UA-GG
G-U--------------------AU-A-GGGAGCAGACCCGGGGAACUGAAACAUCUCAGUACCUG--GAGG
AAGAGAAAGAAA--CCUCGAUUUCCUGA--GU-AG-CGGCGAGCGAAACGGAA-A-GA-GCCCAAACCAGA-
--AA-GCUUG----------------------------------CUU-UCUGGGG--UUG------------
-----UAGG-ACA-------CUCC-UU----------------------UG--GAGUUACAAA--GAAA---
U-------UCUUU-AAAUG------AAUCGA-UCUG------------------------------------
-------GAA-AG-GUCAG-------CCAAAGAAGGUAACAGCCCUGUA---------AUUGAAA----AGG
A------A--UUUCC----------UCCG---GAGU--GU-AU------CCUGAGUACGGCGGAACACGUGG
AAUUCCGUCGGAA-UCCGGGAGG-ACC--AUCUCCCAAGGCUAAAUA--CUC-CCUAG-UGACCGAUAGU-G
AAC-C-AGUACCGUGAGGGAAAG-GUGAAA-AGCAC-CCCGG-AAGGGG-A-GUGAAAUAGAACCUGAAACC
G-UGUGCCUACAA-GUA-GU-CGAA---GCCC-G--------------------------------------
------------------------------------------------------------------------
------------------------------------------------------------------------
------------------------------------------------------------------------
------------------------------------------------------------------------
------------------------------------------------------------------------
------------------------------------------------------------------------
------------------------------------------------------------------------
------------------------------------------------------------------------
--------------------------------------------------------------U--U-AA---
------------------------------------------------------------------------
------------------------------------------------------------------------
------------------------------------------------------------------------
------------------------------------------------------------------------
------------------------------------------------------------------------
------------------------------------------------------------------------
------------------------------------------------------------------------
------------------------------------------------------------------------
------------------------------------------------------------------------
------------------------------------------------------------------------
------------------------------------------------------------------------
------------------------------------------------------------------------
------------------------------------------------UG--G---GUGACGGCGUACCUUU
UGUAGAAUGGACCGGCGAGUUAC-G-AUCCCAUGCAA-GGUUAAGU--G----------GAAGA--------
--CA--CGGA--GCCGCAGCGAAAG-C---------------------------------------------
```

```
------------------------------------------------------------------------
------------------------------------------------------------------------
---------------------------------------------------------------------GAG
UCU------------------------------------GAAUA-------------------------GG-
G-C-GA--------------------AUAUAGUAUGUGGUCGUA-GACCCGAAACC-GUGUGAUCUACCCAU
GUCCAGGGUGAAGGUCAG-GUAACA-CUGACUGGAGGCCCGA--ACCCACGU-AUGUUGAAAAAUGC-GGGG
AUGAGGUGUGGGUAGGGGUGAAAUGCCAAUCGAACACGG-AGAUAGCUGGUUCUCUCCGAAA-UAGCUUUAG
GGCUAGCCUCAA-GAUG-------------------------------------------------------
----------------------AU-----GA-GUACUGGAGGUAGAGC-ACUGAUU-GGA-CUA--GGGGCC
C---CUACC--GGGUUACCGAAUUCAGUCAAACUCCGAAUGCCAGU--------------------------
------------------------------------------------------------------------
------------------UACU-U-AA--------ACUUGGG-AGUCAGA-C-UAUG-G-GUGAUAAGGUUC
AU-A-GUCGAAAG-GGAAAC---AGCCCAGACCGCCA-GCUAAGGUCCCAAAGU--AU-ACG-UU-AA---G
-U-GGA-------AAAGGAUGUGGAGUUGCCCAGACAACCAGGAUGUUGGCUUAGAAGCAGCCA-CCAUUUA
AAGAGUGCGUAAUAGCUCACUGGUC-----GAGUGACUCUGCGCCGAAAAUGUACC-GGGGCU-A-AACGUA
U----------CACCGAA-GCUGCGGAUUGUUCU--U-----------------------------------
------------------------------------------------------------------------
------------------------------------------------------------------------
---------------------------------ACGA-ACAA-UGGUAGGAGAGC-GUUCUAA---------
-----------G-U-GCUGC-GAAGUCA--G-ACCGUGAGGA--C-UGGUGGAGCG-C-UUA-GAAGUGAGA
AUGC-CGGUAU-GAGUA-GCGAAA----AAA-GAGUGA--G---------------AAUCUC-UUU-CACCG
A-AUGCCU-AAGGUUUCC-UGAGGAAGG-CUCGUCCUCUCAGGGUUAGUCGGG-AC-CUAAGCCGAGGCCG-
A-U-AGGC-----GUA--------------------------------------------------------
----------------------------------------GGC-GA-U--GGAC-AACAGGUA-GAUA-UUC
CUGUA----------------------------------CCACC----UCAU--------G-AU--------
------------------UGUUUGA-----------------------------------------------
----------------------------------------------------------GCGAU-GGGGGGAC
GCAGUA--------------GGAUAA-GGAAUGCGC-ACCG----AUGGA------------CGUGUGCGC-
CCAAGC-AGU-GA-----G-A-------------------A------AG------------------UC-AG
A-UAGG--------------------CAAAU-CCGUCUGGC---------AAUUUCA----AGC-UG-U-G-
AUG--GG---------------------------------------------------------------GA
-----GGGAAA-UUGAGUACCGAAG-U-UCC-G-GAUUUCACACUGC-C-AAGAAA-AGCCU-CUAGUU---
--------------------------------------------------------AGAUCA-UA-GGUGCC
CGUACCGCAAACCGAC------------------------------ACAGGUA--GGCGAGGA-GAGAAUCC
UAAGGU-G-AG-CGGG-AGAACUCU-C-GUUAAGGAACUCGGC-AAAAUGACCCCGUAACUUCGG-GAGAAG
GGGUG-CUCC-U-UC---------------------------------------------------------
------------------------------------------------------------------------
------------------------------------------------------------------------
------------------------------------------------------------------------
------------------------------------------------------------------------
------------------------------------------------------------------------
------------------------------------------------------------------------
------------------------------------------------------------------------
------------------------------------------------------------------------
------------------------------------------------------------------------
```

```
------------------------------------------------------------------------
------------------------------------------------------------------------
------------------------------------------------------------------------
------------------------------------------------------------------------
-------------------------------------------------------------------AUAGG
AGG--AGCCGCAGUGAAUAGGCCCAAGCGA-----CUGUUUAGCAAAAACACAGGUCUCUGCGAAGCCG---
-AA-A-------------GGCGAAGU-AUAGGGGCUGA-CACCUGCCCGGUGCUG-G-AAGG-UUAAGGGGA
CGAGUUA-GCUC---UUUG---GAGCGAAGCUUUGAACCGAAGCCC--CAGUAAAC-GGCGGCCGUAACUAU
AACGGUCCUAAGGUAGCGAAAUUCCUUGUCGGG-UAAGUUCCGACCCGCACGAAAGGUGCAACGACU--UGG
GCACUGUCUCAACG----AGA-GACCCGGUGAAAUU--AUA---CUAUGCGUGAAGAUGCGCAUUACCCGCG
ACAGGACGGAAAGA-CCCCGUGGAGC-UUUACU--GUAGCCU-GAUAUUGAAUGUUUGUGCAGCUUGUACAG
GAU-AGGUGGGAGCCUUUG----------AAGCGUG-AGC----------------------------GCUA
--------------------------------------GC-UUACGUGGAG-G--C-AC-CCGUGGGAUA-C
CACC-CUCGCUGUAUGAACAUUCUAACCCAGG--ACCG-U-GA-U-C-------------------------
------------------------------------------------------------------------
------------------------------------------------------------------------
------------------------------------------------------------------------
------------------------------------------------------------------------
------------------------------------------------------CG----GU----------
------------------------------------------------UCGG-----AGACA-GUGUCAGGC
GGGCAGUUUGACUGGGGCGGU-CGCCUCCCA-AAGAGUAAC-GGAGGCGCCCA-AAGGUUCCCUCAGAAUGG
UUGGAAAUCAUUCGUA-GAGUG-UAAAGGCAG-AAGGGAGCUUGACU-GCGA--------------------
--------------GACCUA---CAAGUCGAGCAGGGACGAAA-GUCGGGCUUAGUGAUCCGGU--------
---GG-U-----UCCGC-AUG--------------------GA-AGGGCCAUCGCUCAACGG-AUAAAAGCU
ACCCCGGGG-AUAACAGGCUUAUCUCCCCCAAGAGUUCACAUCGACGGGGAGGUUUGGCACCUCGAUGUCGG
CUCAUCGCAUCCUGGGGCUGUA-GUCGGUCCCAAGGGUUGGGCUGUUCGCCCAUUAAAGCGGUACGCGAGCU
GGGUUCAGAACGUCGUGACAGUUCGGUCCCUAUCCGUC----------------------G-UGG-------
-------------------------------GCGU--UGGAAGUUUGAGAGG-AGC-UGUCCUUAGUACGAG
AGGAC-CGGGAUGGACACACCGC-UGGU-G-UACCAGUUGUUCCGCCAGGAGCAU-G-GCUGGGUAGCUACG
UGUGGUAAGGAUAAGUGCUGAAAGCA-UC--UAAGCAUGAAGCCCCCCUCAA-G-AUG--AAACUUCC-CA-
--UCACU-----------------------------------------------------------------
--------------------------UC--------------------------------------------
------------------------------------------------------------------------
------------------------------------------------------------------------
------------------------------------------------------------------------
------------------------------------------------------------------------
------------------------------------------------------------------------
------------------------------------------------------------------------
-------------------------------------------------GAGUGAG--UA-AGA------UC
CCUCAGAGACGAUGAGGU-UGA-UAGGUC-CGA-GGUGGAAG-CGU-G----------------GC------
----GA-----------CACGUGCAGCUG--A-CGGA-UACUAAUU---GAUCGAG-G-ACUUAUCUA----
------------------------------------------------------------------------
------------------------------------------------------------------------
------------------------------------------------------------------------
------------------------------------A------------
```

```
>LSUb_169963_LISMO/1-2931
---------------------G-GUUAAGUU----AG--AAAGGGCGCAC-GGUGGAUG-C-CUUGGC-ACU
AGG---AGCCGAAGAAGGACGGGA--CUAACACCGAUAUGCUUUGGGGAGCUGUACGUAAGCGU------UG
AUC-CAG-AGAUUUCCGAAUGGGGGAAC--------CC--------AC----UAUC-U-U----------UA
G-U-------------C---G----G----A---UAG----U-A-UCCU-UAC-----GUGA----------
-----------A-----------UACA----------------------------------------UA-GC
G-U--------------------GA-G-GAAGGCAGACCCAGGGAACUGAAACAUCUAAGUACCUG--GAGG
AAGAGAAAGAAA--AAUCGAUUUCCUGA--GU-AG-CGGCGAGCGAAACGGAA-A-GA-GCCCAAACCAAG-
--AA-GCUUG----------------------------------CUU-CUUGGGG--UUG------------
-----UAGG-ACA-------CUCUAUA----------------------CG--GAGUUACAAA--AGAA---
A-------GUUAU-AAAUG------AAGCGG-UCUG------------------------------------
-------GAA-AG-GCCCG-------CCAAAGACGGUAACAGCCCGGUA---------GUUGAAA----UAG
C------U--UUCCC----------UCCA---GAGU--GG-AU------CCUGAGUACGGCGGAACACGUGA
AAUUCCGUCGGAA-UCCGGGAGG-ACC--AUCUCCCAAGGCUAAAUA--CUC-CCUAG-UGACCGAUAGU-G
AAC-C-AGUACCGUGAGGGAAAG-GUGAAA-AGCAC-CCCGG-AAGGGG-A-GUGAAACAGUUCCUGAAACC
G-UGUGCCUACAA-GUA-GU-UAGA---GCCC-G--------------------------------------
------------------------------------------------------------------------
------------------------------------------------------------------------
------------------------------------------------------------------------
------------------------------------------------------------------------
------------------------------------------------------------------------
------------------------------------------------------------------------
------------------------------------------------------------------------
------------------------------------------------------------------------
--------------------------------------------------------------U--U-AA---
------------------------------------------------------------------------
------------------------------------------------------------------------
------------------------------------------------------------------------
------------------------------------------------------------------------
------------------------------------------------------------------------
------------------------------------------------------------------------
------------------------------------------------------------------------
------------------------------------------------------------------------
------------------------------------------------------------------------
------------------------------------------------------------------------
------------------------------------------------------------------------
------------------------------------------------------------------------
------------------------------------------------UG--G---GUGAUAGCGUGCCUUU
UGUAGAAUGAACCGGCGAGUUAC-G-AUUUGUUGCAA-GGUUAAGC--G----------GAAAA--------
--AG--CGGA--GCCGUAGCGAAAG-C---------------------------------------------
------------------------------------------------------------------------
------------------------------------------------------------------------
---------------------------------------------------------------------GAG
UCU------------------------------------GAAUA-------------------------GG-
G-C-GC---------------------AUAAGUAACAGGUCGUA-GACCCGAAACC-AGGUGAUCUACCCAU
```

```
GUCCAGGAUGAAGGUAAG-GUAAUA-CUUACUGGAGGUCCGA--ACCCACGC-ACGUUGAAAAGUGC-GGGG
AUGAGGUGUGGGUAGCGGAGAAAUUCCAAUCGAACUUGG-AGAUAGCUGGUUCUCUCCGAAA-UAGCUUUAG
GGCUAGCCUCGA-GGUA-------------------------------------------------------
----------------------AA-----GA-GUCAUGGAGGUAGAGC-ACUGUUU-GGA-CUA--GGGGCC
C---UUCUC--GGGUUACCGAAUUCAGAUAAACUCCGAAUGCCAUG--------------------------
------------------------------------------------------------------------
------------------UACU-U-AU--------ACUCGGG-AGUCAGA-C-UGCG-A-GUGAUAAGAUCC
GU-A-GUCGAAAG-GGAAAC---AGCCCAGACCACCA-GUUAAGGUCCCCAAAU--AU-AUG-UU-AA---G
-U-GGA-------AAAGGAUGUGGGGUUGCUUAGACAACCAGGAUGUUGGCUUAGAAGCAGCCA-CCAUUGA
AAGAGUGCGUAAUAGCUCACUGGUC-----GAGUGACCCCGCGCCGAAAAUGUACC-GGGGCU-A-AACAUA
U----------UACCGAA-ACUGUGGAUGAACCU--C-----------------------------------
------------------------------------------------------------------------
-----UU-----------------------------------------------------------------
-------------------------------UAGAGG-UUCG-UGGUAGGAGAGC-GUUCUAA---------
-----------G-G-GCGGU-GAAGUCA--G-ACCGGAAGGA--C-UGGUGGAGCG-C-UUA-GAAGUGAGA
AUGC-CGGUAU-GAGUA-GCGAAA----GAA-GGGUGA--G---------------AAUCCC-UUC-CACCG
A-AUAUCU-AAGGUUUCC-UGAGGAAGG-CUCGUCCGCUCAGGGUUAGUCGGG-AC-CUAAGCCGAGGCCG-
A-U-AGGC-----GUA--------------------------------------------------------
----------------------------------------GGC-GA-U--GGAC-AACAGGUA-GAGA-UUC
CUGUA----------------------------------CCAGU----GCUA--------A-UU--------
------------------GUUUAA------------------------------------------------
----------------------------------------------------------CCGAU-GGGGUGAC
ACAGAA--------------GGAUAG-GGAAUCGCA-CG-A----AUGGA------------AAUGUGCGU-
CCAAGC-AGU-GA-----G-U-------------------G------UG------------------AG-AA
G-UAGG--------------------CAAAU-CCGCUUCUCG------CGAAGCAUG----AGC-UG-U-G-
AUG--GG---------------------------------------------------------------GA
-----AGGAAA-UUAAGUACGGAAG-U-UCC-U-GAUUUCACGCUGU-C-AAGAAA-AGCCU-CUAGGA---
--------------------------------------------------------AGAGUA-GU-ACUGCC
CGUACCGCAAACCGAC------------------------------ACAGGUA--GAUGAGGA-GAGAAUCC
UAAGGU-G-AG-CGAG-AGAACUCU-C-GUUAAGGAACUCGGC-AAAAUGACCCCGUAACUUCGG-GAGAAG
GGGUG-CUCUAU-UA--GGGUG--------------------------------------------------
------------------------------------------------------------------------
------------------------------------------------------------------------
------------------------------------------------------------------------
------------------------------------------------------------------------
------------------------------------------------------------------------
------------------------------------------------------------------------
------------------------------------------------------------------------
------------------------------------------------------------------------
------------------------------------------------------------------------
------------------------------------------------------------------------
------------------------------------------------------------------------
------------------------------------------------------------------------
------------------------------------------------------------------------
-------------------------------------------------------------CAA--GCCCGA
```

GAG--AGCCGCAGUGAAUAGGCCCAGGCGA-----CUGUUUAGCAAAAACACAGGUCUCUGCAAAACCG---
-UA-A-------------GGUGACGU-AUAGGGGCUGA-CGCCUGCCCGGUGCUG-G-AAGG-UUAAGAGGA
GUGCUUA-GC-----UUCG-----GCGAAGGUACGAAUUGAAGCCC--CAGUAAAC-GGCGGCCGUAACUAU
AACGGUCCUAAGGUAGCGAAAUUCCUUGUCGGG-UAAGUUCCGACCCGCACGAAAGGCGCAACGAUC--UGG
GCACUGUCUCAACG----AGA-GACUCGGUGAAAUU--AUA---GUACCUGUGAAGAUGCAGGUUACCCGCG
ACAGGACGGAAAGA-CCCCGUGGAGC-UUUACU--GCAACCU-GAUAUGGAAUGUUUGUACCGCUUGUACAG
GAU-AGGUAGGAGCCGAAG----------AGACGUG-UGC----------------------------GCUA
--------------------------------------GC-AUACGAGGAG-G--C-AA-UGGUGGGAUA-C
UACC-CUGGCUGUAUGACCAUUCUAACCCGCC--ACGC-U-UA-G-C-------------------------
------------------------------------------------------------------------
------------------------------------------------------------------------
------------------------------------------------------------------------
------------------------------------------------------------------------
------------------------------------------------------GC----GU----------
------------------------------------------------GGGG-----AGACA-GUGUCAGGU
GGGCAGUUUGACUGGGGCGGU-CGCCUCCUA-AAGAGUAAC-GGAGGCGCCCA-AAGGUUCCCUCAGAAUGG
AUGGAAAUCAUUCGCA-GAGUG-UAAAGGCAC-AAGGGAGCUUGACU-GCGA--------------------
--------------GACUGA---CAAGUCGAGCAGGGACGAAA-GUCGGGCUUAGUGAUCCGGU--------
---GG-U-----UCCGC-AUG--------------------GA-AGGGCCAUCGCUCAACGG-AUAAAAGCU
ACCCCGGGG-AUAACAGGCUUAUCUCCCCCAAGAGUCCACAUCGACGGGGAGGUUUGGCACCUCGAUGUCGG
CUCGUCGCAUCCUGGGGCUGUA-GUCGGUCCCAAGGGUUGGGCUGUUCGCCCAUUAAAGCGGCACGCGAGCU
GGGUUCAGAACGUCGUGAGACAGUUCGGUCCCUAUCCGUC--------------------G-CGG-------
-------------------------------GCGC--AGGAAAUUUGAGAGG-AGC-UGUCCUUAGUACGAG
AGGAC-CGGGAUGGACACACCGC-UGGU-G-UACCAGUUGUUCCGCCAGGAGCAU-C-GCUGGGUAGCUAUG
UGUGGCAGGGAUAAACGCUGAAAGCA-UC--UAAGCGUGAAGCCCCCCUCAA-G-AUG--AGAUUUCC-CA-
--UUUCU-----------------------------------------------------------------
--------------------------UC--------------------------------------------
------------------------------------------------------------------------
------------------------------------------------------------------------
------------------------------------------------------------------------
------------------------------------------------------------------------
------------------------------------------------------------------------
------------------------------------------------------------------------
--------------------------------------------------GGAAAG--UA-AGA------UC
CCUGAAAGAUGAUCAGGU-AGA-UAGGUU-UGG-AGUGGAAG-UGU-A----------------GC------
----GA-----------UACAUGGAGCGG--A-CAAA-UACUAAUC---GAUCGAG-G-ACUUAACCA----
------------------------------------------------------------------------
------------------------------------------------------------------------
------------------------------------------------------------------------
-------------------------------------------------
>LSUb_282458_STAAR/1-2925
---------------------G-AUUAAGUU----AU--UAAGGGCGCAC-GGUGGAUG-C-CUUGGC-ACU
AGA---AGCCGAUGAAGGACGUUA--CUAACGACGAUAUGCUUUGGGGAGCUGUAAGUAAGCUU------UG
AUC-CAG-AGAUUUCCGAAUGGGGAAAC--------CC--------AG----CAUG-AGU-----------U
A-U-------------G---U----C----A---UGU----U-A-UCGA-UAU-----GUGA----------

```
-----------A-----------UACA----------------------------------------UAG-C
A-U--------------------AU-CAGAAGGCACACCCGGAGAACUGAAACAUCUUAGUACCCG--GAGG
AAGAGAAAGAAA--AUUCGAUUCCCUUA--GU-AGCCGGCGAGCGAAACGGGA-A-GA-GCCCAAACCAAC-
--AA-GCUUG----------------------------------CUU-GUUGGGG--UUG------------
-----UAGG-ACA-------CUCUAUA----------------------CG--GAGUUACAAA--GGAC---
G-------ACAUU-AGACG------AAU-CA-UCUG-------------------------------------
-------GAA-AG-AUGAA-------UCAAAGAAGGUAAUAAUCCUGUA---------GUCGAAA----AUG
U------U--GUCUC----------UCUU---GAGU--GG-AU------CCUGAGUACGACGGAGCACGUGA
AAUUCCGUCGGAA-UCUGGGAGG-ACC--AUCUCCUAAGGCUAAAUA--CUC-UCUAG-UGACCGAUAGU-G
AAC-C-AGUACCGUGAGGGAAAG-GUGAAA-AGCAC-CCCGG-AAGGGG-A-GUGAAAUAGAACCUGAAACC
G-UGUGCUUACAA-GUA-GU-CAGA---GCCC-G--------------------------------------
------------------------------------------------------------------------
------------------------------------------------------------------------
------------------------------------------------------------------------
------------------------------------------------------------------------
------------------------------------------------------------------------
------------------------------------------------------------------------
------------------------------------------------------------------------
------------------------------------------------------------------------
--------------------------------------------------------------U--U-AA---
------------------------------------------------------------------------
------------------------------------------------------------------------
------------------------------------------------------------------------
------------------------------------------------------------------------
------------------------------------------------------------------------
------------------------------------------------------------------------
------------------------------------------------------------------------
------------------------------------------------------------------------
------------------------------------------------------------------------
------------------------------------------------------------------------
------------------------------------------------------------------------
------------------------------------------------------------------------
------------------------------------------------UG--G---GUGAUGGCGUGCCUUU
UGUAGAAUGAACCGGCGAGUUAC-G-AUUUGAUGCAA-GGUUAAGC--A----------GUAAA--------
--UG--UGGA--GCCGUAGCGAAAG-C---------------------------------------------
------------------------------------------------------------------------
------------------------------------------------------------------------
---------------------------------------------------------------------GAG
UCU------------------------------------GAAUA-------------------------GG-
G-C-GU----------------------UUAGUAUUUGGUCGUA-GACCCGAAACC-AGGUGAUCUACCCUU
GGUCAGGUUGAAGUUCAG-GUAACA-CUGAAUGGAGGACCGA--ACCGACUU-ACGUUGAAAAGUGA-GCGG
AUGAACUGAGGGUAGCGGAGAAAUUCCAAUCGAACCUGG-AGAUAGCUGGUUCUCUCCGAAA-UAGCUUUAG
GGCUAGCCUCAA-GUG--------------------------------------------------------
----------------------AU-----GA-UUAUUGGAGGUAGAGC-ACUGUUU-GGA-CGA--GGGGCC
C---CUCUC--GGGUUACCGAAUUCAGACAAACUCCGAAUGCCAAU--------------------------
```

```
------------------------------------------------------------------------
------------------UAAU-U-UA--------ACUUGGG-AGUCAGA-A-CAUG-G-GUGAUAAGGUCC
GU-G-UUCGAAAG-GGAAAC---AGCCCAGACCACCA-GCUAAGGUCCCAAAAU--AU-AUG-UU-AA---G
-U-GGA-------AAAGGAUGUGGCGUUGCCCAGACAACUAGGAUGUUGGCUUAGAAGCAGCCA-UCAUUUA
AAGAGUGCGUAAUAGCUCACUAGUC-----GAGUGACACUGCGCCGAAAAUGUACC-GGGGCU-A-AACAUA
U----------UACCGAA-GCUGUGGAUUGUCCU--U-----------------------------------
------------------------------------------------------------------------
------------------------------------------------------------------------
----------------------------------UGG-ACAA-UGGUAGGAGAGC-GUUCUAA---------
-----------G-G-GCGUU-GAAGCAU--G-AUCGUAAGGA--C-AUGUGGAGCG-C-UUA-GAAGUGAGA
AUGC-CGGUGU-GAGUA-GCGAAA----GAC-GGGUGA--G---------------AAUCCC-GUC-CACCG
A-UUGACU-AAGGUUUCC-AGAGGAAGG-CUCGUCCGCUCUGGGUUAGUCGGG-UC-CUAAGCUGAGGCCG-
A-C-AGGC-----GUA--------------------------------------------------------
----------------------------------------GGC-GA-U--GGAU-AACAGGUU-GAUA-UUC
CUGUA----------------------------------CCACC----UAUA--------A-UC--------
------------------GUUUUAA-----------------------------------------------
---------------------------------------------------------UCGAUG-GGGGGGAC
GCAGUA--------------GGAUAG-GCGAAGCGU-GC-G----AUUGG------------AUUGCACGU-
CUAAGC-AGU-AA-----G-G-------------------C------UG------------------AG-UA
U-UAGG--------------------CAAAU-CCGGUACUCA------UUAAGGCUG----AGC-UG-U-G-
AUG--GG---------------------------------------------------------------GA
----GAAGACA-UUGUGUCUUCGAG-U-CGU-U-GAUUUCACACUGC-C-GAGAAA-AGCCU-CUAGAU---
--------------------------------------------------------AGAAAA-UA-GGUGCC
CGUACCGCAAACCGAC------------------------------ACAGGUA--GUCAAGAU-GAGAAUUC
UAAGGU-G-AG-CGAG-CGAACUCU-C-GUUAAGGAACUCGGC-AAAAUGACCCCGUAACUUCGG-GAGAAG
GGGUG-CUCU-U-UA--GGGUUAAC-----------------------------------------------
------------------------------------------------------------------------
------------------------------------------------------------------------
------------------------------------------------------------------------
------------------------------------------------------------------------
------------------------------------------------------------------------
------------------------------------------------------------------------
------------------------------------------------------------------------
------------------------------------------------------------------------
------------------------------------------------------------------------
------------------------------------------------------------------------
------------------------------------------------------------------------
------------------------------------------------------------------------
------------------------------------------------------------------------
------------------------------------------------------------------GCCCAG
AAG--AGCCGCAGUGAAUAGGCCCAAGCGA-----CUGUUUAUCAAAAACACAGGUCUCUGCUAAACCG---
-UA-A-------------GGUGAUGU-AUAGGGGCUGA-CGCCUGCCCGGUGCUG-G-AAGG-UUAAGAGGA
GUGGUUA-GC-----UUCU-----GCGAAGCUACGAAUCGAAGCCC--CAGUAAAC-GGCGGCCGUAACUAU
AACGGUCCUAAGGUAGCGAAAUUCCUUGUCGGG-UAAGUUCCGACCCGCACGAAAGGCGUAACGAUU--UGG
GCACUGUCUCAACG----AGA-GACUCGGUGAAAUC--AUA---GUACCUGUGAAGAUGCAGGUUACCCGCG
```

```
ACAGGACGGAAAGA-CCCCGUGGAGC-UUUACU--GUAGCCU-GAUAUUGAAAUUCGGCACAGCUUGUACAG
GAU-AGGUAGGAGCCUUUG----------AAACGUG-AGC----------------------------GCUA
--------------------------------------GC-UUACGUGGAG-G--C-GC-UGGUGGGAUA-C
UACC-CUAGCUGUGUUGGCUUUCUAACCCGCA--CCAC-U-UA-U-C-------------------------
------------------------------------------------------------------------
------------------------------------------------------------------------
------------------------------------------------------------------------
------------------------------------------------------------------------
------------------------------------------------------GU----GG----------
------------------------------------------------UGGG-----AGACA-GUGUCAGGC
GGGCAGUUUGACUGGGGCGGU-CGCCUCCUA-AAAGGUAAC-GGAGGCGCUCA-AAGGUUCCCUCAGAAUGG
UUGGAAAUCAUUCAUA-GAGUG-UAAAGGCAU-AAGGGAGCUUGACU-GCGA--------------------
--------------GACCUA---CAAGUCGAGCAGGGUCGAAA-GACGGACUUAGUGAUCCGGU--------
---GG-U-----UCCGC-AUG--------------------GA-AGGGCCAUCGCUCAACGG-AUAAAAGCU
ACCCCGGGG-AUAACAGGCUUAUCUCCCCCAAGAGUUCACAUCGACGGGGAGGUUUGGCACCUCGAUGUCGG
CUCAUCGCAUCCUGGGGCUGUA-GUCGGUCCCAAGGGUUGGGCUGUUCGCCCAUUAAAGCGGUACGCGAGCU
GGGUUCAGAACGUCGUGAGACAGUUCGGUCCCUAUCCGUC--------------------G-UGG-------
-------------------------------GCGU--AGGAAAUUUGAGAGG-AGC-UGUCCUUAGUACGAG
AGGAC-CGGGAUGGACAUACCUC-UGGU-G-UACCAGUUGUCGUGCCAACGGCAU-A-GCUGGGUAGCUAUG
UGUGGACGGGAUAAGUGCUGAAAGCA-UU--UAAGCAUGAAGCCCCCCUCAA-G-AUG--AGAUUUCC-CA-
--ACUUC-----------------------------------------------------------------
------------------------------------------------------------------------
------------------------------------------------------------------------
------------------------------------------------------------------------
------------------------------------------------------------------------
------------------------------------------------------------------------
------------------------------------------------------------------------
------------------------------------------------------------------------
---------------------------------------------------GGUUA--UA-AGA------UC
CCUCAUAGAUGAUGAGGU-UAA-UAGGUU-CGA-GGUGGAAG-CAU-G----------------GU------
----GA-----------CAUGUGGAGCUG--A-CGAA-UACUAAUC---GAUCGAA-G-ACUUAAUCA----
------------------------------------------------------------------------
------------------------------------------------------------------------
------------------------------------------------------------------------
------------------------------------A------------
>LSUb_171101_STRR6/1-2902
---------------------G-GUUAAGUU----AA--UAAGGGCGCAC-GGUGGAUG-C-CUUGGC-ACU
AGG---AGCCGACGAAGGACGUGA--CAAACGACGAUAUGCCUUGGGUAGCUGUAAGUAAGCGA------UG
AUC-CAG-GGAUUUCCGAAUGGGGGAAC--------CC--------AA----CAGG-U-------------A
A-U-----------------A----C----C---UGU----U-A-CCCA-CAU-----CU------------
-----------G-----------UUAA------------------------------------------G-G
A-U--------------------GU-GAGGAGGAAGACGCAGUGAACUGAAACAUCUAAGUAGCUG--CAGG
AAGAGAAAGCAA--AAGCGAUUGCCUUA--GU-AG-CGGCGAGCGAAACGGCA-G-GA-GGGCAAACCGAA-
--GA-GUUUA----------------------------------CUC-UUCGGGG--UUG------------
-----UAGG-ACU-------GCAA-UG----------------------UG--GACUCA--------AA---
```

G-------AUUAU-AGAAG------AAU-GA-UUUG------------------------------------
-------GGA-AG-AUCAG-------CCAAAGAGAGUAAUAGCCUCGUA---------UUUAAAA----UAG
U------C--UUUGU----------ACUU---AGCA--GU-AU------CCUGAGUACGGCGGGACACGUGA
AAUCCCGUCGGAA-UCUGGGAGG-ACC--AUCUCCCAACCCUAAAUA--CUC-CCUAG-UGACCGAUAGU-G
AAC-C-AGUACCGUGAGGGAAAG-GUGAAA-AGCAC-CCCGG-GAGGGG-A-GUGAAAUAGAACCUGAAACC
G-UGUGCCUACAA-CAA-GU-UCGA---GCCC-G--------------------------------------
------------------------------------------------------------------------
------------------------------------------------------------------------
------------------------------------------------------------------------
------------------------------------------------------------------------
------------------------------------------------------------------------
------------------------------------------------------------------------
------------------------------------------------------------------------
------------------------------------------------------------------------
--------------------------------------------------------------U--U-AA---
------------------------------------------------------------------------
------------------------------------------------------------------------
------------------------------------------------------------------------
------------------------------------------------------------------------
------------------------------------------------------------------------
------------------------------------------------------------------------
------------------------------------------------------------------------
------------------------------------------------------------------------
------------------------------------------------------------------------
------------------------------------------------------------------------
------------------------------------------------------------------------
------------------------------------------------------------------------
------------------------------------------------UG--G---GUGAGAGCGUGCCUUU
UGUAGAAUGAACCGGCGAGUUAC-G-UUAUGAUGCGA-GGUUAAGU--U----------GAAGA--------
--GA--CGGA--GCCGUAGGGAAAC-C---------------------------------------------
------------------------------------------------------------------------
------------------------------------------------------------------------
---------------------------------------------------------------------GAG
UCU------------------------------------GAAUA-------------------------GG-
G-C-AC---------------------CUUAGUAUCAUGACGUA-GACCCGAAACC-AUGUGACCUACCCAU
GAGCAGGUUGAAGGUGCG-GUAAGA-CGCACUGGAGGACCGA--ACCAGGGC-ACGUUGAAAAGUGC-UUGG
AUGACUUGUGGGUAGCGGAGAAAUUCCAAACGAACUUGG-AGAUAGCUGGUUCUCUCCGAAA-UAGCUUUAG
GGCUAGCGUCGACAUUA-------------------------------------------------------
----------------------GA-----GA-UUCUUGGAGGUAGAGC-ACUGUUU-GGG-UGA--GGGGUC
C---AUCCC--GGAUUACCAAUCUCAGAUAAACUCCGAAUGCCAAU--------------------------
------------------------------------------------------------------------
------------------GAAU-U-AU--------GGUCGGC-AGUCAGA-C-UGCG-A-GUGCUAAGAUCC
GU-A-GUCGAAAG-GGAAAC---AGCCCAGACCACCA-GCUAAGGUCCCAAAAU--AA-UUG-UU-AA---G
-U-GGA-------AAAGGAUGUGGGGUUGCACAGACAACUAGGAUGUUAGCUUAGAAGCAGCUAUUCAUUCA
AAGAGUGCGUAAUAGCUCACUAGUC-----GAGUGACCCUGCGCCGAAAAUGUACC-GGGGCUAA-AACAAU

```
U----------UACCGAA-GCUGUGGAUACCUUU--------------------------------------
------------------------------------------------------------------------
------------------------------------------------------------------------
----------------------------------AUA-GGUA-UGGUAGGAGAGC-GUUCUAU---------
-----------G-U-GUGAU-GAAGGUA--U-ACCGUGAGGA--G-UGCUGGAACG-C-AUA-GAAGUGAGA
AUGC-CGGUAU-GAGUA-GCGAAA----GAC-AGGUGA--G---------------AAUCCU-GUC-CACCG
U-AAGACU-AAGGUUUCC-AGGGGAAGG-CUCGUCCGCCCUGGGUUAGUCGGG-AC-CUAAGGAGAGACCG-
A-A-AGGU-----GUA--------------------------------------------------------
----------------------------------------UCC-GA-U--GGAC-AACAGGUU-GAUA-UUC
CUGUA----------------------------------CUAGA----GUAU--------G-UA--------
------------------------------------------------------------------------
----------------------------------------------------------GUGAU-GGAGGGAC
GCAGUA--------------GGCUAA-CUAAAGCAG-ACGA----UUGGA------------AGAGUCUGU-
CUAAGC-AGU-GA-----G-G-------------------U------GU------------------GA-AU
U-GAGU--------------------CAAAU-GCUUAAUUCU------AUAACAUUG----AGC-UG-U-G-
AUG--GG---------------------------------------------------------------GA
-----GCGAAG-UUUAGUAGCGAAG-U-UAG-U-GACGUCACACUGC-C-AAGAAA-AGCUU-CUAGCGUU-
-------------------------------------------------------UAACCAU-AC-UCUACC
CGUACCGCAAACCGAC------------------------------ACAGGUA--GUCGAGGC-GAGUAGCC
UCAGGU-G-AG-CGAG-AGAACUCU-C-GUUAAGGAACUCGGC-AAAAUGACCCCGUAACUUCGG-GAGAAG
GGGUG-CUGA-C-UU---------------------------------------------------------
------------------------------------------------------------------------
------------------------------------------------------------------------
------------------------------------------------------------------------
------------------------------------------------------------------------
------------------------------------------------------------------------
------------------------------------------------------------------------
------------------------------------------------------------------------
------------------------------------------------------------------------
------------------------------------------------------------------------
------------------------------------------------------------------------
------------------------------------------------------------------------
------------------------------------------------------------------------
------------------------------------------------------------------------
---------------------------------------------------------------------UAA
GUC--AGCCGCAGUGAAUAGGCCCAAGCAA-----CUGUUUAUCAAAAACACAGCUCUCUGCUAAAUCG---
-UA-A--------------GAUGAUGU-AUAGGGGUGA-CGCCUGCCCGGUGCUG-G-AAGG-UUAAGAGGA
GUGCUUA-GC-----GUAA-----GCGAAGGUAUGAAUUGAAGCCC--CAGUAAAC-GGCGGCCGUAACUAU
AACGGUCCUAAGGUAGCGAAAUUCCUUGUCGGG-UAAGUUCCGACCCGCACGAAAGGCGUAAUGAUU--UGG
GCACUGUCUCAACG----AGA-GACUCGGUGAAAUU--UUA---GUACCUGUGAAGAUGCAGGUUACCCGCG
ACAGGACGGAAAGA-CCCCAUGGAGC-UUUACU--GCAGUUU-GAUAUUGAGUGUCUGUACCACAUGUACAG
GAU-AGGUAGGAGUCUAAG----------AGAUCGG-GAC----------------------------GCCA
--------------------------------------GU-UUCGAAGGAG-A--C-GC-UGUUGGGAUA-C
UACC-CUUGUGUUAUGGCCACUCUAACCCAGA--UAGG-U-GA-U-C-------------------------
------------------------------------------------------------------------
```

```
------------------------------------------------------------------------
------------------------------------------------------------------------
------------------------------------------------------------------------
------------------------------------------------------CC----UA----------
------------------------------------------------UCGG-----AGACA-GUGUCUGAC
GGGCAGUUUGACUGGGGCGGU-CGCCUCCUA-AAAGGUAAC-GGAGGCGCCCA-AAGGUUCCCUCAGAAUGG
UUGGAAAUCAUUCGCA-GAGUG-UAAAGGUAU-AAGGGAGCUUGACU-GCGA--------------------
--------------GAGCUA---CAACUCGAGCAGGGACGAAA-GUCGGGCUUAGUGAUCCGGU--------
---GG-U-----UCCGU-AUG--------------------GA-AGGGCCAUCGCUCAACGG-AUAAAAGCU
ACCCUGGGG-AUAACAGGCUUAUCUCCCCCAAGAGUUCACAUCGACGGGGAGGUUUGGCACCUCGAUGUCGG
CUCGUCGCAUCCUGGGGCUGUA-GUCGGUCCCAAGGGUUGGGCUGUUCGCCCAUUAAAGCGGCACGCGAGCU
GGGUUCAGAACGUCGUGAGACAGUUCGGUCCCUAUCCGUC--------------------G-CGG-------
-------------------------------GCGU--AGGAAAUUUGAGAGG-AUC-UGCUCCUAGUACGAG
AGGAC-CAGAGUGGACUUACCGC-UGGU-G-UACCAGUUGUCUUGCCAAAGGCAU-C-GCUGGGUAGCUAUG
UAGGGAAGGGAUAAACGCUGAAAGCA-UC--UAAGUGUGAAACCCACCUCAA-G-AUG--AGAUUUCC-CA-
--UGAUU-----------------------------------------------------------------
--------------------------AU--------------------------------------------
------------------------------------------------------------------------
------------------------------------------------------------------------
------------------------------------------------------------------------
------------------------------------------------------------------------
------------------------------------------------------------------------
------------------------------------------------------------------------
-------------------------------------------------AUAUCAG--UA-AGA------GC
CCUGAGAGAUGAUCAGGU-AGA-UAGGUU-AGA-AGUGGAAG-UGU-G----------------GC------
----GA-----------CACAUGUAGCGG--A-CUAA-UACUAAUA---GCUCGAG-G-ACUUAUCCA----
------------------------------------------------------------------------
------------------------------------------------------------------------
------------------------------------------------------------------------
------------------------------------A------------
>LSUb_292459_SYMTH/1-2946
---------------------A-AUCAAGCU----GG--AAAGGGCACAC-GGCGGAUG-C-CUGGGC-GCC
AGG---CGCCGAUGAAGGACGUGG--CAAGCUGCGAUAAGCCUCGGGGAGCCGCAAGCAGGCGU------AG
AUC-CGG-GGAUCUCCGAAUGGGGCAAC--------CC--------GG----CUGG-CGU-----------A
A-U-------------G---G----C----C---AGU----C-A-CCCA-CAG-----CUGA----------
-----------A-----------UCCA----------------------------------------UAG-G
C-U--------------------GU-GAGG-AGGGCACCGCCCGAACUGAAACAUCUCAGUAGGGC--GAGG
AAAAGAAAUCAA--CCGAGAUUCCCUCA--GU-AG-CGGCGAGCGAACGGGGA-G-GA-GCCUAAACCAGUU
-CCG-GCUGA-------------------------------GC-CGGGGCUGGGG--UUG------------
-----CGGG-ACG-------CCGA-UG----------------------AA--GCGAU-------GCUC-CU
G-------AGCCU-AGCUG------AACAGG-UCUG------------------------------------
-------GAA-AG-GCCGG-------CCACAGAAGGUAAUAGCCCUGUA---------GGCGAAA----GGC
G------A--AGGAC----------CGCU---UGGC--GG-AU------CCCAAGUACCACGGGACACGAGG
AAUCCCGUGGGAA-UCCGGGGGG-ACC--ACCCUCCAAGGCUAAAUA--CGC-CU-GG-CGACCGAUAGU-G
AAC-C-AGUACCGUGAGGGAAAG-GUGAAA-AGCAC-CGCGG-GAGCGG-A-GUGAAAGAGACCCUGAAACC
```

```
G-UGUGCCUACAA-GCA-GU-ACGA---GCCC-C--------------------------------------
------------------------------------------------------------------------
------------------------------------------------------------------------
------------------------------------------------------------------------
------------------------------------------------------------------------
------------------------------------------------------------------------
------------------------------------------------------------------------
------------------------------------------------------------------------
------------------------------------------------------------------------
-------------------------------------------------------------AU--A-UA---
------------------------------------------------------------------------
------------------------------------------------------------------------
------------------------------------------------------------------------
------------------------------------------------------------------------
------------------------------------------------------------------------
------------------------------------------------------------------------
------------------------------------------------------------------------
------------------------------------------------------------------------
------------------------------------------------------------------------
------------------------------------------------------------------------
------------------------------------------------------------------------
------------------------------------------------------------------------
------------------------------------------------AG--G---GUGAGUGCGUGCCUAU
UGAAGAAUGAACCGGCGAGUUAC-G-GUCACUGGCAU-GGUUAAGG--G-C--------CUCAG--------
G-CC--CGGA--GCCGAAGCGAAAG-C---------------------------------------------
------------------------------------------------------------------------
------------------------------------------------------------------------
---------------------------------------------------------------------GAG
UCU------------------------------------GAAGA-------------------------GG-
G-C-GU------------------------AGUCAGUGGCUGUA-GACCCGAAACC-CGGUGAUCUAUCCAU
GGCCAGGGUGAAGCGGGG-GUAAGA-CCUCGUGGAGGCCCGA--ACCGGUCA-GUGUUGAAAAACUG-UCGG
AUGAGCUGUGGAUAGGGGUGAAAAGCCAAACGAACCGGG-AGAUAGCUGGUUCUCCCCGAAA-UAGCUUUAG
GGCUAGCGUCGGCAGAG-------------------------------------------------------
----------------------AA-----GC-UUGCUGGAGGUAGAGC-ACUGAUU-GGG-CUA--GGGGCC
----CUAUC---GGUUACCGAACCCAGUCAAACUCCGAAUGCCAGC--------------------------
------------------------------------------------------------------------
------------------AAGU-A--U--------GGCCGGC-AGUCAGA-C-GGCG-A-GCGCUAAGGUCC
GU-C-GUCAAGAG-GGAAAC---AGCCCAGACCGCCA-GCUAAGGUCCCCAAGU--CU-CUG-CU-AA---G
-U-GGU-------CAAGGAUGUGGCGUUGUAGAGACAACCAGGAUGUUGGCUUAGAAGCAGCCA-UCAUUGA
AAGAGUGCGUAACAGCUCACUGGUC-----GAAUGACGCUGCGCCAUAAAUAC-UC-GGGGCU-A-AGCAGA
G----------CACCGAA-GCUGCGGACUGCCG---------------------------------------
------------------------------------------------------------------------
------------------------------------------------------------------------
---------------------------------UAGG-GCAG-UGGUAGGGGAGC-GUUCCAU---------
-----------G-U-GCGGC-GAAGGUC--G-ACCGGAAGGA--C-GGCUGGAGCG-C-AUG-GAAGUGAGA
```

```
AUGC-CGGUAU-GAGUA-GCGAAA---AGAC-AGGUGA--G---------------AAUCCU-GUC-CACCG
A-AAGCCU-AAGGAUUCC-UGAGCAAGG-CUCGUCCGCUCAGGGUCAGUCGGG-AC-CUAAGCCGAGGCCG-
A-A-UGGC-----GUA--------------------------------------------------------
----------------------------------------GGC-GA-U--GGGA-AACAGGUC-GACA-UUC
CUGUA----------------------------------CCACC----UACG--------G-GU--------
------------------CGCUUGA-----------------------------------------------
----------------------------------------------------------CCGAU-GCGGGGAC
GCAGGA--------------GGCUAG-GGGGACCCG-GCCG----CUGGU------------UGAGCCGGG-
GUAAGC-GGG-UA-----GCG-------------------U------GG------------------CC-CG
G-UAGG--------------------GAAGU-CCGCCGGGCU--------AAACGUG----AGA-CG-C-G-
AUG--CC---------------------------------------------------------------GA
-----CCGAAA-UGAAGUAGGGAAG-C-CCC-U-GACGCCACACUGC-C-GAGAAA-AGCCG-CUAGGG---
--------------------------------------------------------AGACCC-GG-GGUGCC
CGUACCGCAAACCGAC------------------------------ACAGGUA--GGCGGGGA-GAGAAUCC
UCAGGU-G-CG-CGGG-AGAACCCU-C-GCUAAGGAACUCGGC-AAAAUGCAUCCGUAACUUCGG-GAGAAG
GAUGG-CCCU-C-GA-ACCGUGAGC----GGA-----------GAGAC------------------------
------------------------------------------------------------------------
------------------------------------------------------------------------
------------------------------------------------------------------------
------------------------------------------------------------------------
------------------------------------------------------------------------
------------------------------------------------------------------------
------------------------------------------------------------------------
------------------------------------------------------------------------
------------------------------------------------------------------------
------------------------------------------------------------------------
------------------------------------------------------------------------
------------------------------------------------------------------------
------------------------------------------------------------------------
----------------------------------------------------------GCCGCAGAGCGGGG
GAG--GGUUGCAGAGGAGAGGCCCAAGCGA-----CUGUUUACCACAAACACAGGUCAGUGCUAAGCCG---
-GA-A-------------GGCGACGU-AUACUGGCUGA-CGCCUGCCCGGUGCUG-G-AAGG-UUAAGGGGA
AGGGUCA-GGCU---GCAA---GGCCGAAGCUCUGAACCGAAGCCC--CAGUAAAC-GGCGGCCGUAACUAU
AACGGUCCUAAGGUAGCGAAAUUCCUUGUCGGG-UAAGUUCCGACCCGCACGAAUGGCGUAACGACU--UGG
GCACUGUCUCGGCG----AGG-GACCCGGUGAAAUU--GCA---AUACCUGUGAAGAUGCAGGUUACCUGCG
AUUGGACAGAAAGA-CCCCAUGGAGC-UUUACU--GUAGCCU-GGCAUUGGAUUUUGGUGGUGCGUGUACAG
GAU-AGGUGGGAGCCUGCG----------AAGCCGG-GUC----------------------------GCCA
--------------------------------------GA-UUCGGUGGAG-G--C-GC-CGUUGGGAUA-C
CACC-CUCGCAGCAUCGAAGUUCUAACACAGG--CCCC-UGGA-G-C-------------------------
------------------------------------------------------------------------
------------------------------------------------------------------------
------------------------------------------------------------------------
------------------------------------------------------------------------
------------------------------------------------------------------------------GG----GG----------
------------------------------------------------CCGU-----GGACC-GUGCCAGGU
```

GGGCAGUUUGACUGGGGCGGU-CGCCUCCCA-AAAGGUAAC-GGAGGCGCCCA-AAGGUCCGCUCAGGAUGG
UUGGAAAUCAUCCGUG-GAGUG-CAAAGGCAG-AAGCGGGCUUGACU-GCGA--------------------
--------------GACAGA---CAGGUCGAGCAGGGUCGAAA-GACGGGCUUAGUGAUCCGGC--------
---GG-U-----UCCGG-GUG--------------------GA-AGGGCCGUCGCUCAACGG-AUAAAAGCU
ACCCUGGGG-AUAACAGGCUGAUCUCCCCCAAGAGUCCACAUCGACGGGGAGGUUUGGCACCUCGAUGUCGG
CUCAUCGCAUCCUCGGGCUGAA-GUCGGUCCGAAGGGUUCGGCUGUUCGCCGAUUAAAGCGGUACGCGAGCU
GGGUUCAGAACGUCGUGAGACAGUUCGGUCCCUAUCCAUC--------------------G-CAG-------
-------------------------------GCGG--AGGAGAUUUGACGGG-AUC-UGCCCCUAGUACGAG
AGGAC-CGGGGUGGACGGCCCGC-UGGU-G-UACCAGUUAUCCCGCCAGGGGUAC-A-GCUGGGUAGCCAUG
GCCGGAACGGAUAAGCGCUGAAAGCA-UC--UAAGCGCGAAGCCGGCCCGAA-G-AUG--AGAUCUCC-CA-
--UCCCG-----------------------------------------------------------------
--------------------------C---------------------------------------------
------------------------------------------------------------------------
------------------------------------------------------------------------
------------------------------------------------------------------------
------------------------------------------------------------------------
------------------------------------------------------------------------
------------------------------------------------------------------------
-------------------------------------------------AAGGGAG--UA-AGG------CC
CCUGGAAGAUGACCAGGU-AGA-UAGGCC-GGG-AGUGGAAG-UGC-C----------------GU------
----AA-----------GGCAUGGAGCGG--A-CCGG-UACUAAUC---GGCCGAG-G-GCUUGAUUU----
------------------------------------------------------------------------
------------------------------------------------------------------------
------------------------------------------------------------------------
------------------------------------C------------
>LSUb_265311_MESFL/1-2897
---------------------G-----------------UAAGGGCACAU-GGUGAAUG-C-CUUGGA-AAA
UGG---AGCCGAAGAAGGACGUGA--UUACCUGCGAAAAGCAUCGGGGAGCUGGAAGUGAGCUU------UG
AUC-CGG-UGAUAUCCGAAUGGGAAAC---------CC--------AA----UACG-A-U----------UA
A-U-------------C---U----C----G---UAU----U-A-UCCA-UAA-----GUGA----------
-----------A-----------UACA----------------------------------------UA-GC
U-U--------------------AU-GAGA-AGGGAACCUUGGGAACUGAAACAUCUUAGUACCAA--GAGG
AAAAGAAAAUAAA-UAAUGAUUCUGUUA--GU-AG-CGGCGAGCGAAAGCGGA-A-GA-GGCCAAACCAG--
--UC-U---A----------------------------------CGG-GCUGGGG--UUG------------
-----UAGG-ACA-------UCUU-UU----------------------UA--GAGUUACAAA--AUCA---
A-------CAUAU-AGUAG------AAG-UU-ACUG-------------------------------------
-------GGA-AG-UAACG-------GCAUAGAGGGUGAUACCCCGUA----------UACGAAA----UGU
G------U--UGAUC----------UCUU---AGAU--GG-AU------CCUGAGUACGGCGAAACACGUGA
AAUUUUGUCGGAA-UCCGCCGAG-ACC--ACUCGGCAAGCCUAAAUA--CUC-CCAUU-UUACCGAUAGU-G
AAC-C-AGUACCGUGAGGGAAAG-GUGAAA-AGUAC-CCCGA-GAGGGG-A-GUGAAAUAGUUCCUGAAACC
A-UGUGCUUACAA-GAA-GA-UAGA---GCCC-G--------------------------------------
------------------------------------------------------------------------
------------------------------------------------------------------------
------------------------------------------------------------------------
------------------------------------------------------------------------

```
------------------------------------------------------------------------
------------------------------------------------------------------------
------------------------------------------------------------------------
------------------------------------------------------------------------
--------------------------------------------------------------U--U-AA---
------------------------------------------------------------------------
------------------------------------------------------------------------
------------------------------------------------------------------------
------------------------------------------------------------------------
------------------------------------------------------------------------
------------------------------------------------------------------------
------------------------------------------------------------------------
------------------------------------------------------------------------
------------------------------------------------------------------------
------------------------------------------------------------------------
------------------------------------------------------------------------
------------------------------------------------------------------------
------------------------------------------------UG--G---GUGAUAUCGUGCUUUU
UGUAGAAAGAGCCGGCGAGUUAA-U-GUAUCGUGCGA-GGUUAAGC--A----------GAAUA--------
--UG--CGGA--GCCGUAGUGAAAG-C---------------------------------------------
------------------------------------------------------------------------
------------------------------------------------------------------------
---------------------------------------------------------------------GAG
CCU------------------------------------UAAUA-------------------------GG-
G-C-GU----------------------UUAGUACGUUGCAUUA-GACACGAAACC-GGGUGAUCUAGCCAU
GAGCAGGUUGAAGUUUGG-GUAAAA-CCAAAUGGAGGACCGA--ACCGACGU-UCGUUGAAAUGACC-GCGG
AUGACUUGUGGCUAGCGGUGAAAUUCCAAUCGAACCCGG-AGAUAGCUAGUUCUCCCCGAAA-UAUAUUUAA
GUAUAGCGUCGA-GGUU-------------------------------------------------------
----------------------UA-----CC-ACAAUGGAGGUAGAGC-ACUGAAU-CUA-UGA--UGGCCC
C---ACCUA--GGGGUACUGAAUAGAAUUAAACUCCGAAUGCCAUU--------------------------
------------------------------------------------------------------------
------------------GUCG-G-AU--------ACUCGGC-AGUCAGA-A-CAUG-G-GUGAUAAGGUCC
AU-G-CUCGAGAG-GGAAAC---AGCCCAGAUCGUCA-GCUAAGGUCCCUAAAU--UU-AGG-UU-AA---G
-U-GUG-------UAAGGAUGUGGAAUUGCACAGACAGCUAGGAUGUUGGCUUAGAAGCAGCCA-CCAUUUA
AAGAGUGCGUAACAGCUCACUAGUC-----GAGUGAUUCUGCGCCGAAAAUGUACC-GGGGCU-A-AACCUA
A----------UACCGAA-GCUACGGAUUGAUUU--U-----------------------------------
------------------------------------------------------------------------
-----U------------------------------------------------------------------
---------------------------------AAAU-ACAG-UGAUAGGGGAGC-GUUCUAA---------
-----------U-U-GUGAU-GAAGUCA--G-ACUGUGAGGA--C-UGGUGGAACG-A-UUA-GAAGUGAUU
AUGC-CGGCAU-GAGUA-ACGAU-----UGA-AUGUGA--G---------------AAUCAU-UCA-UACCG
U-UUGACC-AAGGUUUCC-UGGGCAAGG-UUCGUCCACCCAGGGUUAGUCAGG-AC-CUAAGGCGAGGCCG-
A-A-AGGC-----GUA--------------------------------------------------------
----------------------------------------GUC-GA-U--GGAC-AACAGGUU-GAUA-UUC
CUGUA----------------------------------CCUGC----UA----------G-UU--------
```

```
------------------A-----------------------------------------------------
----------------------------------------------------------GUGAU-GGAGUGAC
GGAGAA--------------AGGUAGUGUAUCCCAG-GU-G----AUGGA------------UGUCCUGGG-
UUAAGC-ACA-AA-----G-G-------------------C------GG------------------CA-AC
A-UAGG--------------------CAAAU-CCGUGUUGUA------UAAACGCUG-----GU-UA-U-G-
AGGA-GU---------------------------------------------------------------GA
AC---GGUUCG--CCUAGUAACGAA-G-UAC-A-UGACCCUACGCUU-C-CAAGAA-AAGCU-UCUAACUU-
--------------------------------------------------------AAUAAC-UA-GUAGCC
UGUACCUAUAACGAAC------------------------------ACACGUG--GUCAAGGA-GAAAAUCC
UAAGGU-A-AG-CGAG-AUAACUGU-A-GCUAAGGAACUCUGC-AAAAUAACUCCGUAACUUCGG-AAGAAG
GAGUG-CUCA-A-CG---------------------------------------------------------
------------------------------------------------------------------------
------------------------------------------------------------------------
------------------------------------------------------------------------
------------------------------------------------------------------------
------------------------------------------------------------------------
------------------------------------------------------------------------
------------------------------------------------------------------------
------------------------------------------------------------------------
------------------------------------------------------------------------
------------------------------------------------------------------------
------------------------------------------------------------------------
------------------------------------------------------------------------
------------------------------------------------------------------------
--------------------------------------------------------------------CAAG
UUG--AGCCGCAGUGAAGAGGGAGGGGCAA-----CUGUUUAGCAAAAACACAGCUCUCUGCUAAGUCG---
-CA-A-------------GACGAAGU-AUAGGGGGUGA-CGCCUGCCAGUGCCG-G-AAGG-UUAAGAGGA
GAAGUCA-GC-----GCAA-----GCGAAGCUUCGAAUUGAAGCCC--CGGUGAAC-GGCGGCCGUAACUAU
AACGGUCCUAAGGUAGCGAAAUUCCUUGUCAGG-UAAGUUCUGACCCGCACGAAAGGCGUAAUGAUC--CCU
UCGCUGUCUCGGCU----GCA-GACUCGGUGAAAUU--UUA---GUACCGGUGAAGAUGCCGGUUACCCGCA
ACUAGACGGAAAGA-CCCCGUGGAGC-UUUACU--AUAACUU-GAUAUUGAAAUUUGGUAUAACGUGUAGAG
GAU-AGGUGGGAGACUUG-----------AAGCUGG-ACC----------------------------GCUA
--------------------------------------GG-GCCAGUGGAG-U--C-AA-CCUUGGAAUA-C
CACC-CUCGUUAUAUUGGAUUUCUAACUUCGA--CCCG-U-UA-U-C-------------------------
------------------------------------------------------------------------
------------------------------------------------------------------------
------------------------------------------------------------------------
------------------------------------------------------------------------
------------------------------------------------------CG----GG----------
------------------------------------------------UUAA-----GGACA-GUGUCUGGU
GGGUAGUUUGACUGGGGCGGU-CGCCUCCUA-AAAUGUAAC-GGAGGCGCUCA-AAGUUACACUCAGCAUGG
UUGGAAAUCAUGCAUA-GAGCG-CAAAAGUAU-AAGUGUGAUUGACU-GUGA--------------------
--------------GACUUA---CAAGUCGAACAGGUACGAAA-GUAGGAUUUAGUGAUCCGGC--------
---GG-U-----CCCGU-GUG--------------------GA-AGGGCCGUCGCUCAACGG-AUAAAAGUU
ACCCCGGGG-AUAACAGGCUGAUCUCCCCCAAGAGUUCACAUCGACGGGGAGGUUUGGCACCUCGAUGUCGG
```

CUCAUCGCAUCCUGGAGCUGUA-GUCGGUUCCAAGGGUUGGGCUGUUCGCCCAUUAAAGCGGUACGCGAGCU
GGGUUCAGAACGUCGUGAGACAGUUUGGUCCCUAUCUGUU--------------------G-UGG-------
-------------------------------GCGU--AGGAAAAUUGAAGAG-AGC-UGUUCCUAGUACGAG
AGGAC-CGGAAUGGACACACCUC-UGGU-G-CUCCUGUUGUCACGCCAGUGGCAC-A-GCAGGGUAGCUAUG
UGUGGAACGGAUAAUCGCUGAAGGCA-UC--UAAGCGAGAAGCCUCCUUUAA-G-AUG--AAUUUUCC-CA-
--UUUCU-----------------------------------------------------------------
--------------------------UU--------------------------------------------
------------------------------------------------------------------------
------------------------------------------------------------------------
------------------------------------------------------------------------
------------------------------------------------------------------------
------------------------------------------------------------------------
------------------------------------------------------------------------
--------------------------------------------------AGAAUG--UA-AGA------UC
CCUUAUAGACUAUGAGGU-UGA-UAGGAU-GGA-UGUGUAAG-UGU-C----------------GC------
----GA-----------GGCAUUAAGCUA--A-CCAU-UACUAAUA---GAUCGAGAG-AAUUUUAG------
------------------------------------------------------------------------
------------------------------------------------------------------------
------------------------------------------------------------------------
-------------------------------------------------
>LSUb_273068_CALS4/1-2918
---------------------G-UCAAGUGA----GG--UAAGGGCGCAU-GGUGGAUG-C-CCAGGC-ACC
UGG---AGCCGAUGAAGGACGCGG--CAAGCGGCGAAACGCCCCGGGGAGCCGCAAGCGGGCGU------UG
AUC-CGG-GGAUCUCCGAAUGGGAAAC---------CC--------GC----UUGG-G-G----------UA
A-U-------------A---C----C----C---AAG----C-A-UCCG-CUG-----AUGA----------
-----------A-----------UCCA----------------------------------------UAG-U
C-A--------------------GC-G-GA-GGGGAACCGGGCGAACUGAAACAUCUCAGUAGCCC--GAGG
AAAAGAAAGCAA--AAGCGAUUCCCCGA--GU-AG-UGGCGAGCGAAAGGGGA-A-GA-GCCCAAACCUAC-
--CG-CUUAG----------------------------------GCG-GUAGGGG--UU-------------
-----CGGG-CCC-------ACCG-CA---------------------AAA--GUG----------UGCGAG
U-------ACUCU-AGGCG------AACGGA-CCUG------------------------------------
-------GAA-AG-GCCGG-------CCAUAGAGGGUGAAAGCCCCGUA---------GCCGAAA----GGG
G------A--AAGCG----------CACU---GGUG--GG-UA------CCAGAGUACCACGGGAUAGGCG-
-ACCCUGUGGGAA-GGAGGGAGG-ACC--ACCUCCUAAGGCUAAAUA--CUC-C-AGG-UGACCGAUAGC-G
CA--U-AGUACCGUGAGGGAAAG-GUGAAA-AGAAC-CCCGG-GAGGGG-A-GUGAAAUAGAACCUGAAACC
A-UGUGCCCACAA-GCA-GU-CAGA---GCCU-G---GG---------------------------------
------------------------------------------------------------------------
------------------------------------------------------------------------
------------------------------------------------------------------------
------------------------------------------------------------------------
------------------------------------------------------------------------
------------------------------------------------------------------------
------------------------------------------------------------------------
------------------------------------------------------------------------
--------------------------------------------------------------U--A-AU---

```
------------------------------------------------------------------------
------------------------------------------------------------------------
------------------------------------------------------------------------
------------------------------------------------------------------------
------------------------------------------------------------------------
------------------------------------------------------------------------
------------------------------------------------------------------------
------------------------------------------------------------------------
------------------------------------------------------------------------
------------------------------------------------------------------------
------------------------------------------------------------------------
------------------------------------------------------------------------
----------------------------------------------ACCA--G---GUGAUGGCGUACUUUU
UGCAGUACGGGCCGGCGAGUUAU-G-GUAGCGAGCGA-GGUUAAGG--G-UA--------GAGA-------U
A-CC--CGGA--GCCGAAGCGAAAG-C---------------------------------------------
------------------------------------------------------------------------
------------------------------------------------------------------------
---------------------------------------------------------------------GAG
UCC------------------------------------GAAGA-------------------------GG-
G-C-GG----------------------AAAGUUCGCUGCCAUA-GACCCGAAACC-GUGCGACCUACCCAU
GGCCAGGGUGAAGCCGGA-AUAAAA-UCUGGUGGAGGCCCGA--ACCCGUUG-GCCGUGAAAAGCCA-UGGG
AUGAGCUGUGGGUAGCGGUGAAAUGCCAAUCGAGCACGG-AGAUAGCUGGUUCUCCCCGAAA-UAGCUUUAG
GGCUAGCCUCAG-GGU--------------------------------------------------------
----------------------AA-----AG-CCUGUGGAGGUAGAGCCACUGACU-GGG-CUA--GGGGCC
----GA-GA---GGCUACCGAACCCUAUCAAACUCCGAAUGCCACA--------------------------
------------------------------------------------------------------------
------------------GGCA-A-GU--------ACCUGGG-AGUCAGA-C-CACG-A-GUGAUAAGAUCC
GU-G-GUCGAGAG-GGGAAC---AGCCCAGACCCCCA-GCUAAGGUCCCGAAGG--GC-AUG-UU-AA---G
-U-GGG-------AAAGGAAGUGGGACUUCGAAGACAACCGGGAUGUUGGCUUAGAAGCAGCCAUACAUUUA
AAGAGUGCGUAACAGCUCACCGGUC-----GAGAGGUCCUGCGCCGAAAAUG-AAC-GGGGCUCA-AACAUG
C----------CACCGAA-GCUGGGGCUGCGCGU--------------------------------------
------------------------------------------------------------------------
------------------------------------------------------------------------
----------------------------------AAG-CGCG-GGGUAGGGGAGC-GUACUGC---------
-----------G-C-GGGCA-GAAGGCU--G-AGCGUAAGCG--A-GGCUGGACUG-C-GCA-GUAGUGAGA
AUGC-CGGCAU-GAGUA-GCGAGA----GCA-AGGUGA--G---------------AAACCU-UGC-CGCCG
G-AAGCCU-AAGGUUUCC-UGGGGAAGG-GUAAUCCGCCCAGGGUAAGUCGGG-GC-CUAAGCCGAGGCCG-
G-A-AGGC-----GUA--------------------------------------------------------
----------------------------------------GGC-GA-U--GGGA-AACCGGUU-GAUA-UUC
CGGUA----------------------------------CCACC----GAGG--------G-CC--------
------------------GCUUGA------------------------------------------------
----------------------------------------------------------GGGAA-GCGGGGAC
GCAGCG--------------AGGUAG-GCGGAGCGC-GCGG----UUGGU------------AGAGCGCGU-
CCAACC-AGC-CA-------C-------------------CG--A-AGG------------------GC-AC
G-UAGG--------------------CAAAU-CCGCGUGCUG---------GCAGGG----GGC-UG-G-G-
```

```
AAG--GG-----------------------------------------------------------------
-----GAGCCG------CAAGGCGA-A-GCC-G-CCGAGGCAGCUGC-C-AAGAAA-AGCCG-CUACCG---
--------------------------------------------------------AGGCCU-GA-GGUGCC
CGUACCGCAAACCGAC------------------------------ACAGGUA--GGCGAGGA-GAGAAUCC
ACAGGC-G-AG-CGGG-AGAACCCU-C-CUUAAGGAACUCGGC-AAAUUGACCCCGUAACUUCGG-GAGAAG
GGGUG-CCUCCC-UC--AGGCUUAG----AGU-----------UUAUU------------------------
------------------------------------------------------------------------
------------------------------------------------------------------------
------------------------------------------------------------------------
------------------------------------------------------------------------
------------------------------------------------------------------------
------------------------------------------------------------------------
------------------------------------------------------------------------
------------------------------------------------------------------------
------------------------------------------------------------------------
------------------------------------------------------------------------
------------------------------------------------------------------------
------------------------------------------------------------------------
------------------------------------------------------------------------
----------------------------------------------------------GCUC-UAGGCUUGA
GAA--GGCCGCAGAGAAGAGGCCCAAGCGA-----CUGUUUACCAAAAACACAGGUCUCUGCUAAGCCG---
-AA-A-------------GGCGAAGU-AUAGGGGCUGA-CGCCUGCCCGGUGCUG-G-AAGG-UUAAGGGGA
AGGGU----------GAAA----------GCCCUGAACCGAAGCCC--CAGUGAAC-GGCGGCCGUAACUAU
AACGGUCCUAAGGUAGCGAAAUUCCUUGUCGGG-UAAGUUCCGACCUGCACGAAAGGCGUAACGACU--UGG
GCGCUGUCUCGAGG----AGG-GGCCCGGCGAAGUU--GUA---GUGCUCGUGAAGAUGCGAGCUACCCGCG
AUUGGACGGAAAGA-CCCCGUGGAGC-UUUACU--GUAGCCU-GUCACUGGGUCCUGGUAGUUCCUGUACAG
GAU-AGGUGGGAGACUGAG----------AAGGUGG-CGC----------------------------GCCA
--------------------------------------GC-GUCACUGGAG-U--C-GC-CGGUGGGAUA-C
CACC-CUGGGACUACUGGGGUCCUAACCUAUA--GCCG-U-UA-U-C-------------------------
------------------------------------------------------------------------
------------------------------------------------------------------------
------------------------------------------------------------------------
------------------------------------------------------------------------
------------------------------------------------------CG----GC----------
------------------------------------------------GUAG-----GGACA-AUGGCAGGU
GGGCAGUUUGACUGGGGCGGU-CGCCUCCUA-AAAGGUAAC-GGAGGCGCCCA-AAGGUCCCCUCAGCGCGG
ACGGAAAUCGCGCGUC-GAGUG-CAAGGGCAG-AAGGGGGCUUGACU-GUGA--------------------
--------------GACCGA---CGGGUCGAGCAGGGACGAAA-GUCGGGCCUAGUGAUCCGAC--------
---GG-U-----AGCGA-GUG--------------------GG-AGUGCCGUCGCUCAACGG-AUAAAAGCU
ACCCGGGG-AUAACAGGCUGAUCUCCCCCAAGAGUCCACAUCGACGGGGAGGUUUGGCACCUCGAUGUCGG
CUCAUCGCAUCCUGGGGCUGAA-GAAGGUCCCAAGGGUUGGGCUGUUCGCCCAUUAAAGCGGUACGUGAGCU
GGGUUCAGAACGUCGUGAGACAGUUCGGUCCCUAUCCGUC--------------------G-CGG-------
-------------------------------GCGC--AGGAGACUUGAGGGG-AGC-UGCCCCUAGUACGAG
AGGAC-CGGGGCGGACGAGCCGA-UGGU-G-UACCAGUUGCACCGCCAGGUGCAC-A-GCUGGGUAGCCAAG
CUCGUAUGGGAUAAACGCUGAAAGCA-UC--UAAGCGUGAAGCCCACCCCAA-G-AUA--AGGUCUCC-CA-
```

```
--CCUCG-----------------------------------------------------------------
--------------------------U---------------------------------------------
------------------------------------------------------------------------
------------------------------------------------------------------------
------------------------------------------------------------------------
------------------------------------------------------------------------
------------------------------------------------------------------------
------------------------------------------------------------------------
-------------------------------------------------AAGAGGG--UA-AGG------CA
CCUCGAAGAAGACGAGGU-AGA-UAGGCC-GGG-GGUGUAAG-AGC-G----------------GA------
----AA-----------CGCUCUUAGCUG--A-CCGG-UACUAAUA---AGCCGAG-G-ACUUGACCC----
------------------------------------------------------------------------
------------------------------------------------------------------------
------------------------------------------------------------------------
------------------------------------U------------
>LSUb_240292_ANAVT/1-2830
---------------------G-GUCAAGCU----AA--UAAGGGCUAAU-GGUGGAUA-C-CUAGGC-ACA
CAG---AGGCGAAGAAGGACGUGG--UUACCGACGAAAUACUCCGGGGAGUUGGAAGCAAACUA------UG
AGC-CGG-AGAUGUCCGAAUGGGGCAAC--------CC--------UA------------------------
------------------------------------A----AUA-CUAC-CUG-----UUGA----------
-----------A-----------UAUA----------------------------------------UA-GA
C-A--------------------GG-AAAG-AGCCAACCCAGCGAACUGAAACAUCUUAGUAGCUG--GAGG
AAGAGAAAUCAA--AAGAGAUUCCCUGA--GU-AG-UGGUGAGCGAAAGGGGA-A-GA-GCCUAAACCAGU-
--UG-GUUUA----------------------------------CUG-ACUGGGG--UAG------------
-----UGGG-ACA-------GCGA-UA----------------------UC--GAA---------UCUA---
G-------AGAUU-AGACG------AAG-CA-GCU-------------------------------------
-------AAA-UA-CUGCA-------CCAGAGAAAGUGAAAGUCUUGUA---------GUCGAAA----AUU
G------A--AGGAU-----------AGU---AGCU--GA-AU------CCCGAGUAGCAUGGGGCACGAGG
AAUCCCAUGUGAA-UCAGCGAGG-ACC--AUCUCGUAAGGCUAAAUA--CUA-CUGUG-UGACCGAUAGU-G
AAC-C-AGUACCGCGAGGGAAAG-GUGAAA-AGAAC-CCCGC-AAGGGG-A-GUGAAAUAGAACAUGAAACC
A-UUAGCUUACAA-GCA-GU-GGGA---GUCC-G---------------------------------------
------------------------------------------------------------------------
------------------------------------------------------------------------
------------------------------------------------------------------------
------------------------------------------------------------------------
------------------------------------------------------------------------
------------------------------------------------------------------------
------------------------------------------------------------------------
------------------------------------------------------------------------
-------------------------------------------------------------AU--U-AAA--
------------------------------------------------------------------------
------------------------------------------------------------------------
------------------------------------------------------------------------
------------------------------------------------------------------------
------------------------------------------------------------------------
```

```
------------------------------------------------------------------------
------------------------------------------------------------------------
------------------------------------------------------------------------
------------------------------------------------------------------------
------------------------------------------------------------------------
------------------------------------------------------------------------
------------------------------------------------------------------------
------------------------------------------------CG--G---AUGACCGCGUGCCUGU
UGAAGAAUGAGCCGGCGACUUAU-A-GGCACUGGUA--GGUUAAGA--C-G---------AGAA--------
U-GU--CGGA--GCCAAAGGGAAAC-C---------------------------------------------
------------------------------------------------------------------------
------------------------------------------------------------------------
---------------------------------------------------------------------GAG
UCU------------------------------------GAAAA-------------------------GG-
G-C-GA-----------------------UAAUCAGUGUUUAUA-GACCCGAACCC-UGGUGAUCUAACCAU
GGCCAGGAUGAAGCUUGG-GUAACA-CCAAGUGGAGGUCCGA--ACCGACCG-AUGUUGAAAAAUCG-GCGG
AUGAGUUGUGGUUAGGGGUGAAAUGCCAAUCGAACCAGG-AGCUAGCUGGUUCUCCCCGAAA-UGUGUUGAG
GCGCAGCGGUAA-UGA--------------------------------------------------------
----------------------UU-----AU-AGUCGGGGGGUAAAGC-ACUGUUU-CGG-UGC--GGGCUG
G---GA-GA--CCGGUACCAAAUCGAGACAAACUCAGAAUACCCGA--------------------------
------------------------------------------------------------------------
------------------U-GG-A-CA--------CAUUGCC-AGUGAGA-C-GGUG-G-GGGAUAAGCUUC
AU-C-GUCAAGAG-GGAAAC---AGCCCAGACCACCA-GCUAAGGUCCCCAAAU--CA-UCG-CU-AA---G
-U-GAU-------AAAGGAGGUGAGAUUGCCUAGACAACUAGGAGGUUUGCCUAGAAGCAGCCA-CCCUUGA
AAGAGUGCGUAAUAGCUCACUAGUC-----AAGCGAUCUUGCGCCGAAAAUG-AAC-GGGGCU-A-AGCGAU
G----------UACCGAA-GCUGUGGGAUUAACU--A-----------------------------------
------------------------------------------------------------------------
------------------------------------------------------------------------
--------------------------------AACAU-UAAU-CGGUAGGGGAGC-GUUCCGU---------
-----------AGU-AGGAA-GAAGCAG--UAGCGGCAAGCA--G-CUGUGGACGA-G-ACG-GAAGUGAGA
AUGU-CGGCUU-GAGUA-GCGCAA---ACAU-UGGUGA--G---------------AAUCCA-AUG-CCCCG
A-AACCCU-AAGGGUUCC-AGAGCCAGG-UUCGUCCACUCUGGGUUAGUCGGG-AC-CUAAGGCGAGGCCG-
A-A-AGGC-----GUA--------------------------------------------------------
----------------------------------------GUC-GA-U--GGAC-ACAGGGUC-AACA-AUC
CCUGA----------------------------------CUAUG----GUAU-------GG-GA--------
------------------------------------------------------------------------
---------------------------------------------------------GCAUUG-CUAGGGAC
GCAUGAA-------------AGAUAG-CUACGCCCU-GA-U----UGGUU------------UGGGAGGAG-
UUUA--------------------------------------------------------------------
------------------------------------------------------------------------
------------------------------------------------------------------------
----------------CGAACUCCG-A-GUA-G-UGAAGGAUAGUGU-C-AAGAAA-AGCUA-GUAAUGUGA
-------------------------------------------------------UGAACAU-AU-CGUACC
CGUACCCGAAACCGAC------------------------------ACAGGUA--GGGAGGUU-GAGAAUAC
CAAGGG-G-CG-CGAG-AUAACUCU-C-UCUAAGGAACUCGGC-AAAAUGGCCCCGUAACUUCGG-AAGAAG
```

```
GGGUG-CCCA-C-CG---------------------------------------------------------
------------------------------------------------------------------------
------------------------------------------------------------------------
------------------------------------------------------------------------
------------------------------------------------------------------------
------------------------------------------------------------------------
------------------------------------------------------------------------
------------------------------------------------------------------------
------------------------------------------------------------------------
------------------------------------------------------------------------
------------------------------------------------------------------------
------------------------------------------------------------------------
------------------------------------------------------------------------
------------------------------------------------------------------------
-------------------------------------------------------------------AUGAG
GUG--GGUCGCAGUGAAGAGAUCCAGGCGA-----CUGUUUACCAAAAACACAGGUCUCCGCAAACUCG---
-UA-A-------------GAGGAAGU-AUGGGGGCUGA-CGCCUGCCCAGUGCCG-G-AAGG-UUAAGGAAG
UUGGUCA-GGGC---GAAA---GUUUGAAGCUGACGACCGAAGCCC--CGGUGAAC-GGCGGCCGUAACUAU
AACGGUCCUAAGGUAGCGAAAUUCCUUGUCGGG-UAAGUUCCGACCCGCACGAAAGGCGUAACGAUC--UGG
AUGGUGUCUCAGAG----AGA-GACUCGGCGAAAUA--GGA---AUGUCUGUGAAGAUACGGACUGCCUGCA
CCUGGACAGAAAGA-CCCUAUGAAGC-UUUACU--GUAGCCU-GGAAUUGUGUCCGGGCUUCGCUUGCGCAG
GAU-AGGUGGGAAGCGAUG----------AAGUAUU-CCU----------------------------UGUG
-------------------------------------GG-GAAUAUGGA---G--CUAA-CGGUGAGAUA-C
CACU-CUGGCGAAGCUAGGAUUCUAACUUAUU--UCCG-U-GA-U-C-------------------------
------------------------------------------------------------------------
------------------------------------------------------------------------
------------------------------------------------------------------------
------------------------------------------------------------------------
------------------------------------------------------CG----GA----------
------------------------------------------------AAAA-----GGACA-GUUUCAGGU
GGGCAGUUUGACUGGGGCGGU-CGCCUCCUA-AAAGGUAAC-GGAGGCGCGCA-AAGGUUCCCUCAGCACGC
UUGGAAACCGUGCGGC-GAGUG-UAAAGGCAUAAAGGGAGCUUGACU-GCAA--------------------
--------------GACCGA---CAAGUCGAGCAGGUACGAAA-GUAGGCCUUAGUGAUCCGAC--------
---GG-C-----GCAGA-AUG--------------------GA-AUGGCCGUCGCUCAACGG-AUAAAAGUU
ACUCUAGGG-AUAACAGGCUGAUCUCCCCCAAGAGUCCACAUCGACGGGGAGGUUUGGCACCUCGAUGUCGG
CUCAUCGCAACCUGGGGCGGAA-GUACGUCCCAAGGGUUGGGCUGUUCGCCCAUUAAAGCGGUACGUGAGCU
GGGUUCAGAACGUCGUGAGACAGUUCGGUCCAUAUCCGGU--------------------G-CAG-------
-------------------------------GCGU--AAGAGCAUUGAGAGG-AGC-CUUCCUUAGUACGAG
AGGAC-CGGGAAGGACGCACCGC-UGGU-G-UACCAGUUAUUGUACCCGCAGUAG-ACGCUGGGUAGCCAAG
UGCGGAGCGGAUAACCGCUGAAAGCA-UC--UAAGUGGGAAGCCCACCUCAA-G-AUG--AGUGCUCU-CA-
--CUACA-----------------------------------------------------------------
--------------------------CC--------------------------------------------
------------------------------------------------------------------------
------------------------------------------------------------------------
------------------------------------------------------------------------
```

```
------------------------------------------------------------------------
------------------------------------------------------------------------
------------------------------------------------------------------------
-------------------------------------------------AAGUAGG--UA-AGG------UC
ACGGGCAGAACACC-CGU-UUA-UAGGCU-CUA-AGUGGAAG-UGC-A----------------GU------
----AA-----------UGUAUGAAGCUG--A-GGAG-UACUAACA---GACCGAG-G-GCUUGACCU----
------------------------------------------------------------------------
------------------------------------------------------------------------
------------------------------------------------------------------------
------------------------------------C------------
>LSUb_165597_CROWT/1-2814
---------------------C-----------------AAAGGGCUAAC-GGUGGAUA-C-CUAGGC-ACA
CAG---AGGCGAAGAAGGACGUAG--UUACCGACGAUACGCUCCGGGGAGCUGGAAGCAAGCAU------UG
AGC-CGG-AGAUUUCCCAAUGGGGCAAC--------CC--------CA------------------------
------------------------------------A----GAA-CGGU-CAC-----CUGA----------
-----------A-----------UCUA----------------------------------------UA-GG
G-U--------------------GA-UGCG-AGCGAACUGGGCGAACUGAAACAUCUUAGUAACCC--AAGG
AAGAGAAAGCAA--AAGCGAUUCCCCUA--GU-AG-CGGCGAGCGAAGAGGGA-C-AA-GCCUAAACCAUC-
--GA-CUAAG----------------------------------GUU-GAUGGGG--UUG------------
-----UGGG-ACA-------GUGA-CG----------------------UG--GAC---------UGUA---
G-------CGGUU-AAACG------AAG-CA-GCU-------------------------------------
-------GAA-AA-CUGCA-------CCAGAGAAGGUGAAAGUCCUGUA---------GUUGAAA----ACU
U------A--AGCAG-----------CCU---AACU--GU-AU------CCCGAGUAGCACGGAGCCCGUGG
AAUUCCGUGUGAA-UCAGCGAGG-ACC--ACCUCGUAAGGCUAAAUA--CUC-CUGUG-UGACCGAUAGA-G
AAA-C-AGUACCGCGAGGGAAAG-GUGAAA-AGAAC-CCCGG-AAGGGG-A-GUGAAAUAGAACAUGAAACC
G-UUAGCCUACAA-GCA-AU-GGGA---GGAC-G--------------------------------------
------------------------------------------------------------------------
------------------------------------------------------------------------
------------------------------------------------------------------------
------------------------------------------------------------------------
------------------------------------------------------------------------
------------------------------------------------------------------------
------------------------------------------------------------------------
------------------------------------------------------------------------
-------------------------------------------------------------AU--U-UAA--
------------------------------------------------------------------------
------------------------------------------------------------------------
------------------------------------------------------------------------
------------------------------------------------------------------------
------------------------------------------------------------------------
------------------------------------------------------------------------
------------------------------------------------------------------------
------------------------------------------------------------------------
------------------------------------------------------------------------
------------------------------------------------------------------------
------------------------------------------------------------------------
```

```
------------------------------------------------------------------------
------------------------------------------------------------------------
------------------------------------------------CG--U---CUGACCGUGUGCCUGU
UGAAGAAUGAGCCGGCGACUUAC-A-GGCUGUGGCA--GGUUAAAU--G----------GGAUG--------
--AC--CGAA--GCCAAAGUGAAAG-C---------------------------------------------
------------------------------------------------------------------------
------------------------------------------------------------------------
---------------------------------------------------------------------GAG
CCU------------------------------------GACAA-------------------------GG-
G-C-GA-----------------------UAGUCACAGUUUGUA-GACCCGAACCC-GGGUGAUCUAACCAU
GGCCAGGAUAAAGCUAGG-GUAAUA-CCUUGUGGAGGUCCGA--ACCGACUU-CUGUUGAAAAAGGA-GCGG
AUGAGCUGUGGUUAGGGGUGAAAUGCCAAUCGAACCCGG-AGCUAGCUGGUUCUCCCCGAAA-UGCGUUUAG
GCGCAGCGGUUG-UGU--------------------------------------------------------
----------------------AC-----CU-UAUGGGGGGGUAAAGC-ACUAUUU-CGC-UGC--GGGCUG
C---GA-GA--GCGGUACCAAAGCGAGAUAAACUCAGAAUACCCUG--------------------------
------------------------------------------------------------------------
------------------U--A-A-GC--------AUCAACC-AGUGAGA-C-GGUG-G-GGGAUAAGCUUC
AU-C-GUCGAAAG-GGAAAC---AGCCCAGACCACCA-GCUAAGGUCCCCAAAU--AA-AUA-CU-AA---G
-U-GAU-------AAAGGAGGUGGGAGUGCAUAGACAACCAGGAGGUUUGCCUAGAAGCAGCAA-UCCUUGA
AAGAGUGCGUAAUAGCUCACUGGUC-----AAGCGCUCCUGCGCCGAAAAUG-AAC-GGGGCU-A-AGUAUU
U----------UACCGAA-GCUGUGGACUUAAUU--A-----------------------------------
------------------------------------------------------------------------
-----UUAA---------------------------------------------------------------
---------------------------UUGUUAAUUA-UUAA-UUGUAGGGGAGC-GUUCUGU---------
-----------A-U-AGGGU-GAAGCAC--UAGCGGCAAGCA--G-GUGUGGACAG-U-ACA-GAAGUGAGA
AUGU-CGGCUU-GAGUA-GCGAAA---AUAU-AUGUGA--G---------------AAUCGU-AUA-CCCCG
A-AAUCCU-AAGGGUUCC-UCCGGAAGG-CUCGUCCGCGGAGGGUUAGUCGGG-AC-CUAAGGCGAGGCCG-
A-A-AGGC-----GUA--------------------------------------------------------
----------------------------------------GUC-GA-U--GGAA-ACAAGGUU-AAUA-UUC
CUUGA----------------------------------CUGUG----AUGA--------G-GG--------
------------------------------------------------------------------------
----------------------------------------------------------AGUUU-UUAGGGAC
GCAUGA--------------CGAUUA-GUCAUACCC-UG-A----AUGGA------------UUGGGAGGA-
GACUAC-GGU-CU-----------------------------------------------------------
------------------------------------------------------------------------
------------------------------------------------------------------------
----------------------CUG-C-AUG-A-UGAACAAUAGUGC-C-AAGAAA-AGCUA-AUAAACGU-
-------------------------------------------------------UGAACUC-AU-GACACU
CGUACCCGAAACCGAC------------------------------ACAGGUA--GGAAGGUA-GAGAAUAC
UAAGGG-G-CG-CGAG-AUAACUCU-C-UCUAAGGAACUCGGC-AAAAUGACCCCGUAACUUCGG-GAGAAG
GGGUG-CCAC-C-GA---------------------------------------------------------
------------------------------------------------------------------------
------------------------------------------------------------------------
------------------------------------------------------------------------
------------------------------------------------------------------------
```

```
------------------------------------------------------------------------
------------------------------------------------------------------------
------------------------------------------------------------------------
------------------------------------------------------------------------
------------------------------------------------------------------------
------------------------------------------------------------------------
------------------------------------------------------------------------
------------------------------------------------------------------------
------------------------------------------------------------------------
----------------------------------------------------------------------GA
GGU--GGUCGCAGUGAAUAGGCCCAGGCGA-----CUGUUUACCAAAAACACAGGUCUCUGCCAAGUCG---
-UA-A-------------GACGCAGU-AUAGGGGCUGA-CGCCUGCCCAGUGCCG-G-AAGG-UUAAGGAAG
UUGGUUA-GU-----UUCG-----ACGAAGCUAGCGACCGAAGCCC--CGGUGAAC-GGCGGCCGUAACUAU
AACGGUCCUAAGGUAGCGAAAUUCCUUGUCGGG-UAAGUUCCGACCCGCACGAAAGGCGUAACGAUC--UGG
GCACUGUCUCGGAG----AGA-GGCUCGGCGAAAUA--GGA---UUGUCUGUGAAGAUACGGACUACCUGCA
CCUGGACAGAAAGA-CCCUAUGAAGC-UUUACU--GUAGCUU-GGAAUUGGGUUCGGGCUUCCUUUGCGCAG
CAU-AGGUGGGAGGCGUUG----------AAGUUUU-CCU----------------------------CGUG
--------------------------------------GG-GAGAAUGGAG-C--C-AA-CGGUGAGAUA-C
CACU-CUAGGGAGGCUAGAAUUCUAACUUAAG--CCCG-U-AA-U-C-------------------------
------------------------------------------------------------------------
------------------------------------------------------------------------
------------------------------------------------------------------------
------------------------------------------------------------------------
------------------------------------------------------CG----GG----------
------------------------------------------------UUAA-----GAACA-GUUUCAGGU
GGGCAGUUUGACUGGGGCGGU-CGCCUCCUA-AAAGGUAAC-GGAGGCGCGCA-AAGGUUCCCUCAGGCUGG
UUAGAAAUCAGCCGCA-GAGUG-CAAAGGCAG-AAGGGAGCUUGACU-GCGA--------------------
--------------GACAAA---CAAGUCAAGCAGGGACGAAA-GUCGGCCUUAGUGAUCCGAC--------
---GG-C-----ACCGA-GUG--------------------GA-AGGGCCGUCGCUCAACGG-AUAAAAGUU
ACUCUAGGG-AUAACAGGCUGAUCUCCCCCAAGAGUUCAUAUCGACGGGGAGGUUUGGCACCUCGAUGUCGG
CUCAUCGCAGCCUGGGGCUGUA-GUAGGUCCCAAGGGUUGGGCUGUUCGCCCAUUAAAGCGGUACGUGAGCU
GGGUUCAGAACGUCGUGAGACAGUUCGGUCCAUAUCCGGU--------------------G-UAG-------
-------------------------------GCGU--AAGAGCAUUGAGAGG-AGC-CUUCCUUAGUACGAG
AGGAC-CGGGAAGGACGCACCGC-UGGU-G-UACCUGUUAUCGUGCCAACGGUAA-ACGCAGGGUAGCCAAG
UGCGGAGCGGAUAACCGCUGAAAGCA-UC--UAAGUGGGAAGCCCACCUCAA-G-AUG--AUUGCUCU-CA-
--CAGGG-----------------------------------------------------------------
--------------------------UU--------------------------------------------
------------------------------------------------------------------------
------------------------------------------------------------------------
------------------------------------------------------------------------
------------------------------------------------------------------------
------------------------------------------------------------------------
------------------------------------------------------------------------
-------------------------------------------------AACCUGG--UA-AGG------UC
ACGGGAAGACCACC-CGU-UAA-UAGGCA-CUC-GGUGGAAG-UCC-A----------------GC------
```

```
----AA-----------UGGAUGAAGCCA--A-GGUG-UACUAACA---GACCGAG-G-GCUUGACCU----
------------------------------------------------------------------------
------------------------------------------------------------------------
------------------------------------------------------------------------
-------------------------------------------------
>LSUb_1148_SYNYG/1-2883
---------------------G-GUCAAGCU----AC--AAAGGGCUAAC-GGUGGAUA-C-CUUGGC-ACA
CAG---AGGCGAAGAAGGACGUGG--UUACCGACGAAACGCUUCGGGGAGCUGGAAACAAGCAU------UG
AUC-CGG-AGGUGUCCGAAUGGGGCAAC--------CC--------AA------------------------
--G---------------------------------U----GAA-CUAC-CCG-----CUGAA---------
-----------U-----------AAA----------------------------------------AUAG-G
CGG--------------------GA-G-AG-AGCAAACCUAGUGAACUGAAACAUCUUAGUAGCUA--GAGG
AAAAGAAAACAA--AAGUGAUUCUCCAA--GU-AG-CGGCGAGCGAACGGGGA-A-CA-GCCUAAACCAAU-
--CU-UUUCG----------------------------------AAG-AUUGGGG--UCG------------
-----UGGG-ACA-------GCAA-CG----------------------UG--GAC---------UGAG---
U-------AGAUU-AGACG------AAG-CA-GCU-------------------------------------
-------GAA-AA-CUGCA-------CCAGAGGAGGUGAAAGUCCUGUA---------GUCGAAA----AUU
G------A--AUCAG-----------CCU---AGCU--GU-AU------CCCGAGUAGGUCGGAGCUCGUGG
AAUUCCGAUUGAA-UCAGCCGGG-ACC--ACCCGGUAAGGCUAAAUA--CUA-CUGUG-UGACCGAUAGC-G
UA--A-AGUACCGCGAGGGAAAG-GUGAAA-AGAAC-CCCGUUGAGGGG-A-GUGAAAUAGAACAUGAAACC
G-UUAGCUACAA-GCA-AU-GGGA---GCUC-G---------------------------------------
------------------------------------------------------------------------
------------------------------------------------------------------------
------------------------------------------------------------------------
------------------------------------------------------------------------
------------------------------------------------------------------------
------------------------------------------------------------------------
------------------------------------------------------------------------
------------------------------------------------------------------------
-------------------------------------------------------------AU--U-AAA--
------------------------------------------------------------------------
------------------------------------------------------------------------
------------------------------------------------------------------------
------------------------------------------------------------------------
------------------------------------------------------------------------
------------------------------------------------------------------------
------------------------------------------------------------------------
------------------------------------------------------------------------
------------------------------------------------------------------------
------------------------------------------------------------------------
------------------------------------------------------------------------
------------------------------------------------------------------------
------------------------------------------------CG--A---GUGACCGUGUGCCUGU
UGAAGAAUGAGCCGGCGACUUAC-A-GGUUGUGGCA--GGUUAAGG--U-G--------UGUUG--------
C-AC--CGAA--GCCAAAGUGAAAG-C---------------------------------------------
```

```
------------------------------------------------------------------------
------------------------------------------------------------------------
---------------------------------------------------------------------GAG
CCU------------------------------------GAAUA-------------------------GG-
G-C-GU-----------------------UAGUCACAAUUUGUA-GACCCGAACCC-GGGUGAUCUAACCAU
GGCCAGGAUGAAGCUUAG-GUAACA-CUAAGUGGAGGUCCGA--ACCGACUA-AUGUUGAAAAAUUA-GCGG
AUGAGCUGUGGUUAGGGGUGAAAUGCCAAUCGAACCCGG-AGCUAGCUGGUUCUCCCCGAAA-UGUGUUUAG
GCGCAGCGGUUG-UAAA--------------------------------------------------------
----------------------GU-----CA-ACUUGGGGGGUAAAGC-ACUGUUU-CGC-UGC--GGGCGG
C---GA-GA--GCUGUACCAAAGUGAGGCAAACUCAGAAUACCCAA--------------------------
------------------------------------------------------------------------
------------------G--A-A-GA--------AACGACC-AGUAAGA-C-GGUG-G-GGGAUAAGCUUC
AU-C-GUCGAAAG-GGAAAC---AGCCCAGACCACCA-GCUAAGGUCCCCAAAU--CA-UCA-CU-AA---G
-U-GAG-------AAAGGAGGUGGGAGUGCAUUGACAACCAGGAGGUUUGCCUAGAAGCAGCAA-UCCUUAA
AAGAGUGCGUAAUAGCUCACUGGUC-----AAGCGCUCCUGCGCCGAAAAUG-AAC-GGGGCU-A-AGUGAU
G----------UACCGAA-GCUGUGGACUCAGA---------------------------------------
------------------------------------------------------------------------
------------------------------------------------------------------------
-----------------------------------UA-UGAG-UGGUAGGGGAGC-GUUCUGU---------
-----------A-U-AGGGU-GAAGCAC--UAGCGGCAAGCA--G-GUGUGGACAG-U-ACA-GAAGUGAGA
AUGU-CGGCUU-GAGUA-GCGAAA---AUAU-GGGUGA--G---------------AAUCCC-AUA-CCCCG
A-AAUCCU-AAGGGUUCC-UCCGGAAGG-CUCGUCCGCGGAGGGUUAGUCAGG-AC-CUAAGGCGAGGCCG-
A-G-AGGC-----GUA--------------------------------------------------------
----------------------------------------GUC-GA-U--GGAC-AACCGGUC-AAUA-UUC
CGGUA----------------------------------CUGAU----UAUA--------G-AU--------
------------------------------------------------------------------------
----------------------------------------------------------UGUGG-CGGGGGAC
GGAGAA--------------GGCUAG-UGCAACUGG-AA-G----UUGGU------------UACCAGUCC-
--AAGC-AUU-CGA----G-G-------------------CG--A--UG-----------------AGA-GA
C---GG--------------------CGAAA-ACGU--CUUG----------AGCAG----AGG-UG-U-G-
AUG--GG---------------------------------------------------------------GA
------GUCUC-UACGGAGACGAAG-U-GCA-U-GAUGUCAAGCUUC-C-AAGAAA-AGCCC-GAACCACGU
-------------------------------------------------------UAAUUUA-UA-GUCACC
UGUACCCGAAACCGAC------------------------------ACAGGUA--GGAAGGUA-GAGAAUAC
UAAGGG-G-CG-CGAG-GUAACUCU-C-UCUAAGGAACUCGGC-AAAAUUACCCCGUAACUUCGG-GAGAAG
GGGUG-CCUC-C-AG---------------------------------------------------------
------------------------------------------------------------------------
------------------------------------------------------------------------
------------------------------------------------------------------------
------------------------------------------------------------------------
------------------------------------------------------------------------
------------------------------------------------------------------------
------------------------------------------------------------------------
------------------------------------------------------------------------
------------------------------------------------------------------------
```

```
------------------------------------------------------------------------
------------------------------------------------------------------------
------------------------------------------------------------------------
------------------------------------------------------------------------
--------------------------------------------------------------------CAAU
GGA--GGUCGCAGUGAAUAGGCCCAGGCGA-----CUGUUUACCAAAAACACAGGUCUCCGCAAAGUCG---
-UA-A-------------GACGCAGU-AUGGGGGCUGA-CGCCUGCCCAGUGCCG-G-AAGG-UUAAGGAAG
UUGGUCA-GGGA---GUAA---UCCUGAAGCUAGCGACUGAAGCCC--CGGUGAAC-GGCGGCCGUAACUAU
AACGGUCCUAAGGUAGCGAAAUUCCUUGUCGGG-UAAGUUCCGACCCGCACGAAAGGCGUAACGAUC--UGG
GCACUGUCUCGGAG----AGA-GGCUCGGCGAAAUA--GGA---UUGUCUGUGAAGAUACGGACUCCCUGCA
CCUGGACAGAAAGA-CCCUAUGAAGC-UUUACU--GUAGCUU-GGAAUUGGGUUCGGGCUUUGCUUGCGCAG
GAU-AGGUGGGAGACUAUG-----------AAGUUGC-UCU----------------------------UGUG
--------------------------------------GG-AGUAAUGGAG-U--C-AA-CGGUGAGAUA-C
CACU-CUGGUGAGGCUAGAAUUCUAACUUGUU--CCCG-U-UA-U-C-------------------------
------------------------------------------------------------------------
------------------------------------------------------------------------
------------------------------------------------------------------------
------------------------------------------------------------------------
------------------------------------------------------CG----GG----------
------------------------------------------------AGAA-----GGACA-AUUUCAGGU
GGGCAGUUUGACUGGGGCGGU-CGCCUCCUA-AAAGGUAAC-GGAGGCGCGCA-AAGGUUUCCUCAGGCUGG
UUGGAAAUCAGCCGAA-GAGUG-UAAAAGCAG-AAGGAAGCUUGACU-GUGA--------------------
--------------GACUGA---CAAGUCAAACAGGGACGAAA-GUCGGCUUUAGUGAUCCGAC--------
---GG-C-----ACUGA-GUG--------------------GA-AGGGCCGUCGCUCAACGG-AUAAAAGUU
ACUCUAGGG-AUAACAGGCUGAUCUCCGCCAAGAGUUCACAUCGACGCGGAGGUUUGGCACCUCGAUGUCGG
CUCAUCGCAACCUGGGGCGGUA-GUACGUCCCAAGGGUUGGGCUGUUCGCCCAUUAAAGCGGUACGUGAGCU
GGGUUCAGAACGUCGUGAGACAGUUCGGUCCAUAUCCGGU--------------------G-UAG-------
-------------------------------GCGU--AAGAGUAUUGAGAGG-AGA-CUUCCUUAGUACGAG
AGGAC-CGGGAAGUACGCACCUC-UGGU-G-UACCUGUUAUCGUGCCAACGGUAA-ACGCAGGGUAGCCAAG
UGCGGAGUGGAUAACCGCUGAAAGCA-UC--UAAGUGGGAAGCCCACCUCAA-G-AUG--AGUACUCU-CA-
--UGGUG-----------------------------------------------------------------
--------------------------UU--------------------------------------------
------------------------------------------------------------------------
------------------------------------------------------------------------
------------------------------------------------------------------------
------------------------------------------------------------------------
------------------------------------------------------------------------
------------------------------------------------------------------------
-------------------------------------------------AAGCCAG--UA-AGG------UC
ACGGGAAGACUACCC-GU-UGA-UAGGCU-CUA-GAUGGAAG-UCC-A----------------GU------
----AA-----------UGGAUGAAGUCG--A-GGAG-UACUAAUA---GACCGAG-G-GCUUGACCU----
------------------------------------------------------------------------
------------------------------------------------------------------------
------------------------------------------------------------------------
-------------------------------------------------
```

>LSUb_197221_THEEB/1-2869
---------------------G-GUCAAGCU----AC--AAAGAGCCUAC-GGUGGAUA-C-CUAGGC-ACA
CAG---AGGCGAUGAAGGACGUGG--AUACCGACGAUACGCUUCGGGGAGCUGGAAACAAGCUU------UG
AUC-CGG-AGGUCUCCGAAUGGGGCAAC--------CC--------AG------------------------
------------------------------------U----A-A-CUCU-AAC-----GCC-----------
-----------A-----------AUCC-----------------------------------------AU-A
A-U--------------------UA-G-AG-GGCGAACCUGGCGAACUGAAACAUCUUAGUAGCCA--GAGG
AAAAGAAAGAAA--ACUCGAUUCCCUUA--GU-AG-CGGCGAGCGAAAGGGGA-A-CA-GCCUAAACCAAC-
--GG-GCUUG----------------------------------CCC-GUUGGGG--UCG------------
-----UGGG-ACA-------GCAA-CA----------------------UG--GAA---------UUGG---
G-------CGGCU-AGACG------AAG-UA-GUG--------------------------------------
-------GAA-UG-CUACA-------CCAGAGAAGGUGAAAGUCCUGUA---------GUCGAAA----GCU
U------A--ACAAU-----------CCU---AGCU--GU-AU------CCCGAGUAGUCCGGAGCACGUGG
AAUUCCGGAUGAA-UCUGCGAGG-ACC--ACCUCGUAAGGCUAAAUA--CUC-CUGUG-UGACCGAUAGC-G
CA--U-AGUACCGCGAGGGAAAG-GUGAAA-AGAAC-CCCGG-AAGGGG-A-GUGAAAUAGAGCAUGAAACC
G-UAGGCUUACAA-GCA-GU-CGAA---GCUC-G--------------------------------------
------------------------------------------------------------------------
------------------------------------------------------------------------
------------------------------------------------------------------------
------------------------------------------------------------------------
------------------------------------------------------------------------
------------------------------------------------------------------------
------------------------------------------------------------------------
------------------------------------------------------------------------
-------------------------------------------------------------AU--U-UAA--
------------------------------------------------------------------------
------------------------------------------------------------------------
------------------------------------------------------------------------
------------------------------------------------------------------------
------------------------------------------------------------------------
------------------------------------------------------------------------
------------------------------------------------------------------------
------------------------------------------------------------------------
------------------------------------------------------------------------
------------------------------------------------------------------------
------------------------------------------------------------------------
------------------------------------------------------------------------
------------------------------------------------CG--A---GUGACGGCGUGCCUGU
UGAAGAAUGAGCCGGCGACUUAU-A-GGUACUGGCA--GGUUAAGG--C-A---------AAAA--------
U-GC--CGAA--GCCAUAGCGAAAG-C---------------------------------------------
------------------------------------------------------------------------
------------------------------------------------------------------------
---------------------------------------------------------------------GAG
UCU------------------------------------GAAUA-------------------------GG-
G-C-GU-----------------------UUGUCAGUACUUAUA-GACCCGAACCC-GGGUGAUCUAACCAU

```
GGCCAGAAUGAAGCUUGG-GUAACA-CCAAGUGGAGGUUCGC--ACCGACCG-AUGUUGAAAAAUCG-GCGG
AUGAGCUGUGGUUAGGGGUGAAAUGCCAAUCGAACCCGG-AGCUAGCUGGUUCUCCCCGAAA-UAUGUUGAG
GCAUAGCGGUUG-UGA--------------------------------------------------------
----------------------UU-----AU-AGCAGGGGGGUAAAGC-ACUGAUU-CGG-UGC--GGGCUG
C---GA-GA--GCGGUACCAAAUCGAGUCAAACUCUGAAUACCCUG--------------------------
------------------------------------------------------------------------
------------------U-GU-A-CA--------CACAGCC-AGUCAGA-C-GGUG-G-GGGAUAAGCUUC
AU-C-GUCGAGAG-GGAAAC---AGCCCAGACCACCA-GCUAAGGUCCCCAAAU--CA-UCA-CU-AA---G
-U-GGC-------AAAGGAGGUGGGGAUGCUCAGACAACCAGGAGGUUUGCCUAGAAGCAGCCA-UCCUUAA
AAGAGUGCGUAAUAGCUCACUGGUC-----AAGCGUUCCUGCGCCGAAAAUG-AAC-GGGACU-A-AGUGAU
G----------UACCGAA-GCUGUGGACUUACUA--------------------------------------
------------------------------------------------------------------------
------------------------------------------------------------------------
-----------------------------------CG-UAAG-UGGUAGGGGAGC-GUUCUGC---------
-----------ACU-AGUUU-GAAGCAC--UAGC-GUAAGCA--G-GUGUGGACGA-A-GCA-GAAGUGAGA
AUGC-CGGCUU-GAGUA-GCGAAA---ACAU-UGGUGA--G---------------AAUCCA-AUG-CCCCG
A-AACCCC-AAGGGUUCC-UUCGGAAGG-CUCGUCCGCGAAGGGUUAGUCGGG-AC-CUAAGGCGAGGCCG-
A-A-AGGC-----GUA--------------------------------------------------------
----------------------------------------GUC-GA-U--GGAC-AACUGGUC-AAUA-UUC
CAGUA----------------------------------CUCAU----CUUG--------G-UU--------
------------------------------------------------------------------------
----------------------------------------------------------UGUGC-AGAGGGAC
GAAGAA--------------GGCUAG-AUCAGCCGG-AU-G----UUGGU------------UACCGGUUC-
--AACU-GUC-CGA----G-G-------------------UG--U--UG-----------------AGG-AG
C---GG--------------------UGAAA-ACGC--UCUG----------AGCUG----AGG-CA-G-G-
AGU--AC---------------------------------------------------------------GA
-----CCCAC--UACGGUGGGGAAG-U-GAU-U-GAUGUCAUGCUUC-C-AAGAAA-AGCUC-UAAGCACG-
-------------------------------------------------------UUAAGCC-AA-GAUACC
CGUACCCGAAACCGAC------------------------------ACAGGUG--GGGGGGUA-GAGUAUAC
CAAGGG-G-CG-CGAG-AUAACUCU-C-UCUAAGGAACUCGGC-AAAAUGGCCCCGUAACUUCGG-GAGAAG
GGGUG-CCAC-C-GA---------------------------------------------------------
------------------------------------------------------------------------
------------------------------------------------------------------------
------------------------------------------------------------------------
------------------------------------------------------------------------
------------------------------------------------------------------------
------------------------------------------------------------------------
------------------------------------------------------------------------
------------------------------------------------------------------------
------------------------------------------------------------------------
------------------------------------------------------------------------
------------------------------------------------------------------------
------------------------------------------------------------------------
------------------------------------------------------------------------
----------------------------------------------------------------------GA
```

```
GGU--GGUCGCAGUGAAGAGGCCCAGGCGA-----CUGUUUACCAAAAACACAGGUCUCCGCUAAGUCG---
-UA-A-------------GACGAUGU-AUGGGGGCUGA-CGCCUGCCCAGUGCCG-G-AAGG-UUAAGGAAG
CCGGUCA-GC-----GCAA-----GCGAAGCUGGCGACCGAAGCCC--CGGUGAAC-GGCGGCCGUAACUAU
AACGGUCCUAAGGUAGCGAAAUUCCUUGUCGGG-UAAGUUCCGACCCGCACGAAAGGCGUAACGAUC--UGG
GCGCUGUCUCGGAG----AGA-GGCUCGGCGAAAUA--GAA---GUGUCUGUGAAGAUGCGGACUACCUGCA
CCCGGACAGAAAGA-CCCUAUGAAGC-UUUACU--GUAGCCU-GGAAUUGGGUUCGGGCUUUGCUUGCGCAG
GAU-AGGUGGGAGACGUUG----------AAGUAUC-CCU----------------------------UGCG
--------------------------------------GG-GGAUAUGGAG-U--C-GC-CAGUGAGAUA-C
CACU-CUGGCAAGGCUAGAAUUCUAACCUUCU--GCUG-U-UA-G-C-------------------------
------------------------------------------------------------------------
------------------------------------------------------------------------
------------------------------------------------------------------------
------------------------------------------------------------------------
------------------------------------------------------CA----GC----------
------------------------------------------------GAAG-----GAACA-GUUUCAGGU
GGGCAGUUUGACUGGGGCGGU-CGCCUCCUA-AAUGGUAAC-GGAGGCGCGCA-AAGGUUCCCUCAGGCUGG
UCGGAAAUCAGCCGUA-GAGUG-CAAAGGCAU-AAGGGAGCUUGACU-GCAA--------------------
--------------GACCUA---CAAGUCGAGCAGAGACGAAA-GUCGGCCUUAGUGAUCCGAC--------
---GG-U-----UCCGC-GUG--------------------GA-AGGGCCGUCGCUCAACGG-AUAAAAGUU
ACUCUAGGG-AUAACAGGCUGAUCUCCCCCAAGAGUUCACAUCGACGGGGAGGUUUGGCACCUCGAUGUCGG
CUCAUCGCAACCUGGGGCGGUA-GUACGUCCCAAGGGUUGGGCUGUUCGCCCAUUAAAGCGGUACGUGAGCU
GGGUUCAGAACGUCGUGAGACAGUUCGGUCCAUAUCCGGU--------------------G-CAG-------
-------------------------------GCGU--AGGAGCAUUGAGAGG-AGU-CCUCCUUAGUACGAG
AGGAC-CGGGAGGAACGCACCUC-UGGU-G-UACCUGUUAUCGUGCCAACGGUAA-ACGCAGGGUAGCCAUG
UGCGGAGUGGAUAACCGCUGAAAGCA-UC--UAAGUGGGAAGCCCACCUCAA-G-AUG--AGUGCUCC-CA-
--UGGCA-----------------------------------------------------------------
--------------------------U---------------------------------------------
------------------------------------------------------------------------
------------------------------------------------------------------------
------------------------------------------------------------------------
------------------------------------------------------------------------
------------------------------------------------------------------------
------------------------------------------------------------------------
-------------------------------------------------AAGCCAG--UA-AGG------UC
ACGGGUAGAACACCC-GU-UAA-UAGGCU-CCC-GAUGGAAG-UCC-A----------------GU------
----AA-----------UGGAUGAAGUCU--A-GGAG-UACUAAUA---GACCGAG-G-GCUUGACCU----
------------------------------------------------------------------------
------------------------------------------------------------------------
------------------------------------------------------------------------
------------------------------------C------------
>LSUb_91464_SYNS7/1-2882
------------------------UCAAGCU----AC--AAAGGGCUCAC-GGUGGAUA-C-CUAGGC-ACA
CAG---AGGCGAAGAAGGACGUGG--UUACCGACGAUACGCUUUGGGGAGCCGGAAGCAGGCAU------UG
AUC-CAG-AGAUUUCCGAAUGGGGCAAC--------CC--------UA------------------------
------------------------------------G----AAA-CAAC-UUA-----CUGA----------
```

```
-----------A-----------UCUA----------------------------------------UAG-G
U-A--------------------AG-UAUG-AGCGAACGCAGUGAACUGAAACAUCUAAGUAGCUG--UAGG
AAGAGAAAGAAA--ACUCGAUUCCCUUA--GU-AG-CGGCGAGCGAAGCGGGA-A-GA-GCCCAAACCACC-
--GG-UUUUU----------------------------------ACU-GGUGGGG--UUG------------
-----UGGG-AGC-------GCGA-UA----------------------UG--GAA---------UCUA---
G-------CGAGU-AGACG------AAU-CA-CUU-------------------------------------
-------AAG-AG-GUGAA-------CCGUAGAGGGUGAGAGUCCGUA----------GUUGAAA----CUU
U------A--AGGAU-----------ACU---AGCG--UU-AU------CCCGAGUAGGCCGGAACACGAGA
AAUUCCGGUUGAA-UCCACGAGG-ACC--ACCUCGUAAGGCUAAAUA--CUC-CUGUG-UGACCGAUAGU-G
AA--G-AGUACCGCGAGGGAAAG-GUGAAA-AGAAC-CCCGG-GAGGGG-A-GUGAAAUAGAACAUGAAACC
G-UGAGCUUACAA-GCA-AU-UAGA---GCAC-G--------------------------------------
------------------------------------------------------------------------
------------------------------------------------------------------------
------------------------------------------------------------------------
------------------------------------------------------------------------
------------------------------------------------------------------------
------------------------------------------------------------------------
------------------------------------------------------------------------
------------------------------------------------------------------------
-------------------------------------------------------------AU--U-AAA--
------------------------------------------------------------------------
------------------------------------------------------------------------
------------------------------------------------------------------------
------------------------------------------------------------------------
------------------------------------------------------------------------
------------------------------------------------------------------------
------------------------------------------------------------------------
------------------------------------------------------------------------
------------------------------------------------------------------------
------------------------------------------------------------------------
------------------------------------------------------------------------
------------------------------------------------------------------------
------------------------------------------------CG--U---GUGAUAGUGUGCCUGU
UGAAGAAUGAGCCGGCGACUUAU-A-GGGUCUGGCA--GGUUAAGG--A-G--------AAUAU--------
A-UC--CGAA--GCCAAAGGGAAAC-C---------------------------------------------
------------------------------------------------------------------------
------------------------------------------------------------------------
---------------------------------------------------------------------GAG
UCU------------------------------------UAAUA-------------------------GG-
G-C-GA----------------------GUCGUCAGAUUUAUA-GACCCGAACCC-AGGUGAUCUAACCAU
GUCCAGGAUGAAGCUUAG-GUAACA-CUAAGUGGAGGUCCGA--ACCGACUU-CUGUUGAAAAAGGA-GCGG
AUGAGGUGUGGUUAGGGGUGAAAUGCCAAUCGAACCUGG-AGCUAGCUGGUUCUCCUCGAAA-UGUGUUUAG
GCGCAGCGGUCA-UGA--------------------------------------------------------
----------------------UU-----AU-AGUCGGGGGGUAAAGC-ACUGAUU-CGG-UGC--GGGCUG
C---GA-GA--GCGGUACCAAAUCGAGUCAAACUCAGAAUACCCGA--------------------------
```

```
------------------------------------------------------------------------
------------------U-GA-A-CA--------CAUGGCC-AGUAAGA-C-GGUG-G-GGGAUAAGCUUC
AU-C-GUCAAGAG-GGAAAC---AGCCCAGACCACCA-GCUAAGGUCCCCAAGU--CA-AUA-CU-AA---G
-U-GGC-------AAAGGAGGUGGGAAUGCAGAGACAACCAGGAGGUUUGCCUAGAAGCAGCCA-UCCUUGA
AAGAGUGCGUAAUAGCUCACUGGUC-----AAGCGUUCCUGCGCCGAAAAUG-AAU-GGGACU-A-AGUAUU
G----------CACCGAA-GCUGUGGGCUUGUAU--U-----------------------------------
------------------------------------------------------------------------
------------------------------------------------------------------------
---------------------------------UAUA-CAAG-CGGUAGAGGAGC-GUUCUGU---------
-----------GGU-AGUGC-GAAGCAU--UAGC-GAGAGCA--G-GUGUGGACGA-A-GCA-GAAGUGAGA
AUGU-CGGCUU-GAGUA-GCGAAA---ACUA-GGGUGA--G---------------AAUCCC-UAG-CCCCG
A-AAUCCU-AAGGAUUCC-UCCGGAAGG-CUCGUCCGCGGAGGGUUAGUCGGG-AC-CUAAGGCGAGGCCG-
A-C-AGGC-----GUA--------------------------------------------------------
----------------------------------------GUC-GA-U--GGUU-AACAGGUU-AAUA-UUC
CUGUA----------------------------------CCGAU----UGUG--------A-AU--------
------------------------------------------------------------------------
----------------------------------------------------------UGUGC-AGAGGGAC
GGAGAA--------------GGCUAA-GUGUGCCGC-AA-G----UUGGU------------UAGCGGUUU-
--AAGU-AUU-CGA----G-G-------------------UG--A--UG-----------------AGG-AA
C---GG--------------------CGAAA-ACGU--UCUG----------AGCUG----AGG-UG-C-G-
AAU--AG---------------------------------------------------------------GA
------GUUAC-UACGGUAACGAAG-A-CAC-U-GAUGUCCAGCUUC-C-AAGAAA-AGCUC-UAACCACGU
-------------------------------------------------------UAAUUCA-UA-AUUGCC
CGUACCCUAAACCGAC------------------------------ACAGGUA--GGAAGGUU-GAGUAUAC
CCAGGG-G-CG-CGAG-AUAACUCU-C-UCUAAGGAACUCGGC-AAAAUGGCCCCGUAACUUCGG-GAGAAG
GGGUG-CCAG-C-GA---------------------------------------------------------
------------------------------------------------------------------------
------------------------------------------------------------------------
------------------------------------------------------------------------
------------------------------------------------------------------------
------------------------------------------------------------------------
------------------------------------------------------------------------
------------------------------------------------------------------------
------------------------------------------------------------------------
------------------------------------------------------------------------
------------------------------------------------------------------------
------------------------------------------------------------------------
------------------------------------------------------------------------
------------------------------------------------------------------------
----------------------------------------------------------------------GA
GCU--GGCCGCAGUGAAGAGAUCCAGGCGA-----CUGUUUACCAAAAACACAGGUUUCCGCUAAGUCG---
-AA-A-------------GACGAUGU-AUGGGAGCUGA-CGCCUGCCAGUGCCG-G-AAGG-UUAAGGAAG
UAGGUCA-GUCU---AUUC---GUAGGAAGCUCACGACUGAAGCCC--CGGUGAAC-GGCGGCCGUAACUAU
AACGGUCCUAAGGUAGCGAAAUUCCUUGUCGGG-UAAGUUCCGACCCGCACGAAAGGCGUAACGAUC--UGG
AUGCUGUCUCGGAG----AGA-GACUCGGCGAAAUA--GGA---GUGUCUGUGAAGAUACGGACUACCUGCA
```

```
CCCGGACAGAAAGA-CCCUAUGAAGC-UUUACU--AUAGCCU-GGUAUUGGGUUCGGGCCCUGAUUGCGCAG
GAU-AGGUGGGAGACUAUG----------AUCUACU-CCU----------------------------UGUG
--------------------------------------GG-GAGUAGUGAG-U--C-AC-UGGUGAGAUA-C
CACU-CUAUCAGGGCUAGGAUUCUAACUUUGA--GCCA-U-CA-A-C-------------------------
------------------------------------------------------------------------
------------------------------------------------------------------------
------------------------------------------------------------------------
------------------------------------------------------------------------
------------------------------------------------------UG----GC----------
------------------------------------------------CAAG-----AGACA-GUAUCAGGU
GGGUAGUUUGACUGGGGCGGU-CGCCUCCUA-AAAGGUAAC-GGAGGCGCGCA-AAGGUUCCCUCAGGCUGG
UUGGAAAUCAGCCGCA-GAGUG-UAAAGGCAG-AAGGGAGCUUGACU-GCGA--------------------
--------------GAGCUA---CAACUCGAGCAGGUACGAAA-GUAGGCCUUAGUGAUCCGAC--------
---GG-U-----UCCGC-GUG--------------------GA-AGGGCCGUCGCUCAACGG-AUAAAAGUU
ACUCUAGGG-AUAACAGGCUGAUCUCCCCCAAGAGUUCACAUCGACGGGGAGGUUUGGCACCUCGAUGUCGG
CUCAUCGCAACCUGGGGCGGUA-GUACGUCCCAAGGGUUGGGCUGUUCGCCCAUUAAAGCGGUACGUGAGCU
GGGUUCAGAACGUCGUGAGACAGUUCGGUCCAUAUCCGGU--------------------G-CAG-------
-------------------------------GCGU--AAGAACAUUGAGAGG-AGC-CUUCCUUAGUACGAG
AGGAC-CGGGAAGGACGCACCGC-UGGU-G-UACCAGUUAUCGUGCCAACGGUAA-ACGCUGGGUAGCCAAG
UGCGGAGUGGAUAACCGCUGAAAGCA-UC--UAAGUGGGAAGCCACCUCAA-G-AUG--AGUGUUCU-CA-
--UGGAG-----------------------------------------------------------------
--------------------------UU--------------------------------------------
------------------------------------------------------------------------
------------------------------------------------------------------------
------------------------------------------------------------------------
------------------------------------------------------------------------
------------------------------------------------------------------------
------------------------------------------------------------------------
-------------------------------------------------AAUCCAG--UA-AGG------UC
ACGGGAAGAAGACCC-GU-UGA-UAGGUU-CUA-GGUGGAAG-UAC-A----------------GU------
----AA-----------UGUAUGUAGCCG--A-GGAA-UACUAACA---GACCGAG-G-GCUUGUCCU----
------------------------------------------------------------------------
------------------------------------------------------------------------
------------------------------------------------------------------------
------------------------------------U------------
>LSUb_251221_GLOVI/1-2781
------------------------UCAAGCU----AC--ACAGGGCGUAC-GGUGGAUA-C-CUAGGC-ACA
GUC---AGCCGAUGAAGGACGCGG--CUACCGGCGAAACGCGACGGGGAGCAGGAAGCAUGCUU------UG
AUC-CGU-CGGUGUCCGAAUGGGAAAC---------CC--------GG----CAUC-C-U----------UA
A-C-------------A---G----G----G---UGU----C-G-UCC------------------------
-----------G-----------CAA----------------------------------------------
------------------------G-G-AC--GGGAACCAGGCGAAUUGAAACAUCUUAGUAGCCU--GAGG
AAAAUAAAGUAA--AAACGAUUCCCUCA--GU-AG-CGGCGAGCGAACGGGGA-A-GA-GCCCAAACCGAU-
--GG-GUUAC-------------------------------GA-CCU-GUCGGGG--UUG------------
-----UAGG-ACA-------GCUA-UC----------------------UG--GAG---------UGCG---
```

```
U-------UUGCU-AGGCG------AAU-GG-GUUG------------------------------------
-------GAA-AA-CCCGA-------CCAAAGAAGGUGAUAGUCCUGUA---------CUCGAAA----GCA
A------A--CAAAC-----------CCU---GGCU--GA-AU------CCUGAGUAGCACGGGGCACGUGA
AAUCCCGUGUGAA-UCUGCGAGG-ACC--AUCUCGUAAGGCUAAAUA--CUG-UC-UG-UGACCGAUAGC-G
-AU-U-AGUACCGCGAGGGAAAG-GUGAAA-AGAAC-CCCGG-GAGGGG-A-GUGAAAUAGAACAUGAAACC
G-UAUGCCUACAA-GCA-GU-GGGA---GCAC-G--------------------------------------
------------------------------------------------------------------------
------------------------------------------------------------------------
------------------------------------------------------------------------
------------------------------------------------------------------------
------------------------------------------------------------------------
------------------------------------------------------------------------
------------------------------------------------------------------------
------------------------------------------------------------------------
-------------------------------------------------------------AU--U-UAA--
------------------------------------------------------------------------
------------------------------------------------------------------------
------------------------------------------------------------------------
------------------------------------------------------------------------
------------------------------------------------------------------------
------------------------------------------------------------------------
------------------------------------------------------------------------
------------------------------------------------------------------------
------------------------------------------------------------------------
------------------------------------------------------------------------
------------------------------------------------------------------------
------------------------------------------------------------------------
------------------------------------------------CG--U---GUGACCGCGUGCCUGU
UGAAGAAUGAGCCGGCGAGUUAC-G-GUCAGUGGCA--GGUUAAGC--C-G---------AAAA--------
U-GG--CGGA--GCCAAAGGGAAAC-C---------------------------------------------
------------------------------------------------------------------------
------------------------------------------------------------------------
---------------------------------------------------------------------GAG
UCU------------------------------------GAAUA-------------------------GG-
G-C-GC-----------------------UAGUCACUGGUCGUA-GACCCGAAGCC-UGGUGAUCUAACCAU
GGCCAGGAUGAAGCUUGG-GUAACA-CCAAGUGGAGGUCCGA--ACCGACUG-AUGUUGAAAAAUCA-GCGG
AUGAGCUGUGGUUAGGGGUGAAAUGCCAAUCGAACUAGG-CGAUAGCUGGUUCUCCCCGAAA-UGUGUUUAG
GCACAGCGUUAG-GUG--------------------------------------------------------
----------------------UU-----UA-UGGGCGGGGGUAGAGC-ACUGAUU-CGG-UGC--GGGCUG
C---GA-GA--GCGGUACCAAAUCGAGCCAAACUCCGAAUACCGUC--------------------------
------------------------------------------------------------------------
------------------CAAC-A-UA--------GCCUGGC-AGUGAGA-C-UGUG-G-GGGAUAAGCUCC
AU-G-GUCGAGAG-GGAAAC---AGCCCAGACCACCA-ACUAAGGUCCCCAAAU--UG-UAG-CU-CA---G
-U-GAU-------UAAGGAGGUGGGAUUGCACAGACAACCAGGAGGUUGGCCUAGAAGCAGCCA-CCCUUGA
AAGAGUGCGUAAUAGCUCACUGGUC-----AAGCGAUCCUGCGCCGAAAAUG-AAC-GGGGCU-A-AGCUAC
```

```
A----------UACCGAA-GUUGUGGAUUUA-----------------------------------------
------------------------------------------------------------------------
------------------------------------------------------------------------
-------------------------------------------UGGUAGGGGAGC-GUUCCGC---------
-----------AGU-AGGAU-GAAGCGU--CAGCGACAAGCA--G-GCGUGGACGA-G-GCG-GAAGUGAGA
AUGU-CGGCUU-GAGUA-GCGCAA---ACAU-UGGUGA--G---------------AAUCCA-AUG-CUCCG
A-AAGCAC-AAGGUUUCC-UCCGGAAGG-UUCGUCCGCGGAGGGUUAGCCGGG-AC-CUAAGGCGAAGCCG-
A-A-GGGC-----GUA--------------------------------------------------------
----------------------------------------GUC-GA-U--GGAU-AACAGGUG-UAUA-UUC
CUGUG----------------------------------CUGUG----UCGC--------U-GGA-------
------------------------------------------------------------------------
----------------------------------------------------------GUGUC-AAAGGGAC
GCAUGA--------------GGAUAG-GCCAUACCC-UU-A----CUGGA------------UUGGGGAGC-
------------------------------------------------------------------------
------------------------------------------------------------------------
----------------------------------------------------------------------UC
-------------CUACGGGAGCAG-U-UUG-G-CCGAAAGUAGUGC-C-GAGAAA-AGCUU-UGCCACAU-
--------------------------------------------------------UGAAAG-CA-CGCACC
CGUACCCGAAACCGAC------------------------------ACAGGUG--UGCAGGUA-GAGAAUAC
CAAGGA-G-CG-CGAG-AUAACCCU-C-UUUAAGGAACUCGGC-AAAAUAGCCCCGUAACUUCGG-AAGAAG
GGGUG-CCCC-C-GC---------------------------------------------------------
------------------------------------------------------------------------
------------------------------------------------------------------------
------------------------------------------------------------------------
------------------------------------------------------------------------
------------------------------------------------------------------------
------------------------------------------------------------------------
------------------------------------------------------------------------
------------------------------------------------------------------------
------------------------------------------------------------------------
------------------------------------------------------------------------
------------------------------------------------------------------------
------------------------------------------------------------------------
------------------------------------------------------------------------
----------------------------------------------------------------------GA
GGG--GGUCGCAGUGAAAUGGCUCUGGCGA-----CUGUUUACCAAAAACACAGGUCUCUGCAAAGCCG---
-CA-A-------------GGCGACGU-AUAGGGGCUGA-CUCCUGCCCAGUGCCG-G-AAGG-UUACGGAAG
UCGGUUA-GC-----GAAA-----GCGAAGCUGGCGACCUAAGCCC--CGGUGAAC-GGCGGCCGUAACUAU
AACGGUCCUAAGGUAGCGAAAUUCCUUGUCGGG-UAAGUUCCGACCUGCACGAAAGGAGUAACGAUC--AGA
GCGCUGUCUCGAAG----AGG-GACUCGGCGAAAUA--GGA---UUGUCUGUGAAGAUACGGACUACUUGUA
CCCGGACAGAAAGA-CCCUAUGAAGC-UUUACU--CUAGCUU-GAAAUUGGGUUCGGGCUUUGUCUGCGCAG
GAU-AGGCGGGAGCCGGUG----------AUACAUC-UCU----------------------------UGCG
--------------------------------------GG-GGGUGUGGAG-G--C-AA-CGGUGAGAUA-C
CGCU-CUGACAGGGUCUGGAUUCUAACC-------------UA-C-G-------------------------
------------------------------------------------------------------------
```

```
------------------------------------------------------------------------
------------------------------------------------------------------------
------------------------------------------------------------------------
------------------------------------------------------------------------
---------------------------------------------------G-----GGACA-GUUUCAGGU
GGGGAGUUUGACUGGGGCGGU-CGCCUCCUA-AACGGUAAC-GGAGGCGCGCA-AAGGUUCCCUCAGGCUGG
UCGGAAAUCAGCCGCA-GAGUG-UAAAGGCAU-AAGGGAGCUUGACU-GCGA--------------------
--------------GACCUA---CAAGUCGAGCAGAGACGAAA-GUCGGCCUUAGUGAUCCGAC--------
---GG-U-----UCUGC-AUG--------------------GA-AGGGCCGUCGCUCAACGG-AUAAAAGUU
ACUCUAGGG-AUAACAGGCUGAUCUCCCCCAAGAGUCCACAUCGACGGGGAGGUUUGGCACCUCGAUGUCGG
CUCGUCGCAACCUGGGGCGGUA-GUACGUCCCAAGGGUUGGGCUGUUCGCCCAUUAAAGCGGCACGUGAGCU
GGGUUCAGAACGUCGUGAGACAGUUCGGUCCAUAUCCGGU--------------------A-CAA-------
-------------------------------GCGC--AAGAGUGUUGAGAGG-AGC-UCUCCCUAGUACGAG
AGGAC-CGGGAGGGACUGACCGC-UGGU-G-UACCUGUUAUCGUGCCAACGGUAA-ACGCAGGGUAGCUAUG
UCGGGAGUGGAUAACCGCUGAAAGCA-UC--UAAGUGGGAAGCCCACCUCAA-G-AUG--AGCACUCU-CU-
--CCGCA-----------------------------------------------------------------
------------------------------------------------------------------------
------------------------------------------------------------------------
------------------------------------------------------------------------
------------------------------------------------------------------------
------------------------------------------------------------------------
------------------------------------------------------------------------
------------------------------------------------------------------------
----------------------------------------------------AGGG--UA-AGA------CC
ACGGGAAGAUUACC-CGU-UAA-UAGGCU-CUA-GGUGGAAG-CGC-A----------------GC------
----AA-----------UGUGUGCAGCCG--A-GGAG-UACUAAUA---GGUCGAG-G-GCUUGACUU----
------------------------------------------------------------------------
------------------------------------------------------------------------
------------------------------------------------------------------------
-------------------------------------------------
>LSUb_205913_BIFLD/1-3066
---------------------GUGUGUUGCU----UG--CAAGGGCGUAU-GGUGGAUG-C-CUUGGC-AGA
CAG---GACCGAUGAAGGACGUUU--GAGGCUGCGAUAUGCCUCGGGGAGCCGCCAACAGGGCU-----UUG
AUC-CGA-GGAUUUCCGAAUGGGGGAAC--------CC--------AC----CGAC-C-G----------UC
A-U-------------G---G----G----U---CGG----U-A-CCGG-C---------------------
-----------U-----------UCG----------------------------------------------
-----------------------GC-C-GG-GGGGUACGCAGGGAAGUGAAACAUCUCAGUACCUG--CAGG
AAAGGA------------UAUUCCGUGA--GU-AG-UGGCGAGCGAAAGCGGA-U-GAUGGCCAAACCUUGC
-GCG-UGUGA-UACCCG-UCG------------G---GGG--U-UGCGCGCGGGGUGUUG------------
-----CGGG-AUC-------GCGUGCGCCGG--CUCCGACGGGCCGGCCGG--CAGUUGCAGAAAACCA---
U-------GUGUC-AGGGG------AAC-CG-GGUU------------------------------------
-------GAA-UA-CCGGG-------CCGCAGAGGGUGAGGGCCCCGUA---------CCUGAAC----GCG
C------A--UGGUC----------UGCCG-AUCGC--GU-CU------CCCAAGUAGCACGGGUCCCGUGG
AACCCCGUGCGAA-UCCGCCCAG-ACC--GUUGGGUAAGCCUGAAUA--UUC-CUGUC-UGACCGAUAGC-G
AAC-G-AGUACCGUGAGGGAAAG-GUGAAA-AGUAC-CCCGG-GAGGGG-A-GUGAAACAGUCUCUGAAACC
```

```
G-UGCGCCUACAA-UCC-GU-CGGA---GCCC-U--------------------------------------
------------------------------------------------------------------------
------------------------------------------------------------------------
------------------------------------------------------------------------
------------------------------------------------------------------------
------------------------------------------------------------------------
------------------------------------------------------------------------
------------------------------------------------------------------------
------------------------------------------------------------------------
-------------------------------------------------------------UU--U-GU---
------------------------------------------------------------------------
------------------------------------------------------------------------
------------------------------------------------------------------------
------------------------------------------------------------------------
------------------------------------------------------------------------
------------------------------------------------------------------------
------------------------------------------------------------------------
------------------------------------------------------------------------
------------------------------------------------------------------------
------------------------------------------------------------------------
------------------------------------------------------------------------
------------------------------------------------------------------------
------------------------------------------------GG--G---GUGACGGCGUGCCUAU
CGAAAAAUGAGUCUGCGAGUCAGUG-GCAUGUGGCGA-GCAUAACC--C-----------GUGU--------
--GG--GGUA--UGCGUAGCGAAGG-C---------------------------------------------
------------------------------------------------------------------------
------------------------------------------------------------------------
---------------------------------------------------------------------GAG
UCU------------------------------------UAAA--------------------------AG-
G-C-GU---------------------UUGAGUCGCGUGUCCUG-GACCCGAAGCG-GGAUGAUCUAGCCCU
GAGCAGGUUGAAGCGCGG-GUAAGA-CCGCGUGGAGGACCGA--ACCCACCU-GGGUUGAAAACCGG-GGGG
AUGACUUGGGGCUAGGGGUGAAAGGCCAAUCAAAUUCCG-UGAUAGCUGGUUCUCUCCGAAA-UGCAUUUGG
GUGCAGCGUCGG-GUC--------------------------------------------------------
----------------------AU-----UACGUCCCGGGGGUAGAGCUACUGGAU-GCU-UGC--GGGCCC
----GUAUC---GGGUACCAACAGCAACCAAACUCCGAAUACCGGU--------------------------
------------------------------------------------------------------------
------------------GACGCG-UA--------UCCCGGC-AGUGAGU-C-GGCG-G-GGGAUAAGCUUC
GU-C-GUCGAAAG-GGAAAC---AGCCCAGACCGUCG-UCUAAGGUCCCGAAGC--GU-GUG-CU-AA---G
-U-GGG-------AAAGGAUGUGGAGUCGCAUAGACAGCCAGGAGGUUGGCUCAGAAGCAGCCA-CCCUUGA
AAGAGUGCGUAACAGCUCACUGGUC-----UAGUGGUUCCGCGCCGACAAUGUAGC-GGGGCUCA-AGCACA
C----------CACCGAA-GACGCGGCAGUAUUU--------------------------------------
------------------------------------------------------------------------
------------------------------------------------------------------------
-----------------------------------UG-UACU-GGGUAGGAGAGC-GUCCCAU---------
-----------G-A-GGGGC-GAAGCGG--C-CGUGUGAACG-CG-CCGUGGACUU-C-AUG-GGAGUGAGA
```

```
AUGC-AGACAU-GAGUA-GCGAGA----GAC-GGGUGA--A---------------GAUCCC-GUC-CGCUG
G-AUGACC-AAGGGUUCC-AGGGCCACG-UUCAUCGUCCCUGGGUGAGUCGGG-UC-CUAAGGCGAGGCCG-
A-C-AGGC-----GUA--------------------------------------------------------
----------------------------------------GUC-GAAU--GGAUGAACGGGUC-GAUA-UUC
CCGUA----------------------------------CCGGC----UUUC--------A-GC--------
------------------CGCCAAAACCGAGGCUUGGAGUACUAACCUCCGGGUCCGGGCUUACUGCUCCUU
CGGGGGCGGGAUGCCCUGGCCUGUUGGGACC--GAUCCUUGUAGUAGGUCA-------GCGCA-GGAGUGAC
GCAGAA--------------GGGUAG-CCGGCCGCG-GA-G----GUGGU------------CUUCCGUGG-
UCAAGC-ACG-CA-----G-G-------------------G------CG------------------UG-GG
A-CAGG--------------------CAAAU-CCGUCCCGCA------U-GGGUCCG----AGG-UG-C-G-
AUG--AU---------------------------------------------------------------GG
-----GGGCCU--UAUGGCCCGAUA-U-CCG-G-UGAUCCCGGCUGC-C-GAGAAA-AGCUU-CGGCGUCA-
--------------------------------------------------------GGCUCG-AA-GCCGCC
CGUACCCCAAACCGAC------------------------------ACUGGUG--GUCAGGUA-GAGAAUAC
CAAAGC-G-AU-CGAG-CGAAUCCU-G-GUCAAGGAACUCGGC-AAAUCACUCCCGUUCCUUCGG-UUUAAG
GGAGA-CCCC-C-GA--UGGUGAAC----CGC-----------CUCGC------------------------
------------------------------------------------------------------------
------------------------------------------------------------------------
------------------------------------------------------------------------
------------------------------------------------------------------------
------------------------------------------------------------------------
------------------------------------------------------------------------
------------------------------------------------------------------------
------------------------------------------------------------------------
------------------------------------------------------------------------
------------------------------------------------------------------------
------------------------------------------------------------------------
------------------------------------------------------------------------
------------------------------------------------------------------------
--------------------------------------------------------GCGG-GGA--GCUUUU
GGG--GGUGGCACAGACCAGGGGGUAGCGA-----CUGUUUACCAAAAACACAGGUGCAUGCGAAGGCG---
-CA-A-------------GCCGCUGU-AUAUGCACUGA-CGCCUGCCCGGUGCCG-G-AAGG-UUAAGAGGA
UCCGUCA-GCC----UUCG----GGUGAAGCGGUGAAUUCAAGCCC--CGGUAAAC-GGCGGUGGUAACUAU
AACCAUCCUAAGGUAGCGAAAUUCCUUGUCGGG-UAAGUUCCGACCUGCACGAAUGGCGUAACGACU--UCC
CCACUGUCUCGACC----AGG-AGCUCGGCGAAAUU--GCA---GUACGAGUAAAGAUGCUCGUUAAGCGCA
GAAGGACGAAAAGA-CCCCG-GGACC-UUUACU--AUACCUU-GGUAUUGGCAUUCGGUGUGGAUUGUGUAG
CAU-AGGCGGGAGGCCGAG----------AGGCGCG-GUC----------------------------GCCA
--------------------------------------GA-UCGCGCGGAG-C--C-GUAAGGUGAAAUA-C
CGCU-CUGUUCGCAUUGGAUGUCUAACCUCGA--ACAG-U-CA-U-C-------------------------
------------------------------------------------------------------------
------------------------------------------------------------------------
------------------------------------------------------------------------
------------------------------------------------------------------------
------------------------------------------------------CU----GU----------
------------------------------------------------UCAG-----GGACA-GUGCCUGGC
```

```
GGGUAGUUUAACUGGGGCGGU-UGCCUCCCA-AAGAGUAAC-GGAGGCGCCCA-AAGGUCCGCUCAGCCCGG
UUGGCAAUCGGGUGGC-GAGUG-CAAGUGCAC-AAGCGGGCUUGACU-GCGA--------------------
--------------GGACGA---CGGUCCGAGCAGGGACGAAA-GUCGGGACUAGUGAUCCGGU--------
---GC-C-----GGUGU-ACG--------------------GA-CGCGGCAUCGCUCAACGG-AUAAAAGGU
ACCCCGGGG-AUAACAGGCUGAUCAUUCCCAAGAGUCCAUAUCGACGGGAUGGUUUGGCACCUCGAUGUCGG
CUCGUCGCAUCCUGGGGCUGGA-GCAGGUCCCAAGGGUUCGGCUGUUCGCCGAUUAAAGCGGCACGCGAGCU
GGGUUCAGAACGUCGUGAGACAGUUUGGUCUCUAUCCUCU--------------------G-CGC-------
-------------------------------UCGU--UGGAAUGCUGAGGAG-GCC-UGCCCAUAGUACGAG
AGGAC-CUGGGUGGACGAACCUC-UGGU-A-UGCCGGUUGUCACGCCAGUGGCAC-G-GCCGGUUGGCUACG
UUCGGAAGGGAUAACCGCUGAAAGCA-UC--UAAGCGGGAAGCCUGCUCCAA-G-AUA--GCAUUCCG-UG-
--GAACC-----------------------------------------------------------------
--------------------------UU---------CG---------------------------------
------------------------------------------------------------------------
------------------------------------------------------------------------
------------------------------------------------------------------------
------------------------------------------------------------------------
------------------------------------------------------------------------
------------------------------------------------------------------------
-------------------------------------------------AGUUCCU--GU-AGU------CC
CCAUGCAGAACACGUGGU-CGA-UAGGCC-GGA-CGUGGAAG-CCC-C----------------GC------
----GA-----------GGGGUGGAGCCG--A-CCGG-UACUAACG---GACGAAA-G-GCAACACUA----
------------------------------------------------------------------------
------------------------------------------------------------------------
------------------------------------------------------------------------
------------------------------------A------------
>LSUb_281090_LEIXX/1-3107
-------------------------CAAGUU----UC--UAAGAGCAAAC-GGUGGAUG-C-CUUGGC-AUC
UGG---AGCCGAAGAAGGACGUAG--UAAUCUGCGAUAAGCCUCGGGGAGUUGAUAAACGGACU-----UUG
AUC-CGA-GGAUUUCCGAAUGGGGAAAC--------CC--------CG----CUGG-G-CCC---------U
U-U-----------GGGUGAC----C----C---AGU----G-A-CUCC-CGC-----CUGA----------
-----------A-----------UAUA----------------------------------------UA-GG
G-C--------------------GG-GUAG-AGGGAACGUGGGGAAGUGAAACAUCUCAGUACCCA--CAGG
AAGAGAAAGCAA--AAGCGAUUCCGUUA--GU-AG-UGGCGAGCGAAACCGGA-A-GA-GGCUAAACCGAUC
-AUG-UGUGAUAGCCGG-CAG------------G---CGU-UG-CAUGGUCGGGG--UUG------------
-----CGGG-ACU-------UUUC-UGCUGC--UUCUGCCGAACAGCAAGC--GUU---------ACAGA-G
G-------AUCAU-AGGUG------AAU-GG-CAUU------------------------------------
-------GAA-AG-GCCAG-------UCAUAGAGGGUGCCAACCCCGUA---------ACCGAAA----UGG
U------C--CAAUG----------GUGCG--AAGA--GU-AU------CCCAAGUAGCACGGGGCCCGAGA
AAUCUCGUGUGAA-UCUGUCAGG-ACC--ACCUGAUAAGCCUAAAUA--CUC-CCAGA-UGACCGAUAGC-G
GAC-A-AGUACCGUGAGGGAAAG-GUGAAA-AGUAC-CCCGG-GAGGGG-A-GUGAAAUAGUACCUGAAACC
G-UUUGCUUACAA-ACC-GU-UGGA---GCCU-C---C----------------------------------
------------------------------------------------------------------------
------------------------------------------------------------------------
------------------------------------------------------------------------
------------------------------------------------------------------------
```

```
------------------------------------------------------------------------
------------------------------------------------------------------------
------------------------------------------------------------------------
------------------------------------------------------------------------
--------------------------------------------------------------U--U-GU---
------------------------------------------------------------------------
------------------------------------------------------------------------
------------------------------------------------------------------------
------------------------------------------------------------------------
------------------------------------------------------------------------
------------------------------------------------------------------------
------------------------------------------------------------------------
------------------------------------------------------------------------
------------------------------------------------------------------------
------------------------------------------------------------------------
------------------------------------------------------------------------
------------------------------------------------------------------------
-----------------------------------------------AGG--G---GUGACAGCGUGCCUUU
UGAAGAAUGAGCCUGCGAGUUAGUG-AUAUGUGGCGA-GGUUAACC--C-----------GUGA--------
--GG--GGUA--GCCGUAGCGAAAG-C---------------------------------------------
------------------------------------------------------------------------
------------------------------------------------------------------------
---------------------------------------------------------------------GAG
UCU------------------------------------GAAUA-------------------------GG-
G-C-GA---------------------UUCAGUCGCAUGUCCUA-GACCCGAAGCG-AAGUGAUCUAUCCAU
GGCCAGGUUGAAGCGACG-GUAAGA-CGUCGUGGAGGACCGA--ACCCACUU-AGGUUGAAAACUGA-GGGG
AUGAGCUGUGGAUAGGGGUGAAAGGCCAAUCAAACUUCG-UGAUAGCUGGUUCUCUCCGAAA-UGCAUUUAG
GUGCAGCGUUGC-GUG--------------------------------------------------------
----------------------UU-----UC-UUGCCGGAGGUAGAGCUACUGGAU-GGC-CGA--UGGGCC
C---CA-AA--AGGUUACUGACGUCAGCCAAACUCCGAAUGCCGGU--------------------------
------------------------------------------------------------------------
------------------AAGUGA-GA--------GCGCAGC-AGUGAGA-C-GGUG-G-GGGAUAAGCUUC
AU-C-GUCGAGAG-GGAAAC---AACCCAGACCACCA-ACUAAGGUCCCUAAGC--GC-GUG-CU-AA---G
-U-GGG-------AAAGGAUGUGGAGUUGCACAGACAACCAGGAGGUUGGCUUAGAAGCAGCCA-CCCUUGA
AAGAGUGCGUAACAGCUCACUGGUC-----AAGUGAUUCCGCGCCGACAAUGUAAC-GGGGCUCA-AGCACG
C----------CACCGAA-GUUGUGGCAUUAACA--U-UAU-------------------------------
----------------------------------------------UGG-----------------------
-UAGGCUUUCGU------------------------------------------------------------
------------------------GGUCCAGCCGUGU-UGAU-GGGUAGGAGAGC-GUCGUGU---------
-----------G-G-CGAGU-GAAGCGG--C-GGUGUGAACC-AG-CCGUGGACGC-C-ACA-CGAGUGAGA
AUGC-AGGCAU-GAGUA-GCGAAA----GAC-GGGUGA--G---------------AAACCC-GUC-CUCCG
A-AAGACC-AAGGGUUCC-AGGGCCAGG-CUAAUCCGCCCUGGGUAAGUCGGG-AC-CUAAGGCGAGGCCG-
A-C-AGGC-----GUA--------------------------------------------------------
----------------------------------------GUC-GA-U--GGAC-AACGGGUU-GAUA-UUC
CCGUA----------------------------------CCGGC----GAAG--------A-AC--------
```

```
------------------CGUCCAAGCUAAUCCAGUGGUGCUAAGAGUCCUAAUCCUGGACACCGAUCCCUU
CGGGGUGACGGUCCAGGUCUAACGCUCGACC--CCAUGCUGGUGCGGCUAGCGUAU--UAACA-GGUGUGAC
GCAGGA--------------AGGUAG-CUGUACCGG-GC-G----AUGGU------------UGUCCCGGG-
GUAAGG-AUG-UA-----G-G-------------------G------CG------------------AG-AG
A-UAGG--------------------CAAAU-CCGUCUCUCA------CAUAGCCUG----AGA-UC-U-U-
AUG--CA---------------------------------------------------------------UA
-----GCCGU------CAAGGCGAA-U-UCA-G-UGAUCCUAUGCUG-C-CAAGAA-AAGCA-UCGACGCG-
--------------------------------------------------------AGGUUC-CA-GCCGCC
CGUACCCCAAACCGAC------------------------------UCAGGUG--GUCAGGUA-GAGAAUAC
UAAGGA-G-AU-CGAG-AGAAUCGU-G-GUUAAGGAACUCGGC-AAAAUGCCCCCGUAACUUCGG-GAGAAG
GGGGG-CCUG---AG--GCGUGAAG----GGA-----------UUUAC------------------------
------------------------------------------------------------------------
------------------------------------------------------------------------
------------------------------------------------------------------------
------------------------------------------------------------------------
------------------------------------------------------------------------
------------------------------------------------------------------------
------------------------------------------------------------------------
------------------------------------------------------------------------
------------------------------------------------------------------------
------------------------------------------------------------------------
------------------------------------------------------------------------
------------------------------------------------------------------------
------------------------------------------------------------------------
--------------------------------------------------------UCCU-GGA--GCGUUU
GAA--GGCCGCAGAGACCAGUGGGAAGCGA-----CUGUUUACUAAAAACACAGGUCCGUGCCAAGUCG---
-CA-A-------------GACGAUGU-AUACGGACUGA-CGCCUGCCCGGUGCUG-G-AAGG-UUAAGAGGA
AGGGUUA-GC-----UUCG-----GCGAAGCUCAGAAUUUAAGCCC--CAGUAAAC-GGCGGUGGUAACUAU
AACCAUCCUAAGGUAGCGAAAUUCCUUGUCGGG-UAAGUUCCGACCUGCACGAAUGGCGUAACGACU--UCC
CAGCUGUCUCAACC----ACG-AACUCGGCGAAAUU--GCA---CUACGAGUAAAGAUGCUCGUUACGCGCA
GCAGGACGGAAAGA-CCCCG-UGACC-UUUACU--AUAGUUU-GGUAUUGGUGUUCGGUGUGGCUUGUGUAG
GAU-AGGUGGGAGACUUUG----------AAGCGGG-CAC----------------------------GCCA
--------------------------------------GU-GUUCGUGGAG-U--C-AU-UUUUGAAAUA-C
CACU-CUGGUCAUUCUGGAUAUCUAACUUAGA--ACCG-U-AA-U-C-------------------------
------------------------------------------------------------------------
------------------------------------------------------------------------
------------------------------------------------------------------------
------------------------------------------------------------------------
------------------------------------------------------CG----GU----------
------------------------------------------------UCAG-----GGACA-GUGCCUGAU
GGGUAGUUUAACUGGGGCGGU-UGCCUCCCA-AAAAGUAAC-GGAGGCGCCCA-AAGGUUCCCUCAACCUGG
UUGGCAAUCAGGUGUC-GAGUG-UAAGUGCAC-AAGGGAGCUUGACU-GUGA--------------------
--------------GACUGA---CAAGUCGAGCAGGGACGAAA-GUCGGGACUAGUGAUCCGGC--------
---AG-U-----GGCUU-GUG--------------------GA-AGCGCUGUCGCUCAACGG-AUAAAAGGU
ACCUCGGGG-AUAACAGGCUGAUCUUGCCCAAGAGUCCAUAUCGACGGCAUGGUUUGGCACCUCGAUGUCGG
```

```
CUCGUCGCAUCCUGGGGCUGGA-GUAGGUCCCAAGGGUUGGGCUGUUCGCCCAUUAAAGCGGUACGCGAGCU
GGGUUUAGAACGUCGUGAGACAGUUCGGUCCCUAUCCGCU--------------------G-CGC--------
-------------------------------GCGC--AGGAAAUUUGAGAGG-AUC-UGACCCUAGUACGAG
AGGAC-CGGGUUGGACGAACCUC-UGGU-G-UGCCAGUUGUUCCGCCAGGAGCAC-C-GCUGGUUAGCUACG
UUCGGGAUGGAUAACCGCUGAAAGCA-UC--UAAGCGGGAAGCCGGCCUCAA-G-AUG--AGAUUUCC-AC-
--GCC-------------------------------------------------------------------
--------------------------UU--------------------------------------------
------------------------------------------------------------------------
------------------------------------------------------------------------
------------------------------------------------------------------------
------------------------------------------------------------------------
------------------------------------------------------------------------
------------------------------------------------------------------------
-------------------------------------------------CGGGCGA--GA-GGC------UC
CCAGCUAGACUACUGGGU-UGA-UAGGCC-GGA-UGUGGAAG-UGG-G----------------GACU----
-AAAGA-----------CCCAUGGAGCUG--A-CCGG-UACUAAUA---AGCCGAU-A-ACUUGAU------
------------------------------------------------------------------------
------------------------------------------------------------------------
------------------------------------------------------------------------
-------------------------------------------------
>LSUb_203267_TROWT/1-3099
---------------------A-UUCAAGCU----ACUUUAAGAGCAAAU-GGUGGAUG-C-CUUGGU-AUC
UGG---AGCCGAAGAAGGACGUAG--CAAUCUGCGAUAAGCCUCGGGAAGCUGAUAAGCGAGCU-----UUG
AUC-CGA-GGAUUUCCGAAUGGGGAAAC--------CU--------GG----UGGG-G-CGCA-------GC
A-A----------UGCG--AC----C----C---UGU----C-A-CUCC-CGC-----CUGA----------
-----------A-----------UAUA----------------------------------------UAG-G
G-C--------------------GG-GCAG-AGGGAACGUGGGGAAGUGAAACAUCUCAGUACCCA--CAGG
AAGAGAAAGCAA--UAGCGAUUCCGUUA--GU-AG-UGGCGAGCGAAACCGGA-A-CA-GGCCAAACCGAUU
-GAG-UGUUAUAGCCGG-CAG------------G---CGU-UU-CUUGAUCGGGG--UAG------------
-----UGGG-AUC-------CUCU-AUGUGG--UUCUGCCGAACCACACGC--GUA---------GUGCA-G
G-------GGUAU-AGGUG------AAU-GA-UUUU------------------------------------
-------GAA-CA-GUCAA-------CCGAAGAGGGUGAUAGUCCCGUA---------AUCGAAA----UGC
C------C--CGGUG----------CCGCG-AGAGG--AU-AU------CCCAAGUAGCACGGGCCCCGUGA
AAUCCCGUGUGAA-UCUGUCAAG-ACC--ACUUGAUAAGCCUAAAUA--CUA-CCAGA-UAACCGAUAGU-G
GAC-G-AGUACCGUGAGGGAAAG-GUGAAA-AGUAC-CCCGA-GAGGGG-A-GUGAAAUAGUACCUGAAUCC
A-UUUGCUUACAA-ACC-GU-CGGA---GCAGUG---UG---------------------------------
------------------------------------------------------------------------
------------------------------------------------------------------------
------------------------------------------------------------------------
------------------------------------------------------------------------
------------------------------------------------------------------------
-------------------------------------------------------------C-G--------
------------------------------------------------------------------------
------------------------------------------------------------------------
-------------------------------------------------------------AC--U-UU---
```

------------------------------------------------------------------------
------------------------------------------------------------------------
------------------------------------------------------------------------
------------------------------------------------------------------------
------------------------------------------------------------------------
------------------------------------------------------------------------
------------------------------------------------------------------------
------------------------------------------------------------------------
------------------------------------------------------------------------
------------------------------------------------------------------------
------------------------------------------------------------------------
------------------------------------------------------------------------
-------------------------UGU------------------CGCU--U---GUGACGGCGUGCCUUU
UGAAGAAUGAGCCUGCGAGUUAGCG-AUGUGUGGCGA-GAUUAACC--C-----------GUUG--------
--GG--GGUA--GUCGUAGCGAAAG-C---------------------------------------------
------------------------------------------------------------------------
------------------------------------------------------------------------
---------------------------------------------------------------------GAG
UCU------------------------------------GAAUA-------------------------GG-
G-C-GA---------------------AUUAGUCGCAUGUCCUA-GACCCGAAGCG-AAGUGAUCUAUCCAU
GGCCAGGCUGAAGCGAGG-GUAAGA-CCUCGUGGAGGGCCGA--ACCCACUU-AGGUUGAAAACUGA-GGGG
AUGAGCUGUGGAUCGGGGUGAAAGGCCAAUCAAACUUCG-UUAUAGCUGGUUCUCUCCGAAA-UGCAUUUAG
GUGCAGCGUUGC-GUG--------------------------------------------------------
----------------------UU-----UC-UUGCCGGAGGUAGAGCUACUGGAU-GGC-CAA--UGGGCC
C---UA-CA--AGGUUACUGACGUCAGCCAAACUCCGAAUGCCGGU--------------------------
------------------------------------------------------------------------
------------------AAGUUA-GA--------GCGCAGC-AGUGAGG-C-UGCG-G-GGGAUAAGCUUC
GU-A-GCCGAGAG-GGAAAC---AACCCAGACCACCG-AUUAAGGUCCCUAAGU--GU-GUG-CU-AA---G
-U-GGU-------AAAGGAUGUGGAGUUGCACAGACAGCCAGGAGGUUGGCUUAGAAGCAGCCA-CCCUUGA
AAGAGUGCGUAAUAGCUCACUGGUC-----AAGUGAUUCUGCGCCGACAAUGUAAC-GGGGCUAA-AGCACA
U----------CACCGAA-AUUGUGGCAUCGAUA--CACAU-------------------------------
----------------------------------------------AGG-----------------------
-UAGGCUUUCGU------------------------------------------------------------
------------------------AGUCCAGCCGUAU-CGAU-GGGUAGGAGAGC-GUCGUGU---------
-----------G-G-CGAAA-GAAGCGG--C-GAUGUAAAUC-AG-CCGUGGACGC-U-ACA-CGAGUGAGA
AUGC-AGGCAU-GAGUA-GCGAAA----GGC-GGGUGA--A---------------AAACCC-GCCCCCCCG
A-AUAACU-AAGGGUUCC-AGGGCCAGG-CUAAUCCGCCCUGGGUAAGUCGGG-AC-CUAAGGCGAGGCCG-
A-G-AGGC-----GUA--------------------------------------------------------
----------------------------------------GUC-GA-U--GGAC-AACGGGUU-GAUA-UUC
CCGUA----------------------------------CCGGC----AAAG--------A-AC--------
------------------CGCCCAUAUUCCGCGUAGUUUCGUUCAGCAUUUUGCUGUGCUUACGGUACCCCU
UUUACGGUGUGCGGAUAA--------------------------------------GUGUUCA-GGUGUGAC
GCAGGA--------------AGGUAG-CCAAGCCGG-GC-G----AUGGU------------UGACCCGGU-
GUAAGU-GUG-UA-----A-C-------------------C------CG------------------GA-GG
G-UAGG--------------------CAAAU-CCGCCCUCCAU----CUAAAGGGCC----AGG-CA-C-G-

```
ACG--CG---------------------------------------------------------------UA
-----GCGCUU----CGGCGCGAAA-U-UGG-U-GAUCCUAUGCUGA-C-UAGAAA-AGCAU-CGACACA--
--------------------------------------------------------AGGUUC-UA-GCUGCC
CGUACCCCAAACCGAC------------------------------UCAGGUG--GUUAGGUA-GAGAAUAC
CAAGGG-G-AU-CGAG-AGAAUCGU-G-GUUAAGGAACUCGGC-AAAAUGCCUCCGUAACUUCGG-GAGAAG
GAGGG--CCU-G-AG--GCGUGACG----AGA-----------CUAGC------------------------
------------------------------------------------------------------------
------------------------------------------------------------------------
------------------------------------------------------------------------
------------------------------------------------------------------------
------------------------------------------------------------------------
------------------------------------------------------------------------
------------------------------------------------------------------------
------------------------------------------------------------------------
------------------------------------------------------------------------
------------------------------------------------------------------------
------------------------------------------------------------------------
------------------------------------------------------------------------
------------------------------------------------------------------------
----------------------------------------------------------UCUC-CUAGCGCUG
CAG--GGUCACAGAAACCAGUGGGGAGCGA-----CUGUUUAUUAAAAACAUAGGUCUGUGCCAAGUCG---
-UA-A-------------GACGACGU-AUACAGACUGA-CGCCUGCCCGGUGCCG-G-AAGG-UUAAGAGGA
CGGGUCA-GC-----UUUG-----GCGACGCUCGGAAUUUAAGCCC--CGGUAAAC-GGCGGUGGUAACUAU
AACCAUCCUAAGGUAGCGAAAUUCCUUGUCGGG-UAAGUUCCGACCUGCACGAAUGGCGUAACGACU--UCC
CAACUGUCUCAACC----GCG-AACUCGGCGAAAUU--GCA---CUACGAGUGAAGAUGCUCGUUACGCGUA
GCAGGACGGAAAGA-CCCCG-AGACC-UUUACU--AUAGCUU-GGUACUGGUACUUGGUGCGGUUUGUGUAG
GAU-AGGUGGGAGACUGUG----------AAGCGGG-UGC----------------------------GCCA
--------------------------------------GC-ACUCGUGGAG-U--C-GU-UGUUGAAAUA-C
CACU-CUGAUCGCCCCGGGUGUCUAACUUAGA--ACCG-U-AA-U-C-------------------------
------------------------------------------------------------------------
------------------------------------------------------------------------
------------------------------------------------------------------------
------------------------------------------------------------------------
------------------------------------------------------CG----GU----------
------------------------------------------------UCGA-----GGACA-AUGCCUGGU
GGGUAGUUUAACUGGGGCGGU-UGCCUCCCA-AAAGGUAAC-GGAGGCGCCCA-AAGGUUCCCUCAGCCUGG
UCGGUAAUCAGGUGGC-GAGUG-UAAGUGCAC-AAGGGAGCUUGACU-GCGA--------------------
--------------GACUGA---CGGGUCGAGCAGGGACGAAA-GUCGGGACUAGUGAUCCGGC--------
---AG-U-----GGCUU-GUG--------------------GA-AGCGCUGUCGCUCAACGG-AUAAAAGGU
ACCUCGGGG-AUAACAGGUUGGUCCCUCCCAAGAGUCCAUAUCGACGGAGGGGUUCGACACCUCGAUGUCGG
CUCGUCGCAUCCUGGGGCUGUA-GCAGGUCCCAAGGGUUGGGCUGUUCGCCCAUUAAAGCGGUACGCGAGCU
GGGUUUAGAACGUCGUGAGACAGUUCGGUCCCUAUCCGCU--------------------A-CGC-------
-------------------------------GCGU--CGGAAAUUUGAGAGG-AGC-UGACCCUAGUACGAG
AGGAC-CGGGUUGGACGAACCUC-UGGU-G-UACCAGUUGUCCUGCCAAGGGCAC-G-GCUGGUUUGCUACG
UUCGGAACGGAUAACCGCUGAAAGCA-UC--UAAGCGGGAAACCGGCCUCAA-G-AUG--AGAUUUCC-AU-
```

```
--GCCCU-----------------------------------------------------------------
--------------------------UU--------------------------------------------
------------------------------------------------------------------------
------------------------------------------------------------------------
------------------------------------------------------------------------
------------------------------------------------------------------------
------------------------------------------------------------------------
------------------------------------------------------------------------
--------------------------------------------------UGGGCG--AG-AGG------CU
CCCAGUAGAACACUGGGU-UGA-UAGGCU-CGA-UGUGUACG-CAG-G----------------GACU----
-AAAGA-----------CCUGUUUAGCUU--A-CGAG-UACUAAUA---GGCCGAU-U-GCUUGAAAC----
------------------------------------------------------------------------
------------------------------------------------------------------------
------------------------------------------------------------------------
------------------------------------A------------
>LSUb_100226_STRCO/1-3122
---------------------G-GCCAAGUU----UU--UAAGGGCGCAC-GGUGGAUG-C-CUUGGC-ACC
AGG---AACCGAUGAAGGACGUGG--GAGGCCACGAUAGGCCCCGGGGAGUCGUCAACCAGGCU-----UUG
AUC-CGG-GGGUGUCCGAAUGGGGAAAC--------CC--------GG----CAGU-CGU-----------C
A-U-------------G---G----G----C---UGU----C-A-CCCG-CUG-----CUGA----------
-----------A-----------CACA----------------------------------------UAG-G
C-A--------------------GU-GUGG-AGGGAACGCGGGGAAGUGAAACAUCUCAGUACCCG--CAGG
AAGAGAAAACAA--CCGUGAUUCCGGGA--GU-AG-UGGCGAGCGAAACCGGA-U-GA-GGCCAAACCGUAU
-ACG-UGUGAGACCCGG-CAG------------G---GGU-UG-CGUGUGCGGGG--UUG-------------
-----UGGG-AUC-------UCUC-UUUUACG-GUCUGCCGGCCGUGAGAC--GAGUCAGAAA--CCGU---
U-------GAUGU-AGGCG------AAG-GA-CAUGC-----------------------------------
-------GAA-AG-GUCCG-------GCGUAGAGGGUAAGACCCCCGUA---------GUCGAA-----CGU
C------A--ACGGC----------UCGUU-UGAGA--GA-CA------CCCAAGUAGCACGGGGCCCGAGA
AAUCCCGUGUGAA-UCUGGCGGG-ACC--ACCCGCUAAGCCUAAAUA--UUC-CCUGG-UGACCGAUAGC-G
GAU---AGUACCGUGAGGGAAUG-GUGAAA-AGUAC-CGCGG-GAGCGG-A-GUGAAAUAGUACCUGAAACC
G-UGUGCCUACAA-GCC-GU-GGGA---GCGUCG---GG---------------------------------
------------------------------------------------------------------------
------------------------------------------------------------------------
------------------------------------------------------------------------
------------------------------------------------------------------------
------------------------------------------------------------------------
------------------------------------------------------------AU-GGAG-----
------------------------------------------------------------------------
------------------------------------------------------------------------
--------------------------------------------------------------C--U-UG---
------------------------------------------------------------------------
------------------------------------------------------------------------
------------------------------------------------------------------------
------------------------------------------------------------------------
------------------------------------------------------------------------
```

```
------------------------------------------------------------------------
------------------------------------------------------------------------
------------------------------------------------------------------------
------------------------------------------------------------------------
------------------------------------------------------------------------
------------------------------------------------------------------------
------------------------------------------------------------------------
----------------------CUCCA-------------------UCCU--C---GUGACUGCGUGCCUUU
UGAAGAAUGAGCCUGCGAGUUUGCG-GUGUGUUGCGA-GGUUAACC--C-----------GAGU--------
--GG--GGAA--GCCGUAGCGAAAG-C---------------------------------------------
------------------------------------------------------------------------
------------------------------------------------------------------------
---------------------------------------------------------------------GAG
UCC------------------------------------GAAUA-------------------------GG-
G-C-GU--------------------UUUUAGUAGCACGCUCAA-GACCCGAAGCG-GAGUGAUCUAGCCAU
GGGCAGGUUGAAGCGGCU-GUAAGA-GGUCGUGGAGGACCGA--ACCCACCA-GGGUUGAAAACCUG-GGGG
AUGACCUGUGGUUAGGGGUGAAAGGCCAAUCAAACUCCG-UGAUAGCUGGUUCUCCCCGAAA-UGCAUUUAG
GUGCAGCGUCGU-GUG--------------------------------------------------------
----------------------UU-----UC-UUGCCGGAGGUAGAGC-ACUGGAU-AGG-CGA--UGGGCC
C---UA-CC--GGGUUACUGACCUUAGCCAAACUCCGAAUGCCGGU--------------------------
------------------------------------------------------------------------
------------------AAGUGA-GA--------GCGCGGC-AGUGAGA-C-UGUG-G-GGGAUAAGCUCC
AU-G-GUCGAGAG-GGAAAC---AGCCCAGAGCAUCG-ACUAAGGCCCCUAAGC--GU-ACG-CU-AA---G
-U-GGG-------AAAGGAUGUGGAGUCGCACAGACAACCAGGAGGUUGGCUUAGAAGCAGCCA-CCCUUGA
AAGAGUGCGUAAUAGCUCACUGGUC-----UAGUGAUUCCGCGCCGACAAUGUAGC-GGGGCUCA-AGCGUA
C----------CGCCGAA-GUCGUGUCAUUGCAG--CA----------------------------------
------------------------------------------------A-----------------------
-UACGGCCA---------------------------------------------------------------
---------------------------ACGCCGGCUG-UGAU-GGGUAGGGAGC-GUCGUGU----------
-----------G-C-CGGGU-GAAGCAG--C-CGCGGAAGCG-AG-UUGUGGACGG-U-UCA-CGAGUGAGA
AUGC-AGGCAU-GAGUA-GCGAUA----CAA-ACGUGA--G---------------AAACGU-UUG-CGCCG
A-UUGACU-AAGGGUUCC-UGGGUCAAG-CUGAUCUGCCCAGGGUAAGUCGGG-AC-CUAAGGCGAGGCCG-
A-C-AGGC-----GUA--------------------------------------------------------
----------------------------------------GUC-GA-U--GGAU-AACCGGUU-GAUA-UUC
CGGUA----------------------------------CCCGC----UGUG--------A-AG--------
------------------CGUCAAACAUCGAGCAUCGUGAUGCUAAGGCCGUGAAGCCGCCCUGAUCUCUUC
GGAGUUGAGGGGAGUGGUGGAGCCGCCGAAC--CAAGCGGUUAGUAGGUGA-------GUGAU-GGGGUGAC
GCAGGA--------------AGGUAGUCCAUCCCGG-GC-G----GUGGU------------UGUCCCGGG-
GUAAGG-GUG-UA-----G-G-------------------C------CG------------------GU-GU
G-UAGG--------------------UAAAU-CCGCACACCU------U-GUGGCUG----AGA-CC-U-G-
AUG--CC---------------------------------------------------------------GA
-----GCCGAU----UGUGGUGAAG-U-GGA-U-GAUCCUAUGCUGU-C-GAGAAA-AGCCU-CUAGCGA--
--------------------------------------------------------GUUUCA-UG-GCGGCC
CGUACCCUAAACCGAC------------------------------UCAGGUG--GUCAGGUA-GAGAAUAC
CGAGGC-G-UU-CGGG-UGAACUAU-G-GUUAAGGAACUCGGC-AAAAUGCCCCCGUAACUUCGG-GAGAAG
```

```
GGGGG-CCAC-A-CC--UGGUGACG----AGU-----------UUUGC------------------------
------------------------------------------------------------------------
------------------------------------------------------------------------
------------------------------------------------------------------------
------------------------------------------------------------------------
------------------------------------------------------------------------
------------------------------------------------------------------------
------------------------------------------------------------------------
------------------------------------------------------------------------
------------------------------------------------------------------------
------------------------------------------------------------------------
------------------------------------------------------------------------
------------------------------------------------------------------------
------------------------------------------------------------------------
----------------------------------------------------------ACUC-UGAGCUGGG
GGU--GGCCGCAGAGACCAGCGAGAAGCGA-----CUGUUUACUAAAAACACAGGUCCGUGCGAAGCCG---
-UA-A-------------GGCGAUGU-AUACGGACUGA-CGCCUGCCCGGUGCUG-G-AACG-UUAAGGGGA
CCGGUUA-GUCACAUUUCGGUGUGGCGAAGCUGAGAACUUAAGCGC--CAGUAAAC-GGCGGUGGUAACUAU
AACCAUCCUAAGGUAGCGAAAUUCCUUGUCGGG-UAAGUUCCGACCUGCACGAAUGGCGUAACGACU--UCU
CGACUGUCUCAACC----AUA-GGCCCGGUGAAAUU--GCA---CUACGAGUAAAGAUGCUCGUUUCGCGCA
GCAGGACGGAAAGA-CCCCG-GGACC-UUUACU--ACAGUUU-GAUAUUGGUGUUCGGUUCGGCUUGUGUAG
GAU-AGCUGGGAGACUUUG----------AAGCUCG-CAC----------------------------GCCA
--------------------------------------GU-GUGGGUGGAG-U--C-GU-CGUUGAAAUA-C
CAGU-CUGGUCGUGCUGGAUGUCUAACCUGGG--UCCG-U-GA-U-C-------------------------
------------------------------------------------------------------------
------------------------------------------------------------------------
------------------------------------------------------------------------
------------------------------------------------------------------------
------------------------------------------------------CG----GA----------
------------------------------------------------UCAG-----GGACA-GUGUCUGAU
GGGUAGUUUAACUGGGGCGGU-UGCCUCCUA-AAGAGUAAC-GGAGGCGCCCA-AAGGUUCCCUCAGCCUGG
UUGGCAAUCAGGUGUU-GAGUG-UAAGUGCAC-AAGGGAGCUUGACU-GUGA--------------------
--------------GACCGA---CGGGUCGAGCAGGGACGAAA-GUCGGGACUAGUGAUCCGGC--------
---GG-U-----GGCUU-GUG--------------------GA-AGCGCCGUCGCUCAACGG-AUAAAAGGU
ACCCCGGGG-AUAACAGGCUGAUCUUCCCCAAGAGUCCAUAUCGACGGGAUGGUUUGGCACCUCGAUGUCGG
CUCGUCGCAUCCUGGGGCUGGA-GUCGGUCCCAAGGGUUGGGCUGUUCGCCCAUUAAAGCGGUACGCGAGCU
GGGUUUAGAACGUCGUGAGACAGUUCGGUCCCUAUCCGCU--------------------G-UGC-------
-------------------------------GCGU--AGGAGUCUUGAGAAG-GGC-UGUCCCUAGUACGAG
AGGAC-CGGGACGGACGAACCUC-UGGU-G-UGCCAGUUGUCCUGCCAAGGGCAU-G-GCUGGUUGGCUACG
UUCGGGAGGGAUAACCGCUGAAAGCA-UC--UAAGCGGGAAGCCUGCUUCGA-G-AUG--AGGACUCC-CA-
--CCCCC-----------------------------------------------------------------
--------------------------UU--------------------------------------------
------------------------------------------------------------------------
------------------------------------------------------------------------
------------------------------------------------------------------------
```

```
------------------------------------------------------------------------
------------------------------------------------------------------------
------------------------------------------------------------------------
-------------------------------------------------GAGGGGU--UA-AGG------CU
CCCAGUAGACGACUGGGU-UGA-UAGGCC-GGA-UCUGGAAG-CAC-C----------------GU------
----GA-----------GGUGUGGAGGUG--A-CCGG-UACUAAUA---GGCCGAG-G-GCUUGUCCU----
------------------------------------------------------------------------
------------------------------------------------------------------------
------------------------------------------------------------------------
------------------------------------C------------
>LSUb_267747_PROAC/1-3079
--------------------------AAGUU----GU--GUAGUGCGAUC-GGUGGAUG-C-CUUGGC-ACC
AAG---AGCCGAUGAAGGACGUUG--UGACCUGCGAUAAGCCCUGGGGAGUUGGUGAGCGAGCU-----GUG
AUC-CGG-GGGUGUCCGAAUGGGGAAAC--------CUG-------GAAUGUCCGG-AGU-----------A
G-U-------------G---U----C----C---GGU----G-G-CCCU-GCC-----CUGA----------
-----------A-----------UGUA----------------------------------------UA-GG
G-U--------------------GU-G-GG-UGGUAACGCGGGGAAGUGAAACAUCUUAGUACCCG--UAGG
AAGAGAAAACAA--GUGUGAUUCCGUGA--GU-AG-UGGCGAGCGAAAGCGGA-G-GA-GGCUAAACCGUGU
-GUG-UGUUCAAGCCGG-CAG------------G---UGU-UG-CAUGUGCGGGG--UUG------------
-----UGGG-GGC-------CUUG-UGGUUC--UGCUGCCGCAGGAUUGGC--CAGUGAGAA---AUGU---
U-------GCGUGAAGGUG------AAG-CG-UCUG------------------------------------
-------GGA-AG-GCGUA-------CCGGAGUGGGUGAGAGUCCUGUA---------ACUGUAA----GCG
U------G--GCACU-----------GGUG-UGGGG--UU-GC------CCCGAGUAGCGUGGGACUCGUGG
AAUUUCGCGUGAA-UCUGGCGGG-ACC--ACCCGUCAAGCCUAAAUA--CUC-CUUGG-UGACCGAUAGU-G
GAU-G-AGUACCGUGAGGGAAGG-GUGAAA-AGUAC-CCCGG-GAGGGG-A-GUGAAAGAGUUCCUGAAACC
G-GUCGCAUGCAA-GCC-GU-CAGA---GCC-----------------------------------------
------------------------------------------------------------------------
------------------------------------------------------------------------
------------------------------------------------------------------------
------------------------------------------------------------------------
------------------------------------------------------------------------
------------------------------------------------------------------------
------------------------------------------------------------------------
------------------------------------------------------------------------
-------------------------------------------------------------UG--U-GU---
------------------------------------------------------------------------
------------------------------------------------------------------------
------------------------------------------------------------------------
------------------------------------------------------------------------
------------------------------------------------------------------------
------------------------------------------------------------------------
------------------------------------------------------------------------
------------------------------------------------------------------------
------------------------------------------------------------------------
------------------------------------------------------------------------
```

```
------------------------------------------------------------------------
------------------------------------------------------------------------
----------------------------------------------------G---GUGAUGGCGUGCCUUU
UGAAGAAUGAGCCUGCGAGUUAGUG-AUGCGUGGCGA-GGUUAACC--C----------GUGGU--------
--GG--GGGA--GUCGUAGCGAAAG-C---------------------------------------------
------------------------------------------------------------------------
------------------------------------------------------------------------
---------------------------------------------------------------------GAG
UCC------------------------------------GAUAA-------------------------GG-
G-C-GU-----------------------GAGUCGCGUGUUCUA-GACCCGAAGCG-GUGUGAUCUAUCCAU
GGCCAGGAUGAAGCGUCG-GUAAGA-CGUCGUGGAGGUCCGA--ACCCACUU-CAGUUGAAAAUGGA-GGGG
AUGAGUUGUGGAUAGGGGUGAAAGGCCAAUCAAACACCG-UGAUAGCUGGUUCUCCCCGAAA-UGCAUUUAG
GUGCAGCGUUGC-GUG--------------------------------------------------------
----------------------GU-----UC-UUGUUGGAGGUAGAGC-ACUGGAU-GGU-CUA--GGGGGC
U---UA-UC--GGCUUACCGAAAUCAGCCAAACUCCGAAUGCUGGC--------------------------
------------------------------------------------------------------------
------------------AAGU-G-UA--------GCGUGGC-AGUGAGA-C-GGCG-G-GGGAUAAGCUUC
GU-C-GUCGAGAG-GGAAAC---AGCCCAGAUCAUCA-GCUAAGGCCCCUAAGU--GG-UGA-CU-CA---G
-U-GGG-------AAAGGAUGUGGAGUUGCGUAGACAACCAGGAGGUUGGCUUGGAAGCAGCCA-UCCUUGA
AAGAGUGCGUAAUAGCUCACUGGUC-----AAGUGAUUCCGCGCCGAUAAUGUAGC-GGGGCUUA-AGUUAU
C----------CGCCGAA-GCUGUGGCAACCUGU--------------------------------------
------------------------------------------------------------------------
------------------------------------------------------------------------
-----------------------------------GU-GGUU-GGGUAGGGGAGC-GUCGUGU---------
-----------C-A-GUGGU-GAAGCUG--C-CGGGUGACUG-UG-UGGUGGAGUU-G-AUG-CGAGUGAGA
AUGC-AGGCAU-GAGUA-GCGUGU----GAC-GGGUGG--G---------------AAUCCC-GUC-CGCCG
A-UUAUCC-AAGGGUUCC-AGGGUCAAG-UUUGUCUGCCCUGGGUGAGUCGGG-UC-CUAAGGCGAGGCCG-
A-C-AGGC-----GUA--------------------------------------------------------
----------------------------------------GUC-GA-U--GGGU-AACCAGUU-GAUA-UUC
UGGUA----------------------------------CCGGC----UUGU--------C-AC--------
------------------CGUCCGUGUCGAGGUGUGUGAUGCUAAGCAUGUGAGCUUGUCUUGGGAGUGUUU
UUGAUGCUUCUGGGGUGGGUGAGUGUGUGAA--CCGAUCAUGUAGUAGGCAA------GCUGC-GGAGGGAC
GCAGUG--------------AGGUAGCUCAUCCCCG-GU-G----AUGGU------------UGUCCGGGG-
CUAAGU-GUG-UG-----G-A-------------------C------UG------------------UC-UG
G-UAGG--------------------UAAAU-CCGCUGGGCG------UGAAGGUUG----AGG-CA-U-G-
AUG--GG---------------------------------------------------------------GA
-----GCCCAU-GGUUGUGGGUGAG-U-GAG-U-GAUCCUGUACUGU-C-GAGAAA-AGCUU-CGUGAGCGA
-------------------------------------------------------GGUGGCG-GG-UCCGCC
CGUACCCUAAACCGAC------------------------------ACUGGUG--GAUAGGUA-GAGUAUAC
CGAGGC-G-AU-CGAG-UGAAUCGU-G-GUGAAGGAACUCGGC-AAAAUGCCCCCGUAACUUUGG-GAUAAG
GGGGA-CCUG-A-GC--UGUCGUGC----ACC-----------CGUGC------------------------
------------------------------------------------------------------------
------------------------------------------------------------------------
------------------------------------------------------------------------
------------------------------------------------------------------------
```

```
------------------------------------------------------------------------
------------------------------------------------------------------------
------------------------------------------------------------------------
------------------------------------------------------------------------
------------------------------------------------------------------------
------------------------------------------------------------------------
------------------------------------------------------------------------
------------------------------------------------------------------------
------------------------------------------------------------------------
----------------------------------------------------------GGUGUGGUGGCGGU
GAG--GGGCGCAGAGUCCAGGGGGAAACGA-----CUGUUUACCAAAAACACAGGUCCGUGCGAAGUCG---
-UA-A-------------GACGAUGU-AUACGGACUGA-CUCCUGCCCGGUGCUG-G-AAGG-UUAAGGGGA
ACUGUGA-GU-----GUAU-----GCGUAGCGGUGAACUUAAGCCC--CAGUAAAC-GGCGGUGGUAACUAU
AACCAUCCUAAGGUAGCGAAAUUCCUUGUCGGG-UAAGUUCCGACCUGCACGAAUGGAGUAACGAUU--UCC
CUACUGUCUCCACC----AUG-AACUCGGUGAAAUU--GCA---UUACGAGUAAAGAUGCUCGUUACGCGCA
GCAGGACGGAAAGA-CCCCG-GGACC-UUUACU--AUAGUUU-GGUAUUGGUGAUUGGGACGGUUUGUGUAG
GAU-AGGUGGGAGACGUUG----------AUGCGGC-CAC----------------------------GCUA
--------------------------------------GU-GGUGGUGGAG-U--C-GU-UGGUGAAAUA-C
CACU-CUGAUUGUUCUGGUUAUCUCACUGUGG--ACCG-U-GA-U-C-------------------------
------------------------------------------------------------------------
------------------------------------------------------------------------
------------------------------------------------------------------------
------------------------------------------------------------------------
------------------------------------------------------CG----GU----------
------------------------------------------------UCAA-----GGACA-GUGCCUGAU
GGGUAGUUUGACUGGGGCGGU-CGCCUCCUA-AAGGGUAAC-GGAGGCGCCCA-AAGGUUCCCUCAGCCUGG
UUGGUAAUCAGGUGGU-GAGUG-UAAGUGCAC-AAGGGGGCUUGACU-GUGA--------------------
--------------GACUGA---CAGGUCGAGCAGGGACGAAA-GUCGGGACUAGUGAUCUGAC--------
---GG-U-----GGCUU-GUG--------------------GA-UGCGCCGUCACUCAACGG-AUAAAAGGU
ACCCCGGGG-AUAACAGGCUGAUCUUGCCCGAGCGCUCAUAGCGACGGCAUGGUUUGGCACCUCGAUGUCGG
CUCGUCGCAUCCUGGGGCUGGA-GUCGGUCCCAAGGGUUGGGCUGUUCGCCCAUUAAAGCGGCACGCGAGCU
GGGUUCAGAACGUCGUGAGACAGUUCGGUCCCUAUCCGCU--------------------G-CGC-------
-------------------------------GCGG--AGGAAUCUUGAGAAG-GGC-UGUCUUUAGUACGAG
AGGAC-CGAGACGGACUGACCUC-UGGU-G-UGCCAGUUGUUCUGCCAAGAGCAU-G-GCUGGUUGGCUACG
UCGGGUUGUGAUAACCGCUGAAAGCA-UC--UAAGCGGGAAGCACGCUUCAA-G-AUA--AGGGUUCC-CA-
--CAGGA-----------------------------------------------------------------
--------------------------GU----------G---------------------------------
------------------------------------------------------------------------
------------------------------------------------------------------------
------------------------------------------------------------------------
------------------------------------------------------------------------
------------------------------------------------------------------------
------------------------------------------------------------------------
-------------------------------------------------UUCCUGG--UA-AGG------UC
CCCGAGAGAUGAUCGGGU-UGA-UAGGCC-AGA-CGUGGACG-CAU-C----------------GU------
```

```
----AA-----------GGUGUGGAGCUG--A-CUGG-UACUAAUA---GACCGAG-G-ACUUUCAUU----
------------------------------------------------------------------------
------------------------------------------------------------------------
------------------------------------------------------------------------
------------------------------------A------------
>LSUb_272631_MYCLE/1-3122
---------------------A-AGUAAGUG----CC--UAAGGGCGCAU-GGUGGAUG-C-CUUGGC-AUC
GAG---AGCCGAUGAAGGACGUGG--GAGGCUGCGAUAUGCCUCGGGGAGCUGUCAACCGAGCG-----UGG
AUC-CGA-GGAUUUCCGAAUGGGGAAAC--------CC--------AA----CACG-AGU-----------C
A-U-------------G---U----C----G---UGU----U-A-CCCG-UAU-----CUGA----------
-----------A-----------UAUA----------------------------------------UA-GG
G-U--------------------UC-G-GG-AGGGAACGCGGGGAAGUGAAACAUCUCAGUACCCG--UAGG
AGAAGAAAACAA--UUGUGAUUCCGCAA--GU-AG-UGGCGAGCGAACGUGGA-A-UAUGGCUAAACCGCAC
-ACA-UGUCUAACUAGG-UAG------------G---GGU-UG-UGUGUGCGGUG--UUG------------
-----UGGG-AUU-------GGUA-UGUCUCAACUCUACCUGGUUGAGGGG--UAGUCAGAAA--GUGC--C
G-------UGGUU-AGCGG-A----AAU-GG-CCUG------------------------------------
-------GGA-UG-GCCUG-------CCGUAGACGGUGAGAGCCAGUA----------CGCGAAA----GCC
U------G--GCACC----------UGCCUUGUAUC--AA-UU------CCCGAGUAGCAGCGGGCCCGUGG
AAUCUGCUGUGAA-UCUGCCGGG-ACC--ACCCGGUAAGCCUAAAUA--CUU-CUCGA-UGACCGAUAGC-G
GAU-C-AGUACCGUGAGGGAAUG-GUGAAA-AGUAC-CCCGG-GAGGGG-A-GUGAAAUAGUACCUGAAACC
G-UGUGCCUACAA-UCC-GU-CAGA---GCCU-C--------------------------------------
------------------------------------------------------------------------
------------------------------------------------------------------------
------------------------------------------------------------------------
------------------------------------------------------------------------
------------------------------------------------------------------------
------------------------------------------------------------------------
------------------------------------------------------------------------
------------------------------------------------------------------------
-------------------------------------------------------------UU--U-GU---
------------------------------------------------------------------------
------------------------------------------------------------------------
------------------------------------------------------------------------
------------------------------------------------------------------------
------------------------------------------------------------------------
------------------------------------------------------------------------
------------------------------------------------------------------------
------------------------------------------------------------------------
------------------------------------------------------------------------
------------------------------------------------------------------------
------------------------------------------------------------------------
------------------------------------------------------------------------
------------------------------------------------GG--G---GUGAUGGCGUGCUUUU
UGAAGAAUGAGCCUGCGAGUCAGGG-ACAUGUCGCGA-GGUUAACC--C-----------GUGU--------
--GG--GGUA--GCCGCAGCGAAAG-C---------------------------------------------
```

```
------------------------------------------------------------------------
------------------------------------------------------------------------
---------------------------------------------------------------------GAG
UCU------------------------------------GAAUA-------------------------GG-
G-C-GU---AUCACGUGUGAG--CGUGUGUAGUGGCGUGUUCUG-GACCCGAAGCG-GAGUGAUCUACCCAU
GGCCAGGGUGAAGCGCGG-GUAAGA-CCGCGUGGAGGCCCGA--ACCCACUU-AAGUUGAAGAUUGA-GGGG
AUGAGUUGUGGGUAGGGGUGAAAGGCCAAUCAAACUCCG-UGAUAGCUGGUUCUCCCCGAAA-UGCAUUUAG
GUGCAGCGUUGC-GUG--------------------------------------------------------
----------------------GU-----UC-ACCACGGAGGUAGAGCUACUGGAU-GGC-CGA--UGGGCC
C---UA-CU--AGGUUACUGACGUCAGCCAAACUCCGAAUGCCGUG--------------------------
------------------------------------------------------------------------
------------------GUUUAA-AA--------GCGUGGC-AGUGAGA-C-GGCG-G-GGGAUAAGCUCC
GUAC-GUCGAAAG-GGAAAC---AGCCCAGAUCGCCG-GCUAAGGCCCCUAAGC--GU-GUG-CU-AA---G
-U-GGA-------AAAGGAUGUGCAGUCGCAAAGACAACCAGGAGGUUGGCUUAGAAGCAGCCA-CCCUUGA
AAGAGUGCGUAAUAGCUCACUGGUC-----AAGUGAUUGUGCGCCGAUAAUGUAGC-GGGGCUCA-AGCACA
C----------CGCCGAA-GCCGCGGCACAUUCA--C-----------------------------------
------------------------------------------------------------------------
-----CUU----------------------------------------------------------------
----------------------------CUAGGGUGG-AUGU-GGGUAGGGGAGC-GUUCCUC----------
-----------A-U-UCAGC-GAAGCCU--C-CGGGUAACCG--G-UGGUGGAGGG-U-GGG-GAAGUGAGA
AUGC-AGGCAU-GAGUA-GCGAUA---AGGC-AAGUGA--G---------------AACCUU-GCC-CGCCG
U-AAGACC-AAGGGUUCC-UGGGCCAGG-CCAGUCCGCCCAGGGUGAGUCGGG-AC-CUAAGGCGAGGCCG-
A-C-AGGC-----GUA--------------------------------------------------------
----------------------------------------GUC-GA-U--GGAC-AACGGGUU-GAUA-UUC
CCGUA----------------------------------CCCGU----GUGU--------G-UG--------
------------------CGCCCGUGAUGAAUCAGCGGUACUCACCACCCAAAACCGGAUCGACCAUAUCCC
CUUCGGGGGCUAUGGAGGUUCGGGGCUGCGU--GGGAACUUCGUUGGUAGUAGUCAA-GCGAU-GGGGUGAC
GCAGGA--------------AGGUAG-CCGUACCAG-UC-A----GUGGU------------AAUACUGGA-
GCAAGC-CCG-UA-----G-G-------------------G------AG------------------AG-CG
A-UAGG--------------------CAAAU-CCGUCGUUCG------UGAAUCCUG----AGA-GG-U-G-
AUG--CA---------------------------------------------------------------UA
-----GCCGAU-----UGAGGCGAA-U-UCG-G-UAAGCCUCUGCUG-C-CAAGAA-AAGCC-UCUAGCGA-
--------------------------------------------------------GCAUAC-AU-GCGGCC
CGUACCCCAAACCGAC------------------------------ACAGGUG--GUCAGGUA-GAGCAUAC
CAAGGC-G-UA-CGAG-AUAACUAU-G-GUUAAGGAACUCGGC-AAAAUGCCUCCGUAACUUCGG-GAGAAG
GGGGG-CCGG-A-AU-AUCGUGAAC----ACC-----------CUUGC------------------------
------------------------------------------------------------------------
------------------------------------------------------------------------
------------------------------------------------------------------------
------------------------------------------------------------------------
------------------------------------------------------------------------
------------------------------------------------------------------------
------------------------------------------------------------------------
------------------------------------------------------------------------
------------------------------------------------------------------------
```

```
------------------------------------------------------------------------
------------------------------------------------------------------------
------------------------------------------------------------------------
------------------------------------------------------------------------
--------------------------------------------------------GGUG-GGA--GCGGGA
UCC--GGUCGCAGAGACCAGUGAGAAGCGA-----CUGUUUACUAAAAACACAGGUCCGUGCGAAGUCG---
-UA-A-------------GACGAUGU-AUACGGACUGA-CGCCUGCCCGGUGCUG-G-AAGG-UUAAGAGGA
CCUGUUA-ACCC---GAAA---GGGUGAAGCGGAGAAUUUAAGCCC--CAGUAAAC-GGCGGUGGUAACUAU
AACCAUCCUAAGGUAGCGAAAUUCCUUGUCGGG-UAAGUUCCGACCUGCACGAAUGGCGUAACGACU--UCU
CAACUGUCUCAACC----AUA-GACUCGGCGAAAUU--GCA---CUACGAGUAAAGAUGCUCGUUACGUGCG
GCAGGACGAAAAGA-CCCCG-GGACC-UUCACU--ACAACUU-GGUAUUGAUGUUCGGUGCGGUUUGUGUAG
GAU-AGGUGGGAGACUGUG----------AAACUUC-GAC----------------------------GCUA
--------------------------------------GU-UGGGGUGGAG-U--C-GU-UGUUGAAAUA-C
CACU-CUGAUUGUAUUGAACAUCUAACCUCGA--ACCG-UAUA-U-C-------------------------
------------------------------------------------------------------------
------------------------------------------------------------------------
------------------------------------------------------------------------
------------------------------------------------------------------------
------------------------------------------------------CG----GU----------
------------------------------------------------UUAG-----GGACA-GUGCCUGGC
GGGUAGUUUAACUGGGGCGGU-UGCCUCCUA-AAAUGUAAC-GGAGGCGCCCA-AAGGUUCCCUCAACCUGG
UCGGUAAUCAGGUGGC-GAGUG-UAAAUGCAC-AAGGGGGCUUGACU-GCGA--------------------
--------------GACCUA---CACGUCAAGCAGGACGAAA-GUCGGGAUUAGUGAUCCGGC---------
---AC-C-----UCUGA-GUG--------------------GA-AGGGGUGUCGCUCAACGG-AUAAAAGGU
ACCCCGGGG-AUAACAGGCUGAUCUUCCCCAAGAGUCCAUAUCGACGGGAUGGUUUGGCACCUCGAUGUCGG
CUCGUCGCAUCCUGGGGCUGAA-GCAGGUCCCAAGGGUUGGGCUGUUCGCCCAUUAAAGCGGCACGCGAGCU
GGGUUUAGAACGUCGUGAGACAGUUCGGUCUCUAUCCGCC--------------------G-CGC-------
-------------------------------GCGU--CAGAAACUUGAGGAA-AUC-UGUCCCUAGUACGAG
AGGAC-CGGGACGGACGAACCUC-UGGU-A-UACCAGUUGUCUCACCAGGGGCAC-C-GCUGGAUAGCCACG
UUCGGACAAGAUAACCGCUGAAAGCA-UC--UAAGCGGGAAACCUUCUCCAA-G-AUC--AGGUUUCU-UA-
--CCCAC-----------------------------------------------------------------
--------------------------UU--------------------------------------------
------------------------------------------------------------------------
------------------------------------------------------------------------
------------------------------------------------------------------------
------------------------------------------------------------------------
------------------------------------------------------------------------
------------------------------------------------------------------------
-------------------------------------------------GGUGGGA--UA-AGG------CC
CCCCGCAGAACACGGGAU-UGA-UAGGUC-AGA-CCUAGAAG-CUC-A----------------GU------
----AA-----------UGAGUGUAGGGA--A-CUGG-CACUAACU---GGCCGAA-A-GCUUACAAA----
------------------------------------------------------------------------
------------------------------------------------------------------------
------------------------------------------------------------------------
------------------------------------A------------
```

```
>LSUb_243274_THEMA/1-3021
---------------------G-GUCAAGGU----AC--UAAGGGCACGC-GGUGGAUG-C-CUUGGC-GGC
GGG---AGGCGAUGAAGGGCGUGG--CAAGCUGCGAUAAGCCCGGGGGAGCCGCAAGCAGGCGU------UG
AUC-CCG-GGAUUCCCGAAUGGGGAAAC--------CU--------GG----GUGG-C-CG----------U
U-G------------AG---G----C----C---ACU----C-G-CCGCCGAA-------------------
-----------A-----------GGC----------------------------------------------
G-G--------------------UG-C-GA-----CACCGGGGGAAGUGAAACAUCUCAGUACCCC--GAGG
AAAAGAAAUCAA--CCGAGAUUCCCCUA--GU-AG-CGGCGAGCGAACGGGGA-G-GA-GCCCAAACCAGCG
-UGG-UGUCAAAGCCCG-UGG------------G---CGU-UG-CCACGCUGGGG--UUG------------
-----UGGG-ACA-------CGGC-CGGGCGA-GGCCACGGACUCGCCGGG--GAGUUACAAA--UCCC---
U-------CCUCU-AGCCG------AAC-CA-CCUG-------------------------------------
-------GGA-AG-GUGGG-------CCGGAGAGGGUGACAGCCCCGUA---------GGCGAAA----GGG
G------A--GGGAC----------UCCCUGGGCCG--UG-CU------CCCGAGUACCACGGGACACGUGG
AAUCCCGUGGGAA-GCUGGGGGG-ACC--ACCCUCCAAGGCUAAAUA--CUA-CCCGC-CGUCCGAUAGC-G
CAC-C-AGUACCGUGAGGGAAAG-GUGAAA-AGCAC-CCCGG-AAGGGG-A-GUGAAAGAGGACCUGAAACC
G-CGUGCCUACAA-GAA-GU-CGGA---GCCC-C---C----------------------------------
------------------------------------------------------------------------
------------------------------------------------------------------------
------------------------------------------------------------------------
------------------------------------------------------------------------
------------------------------------------------------------------------
------------------------------------------------------------------------
------------------------------------------------------------------------
------------------------------------------------------------------------
--------------------------------------------------------------A--U-UU---
------------------------------------------------------------------------
------------------------------------------------------------------------
------------------------------------------------------------------------
------------------------------------------------------------------------
------------------------------------------------------------------------
------------------------------------------------------------------------
------------------------------------------------------------------------
------------------------------------------------------------------------
------------------------------------------------------------------------
------------------------------------------------------------------------
------------------------------------------------------------------------
------------------------------------------------------------------------
------------------------------------------------------------------------
-----------------------------------------------GGG--G---GUGACGGCGUGCCUUU
UGAUUAAUGAGCCCGCGAGUUGC-C-GUCGGUGGCGA-GGUUAAGC--C-G---------ACGA--------
A-GG--CGUA--GCCGUAGCGAAAG-C---------------------------------------------
------------------------------------------------------------------------
------------------------------------------------------------------------
---------------------------------------------------------------------GAG
UCC------------------------------------GAACA-------------------------GG-
G-C-GC----------------------UCAGUCACCGGCGGCA-GACCCGAAGCC-GGGUGAGCUACCCCU
```

```
GGGCAGGGUGAAGGUGGG-UUAAAA-CCCACUGGAGGCCCGA--ACCGGUGG-ACAGUGAAAAGUCC-UCGG
AUGACCUGGGGGUAGGAGUGAAAAGCUAACCGAACCCGG-UGAUAGCUGGUUCUCCCCGAAA-UGCAUUGAG
GUGCAGCUCGG-GCG---------------------------------------------------------
----------------------GU-----CU-GUGCAGGGGGUAGAGC-ACUGAUG-GGG-CUA--GGGGG-
----GU-UU----CCU-CCGAACCCGUCAAACUCCGAAUCCCUGC---------------------------
------------------------------------------------------------------------
------------------ACAC-A-AA--------GCCCGGG-AGUGAGC-C-CGCG-G-GGGAUAAGCUCC
GC-G-GACGAGAG-GGGAAC---AACCCAGACCGCCG-GCUAAGGGCCCGAAGA--GG-UGG-CU-AA---G
-U-GGU-------AAAGGAUGUGGAGCGCCUAAGACAGCUGGGAUGUUGGCUUAGAAGCAGCCA-UCAUUUA
AAGAGUGCGUAACAGCUCACCAGCC-----GAGGCGCUCUGCACCGAAAAUGUAAC-GGGGCUCA-AGCCAC
C----------CCCCGAA-GCCGCGGGUCAGUAC--C-----------------------------------
------------------------------------------------------------------------
-----UCCG---------------------------------------------------------------
----------------------------CUGGAGGUA-CUGG-CGGUAGGGGAGC-UUUCCGC---------
-----------UGU-AGGGU-GAAGGCG--GACCCGCGAGGG--C-CGCUGGACGA-G-GCG-GAAGUGAGA
AUGC-GGGCAU-GAGUACGCGA-A---AGGA-GGGUGA--G---------------AAUCCC-UCC-CCCCG
U-AAGCCC-AAGGGUACC-UGGGGAAGG-UUCGUCCGCCCAGGGUUAGCCGGGACC-CUAAGGUGAACCCG-
A-A-AGGG-----GUA--------------------------------------------------------
----------------------------------------GCC-GA-A--GGGA-AGCCGGUU-AAUA-UUC
CGGCG----------------------------------CCACC----UGCG--------G-UC--------
------------------GAU---------------------------------------------------
--------------------------------------------------------GGUACGA-GGGGUGAC
GCAGGA--------------GGGUAG-GCGCCGGGG-GC-A----AGUGG------------CAUGCCCCU-
CCAAGC-GGU-UA-----G-G-------------------G------CG-------------AUGACGG-GC
G-UAGG--------------------UAAAU-CCGCGCCCG----------AGCCUG----AGC-CG-U-G-
AUG--GG---------------------------------------------------------------GA
----GGUCCCU-UCGGGGACCAACG-G-CGC-U-GACCCCACACUGC-C-GAGAAA-AACCU-CGCGUACCG
-----------------------------------------------CU-----UGAGACCG-UA-GGUGCC
CGUUCCGCAAACCGAC------------------------------ACAGGUG--GGCGGGCU-GAGAAGGC
UCAGGG-G-AG-CGGG-UUAACCCU-C-GCCAAGGAACUCGGC-AAAUUGGCCCCGUAACUUCGG-GAGAAG
GGGUG-CCGC-G-UU-AGGGUGAAC----CCAGG-GGAAGGGGCAACCCGGAAA------------------
------------------------------------------------------------------------
------------------------------------------------------------------------
------------------------------------------------------------------------
------------------------------------------------------------------------
------------------------------------------------------------------------
------------------------------------------------------------------------
------------------------------------------------------------------------
------------------------------------------------------------------------
------------------------------------------------------------------------
------------------------------------------------------------------------
------------------------------------------------------------------------
------------------------------------------------------------------------
------------------------------------------------------------------------
----------------------------------------------------------CUGGGGGAGCCCGA
```

```
GGC--GGUCGCAGUGACAAGGCCCUGGCGA-----CUGUUUACCAAAAACACAGGUCUCUGCUAACUCGA--
-AA-A-------------GAGGAAGU-AUAGGGACUGA-CGCCUGCCCAGUGCCG-G-AAGG-UUAAGGGGA
GGGGUGAGGCUCCCCU-UGUGGGAGCGUAGCUCCGACCCGAAGCCC--CGGUAAAC-GGCGGCCGUAACUAU
AACGGUCCUAAGGUAGCGAAAUUCCUUGUCGGG-UAAGUUCCGACCUGCAUGAAUGGCGUAACGACU--GGG
GCACUGUCUCGGCG----GGG-GGCCCGGCGAAAUU--UCA---GUCUGGGUGAAGAUGCCCAGUACCCGCG
GCUAGACGGAAAGA-CCCCGUGGAGC-UUUACU--GCAGCCU-GGUAUUGGGCUCUGGUGCAUCGUGUAUAG
CAU-AGGUGGGAGGCUGUG----------AAGCCGC-CUC----------------------------GCCA
--------------------------------------GG-GGCGGUGGAG-C--C-GC-CAGUGGAAUA-C
CACC-CUCGGUGCACUGGAGUCCUAACCUGAC--CCUGUG-AA-G-A-------------------------
------------------------------------------------------------------------
------------------------------------------------------------------------
------------------------------------------------------------------------
------------------------------------------------------------------------
-----------------------------------------------------GCA----GG----------
------------------------------------------------GUGG-----GGACA-GUGCCAGGU
GGGCAGUUUGACUGGGGCGGU-CACCUCCUA-AAAGGUAAC-GGAGGUGUGCA-AAGGUCGGCUCAGGUGGG
UUGGAAAUCCACCGCA-GAGUG-CAAGGGCAU-AAGCCGGCCUGACU-GCGA--------------------
--------------GGCCGA---CAGGCCGAGCAGGGGGGAAA-CCCGGCCCUAGUGACCCGGC--------
---GG-U-----CCCGU-GUG--------------------GA-AGGGCCGUCGAUCAACGG-AUAAAAGUU
ACCCCGGGG-AUAACAGGCUGGUCCCGCCCGAGAGUUCACAUCGACGGCGGGGUUCGGCACCUCGAUGUCGG
CUCAUCCCAUCCUGGGGCUGAA-GCAGGUCCCAAGGGUUGGGCUGUUCGCCCAUUAAAGGGGUACGUGAGCU
GGGUUCAGACCGUCGUGAGACAGGUCGGUCCCUAUCUGCC--------------------G-CGG-------
-------------------------------GCGU--AGGAGGCUUGAGGGG-GGU-CCUCCCUUGUACGAG
AGGAC-CGGGAGGAGGGGGCCUC-UGGU-G-UACCGGCUGUCGCGCCAGCGGCAU-ACGCCGGGUAGCUACG
CCCCUAAGCGAUAACCGCUGAAAGCA-UC--UAAGCGGGAAGCGCGCCCCAA-G-AUU--AGGCCUCC-CA-
--UCCCG-----------------------------------------------------------------
--------------------------UU--------------------------------------------
------------------------------------------------------------------------
------------------------------------------------------------------------
------------------------------------------------------------------------
------------------------------------------------------------------------
------------------------------------------------------------------------
------------------------------------------------------------------------
-------------------------------------------------AAGGGAG--UA-AGG------CC
GGUCCGAGAAGAGGACCU-UGA-UAGGCC-GGG-GGUGUAAG-CGC-C----------------GC------
----AA------------GCGUUGAGCCC--A-CCGG-UACUAAUC---GGCCGAG-G-CCUUGACCU----
------------------------------------------------------------------------
------------------------------------------------------------------------
------------------------------------------------------------------------
------------------------------------C------------
>LSUb_262724_THET2/1-2911
---------------------G-GUCAAGAU----GG--UAAGGGCCCAC-GGUGGAUG-C-CUCGGC-ACC
C-G---AGCCGAUGAAGGACGUGG--CUACCUGCGAUAAGCCAGGGGGAGCCGGUAGCGGGCGU------GG
AUC-CCU-GGAUGUCCGAAUGGGGGAAC--------CC--------GG----CCGG-C-G----------GG
A-A-------------C---G----C----C---GGU----C-A-CCGC-GC--------------------
```

```
-----------U-----------UUU----------------------------------------------
--G--------------------CG-C-GG-GGGGAACCUGGGGAACUGAAACAUCUCAGUACCCA--GAGG
AGAGGAAAGAGA--AAUCGACUCCCUGA--GU-AG-CGGCGAGCGAAAGGGGA-C-CA-GCCUAAACCGUC-
--CG-GCUU---------------------------------G-UCCGGGCGGG--UCG-------------
-----UGGG-GCC-------CUCG-GA----------------------CA--CCG--------AAUCC--C
C-------AGCCU-AGCCG------AAG-CU-GUUG------------------------------------
-------GGA-AG-CAGCG-------CCAGAGAGGGUGAAAGCCCGUA----------GGCGAAA----GGU
G------G--GGGGA----------UAGG---UGAG--GG-UA------CCCGAGUACCCCGUGGUUCGUGG
AGCCAUGGGGGAA-UCUGGGCGG-ACC--ACCGCCUAAGGCUAAGUA--CUC-C-GGG-UGACCGAUAGC-G
CAC-C-AGUACCGUGAGGGAAAG-GUGAAA-AGAAC-CCCGG-GAGGGG-A-GUGAAAUAGAGCCUGAAACC
G-UGGGCUUACAA-GCA-GU-CACG---GCCC-C--------------------------------------
------------------------------------------------------------------------
------------------------------------------------------------------------
------------------------------------------------------------------------
------------------------------------------------------------------------
------------------------------------------------------------------------
------------------------------------------------------------------------
------------------------------------------------------------------------
------------------------------------------------------------------------
--------------------------------------------------------------G--C-AA---
------------------------------------------------------------------------
------------------------------------------------------------------------
------------------------------------------------------------------------
------------------------------------------------------------------------
------------------------------------------------------------------------
------------------------------------------------------------------------
------------------------------------------------------------------------
------------------------------------------------------------------------
------------------------------------------------------------------------
------------------------------------------------------------------------
------------------------------------------------------------------------
------------------------------------------------------------------------
------------------------------------------------GG--G---GUUGUGGCGUGCCUAU
UGAAGCAUGAGCCGGCGACUCAC-G-GUCGUGGGCGA-GCUUAAGC--C----------GUUGA--------
--GG--CGGA--GGCGUAGGGAAAC-C---------------------------------------------
------------------------------------------------------------------------
------------------------------------------------------------------------
---------------------------------------------------------------------GAG
UCC------------------------------------GAACA-------------------------GG-
G-C-GCAAGCGGGCCGCACGCGGCCCGCAAAGUCCGCGGCCGUG-GACCCGAAACC-GGGCGAGCUAGCCCU
GGCCAGGGUGAAGCUGGG-GUGAGA-CCCAGUGGAGGCCCGA--ACCGGUGG-GGGAUGCAAACCCC-UCGG
AUGAGCUGGGGCUAGGAGUGAAAAGCUAACCGAGCCCGG-AGAUAGCUGGUUCUCCCCGAAA-UGACUUUAG
GGUCAGCCUCAG-GCG--------------------------------------------------------
----------------------CU-----GA-CUGGGGCCUGUAGAGC-ACUGAUA-GGG-CUA--GGGGGC
C---CA-CC--AGCCUACCAAACCCUGUCAAACUCCGAAGGUCCC---------------------------
```

------------------------------------------------------------------------
------------------AGGU-G-GA--------GCCUGGG-AGUGAGG-G-CGCG-A-GCGAUAACGUCC
GC-G-UCCGAGCGCGGGAAC---AACCGAGACCGCCA-GCUAAGGCCCCCAAGU--CU-GGG-CU-AA---G
-U-GGU-------AAAGGAUGUGGCGCCGCGAAGACAGCCAGGAGGUUGGCUUAGAAGCAGCCA-UCCUUUA
AAGAGUGCGUAAUAGCUCACUGGUC-----GAGUGGCGCCGCGCCGAAAAUG-AUC-GGGGCUUA-AGCCCA
G----------CGCCGAA-GCUGCGGGUCUGGGG--GA----------------------------------
------------------------------------------------------------------------
------------------------------------------------------------------------
-------------------------------UGACCC-CAGG-CGGUAGGGGAGC-GUUCCCG---------
-----------AUG-CCGAU-GAAGGCC--GACCCGCGAGGG--C-GGCUGGAGGU-A-AGG-GAAGUGCGA
AUGC-CGGCAU-GAGUA-ACGAUA---AAGA-GGGUGA--G--------------AAUCCC-UCU-CGCCG
U-AAGCCC-AAGGGUUCC-UACGCAAUG-GUCGUCAGCGUAGGGUUAGGCGGG-AC-CUAAGGUGAAGCCG-
A-A-AGGC-----GUA--------------------------------------------------------
----------------------------------------GCC-GA-A--GGGC-AGCCGGUU-AAUA-UUC
CGGCC----------------------------------CUUCC----CGCA--------G-GU--------
------------------------------------------------------------------------
----------------------------------------------------------GCGAU-GGGGGGAC
GCUCUA--------------GGCUAG-GGGGACCGG-AGCC----AUGGA------------CGAGCCCGG-
CCAGAA-GCG-CA-----G-G-------------------G------UG------------------GG-AG
G-UAGG--------------------CAAAU-CCGCCUCCCA------ACAAGCUC------UG-CG-U-G-
GUG--GG---------------------------------------------------------------GA
-----AGCCCG-UACGGGUGACAAC-C-CCC-C-GAAGCCAGGGAGC-C-AAGAAA-AGCCU-CUAAGCAC-
--------------------------------------------------------AACCUG-CG-GGAACC
CGUACCGCAAACCGAC------------------------------ACAGGUG--GGCGGGUGCAAGAGCAC
UCAGGC-G-CG-CGGG-AGAACCCU-C-GCCAAGGAACUCUGC-AAGUUGGCCCCGUAACUUCGG-GAGAAG
GGGUG-CUCC-C-UG--GGGUGA-------------------------------------------------
------------------------------------------------------------------------
------------------------------------------------------------------------
------------------------------------------------------------------------
------------------------------------------------------------------------
------------------------------------------------------------------------
------------------------------------------------------------------------
------------------------------------------------------------------------
------------------------------------------------------------------------
------------------------------------------------------------------------
------------------------------------------------------------------------
------------------------------------------------------------------------
------------------------------------------------------------------------
------------------------------------------------------------------------
---------------------------------------------------------------UGAGCCCCG
GGG--AGCCGCAGUGAACAGGCUCUGGCGA-----CUGUUUACCAAAAACACAGCUCUCUGCGAACUCG---
-UA-A-------------GAGGAGGU-AUAGGGAGCGA-CGCUUGCCCGGUGCCG-G-AAGG-UCAAGGGGA
GGGGU----------GCAA----------GCCCCGAACCGAAGCCC--CGGUGAAC-GGCGGCCGUAACUAU
AACGGUCCUAAGGUAGCGAAAUUCCUUGUCGGG-UAAGUUCCGACCUGCACGAAAAGCGUAACGACC--GGA
GCGCUGUCUCGGCG----AGG-GACCCGGUGAAAUU--GAA---CUGGCCGUGAAGAUGCGGCCUACCCGUG

```
GCAGGACGAAAAGA-CCCCGUGGAGC-UUUACU--GCAGCCU-GGUGUUGGCUCUUGGUCGCGCCUGCGUAG
GAU-AGGUGGGAGCCUGUG----------AACCCCC-GCC----------------------------UCCG
--------------------------------------GG-UGGGGGGGAG-G--C-GC-CGGUGAAAUA-C
CACC-CUGGCGCGGCUGGGGGCCUAACCC-UC---------GG-A-U-------------------------
------------------------------------------------------------------------
------------------------------------------------------------------------
------------------------------------------------------------------------
------------------------------------------------------------------------
------------------------------------------------------------------------
------------------------------------------------GGGG-----GGACA-GCGCUUGGC
GGGCAGUUUGACUGGGGCGGU-CGCCUCCUA-AAAGGUAAC-GGAGGCGCCCA-AAGGUCCCCUCAGGCGGG
ACGGAAAUCCGCCGGA-GAGCG-CAAGGGUAG-AAGGGGGCCUGACU-GCGA--------------------
--------------GGCCUG---CAAGCCGAGCAGGGGCGAAA-GCCGGGCCUAGUGAACCGGU--------
---GG-U-----CCCGU-GUG--------------------GA-AGGGCCAUCGAUCAACGG-AUAAAAGUU
ACCCCGGGG-AUAACAGGCUGAUCUCCCCCGAGCGUCCACAGCGGCGGGGAGGUUUGGCACCUCGAUGUCGG
CUCGUCGCAUCCUGGGGCUGAA-GAAGGUCCCAAGGGUUGGGCUGUUCGCCCAUUAAAGCGGCACGCGAGCU
GGGUUCAGAACGUCGUGAGACAGUUCGGUCUCUAUCCGCC--------------------A-CGG-------
-------------------------------GCGC--AGGAGGCUUGAGGGG-GGC-UCUUCCUAGUACGAG
AGGAC-CGGAAGGGACGCACCUC-UGGU-U-UCCCAGCUGUCCCUCCAGGGGCAU-AAGCUGGGUAGCCAUG
UGCGGAAGGGAUAACCGCUGAAAGCA-UC--UAAGCGGGAAGCCCGCCCCAA-G-AUG--AGGCCUCC-CA-
--CGGCG-----------------------------------------------------------------
--------------------------UC--------------------------------------------
------------------------------------------------------------------------
------------------------------------------------------------------------
------------------------------------------------------------------------
------------------------------------------------------------------------
------------------------------------------------------------------------
------------------------------------------------------------------------
-------------------------------------------------AAGCCGG--UA-AGG------AC
CCGGGAAGACCACCCGGU-GGA-UGGGCC-GGG-GGUGUAAG-CGC-C----------------GC------
----GA-----------GGCGUUGAGCCG--A-CCGG-UCCCAAUC---GUCCGAG-G-UCUUGACC-----
------------------------------------------------------------------------
------------------------------------------------------------------------
------------------------------------------------------------------------
-------------------------------------------------
>LSUb_243230_DEIRA/1-2877
---------------------G-GUCAAGAU----AG--UAAGGGUCCAC-GGUGGAUG-C-CCUGGC-GCU
-GG---AGCCGAUGAAGGACGCGA--UUACCUGCGAAAAGCCCCGACGAGCUGGAGAUACGCUU------UG
ACU-CGG-GGAUGUCCGAAUGGGGAAAC--------CC--------AC----CUC---------------GU
A-A---------------------------G---AGG----U-A-UCC------------------------
-----------G-----------CAA----------------------------------------------
--------------------------G-GAU-GGGAACUCAGGGAACUGAAACAUCUCAGUACCUG--AAGG
AGAAGAAAGAGA--AUUCGAUUCCGUUA--GU-AG-CGGCGAGCGAACCCGGA-U-CA-GCCCAAACCGAA-
--AC-GCUUG----------------------------------CGU-UUCGGGG--UUG------------
-----UAGG-ACC-------AGUU-UU----------------------UA--AGAU--------UCAA---
```

```
C-------CCCUCAAGCCG------AAG-UG-GCUG------------------------------------
-------GAA-AG-CUACA-------CCUCAGAAGGUGAGAGUCCUGUA---------GGCGAAC----GAG
C------G--GUUGA----------CUGU----ACU--GG-CA------CCUGAGUAGGUCGUUGUUCGUGA
AACGAUGACUGAA-UCCGCGCGG-ACC--ACCGCGCAAGGCUAAAUA--CUC--CCAG-UGACCGAUAGC-G
CA--U-AGUACCGUGAGGGAAAG-GUGAAA-AGAAC-CCCGG-GAGGGG-A-GUGAAAGAGAACCUGAAACC
G-UGGACUUACAA-GCA-GU-CAUG---GCAC----------------------------------------
------------------------------------------------------------------------
------------------------------------------------------------------------
------------------------------------------------------------------------
------------------------------------------------------------------------
------------------------------------------------------------------------
------------------------------------------------------------------------
------------------------------------------------------------------------
------------------------------------------------------------------------
-------------------------------------------------------------CU--U-AUC--
------------------------------------------------------------------------
------------------------------------------------------------------------
------------------------------------------------------------------------
------------------------------------------------------------------------
------------------------------------------------------------------------
------------------------------------------------------------------------
------------------------------------------------------------------------
------------------------------------------------------------------------
------------------------------------------------------------------------
------------------------------------------------------------------------
------------------------------------------------------------------------
------------------------------------------------------------------------
-------------------------------------------------G--U---GUUAUGGCGUGCCUAU
UGAAGCAUGAGCCGGCGACUUAGAC-CUGACGUGCGA-GCUUAAGU--U-----------GAAA--------
--AA--CGGA--GGCGGAGCGAAAG-C---------------------------------------------
------------------------------------------------------------------------
------------------------------------------------------------------------
---------------------------------------------------------------------GAG
UCC------------------------------------GAAUA-------------------------GG-
G-C-GG--------------------CAUUAGUACGUCGGGCUA-GACUCGAAACC-AGGUGAGCUAAGCAU
GACCAGGUUGAAACCCCC-GUGACA-GGGGGCGGAGGACCGA--ACCGGUGC-CUGCUGAAANAGUC-UCGG
AUGAGUUGUGUUUAGGAGUGAAAAGCUAACCGAACCUGG-AGAUAGCUAGUUCUCCCCGAAA-UGUAUUGAG
GUACAGCCUCGG-AUG--------------------------------------------------------
----------------------UU-----GA-CCAUGUCCUGUAGAGC-ACUCACA-AGG-CUA--GGGGGC
C---UA-CC--AGCUUACCAAACCUUAUGAAACUCCGAAGGGGCAC--------------------------
------------------------------------------------------------------------
------------------GCGU-U-UA--------GUCCGGG-AGUGAGG-C-UGCG-A-GAGCUAACUUCC
GU-A-GCCGAGAG-GGAAAC---AACCCAGACCAUCA-GCUAAGGUCCCUAAAU--GA-UCG-CU-CA---G
-U-GGU-------UAAGGAUGUGUCGUCGCAUAGACAGCCAGGAGGUUGGCUUAGAAGCAGCCA-CCCUUCA
AAGAGUGCGUAAUAGCUCACUGGUC-----GAGUGACGAUGCGCCGAANAUG-AUC-GGGGCUCA-AGUGAU
```

```
C----------UACCGAA-GCUAUGGAUUCAACU--C-----------------------------------
------------------------------------------------------------------------
-----GC-----------------------------------------------------------------
----------------------------GAAGCGAGU-UGUC-UGGUAGGGGAGC-GUUCAGU---------
-----------C-C-GCGGA-GAAGCCA--U-ACCGGAAGGA--G-UGGUGGAGCC-G-ACU-GAAGUGCGG
AUGC-CGGCAU-GAGUA-ACGAUA---AAAG-AAGUGA--G---------------AAUCUU-CUU-CGCCG
U-AAGGAC-AAGGGUUCC-UGGGGAAGG-GUCGUCCGCCCAGGGAAAGUCGGG-AC-CUAAGGUGAGGCCG-
A-A-CGGC-----GCA--------------------------------------------------------
----------------------------------------GCC-GA-U--GGAC-AGCAGGUC-AAGA-UUC
CUGCA----------------------------------CCGAU----CAUG--------U-GGA-------
------------------------------------------------------------------------
----------------------------------------------------------GUGAU-GGAGGGAC
GCAUUA--------------CGCUAUCCAUGCCAA-GC-U-----AUGGC------------UAUGCUGGU-
UGGUAC-GCU-CAA----G-G-------------------G------CG------------------AU-CG
GGUCAG--------------------AAAAU-CUACCGGUCA------C-AUGCCUC----AGA-CG-U-A-
UCG--GG-----------------------------------------------------------------
-----AGCUUC-CUCGGAAGCGAAG-U-UGG-A-AACGCGACGGUGC-C-AAGAAA-AGCUU-CUAAACGU-
-------------------------------------------------------UGAAACA-UG-AUUGCC
CGUACCGCAAACCGAC------------------------------ACAGGUG--UCCGAGUGUCAAUGCAC
UAAGGC-G-CG-CGAG-AGAACCCU-C-GUUAAGGAACUUUGC-AAUCUCACCCCGUAACUUCGG-AAGAAG
GGGUC-CCCA-C-GC---------------------------------------------------------
------------------------------------------------------------------------
------------------------------------------------------------------------
------------------------------------------------------------------------
------------------------------------------------------------------------
------------------------------------------------------------------------
------------------------------------------------------------------------
------------------------------------------------------------------------
------------------------------------------------------------------------
------------------------------------------------------------------------
------------------------------------------------------------------------
------------------------------------------------------------------------
------------------------------------------------------------------------
------------------------------------------------------------------------
-------------------------------------------------------------------UUCGC
GUG--GGGCGCAGUGAAUAGGCCCAGGCGA-----CUGUUUACCAAAAUCACAGCACUCUGCCAACACG---
-AACA-------------GUGGACGU-AUAGGGUGUGA-CGCCUGCCCGGUGCCG-G-AAGG-UCAAGUGGA
GCGGU----------GCAA----------GCUGCGAAAUGAAGCCC--CGGUGAAC-GGCGGCCGUAACUAU
AACGGUCCUAAGGUAGCGAAAUUCCUUGUCGGG-UAAGUUCCGACCUGCACGAAAGGCGUAACGAUC--UGG
GCGCUGUCUCAACG----AGG-GACUCGGUGAAAUU--GAA---UUGGCUGUAAAGAUGCGGCCUACCCGUA
GCAGGACGAAAAGA-CCCCGUGGAGC-UUUACU--AUAGUCU-GGCAUUGGGAUUCGGGUUUCUCUGCGUAG
GAU-AGGUGGGAGCCUGCG----------AAACUGG-CCU----------------------------UUUG
--------------------------------------GG-GUCGGUGGAG-G--C-AA-CGGUGAAAUA-C
CACC-CUGAGAAACUUGGAUUUCUAACCUGAA--A------AA-U-C-------------------------
------------------------------------------------------------------------
```

```
------------------------------------------------------------------------
------------------------------------------------------------------------
------------------------------------------------------------------------
------------------------------------------------------AC----UU----------
------------------------------------------------UCGG-----GGACC-GUGCUUGGC
GGGUAGUUUGACUGGGGCGGU-CGCCUCCCA-AAAUGUAAC-GGAGGCGCCCA-AAGGUCACCUCAAGACGG
UUGGAAAUCGUCUGUA-GAGCG-CAAAGGUAG-AAGGUGGCUUGACU-GCGA--------------------
--------------GACUGA---CACGUCGAGCAGGGAGGAAA-CUCGGGCUUAGUGAACCGGU--------
---GG-U-----ACCGU-GUG--------------------GA-AGGGCCAUCGAUCAACGG-AUAAAAGUU
ACCCCGGGG-AUAACAGGCUGAUCUCCCCCGAGAGUCCAUAUCGGCGGGGAGGUUUGGCACCUCGAUGUCGG
CUCGUCGCAUCCUGGGGCUGAA-GAAGGUCCCAAGGGUUGGGCUGUUCGCCCAUUAAAGCGGCACGCGAGCU
GGGUUCAGAACGUCGUGAGACAGUUCGGUCUCUAUCCGCU--------------------A-CGG-------
-------------------------------GCGC--AGGAGAAUUGAGGGG-AGU-UGCUCCUAGUACGAG
AGGAC-CGGAGUGAACGGACCGC-UGGU-C-UCCCUGCUGUCGUACCAACGGCAC-AUGCAGGGUAGCUAUG
UCCGGAACGGAUAACCGCUGAAAGCA-UC--UAAGCGGGAAGCCAGCCCCAA-G-AUG--AGUUCUCC-CA-
--CUGUU-----------------------------------------------------------------
--------------------------U---------------------------------------------
------------------------------------------------------------------------
------------------------------------------------------------------------
------------------------------------------------------------------------
------------------------------------------------------------------------
------------------------------------------------------------------------
------------------------------------------------------------------------
--------------------------------------------------AUCAGG--UA-AGA------CU
CCCGGAAGACCACCGGGU-UAN-GAGGCC-AGG-CGUGCACG-CAU-A----------------GC------
----AA-----------UGUGUUCAGCGG--A-CUGG-UGCUCAUC---AGUCGAG-G-UCUUGACCA----
------------------------------------------------------------------------
------------------------------------------------------------------------
------------------------------------------------------------------------
------------------------------------C------------
>LSUb_243164_DEHM1/1-2840
---------------------G-GUCAAGCA----GG--UAAGGGCACAU-GGUGGAUG-C-CUUGGU-AUC
AAG---AGCCGAUGAAGGACGUGG--UAA-CUGCGAUAAGCCUCGGUCAGCUGUAA-CAAGCU--------U
AGC-CGG-GGAUUUCCGAAUGGGGCAAC--------CC--------GC----UCAG-G-C-----------A
A-A-------------A---C----C----U---GGG----U-A-CUUC-CAG-----CUCA----------
-----------A-----------CACA----------------------------------------UA-GG
C-U--------------------GG-GUAG-AGGUAAGUGGGGGAAGUGAAACAUCUCAGUACCCC--GAGG
AAAAGAAA---------UUAUUCUCCUA--GU-AG-CGGCGAGCGAACGGGGA-U-GA-GCCCAAA------
--------------------------------------------CAAGCCUGG----UUG------------
-----AAAG-AAA-------UCUG-UG----------------------GC--AGA---------GCAA---
A-------AGAAG-ACCUG------AAGGUC-CCUG------------------------------------
-------GAA-CG-GGCAA-------CCACAGAGGGUGAGAGUCCGUA----------AAUUAAA----GGG
U------G--UAGGC----------UUCU---GAUU--CU-UU------CUUGAGUACCACGAGACUCGUGG
AAUCUCGUGGGAA-UCCGCCGUG-ACC--ACAUGGCAAGGCUAAAUA--CUC--UUGA-UGACCGAUAGU-G
AAC-U-AGUACCGUGAGGGAAAG-GUGAAA-AGCAC-CCCGA-GAGGGG-A-GUGAAAUAGAA-CUGAAACC
```

```
G-UGUGCCUACAA-GCA-GU-CAGA---GGAC----------------------------------------
------------------------------------------------------------------------
------------------------------------------------------------------------
------------------------------------------------------------------------
------------------------------------------------------------------------
------------------------------------------------------------------------
------------------------------------------------------------------------
------------------------------------------------------------------------
------------------------------------------------------------------------
-------------------------------------------------------------GU--A-AG---
------------------------------------------------------------------------
------------------------------------------------------------------------
------------------------------------------------------------------------
------------------------------------------------------------------------
------------------------------------------------------------------------
------------------------------------------------------------------------
------------------------------------------------------------------------
------------------------------------------------------------------------
------------------------------------------------------------------------
------------------------------------------------------------------------
------------------------------------------------------------------------
------------------------------------------------------------------------
-------------------------------------------------C--U---CUGAUGGCGUGACUAU
UGAAGAAUUAUCCGACGAGUUAC-----UCAGUGCAG-GGUUAAGC--A-U---------UAAG--------
U-UG--CGCA--GCCGAAGCGAAAG-C---------------------------------------------
------------------------------------------------------------------------
------------------------------------------------------------------------
---------------------------------------------------------------------GAG
UCU------------------------------------GAAUA-------------------------GG-
G-C-GU---------------------UUUAGUUG--UACUGUA-GACCCGAAGCC-GGGUGAUCUAUCCAU
GGCCAGGGUGAAGUGAAG-GUAAUA-CUUCGCGGAGGCCCGA--ACCAGUCA-ACGUGGCAAAGUUG-UUGG
AUGAGCUGUGGAUAGCGGUUAUAUGCCAAACGAACCCGG-GGAUAGCUGGUUCUCCUCGAAA-UGCUUUAGG
----AGCCUCAG-GCG--------------------------------------------------------
----------------------UU-----CA-UUAAUGGAGGUAGUGC-ACUGGAU-AGA-CUA--GGGGGA
----GA-GA---UCCUGUCA-ACUCUAACAAACAUCGAAUGCCAUU--------------------------
------------------------------------------------------------------------
------------------AAUC-A-AA--------GCCUGGG-AGUGAGA-C-UAUG-G-GGGCUAAGCUCC
AU-A-GUCAAAAG-GGAAAC---AGCCCAGACCAUCA-GCUAAGGUCCCUAAGC--GC-AGG-CU-AA---G
-U-GGG-------AAAGGAAGUAGCAUUGCCUAGACAUCCAGUAGGUUGGCUUAGAAGCAGCCA-CCCUUUA
AAGAGUGCGUAAUAGCUCACUGGUU-----GAGUGGUGUUGCGCCGACAAUCCAAC-GGGGCUUA-A--CUG
U----------CACCGAA-GCUAUGGAUCUACCU--------------------------------------
------------------------------------------------------------------------
------------------------------------------------------------------------
---------------------------------AUGG-UAGA-UGGUAGAGGAGC-GUUCCCU---------
-----------C-G-GCAGU-GAAGCAG--G-AGUGUAAACA--A-CUGUGGAGCU-U-AGG-GAAGUGAGA
```

```
AUGU-CGGAAU-GAGUA-GCG-CA---AGGC-GUGUGA--G---------------AAACAC-GCC-CACCG
U-AAACCU-AAGGUUUCC-UACGCAAGG-UUAAUCCGCGUAGGGUUAGUCGG--AC-CUAAGCCGAGGCCG-
A-A-AGGC-----GUA--------------------------------------------------------
----------------------------------------GGC-GA-U--GGAC-AGCAGGUU-AUUA-UUC
CUGCA----------------------------------CCACC----GUU---------G-UG--------
------------------------------------------------------------------------
----------------------------------------------------------GAGGA-GGGGGGAC
ACAGGA--------------GGAUAG-AUCAGGCAU-GC-A----UUGGA------------CAUCGUGUC-
GCUGAA-CCC-UA-----G-A-------------------C------UU------------------UC-AA
A-UAGG--------------------CAAAU-CCGUUUGGAG------UUAAGUUGA----AGU-UG-U-G-
AAGGUGC---------------------------------------------------------------GG
-----GAGAUC----CAAGCCAAUC-U-GGU-U-GAGUCCAAGCUGU-C-GAGAAA-AGCCU-CUACUGU--
---------------------------------------------------------GAAGA-AU-GGUGAC
CGUACCGCAAACCGAC-------------------------------ACUGGUA--GGUGGGG-UAAAUUCC
CAAGGU-G-AU-CGAG-AUAACCCU-C-GUUAAGGAACUCGGC-AAUCUAGCCCCGUAACUUCGG-GAGAAG
GGGUG-CCCU-C-GC---------------------------------------------------------
------------------------------------------------------------------------
------------------------------------------------------------------------
------------------------------------------------------------------------
------------------------------------------------------------------------
------------------------------------------------------------------------
------------------------------------------------------------------------
------------------------------------------------------------------------
------------------------------------------------------------------------
------------------------------------------------------------------------
------------------------------------------------------------------------
------------------------------------------------------------------------
------------------------------------------------------------------------
------------------------------------------------------------------------
----------------------------------------------------------------------GA
GAG--GGUUACAAUAAAGUGGCUCGAGUGA-----CUGUUUAACAAAGACACAGGUCUCUGCAAAAGCG---
-AA-A-------------GCUGAAGU-AUAGGGGCUGA-CACCUGCCCAGUGCUG-G-AAGG-UUAAGGGGA
GGAGUUA-GG-----GCAA-----CCAAAGCUCUGAACCGAAGCCC--CAGUGAAC-GGCGGCCGUAACUAU
AACGGUCCUAAGGUAGCGAAAUCCCUUGUCGGG-UAAGUUCCGACCCGCACGAAAGGUAUAACAACU--UGA
GCGCUGUCUUAACG----AGG-GACUCGGUGAAAUU--GAA---UUACCCGUGAAGAUGCGGGUUUCCCGCG
ACUGGACAAAAAGA-CCCCGUGGAGC-UUUACU--AUAACUU-GCCAUUGAGUCUGGGUUCAUCAUGUGUAG
GAU-AGGUGGGAGGCUGAG----------AAGCCAG-GGC----------------------------GCUA
--------------------------------------GC-CUUGGUGGAG-C--C-AA-CGUUGAAAUA-C
CACU-CUUGAUGGAUCUAGCCUCUAACUC-UA--G----AACA-A-G-------------------------
------------------------------------------------------------------------
------------------------------------------------------------------------
------------------------------------------------------------------------
------------------------------------------------------------------------
-------------------------------------------------------------C----------
------------------------------------------------UAGA-----GGACA-GUGGCUGGU
```

```
GGAUAGUUUGACUGGGGCGGU-CGCCUCCUA-AAUUGUAAC-GGAGGCGCCCA-AAGGUUCCCUCAGGGUGG
AUGGAAAUCACCCUAA-GAGUG-CAUUGGCAU-AAGGGAGCUUGACU-GCGA--------------------
--------------GACAUA---CAAGUCGAGCAGGGACGAAA-GUCGGCAAAGUGAUCCUAC---------
---GG-C-----ACUGA-GUG--------------------GA-AGGGCCGUGGCUUAUCGG-AUAAAAGCU
ACCCCGGGG-AUAACAGGCUUAUCUUGCCCAAGAGUUCACAUCGACGGCAAGGUUUGGCACCUCUAUGUCGG
CUCGUCGCAUCCUGGGGCUGGA-GUAGGUCCCAAGGGUCCGGCUGUUCGCCGGUUAAAGCGGCACGCGAGCU
GGGUUCAGACCGUCGUGAGACAGGUCGGUCUCUAUCCGUC--------------------G-UGG-------
-------------------------------GCGU--UGGAGAUUUGAGAGG-GUC-UUUCCAUAGUACGAG
AGGAC-CUGGAAGGACAGACCAA-UGGU-G-UGUCAGUUGUUCUGCCAAGGGCAU-U-GCUGAGUAGCCAAG
UCUGGUUUGGAUAAGCGCUGAAAGCA-UC--UAAGUGCGAAGCCGGCCUCAA-G-AUU--AGAUCUCC-CA-
--UCCCC-----------------------------------------------------------------
------------------------------------------------------------------------
------------------------------------------------------------------------
------------------------------------------------------------------------
------------------------------------------------------------------------
------------------------------------------------------------------------
------------------------------------------------------------------------
------------------------------------------------------------------------
-------------------------------------------------AAGGGGG--UA-AGA------CC
GCAGGUAGACUACCUGCU-UGA-UAGGCU---G-UGUGUAAG-UAC-G----------------GU------
----AA-----------CGUAUUAAGCU------CAG-UACUAAU----GGUCGAG-G-GCUUGACC-----
------------------------------------------------------------------------
------------------------------------------------------------------------
------------------------------------------------------------------------
-------------------------------------------------
>LSUb_235279_HELHP/1-2880
---------------------A---AAAGCU----UU--UAAGGGCAGAU-GGUGGAUG-C-CUUGGG-UGA
UAG---AGGCGAUGAAGGACGUAC--UAGACUGCGAUAAGCUAUGGAGAGCUGUCAAGAAGCUU------UG
ACC-CAU-AGAUUUCCGAAUGGGGCAAC--------CCAA-----UAC----CUA---------------GA
G-A---------------------------U---AGG----U-A-UUAC-----------------------
-----------C-----------UUUA---------------------------------------------
------------------------A-U-GG-AGCGAACCUAGCGAAGUGAAACAUCUCAGUAGCUA--GAGG
AAAAGAAAUCAA--ACGAGAUUCUCCUA--GU-AG-CGGCGAGCGAACGGGGA-A-GA-GGGCAAACCAGU-
--AG-CU-------------------------------------UGCUACUGGGG--UUG------------
------AGG-ACU-------GCAA-UA----------------------UC--CAC---------UUGA---
A-------UAAUUGAGCAG------AAG-UA-UUUG------------------------------------
-------GAA-AA-AUACA-------UCACAGAGGGUGAUAAUCCCGUA---------UGCGAAG----GUU
A------U--UCAUA-----------GGU---AGCA--GU-AU------CCAGAGUAGGUCAGGGCACGUGA
AACCCGGACUGAA-GCAGGGGAG-ACC--ACUCUCCAACCCUAAAUA--CUA-CUAUC-ACACCGAUAGC-G
AAC-A-AGUACCGUGAGGGAAAG-GUGAAA-AGAAC-CGCAGUGAGCGG-A-GUGAAAUAGAACCUGAAACC
A-UCUGCCUACAA-UCA-UU-CAGA---GCCC-U---AU---------------------------------
------------------------------------------------------------------------
------------------------------------------------------------------------
------------------------------------------------------------------------
------------------------------------------------------------------------
```

```
------------------------------------------------------------------------
------------------------------------------------------------G-----------
------------------------------------------------------------------------
------------------------------------------------------------------------
--------------------------------------------------------------A--U-UU---
------------------------------------------------------------------------
------------------------------------------------------------------------
------------------------------------------------------------------------
------------------------------------------------------------------------
------------------------------------------------------------------------
------------------------------------------------------------------------
------------------------------------------------------------------------
------------------------------------------------------------------------
------------------------------------------------------------------------
------------------------------------------------------------------------
------------------------------------------------------------------------
------------------------------------------------------------------------
---------------------------A------------------UCAG--G---GUGAUGGACUGCCUUU
UGCAUAAUGAUCCUGCGAGUUGU-G-GUAUCUGGCAA-GGUUAAGC--------------AAAC--------
---G--CGAA--GCCGUAGCGAAAG-C---------------------------------------------
------------------------------------------------------------------------
------------------------------------------------------------------------
---------------------------------------------------------------------GAG
UCU------------------------------------GAAA--------------------------GG-
G-C-GU----------------------UUAGUCAGAUGCUGCA-GACCCGAAGCC-AAGUGAUCUAUCCAU
GGCCAAGUUGAAGCGAGU-GUAAUA-GCUCGUGGAGGACUGA--ACUCGUAC-CCAUUGAAACGGGU-UGGG
AUGAGCUGUGGAUAGGGGUGAAAGGCCAAACAAACUUGG-UGAUAGCUGGUUCUCUUCGAAA-UAUAUUUAG
GUAUAGCCUCAA-GUG--------------------------------------------------------
----------------------AU-----AG-UAAUAGGGGGUAGAGC-UCUGAUU-GGG-CUA--GGGCUG
C---UCACC--GCGGUACCAACCCCUGUCAAACUACGAAUACCUAU--------------------------
------------------------------------------------------------------------
------------------UACC-G-UA--------UCUUGGG-AGUCAGG-C-GGUG-G-GUGAUAAAAUCA
AU-C-GUCAAAAG-GGGAAC---AACCCAGACUACCA-ACUAAGGUCCCAAAGU--UC-UAUUCU-AA---G
-U-GGA-------AAAUGAUGUGAAGUUACUUAGACAACCAGGAGGUUGGCUUAGAAGCAGCCA-UCCUUUA
AAGAAAGCGUAACAGCUCACUGGUC-----UAGUGAUUUUGCGCAGAAAAUAUAAC-GGGGCU-A-AGAUAG
A----------CACCGAA-GUUGUAGAUUGUGCU--G-----------------------------------
------------------------------------------------------------------------
------------------------------------------------------------------------
---------------------------------AUGC-ACAG-UGGUAGAAGAGC-GUUCGUA---------
-----------UCA-GCGUU-GAAGGCA--UACCGGUAAGGA--G-UGCUGGAGCG-A-UGC-GAAGUGAGC
AUGC-AGGAAU-GAGUA-GCGAUA---AAAG-UGGUGA--G---------------AAUCCA-CUU-CGCCG
A-AAAUCU-AAGGUUUCC-UACGCGAUG-CUCGUCAUCGUAGGGUUAGUCGGG-UC-CUAAGACAAGUCCG-
A-A-AGGG-----GUA--------------------------------------------------------
----------------------------------------GUC-GA-U--GGAA-AAUGGGUU-AAUA-UUC
CCAUA----------------------------------CCAAC----AUUA--------G-UGU-------
```

```
------------------------------------------------------------------------
----------------------------------------------------------GCGAU-GGGGGGAC
GCAUAG--------------GGCUAAAUGGCGUUAG-CU-G----AUGGA------------AAUGCUAAU-
UUAAGA-GCG-UA-----G-A-------------------U------UG------------------GA-GG
A-AUGG--------------------AAAAU-CCGCCUCUGU----------AUUUGA----AAC-UC-C-AA
UAGCUUC---------------------------------------------------------------AA
GCGAGCUUCGG-CUCAAUUGGAGUG-C-CAU-U-GAUGCCGUCGUGC-C-AAGAAA-AGCCU-CUAAGUUU-
---------------------------------------------------------AGCUA-AU-GUUGCC
CGUACCGCAAACCGAC------------------------------ACAGGUA--GAUGAGAU-GAGUAUUC
UAAGGC-G-CG-UGAA-AGAACUCU-G-GUUAAGGAACUCUGC-AAACUAGCACCGUAAGUUCGC-AAUAAG
GUGUG-CCUA-C-GC---------------------------------------------------------
------------------------------------------------------------------------
------------------------------------------------------------------------
------------------------------------------------------------------------
------------------------------------------------------------------------
------------------------------------------------------------------------
------------------------------------------------------------------------
------------------------------------------------------------------------
------------------------------------------------------------------------
------------------------------------------------------------------------
------------------------------------------------------------------------
------------------------------------------------------------------------
------------------------------------------------------------------------
------------------------------------------------------------------------
------------------------------------------------------------------------
----------------------------------------------------------------------AA
GUA--GGUCUCAGCAAAGAGUCCCUCCCGA-----CUGUUUACCAAAAACACAGCACUUUGCAAACUCG---
-UA-A-------------GAGGAAGU-AUAAGGUGUGA-CGCCUGCCCGGUGCUC-G-AAGG-UUAAGAGGA
UUAGUUA-GCA----GCAA----UGCGAAGCUUUGAAUUGAAGCCC--GAGUAAAC-GGCGGCCGUAACUAU
AACGGUCCUAAGGUAGCGAAAUUCCUUGUCGGU-UAAAUACCGACCUGCAUGAAUGGCGUAACGAGA--UGG
GAGCUGUCUCAACC----AGG-GAUUCAGUGAAAUU--GUA---GUGGAGGUGAAAAUUCCUCCUACCCGCG
GCAAGACGGAAAGA-CCCCGUGGACC-UUUACU--ACAGCUU-GGCACUGCCGAUGGGAGCAUUAUGCGCAG
GAU-AGGUGGGAGGCUUUG----------AAGUCUU-CAC----------------------------UCUG
--------------------------------------GU-GGAGAUGGAG-C--C-AU-CGUUGAGAUA-C
CACC-CUUAAUGUUUCUGUCUGCUAACUGGCU--UAUG-U-UA-U-C-------------------------
------------------------------------------------------------------------
------------------------------------------------------------------------
------------------------------------------------------------------------
------------------------------------------------------------------------
------------------------------------------------------CA----UA----------
------------------------------------------------AGCA-----GGACA-AUGCCUGGU
GGGUAGUUUGACUGGGGCGGU-CGCCUCCUA-AAAAGUAAC-GGAGGCUUGCA-AAGGUUGGCUCAUUGCGG
UUGGAAAUCGCAAGAA-GAGUG-UAAUGGCAU-AAGCCAGCCUGACU-GUAA--------------------
--------------GACAAA---CAAGUCAAGCAGAGACGAAA-GUCGGUCAUAGUGAUCCGGU--------
---GA-U-----UCUGU-GUG--------------------GA-AGGGUCAUCGCUCAAAGG-AUAAAAGGU
ACCCCGGGG-AUAACAGGCUGAUCUCCCCCAAGAGCUCACAUCGACGGGGAGGUUUGGCACCUCGAUGUCGG
```

CUCAUCGCAUCCUGGGGCUGGA-GUAGGUCCCAAGGGUAUGGCUGUUCGCCAUUUAAAGCGGUACGCGAGCU
GGGUUCAGAACGUCGUGAGACAGUUCGGUCCCUAUCUGCC--------------------G-UGG-------
-------------------------------GCGU--AGGAAAGUUGAGGAG-AGC-UGUCCCUAGUACGAG
AGGAC-CGGGAUGGACACGCCAC-UGGU-G-UACCAGUUGUUCUGCCAAGAGCAU-C-GCUGGGUAGCUACG
CGUGGAUGUGAUAACCGCUGAAAGCA-UC--UAAGCGGGAAGCCAACUCCAA-G-AUA--AACUUUCC-CU-
------------------------------------------------------------------------
------------------------------------------------------------------------
------------------------------------------------------------------------
------------------------------------------------------------------------
------------------------------------------------------------------------
------------------------------------------------------------------------
------------------------------------------------------------------------
------------------------------------------------------------------------
----------------------------------------------------------GA-AGG------UC
GCAGCAAGACUAGCUGCU-UGA-UAGGGU-AGA-UGUGUAAG-CAU-A----------------GU------
----AA-----------UGUGUUUAGCUG--A-CUAC-UACUAAUA---GACCGCUUG-GCUUUUCUU----
------------------------------------------------------------------------
------------------------------------------------------------------------
------------------------------------------------------------------------
------------------------------------U------------
>LSUb_226186_BACTN/1-2858
----------------------------------------AAGGGCGCAU-GGCGGAUG-C-CUUGGC-UCU
CGG---AGGCGAUGAAGGACGUGA--UAAGCUGCGAUAAGCUCUGGGUAGGUGCAAAUAACCUU------UG
AUC-CAG-AGAUUUCCGAAUGGGACAAC--------CC--------GG----CAUU-C-U-----------G
A-A-------------G---G----A----A---UGU----C-A-UCCA-U---------------------
-----------C-----------UUUG---------------------------------------------
-----------------------AU-G-GA-AGCUAACGCAGGGAACUGAAACAUCUUAGUACCUG--UAGG
AAAAGAAAAUAA--UAAUGAUUCCCCUA--GU-AG-UGGCGAGCGAACGGGGA-A-UA-GCCCAAACCACCC
-AUG-UUACG--------------------------------G-CAUGUGUGGGG--UUG------------
-----UAGG-ACC-------ACGA-UG----------------------UC--GCA--------AGACA---
U-------UUGAUGAGUAG------AAU-CC-UCUG------------------------------------
-------GAA-AG-UUGAA--------CCAUAGACGGUGAUAGUCCGGUA---------UACGAAG----UCA
A------A--UUAAG-----------CGU---AGUG--GU-AU------CCUGAGUAGCGCGGGACACGAGA
AAUCUUGCGUGAA-UCUGCCGGG-ACC--AUCCGGUAAGGCUAAAUA--CUC-CCGAG-AGACCGAUAGC-G
AAC-C-AGUACUGUGAAGGAAAG-GUGAAA-AGCAC-UUCGAAUAGAAG-A-GUGAAAUAGUCCCUGAAACC
G-UGCGCCUACAA-GCG-GU-CGGA---GCUG-C--------------------------------------
------------------------------------------------------------------------
------------------------------------------------------------------------
------------------------------------------------------------------------
------------------------------------------------------------------------
------------------------------------------------------------------------
------------------------------------------------------------------------
------------------------------------------------------------------------
------------------------------------------------------------------------
--------------------------------------------------------------U--U-AA---

```
------------------------------------------------------------------------
------------------------------------------------------------------------
------------------------------------------------------------------------
------------------------------------------------------------------------
------------------------------------------------------------------------
------------------------------------------------------------------------
------------------------------------------------------------------------
------------------------------------------------------------------------
------------------------------------------------------------------------
------------------------------------------------------------------------
------------------------------------------------------------------------
------------------------------------------------------------------------
------------------------------------------------GC--A---GUGACGGCGUGCCUUU
UGCAUAAUGAACCUACGAGUUAC-U-UUUUCCGGCAA-GGUUAAGC--A-UC--------UUGA-------G
A-UG--UGCA--GCCGAAGCGAAAG-C---------------------------------------------
------------------------------------------------------------------------
------------------------------------------------------------------------
---------------------------------------------------------------------GAG
UCU------------------------------------GAACA-------------------------GG-
G-C-GU----------------------CGAGUCGGAAGGAGUA-GACGCGAAACC-AAGUGAUCUACCCUU
GGUCAGGUUGAAGGUUAG-GUAACA-CUAACUGGAGGACCGA--ACCGAUAA-GCGUUGAAAAGCUU-CCGG
AUGAACUGAGGGUGGGGGUGAAAGGCUAAUCAAACUUGG-AGAUAGCUCGUACUCCCCGAAA-UGCAUUUAG
GUGCAGCCUUGU-GAG--------------------------------------------------------
----------------------UU-----AC-UAAUGUGAGGUAGAGCGACUGAUA-AGA-UGC--GAGGGC
U---UC-AC--CGCCUAUCAAGUCUUGAUAAACUCCGAAUGCGCAU--------------------------
------------------------------------------------------------------------
------------------UAGUUC-UA--------UCACAGG-AGUGAGG-G-CAUG-G-GUGCUAAGGUCC
AU-G-UCCUAAAG-GAGAAG---AAUCCAGACCAUCA-GCUAAGGUCCCCAAAU--AA-ACA-CU-AA---G
-U-UGAA-----CUAACGAAGUCAGAUUGCUAAGACAGCUAGGAUGUUGGCUUGGAAGCAGCCAUUCAUUUA
AAGAGUGCGUAACAGCUCACUAGUC-----GAGGAGUUUGGCGUGGAUAAUA-AUC-GGGCAUUA-AGUGUU
U----------UACCGAA-GCUAUGGGAUCA-----------------------------------------
------------------------------------------------------------------------
---GUAA-----------------------------------------------------------------
--------------------------------------UGAU-CGGUAGGGGAGC-AUUCCAC---------
-----------UCU-GCGUC-GAAGGUG--A-AGCGUGAGCU--U-UGCUGGAGCG-U-GUG-GAAAAGCAA
AUGU-AGGUAU-AAGUA-ACGAUA---AAGG-GGGUGA--G---------------AAACCC-CCU-CGCCG
A-AAGACU-AAGGUUUCC-UGAUCAACG-CUAAUCGGAUCAGGGUUAGUCGGG-UC-CUAAGGCCCAGCCG-
A-A-CGGU-----GAG--------------------------------------------------------
----------------------------------------GCC-GA-U--GGCAGAACAGGUU-AAUA-UUC
CUGUA----------------------------------CUACC----UUAA--------G-GA--------
------------------------------------------------------------------------
----------------------------------------------------------GUGAC-GUGGAGAC
GGAGCAG-------------UGACAG-CGCCGCGGA-CU-G----ACGGA------------AUAGUCCGU-
UGAAGG-GUG-UA-----G-A-------------------UG--UUGAU------------------CA-UC
C-CAGG--------------------CAAAU-CCGGGAUGAG---------AGUCGA----ACC-UG-A-U-
```

```
AGUAUGG---------------------------------------------------------------AG
C-GUUCUUCGG-AACAAUCCAAUAG-U-GCG-U-GUAAGCAUACUCC-C-AAGAAA-AUCCG-CUAAACUU-
-------------------------------------------------------AAUCCUU-GA-GGUACC
CGUACCGCAAACGGAC------------------------------ACACGUA--GUCGGGUU-GAAUAUAC
UAAGGC-G-CU-UGAG-UGAUUCAC-G-GUUAAGGAACUAGGC-AAAUUGACCCUGUAACUUCGG-GAUAAA
GGGUC-CCAA-C-GA---------------------------------------------------------
------------------------------------------------------------------------
------------------------------------------------------------------------
------------------------------------------------------------------------
------------------------------------------------------------------------
------------------------------------------------------------------------
------------------------------------------------------------------------
------------------------------------------------------------------------
------------------------------------------------------------------------
------------------------------------------------------------------------
------------------------------------------------------------------------
------------------------------------------------------------------------
------------------------------------------------------------------------
------------------------------------------------------------------------
----------------------------------------------------------------------GA
GUU--GGGCGCAGAGAAUAGGUCCAGGCAA-----CUGUUUAACAAAAACACAGGGCUGUGCCAAAUUG---
-AA-A-------------GAUUACGU-AUACAGCCUGA-CACCUGCCCGGUGCUG-G-AAGG-UUAAGAGGA
GAUGUCA-UC-----GCAA-----GAGAAGCAUUGAAUUGAAGCCC--CAGUAAAC-GGCGGCCGUAACUAU
AACGGUCCUAAGGUAGCGAAAUUCCUUGUCGGG-UAAGUUCCGACCUGCACGAAUGGUGUAAUGAUC--UGG
ACACUGUCUCAACC----GUG-AGCUCAGUGAAAUU--GUA---GUAUCGGUGAAGAUGCCGAUUACCCGCG
AUGGGACGAAAAGA-CCCCGUGAACC-UUUACU--AUAGCUU-AACAUUGAAUUUGGGUAAUUGAUGUGUAG
GAU-AGGCCGGAGACUUUG----------AAGCAGG-UAC----------------------------GCCA
---------------------------------------GU-AUUUGUGGAG-U--C-GC-UGUUGAAUA-C
GGCC-CUUUGAUUAUUUGAGUUCUAACUC-GU--------AG-U-U--------------------------
------------------------------------------------------------------------
------------------------------------------------------------------------
------------------------------------------------------------------------
------------------------------------------------------------------------
------------------------------------------------------------------------
------------------------------------------------GCGA-----GGACA-CUGUUUGGU
GGGUAGUUUGACUGGGGUGGU-CGCCUCCAA-AAUAGUAAC-GGAGGCUUCUA-AAGGUGCCCUCAGGACGA
UUGGUAACCGUCCGUA-GAGUG-UAAUGGCAU-AAGGGCGCUUGACU-GGGA--------------------
--------------GACUAA---CAAGUCGAUCAGGUAGGAAA-CUAGAGCAUAGUGAUCCGGU--------
---GU-U-----UCCGU-AUG--------------------GA-AGGGACAUCGCUCAAAGG-AUAAAAGGU
ACUCCGGGG-AUAACAGGCUGAUCCCUCCCAAGAGCUCAUAUCGACGGAGGGGUUUGGCACCUCGAUGUCGG
CUCGUCACAUCCUGGGGCUGGA-GAAGGUCCCAAGGGUUGGGCUGUUCGCCCAUUAAAGUGGCACGCGAGCU
GGGUUCAGAACGUCGUGAGACAGUUCGGUCUCUAUCUAUC--------------------G-UGG-------
-------------------------------GCGU--AUGAAAUUUGCGUGG-CUC-UGACACUAGUACGAG
AGGAC-CGUGUUGGACUGACCGC-UGGU-U-UACCAGUUGUGCCGCCAGGUGCAU-U-GCUGGGUAUCUAAG
UCGGGAAUGGAUAAGUGCUGAAAGCAUUC--UAAGUACGAAGCCA----CAA-G-AUU--AGAUUCUU----
```

```
------------------------------------------------------------------------
------------------------------------------------------------------------
------------------------------------------------------------------------
------------------------------------------------------------------------
------------------------------------------------------------------------
------------------------------------------------------------------------
------------------------------------------------------------------------
------------------------------------------------------------------------
--------------------------------------------------------------GG------GU
CGUUAAAGACGAUGACGU-UGA-UAGGAU-GCA-GGUGUAAA-GGU-G----------------GU------
----AA-----------CAU-CAAAGCCG--A-GCAU-UACUAAUU---GCCCG-------UUCACUU----
------------------------------------------------------------------------
------------------------------------------------------------------------
------------------------------------------------------------------------
------------------------------------U------------
>LSUb_242619_PORGI/1-2884
----------------------------------------AAGGGCAGAC-GGUGGAUG-C-CUAGGC-UCU
CGG---AGGCGAAGAAGGACGUGA--UAAGCUGCGAAAAGCUGCGGGAAUCGGCACAUACGAAU------UG
AUC-CGC-AGAUAUCCGAAUGGGGCAAC--------CC--------GU----CAGG-C------------CA
A-G-----------------G----C----C---UGA----C-A-CAUG-AA--------------------
-----------U-----------UGAU---------------------------------------------
--U--------------------UC-A-UG-AGCGAACGCGGGGAACUGAAACAUCUCAUUACCCG--UAGG
AGAAGAAAACAA--AAGUGAUUCCCUCA--GU-AG-UGGCGAGCGAACGGGGA-U-UA-GCCCAAACCGGCU
-UUG-UUUCG--------------------------------G-CAAAGUCGGGG--UAA------------
-----UAGG-ACU-------UCGA-CAA-------------------AAGU--UUA---------AUGA--U
A-------GAUAU-AGGAG------AAC-CU-ACUG------------------------------------
-------GAA-AG-UAUGG-------CCGAAGAGCAUGAAAGCUGCGUA---------UCCGACA----UAU
C------G--AACAA----------UAGAC-GACGG--AG-CA------CCUGAGUAGCGCGGGACACGAGA
CAUCCUGUGUGAA-UUGGCGGGG-CCC--AUCCCGUAAGGCUAAAUA--CUC-CCGAG-AGACCGAUAGU-G
AAC-C-AGUACCGUGAGGGAAAG-GUGAAA-AGAAC-CUCGAACAGAGG-A-GUGCAAUAGACCCUGAACCC
G-UCUGCCUACAA-GCG-GU-AGGA---GCGC-C--------------------------------------
------------------------------------------------------------------------
------------------------------------------------------------------------
------------------------------------------------------------------------
------------------------------------------------------------------------
------------------------------------------------------------------------
------------------------------------------------------------------------
------------------------------------------------------------------------
------------------------------------------------------------------------
-------------------------------------------------------------AU--U-AA---
------------------------------------------------------------------------
------------------------------------------------------------------------
------------------------------------------------------------------------
------------------------------------------------------------------------
------------------------------------------------------------------------
```

```
------------------------------------------------------------------------
------------------------------------------------------------------------
------------------------------------------------------------------------
------------------------------------------------------------------------
------------------------------------------------------------------------
------------------------------------------------------------------------
------------------------------------------------------------------------
------------------------------------------------GG--U---GUGACUGCGUGCCUUU
UGCAUAAUGAACCUACGAGUUAC-U-GUUUGUGGCAA-GGUUAAUU--G-UUAU------AAUC-----AAG
A-CA--AGGA--GCCGAAGCGAAAG-C---------------------------------------------
------------------------------------------------------------------------
------------------------------------------------------------------------
---------------------------------------------------------------------GAG
UCU------------------------------------UAAAA-------------------------GG-
G-C-GC------------------CCAUUUAGUCACGAGCAGUA-GACGCGAAACC-AAGUGAUCUACCCUU
GGUCAGGUUGAAGGUUAG-GUAACA-CUAACUGGAGGACCGA--AUCGGUAA-GCGUUGAAAAGCUU-UCGA
AUGAACUGAGGGUAGGGGUGAAAGGCUAAUCAAACUUGG-AGAUAGCUCGUACUCCCCGAAA-UGCAUUUAG
GUGCAGCCUGUU-GGA--------------------------------------------------------
-----------------------U-----GU-UAUCAUGAGGUAGAGCGACUGAUU-GGA-UGC--GAGGGU
U---UC-AC--CGCCUAUCAAGUCCAGAUAAACUCCGAAUGCAUGA--------------------------
------------------------------------------------------------------------
------------------UAAU-U-GA--------CCGAUGG-AGUGAGG-G-CAUG-G-GUGCUAAGGUCC
AU-G-UCCGAGAG-GAGAAG---AAUCCGGACCACCG-GCUAAGGUCCCGAAAU--AA-UUG-CU-AA---G
-U-UGUA-----AAAACGAAGUCAAGAUGCAGAGACAGCUAGGAUGUUGGCUUGGAAGCAGCCAUUCAUUUA
AAGAGUGCGUAACAGCUCACUAGUC-----GAGGAUUUUGGCAUGGAUAAUA-CAC-GGGCAU-A-AGCAAU
U----------UACCGAA-GCCGUGGGAUAUAGU--------------------------------------
------------------------------------------------------------------------
------------------------------------------------------------------------
----------------------------------AAU-AUAU-CGGUAGGGGAGC-AUUCCAG---------
-----------C-G-ACGUAGAAGGGAA--A-AGGGCGACUU--U-UUCUGGAGUU-U-CUG-GAAAAGCAA
AUGU-AGGUAU-AAGUA-ACGAUA---AAGG-GGGCGA--G---------------AACCCC-CCU-CGCCG
A-AAGACC-AAGGUUUCC-UGAUCAACG-CUAAUCGGAUCAGGGUUAGUCGGG-GC-CUAAGGAUAAGCCG-
A-A-UGGC-----GAU--------------------------------------------------------
----------------------------------------UCC-GA-U--GGAAGAACCGGUU-AAUA-UUC
CGGUA----------------------------------CUGAU----ACAG--------A-GA--------
------------------------------------------------------------------------
----------------------------------------------------------GCGAU-GUGGUGAC
GGAGAAG-------------UGACAG-UCCGGCCGU-CU-G----ACGGA------------AUAGGCGGU-
UAAAGG-GUG-UA-----G-A-------------------UG--UUGAU------------------CG-GG
G-UAGG--------------------CAAAU-CCGCCCUGAG---------AGUCGA----ACC-UG-A-C-
AGUACCC---------------------------------------------------------------GG
----AGUACAA-GUACGAAGGGAUA-A-GGA-C-GUAAACCGGCUCC-C-AAGAAA-ACCCG-CUAAGCAU-
--------------------------------------------------------AUUUCU-GU-GUUACC
CGUACCGUAAACCGAC------------------------------ACAGGUG--GUUGGGUU-GAGUAUAC
UAAGGC-G-CU-CGAG-AGAUUCGC-G-GUUAAGGAACUAGGC-AAAAUGGUCCUGUAACUUCGG-GAGAAA
```

```
GGACG-CCUG-U-CU---------------------------------------------------------
------------------------------------------------------------------------
------------------------------------------------------------------------
------------------------------------------------------------------------
------------------------------------------------------------------------
------------------------------------------------------------------------
------------------------------------------------------------------------
------------------------------------------------------------------------
------------------------------------------------------------------------
------------------------------------------------------------------------
------------------------------------------------------------------------
------------------------------------------------------------------------
------------------------------------------------------------------------
------------------------------------------------------------------------
--------------------------------------------------------------------CCGG
ACA--GGCCGCAGAAACCAGGCCCAGGCGA-----CUGUUUAACAAAAACACAAGGCUAUGCAAAAAAG---
-CA-A-------------UUUGAGGU-AUAUAGUCUGA-CACCUGCCCGGUGCUG-G-AAGG-UUAAGAGGA
GGAGUCA-UCG----U-CA----AGAGAAGCUCUGAAUUGAAGCCC--CAGUAAAC-GGCGGCCGUAACUAU
AACGGUCCUAAGGUAGCGAAAUUCCUUGUCGGG-UAAGUUCCGACCUGCACGAAUGGUGUAACGAUC--UGG
GCACUGUCUCAACC----GCG-AUCUCGGUGAAAUU--GUA---GUAUCGGUGAAGAUGCCGAUUACCCGCA
ACGGGACGAAAAGA-CCCCGUGAACC-UUUACU--AUAGCUU-UACAUUGUAUUUGGGCAUCAGAUGUGUAG
GAU-AGGCCGGAGGCAGAG----------AAGCGGG-UAC----------------------------GCCA
--------------------------------------GU-AUUCGUGGAG-U--C-GA-UGUUGAAAUA-C
GGCC-CUUUUGAUGUUUGGAUACUAACUC-GC-------G-GC-G-U-------------------------
------------------------------------------------------------------------
------------------------------------------------------------------------
------------------------------------------------------------------------
------------------------------------------------------------------------
------------------------------------------------------------------------
------------------------------------------------GCGA-----GGACA-GUGUAUGGU
GGGUAGUUUGACUGGGGUGGU-CGCCUCCAA-AAGCGUAAC-GGAGGCUUCUA-AAGGUACCCUCAGGCCGA
UUGGUAACCGGUCGCA-GAGUG-UAAUGGCAC-AAGGGUGCUUGACU-GGGA--------------------
--------------GACAAA---CAAGUCGCACAGGUAGGAAA-CUAGAGCAUAGUGAUCCGGU--------
---GG-U-----UCCGC-AUG--------------------GA-AGGGCCAUCGCUCAAAGG-AUAAAAGGU
ACUCCGGGG-AUAACAGGCUGAUCACUCCCAAGAGCUCAUAUCGACGGAGUGGUUUGGCACCUCGAUGUCGG
CUCGUCACAUCCUGGGGCUGGA-GAAGGUCCCAAGGGUUGGGCUGUUCGCCCAUUAAAGUGGCACGCGAGCU
GGGUUCAGAACGUCGUGAGACAGUUCGGUCUCUAUCUGUU--------------------G-UGG-------
-------------------------------GCGC--AGGAAAUUUGCGAGG-GUC-UGACACUAGUACGAG
AGGAC-CGUGUUGGACAGACCCC-UGGU-U-UACCGGUUGUACCGCCAGGUGCAC-C-GCCGGGUAUCCACG
UCUGGUAAGGAUAAGUGCUGAAAGCA-UC--UAAGCACGAAGCCGGCCUCAA-G-AUA--AGAUUUCC-AU-
--AA--------------------------------------------------------------------
------------------------------------------------------------------------
------------------------------------------------------------------------
------------------------------------------------------------------------
------------------------------------------------------------------------
```

```
------------------------------------------------------------------------
------------------------------------------------------------------------
------------------------------------------------------------------------
----------------------------------------------------------AU-AGG------GU
GGUUAAAGACUAUGACCU-UGA-UAGGCU-GCA-GGUGUAUG-AUU-G----------------GU------
----AA-----------CA-AUUAAGCCG--A-GCAG-UACUAAUA---GCCCGAA---ACUUUUGU-----
------------------------------------------------------------------------
------------------------------------------------------------------------
------------------------------------------------------------------------
-------------------------------------------------
>LSUb_269798_CYTH3/1-2883
---------------------A--GAAAAUA----AA--UAAGGGCGUAU-GGGGAAUG-C-CUAGGC-UCU
CAG---AGGCGAAGAAAGACGUGC--CAAGCUGCGAAAAGUUACGGGAUCUGCUGAGGAGAUU------UG
AUC-CGU-AAAUGUCUGAAUGGGGCAAC--------CU--------GG----UAUG-U-U-----------G
A-A-------------G---A----C----A---UAU----C-G-ACCA-----------------------
-----------G-----------CAA----------------------------------------------
------------------------U-G-GU-AGCGAACCCGGUGAACUGAAACAUCUAAGUAACUG--GAGG
AAGAGAAAAUAAC-UAAUGAUUCCCUAA--GU-AG-UGGCGAGCGAACAGGGA-A-GA-GCCCAAACCAAUG
-UUG-UUACG--------------------------------G-CAAUAUUGGGG--UUG------------
-----UAGG-ACC-------GCAU-U-----------------------UA--AGA---------GAAG---
U-------AAUUGAACUGG------AAA-UG-UCUG-------------------------------------
-------GGA-AG-GCAUG-------CCAUAGCGGGUGAGAGCCCUUA----------CAGGUAA----GAU
U------A--UCAUU-------------U--GAGCG--GU-AU------CCUGAGUAGGGCGGGACCGGAGA
AAUCCCGUUUGAA-ACUGCCGGC-ACC--AUCCGGUAAGGCUAAAUA--CUC-CUGAG-AGACCGAUAGU-G
AAC-C-AGUACCGUGAGGGAAAG-GUGAAA-AGUAC-CCUGAAUAAGGG-G-GUGAAAUAGAACCUGAAACC
A-UACGCUUACAA-GCG-GU-CGGA---GCCC-U--------------------------------------
------------------------------------------------------------------------
------------------------------------------------------------------------
------------------------------------------------------------------------
------------------------------------------------------------------------
------------------------------------------------------------------------
------------------------------------------------------------------------
------------------------------------------------------------------------
------------------------------------------------------------------------
--------------------------------------------------------------U--A-AU---
------------------------------------------------------------------------
------------------------------------------------------------------------
------------------------------------------------------------------------
------------------------------------------------------------------------
------------------------------------------------------------------------
------------------------------------------------------------------------
------------------------------------------------------------------------
------------------------------------------------------------------------
------------------------------------------------------------------------
------------------------------------------------------------------------
```

```
------------------------------------------------------------------------
------------------------------------------------------------------------
------------------------------------------------GG--G---GUGACGGCGUGCCUUU
UGCAUAAUGAUCCUACGAGUUACUC-CUCACUGGCAA-GGUUAAGC--A-CU--------UUUA-------G
G-UG--CGUA--GCCGGAGCGAAAG-C---------------------------------------------
------------------------------------------------------------------------
------------------------------------------------------------------------
---------------------------------------------------------------------GAG
UCU------------------------------------GAAUA-------------------------GG-
G-C-GC----------------------GUAGUCAGUGGGGGUA-GACGCGAAACC-UUGUGAUCUACCCAU
GGCCAGGAUGAAGUUGCA-GUAACA-UGCAAUGGAGGUCCGA--ACUGGUAA-ACGUUGAAAAGUUU-UCGG
AUGAGCUGUGGGUAGGGGUGAAAGGCUAAUCAAACUGGG-AAAUAGCUCGUACUCCCCGAAA-UGUAUUUUG
GUACAGCGUCGG-GGUA-------------------------------------------------------
----------------------AA-----GU-UUAGCGGAGGUAGAGCUACCAAUA-GGA-CUA--GGGGGA
G---UC-AA--AUCCUACCAAAUCCUGAUGAACUCCGAAUGCCGCU--------------------------
------------------------------------------------------------------------
------------------AAAU-A--U--------ACCCGGC-AGUGAGG-G-CUUG-G-GUGCUAAGGUCC
AA-G-UCCGAGAG-GGAAAG---AACCCAGACUAACA-GCUAAGGUCCCAAAU--CU-AUC-CU-AA---G
-U-UGAA-----CAAAGGAGGUCCAGUUGCAGAGACAGCCAGGAUGUUUGCUCGGAAGCAGCAAUUCAUUUA
AAGAGUGCGUAACAGCUCACUGGUC-----GAGCGACAGGGCAUCGAUAAUA-AAC-GGGCAUCA-AGGGUA
G----------UACCGAA-GCUUUAGAUUUAUAC--U-----------------------------------
------------------------------------------------------------------------
-----U------------------------------------------------------------------
-------------------------------UAGGAG-UAAG-UGGUAGGGGAGC-AUUCCAA---------
-----------G-G-CCGGU-GAAGGCG--A-AACGCGAGUU--U-UGCUGGAGGG-C-UUG-GAAAAGCAA
AUGU-AGGAAU-AAGUA-ACGAUA---AUAA-GGGUGA--G---------------AAUCCC-UUA-CACCG
A-UAGACU-AAGGUUUCC-UGAUCAACG-UUCGUCGGAUCAGGGUUAGUCGGG-AC-CUAAGGCAAAUCCG-
----UGGC-----GUA--------------------------------------------------------
----------------------------------------GUC-GA-U--GGAC-AACUGGUU-AAUA-UUC
CAGUA----------------------------------CUUGU----GUAU--------A-CU--------
------------------------------------------------------------------------
----------------------------------------------------------GCUAU-GGAGAGAC
GGAGUAG-------------UGAAAA-GUUCGCCUG-GU-G----ACGGA------------AUACCAGGU-
UAAAAC-CCG-UA-----G-A-------------------UA--U-AUU------------------UU-GA
G-UAGG--------------------CAAAU-CCGCUCGAGA---------UGUCGA----AGG-UG-A-U-
AGU--AC---------------------------------------------------------------AA
CAA-AGCUUCG-GCAGCGUUGAUAG-U-GAC-U-GUAAACAGACUUC-C-AAGAAA-AUCUU-CUAUGCUU-
-------------------------------------------------------CAGGUAU-AU-ACGACC
CGUACCGCAAACCGAC------------------------------ACAGGUA--GUCAAGGA-GAGAAUCC
UGAGGU-G-CG-CGAG-UGAUUCAC-G-GCUAAGGAACUCGGC-AAAUUAACCCUGUAACUUCGG-GAGAAG
GGGUG-CUUCCU-CU---------------------------------------------------------
------------------------------------------------------------------------
------------------------------------------------------------------------
------------------------------------------------------------------------
------------------------------------------------------------------------
```

```
------------------------------------------------------------------------
------------------------------------------------------------------------
------------------------------------------------------------------------
------------------------------------------------------------------------
------------------------------------------------------------------------
------------------------------------------------------------------------
------------------------------------------------------------------------
------------------------------------------------------------------------
------------------------------------------------------------------------
---------------------------------------------------------------A--GCGAUG
AGA--AGCCGCAGUGAAAAGUCUCAGGCGA-----CUGUUUAACAAAAACACAUGGCUCUGCGAAAUCG---
-AA-A-------------GAUAAAGU-AUAGGGCCUGA-CACCUGCCCGGUGCCG-G-AAGG-UUAAGAGGG
GGGCUUA-GC-----GUAA-----GCGACGGUCUGAAUCGAAGCCC--CGGUAAAC-GGCGGCCGUAACUAU
AACGGUCCUAAGGUAGCGAAAUUCCUUGUCGGG-UAAGUUCCGACCUGCACGAAUGGUGUAACGAUC--UGA
GAACUGUCUCAGCC----GUG-AGCUCGGUGAAAUU--GUA---GUAGCGGUGAAGAUGCCGCUUACCCGUC
ACGGGACGGAAAGA-CCCCGUGCACC-UUUACU--AUAGCUU-UGCAUUGAUUUGGGUAAAUGAUGUGUAG
GAU-AGGUGGGAGGCUUUG----------AAGCGGG-UAC----------------------------GCUA
--------------------------------------GU-AUUCGUGGAG-C--C-AA-CGUUGAAAUA-C
CACC-CUUUAUUUACUUGGAGUCUAACCC-CU---------GA-A-U-------------------------
------------------------------------------------------------------------
------------------------------------------------------------------------
------------------------------------------------------------------------
------------------------------------------------------------------------
------------------------------------------------------------------------
------------------------------------------------AGGG-----AAACA-GUGCAUGGU
GGGUAGUUUGACUGGGGUGGU-CGCCUCCAA-AAAAGUAUC-GGAGGCUUCUA-AAGGUUCCCUCAGUACGC
UUGGUAACCGUACGUA-GAGUG-CAAUGGCAU-AAGGGAGCUUGACU-GUGA--------------------
--------------GACAAA---CAAGUCGAUCAGGUACGAAA-GUAGAGCAUAGUGAUCCGGU--------
---GG-U-----UCCGC-AUG--------------------GA-AGGGCCAUCGCUCAAAGG-AUAAAAGGU
ACGCCGGGG-AUAACAGGCUGAUCUCCCCCAAGAGCUCACAUCGACGGGGAGGUUUGGCACCUCGAUGUCGG
CUCGUCACAUCCUGGGGCUGGA-GAAGGUCCCAAGGGUUCGGCUGUUCGCCGAUUAAAGUGGCACGCGAGCU
GGGUUCAGAACGUCGUGAGACAGUUCGGUCCCUAUCUGUG--------------------A-UGG-------
-------------------------------GCGU--AGGAAGUUUGAGUGG-ACC-UGACCUUAGUACGAG
AGGAC-CGGGUUGGACUAACCUC-UGGU-G-UAUCUGUUGUGCCGCCAGGUGCAG-U-GCAGAGUAGCCACG
UUGGGACGAGAUAAGCGCUGAAAGCA-UC--UAAGCGCGAAACUCACCACAA-G-AUG--AGACUUCC-AU-
------------------------------------------------------------------------
------------------------------------------------------------------------
------------------------------------------------------------------------
------------------------------------------------------------------------
------------------------------------------------------------------------
------------------------------------------------------------------------
------------------------------------------------------------------------
------------------------------------------------------------------------
----------------------------------------------------------AC-AGG------GA
CGUCAGAGACUAUGACGU-UGA-UAGGCU-GCA-AGUGUAAA-GUC-A----------------GA------
```

```
----AA-----------UG-GCAUAGCUG--A-GCAG-UACUAAUU---ACCCGAU-A-AUUUUCUUU----
------------------------------------------------------------------------
------------------------------------------------------------------------
------------------------------------------------------------------------
------------------------------------G------------
>LSUb_194439_CHLTE/1-2915
------------------------UUAAGUU----AC--UAAGGGCGUAC-GGUGGAUG-C-CUUGGC-ACA
AGA---AGGCGAUGAAGGACGUGG-UUUAACUGCGAAAAGCUGCGGGGAGGCGUAAACAGCCUG------AU
AUC-CGC-AGAUCUCCGAAUGGGGCAAC--------CCA-------CC----UC----------------GA
U-U-------------------------------GAG----G-AAUCCC-----------------------
-----------G-----------UAAU---------------------------------------------
------------------------G-G-GA-GGCUAACCCAGGGAAGUGAAACAUCUCAGUACCUG--GAGG
AAAAGAAAACAA--AAGUGAUUUCCUGA--GU-AG-UGGCGAGCGAAAAGGAA-C-AA-GCCCAAACCGGUG
CCUG-UUUCG--------------------------------G-CAGCGCCGGGG--UUG------------
-----UAGG-ACU-------CCAG-CA----------------------CG--UGA---------CAGC--G
A-------GCGAU-AGCAG------AAC-UA-CCUG------------------------------------
-------GAA-AG-GUAGG-------CCAAAGAAGGUGAAAGCCCUGUA---------AGCGAAA----UCG
C------A--AACUG-----------CAC---AGGA--GG-AU------CCUGAGUAGCACGGGGUACGUGA
GGUCCUGUGUGAA-CCAGCGAGA-ACC--AUCUCGUAAGGCUAAAUA--CUC-UCUUG-UGACCGAUAGU-G
AAC-A-AGUACCGUAAGGGAAAG-GUGAAA-AGUAC-CCUUAAGAAGGG-A-GUGAAAUAGUACCUGAAACC
G-UACGCUUACAAAACU-GU-AGGA---GCUA-A---CC---------------------------------
------------------------------------------------------------------------
------------------------------------------------------------------------
------------------------------------------------------------------------
------------------------------------------------------------------------
------------------------------------------------------------------------
------------------------------------------------------------CA-C--------
------------------------------------------------------------------------
------------------------------------------------------------------------
--------------------------------------------------------------G--C-AA---
------------------------------------------------------------------------
------------------------------------------------------------------------
------------------------------------------------------------------------
------------------------------------------------------------------------
------------------------------------------------------------------------
------------------------------------------------------------------------
------------------------------------------------------------------------
------------------------------------------------------------------------
------------------------------------------------------------------------
------------------------------------------------------------------------
------------------------------------------------------------------------
------------------------------------------------------------------------
-------------------------GUG-------------------GGG--U---GUGACUGCGUGCCUUU
UGCAUAAUGAGCCUACGAGUUACUG-GUGUGUUGCGA-GGUUAAGG--U-U--------UUCAG--------
A-AC--CGGA--GCCGCAGCGAAAG-C--------------------------------------------
```

```
------------------------------------------------------------------------
------------------------------------------------------------------------
---------------------------------------------------------------------GAG
UUC------------------------------------GAACA-------------------------GA-
G-C-GG-----------------------GAGUAACAUGCCGUA-GACGCGAAACC-AGGUGAUCUAUCCUU
GAUCAGGAUGAAGUAUGG-GUAAAA-CCAUAUGGAGGUCCGA--ACCAGUGU-GGGUUGAAAACCGC-UUGG
AUGAAUUGAGGAUAGGGGUGAAAGGCCAAUCAAACUUGG-UGAUAGCUCGUACUCCCCGAAA-UGUCUUGAG
GGACAGCCUCGG-GGUC-------------------------------------------------------
----------------------GA-----GU-CUGCCGGAGGUAGAGCUACCAAUU-GGG-CUA--GGGCUG
U---CA-CA--ACGGUACCAAACCCAGAUGAACUCCGAAUGCCGGU--------------------------
------------------------------------------------------------------------
------------------AUGA-C-AU--------ACCCGGG-AGUGAGG-G-CAUG-A-GCGAUAAGGUCC
AU-G-UCCGAGAG-GGAAAC---AACCCAGAGCCACA-GCUAAGGCCCCCAAAU--UC-ACG-CU-AA---G
-U-GUAU-----UAAAGAUUGUUUGAUUGCCCAGACAGCUAGGAUGUUGGCUUAGAAGCAGCCA-CCAUUUA
AAGAGUGCGUAAUAGCUCACUAGUC-----AAGCGAUCAUGCGUCGAUAAUA-CUC-GGGACU-A-AGCGUG
G----------UGCCGAA-GCUUGGUGAUUUCUU--U-------------------------------------
------------------------------------------------------------------------
---AGGCUA---------------------------------------------------------------
---------------------------CGGUCUAAAU-GAAU-CGGUAGGGGAGC-AUUCCAG---------
-----------U-U-GCGUU-GAAGGUG--A-GUGGUAACGC--U-UGCUGGAGCG-G-CUG-GAAACGAAC
AUGU-AGGCAU-GAGUA-GCGAUA---AACA-AAGUGA--G---------------AAACUU-UGU-CACCG
A-AAAUCU-AAGGUUUCC-UGAGCAAUG-UUAAUCAUCUCAGGGUUAGUCGGG-AC-CUAAGGUGAGGCCG-
A-A-AGGC-----GUA--------------------------------------------------------
----------------------------------------GCC-GA-U--GGAA-AACUGGUU-AAUA-GUC
CAGUA----------------------------------CCCAA----UCAU--------G-AA--------
------------------GGUUAUUA----------------------------------------------
----------------------------------------------------------CCUAU-GGGGUGAC
GCAGAAG-------------UGAAAG-GAGAGCCAC-CG-G----ACGGA------------AGUGGUGGU-
UUAAGU-GCG-UA-----G-G-------------------C------UG------------------AA-GC
G-UAGG--------------------CAAAU-CCGCGCUUCA------UCAAGGCCG----AAA-AA-C-G-
AAC--AG---------------------------------------------------------------GG
-----CGCUCG-UAAGAGCCACAAC-U-CUC-C-CUAAUCCAGCUGC-C-GAGAAA-AGCCU-CUCUGGGA-
--------------------------------------------------------CGACAU-GA-CUGGCC
CGUACCGUAAACCGAC------------------------------ACAGGUA--GAUGGGAU-GAGUAUUC
UAAGGU-G-CU-CGAG-UGAGACGC-A-GCUAAGGAACUCGGC-AAAAUAGCCCCGUAACUUCGG-GAGAAG
GGGUG-CCUA-G-AU---------------------------------------------------------
------------------------------------------------------------------------
------------------------------------------------------------------------
------------------------------------------------------------------------
------------------------------------------------------------------------
------------------------------------------------------------------------
------------------------------------------------------------------------
------------------------------------------------------------------------
------------------------------------------------------------------------
------------------------------------------------------------------------
```

```
------------------------------------------------------------------------
------------------------------------------------------------------------
------------------------------------------------------------------------
------------------------------------------------------------------------
---------------------------------------------------------------------UUU
CUA--GGCCGCAGUGAAAUGGGUCAAGCGA-----CUGUUUAACAAAAACACAUCUCUCUGCCAAGACGA-U
UUA-A-------------GUCGAUGU-AUAGGGAGUGA-CACCUGCCCGGUGCUG-G-AAGG-UUAAGGGGA
UCUGUCA-GC-----GCAA-----GUGAAGCAGUGAACCGAAGCCC--CAGUAAAC-GGCGGCCGUAACUAU
AACGGUCCUAAGGUAGCGAAAUUCCUUGUCGGG-UAAGUUCCGACCUGCACGAAUGGUGUAACGAUU--UGA
CCACUGUCUCGGCU----GCG-CGCUCGGCGAAAUU--GUA---GUGCCGGUGAAGAUGCCGGCUACUCGCA
UCUGGACGGAAAGA-CCCUAUGCACC-UUUACU--AUAACCU-GGCAUUGGGCUUGAGCAUGAUUUGUGUAG
AAU-AGGUGGGAGACUUUG----------AAGCGGG-GAC----------------------------GUCA
--------------------------------------GU-UUCCGUGGAG-U--C-GCAAUGUGAAAUA-C
CACC-CUGGUUGUGUUUGAGUCCUAACCUCGA--CGAG-CAAA-U-C--------------------------
------------------------------------------------------------------------
------------------------------------------------------------------------
------------------------------------------------------------------------
------------------------------------------------------------------------
------------------------------------------------------CU----CG----------
------------------------------------------------UCAG-----GGACC-AUGUCAGGC
GGGUAGUUUGACUGGGGCGGU-CGCCUCCUA-AAAAGUAAC-GGAGGCUUGCA-AAGGUUCACUCAGUUCCG
UCGGUAAUGGAACGUA-GAGCA-UAAUGGUAG-AAGUGAGCUUGACU-GUGA--------------------
--------------GACCGA---CAGGUCGAGCAGGAACGAAA-GUUGGCCAUAGUGAUCCGGU--------
---GG-U-----CCCGC-AUG--------------------GA-AGGGCCAUCGCUCAUAGG-AUAAAAGGU
ACGCUAGGG-AUAACAGGCUGAUCCCCGCCGAGAGUUCAUAUCGACGUGGGGGUUUGGCACCUCGAUGUCGG
CUCAUCACAUCCUGGGGCUGGA-GAAGGUCCCAAGGGUCCGGCUGUUCGCCGGUUAAAGUGGUACGUGAGCU
GGGUUCAGAACGUCGUGAGACAGUUCGGUCCCUAUCCGAU--------------------G-CGA-------
-------------------------------GCGC--AGGAAAUUUGAGAGG-GCU-UGUCCUUAGUACGAG
AGGAC-CGGGACGAACAAACCUC-UGGU-G-CACCAGUUGUCACGCCAGUGGCAU-G-GCUGGGUAGCUAUG
UUUGGUUGAGCUAAGCGCUGAAAGCA-UC--UAAGUGCGAAACUCGCCUCAA-G-AUG--AGAUUUCC-CC-
--A---------------------------------------------------------------------
------------------------------------------------------------------------
------------------------------------------------------------------------
------------------------------------------------------------------------
------------------------------------------------------------------------
------------------------------------------------------------------------
------------------------------------------------------------------------
------------------------------------------------------------------------
----------------------------------------------------------UC-AGG------CU
CCAGGAAGAUGACCUGGU-CGA-UAGGCU-ACA-GGUGUACG-CAC-A----------------GU------
----AA-----------UGUGUUUAGCCG--A-GUAG-UACUAAUA---AGCCGAUCG-ACUUAACCA----
------------------------------------------------------------------------
------------------------------------------------------------------------
------------------------------------------------------------------------
------------------------------------U------------
```

```
>LSUb_190304_FUSNN/1-2908
---------------------G-UUAAAAUA----AU--UAAGGGCACAC-AAAGGAUG-C-CUAGGU-AGU
AAG---AGCCGAUGAAGGACGUGG--UAAGCUGCGAUAAGCCUAGAUAAGUUGCAAUCGAACGU-----AAG
AGU-CUA-GGAUUUCCGAAUGGAGCAAU--------CU--------AU----UAAG-A-U-----------G
G-A-------------G---U----C----U---UAA----U-A-C--------------------------
-----------G-----------AAA----------------------------------------------
----------------------------GA--GGGAACCGCGUGAACUGAAACAUCUAAGUAACGC--GAGG
AAAAGAAAGUAA--AAACGAUACCCAAA--GU-AG-CGGCGAGCGAACUGGGU-C-AA-GCCUAAACCUUAA
-AUA-UGUCA-AGGAUA-CAG------------C---CGU-UG-UAUUUAAGGGG--UAG------------
-----AGGG-ACA-------AAGU-AGUGAAG-AACUGUAAGAUAUUCAAU--AUA---------GUGU--A
U-------UGAUGAAUUAG------AAU-UG-UCUG------------------------------------
-------GAA-AG-AUGAA-------CCGCAGAAGGUGAAAGUCCUGUA---------UAAGUAA----AUC
C------U--UACAC----------AUAU--AACUU--UG-CU------CCCAAGUAACAUGGAACACGAGG
AAUUCUGUGUGAA-UCAGUGAGG-ACCAAAUCUCAUAAGGCUAAAUA--CUC-UU-AC-UAACCGAUAGC-G
CA--U-AGUACCGUGAGGGAAAG-GUGAAA-AGAAC-CCCUG-GAGGGG-A-GUGAAAUAGAACCUGAAAUU
G-UGUGCUUACAA-GCG-GU-CAGA---GCCC----------------------------------------
------------------------------------------------------------------------
------------------------------------------------------------------------
------------------------------------------------------------------------
------------------------------------------------------------------------
------------------------------------------------------------------------
------------------------------------------------------------------------
------------------------------------------------------------------------
------------------------------------------------------------------------
--------------------------------------------------------------A--U-UU---
------------------------------------------------------------------------
------------------------------------------------------------------------
------------------------------------------------------------------------
------------------------------------------------------------------------
------------------------------------------------------------------------
------------------------------------------------------------------------
------------------------------------------------------------------------
------------------------------------------------------------------------
------------------------------------------------------------------------
------------------------------------------------------------------------
------------------------------------------------------------------------
------------------------------------------------------------------------
-------------------------------------------------G--G---GUGAUGGCGUGCCUUU
UGGAGAAUGAUCCUGCGAGUUAC-G-UUAAACGGCGA-GGUUAAG--------------UAUAA--------
------CGGA--GCCGAAGGGAAAC-C---------------------------------------------
------------------------------------------------------------------------
------------------------------------------------------------------------
---------------------------------------------------------------------AAG
UCU------------------------------------UAAUA-------------------------GG-
G-C-GA----------------------AUUAGUCGUUUGGCGUA-GACGCGAAACC-UGGUGAUCUAAACCU
```

```
GUCCAGGAUGAAGCUGUG-GUAAGA-CACAGUGGAGGUCCUA--ACCCACCG-CCGUUGAAAAGUUG-GGGG
AUGAGGUAGGUUUAGGGGUGAAAAGCCAAUCGAACCAGG-AGAUAGCUCGUUCUCUCCGAAA-UGCAUCUAG
GUGCAGCCUUGA-GUG--------------------------------------------------------
----------------------UU-----CA-AUUAUGGGGGUAGAGC-ACUGAAU-GAU-CUA--GGGGGC
----AUAUU---GCUUACUGAAAUCAAUCAAACUCCGAAUACCAUA--------------------------
------------------------------------------------------------------------
------------------AUUU-A-UA--------GCUCAGG-AGUGAGA-C-UAUG-G-GAGUUAACUUCC
AU-U-GUCAAAAG-GGAAAC---AACCCAGACCACCA-GCUAAGGUCCCUAAUU--AU-AA--CU-AA---G
-U-GGG-------AAAGGAGGUGGAGAUUCACAAACAACUAGGAGGUUGGCUUAGAAGCAGCCAUACCUUUA
AAGAGUGCGUAAUAGCUCACUAGUC-----GAGAGUCUCUGCGCCGACAAUGUAAC-GGGGCU-A-AGUUAU
A----------AACCGAA-GCUGUGGAAUCCUUU--------------------------------------
------------------------------------------------------------------------
------------------------------------------------------------------------
------------------------------------U-GGAU-UGGUAGGAGAGC-GUUCUGU---------
-----------AGG-CCGUU-GAAGAAG--A-AGGGUAACCG--A-CUUUGGAGGU-A-UCA-GAAGUGAGA
AUGC-AGGAAU-AAGUA-GCGAGA---AAGG-GGGCGA--G---------------AAUCCU-CCU-CGCCG
G-AAGACC-AAGGUUUUC-AGGGUAAAG-CUUGUCUUCCCUGAGUAAGCCGGG-AC-CUAAGCCCAGGCUA-
U-A-AUGC-----GUA--------------------------------------------------------
----------------------------------------GGC-GAAU--GGAA-AACAGAUU-AAUA-UUU
CUGUG----------------------------------CCAGU----CAUG--------U-AUU-------
------------------------------------------------------------------------
----------------------------------------------------------GUGAU-GGAGGGAC
GCAGAA--------------GGGUAU-GCGCGCGGA-CG-A----ACGGA------------AGUGUCCGU-
AGAAGU-AUG-UA-----G-G-------------------G------UG-----------------ACU-UA
G-UAGG--------------------UAAAU-CCAUUAAGUU--------AAAUCUG----AGG-UA-U-G-
AUA--UA---------------------------------------------------------------CA
-----GUCG------UAAGAUGAAU-G-CGC-A-AAUCCCACGCUGC-C-AAGAAA-AGCUU-CUAACGUU-
--------------------------------------------------------AAUAUA-UG-ACUGCC
CGUACUGUAAACCGAC------------------------------ACAGGUG--GUCAGGAU-GAGAAAUC
UAAGGC-G-GA-CAGG-CUAACUCU-C-GUUAAGGAACUCUGC-AAAAUAACCUCGUAACUUCGG-GAGAAG
AGGAG-CCCU-U-GU--GUGUGAGU----AUA-----------CACGC------------------------
------------------------------------------------------------------------
------------------------------------------------------------------------
------------------------------------------------------------------------
------------------------------------------------------------------------
------------------------------------------------------------------------
------------------------------------------------------------------------
------------------------------------------------------------------------
------------------------------------------------------------------------
------------------------------------------------------------------------
------------------------------------------------------------------------
------------------------------------------------------------------------
------------------------------------------------------------------------
------------------------------------------------------------------------
--------------------------------------------------------GAUACAAA--GCGCAC
```

```
GAG--GGUCGCAGUGAAGAGGCUCAAGCAA-----CUGUUUAACAAAAACACAGGUCUAUGCUAAGCUG---
-UA-A-------------GGCGAUGU-AUAUGGGCUGA-CACCUGCCCAGUGCUG-G-AAGG-UUAAGAGGA
GGAGU----------GAGA----------GCUCCGAAUUGAAGCCC--CAGUGAAC-GGCGGCCGUAACUAU
AACGGUCCUAAGGUAGCGAAAUUCCUUGUCGGG-UAAGUUCCGACCUGCACGAAUGGUGUAAUGAUU--UGA
GCGCUGUCUUGACG----GGA-GGCCUGGUGAAAUU--GUA---UUACCGGUGAAGAUACCGGUUACCUACA
GUAGGACGGAAAGA-CCCCAUGGAGC-UUUACU--GUAGCUU-GGUAUUGGGUUUUGGCAUUGCAUGUAUAG
GAU-AGUUGGGAGACUAUG----------AUGAUAU-GGC----------------------------GCUA
--------------------------------------GC-UGUAUCGGAG-U--C-AU-CGGUGGAAUA-C
CAAC-CAUUCAAUGCUGAAAUUCUAAUCU-GU--GGU--U-UG-U-A-------------------------
------------------------------------------------------------------------
------------------------------------------------------------------------
------------------------------------------------------------------------
------------------------------------------------------------------------
-------------------------------------------------------G----CC----------
------------------------------------------------ACGG-----AGACA-GUGCUAGGU
GGGCAGUUUGACUGGGGCGGU-CGCCUCCGA-AAGAGUAAC-GGAGGCGUUCA-AAGGUUCUCUCAGGUUGG
AUGGAAAUCAACCAUA-GAGUG-CAAUGGCAU-AAGAGAGCUUGACU-GCAA--------------------
--------------GACUGA---CGGGUCGAGCAGAUGCGAAA-GCAGGACAUAGUGAUCCGGC--------
---GA-U-----UCCGA-AUG--------------------GA-AGGGUCGUCGCUCAACGG-AUAAAAGCU
ACCCUGGGG-AUAACAGGCUGAUCCUACCCGAGAGUCCAUAUCGACGGUAGGGUUUGGCACCUCGAUGUCGG
CUCAUCGCAUCCUGGGGCUGGA-GAAGGUCCCAAGGGUUGGGCUGUUCGCCCAUUAAAGCGGUACGUGAGCU
GGGUUCAGAACGUCGUGAGACAGUUCGGUCCCUAUCCACU--------------------G-UAG-------
-------------------------------GCGU--UAGAAUAUUGAGAAG-ACC-UGUCCUUAGUACGAG
AGGAC-CGGGAUGGACAAACCUC-UGAU-G-UACCAGUUGUCACGCCAGUGGCAC-A-GCUGGGUAGUCACG
UUUGGAAUAGAUAACCGCUGAAAGCA-UC--UAAGCGGGAAACUAACUUCAA-G-AUA--AGUAUUCU-U--
------------------------------------------------------------------------
------------------------------------------------------------------------
------------------------------------------------------------------------
------------------------------------------------------------------------
------------------------------------------------------------------------
------------------------------------------------------------------------
------------------------------------------------------------------------
------------------------------------------------------------------------
----------------------------------------------------------UA-AGA------UA
CCUUCGAGCCUAGGAGGU-UGA-UAGGUU-GGG-GGUGUAAG-UAC-A----------------GC------
----AA-----------UGUAUUUAGCUG--A-CCAA-UACUAAUU---AUCGAA--G-UUUUAAUCU----
------------------------------------------------------------------------
------------------------------------------------------------------------
------------------------------------------------------------------------
------------------------------------A------------
>LSUb_243276_TREPA/1-2947
----------------------------GCG----AA--UAGUGGUUUAC-GGUGGAUG-U-CUUGGA-GUU
GUC---AGGCGAUGAAGGUCGUGA--UAAGCUGCGAAAAGCCUCGGGGAGGAGCACAUGUCCUG------UG
AUC-CGG-GGAUGACCGAAUGGGGUAAC--------CC--------GA----CAGG-G-U-----------A
A-A-------------G---C----C----U---UGU----C-A-UUGC-CUU-----CCUGA---------
```

```
-----------A-----------UGAA---------------------------------------UAGG-G
A-G--------------------GG-U-AAGGCGAAACUGGGUGAACUGAACCAUCUAAGUAACUU--G-GG
AAAAGAAAUCAA--GAGAGAUUCCGAAA--GU-AG-UGGCGAGCGAAAUUGGA-G-GA-GCCUAAACCUG--
--UG-UCUAA-------------------------------CAGGGGUUGUAGGG--CCG------------
----CGCGG-GCU-------UGCGUUC---------------------GGU--GGGUGAAAUA-AUCCG---
G-------CCUAU-AGCAG-A----AAG-GU-UUUG------------------------------------
-------GGA-AA-GCCUG-------ACAGAGAGGGUGAAAUCCCGUA----------UGCGGAA----UGG
G------G--CGGAC----------CUGCU-GGUGC--GG-UA------CCUGAGUACGGCGGGACACGAGG
AAUCCUGUCGGAA-UCUGGGUCG-ACC--ACGAUCUAAGGCUAAAUA--CUC-GACAA-CUACCGAUAGU-G
GAC-A-AGUACCGUGAGGGAAAG-AUGAAA-AGAAC-CCCGGUGAGGGG-A-GUGAAAUAGAACCUGAAACC
G-UAAACCAACAA-GAU-GU-UACA---GCCU----------------------------------------
------------------------------------------------------------------------
------------------------------------------------------------------------
------------------------------------------------------------------------
------------------------------------------------------------------------
------------------------------------------------------------------------
------------------------------------------------------------------------
------------------------------------------------------------------------
------------------------------------------------------------------------
--------------------------------------------------------------G--U-GA---
------------------------------------------------------------------------
------------------------------------------------------------------------
------------------------------------------------------------------------
------------------------------------------------------------------------
------------------------------------------------------------------------
------------------------------------------------------------------------
------------------------------------------------------------------------
------------------------------------------------------------------------
------------------------------------------------------------------------
------------------------------------------------------------------------
------------------------------------------------------------------------
------------------------------------------------------------------------
-------------------------------------------------G--G---GUGGUAGCGUGCCUUU
UGUAGAAUGAGCCUGCGAGUUAC-G-GUGUGCAGCAA-GGUUAAGG--G-AU--------AGGA-------G
U-CC--UGGA--GCCGGAGGGAAAC-C---------------------------------------------
------------------------------------------------------------------------
------------------------------------------------------------------------
---------------------------------------------------------------------GAG
UCU------------------------------------UAAAA-------------------------GG-
G-C-GU--------------------GGUGAGUUGUACGUCGUA-GACCCGAAGCCAGGGUGAUCUAGUUAU
GAGCAGGUUGAAACAGGG-GUGAAG-CCUUGUGGAGGACCGA--ACUAUAAU-CUGUUAAAAAAGGU-AUGG
AUGACUUGUGACUAGGAGUGAAAGGCUAAACAAACCUGG-AGAUAGCUGGUUCUCCCCGAAA-UGCCUUUAG
GGACAGCCUUAU-ACA--------------------------------------------------------
-----------------------A-----AA-CUGUCGGAGGUAAAGC-ACUGGAU-GGG-CUA--GGGGGU
U---UC-AU--CGCCUACCAAACCCAAUCAAACUCUGAAUGCCGGC--------------------------
```

```
------------------------------------------------------------------------
------------------AGUC-A-AC--------GUGUGGG-AGUGAGA-C-UGCG-U-GCGACAAGGUUC
GU-A-GUCGAGAG-GGAAAC---AGCCCAGACCGUCA-GCUAAGGUCCCGAAAU--AC-CGC-UU-GA---G
-U-GUG-------AAAUGAAGUGUGGGUACCUGGACAGCCAGGAGGUUGGCUUAGAAGCAGCCAUUCCUUGA
AAGAGUGCGUAAUAGCUCACUGGUC-----GAGUACGCAUGCGCAGAUAAUGUAUC-GGGGCU-A-AGCGGU
A----------UACCGAA-GCUACGGGUCUUGCA--U-----------------------------------
------------------------------------------------------------------------
-----UUU----------------------------------------------------------------
-------------------------------UGGUGC-AAGG-CGGUAGGGGAGC-AUUCCAU---------
-----------GUA-CUGAU-GAAGGAA--UAUCCGGGAGGA--G-UUCUGGAGGG-G-AUG-GAAGAGAGA
AUGC-AGGUAU-AAGUACACGAAA---AGGA-GGGUGA--G----------------AUUCCU-UCC-CGCCG
A-AAACCU-AAGGUUUCC-UGGGUGAAG-GUCAUCUGCUCAGGGUAAGUCGGC-CC-CUAAGGCGAGGACG-
A-G-GGUC-----GUA--------------------------------------------------------
----------------------------------------GUC-GA-U--GGGA-AUCCGGUU-UAUA-UUC
CGGAA----------------------------------CCUCU----UGCA--------A-UU--------
------------------------------------------------------------------------
----------------------------------------------------------UCGAU-GGCAGGAC
GCGUGA--------------GGUGAAGCCCGGCCAA-AG-A----UUGGU------------AGUUUUGGU-
CUAAGU-AUC-CGA----G-C-------------------CGUUUUAAG------------------AG-CG
A-UAGG--------------------CAAAU-CCGUCGUUCGA----GGUAAGGUGC----GAG-UG-C-G-
ACU--GG---------------------------------------------------------------AG
C----GAUGAG----CGAAGGGAAG-C-AGG-U-GUAGUCAUGGCGA-C-GGGAAAUACUGU-CUAAGGUU-
--------------------------------------------------------AGGUUG-CA-AGAGAC
CGUACCGCAAACCGAC------------------------------ACAGGUA--GGUAGGAU-GAGUAAUC
UAAGGC-G-CU-CGAG-AGAACUCG-C-GUCAAGGAACUCGGC-AAAAUACACACGUAACCUCGG-GAGAAG
UGUGA-CCCU-U-GC--CU-----------------------------------------------------
------------------------------------------------------------------------
------------------------------------------------------------------------
------------------------------------------------------------------------
------------------------------------------------------------------------
------------------------------------------------------------------------
------------------------------------------------------------------------
------------------------------------------------------------------------
------------------------------------------------------------------------
------------------------------------------------------------------------
------------------------------------------------------------------------
------------------------------------------------------------------------
------------------------------------------------------------------------
------------------------------------------------------------------------
-------------------------------------------------------------------UUGGU
GAG--GGUGGCAGAAAGCAGGUCCAGGCGA-----CUGUUUAUCAAAAACAUAGCCAUCUGCAAAUCAG---
-UA-A-------------UGAGACGU-AUAGGUGGUGA-CACCUGCCCGGUGCUG-G-AAGG-UUAAGAGGA
GAGGUUC-GUGG---UAAC-----ACAACGCUUUGAAUUGAAGCCC--CAGUAAAC-GGCGGCCGUAACUAU
AACGGUCCUAAGGUAGCGAAAUUCCUUGUCGGG-UAAGUUCCGACCCGCACGAAUGGUGUAACGACU-CUGG
ACACUGUCUCGACG----CGA-GACUCGGUGAAAUU--UAU---GUACCGGUAAAGAAGCCGGUUACCCAUA
```

```
GUUAGACGGAAAGA-CCCCGUGAACC-UUCACC--GUAGCUU-ACUAUUGGAACUUGGUUUACCAUGUGUAG
UAU-AGGUGGGAGACAGAG----------AAGCUUG-GCC----------------------------GUCA
--------------------------------------GG-UUAGGCGGAG-U--C-AA-CAGUGAAAUA-C
CACC-CUUGGUACGUCAGGUUUCUAACCUUUG--GCCG-UGGA-U-C-------------------------
------------------------------------------------------------------------
------------------------------------------------------------------------
------------------------------------------------------------------------
------------------------------------------------------------------------
------------------------------------------------------CG----GC----------
------------------------------------------------AAAG-----GGACC-GUGGUAGGU
GGGCGGUUUGACUGGGGCGGU-CGCCUCCUA-AAAGGUAAC-GGAGGUGCGCG-AAGGUCUCCUCACACCGG
UUGGAAAUCGGUGCGC-GAGUG-UAAAGGCAC-AAGGAGGCUUAACU-GCGA--------------------
--------------GACCGA---CAGGUCGAGCAGAUACGAAA-GUAGGUCUUAGUGAUCUGGC--------
---GG-U-----AGCGU-GUG--------------------GA-AGCGCCGUCACUUAACGG-AUAAAAGGU
ACUCCGGGG-AUAACAGGCUGAUUUCCCCAAGAGUUCACAUCGACGGGAAAGUUUGGCACCUCGAUGUCGG
CUCAUCGCAUCCUGGGGCUGAA-GCAGGUCCCAAGGGUUUGGCUGUUCGCCAAUUAAAGCGGUACGUGAGCU
GGGUUCAGAACGUCGCGAGACAGUUCGGUCCCUAUCUGCU--------------------A-UGG-------
-------------------------------GCGU--UGGAUAUGUGAGAGG-AGC-UGCUUUUAGUACGAG
AGGAC-CGAAGUGGACGAACCUC-UGGU-G-UACCAGUUAUCCUGCCAAGGGUAC-GUGCUGGGUAGCUAUG
UUCGGAAGGGAUAACCGCUGAAGGCA-UC--UAAGUGGGAAGCCCGCCUCAA-G-AUU--ACAUAUCC-CU-
--GAAGG-----------------------------------------------------------------
--------------------------UU---------GA---------------------------------
------------------------------------------------------------------------
------------------------------------------------------------------------
------------------------------------------------------------------------
------------------------------------------------------------------------
------------------------------------------------------------------------
------------------------------------------------------------------------
-------------------------------------------------CCUUCCU--GA-AGA------CU
CCUUGCACACUACAAGGU-CGA-UAGGCU-GGA-GGUCUACG-UAC-C----------------GU------
----AA-----------GGUAUUAAGCCG--A-CCAG-UACUAAUA---AGUCGUGAG-GCUUGACCA----
------------------------------------------------------------------------
------------------------------------------------------------------------
------------------------------------------------------------------------
------------------------------------U------------
>LSUb_227941_CHLCV/1-2939
---------------------G-ACCAAGUU----GU--UAAGAGCUAUU-GGCGGAUG-C-CUUGGC-AUU
GAC---AGGCGAUGAAGGAUGCGU--UUACCUGCAGUAAUCUUCGGCGAGCUGGUAUAAAGCUA------UG
ACC-CGG-AGGUAUCCGAAUGGGGUAAC--------CC--------GG----UAGA-U-U-----------A
A-U-------------C---G----U----C---UAC----C-A-UUAU-AUA-----CUGA----------
-----------A-----------UACA----------------------------------------UA-GG
U-A--------------------UA-U-AAAGCGACACCUGCUGAACUGAAACAUCUUAGUAAGCA--GAGG
AAAAUAAAUCAA---AGAGAUUCCCUAA--GU-AG-CGGCGAGCGAAAGGGGA-G-AA-GACCAAACCACG-
--CC-UU-------------------------------------UGG-UGUGGG--UUG-------------
-----UAGG-GUC-------GAUAACA----------------------UG--AGA---------UCUU--A
```

```
A-------GUUUU-AGUUG------AAUACU-UCUG------------------------------------
-------GAA-AG-UAGAA-------CGAUACAGGGUGAUAGUCCCGUA---------AACGAAA----GAA
C------A--AGAGA----------CUCU---AUUC--GA-UG------CCUGAGUAGGGCUAGACACGUGA
AACCUAGUCUGAA-UCUGGGGAG-ACC--ACUCUCCAAGUCUAAAUA--CUA-GUCAA-UGACCUAUAGU-G
AAC-C-AGUACUGUGAAGGAAAG-GUGAAA-AGAAC-CCUUGUUAAGGG-A-GUGAAAUAGAACCUGAAACC
A-GUAGCUUAUAA-GCG-GU-CGGA---GACC-U---AU---------------------------------
------------------------------------------------------------------------
------------------------------------------------------------------------
------------------------------------------------------------------------
------------------------------------------------------------------------
------------------------------------------------------------------------
------------------------------------------------------------AG-CUUC-----
------------------------------------------------------------------------
------------------------------------------------------------------------
--------------------------------------------------------------C--U-CG---
------------------------------------------------------------------------
------------------------------------------------------------------------
------------------------------------------------------------------------
------------------------------------------------------------------------
------------------------------------------------------------------------
------------------------------------------------------------------------
------------------------------------------------------------------------
------------------------------------------------------------------------
------------------------------------------------------------------------
------------------------------------------------------------------------
------------------------------------------------------------------------
------------------------------------------------------------------------
----------------------GAAGC-------------------CAUG--G---UUGACGGCGUGCCUUU
UGCAUGAUGAGCCAGGGAGUUAA-G-UUAAACGGCGA-GGUUAAGG--G-AU--------CUAC-------A
U-UC--CGGA--GCCGAAGCGAAAG-C---------------------------------------------
------------------------------------------------------------------------
------------------------------------------------------------------------
---------------------------------------------------------------------GAG
UUU------------------------------------UAAUA-------------------------GA-
G-C-GU----------------------AUAGUCGUUUGAUUUA-GACACGAAACC-AAGUGAGCUAUUUAU
GACCAGGUUGAAGCAUGG-GUAAGA-CCUUGUGGAGGACCGA--ACCAGUAC-AUGUUGAAAAAUGU-UUGG
AUGAGUUGUGAAUAGGGGUGAAAGGCCAAUCAAACUUGG-AGAUAUCUUGUUCUCUCCGAAA-UAACUUUAG
GGUUAGCCUCGG-AUA--------------------------------------------------------
----------------------UU-----AAGCUUUGGGGGGUAGAGC-ACUGAAU-UCU-AGC--GGGGGC
C---UA-CC--GGCUUACCAACGGAAAUCAAACUCCGAAUACCAAA--------------------------
------------------------------------------------------------------------
------------------A--G-C-GA--------GUCCGGG-AGAUAGA-C-AGCG-G-GGGCUAAGCUUC
GU-U-GUCGAGAG-GGGAAC---AGCCCAGACCGCCG-AUUAAGGUCCCAAAUU--UU-AUG-CU-AA---G
-U-GAG-------UAAGGAAGUGAUGAUUCUAAGACAGUUGGAAUGUUGGCUUAGAGGCAGCAA-UCAUUUA
AAGAGUGCGUAACAGCUCACCAAUC-----GAGAAUCAUCGCGCCAAUAAUA-AUC-GGGGCUCA-AGCAUA
```

```
A----------AACCGAC-AUCGCGGGUGUAUAU--U-----------------------------------
------------------------------------------------------------------------
------------------------------------------------------------------------
-------------------------------UAUAUA-UACG-CGGUAGGAGAGC-GUAGUAU---------
-----------UCA-GCAGU-GAAGGUA--U-ACCGUAAGGA--G-UGCUGGAGCG-G-AUA-CUAGUGAAG
AUCCAUGGCAU-AAGUA-ACGAUA---AAGG-AAGUGA--A---------------AAUCUU-CCU-CGCCG
U-AAGCAC-AAGGUUUCC-AGGGUCAAG-CUCGUCUUCCCUGGGUUAGUCGGC-CC-CUAAGUCGAGGCAC-
A-A-AUGC-----GUA--------------------------------------------------------
----------------------------------------GAC-GA-U--GGAGCAACAGGUUAAAUA-UUC
CUGUA----------------------------------CCACC----UAAA--------A-C---------
------------------UUUA--------------------------------------------------
----------------------------------------------------------GCAAU-GGAAUGAC
GGAGUA--------------CGUUAA-GCACGCGGA-CG-A----UUGGA------------AAUGUCCGU-
AUCACA-AUG-A------G-A-------------------C------UG------------------GU-UA
G-UAGG--------------------UAAAU-CCGCUAACAU--------AAGGUUA----GGU-UG-U-G-
GUUAAGG---------------------------------------------------------------GA
-----AAUCUU-CGGAGGAACUGAU-G-GUG-U-AGCGCAAUGCUUU-C-AAGAAAUAAUUU-CUAGCU---
----------------------------------------------------------GUUG-AU-GGUGAC
CGUACC-UAAACCGAC------------------------------ACAGGUG--UGCGAGAU-GAGUAUUC
UAAGGC-G-CG-CGAG-AUAACUUU-C-GCUAAGGAACUCGGC-AAAUUAUCCCCGUAACUUCGG-AAGAAG
GGGAG-CCUU-C-UA--GGGUGAUU----ACC-----------UUUUC------------------------
------------------------------------------------------------------------
------------------------------------------------------------------------
------------------------------------------------------------------------
------------------------------------------------------------------------
------------------------------------------------------------------------
------------------------------------------------------------------------
------------------------------------------------------------------------
------------------------------------------------------------------------
------------------------------------------------------------------------
------------------------------------------------------------------------
------------------------------------------------------------------------
------------------------------------------------------------------------
------------------------------------------------------------------------
--------------------------------------------------------GGUA-UGA--GCCCCG
GGA--GGCCGCAGAGAAAUGGCCCAGGCGA-----CUGUUUAACAAAAACACAGCACUAUGCAAACCUC---
-UA-A-------------GGGGAAGU-AUAUGGUGUGA-CGCCUGCCCAAUGCCA-A-AAGG-UUAAAGGGA
UAUGUCA-GCC----GUAA----GGCAAAGCAUUGAACCCAAGCCC--UGGUGAAU-GGCCGCCGUAACUAU
AACGGUGCUAAGGUAGCGAAAUUCCUUGUCGGG-UAAGUUCCGACCUGCACGAAUGGUGUAACGAUC--UGG
GCACUGUCUCAGCG----AAA-GACUCGGUGAAAUU--GUA---GUAGCAGUGAAGAUGCUGUUUACCCGCA
AAAGGACGAAAAGA-CCCCGUGAACC-UUUACU--GUACUUU-GGUAUUGAUUCUUGAUUUGUUAUGUGUAG
CAU-AGCCAGGAGACUGUG----------AACACUC-UUC----------------------------GUCA
--------------------------------------GG-AGAGUGGGAG-U--C-AU-CGUUGAAAUA-C
UGGU-CUUAACAAGUUGGGAGUCUAACAUAAU--UCCA-UGAA-U-C-------------------------
------------------------------------------------------------------------
```

------------------------------------------------------------------------
------------------------------------------------------------------------
------------------------------------------------------------------------
------------------------------------------------------UG----GA----------
------------------------------------------------AUAU-----GGACA-UUGCCAGAC
GGGCAGUUUUACUGGGGCGGU-AUCCUCCUA-AAAAGUAAC-GGAGGAGCCCA-AAGCUUAUUUCAUCGUGG
UUGGCAAUCACGAGUA-GAGCG-UAAAGGUAU-AAAAUAGGUUGACU-GCAA--------------------
--------------GACCUA---CAAGUCAAGCAGAGACGAAA-GUCGGGCUUAGUGAUCCGGC--------
---GG-U-----GGAAA-GUG--------------------GA-AUCGCCGUCGCUUAACGG-AUAAAAGGU
ACUCCGGGG-AUAACAGGCUGAUCGCCACCAAGAGUUCAUCGACGUGGCGGUUUGGCACCUCGAUGUCGG
CUCAUCGCAUCCUGGGGCUGGA-GAAGGUCCCAAGGGUUUGGCUGUUCGCCAAUUAAAGCGGUACGCGAGCU
GGGUUCAAAACGUCGUGAGACAGUUUGGUCUCUAUCCUUU--------------------G-UGG-------
-------------------------------GCGC--AGGAUACUUGAAAGG-AGC-UGUUCCUAGUACGAG
AGGAC-CGGAAUGGACGAACCAA-UGGU-G-UGUCGGUUGUUUGCCAAAAGCAU-A-GCCGAGUAGCUACG
UUCGGAAAGGAUAAGCAUUGAAAGCA-UC--UAAAUGCCAAGCCUCCCUUAA-G-AUA--AGGUAUCC-CU-
--A---------------------------------------------------------------------
------------------------------------------------------------------------
------------------------------------------------------------------------
------------------------------------------------------------------------
------------------------------------------------------------------------
------------------------------------------------------------------------
------------------------------------------------------------------------
------------------------------------------------------------------------
----------------------------------------------------------UG-AGA------CU
CCAUAUAGACUAUGUGGU-UGA-UAGGUU-GGG-UGUGUACG-CAC-A----------------GU------
----AA-----------UGUGUUUAGCUA--A-CCAA-UACUAAUA---AGUCCAUUG-ACUUGGUUU----
------------------------------------------------------------------------
------------------------------------------------------------------------
------------------------------------------------------------------------
------------------------------------U------------
>LSUb_243090_RHOBA/1-2891
---------------------G-GCCAAUCU----AU--UAAGGGCACAU-GGGGGAUG-U-CUUGGC-GUU
AGA---AGG---AGAAGGGCGUGG--AAGUCUGCGAAAAGCCUGGGGAAGUUGACACACAAGCG-----AUG
AUC-CCG-GGUUCCCGA-----------------------------CC------------------------
------------------------------------A----C-A-GCAC-ACU-----GA------------
-----------A-----------UACA----------------------------------------UAG-G
U-G--------------------UG-U-UUGUAACAACCUGGUGAACUGAAACAUCUAAGUAACCA--GAGG
AACAGAAAGAAA--AAUCGAUUCCGUCA--GU-AG-CGGCGAGCGAAAUCGGA-A-UA-GCCCAAACCGAGG
-AGG-U-----------------------------------UU-CCUUCUCGGGG--UUG------------
-----UAGG-ACU-------UCAA-UA----------------------UG--GGA---------UUGAG-A
C-------GUGUU-AGUGA------AAC-GA-CUUG------------------------------------
-------GAA-CG-GUCGG-------CCACAGAGGGUGACAGCCCCGUA---------CGCGAAA----AUA
U------U--GUCUC----------ACCU---UAGA--AG-CA------CCUGAGUAGGUCCAGUCACGUGA
AACCUGGACUGAA-UCCGCGGGG-ACC--AUCCCGUAAGGCUAAAUA--CUC-UCUAA-CGACCGAUAGC-G
AAC-C-AGUAGCGUGAGUGAAUG-GUGAGA-AGAAC-CCCUGCUAGGGG-A-G-GAAUGUGAACCUGAAACC

```
A-UGUGCCUACAA-GCG-GU-CGGA---GCAC-U---A----------------------------------
------------------------------------------------------------------------
------------------------------------------------------------------------
------------------------------------------------------------------------
------------------------------------------------------------------------
------------------------------------------------------------------------
------------------------------------------------------------------------
------------------------------------------------------------------------
------------------------------------------------------------------------
-------------------------------------------------------------UU--U-UA---
------------------------------------------------------------------------
------------------------------------------------------------------------
------------------------------------------------------------------------
------------------------------------------------------------------------
------------------------------------------------------------------------
------------------------------------------------------------------------
------------------------------------------------------------------------
------------------------------------------------------------------------
------------------------------------------------------------------------
------------------------------------------------------------------------
------------------------------------------------------------------------
------------------------------------------------------------------------
-----------------------------------------------AUG--U---GUGACGGCGUGCCUUU
UGCAUAAUGAUCCGGCGAGUUGU-G-GUAAUCGGCUU-GGUUAAGU--U-C--------UGACG--------
G-AA--CGAA--GCCCAAGGGAAAC-C---------------------------------------------
------------------------------------------------------------------------
------------------------------------------------------------------------
---------------------------------------------------------------------AAG
UCU------------------------------------GAAUA-------------------------GG-
G-C-GU-----------------------UGGUCGAUUGCUGCA-GACGCGAAACCUGUGUGAUCUACCCAU
GGGCAGGUUGAAGCGCGG-GUUAAA-CUGCGUGGAGGACCGA--ACCCACUU-AGGUUGAAAACUGA-GGGG
AUGACCUGUGGGGAGGACUGAAAGUGUAAUCAAACCAGG-AGAUAGCUCGUUCUCUCCGAAA-UAUCUUGAG
GGAUAGCGUCGA-GUAA-------------------------------------------------------
----------------------AU-----UAGUGACCGGGGGUAGAGAUACUGAAU-CGG-AUG--GGGGCC
C---UUCCA--AGGGUACCCCACCGAACCAAACUCCGAAUACCGGU--------------------------
------------------------------------------------------------------------
------------------UCACCU-UA--------CCUCGGC-AGUCAGU-C-CCUG-G-GGGAUAAGCUCC
AG-G-GUCGAGAG-GGAAAC---AACCCAGAUCGCCU-GCUAAGGUCCCGAAGU--AC-UA--CU-UA---G
-U-CAC-------UAAGGAAGUUGAAGUGCCGUGACAGCUGGGAUGUUGGCUUAGAAGCAGCCA-CCAUUUA
AAAAGUGCGUAACAGCUUACCAGUC-----GAGCAUUUCUGCGCCGAUAAUG-AAC-GGGAGU-A-AGUAGU
A----------CACCGAA-GCAGCGGAUUCUAGA--U-----------------------------------
------------------------------------------------------------------------
------------------------------------------------------------------------
-------------------------------UUAUCU-AGAG-UGGUAGGAGAGC-GCCGAUG---------
-----------A-U-CAGAUACGAGCCG--U-ACGGUAACGA--G-CGGUGUCGGG-G-UCAUCGGGUGAUU
```

```
AUGC-CGGAAU-GAGUA-ACGAUA---AAAC-AGGUGA--G---------------AAUCCU-GUU-CGCCG
A-AAGCCU-AAGGUUUCC-UGGGGAAGG-UAAAUCCGCCCAGGGUUAGCCGGU-AC-CUUAGUCGAGGCCG-
A-A-AGGC-----GUA--------------------------------------------------------
----------------------------------------GAC-GA-U--GGAU-AGAAGGUC-AAUA-UUC
CUUCG----------------------------------CCACA----UGGG-------UA-CU--------
------------------------------------------------------------------------
----------------------------------------------------------UCGAU-GCGGGGAC
GGCGUC--------------CAGAAA-GUGAGCGGU-ACGA----AUAGA------------AAUGUCCGU-
AGCGAA-AAC-UGA----G-G-------------------U--------------------------GU-UG
G-GGGA--------------------UAAAUGACCUGACAU---------AAGCCAA----GGU-UG-A-G-
CUC--GA-----------------------------------------------------------------
-----GAGCAU--CCAAGUUCUCGA-A-GUC-A-UUGGAAGGACGUC-C-AAGAAA-AGCCG-CUAAGUU--
--------------------------------------------------------AUGUGC-CC-UGUGAC
CGUACUA-AAACUGAC------------------------------ACAGGUA--GGCGAGAU-GAGAAUUC
UAAGGC-G-CU-CGGG-AGAACGGU-G-GUUAAGGAACUCUGC-AAAAUGGCCCCGUAAGUUCGC-GAUAAG
GGGCG-CCUC-C-GA--CGGUU---------------------------------------------------
------------------------------------------------------------------------
------------------------------------------------------------------------
------------------------------------------------------------------------
------------------------------------------------------------------------
------------------------------------------------------------------------
------------------------------------------------------------------------
------------------------------------------------------------------------
------------------------------------------------------------------------
------------------------------------------------------------------------
------------------------------------------------------------------------
------------------------------------------------------------------------
------------------------------------------------------------------------
------------------------------------------------------------------------
---------------------------------------------------------------CACGCUGUU
GGA--GGCCACAGAAAAUCGGCUCUAACGA-----CUGUUUAUCAAAAACACAGGUCUCUGCUAACACG---
-CA-A-------------GUGGAUGU-AUAGAGACUGA-CGCCUGCCCGGUGUUG-G-UAGG-UUAAGGAAG
ACGGUUA-GCUC---UUCG---GAGCGAAGCUGGCGACCGAAGCCC--CAAUAAAC-GGCGGCCCUAACUAU
GAGGGUCCUAAGGUAGCGAAGUUCCUUGUCGGG-UAAGUUCCGACCUGCAUGAAAGGCGUAACGAUU--GGA
GCACUGUCUCAACC----ACC-GACCCGGUGAAAUU--GUA---GUUGUGGUGAAGAUGCCACAUACCCGCG
GUUAGACGGAAAGA-CCCCGUGAACC-UUUACU--GUAGGCU-GCUAUUGGGUUGAGAUAUGUCUUGUGUAG
GAU-AGCUGGGAGACUUUG----------AACUUUG-CAC----------------------------GCUA
--------------------------------------GU-GUAGAGGGAG-U--C-GU-UGUUGAAAUA-C
CAGC-CUGGUCAUAUUUUAAUUCUAACUU-UGAUCCCC-UGAA-A-C-------------------------
------------------------------------------------------------------------
------------------------------------------------------------------------
------------------------------------------------------------------------
------------------------------------------------------------------------
------------------------------------------------------GG----GG----------
------------------------------------------------CAAA-----GGACA-GUGUCAGUC
```

GGGCAGUUUGACUGGGGCGGU-CUCCUCCCA-AAGAGUAAC-GGAGGAGUGCA-AAGGUACACUCAGCCUGG
UCGGCAAUCAGGCGUA-GAGCG-UAAAGGUAG-AAGUGUGCUUGACU-GCGA--------------------
--------------GACUCA---UAAGUCGAGCAGGAACGAAA-GUUGGCCUUAGUGAUCCGGU--------
---AA-U-----CCCGA-AUG--------------------GA-AGGGUUAUCGCUCAUCAG-AUAAAAGGU
ACUCCGGGG-AUAACAGGCUUAUCGCAUCCGAGCGUCCACAGCGGCGAUGCGGUUUGGCACCUCGAUGUCGG
CUCAUCACAUCCUGGGGGUGUA-GAAGCUCCCAAGGGUUUGGCUGUUCGCCAAUGAAAGUGGUACGUGAGCU
GGGUUCAGACCGUCGUGAGACAGGUCGGUCCCUAUCUGCC--------------------G-UGG-------
-------------------------------GCGU--UCGAAUCUUGAUGGG-AUU-CAACUUUAGUACGAG
AGGAU-UUAGUUGGACGAAGCUC-UGGU-G-UACCAGUUGUCGCGCUAGCGGCAC-G-GCUGGAUAGCUAAC
UUCGGAAAGGAUAAACGCUGAAAGCA-UC--UAAGCGUGAAGCCCGCCCAGA-G-AUA--AGGAUUCG-UA-
--GACAU-----------------------------------------------------------------
--------------------------UA--------------------------------------------
------------------------------------------------------------------------
------------------------------------------------------------------------
------------------------------------------------------------------------
------------------------------------------------------------------------
------------------------------------------------------------------------
------------------------------------------------------------------------
----------------------------------------------------GUCG--AA-AGC------CC
CCUGGAAGACUACCAGGU-UGA-UAGGUU-GGA-CGUGUAAG-UGC-A----------------GU------
----AA-----------UGCAUUAAGCAA--A-CCAA-UACUAACG---GGCGAAA---GUUUGGCCA----
------------------------------------------------------------------------
------------------------------------------------------------------------
------------------------------------------------------------------------
-------------------------------------------------
>LSUa_272557_AERPE/1-3083
----------------------------GGC----AC--CUAUGCCGCCC-GGGGGAUG-G-CUCGGC-UCG
G-G---CGCCGAGGAAGGGCGUGG--CAAGCUGCGAUAAGCCCGGGGUAGGCGCAGGCAGCCGU------CG
AAC-CCG-GGAUCCCCGAAUGGGACU-U--------CCC-------GC----UGCG-G-GCG---------A
A-G-----------AGC---C----C----G---CAG----C-G-CCCC-C---------------------
-----------G-----------UUU----------------------------------------------
-----------------------GG-G-GG-CGGGAACCCCCCGAAAGGAAGCAUCUGAGUAGAGG--GAGG
AGAAGAAACCAA--CAGGGAUCCCCUGA--GU-AG-GGGCGACCGAAAGGGGG-A-GA-GCCCAAACCAAUC
CCCC-ACGGGA-------GGA------------C---CGU-GGGGGGAUGAGGGG--UUG------------
C-GG--CCC-UGC-G-----G------CUGGGGCUUCGAGCCCGGCGGGC--CGA------------CC--C
A-------CGGUA--GCCG------AAGUGG-GCUG------------------------------------
-------GAA-UG-CCCCG-------CCGUAGAGGGUGAUAGCCCCGUA---------GGC---G----AAA
C------CGUGGGGG-CCC-----------------GUGC-CGCAG--GGCAGAGUACCACGGCUUG---GU
AUUGCCGUGGGAA-GCUGGGAGGAACC--GACUUCCAAGGCUAAAUA--CGU-C-CCG-AGACCGAUAGC-G
AAC-UAAGUACCGUGAGGGAAAG-CUGAAA-AGAAC-CCCGG-GAGGGG-G-GUG-AAAAGAGCCUGAAACC
G-GGCGGCGACAU-AC--GG-UGCG---ACCCGAAA-GGGGUG-----------------------------
------------------------------------------------------------------------
------------------------------------------------------------------------
------------------------------------------------------------------------
------------------------------------------------------------------------

```
------------------------------------------------------------------------
------------------------------------------------------------AA-ACCCCC-G-
---AA-----------GG-A-------------------A------ACC-----------------CGG---
------------------------------------------------------------------------
--------------------------------------------------------------G--C-GA---
------------------------------------------------------------------------
-------------------------------------------------C-CG------GG--------GAG
UACGA-------------------------------------------------------------------
------------------------------------------------------------------------
------------------------------------------------------------------------
------------------------------------------------------------------------
------------------------------------------------G-----------------------
------------------------------------------------------------------------
------------------------------------------------------------------------
------------------------------------------------------------------------
------------------------------------------------------------------------
------------------------------------------------------------------------
-------------------GGGGAU-GG-----------------ACCGG--G---GUCGCACCGUCCGUCU
UGAAACACGGCCGGGGAGUUCA-C-GGCCGUGGCGA-GCUUAAGG--G-G---------GUCAA--------
C-CC--CGUA--GGCGCAGGGAAAC-CGA-----------------CAG----------------------G
CCCG-CAGCCGG----G-------------------------------------------------------
---GG-------------------------------------------------------------------
------------CGACC--CCGG------------------------------------CGA-GGGGCG-GG
GUCC-----------------------------------UAAUA-------------------------GG-
G-C-CC----------------------GGAGUCACGGCCGUGA-GACCAGAAACC-GGGCGAUCUAGGCGG
GGGCAGGGCGAAGCCGGG-GGAAAC-CCCGGUGGAGGCCCGC--AAGGGUGCUGACGUGCAAAUCGC-UCCC
CUGACCCCCGUCUAGGGGUGAAAGGCCAAUCUAGCCCGGAAGAUAGCUGGUUCCCGCCGAAG-UGGCUCGCA
GGCCAGCCCCGC-CGGA-------------------------------------------------------
----------------------GG-----CCGGGCCGUGGGGUAGAGCUACUGAUU-GGG-CGUG-CAGGGG
G-C-GUAAA-GCCC-C-CGGCGCCCAGUCAAACUCCGAACCUGCGG--------------------------
------------------------------------------------------------------------
------------------C-CG-C--CG-UAGA--UGGCGGG-AGUGGGGUU-CCCC-G-GGG-UAAGCCCG
GG-G-GCCGAGAG-GGGAAC---AACCCAGACCGG-G-GUUAAGGCCCCUAAGU--GC-CGG-CU-AA---G
-U-GCCAAG--CCAAAGGGCGUCCCCCGCCUAAGACAGCGGGGAGGUGGGCCUAACAGCAGCCA-UCCUCUA
AGGAGUGCGUAACAGCUCACCCGCC-----GAGGCGGGGGGCCCCGAAGAUUGGUC-GGGGCUUA-AGCCGG
C----------CGCCGAG-ACCCCGGGCCGCGGC--U-----------------------------------
------------------------------------------------------------------------
-----CCGAUG-------------------------------------------------------------
-------------------------------GAGCCG-CGUG-CGGUAGGCGGGC-GUCGGGG---------
-----------U-G-GCGUA-GAAGCCG--G-GCCGCGAGGU--C-CGGUGGAGCC-G-CCC-CGAGCGCGG
AUCC-CGGCGG-UAGUA-ACAGCG--AAGCG-GGGUGA--G---------------AAUCCC-CGC-CGCCG
GAAAGGGC-AAGGGUUCC-CCGGCAAUG-GUCGUCAGCCGGGGGUCAGCCGG--UC-CUAACCGGGACCGU-
AAC-ACGG---A-UCC--------------------------------------------------------
----------------------------------------CGG-GA-A-GGGGA-AACGGGUU-AACA-UUC
CCGUG----------------------------------CCGCG----GGGG--------U-AC--------
```

```
------------------GCUUC-------------------------------------------------
-------GCGG--CAAC---------GC-AA---GGC--------------------CCCA-C-CCCC-GAC
GCCUCGG-------------GGUAGG-CGGACCGGG-GG-CCCCC-UUAGG--------GGGGCGACCCGGC
UAAC-C-UGC-CG-----A-A-------------------GG-----CC----------------GGGG-AG
U-G-CC--------------------GUAAC-GGCGAGAACC---------GGCCGA----AGG-GG-GGA-
AUG--GC-----------------------------------------------------------------
-----CCGCC------UAUGGCGGU-U-CCG-C-C-GACCCCUGGACCC-UUGAAA-A-GGG-GGUGGGGAA
--------------------------------------------------------GGAUCC-CC-CGCGCC
CGUACCGAGAACCGAC------------------------------ACAGGUGC-CCCUGGGU-GAGCAGCC
CAAGGC-GUCU-GGGGGCUAACCCG-G-GCCAGGGAACUCGGC-AAAUUGGCCCCGUAACUUCGG-GAGAAG
GGGUG-CCUG-C-GG--UCCU----------GGG-GCUCGAACCCU--------------------------
------------------------------------------------------------------------
------------------------------------------------------------------------
------------------------------------------------------------------------
------------------------------------------------------------------------
------------------------------------------------------------------------
------------------------------------------------------------------------
------------------------------------------------------------------------
------------------------------------------------------------------------
------------------------------------------------------------------------
------------------------------------------------------------------------
------------------------------------------------------------------------
------------------------------------------------------------------------
------------------------------------------------------------------------
------------------------------------------------------------------GGGACC
GCA--GGUCGCAGUGCCUAGGGGGGCCUGA-----CUGUUUAACAAAAACAUAGCUCCCCGCUAGCCCG---
-AA-A-------------GGGUGUGU-ACGGGGGGUGA-GCCCUGGCCACUGGCG-G-UCCG-UGAAACCGG
GG-------------UACA-------------ACCCGGCGAAGCGC--CGCUGAAG-GCCGGGAGUAACCCU
GACUCUCUUAAGGUAGCCAAAUGCCUUGCCGGG-UAAGUUCCGGCGCGCAUGAACGGGUCAACGAGG--UCC
CCACUGUCCUGGCC----CGG-GGCCCCGUGAACCC--CCU---GAGCCGGUGCACAGUCCGGCAUCCCCCC
GCAGGGAGAGAAGA-CCCCGUGGAGC-UUUACC--GCAGCCC-GGCGUUGGCCCCCGGGCGGGGGUGCGUAG
CGU-AGGUGGGAGCCGAUG----------AGCCACC-CUC----------------------------UCCG
--------------------------------------GG-GGGUGGGGAG-G--C-GU-CCAUGAAACA-C
CACC-CACCCCCGCCCGGGGGCCUAACCC-CG--CCC--C-GG-G-U-------------------------
------------------------------------------------------------------------
------------------------------------------------------------------------
------------------------------------------------------------------------
------------------------------------------------------------------------
-------------------------------------------------------G----GG----------
------------------------------------------------CGGG-----GGACA-GCGUCGGGU
GGGCGGUUUGGCUGGGGCGGCACGCCCGCGA-AGGGGUAAC-ACGGGCGCCCA-AAGGUCGGCUCAGGCGGG
UCAGAACUCCGCCGUA-GAGUG-CAAGGGCAA-AAGCCGGCCUGACC-GGAC--------------------
--------------CCCUGAAAGCAAGGGGUCCGGCCGGGAAA-CCGUGGCCUAGCGCACGCUC--------
---GU-G-----CCCCC-UCG--------------------GU-GGGGGCCGGGCACGACAG-AA--AAGUU
ACCCCGGGG-AUAACAGAGUCGUCGCGGGCGAGAGCCCCCAUCGACCCCGCGGUUUGCUACCUCGAUGUCGG
```

CUCUUCCCAUCCUGGGGGUGCA-GCAGCCCCCAAGGGUGGGGCUGCCCGCCCAUUAAAGGGGAACGUGAGCU
GGGUUCAGACCGUCGUGAGACAGGUCGGUCUCUACCCGCG--------------------G-GGG-------
-------------------------------GUGU--GGGCCGCCUGAGGGGAAGG-UGCCCUUAGUACGAG
AGGAA-CGGGGCGCCGCGGCCUC-UGGU-G-UACCGGUUGUCC-UGGU-GGGCAA-C-GCCGGGCAGCUAAG
CCGUA-GGGGGUAACCGCUGAAAGCA-UC--UAAGCGGGAACCCCUCCCCGA-A-AAG--AGGCGGCC-GU-
--UGGGCGCCGGC-CUAGU------------GCCG-------------------------------------
------------------------------------------------------------------------
------------------------------------------------------------------------
------------------------------------------------------------------------
------------------------------------------------------------------------
------------------------------------------------------------------------
------------------------------------------------------------------------
------------------------------------------------------------------------
------------------------------------------------GCGUCCA--CG-AGG------GC
UCCCGUAGAAGACGGGGU-UGA-UGGGGC-GGG-GGUGUGAG-CCC-C--------GAG-----AGGGCUU-
AGCCCU--------CGAGGGGUGUAGCCC--G-CCGC-UCCCAAUC---GCCCGAGGC-CG-----------
------------------------------------------------------------------------
------------------------------------------------------------------------
------------------------------------------------------------------------
-------------------------------------------------
>LSUa_415426_HYPBU/1-3084
----------------------UCAC--GGC----AC--GGACGCCGCCC-GGUGGAUG-G-CUCGGC-UCG
G-G---CGCCGACGAAGGGCGCGG--CAAGCGGCGAUAUGCCCGGGGUAGGCGCAGGCAGCCGU------GG
AAC-CCG-GGAUCCCCGAAUGGGGCC-U--------CCC-------GC----CACG-G-GCG---------A
A-U-----------AGC---C----C----G---UGG----C-G-CUCG-GGA-----AACC----------
-----------G-----------UUA-----------------------------------------GGGAG
U-A--------------------CC-G-AG-UGGGAACCCCCCGAACGGAAACAUCUUAGUAGGGG--GAGG
AAAAGAAAGCAA--CAGCGAUCCCCUGA--GU-AG-GGGCGACCGAAAGGGGG-A-GA-GCCCAAACCAAUC
CUCC-GCGGGA-------UAA------------C---CGC-GGAGGGAUGAGGGG--UUG------------
C-GG--CCC-CGC-G-----G------CUGGGGCUUCGAGCCCAGCGGGC--CGA------------CC--C
C-------GGGUA--GCCG------AAGUGG-GCUG------------------------------------
-------GAA-AG-CCCCG-------CCGUAGAGGGUGACAGCCCCGUA---------GGC---U----AAA
C------CCGGGGGG-CCC-----------------GCGC-CGCGG--GGCAGAGUACCACGGCUUG---GU
UCUGCCGUGGGAA-GCUGGGAGUCACC--GACUCCCAAGGCUAAAUA--CGU-C-CCG-AGACCGAUAGC-G
AAC-U-AGUACCGUGAGGGAACG-CUGAAA-AGCAC-CCCGG-AAGGGG-G-GUG-AAAAGAGCCUGAAACC
G-GGCGGCUAAGC-AC--GG-GGCG---GCCCGAAA-GGGAGG-----------------------------
------------------------------------------------------------------------
------------------------------------------------------------------------
------------------------------------------------------------------------
------------------------------------------------------------------------
------------------------------------------------------------------------
------------------------------------------------------------AG-GCCCCCCG-
---AA-----------GG-A-------------------A------ACC-----------------GGC---
------------------------------------------------------------------------
--------------------------------------------------------------G--C-GA---

```
------------------------------------------------------------------------
-------------------------------------------------G-CC------GG--------GAG
UACGA-------------------------------------------------------------------
------------------------------------------------------------------------
------------------------------------------------------------------------
------------------------------------------------------------------------
------------------------------------------------G-----------------------
------------------------------------------------------------------------
------------------------------------------------------------------------
------------------------------------------------------------------------
------------------------------------------------------------------------
------------------------------------------------------------------------
-------------------GGGGGC-GG-----------------ACCGG--G---GUCGCCCCGUCCGUCU
UGAAACACGGCCGGGGAGUUCA-C-GGCCGUGGCGA-GCCUAAGG--G-G---------UUCAC--------
C-CC--CGAA--GGCGCAGGGAAAC-CGA-----------------CGA----------------------G
CCCG-CAGCCGG----G-------------------------------------------------------
---GG-------------------------------------------------------------------
------------U-UCC--CCGG------------------------------------CGA-GGGGCG-GG
GUCC-----------------------------------GAAA--------------------------GG-
G-C-CC----------------------GGAGUCACGGCCGUGA-GACCAGAAACC-GGGCGAUCUAGGCGG
GGGCAGGGUGAAGCCGGG-CGAAAG-CCCGGUGGAGGCCCGA--AGGGGUUCUGACGUGCAAUUCGU-UCCC
CUGACCCCCGUCUAGGGGUGAAAGGCCAAUCUAGCCCGG-UGAUAGCUGGUUCCCGCCGAAG-UGCCUCGCA
GGGCAGCCCGC-CGGA--------------------------------------------------------
----------------------GG-----CCGGGCCGCGGGGUAGAGC-ACUGAUU-GGG-GGUG-CAGGGG
G-C-GUAAU-GCCC-C-CGGCCCCCUAUCAAACUCCGAACCCGCGG--------------------------
------------------------------------------------------------------------
------------------C-CG-C--CG-UAGA--UGGCGGG-AGUGGGGCU-CCCC-G-GGG-UAAGCCCG
GG-G-GCCGAGAG-GGGAAC---AACCCAGACCGG-G-GUUAAGGCCCCCAAGG--GC-CGG-CU-AA---G
-U-GCCAA---CGAAAGGGCGUCCUCGGCCUCAGACAGCGGGGAGGUGGGCCUAACAGCAGCCA-UCCUCUA
AGGAGUGCGUAACAGCUCACCCGCC-----GAGGCCGAGGGCCCCGAAGAUAUGUC-GGGGCUUA-AGCCGG
C----------CGCCGAA-ACCCCGGGGGCGGGC--C-----------------------------------
------------------------------------------------------------------------
-----GCUAUG-------------------------------------------------------------
------------------------------CGGCCCG-CAUC-CGGUAGGCGGGC-GUCGGGG---------
-----------C-G-GCGCA-GAAGGCG--G-GCCGUGAGGC--C-CGCUGGAGCC-G-CCC-CGAGUGCGG
AUCC-CGGCGG-UAGUA-ACAGCG--AAGAG-GGGUGA--G---------------AAUCCC-CUC-CGCCG
GAAAGGGC-AAGGGUUCC-UCGGCAACG-GUCGUCGGCCGAGGGUGAGCCGG--CC-CUAACCGGGGCCGU-
AAC-ACGG---A-CCC--------------------------------------------------------
----------------------------------------CGG-GA-A-AGGGA-AACGGGUU-AAUA-UUC
CCGUG----------------------------------CCGCG----GGGGUAGG-------U--------
------------------------------------------------------------------------
-------GCGG--CAAC--------GC-AA-----GC--------------------CCCG-C-CUCC-GAC
GCCUCCG-------------GGUAGG-CCGACCGGG-GG-CGGCC-GAAGG--------GCCGCGACCCGG-
CUAA-C-CGC-UG-----A-A-------------------GG-----CC----------------GGGG-AG
U-G-CC--------------------GUAAU-GGCGAGAACC---------GGCCGA----AGG-CG-GGA-
```

```
AUG--GG-----------------------------------------------------------------
-----GGGCC---GUUAGGCCCCUU-C-GGC-C-G-AUCCCGGGGGCCC-UUGAAA-A-GGG-GGCGGGGAA
--------------------------------------------------------CGAUCC-CC-CGCGCC
CGUACCGAGAACCGAC------------------------------ACAGGUGC-CCCUGGGU-GAGAAGCC
CCAGGC-GUCU-GGGGGGCAACCCG-G-GCCAGGGAACUCGGC-AAAUUGGCCCCGUAACUUCGG-GAGAAG
GGGUG-CCCG-C-GG--UCUG----------GGG-GUAG---------------------------------
------------------------------------------------------------------------
------------------------------------------------------------------------
------------------------------------------------------------------------
------------------------------------------------------------------------
------------------------------------------------------------------------
------------------------------------------------------------------------
------------------------------------------------------------------------
------------------------------------------------------------------------
------------------------------------------------------------------------
------------------------------------------------------------------------
------------------------------------------------------------------------
------------------------------------------------------------------------
------------------------------------------------------------------------
----------------------------------------------------------ACCCCC--UGGACC
GCG--GGUCGCAGUGCCUAGGGGGGCCUGA-----CUGUUUAAUAAAAACAUAGGUCCCCGCAAGCCCG---
-AA-A-------------GGGUGUGU-ACGGGGGCCGA-AUCCUGGCCACUGGCG-G-UCCG-UGAAACCGG
GG-------------UUCA-------------ACCCGGCGAAGCGC--CGCUGAAG-GCCGGGAGUAACUCU
GACUCUCUUAAGGUAGCCAAAUGCCUUGCCGGG-UAAGUUCCGGCGCGCAUGAAUGGAUCAACGAGG--UCC
CCACUGUCCUGGCC----CGG-GGCCCCGUGAACCC--ACG---GAGCCGGUGCACAGGCCGGCAUCCCCCC
GCAGGGCGAGAAGA-CCCCGUGGAGC-UUUACC--GCAGCCU-GGGGUUGCCCUUCGGGCCUGGGUGCGUAG
CGU-AGGUGGGAGCCGAUG----------AGCCACC-CUC----------------------------UCCG
--------------------------------------GG-GGGUGGGGAG-G--C-GC-CAAUGAAACA-C
CACC-CACCCAGGCCCGGGGGGCUUACCC-CC--GGC--G-GG-A-A-------------------------
------------------------------------------------------------------------
------------------------------------------------------------------------
------------------------------------------------------------------------
------------------------------------------------------------------------
-------------------------------------------------------G----CC----------
------------------------------------------------GGGG-----GGACA-GCCCCAGGU
GGGCGGUUUGGCUGGGGCGGCACGCCCGCGA-AAAGGUAAC-ACGGGCGCCCA-AAGGUCGGCUCAGGCGGG
UCAGAGCUCCGCCGUA-GAGUG-CAAGGGCAA-AAGCCGGCCUGACC-GGAC--------------------
--------------CCCUAAAAGCAGGGGGUCCGGCCGGGAAA-CCGCGGCCUAGCGAACGCUC--------
---GU-G-----CCCCCCUCG--------------------GU-GGGGGCCGGGCAUGACAG-AA--AAGUU
ACCCCGGGG-AUAACAGAGUCGUCGCGGGCGAGAGCUCCCAUCGACCCCGCGGUUUGCUACAUCGAUGUCGG
CUCUUCCCAUCCUGGGGGUGCA-GCAGCCCCCAAGGGUGGGGCUGCCCGCCCAUUAAAGGGGAACGUGAGCU
GGGUUUAGACCGUCGUGAGACAGGUCGGACUCUACCCGCG--------------------G-GGG-------
-------------------------------GUGU--GGGCCGCCUGAGGGGAAGG-UGCCCCUAGUACGAG
AGGAA-CGGGGCGCCGCGGCCUC-UAGU-G-UACCGGUUGUCC-GGCA-GGGCAG-C-GCCGGGUAGCCACG
CCGUA-GGGGGUAACCGCUGAAAGCA-UC--UAAGCGGGAACCCCUCCCCGA-A-AAG--AGGCGGCC-GC-
```

```
--CAGCCCGGGGC-CA----------------CCC-------------------------------------
------------------------------------------------------------------------
------------------------------------------------------------------------
------------------------------------------------------------------------
------------------------------------------------------------------------
------------------------------------------------------------------------
------------------------------------------------------------------------
------------------------------------------------------------------------
-------------------------------------------------GGGCUGG--CC-CGG------GC
UCCCGUAGAAGACGGGGU-UGA-UGGGGC-GGG-GGUGUAAG-CCC-C--------GAG-----GGCGUUA-
--GGCC--------CGAGGGGUUCAGCCC--G-CCGC-UCCCAAUC---GCCCGGA----------------
------------------------------------------------------------------------
------------------------------------------------------------------------
------------------------------------------------------------------------
-------------------------------------------------
>LSUa_399550_STAMF/1-3092
------------------------------------C--CGACGCCGCCC-GGUGGAUG-G-CUCGGC-UCG
G-G---CGCCGAGGAAGGCCGUGG--CAAGCUGCGAUAAGCCCGGGGUAGGCGCAUGCAGCCGU------UG
AAC-CCG-GGAUCGCCGAAUGGGACC-U--------CCU-------GC----CACG-G-GCG---------A
A-G-----------AGC---C----C----G---UGG----C-G-UCCG-GGA-----AACCG---------
-----------G-----------GAU----------------------------------------GGGA-G
U-A--------------------CC-G-GA-CGGGAACCCCCCGAACGGAAACAUCUUAGUAGGGG--GAGG
AGGAGAAACCACC-ACGGGAUCCCCUGA--GU-AG-GGGCGACCGAAAGGGGG-A-GA-GCCCAAACCAAUC
CUCC-ACGGGA-------UAA------------C---CGU-GGAGGGAUGAGGGG--UUG------------
A-UG--GGU-CUC-C-----G------GC-GGGGCUUGAGCCCCAGAGGGC--CUA-----------UC--C
A-------CGGUA--GCCG------AAGUGG-CCUG------------------------------------
-------GAA-UG-GCCCG-------CCGUAGAGGGUGAAAGCCCCGUA---------GGC---U----AAA
C------CGUGGAGG-CCC-----------------UCGC-CGGAG--ACCAGAGUACCACGGCUUG---GU
UUUGCCGUGGGAA-GCCGGGAGGCACC--GGCUUCCAAGGCUAAAUA--CGU-C-CCG-AGACCGAUAGC-G
UAC-UAAGUACCGUGAGGGAAAG-CUGAAA-AGUAC-CCUGG-AAAGGG-G-GUG-AAAAGAGCCUGAAACC
G-GGCGGCUACAU-AC--GG-CGCA---GCCUGGAA-GAGAUG-----------------------------
------------------------------------------------------------------------
------------------------------------------------------------------------
------------------------------------------------------------------------
------------------------------------------------------------------------
------------------------------------------------------------------------
------------------------------------------------------------AG-GUCCUCCA-
---AA-----------GG-A-------------------A------ACC-----------------GGG---
------------------------------------------------------------------------
--------------------------------------------------------------G--C-GA---
------------------------------------------------------------------------
-------------------------------------------------C-CC------GG--------GAG
UACGA-------------------------------------------------------------------
------------------------------------------------------------------------
------------------------------------------------------------------------
```

```
------------------------------------------------------------------------
------------------------------------------------G-----------------------
------------------------------------------------------------------------
------------------------------------------------------------------------
------------------------------------------------------------------------
------------------------------------------------------------------------
------------------------------------------------------------------------
-------------------GAGGAC-UG-----------------AUCAG--G---GUUGCGCCGUCCGUCU
UGAAACACGGCCGGGGAGUCCA-C-GGCAGUGGCGA-GCCUAAGG--G-GG--------ACAAA-------G
C-CC--CGUA--GGCGUAGGGAAAC-CGA-----------------CAA----------------------G
CCCG-CAACCGG----G-------------------------------------------------------
---CG-------------------------------------------------------------------
--------------CAA--GGCG------------------------------------UGA-GGGGCG-GG
GUCC-----------------------------------UCAUA-------------------------GG-
G-C-CC----------------------GGAGUCACUGCCGUGG-GACCAGAAACC-GGGCGAUCUAGGCGG
GGGCAGGGUGAAGCCGGG-GGAAAC-CCCGGUGGAGGCCCGC--AAGGGUUCUGACGUGCAAUUCGU-UCCC
CUGACCCCCGUCUAGGGGCAAAAGACCAAUCUAGCCCGG-UGAUAGCUGGUUCCCGCCGAAG-UGGCUCUCA
GGCCAGCCCCGC-CAGA-------------------------------------------------------
----------------------GG-----CU-GGCUACGGGGUAGAGCUACGGAUU-GGG-GGUG-CAGGGG
G-C-GA-AA-GCCC-C-CGGCCCCCUAUCCAACUCCGAACCCGUGG--------------------------
------------------------------------------------------------------------
------------------C-CG-C--CG-CAGA--AGGCGGG-AGUGGGGCU-CCCC-G-GGG-UAAGCCCG
GG-G-GCCGAGAG-GGGAAC---AACCCAGACCGG-G-GUUAAGGCCCCAAAGU--GC-CGG-CU-AA---G
-U-GCCAAGC-CCAAAGGGCGUCCCGCGGCUUAGACAGCGGGGAGGUGGGCCUAACAGCAGCCA-UCCUCUA
AGGAGUGCGUAACAGCUCACCCGCC-----GAGCCGCGGGGCCCCGAAGAUUGGUC-GGGGCUCA-AGCCGG
C----------CGCCGAG-ACCCCGGGGCACGGC--U-----------------------------------
------------------------------------------------------------------------
-----CCGAUG-------------------------------------------------------------
------------------------------GAGCCGU-GAUC-CGGUAGGCGGGC-GCCGGGU----------
-----------C-G-GGGCA-GAAGCCG--G-GCCGAGAGGU--C-CGGUGGACCC-G-AUC-CGGGUGUGG
AUCC-CGGCGG-CAGUA-GCAGCG--AAGAG-GGGUGG--A---------------AAUCCC-CUC-CGCCG
GAAAGGGC-CAGGGUUCC-UCGGCAACG-GUCGUCGGCCGAGGGUUAGCCGG--CC-CUAACCCGGGCCGU-
AAC-ACGG---A-CCC--------------------------------------------------------
----------------------------------------GGG-GA-A-GGGGA-AACGGGUU-AAUA-UUC
CCGUG----------------------------------CCGCG----GGGGUAGA-------C--------
------------------------------------------------------------------------
-------GCGG--CAAC--------GC-AA-----GC--------------------CCGG-C-CUCC-GAC
GCCUCCG-------------GAUAGG-CGGACCGGG-GG-CACCC-GAAUG--------GGUGCGACCCGG-
CUAA-C-CGG-UG------A-A-------------------GG-----CC----------------CUGG-AG
A-A-CC--------------------GUAAU-GGUGAGAAGG---------GGCCGA----AGC-CG-GGA-
AUG--GG-----------------------------------------------------------------
-----ACCCC---GUAUGGGGUCUU-C-CGC-U-G-ACUCCGGGGGCCC-AUGAAA-A-GGG-GGCCGGGAA
--------------------------------------------------------CGAUCC-CC-CGCGUC
CGUACCGAGAACCGAC------------------------------ACAGGUGC-CCCUGGGU-GAGUAGCC
CCAGGC-GUCU-GGGGGAUAACCCG-G-GCCAGGGAACUCGGC-AAAUUGGCCCCGUAACUUCGG-GAGAAG
```

```
GGGUG-CCCG-C-GG--UCUG---G----AGGCA-A------------------------------------
------------------------------------------------------------------------
------------------------------------------------------------------------
------------------------------------------------------------------------
------------------------------------------------------------------------
------------------------------------------------------------------------
------------------------------------------------------------------------
------------------------------------------------------------------------
------------------------------------------------------------------------
------------------------------------------------------------------------
------------------------------------------------------------------------
------------------------------------------------------------------------
------------------------------------------------------------------------
------------------------------------------------------------------------
----------------------------------------------------------AAACCC--UGGACC
GCG--GGUCGCAGUGCCUAGGGGGGCCUGA-----CUGUUUAAUAAAAACAUAGGUCCCCGCAAGCCCG---
-AA-A-------------GGGUGUGU-ACGGGGGCUGA-AUCCUGGCCACUGGCG-G-UCCG-UGAAACCGG
GG-------------UACA-------------ACCCGGCGAAGCGC--CGCUGAAG-GCCGGGAGUAACUCU
GACUCUCUUAAGGUAGCCAAAUGCCUUGCCGGG-UGAGUUCCGGCGCGCAUGAAUGGAUCAACGAGG--UCC
CCACUGUCCUGGCC----CGG-GGCCCCGCGAAUAC--ACG---GAGCCGGUGCACAGGCCGGCAACCCCCC
GCAGGGCGAUAAGA-CCCCGUGGAGC-UUCACC--GCAGCCU-GGCGUUGGCCUCCGGGCUGCGGCGCGUAG
CGU-AGGUGGGAGCCAGUG----------AACCCGC-CCC----------------------------UCCG
--------------------------------------GG-GGCGGGGGAG-G--C-GC-CAAUGAAACA-C
CACC-CACCGCAGCCCGGAGGCCUUACCC-CC--GGG--G-AG-A-U-------------------------
------------------------------------------------------------------------
------------------------------------------------------------------------
------------------------------------------------------------------------
------------------------------------------------------------------------
-------------------------------------------------------C----CC----------
------------------------------------------------GGGG-----GGACA-GCGUCAGGU
GGGCGGUUUGGCUGGGGCGGCACGCCCGCGA-AAAGGUAAC-ACGGGCGCCCA-AAGGUCGGCUCAGGCGGG
UCAGAGCUCCGCCGUA-GAGUG-CAAGGGCAA-AAGCCGGCUUGACC-GGAC--------------------
--------------CCCGGAACACAACGGGUCCGGCCGGGAAA-CCGCGGCCUAGCGAACGCUC--------
---GU-G-----CCCCCCUCG--------------------GU-GGGGGCCGGGCAUGACAG-AA--AAGUU
ACCCCGGGG-AUAACAGAGUCGUCGCGGGCGAGAGCUCCCAUCGACCCCGCGGUUUGCUACAUCGAUGUCGG
CUCUUCCCAUCCUGGGAGUGCA-GCAGCUCCCAAGGGUGGGGCUGCCCGCCCAUUAAAGGGGAACGUGAGCU
GGGUUUAGACCGUCGUGAGACA-GUCGGACUCUACCCGCG--------------------G-GGG-------
-------------------------------GUGU--GGGCCGCCUGAGGGGAAGG-UGCCCUCAGUACGAG
AGGAA-CGGGGCGCCGCGGCCAA-UGGU-G-UACCGGUUGUCC-GGCA-GGGCAU-A-GCCGGGCAGCCACG
CCGUA-GGGGGUAACCGCUGAAAGCA-UC--UAAGCGGGAACCCCUCCCCGA-A-AAG--AGGCGGCC-GU-
--CCGCCGAAG-------------------------------------------------------------
-------------------------------------------G----------------------------
------------------------------------------------------------------------
----------------------GGUUU-C-------------------------------------------
------------------------------------------------------------------------
```

```
------------------------------------------------------------------------
------------------------------------------------------------------------
------------------------------------------------------------------------
---------------------------------------------CU-CUUCGGCG--GG-AGG------GC
UCCCGUAGAAGACGGGGU-UGA-UGGGGC-GGG-GGUGUAAG-CCC-C--------GAG-----GGAGCAUU
GUCUCC--------CAAGGGGUUUAGCCC--G-CCGC-UCCCAAAA---GCCCGACGC-CGCA---------
------------------------------------------------------------------------
------------------------------------------------------------------------
------------------------------------------------------------------------
-------------------------------------------------
>LSUa_453591_IGNH4/1-3026
------------------------------C----GC--GUAAGCCGCCC-GGUGGAUG-G-CUCGGC-UCG
G-G---CGCCGACGAAGGGCGUGG--CAAGCUGCGAUAAGCCCGGGGGAGGCGCACGCAGCCGU------CG
AAC-CCG-GGAUCCCCGAAUGGGACU-U--------CCU-------GC----CGGG-G-GCG---------A
A-C-----------AGC---C----C----C---CGG----C-G-CGGG-G---------------------
-----------A-----------GAA----------------------------------------------
------------------------C-C-CG-CGGGAACCCCCCGAACGGAAACAUCUUAGUAGGGG--GAGG
AAGAGAAACCACC-ACGGGAUCCCCCGA--GU-AG-GGGCGACCGAAAGGGGG-G-UA-GCCCAAACCAAUU
CCCC-ACGGGA-------UAA------------C---CGU-GGGGAGAUGAGGGG--UGG------------
A-UG--CCC-CGC-G-----G------CU--CCCCCUCGAGGGGAGGGGCC--CGA-----------CC--C
U-------CGGUA--CCCG------AACUCC-GGUG------------------------------------
-------GAA--C-CGGGG-------CCGUAGAGGGUGACAGCCCCGUA---------GGG---G----AAA
C------CGAGGGG-GCC------------------CUGC-CGCGG--GGAUGAGUACCACGGCUUG---GU
AUUGCCGUGGGAA-GCUGGGUGGCACC--AGCAUCCAAGGCUAAACA--CGU-C-CCG-AGACCGAUAGC-G
AAC-UAGGUACCGUGAGGGAAAG-CUGAAA-AGCAC-CCCUG-GCGGGG-G-GUG-AAAAGAGCCUGAAACC
G-GGCGGUCACAC-AG--GG-UGCG---GCCCGCAA-GGGAUG-----------------------------
------------------------------------------------------------------------
------------------------------------------------------------------------
------------------------------------------------------------------------
------------------------------------------------------------------------
------------------------------------------------------------------------
------------------------------------------------------------AA-GCC-CCGG-
---AA-----------GG-A-------------------A------ACC-----------------CGG---
------------------------------------------------------------------------
--------------------------------------------------------------G--C-GA---
------------------------------------------------------------------------
-------------------------------------------------C-CG------GG--------GAG
UACGU-------------------------------------------------------------------
------------------------------------------------------------------------
------------------------------------------------------------------------
------------------------------------------------------------------------
------------------------------------------------G-----------------------
------------------------------------------------------------------------
------------------------------------------------------------------------
------------------------------------------------------------------------
```

------------------------------------------------------------------------
------------------------------------------------------------------------
-------------------GGU-GC-AG-----------------ACCGG--G---GUCGCACCUUUCGUCU
UGAAACACGGGCCGGGAGUGCA-C-GGCCGUGGCGA-GGCUAAGG--G-G---------UUCAC--------
C-CC--CGUA--ACCGCAGGGAAAC-CGA-----------------CGA----------------------G
CCCG-CAGGGGC----U-------------------------------------------------------
------------------------------------------------------------------------
--------------CAU--GCCC------------------------------------CGA-GGGGCG-GG
GUCC-----------------------------------GCAA--------------------------GG-
G-C-CC----------------------GGAGUCACGGUCGUGC-GACCCGAAACC-GGGCGAUCUAGGCGG
GGGCAGGGUGAAGCCGGG-CGAAAG-CCCGGUGGAGGCCCGC--AAGGGUUCUGACGUGCAAUUCGU-UCCC
CUGACCUCCGCCUAGGGGUGAAAGGCCAAUCUAGCUCGG-UGAUAGCUGGUUCCCGCCGAAG-UGGGUCUAA
GCCCAGCGUCCC-CGGA-------------------------------------------------------
----------------------GG-----CG-GGCCACGGGGUAGAGCUACUGAUG-GGA-GGUG-CAGGGG
GGC-GU-AG-GCCC-C-CGGCCUCCCGUCAAACUCCGAACCCGUGG--------------------------
------------------------------------------------------------------------
------------------C-CG-C--CG-UAGA--AGGGGGC-AGUGGGGCG-CCCC-G-GGG-UAAGCCCG
GG-G-GCCGAGAG-GGGAAC---AACCCAGACCGG-G-GUUAAGGCCCCCAAGU--GC-CGG-CU-AA---G
-U-GCCAAC--CGAAAGGGCGUCUCGGGCCUCAGACAGCGGGGAGGUGGGCCUAACAGCAGCCA-CCCUCUA
AGGAGUGCGUAACAGCUCACCCGCC-----GAGGCCCGAGGCCCCGAAGAUUGGUC-GGGGCUUA-AGCCGG
C----------CGCCGAG-ACCCCGGGCCCCCCU--C-----------------------------------
------------------------------------------------------------------------
-----CCGGUG-------------------------------------------------------------
-------------------------------GGAGGG-GGUG-CGGUAGGCGGGC-GUCGGGG---------
-----------C-G-GCCCA-GAAGGCG--G-GCCGUGAGGU--C-CGCUGGAGCC-G-CUC-CGAGUGCCG
AUCC-CGGCGG-UAGUA-ACAGCG--AAGAG-GGGUGA--G---------------AAUCCC-CUC-CGCCG
GAAAGGGC-AAGGGUUCC-CCGGCAACU-GUCAUAGGCCGGGGGUGAGUCGG--UC-CUAACCCGCCUCCC-
AAC-UGGA---G-CGC--------------------------------------------------------
----------------------------------------GGG-GA-A-AGGGA-AGCCGGUU-AAUA-UUC
CGGCA----------------------------------CCGCG----GGGGUAGG-------U--------
------------------------------------------------------------------------
-------GCGG--CAAC--------GC-AA-----GC--------------------CCCA-C-CUCC-GAC
GCCUCGG-------------GGUAGG-CGGACCGGG-GC-------UGUC-------------G-CCCCGG-
CUAA-C-C-C-CG-----A-A-------------------GG-----CC----------------CUGG-AG
G-G-CC--------------------GUAAC-GGCGAGAAGG---------GGCCGA----AGG-CG-GGA-
AGG--GG-----------------------------------------------------------------
-----CCCCC---GUAUGGGGGCUU-C-CGC-C-G-AUCCCUGGGGCCC-AUGAAA-A-GGG-GGUGGGGAA
--------------------------------------------------------CGAUCC-CC-CGCGCC
CGUACCGAGAACCGAC------------------------------GCAGGUGC-CCCUGGGU-GAGAAGCC
CAAGGC-GGCU-GGGGGUUAACCCG-G-GCCAGGGAACUCGGC-AAAUUGGCCCCGUAACUUCGG-GAGAAG
GGGUG-CCUG-C-GG--UCCU----------GGG-GCCC---------------------------------
------------------------------------------------------------------------
------------------------------------------------------------------------
------------------------------------------------------------------------
------------------------------------------------------------------------

```
------------------------------------------------------------------------
------------------------------------------------------------------------
------------------------------------------------------------------------
------------------------------------------------------------------------
------------------------------------------------------------------------
------------------------------------------------------------------------
------------------------------------------------------------------------
------------------------------------------------------------------------
------------------------------------------------------------------------
----------------------------------------------------------ACCCCU--GGGACC
GCA--GGUUGCAGUGCCUAGGGGGGCCUGA-----CUGUUUAAUAAAAACAUAGGUCCCCGCAAGCCCG---
-AA-A-------------GGGUGUGU-ACGGGGGCCGA-AUCCUGGCCACUGGCG-G-UCCG-UGAAACCGG
GG-------------UUCA-------------ACCCGGCGAAGCGC--CGCUGAAG-GCCGGGAGUAACUCU
GACUCUCUUAAGGUAGCCAAAUGCCUUGCCGGG-UAAGUUCCGGCGCGCAUGAAUGGAUCAACGAGG--UCC
CCACUGUCCCGGCC----CGG-GGCCCCGUGAACCC--CCU---GAGCCGGUGCACAGUCCGGCAACCCCCC
GUAGGGAGAGAAGA-CCCCGUGGAGC-UUUACU--GCAGCCC-GGCGUUGCCCCCCGGGUGGGGGUGCGUAG
CGU-AGGUGGGAGCCGGUG----------AACCGGC-CCC----------------------------UCCG
--------------------------------------GG-GGUCGGGGAG-G--C-GC-CCAUGAAACA-C
CACC-CACCCUCACCCGGGGGGCUUACCC-CG--GG---G-GA-A-G-------------------------
------------------------------------------------------------------------
------------------------------------------------------------------------
------------------------------------------------------------------------
------------------------------------------------------------------------
------------------------------------------------------------CC----------
------------------------------------------------CGGG-----GGACA-GCGCCGGGU
GGGCAGUUUGGCUGGGGCGGCACGCCCGCGA-GAGGGUAAC-ACGGGCGCCCU-AAGGUCGGCUCAGGCGGG
UCAGAAACCCGCCGUA-GAGUG-CAAGGGCAA-AAGCCGGCCUGACC-GGAC--------------------
--------------CCCUAAAGGCAGGGGGUCCGGCCGGGAAA-CCGCGGCCUAGCGAACGCUC--------
---GU-G-----UCCCCCCUG--------------------GU-GGGGGCCGGGCAUGACAG-AA--AAGUU
ACCCCGGGG-AUAACAGAGUCGUCGCGGGCGAGAGCCCACAUCGACCCCGCGGUUUGCUACCUCGAUGUCGG
CUUUUCCCAUCCUGGGGGUGCA-GCAGCCCCCAAGGGUGGGGCUGCCCGCCCAUUAAAGGGGAACGUGAGCU
GGGUUUAGACCGUCGUGAGACAGGUCGGACUCUACCCACG--------------------G-GGG-------
-------------------------------GUGU--CGGCCGCCUGAGGGGAAGG-UGCCCUCAGUACGAG
AGGAA-CGGGGCGCCCCGGCCUC-UGGU-G-UACCGGUUGUCC-GGCA-GGGCAG-A-GCCGGGCAGCUACG
CCGGA-AGGGGUAACCGCUGAAAGCA-UC--UAAGCGGGAACCCCACCCCGA-A-AAG--AGGCGGCC-AU-
------------------------------------------------------------------------
------------------------------------------------------------------------
------------------------------------------------------------------------
------------------------------------------------------------------------
------------------------------------------------------------------------
------------------------------------------------------------------------
------------------------------------------------------------------------
------------------------------------------------------------------------
----------------------------------------------------------UG-AGG------GC
UCCCGUAGAAGACGGGGU-UGA-UGGGGC-GGG-GGUGUAAG-CCC-C--------GAG-----GGGU----
```

```
--UUCC--------CGAGGGGCG-AGCCC--G-CCGC-UCCCAAUC---GCCCGAGGC-CGUUAGC------
------------------------------------------------------------------------
------------------------------------------------------------------------
------------------------------------------------------------------------
-------------------------------------------------
>LSUa_410359_PYRCJ/1-2963
----------------------CUCC-GCAC----GG--CAAAGCCGCCC-GGUGGAUG-G-CUCGGC-UCG
G-G---CGCCGAAGAAGGGCGUGG--CAAGCUGCGAUAAGCCCGGGGUAGCCGCAGGCAGGCUU------AG
AAC-CCG-GGAUCCCCGAAUGGGGCU-U--------CCU-------GC----CGGG-G-CCG---------A
A-U-----------AGG---C----C----C---CGG----C-G-CCCC-----------------------
-----------G-----------UAA----------------------------------------------
------------------------G-G-GG-CGGGAACGCGGGGAAAGGAAACAUCUUAGUACCCG--CAGG
AAGGGAAGCCAA--CAGGGACCCCCUGA--GU-AG-GGGCGACCGAAAGGGGG-A-UA-GCCCAAACCAAAU
CCUC-GCGGGA-------CAA------------C---CGU-GGGGAGAUGUGGGG--CUU------------
G-GG--CCC-GGG-C-----A-------------------------ACCGC--CGG-----------CG--G
G-------CGGUA--GCCG------AAGUGG-GCUG------------------------------------
-------GAA-UG-CCCCG-------CCGUAGAGGGUGAUAGCCCCGUA---------GGC---G----AAA
C------CGCCCGUG-GCGGAG--------------UCCC-GGGGU--CCCGGAGUACCUCGGCUUA---GU
UUUGCCGGGGGAA-CGCGCCGGCCACU--GGCCGGCAAGGCUAAGCA--CGU-C-CCG-AGUCCGAUAGC-G
CAC-U-AGUACCGUGAGGGAAAG-CUGAAA-AGAAC-CCCGG-AAGGGG-G-GUG-AAAAGAGCCUGAAACC
G-GGCGGCUACAG-UG--GG-GCAG---GCCCGAAA-GGAUG------------------------------
------------------------------------------------------------------------
------------------------------------------------------------------------
------------------------------------------------------------------------
------------------------------------------------------------------------
------------------------------------------------------------------------
------------------------------------------------------------CC-CCCUCCCG-
---AA-----------GG-A-------------------A------ACC-----------------CCG---
------------------------------------------------------------------------
--------------------------------------------------------------G--U-GA---
------------------------------------------------------------------------
-------------------------------------------------C-GG------GG--------GAG
UACGA-------------------------------------------------------------------
------------------------------------------------------------------------
------------------------------------------------------------------------
------------------------------------------------------------------------
------------------------------------------------G-----------------------
------------------------------------------------------------------------
------------------------------------------------------------------------
------------------------------------------------------------------------
------------------------------------------------------------------------
------------------------------------------------------------------------
-------------------GGAGGG-GG-----------------UCCAG--G---GUCUGCCCUUACGUCU
AGAAACACGGGCCGGGGAGUUCA-C-GGCCGUGGCGA-GCCUAAGG--G-G--------UUCAA--------
C-CC--CGGA--GGCGUAGGGAAAC-CGA-----------------CA-----------------------G
```

```
CCCG-CAGCGGG----G-------------------------------------------------------
------------------------------------------------------------------------
--------------CAA--CCCG------------------------------------CGA-GGGGCG-GG
GUCU-----------------------------------UAAA--------------------------GG-
G-C-CC----------------------GUAGUCACGGCCGUGA-GACCAGAAACC-GGGCGAUCUAGCCCU
GGGCAGGGUGAAGCGGGG-CGAAAG-CCCCGUGGAGGCCCGA--AGGGGUUCUGAUGUGCAAAUCGU-UCCC
AUGACCUGGGGCUAGGGGCAAAAGACCAAUCAAGCCCGG-UGAUAGCUGGUUCCCCCCGAAG-CGGGUCUCA
GCCCGGCCUCCC-CGGA-------------------------------------------------------
----------------------GG-----CG-GCCGGCGGGGUAGAGU-ACUGAUC-GGG-GGUG-CGGGAG
C-C-GA-AA-GGCU-C-CGGCCCCCGGUCAAACUCCGAACCUGCCA--------------------------
------------------------------------------------------------------------
------------------G-CG-C--CG-UAGA--AGGGGGG-AGGCGGGGG-CGGU-G-GGG-UAAGCCUC
CG-C-UCCGAGAC-GGGAAC---AACCGAGACCGG-G-GUUAAGGCCCCCAAGU--GC-GGG-CU-UA---G
-U-GUCAAUC-UAAAAGGGCGUCCCCCGCCCAAGACAGCGGGGCCGUGGGCCUAACAGCAGCCA-UCGGCUA
AGCAACGCGUAACAGCGGACCCGCC-----GAGGCGGGGGGCCCCGAAGAU-GUACAGGGACU-A-AGCCCG
C----------CGCCGAG-ACCCCGGCCCGCGGG--C-----------------------------------
------------------------------------------------------------------------
-----C-GUUG-------------------------------------------------------------
--------------------------------GCCCG-CGUG-GGGUAGGGGGGC-GCGGCCG---------
-----------U-G-GGGCA-GAAGCCG--G-GCCGUGAGGU--C-CGGUGGACCC-G-CGG-CCGACGAAG
AUCC-CGGCGG-UAGUA-GCAGCG--AAGAG-GGGUGA--G---------------AAGCCC-CUC-CGCCG
GAAAGGAC-CAGGGUUUC-CUGGCAACU-UCAAUAGGCCAGGAGUUAGCCGG--UC-CUAAGGCGGGGCCU-
AAU-AGGC---A-CCC--------------------------------------------------------
----------------------------------------GCC-GA-A-AGGGA-AACGGGUU-AAUA-UUC
CCGUG----------------------------------CCGCG----GGGGUAGGU---U-CU--------
------------------------------------------------------------------------
-------GCGG--CAAC--------GC-AG-----GC--------------------CCCG-U-CCCC-GAC
GCCUCGG-------------GAUAGG-GCGGGCGGG-AC-------UGCC-------------G-UCCCGC-
UUAA-C-CGU-CG-----A-A-------------------GG-----CC----------------GGGG-AG
U-G-CC--------------------GUAAU-GGCGAGAACC---------GGCCGA----AGG-CG-GGA-
AUA--GC-----------------------------------------------------------------
-----CGGGG---GUUUCCCCGGUC-C-GCC-C-G-ACUCCUGGGGCCC-GUGAAA-A-GGG-GACGGGGAA
--------------------------------------------------------CGAGCC-CC-CGCGCC
CGUACCGAGAACCGAC------------------------------GCAGGUGC-UCCUGGGU-GAGAAGCC
CAAGGC-GGCU-CGGG-UGACCCCG-G-GCCAGGGAACUCGGC-AAAUUGGCCCCGUAACUUCGG-GAGAGG
GCGCG-GCGG-A-GG--AUUU----------AAU-U------------------------------------
------------------------------------------------------------------------
------------------------------------------------------------------------
------------------------------------------------------------------------
------------------------------------------------------------------------
------------------------------------------------------------------------
------------------------------------------------------------------------
------------------------------------------------------------------------
------------------------------------------------------------------------
------------------------------------------------------------------------
```

```
------------------------------------------------------------------------
------------------------------------------------------------------------
------------------------------------------------------------------------
------------------------------------------------------------------------
----------------------------------------------------------UAUUCC--CGAGGG
GCA--GGUCGCAGUGGCAAGGGGGACCUGA-----CUGUUUAACAAAAACAUAGGUCCCCGCGAGCCCG---
-UA-A-------------GGGUGUGU-ACGGGGGCUGA-AUCCUGGCCACUGGCG-G-UACG-UGAACCCCG
GG-------------UACA-------------ACCGGGCGAAGCGC--CGCUGAAG-GCCGGGGGUAACUCU
GACCCUCUUAAGGUAGCCAAAUGCCUUGCCGGG-UAAGGGCCGGCGUGCAUGAAUGGAUCAACGAGG--UCC
CCACUGUCCCGGCC----CGG-GGCCCGGCGAACCC---AC---CUCCAGGUGCACAGUCCUGGGACCCCCG
ACGGGGCGAGAAGU-CCCUAUGGAGC-UUCACA--GCAGCCU-GUCGUUGCGGGGGGGCGGGGGGUGCAGAG
CGU-AGGUGGGAGCGAU-G-----------AAACGGG-GUC----------------------------UCCG
--------------------------------------GG-CCCCGUGGAU-G--C-GA-CCCUGGAACA-C
CACC-CACUCUCCGCCCCUCCGCUUACCC-GC--C------GC-A-A-------------------------
------------------------------------------------------------------------
------------------------------------------------------------------------
------------------------------------------------------------------------
------------------------------------------------------------------------
-------------------------------------------------------------G----------
------------------------------------------------GCGG-----GGACA-GCGGCAGGC
GGGCUGUUCGGCUGGGGCGGCACACCCCUGA-AAAGAUAUC-GGGGGUGCCCA-AAGCUCGGCUCAGGCGGG
UCAGAAAUCCGCCGUA-GAGUG-UAAGGGCAA-AAGCCGGGCUGACU-GGGC--------------------
--------------CCUUGAACGCAAGGGGCCCAGGCGGGAAA-CCGGGGCCUAGAGAACGCUC--------
---GU-G-----CCCCCACCA--------------------GU-GGGGGCCGGGCAUGACAG-AA--AAGUU
ACCCUAGGA-AUAACCGGCUCGUCGCGGGUGAGAGUCCCCAUCGACCCCGCGGUUUGGUACCCAGACGUCGU
CUCUUCCCAUCCUGGCGGUGCA-GCAGCCGCCAAGGGUGGGGCUGCCCGCCCAUUAAAGGGGAACGUGAGAU
GGGUUCAGACCGUCGCGAGACAGGUCGGUCUCUACCUGUC--------------------G-GGG-------
-------------------------------GCGC--UGGCCGCCUGAGGGGAAGG-UGCCCUCAGUACGAG
AGGAA-CGGGGCGCCGCGGCCUC-UAGU-G-UACCGGUUGUCC-GGCA-GGGCAC-U-GCCGGGCAGCCACG
CCGUG-GGGGAUAACCGCUGAAAGCA-UC--UAAGCGGGAAG------------------------------
------------------------------------------------------------------------
------------------------------------------------------------------------
------------------------------------------------------------------------
------------------------------------------------------------------------
------------------------------------------------------------------------
------------------------------------------------------------------------
------------------------------------------------------------------------
------------------------------------------------------------------------
------------------------------------------------------------------------
-CCCGUAGAAGACGGGGU-UGA---GGGG-GGC-GGUGUAAC-CCC-C--------GAG-----GGU-----
--UUCC--------CGAGGGGAG-AGCCG--G-CCCC-UCCCAUC---GCCCGA------------------
------------------------------------------------------------------------
------------------------------------------------------------------------
------------------------------------------------------------------------
-------------------------------------------------
```

```
>LSUa_397948_CALMQ/1-3030
----------------------ACCC-GUGC----CG--CGAAGCCGUCC-GGUGGAUG-G-CUCGGC-UCG
G-G---CGCCGAGGAAGGGCGUGG--CAAGCGGCGAUAUGCCCGGGGUAGCCGCAGGCAGGCGU------UG
AUC-CCG-GGAUCCCCGAAUGGGGCU-U--------CC--------UA----CCUG-G-GCCG--------A
A-U----------AGGC---U----C----A---GGU----G---CCCC-----------------------
-----------G-----------UAG----------------------------------------------
------------------------G-G-GG-UGGGAACGCGGGGAACGGAAACAUCUUAGUACCCG--CAGG
AGGGGAAACCAA--AAGGGACCCCCUGA--GU-AG-GGGCGACCGAAAGGGGG-U-UA-GCCCAAACCGAAC
CCCC-AUGGGG-------UAA------------C---CGU-GGGGGGAUGUGGUG--UUU------------
G-GG--CUU-GGG-U-----A--------------------------CCGC--CGA-----------GG--C
A-------GCCUA--GCCG------AAGUGG-GCUG------------------------------------
-------GAA-AG-CCCCG-------CCGUAGAGGGUGAUAGCCCCGUA---------GGC---U----AAA
G------GUUGCUGG-CGG-AU--------------UCCC-AAUGU--CCCGGAGUACCACGGCUUG---GU
UUUGCCGUGGGAA-UACGGGGGCCUCC--AGCCCCUAAGGCUAAGUA--CGU-C-CCG-AGUCCGAUAGC-G
AAC-A-AGUACCGUGAGGGAAAG-CUGAAA-AGAAC-CCCGC-GAGGGG-U-GUG-AAAAGAGCCUGAAACC
G-GACGGUUAAAG-GU--GC-GGCG---GCCCUCAA-GGAUA------------------------------
------------------------------------------------------------------------
------------------------------------------------------------------------
------------------------------------------------------------------------
------------------------------------------------------------------------
------------------------------------------------------------------------
------------------------------------------------------------AC-CCCACCUG-
---AA-----------GG-A-------------------A------ACC-----------------GGG---
------------------------------------------------------------------------
--------------------------------------------------------------G--U-GA---
------------------------------------------------------------------------
-------------------------------------------------C-CC------GG--------GAG
UACGA-------------------------------------------------------------------
------------------------------------------------------------------------
------------------------------------------------------------------------
------------------------------------------------------------------------
------------------------------------------------G-----------------------
------------------------------------------------------------------------
------------------------------------------------------------------------
------------------------------------------------------------------------
------------------------------------------------------------------------
------------------------------------------------------------------------
-------------------GGUGGG--G-----------------UACGG--G---GUCGCCGCUUACGUCC
AGAAACACGAGCCAGGGAGUUCC-C-AUCUGUGGCGA-GUCUAGG--G-G---------GUUAA--------
C-CC--CGGA--GGCGUAGGGAAAC-CGA-----------------AG-----------------------G
CCCG-CAGUCGG----G-------------------------------------------------------
---U--------------------------------------------------------------------
--------------UAA--CCGA------------------------------------CGA-GGGGCU-GG
GUCC-----------------------------------GUUAA-------------------------GG-
G-C-CC----------------------GUAGCCACAGGUGGGA-GACCAGAAACC-GAGCGAUCUAGCCCG
```

```
GGGUAGGGUGAAGCGGGG-CGAAAG-CCCGUGGAGGCCCGA-UUGGGGUUCUGACGUGCAAAUCGU-UCCC
GUGACCCUGGGCUAGGGGCAAAAGACCAAUCAAGCUCGG-UGAUAGCUGGUUCCCCCGAAG-CAGCUCUUA
GGCUGGCUCCC-UGGA--------------------------------------------------------
----------------------GG-----GU-GCUCAUGGGGUAGAGC-ACUGAUC-CGG-GAUG-CAGGGG
C-C-GA-AA-GGCU-C-CGGUCCCGGGUCAAACUCCGAACCCAUGA--------------------------
------------------------------------------------------------------------
------------------G-CC-C--CA-UUGA--AGGGGGG-AGACGGGUU-CCCCGG-GGG-UAAGCUUG
GG-G-GCCGAGAG-GGAAAC---AACCCAGACCGG-G-GUUAAGGCCCCAAAGU--GC-GGG-CU-AA---G
-U-GUCAAUC-UAGAAGGGUGUCUCCCACCGAAGACGGCGGACCGUAGGCCCAGCAGCAGCCA-UCGGCUA
AGCAACGCGUAACAGCGGACCCGCU-----GAGGUGGGAGGCCCCGAAGAC-GCACAGGGACUCA-AGCCCG
C----------CGCCGAG-ACCCCGGCCCACGGA--U-----------------------------------
------------------------------------------------------------------------
-----C-AUUG-------------------------------------------------------------
--------------------------------AUCCG-CGUG-GGGUAGGGGGC-GUCGCCG----------
-----------U-G-GGGCG-GAAGCCG--G-GCCGUGAGGU--C-CGGUGGACCC-G-CGG-UGAAUGAGG
AUCC-UGGCGG-UAGUA-GCAGCG--AAGAG-GGGUGG--G---------------AUACCC-CUC-CGCCG
GAAAGGCC-CAGGGUUCC-AUGGUAAUG-UUCAUCAGCCAUGGGUUAGCCGG--UC-CUAAGGUGGGGCCU-
AAU-UGGC---A-CCC--------------------------------------------------------
----------------------------------------ACC-GA-A-AGGGA-AACGGGUU-AAUA-AUC
CCGUG----------------------------------CCGUG----GGGG--------U-AG--------
------------------GUUU--------------------------------------------------
-------GCGG--CAAC---------GC-AG----GC--------------------CCCA-U-UCCC-GAC
GUCUCAG-------------GAUAGG-GUGAGUGGG-GC-------UGCC-------------G-CCCAU-
UUAA-C-CGU-CG-----A-A-------------------GA-----CC----------------GGGG-AG
U-G-CC--------------------GUCAU-GGCGAGAACC---------GGUUGA----AGA-CG-GUA-
AUA--GC-----------------------------------------------------------------
-----CGGGC--CUUGAGCCCGGUU-C-ACC-C-G-ACUCCUGGGGCCC-AUGAAA-A-GGG-AAUGGGGAA
--------------------------------------------------------CGAGCC-CC-CACGAC
CGUACCGAGAACCGAC------------------------------ACAGGU-C-GCCUGGGU-GAGAAGCC
CAAGGC-GGGU-CGGG-UCACGCCA-G-GCUAGGGAACUCGGC-AAAUUAGCCCUGUAACUUCGG-GAUAAA
GGGUG-CCCG-C-GG--UCU-----------GGG-UA-----------------------------------
------------------------------------------------------------------------
------------------------------------------------------------------------
------------------------------------------------------------------------
------------------------------------------------------------------------
------------------------------------------------------------------------
------------------------------------------------------------------------
------------------------------------------------------------------------
------------------------------------------------------------------------
------------------------------------------------------------------------
------------------------------------------------------------------------
------------------------------------------------------------------------
------------------------------------------------------------------------
------------------------------------------------------------------------
----------------------------------------------------------AAACCC--GGGACC
```

```
GUG--GGUCGCAGUGGCAAGGGGGACCUGA-----CUGUUUAACAAAAACAUAGGUCCCCGCGAGCCCG---
-UA-A-------------GGGUUUGU-ACGGGGGCUGA-AUCCUGGCCACUGGCG-G-UACG-UGAAACCCG
GG-------------UUCA-------------ACCGGGCUAAGCGC--CGCUGAAG-GCCGGGGGUAACUCU
GACCCUCUUAAGGUAGCCAAAUGCCUUGCCGGG-UAAGUUCCGGCGUGCAUGAAUGGAUCAACGAGG--UCC
CCACUGUCCCAGCC----UGG-UGCCCGGCGAACCC--ACC----UCCAGGUGCAGAGUCCUGGGACCCCCG
ACGGGGCGAGAAGA-CCCUGUGGACC-UUUACU--GCAGCCC-GUCGUUGGGGUGGGGGAGGGGGUGCAUAG
UGU-AGGUGGGAGCGAU-G-----------AACCAUG-GUC----------------------------UCCG
--------------------------------------GG-CCGUGGGGAU-G--C-GU-CAAUGAAACA-C
CACC-CAUCUCCUCCUCUGCCCCUAACCC-UG--GG-----GU-U-C-------------------------
------------------------------------------------------------------------
------------------------------------------------------------------------
------------------------------------------------------------------------
------------------------------------------------------------------------
------------------------------------------------------------AC----------
------------------------------------------------CCGG-----GGACA-GCGGCGGGU
UGGCAGUUUGGCUGGGGCGGCACACCUCUGA-AAAGAUAUC-AGGGGUGCCCU-AAGCUCCGCUCAGGUCGG
UCAGAAA----CCGUA-GAGGG-UAAGGCCAA-AUGCGGGGCUGACU-GCCC--------------------
--------------CCUUGAACGCUAGGGGGGCAGGGGGGAAA-CCCGGGCCUAGCGAACGCUC--------
---GU-G-----CCCCCACUG--------------------GA-GGGGGCCGGGCAUGUCGG-AA--AAGGU
ACCCCAGGA-GUAACCGGCUCGUCGCGGGUGAGAGUUCCCAUCGACCCCGCGGUUUGGUACCCAGACGUCGU
CUCUUCCCAUCCUGGCGGUGCA-GCAGCCGCCAAGGGUGGGGCUGCCCGCCCAUUAAAGGGGAACGUGAGAU
GGGUUCAGACCGUCGUGAGACAGGUCGGUCUCUACCUGUC--------------------G-GGG-------
-------------------------------GUGC--UGGCCGCCUGAGGGGAAGG-UGCCCUCAGUACGAG
AGGAA-CGGGGCGCCGCGGCCUC-UAGU-C-UACCGGUUGUCC-GGCA-GGGCAG-U-GCCGGGCAGCCACG
CCGCU-UGGGGUAACCGCUGAAGGCA-UC--UAAGCGGGAAACCCGCCCCUA-A-AAU--AGGCGGCC-GC-
--CGCCUGACCGG-GGUUA------------CCCGGUU----------------------------------
-------------------------G----------------------------------------------
------------------------------------------------------------------------
------------------------------------------------------------------------
------------------------------------------------------------------------
------------------------------------------------------------------------
------------------------------------------------------------------------
------------------------------------------------------------------------
-------------------------------------------------GGGGUGG--UU-AGG------GC
UCCCGUAGAAGACGGGGU-UGA-UGGGGU-GGU-GGUGUAUC-CCC-C--------GAG-----GGUU----
--UUCC--------CGAGGGGGG-AGCCU--G-CCAC-UCCCAAUC---GCCCGAUG----------------
------------------------------------------------------------------------
------------------------------------------------------------------------
------------------------------------------------------------------------
-------------------------------------------------
>LSUa_368408_THEPD/1-3059
-----------------------------------CG--CUAAGCCACCC-GGUGGAUG-G-CUCGGC-UCG
G-G---CGCCGAGGAAGGCCGUGG--CAAGCGGCGAUACGCCCCGGGGAGCCGCAUGCAGGCUU------CG
AUC-CGG-GGAUCGCCGAAUGGGACC-U---------CCU-------GC----CGUG-G-G----------UC
A-A-------------C---C----C----A---CGG----C-G-CUCG-GGA-----AACCC---------
```

```
-----------G-----------CAA----------------------------------------GGGGAG
U-A--------------------CC-G-AG-CGGGAACCCCCCCAACGGAAACAUCUUAGUAGGGG--GAGG
AGAAGAAACCAA--UUGGGAUCCCCUGA--GU-AG-GGGCGACCGAAAGGGGG-A-GA-GCCCAAACCGAAC
GUCC-AUGCGA-------AGA------------G---CAU-GGACGGAUGUGGGG--UUG------------
C-AG--GGC-CCC-G-----C------GU-----------------CCCCC--CUA-----------GC--C
C-------GGAUA--GCCG------AAGUCG-GCUG------------------------------------
-------GAA-AG-CCGCG-------CCGUAGAGGGUGACAGCCCCGUA---------GGC---U----AAA
U------CCGGGUGG-GGG-AU--------------GGCG-GGUGU--CCCUGAGUACCACGGCUUG---GU
UUUGCCGUGGGAA-GCUGGGGGGCACC--GACCUCCAAGGCUAAAUA--CGU-C-CCG-AGACCGAUAGC-G
AAC-UAAGUACCGUGAGGGAAAG-CUGAAA-AGCAC-CCCGG-AAGGGG-G-GUG-AAAAGAGCCUGAAACC
G-GGUGGCGACAG-GU--GG-CGCG---GCCCGAAA-GGGUAG-----------------------------
------------------------------------------------------------------------
------------------------------------------------------------------------
------------------------------------------------------------------------
------------------------------------------------------------------------
------------------------------------------------------------------------
-------------------------------------------------------------AU-CCUCCCG-
---AA-----------GG-A-------------------A------ACC-----------------CGG---
------------------------------------------------------------------------
--------------------------------------------------------------G--C-GA---
------------------------------------------------------------------------
-------------------------------------------------C-CG------GG--------GAG
UACGA-------------------------------------------------------------------
------------------------------------------------------------------------
------------------------------------------------------------------------
------------------------------------------------------------------------
------------------------------------------------G-----------------------
------------------------------------------------------------------------
------------------------------------------------------------------------
------------------------------------------------------------------------
------------------------------------------------------------------------
------------------------------------------------------------------------
-------------------GGGAGG-GG-----------------ACCGG--G---GUCGCGCCUUACGUCU
AGAAACACGGGCCGGGGAGUUCA-C-GGCUGUGGCGA-GCCUAAGG--G-GU--------UCAA-------G
C-CC--CGAA--GGCGUAGGGAAAC-CGA-----------------CAA----------------------G
CCCG-UAGCCAG----C-------------------------------------------------------
---CG-------------------------------------------------------------------
------------CAUGG--CUGG------------------------------------UGU-GGGGCG-GG
GUCU-----------------------------------GAAA--------------------------GG-
G-C-CC----------------------GUAGUCACAGCCGUGA-GACCAGAAACC-GAGCGAUCUAGGCCG
GGGCAGGGUGAAGCCCGG-CGAAAG-CCGGGUGGAGGCCCGA--AGGGGUUCUGACGUGCAAUUCGU-UCCC
AUGACCUCGGCCUAGGGGCAAAAGACCAAUCAAGCUCGG-UGAUAGCUGGUUCCCCCCGAAG-CGGAUCCCA
GUCCGGCCCGCC-CUGA-------------------------------------------------------
----------------------GG-----UU-GCCGGUUGGGUAGGGC-ACUGAUU-GGG-AGUC-CGGGGA
C-C-GA-AA-GGUC-C-CAGCUCCCUUUCAAACCCCGAACCGACCG--------------------------
```

```
------------------------------------------------------------------------
------------------G-CA-C--CG-UAGA--UGGGCGG-AGACGGUU-CUGG-C-GGA-UAAGCCGC
CA-G-GCCGAGAG-GGGAAC---AACCCAGACCGG-G-GUUAAGGCCCCCAAGU--GC-CGG-CUAAA---G
-U-GUCAAGC-CAGAAGGGUGUCCCCCGCCUUAGACAGCGGGGCCGUAGGCUUAGAAGCAGCCA-UCGGCUA
AGAAGUGCGUAACAGCUUACCCGCC-----GAGGCGGGGGGCCCCGAAGAU-UACC-GGGACU-A-AGCCGG
C----------CGCCGAG-ACCCCGGGGCACCCC--G-----------------------------------
------------------------------------------------------------------------
-----C-AUUG-------------------------------------------------------------
-------------------------------CGGGGU-GAUC-CGGUAGGGGGGC-GUCCAGG---------
-----------U-G-GACUG-GAAGCCG--G-GCCGUGAGGU--C-CGGUGGAUCC-G-CCU-GGAAUGAAA
AUCC-CGGCGG-UAGUA-AGAGCG-CAAGAG-GGGUGA--G---------------AAUCCC-CUC-CGCCG
A-AAGGGC-AAGGGUUCC-UCAGCAACG-GUCGUCGGCUGAGGGUUAGCCGG--UC-CUAAGGCGGCCCUU-
AAC--CGG---U-GCC--------------------------------------------------------
----------------------------------------GCC-GA-A-AGGGA-AAGGGGUU-AAUA-UUC
CCCUG----------------------------------CCGUG----GGGU-ACGC---U-UU--------
------------------------------------------------------------------------
-------GCGG--CAAC--------GC-GA-----GC--------------------CCCG-C-UCCU-GAC
GCCUCGG-------------GAUAGG-GGGAGUGGG-AC-------UGCC-------------G-UCCCAU-
CCAA-G-CGC-UG-----A-A-------------------GC-----CC----------------GCUG-AG
U-G-CC--------------------GUCAU-GGCGAGAAGC---------GGGUGA----AGG-CG-CGA-
UGG--GC-----------------------------------------------------------------
-----CCGCC---GUUAGGCGGGUU-C-CCC-C-G-ACUCCUGGGGCCC-GUGAAA-A-GGG-AGCGGGGA-
--------------------------------------------------------AGGAGC-CC-CACGCC
CGUACCGAGAACCGAC------------------------------ACAGGUGC-CCCUGGGU-GAAAAGCC
CAAGGC-GUGG-CGGG-GUAACCCA-G-GCUAGGGAACUCGGC-AAAUUAGCCCCGUAACUUCGG-GAGAAG
GGGUG-CCUG-C-GG--UCUU---G----GGGCA-U------------------------------------
------------------------------------------------------------------------
------------------------------------------------------------------------
------------------------------------------------------------------------
------------------------------------------------------------------------
------------------------------------------------------------------------
------------------------------------------------------------------------
------------------------------------------------------------------------
------------------------------------------------------------------------
------------------------------------------------------------------------
------------------------------------------------------------------------
------------------------------------------------------------------------
------------------------------------------------------------------------
------------------------------------------------------------------------
----------------------------------------------------------ACCCCU--GGGACC
GCA--GGUCGCAGUGACAAGGGGGACCCGA-----CUGUUUAAUAAAAACAUAGGUCCCCGCGAGCCCG---
-AA-A-------------GGGUGAGU-ACGGGGGCUGA-AUCCUGGCCACUGGCG-G-UACG-UGAAACCCG
GG-------------UACA-------------ACCGGGCGAAGCGC--CGCCGAAG-GCCGGGAGUAACUCU
GACUCUCUUAAGGUAGCCAAAUGCCUUGCCGGG-UAAGUUCCGGCGCGCAUGAAUGGAUCAACGAGG--UCC
CCACUGUCCCAGCC----UGG-GACCCGCUGAACCC---GC---AACCAGGUGCAGAGUCCUGGGAGUCCCG
```

```
GUGGGGCGAGAAGA-CCCCGUGGAGC-UUCACA--GCAGCCU-GGCGUUGAGACACGGCUGCGGGUGCGUAG
CGU-AGGCGGGAGCGAU-G----------AACCUGG-CCC----------------------------UCCG
--------------------------------------GG-GUCAGGGGAU-G--C-GG-CCAUGAAACA-C
CGCU-CACCUGCGGCUGUGUCCCUAACCC-CG--GG-----GU-U-U-------------------------
------------------------------------------------------------------------
------------------------------------------------------------------------
------------------------------------------------------------------------
------------------------------------------------------------------------
------------------------------------------------------------CC----------
------------------------------------------------CGGG-----GGACA-GCGCCAGGU
GGGCUGUUCGGCUGGGGCGGCACACCCCUGA-GAAGGUAUC-AGGGGUGCCCA-AAGCUCGGCUCAGGCGGG
UCAGAACUCCGCCGUA-GAGGG-CAAGGCCAA-AAGCCGGGCUGACU-GCGC--------------------
--------------CCUUAAACGCGUGGGGCGCAGGCGGGAAA-CCGGGGCCUAGCGAACGCUC--------
---GU-G-----CCCCCUUCG--------------------GU-GGGGGCCGGGCAUGACAG-AA--AAGUU
ACCCCGGGA-GUAACCGGCUCGUCGCGGGCGAGAGUUCACAUCGACCCCGCGGUUUGGUACCCAGACGUCGU
CUCUUCCUAGCUUGGCCCUGCA-GCAGGGGCCAAGAGUGGGGCUGCUCGCCCAUUAAAAGGGAACGUGAGAU
GGGUUUAGACCGUCGCGAGACAGGUCGGACUCUACCUGCC--------------------G-GGA-------
-------------------------------CCGU--UGGCCGCCUGAGGGGAUGG-UCCGCACAGUACGAG
AGGAA-CUGCGGGCCGUGGCCUC-UAGU-G-UACCGGUUGUCC-GGCA-GGGCAC-U-GCCGGGCAGCCACG
CCGCA-AGGGGUAACCGCUGAAGGCA-UC--UAAGCGGGAAACCCACCCCGA-G-ACG--AGGCGGCC-GC-
--UCCCGAUCC-------------------------------------------------------------
----------------------------------------GUUGC-UC------------------------
------------------------------------------------------------------------
---------------------CCGGCGACGGGG---------------------------------------
------------------------------------------------------------------------
------------------------------------------------------------------------
------------------------------------------------------------------------
------------------------------------------------------------------------
-----------------------------------------GCUUUG-GGUCGGGA--CG-AGG------GC
ACCCGUAGAAGACGGGGU-UGA-UGGGGU-GGC-GGUGUAUG-CGC-C--------GAG-----GG------
--UUUC--------CGAGGCGCG-AGCCG--G-CCAC-UCCCAAUA---GCCCGAGGC--------------
------------------------------------------------------------------------
------------------------------------------------------------------------
------------------------------------------------------------------------
-------------------------------------------------
>LSUa_399549_METS5/1-3044
----------------------GCUA-GGGA----CA--AGAAGUCGUCC-GGUGGAUG-G-CUCGGC-UCA
G-G---GGCCUAAGAAGGGCGCGG--CAAGCGGCGAAAUGCCCAGGGGAGGCGCACGCAGCCUU------AG
AUC-CUG-GGAUCCCGAAUGGGAAU-U---------CCU-------GC----CUGG-A-GUU---------A
A-C-----------AGC---U----C----C---AGG----C-G-UCCC-----------------------
-----------G-----------UAA----------------------------------------------
------------------------G-G-GA-CGGGAACCCCCCGAACGGAAGCAUCUUAGUAGGGG--GAGG
AGGAGAAAUCAA--UUGAGAUUCCCUGA--GU-AG-GGGCGACCGAAAGGGGA-A-CA-GCCCAAACCAAAU
CCGC-UAGUGA-------CGA------------G---CUG-GCGGAGAUGUGGUG--UUU------------
C-UG--GCC-CUG-G-----G------UC--A-GGG-GCAACCCAGCUCC--GGU------------GU--C
```

```
U-------GGAUA--GCCG------AAUUCA-CCUG------------------------------------
-------GAA-AG-GUGAG-------CCAUAGAGGGUGACAGCCCCGUA---------GGC---G----AAA
U------CCAGGCGG-GAG-----------------GUGG-CCCAG--GGCAGAGUACCAUCCCCUG---GU
UUGGGGGUGGGAA-GAUGGGAGACACG--UGCUUCCAAGGCUAAAUA--CUU-C-CUG-AGACCGAUAGC-G
AAC-UCAGUACUGUGAAGGAAAG-CUGAAA-AGCAC-CCCGG-AAGGGG-A-GUG-AAAAGUGCCUGAAACC
G-GGCGAUUACAC-AG--GG-CAAG---GCCCUGAA-GGGCUG-----------------------------
------------------------------------------------------------------------
------------------------------------------------------------------------
------------------------------------------------------------------------
------------------------------------------------------------------------
------------------------------------------------------------------------
------------------------------------------------------------AC-UUCCCCCG-
---AA-----------GG-A-------------------A------AGG-----------------GAC---
------------------------------------------------------------------------
--------------------------------------------------------------G--C-AA---
------------------------------------------------------------------------
-------------------------------------------------G-UC------CU--------CAG
UACGA-------------------------------------------------------------------
------------------------------------------------------------------------
------------------------------------------------------------------------
------------------------------------------------------------------------
------------------------------------------------G-----------------------
------------------------------------------------------------------------
------------------------------------------------------------------------
------------------------------------------------------------------------
------------------------------------------------------------------------
------------------------------------------------------------------------
-------------------GGGGAA-UG-----------------ACCAG--G---GUCUUGCUUUUCGUCU
UGAAACACGGGCCGGGGAGGUCA-C-GUCAGUGGCGA-GCCUAAGG--U-G--------GUCAG--------
C-AC--CGAA--GGCAAAGGGAAAC-CGA-----------------GG-----------------------A
UCCG-CAACCCG----G-------------------------------------------------------
---GA-------------------------------------------------------------------
--------------AAC--CGGG------------------------------------UGA-GGGAUC-GG
GUCU-----------------------------------GUCA--------------------------GG-
G-C-CU----------------------UGAGUCACUGGCGUGA-AACUAGAAACC-GGGCGAUCUAGACCU
GGGCAGGUCGAAGUCGGG-GGAAAC-CCCGAUGGAGGACCGAAUAGGGGUUCUGACGUGCAAUUCGU-UCCC
UUGACCUGGGUCUAGGGGCAAAAGACCAAUCUAGCCCGG-UGAUAUCUAGUUCCCUCCGAAA-UGCGUCCUA
GCGCAGCCUCCC-UGGA-------------------------------------------------------
----------------------GG-----UU-GCUCAUGGGGUAGAGAUACUGAUU-GGG-GGC--CUC-UG
A-C-GA-AA-GUCA-G-GGGAACCCAGUCAAACUCCGAACCCAUGG--------------------------
------------------------------------------------------------------------
------------------G-CG-C--CA-GAGA--AGGGGGG-AGUGGAU-C-ACCC-G-GCG-UAAGGUUG
GG-U-GGUAAGAG-GGGAAC---AACCCAGACCCG-G-GUUAAGGUCCCCAAGU--AC-UGG-CU-AA---G
-U-GCGAAUC-UGGAAGGGCGUCCCAUGCCUAAGACAGCGGGAAGGUGGGCCCAGCAGCAGCCA-UCCUCUA
AGGAGUGCGUAACAGCUCACCCGCC-----GAGGCCUGGGGCCCUGCAGACUAGUC-GGGGCUCA-AGCCAG
```

```
U----------CACCGAU-ACCCGGGGGGUGGCG--C-----------------------------------
------------------------------------------------------------------------
-----UCAUUG-------------------------------------------------------------
--------------------------------AGUGU-CACC-CGGUAGGGGGC-GCCGCGG----------
-----------U-G-GGCGA-GAAGGUG--G-GCCGUGAGGU--C-CACUGGACCC-U-CCG-CGGGUGCAG
AUCC-CGGCGG-CAGUA-AUAGCG--AAGGA-GGGUGA--G---------------AAUCCC-UCC-CGCCG
A-AAGGGC-AAGGGUUUC-CCGGCAAUG-UUCGUCAACCGGGAGUUAGCCGG--UC-CUAAGGCAGGGCCU-
AAC-UGGU---A-CCU--------------------------------------------------------
----------------------------------------GUC-GA-A-AGGGA-AAGGGGUU-AAUA-UUC
CCCUG----------------------------------CCACG----GGGGUAGG-------U--------
------------------------------------------------------------------------
-------GCGG--CAAC--------GC-UG----CGG--------------------GGAU-C-UCCU-GAC
GGAUCGA-------------GGUACG-GGGAGUGGG-GC-------CAUC-------------G-CCCCAU-
CCAA-G-UAC-CG-----A-A-------------------GU-----CC----------------CUGG-AG
A-G-CC--------------------GUAAU-GGUGAGAAGG---------GGACGA----AGG-CU-CGA-
UGG--GC-----------------------------------------------------------------
-----CUUCC---GUUAGGAGGGUU-C-UCC-G-G-AUCCUUGGUCCCC-AUGAAA-A-GGG-AGAUCCCAA
--------------------------------------------------------CGAUCC-UC-CGUGAC
CGUACCCAGAACCGAC------------------------------ACUGGUGC-CCCUGGGU-GAGAAGCC
CAAGGC-GUCU-GGGGGGUAACUCA-G-GCUAGGGAACUCGGC-AAAAUGGUCCCGUACCUUCGG-AAGAAG
GGAUG-CCUA-C-CA--UUGU----------AGU--------------------------------------
------------------------------------------------------------------------
------------------------------------------------------------------------
------------------------------------------------------------------------
------------------------------------------------------------------------
------------------------------------------------------------------------
------------------------------------------------------------------------
------------------------------------------------------------------------
------------------------------------------------------------------------
------------------------------------------------------------------------
------------------------------------------------------------------------
------------------------------------------------------------------------
------------------------------------------------------------------------
------------------------------------------------------------------------
----------------------------------------------------------UACGCU--GCAGUG
GUA--GGUCGCAGUGACCAGGGGGAUCUGA-----CUAUUUAAUAAAAAUAUAGGUCCCCGCUAGCCCG---
-AA-A-------------GGGUGUGA-ACGGGGGCUAA-AUCCUGGCCACUGGCG-G-UCGG-UUAAACCCG
GG-------------UCCA-------------ACCGGGCGAAGCCC--CGCUGAAG-GCCGGGGGUAACUCU
GACCCUCUCAAGGUAGCCAAAUGCCUUGCCGGG-CAAGUUCCGGCGCGCAUGAAUGGAUCAACGUGA--UCC
CCACUGUCCCAGCC----UGG-GGCCCCAUGACCGC--CCA---GAGUGGGUUCACAGUCCCGCAACUCCUC
ACACCGAGAGAAGA-CCCCGUGGAGC-UUCACC--GCAGCCU-GGCGCUGGUUCUCGGGCGUUCAUGCGUAG
CGU-AGGUGGGAGGCGUCG----------AAGUCGU-CCC----------------------------UUCG
--------------------------------------GG-GGCGAUGGAG-C--C-GC-AAGUGAAACA-C
CACC-CAUGAAUGCUCGGGAACCUAACCU------------GC-G-A-------------------------
------------------------------------------------------------------------
```

```
------------------------------------------------------------------------
------------------------------------------------------------------------
------------------------------------------------------------------------
------------------------------------------------------------------------
--------------------------------------------------AG-----GGACA-ACGUCAGGU
GGGCGGUUCGGCUGGGGCGGCACUCCCGCGA-AAAGAUAAC-ACGGGAGCCCA-AAGGUCAGCUCAGGCGGU
ACAGAACGCCGCCGUA-GAGUG-CAAGGCAA-AAGCUGGCUGACG-AGAC----------------------
--------------CCUUCAAAGUACGGGGUCUCGACGCGAAA-GCGCGGCCUAGCGAACGCUC--------
---AU-G-----CCCCCACAC--------------------GU-GGGGGCUGGGCAUGACAG-AA--AAGUU
ACCCCGGGG-AUAACAGGGUCGUCGCGGGCGAGAGCUCCCAUCGACCCCGCGGUUUGCUACAUCGAUGUCGG
CUCUUCCCACCCUGGAGGUGCA-GCUGCCUCCAAGGGUAGGGCUGCCCGCCCGUUAAAGGGGAACGUGAGCU
GGGUUUAGACCGUCGCGAGACAGGUCGGACUCUAAGGGUG--------------------A-GGG-------
-------------------------------GUGC--ACGCCGCCUGAGGGGAAGG-UACUCCUAGUACGAG
AGGAA-CAGGGUAUCGGGGCCUC-UAGU-U-UGCCGGUUGUCC-GAUA-GGGCAG-C-GCCGGGCAGCCACG
CCCUA-GGGGGUAACCGCUGAAAGCA-UC--UAAGCGGGAACCCCUCCCCGA-A-AAG--AGGCGGCG-GU-
--GUAUUGGCCGC-AA----------------GCA-------------------------------------
------------------------------------------------------------------------
------------------------------------------------------------------------
------------------------------------------------------------------------
------------------------------------------------------------------------
------------------------------------------------------------------------
------------------------------------------------------------------------
------------------------------------------------------------------------
-------------------------------------------------GCAUGCG--GG-AGC------CC
UCCCCAAGAACAGGGGGU-UGA-UGGGGU-GGG-GGUGUAAG-CCU-C--------GAG-----GGCGA---
-AAGUC--------CGAGGGGUUCAGCCU--G-CCAC-UCCCAAUC---GGGCGAGUC-CAGCGC-------
------------------------------------------------------------------------
------------------------------------------------------------------------
------------------------------------------------------------------------
-------------------------------------------------
>LSUa_330779_SULAC/1-3022
-----------------------------------GC--CGAAGCCUCCC-GGUGGAUG-G-CUCGGC-UCG
G-G---CACCGAAGAAGGGCGCGG--CAAGCAGCGAAAUGCUCGGGUGAGGCGCAAGCAGCCGU------UG
ACC-CCG-AGGUCCCCUAAUGGGAUA-U--------CCU-------GC----CGG---------------GU
U-U---------------------------C---CGG----C-G-CUCC-CGG-----U-------------
-----------U-----------UAUA--------------------------------------------A
C-U--------------------GG-G-AG-UGGGAACCCCCCGAACGGAAACAUCUUAGUAGGGG--GAGG
AAAAGAAAUCAA--UUGAGAUCCCCUGA--GU-AG-GGGCGACCGAAAGGGGG-A-CA-GCCCAAACUAAAC
CUGC-CGAUGA-------UAA------------G---UCG-GUGGGGAUGUGGUG--UUA------------
C-GA--CCU-CUA-G-----C------CU--GAGGUUCGACCUCGGCUUUC--CUA-----------AC--C
U-------AUCUA--GCCG------AACUCC-CCUG------------------------------------
-------GAA-CG-GGGGG-------CCAUAGAGGGUGAAAGCCCCGUA---------GGC---U----AAA
G------AUAGGUGG-AAA-----------------GUGG-CUAGA--GGUAGAGUACCAUCCCCUG---GU
UUGGGGGUGGGAA-GUUAGGGGACACG--UGCCUCUAAGGCUAAAUA--UGU-C-CCG-AGACCGAUAGC-A
AAC-UAAGUACCGUGAGGGAAAG-CUGAAA-AGAAC-CCCGG-AAGGGG-GAGUG-CAAAGAGCCUGAAACC
```

```
G-GGUGGUUAUAC-AG--GG-UGUG---GCUCGAAA-GAAGUG-----------------------------
------------------------------------------------------------------------
------------------------------------------------------------------------
------------------------------------------------------------------------
------------------------------------------------------------------------
------------------------------------------------------------------------
------------------------------------------------------------AA-CCCUUCCG-
---AA-----------GG-A-------------------A------AGG-----------------GGC---
------------------------------------------------------------------------
--------------------------------------------------------------G--C-AA---
------------------------------------------------------------------------
-------------------------------------------------G-CC------CU--------UAG
UACGA-------------------------------------------------------------------
------------------------------------------------------------------------
------------------------------------------------------------------------
------------------------------------------------------------------------
------------------------------------------------G-----------------------
------------------------------------------------------------------------
------------------------------------------------------------------------
------------------------------------------------------------------------
------------------------------------------------------------------------
------------------------------------------------------------------------
-------------------GAAGGG-CG-----------------AUCGG--G---GUCACGCCUUUCGUCU
UGAAACACGGGCCGGGGAGUUCA-C-AUCAGUGGCGA-GCUUAAGG--A-G--------AUUAU--------
C-UC--CGAA--GGCAUAGGGAAAC-CAA-----------------GU-----------------------G
CCCG-CAGCCUA----G-------------------------------------------------------
---U--------------------------------------------------------------------
--------------UUC--UAGG------------------------------------CGA-GGGGCA-GG
GUCU-----------------------------------GUCA--------------------------GG-
G-C-CU----------------------GAAGCCACUGAUGUGA-GGCUAGAAACC-GGGCGAUCUAGUCCG
GGGCAGGCUGAAGGUGGG-GGAAAC-CCCACUGGAGGGCCGAAUAGGGGUUCUGACGUGCAAUUCGU-UCCC
UUGACCUCGGACUAGGGGCAAAAGACCAAUCUAGCCCGG-UGAUAGCUAGUUCCCCCCGAAA-UGCGUCCUA
GCGCAGCCUCCC-UAAA-------------------------------------------------------
----------------------GG-----CA-GCUCGCGGGGUAGAGUGACAGAUC-GGG-GGC--UCC-AG
G-C-GA-AA-GCCU-G-GGGCUUCCGGUCUAACUCCGAACCCACGA--------------------------
------------------------------------------------------------------------
------------------G-CG-C--CG-AAGA--AGGGGGG-AGUGGGU-C-ACUC-G-GCG-UAAGGUUG
GG-U-GGCAAAAG-GGGAAC---AACCCAGACCUG-G-GUUAAGGCCCCAAAGU--CC-CGG-CU-AA---G
-U-GCCAA---CGAAAAGGCGUCUCCAGCCUUAGACAGCGGGAAGGUGGGCCCAGCAGCAGCCA-UCCUCUA
AGGAGUGCGUAACAGCUCACCCGCC-----GAGGCUGGAGGCCCUAAAGAUUGGUC-GGGGCUCA-AGCCGG
G----------CGCCGAG-ACCCAGGAGGGGGUC--U-----------------------------------
------------------------------------------------------------------------
-----CUACUA-------------------------------------------------------------
--------------------------------GAGAU-CCUC-UGGUAGGGGGGC-GCUGUGA---------
-----------U-G-GGGUA-GAAGGUG--G-GUCGUGAGAU--C-CACUGGACCC-G-UCA-CAGGUGCAG
```

```
AUCC-CGGCGG-UAGUA-ACAGCG--AAGGG-GGGUGA--G---------------AAUCCC-CCU-CGCCG
G-AAGGGC-AAGGGUUUC-CCGGCAACG-UUCGUCAGCCGGGAGUUAGCCGG--UC-CUAAGGUAGGGCCU-
AAU-AGGU---A-CCU--------------------------------------------------------
----------------------------------------ACC-GA-A-AGGGA-AAGGGGUU-AACA-UUC
CCCUG----------------------------------CCUCC----CGGGUAGG-------U--------
------------------------------------------------------------------------
-------GCGG--UAAC--------GC-AA-----GC--------------------CAGA-C-UCCU-GAC
GGAUUGG-------------GGUAGG-GAGAGUAGG-AC-------CACC-------------G-UCCUAC-
CCAA-G-CAC-UC-----A-A-------------------GC-----CC----------------UUGG-AG
A-G-CC--------------------GUAAC-GGUGAGAAGA---------GGGCGA----AGG-UG-UGA-
UGG--GC-----------------------------------------------------------------
-----CUUCC---GUUAGGAGGGUU-C-UCC-U-G-AUCCCUAGUCCCC-AUGAAA-A-GGG-AGUCUGGAA
--------------------------------------------------------CGAUCC-CG-GGAGAC
CGUACCUAGAACCGAC------------------------------ACUGGUGC-CCCUGGGU-GAGAAGCC
CAAGGC-GUCU-GAGGGGUAACCCA-G-GCUAGGGAACUCGGC-AAAUUAGCCCCGUAACUUCGG-GAGAAG
GGGUG-CCUA-U-CG--UGGU---U----U------------------------------------------
------------------------------------------------------------------------
------------------------------------------------------------------------
------------------------------------------------------------------------
------------------------------------------------------------------------
------------------------------------------------------------------------
------------------------------------------------------------------------
------------------------------------------------------------------------
------------------------------------------------------------------------
------------------------------------------------------------------------
------------------------------------------------------------------------
------------------------------------------------------------------------
------------------------------------------------------------------------
------------------------------------------------------------------------
----------------------------------------------------------AACAAA--GCCACG
AUA--GGUCGCAGUGACCAGAGGGACCUGA-----CUGUUUAAUAAAAACAUAGGUCCCCGCUAGCCCG---
-AA-A-------------GGGUGUGU-ACGGGGGCUAA-AUCCUGGCCACUGGUG-G-UUGG-UUAAAUCCG
GG-------------UUCA-------------ACCGGGCGAAGCCC--CACCGAAG-GCCGGGGGUAACUCU
GACCCUCUUAAGGUAGCCAAAUGCCUUGCCGGG-UAAGUUCCGGCGCGCAUGAAUGGAUCAACGCGG--UCC
CUACUGUCCCAGCC----UGG-GGCCUCGUGAACGC--CCU---GAGCCGGUGCACAGUCCGGCAUCUCCCU
ACACCGAGAGAAGA-CCCCGUGGAGC-UUCACC--GCAGCCU-GGCGUUGUCCCUCGGGCGUUUAUGCGUAG
AGU-AGGUGGGAGGGGUCG----------AACCUGU-CCU----------------------------UUCG
--------------------------------------GG-GGCAGGGGAC-C--C-GA-AAGUGAAACA-C
CACC-CAUGGACGCUCGAGGGACUAACCU-CU--C------GA-A-A-------------------------
------------------------------------------------------------------------
------------------------------------------------------------------------
------------------------------------------------------------------------
------------------------------------------------------------------------
-------------------------------------------------------------G----------
------------------------------------------------AGAG-----GAACA-ACGUCAGGU
```

```
GGGCGGUUCGGCUGGGGCGGCACUCCCGCGA-AAAGAUAAC-ACGGGAGCCCA-AAGGUCGGCUCAGGCGGU
ACAGAACGCCGCCGUA-GAGCG-CAAGGGCAA-AAGCCGGCCUGACG-UGAC--------------------
--------------CCUUCCAAGUACGGGGUCACGACGCGAAA-GCGGGGCCUAGCGAACGCUC--------
---GU-G-----CCCCCACAC--------------------GU-GGGGGCCGGGCAUGACAG-AA--AAGUU
ACCCCGGGG-AUAACAGGGUCGUCGCGGGCGAGAGCUCACAUCGACCCCGCGGUUUGCUACAUCGAUGUCGG
CUCUUCCCACCCUGGAGGUGCA-GCUGCCUCCAAGGGUAGGGCUGCCCGCCCGUUAAAGGGGAGCGUGAGCU
GGGUUUAGACCGUCGCGAGACAGGUCGGACUCUAAGGGUA--------------------G-GGA-------
-------------------------------GUGC--GGACCGCUUGAGGGGAAGG-AACCCCUAGUACGAG
AGGAA-CAGGGUUCCGGGGCCUC-CAGU-U-UACCGGUUGUCC-GGUA-GGGCAA-U-GCCGGGCAGCCGCG
CCCUG-AGGGGUAACCGCUGAAAGCA-UC--UAAGCGGGAACCCCUCCCCUA-A-AAG--AGGCGGUC-AG-
--GCGUUA----------------------------------------------------------------
------------------------------------------------------------------------
------------------------------------------------------------------------
------------------------------------------------------------------------
------------------------------------------------------------------------
------------------------------------------------------------------------
------------------------------------------------------------------------
------------------------------------------------------------------------
-----------------------------------------------------GCC--GG-GGC------CU
UCCCCUAGAAGAGGGGGU-UGA-UGGGGU-GGG-GAUGUAAG-CUC-C--------AAG-----GUUGU---
-AAGAC--------CGAGGAGUUUAGUCC--G-CCAC-UCCCAAUC---AGGCUCCCC-CC-----------
------------------------------------------------------------------------
------------------------------------------------------------------------
------------------------------------------------------------------------
-------------------------------------------------
>LSUa_273063_SULTO/1-3030
----------------------CC---AGGG----GC--UGACGCCACCC-GGUGGAUG-G-CUCGGC-UUG
G-G---CGCCGAAGAAGGGCGCGG--CAAGCUGCGAAAUGCCCGGGUUAGGCGCAAGCAGCCGU------UG
ACC-CCG-GGAUCCCCGAAUGGGAUA-U--------CCU-------GC----CGG---------------GU
U-A---------------------------C---CGG----C-G-CUCC-CAG-----C-------------
-----------U-----------CCU---------------------------------------------G
CAU--------------------GG-G-AG-CGGGAACCCCCCGAACGGAAACAUCUUAGUAGGGG--GAGG
AAAAGAAAUCAAA-UCGAGAUCCCCUGA--GU-AG-GGGCGACCGAAAGGGGG-A-GA-GCCCAAACCAAAC
CUGC-CGGUGA-------UAA------------G---CCG-GUGGGGAUGUGGUG--UUA------------
U-UG--GCC-UCU-G-----G------CCU-GGGGUUCGACCCAACCUCC--CUA------------GC--C
C-------UCUGA--GCCG------AACUCC-CCUG------------------------------------
-------GAA-UG-GGGGG-------CCAUAGAGGGUGAUAGCCCCGUA---------GGC---G----AAG
G------A-GGGUGG-GAG-----------------GUGG-CCAGA--GGCAGAGUACCAUCCCCUG---GU
UUGGGGGUGGGAA-GUUGGGGGACACG--UGCCUCCAAGGCUAAAUA--CGU-C-CCA-AGACCGAUAGC-G
AAC-UUAGUACCGUGAGGGAAAG-CUGAAA-AGAAC-CCCGG-AAGGGG-GAGUG-AAAAGAGCCUGAAACC
G-GGUGGUUACAG-AG--GG-CAGG---GCUCGAAA-GGGGUG-----------------------------
------------------------------------------------------------------------
------------------------------------------------------------------------
------------------------------------------------------------------------
------------------------------------------------------------------------
```

```
------------------------------------------------------------------------
------------------------------------------------------------AG-UCCACCCG-
---AA-----------GG-A-------------------A------AGG-----------------GCU---
------------------------------------------------------------------------
--------------------------------------------------------------G--U-GA---
------------------------------------------------------------------------
-------------------------------------------------A-GC------CC--------GAG
UACGA-------------------------------------------------------------------
------------------------------------------------------------------------
------------------------------------------------------------------------
------------------------------------------------------------------------
------------------------------------------------G-----------------------
------------------------------------------------------------------------
------------------------------------------------------------------------
------------------------------------------------------------------------
------------------------------------------------------------------------
------------------------------------------------------------------------
-------------------GGUGGA-UG-----------------ACCGG--G---GUCCUGCCUUUCGUCU
UGAAACACGGGCCGGGGAGUUCA-C-GCCAGUGGCGA-GCUUAAGG--G-G--------GUUAA--------
C-CC--CGAA--AGCGUAGGGAAAC-CGA-----------------GC-----------------------G
CCCG-CAACCCG----G-------------------------------------------------------
---GA-------------------------------------------------------------------
--------------AAC--CGGG------------------------------------UGA-GGGGCG-AG
GUCC-----------------------------------GUCA--------------------------GG-
G-C-CU----------------------GGAGUCACUGGCGUGA-GGCUAGAAACC-AGGCGAUCUAGUCCG
GGGCAGGCCGAAGUCGGG-GGAAAC-CCCGAUGGAGGGCCGAAUAGGGGUUCUGACGUGCAAUUCGU-UCCC
CUGACCCCGGACUAGGGGCAAAAGACCAAUCUAGCCUGG-UGAUAGCUAGUUCCUCCCGAAA-UGCGUCUUA
GCGCAGCCUCCC-UGGA-------------------------------------------------------
----------------------GG------UC-GCCCGCGGGUAGAGUGACAGAUC-GGG-GGU--UCC-AG
A-C-GA-AA-GUCU-G-GGGCUCCCGGUCUAACUCCGAACCCACGG--------------------------
------------------------------------------------------------------------
------------------G-CA-C--CG-AAGA--AGGGGGG-AGUGGGU-C-ACCC-G-GCG-UAAGGUUG
GG-U-GGCAAAAG-GGGAAC---AACCCAGACUCG-G-GUUAAGGCCCCUAAGU--GC-CGG-CU-AA---G
-U-GCCAA---UCAAAAGGCGUCCCCAGCCUUAGACAGCGGGAAGGUGGGCCCAGCAGCAGCCA-UCCUCUA
AGGAGUGCGUAACAGCUCACCCGCC-----GAGGCUGGGGGCCCUGAAGAUUGGUC-GGGGCUCA-AGCCGG
C----------CGCCGAG-ACCCGAGGGUAGGU--C-----------------------------------
------------------------------------------------------------------------
-----CCACUG-------------------------------------------------------------
--------------------------------GGACC-UACU-CGGUAGGGAGGC-GCUGCGG---------
-----------U-A-GGGCA-GAAGGUG--G-GCCGAAAGGU--C-CACUGGACCU-A-CCG-CAGGUGCAG
AUCC-CGGCGG-CAGUA-GCAGCG--AGUAG-GGGUGA--G---------------AAUCCC-CCA-CGCCG
G-AAGGGC-CAGGGUUUC-CCGGCAAUG-GUCGUCAGCCGGGAGUUAGCCGG--UC-CUAAGGCGGGGCCU-
AAC-UGGU---A-CCC--------------------------------------------------------
----------------------------------------GCC-GA-A-AGGGA-AAGGGGUU-AAUA-UUC
CCCUG----------------------------------CCUCC----CGGGUAGG-------U--------
```

```
------------------------------------------------------------------------
-------GCGG--CAAC--------GC-AA-----GC--------------------CCGG-C-UUCU-GAC
GGAUCGG-------------GCUAGG-GGGAGCGGG-GC-------CACC-------------G-CCCCGC-
CCAA-G-CGC-UU-----A-A-------------------GC-----CC----------------CUGG-AG
A-A-CC--------------------GUAAU-GGUGAGAAGG---------GGGCGA----AGG-CG-UGA-
UGG--GC-----------------------------------------------------------------
-----CUGCC---GUUAGGCGGGUU-C-CCC-U-G-AUGCCUGGUCCCC-AUGAAA-A-GGA-AGCCGGGAA
--------------------------------------------------------CGAUCC-CG-GGAGAC
CGUACCGAGAACCGAC------------------------------ACAGGUGC-CCCUGGGU-GAGAAGCC
CAAGGC-GUCU-GGGGGGUAACCCA-G-GCUAGGGAACUCGGC-AAAUUAGCCCCGUAACUUCGG-GAGAAG
GGGUG-CCUA-U-CA--UGGU---C----GU-----------------------------------------
------------------------------------------------------------------------
------------------------------------------------------------------------
------------------------------------------------------------------------
------------------------------------------------------------------------
------------------------------------------------------------------------
------------------------------------------------------------------------
------------------------------------------------------------------------
------------------------------------------------------------------------
------------------------------------------------------------------------
------------------------------------------------------------------------
------------------------------------------------------------------------
------------------------------------------------------------------------
------------------------------------------------------------------------
----------------------------------------------------------UAAACG--GCCAUG
AUA--GGUCGCAGUGACAAGGGGGACCUGA-----CUGUUUAAUAAAAACAUAGGUCCCCGCUAGCCCG---
-UA-A-------------GGGUGAGU-ACGGGGGCUAA-AUCCUGGCCACUGGCG-G-UUGG-UGAAACCCG
GG-------------UACA-------------ACCGGGCGAAGCCC--CGCUGAAG-GCCGGGGGUAACUCU
GACCCUCUUAAGGUAGCCAAAUGCCUUGCCGGG-UAAGUUCCGGCGCGCAUGAAUGGAUCAACGAGG--UCC
CCACUGUCCCAGCC----UGG-GGCCCCGCGAACGC--CCU---GAGCCGGUGCACAGUCCGGCACCCCCCU
ACACCGAGAGAAGA-CCCCGUGGAGC-UUCACC--GCAGCCU-GGCGUUGUCCUCCGGGCGCUCAUGCGUAG
AGU-AGGUGGGAGGGGUCG----------AAGCCGC-UCC----------------------------UUCG
--------------------------------------GG-GGCGGUGGAC-C--C-GA-AAGUGAAACA-C
CACC-CAUGAGUGCUCGGAGGACUAACCU-C----------GA-A-A-------------------------
------------------------------------------------------------------------
------------------------------------------------------------------------
------------------------------------------------------------------------
------------------------------------------------------------------------
------------------------------------------------------------------------
-------------------------------------------------GGG-----GGACA-UCGUCAGGU
GGGCGGUUCGGCUGGGGCGGCACUCCCGCGA-AAAGAUAAC-ACGGGAGCCCA-AAGGUCGGCUCAGGCGGU
ACAGAACGCCGCCGUA-GAGUG-CAAGGGCAA-AAGCCGGCCUGACG-GGAC--------------------
--------------CCUCCAAAGUACGGGGUCCCGACGCGAAA-GCGGGGCCUAGCGAACGCUC--------
---GU-G-----CCCCCACAC--------------------GU-GGGGGCCGGGCAUGACAG-AA--AAGUU
ACCCCGGGG-AUAACAGGGUCGUCGCGGGCGAGAGCUCACAUCGACCCCGCGGUUUGCUACAUCGAUGUCGG
```

CUCUUCCCACCCUGGGGGUGCA-GCUGCCCCCAAGGGUAGGGCUGCCCGCCCGUUAAUGGGGAGCGUGAGCU
GGGUUUAGACCGUCGCGAGACAGGUCGGACUCUAAGGGUA--------------------G-GGG-------
-------------------------------GUGC--UGGCCGCCUGCGGGGAAGG-UACCCCUAGUACGAG
AGGAA-CAGGGUGCCGGGGCCUC-CAGU-U-UACCGGUUGUCC-GGUA-GGGCAG-C-GCCGGGCAGCCGCG
CCCUG-AGGGGUAACCGCUGAAAGCA-UC--UAAGCGGGAACCCCUCCCCUA-A-AAG--AGGCGGCC-AG-
--GCCGCA----------------------------------------------------------------
------------------------------------------------------------------------
------------------------------------------------------------------------
------------------------------------------------------------------------
------------------------------------------------------------------------
------------------------------------------------------------------------
------------------------------------------------------------------------
------------------------------------------------------------------------
---------------------------------------------------AGGCC--GG-GGG------CC
UCCCCCAGAAGAGGGGUU-UGA-UGGGGU-GGG-GGUGUAAG-CCC-C--------AAG-----GGU-----
-UUUCC--------CGAGGGGUUUAGCCC--G-CCAC-UCCCAAUA---GGCCCCUCC-CUUGG--------
------------------------------------------------------------------------
------------------------------------------------------------------------
------------------------------------------------------------------------
-------------------------------------------------
>LSUa_190192_METKA/1-3094
----------------------CG---GGUC----GG--CUAAGCCGUCG-GGUGGAUG-C-CUUGGC-UCG
G-G---CGCCGAGGAAGGCCGUGG--CAAGCUGCGAUAAGCCCGGGCGAGGCGCAUGCAGCCGU------GG
AAC-CCG-GGAUCGCCGAAUGGGACC-U--------CUC-------GC----CGGG-G-CC---------GA
A-A------------GG---U----C----C---CGG----C-A-CUCC-GGG-----GGGAA-ACCC-CGG
GCC--------C-----------UCGU--------------GGC------------C--CGGGGGAGUAC-C
C-C--------------------CG-G-AG-GGGGAACCCGCCGAACUGAAACAUCUUAGUAGGCG--GAGG
AAAAGAAAGCAAC-ACGCGAUGCCGUGA--GU-AG-GGGCGACCGAAAACGGC-A-GA-GGGCAAACUGAAC
ACCC-GCCCGA--------AA------------G---GGC-GGGUGGAUGUGGGG--UUG------------
C-AG--GCC-CGG-G-----C------CGCCUAUGUG--------GCCGUC--UCC-----------CC--C
G-------GAACG--GCCG------AAGUGC-CCUG------------------------------------
-------GAA-CG-GGCCG-------CCGGAGAGGGUGACAGCCCCGUA---------GGC---C----GCC
G------GGGGACGG-CCG-----------------CUGG-CCCGG--GCCUGAGUACCGUCGGUUG---GA
UAUCCGGCGGGAA-UCUGGGGGACAUC--GGCCCCCAACCCUAAAUA--CGU-C-CCG-AGUCCGAUAGC-G
AAC-A-AGUACCGUGAGGGAAAG-GUGAAA-AGAAC-CCCGG-AAGGGG-A-GUG-AAAAGAGCCUGAAACC
C-GACGGCGAUAG-UCU-GC-CGGG---GCCCGCAA-GGGAUG-----------------------------
------------------------------------------------------------------------
------------------------------------------------------------------------
------------------------------------------------------------------------
------------------------------------------------------------------------
------------------------------------------------------------------------
------------------------------------------------------------AA-GCCCCCCG-
---AA-----------GG-A-------------------A------ACC-----------------CCG---
------------------------------------------------------------------------
--------------------------------------------------------------G--U-GA---

```
------------------------------------------------------------------------
-------------------------------------------------C-GG------GG--------GAG
UACGA-------------------------------------------------------------------
------------------------------------------------------------------------
------------------------------------------------------------------------
------------------------------------------------------------------------
------------------------------------------------G-----------------------
------------------------------------------------------------------------
------------------------------------------------------------------------
------------------------------------------------------------------------
------------------------------------------------------------------------
------------------------------------------------------------------------
-------------------GGGGGC-CG-----------------ACCGG--G---GUCCCGGCGUUCGUUU
UGAAACACGAGGCGGGGAGUCCG-C-GGCCGCGGCGA-GUCUAAGG--G-G--------UUCAA-------A
C-CC--CGAA--GGCGUAGGGAAAC-CGA-----------------CAC----------------------G
CCCG-CAGCCCG----G-------------------------------------------------------
---CCG------------------------------------------------------------------
-----------UUAGGC--CGGG------------------------------------CCA-GGGGCG-GG
GUCC-----------------------------------UAAUA-------------------------GG-
G-C-CC----------------------GCAGCCGCGGCCGCGG-GACCCGAAACC-GGCCGAUCUAGCCCG
GGGCAGGGUGAAGCGGCC-CUAAAG-GGCCGUGGAGGCCCGC-UAGGGGUGCUGUAGUGCAAAACGC-UCCC
GUGACCCCGGGCUAGGGGUGAAAGGCCAAUCGAGGCCGG-AGACAGCUGGUUCCCCCCGAAA-CGGCCCGCA
GGUCGGCCCGGC-CGGA-------------------------------------------------------
----------------------GG-----UA-GGUGGCGGGGUAGAGC-ACUGAUU-CCC-CGGU-UAGGGG
C-C-GA-GA-GGCC-C-CGCCGGGGAGUCAAACUCCGAACCCGCCA--------------------------
------------------------------------------------------------------------
------------------C-CG-C--CG-UAGA--AGGCCGG-AGUUGGG-G-CCGG-C-GGG-UAAGCCGC
CG-G-UCUGAGAG-GGGAAU---AACCCAGACCGG-G-GUUAAGGCCCCAAAGU--GC-CGG-CU-AA---G
-U-GUUAAAU-AGAAAGGGAGUCCCCGGCCGAAGACAGCGGGGAGGUAGGCUUAGAAGCAGCCA-UCCUUUA
AAGAGUGCGUAACAGCUCACCCGUC-----GAGGUCGGGGGCCCCGAAAAU-GGAC-GGGGCUGA-AGCCGG
C----------CGCCGAG-ACCCCGGGGCCGCGG--G-----------------------------------
------------------------------------------------------------------------
-----CCGAUG-------------------------------------------------------------
-------------------------------GCCCGC-GGGC-CGGUAGGGGGGC-GCCCCGG---------
-----------C-G-GCUCA-GAAGCCG--G-GCCGUGAGGU--C-CGGUGGAGCC-G-UCG-GGGACGAGA
AUCC-CGCCGG-UAGUA-GCAGCA--AAGCG-GGGUGA--G---------------AAUCCC-CGC-CGCCG
G-AGGGGC-CAGGGUUCC-UCGGCAAUG-UUCGUCAGCCGAGGGUAAGUCGG--CC-CUAAGGCCGGCGGU-
AAC-ACCG---A-CCG--------------------------------------------------------
----------------------------------------GCC-GA-A-AGGGA-AACGGGUU-AAUA-UUC
CCGUA----------------------------------CCGCG----GGGGUACG-------C--------
------------------------------------------------------------------------
-------CCGG--CAAC--------GG-GA----AGC--------------------CCGG-C-GGCC-GAC
GCCUCGG-------------GGUAGG-CCGGGCGG-GC--------UGUC-------------G-CCCCGU-
CCAA-C-CGG-UG-----A-A-------------------GG-----CC----------------GGGG-AG
U-C-CC--------------------GUAAU-GGGGAGAACC---------GGCCGA----AGC-CG-GGA-
```

```
AGG--GC-----------------------------------------------------------------
-----CGCCC--GUUAUGGGCGGUC-C-GGC-C-G-AUCCCCGGGGCCC-GUGAAA-A-GGC-CGCCGGGAA
--------------------------------------------------------GGAUCC-CC-CGCGAC
CGUACCGAGAACCGAC------------------------------ACAGGUGC-CCCUGGGU-GAGAAGCC
UAAGGC-GCGG-CGGG-GUAACCCG-G-CCGAGGGAAAUCGGC-AAAUUGGCCCCGUAACUUCGG-GAGAAG
GGGUG-CCCG-C-GG--CCCUGACC----CCGAG-GGUCGAACCUCGGGGGA--------------------
------------------------------------------------------------------------
------------------------------------------------------------------------
------------------------------------------------------------------------
------------------------------------------------------------------------
------------------------------------------------------------------------
------------------------------------------------------------------------
------------------------------------------------------------------------
------------------------------------------------------------------------
------------------------------------------------------------------------
------------------------------------------------------------------------
------------------------------------------------------------------------
------------------------------------------------------------------------
------------------------------------------------------------------------
------------------------------------------------------------------GGGGCC
GCG--GGUCGCAGUGACUAGGGGGGGCCGA-----CUGUUUAAUAAAAACAUAGGUCCCGGCUAGCCCG---
-AA-A-------------GGGCUGGU-ACCGGGGCCGA-CGCCUGCCCAGUGCCG-G-UACG-UGAAGCCCG
GG-------------UACA-------------ACCGGGUGAAGCGC--CGGUAAAC-GGCGGGGGUAACUAU
AACCCUCUUAAGGUAGCGAAAUGCCUUGCCGGU-UAAGUACCGGCUUGCAUGAAUGGCGUAACGAGC--CCC
CCACUGUCCCCGGC----CGG-GACCCGGUGAACCC--GCC---AUCCCCGUGCAGAGGCGGGGGACCCCCG
GCGGGAAGCGAAGA-CCCCGUGGAGC-UUUACC--GCAGCCU-GCCGUUGGGGCACGGCUGCGGGUGUAUAG
CGU-AGGUGGGAGCCGUUAC-------UCAGGGGCG-GCC----------------------------GCCA
--------------------------------------GGCCGUCCCCGAG-G--C-GC-CCAUGGAACA-C
CACC-CUCCCGCGGCUGUGCCCCUAACCC-GG--GCC--G-AG-A-A-------------------------
------------------------------------------------------------------------
------------------------------------------------------------------------
------------------------------------------------------------------------
------------------------------------------------------------------------
-------------------------------------------------------G----GC----------
------------------------------------------------CCGG-----GGACA-GCGGUUGGU
GGGCGGUUUGGCUGGGGCGGCACGCCCCCGA-GAAGGCAUC-GGGGGCGCCCA-AAGGUCGGCUCAGGCGGG
UCAGAAAUCCGCCGUA-GAGUG-CAAGGGCAA-AAGCCGGCCUGACU-GGGU--------------------
--------------CCCGCACAAUAAGGGACCCAGGCGGGAAA-CCGUGGCCUAGCGAACCCCG--------
---GU-G-----CCGCC-CCG--------------------GU-GGCGGCCCGGGAUGACAG-AA--AAGCU
ACCCCGGGG-AUAACAGGGUGGUCGCGGGCAAGAGCCCACAUCGACCCCGCGGCUUGCUACCUCGAUGUCGG
CUCUCCCCAUCCUGGCGGUGCA-GCAGCCGCCAAGGGUGGGGUUGUUCGCCCAUUAAAGGGGAACGUGAGCU
GGGUUUAGACCGUCGUGAGACAGGUCGGUCGCUAUCUGCC--------------------G-GGG-------
-------------------------------GUGU--UGGCCGCCUGAGGGGAAGG-ACCGCCUAGUACGAG
AGGAA-CGGCGGUCCGCGGCCUC-UGGU-G-GACCGGCUGUCC-GGCA-GGGCA--A-GCCGGGUAGCUACG
CCGUA-UCCGAUAAGGGCUGAAAGCA-UC--CAGGCCCGAAGCGGUCCCCGA-A-AAU--AGGCGGCC-GU-
```

------------------------------------------------------------------------
------------------------------------------------------------------------
------------------------------------------------------------------------
------------------------------------------------------------------------
------------------------------------------------------------------------
------------------------------------------------------------------------
------------------------------------------------------------------------
------------------------------------------------------------------------
----------------------------------------------------------UG-AGG------CC
UCGGGUAAAAGACCCGGU-UGA-UGGGGC-GGG-GGUGUAAG-GCC-C--------AAG-----GGGCCGUU
GGGCCC--------CGAGGGCCUGAGCCC--A-CCGC-UCCCAAUC---GGCCGAUGG-ACCCGGC------
------------------------------------------------------------------------
------------------------------------------------------------------------
------------------------------------------------------------------------
-------------------------------------------------
>LSUa_69014_THEKO/1-3027
----------------------CCGG-UGGC----AA--CUAAGCCGCCU-GGUGGAUG-G-CUCGGC-UCG
GGG---CGCCGACGAAGGGCGUGG--CAAGCUGCGAUAAGCCCCGGCGAGGCGCAGGCAGCCGU------CG
AAC-CGG-GGAUGCCCGAAUGGGACC-U--------CCC-------GC----GGC---------------UU
U-U---------------------------G---CCG----C-A-CUCC-C---------------------
-----------A-----------GUC----------------------------------------------
-----------------------GG-G-AG-GGGGAACGCGGGGAAUUGAAACAUCUUAGUACCCG--CAGG
AAAAGAAAGCAA--AAGCGAUGCCGUUA--GU-AG-GGGCGACCGAAAGCGGC-A-CA-GGGCAAACUGAAC
CCCG-GGCCGA--------AA------------G---GUU-CGGGGGAUGUGGUG--UUG------------
U-AG--GGC-CCG-C-----G------GA-----------------GGAGC--CCU-----------GG--G
G-------GCGAA--GCCG------AAGUCC-GCUG------------------------------------
-------GAA-CG-CGGCG-------CCGGAGAGGGUGAAAGCCCCGUA---------GGC---G----UAA
G------CCCCCGAG-CUC-----------------CUGC-GGUGU--CCCUGAGUACCGUCGGUCG---GA
UAUCCGGCGGGAA-GCUGGGAGGCAUC--GGCUCCCAACCCUAAAUA--CGU-C-CCG-AGACCGAUAGC-G
AAC-U-AGUACCGUGAGGGAAAG-CUGAAA-AGCAC-CCCGG-GAGGGG-G-GUG-AAAAGAGCCUGAAACC
A-GGCGGCGAUAG-GAG-GG-UGCG---GCCCGAAA-GGGUUG-----------------------------
------------------------------------------------------------------------
------------------------------------------------------------------------
------------------------------------------------------------------------
------------------------------------------------------------------------
------------------------------------------------------------------------
------------------------------------------------------------AC-CCUCCCCG-
---AA-----------GG-A-------------------A------ACA-----------------CGG---
------------------------------------------------------------------------
--------------------------------------------------------------G--C-GA---
------------------------------------------------------------------------
-------------------------------------------------C-CG------UG--------GAG
UACGA-------------------------------------------------------------------
------------------------------------------------------------------------
------------------------------------------------------------------------

```
------------------------------------------------------------------------
------------------------------------------------G-----------------------
------------------------------------------------------------------------
------------------------------------------------------------------------
------------------------------------------------------------------------
------------------------------------------------------------------------
------------------------------------------------------------------------
-------------------GGGAGG-UG-----------------ACCGG--G---GUUGCACCGUCCGUCU
UGGAUCACGGGGCAGGGAGUUCA-C-CCGAGCGGCGA-GGUUAAGG--G-G--------GUCAA--------
C-CC--CGAA--GCCGCAGGGAAAC-CGA-----------------CAG----------------------G
UCCG-CAGCCCG----U-------------------------------------------------------
------------------------------------------------------------------------
----------------A--AGGG------------------------------------CGA-GGGACG-GG
GUGU-----------------------------------GAAA--------------------------GC-
G-C-CC----------------------GGAGUCGCUCGGUGA-GACCCGAAGCC-GGUCGAUCUAGCCCG
GGGCAGGGUGAAGUCCCU-CAACAG-AGGGAUGGAGGCCCGC-UAGGGGUGCUGACGUGCAAUUCGC-UCCC
GUGACCCCGGGCUAGGGGUGAAAGGCCAAUCGAGGCCGG-CGAUAGCUGGUUCCCGCCGAAU-UAUCCCCCA
GGAUAGCCCGGC-CGGA-------------------------------------------------------
----------------------GG-----UA-GGCAGUGGGGUAGAGC-ACUGAUU-GGG-GGUU-UAGGGG
G-C-GA-AA-GCCU-C-CGGCUCCCUGUCAAACUCCGAACCCACUG--------------------------
------------------------------------------------------------------------
------------------C-CG-C--CG-UAGA--UGGCCGG-AGUAGGG---GGGC-G-GUG-UAAGCC-G
UC-A-CCCGAGAG-GGGAAC---AACCCAGACCGG-G-GUUAAGGCCCCAAAGU--GC-CGG-CU-AA---G
-U-GUUACU--CCAAAGGGUGUCCCUGGCCUUAGACAGCGGGGAGGUAGGCUUAGAAGCAGCCA-UCCUUUA
AAGAGUGCGUAACAGCUCACCCGUC-----GAGGUCAGGGGCCCCGAAAAU-GGAC-GGGGCUUA-AGCCGG
C----------CGCCGAG-ACCCCGGCGCACGGA--C-----------------------------------
------------------------------------------------------------------------
-----CGAUUG-------------------------------------------------------------
-------------------------------GUCCGU-GAUC-GGGUAGGCGGGC-GUGCCGA----------
-----------U-G-GGGCG-GAAGCCG--G-GCCGUGAGGU--C-CGGUGGACCC-G-UCG-GUAUUGCGG
AUCC-UGCCGG-GAGUA-GCAGCA--UAGCC-GGGUGA--G---------------AAUCCC-GGC-CGCCG
U-AGGGGC-CAGGGUUCC-ACGGCAAUG-UUCGUCAGCCGUGGGUUAGUCGG--UC-CUAACCCGACCCGU-
AAC-UCGG---C-GUC--------------------------------------------------------
----------------------------------------GGG-GA-A-AGGGA-AACGGGUU-UAUA-UUC
CCGUA----------------------------------CCGCG----GUGGUAGG-------U--------
------------------------------------------------------------------------
-------GCGG--CAAC--------GC-AA-----GC--------------------CCGA-G-GGGU-GAC
GCCUCGG-------------GGUAGG-CGGACCGGU-C---------CACA-------------A-GGCCGG-
CUAA-G-CGU-AU-----A-A-------------------GC-----CC----------------GGGG-AG
U-G-CC--------------------GUAAU-GGCGAGAACC----------GGGUGA----AAG-CG-CGA-
AUG--GC-----------------------------------------------------------------
-----CUCCC---GUAAGGGGGGUU-C-CGC-C-G-AUCCCUGGGGCCC-GUGAAA-A-GCC-CCUCGGGAA
--------------------------------------------------------CGAUCC-AC-CGCGAC
CGUACCGAGAACCGAC------------------------------ACUGGUGC-CCCUGGGU-GAGAAGCC
UAAGGC-GUGU-CGGGGGAAACCCA-G-CCGAGGGAACUCGGC-AAAUUGGCCCCGUAACUUCGG-GAGAAG
```

```
GGGUG-CCUG-C-GG--GUG----------------------------------------------------
------------------------------------------------------------------------
------------------------------------------------------------------------
------------------------------------------------------------------------
------------------------------------------------------------------------
------------------------------------------------------------------------
------------------------------------------------------------------------
------------------------------------------------------------------------
------------------------------------------------------------------------
------------------------------------------------------------------------
------------------------------------------------------------------------
------------------------------------------------------------------------
------------------------------------------------------------------------
------------------------------------------------------------------------
--------------------------------------------------------------UG--UAACCC
GCA--GGUCGCAGUGACUAGGGGGGCCCGA-----CUGUUUAGUAAAAACACAGGUCCCAGCUAGCCCG---
-AA-A-------------GGGUUUGU-ACUGGGGCCGA-CGCCUGCCCAGUGCCG-G-UAUG-UGAAGCCCG
GG-------------UACA-------------ACCGGGUGAAGCAC--CGGUAAAC-GGCGGGGGUAACUAU
AACCCUCUUAAGGUAGCGAAAUUCCUUGUCGGU-UAAAUGCCGACCUGCAUGAAUGGCGUAACGAGG--UCC
CCACUGUCCCCGGC----UGG-GGCCCGGCGAAACC---UC---UGCCAGGCGCAUAUGCCUGGGACCUCCG
GUGGGAAGCGAAGA-CCCCAUGGAGC-UUUACU--GCAGCCU-GCCGUUGCCGUACGGCGGGGGGUGCGCAG
CGU-AGGCGGGAGGCGUCG----------AAGCCCG-CCC----------------------------UCCG
--------------------------------------GG-GCGGGUGGAG-C--C-GC-CCAUGAGACA-C
CGCC-CACCCUCUGCCGUACGGCUAACCC-GC---------GG-A-A-------------------------
------------------------------------------------------------------------
------------------------------------------------------------------------
------------------------------------------------------------------------
------------------------------------------------------------------------
------------------------------------------------------------------------
------------------------------------------------GCGG-----GGACA-GCGGUAGGU
GGGCAGUUUGGCUGGGGCGGCACACCCUCGA-AAAGGUAUC-GAGGGUGCCCU-AAGGUCGGCUCAGGCGGG
UCAGGAAUCCGCCGUA-GAGUG-CAAGGGCAA-AAGCCGGCCUGACU-GGAC--------------------
--------------CCGUAACAGAGGCGGGUCCAGCCGCGAAA-GCGUGGCCUAGCGAACCCCU--------
---GU-G-----CCUCC-CCG--------------------GU-GGGGGCCAGGGAUGACAG-AA--AAGCU
ACCCUGGGG-AUAACAGAGUCGUCUCGGGCGAGAGCCCAUAUCGACCCCGAGGCUUGCUACCUCGCUGUCGG
CUCUUCCCAUCCUGGCCCUGCA-GCAGGGGCCAAGGGUGGGGGUGUUCACCCAUUAAAGGGGAACGUGAGCU
GGGUUUAGACCGUCGUGAGACAGGUCGGAUGCUAUCUACC--------------------G-GAG-------
-------------------------------GUGU--UGGCCGCCUGAGGGGAAGG-CUCCCCCAGUACGAG
AGGAA-CAGGGAGCCGCGGCCUC-UGGU-C-UACCGGUUGUCC-UACA-GGGCAU-A-GCCGGGCAGCUACG
CCGCA-GCCGAUAAGGCCUGAAAGCA-UC--UAAGGCCGAAGCGGCCCCCGA-A-AAC--AGGCGGCC-AC-
--UCCUGGGCG-------------------------------------------------------------
-----------------------------------------UUGGGCC------------------------
------------------------------------------------------------------------
---------------------UGGGCGACCGGU---------------------------------------
------------------------------------------------------------------------
```

```
------------------------------------------------------------------------
------------------------------------------------------------------------
------------------------------------------------------------------------
-----------------------------------------CCG-UU-GCCCAGGA--CG-AGG------GC
CCGGGAAGAAGACCCGGU-UGA-UGGGGC-GGG-GAUGUAAG-CGG-G--------AAG-----GG------
--AAAC--------CGACCCGUUCAGUCU--G-CCGC-UCCCAACA---GCCCGAGUG-CUACCGGGC----
------------------------------------------------------------------------
------------------------------------------------------------------------
------------------------------------------------------------------------
-------------------------------------------------
>LSUa_243232_METJA/1-2989
-----------------------------------------AGCCCACCU-GGUGGAUG-G-CUCGGC-UCG
GGG---CGCCGAGGAAGGGCGUGG--CAAGCUGCGAUAAGCCCGGGGGAGGCGCAGGCAGCCGU------GG
AAC-CCG-GGAUCCCCGAAUGGGACU-U--------CCU-------GC----CCC---------------AU
U-U---------------------------G---GGG----C-G-CUCC-C-G-------------------
-----------U-----------UAG----------------------------------------------
------------------------G-G-AG-CGGGAACGCGGGGAAAAGAAGCAUCCGAGUACCCG--CAGG
AAAAGAAACCAA--CAGGGAUGCCGGGA--GU-AG-GGGCGACCGAAACCGGC-A-CA-GGGCAAACCGAAU
CCCU-AUCCGU--------AA------------G---GGU-AGGGAGAUGUGGAG--UUG------------
C-AG--GGC-CCC-C-----A------AU----------------AUAGAC--CCC-----------CA--C
U-------GGGAA--GCCG------AAGUCC-CCUG------------------------------------
-------GAA-UG-GGGCG-------CCAUAGAGGGUGAAAGCCCGUA----------GGC---G----UAA
C------CAGUUGGG-GGU-----------------CUGG-GGUGU--CCCUGAGUACCGCGCGUUG---GA
UAUCGCGCGGGAA-GCUGGGAGACAUU-AGGCUUCCAACCCUAAAUA--CGU-C-CCG-AGACCGAUAGC-G
AAC-U-AGUACCGUGAGGGAAAG-CUGAAA-AGCAC-CCCUU-GCGGGG-G-GUG-AAAAGAGCCUGAAACC
A-GGUGGGUACGG-AAU-GG-CACG---GCCCGAAA-GGU--------------------------------
------------------------------------------------------------------------
------------------------------------------------------------------------
------------------------------------------------------------------------
------------------------------------------------------------------------
------------------------------------------------------------------------
------------------------------------------------------------AA-CCACCCCG-
---AA-----------GG-A-------------------A------ACU-----------------CCC---
------------------------------------------------------------------------
--------------------------------------------------------------G--C-GA---
------------------------------------------------------------------------
-------------------------------------------------G-GG------AG--------GAG
UACGA-------------------------------------------------------------------
------------------------------------------------------------------------
------------------------------------------------------------------------
------------------------------------------------------------------------
------------------------------------------------G-----------------------
------------------------------------------------------------------------
------------------------------------------------------------------------
------------------------------------------------------------------------
```

------------------------------------------------------------------------
------------------------------------------------------------------------
-------------------GGGUGGCAU-----------------GCCGG--G---GUCGUGCCGUCCGUUU
CGAAAAACGGGCCGGGGAGUGUA-C-GGGUGUGGCGA-GCCUAAGG--G-G--------UUCAA--------
C-CC--CGGA--GGCGUAGGGAAAC-CGA-----------------CAU----------------------G
CCCG-CAACCCU----U-------------------------------------------------------
------------------------------------------------------------------------
----------------A--UGGG------------------------------------UGA-GGGGCG-GG
GUCU-----------------------------------UAAU--------------------------GG-
G-C-CC----------------------GGAGUCACACCCGUAC-GACCCGAAACC-GGGCGAUCUAGGCCG
GGGUAGGGUGAAGCCCCU-CGCCAG-AGGGGUGGAGGCCCGC--AGGGGUGUUACCGCGCAAAGUGC-UCCU
CUGACCCCGGUCUAGGGGUGAAAAGCCAAUCGAGCCCGG-AGAUAGCUGGUUCCCCCCGAAA-UAACUCGCA
GGUUAGCCGGGG-GUUA-------------------------------------------------------
----------------------GG-----UA-GAUGGCGGGGUAGAGCCACGGAUA-GGG-UGUU-UAGGGG
G-C-GA-GA-GCCC-U-CGGCACCCUGUCAAACUCCGAACCCGUCA--------------------------
------------------------------------------------------------------------
------------------U-CG-C--CG-UAGC--CCCCGAG---UGAGG-G-CAUA-C-GGG-UAAGCCGU
AU-G-UCCGAGAG-GGGAAC---AACCCGGACCCG-G-GUUAAGGCCCCUAAGU--GC-CGG-CU-AA---G
-U-GUAAAU--GAGAAGGGAGUCCCUGGCCUAAGACAGCGGGGAGGUUGGCUUAGAAGCAGCCA-UCCUUUA
AAGAGUGCGUAACAGCUCACCCGUC-----GAGGUCAGGGGCCCCGAAGAU--AAC-GGGGGCUA-AGCCGG
C----------CGCCGAG-ACCCGGGGGGGCUGA--------------------------------------
------------------------------------------------------------------------
------------------------------------------------------------------------
--------------------------------AAAGC-CAUC-CGGUAGGGGGGC-GUCCCGC---------
-----------G-G-GGGUA-GAAGCUC--G-GCCGUGAGGU--C-GGGUGGACCC-C-GUG-GGAACGAGA
AUCC-CGGCAG-UAGUA-ACAGCA--AAGUG-GGGUGA--G---------------AAUCCC-CAC-CGCCG
A-AGGGGC-CAGGUUUCC-ACAGCAACG-GUCGUCAGCUGUGGGUUAGCCGG--UC-CUAACCCCCGGGGU-
AAU-UCCC---U-GGG--------------------------------------------------------
----------------------------------------GGG-GA-A-AGGGA-AGCGGGUU-AAUA-UUC
CCGCG----------------------------------CCACC----GGGGUACG-------U--------
------------------------------------------------------------------------
-------GCGG--CAAC--------GC-AA-----GG--------------------CCAG-C-UCCU-GAC
GCUUCGG-------------GGUAGG-CCGACCACC-CC-------CGUC-------------G-GGGUGG-
CCAA-G-CGC-AU-----A-A-------------------GC-----CC----------------GGGG-AG
U-G-CC--------------------GUAAU-GGCGAGAACC---------GGGCAA----AAG-CG-UGA-
UGG--GC-----------------------------------------------------------------
-----CCUCC---GUUAGGAGGGUU-C-GGC-U-G-AGCCCUGGAGCCC-GUGAAA-A-GGG-AGCUGGCAA
--------------------------------------------------------GGAUCC-CC-GGUGAC
CGUACCCAGAACCGAC------------------------------ACAGGUGC-CCCUAGGC-GAGUAUCC
UAAGGC-GUGU-CGGG-AGAAUCCG-G-CCCAGGGAAGUCGGC-AAAUUGGCCCCGUAACUUCGG-GAGAAG
GGGUG-CCUG-C-GG--UCUU----------CUC-UA-----------------------------------
------------------------------------------------------------------------
------------------------------------------------------------------------
------------------------------------------------------------------------
------------------------------------------------------------------------

```
------------------------------------------------------------------------
------------------------------------------------------------------------
------------------------------------------------------------------------
------------------------------------------------------------------------
------------------------------------------------------------------------
------------------------------------------------------------------------
------------------------------------------------------------------------
------------------------------------------------------------------------
------------------------------------------------------------------------
----------------------------------------------------------AGUGAG--GGGACC
GCA--GGUCGCAGUGGCCAGGGGGGUCCGA-----CUGUUUAAUAAAAACACAGGUCUUGGCUAGCCCG---
-UA-A-------------GGGUGUGU-ACCAAGGCCGA-CGCCUGCCCAGUGCCG-G-UACG-UGAAACCCG
GG-------------UACA-------------ACCGGGCGAAGCGC--CGGUAAAC-GGCGGGGGUAACUAU
AACCCUCUUAAGGUAGCGAAAUUCCUUGUCGGG-UAAGUUCCGACCUGCAUGAAUGGCGUAACGAGA--CCC
CCACUGUCCCGGGC----CGG-AACCCGGUGAACCU--ACC---AUUCCGGUGCAAAGGCCGGAGACCCCCA
GUGGGAAGCGAAGA-CCCCGUGGAGC-UUUACU--GCAGCCU-GUCGUUGGGGCAUGGCCGUGGGUGCACAG
CGU-AGGUGGGAGCCGUCG----------AAGCCAC-CCC----------------------------UCCG
--------------------------------------GG-GGUGGUGGAG-G--C-GC-CCAUGGGACA-C
CACC-CACCCAUGGCCAUGUCCCUAACCC-C--------G-UA-A-A-------------------------
------------------------------------------------------------------------
------------------------------------------------------------------------
------------------------------------------------------------------------
------------------------------------------------------------------------
------------------------------------------------------------------------
-------------------------------------------------GGG-----GGACA-CCGGCAGGU
GGGCAGUUUGGCUGGGGCGGCACCCCCCUGA-AAAGGCAUC-AGGGGGGCCCA-AAGGUCGGCUCAGGCGGG
UCAGAACUCCGCCGUG-GAGUG-CAAGGGCAA-AAGCCGGCCUGACU-UGGU--------------------
--------------CGGUAAAAGAGGCCGACCAAGAGGCGAAA-GCCGGGCCUAGCGAACCCCU--------
---GU-G-----CCUCA-CCG--------------------AU-GGGGGCCAGGGAUAACAG-AA--AAGCU
ACCCCGGGG-AUAACAGAGUUGUCGCGGGCAAGAGCCCAUAUCGACCCCGCGGCUUGCUACAUCGAUGUCGG
UUCUUCCCAUCCUGGGCCUGCA-GCAGGGCCCAAGGGUGGGGCUGUUCGCCCAUUAAAGGGGAUCGUGAGCU
GGGUUUAAACCGUCGUGAGACAGGUUGGUUGCUAUCUGCU--------------------G-GGG-------
-------------------------------GUGU--UGGCCGCCUGAGGGGAAGG-UGGCUCUAGUACGAG
AGGAA-CGAGCCGCCGGCGCCUC-UGGU-C-UACCGGUUGUCC-GACA-GGGCAU-U-GCCGGGCAGCUACG
CGCUA-AGGGAUAAGGGCUGAAGGCA-UC--UAAGCCCGAAACCCUCCCCGA-A-AAU--AGGCGGCC-AG-
--UCCCU-----------------------------------------------------------------
------------------------------------------------------------------------
------------------------------------------------------------------------
------------------------------------------------------------------------
------------------------------------------------------------------------
------------------------------------------------------------------------
------------------------------------------------------------------------
------------------------------------------------------------------------
-------------------------------------------------UCGGGGA--CG-AGG------GC
UCUCCUAUAAGAGGAGGU-UGA-UAGGCC-GGG-GGUGUAAG-CGC-C--------GAG-----GGCU----
```

```
-UUGCC--------CGAGGCGUUCAGCCC--G-CCGG-UACUAAUC---GCCCAAGGG-CCCGGCAG-----
------------------------------------------------------------------------
------------------------------------------------------------------------
------------------------------------------------------------------------
-------------------------------------------------
>LSUa_419665_META3/1-2964
----------------------CCACAUAUC----UA--UUACCCUAUCU-GGGGGAUG-G-CUUGGC-UUG
GAA---CGCCUAUGAAGGACGUGG--UAAGCUGCGAUAAGCUCAUGCGAGGCGCAUACAGCCGU------AG
AAC-AUG-AGAUUUCCGAAUGGGACU-U--------CCU-------GC----UC----------------UU
U-U-------------------------------GAG----C-G-AUUC-----------------------
-----------G-----------UAA----------------------------------------------
------------------------G-A-AU-CGGGAACGCGGGGGAUUGAAGCAUCUUAGUACCCG--CAGG
AAGAGAAAUCAA--ACGAGAUUCCGUUA--GU-AG-AGGCGAUUGAACACGGA-A-CA-GGGCAAACUGAAU
CCCG----------------------------------CA-AGGGAGAUGUGGUG--UUA------------
U-AG--GAA-ACU-C-----U------U------------------ACGAC--UUA-----------GG--A
A-------AGGAU--GCCG------AAGUUG-UCUG------------------------------------
-------GAA-AG-GCACA-------UCAUAGAGGGUGAUAAUCCCGUA---------GGU---A----UAA
C------UGACCUGA-GU------------------UUGA-GAUUU--UCCUGAGUACCGUGCGUUG---GA
UAUCGCGCGGGAA-UUUGGGAGGCAUC--GACUUCCAACCCUAAAUA--CGU-U-CCA-AGACCGAUAGU-G
CAC-U-AGUACCGUGAGGGAAAG-CUGAAA-AGCAC-CCUUA-ACCGGG-U-GUG-AAAAGUGCCUGAAACC
A-GAUAGGUAUCG-UAU-GG-CAUG---GCUCCAAA-GGC--------------------------------
------------------------------------------------------------------------
------------------------------------------------------------------------
------------------------------------------------------------------------
------------------------------------------------------------------------
------------------------------------------------------------------------
------------------------------------------------------------AA-CUAUUCCG-
---AA-----------GG-A-------------------A------UCC-----------------AUU---
------------------------------------------------------------------------
--------------------------------------------------------------G--U-AA---
------------------------------------------------------------------------
-------------------------------------------------A-AU------GG--------UAG
UACGA-------------------------------------------------------------------
------------------------------------------------------------------------
------------------------------------------------------------------------
------------------------------------------------------------------------
------------------------------------------------G-----------------------
------------------------------------------------------------------------
------------------------------------------------------------------------
------------------------------------------------------------------------
------------------------------------------------------------------------
------------------------------------------------------------------------
-------------------GAAUAG-AG-----------------UCCAG--A---GUUGUGUCGUCCGUUU
CGAAAAACGGGCCGGGGAGUAUA-U-UGUUGUGGCGA-GCCUAAGA--C-C--------UUUAU--------
G-GU--CGGA--GGCGUAGGGAAAC-CAA-----------------CAA----------------------G
```

```
UCCG-CAAGCUU----U-------------------------------------------------------
------------------------------------------------------------------------
--------------------UGC------------------------------------UGA-GGGACG-GG
GUC------------------------------------UUAAG--------------------------G-
G-C-CC----------------------GGAGUCACAGCGAUAU-GACCCGAAACC-GGGCGAUCUAGGCCA
GGGCAAGGUGAAGUCCCU-CAUCAG-AGGGAUGGAGGCCUGC--AGAGGUGCUGCCGUUCGAAGCGC-UCUU
CUGACCUUGGUCUAGGGGUGAAAGGCCAAUCGAGCCCGG-AGAUAGCUGGUUCCCCCCGAAG-UAACCCUAA
GGUUAGCCGGAG-GUUA-------------------------------------------------------
----------------------GG-----UA-GAUGGCAGGGUAGAGC-ACUGAUA-GAG-GGGU-UAGGGG
A-C-GA-AA-GUCC-U-CGCCUUUCUGUCAAACUCCGAACCUGUUA--------------------------
------------------------------------------------------------------------
------------------U-CG-C--UG-CAGC--CUCUGAG-AA--AGG-G-CAUA-C-GGG-UAAGCUGU
AU-G-UCCGAGAC-GGGAAU---AGCCGCGACUCA-G-GUUAAGGUCCCUAAGU--AC-UGA-UU-AA---G
-U-GUGAAC--ACGAAGGGCGUCCUUGGCCUAAGACAGCAGGGAGGUUGGCUUAGAAGCAGCCA-UCCUUAA
AAGAGUGCGUAACAGCUCACCUGUC-----GAGGUCAGGGGCCCCGAAAAU-GGAC-GGGGCU-A-AAUCAG
U----------UACCGAG-ACCUGAGGGCACCGA--------------------------------------
------------------------------------------------------------------------
------------------------------------------------------------------------
--------------------------------AAGGU-GAUC-CGGUAGGGGGGC-GUUCUGU---------
-----------A-A-GGGCA-GAAGUUU--G-GCUGUGAAGU--C-AAAUGGACCU-U-GUA-GAAAUGAAG
AUCC-CGGUAG-UAGUA-ACAGCA-CAAGUG-GAGUGA--G---------------AAUCUC-CAC-CGCCG
A-AGGGGC-AAGGGUUUC-ACAGCAAUG-UUCGUCAGCUGUGAGUAAGCCGG--UC-CUAACUUCCGAGGU-
AAC-UCCU---U-UGG--------------------------------------------------------
----------------------------------------AAG-GA-A-AGGGA-GACAGGUU-AAUA-UUC
CUGUG----------------------------------CCAUC----UAAAUACA-------C--------
------------------------------------------------------------------------
-------GCGG--CAAC--------GC-AA-----GG--------------------UUAG-U-UUCU-GAC
GCUUCAG-------------GGUAUG-CUAACUACU-CU-------UGUC-------------U-GGGUAG-
UCAA-A-UGU-AU-----A-A-------------------GU-----CU----------------GGGG-AG
A-G-UU--------------------GUAAU-AACGAGAACC---------AGAUGA----AAG-CA-UGA-
UGA--UC-----------------------------------------------------------------
-----CAUCC---UUUAGGAUGGUU-U-GGC-A-G-AUCUCUGGAGCCC-GUGAAA-A-GGG-AACUAUCAU
--------------------------------------------------------GGAUUU-UA-GAUGAC
CGUACCCAGAACCGAC------------------------------ACUGGUGC-CCCUAGGU-GAGUAUCC
UAAGGC-GUAU-CGGA-UGAAUCUA-G-UCGAGGGAAGUCGGC-AAAAUGGCUCCGUAACUUCGG-GAGAAG
GAGUG-CCAG-U-GA--CCUUGUC---------------------------A--------------------
------------------------------------------------------------------------
------------------------------------------------------------------------
------------------------------------------------------------------------
------------------------------------------------------------------------
------------------------------------------------------------------------
------------------------------------------------------------------------
------------------------------------------------------------------------
------------------------------------------------------------------------
------------------------------------------------------------------------
```

```
------------------------------------------------------------------------
------------------------------------------------------------------------
------------------------------------------------------------------------
------------------------------------------------------------------------
------------------------------------------------------------------AGGGUC
GCU--GGUCGCAGUGACCAGGGAGGUCCGA-----CUGUUCAAUACAAACACAGGUCUUAGCUAGUCUG---
-AA-A-------------AGAUGUGU-ACUAAGGCCGA-CGCCUGCCCAGUGCUG-G-UACG-UGAACCCCG
GG-------------UUCA-------------ACCGGGCGAAGCGC--CAGUAAAC-GGCGGGGGUAACUAU
AACCCUCUUAAGGUAGCGAAAUUCCUUGUCGGG-UAAGUUCCGACCUGCAUGAAUGGCGUAACGAGA--CCU
CCACUGUCCCCGAC----UAG-AAUCCGGUGAACCU--ACC---AUUCCGGUGCAAAGGCCGGAGACUUCCA
GUGGGAAGCGAAGA-CCCCGUGGAGC-UUUACU--GCAGCCU-GUCGUUGGGGCAUGAUUGUGGGUGUACAG
UGU-AGGUGGGAGCCAUCG----------AAACCUU-CUC----------------------------GCUA
--------------------------------------GG-GAAGGUGGAG-G--C-GU-CCAUGGGACA-C
CACC-CUCCUAUGAUCAUGUCCCUUACCC-A----------AG-U-A-------------------------
------------------------------------------------------------------------
------------------------------------------------------------------------
------------------------------------------------------------------------
------------------------------------------------------------------------
------------------------------------------------------------------------
-------------------------------------------------UGG-----GGACA-CCGGUAGGU
GGGCAGUUUGGCUGGGGCGGUACCCUCCUAA-AAAUGCAUC-AGGAGGGCCCA-AAGGUUGGCUCAAGCGGG
UCAGGACUCCGCUGUU-GAGUG-UAAGGGCAA-AAGCCAGCCUGACU-UAGU--------------------
--------------UGUCCACAAUACGCAACUAAGAGACGAAA-GUCGGGCCUAACGAACCCCU--------
---GU-G-----CCUCA-CCG--------------------AU-GGAGGCCAGGGAUGACAG-AA--AAGCU
ACCCCGGGG-AUAACAGAGUUGUCGCGGGCAAGAGCCCAUAUCGACCCCGCGGCUUGCUACCUCGAUGUCGG
UUUUUCCCAUCCUGGAUCUGCA-GCAGGAUCCAAGGGUGGGGCUGUUCGCCCAUUAAAGGGGAUCAUGAGCU
GGGUUUAGACCGUCGUGAGACAGGUUGGUUGCUAUCUGCU--------------------G-GAA-------
-------------------------------GUGU--UGGCUGUCUGAGGGAAAGG-UGGUUCUAGUACGAG
AGGAA-CGAGCCGCCGGUGCCUC-UAGU-U-UAUCGGUUGUCC-GACA-GGGCAU-U-GCCGAGCCGCCACG
CACCA-AGGGAUAAGAGCUGAAAGCA-UC--UAAGCUCGAAGUCCAUCCUGA-A-AAC--AGACAGCC-GU-
--CCUUC-----------------------------------------------------------------
------------------------------------------------------------------------
------------------------------------------------------------------------
------------------------------------------------------------------------
------------------------------------------------------------------------
------------------------------------------------------------------------
------------------------------------------------------------------------
------------------------------------------------------------------------
-------------------------------------------------UAGGAGG--CG-AGG------GG
UCCCAUAGAAGAUGGGUU-UGA-UAGGCU-AGG-AAUGUAAG-CAU-C--------AAG-----GU------
--UUUC--------CGAGAUGUUUAGUUC--G-CUAG-UACUAACC---CCCCAAGAG-AUA----------
------------------------------------------------------------------------
------------------------------------------------------------------------
------------------------------------------------------------------------
-------------------------------------------------
```

```
>LSUa_1368239_BATHA/1-3058
C-------------------CG-ACAUG-----------CUACGCCGUCU-GGUGGAUG-A-CCAGGC-UUG
A-G---CGCUGAGGAAGGGCGCGG--CAAGCAGCGAAAUGUCCCGGGUAGCUGCAUGCAAGCUU------AG
AUC-CGG-GGGUGCCUGAAUGAGACU-U--------CUU-------GC----GACG-G-AC---------UC
C-C-------ACG---------------------GGA----C-U-CCGU-----------------------
-----------U-----------GCG----------------------------------------------
--UUCCGUA--------------AG-G-AA-CGGGAACUCCCUGAACGGAAACAUCUUAGUAAGGG--AAGG
AAAAGAAACCAA--CCGGGAUUCCCUAA--GU-AG-C-GUGAGCGAAAGGGGA-A-GA-GUCCAAACCGAAC
CCCG-CAGGGA--------AA------------C---CUG-AGGGGGAUGUAGUG--UUG------------
C-GG--GCC-CAG-CC------------------------------GCCUU--CUA-----------GC--C
G-------GCGAA--GCCG------AAGUGG-CCUG------------------------------------
-------GAA-AG-GCCUG-------CCAUAGAGGGUGAUAGCCGUUA----------GGC---G----UAA
A------CCGGAAGG-AGUUG---------------GGCUGGG-GU--CCCUGAGUACCAUACCCUG---GA
UUCGGUGUGGGAA-GCUGGAGGCCACC--AACCUCCAAGACUAAAUA--CGU-C-UCA-AGACCGAUAGC-G
CAC-U-AGUACGGUGACAGAAAG-CUGAAA-AGUAC-CCCGG-AAGGGG-G-GUG-AAAAGAGCCUGAAACC
A-GACGGUUACAG-ACG-UU-UGCA---GCCCGGAA-GGGUAG-----------------------------
------------------------------------------------------------------------
------------------------------------------------------------------------
------------------------------------------------------------------------
------------------------------------------------------------------------
------------------------------------------------------------------------
------------------------------------------------------------AU-CCCUCCUA-
---AA-----------GG-AACCUGCGGCGACGCAGCAGU--------------------------------
------------------------------------------------------------------------
------------------------------------------------------------------------
------------------------------------------------------------------------
------------------------------------------------------------------------
-ACGA-------------------------------------------------------------------
------------------------------------------------------------------------
------------------------------------------------------------------------
------------------------------------------------------------------------
------------------------------------------------G-----------------------
------------------------------------------------------------------------
------------------------------------------------------------------------
------------------------------------------------------------------------
------------------------------------------------------------------------
------------------------------------------------------------------------
-------------------GGAGGG-GG-----------------ACCAG--G---GUUGCAAGUUCCGUCU
AGAAACACGGGCCGGGGAGUUCA-A-GGUCGUGGCAA-GCCUAAAG--GCU--------GACAA--------
G-CC--UGCA--AGCAUAGGGAAAC-CAA-----------------CAU----------------------G
CGCG-CAA-CCA----G-------------------------------------------------------
---C--------------------------------------------------------------------
--------------CAAGGCUGG------------------------------------CGA-GGCGCG-GG
GUCU-----------------------------------GAAA--------------------------GG-
G-C-CC----------------------GGAGUCACGGCUUUGA-CACUAGAAACC-GGACGAUCUAGCCCU
```

GGGCAGGGCGAAGUCAGG-CGAAAG-CCUGGUGGAGGCCCGA--AGGGGUGCUGACGUGCAAUUCGU-UCCU
CUGACCUGGGGCUAGGGGCGAAAGACCAAUCUAGUCCGG-UGAUAGCUAGUUCCCUCCGAAG-UGGGUCGCA
GCCCAGCCCCAA-GCGA-------------------------------------------------------
----------------------GG-----UA-GCCGAUGGGGUAGAGC-ACUGAUU-GGG-AAGU-AAGGGG
U-C-GA-AA-GGCC-U-CGCUUUUCUUUCAAACUCCGAAUCCAUCG--------------------------
------------------------------------------------------------------------
------------------G-UG-C--CG-UAGA--CCUUGGG-AGACGGGUU-CUGG-G-GGG-UAA-GCCC
CC-AGUCCGAGAG-GGGGAC---AACCCAGACUGA-G-GUUAAGGCCCCAAAGU--GC-UGG-CU-AA---G
-U-GUCAAA--CAAAAAGGCGUCAUCAGCCCCAGACAGCGGGGAGGUAGGCUUAGAAGCAGCCA-CCCUUUA
AAGAAAGCGUAACAGCUCACCCGCC-----GAGGCUGAUGGCCCUGAAAAU-GGAC-GGGGCU-A-AGCCAG
C----------CGCCGAU-ACCCCAGGGUGCAGCU-U-----------------------------------
------------------------------------------------------------------------
-----AG-----------------------------------------------------------------
-------------------------------CGCUGC-AAUC-CGGUAGGAGGGC-GUCCCGA---------
-----------C-U-GGGUG-GAAGCCG--G-GCCGUGAGGU--C-CGGUGGACCG-GUUCG-GGAUUGCAU
AUCC-CGGUGG-UAGUA-ACAGCA--AAGAG-AAGUGA--G---------------AAUCUU-CUC-CGCCG
A-AAGGGC-AAGGGUUCC-CCAGCAAUGCGUCGUCAGCUGGGGGUAAGUCGG--UC-CUAAGGCCGAUCUU-
AAC-UGAG---G-AG---------------------------------------------------------
----------------------------------------GCC-GA-A-AGGGA-AGCCGGUU-AAGA-UUC
CGGCA----------------------------------CCACG----GAGGUACA-------C--------
------------------------------------------------------------------------
-------GCGG--CGAC--------GC-AA-----GC--------------------CCAU-C-AUCU-GAC
GCUUCGG-------------GAUAGG-CUGAGCAGG-GC-------UGUC-------------G-CCUUGU-
UUAA-C-CGU-AU-----A-A-------------------GC-----CU----------------GGGG-AG
U-G-CC--------------------GUAAU-GGCGAGAACC---------AGGCGA----AGG-CG-GGA-
AUA--GC-----------------------------------------------------------------
-----CGCCCGUUU-UAGGGCGGUU-C-AGC-U-G-AUUUCUGGAGCCC-UUGAAA-A-GGA-UGUUGGGAA
--------------------------------------------------------GGAUCC-UC-UGUGAC
CGUACCCAGAUCCGAC------------------------------ACAGGUGU-CCCUAGGC-GAGGAACC
UAAGGC-GUGU-CGGGUGCACACCG-A-GCGAGGGAACUCGGC-AAAUUAGCCCCGUAACUUCGG-GAUAAG
GGGUG-CCUG-C-GG--UUUU--------------------------GG-CU--------------------
------------------------------------------------------------------------
------------------------------------------------------------------------
------------------------------------------------------------------------
------------------------------------------------------------------------
------------------------------------------------------------------------
------------------------------------------------------------------------
------------------------------------------------------------------------
------------------------------------------------------------------------
------------------------------------------------------------------------
------------------------------------------------------------------------
------------------------------------------------------------------------
------------------------------------------------------------------------
------------------------------------------------------------------------
----------------------------------------------------------UCGGCU--GGGACU

```
GCA--GGUCGCAGUGGCAAAGGGGUCUCGA-----CUGUUUAAUAAAAACACAGGAAGCCGCAAGCCCG---
-AA-A-------------GGGUGUGA-ACGGCUUCUGA-AUCCUGGCCAGUACAG-G-UACC-UGAAACCCG
GU-------------UUCA-------------ACCGGAGCAAGGGC--CUGCAAAC-GCCGGGAGUAACUCU
GACUCUCUCAAGGUAGCCAAAUGCCUUGUCGGG-UAAGUUCCGACGCGCAUGAAUGGAGCAACGAGG--ACC
CCGCUGUCCCCGCU----UGG-UACCCGGUGAACCC---ACGUAAUGUGGACGAACAGUCCACGGACUCCCA
GCGGGAAAAGAAGA-CCCCGUGGAGU-UUUACU--GCAGCCU-GUCGCUGGGAUACGGCUGCGGAUGCAGAG
UGU-AGGCGGGAGCCUUUA----------AUCCCGGUCUC----------------------------UCCG
--------------------------------------GGGGCUGGGGGAG-G--C-GC-CCAUGGAACA-C
CGCC-CUUUCGCAUCUGUGUCUCUUACCG-GC--G------CU-U-C-------------------------
------------------------------------------------------------------------
------------------------------------------------------------------------
------------------------------------------------------------------------
------------------------------------------------------------------------
-------------------------------------------------------G----GC----------
------------------------------------------------GCCG-----GGACA-ACGGUUGGU
GGGCAGUUUGGCUGGGGCGGCACCCCCUUGA-AAAAAUAUC-GAGGGGGCCCA-AAGGUCGGCUCAGGCGGG
ACAGAAACCCGCCGCA-GAGUG-CAAAGGCAA-AAGCCGGUCUGACU-GGAU--------------------
--------------CCUCCAAGGCAAGGGAUCCAGAGGCGAAA-GCCUGGCUUAGCGAACAUCC--------
---AG-A-----UCCUCUUUG--------------------GU-GGGGGUUGGAUAUGUCAG-AA--AAAUU
ACCCCGGGG-GUAACAGGCUUGUCGCGGGCGAGAGUUCACAUCGACCCCGCGGCUUGGUACCUCGAUGUCGG
CUCUUCCUAUCCUGGUCCUGCA-UAAGGGGCCAAGGGUGAGGCUGCUCGCCUAUUAAAAGGGAACGUGAGCU
GGGUUUAGACCGUCGUGAGACAGGUCGGACUUUAUCUGCU--------------------G-GGA-------
-------------------------------GUGC--UGGUCGCCUGAGGGGAAAG-GGUUCCCAGUACGAG
AGGAA-CGGAACGUUGUAGCCCA-UAGU-C-UACCGGUUGUUC-GACA-GGGCAC-U-GCCGGGCAGCCACG
CUGUC-GCGGAUAAGAGCUGAAGGCA-UC--UAAGCUCGAAGCCGCUCCCGA-A-AAU--AGGCGGCC-AU-
--UCCAUG-A-CA-C---------------------------------------------------------
------------------------------------------------------------------------
------------------------------------------------------------------------
----------------GAGACUUGG-CGACAGG--CCUC-A--G-UCAU-----------------------
------------------------------------------------------------------------
------------------------------------------------------------------------
------------------------------------------------------------------------
------------------------------------------------------------------------
-------------------------------------------------------GGACG-AGG------GC
UCCCAUAGAAGAUGGGUU-UGA-UGGAGC-CGG-GGUGCACG-CAU-C--------AAG-----GC------
-UUUGG-------CCGAGGUGUUUAGCCC--G-CGGC-CCCCAACC---GCCCGAGG--UGUCGGG------
------------------------------------------------------------------------
------------------------------------------------------------------------
------------------------------------------------------------------------
-------------------------------------------------
>LSUa_224325_ARCFU/1-2932
---------------------------CGGC----GA--AUAUCCCGGCC-GGUGGAUG-G-CUCGGC-UCG
G-G---CGCCGACGAAGGGCGUGC--CAAGCUGCGAAAAGCCCGGGGGAGGCGCAAGGAGCCUU------AG
AAC-CCG-GGAUUCCCGAAUGGGAUA-U--------CCU-------GU----CCC---------------UU
C-G---------------------------G---GGA----C-G-CUCC-C---------------------
```

```
-----------U-----------UAC----------------------------------------------
-----------------------GG-G-AG-CGGGAACGCGGGGAAAGGAAACAUCUGUGUACCCG--CAGG
AAAAGAAAGCAA--ACGCGAUGCCGUGA--GU-AG-GGGCGACCGAAAGCGGC-A-CA-GCCCAAACCGAAC
AUCC-GGGGGU--------AA------------C---UCC-GGAUGGAUGUGGUG--UUG-------------
U-AG--GAC-CCU-G-----C------C------------------UAAUG--CCG-----------UG--G
A-------ACGAA--GCCG------AAGUGG-UCUG------------------------------------
-------GAA-CG-GCCCG--------CCGUAGAGGGUGAUAGCCCGUA----------GGC---G----UAA
G------UUCCACGG-UUACAA--------------GGCG-GGGUA--UCCUGAGUACCGUGGGUUG---GA
AUUCUCGCGGGAA-GCUGGGGGUCAUC--AACCCCCAAGGCUAAAUA--CGU-C-CCG-AGUCCGAUAGC-G
CAA-U-AGUAGGGUGACCGAAAG-CUGAAA-AGAAC-CCCGAGAAGGGG-A-GUG-AAAAGAGCCUGAAAUC
G-GCCGGGGAUAG-UGA-GC-AGUG---GCUCGAAA-GGU--------------------------------
------------------------------------------------------------------------
------------------------------------------------------------------------
------------------------------------------------------------------------
------------------------------------------------------------------------
------------------------------------------------------------------------
------------------------------------------------------------AA-UCCCGGCA-
---AA-----------GG-A-------------------A------AGU-----------------GCU---
------------------------------------------------------------------------
--------------------------------------------------------------G--A-GA---
------------------------------------------------------------------------
-------------------------------------------------G-GC------GC--------GAG
UACGA-------------------------------------------------------------------
------------------------------------------------------------------------
------------------------------------------------------------------------
------------------------------------------------------------------------
------------------------------------------------G-----------------------
------------------------------------------------------------------------
------------------------------------------------------------------------
------------------------------------------------------------------------
------------------------------------------------------------------------
------------------------------------------------------------------------
-------------------CCGGGACAU-----------------GCCGG--A---GUCACUGCGUACGUUC
UGAAGAACGGGCCAGGGAGUGUA-C-GGCAGGGGCGA-GGCUAAGG--A-G--------UGAAA--------
C-UC--CGAA--GCCGGAGGGAAAC-CGA-----------------CA-----------------------G
CCCG-CAGCUU-------------------------------------------------------------
------------------------------------------------------------------------
--------------------AUG------------------------------------CGA-GGGGCG-GG
GUG------------------------------------UGCU---------------------------C-
G-C-CU----------------------GUAGUCCCUGCCGUAC-GACCCGAAGCC-GGGCGAUCUAGGCGG
GGGCAGGGUGAAGCGGUG-CGAAAG-CGCCGUGGAGGCCCGC--AGGGGUGUUGAUGUGCAAAUCGC-UCCU
CUGACCCCCGUCUAGGGGUGAAAGUCCAAUCGAGCCCGG-GGAUAGCUGGUUCCCCCCGAGA-CAUACCGCA
GUAUGACCCGUC-CGGA-------------------------------------------------------
----------------------GG-----UU-GGCGGUGGGGUAGAGC-ACUGAUA-GGA-GGAU-CCGGGG
G-G-UG-AC-ACCC-U-CGCCCUCCUGUCAAACUCCGAAUCCACCG--------------------------
```

```
------------------------------------------------------------------------
------------------C-CG-C--CG-UAGA--UGGGCGG-AGUCAGG-G-GCAG-G-GGG-UAAGCUCC
UG-U-UCCGAGAG-GGAGAC---AACCCAGACCGG-G-GUUAAGGCCCCUAAGU--GC-CGG-CU-AA---G
-U-GUAAACG-AUAAAGGAGGUCCCGGGUCGAAGACAGCGGGGAGGUAGGCUUAGAAGCAGCCA-CCCUUUA
AAGAGUGCGUAACAGCUCACCCGCC-----GAGACUCGGGGCGCCGAAAAU-GGAC-GGGGCUCA-AGCCGG
C----------CGCCGAU-ACCCCGGGGCUCCGA--------------------------------------
------------------------------------------------------------------------
------------------------------------------------------------------------
--------------------------------AAGGA-GAUC-CGGUAGGGGGGC-GUGCCGA---------
-----------U-G-GCGCAGAAGCCUG--A-GGCGUAAGCU--C-GGGUGGAGCC-G-UCG-GUAACGAAU
AUCC-UGGCGG-UAGUA-GCAGCA--UAGUC-GGGUGA--G---------------AAUCCC-GAC-CGCCG
G-AAGGGC-AAGGGUUCC-ACGGCAAUG-UUCGUCAGCCGUGGGUGAGUCGG--UC-CUAAGGUGGCCCGU-
AAC-UCGG---C-GCC--------------------------------------------------------
----------------------------------------GCC-GA-AAAGGGA-AGCCGGUU-AAUA-UUC
CGGCA----------------------------------CCGCU----GCCC--------G-CU--------
------------------------------------------------------------------------
-------GCCUU------------------------U--------------------AUGG-C-ACCC-GAC
GCCUCGG-------------GAUAGG-CCGGGCGGGAGC-------UGCC-------------A-GCCCGU-
CCAA-G-CGU-CG-----A-A-------------------GC-----CC----------------CCGG-AG
A-U-CC--------------------GUAAU-GGAGAGAAGG---------GGGUGA----AGG-CG-UGA-
UG---GC-----------------------------------------------------------------
----------------G-UA-A-GC-C-GGC-C-G-ACUCCUGGGGCCC-GUGAAA-A-GGG-GGGC-----
---------------------------------------------------------------AG-UUGGAC
CGUACCGAGAACCGAC------------------------------ACAGGUG--CCCUGGGU-GAGUAGCC
UAAGGC-GUGG-CGGA-ACAUUCCG-G-CCGAGGGAAUUCGGC-AAGUUGGCCCCGUAUCUUUGG-UAGAAG
GGGUG-CCUG-C-GG--UUU------------------------AG--------------------------
------------------------------------------------------------------------
------------------------------------------------------------------------
------------------------------------------------------------------------
------------------------------------------------------------------------
------------------------------------------------------------------------
------------------------------------------------------------------------
------------------------------------------------------------------------
------------------------------------------------------------------------
------------------------------------------------------------------------
------------------------------------------------------------------------
------------------------------------------------------------------------
------------------------------------------------------------------------
------------------------------------------------------------------------
------------------------------------------------------------------CAGACC
GCA--GGUCGCAGUGACUAGGGGGGAGCGA-----CUGUUUACUAAAAACACAGGGGGCUGCAAUCCGG---
-CA-A-------------CGGUUUGU-ACAGCCCCUGA-GUCCUGCCCAGUGCGG-G-UACC-UGAAACCAG
GG-------------UACA-------------ACCGGGCGAAGGGC--CCGUAAAC-GGCGGGGGUAACUAU
GACCCUCUUAAGGUAGCGUAAUACCUUGCCGCU-UAAUUGGCGGCUUGCAUGAACGGAUUAACGACU--CCC
CCACUGUCCCCGGC----CGG-AAGCCGGCGAACCC--GAC---AUCCCAGUGAAGAGUCUGGGGACCCCCG
```

```
UUGGGAAGCGAAGA-CCCUGUGGAGC-UUUACU--GCAGCCU-GCCGUUGCUCCACAGCUGGGGGUGCGCAG
GGU-AGGCGGAGGCUUCG-----------AAGCCCC-GUC----------------------------UCCG
--------------------------------------GG-CGGGGUGGAG-C--C-GU-CGAUGAGACA-C
CGCC-CACCUCCAGCUGUGGAGCUAACUC-C--------C-GC-U-G-------------------------
------------------------------------------------------------------------
------------------------------------------------------------------------
------------------------------------------------------------------------
------------------------------------------------------------------------
------------------------------------------------------------------------
-------------------------------------------------GGA-----GGACA-UCGGUAGGU
GGGCAGUUUGGGUGGGGCGCCACUCCCCCGA-AAAGGUAUC-GGGGGAGCCCA-AAGGUCGGCUCAGGCGGG
UCAGAAAUCCGCCGUA-GAGUG-CAAGGGCAA-AAGCCGGCCUGACG-UGAA--------------------
--------------CCCGCAAAAUAAGGGUUCACGAGCCGAAA-GGCGGGCCUAGCGAACCACC--------
---CU-G-----GCCUU-UUG--------------------GU-GAGGCCGGGUGACGACAG-AA--AAGCU
ACCCCAGGG-AUAACAGAGUUGUCCCCGGCGAGAGUACAUAUCGACCCGGGGGCUUGCUACCUCGAUGUCGG
CUCUCCCCAUCCUGGCCGUGCA-GCAGCGGCCAAGGGUGAGGUUGUUCGCCUAUUAAAGGGGAUCGUGAGCU
GGGUUUAGACCGUCGUGAGACAGGUCGGUUGCUAUCUAAC--------------------G-GGG-------
-------------------------------GUGUC-AGGGCGGCUGAGGGGAAGG-AGGCUCUAGUACGAG
AGGAA-CGAGCCUCCGGCGCCAC-UGGU-U-UACCGGUUGUCC-GGCA-GGGCAU-C-GCCGGGCAGCUACG
CGCCACGCGGUUAAAGGCUGAAAGCA-UC--UAAGCCUGAAACCGCCCCCGA-A-AAG--AGCCGCCC-AU-
------------------------------------------------------------------------
------------------------------------------------------------------------
------------------------------------------------------------------------
------------------------------------------------------------------------
------------------------------------------------------------------------
------------------------------------------------------------------------
------------------------------------------------------------------------
------------------------------------------------------------------------
----------------------------------------------------------UA-AGG------GC
UCCCGUAAAAGACGGGGU-UGA-UAGGAC-CGG-GGUGUAAG-CGG-C--------GAG-----CU------
--UCGG--------CGAGCCGUUCAGCCC--G-CGGU-CACUAAUC---GCCCGC-GC-CG-----------
------------------------------------------------------------------------
------------------------------------------------------------------------
------------------------------------------------------------------------
-------------------------------------------------
>LSUa_259564_METBU/1-2917
----------------------ACUC-UGGU----AA--UUAUGCCGUCA-GGUGGAUG-G-CUCGGC-UCA
A-G---CGCUGAUGAAGGACGUGC--CAAGCUGCGAUAAGCUCAGGGUAGGUGCAUGGAACCUA------UG
AAC-CUG-AGAUUUCCUAAUGAGACC-U--------CUU-------AG----CAC---------------AU
U-U---------------------------G---UGC----U-G-AUCA-----------------------
-----------G-----------UAA----------------------------------------------
------------------------U-G-AU-CGGGAACGCCCCGAAUUGAAACAUCUUAGUAGGGG--CAGG
AAGAGAAAUCGA---AGAGAUGCCGUUA--GU-AA-UGGCGAAUGAACACGGC-A-CA-GUUCAAACUGAAU
CCCU--CCAGU--------AA------------U---GUG-AGGGAGAUGUGGUG--UUA------------
U-AG--GAC-UUU-C-----U------AA----------------GAUUCC--UAU-----------CU--U
```

```
U-------UCUUA--GCUG------AACUUU-UCUG------------------------------------
-------GAA-CG-UUAGA-------CCAUAGAGUGUGAUAGUCACGUA---------AGC---A----AUG
A------AAAGAGGA-AU------------------UGGA-AAUGU--UCCUGAGUAUCGUGUGUUG---GA
AAUCACGCGAGAA-UUUGGGAGGCACC--AACUUCCAAAACUAAAUA--CGU-C-UUG-AGACCGAUAGC-G
AAA-U-AGUAGGGUGACCGAAAA-CUGAAA-AGUAC-CCCCAAAAGGGA-G-GUU-AAAAGUGCCUGAAACC
U-GAUGGUGAUGG-AGC-GA-UAUG---GCAUGAAA-GGAU-------------------------------
------------------------------------------------------------------------
------------------------------------------------------------------------
------------------------------------------------------------------------
------------------------------------------------------------------------
------------------------------------------------------------------------
------------------------------------------------------------CU-UUAGUUCG-
---AA-----------GG-A-------------------A------UCA-----------------ACC---
------------------------------------------------------------------------
--------------------------------------------------------------G--C-GA---
------------------------------------------------------------------------
-------------------------------------------------G-GU------UG--------UAG
UACGA-------------------------------------------------------------------
------------------------------------------------------------------------
------------------------------------------------------------------------
------------------------------------------------------------------------
------------------------------------------------G-----------------------
------------------------------------------------------------------------
------------------------------------------------------------------------
------------------------------------------------------------------------
------------------------------------------------------------------------
------------------------------------------------------------------------
-------------------UUCUAA--U-----------------GCCGA--U---GUCAUAUCAUACGUUU
UGAAGAACGGGCCAGGGAGUGUA-A-CUAAGUGGCGAUGGCUAACC--U-U---------UUAU--------
C-UG--GGAA--GCCGAAGCGAAAG-CAA-----------------CAU----------------------G
CGUG-CAACCUU----U-------------------------------------------------------
------------------------------------------------------------------------
----------------A--CGGG------------------------------------UGA-GACGCG-GC
GUAU-----------------------------------ACAA--------------------------GU-
G-C-GU----------------------GGAGUCACUAAGUUAC-GACCCGAAGCC-GGGUGAUCUAGGCGU
GGGCAGGUUGAAGCGUGG-CGAAAG-CUACGUGGAGGACCGC-AAGCGGUAUUGAUCUGCAAAUCAU-UCGU
GUGACCUGCGUCUCGGAGUGAAAGGCUAAUCUAACCCGG-CAUCAGCUGGUUCCUUCCGAAA-CAUGUCGUA
GCAUGACCUAAC-UGGA-------------------------------------------------------
----------------------GA-----UA-GUCGGUGGAGUAGAGC-ACUGAUU-GAU-GGCC-CUGGGG
A-A-GA-AA-UUCC-U-CAUCCGUCUGUCAAACUCCGAACCCACCG--------------------------
------------------------------------------------------------------------
------------------U-CG-U--CG-UAGA--AGGUUGG-AGUCCGG-A-CUACUG-GGG-UAAGCCUG
UA-G-UCCGUAAG-GGAGAC---AACCCAGCCCGU-G-GUUAAGGUCCCUAAGU--GU-CGA-CU-AA---G
-U-GUUAAUA-CUAAAGGGCGUCCCAAGCCCUAGACAGCUGGAAGGUUAGCUUAGAAGCAGCUA-UCCUUUA
AAGAGUGCGUAACAGCCCACCAGUC-----GAGGUUUGGGGCCCCGAAAAU-GGAC-GGGGCUCA-AGUCGA
```

```
C----------CACCGAU-ACCACGGAGUACCGU--------------------------------------
------------------------------------------------------------------------
------------------------------------------------------------------------
--------------------------------AAGGU-AAUC-UCGUAGGAAGGC-GUUGCGU---------
-----------U-U-GGGUA-GAAGCAG--G-GCUGUGAAGU--C-CUGUGGACCG-A-GCG-GAAACGAAA
AUCC-UGGUAA-UAGUA-GCAGCA--UAGUU-AGGUGA--G---------------AAUCCU-AAC-CGCCG
A-AGGGGC-UAGGUUUCC-UCGGCAAUG-AUCGUCAGCCGAGGGUUAGUCGG--UC-CUAAGUUGUACCGU-
AAU-UCGA---G-UAC--------------------------------------------------------
----------------------------------------AUC-AA-A-AGGGA-AACAGGUU-AAUA-UUC
CUGUA----------------------------------CUAUU----CAAC--------------------
------------------------------------------------------------------------
---------------------------------------------------------AAUA-A-AACU-GAC
GUUUUGG-------------GGCAGG-CUGAGCGGC-GC-------UGUC-------------G-CGCCGU-
UUAA-G-CAU-CU-----A-A-------------------AU-----CU----------------GUGG-AG
A-G-CC--------------------GUAAU-GGCGAGAAGC----------AGACGA----AUA-UG-UUA-
UG---AC-----------------------------------------------------------------
----------------G-AA-A-GU-C-GGC-U-G-CACCCCGGAACCC-GUGAAA-A-GGG-AGU------
---------------------------------------------------------------UG-AAUAUC
CGUACCGAGAUCUGAC------------------------------ACAGGUGC-CCCUAGCU-GAAAAGGC
UAAGGC-GUGU-CGGA-AUAAUUUG-G-CUAAGGGAAUUCGGC-AAAUUGGCUCCGUAACUUCGG-GAGAAG
GAGUG-CCUG-C-CG--UGA----------------------------------------------------
------------------------------------------------------------------------
------------------------------------------------------------------------
------------------------------------------------------------------------
------------------------------------------------------------------------
------------------------------------------------------------------------
------------------------------------------------------------------------
------------------------------------------------------------------------
------------------------------------------------------------------------
------------------------------------------------------------------------
------------------------------------------------------------------------
------------------------------------------------------------------------
------------------------------------------------------------------------
------------------------------------------------------------------------
-------------------------------------------------------------------AGACG
GCA--GGUCGCAGUGACCAGGGAACUCUAA-----CUGUCUAAUAUCAACAUAGGAGAUCGCAAACCCG---
-UA-A-------------GGGCUAGU-ACGAUCUCUGA-AUCCUGCUCAGUGCAG-G-UACC-UGAAACUUC
GG-------------UUCA-------------ACGGAAUGAAGGGC--CUGUAAAC-AGCGGGGGUAACUAU
GACCCUCUUAAGGUAGCGUAGUACCUUGUCGCU-UAAUUGGCGACUUGCAUGAAUGGAUCAAUGAGA--GUU
CUACUGUCCCUAGC----CAG-AGUCCGGUGAAGCU--UAC---AUUCUAGUGCAGAGUCUAGAGACCUCUA
GGGGGAAGUGAAGA-CCCCGUGGAGC-UUUACU--GCAGCCU-GUCGCUGGGUUGUGAUUUUGGAUGUACAG
UGU-AGGUAGGAGACGUCG----------AAGCCGG-UGC----------------------------GCCA
--------------------------------------GC-AUCGGUGGAG-U--C-GC-AAUUGGGACA-C
UACC-CUUCCAAAGUUACGACCCUCACUC-C----------GU-G-A-------------------------
------------------------------------------------------------------------
```

```
------------------------------------------------------------------------
------------------------------------------------------------------------
------------------------------------------------------------------------
------------------------------------------------------------------------
-------------------------------------------------GGA-----GGACC-CCGAUAGGU
GGGCAGUUUGCCUGGGGCGGGACGCCCUUGA-AAAGAUAUC-AAGGGCGCGCA-AUGGUUAACUCAAGUGGG
UCGGAAACCUACUGAA-GAGUG-CAAGAGCAU-AAGUUAGCCUGACG-UGAU--------------------
--------------UCAGCACAGUAGUGGAUCACGAGACGAAA-GUCGGUUCUAGCGAACCUUU--------
---AU-G-----CCCGC-UUG--------------------GU-GCGGGCUAAAGAUGACAG-AA--AAGUU
ACCCCGGGG-AUAAUUGAGUCGUUGCCGGCAAGAGCACAUAUCGACCCGGCAGCUUGCUACCUCGAUGUCGG
UUCUUUCCAUCCUGGCUGUGCA-GCAGCAGCCAAGGGUGAGGUUGUUCGCCUAUUAAAGGAGAUCGUGAGCU
GGGUUUAGACCGUCGUGAGACAGGUCGGUUACUAUCUACU--------------------A-GAG-------
-------------------------------GUGU--AUGUGGUCUGAGGGUAAGU-UGCUUUUAGUACGAG
AGGAA-CAGAGCAACGGCGCCAC-UGGU-C-GAUCGGUUGUCU-GACA-AGGCAC-U-GCCGAGCAGCUACG
CGCUA-AGGAAUAAGAGCUGAAUGCA-UC--UAAGCUCGAAAUCUAACCUAA-A-A-A--GAGACCAC-UU-
------------------------------------------------------------------------
------------------------------------------------------------------------
------------------------------------------------------------------------
------------------------------------------------------------------------
------------------------------------------------------------------------
------------------------------------------------------------------------
------------------------------------------------------------------------
------------------------------------------------------------------------
----------------------------------------------------------UA-AGG------GU
UCCGGUAAAAGACCGGAU-CGA-UAGGAA-UGG-GAUGUAAG-CAC-C--------AAG-----GC------
----AA--------CGAGGUGUUCAGUCC--G-CAUU-UACUAAU----AUCCGUGCC-UC-----------
------------------------------------------------------------------------
------------------------------------------------------------------------
------------------------------------------------------------------------
-------------------------------------------------
>LSUa_188937_METAC/1-2899
----------------------------AGG----AA--UUAUGCUAUCA-GGUGGAUG-G-CUCGGC-UCA
A-G---AGCUUAUGAAGGACGUGC--CAAGCUGCGAUAAGCCCGGGGUAGGUGCAUGGAGCCAA------UG
AAC-CCG-GGAUUUCCGAAUGGGACC-U--------CU--------CC-----------------------G
U-A---------------------------------G----U-G-AUCA-----------------------
-----------U-----------UCA----------------------------------------------
------------------------U-G-AU-CGGGAACGCUCCGAAUUGAAACAUCUCAGUAGGAG--CGGG
AAAAGAAAUCAA--UUGAGAUGCCGUAA--GU-AA-CGGCGAGUGAAACCGGC-A-CA-GUUCAAACCGAAU
CCCU----------------------------------UC-GGGGACAUGUGGUG--UUG------------
U-AG--GGC-UGU-U-----C-------------------------AAACG--UCU-----------GG--C
G-------GUAAA--AUCG------AACUUG-CCUG------------------------------------
-------GAA-CG-GCAGA-------CCAUAGAGUGUGACAGUCACUUA---------GAU---G----UAC
G------CCAAAAGA-CC------------------GGAA-CAUGU--CCCUGAGUACCGUGCGUUG---GA
UAUCGUGCGUGAA-UCUGGGGGGCACC--AACCUCCAAAACUAAAUA--CUC-C-UUG-AGACCGAUAGC-G
AAA-U-AGUAGGGUGACCGAACG-CUGAAA-AGUAC-CCCGAGAAGGGA-G-GUG-CAAAGUGCCUGAAACC
```

U-GAUAGUGAUGG-AAC-GA-UACG---GCAUGAAA-GGAU-------------------------------
------------------------------------------------------------------------
------------------------------------------------------------------------
------------------------------------------------------------------------
------------------------------------------------------------------------
------------------------------------------------------------------------
------------------------------------------------------------CU-CUGGCAUG-
---AA-----------GG-A-------------------A------UCA-----------------UUC---
------------------------------------------------------------------------
--------------------------------------------------------------G--C-GA---
------------------------------------------------------------------------
-------------------------------------------------G-GA------UG--------UAG
UACGA-------------------------------------------------------------------
------------------------------------------------------------------------
------------------------------------------------------------------------
------------------------------------------------------------------------
------------------------------------------------A-----------------------
------------------------------------------------------------------------
------------------------------------------------------------------------
------------------------------------------------------------------------
------------------------------------------------------------------------
------------------------------------------------------------------------
-------------------UGUCAC--U-----------------GCCAG--U---GUCGUAUCUUACGUUU
UGAAGAACGGCCAGGGAGUGUA-U-CUUGGUGGCAAUGGCUAACC--U-U----------UUAU--------
C-UG--GGCA--GCCGAAGCGAAAG-CAA-----------------CAU----------------------G
UGCG-CAGCCUU----U-------------------------------------------------------
------------------------------------------------------------------------
--------------------UGG------------------------------------CGA-GGCACG-AC
GUAU-----------------------------------ACAA--------------------------GU-
G-C-GU----------------------GGAGUCACCGGGGUAC-GACCCGAAGCC-GGGUGAUCUAGGCGU
GGGCAGGUUGAAGCGUGG-CGAAAG-CUACGUGGAGGACCGU-AAACGGUAUUGAUCUGCAAAUCAU-UCGU
GUGACCUGCGUCUCGUAGUGAAAUGCUAAUCUAACCCGG-CAUCCGCUGGUUCCUUCCGAAA-CAUGUCGCA
GCAUGACCUGAC-UUGA-------------------------------------------------------
----------------------GA-----UC-GUCGGUGGAGUAGAGC-ACUGAUU-GGC-GAUU-CCGGGG
G-A-GA-AA-UCCC-U-CGCUCGCUUGUCAAACUCCAAACCCCCG--------------------------
------------------------------------------------------------------------
------------------U-CA-U--CC-UAGA--AAGUCGG-AGUCCGG-A-CUGCUG-GGG-UAAGCUUG
UA-G-UCCGUAAG-GGAGAC---AACCCAGCCCGU-G-GUUAAGGUCCCCAAGU--GU-CGA-CU-AA---G
-U-GUUAAUA-CUAAAGGGCGUCCCAAGCCCCAGACAGCUGGAAGGUUAGCUUAGAAGCAGCUA-CCCUUCA
AAGAGUGCGUAACAGCUCACCAGUC-----GAGGUUUGGGGCCCCGAAAAU-UGAC-GGGGCUCA-AGUCGA
C----------CACCGAU-ACCACGGAGUACCGC--------------------------------------
------------------------------------------------------------------------
------------------------------------------------------------------------
--------------------------------AAGGU-AAUC-UCGUAGGAAGGC-GUUGUGU---------
-----------U-C-GGGUC-GAAGCAG--G-GCUGUGAAGU--C-CUGUGGACCG-U-UCA-CAAACGAAA

AUCC-UGGUAA-UAGUA-GCAGCA--AAGCA-GGGUGA--G---------------ACUCCC-UGC-CGCCG
A-AGGGGC-CAGGUUUCC-UCGGCAAUG-UUCGUCAGCCGAGGGUUAGUCGG--UC-CUAAGACGUACCGU-
AAU-UCGA---G-UAC--------------------------------------------------------
----------------------------------------GCC-GA-A-AGGGA-AACAGGUU-AAUA-UUC
CUGUA----------------------------------CCAUU----UAGC--------------------
------------------------------------------------------------------------
---------------------------------------------------------GUUU-A-GUCU-GAC
GUCUCAG-------------GGCAAG-CCGAGCGGC-GU-------CGUC-------------G-CGCCGU-
CUAA-G-CAG-CG-----A-A-------------------CC-----CC----------------GUGG-AG
A-G-CC--------------------GUAAU-GGCGAGAAGC---------GGGCGA----AUC-UG-UAA-
UG---GC-----------------------------------------------------------------
----------------A-CA-A-GU-C-GGC-U-U-CACCCCGGGACCC-GUGAAA-A-GGA-CAGC-----
---------------------------------------------------------------UA-AAUGUC
CGUACCGAGAACUGAC------------------------------ACAGGUGC-CCCUAGCU-GAAAAGGC
UAAGGC-GUGU-CGGA-GUAAUUCA-G-UUAAGGGAAUUCGGC-AAAUUAGCUCCGUAACUUCGG-GAUAAG
GAGUG-CCUG-C-UG--UGU----------------------------------------------------
------------------------------------------------------------------------
------------------------------------------------------------------------
------------------------------------------------------------------------
------------------------------------------------------------------------
------------------------------------------------------------------------
------------------------------------------------------------------------
------------------------------------------------------------------------
------------------------------------------------------------------------
------------------------------------------------------------------------
------------------------------------------------------------------------
------------------------------------------------------------------------
------------------------------------------------------------------------
------------------------------------------------------------------------
-------------------------------------------------------------------AGACA
GCA--GGUCGCAGUGACCAGGGGGCUCUAA-----CUGUCUAAUACCAACAUAGGAGAUUGCAAACCCG---
-UA-A-------------GGGCUAGU-ACAAUCUCUGA-AUCCUGCUCAGUGCAG-G-UACC-UGAAACCCC
GG-------------UUCA-------------ACGGGAAGAAGGGC--CUGUAAAC-AGCGGGGGUAACUAU
GACCCUCUUAAGGUAGCGUAGUACCUUGUCGCU-UAAUUGGCGACUUGCAUGAAUGGAUCAAUGAGA--GCC
CUACUGUCCCUAAC----UGG-AGUCCGGUGAAGCU--UAC---AUUCUAGUGCACAGUCUAGAGACCUCUA
GGGGGAAGUGAAGA-CCCCGUGGAGC-UUUACU--GCAGCCU-GUCGUUGGGUUGUGGUUCUGGAUGUACAG
UGU-AGGUAGGAGGCAUCG----------AAGCGGG-UGC----------------------------GCCA
--------------------------------------GC-AUUCGUGGAG-C--C-GU-CCUUGGGACA-C
UACC-CUUCCGGGACUACGACCCUAACUC-U----------GA-G-A-------------------------
------------------------------------------------------------------------
------------------------------------------------------------------------
------------------------------------------------------------------------
------------------------------------------------------------------------
------------------------------------------------------------------------
-------------------------------------------------AGA-----GGACC-CCGAUAGGU

```
GGGCAGUUUGCCUGGGGCGGGACGCCCUUGA-AAAGAUAUC-AAGGGCGCGCA-AUGGUUGACUCAAGCGGG
UCGGAAACUCGCUGAA-GAGUG-CAAGAGCAU-AAGUCAGCCUGACG-UUAU--------------------
--------------UCAGCACAGCAGUGGAUGACGAGACGAAA-GUCGGCUCUAGCGAACUUUU--------
---GA-G-----CCUCC-UUG--------------------GU-GGGGGCCAAAAUGACAG-AA--AAGUU
ACCCCGGGG-AUAAUUGAGUCGUUGCCGGCAAGAGUACAUAUCGACCCGGCAGCUUGCUACCUCGAUGUCGG
UUCUUUCCAUCCUGGCCGUGCA-GCAGCGGCCAAGGGUGAGGUUGUUCGCCUAUUAAAGGAGAUCGUGAGCU
GGGUUUAGACCGUCGUGAGACAGGUCGGUUACUAUCUACU--------------------A-GAG-------
-------------------------------GUGC--CCGAGGUCUGCGGGUAAGC-UGCUUUUAGUACGAG
AGGAA-CCAAGCAGCGGCGCCAC-UGGU-G-UACCGGUUGUCC-GACA-GGGCAU-C-GCCGGGCAGCUACG
CGCUC-AGGAAUAAGAGCUGAAUGCA-UC--UAAGCUCGAACUCUGACCUAA-A-A-A--GAGACCUC-UU-
------------------------------------------------------------------------
------------------------------------------------------------------------
------------------------------------------------------------------------
------------------------------------------------------------------------
------------------------------------------------------------------------
------------------------------------------------------------------------
------------------------------------------------------------------------
------------------------------------------------------------------------
----------------------------------------------------------UA-AGG------AU
UCCGGUAAAAGACCGGUU-UGA-UAGAAA-CGG-GAUGUAAG-CAC-C--------AAG-----GC------
----AA--------CGAGGUGUUCAGUCC--G-CGUU-UACUAACC---AUUCGUUCC-UUUCACUU-----
------------------------------------------------------------------------
------------------------------------------------------------------------
------------------------------------------------------------------------
-------------------------------------------------
>LSUa_192952_METMA/1-2893
---------------------------UAGG----AA--UUAUGCUAUCA-GGUGGAUG-G-CUCGGC-UCA
A-G---AGCUGACGAAGGACGUGC--CAAGCUGCGAUAAGCCCGGGGUAGGUGCAUGGAUCCAA------UG
AAC-CCG-GGAUCUCCGAAUGGGACC-U--------CU--------CC-----------------------A
U-A---------------------------------G----U-G-AUCA-----------------------
-----------G-----------UCA----------------------------------------------
------------------------U-G-AU-CGGGAACGCUCCGAAUUGAAACAUCUCAGUAGGAG--CUGG
AAAAGAAAUCAA--ACGAGAUGCCGUAA--GU-AA-CGGCGAGUGAAACCGGC-A-CA-GUUCAAACCGAAU
CCCU----------------------------------UC-GGGGAAAUGUGGUG--UUG------------
U-AG--GGC-UGU-U-----C-------------------------AAAAG--UCU-----------CG--C
G-------GCAAA--ACCA------AACUG-CCUG-------------------------------------
-------GAA-CG-GCAGA-------CCAUAGAGUGUGACAGUCACGUA---------GGU---G----UAA
G------CCAGAAGA-CG------------------GGAA-CAUGU--CCCUGAGUACCGUGCGUCG---GA
UAUCGUGCGGGAA-UCCGGGGGGCACC--AACCUCCAAAACUAAAUA--CUC-C-UUG-AGACCGAUAGC-G
AAA-U-AGUAGGGUGACCGAACG-CUGAAA-AGUAC-CCCGAGAAGGGA-G-GUG-CAAAGUGCCUGAAACC
U-GAUAGUGAUGG-AAC-GA-UACG---GCAUGAAA-GGAU--------------------------------
------------------------------------------------------------------------
------------------------------------------------------------------------
------------------------------------------------------------------------
------------------------------------------------------------------------
```

```
------------------------------------------------------------------------
------------------------------------------------------------CU-UUGAUAUG-
---AA-----------GG-A-------------------A------CCA-----------------CUC---
------------------------------------------------------------------------
--------------------------------------------------------------G--C-GA---
------------------------------------------------------------------------
-------------------------------------------------G-AG------UG--------UUG
UACGA-------------------------------------------------------------------
------------------------------------------------------------------------
------------------------------------------------------------------------
------------------------------------------------------------------------
------------------------------------------------A-----------------------
------------------------------------------------------------------------
------------------------------------------------------------------------
------------------------------------------------------------------------
------------------------------------------------------------------------
------------------------------------------------------------------------
-------------------UAUCAC--U-----------------GCCAG--U---GUCGUAUCUUACGUUU
UGAAGAACGGGCCAGGGAGUGUA-U-CUUGAUGGCAAUGGCUAACC--U-U---------UUAA--------
U-UG--GGCA--GCCGAAGCGAAAG-CAA-----------------CAU----------------------G
UGCG-CAACCCU----U-------------------------------------------------------
------------------------------------------------------------------------
----------------A--CGGG------------------------------------UGA-GGCACG-AC
GUAU-----------------------------------ACAA--------------------------GU-
G-C-GU----------------------GGAGUCAUCGGGGUAC-GACCCGAAGCC-GGGUGAUCUAGGCGU
GGGCAGGUUGAAGCGUGG-CGAAAG-CUACGUGGAGGACCGC-AAGCGGUAUUGAUCUGCAAAUCAU-UCGU
GUGACCUGCGUCUCGUAGUGAAAUGCUAAUCUAACCCGG-CAUCCGCUGGUUCCUUCCGAAA-CAUGUCGCA
GCAUGACCUGAC-UGGA-------------------------------------------------------
----------------------GA-----UU-GUCGGUGGAGUAGAGC-ACUGAUU-GGC-GGUU-CCGGGG
G-G-GA-AA-CCCC-U-CGCCCGCUUGUCAAACUCCAAACCCACUG--------------------------
------------------------------------------------------------------------
------------------U-CA-U--CA-AAGA--UAGUCGG-AGUCCGG-A-CUGCUG-GGG-UAAGCUUG
UA-G-UCCGUAAG-GGAGAC---AACCCAGCCCGU-G-GUUAAGGUCCCCAAGU--GU-CGA-CU-AA---G
-U-GUAAACA-CUAAAGGGCGUCCCAAGCCCAAGACAGCUGGAAGGUUAGCUUAGAAGCAGCUA-CCCUUCA
AAGAGUGCGUAACAGCUCACCAGUC-----GAGGUUUGGGGCCCCGAAAAU-UGAC-GGGGCUCA-AGUCGA
C----------CACCGAU-ACCACGGAGUACCGU--------------------------------------
------------------------------------------------------------------------
------------------------------------------------------------------------
--------------------------------AAGGU-AAUC-UCGUAGGAAGGC-GUUGUGU---------
-----------U-C-GGGCU-GAAGCAG--G-GCUGUGAAGU--C-CUGUGGACCG-U-UCA-CAAACGAAA
AUCC-UGGUAA-UAGUA-GCAGCA--UAGCA-GGGUGA--G---------------AAUCCC-UGC-CGCCG
A-AGGGGC-CAGGUUUCC-UCGGCAAUG-UUCGUCAGCCGAGGGUUAGUCGG--UC-CUAAGACGUACCGU-
AAU-UCGA---G-UAC--------------------------------------------------------
----------------------------------------GCC-AA-A-AGGGA-AACAGGUU-AAUA-UUC
CUGUA----------------------------------CCAUU----UAGC--------------------
```

```
------------------------------------------------------------------------
---------------------------------------------------------ACUG-A-GUCU-GAC
GUCUCAG-------------GGCAGG-CCGAGCGGC-GC-------CGUC-------------G-CGCCGU-
CUAA-G-CAG-CG-----A-A-------------------UC-----CC----------------GUGG-AG
A-G-CC--------------------GUAAU-GGCGAGAAUC-----------GGGCGA----AUC-UG-UAA-
UG---GC-----------------------------------------------------------------
----------------U-UA-A-GU-C-GGC-U-G-CACCCAGGGACCC-GUGAAA-A-GGC-AGC------
---------------------------------------------------------------UA-AAUGUC
CGUACCGAGAACUGAC------------------------------ACAGGUGC-CCCUAGCU-GAAAAGGC
UAAGGC-GUGU-CGGA-GUACUUCA-G-UUAAGGGAAUUCGGC-AAAUUAGCUCCGUAACUUCGG-GAUAAG
GAGUG-CCUG-C-UG--UGA----------------------------------------------------
------------------------------------------------------------------------
------------------------------------------------------------------------
------------------------------------------------------------------------
------------------------------------------------------------------------
------------------------------------------------------------------------
------------------------------------------------------------------------
------------------------------------------------------------------------
------------------------------------------------------------------------
------------------------------------------------------------------------
------------------------------------------------------------------------
------------------------------------------------------------------------
------------------------------------------------------------------------
------------------------------------------------------------------------
-------------------------------------------------------------------AGACA
GCA--GGUCGCAGUGACCAGGGGGCUCUAA-----CUGUCUAAUACCAACAUAGGAGAUUGCAAACCCG---
-UA-A-------------GGGCUAGU-ACAAUCUCUGA-AUCCUGCUCAGUGCAG-G-UACC-UGAAACCCC
GG-------------UUCA-------------ACGGGAAGAAGGGC--CUGUAAAC-AGCGGGGGUAACUAU
GACCCUCUUAAGGUAGCGUAGUACCUUGUCGCU-UAAUUGGCGACUUGCAUGAAUGGAUCAAUGAGA--GCC
CUACUGUCCCUAAC----UGG-AAUCCGGUGAAGCU--UAC---AUUCUAGUGCACAGUCUAGAGACCUCUA
GGGGGAAGUGAAGA-CCCCGUGGAGC-UUUACU--GCAGCCU-GUCGCUGGGUUGUGGUUCUGGAUGUACAG
AGU-AGGUAGGAGGCGUCG----------AAGCGUG-UGC----------------------------GCCA
--------------------------------------GC-AUGCGUGGAG-C--C-AC-AAUUGGGACA-C
UACC-CUUCUGGGACUACGACCCUAACUC-U----------GC-G-A-------------------------
------------------------------------------------------------------------
------------------------------------------------------------------------
------------------------------------------------------------------------
------------------------------------------------------------------------
------------------------------------------------------------------------
-------------------------------------------------AGA-----GGACC-CCGAUAGGU
GGGCAGUUUGCCUGGGGGCGGGACGCCCUUGA-AAAGAUAUC-AAGGGCGCGCA-AUGGUUGACUCAAGUGGG
UCGGAAACUCACUGAA-GAGUG-CAAGAGCAU-AAGUCAGCCUGACG-UUAU--------------------
--------------UCAGCACAGCAGUGGAUGACGAGACGAAA-GUCGGUUCUAGCGAACUUUU--------
---GA-G-----CCUCC-UUG--------------------GU-GGGGGCCAAAAAUGACAG-AA--AAGUU
ACCCCGGGG-AUAAUUGAGUCGUUGCCGGCAAGAGUACAUAUCGACCCGGCAGCUUGCUACCUCGAUGUCGG
```

```
UUCUUUCCAUCCUGGCCGUGCA-GCAGCGGCCAAGGGUGAGGUUGUUCGCCUAUUAAAGGAGAUCGUGAGCU
GGGUUUAGACCGUCGUGAGACAGGUCGGUUACUAUCUACU--------------------A-GAG-------
-------------------------------GUGC--ACAAGGUCUGCGGGUAAGC-UGCUUUUAGUACGAG
AGGAA-CCAAGCAGCGGCGCCAC-UGGU-G-UACCGGUUGUCU-GACA-AGGCAU-C-GCCGGGCAGCUACG
CGCUA-AGGAAUAAGAGCUGAAUGCA-UC--UAAGCUCGAAAUCUGACCUAA-A-A-A--GAGACCUU-UU-
------------------------------------------------------------------------
------------------------------------------------------------------------
------------------------------------------------------------------------
------------------------------------------------------------------------
------------------------------------------------------------------------
------------------------------------------------------------------------
------------------------------------------------------------------------
------------------------------------------------------------------------
----------------------------------------------------------UA-AGG------AU
UCCGGUAGAAGACCGGUU-UGA-UAGAAA-CGG-GAUGUAAG-CAC-C--------AAG-----GC------
----AA--------CGAGGUGUUCAGUCC--G-CGUU-CACUAACU---AUCCGUUCC--------------
------------------------------------------------------------------------
------------------------------------------------------------------------
------------------------------------------------------------------------
-------------------------------------------------
>LSUa_269797_METBF/1-2895
------------------------------G----AA--UUAUGCUAUCA-GGUGGAUG-G-CUCGGC-UCA
A-G---AGCUGAUGAAGGACGUGC--CAAGCUGCGAUAAGCCCGGGGUAGGUGCAAGGAGCCUA------UG
AAC-CCG-GGAUUUCCGAAUGGGACC-U--------CU--------CC-----------------------A
U-A---------------------------------G----U-G-AUCA-----------------------
-----------G-----------UAA----------------------------------------------
------------------------U-G-AU-CGGGAACGCUCCGAAUUGAAACAUCUCAGUAGGAG--CAGG
AAAAGAAAUCAA--CCGAGAUACCGUGA--GU-AA-CGGCGAGUGAAAACGGC-A-CA-GUUCAAACCGAAU
CCCU-----------------------------U---CAC-AGGGAAAUGUGGUG--UUG------------
U-AG--GGC-UGU-U-----C-------------------------AAACG--UCU-----------GG--C
G-------GCUAA--ACAG------AACUUG-CCUG------------------------------------
-------AAA-CG-GCAGA-------CCGUAGAGUGUGACAGUCACGUA---------UGU---G----UAA
A-----CCAAAAGA-UC-------------------GGAA-CAUGU--CCCUGAGUACCGUGCGUUG---GA
UAUCGUGCGGGAA-UCUGGGGGGCACC--AACCUCCAAAACUAAAUA--CUA-C-UUG-AGACCGAUAGC-G
AAA-U-AGUAGGGUGACCGAACG-CUGAAA-AGUAC-CCCGAGAAGGGA-G-GUG-CAAAGUGCCUGAAACC
U-GAUAGUGAUGG-AAC-GA-UACG---GCAUUAAA-GGAA-------------------------------
------------------------------------------------------------------------
------------------------------------------------------------------------
------------------------------------------------------------------------
------------------------------------------------------------------------
------------------------------------------------------------------------
------------------------------------------------------------CU-CUGACAUG-
---AA-----------GG-A-------------------A------ACA-----------------UUC---
------------------------------------------------------------------------
--------------------------------------------------------------G--C-GA---
```

```
------------------------------------------------------------------------
-------------------------------------------------G-AA------UG--------CUG
UACGA-------------------------------------------------------------------
------------------------------------------------------------------------
------------------------------------------------------------------------
------------------------------------------------------------------------
------------------------------------------------A-----------------------
------------------------------------------------------------------------
------------------------------------------------------------------------
------------------------------------------------------------------------
------------------------------------------------------------------------
------------------------------------------------------------------------
-------------------UGUCAC--U-----------------GCCAG--U---GUCGUAUCUUACGUUU
UGAAGAACGGGCCAGGGAGUUUA-U-CUCAGUGGCAAUGGCUAACC--U----------UUAUU--------
--UG--GGCA--GCCGAAGCGAAAG-CAA-----------------CAU----------------------G
UGCG-CAGCUCU----C-------------------------------------------------------
------------------------------------------------------------------------
-------------------CGAG------------------------------------CGA-GGCACG-AC
GUAU-----------------------------------ACAA--------------------------GU-
G-C-GU----------------------GGAGUCACUGGGGUAA-GACCCGAAGCC-GGGUGAUCUAGGCGU
GGGCAGGUUGAAGCGUGG-CGAAAG-CUACGUGGAGGACCGC-AAGCGGUAUUGAUCUGCAAAUCAU-UCGU
GUGACCUGCGUCUCGUAGUGAAAUGCUAAUCUAACCCGG-CAUCCGCUGGUUCCUUCCAAAA-CAUGUCGCA
GCAUGACCUGAC-UGGA-------------------------------------------------------
----------------------GA-----UC-GUCGGUGGAGUAGAGC-ACUGAUU-GGC-GAUU-CCGGGG
G-G-GA-AA-CUCC-U-CGCUCGCCUGUCAAACUCCAAACCCACCG--------------------------
------------------------------------------------------------------------
------------------U-CA-U--UG-UAGA--AAGUCGG-AGUCCGG-A-CUGCUG-GGG-UAAGCUUG
UA-G-UCCGUAAG-GGAGAC---AACCCAUCCCGU-G-GUUAAGGUCCCCAAGU--GU-CGA-CU-AA---G
-U-GUUAAUA-CUAAAGGGCGUCCCAAGCCCAAGACAGCUGGAAGGUUAGCUUAGAAGCAGCUA-CCCUUUA
AAGAGUGCGUAACAGCUCACCAGUC-----GAGGUUUGGGGCCCCGAAAAU-UGAC-GGGGCUCA-AGUCGA
C----------CACCGAU-ACCACGGAGUACCGC--------------------------------------
------------------------------------------------------------------------
------------------------------------------------------------------------
--------------------------------AAGGU-AAUC-UCGUAGGAAGGC-GUUGCGU---------
-----------U-C-GGGCC-GAAGCAG--G-GCUGUAAAGU--C-CUGUGGACCG-U-UCG-CAAACGAAA
AUCC-UGGUAA-UAGUA-GCAGCA--AAGCA-GGGUGA--G---------------AAUCCC-UGC-CGCCG
A-AGGGGC-CAGGUUUCC-UCGGCAAUG-AUCGUCAGCCGAGGGUUAGUCGG--UC-CUAAGACGUACCGU-
AAC-UCGA---G-UAC--------------------------------------------------------
----------------------------------------GCC-GA-A-AGGGA-AACAGGUU-AAUA-UUC
CUGUA----------------------------------CCAUU----UAGC--------------------
------------------------------------------------------------------------
---------------------------------------------------------AAUC-C-GUCU-GAC
GUCUCUG-------------GGCAGA-CCGAGCGGC-GU-------CGUC-------------G-CGCCGU-
CUAA-G-CAG-CG-----A-A-------------------CC-----CC-----------------GUGG-AG
A-G-CC--------------------GUAAU-GGCGAGAAUC---------GGGCGA----AUC-UG-UUA-
```

```
UG---GC-----------------------------------------------------------------
----------------U-AA-A-GU-C-GGU-C-G-CACCCCGGGACCC-GUGAAA-A-GGC-CGC------
---------------------------------------------------------------UA-AAUGUC
CGUACCAAGAUCUGAC------------------------------ACUGGUGC-CCCUAGCU-GAAAAGGC
UAAGGC-GUGU-CGGA-GUACUUCA-G-UUAAGGGAAUUCGGC-AAAUUAGCUCCGUAACUUCGG-GAUAAG
GAGUG-CCUG-C-UG--UGG----------------------------------------------------
------------------------------------------------------------------------
------------------------------------------------------------------------
------------------------------------------------------------------------
------------------------------------------------------------------------
------------------------------------------------------------------------
------------------------------------------------------------------------
------------------------------------------------------------------------
------------------------------------------------------------------------
------------------------------------------------------------------------
------------------------------------------------------------------------
------------------------------------------------------------------------
------------------------------------------------------------------------
------------------------------------------------------------------------
-------------------------------------------------------------------AGACA
GCA--GGUCGCAGUGACCAGGGGGCUCUAA-----CUGUCUAAUACCAACAUAGGAGAUUGCAAACCCG---
-UA-A-------------GGGCUAGU-ACAAUCUCUGA-AUCCUGCUCAGUGCAG-G-UACC-UGAAACCCC
GU-------------UUCA-------------ACGGGACGAAGGGC--CUGUAAAC-AGCGGGGGUAACUAU
GACCCUCUUAAGGUAGCGUAGUACCUUGUCGCU-UAAUUGGCGACUUGCAUGAAUGGAUCAAUGAGA--GCC
CUACUGUCCCUAAC----UGG-AAUCCGGUGAAGCU--UAC---AUUCUAGUGCACAGUCUAGAGACCUCUA
GGGGGAAGUGAAGA-CCCCGUGGAGC-UUUACU--GCAGCCU-GUCGCUGGGUUGUGGUUCUGGAUGUACAG
AGU-AGGUAGGAGGCAUCG----------AAGCGUA-UGC----------------------------GCCA
--------------------------------------GC-AUACGUGGAG-C--C-GC-AAUUGGGACA-C
UACC-CUUCUGGGACUACGACCCUAACUC-U----------GA-G-A-------------------------
------------------------------------------------------------------------
------------------------------------------------------------------------
------------------------------------------------------------------------
------------------------------------------------------------------------
------------------------------------------------------------------------
-------------------------------------------------AGA-----GGACC-CCGAUAGGU
GGGCAGUUUGCCUGGGGCGGGACGCCCUUGA-AAAGAUAUC-AAGGGCGCGCA-AUGGUUGACUCAAGCGGG
UCGGAAACUCGCUGAA-GAGUG-CAAGAGCAU-AAGUCAGCCUGACG-UUAU--------------------
--------------UCAGCAUAGCAGUGGAUGACGAGACGAAA-GUCGGCUCUAGCGAACUUUU--------
---GA-G-----CCUCC-UUG--------------------GU-GGGGGCCAAAAAUGACAG-AA--AAGUU
ACCCCGGGG-AUAAUUGAGUCGUUGCCGGCAAGAGUACAUAUCGACCCGGCAGCUUGCUACCUCGAUGUCGG
UUCUUUCCAUCCUGGCCGUGCA-GCAGCGGCCAAGGGUGAGGUUGUUCGCCUAUUAAAGGAGAUCGUGAGCU
GGGUUUAGACCGUCGUGAGACAGGUCGGUUACUAUCUACU--------------------A-GAG-------
-------------------------------GUGC--ACGAGGUCUGCGGGUAAGC-UGCUUUUAGUACGAG
AGGAA-CCAAGCAGCGGCGCCAC-UGGU-G-UACCGGUUGUCU-GACA-AGGCAU-A-GCCGGGCAGCUACG
CGCCA-AAGAAUAAGAGCUGAAUGCA-UC--UAAGCUCGAACUCUCACCUAA-A-A-A--GAGACCUC-AU-
```

```
------------------------------------------------------------------------
------------------------------------------------------------------------
------------------------------------------------------------------------
------------------------------------------------------------------------
------------------------------------------------------------------------
------------------------------------------------------------------------
------------------------------------------------------------------------
------------------------------------------------------------------------
----------------------------------------------------------UA-AGG------AU
UCCGGUAGAAGACCGGUU-UGA-UAGAAA-CGG-GAUGUAAG-CAC-C--------AAG-----GC------
----AA--------CGAGGUGUUCAGUCC--G-CGUU-UACUAACU---AUCCGUUCC-UUUCU--------
------------------------------------------------------------------------
------------------------------------------------------------------------
------------------------------------------------------------------------
-------------------------------------------------
>LSUa_349307_METTP/1-2886
----------------------------------------UAUGCCAGCC-GGGGGAUG-G-CUCGGC-UCG
A-G---CGCUGAUGAAGGGCGUGC--CAAGCUGCGAUAAGCCCCGGCGAGGCGCAGGGAGCCUU------UG
AAC-CGG-GGAUGCCCUAAUGGGAAA-U--------CCC-------AG----UU-----------------U
C-A-------------------------------UGC----U-G-AUCC-----------------------
-----------G-----------UAA----------------------------------------------
------------------------G-G-AU-CGGGAACGCCCCGGAUUGAAACAUCUUAGUAGGGG--CAGG
AGAAGAAAUCAAA-ACGAGAUGCCGUCA--GU-AA-CGGCGAGUGAACGCGGC-A-GA-GCCCAAACCGAAU
CCCG----------------------------------AA-AGGGAGAUGUGGUG--UGA------------
U-AA--GAC-CGC-U-----C------AU----------------AACGGU--CUG-----------GC--A
A-------GGGAA--UCCG------AAGUGG-CCUG------------------------------------
-------GAA-CG-GCCCG-------CCAAAGAGUGUGAUAGCCACGUA---------GGA---G----CAA
C------CAAGCCGG-AUC-----------------CGAG-CGGCG--UCUUGAGUACCGUGAGUCG---GA
UAUCUCGCGGGAA-GUUGGGGGUCACC--AACCUCCAAGGCUAAAUA--CGU-C-UCG-AGACCGAUAGC-G
CAG-U-AGUAGGGUGACCUAAAG-CUGAAA-AGAAC-CCUGGAUAGGGG-U-GUG-AAAAGAGCCUGAAAUC
G-GCUGGCGAUGG-UGC-GU-CACG---GCAUGUGA-GGAC-------------------------------
------------------------------------------------------------------------
------------------------------------------------------------------------
------------------------------------------------------------------------
------------------------------------------------------------------------
------------------------------------------------------------------------
------------------------------------------------------------CU-UCAGGCCG-
---AA-----------GG-A-------------------G------GUC-----------------GUC---
------------------------------------------------------------------------
--------------------------------------------------------------G--C-GA---
------------------------------------------------------------------------
-------------------------------------------------G-GC------GA--------CAG
UACGA-------------------------------------------------------------------
------------------------------------------------------------------------
------------------------------------------------------------------------
```

```
------------------------------------------------------------------------
------------------------------------------------G-----------------------
------------------------------------------------------------------------
------------------------------------------------------------------------
------------------------------------------------------------------------
------------------------------------------------------------------------
------------------------------------------------------------------------
-------------------GCCUGA--U-----------------GCCGG--U---GUCGUGACUUACGUUU
UGAAGAACGGACCAAGGAGUGCA-U-CUGGCUGGCAA-GGUUAAUC--U-C---------GAAA--------
G-AG--AGUA--GCCGAAGCGAAAG-CGA-----------------CAU----------------------G
CGCG-CAUCGA-------------------------------------------------------------
------------------------------------------------------------------------
-------------------UGAG------------------------------------AGA-GGCGCG-GC
GUAU-----------------------------------GAAA--------------------------GU-
G-C-GC----------------------GGAGUCAGCUGGAUGC-GACCCGAAGCC-GGGCGAUCUAGGCGU
GGGCAGGUUGAAGCGUGG-CGAAAG-CUGCGUGGAGGACCGC-UAGCGGUGCUGAUAUGCAAAUCGC-UCGC
GUGACCUGCGUCUAGGGGUGAAAGUCCAAUCGAGCCCGG-AAAUAGCUGGUUCCUCCCGAAA-CAUGUCGCA
GCAUGACCUGGC-CGGA-------------------------------------------------------
----------------------GG-----CA-GUCGGCGGAGUAGAGC-ACUGAUU-GGG-GAUC-CCGGGC
G-G-GA-AA-CCGC-U-CGCUCUCCUGUCAAACUCCCAACACGCCG--------------------------
------------------------------------------------------------------------
------------------U-CG-C--CU-UAGA--UGGCCGGAAGUCCGG-G-CUGG-U-GGG-UAAGCUAC
UA-G-UCCGUGAG-GGAGAC---AACCCAGACCGU-G-GUUAAGGUCCCAAAAU--GU-CGG-CU-AA---G
-U-GUUAACA-CUGAAGGGCGUCCUAGACCUAAGACAACUGGGAGGUUAGCUUAGAAGCAGCUA-CCCUUUA
AAGAGUGCGUAACAGCUCACCAGUC-----GAGGCCUAGGGCCCCGAAAAU-GGAC-GGGGCU-A-AGCCGA
C----------UACCGAU-ACCACGGAGCGCCGC--------------------------------------
------------------------------------------------------------------------
------------------------------------------------------------------------
--------------------------------AAGGC-GAUC-UGGUAGGGAGGC-GCCGCGC---------
-----------G-U-GGGUG-GAAGCAA--C-UCCGUGAGGA--G-UCGUGGACCA-C-GCG-GGGACGAAA
AUCU-UGGUAG-CAGUA-GCAGCA--AAGUC-GGGUGA--G---------------AAUCCC-GAC-CGCCG
U-AAGGGC-CAGGGUUCC-UGGGCAACG-AUCGUCAGCCCAGGGUUAGUCGG--UC-CUAAGGUGUGCCGU-
AAC-UCGA---G-CAC--------------------------------------------------------
----------------------------------------AUC-AA-A-UGGGA-AACAGGUU-AAUA-UUC
CUGUA----------------------------------CCUAC----CGGC--------------------
------------------------------------------------------------------------
---------------------------------------------------------ACUA-A-AACC-GAC
GCUUCCG-------------GAUAGC-CUGA-CACC-CG-------CGCC-------------A-GGGUGC-
UCAA-G-CGC-CG-----A-A-------------------GU-----CC----------------GCGG-AG
A-G-CC--------------------GUCAU-GGCGAGAAAC---------GGACGA----AUG-CG-UGA-
UG---AC-----------------------------------------------------------------
----------------G-CA-A-GC-A-GGC-A-A-AUUCCGGGAGCCC-GUGAAA-A-GGG-AG-------
---------------------------------------------------------------CC-GGAGGC
CGUACCGAGAACCAAC------------------------------ACUGGUGC-CCCGAGCU-AAGAAGGC
UCAGGC-GUGU-CGGAAGUAAUCCG-G-CUAAGGGAAUUCGGC-AAAUUAGCCACGUAACUUCGG-GAUAAG
```

```
UGGUG-CCUG-C-UU--CGC----------------------------------------------------
------------------------------------------------------------------------
------------------------------------------------------------------------
------------------------------------------------------------------------
------------------------------------------------------------------------
------------------------------------------------------------------------
------------------------------------------------------------------------
------------------------------------------------------------------------
------------------------------------------------------------------------
------------------------------------------------------------------------
------------------------------------------------------------------------
------------------------------------------------------------------------
------------------------------------------------------------------------
------------------------------------------------------------------------
-------------------------------------------------------------------AAGAA
GCA--GGUCGCAGUGACGAGGGGGCUCUGA-----CUGUCUAAUAACAACAUAGGUGGUUGCAGACCCG---
-AA-A-------------GGGCAAGU-ACAACCACUGA-AUCCUGCCCAGUGCGG-G-UACC-UGAAAACCC
GG-------------UACA-------------ACGGGAAGAAGGGC--CCGUCAAC-GGCGGGGGUAACUAU
GACCCUCUUAAGGUAGCGUAGUACCUUGCCGCU-UAAUUGGCGGCUCGCAUGAAUGGAUCAACGAGA--GCC
CCACUGUCCCUAGC----CGG-AACCCGGUGAACCU--UAC---GUCCCAGUGCAGAGUCUGGGGACCCCUG
AUGGGAAGUGAAGA-CCCCGUGGAGC-UUUACU--GCAGCCU-GCCGCUGGGUUGUGGUUCUGGAUGUAGAG
CAU-AGGUAGGAGGCGUCG----------AAACCCG-GGC----------------------------GCCA
--------------------------------------GC-UCGGGCGGAG-C--C-AC-CACUGGAAUA-C
UACC-CUUCUGGAACUACCACCCUAACUC-C----------GA-A-A--------------------------
------------------------------------------------------------------------
------------------------------------------------------------------------
------------------------------------------------------------------------
------------------------------------------------------------------------
------------------------------------------------------------------------
-------------------------------------------------GGA-----GGACA-CCGGUAGGU
GGGCAGUUUGGGUGGGGCGCCACUCCCUAGA-AAAGAUAUC-UAGGGAGCCCU-AAGGUCGACUCAGUUGGG
UCGGAAACCCAACAAA-GAGUG-UAAAAGCAU-AAGUCGGCCUGACG-UCAU--------------------
--------------CCCGCAUAGCAAAGGAUGACGAGACGAAA-GUCGGGUUUAGCGAACCUCU--------
---CA-G-----CCCCC-UUG--------------------GU-GGGGGCGUUAGAUGACAG-AA--AAGCU
ACCCCGGGG-AUAACAGAGUCGUCACCGGCAAGAGCACAUAUCGACCCGGUGGCUUGCUACCUCGAUGUCGG
UUCUCUCCAUCCUGGCCGUGCA-GCAGCGGCCAAGGGUGAGGUUGUUCGCCUAUUAAAGGAGAUCGUGAGCU
GGGUUUAGACCGUCGUGAGACAGGUCGGUUACUAUCUGUC--------------------A-GGG-------
-------------------------------GUGCU-UCGCGGUCUGAGGGCAGGC-UGGCUUUAGUACGAG
AGGAA-CAAGCCAGCGGCGCCAC-UGGU-C-GAUCGGUUGUCU-GGCA-AGGCAC-G-GCCGAGCAGCUACG
CGCCACAGGGAUAAGGGCUGAAAGCA-UC--UAAGCCCGAAACCCAACCCGA-A-A-A--GAGACCGC-UC-
------------------------------------------------------------------------
------------------------------------------------------------------------
------------------------------------------------------------------------
------------------------------------------------------------------------
------------------------------------------------------------------------
```

------------------------------------------------------------------------
------------------------------------------------------------------------
------------------------------------------------------------------------
----------------------------------------------------------CA-AGG------GC
CCUCGUAAAAGACGAGUU-CGA-UAGAGU-CGG-GGUGUACG-CAC-C--------AAG-----GC------
----AA--------CGAGGUGCUCAGCCC--G-CGAC-CACUAACC---GCC--------------------
------------------------------------------------------------------------
------------------------------------------------------------------------
------------------------------------------------------------------------
-------------------------------------------------
>LSUa_410358_METLZ/1-2886
------------------------------U----UC--UUAUGCCGGUU-AGUGAAUG-G-UUCGGU-UUG
A-G---AGCUGAUGAAGGGCGUGC--CAAGCAGCGAAAUGCCCCGGGAAGACGCAAGGAGUCUG------UG
AUC-CGG-GGAUCCCCUAAUAGGACC-U--------CCC-------CA----CGA---------------UU
U-A---------------------------U---CGU----G-A-AUCC-----------------------
-----------G-----------UAA----------------------------------------------
------------------------G-G-AU-GGGGAACCCCCCGAAGUGAAACAUCUCAGUAGGGG--GAGG
AAGAGAAAUCAA--UUGAGAUUACGUUA--GU-AG-AGGCGAUCGAACCCGUA-A-UA-GUCUAAACUGAAU
CCCG----------------------------------CA-AGGGACAUGUAGUG--UAU------------
G-GC--CUU-CGU-U-----G-------------------------UAGUA--UGU-----------GU--U
U-------AUCUA--AUGA------AAGUAG-CUUG------------------------------------
-------GAA-UG-GCUUA-------CCUUAGAGGGUGACAGUCCCGUA---------CGU---G----GUG
G------UAAACACA-GC------------------UUAG-AAGUG--UCCUGAGUAGCGCGGGUUG---GA
AUUCCCACGUGAA-UAUGGGGGUCACC---ACCUCCAAGACUAAAUA--CUC-C-UCA-AAACCGAUAGC-G
AAA-U-AGUAGCGUGAGCGAAAA-CUGAAA-AGUAU-CCCUGAAAAGGA-A-AUG-AAAAGUGCCUGAAACU
A-ACCGGUGAUCG-UGU-GU-UACG---GCGUGCAA-GGAU-------------------------------
------------------------------------------------------------------------
------------------------------------------------------------------------
------------------------------------------------------------------------
------------------------------------------------------------------------
------------------------------------------------------------------------
------------------------------------------------------------CU-UCCGAUCG-
---AA-----------AG-A-------------------A------CCC-----------------GUC---
------------------------------------------------------------------------
--------------------------------------------------------------G--C-GA---
------------------------------------------------------------------------
-------------------------------------------------G-GC------GG--------UAG
UAUGA-------------------------------------------------------------------
------------------------------------------------------------------------
------------------------------------------------------------------------
------------------------------------------------------------------------
------------------------------------------------G-----------------------
------------------------------------------------------------------------
------------------------------------------------------------------------
------------------------------------------------------------------------

```
------------------------------------------------------------------------
------------------------------------------------------------------------
-------------------AUUGGU--U-----------------GCCGA--C---GUCGUAACUUACGUUU
UGAAGAACGGGCCAAAGAGUGUA-U-UCCGCUGGCGA-GGUUAACC--------------GGAA--------
---G--GGCA--GCCGCAGCGAAAG-CGA-----------------CAA----------------------G
CGCG-UAGUU--------------------------------------------------------------
------------------------------------------------------------------------
---------------------UA------------------------------------CCU-GGCGCG-GC
GUAC-----------------------------------UAAUA-------------------------GU-
G-C-GU----------------------GGAGUCAACGGGAUAC-GACCUGAAACC-UGGUGAUCUAUGCCU
GAGCAGGUUGAAGCGUGC-CGAAAG-GUGCGUGGAGGACCGC-AUGCGGUAUUGAUAUGCAAAUCAU-UCGC
GUGACUUGGGUAUAGGAGUGAAAGAUUCAUCGAACCGGG-UGUCAGCUGGUUCCUCUCGAAA-CAUGUCUAA
GCAUGACCUCGA-CCGA-------------------------------------------------------
----------------------GA-----AU-GAUAGUGAGGUAGAGC-ACCUAUU-GGA-AAUC-CCGGGG
A-A-GA-AA-UUCC-U-CGCUUUCCUGUAAAACUCCAAACUCCCUA--------------------------
------------------------------------------------------------------------
------------------U-CU-U--UG-UAGA--CGUCGGGCAGUCCGG-G-CUAC-G-GGG-UAAGCUUG
UA-G-UCCGUAAG-GGAGAC---AACCCAGACCAU-G-GUUAAUGCCCCCCAAU--GU-AGG-CU-AA---G
-U-GUCAACA-CUGAAUGAAGUCCUAGGUCAAAGACAACUGGAAGGUGAGCUCAGAAGCAGCUA-UCCUUUA
AAAAGUGCGUAACAGCUUACCAGUC-----AAGAUUUAGGGCGCAGAAAAU-GGAC-GGGGCUCA-AGCCUA
C----------UGCAGAU-ACCAUGGCGCACGUU--U-----------------------------------
------------------------------------------------------------------------
-----UUG----------------------------------------------------------------
--------------------------------AACGU-GAUC-GAGUAGAGAGGC-GUCCUGU---------
-----------A-U-GGGCU-GAAGCAG--G-GGUGUAAGUU--C-CUGUGGACCG-U-AUA-GGAAUGAGA
AUUU-UGGCAG-UAGUA-GAAGCA--AAGAA-UGGUGA--G---------------AAUCCA-UUC-CGCCG
A-AGGGGC-UAGGUUUCC-UCGGCAAUG-UUCGUCAGCCGAGGGUUAGUCGG--UC-CUAAGCUGCAUCAU-
AAG-UUGC---A-UGC--------------------------------------------------------
----------------------------------------AGC-AA-A-AGGGA-AACAGGUU-AAUA-UUC
CUGUA----------------------------------CCGGC----GUUA--------------------
------------------------------------------------------------------------
-----------------------------------------------------------AU-A-UACU-GAC
GCAUCGG-------------GUUAUG-CAUAGCGGG-AC-------CGUC-------------G-UCCCGU-
CCAA-U-CGU-AU-----A-A-------------------AC-----UC----------------UUGG-AG
U-A-CC--------------------GUAAU-GGUGAGAAUU---------GAGUGA----AAG-CG-UGA-
UG---GC-----------------------------------------------------------------
----------------G-UA-A-GU-G-UGU-A-A-ACACCUGGUGCCC-GUGAAA-A-GGU-AAC------
---------------------------------------------------------------GC-AAUGUC
CGUACCGAGAACCGAC------------------------------ACAGGUGC-CCCUAGUU-GAAAAGAC
UCAGGC-GUGU-CGGAAUUAUUUGU-G-GUAAGGGAACUCGGC-AAAAUGGCUCCGUACCUUCGG-AACAAG
GAGUG-CCUG-C-CU--AGU----------------------------------------------------
------------------------------------------------------------------------
------------------------------------------------------------------------
------------------------------------------------------------------------
------------------------------------------------------------------------
```

```
------------------------------------------------------------------------
------------------------------------------------------------------------
------------------------------------------------------------------------
------------------------------------------------------------------------
------------------------------------------------------------------------
------------------------------------------------------------------------
------------------------------------------------------------------------
------------------------------------------------------------------------
------------------------------------------------------------------------
------------------------------------------------------------------GAUUAG
GCA--GGUCGCAGUGACCAGGGAGCUCUAA-----CUGUCUAACAACAACAUAGCAGACUGCAACUCCG---
-UA-A-------------GGACUAGU-AUAGUCUGUGA-AUCCUGCCCAGUGCGG-G-UAUC-UGAACACUG
AU-------------UACA-------------AUCGGACGAAGGAC--CCGUAAAC-GGCGGGGGUAACUAU
GACCCUCUUAAGGUAGCGUAGUACCUUGUCGCU-UAAUUGGCGACUUGCAUGAAUGGAUUAAUAAGG--GCU
CUACUGUCCCUACC----GCA-AAUCCGUUGAACCU--UUU---AUCCUUGUGCAGAGGCAAGGGACUCCCA
UCGGGAAGUGAAGA-CCCCAUGGAGC-UUUACU--GCAGCCU-GUCGUUGACUUACGGUAUUACAUGCGCAG
CGU-AGAUAGUAGGUGUUA----------AUGCUCA-UAU----------------------------UCCG
--------------------------------------GU-AUGAGUGGAG-C--C-AC-AGUUGAGACA-C
UAUC-CUUGUUCUGCUGUAAUUCUCACUC-------------U-U-U-------------------------
------------------------------------------------------------------------
------------------------------------------------------------------------
------------------------------------------------------------------------
------------------------------------------------------------------------
------------------------------------------------------------------------
--------------------------------------------------GA-----GGACA-UCGGUAGGU
AGGCAGUUUGGGUGGGGCGCCACACCCUCGA-AAAAAUAUC-AAGGGUGCCCU-AUGGUCAACUCAUGUGAG
UCAGCAACUCACAGGA-GAGUGUCAAGAGCAU-AAGUUGGCUUGACG-UGAU--------------------
--------------UUCGCAUAGCAAGAAAUCACGAGUGGAAA-CACGGGUCUAACGAACCAAU--------
---AC-G-----CCUUU-UUG--------------------GU-GAGGGCUAUUGACUACAG-AA--AAGCU
ACCCUGGGG-AUAACAGAGUCGUCGCCGGCAAGAGCACAUAUCGACCCGGCGGCUUGCUACCUCGAUGUCGG
UUCUUUCCAUCCUGGCUGUGCA-GCAGCAGCCAAGGGUGAGGUUGUUCGCCUAUUAAAGGGGAUCGUGAGCU
GGGUUCAGACCGUCGUGAGACAGGUCGGUUACUAUCUGAU--------------------G-GGA-------
-------------------------------GUGU--UCGAAAUCUGAGAGCAAGA-AUGAAAUAGUACGAG
AGGAA-CUUUCAUUCGUCACCUC-UGGU-G-UAUCGGUUGUCU-GAUA-AGGCAG-U-GCCGAGCAGCUACG
UGGCA-CGAGAUAAAAACUGAAAGCA-UC--UAAGUUUGAAACUUCGCUUGA-A-A-A--GAGAUUUC-GU-
------------------------------------------------------------------------
------------------------------------------------------------------------
------------------------------------------------------------------------
------------------------------------------------------------------------
------------------------------------------------------------------------
------------------------------------------------------------------------
------------------------------------------------------------------------
------------------------------------------------------------------------
----------------------------------------------------------UAUAGA------GC
ACCGGUAAAAGACCGGGU-UAA-UAGGCU-UGG-GAUGUACG-CAC-G--------AAG-----GU------
```

```
----AA--------CGACGUGUUUAGUCC--G-CAAG-UACUAA-C---GCUCAA-----------------
------------------------------------------------------------------------
------------------------------------------------------------------------
------------------------------------------------------------------------
-------------------------------------------------
>LSUa_368407_METMJ/1-2898
----------------------------------------UAUGCCUAUC-AGUGGAUG-G-CUCGGU-UCG
A-G---UGCCGAAGAAGGGCGUGC--CAAGCUGCGAUAAGCUCCGGGAAGACGCAUGGAGUCUA------CG
AUC-CGG-AGAUUCCCUAAUGAGACU-U--------CUU-------AA----CAC---------------UU
C-G---------------------------G---UGU----U-G-AUCC-----------------------
-----------G-----------UAA----------------------------------------------
------------------------G-G-AU-CGGGAACCCCCCGAAUUGAAACAUCUUAGUAGGGG--GAGG
AAAAGAAAUCAA--CCGAGAUGUCGUUA--GU-AG-AGGCGAUCGAACACGAC-A-GA-GUUCAAACUGAAU
CCCG----------------------------------CA-AGGGAGAUGUGGUG--UAG------------
U-AA--GCC-UAC-C-----A------U------------------CUGGA--UGC-----------CA--A
G-------GUCAA--ACGG------AAAUCC-UCUG------------------------------------
-------GAA-UG-AGGUG-------CCAAAGAGGGUGAUAGCCCGUA----------CGU---G----CAA
G------CUGAGGUG-UC------------------CUGG-UAGUG--UCUUGAGUACCGCGGGUCG---GA
AUUCUCGCGGGAA-UCUGGGGGUCAUC--AACCUCCAAAACUAAAUA--CUA-C-UCG-AAACCGAUAGC-G
UAU-U-AGUAGCGUGAGCGAAGG-CUGAAA-AGUAA-CCCUUGAAAGGU-G-GUU-AAAAGCGCCUGAAACU
G-AUAGGUGAUAG-UGA-GU-UACG---GCGCGAAA-GGAU--------------------------------
------------------------------------------------------------------------
------------------------------------------------------------------------
------------------------------------------------------------------------
------------------------------------------------------------------------
------------------------------------------------------------------------
------------------------------------------------------------CU-UCCACGUG-
---AA-----------GG-A-------------------A------CCC-----------------GUC---
------------------------------------------------------------------------
--------------------------------------------------------------G--C-GA---
------------------------------------------------------------------------
-------------------------------------------------G-GC------GG--------UAG
UACGG-------------------------------------------------------------------
------------------------------------------------------------------------
------------------------------------------------------------------------
------------------------------------------------------------------------
------------------------------------------------G-----------------------
------------------------------------------------------------------------
------------------------------------------------------------------------
------------------------------------------------------------------------
------------------------------------------------------------------------
------------------------------------------------------------------------
-------------------CGUGGU--U-----------------GCCGG--U---GUCGUAACGUACGUUU
UGAAGAACGGGCCAGAGAGUUUA-U-UCUGUAGGCGAAGGUUAACC-------------CAAAA--------
---G--GGGA--GCCGAAGGGAAAC-CAA-----------------CAG----------------------G
```

```
UCCG-UAGCCUU----C-------------------------------------------------------
------------------------------------------------------------------------
--------------------GGG------------------------------------CAU-GGGACG-GC
GUAU-----------------------------------UAAAA-------------------------GU-
G-C-GU----------------------GGAGUCUGCAGGAUAA-GACCCGAAGCC-UAGCGAUCUAUGCGU
GGGCAGGUUGAAGCGUGC-CGAAAG-GCGCGUGGAGGACCGC-AAGCGGUUUUGAUAUGCAAAUCAU-UCGU
GUGACCUGCGUAUAGGAGUGAAAGAUUAAUCGAGCUGGG-CAUCCGCUGGUUCCUCUCGAAA-CAUGCCGUA
GCAUGACCUGGC-UGGA-------------------------------------------------------
----------------------GA-----UA-GACGGUGAGGUAGAGC-ACUGAUU-GGG-GGAG-CUGGGG
G-A-GA-AA-UCCC-U-CACCCUCCUGUCAAACUCCAAAUUCCCCG--------------------------
------------------------------------------------------------------------
------------------U-CG-U--CG-AAGA--AGGCUGGCAGUCCGG-A-CUAC-G-GGG-UAAGCUUG
UA-G-UCCGUAAG-GGAGAC---AACCCAGACCGU-G-GUUAAUGUCCCUCAAU--GU-AGG-CU-AA---G
-U-GUCAACA-CUGAAAGGCGUCCCAGGCCAAAGACAACUGGAAGGUGAGCUCAGAAGCAGCUA-UCCUUUA
AAAAGUGCGUAACAGCUUACCAGUC-----GAGGUUUGGGGCACUGAAAAU-GGAC-GGGGCUAA-AGCCUA
C----------UACAGAU-ACCACGGACCACACU--U-----------------------------------
------------------------------------------------------------------------
-----U------------------------------------------------------------------
--------------------------------CGUGU-GAUG-GCGUAGAGAGGC-GUGCUGC---------
-----------C-U-GGGCA-GAAGCAG--G-GGCGUGAGCU--C-CUGUGGACCG-G-GUA-GUAUUGAGA
AUUC-UGGCAU-UAGUA-GAAGCA--AAGUU-GGGUGA--G---------------AAUCCC-AAC-CGCCG
G-AGGGGC-UAGGUUUCC-UCGGCAAUG-UUCGUCAGCCGAGGGUUAGUCGG--UC-CUAAGAUGUGUCGU-
AAU-UCGA---G-CAC--------------------------------------------------------
----------------------------------------GUC-AA-C-AGGGA-AACAGGUU-AAUA-UUC
CUGUA----------------------------------CCAUU----CAUCCGUA----------------
------------------------------------------------------------------------
-------GCAGUUU--------------------------------------------ACU-G-CCCU-GAC
GCUUCGG-------------GAUAUG-CCGGGCAGG-GU-------CGUC-------------G-CCCUGU-
CCAA-G-CGU-UC-----A-A-------------------GC-----UC----------------GUGG-AG
U-A-CC--------------------GCCAU-GGUGAGAAGC---------GAGUGA----AGG-CG-UGA-
UG---GA-----------------------------------------------------------------
----------------G-UA-A-UU-C-GGC-A-A-AUUCCUGGAGCCC-GUGAAA-A-GGG-GGA------
---------------------------------------------------------------UG-AAUGUC
CGUACCGAGAUCUGAC------------------------------ACAGGUGC-CCCUAGCU-GAAAAGGC
UAAGGC-GUGU-CGGU-UUAAUCGU-G-UUAAGGGAACUCGGC-AAAUUGGCCCCGUAACUUUGG-GAUAAG
GGGUG-CCUG-C-CU--GGC----------------------------------------------------
------------------------------------------------------------------------
------------------------------------------------------------------------
------------------------------------------------------------------------
------------------------------------------------------------------------
------------------------------------------------------------------------
------------------------------------------------------------------------
------------------------------------------------------------------------
------------------------------------------------------------------------
------------------------------------------------------------------------
```

```
------------------------------------------------------------------------
------------------------------------------------------------------------
------------------------------------------------------------------------
------------------------------------------------------------------------
------------------------------------------------------------------GAUCAG
GCA--GGUCGCAGUGACCAGGGGGCUCUGA-----CUGUCUAAUAACAACAUAGGUGACUGCAACUCGG---
-AA-A-------------CGACUAGU-AUAGUCACUGA-UUCCUGCCCAGUGCGA-G-UAUC-UGAACACCG
GG-------------UUCA-------------ACCGGACGAAGGAC--UCGUAAAC-GGCGGGGGUAACUAU
GACCCUCUUAAGGUAGCGUAGUACCUUGUCGCU-UAAUUGGCGACUUGCAUGAAUGGAUUAACGAGA--GCC
CUACUGUCCCUAAC----ACG-AAGCCGUUGAACAU--UUU---AUCCUAGUGCAGAGACUAGGGACUCCAG
AUGGGAAGUGAAGA-CCCCGUGGAGC-UUUACU--GCAGCCU-GUCGCUGUGUUACGGUAUUCCUUGCGCAG
CGU-AGAUGGGAGGCGUCG----------AUCCACC-CCU----------------------------UCCG
--------------------------------------GG-GGGUGGGGAG-C--C-AA-CAAUGAGACA-C
CAUC-CUUGGUUUACUGUAACACUAACUC-A----------AA-C-C-------------------------
------------------------------------------------------------------------
------------------------------------------------------------------------
------------------------------------------------------------------------
------------------------------------------------------------------------
------------------------------------------------------------------------
-------------------------------------------------UGA-----GGACA-ACGAUAGGU
AGGCAGUUUGGGUGGGGCGCCACACCCUCGA-AAAAAUAUC-AAGGGUGCCCU-AUGGUCAACUCAUGUGAG
UCAGAAACUCACAGGA-GAGUGUCAAGAGCAU-AAGUUGGCCUGACG-CGAU--------------------
--------------UUCGCAUAGCAAGAAAUCGCGAGAGGAAA-CUCGGGUCUAACGAACCAAU--------
---GA-G-----CCUUA-UUG--------------------AU-GAGGGCCAUUGACGACAG-AA--AAGCU
ACCCCGGGG-AUAACAGAGUCGUCGCCGGCAAGAGCACAUAUCGACCCGGCGGCUUGCUACCUCGAUGUCGG
UUCUUUCCAUCCUGGCCGUGCA-GCAGCGGCCAAGGGUGAGGUUGUUCGCCUAUUAAAGGGGAUCGUGAGCU
GGGUUUAGACCGUCGUGAGACAGGUCGGUUACUAUCUAUC--------------------U-GGA-------
-------------------------------GUGU--AAGGAGUCUGAAGGUAAGA-UGGAAAUAGUACGAG
AGGAA-CUUUCCGUCGUCGCCUC-UGGU-C-GAUCGGUUGUCC-GACA-GGGCAG-U-GCCGAGCAGCUACG
CGAUA-AGGGAUAAAAGCUGAAAGCA-UC--UAAGCUUGAAACCUGACCCGA-A-A-A--GAGACUCC-GU-
------------------------------------------------------------------------
------------------------------------------------------------------------
------------------------------------------------------------------------
------------------------------------------------------------------------
------------------------------------------------------------------------
------------------------------------------------------------------------
------------------------------------------------------------------------
------------------------------------------------------------------------
----------------------------------------------------------UG-AGG------AC
ACGGGUAAAAGACCCGCU-UGA-UAGGUC-CGG-GAUGUAAG-CAC-G--------AAG-----GC------
----AA--------CGACGUGUUCAGUCC--G-CGGA-UACUAA-C---GUC--------------------
------------------------------------------------------------------------
------------------------------------------------------------------------
------------------------------------------------------------------------
-------------------------------------------------
```

```
>LSUa_456442_METB6/1-2899
----------------------------------------UAAACCUACC-AGUGGAUG-G-CUCGGU-UCA
A-G---UGCCGAAGAAGGGCGUGC--CAAGCUGCGAUAAGCUCCGGGUAGACGCAAGGAAUCUA------UG
AUC-CGG-AGAUGCCCUAAUAGGACA-U--------CCU-------GA----CGCU---------------U
U-A----------------------U----G---CGU----C-U-UUCC-----------------------
-----------G-----------UAA----------------------------------------------
------------------------G-G-AA-AGGGAACGCCCCGAAUUGAAACAUCUCAGUAGGGG--CAGG
AGAAGAAAUCAA--CCGAGAUGUCGUUA--GU-AG-AGGCGAUCGAACACGAC-A-GA-GCUCAAACCGAAU
CCCG----------------------------------CA-AGGGAGAUGUGGUG--UUA------------
U-AA--GCC-CAC-C-----G------U------------------ACGAU--GCC-----------GG--A
U-------GUGAA--GCGG------AAGUUG-CCUG------------------------------------
-------GAA-GG-GCACA-------CUACAGAGGGUGAAAGUCCCGUA---------CGC---A----GAA
G------CAGACGGU-GU------------------UAGG-UGGUG--UCUUGAGUACCGCGGGUUG---GA
AUUCUCGCGGGAA-UUCAGGGGUCAUC--AACCCCUAAAGCUAAAUA--CAC-C-UUG-AAACCGAUAGC-G
AAU-U-AGUAGCGUGAGCGAAAG-CUGAAA-AGUAA-CCCUGAAAAGGU-G-GUU-AAAAGUGCCUGAAAUU
G-GUAGGUGAUCG-CGG-GU-UGCG---GCAUGAAA-GGAU-------------------------------
------------------------------------------------------------------------
------------------------------------------------------------------------
------------------------------------------------------------------------
------------------------------------------------------------------------
------------------------------------------------------------------------
------------------------------------------------------------CU-UCCUUAUG-
---AA-----------GG-A-------------------A------CCC-----------------GUC---
------------------------------------------------------------------------
--------------------------------------------------------------G--C-GA---
------------------------------------------------------------------------
-------------------------------------------------G-GC------GG--------UAG
UACGG-------------------------------------------------------------------
------------------------------------------------------------------------
------------------------------------------------------------------------
------------------------------------------------------------------------
------------------------------------------------G-----------------------
------------------------------------------------------------------------
------------------------------------------------------------------------
------------------------------------------------------------------------
------------------------------------------------------------------------
------------------------------------------------------------------------
-------------------UAAGGU--U-----------------GCCAG--U---GUCGCAACGUACGUUU
UGAAGAACGGGCCAGAGAGUUUA-U-UCUGUUGGCGAAGGUUAACC-------------CACAA--------
---G--GGGA--GCCAUAGGGAAAC-CAA-----------------CAA----------------------G
UCCG-UAGCAGC----A-------------------------------------------------------
------------------------------------------------------------------------
--------------------AUG------------------------------------CGU-GGGACG-GC
GUAU-----------------------------------UAAA--------------------------GU-
G-C-GU----------------------GGAGUCAGCAGGAUAA-GACCCGAAGCC-GAGUGAUCUAUGCGU
```

```
UGGCAGGUUGAAGCGUGA-CGAAAG-UUGCGUGGAGGACCGC-AAGCCGUAUUGAUAUGCAAAUCAU-UGGU
GUGACCUACGUAUAGGAGUGAAAGAUUAAUCGAACCCGG-CAUCAGCUGGUUCCUCCCUAAA-CAUGUCGCA
GCAUGACCUAAC-CGGA-------------------------------------------------------
----------------------GA-----UA-GUCGGUGAGGUAGAGC-ACUGAUU-GGG-GGAG-CUGGGC
G-G-GA-AA-CUGC-U-CACCCCCCUGUCAAACUCCAAACUCCCCG--------------------------
------------------------------------------------------------------------
------------------A-CA-U--CC-UAGA--UGGUUGGAAGUCCGC-A-UUAC-G-GGG-UAAGCUUG
UA-A-UGCGUAAG-GGAAAC---AACCCAGACCGU-G-GUUAAUGUCCCUCAAU--GU-AGG-CU-CA---G
-U-GUCAACA-CUGAAAUUGGUCCUAGGUCAAAGACAACUGGAAGGUGAGCUCAGAAGCAGCUA-UCCUUUA
AAAAGUGCGUAACAGCUUACCAGUC-----AAGAUUUAGGGCGGUGAAAAU-GGAC-GGGGCUAA-AGCCUA
C----------UACAGAG-ACCACGGAGCACCGA--------------------------------------
------------------------------------------------------------------------
------------------------------------------------------------------------
--------------------------------AAGGU-GAUC-UGGUAGGGAGGC-GCUCUGC---------
-----------G-U-GGGUA-GAAGCUG--G-GGCGUGAGUU--C-CAGUGGACCG-C-GUA-GUGAUGAGA
AUUC-UGGCAG-UAGUA-GAAGCA--UAGUU-GGGUGA--G---------------AAUCCC-AAC-CGCCG
C-AGGGGC-UAGGUUUCC-UCGGCAACG-UUCGUCAGCCGAGGGUUAGUCGG--UC-CUAAGGUGCGUCGU-
AAC-UCGA---A-CGU--------------------------------------------------------
----------------------------------------ACC-GA-A-UGGGA-AACAGGUU-AAUA-UUC
CUGUA----------------------------------CCAUG----UGGUUUUA-------A--------
------------------------------------------------------------------------
-------GUC--------------------------G--------------------CAAG-A-CCCU-GAC
GCUUUGG-------------GAUAUG-CUCAGCGGG-AC-------UGUC-------------A-UCCCGU-
CCAA-G-CAC-AU-----A-A-------------------AC-----UG----------------GUGG-AG
U-A-CC--------------------GUAAU-GGUGAGAAUC---------CGGUGA----AAG-UG-UGA-
UG---GG-----------------------------------------------------------------
----------------G-UA-A-CU-A-AGC-A-A-AUUCCUUGAGCCC-AUGAAA-A-GGG-AAU------
---------------------------------------------------------------CA-CAUGUC
CGUACCGAGAACCGAC------------------------------ACAGGUGC-CCCUCGCU-GAAGAGGU
GUAGGC-GUGU-CGGA-UUUAAUCG-G-UUAAGGGAACUCGGC-AAAUUGGCCCCGUAACUUUGG-GAUAAG
GGGUG-CCUG-C-CU--GGA----------------------------------------------------
------------------------------------------------------------------------
------------------------------------------------------------------------
------------------------------------------------------------------------
------------------------------------------------------------------------
------------------------------------------------------------------------
------------------------------------------------------------------------
------------------------------------------------------------------------
------------------------------------------------------------------------
------------------------------------------------------------------------
------------------------------------------------------------------------
------------------------------------------------------------------------
------------------------------------------------------------------------
------------------------------------------------------------------------
------------------------------------------------------------------GAUCAG
```

```
GCA--GGUCGCAGUGACCGGGGCGCACUAA-----CUGUCUAAUAACAACAUAGGUGACUGCAACUCCG---
-AA-A-------------GGACUAGU-AUAGUCACUGA-UUCCUGCCCAGUGCGA-G-UAUC-UGAAUACCG
GG-------------UUCA-------------ACCGGACGAAGGAC--UCGUAAAC-GGCGGGGGUAACUAU
GACCCUCUUAAGGUAGCGUAGUACCUUGUCGCU-UAAUUGGCGACUUGCAUGAAUGGAUUAAUGAGU--GCG
CUACUGUCCCUAAC----ACG-AAUCCGUUGAACCU--UUU---GUUCUAGUGCAGAGACUAGAGACUCCUG
AUGGGAAGUGAAGA-CCCCGUUGAGC-UUUACU--GCAGCCU-GUCGUUGAAUUACGAUAAUUCUUGCGCAG
UGU-AGAUGGGAGGCGUUA----------UAUCCAU-UGC----------------------------UUUG
--------------------------------------GU--AAUGGGGAG-C--C-AA-UGAUGAGACA-C
CAUC-CUAGUUUUGUUGUAGUUCUCACUC------------UA-A-C-------------------------
------------------------------------------------------------------------
------------------------------------------------------------------------
------------------------------------------------------------------------
------------------------------------------------------------------------
------------------------------------------------------------------------
--------------------------------------------------GA-----GGACA-CCGGUAGGU
GGGCAGUUUGGGUGGGGCGCCACACCCUCGA-AAAAAUAUC-AAGGGUGCCCC-AAGGUCAACUCAUGUGAG
UCAGAAACUCACAGAA-GAGUGUCAAGAGCAU-AAGUUGGCCUGACG-CGAU--------------------
--------------CACGCAUAGCAAGUGAUCGCGAGAGGAAA-CUCGGGUCUAACGAACCAAU--------
---AU-A-----GCCUU-UUG--------------------GU-GAGGCUUAUUGACUACAG-AA--AAGCU
ACCACGGGG-AUAACAGAGUCGUCGCCGGCAAGAGCACAUAUCGACCCGGCGGCUUGCUACCUCGAUGUCGG
UUCUUUCCAUCCUGGCUGUGCA-GCAGCAGCCAAGGGUGAGGUUGUUCGCCUAUUAAAGGGAUCGUGAGCU
GGGUUUAGACCGUCGUGAGACAGGUCGGUUACUAUCUAUC--------------------A-GGA-------
-------------------------------GUGU--UAGAAGUCUGAAGGUAAGA-AUGAAAUAGUACGAG
AGGAA-CUUUCAUUCGGCGCCUC-UAGU-C-GAUCGGUUGUCU-GACA-AGGCAG-U-GCCGAGCAGCCACG
CGCCA-AGGGAUAAAAGCUGAAAGCA-UC--UAAGCUUGAAACCCGGCCUAA-A-A-A--GAGACUUC-GU-
------------------------------------------------------------------------
------------------------------------------------------------------------
------------------------------------------------------------------------
------------------------------------------------------------------------
------------------------------------------------------------------------
------------------------------------------------------------------------
------------------------------------------------------------------------
------------------------------------------------------------------------
----------------------------------------------------------UACAGG------AC
AUGGGUAAAAGACCCGUU-UGA-UAGGCU-UCG-GAUGUACG-CAC-G--------AAG-----GC------
----AA--------CGACGUGUUCAGUCC--G-GAAG-UACUAA-C---GUCCAAACA--------------
------------------------------------------------------------------------
------------------------------------------------------------------------
------------------------------------------------------------------------
-------------------------------------------------
>LSUa_323259_METHJ/1-2910
------------------------------U----UC--UUACGCCUGUC-AGUGGAUG-G-CUCGGU-UCG
G-G---UGCCGACGAAGGGCGUGC--CAAGCUGCGAUAAGCUCCGGGUAGACGCAUGGAGUCAU------UG
AAC-CGG-AGAUCCCCGAAUAGGACA-U--------CCA-------GA----UGC-------------A-UA
A-A---------------------------G---CAU----CAU-UCCG-----------------------
```

```
-----------C-----------CAU----------------------------------------------
------------------------C-G-GAAAGGGAACGCCCCGAAUUGAAACAUCUUAGUAGGGG--CAGG
AGAAGAAACCAA--CCGGGAUGUCGUUA--GU-AG-AGGCGAUCGAACACGAC-A-GA-GUUCAAACCGAAU
CCUU-------------------------------------CGGGAGAUGUGGUG--UAU------------
G-GA--CCG-CGG-C-----A-------------------------UAAGA--UUU-----------GC--A
U-------UCAGA--ACUG------AAGUUA-CCUG------------------------------------
-------GAA-CG-GUAUA-------CCAGAGAGGGUGACAGUCCCGUA---------UGU---G----UAU
G------CAUGCAGA-CU------------------UUAG-CGGUA--UCCUGAGUAGCGUGGGUCG---GA
AUUCCCGCGUGAA-UGUGGGGGUCAUC--AACCUCCAAAACUAAAUA--CUC-C-CCG-AAACCGAUAGC-G
UAG-U-AGUAGCGUGAGCGAAAG-CUGAAA-AGCAA-CCCUGGAAAGGU-G-UUG-AAAAGUGCCUGAAACU
G-ACAGGUUAUCG-CGU-GA-UACG---GCACGAAA-GGAU-------------------------------
------------------------------------------------------------------------
------------------------------------------------------------------------
------------------------------------------------------------------------
------------------------------------------------------------------------
------------------------------------------------------------------------
------------------------------------------------------------CU-UCAAAGCG-
---AA-----------GG-A-------------------A------CGA-----------------GUU---
------------------------------------------------------------------------
--------------------------------------------------------------G--C-GA---
------------------------------------------------------------------------
-------------------------------------------------G-AC------UC--------AAG
UACGG-------------------------------------------------------------------
------------------------------------------------------------------------
------------------------------------------------------------------------
------------------------------------------------------------------------
------------------------------------------------G-----------------------
------------------------------------------------------------------------
------------------------------------------------------------------------
------------------------------------------------------------------------
------------------------------------------------------------------------
------------------------------------------------------------------------
-------------------UUUUGU--U-----------------GCCGG--U---GUCGUAUCGUACGUUU
UGAAGAACGGGCCAGAGAGUUUA-U-UCUGUUGGCGACGGUUAAUU--U----------CUAUG--------
--AA--AGAA--GCCAGAGGGAAAC-CAA-----------------CAA----------------------G
UCCG-CAGCCGU----A-------------------------------------------------------
------------------------------------------------------------------------
--------------------AGG------------------------------------UCA-GGGACG-AC
GUAC-----------------------------------UACCA-------------------------GU-
G-C-GU----------------------GGAGUCAGCAGGAUAA-GACCCGAAGCC-CGGCGAUCUAUGCGU
GGGCAGGUUGAAGCGUGC-CGAAAG-GUGCGUGGAGGACCGC-AAGCGGUUUUGAUAUGCAAAUCAU-UCGC
GUGACCUGCGUAUAGGAGUGAAAGAUUAAUCGAGCCGGG-CAUCAGCUGGUUCCUCUCGAAA-CAUGCCGUA
GCAUGACCUGAU-CUGA-------------------------------------------------------
----------------------GA-----UC-GACAGUGAGGUAGAGC-ACUGAUU-GGG-GAAG-CUGGGG
A-A-GA-AA-UUCC-U-CACUCUCCUGUCAAACUCCAAAUUCCCUG--------------------------
```

```
------------------------------------------------------------------------
------------------U-CA-U--CA-GCGA--CGAUCGGAAGUCCGC-A-UUAC-G-GGG-UAAGCUUG
UA-A-UGCGUAAG-GGAGAC---AACCCAGACCGU-G-GUUAAUGUCCCUCAGU--GC-AGG-CU-CA---G
-U-GUAAACA-CUGAAAGUAGUCCUGGGUCAAAGACAACUGGGAGGUGAGCUUAGAAGCAGCUA-CCCUUUA
AAAAGUGCGUAACAGCUUACCAGUC-----AAGAUUCAGGGCGCUGAAAAU-GGAC-GGGGCUUA-AGCCUG
C----------CACAGAU-ACCACGGACCAUACG--A-----------------------------------
------------------------------------------------------------------------
-----AU-----------------------------------------------------------------
--------------------------------UGUAU-GAUG-GCGUAGAGAGGC-GUCCUGC---------
-----------A-U-GGGCG-GAAGCAG--G-GUUGCAAGAU--C-CUGUGGACCG-U-GCA-GGAAUGAAA
AUUC-UGGCAG-UAGUA-GAAGCU-UAAGAU-UGGUGA--G---------------AAUCCA-AUC-CGCCG
C-AGGGGC-UAGGUUUCC-UCGACAAUG-UUCGUCAGUCGAGGGUUAGUCGG--UC-CUAAGACGUACCGU-
AAU-UCGA---G-UAC--------------------------------------------------------
----------------------------------------GCC-GA-A-AGGGA-AACAGGUU-AAUA-UUC
CUGUA----------------------------------CCUGU----AUCA--------------------
------------------------------------------------------------------------
---------------------------------------------------------AACA-A-UCCU-GAC
GCUUUCG-------------GAUAGG-CAUUGCGGA-GU-------CAUC-------------G-CUCCGU-
CUAA-GCCGG-AU-----A-U-------------------GA-----AA----------------UUGG-AG
U-A-CC--------------------GUAAU-GGUGAGAAUU----------UUUCAA----AAC-GG-UGA-
UG---GG-----------------------------------------------------------------
----------------G-UA-A-CA-A-UGC-U-G-AUUCCAGGAGCCC-GUGAAA-A-GGG-UG-------
---------------------------------------------------------------AU-ACGGUC
CGUACCGAGAACUGAC------------------------------ACAGGUGC-CCCUCGCU-GAGAAGGC
GAAGGC-GUGU-CGGGAGUAAAUGU-G-UUAAGGGAACUCGGC-AAAUUAGCCCUGUAACUUGG-GAUAAG
GGGUG-CCUG-C-CC--AGC----------------------------------------------------
------------------------------------------------------------------------
------------------------------------------------------------------------
------------------------------------------------------------------------
------------------------------------------------------------------------
------------------------------------------------------------------------
------------------------------------------------------------------------
------------------------------------------------------------------------
------------------------------------------------------------------------
------------------------------------------------------------------------
------------------------------------------------------------------------
------------------------------------------------------------------------
------------------------------------------------------------------------
------------------------------------------------------------------------
------------------------------------------------------------------GAUUGG
GCA--GGUCGCAGUGACAAGAUCGCUCCGA-----CUGUCUAAUAACAACAUAGCAGACUGCAACUCCG---
-AA-A-------------GGACUCGU-AUAGUCUGUGA-UUCCUGCCAGUGCGA-G-UAUC-UGAACACCG
GG-------------UUCA-------------ACCGGACGAAGGAC--UCGUAAAC-GGCGGGGGUAACUAU
GACCCUCUUAAGGUAGCGUAGUACCUUGUCGCU-UAAUUGGCGACUUGCAUGAAUGGAAUAACGAGA--GCG
AUACUGUCCCUAAC----ACA-UACCCGUUGAACCU--UUU---GUACUAGUGCAGAGACUAGUGACUCCUU
```

```
AUGGGAAGUGAAGA-CCCCGUGGAGC-UUUACU--GCAGCCU-GUCGUUGGGUUACGGUAUUACCUGCGCAG
GGU-AGAUGGGAGACGUCG----------AUCCAAC-CCU----------------------------UGUG
--------------------------------------GG-GGUUGGUUAG-U--C-AC-CGAUGAGACA-C
CAUC-CUGGUUUUAUUGUAAUUCUAACUG-A----------UU-U-A-------------------------
------------------------------------------------------------------------
------------------------------------------------------------------------
------------------------------------------------------------------------
------------------------------------------------------------------------
------------------------------------------------------------------------
-------------------------------------------------UCA-----GGACA-UCGAUAGGU
AGGCAGUUUGGGUGGGGCGCCACACCCUCGA-AAAAAUAUC-AAGGGUGCCCU-AAGGUCAACUCAAGCGGG
UCAGAAACCCGCUGAA-GAGUGUCAAGAGCAA-AAGUUGGCCUGACG-CGAU--------------------
--------------UUCGCAUAGCAAGAAAUCGCGAGAGGAAA-CUCGGGUCUAACGAACCAAU--------
---AC-G-----CCCUG-UUG--------------------AU-GAGGGCUAUUGACUACAG-AA--AAGCU
ACCCCGGGG-AUAACAGAGUCGUCGCCGGCAAGAGCACAUAUCGACCCGGCGGCUUGCUACCUCGAUGUCGG
UUCUUUCCAUCCUGGCUGUGCA-GCAGCAGCCAAGGGUGAGGUUGUUCGCCUAUUAAAGGGGAUCGUGAGCU
GGGUUUAGACCGUCGUGAGACAGGUCGGUUACUAUCUAUA--------------------A-GGA-------
-------------------------------GUGU--UGGAAGUCUGAAGGCAAGA-AUGAAAUAGUACGAG
AGGAA-CUUUCAUUCGGCGCCAC-UGGU-C-UACCGGUUGUCU-GACA-AGGCAA-C-GCCGGGCAGCUACG
CGCCA-AGAGAUAAAAGCUGAAAGCA-UC--UAAGCUUGAAACUCAGCCUGA-ACA-A--GAGACUUC-AU-
------------------------------------------------------------------------
------------------------------------------------------------------------
------------------------------------------------------------------------
------------------------------------------------------------------------
------------------------------------------------------------------------
------------------------------------------------------------------------
------------------------------------------------------------------------
------------------------------------------------------------------------
----------------------------------------------------------GA-AAG------AC
ACGGGUAAAAGACCCGGU-UGA-UAGGUU-CGG-GAUGUACG-CAC-G--------AAG-----GC------
----AA--------CGACGUGUUCAGUCC--G-CGAA-UACUAAUC---GUCUUAAUA-ACGUC--------
------------------------------------------------------------------------
------------------------------------------------------------------------
------------------------------------------------------------------------
-------------------------------------------------
>LSUa_187420_METTH/1-3035
----------------------CU------U----AU--UUAUGCCGUCU-GGGGGAUG-G-CUUGGC-UUG
AGU---CGCUGAUGAAGGCCGUGG--CAAGCUGCGAUAAGCCCAGGGGAGGAGCAUGCAUCCUU------GG
AUC-CUG-GGAUUGCCGAAUGGGACU-U--------CCU-------GG----CCGU-C-CCUGCAC-----U
C-U-------GUGCGGG---G----G----C--UGGU----C-A-CUCC-CCU-----U-------------
-----------U-----------UCU---------------------------------------------U
G-G--------------------GG-G-AG-GGGGAACCCGCUGAACUGAAACAUCUUAGUAGGCG--GAGG
AAGAGAAAGCAAU-AUGCGAUGCCGUGA--GU-AA-UGGCGAAUGAAAGCGGU-G-GA-GGACAAACUGAAU
CCAU-CAUGGU--------GA------------C---GUG-GUGGAGAUGUGGUG--UUG------------
C-GG--ACC-CCC-U-----G------CU-----------------UAGGG--GUU-----------CC--A
```

```
G-------GUGUG--GUUGGGUUGAACUUGG-GCUG------------------------------------
-------GAA-UGCCCGGG-------CCGUAGAGGGUUAAAGCCCGUA----------GAC---U----UGC
U------GUGCCUGG-CCCUG---------------CGGG-GGUGU--UCCUGAGUAGCGUCCAUUG---GA
UAUUGGGCGUGAA-GCUGGGAGGCAUC--GACUCCUAAUCCUAAACA--CGU-C-UCA-AGUCCGAUAGC-G
AAC-U-AGUACCGUGAGGGAAAG-CUGAAA-AGUAC-CCCUGAUAGGGG-U-GUG-AAAAGUGCCUGAAACC
A-GGCGGUGACAG-CCC-GG-CACG---GCAUGGAA-GGAAUG-----------------------------
------------------------------------------------------------------------
------------------------------------------------------------------------
------------------------------------------------------------------------
------------------------------------------------------------------------
------------------------------------------------------------------------
-----------------------------------------------------------AGGCUUGCCCUG-
---UA-----------AG-A-------------------A------ACC-----------------AUG---
------------------------------------------------------------------------
--------------------------------------------------------------G--C-AA---
------------------------------------------------------------------------
-------------------------------------------------C-AU------GG--------GAG
UAUGU-------------------------------------------------------------------
------------------------------------------------------------------------
------------------------------------------------------------------------
------------------------------------------------------------------------
------------------------------------------------G-----------------------
------------------------------------------------------------------------
------------------------------------------------------------------------
------------------------------------------------------------------------
------------------------------------------------------------------------
------------------------------------------------------------------------
-------------------GGCUGGCUG-----------------AUCAG--U---GUCGUGUCAUCCGUCU
UGAAACACGGGCCAGGGAGUUUA-GUGGUUGUGGCGA-GACUAAGA--A-GU-------GUGUC-------G
C-UU--UGAA--GUCGUAGGGAAAC-CGA-----------------CAG----------------------G
UCCG-CAGCAGC----A-------------------------------------------------------
---UC-------------------------------------------------------------------
--------------UGU--GCUG------------------------------------UGA-GGGACG-GG
GUCU-----------------------------------UAAUA-------------------------GG-
G-C-CU----------------------GGAGUCACAGCCCUAA-AACCCGAAGCC-GGUCGAUCUAGCCCU
GGGUAGGAUGAAGUCGCU-CUUACG-AGUGAUGGAGGUCCGC--AGGGGUGUUGUCGUGCGAAACAU-UCCU
CUAACCUGGGGUUAGUGGUGAAAGGCCAAUCAAGGCCGG-UGACAGCUGGUUCCACCCGAAA-UGGCUCGUA
GGCCAGCCUGAC-UGGA-------------------------------------------------------
----------------------GG-----UU-GGUGGCGGGGUAGAGC-ACUUAUU-GGG-UGUU-UAGGGG
G-A-GA-AA-UCCC-U-CGGCAUCCUGUAAAACUCCGAACUCGUCA--------------------------
------------------------------------------------------------------------
------------------C-CG-C--UG-UUGA--AGGUUGG-AGUCAGG-G-GCGC-G-GGG-UAAGCCUG
UG-U-CCCGAGAG-AGGAAC---AACUCAGACUGG-G-GUUAAGGUCCCUAAAU--GC-CGG-CU-AA---G
-U-CU---------AAGGGGGUCUUUGGCCCUAGACAAUGGGAAGGUGGGCUUAGAAGCAGCCA-UCCUUUA
AAGAGUUCGUAACAGAUCACCCAUC-----GAGGUCAAAGGCACCGAAAAU-GGAG-GGAAUUA-AGCCGG
```

C----------UACCGAU-ACCUCAGAGCACCAC--U-----------------------------------
------------------------------------------------------------------------
-----U--UUG-------------------------------------------------------------
-------------------------------UGUGGU-GGUC-UUGUAGGGUGGC-GUCCCGU---------
-----------U-G-GGGUU-GAAGUGG--G-GGCGUGAGCU--C-CUGUGGACCC-U-GCG-GGAAUGAGG
AUCC-UGGUAG-UAGUA-GCAGCA--AAGUG-AGGUGA--G---------------AAUCCU-UAC-CGCCG
G-AGGGGC-UAGGGUUCC-UUGGCAAUG-UUCGUCAGCCAAGGGUUAAUCGG--UC-CUAAGGCCAUGGGU-
AAUGUCCA---U-GUG--------------------------------------------------------
----------------------------------------GUC-GA-A-AGGGU-AACAGGUU-AAUA-UUC
CUGUA----------------------------------CGGUC----CAGGUACU-------U--------
------------------------------------------------------------------------
-------GCGG--UGAC--------GCUGG-----GU--------------------UGGG-C-UUCU-GAC
GCUUCGG-------------GGUAGG-CUGAGCGGG-AU-------UUUC-------------G-UCCUGU-
UUAA-G-GGU-UG-----A-A-------------------GC-----CU----------------GGGG-AG
A-G-CC--------------------GUAAU-GGCGAGAACU-------AUUGGUGA----AGG-CC-UGA-
AUA--GC-----------------------------------------------------------------
-----CACUCC-UUGUUGGGUGGUU-C-GGC-U-G-UGCCCUGGAGUCC-UUGAAA-A-GGG-AGUCCUUCU
-U-----------------------------------------------------GGGAUCC-UG-GAUCGC
CGUACCGAGAUCCGAC------------------------------ACUGGUGC-CCCUAGCU-GAGUAGGC
UAAGGC-GUGU-UGGG-GUAACCUG-G-CUAAGGGAAAUCGGC-AAAUUAGCCCCGUAACUUUGG-GAGAAG
GGGUG-CCAG-C-CA--UGU----------------------------------------------------
------------------------------------------------------------------------
------------------------------------------------------------------------
------------------------------------------------------------------------
------------------------------------------------------------------------
------------------------------------------------------------------------
------------------------------------------------------------------------
------------------------------------------------------------------------
------------------------------------------------------------------------
------------------------------------------------------------------------
------------------------------------------------------------------------
------------------------------------------------------------------------
------------------------------------------------------------------------
------------------------------------------------------------------------
-------------------------------------------------------------------AGAUG
GCU--GGUCGCAGUGACAGGGGGGGCCCGA-----CUGUUUAAUAAAAACAUAGCUCCUAGCUAGCCCG---
-UG-A-------------GGGUGUGU-ACUGGGGGCGA-CACCUGCCCAGUGCCG-G-CACG-UGAAGCCCU
GG-------------UUCA-------------ACGGGGUGAAGCGC--CGGUAAAC-GGCGGGGGUAACUAU
AACCCUCUUAAGGUAGCGAAAUGCCUUGCCGGA-UAAGUACCGGCCUGCAUGAAUGGUUGAACGAGG--UCC
CUACUGUCCCUAGC----CAG-GACCUGGUGAAGCU--GCU---GUUCUGGUGCACAAGCCAGAGACUCCCA
GUGGGAAGCGAAGA-CCCCGUAGAGC-UUUACU--GCAGUCU-GCUGUUGGGGCUUGGUCAUGGGUAUGCAG
UGU-AGGUGGGAGGUGUCG----------AUGCCAUGGUC----------------------------GCCA
--------------------------------------GGCCGUGGUGGAG-C--C-GG-UCAUGAGACA-C
CACC-UUCCUGUGACUGUGUCUCUAACCC-CU--CG-----UU-U-U-------------------------
------------------------------------------------------------------------

------------------------------------------------------------------------
------------------------------------------------------------------------
------------------------------------------------------------------------
------------------------------------------------------------UG----------
------------------------------------------------GGGG-----GGACA-UCGGUAGAU
GGGCAGUUUGGCUGGGGCGGCACGCGCUUGA-AAUGGUAUC-AAGCGCGCCCU-AAGGUCGGCUCAGGCGGG
ACAGAGAUCCGCUGUA-GAGUG-UAAGGGCAU-AAGCCGGCUUGACU-GUGC--------------------
--------------UCCUACUAGUAGGGGGUGCAGGUGCGAGA-GCAGGGCCUAGCGAACCCCA--------
---GA-G-----UCCUCGUCG--------------------GU-GGGGGCCUGGGAUGACAG-AA--AAGCU
ACCUCGGGG-AUAACUGGGUGGUCGCAGGCAAGAGCCCAUAUCGACCCUGCGGCUUGCUACUUCGAUGUCGG
UUCUUUCCAUCCUGGGUGUGCA-GCAGCACCAAAGGGUGGGGUUGUUCGCCCAUUAAAGGGGAACGUGAGCU
GGGUUUAGACCGUCGUGAGACAGGUUGGUUGCUAUCUACU--------------------G-GGA-------
-------------------------------GUGUG-UGGUUGCCUGAGGGGAAGG-UGGUUCCAGUACGAG
AGGAA-CGGACCGUCGGCGCCUC-UGGU-U-UACCGGUUAUCC-GAGU-GGGUAU-U-GCCGGGCGGCUACG
CGCUAUGAUUAUAAAGGCUGAAGGCA-UC--UAAGCCUGAGGUUUUCCCUGA-A-AAU--AGGCGGCU-UG-
------------------------------------------------------------------------
------------------------------------------------------------------------
------------------------------------------------------------------------
------------------------------------------------------------------------
------------------------------------------------------------------------
------------------------------------------------------------------------
------------------------------------------------------------------------
------------------------------------------------------------------------
----------------------------------------------------------U--GGA------CC
GCGGGUAGAAGACCUGUU-UGU-UGGGGC-GGG-GGUGUGAG-CUU-C--------GAG-----GCCUGUUU
UUGGGU--------CGAGUUGUUUAGCCU--G-CCGU-UUCCAAGG---UUUUUUGUC-CCU----------
------------------------------------------------------------------------
------------------------------------------------------------------------
------------------------------------------------------------------------
-------------------------------------------------
>LSUa_339860_METST/1-2994
----------------------------------------UAUGCCAUCU-GGGGGAUG-G-CUAGGC-UUG
A-GAUUAACUUAUGAAGGUCGUGG--CAAGCUGCGAUAAGCCUGGGCGAGAAGCAUGCAUUCAU------UG
AAC-CUG-GGAUUGCCGAAUGAGAUU-C--------UU--------AU----UACU-C-U----------UU
A-C-------------C---G----A----G---UAA----U-G-AUCC-AC--------------------
-----------A-----------CUU----------------------------------------------
--G--------------------UG-G-AU-UGGGAACCCACUGAAUUGAAACAUCUUAGUAGGUG--GAGG
AAGAGAAAACAA--UUGUGAUGCCGUUA--GU-AA-CGUCGAGCGAAAACGGU-A-UA-GGACAAACUGAAU
CCCU-ACUAGA-------AAU------------U---AAU-AGGGAGAUGUGGUG--UUG------------
A-AG--AUU-CAA-AAUAUUU------AU-----------------GGUGG--CCU-----------GU--U
A-------UUAUC--CUUG--UUUAAUUAUU-GUUG------------------------------------
-------GAA-UACUUUAA-------CCGUAGAGGGUGAUAGUCCCGUA---------GAC---U-U-UGGG
G------UAAUUUGG-UCAUU---------------UUUU-UUUGA--AUUCGAGUAGCGUUCAUUG---GA
AAUUGGGCGUGAA---UGGGAGGCAUC--GACUUCCAAUUCUAAACA--UUA-C-UCA-AGACCGAUAGU-G
UAU-UUAGUACCGCGAGGGAAUG-CUGAAA-AGAAC-CCUU--UUAGGG-G-GUG-AAAAGUACCUGAAACC

```
A-GAUGGUGACAG-CCU-GG-CAUG---GCAUGAAA-GGAAUG-----------------------------
------------------------------------------------------------------------
------------------------------------------------------------------------
------------------------------------------------------------------------
------------------------------------------------------------------------
------------------------------------------------------------------------
------------------------------------------------------------AU-AAUUCAUG-
---AA-----------U--U-------------------U------CUU-----------------CUG---
------------------------------------------------------------------------
--------------------------------------------------------------G--C-AA---
------------------------------------------------------------------------
-------------------------------------------------C-AC------UU--------GAA
-AUUC-------------------------------------------------------------------
------------------------------------------------------------------------
------------------------------------------------------------------------
------------------------------------------------------------------------
------------------------------------------------U-----------------------
------------------------------------------------------------------------
------------------------------------------------------------------------
------------------------------------------------------------------------
------------------------------------------------------------------------
------------------------------------------------------------------------
-------------------UUGUCUUUG-----------------ACUAG--U---GUUGUGUCAUUCGUCU
UGAAACACGAGCCAGGGAGUUUG-U-AGUUGUGGCGA-GGCUAAAA--A-GC--------UUAA-------G
C-UU--UGUA--GUCGCAGGGAAAC-CGA-----------------GUA----------------------A
GACG-UAACUU-------------------------------------------------------------
------------------------------------------------------------------------
---------------------UU------------------------------------UGU-GUGGUU-CU
GAUU-----------------------------------AUAA--------------------------AA-
U-G-UC----------------------AUAGUCACAGCUUUAA-AACCCGAAGCC-GGUCGAUCUAUCCGU
GAGCAGGAUGAAAUCGCU-CAUACG-AGUGAUGGAGGUUCGC--AGAGUUGUUGUUCUGCAAAACAC-UCUU
AUGACUUGCGGUUAGUGGUGAAAGGUCAAUCAAGAUCGG-UGACAGCUGGUUCCUAUCGAAA-UAGCUCCAA
GGCUAGACUGAU-UGGA-------------------------------------------------------
----------------------GA-----UA-GCUGGCAAGGUAGAGC-ACUGAUU-GAG-UGUU-UAGGGU
G-G-GA-AA-CUAC-U-CGGCAUUCUGUAAAACUCCAAACUUGCUA--------------------------
------------------------------------------------------------------------
------------------G-CA-U--CG-UAGA--AGAUCGG-AUCGAGG-G-GCGU-G-GGG-UAAGCCUG
UG-U-CCCGAAAG-AGAAAC---AACUCAGACUUU-G-GUUAAGGUCCCCAAGU--GC-CGG-AU-UA---G
-U-UU---------AAGGGAGUCAUUAGCCCUAGACAGUGGGGAGGUGGGCUUAGAAGCAGCCA-CCCUUUA
ACGAGUUCGUAACAGAUCACCCAUC-----GAGGUUAAUGGCAUCUAAAAU-GGAA-GGGG-UUA-AGUCGG
C----------CACCGAU-ACCAAUGAACAUAUA--C-----------------------------------
------------------------------------------------------------------------
-----U--UUU-------------------------------------------------------------
-------------------------------GUAUAU-GAUU-UCGUAGAUAGGC-GUCCUGU---------
-----------U-G-GAUUU-GAAGCUG--G-GGUGUGAAUU--C-CAGUGGAUCU-U-GCA-GGAAUGUGG
```

AUCC-UGGUAG-UAGUA-GCAGCA--AAGUA-AAGUGU--G---------------AAUCUU-UAU-CGCCG
A-AGGGGC-UAGGGUUCC-UUGGCAAUG-UUCGUCAGCCAAGGGUUAGGCGG--UC-CUAAGGGUAACGGU-
AAU-UCCA---G-UUA--------------------------------------------------------
----------------------------------------UUC-GA-A-AGGGA-AACAGGUU-AAUA-UUC
CUGUC----------------------------------CAACA----CAAUU-UA-------C--------
------------------------------------------------------------------------
-------ACGG--CAAC--------GUUUA---------------------------UGUU-G-GAUU-AAC
ACUUCGU-------------GAUAUG-CCUUGUAAG-AU-------UGUC-------------G-UCUUAU-
UCAA-AUUUU-AG-----A-A-------------------GC-----UA----------------AGGG-AG
A-A-CU--------------------GUAAU-AGUGAGAACU---------UGGUGU----AUU-UA-UGA-
UGA--GU-----------------------------------------------------------------
-----CUCUCG-UUUUUUUGAGAUU-U-GGU-U-G-AUUCAUGGAGUCC-UUGAAA-A-GAA-AAUAUCCAU
------------------------------------------------U-----UAUUUGAU-UG-UGUUUC
CGUACCGAGAUCCGAC------------------------------ACAGGUGC-CCCUAGUU-UAGUAGAC
UUAGGC-GUGU-UAGG-GUAAACCA-G-CUAAGGGAAUUCGGC-AAUUUAGCUCCGUAACUUUGG-GAUAAG
GAGUG-CCAG-C-UA--CGU----------------------------------------------------
------------------------------------------------------------------------
------------------------------------------------------------------------
------------------------------------------------------------------------
------------------------------------------------------------------------
------------------------------------------------------------------------
------------------------------------------------------------------------
------------------------------------------------------------------------
------------------------------------------------------------------------
------------------------------------------------------------------------
------------------------------------------------------------------------
------------------------------------------------------------------------
------------------------------------------------------------------------
------------------------------------------------------------------------
------------------------------------------------------------------GUAGUA
GCU--GGUCGCAGUGACUAGGGAGGUUCGA-----CUGUUUAAUAAAAACAUAGCUCCCAGCUAGCCCG---
-UA-A-------------GGGUGUGU-ACUGGGGGCGA-CACCUGCCCAGUGCUG-G-CACG-UGAAGCUGG
GG-------------UUCA-------------ACCUGGUGAAGCGC--CAGUAAAC-GGCGGGGGUAACUAU
AACCCUCUUAAGGUAGCGAAAUGCCUUGCCGGA-CAAGUACCGGCCUGCAUGAAUGGUAGAACGAAA--UCU
CCACUGUCCCUAGC----UGG-AACCUAGUGAACCU--GCU---AUUCUGGUGCACAAGCCAGAGACUUCUA
UUGGGAAGCGAAGA-CCCCGUAGAGU-UUUACU--GCAGUCU-GCUAUUGAGGCUUGGUUGCAAGUAUGCAG
CGU-AAGUAGGAGACUUCG----------ACGGUAU-GUC----------------------------GUCA
--------------------------------------GG-CAUAUCUGAG-U--C-AU-UCAUGAGACA-C
UACU-UUCUUGUGACUGUGUUUCUAAGCU-A--------U-UU-U-U-------------------------
------------------------------------------------------------------------
------------------------------------------------------------------------
------------------------------------------------------------------------
------------------------------------------------------------------------
------------------------------------------------------------------------
-------------------------------------------------UUA-----GGACA-UUGGUAGAU

```
GGGCAGUUCGGCUGGGGCGGUACGCGUUUGA-AAAGAUAUC-AAACGCGCCCA-AAGGUUGGUUCAAGUGGG
ACAGAGAUCCACUGUA-GAGUG-UAAGGGCAU-AAGCCAGCCUGACU-GUAU--------------------
--------------UUCGAACAGUAAGAAAUUCAGAGGCGAAA-GCCGGGCCUAACGAACCUCA--------
---AU-G-----UCCUCGUCG--------------------AU-GGGGGCUUGAGAUGACAG-AA--AAACU
ACCUCGGGG-AUAACUGGGUGGUCGCAGGCAAGAGCCCAUAUCGACCCUGCGGCUUGCUACUUCGAUGUCGG
UUCUUUCCAUCCUGGGUGUGCA-GCAGCACCCAAGGGUGGGGCUGUUCGCCCAUUAAAGGGGAACGUGAGCU
GGGUUUAGACCGUCGUGAGACAGGUUGGUUGCUAUCUAAU--------------------A-GAA-------
-------------------------------GUGUU-UUGUUGCUUGAGCGGAAGG-UGGUUCCAGUACGAG
AGGAA-CGAGCCGUCGGUGUCUC-UGGU-C-UACCAGUUGUCU-GAUA-AGGCAU-C-GCUGGGCAGCUACG
CACUAUGAUAAUAAAGGCUGAAAGCA-UC--UAAGCCUGAGGUAUUCCGUGA-A-AAU--AAGCAGCU-UU-
------------------------------------------------------------------------
------------------------------------------------------------------------
------------------------------------------------------------------------
------------------------------------------------------------------------
------------------------------------------------------------------------
------------------------------------------------------------------------
------------------------------------------------------------------------
------------------------------------------------------------------------
---------------------------------------------------------UUU-AGG------GU
AUGAGUAGAAGACUUGUU-UGA-UGGGAU-UGG-GAUGUAAG-CUU-C--------GAG-----AUUCUUU-
-GUUUU--------CGAGUUGUUAAGUCC--G-CAAU-UUCCAACA---CCCGAUUUU-AAUACAAU-----
------------------------------------------------------------------------
------------------------------------------------------------------------
------------------------------------------------------------------------
-------------------------------------------------
>LSUa_374847_KORCO/1-2991 CP000968.1, complete genome; RANGE 1328189:133021
------------------------GCUGGCA----CG--CUACGCCGCCU-GGUGGAUG-G-CUCGGC-UCG
G-G---CGCCGAGGAAGGGGGUGG--CAAGCUGCCAUAAGCCCCGGGGAGCCGCAGGCAGGCUU------CG
AUC-CGG-GGAUCCCCGAAUGGCGAU-G--------CCG-------CC----GAG-----------------
----------------------------------AGG----C-G-CUCC-CGG-----C-------------
------------------------------------------------------------------------
--A--------------------GG-G-AG-CGGGAACCCCCCGAAUUGAAGCAUCUCAGUAGGGG--GAGG
AGGAGAAACCAA--CCGGGAUGCCCUGA--GU-AC-CGGCGAGGGAAAGGGGC-U-GA-GCUCAAACCGAAC
UCCC-GCUCGC--------GA------------G---AGC-GGGUGGAUGUGGGG--UUG------------
C-GG--GGG-GAU-C-----------------------------GGGAGCC--GCU-----------GG--G
A-------GGGAA--GCCG------AAGUGG-CCUG------------------------------------
-------GAA-CG-GCCCG-------CCAUAGAGGGUGAGAGCCCCGUA---------GGC---G----UAA
C------CUACCAGC-GGU-----------------CCGG-UCCUC--UCCCGAGUACCGUGGGUUG---GA
AUACCCGCGGGAA-UCCGGGGGUCAUC--GACCUCCAAAGCUAAAUA--CGU-C-CCG-AGACCGAUAGC-G
AAC-UGAGUACCGUGAGGGAAAG-CUGAAA-AGCAC-CGCUG-GCGCGG-G-GUG-AAAAGAGCCUGAAACC
A-GGCGGUCAUAG-GA--GC-GGCG---GCCCUCAA-GGGCUG-----------------------------
------------------------------------------------------------------------
------------------------------------------------------------------------
------------------------------------------------------------------------
------------------------------------------------------------------------
```

```
------------------------------------------------------------------------
-----------------------------------------------------------AAG---CUCCCG-
---AA-----------GG-A-------------------A------AAU-----------------GCC---
------------------------------------------------------------------------
--------------------------------------------------------------G--C-GA---
------------------------------------------------------------------------
-------------------------------------------------G-GC------AU--------GAG
UACGA-------------------------------------------------------------------
------------------------------------------------------------------------
------------------------------------------------------------------------
------------------------------------------------------------------------
------------------------------------------------G-----------------------
------------------------------------------------------------------------
------------------------------------------------------------------------
------------------------------------------------------------------------
------------------------------------------------------------------------
------------------------------------------------------------------------
-------------------GGAGCG--G-----------------ACCGG--G---GUCGCCCCGUACGUCU
UGAAACACGCGCCGGGGAGUUCA-C-GGGCGUGGCGA-GCCUAAGG--G-C--------UUUAA--------
G-CC--CGAA--GGCGGAGCGAAAG-CGA-----------------GGA----------------------C
CCCG-CAGCGAU----G-------------------------------------------------------
---CG-------------------------------------------------------------------
------------------------------------------------------------AG-GGGGUC-GC
CUCC-----------------------------------GAAA--------------------------GG-
G-G-GU----------------------UCAGUCACGCUCGUGA-GACCAGAAGCC-GGGCGAUCUACUCCU
GGGCAGGGCGAAGCCGGGUCGCGAG-CCCGGUGGAGGCCCGA-UAGGGAUGGUGAGGUGCAAAUCCC-UCCC
GUGACCUGGGAGUAGGGGCAAAAGACCAAUCUAGCCCGG-CAAUAGCUGGUUCCCGCCGAAG-CCUCCCGCA
GGAGGAUCUCCG-CGGA-------------------------------------------------------
----------------------GG-----GU-GGGGGCGUGGUAGGGC-AACGAUU-GGG-UGGG-UAAGGG
G-C-UC----CGCC-C-CGCCACCCUCUCGAACCCCGAAUGCGCCU--------------------------
------------------------------------------------------------------------
------------------C-CC-C--CG-UAGA--AGCGGGG-AGCCGGG---CGGCGG-GGG-GAAGCUCC
GU-C-GCCGAGAG-GGGAAC---AACCCGGACCGG-G-GUUAAGGUCCCCAAGU--CC-UGG-CU-AA---G
-U-GCCGAAAUCCGAAGGGCGUCUCCCUCCUAAGACAGCGGGGAGGUAGGCUCAGAAGCAGCCA-UCCUUUA
AAGAGUGCGUAAAAGCUCACCCGCC-----GAGGAGGGAGGCCCCCAAAAUGAGUC-GGGGCUGA-AGCCAG
G----------CACCGAU-ACCCCGGCAGCGUCC--U-----------------------------------
------------------------------------------------------------------------
-----G------------------------------------------------------------------
-------------------------------GAAGGG-CGCU-GGGUAGGCGGGC-GCGCACA---------
-----------C-C-GGGCG-GAAGCCG--G-GGCGUGAGCU--C-CGGUGGACCG-G-UGU-GUGUUGCAA
AUCC-CGGCGG-UAGUA-GCAGCG--AAGUG-GGGUGA--G---------------AAUCCC-CAC-GGCCG
A-AAGCGC-CAGGGUUCG-CCGGCAAUGCUUCGUCAGCCGGCGGUUAGCCGG--UC-CUAAGGCGGCACCC-
AGC-UGGU---A-GCC--------------------------------------------------------
----------------------------------------GCC-GA-A-AGGGA-AGCCGGUU-AAUA-UUC
CGGCG----------------------------------CCACC----GGGGUAG-----------------
```

```
------------------------------------------------------------------------
-------GCGG--CAAC--------GC-AG-----GC--------------------CUCA-C-CCCU-GAC
GCCUCGG-------------GCUAGG-CUGAGCGGGGGC-C-----CGCUGAG----GCGGGCG-ACCCGU-
CCAA-G-CGC-UU-----A-A-------------------GG-----CC----------------GCCG-AG
A-A-CC--------------------GUAAU-GGUGAGAAGC---------GGCUGA----AGG-CG-CGA-
UGG--GG-----------------------------------------------------------------
-----GCCCC---UUUGGGGGCCUU-C-GGC-U-C-AUGCCUGGGGACC-UUGAAA-A-GGG-GGUGGGGAG
--------------------------------------------------------CGAUCC-CC-GGUGCC
CGUACCGAGAUCCGAC------------------------------ACAGGUGC-GCUGGCUG-AGAAGGCC
UAAGGC-CUAU-CGGUCCUAACGCG-G-GAGAGGGAACUCGGC-AAAUUGGCCCCGUAACUUCGG-GAGAUG
GGGUG-CCUG-C-GU--UCCU-----------------------------CC--------------------
------------------------------------------------------------------------
------------------------------------------------------------------------
------------------------------------------------------------------------
------------------------------------------------------------------------
------------------------------------------------------------------------
------------------------------------------------------------------------
------------------------------------------------------------------------
------------------------------------------------------------------------
------------------------------------------------------------------------
------------------------------------------------------------------------
------------------------------------------------------------------------
------------------------------------------------------------------------
------------------------------------------------------------------------
----------------------------------------------------------GAAAGG--GGGUCC
GCA--GGUCGCAGUGACAAGGGGCGCCCGA-----CUGUUUAACAAAAACAUAGUUCCCCGCGAGCCCG---
-CA-A-------------GGGUGUGU-ACGGGGGAUGA-GGCCUGGCCAAUGCCG-G-UACC-UGAAGCCCC
GG-------------UACA-------------ACGGGGUGAAGGGC--CGGUGAAA-GCCGGGAGUAACUCU
GACUCUCUUAAGGUAGCCAUAUGCCUUGCCGAA-UAAAAAUCGGCGUGCAUGAAUGGCUUAACGAGG--UGC
CCGCUGUCCCCUCC----CGC-GGGCCGGUGAACCC---GC---AUCCCGGUGUACAGACCGGGACGCCCCA
ACGGGGAGAGAAGA-CCCUGUGGAGC-UUCACU--GCAGCCU-GGUCUUGUGGUACAGCUGGGGAUGCAGAG
GGU-AGGCGGGAGCGAUGUGG----CCA---------CCC----------------------------UCCA
--------------------------------------GGGGGUGGCCGAA-G--C-GA-CGAUGGAACA-C
CGCC-CUUCUCCAGCUGUACCACUAACCG-CU----------------------------------------
------------------------------------------------------------------------
------------------------------------------------------------------------
------------------------------------------------------------------------
------------------------------------------------------------------------
------------------------------------------------------------CC----------
------------------------------------------------GGCG-----GGACA-GGGCCAGGU
GGGCAGUUUGGCUGGGGUGGCACGCCCCUGC-AAAGGUAUC-AGGGGCGCCCU-AAGGCGAGCUCAGGCGGG
ACAGAAACCCGCCGUA-GAGGG-CAAGGCCAA-AAGCUCGCCUGACU-GGAU--------------------
--------------CCCUAAACGCGUGGGAUCCAGGGGUGAAA-GCCGGGCCUAGCGAACACUC--------
---GA-G-----CCCUC-UCG--------------------GU-GAGGGCCGGGUACGACAA-AA--AAGUU
ACCCCAGGG-GUAACGGAGCUGUGGCGGGCGAGAGUCCCCAUCGGCCCCGCCGCUUGCUUCCCCGACGUCGG
```

```
CACUGCCCAUCCUGGGGGUGCA-GCAGCCUCCAAGGGUGGGGAUGUUCGCCCAUUAAAGGGGAACGUGAGCU
GGGUUCAGAACGUCGCGAGCCAGUUCGGUUUCUCCCUGUU--------------------G-GGG-------
-------------------------------CUGCA-AGGCCGCCUGAGGGGAAGG-GACCCUUAGUACGAA
AGGAA-CGGGGUGUCGGGGCCUC-CGGU-G-UACCGGUCGUCC-GAAG-GGGCGA-C-GCCGGGUAGCCGCG
CCCCA-GCCGAUAAGGGCUGAAAGCA-UC--UAAGCCCGAAGCGGCCCCCGA-A-AAU--AGGCGGCC-UC-
--GCCCC-----------------------------------------------------------------
------------------------------------------------------------------------
------------------------------------------------------------------------
------------------------------------------------------------------------
------------------------------------------------------------------------
------------------------------------------------------------------------
------------------------------------------------------------------------
------------------------------------------------------------------------
-------------------------------------------------UCGUGGG-GUU-AGG-------GC
GCGGGUAGAAGACCCGUU-CGA-UGGGGG-GGU-GGUGUAAG-GCA-G--------GAG-------------
CUUCGG-------CGGACUGCCUCAGCCU--A-CCCC-UCCCAA-A---GCCCGAGCG--------------
------------------------------------------------------------------------
------------------------------------------------------------------------
------------------------------------------------------------------------
-------------------------------------------------
>LSUa_272569_HALMA/1-2923
----------------------GU---UGGC----UA--CUAUGCCAGCU-GGUGGAUU-G-CUCGGC-UCA
G-G---CGCUGAUGAAGGACGUGC--CAAGCUGCGAUAAGCCAUGGGGAGCCGCACGGAGGCGA------AG
AAC-CAU-GGAUUUCCGAAUGAGAAU-C--------UCU-------CU------------------------
------------------------------------A----ACA-AUUG-C---------------------
-----------U-----------UCGC---------------------------------------------
-----------------------GC-A-AU-GAGGAACCCCGAGAACUGAAACAUCUCAGUAUCGG--GAGG
AACAGAAAACGCA-AUGUGAUGUCGUUA--GU-AA-CCGCGAGUGAACGCGAU-A-CA-GCCCAAACCGAAG
CCCU----------------------------------CA-CGGGCAAUGUGGUG--UC-------------
A-GG--GCU-ACC-U-----C------U------------------CAUCA--GCC-----------GA--C
C-------GUCUC--GACG------AAGUCU-CUUG------------------------------------
-------GAA-CA-GAGCG-------UGAUACAGGGUGACAACCCCGUA---------CUC---G----AGA
C------CAGUACGC-UG------------------UGCG-GUAGU--GCCAGAGUAGCGGGGGUUG---GA
UAUCCCUCGCGAA-UAACGCAGGCAUC--GACUGCGAAGGCUAAACA--CAA-C-CUG-AGACCGAUAGU-G
AAC-A-AGUAGUGUGAACGAACG-CUGCAA-AGUAC-CCUCAGAAGGGA-G-GCGAAAUAGAGCAUGAAAUC
A-GUUGGCGAUCG-AGC-GA-CAGG---GCAUACAA-GG---------------------------------
------------------------------------------------------------------------
------------------------------------------------------------------------
------------------------------------------------------------------------
------------------------------------------------------------------------
------------------------------------------------------------------------
------------------------------------------------------------UC-CCUUGACG-
---AA-----------UG-A-------------------C------CGA-----------------CGC---
------------------------------------------------------------------------
--------------------------------------------------------------G--C-GA---
```

```
------------------------------------------------------------------------
-------------------------------------------------G-CG------UC--------CAG
UAAGA-------------------------------------------------------------------
------------------------------------------------------------------------
------------------------------------------------------------------------
------------------------------------------------------------------------
------------------------------------------------C-----------------------
------------------------------------------------------------------------
------------------------------------------------------------------------
------------------------------------------------------------------------
------------------------------------------------------------------------
------------------------------------------------------------------------
-------------------UCACGGAA------------------GCCGA--U---GUUCUGUCGUACGUUU
UGAAAAACGAGCCAGGGAGUGUG-U-CUGCAUGGCAA-GUCUAACC--G-G--------AGUAU--------
C-CG--GGGA--GGCACAGGGAAAC-CGA-----------------CAU----------------------G
GCCG-CAGGGCU----U-------------------------------------------------------
------------------------------------------------------------------------
-------------------UGCC------------------------------------CGA-GGGCCG-CC
GUCU-----------------------------------UCAA--------------------------GG-
G-C-GG----------------------GGAGCCAUGUGGACAC-GACCCGAAUCC-GGACGAUCUACGCAU
GGACAAGAUGAAGCGUGC-CGAAAG-GCACGUGGAAGUCUGU-UAGAGUUGGUGUCCUACAAUACCC-UCUC
GUGAUCUAUGUGUAGGGGUGAAAGGCCCAUCGAGUCCGG-CAACAGCUGGUUCCAAUCGAAA-CAUGUCGAA
GCAUGACCUCCG-CCGA-------------------------------------------------------
----------------------GG-----UA-GUCUGUGAGGUAGAGCGACCGAUU-GGU-GUGU-CCGCCU
C-C-GA-GA-GGAGUC-GGCACACCUGUCAAACUCCAAACUUACAG--------------------------
------------------------------------------------------------------------
------------------A-CG-C--CGUUUGA--CGCGGGG-AUUCCGG-U-GCGC-G-GGG-UAAGCCUG
UG-U-ACCAGGAG-GGGAAC---AACCCAGAGAUA-G-GUUAAGGUCCCCAAGU--GU-GGA-UU-AA---G
-U-GUAAUCCUCUGAAGGUGGUCUCGAGCCCUAGACAGCCGGGAGGUGAGCUUAGAAGCAGCUA-CCCUCUA
AGAAAAGCGUAACAGCUUACCGGCC-----GAGGUUUGAGGCGCCCAAAAU-GAUC-GGGACUCA-AAUCCA
C----------CACCGAG-ACCUGUCCGUACCAC--U-----------------------------------
------------------------------------------------------------------------
-----C--AUA-------------------------------------------------------------
--------------------------------CUGGU-AAUC-GAGUAGAUUGGC-GCUCUAA----------
-----------U-U-GGAUG-GAAGUAG--G-GGUGAAAACU--C-CUAUGGACCG-A-UUA-GUGACGAAA
AUCC-UGGCCA-UAGUA-GCAGCG-AUAGUC-GGGUGA--G---------------AACCCC-GAC-GGCCU
A-AUGGAU-AAGGGUUCC-UCAGCACUG-CUGAUCAGCUGAGGGUUAGCCGG--UC-CUAAGUCAUACCGC-
AAC-UCGA---C-UAU--------------------------------------------------------
----------------------------------------GAC-GA-AAUGGGA-AACGGGUU-AAUA-UUC
CCGUG----------------------------------CCACU----AUGC--------------------
------------------------------------------------------------------------
--------------------------------------------------------AGUGA-A-AGUU-GAC
GCCCUGG-------------GGUCGA-UCACGCUGG-GC-------AUUC-------------G-CCCAGU-
CGAA-C-CGU-CC-----A-A-------------------CU-----CC----------------GUGG-A-
A-G-CC--------------------GUAAU-GGCAGGAAGC---------GGACGA----ACG-GC-GGU-
```

```
AUA--GG-----------------------------------------------------------------
----------------G-AA-A--C-G-UGA-U-U-CAACCUGGGGCCC-AUGAAA-A-GAC-GAGC-----
---------------------------------------------------------------AU-AGUGUC
CGUACCGAGAACCGAC------------------------------ACAGGUGU-CCAUGGCG-GCGAAAGC
CAAGGC-CUGU-CGGGAGCAACCAA-C-GUUAGGGAAUUCGGC-AAGUUAGUCCCGUACCUUCGG-AAGAAG
GGAUG-CCUG-C-UC--CGG----------------------------------------------------
------------------------------------------------------------------------
------------------------------------------------------------------------
------------------------------------------------------------------------
------------------------------------------------------------------------
------------------------------------------------------------------------
------------------------------------------------------------------------
------------------------------------------------------------------------
------------------------------------------------------------------------
------------------------------------------------------------------------
------------------------------------------------------------------------
------------------------------------------------------------------------
------------------------------------------------------------------------
------------------------------------------------------------------------
------------------------------------------------------------------AACGGA
GCA--GGUCGCAGUGACUCGGAAGCUCGGA-----CUGUCUAGUAACAACAUAGGUGACCGCAAAUCCG---
-CA-A-------------GGACUCGU-ACGGUCACUGA-AUCCUGCCCAGUGCAG-G-UAUC-UGAACACCU
CG-------------UACA-------------AGAGGACGAAGGAC--CUGUCAAC-GGCGGGGGUAACUAU
GACCCUCUUAAGGUAGCGUAGUACCUUGCCGCA-UCAGUAGCGGCUUGCAUGAAUGGAUUAACCAGA--GCU
UCACUGUCCCAACG----UUG-GGCCCGGUGAACUG--UAC---AUUCCAGUGCGGAGUCUGGAGACACCCA
GGGGGAAGCGAAGA-CCCUAUGGAGC-UUUACU--GCAGGCU-GUCGCUGAGACGUGGUCGCCGAUGUGCAG
CAU-AGGUAGGAGACACUAC-------ACAGGUACC-CGC----------------------------GCUA
--------------------------------------GCGGGCCACCGAG-U--C-AA-CAGUGAAAUA-C
UACC-CGUCGGUGACUGCGACUCUCACUC-C----------GG-G-A-------------------------
------------------------------------------------------------------------
------------------------------------------------------------------------
------------------------------------------------------------------------
------------------------------------------------------------------------
------------------------------------------------------------------------
-------------------------------------------------GGA-----GGACA-CCGAUAGCC
GGGCAGUUUGACUGGGGCGGUACGCGCUCGA-AAAGAUAUC-GAGCGCGCCCU-AUGGCUAUCUCAGCCGGG
ACAGAGACCCGGCGAA-GAGUG-CAAGAGCAA-AAGAUAGCUUGACA-GUGU--------------------
--------------UCUUCCAACGAGGAACGCUGACGCGAAA-GCGUGGUCUAGCGAACCAAU--------
---UA-G-----CCUGC-UUG--------------------AU-GCGGGCAAUUGAUGACAG-AA--AAGCU
ACCCUAGGG-AUAACAGAGUCGUCACUCGCAAGAGCACAUAUCGACCGAGUGGCUUGCUACCUCGAUGUCGG
UUCCCUCCAUCCUGCCCGUGCA-GAAGCGGGCAAGGGUGAGGUUGUUCGCCUAUUAAAGGAUGUCGUGAGCU
GGGUUUAGACCGUCGUGAGACAGGUCGGCUGCUAUCUACU--------------------G-GGU-------
-------------------------------GUGUA-AUGGUGUCUGACAAGAACG-ACCGUAUAGUACGAG
AGGAA-CUACGGUUGGUGGCCAC-UGGU-G-UACCGGUUGUUC-GAGA-GAGCACGU-GCCGGGUAGCCACG
CCACA-CGGGGUAAGAGCUGAACGCA-UC--UAAGCUCGAAACCCACUUGGA-A-A-A--GAGACACC-GC-
```

------------------------------------------------------------------------
------------------------------------------------------------------------
------------------------------------------------------------------------
------------------------------------------------------------------------
------------------------------------------------------------------------
------------------------------------------------------------------------
------------------------------------------------------------------------
------------------------------------------------------------------------
----------------------------------------------------------CG-AGG------UC
CCGCGUACAAGACGCGGU-CGA-UAGACU-CGG-GGUGUGCG-CGU-C--------GAG-----GU------
----AA--------CGAGACGUUAAGCCC--A-CGAG-CACUAACA---GACCAAAGC-CAUCAU-------
------------------------------------------------------------------------
------------------------------------------------------------------------
------------------------------------------------------------------------
-------------------------------------------------
>LSUa_64091_HALSA/1-2906
----------------------G----UGGC----UA--CUGUGCCACCU-GGUGGAUA-G-CUCGGC-UCG
G-A---UGCCGACGAAGGACGUGC--CAAGCUGCGAUAAGCCUGAGGGAGCCGCACGGAGGCUA------AG
AAC-UCA-GGAUCUCCUAAUGGGAAU-C--------CCU-------AU------------------------
------------------------------------A----ACA-AUUG-C---------------------
-----------C-----------UUGC---------------------------------------------
-----------------------GC-A-AU-GGGGAACGGCCGGAAUUGAAGCAUCUCAGUACGGC--CAGG
AAGAGAAAUCGA--AUGAGACGCCGUUA--GU-AA-UGGCGAAUGAACGCGGC-A-CA-GUCCAAACCGAAG
CCUU-------------------------------------CGGGCAAUGUGGUG--UU-------------
C-GG--ACU-GAC-U-----U------U------------------CAUCG--UUU-----------GA--C
C-------GUUCG--UGUG------AAGUCU-CCUG------------------------------------
-------AAA-CG-GAGCG-------CGAUACAGGGUGACAGCCCCGUA---------UCA---C----GGA
C------CAGUACGA-CG------------------UGCG-UCAGC--UCCAGAGUAGCGGGGGUUG---GA
AAUCCCUCGUGAA-UGUGGCAGGCAUC--GACUGCCAAGACUAAGUA--CUC-U-CCG-AGACCGAUAGU-G
AAC-A-AGUAGUGUGAACGAACG-CUGAAA-AGCAC-CCCACAAAGGGG-G-GUGAAAUAGGGCUUGAAAUC
A-GGUGGCGAUGG-AGC-GA-CGGG---GCAUAAAA-GG---------------------------------
------------------------------------------------------------------------
------------------------------------------------------------------------
------------------------------------------------------------------------
------------------------------------------------------------------------
------------------------------------------------------------------------
------------------------------------------------------------CC-UCUCUGGG-
---AA-----------CG-A-------------------C------UUG-----------------AGU---
------------------------------------------------------------------------
--------------------------------------------------------------G--C-AA---
------------------------------------------------------------------------
-------------------------------------------------A-CU------CA--------UGG
UAGGA-------------------------------------------------------------------
------------------------------------------------------------------------
------------------------------------------------------------------------

```
------------------------------------------------------------------------
------------------------------------------------C-----------------------
------------------------------------------------------------------------
------------------------------------------------------------------------
------------------------------------------------------------------------
------------------------------------------------------------------------
------------------------------------------------------------------------
-------------------CUGAGAGGA-----------------GCCGA--U---GUUCCGUCGUACGUUU
UGAAAAACGAGCCAGGGAGUGUG-C-CUGUUUGACGA-GUCUAACC--G-G--------AGUAU--------
C-CG--GGAA--GGCGUAGGGAAAC-CAA-----------------UAU----------------------G
GCCG-CAGCAU-------------------------------------------------------------
------------------------------------------------------------------------
---------------------UG-------------------------------------CGA-GGGCCA-CC
GUGU-----------------------------------UCAA--------------------------GC-
G-C-GG----------------------GGAGUCAAACGGGCAC-GACCCGAAACC-CGGUGAUCUACGCGU
GGGCAAGGUGAAGCAUGG-CGAAAG-CCAUGUGGAGGCCUGU-UAGGGUUGGUGUCUUUCAACACCC-UCCC
GUGACCUACGUGUAGGGGUGAAAGGCCCAUCGAACCGGG-CAACAGCUGGUUCCAACCGAAA-CAUGUCGAA
GCAUGACCUCUG-CCGA-------------------------------------------------------
----------------------GG-----UA-GUUCGUGGGGUAGAGCGACCGAUU-GGG-GAGU-UCAACU
C-C-GA-GA-GGAGUU-GUCUCCCCUGUCAAACUCCAAACCUACGG--------------------------
------------------------------------------------------------------------
------------------A-CG-C--CG-UCGA--CGCAGGG-AAUCCGG-U-GUGC-G-GGG-UAAGCCUG
UG-C-ACCGUGAG-GGAGAC---AACCCAGAGUUA-G-GUUAAGGUCCCAAAGU--GC-GAG-CU-AA---G
-U-GCGA----UUGAAGGUGGUCUCGAGCCCUAGACAGCCGGGAGGUGAGCUUAGAAGCAGCUA-CCCUCUA
AGAAAAGCGUAACAGCUUACCGGCC-----GAGGUUCGAGGCGCCCAAAAU-GAUC-GGGGCUCA-AGUUCG
C----------CACCGAG-ACCUAACGGCACGGG--U-----------------------------------
------------------------------------------------------------------------
-----A--AC--------------------------------------------------------------
--------------------------------ACCGU-GAUC-CAGUAGGUUGGC-AUUCUGU---------
-----------U-C-GGGUG-GAAGCUC--G-GGUGAGAACU--C-GAGUGGACCG-A-GUG-GAAAAGAAA
AUCC-UGGCCA-UAGUA-GCAGCG-UUAGUC-GGGUAA--G----------------AUCCC-GAU-GGCCG
A-AAGAGC-AAGGGUUCC-UCGGCAAUG-CUUAUCAGCCGAGGGUUAGCCGA--UC-CUAAGGCCCGUCGU-
AAU-UCGA---G-CGG--------------------------------------------------------
----------------------------------------GUC-AA-A-AGGGA-AACUGGUU-AAUA-UUC
CAGUG----------------------------------CCACC----GUAC--------------------
------------------------------------------------------------------------
--------------------------------------------------------AUUGA-A-AGUC-GAC
GCCUCGG-------------AGCAGC-UUGAGCCGG-GC-------AUUC-------------G-CCCGGU-
CGAA-C-CGU-CG------A-A-------------------GU-----UC----------------GUGG-A-
A-G-CC--------------------GUAAU-GGCAGGAAGC---------GAACGA----ACG-UC-GGA-
ACA--GG-----------------------------------------------------------------
----------------G-AA-A--C-U-CAA-G-U-CAAUCUGGGGCCC-GUGAAA-A-GGC-GAGU-----
---------------------------------------------------------------AC-GGUGUU
CGUACCGAGAUCCGAC------------------------------ACAGGUGC-UC-UGGCA-GAGGAAGC
CAAGGC-CUGU-CGGGAAUAACCGA-C-GUUAGGAAUUCGGC-AAGUUAGUCCCGUAAGUUCGC-GAUAAG
```

```
GGAUG-CCUG-C-CU--CGC----------------------------------------------------
------------------------------------------------------------------------
------------------------------------------------------------------------
------------------------------------------------------------------------
------------------------------------------------------------------------
------------------------------------------------------------------------
------------------------------------------------------------------------
------------------------------------------------------------------------
------------------------------------------------------------------------
------------------------------------------------------------------------
------------------------------------------------------------------------
------------------------------------------------------------------------
------------------------------------------------------------------------
------------------------------------------------------------------------
------------------------------------------------------------------AAUGAG
GCA--GGUCGCAGUGACUCGGAGGCUCCGA-----CUGUCUAAUAACAACAUAGGUGACCGCAAAUCCG---
-CA-A-------------GGACGCGU-ACGGUCACUGA-AUCCUGCCCAGUGCGG-G-UAUC-UGAACACCC
AG-------------UACA-------------AUGGGGCGAAGGAC--CCGUUAAC-GGCGGGGGUAACUAU
GACCCUCUUAAGGUAGCGUAGUACCUUGCCGCU-UCAGUAGCGGCUUGCAUGAAUGGAUCAACGAGA--GCC
UCACUGUCCCAACG----UUG-GGCCCGGUGAACUG--UAC---GUUCCAGUGCGGAGUCUGGAGACCCCCA
AGGGGAAGCGAAGA-CCCUAUAGAGC-UUUACU--GCAGGCU-GUCGCUGGGACACGGUCGCUGAUGUGCAG
AGU-AGGUAGGAGACGUUAC-------ACAGGUACG-UGC----------------------------GCUA
--------------------------------------GCACGCCACCGAG-U--C-AC-ACAUGAAACA-C
UACC-CGUCAGUGACUGUGACCCUCACUC-C----------GG-G-A-------------------------
------------------------------------------------------------------------
------------------------------------------------------------------------
------------------------------------------------------------------------
------------------------------------------------------------------------
------------------------------------------------------------------------
-------------------------------------------------GGA-----GGACA-CCGGUAGCC
GGGCAGUUUGACUGGGGCGGUACGCGCUUGA-AAAGAUAUC-GAGCGCGCCCU-AAGCCUAUCUCAGCCGAG
UCAGAGACUCGGCGAA-GAGUG-CAAGAGCAU-AAGAUAGGCUGACA-GUGU--------------------
--------------CCUACACAACGAGGGACGCUGACGCGAAA-GCGUGGUCUAGCGAACCAAU--------
---UA-G-----GCUGC-UUG--------------------AU-GCGGCCAAUUGCUGACAG-AA--AAGCU
ACCUUAGGG-AUAACAGAGUCGUCACUCGCAAGAGCACAUAUCGACCGAGUGGCUUGCUACCUCGAUGUCGG
UUCCCUCCAUCCUGCCCGUGCA-GAAGCGGGCAAGGGUGAGGUUGUUCGCCUAUUAAAGGAGGUCGUGAGCU
GGGUUUAGACCGUCGUGAGACAGGUCGGCUGCUAUCUAUU--------------------G-GGG-------
-------------------------------GUGUU-UUGGUGCUUGACAGGAACG-UUCGUAUAGUACGAG
AGGAA-CUACGAACGGGUGCCAC-UGGU-G-UAUCGGUUGUCC-GAGA-GGGCAUGU-GCCGAGCAGCUACG
CACCA-CGGGGUAAGAGCUGAAUGCA-UC--UAAGCUCGAAACCCACCUGGA-A-A-A--GAAGCACC-AC-
------------------------------------------------------------------------
------------------------------------------------------------------------
------------------------------------------------------------------------
------------------------------------------------------------------------
------------------------------------------------------------------------
```

```
------------------------------------------------------------------------
------------------------------------------------------------------------
------------------------------------------------------------------------
----------------------------------------------------------UG-AGA------CC
GCUCGUAGAAGACGAGUU-CGA-UAGACU-UGG-GGUGUACG-CGU-C--------GAG-----GC------
----AA--------CGAGACGUUUAGCCC--G-CGAG-UACUAACA---GGUCAAUGC-CACAC--------
------------------------------------------------------------------------
------------------------------------------------------------------------
------------------------------------------------------------------------
-------------------------------------------------
>LSUa_348780_NATPD/1-2906
----------------------G----UGGC----UA--CUGUGCCAUCU-GGUGAAUA-G-CUCGGC-UCA
G-G---CGCCGACGAAGGACGUGU--CAAGCUGCGAUAAGCCUCAGGGAGCCGCACGAAGGCAA------AG
AAC-UGA-GGAUCUCCUAAUGGGAAU-C--------CCC-------AC------------------------
--C---------------------------------G----CAA-UUGC-----------------------
-----------U-----------UCGC---------------------------------------------
------------------------G-C-AAUGGGGAACGCCCCGGAUUGAAACAUCUUAGUAGGGG--CAGG
AAGAGAAAGCGA--AUGCGAUGUCGUUA--GU-AA-CCGCGAGUGAACGCGAC-A-CA-GUCCAAACCGAAG
CCUU-------------------------------------CGGGCAAUGUGGUG--UU-------------
C-GG--ACU-GAC-U-----C------C------------------CAUCA--GCA-----------GA--A
C-------GUUUC--CGAG------AAGUCU-CCUG------------------------------------
-------GAA-CG-GAGCG-------CGAUACAGGGUGACAGCCCCGUA---------UCG---G----AAG
C------AAGUAUGC-UG------------------UGCG-UCAGC--UCCAGAGUAGCGGGGAUUG---GA
AAUUCCUCGUGAA-UUCGGCAGACAUC--GUCUGCCAAGACUAAAUA--CGU-C-CUG-AGACCGAUAGC-G
AAC-A-AGUAGCGUGAGCGAACG-CUGAAA-AGCAC-CCCGAGAAGGGA-G-GUGCAAUAGGGCUUGAAAUC
A-GAUGGCGAUCG-AGC-GA-CGGG---GCACAAAA-GG----------------------------------
------------------------------------------------------------------------
------------------------------------------------------------------------
------------------------------------------------------------------------
------------------------------------------------------------------------
------------------------------------------------------------------------
------------------------------------------------------------UC-ACUUGGAA-
---AA-----------CG-A-------------------C------CGA-----------------GGC---
------------------------------------------------------------------------
--------------------------------------------------------------G--C-GA---
------------------------------------------------------------------------
-------------------------------------------------G-CC------UC--------CAG
UAGGA-------------------------------------------------------------------
------------------------------------------------------------------------
------------------------------------------------------------------------
------------------------------------------------------------------------
------------------------------------------------C-----------------------
------------------------------------------------------------------------
------------------------------------------------------------------------
------------------------------------------------------------------------
```

```
------------------------------------------------------------------------
------------------------------------------------------------------------
-------------------CCAAGUGAA-----------------GCCGG--U---GUUCCGUCGUACGUUU
UGAAAAACGAACCAGGGAGUGUA-U-UUGUUUGGCGA-GUCUAACC--C-G--------UUCAU--------
C-GG--GGAA--GGCGUAGGGAAAC-CGA-----------------CAU----------------------G
GCCG-CAGCAU-------------------------------------------------------------
------------------------------------------------------------------------
---------------------UG------------------------------------CCA-GGGCCG-CC
GUGU-----------------------------------UCAA--------------------------GC-
G-C-GG----------------------GGAGUCAAACGGAUAC-GACCCGAAACC-GGGUGAUCUAUGCGU
GGACAGGACGAAGCGUGG-CGAAAG-CUACGUGGAGGUCCGU-UAGAGUUGGUGUCCUACAAUACCC-UCUC
GUGAUCUACGUAUAGGGGUGAAAGGCCCAUCGAACCCGG-AAACAGCUGGUUCCGACCGAAA-CAUGUCGAA
GCAUGACCUCUG-CCGA-------------------------------------------------------
----------------------GG-----UA-GUCCGUGGGGUAGAGAGACUGAUU-GGA-GGCC-CCGCCC
C-C-GA-GA-GGGGUC-GGACCUCCUGUCAAACUCCGAACUCACGG--------------------------
------------------------------------------------------------------------
------------------A-CG-C--CG-UGGA--CGCAGGG-AGUCCGC-U-GUGC-G-GGG-UAAGCCUG
UG-C-AGCAUGAG-GGAGAC---AACCCAGAGCUG-G-GUUAAGGUCCCCAAGU--GU-GGA-CU-AA---G
-U-GCGA----UUGAAGGUAGUCCCGAGCCCUAGACAGCCGGGAGGUGAGCUUAGAAGCAGCUA-CCCUCUA
AGAAAAGCGUAACAGCUUACCGGCC-----GAGGUUCGGGGCGCCCAAAAU-GAUC-GGGGCUCA-AGUCCA
C----------CACCGAG-ACCUAGCGGCACGCC--U-----------------------------------
------------------------------------------------------------------------
-----U--AC--------------------------------------------------------------
--------------------------------AGCGU-GGUU-CCGUAGGUCGGC-ACUCCGG----------
-----------U-U-GGGUG-GAAGCAC--G-GACGAGAGUU--C-GUGUGGACCG-A-CUG-GUGAAGAUA
AUCC-UGGUUG-CAGUA-GCAGCG-AUAGUC-GGGUGU--G---------------AAUCCU-GAC-GGCCU
A-AUGAGC-AAGGGUUCC-UCGGCAAUG-UUUGUCAGCCGAGGGUUAGCCGG--UC-CUAAGACUCGUCGU-
AAU-UCGA---G-CGA--------------------------------------------------------
----------------------------------------GUC-GA-AGUGGGA-AACUGGUU-AAUA-UUC
CAGUG----------------------------------CCACU----AUGC--------------------
------------------------------------------------------------------------
---------A-----------------------------------------------UUGA-A-CGCC-GAC
GCCUUGG-------------GGUCGA-CCGAACCGG-GC--------CUUC-------------G-CCCGGU-
CGAA-C-UGU-CC-----A-A-------------------AU-----CC----------------GUGG-A-
A-G-CC--------------------GUAAU-GGCACGAAGC---------GGACGA----ACG-GC-AGG-
AUA--GC-----------------------------------------------------------------
----------------G-CA-A--G-U-CGG-U-U-CAACCUGGGGCCC-GUGAAA-A-GGC-AAGC-----
---------------------------------------------------------------AU-AGUGUC
CGUACCGAGAUCCGAC------------------------------ACAGGUGC-UCAUGCCG-GCGAAAGG
CAAGGC-CUGU-CGGGAUCAACCGA-C-GUUAGGGAAUUCGGC-AAUCUGGCCCCGUAAGUUCGC-GAUAAG
GGGUG-CCUG-C-CU--CGG----------------------------------------------------
------------------------------------------------------------------------
------------------------------------------------------------------------
------------------------------------------------------------------------
------------------------------------------------------------------------
```

```
------------------------------------------------------------------------
------------------------------------------------------------------------
------------------------------------------------------------------------
------------------------------------------------------------------------
------------------------------------------------------------------------
------------------------------------------------------------------------
------------------------------------------------------------------------
------------------------------------------------------------------------
------------------------------------------------------------------------
------------------------------------------------------------------AAAGAG
GCA--GGUCGCAGUGACUCGGACGCUCCGA-----CUGUCUAGUAACAACAUAGGUGACCGCAAAUCCG---
-CA-A-------------GGACUCGU-ACGGUCACUGA-AUCCUGCCCAGUGCGG-G-UAUC-UGAACACCU
CG-------------UACA-------------AGAGGACGAAGGAC--CCGUCAAC-GGCGGGGGUAACUAU
GACCCUCUUAAGGUAGCGUAGUACCUUGCCGCU-UCAGUAGCGGCUUGCAUGAAUGGAUUAACGAGA--GCG
UCGCUGUCCAACG-----UUG-GGCCCGGUGAACCA--UAC---GUUCCAGUGCGGAGUCUGGAGACACCCA
GGGGGAAGCGAAGA-CCCUAUGGAGC-UUUACU--GCAGGCU-GUCGCUGGGACAUGGUCGCUGAUGUGCAG
GAU-AGGUAGGAGACGUUAC-------ACAGGUGCG-UGC----------------------------GCCA
--------------------------------------GCACGUCGCCGAG-U--C-AA-CACUGAAAUA-C
UACC-CGUCAGUGACUGUGACCCUCACUC-C----------GG-G-A-------------------------
------------------------------------------------------------------------
------------------------------------------------------------------------
------------------------------------------------------------------------
------------------------------------------------------------------------
------------------------------------------------------------------------
-------------------------------------------------GGA-----GGACA-CCGAUAGCC
GGGCAGUUUGACUGGGGCGGUACGCGCUCGA-AACAAUAUC-GAGCGCGCCCU-AAGUCCAUCUCAGUCGGG
UCGGAGACCCGACGAA-GAGUG-CAAGAGCAA-AAGAUGGAUUGACA-GUGU--------------------
--------------UCUUCCAACGAGGAACGCUGACGCGAAA-GCGUGGUCUAGCGAACCGAU---------
---GA-G-----CCUGC-CCG--------------------AU-GCGGGCCAUUGAUGACAG-AA--AAGCU
ACCCUAGGG-AUAACAGAGUCGUCACUCGCAAGAGCACAUAUCGACCGAGUGGCUUGCUACCUCGAUGUCGG
UUCCCUCCAUCCUGCCCGUGCA-GAAGCGGGCAAGGGUGAGGUUGUUCGCCUAUUAAAGGAGGUCGUGAGCU
GGGUUUAGACCGUCGUGAGACAGGUCGGCUGCUAUCUACU--------------------G-GGU-------
-------------------------------GUGU--CUGACGCUUGACAGGAACA-ACCGUAUAGUACGAG
AGGAA-CUACGGUUGGGUGCCAC-UGGU-U-UACCGGUUGUCC-GAGA-GGGCAC-U-GCCGGGCAGCUACG
CACCA-CAGGGUAACGGCUGAAUGCA-UC--UAAGCCGGAAACCUACCUGGA-A-A-A--GAAGCGUC-GC-
------------------------------------------------------------------------
------------------------------------------------------------------------
------------------------------------------------------------------------
------------------------------------------------------------------------
------------------------------------------------------------------------
------------------------------------------------------------------------
------------------------------------------------------------------------
------------------------------------------------------------------------
----------------------------------------------------------CG-AGG------UC
UCUCGUAGAAGACGAGUU-CGA-UAGACU-UGG-GGUGUACG-CGU-C--------GAG-----GC------
```

```
----AA--------CGAGACGUUCAGCCC--G-CGAG-UACUAACA---GACCGAAGC-CACCA--------
------------------------------------------------------------------------
------------------------------------------------------------------------
------------------------------------------------------------------------
-------------------------------------------------
>LSUa_309800_HALVD/1-2910
----------------------CAAC-UGGC----UA--CUGUGCCAGCU-GGUGGAUA-G-CUCGGC-UCG
A-G---AGCCGAUGACGGACGUGC--CAAGCUGCGAUAAGCCUCAGGGACCCGCAUGGAGGGAA------AG
AAC-UGA-GGAUCUCCGAAUGGGAAU-C--------CCC-------AC------------------------
------------------------------------G----C-A-AUUG-C---------------------
-----------C-----------UCGC---------------------------------------------
-----------------------GC-A-AU-GGGGAACGUUCCGAAUUGAAACAUCUUAGUAGGAA--CAGG
AAAAGAAAGCAAU-CCGCGAUGUCGUUA--GU-AA-CGGCGAGUAAACGCGAC-A-CA-GUCCAAACCGAAG
CCUU-------------------------------------CGGGCAAUGUGGUG--UA-------------
C-GG--ACU-ACC-C-----A------U------------------AACGG--ACC-----------UG--G
G-------UCUUA--UGUG------AAGUCU-CCUG------------------------------------
-------GAA-CG-GAGCG-------UGAUACAGGGUGAAAACCCGUA----------ACA--------UAA
G------AACCAGAU-CC------------------CGGG-GUUGU--UCCAGAAUAGCGGGGGUUG---GA
UAUCCCUCGUGAA-UUUCGCAGGCAUC--AACUGCGAAGACUAAACA--CUC-C-UCG-AGACCGAUAGC-G
AAC-A-AGUAGCGUGAGCGAACG-CUGAAA-AGCAC-CCCGAACAGGGC-G-GUGCAAUAGGGCUUGAAAUC
A-GUUGGCGAUAG-AGC-GA-CGUG---GCAUACAA-GG---------------------------------
------------------------------------------------------------------------
------------------------------------------------------------------------
------------------------------------------------------------------------
------------------------------------------------------------------------
------------------------------------------------------------------------
------------------------------------------------------------UC-ACUCGACG-
---AA-----------AG-A-------------------A------CUG-----------------GGA---
------------------------------------------------------------------------
--------------------------------------------------------------G--C-GA---
------------------------------------------------------------------------
-------------------------------------------------U-CC------CG--------UAG
UAUGA-------------------------------------------------------------------
------------------------------------------------------------------------
------------------------------------------------------------------------
------------------------------------------------------------------------
------------------------------------------------C-----------------------
------------------------------------------------------------------------
------------------------------------------------------------------------
------------------------------------------------------------------------
------------------------------------------------------------------------
------------------------------------------------------------------------
-------------------UCGAGUGAA-----------------GCCGA--U---GUUACGUCGUACGUUU
UGAAAAACGAACCAUGGAGUGCA-U-CUGACUGGCGA-GUCUAAGG--G-G--------UGAAC--------
C-CC--CGGA--GGCAUAGGGAAAC-CGA-----------------UAC----------------------G
```

```
GCCG-CAGUCUU------------------------------------------------------------
------------------------------------------------------------------------
--------------------UGA------------------------------------CCA-GGGCCA-CC
GUCU-----------------------------------UCAA--------------------------GG-
G-C-GG----------------------GGAGCCAACCGGAUGC-GACCCGAAACC-AAGUGAUCUACGCCG
GAGCAGGGCGAAGCGUGG-CGAAAG-CUGCGUGGAGGCCCGU-UAGAGUUGGUGUCCUACAAUACCC-UCUC
GUGACUCUGGUGUAGGGGUGAAAGGCCCAUCGAACUUGG-CAACAGCUGGUUCCAACCGAAA-CAUGUCGAA
GCAUGACCUCUG-AUGA-------------------------------------------------------
----------------------GA-----UA-GUUCGUGAGGUAGAGCGACCGAUU-GGG-AAUC-CGGACU
C-C-GA-GA-GGAGUU-CGCUUCCCUGUCAAACUCCGAACUUACGA--------------------------
------------------------------------------------------------------------
------------------A-CG-U--CG-UAGA--CUCAGGG-AGUCCGG-U-GUGC-G-GGG-UAAGCUUG
UG-C-ACCGUGAG-GGAGAC---AACCCAGAGCCA-G-GUUAAGGACCCAAAGU--GU-GGA-UU-AA---G
-U-GUGA----UUGAAGAUUGUCUCGAGCCCUAAACAGCCGGGAGGUGAGCUUAGAAGCAGCUA-CCCUCUA
AGAAUAGCGUAACAGCUUACCGGCC-----GAGGUUCGAGGCGUCGAAAAU-GAUC-GGGACUCA-AAUCCA
C----------CUCCGAG-ACCUGGCGGCGUGUC--U-----------------------------------
------------------------------------------------------------------------
-----U--AU--------------------------------------------------------------
--------------------------------GACAC-GAUU-CCGUAGGUUGGC-AUACUGC---------
-----------A-U-GGGUG-GAAGCAC--G-GGUGAGAACU--C-GUGUGGACCG-U-GCA-GCAACGAAA
AUCA-UGGUCU-CAGUA-GCAGCG-AGAGUC-GGGUGU--G---------------AACCCC-GAC-GGCCU
U-ACGCGC-AAGGGUUCC-UCGGCACUG-UCAAUCAGCCGAGGGUUAGUCGA--UC-CUAAGUCUAACCGU-
AAU-UCGA---A-UUA--------------------------------------------------------
----------------------------------------GAU-GA-A-AGGGA-AACUGGUU-AAUA-UUC
CAGUA----------------------------------CCUUC----UUGC--------------------
------------------------------------------------------------------------
--------------------------------------------------------ACCAA-A-CGUC-GAC
GCUUAG--------------GAAACA-CCGAGCGGG-GC-------CUUC-------------G-CCCCGU-
CGAA-U-CAC-CG-----A-A-------------------AU-----UC----------------GUGG-A-
A-G-CC--------------------GUAAC-GGCACGAAGC---------GAACGA----ACG-GU-GAG-
AUA--GG-----------------------------------------------------------------
----------------G-AA-A--C-U-CGG-C-G-GUACCUAGAGCCC-GUGAAA-A-GAC-AAGC-----
---------------------------------------------------------------AA-GAAGCU
CGUACCGAGAUCCGAC------------------------------ACAGGUGC-GC-UGGCA-GAGCAAGC
CAAGGC-CUGU-CGGGAUCAACCGA-C-GUUAGGGAAUUCGGC-AAGUUAGUCCCGUAAGUUCGC-GAUAAG
GGAUG-CCUG-C-UC--UUG----------------------------------------------------
------------------------------------------------------------------------
------------------------------------------------------------------------
------------------------------------------------------------------------
------------------------------------------------------------------------
------------------------------------------------------------------------
------------------------------------------------------------------------
------------------------------------------------------------------------
------------------------------------------------------------------------
------------------------------------------------------------------------
```

```
------------------------------------------------------------------------
------------------------------------------------------------------------
------------------------------------------------------------------------
------------------------------------------------------------------------
-------------------------------------------------------------------ACCGA
GCA--GGUCGCAGUGACUCGGACGCUCCGA-----CUGUCUAGUAACAACAUAGGUGACCGCAAAUCCG---
-CA-A-------------GGACUCGU-ACGGUCACUGA-AUCCUGCCCAGUGCGG-G-UAUC-UGAACACCC
AU-------------UACA-------------AUGGGACGAAGGAC--CCGUCAAC-GGCGGGGGUAACUAU
GACCCUCUUAAGGUAGCGUAGUACCUUGCCGCU-UCAGUAGCGGCUUGCAUGAAUGGAUCAACGAGA--GCG
UCACUGUCCCAACG----UUG-GGCCCGGUGAACAG--UAC---GUUCCAGUGCGGAGUCUGGAGACCCCCA
AGGGGAAGCGAAGA-CCCUAUAGAGC-UUUACU--GCAGGCU-GUCGCUGAGACGUGGUCGCUAAUGUGCAG
CAU-AGGUAGGAGCCAUUAC-------ACAGGUACC-CGC----------------------------GCCA
--------------------------------------GCGGGCCACCGAG-G--C-AG-CAUUGAAAUA-C
UACC-CGUUAGUGACUGCGACUCUCACUC-C----------UG-G-C-------------------------
------------------------------------------------------------------------
------------------------------------------------------------------------
------------------------------------------------------------------------
------------------------------------------------------------------------
------------------------------------------------------------------------
-------------------------------------------------GGA-----GGACA-CCGGUAGCC
GGGCAGUUUGACUGGGGGCGGUACGCGCUCGA-AAAGAUAUC-GAGCGCGCCCA-AAGAUUCCUCACGCGAG
UCGGGAACUCGUGGAA-GAGCG-CAAGAGCAA-ACGGAAGUCUGACA-GUGA--------------------
--------------CAUUCCCAACGAGUGUCGCUGACGCGAAA-GCGUGGUCUAGCGAACCUAC--------
---GA-G-----GUUCA-UUC--------------------AU-GGGACCCGUAGAUGACAG-AA--AAGCU
ACCUUAGGG-AUAACAGAGUCGUCACCGGCAAGAGCACAUAUCGACCCGGUGGCUUGCUACCUCGAUGUCGG
UUCCCUCCAUCCUGCCUGUGCA-GAAGCAGGCAAGGGUGAGGUUGUUCGCCUAUUAAAGGAGGUCGUGAGCU
GGGUUUAGACCGUCGUGAGACAGGUCGGCUGCUAUCUAUU--------------------G-GGG-------
-------------------------------GUGUU-AAGGAACUUGACGUGAACG-UUCGUAUAGUACGAG
AGGAA-CUACGAAUGGGUGCCAC-UGGU-G-UACCGGUUGUCC-GAGA-GGGCAGAU-GCCGGGCAGCUACG
CACCA-CGGGGUAAGAGCUGAACGCA-UC--UAAGCUCGAAACCCACACGGA-A-A-A--GAAGUUCC-AC-
------------------------------------------------------------------------
------------------------------------------------------------------------
------------------------------------------------------------------------
------------------------------------------------------------------------
------------------------------------------------------------------------
------------------------------------------------------------------------
------------------------------------------------------------------------
------------------------------------------------------------------------
----------------------------------------------------------UG-AGG------AC
ACUCGUAGAAGACGAGUU-CGA-UAGACU-CGG-GGUGUACG-CAA-C--------GAG-----GU------
----AA--------CGAGUUGUUAAGCCC--G-CGAG-CACUAACA---GUCCAA-GC-CACAUUC------
------------------------------------------------------------------------
------------------------------------------------------------------------
------------------------------------------------------------------------
-------------------------------------------------
```

```
>LSUa_416348_HALLT/1-2903
------------------------------------A--CUGUGCCAGCU-GGUGAAUG-G-CUCGGC-UCG
A-G---AGCCGAUGAAGGACGUGC--CAAGCUGCGAAAAGCUCAUGGGACCCGCAUGGAGGGGA------AG
AAC-AUG-AGAUAUCCGAAUCGGAAU-C--------CGU-------UU------------------------
------------------------------------A----C-A-AUUG-C---------------------
-----------C-----------ACGC---------------------------------------------
-----------------------GC-A-AU-AGGGAACUCCCUGAAUUGAAACAUCUCAGUAAGGG--AAGG
AAAAGAACGCAA--UAGCGAUGUCGUCA--GU-AA-CCGCGAGUGAACGCGAC-A-CA-GCCCAAACCGAAG
GUCU----------------------------------UC-GGACCAAUGUGGUG--UU-------------
C-GG--GUU-GAC-A-----A------U------------------CAAUU--CUC-----------GA--C
A-------UUUCA--UCUG------AAGUCU-CCUG------------------------------------
-------GAA-UG-GAGCG-------UGAAACAGGGUGACAACCCCGUA---------AGA---U----GAG
A------CAGUACGA-AG-------------------GCG-UCAAU--ACCAGAGUAGCAGGGGUCG---GA
UAUCCUCUGUGAA-CGACUCAGGCAUC--CCCUGAGAAGGCUAAACA--CUC-C-UCG-AGACCGAUAGC-G
AAC-A-AGUAGCGUGAGCGAACG-CUGAAA-AGCAC-CCCGAGAAGGGC-G-GUGCAAUAGGGCCUGAAAUC
A-GUUGGCGAUCG-AGC-GA-CAGG---GCGUACAA-GG---------------------------------
------------------------------------------------------------------------
------------------------------------------------------------------------
------------------------------------------------------------------------
------------------------------------------------------------------------
------------------------------------------------------------------------
------------------------------------------------------------CG-UCUCUUAA-
---AA-----------CG-A-------------------G------CGA-----------------GAA---
------------------------------------------------------------------------
--------------------------------------------------------------G--C-GA---
------------------------------------------------------------------------
-------------------------------------------------U-UC------UC--------UAG
UAGGA-------------------------------------------------------------------
------------------------------------------------------------------------
------------------------------------------------------------------------
------------------------------------------------------------------------
------------------------------------------------A-----------------------
------------------------------------------------------------------------
------------------------------------------------------------------------
------------------------------------------------------------------------
------------------------------------------------------------------------
------------------------------------------------------------------------
-------------------GAGAGACGA-----------------GCCGG--C---GUCGUGUCGUGCGUUU
UGAAAAACGACCCAGGGAGUGCA-C-UUGAUUGGCGA-GUCUAACC--C-G--------UGAAC--------
C-GG--GGAA--GGCGCAGGGAAAC-CGA-----------------UAC----------------------G
GCCG-CAGCACA----U-------------------------------------------------------
------------------------------------------------------------------------
----------------U--CGUG------------------------------------UGA-GGGCCA-CC
GUGU-----------------------------------UCAA--------------------------GC-
G-C-GG----------------------GGAGUCAAUUGGGUGC-GACCCGAAACC-AGGCGAUCUAUACAC
```

```
AGGCAGGGUGAAGCGUGG-CGAAAG-CCACGUGGAGGCCCGU-UAGAGUUGGUAUUCUACAAUAUCC-UCUC
GUGAUCUGUGCAUAGGGGUGAAAGGCCCAUCGAGCCUGG-CAACAGCUGGUUCCAACCGAAA-CAUGUCGAA
GCAUGACCUCAU-CCGA-------------------------------------------------------
----------------------GA-----UA-GCCCGCGGGGUAGAGCUACCGAUU-GGA-UGAC-CCGCCU
C-C-GA-GA-GGAGUC-GGCCAUCCUGUCGAACUCCAAACCCGCAG--------------------------
------------------------------------------------------------------------
------------------G-CG-U--UG-UAGA--CGAUGGG-AGUCCGG-U-GCGC-G-GGG-UAAGCCUG
UG-U-ACCGUGAG-GGGAAC---AACCCAGAGCCG-G-GUUAAGGUCCCAAAGU--GU-AGA-UU-AA---G
-U-GCGA-----CGAAGGUGGUCUCAAGCCCUAAACAGCCGGGAGGUGAGCUUAGAAGCAGCUA-CCCUCUA
AGAAAAGCGUAACAGCUUACCGGCC-----GAGGUUUGAGGCGCCCAAAAU-GAUC-GGGGCUCA-AAUCUA
C----------CACCGAG-ACCUGGCCGUGCCCU--U-----------------------------------
------------------------------------------------------------------------
-----G--ACA-------------------------------------------------------------
--------------------------------GGGGC-AACC-GCGUAGGUUGGC-GUUCUGU---------
-----------U-C-GGAUG-GAAGCAC--G-GGUGAGAACU--C-GUGUGGACCG-U-UCA-GAAACGACA
AUCC-UGGUCA-CAGUA-GCAGCG-AUAGUC-GGGUGA--G---------------AACCUC-GAC-GGCCU
G-AAGAGC-AAGGGUUCC-UCGGCACUG-CCACUCAGCCGAGGGUCAGCCGG--UC-CUAAGAUGCACCGU-
AAC-UCGA---C-UGC--------------------------------------------------------
----------------------------------------AUC--A-A-CGGGA-AACUGGUU-AAGA-UUC
CAGUG----------------------------------CCAUC----GUGC--------------------
------------------------------------------------------------------------
--------------------------------------------------------AGUAA-A-CGUC-GAC
GCCGUGU-------------GGAACG-CUGAGCCGG-GC--------UUUC-------------G-CCCGGU-
CGAA-U-CGU-GG-----A-A-------------------AC-----UC----------------GUGG-A-
A-G-CC--------------------GUCAC-GGCACGAAGC---------GAGCGA----AAC-GC-GAG-
AUA--GC-----------------------------------------------------------------
----------------G-CA-A--G-U-CAG-C-C-GUACAUAGGGCCC-GUGAAA-A-GAC-AAGC-----
---------------------------------------------------------------AC-GAUGUC
CGUACCAAGAUCCGAC------------------------------ACAGGUGC-UC-UGGCA-GCGCAAGC
CAAGGC-CUGU-CGGGAG-AACCGA-C-GUUAGGGAAUUCGGC-AAGUUAGUCCCGUACCUUCGG-AAGAAG
GGAUG-CCUG-C-UC--UCG----------------------------------------------------
------------------------------------------------------------------------
------------------------------------------------------------------------
------------------------------------------------------------------------
------------------------------------------------------------------------
------------------------------------------------------------------------
------------------------------------------------------------------------
------------------------------------------------------------------------
------------------------------------------------------------------------
------------------------------------------------------------------------
------------------------------------------------------------------------
------------------------------------------------------------------------
------------------------------------------------------------------------
------------------------------------------------------------------------
-------------------------------------------------------------------ACGGA
```

```
GCA--GGUCGCAGUGACUCGGGCGCUCCGA-----CUGUCUAAUAACAACACAGGUGACCGCAAAUCCG---
-CA-A-------------GGACUCGU-ACGGUCACUGA-AUCCUGCCAGUGCGG-G-UAUC-UGAACACCU
CG-------------UACA-------------AGAGGACGAAGGAC--CCGUCAAC-GGCGGGGUAACUAU
GACCCUCUUAAGGUAGCGUAGUACCUUGCCGCU-UCAGUAGCGGCUUGCAUGAAUGGAUAAACGAGA--GCG
CCACUGUCCCAACG----UUG-GACCCGGUGAACUG--UAC---GUUCCAGUGCGGAGUCUGGAGACCCCCA
AGGGGAAGCGAAGA-CCCUAUAGAGC-UUUACU--GCAGGCU-GUCGCUGAGACGUGGUCGCCGAUGUGCAG
CAU-AGGUAGGAGUCAUUAC-------ACAGGUACC-CGC----------------------------GCCA
--------------------------------------GCGGGCCACCGAG-A--C-AA-CAUUGAAAUA-C
UACC-CGUCGGUGACUGCGACUCUCACUC-C----------GG-G-A-------------------------
------------------------------------------------------------------------
------------------------------------------------------------------------
------------------------------------------------------------------------
------------------------------------------------------------------------
------------------------------------------------------------------------
-------------------------------------------------GGA-----GUACA-CCGGUAGCC
GGGCAGUUUGACUGGGGCGGUACGCGCUCGA-AAAGAUAUC-GAGCGCGCCCG-AAGAUCAUCUCAGCCGGG
UCGGAAAUCCGGCGAA-GAGCG-CAAGAGCAC-AAGAUGGUCUGACA-GUGU--------------------
--------------CAUUCCCAACGAGUGACGCUGACGCGAAA-GCGUGGUCUAGCGAACCUAC--------
---GA-G-----GCUCC-GGA--------------------AU-GGGGCCCGUAGAUGACAG-AA--AAGCU
ACCUUAGGG-AUAACAGAGUCGUCACCCGCAAGAGCACAUAUCGACCGGGUGGCUUGCUACCUCGAUGUCGG
UUCCCUCCAUCCUGCCCGUGCA-GAAGCGGGCAAGGGUGAGGUUGUUCGCCUAUUAAAGGAGGUCGUGAGCU
GGGUUUAGACCGUCGUGAGACAGGUCGGCUGCUAUCUAUU--------------------G-GGG-------
-------------------------------GUGUC-AAGGUACUUGACGUGAACG-AACGUAUAGUACGAG
AGGAA-CUACGUUUGGUCGCCAC-UGGU-G-UACCGGUUGUGU-GAGA-ACGCAA-U-GCCGGGUAGCCACG
CGACA-CGGGGUAAGAGCUGAACGCA-UC--UAAGCUCGAAACCCACAUGGA-A-A-A--GAAGUACC-GA-
------------------------------------------------------------------------
------------------------------------------------------------------------
------------------------------------------------------------------------
------------------------------------------------------------------------
------------------------------------------------------------------------
------------------------------------------------------------------------
------------------------------------------------------------------------
------------------------------------------------------------------------
----------------------------------------------------------UG-AGG------UC
ACUCGUAGAAGACGAGUU-CGA-UAGACU-CGG-GAUGUACG-CAC-C--------GAG-----GC------
----AA--------CGAGGUGUUGAGUCC--A-CGAG-CACUAACA---GACCGAAGC-CACAGUC-------
------------------------------------------------------------------------
------------------------------------------------------------------------
------------------------------------------------------------------------
-------------------------------------------------
>LSUa_362976_HALWD/1-2918
----------------------UA-C-UGGC----UA--CUGUACUGGCU-GGUGAAUG-G-CUCGGC-UCG
A-G---AGCCGAUGACGGACGUGC--CAAGCUGCGAAAAGCCUGAGGGAGCCGCACGGAGGCUA------AG
AAC-UCA-GGAUCUCCGAAUGGGAAU-C--------CCC-------AU------------------------
------------------------------------U----GCG-AUUA-C-G------AG------------
```

```
-----------G-----------UAU---------------------------------------------C
U-C--------------------GU-A-AU-CGGGAACGCCCCGAAUUGAAACAUCUUAGUAGGGG--CAGG
AACAGAAAGCAA--CCGCGAUAUCGUUA--GU-AA-CCGCGAGUGAACACGAC-A-CA-GUCCAAACCGAAU
CCGC-------------------------------------AAGGAAAUGUGGUG--UUU------------
C-GG--ACC-AAC-A-----A------U------------------CACGG--ACC-----------GC--A
C-------GAUAC--CGAG------AAAUCU-CCUG------------------------------------
-------GAA-CG-GAGUA-------CGACACAGGGUGACAGUCCGUA----------UC----G----UCA
U------CACGCGAU-CC------------------CGCG-UUGAC--UCCAGAGUCGCAAGGGUUG---GC
UAUCCCUUGUUAA-UUCCGCAGACAUC--AACUGCGAAGACUACACA--CUC-C-UCG-AGACCGAUAGC-G
AAC-A-AGUAGCGUGAGCGAACG-CUGCAA-AGCAC-CCCGAACAGGGC-G-GUGCAAUAGGGCCUGAACUC
A-GUCAGUGAUAG-AAC-GA-CGAG---GCAUACAA-GG---------------------------------
------------------------------------------------------------------------
------------------------------------------------------------------------
------------------------------------------------------------------------
------------------------------------------------------------------------
------------------------------------------------------------------------
------------------------------------------------------------UC-CCACUCAC-
---AA-----------CG-A-------------------C------CUG-----------------AGA---
------------------------------------------------------------------------
--------------------------------------------------------------G--C-AA---
------------------------------------------------------------------------
-------------------------------------------------U-CU------CA--------CUG
UAGAA-------------------------------------------------------------------
------------------------------------------------------------------------
------------------------------------------------------------------------
------------------------------------------------------------------------
------------------------------------------------C-----------------------
------------------------------------------------------------------------
------------------------------------------------------------------------
------------------------------------------------------------------------
------------------------------------------------------------------------
------------------------------------------------------------------------
-------------------GAGUGGGAA-----------------GCCGA--U---GUCGCGUCGUACGUUU
UGAAAAACGACCCAGGGAGUGCA-U-CCACCCGGCAA-GUCUAACU--G-G--------UCCAC--------
C-CA--GGAA--GACGCAGGGAAAC-CGA-----------------UAU----------------------G
ACCG-CAGUCAU----G-------------------------------------------------------
---AC-------------------------------------------------------------------
------------UACUC--AUGA------------------------------------CCA-GGGUCA-CC
GUAC-----------------------------------UUAC--------------------------AU-
G-C-GG----------------------GGAGUCGCGUGGAUGC-GACCCGAAACC-GAGUGAUCUACGCGC
GGGCAAGGUGAAGCGUGC-CGAAAG-GUGCGUGGAAGCCUGU-UAGGGAUGGUGUCCUACAAUACCC-UCCC
GUGACCCACGUGUAGGGGUGAAAGGCCCAUCGAACUCGG-CAACAGCUGGUUCCAGCCGACA-CAUAUCGAA
GUAUGUCCUCAG-CCGA-------------------------------------------------------
----------------------GG-----UA-GUCUGCGAGGUAGAGCUACCGAUU-GGA-UGUC-CCGCCU
C-C-GA-GA-GGAGUC-GGCCAUGCUGUCAAACUCCAAACUUGCAG--------------------------
```

```
------------------------------------------------------------------------
------------------A-CG-C--CG-UAGA--CGCUGGG-AAUCCGG-A-CUGC-G-GGG-UAAGCCUG
UG-G-UCCAUGAG-GGAGAC---AACCCAGAGCAG-G-GUUAAGGUCCCCAAGU--GU-GGA-UU-AA---G
-U-GCAA----CUGAAGGUUGUCUCGAGCCCUAAACAGCCGGGAGGUGAGCUUAGAAGCAGCUA-CCCUCUA
AGAAAAGCGUAAUAGCUUACCGGCC-----GAGGUUCGAGGCGCCCAAAAU-GAUC-GGGGCUCA-AAUCCA
C----------CACCGAG-ACCCUGCAAUACUUA--U-----------------------------------
------------------------------------------------------------------------
-----U--AU--------------------------------------------------------------
--------------------------------CAAGU-AACU-UCGUAGG-UGGC-GUACUGA---------
-----------C-C-GGAUG-GAAGUAC--G-GGCGAGAGCU--C-GUAUGGACCG-C-UCA-GUAACGAUA
AUCC-UGGUCA-CAGUA-GCAGCA-AUAGUC-GGGUGU--G---------------ACCCCC-GAC-GGCCU
A-ACGAGC-AAGGGUUCC-UCGGCACUG-CUAAUCAGCCGAGGGUUAGCCGG--UC-CUAAGUCUGACCGU-
AAU-UCGA---C-UCA--------------------------------------------------------
----------------------------------------GAC-AA-ACAGGGA-AACUGGUU-AAUA-UUC
CAGUG----------------------------------CCGUC----GUGU--------------------
------------------------------------------------------------------------
----------------------------------------------------------CUU-A--UCG-AAC
GCUUUCG-------------GAAACA-GUGAACGGG-GC-------CAUC-------------G-CCCCGU-
CGAA-U-CAU-GG-----A-A-------------------UC-----UC----------------GUGG-A-
U-G-CC--------------------GUAAC-GGCACGAAGC---------GGGAGA----AUC-AU-GAG-
ACC--GC-----------------------------------------------------------------
----------------G-CA-A------GUC-A-C-UGGUACCAAGAGC-CCGUGA-A-AAA-ACAC-----
---------------------------------------------------------------AC-GACGUC
CGUACCAAGAUCCGAC------------------------------ACAGGUGC-UC-AGGCA-GCGAAAGC
CAUGGC-CCGU-CGGGACCAACCGG-C-GUUAGGGAAUUCGGC-AAGUUAGUCCCGUACCUUCGG-AAGAAG
GGAUG-CCUG-C-UC--UUG----------------------------------------------------
------------------------------------------------------------------------
------------------------------------------------------------------------
------------------------------------------------------------------------
------------------------------------------------------------------------
------------------------------------------------------------------------
------------------------------------------------------------------------
------------------------------------------------------------------------
------------------------------------------------------------------------
------------------------------------------------------------------------
------------------------------------------------------------------------
------------------------------------------------------------------------
------------------------------------------------------------------------
------------------------------------------------------------------------
-------------------------------------------------------------------ACCGA
GCA--GGUCGCAGUGACUCGGACGCUCCGA-----CUGUCUAGUAACAACAUAGGUGACCGCAAAUCCG---
-CA-A--------------GGACUCGU-ACGGUCACUGA-AUCCUGCCAGUGCAG-G-UAUC-UGAACACCU
CG-------------UACA-------------AGAGGACGAAGGAC--CUGUUAAC-GGCGGGGGUAACUAU
GACCCUCUUAAGGUAGCGUAGUACCUUGCCGCU-UCAGUAGCGGCUUGCAUGAAUGGAUCAACGAGA--GCG
UCACUGUCCCAACG----CUG-GACCCGGUGAACUG--UAC---GUUCCAGUGCGGAGUCUGGAGACCCCCA
```

```
AGGGGAAGCGAAGA-CCCUAUAGAGC-UUUACU--GCAGGCU-GUCGCUGAGACAUGGUCGCUGAUGUGCAG
CAU-AGGUAGGAGCCGAUAC-------ACAGGUAUC-CGC----------------------------GCCA
--------------------------------------GCGGAUCACCGAG-G--C-AC-UCAUGAAAUA-C
UACC-CGUCAGUGACUGUGACUCUCACUC-C----------GG-G-A-------------------------
------------------------------------------------------------------------
------------------------------------------------------------------------
------------------------------------------------------------------------
------------------------------------------------------------------------
------------------------------------------------------------------------
-------------------------------------------------GGA-----AAACA-CCGGUAGCC
GGGCAGUUUGACUGGGGCGGUACGCGCUCGA-AAAGAUAUC-GAGCGCGCCCA-AAGGUCUCCUCACGUGAG
UCGGAAACUCACAGAC-GAGCG-CAAGAGCAC-ACGGAGGCCUGACA-GUGA--------------------
--------------UCUUCCAACGAGGAUCGCUGACGCGAAA-GCGUGGUCUAGCGAACCAAC--------
---GA-G-----GUUCA-GAG--------------------AU-GGGACCCGUUGAUGACAG-AA--AAGCU
ACCUUAGGG-AUAACAGAGUCGUCACUCGCAAGAGCACAUAUCGACCGAGUGGCUUGCUACCUCGAUGUCGG
UUCCCUCCAUCCUGCCUGUGCA-GACGCAGGCAAGGGUGAGGUUGUUCGCCUAUUAAAGGAGGUCGUGAGCU
GGGUUUAGACCGUCGUGAGACAGGUCGGCUGCUAUCUAUU--------------------G-GGG-------
-------------------------------GUGU--AAGGUACUUGCGAGGAACG-CUCGUAUAGUACGAG
AGGAA-CUACGAGUGGGUGCCAC-UGGU-G-UAUCGGUUGUCC-GACA-GGGCAGUU-GCCGAGCAGCCACG
CACCA-CAGGGUAAGAGCUGAAUGCA-UC--UAAGCUCGAAACCUCCCUCAA-A-A-A--GAAGUACC-AU-
------------------------------------------------------------------------
------------------------------------------------------------------------
------------------------------------------------------------------------
------------------------------------------------------------------------
------------------------------------------------------------------------
------------------------------------------------------------------------
------------------------------------------------------------------------
------------------------------------------------------------------------
----------------------------------------------------------GA-AGG------AU
CCUUCUAGAAGAGAAGGU-AGA-UAGACU-CGG-GGUGUACG-CAU-C--------GAG-----GC------
----AA--------CGAGAUGUUGAGCCC--G-CGAG-CACUAACC---AUCCAA-GC-CAUAAUC------
------------------------------------------------------------------------
------------------------------------------------------------------------
------------------------------------------------------------------------
-------------------------------------------------
>LSUa_228908_NANEQ/1-2955
------------------------------------C--UGACCCCGCCC-GGUGGAUG-C-CUGGGC-UCG
G-G---AGCCGAAGAAGGGCGUGG--CAAGCUGCGAUAAGCCAGGGGGAGGCGCACGCAGCCGU------UG
AUC-CCU-GGAUCCCCGAAUGGGACC-U--------CCU-------GC----CCGG-G-GG----------U
U-A------------AC---C----C----C---GGG----C-G-CCCC-----------------------
-----------G-----------AUAA---------------------------------------------
------------------------G-G-GG-CGGGAACGCGGGAAGUGAAACAUCUCAGUACCCG--CAGG
AAAAGAAAGAGAA-AUUCGAUGGCGGGA--GU-AG-GGGCGACCGAAACCGCC-G-UA-GCCCAAACUGAAU
CCCG-UAGGGA--------AA------------C---CUA-CGGGAGAUGUGGGG--UUGU-----U-----
G-GG--GGU-GGC-G-----G------C------------------UUUCC--CUC-----------CC--G
```

```
C-------GGGAA--CCCG------AACUCC-GGUG------------------------------------
-------GAA-C--CGGGG-------CCGUAGAGGGUGACAGCCCCGUA---------GGG---G----UAA
C------CGCGGAGG-GAGU----------------GCCG-CCACU--UCCAGAGUACCGCGCCUCG---GA
UAUGGCGCGGGAA-GCUGGGGGACACA--CGCCCCCAAGGCUAAAUA--CUC-C-CCG-AGUCCGAUAGC-G
GAC-U-AGUAGGGUGACCGAAAG-GUGAAA-AGAAC-CCGUG-GAAGGG-A-GUG--AAAGAGCCUGAAACC
G-GGCGGGGAUAG-UCA-GG-UGGG---GCCGGUUA-AGCCCC-----------------------------
------------------------------------------------------------------------
------------------------------------------------------------------------
------------------------------------------------------------------------
------------------------------------------------------------------------
------------------------------------------------------------------------
------------------------------------------------------------CG-AAG------
-----------------G-A-------------------A------CCC-----------------CCG---
------------------------------------------------------------------------
--------------------------------------------------------------G--C-GA---
------------------------------------------------------------------------
-------------------------------------------------C-GG------GG--------UAG
UA----------------------------------------------------------------------
------------------------------------------------------------------------
------------------------------------------------------------------------
------------------------------------------------------------------------
------------------------------------------------------------------------
------------------------------------------------------------------------
------------------------------------------------------------------------
------------------------------------------------------------------------
------------------------------------------------------------------------
------------------------------------------------------------------------
----------------------CUU-GC-----------------UAGCC--G---GUUCCACCGUACGUUC
GGGAGCACGUGCGGGGGAGUGUG-C-CCGAGUGGCGA-GGUUAAGG--G-G--------AUAAA--------
C-CC--CGAA--GCCGUAGGGAAAC-CGA-----------------AG-----------------------G
CCCG-CAGCCC-------------------------------------------------------------
------------------------------------------------------------------------
--------------UUA--UGGG------------------------------------CGA-GGGGCC-GC
GUCC-----------------------------------UAUA--------------------------CG-
G-G-CG----------------------UAAGUCACUCGGGCAC-GACCAGAAGCC-CGGCGAUCUACCCCG
GGGCAGGGUGAAGCGGGG-U-UUUU-CCCCGUGGAGGCCCGA--AGGGGUGGUGGC---CUAUCCCC-UCCU
CUGACCCCGGGGUAGGGGCGAUAUACCAAUCGAGCCGGG-UGAUAGCUGGUUCCCGCCGAAA-UGGGCUGCA
GUCCAGCUCGG-CGGA--------------------------------------------------------
-----------------------G-----GC-ACCCUCGGGGUAGAGCGACGGAUU-CCC-GUG--GCAAGG
G-G-GU-UA-ACCC-C-ACCACGGGAGUCCAACUCCGAACCCG-AG--------------------------
------------------------------------------------------------------------
------------------G-GG-C--CG-UAGA--UGCCGGG-AGACGGG-G-GCGG-C-GGA-UAAGCCGC
CG-U-UCCGAGAC-GGGAAC---AACCGAGACCGG-G-GUUAAGGCCCCUAAGU--GC-CGG-CU-AA---G
-U-GGC-----UGGAAGGGUGUCCGCUCCCUAAGACACCGGGGAUGUGGGACUAGUAGUGUCCA-UCAUCUA
AGGAGUGUGCAACAACUCACCCGGC-----GAGGGAGCGGGCCCCGCAGAU-GGAC--GGGCCUA-AGCCGG
```

```
C----------CGCCGAG-ACCCCGGGCCGGGGA--U-----------------------------------
------------------------------------------------------------------------
-----CCGAUG-------------------------------------------------------------
-------------------------------GAUCCC-CGUG-CGGUAGGCGGGC-GUCCCGG---------
-----------U-G-GCCUA-GAAGGCG--G-GCCGUGAGGU--C-CGCUGGAGCC-G-CCG-GGAAUGAGA
AUCC-CCGCGU-GAGUA-GGAGCA--AAGAG-GGGUGA--G---------------AAGCCC-CUC-CGCCG
AAAGGGAA-AAGGGUUCC-CUGGCAAUG-ACCAUCAGCCAGGGGUGAGUCGG--UC-CUAAGGCCGGCGGU-
AAC-UCCG---C-UCG--------------------------------------------------------
----------------------------------------GCC-GA-A-AGGGA-AAGGGGUU-AAUA-UUC
CCCUA----------------------------------CCGCC----CGGGUAGG-------C--------
------------------------------------------------------------------------
-------GCGG--CAAC--------GCCAA-----GC--------------------CCUA-C-UCCC-GAC
GCCUGGG-------------GCUAGG-CCGACCGCG-GA-------UUUC-------------C-CGCGGC-
CAAC-C-GGG-AU-----A-G-------------------GG-----CC----------------GGGG-AG
U-G-CC--------------------AUUAU-GGCGAGAACC---------GGCCAA----ACC-CG-GGA-
CGG--GG-----------------------------------------------------------------
-----CCCCC----GUAUGGGGGCU-U-CGG-C-C-GAGCCCUGGGCCC-AUGAAA-A-GGG-AGUAGGGAA
--------------------------------------------------------CGAUCC-CG-GGCGAC
CGUACCGAGAACCGUG------------------------------ACAGGU-U-CCCUGGGU-GAGCAGCC
UAAGGC-GGUG-GGGU-AUAACCUG-G-CCGAGGGAAAUCGGC-AAAUUGGCCCCGUAUCUUCGG-UAGAAG
GGGUG-CCUG-C-GG--UCGU---G----GGGU---------------------------------------
------------------------------------------------------------------------
------------------------------------------------------------------------
------------------------------------------------------------------------
------------------------------------------------------------------------
------------------------------------------------------------------------
------------------------------------------------------------------------
------------------------------------------------------------------------
------------------------------------------------------------------------
------------------------------------------------------------------------
------------------------------------------------------------------------
------------------------------------------------------------------------
------------------------------------------------------------------------
------------------------------------------------------------------------
----------------------------------------------------------AAUCCU--GCGACC
GCA--GGUCGCAGUGACAAGGGCGGGCCGA-----CUGUUUACUAAAAACAUAGCCUCCGGCAAGCCCG---
-AA-A-------------GGGUGUGU-ACCGGAGGUGA-CGCCUGCCCAGUGCCG-G-UACC-UGAACCCCC
CG-------------UCCA-------------AGGGGGCUAAGGGC--CGGUAAAC-GGCGGGAGUAACUGU
AACUCUCUUAAGGUAGCGUAGUCCCUUGCCGGU-UAAAUGCCGGCUUGC--GAACGGCGUAACGAGC--UCG
CCACUGUCCUCGGC----CAG-GGCCCCCUGAAAAC--GCC---GUUCCCGUGGAUAGGCGGGAGAGGCCCG
GUGGGAAGAGAAGA-CCCCGUGAAGC-UUAACC--GCAGCCU-GUCGUUGCCCCGUUGUCCCCCACGCUCAG
CGU-AGCGGGGAGCGAU-G----------AACCGGC-CCC----------------------------UCCG
--------------------------------------GG-GGUCGGGGAU-G--C-GC-CCAUGAGACA-C
CCGC-CGUGGGGGACAACGGGGCUAACCC-C----------GA-A-A-------------------------
------------------------------------------------------------------------
```

```
------------------------------------------------------------------------
------------------------------------------------------------------------
------------------------------------------------------------------------
------------------------------------------------------------------------
-------------------------------------------------GGG-----GGACA-GCGGCAGGU
GGGCGGUUUGGCUGGGGCGGCACACCGCCGA-AAAGAUAUC-GGCGGUGCCCA-AAGGUCGGCUCAGGCGGG
UCAGAAAUCCGCCGUA-GAGUG-CAAGGGCAA-AAGCCGGCCUGACA-CGGC--------------------
--------------CCUUUA---CAAGGGGCCGUGGCGGGAAA-CCGCGGCCUAGCGAACCCCG--------
---GU-A-----GCCCC-CGU--------------------GU-GGGGCUUCGGGACGACGG-AA--AAGCU
ACUCCGGGG-GUAACAGCUUGGUCGCGGGCGAAAGUUCCCAUCGACCCCGCGGUUUGAGACCUCGAUGUCGG
CUCUUCUCACCCUGGCCGUGCA-GCAGCGGCCAAGGGUGGGGGUGUUCACCCAUUAAAGAGGAACGUGAGCU
GGGUUUAGUACGCCGCGAGGCAGUGUGGAUUCUAUCUACC--------------------G-GGC-------
-------------------------------CUGU--UCGGCGGCUGAGGGGAAGG-ACGGCCUAGUACGAA
AGGAA-CGGCCGUCCGGCGCCUC-UAGU-C-GACGGGGUGUCC-GUAU-GGGCAG---CCCCGGCAGCCACG
CGCUA-GGGGGUAAGGGCUGAAUGCA-UC--UAAGCCCGAACCCCUCCCCGA-A-AAG--AGCCGCCG-AU-
------------------------------------------------------------------------
------------------------------------------------------------------------
------------------------------------------------------------------------
------------------------------------------------------------------------
------------------------------------------------------------------------
------------------------------------------------------------------------
------------------------------------------------------------------------
------------------------------------------------------------------------
----------------------------------------------------------C---CC------CC
UCGGGUAGAAGACCCGGU-UGA-UGGGGG-GGC-GGUGGAAG-CCC-C--------GAG-----GG------
--UUUC--------CGAGGGGCG-AGCCG--G-CCCC-UCCCAAUC---GGGGGGUGG-UCCUGCG------
------------------------------------------------------------------------
------------------------------------------------------------------------
------------------------------------------------------------------------
-------------------------------------------------
>LSUa_1072681_HALSG/1-2928 AGNT01000024.1, Contig12, whole genome shotgun s
--------------------------UCCUA----CU--CAAGUCUAGAU-AGUGGAUG-G-CUCGGU-UCG
A-GA--AGCCGAUGAAGGACGUGG--UAAGCUGCGAUAUGCUUGGGCGAGGUGCAUGCAGCCUU------AG
AGC-CCA-AGAUUUCCUAAUGCGACU-UC-----------------------------------------GU
A-G-----------GUC---G----C----U--UCGG----C-G-AUCU-GGC-------------------
-CC--------G-----------UCA----------------------------------------------
------------------------G-G-GU-CGGGAACGCGGUGAACUGAAACAUCUAAGUAGCCG--CAGG
AAAAUAAAGAAAC-AUUCGAUUCCCUUA--GU-AG-CGGCGAGCGGAAGGGGA-AUUA-GCGCAAACUGAAC
GCCU-UGAAGU--------GA------------U---UCA-AGGUGGAUGUGGUU--CGCCCGUCUU-----
U-GG--AAC-CGA-CUU---U------CU-----------------CAGGUG--------------------
-------------------------AACUCU-GCUG------------------------------------
-------GAA-UG-CAGGA-------CCAUAGAGAGUGACAGUCUCGUA---------ACC---U----ACA
G------AAAGAACG-UUCU----------------UUGA-CGUGU--GCAUGAGUACCAUGGGUUG---GA
AAUCCCGUGGGAA-AACGGAAGGCACA--UUCUUCCAACGCUAAAUA--CUU-C-UCG-AGUCCGAUAGC-G
UAC-UAAGUACCGUGAGGGAAAG-CUGAAA-AGUAC-CCCUAGAAAGGG-G-GUU-AAAAGAGCCUCAAACU
```

```
A-UCUAGUGUUAG-GCUAGU-UAUG---GCAUUCAA-GCGUC------------------------------
------------------------------------------------------------------------
------------------------------------------------------------------------
------------------------------------------------------------------------
------------------------------------------------------------------------
------------------------------------------------------------------------
------------------------------------------------------------GA-UACAGGUG-
---AA-----------AG-A-------------------A------UCU-----------------GCC---
------------------------------------------------------------------------
--------------------------------------------------------------GA---GA---
------------------------------------------------------------------------
-------------------------------------------------G-GC------UU--------UAG
UAUGA-------------------------------------------------------------------
------------------------------------------------------------------------
------------------------------------------------------------------------
------------------------------------------------------------------------
------------------------------------------------A-----------------------
------------------------------------------------------------------------
------------------------------------------------------------------------
------------------------------------------------------------------------
------------------------------------------------------------------------
------------------------------------------------------------------------
-------------------UCCUGU-GG-----------------AGCAG--U---GUCAUAGCAUCCGUUU
UGAAAAACGGGGCAGGGAGUUUG-U-CGCCGUGGCAA-GACUAAGU--C-G--------UUUAC-------G
C-GA--UGGA--GUCGUAGCGAAAG-CAA-----------------GAA----------------------G
CCCG-CAGCAG-------------------------------------------------------------
------------------------------------------------------------------------
--------------UUU--UCUG------------------------------------UGA-GGGGCC-UC
GUGU-----------------------------------UAAC--------------------------GC-
G-C-GA----------------------GAAGUCACCGCGACAU-GAUCCGAAACC-GGUCGAUCCUGGCCU
GGUCCGGCUGAAGCGGCG-CGAAAG-CUCCGUGGAAGGCCUC--ACCGUUGCUGCGUUACAAG-CGC-UCGG
GUGAACUGGGUCAAGGGGUGAAAUGUCAAUCGAGAUCGG-UGAUGGCUGGUUCUUCUCGAAA-UGGAUCGAA
GUCCAGCCUCGA-GCGA-------------------------------------------------------
----------------------GA-----CG-GCCUAAGAGGUAAAGCUACUGAUU-CUG-GGUU-CUGGAG
GUU-UA-----CCUCC-AGACCCGGAGUCAAACUCUGAACUCUUAG--------------------------
------------------------------------------------------------------------
------------------G-CG-U--UG-CAGA--CCUCGGG-AGUGAGG-G-ACAG-U-GCG-UAAGGUAC
UG-U-UCCUAGAG-GGAAAC---AACCCAGAGCAU-G-GUUAAGGUCCCACAAU--UU-CUG-ACUAA---G
-U-GUCAUUG-----AGGAUGUCCUGGUUCUCAGACAGCGGGGAGGUUAGCUUAGAAGCAGCUA-CCCUUUA
AAGAUAGCGUAACAGCUCACCCGUU-----GAGAGCUGGGGCGCCGUAAAU-GGAC-GGGGCUCA-AGUCAG
A----------AACCGAU-ACCAUGUC---------------------------------------------
------------------------------------------------------------------------
------------------------------------------------------------------------
-------------------------------CGCGAG-AGCG-GGUUAGAGAAGC-GAUCUGC----------
-----------A-U-GGGUG-GAAGCAU--G-GCCGUGAGGU--C-GUGUGGACCG-U-GUA-GAUACGAGA
```

```
AUCC-UGCCGG-CAGUA-UCAUCAUUAAGAA-CAGUGA--G---------------AACCUG-UUC-CACCG
U-AAGAGU-AAGGGUUCC-UCUGCAAUG-AUUAUCAGCAGAGGGUUAGUCGG--UC-CUAAUCCACACCGU-
AAU-UCGA---A-UGU--------------------------------------------------------
----------------------------------------GGA-GA-AUAGGGA-AACAGGUU-AAUA-UUC
CUGUA----------------------------------CC----------------------U--------
------------------------------------------------------------------------
----------------------------AAUC-------------------------ACAU-U-UUCU-GAC
GUUUUGG-------------GAUAGC-GGAAGCUGC-GC-------CACC-------------G-CGCAGU-
CUAA-G-----CA-----A-G-------------------UA-----AU----------------GCGG-AC
G-A-GC--------------------GUAAU-GCAUCGACUC-------C---GUGA----ACU-UG-UGA-
AUG--GC-----------------------------------------------------------------
-----CCGGGA-AAAAUCCCAGGUU-U-CCG-C-A-AGUCCUGGAAUCC-UUGAAA-A-AGU-G--------
---------------------------------------------------------------UG-AAAGAC
CGUACCCAGAACCAGC------------------------------ACAGGUAC-UC-UGGCU-GAGUAGGC
UAAGGU-GUUC-GGGU-AUAACGCA-G-CUUAGGGAAUUCGGC-AAGUUAGCCCCGUACCUUCGG-UAGAUG
GGGUG-CCUA-U-CC--ACUU---------------------------------------------------
------------------------------------------------------------------------
------------------------------------------------------------------------
------------------------------------------------------------------------
------------------------------------------------------------------------
------------------------------------------------------------------------
------------------------------------------------------------------------
------------------------------------------------------------------------
------------------------------------------------------------------------
------------------------------------------------------------------------
------------------------------------------------------------------------
------------------------------------------------------------------------
------------------------------------------------------------------------
------------------------------------------------------------------------
------------------------------------------------------------------GAUUGG
AUA--GGUUGCAAUUACUAGGGGAGCUCGA-----CUGUUUACUAAAAACACAGCCGACUGCUACUCCG---
-AA-A-------------GGACUUGU-AUAGUCGGCGA-CUCUUGCCCAGUGCCA-G-CACC-UGAAACCUC
CG-------------UACA-------------AGGAGAAGAAGGGC--UGGUAAAC-GGCGGCGGUAACUAU
GACCGUCUUAAGGUAGCGUAGUCCCUUGCCAAU-UAAUUGUUGGCCCGCAUCAAUAGAGUAACGAGG--CUC
CUACUGUCCCAAGC----UGU-GGCCCGGUGAACCC--GCC---GUACCGGCGCAUAGGCCGGUGACUCCUG
GUGGGAAGCGAAGA-CCCUUUGAACC-UUUACU--GCAGUCU-AUUGUUGCGGUGGGGUUCCAAGUGUGAAG
UGU-AGGUAGUAGACAUUAC-------AGAGACGCA-GUC----------------------------GCAA
--------------------------------------GGCUGUCGCCCAG-U--C-GA-CUAUGGGACA-C
UACC-CUCUUGGAACCCCGCUGCUUACUU-CU--AC----------G-------------------------
------------------------------------------------------------------------
------------------------------------------------------------------------
------------------------------------------------------------------------
------------------------------------------------------------------------
------------------------------------------------------------------------
-------------------------------------------------GAA-----GGACA-UCAAUAGCU
```

```
GGGCAGUUUAGCUGGGGCGGCAUCCCUCUGA-UAACGAAUC-AGAGGGGUCCA-AAAGUAGGCUCAACCCGU
UCAGAAAUCGGGUGUA-GAGGA-CAAAGCCAA-AAGCCUGCUUGACU-GGAA--------------------
--------------UCCUGAUAGUAGGGUUUCCAGGCGCAAA-GCGUGGCUUAGCGAACGCCU---------
---AU-AGCCAAAAUCU-UUU--------------------GU-GAGGCUUAGGCCUGACAA-GA--AAGGU
ACUCAAGGG-AUAACUGGAUUGUCUGAGGCGCGAGUUCACAUCGACCCUCAGGCUUGUUACUUCGAUGUCGG
CUCUCCGUAUCCUGGCUGUGCA-ACAGCAGCCAAGGGUGAGGGUGUUCACCUAUUAAAACGGAUCGUGAGCU
GGGUUUAGAUCGGCGUGAGCCAGAUCUGUUGCUAUCUGCC--------------------A-GGA-------
-------------------------------GUGUGUUUGUGGUCUGAGAGGAAAG-UCCGGACAGUACGAA
AGGAA-CCCCGGACUGAGGCCUC-UGGU-G-UAUCUGUCAUCC-GGAA-GGGUGU-U-GCAGAGUAGCUACG
CCUCGCAGAUGUAAGAGCUGAAAGCA-UC--UAAGCUCGAAAGUCUCCUCAA-A-AAG--AGACCGCG-UC-
------------------------------------------------------------------------
------------------------------------------------------------------------
------------------------------------------------------------------------
------------------------------------------------------------------------
------------------------------------------------------------------------
------------------------------------------------------------------------
------------------------------------------------------------------------
------------------------------------------------------------------------
----------------------------------------------------------CG-----------C
GCCCAUACAAGAU-GGGU-UGA-UAGAGU-GGC-GGUGUAAG-AUC-C---------AAG-----GCAA----
--------------CGAGGAUUUCAGCCU--A-CCAC-UACUAAGA---GCGAAUCUC--------------
------------------------------------------------------------------------
------------------------------------------------------------------------
------------------------------------------------------------------------
-------------------------------------------------
>LSUa_1577684_NANOP/1-2926 CP010514.1, complete genome; RANGE 572395:569490
----------------------------UCU----CA--GGAACCCGCUC-GGUGGAUG-C-CUCGGC-UCG
G-A---AGCCGAAGAAGGGCGUGG--CAAGCUGCGAAAAGCUGGAGGGAGGUGCAUGCAACCGC------UG
AUC-UCC-AGAUUCCCGAAUGGGAUA-U--------CCU-------GC----CGA---------A-------
----------------------------------AGG----C-A-UCCC-----------------------
-----------G-----------UAA----------------------------------------------
------------------------G-G-GA-GGGGAACGCGGGGAAGGGAAGCAUCUUAGUACCCG--CAGG
AAUAAAAAACAAA-UCGUGAUUUCCCGA--GU-AG-GGGCGACCGAAAGGGAA-G-AA-GCCCAAACCGAAU
CCUC-UAAAGU--------AA------------U---UUA-GAGGAGAUGUGGGG--UAAAGG-AGC-----
G-GG--AGC-UUC-CC----G------UUAUUAUAUUC----------------------------------
---------------------UCAGAACUCC-GGUG------------------------------------
-------AAA-GUCCGGGG-------CCGUAGAGGGUGACAGUCCCGUA---------UGG---G----AAU
A------UAAUAACG-GGA-----------------UUUC-UCGCU--UCCGGAGUACCGGCGGUUG---AA
UUUCCGUCGGGAA-UUUGGGGGAAACU--AGUCACCAAGGCUAAAUA--CUU-U-CCG-AGUCCGAUAGC-G
UAU-U-AGUAGGGCGACCGAAAG-GUGAAA-AGAAC-CCUUG-GUAGGG-G-GUG-AAAAGAGCCUGAAACC
G-GGCGGGGAUUG-UCU-GG-UAGG---GCUAGUUA-AGC--------------------------------
------------------------------------------------------------------------
------------------------------------------------------------------------
------------------------------------------------------------------------
------------------------------------------------------------------------
```

------------------------------------------------------------------------
---------------------------------------------------------------ACCUUAUC-
---UG-----------GG-A-------------------A------AAU-----------------CUG---
------------------------------------------------------------------------
--------------------------------------------------------------G--U-GA---
------------------------------------------------------------------------
-------------------------------------------------C-GG------GU--------GAG
UACCA-------------------------------------------------------------------
------------------------------------------------------------------------
------------------------------------------------------------------------
------------------------------------------------------------------------
------------------------------------------------G-----------------------
------------------------------------------------------------------------
------------------------------------------------------------------------
------------------------------------------------------------------------
------------------------------------------------------------------------
------------------------------------------------------------------------
-------------------AUAAGGUGC-----------------UAGCU--A---GUUCUACCGUGAGUUU
UGAAGAACUUGCGGGGGAGUAUA-C-CCGAGUGGCGA-GGUUAAGG-------------UUUAU--------
--AC--CGAA--GCCAUAGGGAAAC-CGA-----------------AA-----------------------G
CCCG-UAACCUG----G-------------------------------------------------------
---U--------------------------------------------------------------------
--------------UUC--CAGG------------------------------------UGA-GGGGCA-CC
GUCC-----------------------------------CAUAA-------------------------GG-
G-CGGU----------------------UUAGUCACUCGGGUAU-GACCGGAAACC-GGGCGAUCUAUCCCG
GGGCAGGGCGAAGCGGGG-----UU-UCCCGUGGAGGCCCGA--AGAGGUG-----GGGCUUACCUC-UCUC
AUGACCUCGGGGUAGAGGUGAAAAGCCACUCGGGCCCGG-UCAUAGCCGGUCCCCGCCGAAA-CUGACCCUA
GCCAGGUCUCGG-GUUA-------------------------------------------------------
----------------------GG-----AU-CCCUAUGGUGUAGAGAGACGGAUU----------------
--C-UGUAC-GGGUUU-CCGUAUGGAGUCCAACUCCGAAGCCAUAG--------------------------
------------------------------------------------------------------------
------------------G-GU-C--CG----U--AGCCCGGGAAACGGG-A-AUUG-C-GGG-AAAGCCGC
AA-U-UCCGAGAC-GGGAAC---AACCGAGACCGA-G-GUUAAGGCCCCAAAGU--GC-UGG-CU-GA---A
-C-GAC-----GUUGAAGGAGUCUGUCCCCUAAGACAUCGGGGAUGUGGGCCUAAUAGUGGCUA-UCAUCUA
AGGAGUGUGUAACAACUCACCCGAC-----GAGGGGAUAGGCUCCGAAAAC-GGAC--GGGUCUA-AGCCAG
C----------CGCCGAG-ACCUCGGUCCGCUGG--C-----------------------------------
------------------------------------------------------------------------
-----AU-----------------------------------------------------------------
-------------------------------UGCCAG-CGUG-AGGUAGGCGGGC-GUCCCGA---------
-----------U-G-GCCUA-GAAGGGU--U-CCCGAAAGGA--A-GCCUGGAGCC-G-UUG-GGAAUGCAG
AUCC-CCGCGG-UAGUA-AGAAU---CAGAC-GGGUGA--G---------------AAUCCC-GUC-GGCCG
A-AGGGGGAAAGGUUUCC-CAGGCAAUG-AAGAUCAGCCUGGGGUUAGUCGA--UC-CUAAUCCAGGCUCC-
GAG-AGGA---G-UCU--------------------------------------------------------
----------------------------------------GGA-GA-A-AGGGA-AAGGGGUU-AAUA-UUC
CCCUA----------------------------------CCGCC----CCGAUAGA-------U--------

```
------------------------------------------------------------------------
-------GCGG--UAAC--------GC-AACAUCCUU-------------------UCCCG-A-CCUG-GAA
GGCUAGG-------------CAUACA-CGGGGGUGGGGU-AAAACCCACU---------------UACUGUG
CUAA-GGGGG-AU-----A-A-------------------GA-----CC----------------GGGG-AG
U-G-CC--------------------GUCAU-GGCGAGAACC---------GGUCUA----AAC-CC-UGA-
UGG---------------------------------------------------------------------
------------GCCCGAAAGGGUU-A-UGC-U-G-AGGCCUUCCGGCA-AUGAAA-A-GGG-AAAGGAUG-
--------------------------------------------------------CGAUUC-GG-GGCGAU
CGUACCGAAGUCCAGA------------------------------AUAGGU-C-CCCUGGGU-GAGCAGCC
UAAGGC-CGUG-GGGG-UCAAGCCG-G-UCUAGGGAACUUGGC-AAAUUGGUCCCGUAACUUCGG-GAGAAG
GGAUG-CCUG-C-GG--UGU----------------------------------------------------
------------------------------------------------------------------------
------------------------------------------------------------------------
------------------------------------------------------------------------
------------------------------------------------------------------------
------------------------------------------------------------------------
------------------------------------------------------------------------
------------------------------------------------------------------------
------------------------------------------------------------------------
------------------------------------------------------------------------
------------------------------------------------------------------------
------------------------------------------------------------------------
------------------------------------------------------------------------
------------------------------------------------------------------------
-------------------------------------------------------------------GGACC
GCA--GGUCGCAGUGACAAGGGCGGGCGGA-----CUGUUUACCAAAAACAUAGCUCCCAGCAAGCCCG---
-AA-A-------------GGGUUUGU-ACUGGGGGUGA-CGUCUGCUCAGCACCG-G-UAUC-UGAAUCCCC
CG-------------UUCA-------------AGGGGGGUAAGGAC--CGGUAAAC-GGCGGGGGUAACUCU
GACCCUCUUAAGGUAGCGUAAUGCCUUGCCAGU-UAAAUGCUGGCUC-UGCGAACGACGUAACUAGC--CCG
UCACUGUCCCGGGC----CGG-CGCCCCCUGAAA---AGCU---AUCCUGGUGAGAAGGCCAGGGACUUCCG
AUGGGUAAGAAGGA-CCCCGUGAAGC-UUAACC--GUAGCCU-GUUGUUGCUCUGAAGAACCUGGUGUGUAG
GGU-AGCGGGGAGCAAU--------G---AAUUCCU-GCC----------------------------GCUA
--------------------------------------GG-UAGGAAGGAU-G--C-GA-CGAUGAAACA-C
CCGC-CUCUAGGUUCUUCAGGGCUAACCC-GA--A-------------------------------------
------------------------------------------------------------------------
------------------------------------------------------------------------
------------------------------------------------------------------------
------------------------------------------------------------------------
------------------------------------------------------------------------
-------------------------------------------------GGG-----GGACA-GCAGCAGGU
GGGCGGUUUGGCUGGGGCGGCACCCUGCCGA-AAAGGUAUC-GGCAGGGUCCAUAAGGUCAGCUCAGGCGGG
UCAGGAAUCCGCCGUA-GAGGA-CAAGGCCAA-AAGCUGGCUUGACU-GUGC--------------------
--------------CCUGACCAUCAAGGGGCACAGGGGGGAAA-CCCGGCCUAGCGAACCCCG---------
---UA-C-----UCUCC-CUA--------------------GU-GGGGGUACGGGCUGGCGG-AA--AAGCU
ACUCCGGGG-GUAACAGCGCCGUCGCGGGCGAGAGCCCCCAUUGACCCCGCGGUUUGCGACAUCGAUGUCGU
```

```
CUCUCCCAUCCUGGUGGUGCA-GAAGCCGCCAAGGGUGGGGGUGUCCACCCAUUAAAGGGGAUCGUGAGAU
GGGUUCAGAACGCCGCGAGGCAGUUCGGUUA-UAACUGUC--------------------G-GAA-------
-------------------------------GUGC--UGGCUGCCUGAGGGGAAGG-UCCACCUAGUACGAA
AGGAA-CGGUGGGCCGGGGCCUC-UGGU-A-UACGGGCUGUCC-GUGA-AGGGCA-U-GCCCGGUAGCUACG
CCCUA-GGGGAUAAGCCCUGAAAGCA-UC--UAAGGGUGAAGCCCUCCUCAA-A-AAU--AGGCAGCC-GU-
------------------------------------------------------------------------
------------------------------------------------------------------------
------------------------------------------------------------------------
------------------------------------------------------------------------
------------------------------------------------------------------------
------------------------------------------------------------------------
------------------------------------------------------------------------
------------------------------------------------------------------------
----------------------------------------------------------UA-AGG------GG
UCGGAUAAAAGAUCCGAC-U---CAGAGC-GGC-GGUGUAAG-CUC-C--------GAA-------------
---------------AAGGAGUUCAGCCG--A-CCGC-UACUAAAU---GUUUGAAAA-UCUG---------
------------------------------------------------------------------------
------------------------------------------------------------------------
------------------------------------------------------------------------
-------------------------------------------------
>LSUa_263820_PICTO/1-2893
----------------------AUUGUGUAC----UG--CUAAACUGUCU-GGGGGAUG-G-UUUGGC-UUG
A-G---AGCCGAAGAAGGACGUGC--CAAGCUGCGAAAUGCUCUGGGGAGGCGCAUGGAGCCUU------AG
AUC-CAG-AGAUCUCCGAAUGGGACU-U--------CCU-------GA------------------------
--U---------------------------------U----C-A-CUCC-----------------------
-----------G-----------UAA----------------------------------------------
------------------------G-G-AG-AGGGCACCCCGGGAAUCGAAACAUCCUAGUACCGG--GAGG
AAAAGAAAUCAA--UCGAGAUACCGUGA--GU-AA-AGGCGAUCGAAAGCGGU-A-GA-GGGCAAACCGAAU
CCUA--CCCGA--------AA------------G---GGC-AAGGAGAUGUGGAG--UA-------------
U-GG--ACU-UUC-C-----C------C------------------AACGC--CUC-----------UU--U
U-------AUCGA--UUCG------AAU-AA-GCUG------------------------------------
-------GGA-AG-CUUAG-------CCAUAGAGGGUGAUAGCCCGUA----------GAA---G----AAG
A------UAAAAAGG-CCAUGA--------------GGGG-AAGUA--UCCUGAGUACCAUGCGUCG---UU
AUUCGCGUGGGAA-UACGGGUGGCAUG--AACAUCCAACCCUAAAUA--CAU-C-UCA-AGUCCGAUAGC-G
AAC-A-AGUACCGUGAGGGAAAA-CUGAAA-AGCAA-CCCGG-AAGGGC-U-GUG-AAAAGAGCCUGAAACC
A-GACAGUGAAAA-CUU-UA-UAGG---GCACUAAA-GGGGU------------------------------
------------------------------------------------------------------------
------------------------------------------------------------------------
------------------------------------------------------------------------
------------------------------------------------------------------------
------------------------------------------------------------------------
------------------------------------------------------------GA-AC-------
------------------------------------------------------------------------
------------------------------------------------------------------------
--------------------------------------------------------------C--G-AA---
```

------------------------------------------------------------------------
------------------------------------------------------------------------
------------------------------------------------------------------------
------------------------------------------------------------------------
------------------------------------------------------------------------
------------------------------------------------------------------------
------------------------------------------------------------------------
------------------------------------------------------------------------
------------------------------------------------------------------------
------------------------------------------------------------------------
------------------------------------------------------------------------
------------------------------------------------------------------------
----------------------AGG-UG-----------------ACCGG--U---GUCCUAUAGUCCGUGU
UGAAGAACGGGCCAGGGAGUUGC-G-ACGGUUGGCAA-GGCUAAUU--C----------UUAAA--------
--GA--AGGA--GCCGUAGCGAAAG-CGA-----------------UU-----------------------A
CCCG-CACAGC-------------------------------------------------------------
------------------------------------------------------------------------
--------------------AAU------------------------------------GGA-GGGGUA-GC
GUAC-----------------------------------CCAA--------------------------GU-
G-C-GU----------------------UUAGUCAGCCGUGGCA-GACCCGAAGCC-GCUCGAUCUACGCCU
GAGUAGGUUGAAGCUUAG-CGAAAG-CUAGGUGGAGGACCGA--ACCGGUUCUGAUGUGCAAAUCGU-UCGG
AUGACUUGGGUGUAGGGGUUAAAGGCUAAUCUAGAGCGG-CAAUAGCGGGUUCCCCCCGAAA-CUACCCGCA
GGUAGACCUCCA-UGGA-------------------------------------------------------
----------------------GA-----UU-UCGGAUGAGGUAGAGCGACCGAUU-GGC-CGGU-UAGUAG
U-C-GA-AA-GACU-G-CGCCGGCCUAUCAAACUCCGAACUUGUCC--------------------------
------------------------------------------------------------------------
------------------G-AA-U--CG-UAGA--UAUGGGG-UGCUAGG-G-CAUA-G-GGA-UAAGCUUU
AU-G-CCCGUGAA-GGGAAC---AACCUAGACCUG-G-AUUAAGGCCUCUAAAU--AC-UGG-CU-AA---G
-U-GCGA----UCUAAAAAGGUUUGCGGCCGAAGACAGUGGGGAGGUUGGCUUAGAAGCAGCCA-UCCUUUA
AAGAGUGCGUAACAGCUCACUCACC-----GAGGCUACAUGCCUUGAAGAU-GGAA-GAGGCUAA-AGCCAG
U----------UGCCGAG-AUCCAGGACCACCGC--------------------------------------
------------------------------------------------------------------------
------------------------------------------------------------------------
--------------------------------AAGGU-GAUG-UGGUAGGGGGC-GUGCUGU----------
-----------A-U-GGGAC-GAAGCUU--C-UUCGAGAGGA--G-GAGUGGACCG-U-ACA-GUAUCGCGG
AUCC-UGGUAA-AAGUA-GCAGUC--AAGAG-UGGUGA--G---------------AAUCCA-CUC-CGCCG
A-AAGGGC-AAAGGUUCC-UUGGCAAUG-UUCGUCAGCCGAGGGUUAGCCGA--UC-CUAAGGCCAUUCCU-
AAC-AGGA---A-AUG--------------------------------------------------------
----------------------------------------GUC-GA-A-AGGGA-AGCUGGUU-AAUA-UUC
CAGCG----------------------------------CCUGA----GGUGUUC--------U--------
------------------------------------------------------------------------
-------GC------------------------------------------------CAUG-C-AUGU-GA-
GCUUCGG-------------GAUAGG-AGGAGCACA-CC-------CGUC-------------A-GUGUGU-
UGGU-G-CAU-AU-----A-A-------------------GC-----AG----------------AUGG-AG
A-G-UU--------------------GUAAU-GACGUAAAGU---------CUGCGA----AGA-UG-UAU-

CU----G-----------------------------------------------------------------
-----UCCCC----UUAUGGGGAUU-U-CUC-U-A-AUUCCUGGAGCCC-AUGAAA-A-GCA-UGCAUGG--
--------------------------------------------------------GUCAAG-CC-CCAGAU
CGUACCAAGAACCGAC------------------------------ACAGGUGC-CCCUGGAU-GAGAAGUC
CAAGGC-GUUU-GGGA-AUAAUGGA-C-GCGAGGGAACUCGGC-CAAAUAGCUCCGUAUCUUAGG-UAGAAG
GAGUG-CCUA-U-CC--AGA----------------------------------------------------
------------------------------------------------------------------------
------------------------------------------------------------------------
------------------------------------------------------------------------
------------------------------------------------------------------------
------------------------------------------------------------------------
------------------------------------------------------------------------
------------------------------------------------------------------------
------------------------------------------------------------------------
------------------------------------------------------------------------
------------------------------------------------------------------------
------------------------------------------------------------------------
------------------------------------------------------------------------
------------------------------------------------------------------------
------------------------------------------------------------------GAGAGG
AUA--GGUCGCAGUGGCAAGGGGACCCCGA-----CUGUUUACCAAAAACACAGAUCACUGCUAGUCCG---
-UA-A-------------GGAUGUGU-AUAGUGGUCGA-AACCUGCCCAGUGCUG-G-UACC-UGAAAGCUC
CG-------------CUCA-------------AGGAGAAGAAGGGC--CAGUAAAC-GGCGGGGGUAACUAU
GACCCUCUCAAGGUAGCGUAAUACCUAGCCGCU-UAAUUGGCGGCUUGCAUGAA-GGUUCAACGAGA--GUC
CCACUGUCCCCGCG----UUC-AGCCCCGUGAAAUU--AAU---GUACUGGUGCACAAUCCAGUCUCUCCCA
CGUGAAAGCGAAGU-CCCCGUGGAGC-UUUACU--GCAGCCU-GUAGCUGUGACGCGAUUCUCAAUGCGUAG
GGU-AGGAAGGAGCCAUCG----------AAGUUCC-GGU----------------------------UGUG
--------------------------------------GC-CGGAAUGGAG-G--C-GA-CGAUGAAACA-C
UUCC-CUUUGGGAGUUGCGUCACUAACCU-U----------GC-A-A-------------------------
------------------------------------------------------------------------
------------------------------------------------------------------------
------------------------------------------------------------------------
------------------------------------------------------------------------
------------------------------------------------------------------------
-------------------------------------------------AAG-----GGACA-ACUAUUGGU
GGGCAGUUUGGGUGGGGCGCCACGCCUCUAA-CAAUCCAAC-AGAGGCCUCCA-AAGGUCAGCUCAGGAGGG
UCAGAAAUCCUCCGUA-GAGUA-UAAAGGCAA-AAGCUGGCUUGACA-GUGU---------------------
--------------UCCAGACAAUAAGGAACGCUGAUGCGAAA-GCAGGGCUUAGCGAACCACC--------
---UU-GUC--UCCCUC-UUC--------------------AU-GAGGGCAGGUGAUAAGAG-AG--AAGUU
ACCCCAGGG-AUAACUGAGUUGUCCUCGGCAAGAGUACACAUCGACCCGAGGGUUUGCUACUUCGAUGUCGU
CUGUUCCUAUCCUGGCGCUGCA-UAAGACGCCAAGGGUGGGGCUGUUCGCCCAUUAAAAGGGAUCCUGAGAU
GGGUUCACUACGUCGCGAGACAGUAGGGUUGCUUCUCCGU--------------------G-GGA-------
-------------------------------GUGC--UCGAUGCCUGAAGGGAAGG-AGCCUUUAGUACGAG
AGGAA-CGGGGGUUCGUGGCCUC-UGGU-U-UACCGGUUGUCC-GACA-GGGCAGUU-GCCGGGUAGCUACG
CCAUACGCGAAUAAAGGCUGAAAGCA-UC--UAAGCCUGAAGUCGCCCCUGA-A-A-A--GAGGCAUC-GU-

------------------------------------------------------------------------
------------------------------------------------------------------------
------------------------------------------------------------------------
------------------------------------------------------------------------
------------------------------------------------------------------------
------------------------------------------------------------------------
------------------------------------------------------------------------
------------------------------------------------------------------------
--------------------------------------------------------UAUC-AGG------UC
CCUCCUAAAAGAGGAGCU-UGA-UAGGAC-CGG-GAUGUAAG-UGC-U--------GAG-----CU------
--UUUG--------CGAGGCAUUAAGUCC--A-CGGU-UACUAAAA---GACCG------------------
------------------------------------------------------------------------
------------------------------------------------------------------------
------------------------------------------------------------------------
-------------------------------------------------
>LSUa_74969_FERRA/1-2900 CP015363.1, complete genome; RANGE 1819037:1821936
----------------------AUCAUA------------CAAACUGCUU-GGGGAAUG-GUUUUGGU-UUG
A-G---AACCGAUGAAGGACGUGC--CAAGCUGCGAUAUGCCUUGGGGAGGUGCAUGGAGCCAU------UG
AUC-CAA-GGAUUUCCGAAUGGGACU-U--------CCU-------GU------------------------
--U---------------------------------UA--AC-A-CUCC-----------------------
-----------G-----------AAA----------------------------------------------
------------------------G-G-AG-AGGGAACCCCGGGAAUUGAAACAUCUUAGUACCGG--GAGG
AAGAGAAAUCAA--UAGAGAUACCGUUA--GU-AA-AGGCGAUCGAAAGCGGU-A-GA-GGGCAAACCGAAU
CUUU--UCUGU--------AA------------A---GAA-AUAGAUAUGUGGAG--UA-------------
C-GG--AUC-UUU-C-----U------C------------------AAUGC--CUA-----------CU--U
U-------AGGAA--UUCU------AAA-AA-GUUG-------------------------------------
-------GAA-AG-CUUUA-------CCAGAGAGGGUGAUAGUCCCCUG---------GAA---G----UAA
C------UAAAGAGG-CUACGA--------------GGAA-AGAUA--UCCUGAGUACCAUGCGUCA---UU
AUUCGCGUGGGAA-UAUGGGUGGCAUG--AACAUCCAACCCUAAAUA--UUA-C-UCA-AGUCCGAUAGC-A
AAG-A-AGUACCGUGAGGGAAAA-CUGAAA-AGCAA-CCCGG-AAGGGC-U-GUA-AAAAGAGCCUGAAACC
A-AGCAGUGAAAA-CUU-UA-UAGG---GCACUAAA-GGAAU------------------------------
------------------------------------------------------------------------
------------------------------------------------------------------------
------------------------------------------------------------------------
------------------------------------------------------------------------
------------------------------------------------------------------------
------------------------------------------------------------GA-UC-------
------------------------------------------------------------------------
------------------------------------------------------------------------
--------------------------------------------------------------UCGA-AA---
------------------------------------------------------------------------
------------------------------------------------------------------------
------------------------------------------------------------------------
------------------------------------------------------------------------
------------------------------------------------------------------------

```
------------------------------------------------------------------------
------------------------------------------------------------------------
------------------------------------------------------------------------
------------------------------------------------------------------------
------------------------------------------------------------------------
------------------------------------------------------------------------
------------------------------------------------------------------------
----------------------GAG-GG-----------------ACCGG--U---GUUCUAUAGUCCGUGU
UGAAGAACGGGCCAGGGAGUUGU-G-AUGGAUGGCGA-GGCUAAUU--C----------UAAAA--------
--GA--AGCA--GCCGUAGCGAAAG-CGA-----------------UU-----------------------A
CUCG-CACAGC-------------------------------------------------------------
------------------------------------------------------------------------
--------------------AAU------------------------------------GGA-GGAGUA-AC
GUA------------------------------------AUAA--------------------------AU-
G-C-GU----------------------UUAGUCAUCUGUGGCA-GACCCGAAGCC-ACUCGAUCUAUACCU
GAGCAGGUUGAAGUUUAG-CGAAAG-CUAGAUGGAGGACCGA--ACCAGUUCUGAUGUGCAAAUCGU-UUGG
AUGACUUGGGUAUAGGGGUUAAAGGCUAAUCUAGAGUGG-CAAUAGCGGGUUCCCCCCAAUA-CUACCCGCA
GGUAGACCUUGG-UGGA-------------------------------------------------------
----------------------GA-----UG-UCUGGUGAGGUAGAGCGACCGAUU-GGG-UGGC-CAGCAG
U-A-GA-AA-UACU-G-CACCACCUUGUCAAACUCCGAACUUGCUA--------------------------
------------------------------------------------------------------------
------------------G-AA-U--CG-AAGA--UACCAGG-UGCUAGG-G-CGUA-G-GGA-UAAGCUUU
AC-G-CCCGUGAC-GGAAAC---AACCGAGACUAA-U-GUUAAGGUCUCUAAA---AU-UGG-CU-AA---G
-U-GCGA----UUUAAAGAGGUUAAUGACCGAAGACAGUGUGGAGGUUAGCUUAGAAGCAGCUA-UCCUUUA
AAGAGUGCGUAACAGCUCACUCACC-----GAGGUUAUUUGCCUCCAAGAU-GGAA-GAGGCUAA-AGCCAG
U----------UACCGAU-ACAUUAGACCACCGA--------------------------------------
------------------------------------------------------------------------
------------------------------------------------------------------------
--------------------------------AAGGU-GAUG-UGGUAGGGGGGC-GUGCUAU---------
-----------A-U-GGAUA-GAAGUUU--C-UUCGAGAGGA--G-GGAUGGACCG-U-AUA-GUAUCGCGG
AUCC-UGGUAA-AAGUA-GCAGAA--UAGAA-UGGUGA--A---------------AAUCCA-UUC-CGCCG
A-AAGGGC-AAAGGUUCC-UUGGCAAUG-UUCGUCAGCCGAGGGUUAGCCGA--UC-CUAAGGUCACCCCU-
AAC-AGGA---A-GUG--------------------------------------------------------
----------------------------------------AUC-GA-A-AGGGA-AGCUGGUU-AAUA-UUC
CAGCG----------------------------------CCUCA----GGUGUU---------U--------
------------------------------------------------------------------------
-------GC------------------------------------------------CAUA-U-AUAU-AC-
GCUUUGG-------------GAUAGA-GAGUGUAUG-CU-------UGUC-------------U-ACGUAU-
GGGU-A-UUU-AU-----A-A-------------------GU-----AA----------------AUGG-AG
A-G-UU--------------------GUAAU-GACGAGAAGU---------UUACGA----AGU-AA-UAU-
CU----G-----------------------------------------------------------------
-----UUCCC----GUAUGGGAAUU-U-UUU-C-A-AGUCCCGGAGCCU-AUGAAA-G-AUA-UAUAUGG--
--------------------------------------------------------GUCAAG-CC-UGAGUU
CGUACCAAGAACCGAC------------------------------ACAGGUGC-CCCUAGAU-GAAUAGUC
UCAGGC-GUUU-GGGA-AUAAUGGA-U-GUGAGGGAACUCGGC-CAAAUAGUCCCGUAUCUUAGG-UAGAAG
```

```
GGAUG-CCUA-U-CC--AGA----------------------------------------------------
------------------------------------------------------------------------
------------------------------------------------------------------------
------------------------------------------------------------------------
------------------------------------------------------------------------
------------------------------------------------------------------------
------------------------------------------------------------------------
------------------------------------------------------------------------
------------------------------------------------------------------------
------------------------------------------------------------------------
------------------------------------------------------------------------
------------------------------------------------------------------------
------------------------------------------------------------------------
------------------------------------------------------------------------
------------------------------------------------------------------AAUAGG
AUA--GGUCGCAGUGGCAAGGGGACCCCGA-----CUGUUUACCAAAAACACAGACCGCUGCUAGUCCG---
-UA-A-------------GGAUGUGU-AUAGCGGCCGA-AACCUGCCCAGUGUUG-G-UACC-UGAAAGCUC
UU-------------UUCA-------------AGGAGAAGAAGGGC--CAAUAAAC-GGCGGGGGUAACUAU
GACCCUCUCAAGGUAGCGUAAUACCUAGCCGCU-UAAUUGGCGGCUCGCAUGAA-GGUUCAACGAGA--GUC
CCACUGUCCCCACA----UUC-AGCCCCGUGAAAUU--AAU---GUACUGGUGCACAAUCCAGUCUCUCCCA
CAUGAAAGCGAAGU-CCCCGUGGAGC-UUUACU--GCAGCCU-GUAGCUGUGACGCGAUUUUUAAUGCGUAG
GGU-AGGAAGGAGCCGUCG----------AUGUUCU-GGU----------------------------CGUG
--------------------------------------GC-CAGAAUGGAG-G--C-AA-CGAUGAAACA-C
UUCC-CUUUAAAAGUUGCGUCACUAACCU-U----------GA-AGA-------------------------
------------------------------------------------------------------------
------------------------------------------------------------------------
------------------------------------------------------------------------
------------------------------------------------------------------------
------------------------------------------------------------------------
-------------------------------------------------AAG-----GGACA-ACUAUUGGU
AGGCAGUUUGGGUGGGGCGCCACGCCUCUAA-GAAGAAAAC-AGAGGUCUCCA-AUGGUUAGCUCAGAAGGG
UCAGAAAUCCUUCAAA-GAGUA-UAAGGGUAA-AAGCUAGCUUGACU-GUGU--------------------
--------------UCCAGCCAAUAGGGAACGCAGAUGCGAAA-GCAGGGCCUAACGAACCACC--------
---UA-GU---UCCCUC-UUU--------------------AU-GAGGGCAGGUGAUAAGAG-AG--AAGUU
ACCCCAGGG-AUAACUGAGUUGUCCUCGGUAAGAGUACACAUCGACCCGAGGGUUUGCUACUUCGAUGUCGU
CUGUUCCUAUCCUGGUGCUGCA-UAAGACACCAAGGGUGGGGCUGUUCGCCCAUUAAAAGGGAUCCUGAGAU
GGGUUCACUACGUCGCGAGACAGUAGGGUUGCUUCUCUGU--------------------G-GGA-------
-------------------------------GUGU--UCAAUGUCUGAAGGGAAGG-AGCCUUUAGUACGAG
AGGAA-CGGGGGUUCGUGACCUC-UAGU-U-UAUCGGUUGUCC-GACA-GGGCAGUU-GCCGAGUAGCCACG
UCAUACGCGAAUAAAAGCUGAAAGCA-UC--UAAGCUUGAAGUCGCCCCUGA-A-A-A--UAGACAUU-GU-
------------------------------------------------------------------------
------------------------------------------------------------------------
------------------------------------------------------------------------
------------------------------------------------------------------------
------------------------------------------------------------------------
```

```
------------------------------------------------------------------------
------------------------------------------------------------------------
------------------------------------------------------------------------
--------------------------------------------------------U-UG-AGA------UC
ACUCUUAAAAGAAGAGGU-UGA-UAGAGU-CGG-GAUAUAAG-UAC-C--------AAG-----CU------
--UUUG--------CGAGGUAUUAAGUCC--A-CGAC-UACUAAUU---GAUCGAAAA-UCUACA-------
------------------------------------------------------------------------
------------------------------------------------------------------------
------------------------------------------------------------------------
------------------------------------A------------
>LSUa_273116_THEVO/1-2909
---------------------------GCUC----UG--CUAAUCUGCCU-AGAGGAUG-G-CUUGGU-UCG
G-G---CGCCGAAGAAGGACGUGC--CAAGCUGCGAUAUGCCUUGGGGAGGCGCAUGGAGCCAU------AG
AUC-CAA-GGAUUUCCGAAUGGGACU-U--------CCU-------GC----C-----------------GU
A-A--------------------------------GG----C-A-CUCC-----------------------
-----------G-----------AAA----------------------------------------------
------------------------G-G-AG-AGGGAACCCGGGGAAUUGAAACAUCUUAGUACCCG--GAGG
AAAAGAAAUCAA--UCGAGAUACCGUUA--GU-AA-AGGCGAUCGAAAGCGGU-A-AA-AGGCAAACCGAAU
AGCC-CUUCGA--------AA------------G---AAG-GGAAAGAUGUGGAG--UU-------------
U-GG--UCU-UUC-U-----C------U------------------AAUGC--CUC-----------CU--G
A-------AGCGA--GAUG------AA-UCU-UCUG------------------------------------
-------GAA-AG-AAGAG-------CCAAAGAAGGUGAUAGCCCUGUA---------AUC---G----AAG
C------UUCAGAAG-CUAUAA--------------GGGG-AAGUA--ACCAGAGUACCAUGCGUCG---UU
UUUCGCGUGGGAA-UGUGGGUGGCACU--AACAUCCAACCUUAAAUA--CGU-C-CCG-AAUCCGAUAGC-G
AAC-UAAGUACCGUGAGGGAAAG-CUGAAA-AGUAA-CCCGA-AAGGGU-G-GUG-AAAAGAGCCUGAAACU
A-GGCAGAGAUAA-CCU-UG-UAGG---GCAGUAAA-GAGGUG-----------------------------
------------------------------------------------------------------------
------------------------------------------------------------------------
------------------------------------------------------------------------
------------------------------------------------------------------------
------------------------------------------------------------------------
-------------------------------------------------------------A-UACCGUU--
------------------------------------------------------------------------
------------------------------------------------------------------------
-----------------------------------------------------------------A-AC---
------------------------------------------------------------------------
------------------------------------------------------------------------
------------------------------------------------------------------------
------------------------------------------------------------------------
------------------------------------------------------------------------
------------------------------------------------------------------------
------------------------------------------------------------------------
------------------------------------------------------------------------
------------------------------------------------------------------------
------------------------------------------------------------------------
```

```
------------------------------------------------------------------------
------------------------------------------------------------------------
-------------------AACGGU-GG-----------------AUCGC--U---GUCCUAUUGUCCGUGU
UGAAGAACGGGCCAGGGAGUUCU-G-ACGGGUGGCAA-GGUUAAUC--C----------UUAAA--------
--GG--AGAA--GCCGUAGCGAAAG-CGA-----------------UU-----------------------A
CCCG-CACAGC-------------------------------------------------------------
------------------------------------------------------------------------
--------------------AAU------------------------------------GGA-GGGGUA-GC
GUG------------------------------------GUAAA--------------------------C-
G-C-GU----------------------UUAGUCACUCGUGAGA-GACCCGAAGCC-GGUCGAUCUACACCU
GAGUAGGUUGAAGCUCAG-UGAAAG-CUGGGUGGAGGACCGA--ACCUAUUCUGAUGUGCAAAUCGU-UUGG
AUGACUUGGGUGUAGGGGUUAAAGGCCAAUCUAGGCCGG-CAAUAGCGGGUUCCCCCCGAUA-CUACCCGCA
GGUAGACCUCGA-CGGA-------------------------------------------------------
----------------------GA-----UU-CUCGACGAGGUAGAGCGACCGAUU-GGU-UGGU-UAGCAG
U-C-GA-AA-GGCU-G-CGCCGACUUGUCAAACUCCGAACUUGCCG--------------------------
------------------------------------------------------------------------
------------------A-GA-U--CG-UAGA--UGUCGGG-UGCUAGG-G-GGCA-G-GGA-UAAGCUUU
GC-U-UCCGUGAU-GGAAAC---AACCAAGACGAG-G-GUUAAGGUCCCUAAAU--UC-UGG-UU-AA---G
-U-GC------ACUAAAUAGGUUUGUGGCCAAAGACAGCGGGGAGGUAGGCUUAGAAGCAGCCA-CCCUUUA
AAGAGUGCGUAACAGCUCACUCGCC-----GAGGUCACAUGCCUAGAAGAU-GGAA-GGGGCUAA-AACCAG
A----------UACCGAG-ACCUUCGAGCACCGA--------------------------------------
------------------------------------------------------------------------
------------------------------------------------------------------------
--------------------------------AAGGU-GAUC-UGGUAGGGGGGC-GUGCCAU---------
-----------G-U-GGGCA-GAAGUUU--C-UUCGAGAGGA--G-GAAUGGACCG-C-AUG-GUAUCGCGG
AUCC-UGGUAA-AAGUA-GCAGAG--AAGAA-CCGUGA--G---------------AAUCGG-UUC-CGCCG
A-AAGGGCUAAGGGUUCC-UUGGCAAUG-UUCGUCAGCCGAGGGUUAGUCGA--UC-CUAAGGUCAUACCU-
AAC-AGGA---U-AUG--------------------------------------------------------
----------------------------------------ACC-GA-A-AGGGA-AGCCGGUU-AAUA-UUC
CGGCA----------------------------------CACUG----AGCG--------------------
------------------------------------------------------------------------
----------------U--------UU-UG-----CC---------------------AUG-U-GUGA-GAA
GGUUCGA-------------GGUAGG-AGCGGUACG-GG-------UGCC-------------A-ACGUAU-
UUCU-G-CCU-AU-----A-A-------------------GC-----GG----------------AUGG-AG
A-G-UC--------------------GUAAU-GACGAGAAGU---------UCGUGA----AAG-GG-UGG-
AA----G-----------------------------------------------------------------
-----CUCCC----GUUUGGGAGUC-G-CUU-C-G-AUCCUUGGAUCCC-AUGAAA-A-UCA-UACAUGG--
--------------------------------------------------------GUCAAA-CU-CAGUAU
CGUACCAAGAACCGAC------------------------------ACAGGUGC-CCCUAGGU-GAGAAGCC
UAAGGC-GUUU-UGGG-AUAAUGGA-C-GCGAGGGAAAUCGGC-AAAAUAGCUCCGUAUCUUCGG-UAUAAG
GAGUG-CCUA-U-UC--CGC----------------------------------------------------
------------------------------------------------------------------------
------------------------------------------------------------------------
------------------------------------------------------------------------
------------------------------------------------------------------------
```

```
------------------------------------------------------------------------
------------------------------------------------------------------------
------------------------------------------------------------------------
------------------------------------------------------------------------
------------------------------------------------------------------------
------------------------------------------------------------------------
------------------------------------------------------------------------
------------------------------------------------------------------------
------------------------------------------------------------------------
------------------------------------------------------------------GAUGGA
AUA--GGUCGCAGUGGCGAAGGGACUCCGA-----CUGUUUACCACAAACACAGAUCGCUGCUAGUCCG---
-UA-A-------------GGAUGUGU-AUAGCGGUUGA-AACCUGCCCAGUGCUG-G-UACC-UGAAAGCCC
CG-------------CACA-------------AGGGGAAGAAGGGC--CAGUAAAC-GGCGGGGGUAACUAU
GACCCUCUUAAGGUAGCGUAAUACCUAGCCGCU-UAAUUGGCGGCUUGCAUGAA-GGUUCAACGUGG--GUC
CCACUGUCCCCGCG----UUC-AGCCCAGUGAAAUU--GAU---GUACUGGUGCACAAUCCAGUCUCUCCCA
CGUGAAAGCGAAGU-CCCCGUGGAGC-UUUACU--GCAGCCU-GUAGCUGUGAUGCGAUCCUGAAUGCGUAG
CGU-AGGAAGGAGCCGUCG----------AAGCUUC-GGU----------------------------UUUG
--------------------------------------GC-CGAAGUGGAG-G--C-GC-CAAUGAAACA-C
UUCC-CUUUCGGGAUUGCGUCACUAACCU-C----------GU-U-A-------------------------
------------------------------------------------------------------------
------------------------------------------------------------------------
------------------------------------------------------------------------
------------------------------------------------------------------------
------------------------------------------------------------------------
-------------------------------------------------GAG-----GGACA-ACUAUUGGU
GGGCAGUUUGGGUGGGGCGCCACGCCCCUAA-CAAUGUAAC-AGGGGCCCCCA-AAGGUCAGCUCAGGAGGG
UCAGAAAUCCUCCGUA-GAGUG-UAAAAGCAA-AAGCUGGCUUGACU-GUGU--------------------
--------------UGCAGACAACUAGCAACGCAGAUGCGAAA-GCAGGGUUUAGCGAACCACC--------
---CA-GU---UCCUCC-UUA--------------------GU-GGGGGCGGGUGAUAAGAG-AG--AAGUU
ACCCCAGGG-AUAACUGAGUCGUCCUCGGCAAGAGUACACAUCGACCCGAGGGUUUGCUACUUCGAUGUCGU
CUGUUCCUAUCCUGGUGCUGCA-UAAGGUGCCAAGGGUGGGGCUGUUCGCCCAUUAAAAGGGAUCCUGAGAU
GGGUUCACUACGUCGCGAGACAGUAGGGUUGCUUCUCCGU--------------------G-GGA-------
-------------------------------GUGC--UCGAUGUCUGAGGGGAAGG-GGCCUUUAGUACGAG
AGGAA-CAAGGGCUCGUGACCUC-UAGU-U-UACCGGUUGUCC-GGCA-GGGCAU-C-GCCGGGUAGCCACG
UCAUACGCGGAUAAGAGCUGAAGGCA-UC--UAAGCUCGAAGCCGCCCCCGA-A-AAU--AGACAUCG-U--
--CAUU------------------------------------------------------------------
------------------------------------------------------------------------
------------------------------------------------------------------------
------------------------------------------------------------------------
------------------------------------------------------------------------
------------------------------------------------------------------------
------------------------------------------------------------------------
------------------------------------------------------------------------
-------------------------------------------------------------AGG------CC
GCCUCUAAAAGAGAGGUU-UGA-UAGGGC-CGG-GAUGUAAG-GAU-C--------GAG-----CU------
```

```
--UUUG--------CGAGAUUUUAAGUCC--A-CGGC-UACUAAAG---GCCGAAAGC-ACAAUC-------
------------------------------------------------------------------------
------------------------------------------------------------------------
------------------------------------------------------------------------
-------------------------------------------------
>LSUa_414004_CENSY/1-2987 DP000238.1, complete genome; RANGE 1200795:119779
------------------------CCGGUGU----AA--AGACGGCAAUA-GGAGGAUU-G-CUAGGC-UUG
A-G---GAGAGAAGAAGGACGUGG--CAAGCUGCGAUAAGCUCGGGGUAGGUGCACGCACCCGU------UG
AUC-CCG-AGAUUUCCGAAUAGGAAU-C--------CU--------------------------------GU
A-U----------------------------------------A-CUCC-G---------------------
--C--------G-----------CAA----------------------------------------------
--G--------------------CG-G-AG-GUCGAACCGUGGGAAUUGAAGCAUCUUAGUACCAC--GAGG
AAGAGAAAUCAA--UAGAGAUUUCCCAA--GU-AG-AGGCGAUCGAAAAGGAA-A-GA-GCCCAAACUGAAU
CUGC-CGCGGU--------AA------------C---GCG-GCCGAGAUGUGGUG--UUU------------
U-GG--UGC-GGC-GC------------------------------AAUGG--CCC-----------CA--G
G-------AAAUA--GUUG------AAGUGU-UCUG------------------------------------
-------AAA-UG-UUCCG-------GCUUAGAGGGUGAUACCCCCUA----------GAC---G----AAG
U------GGAAUGGG-GC------------------AGGG-CCAGC--ACCCGAGUAGUUGUCCUUG---GC
AGUGGGCAGCGAA-GAUUGGUGAAAGU--AGCAUCAAAGGCUAAAUA--UCU-C-UCA-AGACCGAUAGU-G
UAC-U-AGUACCGUGAGGGAAAG-CUGAAA-AGUAC-CCCGG-AAGGGG-G-GUG-AAAAGUGCAUGAAACC
U-AUUGCUUACAG-ACG-UG-CAUG---GCAUGAAA-GACGAU-----------------------------
------------------------------------------------------------------------
------------------------------------------------------------------------
------------------------------------------------------------------------
------------------------------------------------------------------------
------------------------------------------------------------------------
-----------------------------------------------------------UUUUUCGGACCG-
---GA-----------GG-U-------------------U------CCA-----------------GCA---
------------------------------------------------------------------------
--------------------------------------------------------------A--UGGA---
------------------------------------------------------------------------
-------------------------------------------------G-CU------GA--------AAG
UCCGG-------------------------------------------------------------------
------------------------------------------------------------------------
------------------------------------------------------------------------
------------------------------------------------------------------------
------------------------------------------------A-----------------------
------------------------------------------------------------------------
------------------------------------------------------------------------
------------------------------------------------------------------------
------------------------------------------------------------------------
------------------------------------------------------------------------
-------------------GCUCAUGUC-----------------GUCAG--U---GUCAUGCGUUCCGUCU
CGAAACACGGGCCAGGGAGCUUG-C-UGUCAUGGCGA-GGUUAAGC-------------CUAGA-------A
A-GG--CGUA--GCCGAAGGGAAAC-CGA-----------------AU-----------------------U
```

```
CUCG-CAGCAU-------------------------------------------------------------
------------------------------------------------------------------------
---------------------UG------------------------------------UGA-GGAGAA-GC
GUGU-----------------------------------GAAA--------------------------GU-
G-C-GU----------------------UAAGUCAUGGCGGUAA-GGCUAGAAGCC-AGUCGAUCUAUUCCU
GAGCAAGAUGAAGGUGGG-CGAAAG-CCCACUGGAGGUCUGC--AGCAGUUCUGACGUGCAACUCGU-UUGU
GUGACUUGGGAAUAGGGGUCAAAAACCAAUCUAGACUGG-CGCUUGCUAGUUCCAACCGAAG-CGUCUCGCA
GGGCGUGCUAUG-UGGA-------------------------------------------------------
----------------------GA-----UG-GUGUUUCCGGUAGAGC-ACUGAUA-GGG-UGGC-GCGGGG
A-A-GA-GA-UUCC-U-CACCAUCCAUUCAAACUCCGAACGGAUAC--------------------------
------------------------------------------------------------------------
------------------A-CA-U--CG-CAGA--AGCAUGG-ACACGGGUU-CGUGUG-ACG-UAAGGUUC
AC-G-ACCGAGAG-GGGUUU---AACCCAGACUGG-A-GUUAAGGUCCCCAAAU--UU-CUG-CU-AA---G
-U-GUCAAG--CCAAAGUGCGUGUCAUCGCCAAGACAGCAGGGAGGUAAGCUCAGAAGUAGCUA-UCCUUCA
AAAAGUGUGUAACAACUUACCUGCC-----GAGCGAGGGCGCCACAAAAAC-GGAC-GGGGCUAA-AGCAGA
G----------UACCGAU-ACUCCAGACGUGCAC--C-----------------------------------
------------------------------------------------------------------------
-----GUGAUA-------------------------------------------------------------
--------------------------------GGUGC-ACCG-UGGUAGGUUGGC-GUAGUGA---------
-----------U-U-GGGAC-GAAGCAG--G-GCGGUGACGU--C-CUGUGGACCG-A-UUU-CUAUUGCAG
AUCC-UGGUGG-CAGUA-ACAGCA--UAGUG-CGGUGA--G---------------AAUCCG-CAC-CACCG
U-AAGCGC-AAGGGUUUC-CAGGCAAUGCGUCGUCAGCCUGGAGUUAGUCGA--UC-CUAAGGACUGCCUC-
AAC--AGA---G-UAG--------------------------------------------------------
----------------------------------------UCC-GA-A-UGGGA-AACCAGUU-AAUA-UUC
UGGUA----------------------------------CCUUG----CAGG--------------------
------------------------------------------------------------------------
-------------GAGC--------GUUUGA--------------------------ACCG-C-AUGG-UUC
GCUUCCG-------------GAUAGG-GGGAACAUG-AC-------CGUC-------------G-UCGUGU-
CGAA-CCGUU-CG-----A-G-------------------GG-----CC----------------UGGG-AG
U-G-UC--------------------GUAAU-GACGAGAACA---------GGCCCG----AGG-CG-GGA-
GUG--GC-----------------------------------------------------------------
-----UUUGC---GUUAGCAGAGUU-U-UCC-C-G-AUUCCAGGAGACC-AUGAAA-G-ACC-GUGCGGCUA
--------------------------------------------------------GCAAAC-UG-CAAGUU
CGUACCCAGAUCCGAC------------------------------ACAGGUGC-GC-UGGUU-UAGUAAAC
UAAGGU-GCUA-CGGGUAUACCGUAUG-GCGAGGGAAUUCGGC-AAAUUAGUCCUGUAGCUUAGG-UAUAAG
GGAUG-CCUG-C-AG--CUGU--------------------------CAUGA--------------------
------------------------------------------------------------------------
------------------------------------------------------------------------
------------------------------------------------------------------------
------------------------------------------------------------------------
------------------------------------------------------------------------
------------------------------------------------------------------------
------------------------------------------------------------------------
------------------------------------------------------------------------
------------------------------------------------------------------------
```

```
------------------------------------------------------------------------
------------------------------------------------------------------------
------------------------------------------------------------------------
------------------------------------------------------------------------
----------------------------------------------------------GGAAUG--GCGGCA
GCA--GGUUGCAGUGACAAGGGGGUUUCGA-----CUGUUUAAUAAAAACACAGGCGACUGCUAGCCCG---
-AA-A-------------GGGUAUGU-AUAGUCGCUGA-AUCCUGGCCGGUGCCG-G-UAUC-UAAAACUUG
GG-------------UUCA-------------ACCGAGCUAAGGAC--CGGUUAAC-GCCGGGAGUAACUCU
GACUCUCUUAAGGUAGCCAAAUGCCUUGUCGGG-UAAGUUCCGACGUGCAUGAAUGGAACAACGAGA--GCC
CCACUGUCCCCGCC----UAC-AACCCGGUGAAGCCACAUA---AGGUGGACGAACAGUCCAUCAUCUUCCA
UCGGGGAGAGAAGA-CCCCGUGGAGU-UUUACU--GCAGCUU-GUGCUUGUGGUAUAGCAGGGAGUGCAUAG
CGU-AUCUGGGAGUCAUUUCCUGGCG-----------UUC----------------------------UUUG
--------------------------------------GG-ACGUCAGGGG-A--C-GC-CGAUGUAACA-C
CAGA-CAUUUCCUGCAGUGCCACUUACCC-G----------GU-G-A-------------------------
------------------------------------------------------------------------
------------------------------------------------------------------------
------------------------------------------------------------------------
------------------------------------------------------------------------
-------------------------------------------------------A----GA----------
------------------------------------------------CCGG-----GGACA-UGUGCAGGU
GGGCAGUUCGGCUGGGGCGGCACCCCUUUGA-AAAGAUAUC-AAAGGGGCCCA-AAGUUUAGUUCAAGCGGG
UCAGAAAUCCGCUGAA-GAGGG-CAAAGCCAA-AAACUAGACUGAAU-GGAU--------------------
--------------UUCCAAACGCACGGAAUUCAUAGACGAAA-GUCUGGCUUAGCGAUCCUUC--------
---GU-G-----UGCCCACUA--------------------UU-GGGGCCCGGAGGUGACAG-AA--AAAUU
ACCCCAGGG-AUAACAGGCUCGUCGCGGGCGAGAGCUCCUAUCGACCCCGCGGUUUGGUACCUCGAUGUCGG
CUUUUCCUAUCCUGGUCCUGCA-GCAGGGGCCAAGGGUGAGGCUGCUCGCCUAUUAAAAGGGAACAUGAGCU
GGGUUUAGACCGUCGUGAGACAGGUCGGCCUCUGCCUGAU--------------------G-GAA-------
-------------------------------GCGC---GAAUAUCUGAGGGGAAGU-CGCUCUUAGUACGAG
AGGAA-CAGAGAGGCGUGGCCAC-UGGU-C-UACCGGUUGUCC-GACA-GGGCAG---GCCGGGCAGCUAAG
CCAUC-UAGGCUAAGUGCUGAAAGCA-UC--UAAGCACGAAUCCUAUCCCGA-G-ACA--AGAUAUUC-CU-
--UCUUCGG---------------------------------------------------------------
------------------------------------------------------------------------
------------------------------------------------------------------------
------------------------------------------------------------------------
------------------------------------------------------------------------
------------------------------------------------------------------------
------------------------------------------------------------------------
------------------------------------------------------------------------
---------------------------------------------------------AUGAAGG------UU
CGCAGAAGAAGACUGCGU-UGA-UAGGAA-GGU-GGUGUAUC-CCA-C--------AAG-------------
CUUCGG--------CGAGUGGUG-AGCCA--G-CCUU-UACUAAUC---GACCAAAAC-ACCGGUUCAG---
------------------------------------------------------------------------
------------------------------------------------------------------------
------------------------------------------------------------------------
-------------------------------------------------
```

```
>LSUa_436308_NITMS/1-2984 CP000866.1, complete genome; RANGE 896089:893106
-----------------------GCCGGUGU----AA--AGACGGUAAUA-GGUGGAUU-G-CUAGGC-UUG
A-G---GAGAGAUGAAGGACGUGG--CAAGCUGCGAUAAGCUCGGGGUAGGCGCACGCAUCCUA------UG
AUC-CCG-AGAUGUCCGAAUGAGAAU-CUUACACUUUU--------------------------------GU
G-U----------------------------------------A-UUCC-AGUACAC---------------
--U--------A-----------GUA----------------------------------------------
--U--------------------UG-G-AA-GUCGAACCGUGGGAAUUGAAGCAUCUUAGUACCAC--GUGG
AAAAGAAAUCAA--UUGAGAUUUCCCAA--GU-AG-AGGCGAUCGAAAAGGAA-A-UA-GCCCAAACUGAAU
CCUU-CG------------------------------------GGAGAUGUGGUG--UUU------------
U-GG--UAU-GAU-AA------------------------------UAUGGAAUCU-----------UG--A
A-------ACACA--GCUG------AAAUGU-UCUG------------------------------------
-------AAA-AG-UACUG-------ACAUAGAGGGUGAUAUCCCCGUA---------AGC---G----AAA
U------GGGAAGAG-UC------------------CUAU-UCA-U--ACCCGAGUAGUUGUCCUUG---GU
AGUGGGCAAUGAA-UACAGGUGAAAGU--AGCAUCUGAGGCUAAAUA--UCU-C-UCA-AGACCGAUAGU-G
GAC-U-AGUACCGUGAGGGAAAG-CUGAAA-AGUAC-CCCGG-AAGGGG-G-GCG-AAAAGUGCAUGAAACC
U-AUUACUUACAG-ACG-UG-CAUG---GCAUGGAA-GGUGAU-----------------------------
------------------------------------------------------------------------
------------------------------------------------------------------------
------------------------------------------------------------------------
------------------------------------------------------------------------
------------------------------------------------------------------------
-----------------------------------------------------------UUUUUCAUUUUG-
---GA-----------GA-U-------------------U------CUA-----------------GCA---
------------------------------------------------------------------------
--------------------------------------------------------------A--UAGA---
------------------------------------------------------------------------
-------------------------------------------------G-UU------GA--------AGA
UUUGA-------------------------------------------------------------------
------------------------------------------------------------------------
------------------------------------------------------------------------
------------------------------------------------------------------------
------------------------------------------------U-----------------------
------------------------------------------------------------------------
------------------------------------------------------------------------
------------------------------------------------------------------------
------------------------------------------------------------------------
------------------------------------------------------------------------
-------------------GUUCAAGUC-----------------AUCAG--U---GUCAUGCGUUCCGUCU
CGAAACACGGGCCAGGGAGCUAA-C-UGUCAUGGCAA-GCUUAAAU-------------CUUUA-------A
A-GA--UGUA--GGCGAAGGGAAAC-CGA-----------------GU-----------------------U
CUCG-CAGCAU-------------------------------------------------------------
------------------------------------------------------------------------
---------------------UG------------------------------------UGA-GGAGAA-GC
GUGU-----------------------------------GAAA--------------------------GU-
G-C-GU----------------------AGAGUCAUGGCGGCUA-GGCUAGAAGCC-AGUCGAUCUAUUCCU
```

```
GGGCUAGAUGAAGGCGGG-CGAAAG-CCCGCUGGAGGUCUUA--AGACGUUCUGACGUGCAAAUCGU-UGUU
GUGACCUGGGAAUAGGGGUCAAAAACCAAUCUAGACUGG-CGCUCGCUAGUUCCAACCGAAG-CGUCUCGCA
GGGCGUGCUUUG-UGGA-------------------------------------------------------
----------------------GA-----UA-GUGUUUCCGGUAGAGC-ACUGAUA-GAG-CGGC-GCGGGG
G-A-GA-AA-UCCC-U-CACCGCUCAUUCAAACUCCGAACGGAUAU--------------------------
------------------------------------------------------------------------
------------------A-CA-U--CG-UAGA--AGCAAGG-ACACGGGUU-CAUGUG-GCG-UAAGGCUC
AU-G-ACCGAAAG-GGGUUU---AACCCAGACUGA-A-GUUAAGGUCCCCAAAU--UU-UUG-CU-AA---G
-U-GUCAAG--CUAAAGAGCGUGUUCUCGCCAAGACAGCAGGGAGGUAAGCUCAGAAGCAGCUA-UCCUUUA
AAAAGUGUGUAACAACUUACCUGCC-----GAGGGGGGAUGCCUCAAAAAC-GGAC-GGGGCUAA-AGCAAA
G----------UACCGAU-ACUUUAGAUAAACAU--C-----------------------------------
------------------------------------------------------------------------
-------AAUG-------------------------------------------------------------
--------------------------------GAUGU-UUCG-UGGUAGGUUGGC-GUAGUGA---------
-----------U-U-GGGUC-GAAGCAG--G-GCGGUAACGU--C-CUGUGGACCG-G-UUU-CUAUUGCAG
AUCC-UGGCGG-CAGUA-ACAGCA--UAGUA-UGGUGA--G---------------AAUCCA-UAC-CACCG
U-AAGCGC-AAGGGUUUC-CAGGCAAUGCGUUAUCAGCCUGGAGGUAGUCGA--UC-CUAAGAUGUACCUC-
AAC--AGA---G-UAC--------------------------------------------------------
----------------------------------------ACC-GA-A-UGGGA-AACUAGUU-AAUA-UUC
UAGUA----------------------------------CCUUG----CAUG--------------------
------------------------------------------------------------------------
-------------GAGC--------GUUUGA--------------------------GCUA-U-AUGG-UUU
GCUUCCG-------------GAUAGG-GGUUGUACA-AU-------CGUC-------------G-UUGUAU-
CUAA-CUAUU-CG-----A-G-------------------AA-----CU----------------GGGG-AG
U-G-UC--------------------GUAAU-GACGAGAACC---------AGUUCG----AGG-UA-GGA-
AUG--GC-----------------------------------------------------------------
-----UCCGC---GUUAGCGGAGUU-U-UCU-C-G-AUUCCAGGAGACC-AUGAAA-G-ACU-GUGUAGUUA
--------------------------------------------------------GCAAAG-UG-CAAGCU
CGUACCCAGAACCGAC------------------------------ACAGGUGC-GC-UGGCU-UAGUAAGC
UAAGGU-GCUA-CGGGUAUACCGUAUG-GCGAGGGAAUUCGGC-AAAUUAGUCCUGUAGCUUAGG-UAUAAG
GGAUG-CCUG-C-GA--UCUU--------------------------CAUGA--------------------
------------------------------------------------------------------------
------------------------------------------------------------------------
------------------------------------------------------------------------
------------------------------------------------------------------------
------------------------------------------------------------------------
------------------------------------------------------------------------
------------------------------------------------------------------------
------------------------------------------------------------------------
------------------------------------------------------------------------
------------------------------------------------------------------------
------------------------------------------------------------------------
------------------------------------------------------------------------
------------------------------------------------------------------------
----------------------------------------------------------GGAAUG--GGGACC
```

```
GCA--GGUUGCAAUGACAAGGGGGUUUCGA-----CUGUUUAAUAAAAACACAGGCGACUGCUAGUCCG---
-AA-A-------------GGAUUUGU-AUAGUCGCUGA-AUCCUGGCCGGUGCCG-G-UACC-UAAAACUUG
GG-------------UUCA-------------ACCGAGCUAAGGGC--CGGUUAAC-GCCGGGAGUAACUCU
GACUCUCUUAAGGUAGCCAAAUGCCUUGUCGGG-UAAGUUCCGACGCGCAUGAAUGGAACAACGAGA--GCC
CCACUGUCCCCGCC----UAC-AACCCGGUGAAGCCACAUA---AGGUGGACGAACAGUCCAUCGCCUUCCA
UCGGGGAGAGAAGA-CCCCGUGGAGU-UUUACU--GCAGCUU-GUGCUUGUGGUAUAGUAAGAAUUGCACAG
GGU-AUCUGGGAGCCA-UGCUUAACA-----------CUC----------------------------UCCG
--------------------------------------GG-GUGUUGGGAA-G--C-AA-CGAUGUAACA-C
CAGA-CUCUUUUUGCAGUACCACUUAUCC-A----------GU-G-A-------------------------
------------------------------------------------------------------------
------------------------------------------------------------------------
------------------------------------------------------------------------
------------------------------------------------------------------------
-------------------------------------------------------A----GA----------
------------------------------------------------CCGG-----AGACA-CGUGCAGGU
GGGCAGUUCGGCUGGGGCGGCACCCCUUUGA-AAAGAUAUC-AAAGGGGCCCA-AAGUUGAGUUCAAACGGG
GCAGAAAUCCGUUGAA-GAGGG-CAAAGCCAA-AAACUCGACUGAAU-GGAU--------------------
--------------UUCCAAACGCACGGAAUUCAUAGACGAAA-GUCUGGCUUAGCGAUCCUUC--------
---GU-G-----UGCCCACUA--------------------UU-GGGGCCCGGAGGUGACAG-AA--AAAUU
ACCCCAGGG-AUAACAGGCUCGUCGCGGGCGAGAGCUCCUAUCGACCCCGCGGUUUGGUACCUCGAUGUCGG
CUUUUCCUAUCCUGGUCCUGCA-GCAGGGGCCAAGGGUGAGGCUGCUCGCCUAUUAAAAGGGAACAUGAGCU
GGGUUUAGACCGUCGUGAGACAGGUCGGUCUCUGCCUGAU--------------------G-GAA-------
-------------------------------GCGU---GAAUAACUGAGGGGAAGU-UGCUCCGAGUACGAG
AGGAA-CAGAGCAGCGUGGCCAC-UGGU-U-UACCGGUUGUCU-GACA-AGGCAG---GCCGGGCAGCUAAG
CCGUU-UAGGCUAAGUGCUGAAAGCA-UC--UAAGCACGAAUCCUAUCCCGA-G-ACA--AGUUAUUC-UU-
--UCGUAAG---------------------------------------------------------------
------------------------------------------------------------------------
------------------------------------------------------------------------
------------------------------------------------------------------------
------------------------------------------------------------------------
------------------------------------------------------------------------
------------------------------------------------------------------------
------------------------------------------------------------------------
---------------------------------------------------------AUGAAGG------UC
GGCAGCAGAAGACUGCGU-UGA-UAGGAA-GGU-GGUGUAGA-UCA-C--------AAG-------------
CUUCGG--------CGAGUGGUG-AGCCA--G-CCUU-UACUAAUA---GACCAACAC-AUCGGU-------
------------------------------------------------------------------------
------------------------------------------------------------------------
------------------------------------------------------------------------
-------------------------------------------------
>LSUASG_NODE_665_length_1073_cov_95.176413_g493_i0_REV/1-1073
------------------------------------------------------------------------
------------------------------------------------------------------------
------------------------------------------------------------------------
------------------------------------------------------------------------
```

```
------------------------------------------------------------------------
------------------------------------------------------------------------
AAAAGAAAACAA--UAGUGAUUCCGUGA--GU-AA-CGGCGAGCGAAAGCGGA-U-CA-GAUCAAACCGAAU
CCCU-AGGGA---------AA------------C---CUA-GUGGAAAUGUGGUG--UGU------------
G-GA--CCU-CGU-ACGAC----------------------------------CUC-----------CC--U
A-------GAAAA--UAUG------AAUACU-CCUG------------------------------------
-------GAA-AG-UUGAG-------CCAGAGAGCGUGAUAGCCGCGUA---------AUA---G----UAA
U------CUAGAAGG-UUGC----------------UGCG-AGAGU--UCCCGAGUAGUGCGGGAUG---GU
UUUCUCGUACGAA-UGUACCAGAUACA--CGCUGGUAAAUCUAAAUA--UAU-C-CCA-AGACCGAUAGU-G
AAA-U-AGUAGCGUGAGCGAAAG-AUGAAA-AGUAA-CCCGG-AAGGGC-C-GUG-AAAAGUGCUUGAAACC
A-AAUAGCUUAGU-AAG-GA-UGCA---GCCUGAAA-GGAUAG-----------------------------
------------------------------------------------------------------------
------------------------------------------------------------------------
------------------------------------------------------------------------
------------------------------------------------------------------------
------------------------------------------------------------------------
------------------------------------------------------------AU-UCUGAACG-
---AA-----------UG-A-------------------G------UUC-----------------CG----
------------------------------------------------------------------------
-------------------------------------------------------------------GU---
------------------------------------------------------------------------
-------------------------------------------------G-AC------GG--------AAU
GUCAG-------------------------------------------------------------------
------------------------------------------------------------------------
------------------------------------------------------------------------
------------------------------------------------------------------------
------------------------------------------------A-----------------------
------------------------------------------------------------------------
------------------------------------------------------------------------
------------------------------------------------------------------------
------------------------------------------------------------------------
------------------------------------------------------------------------
-------------------GUUUAGAGG-----------------ACUAG--G---GUUGCAUCAUCAGUCU
GGAAUCAUUGGCCAGGGAGUUUA-U-AGUAGGGGCAA-GCUUAAGC--G-U--------CACCC-------G
G-CG--CGUA--GGCACAGGGAAAC-CAA-----------------UAU----------------------U
CUUG-CAACCUA----A-------------------------------------------------------
---UUAUCACAUUGGGAUGAUCUAUCC------GAUAACGGGUUGGCUCUGCCUCCC--AAAAUCGCUUAGU
GAGUCCCUCUAAGGGAU--UAGG------------------------------------CCA-GAAGAA-UG
GUCU-----------------------------------UAAA--------------------------GG-
G-C-CU----------------------AAAGUCUCUGCUGUAA-UACCCGAAGCC-GGUCGAUCUAUUCCU
GGCCAAGAUGAAGUCGUU-CGAAAG-GAUGAUGGAAGUCUGC--AGUAGUGCUGACGUGCAAUUCGC-UUAC
AUGAGCUGGGAAUAGGAGCAAAAGGCUAAUCUAGGACGG-UGAUAGCUGGUUCUCCCCGAAG-UUGGUGUCA
GCCCAGCCUCGA-GUGA-------------------------------------------------------
----------------------GG-----UA-GCCCGUGUUGUAGAGC-ACUAAUU-GUG-GAGU-AUGGGG
G-A-GA-AA-UUCC-U-CGCUUCGCUCUUAAACUCCGAAGAUGCGG--------------------------
```

------------------------------------------------------------------------
------------------G-CG-C--CA-UAGA--CCUCGGG-AGACGGG-U-UACCGG-GCG-UAAAGUCC
GG-U-GCCGAGAA-GGGAAC---AACCUAGACUUG-G-GUUAAGGUCCCUAAGU--GC-AGG-UU-AA---G
-U-GUUAAA--CGAAAGGGCGUUAUAUUUCUAAAACAGCAAAAGGAUAGGUAUAGAAGUA------------
------------------------------------------------------------------------
------------------------------------------------------------------------
------------------------------------------------------------------------
------------------------------------------------------------------------
------------------------------------------------------------------------
------------------------------------------------------------------------
------------------------------------------------------------------------
------------------------------------------------------------------------
------------------------------------------------------------------------
------------------------------------------------------------------------
------------------------------------------------------------------------
------------------------------------------------------------------------
------------------------------------------------------------------------
------------------------------------------------------------------------
------------------------------------------------------------------------
------------------------------------------------------------------------
------------------------------------------------------------------------
------------------------------------------------------------------------
------------------------------------------------------------------------
------------------------------------------------------------------------
------------------------------------------------------------------------
------------------------------------------------------------------------
------------------------------------------------------------------------
------------------------------------------------------------------------
------------------------------------------------------------------------
------------------------------------------------------------------------
------------------------------------------------------------------------
------------------------------------------------------------------------
------------------------------------------------------------------------
------------------------------------------------------------------------
------------------------------------------------------------------------
------------------------------------------------------------------------
------------------------------------------------------------------------
------------------------------------------------------------------------
------------------------------------------------------------------------
------------------------------------------------------------------------
------------------------------------------------------------------------
------------------------------------------------------------------------
------------------------------------------------------------------------
------------------------------------------------------------------------
------------------------------------------------------------------------

```
------------------------------------------------------------------------
------------------------------------------------------------------------
------------------------------------------------------------------------
------------------------------------------------------------------------
------------------------------------------------------------------------
------------------------------------------------------------------------
------------------------------------------------------------------------
------------------------------------------------------------------------
------------------------------------------------------------------------
------------------------------------------------------------------------
------------------------------------------------------------------------
------------------------------------------------------------------------
------------------------------------------------------------------------
------------------------------------------------------------------------
------------------------------------------------------------------------
------------------------------------------------------------------------
------------------------------------------------------------------------
------------------------------------------------------------------------
------------------------------------------------------------------------
------------------------------------------------------------------------
------------------------------------------------------------------------
------------------------------------------------------------------------
------------------------------------------------------------------------
------------------------------------------------------------------------
------------------------------------------------------------------------
------------------------------------------------------------------------
------------------------------------------------------------------------
------------------------------------------------------------------------
------------------------------------------------------------------------
------------------------------------------------------------------------
------------------------------------------------------------------------
------------------------------------------------------------------------
------------------------------------------------------------------------
------------------------------------------------------------------------
-------------------------------------------------
>LSUASG_NODE_558_length_1127_cov_98.925000_g407_i0146_5'/1-981
AGCUUGUUACGCUCUUCUUCGGUGCCCAAGCCAAGAU--AUCCACCAAAU-AGCGGAGG-C-GUUAUG-GCG
U-C---UACUGAAGGAGAGCGUAA--CAAGCUGCGAUAAGCUUGGGUUAGUCGCAUGUAGACGU------AG
ACC-CCA-AGAUUCUCGAAUAGGACA-UCCG-----CUU-------GC----GGC----------C------
----------------------------------AAA----U-A-AUCC-GCAAGA---------------U
UCC--------C-----------UCG----------------------------------------------
--U--------------------GG-G-AA-GGGGAACCCUGAGAAGUAAAACAUUUUAGUAUCAG--GAGG
AAAAGAAAACAA--UAGUGAUUCCGUGA--GU-AA-CGGCGAGCGAAAGCGGA-U-CA-GAUCAAACCGAAU
CCCU-AGGGA---------AA------------C---CUA-GUGGAAUGUGGUG--UGU-------------
G-GA--CCU-CGU-ACGAC----------------------------------CUC-----------CC--U
```

```
A-------GAAAA--UAUG------AAUACU-CCUG------------------------------------
-------GAA-AG-UUGAG-------CCAGAGAGCGUGAUAGCCGCGUA---------AUA---G----UAA
U------CUAGAAGG-UUGC----------------UGCG-AGAGU--UCCCGAGUAGUGCGGGAUG---GU
UUUCUCGUACGAA-UGUACCAGAUACA--CGCUGGUAAAUCUAAAUA--UAU-C-CCA-AGACCGAUAGU-G
AAA-U-AGUAGCGUGAGCGAAAG-AUGAAA-AGUAA-CCCGG-AAGGGC-C-GUG-AAAAGUGCUUGAAACC
A-AAUAGCUUAGU-AAG-GA-UGCA---GCCUGAAA-GGAUAG-----------------------------
------------------------------------------------------------------------
------------------------------------------------------------------------
------------------------------------------------------------------------
------------------------------------------------------------------------
------------------------------------------------------------------------
------------------------------------------------------------AU-UCUGAACG-
---AA-----------UG-A-------------------G------UUC-----------------CG----
------------------------------------------------------------------------
-------------------------------------------------------------------GU---
------------------------------------------------------------------------
-------------------------------------------------G-AC------GG--------AAU
GUCAG-------------------------------------------------------------------
------------------------------------------------------------------------
------------------------------------------------------------------------
------------------------------------------------------------------------
------------------------------------------------A-----------------------
------------------------------------------------------------------------
------------------------------------------------------------------------
------------------------------------------------------------------------
------------------------------------------------------------------------
------------------------------------------------------------------------
-------------------GUUUAGAGG-----------------ACUAG--G---GUUGCAUCAUCAGUCU
GGAAUCAUUGGCCAGGGAGUUUA-U-AGUAGUGGCGA-GCUUAAGU--G-U--------CUUAC-------G
A-CA--CGGA--GGCGCAGGGAAAC-CAA-----------------UAU----------------------U
UCUG-CACCUUG----A-------------------------------------------------------
---UUAUCACAUUAAAAUUAUCUAUUC-------GUCCGAGAAAGGUUCUGCCGAUC--AACGACGCUUAGU
GCGCUUUUCUAAGGAAU--CAAG------------------------------------CUA-GAGAAA-UG
GUCU-----------------------------------CAAA--------------------------GG-
G-C-CU----------------------AUAGUCUCUGCUAUAA-CACCCGAAGCC-GGUCGAUCUAUUCCU
GGGCAAGAUGAAGUCGUU-CGAAAG-GAUGAUGGAGGUCUGC--AGUAGUGCUGACGUGCAAUUCGC-UU--
------------------------------------------------------------------------
------------------------------------------------------------------------
------------------------------------------------------------------------
------------------------------------------------------------------------
------------------------------------------------------------------------
------------------------------------------------------------------------
------------------------------------------------------------------------
------------------------------------------------------------------------
------------------------------------------------------------------------
```

```
--------------------------------------------------------------------------------
--------------------------------------------------------------------------------
--------------------------------------------------------------------------------
--------------------------------------------------------------------------------
--------------------------------------------------------------------------------
--------------------------------------------------------------------------------
--------------------------------------------------------------------------------
--------------------------------------------------------------------------------
--------------------------------------------------------------------------------
--------------------------------------------------------------------------------
--------------------------------------------------------------------------------
--------------------------------------------------------------------------------
--------------------------------------------------------------------------------
--------------------------------------------------------------------------------
--------------------------------------------------------------------------------
--------------------------------------------------------------------------------
--------------------------------------------------------------------------------
--------------------------------------------------------------------------------
--------------------------------------------------------------------------------
--------------------------------------------------------------------------------
--------------------------------------------------------------------------------
--------------------------------------------------------------------------------
--------------------------------------------------------------------------------
--------------------------------------------------------------------------------
--------------------------------------------------------------------------------
--------------------------------------------------------------------------------
--------------------------------------------------------------------------------
--------------------------------------------------------------------------------
--------------------------------------------------------------------------------
--------------------------------------------------------------------------------
--------------------------------------------------------------------------------
--------------------------------------------------------------------------------
--------------------------------------------------------------------------------
--------------------------------------------------------------------------------
--------------------------------------------------------------------------------
--------------------------------------------------------------------------------
--------------------------------------------------------------------------------
--------------------------------------------------------------------------------
--------------------------------------------------------------------------------
--------------------------------------------------------------------------------
--------------------------------------------------------------------------------
--------------------------------------------------------------------------------
--------------------------------------------------------------------------------
--------------------------------------------------------------------------------
--------------------------------------------------------------------------------
```

```
------------------------------------------------------------------------
------------------------------------------------------------------------
------------------------------------------------------------------------
------------------------------------------------------------------------
------------------------------------------------------------------------
------------------------------------------------------------------------
------------------------------------------------------------------------
------------------------------------------------------------------------
------------------------------------------------------------------------
------------------------------------------------------------------------
------------------------------------------------------------------------
------------------------------------------------------------------------
------------------------------------------------------------------------
------------------------------------------------------------------------
------------------------------------------------------------------------
------------------------------------------------------------------------
------------------------------------------------------------------------
------------------------------------------------------------------------
------------------------------------------------------------------------
------------------------------------------------------------------------
------------------------------------------------------------------------
------------------------------------------------------------------------
------------------------------------------------------------------------
------------------------------------------------------------------------
------------------------------------------------------------------------
------------------------------------------------------------------------
------------------------------------------------------------------------
------------------------------------------------------------------------
------------------------------------------------------------------------
-------------------------------------------------
>LSUASG_NODE_962_length_959_cov_159.913377_g720_i0_REV/1-959
------------------UAGACGCCAAUAC----GC--CUCCGCUAUUU-GGUGGAUA-U-CUUGGC-UUG
G-G---AACCGAUGAAGAGCGUGA--CAAGCUGCGAUAAGCUCGGGUUAGUCGCAUGAAGACGC------UG
ACC-CCG-AGAUACUCGAAUAGGACA-UCCG-----CUU-------GC----GAA----------C------
----------------------------------CAA----U-A-AUUC-GCAAGA---------------U
UCC--------C-----------UCG----------------------------------------------
--U--------------------GG-G-AA-GGGGAACCCUGAGAAGUAAAACAUUUUAGUAUCAG--GAGG
AAAAGAAAACAA--UAGUGAUUCCGUGA--GU-AA-CGGCGAGCGAAAGCGGA-U-CA-GAUCAAACCGAAU
CCCU-AGGGA---------AA------------C---CUA-GUGGAAAUGUGGUG--UGU------------
G-GA--CCU-CGU-ACGAC----------------------------------CUC-----------CC--U
A-------GAAAA--UAUG------AAUACU-CCUG------------------------------------
-------GAA-AG-UUGAG-------CCAGAGAGCGUGAUAGCCGCGUA---------AUA---G----UAA
U------CUAGAAGG-UUGC----------------UGCG-AGAGU--UCCCGAGUAGUGCGGGAUG---GU
UUUCUCGUACGAA-UGUACCAGAUACA--CGCUGGUAAAUCUAAAUA--UAU-C-CCA-AGACCGAUAGU-G
AAA-U-AGUAGCGUGAGCGAAAG-AUGAAA-AGUAA-CCCGG-AAGGGC-C-GUG-AAAAGUGCUUGAAACC
```

```
A-AAUAGCUUAGU-AAG-GA-UGCA---GCCUGAAA-GGAUAG-----------------------------
------------------------------------------------------------------------
------------------------------------------------------------------------
------------------------------------------------------------------------
------------------------------------------------------------------------
------------------------------------------------------------------------
------------------------------------------------------------AU-UCUGAACG-
---AA-----------UG-A-------------------G------UUC-----------------CG----
------------------------------------------------------------------------
-------------------------------------------------------------------GU---
------------------------------------------------------------------------
-------------------------------------------------G-AC------GG--------AAU
GUCAG-------------------------------------------------------------------
------------------------------------------------------------------------
------------------------------------------------------------------------
------------------------------------------------------------------------
------------------------------------------------A-----------------------
------------------------------------------------------------------------
------------------------------------------------------------------------
------------------------------------------------------------------------
------------------------------------------------------------------------
------------------------------------------------------------------------
-------------------GUUUAGAGG-----------------ACUAG--G---GUUGCAUCAUCAGUCU
GGAAUCAUUGGCCAGGGAGUUUA-U-AGUAGUGGCGA-GCUUAAGU--G-U--------CUUAC-------G
A-CA--CGGA--GGCGCAGGGAAAC-CAA-----------------UAU----------------------U
UCUG-CACCUUG----A-------------------------------------------------------
---UUAUCACAUUAAAAUUAUCUAUUC-------GUCCGAGAAAGGUUCUGCCGAUC--AACGACGCUUAGU
GCGCUUUUCUAAGGAAU--CAAG------------------------------------CUA-GAGAAA-UG
GUCU-----------------------------------CAAA--------------------------GG-
G-C-CU----------------------AUAGUCUCUGCUAUAA-CACCCGAAGCC-GGUCGAUCUAUUCCU
GGGCAAGAUGAAGUCGUU-CGAAAG-GAUGAUGGAGGUCUGC--AGUAGUGCUGACGUGCAAUUCGC-UU--
------------------------------------------------------------------------
------------------------------------------------------------------------
------------------------------------------------------------------------
------------------------------------------------------------------------
------------------------------------------------------------------------
------------------------------------------------------------------------
------------------------------------------------------------------------
------------------------------------------------------------------------
------------------------------------------------------------------------
------------------------------------------------------------------------
------------------------------------------------------------------------
------------------------------------------------------------------------
------------------------------------------------------------------------
------------------------------------------------------------------------
```

```
------------------------------------------------------------------------------------
------------------------------------------------------------------------------------
------------------------------------------------------------------------------------
------------------------------------------------------------------------------------
------------------------------------------------------------------------------------
------------------------------------------------------------------------------------
------------------------------------------------------------------------------------
------------------------------------------------------------------------------------
------------------------------------------------------------------------------------
------------------------------------------------------------------------------------
------------------------------------------------------------------------------------
------------------------------------------------------------------------------------
------------------------------------------------------------------------------------
------------------------------------------------------------------------------------
------------------------------------------------------------------------------------
------------------------------------------------------------------------------------
------------------------------------------------------------------------------------
------------------------------------------------------------------------------------
------------------------------------------------------------------------------------
------------------------------------------------------------------------------------
------------------------------------------------------------------------------------
------------------------------------------------------------------------------------
------------------------------------------------------------------------------------
------------------------------------------------------------------------------------
------------------------------------------------------------------------------------
------------------------------------------------------------------------------------
------------------------------------------------------------------------------------
------------------------------------------------------------------------------------
------------------------------------------------------------------------------------
------------------------------------------------------------------------------------
------------------------------------------------------------------------------------
------------------------------------------------------------------------------------
------------------------------------------------------------------------------------
------------------------------------------------------------------------------------
------------------------------------------------------------------------------------
------------------------------------------------------------------------------------
------------------------------------------------------------------------------------
------------------------------------------------------------------------------------
------------------------------------------------------------------------------------
------------------------------------------------------------------------------------
------------------------------------------------------------------------------------
------------------------------------------------------------------------------------
------------------------------------------------------------------------------------
------------------------------------------------------------------------------------
------------------------------------------------------------------------------------
```

```
------------------------------------------------------------------------
------------------------------------------------------------------------
------------------------------------------------------------------------
------------------------------------------------------------------------
------------------------------------------------------------------------
------------------------------------------------------------------------
------------------------------------------------------------------------
------------------------------------------------------------------------
------------------------------------------------------------------------
------------------------------------------------------------------------
------------------------------------------------------------------------
------------------------------------------------------------------------
------------------------------------------------------------------------
------------------------------------------------------------------------
------------------------------------------------------------------------
------------------------------------------------------------------------
------------------------------------------------------------------------
------------------------------------------------------------------------
------------------------------------------------------------------------
------------------------------------------------------------------------
------------------------------------------------------------------------
------------------------------------------------------------------------
------------------------------------------------------------------------
------------------------------------------------------------------------
-------------------------------------------------
>LSUASG_NODE_399_length_1227_cov_20.866102_g282_i0_REV/1-1227
------------------------------------------------------------------------
-----------------------------GCUGCGAUAAGCUCGGGUUAGUCGCACGUAGACGU------UG
ACC-CCG-AGAUCCCCGAAUAGGACA-UCCG-----CUU-------GC----GGC----------C------
----------------------------------CAU----A-A-AUCC-GCAAGA---------------C
UCC---------CAUAUCCAAUAGGUG---------------------------------------------
--A--------------------GG-G-AG-AGGGAACCCAGGGAAGUAAAACAUUUUAGUACUUG--GAGG
AAAAGAAACCAA--UAGGGAUUCCGUUA--GU-AA-CGGCGAGCGAAAGCGGA-A-UA-GGUCAAACUGAAU
CCGU-AAGAGA-------CAA------------U---CUU-ACGGAGAUGUGGUG--UAU------------
G-GA--CCU-CGU-ACGGC----------------------------------CUC-----------CU--U
A-------GGGAA--GGUG------AAUAGG-CCUG------------------------------------
-------GAA-UG-GCCAG-------CCAUAGAGCGUGACAGCCGCGUA---------ACU---A----AAA
U------CUAAUAGG-CUGA----------------UGCG-AGGUA--UCCCGAGUAGUGCGGGAUA---GU
UUUCUCGUACGAA-CCUGCCUGACACA--CACAGGCAAACCUAAAUA--UUC-C-CCA-AGACCGAUAGC-G
UAA-U-AGUAGCGUGAGCUAAAG-AUGAAA-AGUAA-CCCGG-AAGGGU-C-UUG-AAAAGUGCUUGAACCC
A-AACAGCUUAAU-AAG-GA-UGGG---GCCUGAAA-GGACAG-----------------------------
------------------------------------------------------------------------
------------------------------------------------------------------------
------------------------------------------------------------------------
------------------------------------------------------------------------
```

```
------------------------------------------------------------------------
------------------------------------------------------------AU-CCUCCCCG-
---AA-----------CG-U-------------------G------UUG-----------------CG----
------------------------------------------------------------------------
-------------------------------------------------------------------GU---
------------------------------------------------------------------------
-------------------------------------------------G-AC------GC--------AAC
GACGG-------------------------------------------------------------------
------------------------------------------------------------------------
------------------------------------------------------------------------
------------------------------------------------------------------------
------------------------------------------------A-----------------------
------------------------------------------------------------------------
------------------------------------------------------------------------
------------------------------------------------------------------------
------------------------------------------------------------------------
------------------------------------------------------------------------
-------------------GGGGAGAGG-----------------ACUAG--G---GUUCCAUCAUCAGUCU
GGAAUCAUUGGCCAGGGAGUUUG-U-AGUAGUGGCGA-GCUUAAGC--G-A--------CUUAC-------G
U-CG--CGUA--GGCGAAGGGAAAC-CAA-----------------UAU----------------------U
CUCG-CAAGCCC----G-------------------------------------------------------
---UUAUCACAUCAGGGUGAUCUACCG-------GUUAGAGGAAGGCUCUGCCUUAC--AAUAACUAGUAGU
GAAUCCUCCUAAGGAAC--GGGU------------------------------------CCA-GGAGAA-CG
GUCU-----------------------------------CAAA--------------------------GG-
G-C-CU----------------------AAAGUCACUGCUAUAA-UACCCGAAGCC-GGUCGAUCUAUUCCU
GGCCAAGAUGAAGUCGCU-CGAAAG-GGCGAUGGAGGUCUGC--AGUAGUGCUGAUGUGCAAUUCGC-UUAC
AUGAGCUGGGAAUAGGAGCAAAGGCUAAUCUAGGACGG-UGAUAGCUGGUUCUCCCCGAAG-UUGGUGUCA
GCCCAGCCUCGA-GUGA-------------------------------------------------------
----------------------GG-----UA-GCCCGUGUUGUAGAGC-ACUAAUU-GUG-GAGU-AUGGGG
G-A-GA-AA-UUCC-U-CGCUUCGCUCUUAAACUCCGAAGAUGCGG--------------------------
------------------------------------------------------------------------
------------------G-CG-C--CA-UAGA--CCUCGGG-AGACGGG-U-UACCGG-GCG-UAAAGUCC
GG-U-GCCGAGAA-GGGAAC---AACCUAGACUUG-G-GUUAAGGUCCCUAAGU--GC-AGG-UU-AA---G
-U-GUUAAA--CGAAAGGGCGUUAUAUUUCUAAAACAGCAAAAGGAUAGGUAUAGAAGUA------------
------------------------------------------------------------------------
------------------------------------------------------------------------
------------------------------------------------------------------------
------------------------------------------------------------------------
------------------------------------------------------------------------
------------------------------------------------------------------------
------------------------------------------------------------------------
------------------------------------------------------------------------
------------------------------------------------------------------------
------------------------------------------------------------------------
------------------------------------------------------------------------
```

```
------------------------------------------------------------------------
------------------------------------------------------------------------
------------------------------------------------------------------------
------------------------------------------------------------------------
------------------------------------------------------------------------
------------------------------------------------------------------------
------------------------------------------------------------------------
------------------------------------------------------------------------
------------------------------------------------------------------------
------------------------------------------------------------------------
------------------------------------------------------------------------
------------------------------------------------------------------------
------------------------------------------------------------------------
------------------------------------------------------------------------
------------------------------------------------------------------------
------------------------------------------------------------------------
------------------------------------------------------------------------
------------------------------------------------------------------------
------------------------------------------------------------------------
------------------------------------------------------------------------
------------------------------------------------------------------------
------------------------------------------------------------------------
------------------------------------------------------------------------
------------------------------------------------------------------------
------------------------------------------------------------------------
------------------------------------------------------------------------
------------------------------------------------------------------------
------------------------------------------------------------------------
------------------------------------------------------------------------
------------------------------------------------------------------------
------------------------------------------------------------------------
------------------------------------------------------------------------
------------------------------------------------------------------------
------------------------------------------------------------------------
------------------------------------------------------------------------
------------------------------------------------------------------------
------------------------------------------------------------------------
------------------------------------------------------------------------
------------------------------------------------------------------------
------------------------------------------------------------------------
------------------------------------------------------------------------
------------------------------------------------------------------------
------------------------------------------------------------------------
------------------------------------------------------------------------
------------------------------------------------------------------------
```

```
------------------------------------------------------------------------
------------------------------------------------------------------------
------------------------------------------------------------------------
------------------------------------------------------------------------
------------------------------------------------------------------------
------------------------------------------------------------------------
------------------------------------------------------------------------
------------------------------------------------------------------------
------------------------------------------------------------------------
------------------------------------------------------------------------
------------------------------------------------------------------------
------------------------------------------------------------------------
------------------------------------------------------------------------
------------------------------------------------------------------------
------------------------------------------------------------------------
------------------------------------------------------------------------
------------------------------------------------------------------------
------------------------------------------------------------------------
------------------------------------------------------------------------
-------------------------------------------------
>LSUa_186497_PYRFU/1-3048
----------------------GGGCUAGGA----CG--CUAAGCCGCCC-GGUGGAUG-G-CUCGGC-UCG
GGG---CGCCGACGAAGGGCGUGG--CAAGCUGCGAUAAGCCCCGGCGAGGCGCAGGCAGCCGU------CG
AAC-CGG-GGAUUCCCGAAUGGGACC-U--------CCC-------GC----GGC---------------UU
A-U---------------------------G---CCG----C-A-CUCC-GGG-----G-------------
-----------U-----------UUAU--------------------------------------------C
C-C--------------------CG-G-AG-GGGGAACGCGGGGAAUUGAAACAUCUUAGUACCCG--CAGG
AAAAGAAAGCAA--AAGCGAUGCCGUGA--GU-AG-GGGCGACCGAAAACGGC-A-GA-GGGCAAACUGAAC
CCCG-GGCCGA-------CGA------------G---GUU-CGGGGGAUGUGGGG--UUG------------
U-AG--GGC-CCC-C-----G------UAU---------------GAGACC--CUC-----------GC--G
G-------GUGAA--GCCG------AAGUCC-GCUG------------------------------------
-------GAA-CG-CGGCG-------CCGGAGAGGGUGAUAGCCCCGUA---------GGC---G----UAA
G------CCCGCAGG-GU------------------CUCG-GGGGA--CCCUGAGUACCGUCGGUUG---GA
UAUCCGGCGGGAA-GCUGGGAGGCAUC--GGCUCCCAACCCUAAAUA--CGU-C-CCG-AGACCGAUAGC-G
AAC-U-AGUACCGUGAGGGAAAG-CUGAAA-AGCAC-CCCGG-GAGGGG-G-GUG-AAAAGAGCCUGAAACC
G-GGCGGCGAUAG-GAG-GG-UGCG---GCCCGAAA-GGAAUG-----------------------------
------------------------------------------------------------------------
------------------------------------------------------------------------
------------------------------------------------------------------------
------------------------------------------------------------------------
------------------------------------------------------------------------
------------------------------------------------------------AG-CCUCCCCG-
---AA-----------GG-A-------------------A------ACC-----------------GCG---
------------------------------------------------------------------------
--------------------------------------------------------------G--C-GA---
```

```
------------------------------------------------------------------------
-------------------------------------------------C-GC------GG--------GAG
UACGA-------------------------------------------------------------------
------------------------------------------------------------------------
------------------------------------------------------------------------
------------------------------------------------------------------------
------------------------------------------------G-----------------------
------------------------------------------------------------------------
------------------------------------------------------------------------
------------------------------------------------------------------------
------------------------------------------------------------------------
------------------------------------------------------------------------
-------------------GGGAGG-GG-----------------ACCGG--G---GUUGCACCGUCCGUCU
UGAAACACGGGGCAGGGAGUUCG-C-GGCCGUGGCGA-GGUUAAGG--G-GG--------UUAA-------G
C-CC--CGUA--GCCGCAGGGAAAC-CGA-----------------CAU----------------------G
CCCG-CAGCCGG----G-------------------------------------------------------
---CU-------------------------------------------------------------------
------------UAUGC--CCGG------------------------------------UGA-GGGGCG-GG
GUGC-----------------------------------GAAA--------------------------GC-
G-C-CC----------------------GGAGUCACGGCCGCGA-GACCCGAAACC-GGUCGAUCUAGCCCG
GGGCAGGGUGAAGUCCCU-CAACCG-AGGGAUGGAGGCCCGC-UAGGGGUGCUGAUGUGCAGUUCGC-UCCC
GUGACCCCGGGCUAGGGGUGAAAGGCCAAUCGAGGCCGG-AGAUAGCUGGUUCCCGCCGAAU-CAUCCCGCA
GGAUGGCCUCCC-CGGA-------------------------------------------------------
----------------------GG-----UA-GGCGGUGGGGUAGAGC-ACUGAUU-GGA-GGUG-CAGGGG
G-C-GA-AA-GCCC-C-CGGCCUCCUGUCAAACUCCGAACCCACCG--------------------------
------------------------------------------------------------------------
------------------C-CG-C--CG-UAGA--UGGGGGG-AGUAGGG---UGGC-G-GUG-UAAGCC-G
UC-C-ACCGAGAG-GGGAAC---AACCCAGACCGG-G-GUUAAGGCCCCAAAGU--GC-CGG-CU-AA---G
-U-GUUACU--CCAAAGGGUGUCCCGGGCCUUAGACAGCGGGGAGGUAGGCUUAGAAGCAGCCA-UCCUUUA
AAGAGUGCGUAACAGCUCACCCGUC-----GAGGUCCGGGGCCCCGAAAAU-GGAC-GGGGCUCA-AGCCGG
C----------CGCCGAG-ACCCCGGCGCACGGA--C-----------------------------------
------------------------------------------------------------------------
-----CGAUUG-------------------------------------------------------------
-------------------------------GUCCGU-GAUC-GGGUAGGCGGGC-GUGCCGG---------
-----------U-G-GCGUA-GAAGCCG--G-GCCGUAAGGU--C-CGGUGGAGCC-G-CCG-GUAUCGCGG
AUCC-UGCCGG-GAGUA-GCAGCG--UAGUC-GGGUGA--G---------------AAUCCC-GAC-CGCCG
G-AGGGGC-CAGGGUUCC-ACGGCAAUG-GUCGUCAGCCGUGGGUUAGUCGG--UC-CUAACCCCGCCCGU-
AAC-UCGG---C-GCG--------------------------------------------------------
----------------------------------------GGG-GA-A-AGGGA-AACGGGUU-UAUA-UUC
CCGUA----------------------------------CCGCG----GGGGUAGG-------U--------
------------------------------------------------------------------------
-------GCGG--CAAC--------GC-AA-----GC--------------------CCGG-A-GGGU-GAC
GCCUCGG-------------GGUAGG-CGGACCGGC-C-------GAUGA---------------GGCCGG-
CUAA-G-CGU-AU-----A-A-------------------GC-----CC----------------GGGG-AG
U-G-CC--------------------GUAAU-GGCGAGAACC---------GGGUGA----AAG-CG-CGA-
```

AUGG-CC-----------------------------------------------------------------
-----CCCCC---GUUAGGGGGGUU-C-CGC-C-G-AUCCCUGGGGCCC-GUGAAA-A-GCC-CUCCGGGAA
-U----------------------------------------------------UCCGAUCC-CC-CGCGAC
CGUACCGAGAACCGAC------------------------------ACAGGUGC-CCCUGGGU-GAGAAGCC
UAAGGC-GUGU-CGGGGGAAACCCG-G-CCGAGGGAACUCGGC-AAACUGGCCCCGUAACUUCGG-GAGAAG
GGGUG-CCUG-C-GG--GUG----------------------------------------------------
------------------------------------------------------------------------
------------------------------------------------------------------------
------------------------------------------------------------------------
------------------------------------------------------------------------
------------------------------------------------------------------------
------------------------------------------------------------------------
------------------------------------------------------------------------
------------------------------------------------------------------------
------------------------------------------------------------------------
------------------------------------------------------------------------
------------------------------------------------------------------------
------------------------------------------------------------------------
------------------------------------------------------------------------
--------------------------------------------------------------CG--UAACCC
GCA--GGUCGCAGUGACUAGGGGGGCCCGA-----CUGUUUAAUAAAAACACAGGUCCCAGCUAGCCCG---
-AA-A-------------GGGUUUGU-ACUGGGGCCGA-CGCCUGCCCAGUGCCG-G-UAUG-UGAAGCCCG
GG-------------UACA-------------ACCGGGUGAAGCAC--CGGUAAAC-GGCGGGGGUAACUAU
AACCCUCUUAAGGUAGCGAAAUUCCUUGUCGGU-UAAAUGCCGACCUGCAUGAAUGGCGUAACGAGG--UCC
CCGCUGUCCCCGGC----CGG-GGCCCGGCGAAACC---UC---UGCCUGGCGCGCAUGCCAGGGACCCCCG
GUGGGAAGCGAAGA-CCCCAUGGAGC-UUUACU--GCAGCCU-GCCGUUGCCACACGGCGGGGGGUGCGCAG
CGU-AGGCGGGAGGCGUCG----------AAGCCCG-GCC----------------------------UCCG
--------------------------------------GG-UCGGGUGGAG-C--C-GU-CCAUGAGACA-C
CGCC-CACUCCCCGCUGUGUGGCUAACCC-CC---------GA-A-A-------------------------
------------------------------------------------------------------------
------------------------------------------------------------------------
------------------------------------------------------------------------
------------------------------------------------------------------------
------------------------------------------------------------------------
------------------------------------------------GGGG-----GGACA-GCGGUAGGU
GGGCAGUUUGGCUGGGGCGGCACGCCUCCGA-AAAGGUAUC-GGAGGCGCCCU-AAGGUCGGCUCAGGCGGG
UCAGGAAUCCGCCGUA-GAGUG-CAAGGGCAA-AAGCCGGCCUGACU-GGAC--------------------
--------------CCGUAACAGAGGCGGGUCCAGCCCCGAAA-GGGUGGCCUAGCGAACCCCU--------
---GU-G-----CCUCC-CCG--------------------GU-GGGGGCCAGGGAUGACAG-AA--AAGCU
ACCCUGGGG-AUAACAGAGUCGUCUCGGGCGAGAGCCCAUAUCGACCCCGAGGCUUGCUACCUCGCUGUCGG
CUCUUCCCAUCCUGGCCCUGCA-GCAGGGGCCAAGGGUGGGGGUGUUCACCCAUUAAAGGGGAACGUGAGCU
GGGUUUAGACCGUCGUGAGACAGGUCGGAUGCUAUCUACC--------------------G-GGG-------
-------------------------------GUGU--UGGCCGCCUGAGGGGAAGG-UGCCCCUAGUACGAG
AGGAA-CAGGGUGCCGCGGCCUC-UGGU-C-UACCGGUUGUCC-UCCC-GGGCAU-C-GCCGGGCAGCUACG
CCGCA-GCCGAUAAGGCCUGAAAGCA-UC--UAAGGCCGAAGCGGCCCCCGA-A-AAU--AGGCGGCC-GU-

```
--UCCCUGGCGCU-----------------------------------------------------------
------------------------------------------------------------------------
------------------------------------------------------------------------
------------------------------------------------------------------------
------------------------------------------------------------------------
------------------------------------------------------------------------
-----------------------------------------UGGGCCUGG-----GCGA----CCGG-CCCA
----------------------------U-------------------------------------------
---------------------UGCCA------------------------GGGA----CG-AGG------GC
UCGGGUAGAAGACCCGGU-UGA-UAGGGC-GGG-GAUGUAAG-CGG-G--------AAGG----GU------
---CAA-------CCGACCCGCUUAGUCC--G-CCGC-CCCUAAUC---GCCCGAGGU-CCUGCCC------
------------------------------------------------------------------------
------------------------------------------------------------------------
------------------------------------------------------------------------
-------------------------------------------------
>LSUASG_2053491_THOR/1-3058 PRDH01000034.1:17074-20131 Candidatus Thorarcha
-----------------------------------GC-UAUACGGCCAGU-GGUGAAUG-A-CUAGGU-UGG
C----UAGCUGAUGAAGGGCGUGA--CAAGCUGCGAUAUGCCGCGGGUAGACGCAUGAAGUCUU------UG
AUC-CGC-GGAUACCUGAAUGGGACU-U--------CCUC------AUA---UGGG--------------UU
AAUACCCCAUAUGGAUC--------C-------GGGGU---C-A-UUCC-----------------------
------------------------------------------------------U-----------------
--------------------------G-GAUUGGGAACGCUCGGAAUUGAAGCAUCUUAGUACGGG--CAGG
AAAAGAAAACAA--UUGUGAUUCCCCUA--GU-AA-CAGCGAGCGAACGGGGA-A-UA-GGCCAAACCGAAU
CCCU-GUGCGA-------UGA-GUA--------C-------AGGGGGAUGUGGUG--UGGGGACCAC-----
--GAACAAU-UAC----------------------------------------CUA--------AGCAU---
--------ACGAA--GCCG------AGGUUG-UCUG------------------------------------
-------GAA-CG-ACAUG-------GCGAAGAGGGUGACACCCCGUA----------GGC---G----UAA
G---AUGCCGGUAUGUUUGUGG-------------------------AACCAGAGUAGCGCGGGUUG---GA
UGUCUCGCGUGAA-AACCGCAGGUACA--AUCUGCGAAGCCUAAAUACUCAGCC-----AAACCGAUAGA-A
AAC-UAUGUAGCGCGAGCGAAAG-UUGAAA-AGCGC-CGCGGGAGCGGG---GUG-AAAAGCACCUGAAACC
A-CUGGUUUACUCAAAG----GAUG-CUGAUUACAA-GAUUAG-----------------------------
------------------------------------------------------------------------
------------------------------------------------------------------------
------------------------------------------------------------------------
------------------------------------------------------------------------
------------------------------------------------------------------------
-------------------------------AU------------------------------ACUCGAGG-
---AA-----------CG-A-------------------A--------------------------------
------------------------------------------------------------------------
----------------------------------------------------------GCAUGAGU-GA---
------------------------------------------------------------------------
---------------------------------------------UUGUG-------------------UUG
UAGGA-------------------------------------------------------------------
------------------------------------------------------------------------
------------------------------------------------------------------------
```

```
------------------------------------------------------------------------
------------------------------------------------------------------------
------------------------------------------------------------------------
------------------------------------------------------------------------
------------------------------------------------------------------------
------------------------------------------------------------------------
------------------------------------------------------------------------
-------------------CUUGAGUGG-----------------AUCGA--G---UCAGCAUCAUCCGUCU
UGAAACACGGGCCAGGGAGUUCA-U-GGGCGUGGCAA-GGAUAAGG-CG------------------AAGAG
C-----CGUA--AACGAAGGGAAAC-CGA-----------------UAU----------------------G
CCCG-CAGCCUU------------------------------------------------------------
------------------------------------------------------------------------
-------------------CGGG------------------------------------CGA-GGGGCG-GC
GUGU-----------------------------------GAAAGU------------------------GC-
G-C-------------------------GAAGUCACUCUCAUGA-UACGAGAAGCC-AGUCGAUCUAGGCUU
GGCCAGGGUGAAGCCGGG-CGAAAG-CUCGGUGGAGGUCCGA--UAGGGUUAUGACGUGCAACUCUU-UCCG
UUGAGCUGAGCGUAGGGGUAAAAGGCUUAUCUAGACUGG-UGAUUGCUCGUUCUCUGCGAAG-UUGCUCGCA
GGCAAGCCCCGC-UUGA-------------------------------------------------------
----------------------GG-----UA-UCACGUGGUGUAGAGC-ACUUUCG-GAUGUGCCCGGGGGG
C---GUCAA-GCCC-U-UCACCAUCCUGAAAACUCCGAAGCUGCGUGAA-----------------------
------------------------------------------------------------------------
-----------------------C--CG-UAGA--CAGCGGG-AGACGGG---UGCCCG-GGG-UAAGCUCG
GCAG-GCCGAGAU-GGGGAG---AACCGAGAGUAC-G-GUUAAGGUCCCUAAGU--AC-CAG-AU-AA---G
-U-GCGGUG--AAGAAGGGCGUCGAGAGCCCAAGACAGUCUGGGGGUGAGCUUAGAAGCAGCUA-UCCCCGA
ACGAAAGCGUAACAGCUCACAGACC-----GAGGCUCUCGGCCUUGAAAAU-GGAC-GGGGCUAA-AUCUGG
U----------CACCGAG-ACCGUACGGCACACG--------------------------------------
------------------------------------------------------------------------
------------------------------------------------------------------------
------------------------GGUGAUUCUGUGU-GAUC-CGUUAGCAGAGC-GAUCCGG---------
----------CU---GGGUA-GAAGUGG--G-GCUGUGAAGU--C-CCGUGGACCG-A-CCG-GAUUUGAUG
AUCC-UGGCAG-UAGUA-ACAGCG--AAGCA-UGGUGA--G---------------AAUCCA-UGU-CACCG
A-AAGGGA-AAGGGUUCU-UCGGCGAUGCGUCAUCAGCCGAAGUUUAGUCGA--UC-CUAAGUGCCGCCGU-
AAC-UCGA---GCGUC--------------------------------------------------------
------AC------------------------------------GA-A-AGGGA-AUCAGGUC-AAGA-AUC
CUGAA----------------------------------CCUCG----GAAGUACG------AUU-------
--------------C---------------------------------------------------------
-------GCGG--CAAC--------GCAAAGC--CGC-------------------ACUUG-U------GAC
UCAUCUG-------------GACGAG-CCUUACACA-GU-------UGUCG--------------CUGUGU-
UUAACACCAU--------A-A-------------------GC-----CU----------------ACUG-AG
U-A-CU--------------------GUAAU-AGUGAGAAGC---------AGGCGAA---AGG----UGGA
AUA--GC--------------------------------------------------------------AGA
UGG---------UCGGACAUCUGUU-U-GGCCC---AUUUCUGGUGACC-UUGAAA-A-----ACGAGUGCG
------------------------------------------------------GAAGAAUCUUC-CGAGUC
CGUACCAAGAUCCGAC------------------------------ACAGGUU--CCCUGGCU-GAGUAGGC
CAAGGU-GUAUCCAGG-UUAACCCU-C-CUGAGGGAAAUCGGC-CAAUUAGCAGCGUAACUUCGG-GAUAUG
```

```
CUGUG-CCUG---GA-----------UUAGUGCA--------------------------------------
------------------------------------------------------------------------
------------------------------------------------------------------------
------------------------------------------------------------------------
------------------------------------------------------------------------
----------------------------------------A-------------------------------
------------------------------------------------------------------------
------------------------------------------------------------------------
------------------------------------------------------------------------
------------------------------------------------------------------------
------------------------------------------------------------------------
------------------------------------------------------------------------
------------------------------------------------------------------------
------------------------------------------------------------------------
------------------------------------------------------------------GCUUUU
CCA--GGUCUUAGUGACUAGAGGGUCCCAA-----CUGUUUAAUACAAACAUAGCUGACUGCGAGCGCG---
-AA-A-------------GCGUUAGU-ACAGUCAGUGA-AUCCUGGUCUGUGCCC-G-UAUGUUAAAGCUCG
GG-----------A------------------ACGGGUCGAAGCAC--GGGUUAAC-GCCGGGACUAACUAU
GAGUCUCUUAAGGUAGCCAAAUGCCUUGUCGGU-UAAAUGCCGACGUGCAUGAAGGGAGCAACGAGG--UCC
CUACUGUCCCCAGG----AGG-AACCGGGUGAACUU---AC---ACCGUACCGAACAGUGUACGUAGCGCCA
AUGGGAAAAGAAGU-CCCCGCGGAGG-UUUACUACGCAGUCU-GUCGUUGCUCGGCUGGUGCCGAUGCAGAG
CGU-AGUCAGGAGC-------------GUUGAAGCACAUU----------------------------CUCC
--------------------------------------GGGAUGUGUGAAU-G--C-AA-AGUUGUAACA-C
UGAC-CAUUUGUGCUGACCGAGCUAACCUAGG-------CCAAGC---------------------------
------------------------------------------------------------------------
-----------------------------------------------------------------G------
------------------------------------------------------------------------
------------------------------------------------------------------------
------------------------------------------------------UG----------------
-----------------------------------------------UCUAG-----GGACA-CCGGCAGAU
AGGCAGUUGGGCUGGGGCGGCACAGGGCUGA-CAAGAGAUC-AGCCCUGACCA-AUGCUGAGCUCAGAGGCG
ACGGAAACGUCUCGUU-GAGAU-UAAGGGCAU-AAGCUCGGCUGACA-GGAU--------------------
--------------CCCGAAACGCAACGGAUCCUGGGUAGAAA-UACUGGCCUAACGAACCUCA--------
---AU------AACGCCAAUU--------------------GU-AAGUGAUUGAGAUGACAG-AA--AACUU
ACCCCAGGG-GUAACAGAGUUGUCGCGUGCAAGAGUUCAUAUCGACCACGCGGCUUGCUUCCUCGAUGUCGG
CUCUUGGUAUCCUCGGGUUGCA-UAAGGCCCGAAGGGUGCGGGUGCUCACCGAUUAAAACCGAACGUGAGCU
GGGUUUAGACCGUUGCGAGACAGGUUGGUUUUUAUCUAUU--------------------G-GCG-------
-------------------------------CUGU--UGGAUAUCUGAGGCCAAGA-UGAUACCGGUACGAG
AGGAA-CGUAUUGUCGCGACCUA-UGGU-G-UAUCCGUUGUCC-GACA-GGGCAC-U-GCGGAGUAGCUAAG
UCGUA-AUGGAUAAACGCUGAAUGCA-UC--UAAGCGUGAAGUCAAAGCCGA-A-AAU--AGAUAUCC-AC-
--UUGCUAGACGAUCUUA------------------------------------------------------
-----------------------------------------------------------------------A
UGAC--------------------------------------------------------------------
--------------GA--------------------------------------------------------
------------------------------------------------------------------------
```

```
------------------------------------------------------------------------
---------------------------------------UGGGACUGGUG-----ACAG----UAACGUCGA
------------------------------------------------------------------------
--------------------GUCUA-------------------------GGAA----CG-AGG------AC
UCUCGUAGAAGACGAGGU-AAA-UAGGCA-GGG-GGUGUAAG-CAC-C--------AAG-------------
CUCCGG--------CGAGGUGUUCAGCUC--A-CCUG-UACUAAU----GUCCGAGGC--------------
------------------------------------------------------------------------
------------------------------------------------------------------------
------------------------------------------------------------------------
-------------------------------------------------
>LSUASG_1837170_THOR/1-1057 MEHG01000018.1:1-1057 Candidatus Thorarchaeota
------------------------------------------------------------------------
------------------------------------------------------------------------
------------------------------------------------------------------------
------------------------------------------------------------------------
------------------------------------------------------------------------
------------------------------------------------------------------------
------------------------------------------------------------------------
------------------------------------------------------------------------
------------------------------------------------------------------------
------------------------------------------------------------------------
------------------------------------------------------------------------
------------------------------------------------------------------------
------------------------------------------------------------------------
------------------------------------------------------------------------
------------------------------------------------------------------------
------------------------------------------------------------------------
------------------------------------------------------------------------
------------------------------------------------------------------------
------------------------------------------------------------------------
------------------------------------------------------------------------
------------------------------------------------------------------------
------------------------------------------------------------------------
------------------------------------------------------------------------
------------------------------------------------------------------------
------------------------------------------------------------------------
------------------------------------------------------------------------
------------------------------------------------------------------------
------------------------------------------------------------------------
------------------------------------------------------------------------
------------------------------------------------------------------------
------------------------------------------------------------------------
------------------------------------------------------------------------
------------------------------------------------------------------------
------------------------------------------------------------------------
------------------------------------------------------------------------
```

```
------------------------------------------------------------------------
------------------------------------------------------------------------
------------------------------------------------------------------------
------------------------------------------------------------------------
------------------------------------------------------------------------
------------------------------------------------------------------------
------------------------------------------------------------------------
------------------------------------------------------------------------
------------------------------------------------------------------------
------------------------------------------------------------------------
------------------------------------------------------------------------
------------------------------------------------------------------------
------------------------------------------------------------------------
------------------------------------------------------------------------
------------------------------------------------------------------------
------------------------------------------------------------------------
------------------------------------------------------------------------
------------------------------------------------------------------------
------------------------------------------------------------------------
------------------------------------------------------------------------
------------------------------------------------------------------------
------------------------------------------------------------------------
------------------------------------------------------------------------
------------------------------------------------------------------------
------------------------------------------------------------------------
------------------------------------------------------------------------
------------------------------------------------------------------------
------------------------------------------------------------------------
------------------------------------------------------------------------
------------------------------------------------------------------------
------------------------------------------------------------------------
------------------------------------------------------------------------
------------------------------------------------------------------------
------------------------------------------------------------------------
------------------------------------------------------------------------
------------------------------------------------------------------------
------------------------------------------------------------------------
------------------------------------------------------------------------
------------------------------------------------------------------------
------------------------------------------------------------------------
------------------------------------------------------------------------
------------------------------------------------------------------------
------------------------------------------------------------------------
------------------------------------------------------------------------
------------------------------------------------------------------------
```

```
------------------------------------------------------------------------
------------------------------------------------------------------------
------------------------------------------------------------------------
------------------------------------------------------------------------
------------------------------------------------------------------------
------------------------------------------------------------------------
------------------------------------------------------------------------
------------------------------------------------------------------------
------------------------------------------------------------------------
------------------------------------------------------------------------
------------------------------------------------------------------------
------------------------------------------------------------------------
--------------------------------ACGGGUCGAAGCAC--AGGUUAAC-GCCGGGACUAACUAU
GAGUCUCUUAAGGUAGCCAAAUGCCUUGUCGGU-UAAAUGCCGACGUGCAUGAAGGGAGCAAUGAGG--UCC
CCACUGUCCCCAGG----AGG-GCUCCGGUGAACCU---AC---ACCAUACCGAACAGUGUAUGUAGCGCCA
AUGGGAAGAGAAGU-CCCUGCGGAGG-UUUACUACGCAGUCU-GUCAUUGCCAUUUCUGGUAUCAUACAGAG
UAU-AGUCAGGAGC-------------GACGAACCACACU----------------------------CUCC
--------------------------------------GGGGUGUGGGAAU-G--C-GA-CCGUGUAAUA-C
UGAC-UAUGUUGUUGGGGAUGGCUAACCUAGA-------CCAAGU---------------------------
------------------------------------------------------------------------
------------------------------------------------------------------------
------------------------------------------------------------------------
------------------------------------------------------------------------
-------------------------------------------------------G----AG----------
-----------------------------------------------UCUAG-----GGACA-CUGGCAGAU
AGGCAGUUGGGCUGGGGCGGCACAGGGCUGA-CAAGAGAUC-AGCCCUGACCA-AUGCUGAACUCAGAGGCG
AUGGAAACGUCUCGUU-GAGAU-CAAGGGCAU-AAGUCCGGUUGAUA-GGAU--------------------
--------------CCCGAAAAGCAACGGGUCCUGGGUGGAAA-CACUGGCCUAGCGAACCUUC--------
---AC-U-----AGCCA-AUU--------------------GU-GAGUGGUGGAGAUGACAA-AA--AACUU
ACCCCAGGG-GUAACAGAGUUGUCGCGUGCAAGAGUUCAUAUCGACCACGCGGCUUGCUUCCUCGAUGUCGG
CUCUUGGUAUCCUCGGGUUGCA-CAAGGCCCGAAGGGUGCGGGUGCUCACCGAUUAAAACCGAACGUGAGCU
GGGUUUAGACCGUUGCGAGACAGGUUGGUUUCUAUCUAUU--------------------G-GUA-------
-------------------------------CUGU--CGGAUGUCUGAUGGCAAGA-UGAUUCCGGUACGAG
AGGAA-CGAAUUGUCGCGACCUA-UGGU-G-UAUCGAUUGUCU-GAGA-AGGCAC-U-GUCGAGUAGCUAAG
UCGUA-AUGGAUAAACGCUGAAUGCA-UC--UAAGCGUGAAGCCAGACCAGA-A-AAU--AGACAUCC-AC-
--UUGC-UAGACG------AUCU-------------------------------------------------
-----------------------------------------------------------------------G
CGAA--------------------------------------------------------------------
-------AGAC---GA--------------------------------------------------------
------------------------------------------------------------------------
------------------------------------------------------------------------
---------------------------------------CGGGACUGGUG-----ACAG----UAACGUCGA
----------------------------A-------------------------------------------
--------------------GUCUA-------------------------GGAA----CG-AGG------AC
UCUUGUAGAAGACAAGGU-AAA-UAGGGC-GGG-GGUGUAAG-CAC-C--------AAG-------------
```

```
CUUCG--------GCGAGGUGUUCAGCUC--A-CCGC-UACUAAU----GUCCGA-----------------
------------------------------------------------------------------------
------------------------------------------------------------------------
------------------------------------------------------------------------
-------------------------------------------------
>LSUASG_Freya_rrl_bb_B40_Guay15_scaffold_00224/1-3397
---------------------------------------CUAUGCCGCCC-GGUGGAUG-G-CUCGGC-UCG
G-G---CGCUGACGAAGGCCGUGG--CAAGCUGCGAUAAGCCCCGGGUAGGCGCAGGCAGCCGU------UG
AUC-CGG-GGAUUGCCGAAUGGGACC-U--------CCU-------GC----CAGGGG-GG---------AU
A-A------------CC---C----C----C---UGG----C-G-UUCC-CGU-----GCGG----------
-----------G-----------UGAC--------------GAG------------C--CCG---------C
A-U--------------------GG-G-AA-CGGGAACCCGCCGAAUUGAAACAUCUUAGUAGGCG--GAGG
AAAAGAAACCAA--AAGGGAUCCCGGGA--GU-AG-GGGCGACCGAAACCGGG-A-GA-GCCCAAACCAAAC
CCCC-GGAGGA--------AA------------C---UCC-GGGGGGAUGUGGUG--UUG------------
C-AG--GAC-CCU-G-----C------CGCC------------------GC--CUC-----------UU--G
G-------GGGAA--GCCG------AACUCC-GGUG------------------------------------
-------GAAGUC-CGGGG-------CCGUAGAGGGUGAUAGCCCCGUA---------GGC---G----UAA
C------CCCGGGG-CGA------------------UGGG-CGGGGUAUCCUGAGUACCGCGGGUCG---GA
UAUCCCGCGGGAA-UCUGGCGGGCACC--GACCGCCAAGGCUAAAUA--CGU-C-CCG-AGACCGAUAGC-G
CAC-U-AGUACCGUGAGGGAAAG-CUGAAA-AGUAC-CCCGG-AAGGGG-G-GUG-AAAAGAGCCUGAAACC
G-GGCGGUGAUAU-AAG-GG-UGCG---GCCCGAAA-GGGUAG-----------------------------
------------------------------------------------------------------------
------------------------------------------------------------------------
------------------------------------------------------------------------
------------------------------------------------------------------------
------------------------------------------------------------------------
------------------------------------------------------------AA-CCUCCCCG-
---AA-----------GG-A-------------------A------ACC-----------------GGG---
------------------------------------------------------------------------
--------------------------------------------------------------G--C-GA---
------------------------------------------------------------------------
-------------------------------------------------C-CC------GG--------GAG
UACGA-------------------------------------------------------------------
------------------------------------------------------------------------
------------------------------------------------------------------------
------------------------------------------------------------------------
------------------------------------------------G-----------------------
------------------------------------------------------------------------
------------------------------------------------------------------------
------------------------------------------------------------------------
------------------------------------------------------------------------
------------------------------------------------------------------------
-------------------GGGAGG-GG-----------------ACCAG--G---GUCGCACCGUCCGUCU
UGAAACACGGGCCGGGGAGUCCG-U-CCGAGUGGCGA-GCCUAAGG--G-G--------AUCAU--------
C-CC--CGAA--GGCUCAGGGAAAC-CAA-----------------CAU----------------------G
```

```
CCCG-CAGCCGG----G-------------------------------------------------------
---GGG---CACUCGGGUAGUGAAAGCCGAAAGGAUUCCAGGUUAAUCCUGGGAUCCGAGUAGGUGAGUAGC
UAUCC------GAGUCC--UCGG------------------------------------UCA-GGGGCG-GG
GUCU-----------------------------------GACAA-------------------------GG-
G-C-CC----------------------GGAGUCACUCGGACGG-GACCAGAAGCC-GAGCGAUCUAGGCCG
GGGUAGGGUGAAGCGGGG-CGAAAG-CCCCGUGGAGGCCCGA--AGGGGUGCUGACGUGCAAUUCGC-UCCC
AUGACCCCGGUCUAGGGGCAAAAGACCAAUCAAGCUCGG-CGAUAGCUGGUUCCCCCCGAAG-CUGGUCGCA
GCCAGGCUCGGC-CGGA-------------------------------------------------------
----------------------GG-----UA-GCCGGCGGGGUAGAGC-ACUGAUU-GGA-GGUA-CCGGGG
G-C-GA-AA-GCCC-U-CGGCCUCCUUUCAAACUCCGAACCCGUCG--------------------------
------------------------------------------------------------------------
------------------G-CG-C--CG-UAGA--UGGCCGG-AGACGGG-C-CUAG-G-GGG-UAAGCUCC
UA-G-GCCGAGAG-GGGAAC---AACCCAGACCCG-G-GUUAAGGCCCCAAAGU--GC-CGG-CU-AA---G
-U-GUGAAU--CCAAAGGGCGUCCCCAGCCUCAGACAGUGGGGAGGUAGGCUUAGAAGCAGCCA-UCCUUUA
AAGAGUGCGUAACAGCUCACCCACC-----GAGGCUGGGGGCCCCGAAAAU-GGAC-GGGGCUUA-AGCCGG
C----------CGCCGAG-ACCCGGGGGCACGGC--C-----------------------------------
------------------------------------------------------------------------
-----GCUUUG-------------------------------------------------------------
------------------------------CGGCCGU-GAUC-CGGUAGGGGGGC-GUCCCGA---------
-----------U-G-GGGUA-GAAGCUG--G-GCCGUGAGGU--C-CAGUGGACCC-G-UCG-GGAACGAAG
AUCC-CGGCGG-UAGUA-ACAGCA--AAGUC-GGGUGA--G---------------AAUCCC-GAC-CGCCG
G-AGGGGC-CAGGGUUCC-UCGGCAAUGCGUCGUCAGCCGAGGGUUAGUCGG--UC-CUAAGGCCGGGGGU-
AAC-ACCU---A-CCG--------------------------------------------------------
----------------------------------------GCC-GA-A-AGGGA-AGCAGGUU-UAUA-UUC
CUGCA----------------------------------CCGUG----GGGGUACA----A-UC--------
------------------------------------------------------------------------
-------GCGG--CAAC--------GC-AA-----GC---------------------CCGG-U-UCCC-GAC
GCUUCGG-------------GGUAGG-CCGACCGGG-AC-------UGCC--------------G-UCCGG-
CUAA-CUCCC-CG-----A-A-------------------GC-----CC----------------GGGG-AG
U-G-CC--------------------GUAAU-GGCGAGAACC---------GGGCGA----AGG-GA-GGA-
AUG--GC-----------------------------------------------------------------
-----CUGCC--GUUAUGGCGGGUU-C-GGC-C-G-AUCCCUGGAGCCC-GUGAAA-A-GGG-AACCGGGAA
--------------------------------------------------------GGAUCC-CC-CACGAC
CGUACCGAGAACCGAC------------------------------ACUGGUGC-CCCUAGGU-GAGAAGCC
UCAGGC-GUGU-CGGGCUGAACCCG-G-GCUAGGGAACUCGGC-AAAUUGGCCCCGUAACUUCGG-GAGAAG
GGGUG-CCUG-G-GG--UCUUGGCC-------------------------GG--------------------
------------------------------------------------------------------------
------------------------------------------------------------------------
------------------------------------------------------------------------
------------------------------------------------------------------------
------------------------------------------------------------------------
------------------------------------------------------------------------
------------------------------------------------------------------------
------------------------------------------------------------------------
------------------------------------------------------------------------
```

------------------------------------------------------------------------
------------------------------------------------------------------------
------------------------------------------------------------------------
------------------------------------------------------------------------
----------------------------------------------------------AAGGCU--GGGACC
CCA--GGUCGCAGUGACUAGGGGGGCCUGA-----CUGUUUAAUAAAAACGUAGCUCCCCGCUAGCCCG---
-AA-A-------------GGGUGUGU-ACGGGGGGUGA-GGCCUGGCCAGUGGCG-G-UCCG-UGAAACCCG
GG-------------UCCA-------------ACCGGGUUAAGCGC--CGCUAAAC-GCCGGGAGUAACUAU
GACUCUCUUAAGGUAGCCAAAUGCCUUGCCGGA-UAAGUACCGGCGUGCAUGAACGGCUUAACGAGG--UCC
CCACUGUCCCAGCC----CGG-GGCCCGGUGAACCC---GC---ACCCUGGUGAACAGACCAGGGACCCCCG
ACGGGGAGAGAAGA-CCCCGUGGAGC-UUCACC--GCAGCCU-GGUCCUGGGUUACGGUUGUGGGUGCAUAG
CGU-AGGCGGGAGCGAU-----------GAAUCCGC-CUC----------------------------UCCG
--------------------------------------GG-GGCGGAGGAU-G--C-GC-CCAUGGAACA-C
CGCC-CACCCACGACCGUAACCCUUACCG-GG-------G-CG-A-A-------------------------
------------------------------------------------------------------------
------------------------------------------------------------------------
------------------------------------------------------------------------
------------------------------------------------------------------------
-------------------------------------------------------A----GU----------
------------------------------------------------CCCG-----GGACA-CGGCCAGGU
GGGCGGUUUGGCUGGGGCGGCACGCCCCCGA-AAAGGUAUC-GGGGGCGCCCA-AAGGCCGGCUCAGGCGGG
UCAGAACUCCGCUGUA-GAGUG-CAAGGGCAA-AAGCCGGCCUGACU-GGAC--------------------
--------------CCCGCACAGCAAGGGGUCCAGGGGUGAAA-GCCUGGCCUAGCGAUCCUUC--------
---GU-G-----CCCCAUCG---------------------GU-GGGGGCCGGAGGUGACAG-AA--AAGUU
ACCCCGGGG-GUAACAGGCCCGUCGCGGGCGAGAGUUCACAUCGACCCCGCGGCUUGGUACCUCGAUGUCGG
CUCUUCCCAUCCUGGGGGUGCA-GCAGCCCCCAAGGGUGGGGCUGUUCGCCCAUUAAAGGGGAACGUGAGCU
GGGUUUAGACCGUCGCGAGACAGGUCGGACUCUACCUGUC--------------------G-GGG-------
-------------------------------GUGU--UGGCCGCCUGAGGGUAAGG-UGCCCUAAGUACGAG
AGGAA-CGGGGCGCCGGGGCCUC-UGGU-G-UACCGGUUGUCC-GGCA-GGGCAU-C-GCCGGGCAGCCACG
CCCCA-ACCGAUAAGGGCUGAAAGCA-UC--UAAGCCCGAAGCGGGUCCCGA-A-AAG--AGGCGGCC-GU-
--UCUCCGGCGCAAAUA-------------------------------------------------------
------------------------------------------------------------------------
------------------------------------------------------------------------
------------------------------------------------------------------------
------------------------------------------------------------------------
------------------------------------------------------------------------
---------------------------------------CCCGGGGCGG------GCGA---CCGCAA-CGG
------------------------------------------------------------------------
---------------------GUGCC-----------------------GGGGA----CG-AGG------GC
CCCCGUAGAAGACGGGGU-UGA-UGGGCC-CGG-GGUGUAAG-UCG-G--------AAG-----GC------
UAACGC--------CGACCGAUUGAGCCC--G-CGGG-UCCAAUC---GCCCGAG-----------------
------------------------------------------------------------------------
------------------------------------------------------------------------
------------------------------------------------------------------------
-------------------------------------------------

```
>LSUASG_115547_ODIN/1-3038 KX764886.1 Uncultured archaeon clone ODINRP19_13
------------------------------C----GC--UUAAGCCUUCU-GGUGGAUG-G-CUCGGC-UUG
G-G---AGUCGAUGAAGGCCGUGG--CAAGCUGCGAUAAGCUCCGGGUAGGCGCAUGCAGCCUU------CG
AUC-CGG-AGAUCGCCGAAUGGGACU-U--------CCC-------GC----CAAA-C-CCUAA------UA
A-G-------GUU---------------------UGG----C-A-CUCC-----------------------
-----------C-----------GUA----------------------------------------------
--U--------------------GG-G-AG-GGAGAACCCCCCGAACUGAAGCAUCUUAGUAGGGG--GAGG
AAGAGAAACCAA--UAGGGAUUCCCUGA--GU-AU-GGGCGACUGAAAGGGGA-A-AA-GGGCAAACCGAAU
CCUU-UAAGGA--------AA------------C---UUA-AAGGAGAUGUGGAG--UUC-------------
A-GG--GCC-CCG-CC------------------------------UCUAC--CUC-----------UU--C
U-------GAGAA--GUCG------AAUUCA-UCUG------------------------------------
-------GAA-AG-AUGAA-------CCAUAGAGGGUGAUAGUCCCGUA---------GAC---G----UAA
U------CAGAAAGG-UACUG---------------GGCG-GGUGU--CCCAGAGUACCGUGGGUCG---GU
AUUCCCGCGGAAA-GAUGGGAGGCACC--AACUCCCAACCCUAAAUA--CUA-C-CCA-AGACCGAUAGC-G
CAA-U-AGUACCGUGAGGGAAAG-CUGAAA-AGUAC-UCCGG-UAGGAG-G-GUG-AAAAGUGUCUGAAACC
A-GAAGGUUAAAG-AGG-GG-CGUG---GCCUGAAA-GGGUCG-----------------------------
------------------------------------------------------------------------
------------------------------------------------------------------------
------------------------------------------------------------------------
------------------------------------------------------------------------
------------------------------------------------------------------------
------------------------------------------------------------AG-UCUUCCCG-
---AA-----------GG-A-------------------A--------------------------------
------------------------------------------------------------------------
------------------------------------------------------------------------
------------------------------------------------------------------------
------------------------------------------------------------------------
-ACGA-------------------------------------------------------------------
------------------------------------------------------------------------
------------------------------------------------------------------------
------------------------------------------------------------------------
------------------------------------------------G-----------------------
------------------------------------------------------------------------
------------------------------------------------------------------------
------------------------------------------------------------------------
------------------------------------------------------------------------
------------------------------------------------------------------------
-------------------GGAAGA-GG-----------------ACCAG--G---GUCGCGCCUUCCGUCU
CGAAUCACGGGCCGGGGAGUUUA-U-ACUCGUGGCGA-GUCUAAGG--A-G--------AUCAU--------
C-UC--CGGU--GACGGAGGGAAAC-CGA-----------------UAA----------------------G
CUCG-CAGCCGG----C-------------------------------------------------------
---U--------------------------------------------------------------------
--------------UGA--CCGG------------------------------------UGA-GGAGCA-GG
GUCU-----------------------------------GAAA--------------------------GG-
G-C-CC----------------------GGAGUCACGAGUAUAA-AACCCGAAACC-AGGCGAUCUAUCCCU
```

GAGUAAGUCGAAGCCAAG-GGAAAC-CUUGGUGGAGGACUGA--AGGGGUGCUGACGUGCAAUUCGC-UCCC
AUAACUUGGGGAUAGGGGCAAAAGACCAAUCUAGCCUGG-UGAUAGCUGGUUCCCUCCGAAGUUGGGUCCCA
GCCAAGCCCAGU-CAGA-------------------------------------------------------
----------------------GG-----UA-GCCUGUGAGGUAGAGC-ACUAAUU-GGG-AAGU-UAGGAG
G-C-GC-GA-GCCU-U-CGCUUCCCUUUUAAACUCCGAACUUGCAG--------------------------
------------------------------------------------------------------------
------------------G-CG-C--CG-UAGA--AGACUGG-AGACGGGUU-UCAG-G-GGG-UAAGCUCC
UG-A-ACCGAGAG-GGGAAC---AACCCAGACUGG-G-GUUAAGGUCCCAAAGU--GU-CGG-CU-AA---G
-U-GUAAAA--ACGAAGGGCGUCCGCAUCCUUAGACAGUGGGAAGGUAGGCUUAGAAGCAGCCA-UCCUUUA
AAGAGUGCGUAAUAGCUCACCCACC-----GAGGAUGCGGGCCCCGAAAAU-GGAC-GGGGCUUA-AGCCGG
C----------CACCGAU-ACCCCAGAGCACAAC--A-----------------------------------
------------------------------------------------------------------------
-----UA-----------------------------------------------------------------
-------------------------------UGUUGU-GAUC-UGGUAGGAGGGC-GUUCCGG---------
-----------U-U-GGGUA-GAAGCUG--G-GCCGUGAGGU--C-CAGUGGACCG-G-CCG-GAAGUGCAG
AUCC-CGGUAG-UAGUA-ACAGCG--CAGCC-GAGUGA--G---------------AAUCUC-GGC-CGCCG
A-AAGGGC-AAGGGUUCC-UCGGCGAUGCGUCAUCAGCCGAGGGUAAGUCGG--UC-CUAAGACACUCCGU-
AAC-UCGA---G-AGU--------------------------------------------------------
----------------------------------------GUC-GA-A-AGGGA-AACCGGUU-UAUA-UUC
CGGUA----------------------------------CCAUG----GUGGUACG-------U--------
------------------------------------------------------------------------
-------GUGG--CGAC--------AC-AA-----GC--------------------CUGU-U-UUCC-GAC
GUUUCG--------------GAUAGG-UCGAGCAGG-GU-------UUUC-------------G-CCCUGU-
CUAA-G-CGC-AU-----A-A-------------------CC-----UU----------------GGGG-AG
U-G-CC--------------------GUAAA-GGCGAGAACC----------AAGAUA----AAG-CG-CGA-
AUG--GG-----------------------------------------------------------------
-----CUCCCG-UUAUAGGGGGCUU-C-GAU-C-G-AGUCCGAGAACCC-GUGAAA-A-GGA-AAACAGGAA
--------------------------------------------------------GGAUCC-AC-CAUGCC
CGUACCAAGAACCGAC------------------------------ACAGGUGC-CCCUAGGU-GAGGAGCC
UAAGGU-GUAU-CAGG-GUAAUCUA-G-GCAAGGGAAAUCGGC-AAACUAGCCCCGUACCUUCGG-AAGAAG
GGGUG-CCUG-G-GG--UCUU--------------------------GAGCU--------------------
------------------------------------------------------------------------
------------------------------------------------------------------------
------------------------------------------------------------------------
------------------------------------------------------------------------
------------------------------------------------------------------------
------------------------------------------------------------------------
------------------------------------------------------------------------
------------------------------------------------------------------------
------------------------------------------------------------------------
------------------------------------------------------------------------
------------------------------------------------------------------------
------------------------------------------------------------------------
------------------------------------------------------------------------
----------------------------------------------------------CAGCUC--GGGAUC

```
CUA--GGUCGCAGUGACUAGGGGGUGCUGA-----CUGUUUAAUAAAAACAUAGGUUUCCGCUAGCCCG---
-UA-A-------------GGGUGUGU-ACGGAAGCUGA-AUCCUGGCCAGUGGCG-G-UCCG-UGAAAGCCG
GG-------------UUCA-------------ACCGGAAGAAGCGC--CGCUAAAC-GCCGGGAGUAACUAU
AACUCUCUUAAGGUAGCCAAAUGCCUUGUCGGA-UAAGUACCGACGUGCAUGAAGGGAACAACGAGC--GCC
CCGCUGUCCCUGCC----UAG-AACCUGGUGAACCU---AC---AGCCUAGCGAACAGUCUAGGGACCCCCA
GCGGGAAAAGAAGA-CCCCGUGGAGC-UUUACU--GCAGCUU-ACAGCUGGGGUACGGUUGUGGAUGCAGAG
UGU-AGGUGGGAGCCGUCG----------AUCCUAGCUC-----------------------------UUCG
--------------------------------------GGGGCUAGGGGAG-G--C-GU-CAAUGGAACA-C
CACC-CUUCUAUGACUGUAUCCCUAAUCC-GA--G------GU-U-U-------------------------
------------------------------------------------------------------------
------------------------------------------------------------------------
------------------------------------------------------------------------
------------------------------------------------------------------------
-------------------------------------------------------G----CC----------
------------------------------------------------UCGG-----UGACA-CCUGUAGGU
GGGCAGUUUGGCUGGGGCGGCACGCCUCUGA-AAAGAUAUC-AGAGGCGCCCA-AAGGUCGGCUCAGGCGGG
UCAGAAGUCCGCCGUA-GAGGG-CAAGGCCAA-AAGCCGGCCUGACA-GGAU--------------------
--------------UCUGAAUCGAAAGGAAUCCUGAGUCGAAA-GACUGGCCUAGCGAUCCUUC--------
---AU-G-----UCCCUCUUG--------------------GU-GAGGGCUGGAGGUGUCAG-AA--AAGUU
ACCCCGGGG-AUAACAGAGUCGUCGCGGGCAAGAGUCCACAUCGACCCCGCGGCUUGCUACCUCGAUGUCGG
UUCUUCCUAUCCUGGUGGUGCA-GCAGCCACCAAGGGUGAGGUUGUUCGCCUAUUAAAGGAACGUGAGCU
GGGUUUAGACCGUCGUGAGACAGGUCGGAUUUUAUCUGCU--------------------G-GGG-------
-------------------------------GUGC--UGGCUAUCUGAGGGUAAGG-UGCCUUCAGUACGAG
AGGAA-CGAGGCGCCGCGGCCAC-UAGU-U-UACCGGUUGUCC-GAGC-GGGCAA-U-GCCGGGCAGCCACG
CCGUA-AGCGGUAAGGGCUGAAAGCA-UC--UAAGCCUGAAACGCGGCCCGA-A-AAG--AGAUAGCC-AU-
--UCCCUGAUC-G-------CGGG-------------------------------------------------
-----------------------------------------------------------------------G
UAA---------------------------------------------------------------------
-------CCCG-------------------------------------------------------------
------------------------------------------------------------------------
------------------------------------------------------------------------
---------------------------------------ACCAUAAUAG------GCGA---CUAUUAUGGU
----------------------------U-------------------------------------------
---------------------GAUC------------------------GGGGA----CG-AGG------GC
UCCGGUAGAAGACCGGGU-UGA-UAGAGC-GGG-UGUGUACG-CUA-G--------AAG-----GC------
-UUCGG-------CCGACUAGCUUAGCAU--G-CCGC-UACUAAUC---GCCCGAGC---------------
------------------------------------------------------------------------
------------------------------------------------------------------------
------------------------------------------------------------------------
-------------------------------------------------
>LSUASG_1936271_ODIN/1-3051 MDVT01000007.1, OdinLCB4_contig000007, whole ge
------------------------CAUGUGC----GC--UUAAGCCUUCU-GGUGGAUG-G-CUAGGC-UUG
G-G---AGUCGAUGAAGGCCGUGG--CAAGCUGCGAUAAGCUCCGGGUAGGCGCAUGCAGCCUU------CG
AUC-CGG-AGAUCGCCGAAUGGGACU-U--------CCC-------GC----CAAG-C-CCUAA------UA
A-G-------GUU--------------------UGG----C-A-CUCC------------------------
```

```
-----------C-----------UUA----------------------------------------------
--U--------------------GG-G-AG-GGAGAACCCCCCGAACUGAAGCAUCUUAGUAGGGG--GAGG
AAGAGAAACCAA--UAGGGAUUCCCUGA--GU-AG-GGGCGACUGAAAGGGGA-A-AA-GGGCAAACCGAAU
CCUU-UAAGGA--------AA------------C---UUA-AAGGAGAUGUGGAG--UUC------------
A-GG--GCC-UCG-CC------------------------------UCUAC--CUC-----------UU--C
U-------GAGAA--GUCG------AAUUCA-UCUG------------------------------------
-------GAA-AG-AUGAA-------CCAUAGAGGGUGAUAGUCCCGUA---------GAC---G----UAA
U------CAGAAAGG-UAUUU---------------GGUG-GGUGU--CCCAGAGUACCGUGGGUCG---GU
AUUCCCGCGGAAA-GAUGGGAGGCACC--AACUCCCAACCCUAAAUA--CUA-C-CCA-AGACCGAUAGC-G
CAA-U-AGUACCGUGAGGGAAAG-CUGAAA-AGUAC-UCCGG-UAGGAG-G-GUG-AAAAGCGUCUGAAACC
A-GAAGGUUAAAG-AGG-GG-CGUG---GCCUGAAA-GGGUAG-----------------------------
------------------------------------------------------------------------
------------------------------------------------------------------------
------------------------------------------------------------------------
------------------------------------------------------------------------
------------------------------------------------------------------------
------------------------------------------------------------AG-UCUUCCCG-
---AA-----------GG-A-------------------A--------------------------------
------------------------------------------------------------------------
------------------------------------------------------------------------
------------------------------------------------------------------------
------------------------------------------------------------------------
-ACGA-------------------------------------------------------------------
------------------------------------------------------------------------
------------------------------------------------------------------------
------------------------------------------------------------------------
------------------------------------------------G-----------------------
------------------------------------------------------------------------
------------------------------------------------------------------------
------------------------------------------------------------------------
------------------------------------------------------------------------
------------------------------------------------------------------------
-------------------GGAAGA-GG-----------------ACCAG--G---GUCGCGCCUUCCGUCU
CGAAUCACGGGCCGGGGAGUUUA-U-AAUCGUGGCGA-GUCUAAGG--A-G--------UUCAU--------
C-UC--CGGU--GACGAAGGGAAAC-CGA-----------------UAA----------------------G
CUCG-CAGCCGG----C-------------------------------------------------------
---U--------------------------------------------------------------------
--------------UGA--CCGG------------------------------------UGA-GGAGCA-GG
GUCU-----------------------------------GAAA--------------------------GG-
G-C-CC----------------------GGAGUCACGAUUAUAA-AACCCGAAACC-AGGCGAUCUAUCCCU
GAGUAAGUCGAAGCCAAG-GGAAAC-CUUGGUGGAGGACUGA--AGGGGUGCUGACGUGCAAUUCGC-UCCC
AUAACUUGGGGAUAGGGGCAAAAGACCAAUCUAGCCUGG-UGAUAGCUGGUUCCCUCCGAAGUUGGGUCCCA
GCCAAGCCCAGU-CAGA-------------------------------------------------------
----------------------GG-----UA-GCCUGUGAGGUAGAGC-ACUAAUU-GGG-AAGU-UAGGAG
G-C-GC-GA-GCCU-U-CGCUUCCCUUUUAAACUCCGAACUUGCAG--------------------------
```

```
------------------------------------------------------------------------
------------------G-CG-C--CG-UAGA--AGACUGG-AGACGGGUU-UCAG-G-GGG-UAAGCUCC
UG-A-ACCGAGAG-GGGAAC---AACCCAGACUGG-G-GUUAAGGUCCCAAAGU--GU-CGG-CU-AA---G
-U-GUAAAA--ACGAAGGGCGUCCGUAUCCUUAGACAGUGGGAAGGUAGGCUUAGAAGCAGCCA-UCCUUCA
AAGAGUGCGUAAUAGCUCACCCACC-----GAGGAUGCGGGCCCCGAAAAU-GGAC-GGGGCUAA-AGCCGG
C----------CACCGAU-ACCCCAGAGCACAAC--G-----------------------------------
------------------------------------------------------------------------
-----UA-----------------------------------------------------------------
-------------------------------CGUUGU-GAUC-UGGUAGGAGGGC-GUUCCGG---------
-----------U-U-GGGUA-GAAGCUG--G-GCCGUGAGGU--C-CAGUGGACCG-A-UCG-GAAGUGCAG
AUCC-CGGUAG-UAGUA-ACAGCG--CAGCC-GAGUGA--G---------------AAUCUC-GGC-CGCCG
A-AAGGGC-AAGGGUUCC-UCGGCGAUGCGUCAUCAGCCGAGGGUAAGUCGG--UC-CUAAGACACUCCGU-
AAC-UCGA---G-AGU--------------------------------------------------------
----------------------------------------GUC-GA-A-AGGGA-AACCGGUU-UAUA-UUC
CGGUA----------------------------------CCAUG----GUGGUACG-------U--------
------------------------------------------------------------------------
-------GUGG--CGAC--------AC-AA-----GC--------------------CUGU-U-UUCC-GAC
GUUUCG--------------GAUAGG-UCGAGCAGG-GU-------CUUC-------------G-CCCUGU-
CUAA-G-CGC-AU-----A-A-------------------CC-----UU----------------GGGG-AG
A-G-CC--------------------GUAAC-GGCGAGAACC---------AAGAGA----AAG-CG-CGA-
AUG--GG-----------------------------------------------------------------
-----CUCCCG-UUAUAGGGGGCUU-C-GAU-C-G-AGUCCGAGAACCC-GUGAAA-A-GGA-AAACAGGAA
--------------------------------------------------------GGAUCC-AC-CAUGCC
CGUACCAAGAACCGAC------------------------------ACAGGUGC-CCCUAGGU-GAGGAGCC
UAAGGU-GUAU-CAGG-GUAAUCUA-G-GCAAGGGAAAUCGGC-AAAUUAGCCCCGUACCUUCGG-AAGAAG
GGGUG-CCUG-G-GG--UCUU--------------------------GAGCU--------------------
------------------------------------------------------------------------
------------------------------------------------------------------------
------------------------------------------------------------------------
------------------------------------------------------------------------
------------------------------------------------------------------------
------------------------------------------------------------------------
------------------------------------------------------------------------
------------------------------------------------------------------------
------------------------------------------------------------------------
------------------------------------------------------------------------
------------------------------------------------------------------------
------------------------------------------------------------------------
------------------------------------------------------------------------
----------------------------------------------------------UAGCUC--GGGAUC
CUA--GGUCGCAGUGACUAGGGGGUGCUGA-----CUGUUUAAUAAAAACAUAGGUUUCCGCUAGCCCG---
-UA-A-------------GGGUGUGU-ACGGAAGCUGA-AUCCUGGCCAGUGGCG-G-UCCG-UGAAAGCCG
GG-------------UUCA-------------ACCGGAAGAAGCGC--CGCUAAAC-GCCGGGAGUAACUAU
AACUCUCUUAAGGUAGCCAAAUGCCUUGUCGGA-UAAGUACCGACGUGCAUGAAGGGAACAACGAGC--GCC
CCGCUGUCCCUGCC----UAG-AACCUGGUGAACCU---AC---AGCCUAGCGAACAGUCUAGGGACCCCCA
```

```
GCGGGAAAAGAAGA-CCCCGUGGAGC-UUUACU--GCAGCUU-ACAGCUGGGGUACGGUUGUGGAUGCAGAG
UGU-AGGUGGGAGCCGUCG----------AUCUUAACCUC----------------------------UUCG
--------------------------------------GGGGUUAAGGGAG-G--C-GC-CAAUGGAACA-C
CACC-CUUCUAUGACUGUAUCCCUAAUCC-GA--G------GU-U-C-------------------------
------------------------------------------------------------------------
------------------------------------------------------------------------
------------------------------------------------------------------------
------------------------------------------------------------------------
-------------------------------------------------------G----CC----------
------------------------------------------------UCGG-----UGACA-CCUGUAGGU
GGGCAGUUUGGCUGGGGCGGCACGCCUCUGA-AAAGAUAUC-AGAGGCGCCCA-AAGGUCGGCUCAGGCGGG
UCAGAAGUCCGCCGUA-GAGGG-CAAGGCCAA-AAGCCGGCCUGACA-GGAU---------------------
--------------UCUGAAUCGAAAGGAAUCCUGAGUCGAAA-GACUGGCCUAGCGAUCCUUC--------
---AU-G-----UCCCUCUUG--------------------GU-GAGGGCUGGAGGUGUCAG-AA--AAGUU
ACCCCGGGG-AUAACAGAGUCGUCGCGGCAAGAGUCCACAUCGACCCCGCGGCUUGCUACCUCGAUGUCGG
UUCUUCCUAUCCUGGUGGUGCA-GCAGCCACCAAGGGUGAGGUUGUUCGCCUAUUAAAAGGGAACGUGAGCU
GGGUUUAGACCGUCGUGAGACAGGUCGGAUUUUAUCUGCU--------------------G-GGG-------
-------------------------------GUGC--UGGCUAUCUGAGGGUAAGG-UGCCUUCAGUACGAG
AGGAA-CGAGGCGCCGCGGCCAC-UAGU-U-UACCGGUUGUCC-GAGC-GGGCAA-U-GCCGGGCAGCCACG
CCGUA-AGCGGUAAGGGCUGAAAGCA-UC--UAAGCCUGAAACGCGGCCCGA-A-AAG--AGAUAGCC-AU-
--UCCCUGAUC-G------CGGG-------------------------------------------------
-----------------------------------------------------------------------U
UAA---------------------------------------------------------------------
-------CCCG-------------------------------------------------------------
------------------------------------------------------------------------
------------------------------------------------------------------------
---------------------------------------ACCAUAAUAG------GCGA---CUAUUAUGGU
----------------------------U-------------------------------------------
---------------------GAUC------------------------GGGGA----CG-AGG------GC
UCCGGUAGAAGACCGGGU-UGA-UAGAGC-GGG-UGUGUACG-CUA-G--------AAG-----GC------
-UUCGG-------CCGACUAGCUUAGCAU--G-CCGC-UACUAAUC---GCCCGAGCG-UCGUGC-------
------------------------------------------------------------------------
------------------------------------------------------------------------
------------------------------------------------------------------------
-------------------------------------------------
>LSUASG_KP308697.1/2181-5234 Uncultured archaeon clone RBG_16_scaffold_3034
---------------------G-ACAUG-----------CCACGCCAUCU-GGGGGAUA-G-CUUGGC-UCG
G-A---CGCCGAAGAAGGGCGUAG--CAAGCCGCGAUAGGCCUCGGGGAGGUGCAUGCAACCUU------CG
AUC-CGA-GGAUCCCGAAUAGGACU-U---------CCA-------GC----CGCG-G-CU---------UC
G-G-------CCG---------------------CGG----C-G-CUCC-----------------------
-----------C-----------ACA----------------------------------------------
--U--------------------GG-G-AG-CGGGAACCCCCUGAAUGGAAACAUCUUAGUAAGGG--GAGG
AAGAGAAACCAA--UCGGGAUUCCCCUA--GU-AG-CAGCGAGCGAAAAGGGA-A-CA-GUUCAAACCGAAU
CCCG-CGGGGA--------GA------------C---CCG-CGGGGGAUGUGGUG--UUG------------
U-GG--GCC-UGG-CC------------------------------UCCUC--CUA-----------AC--C
```

```
G-------ACGAA--GCCG------AAGUGA-CCUG------------------------------------
-------GAA-CG-GUCCG-------CCGCAUAGGGUGAUAGCCCCGUA---------GGC---G----UAA
G------UCGGAAGG-AGUUG---------------GGCU-GG-GU--UCCUGAGUACCAAGCCUCG---AU
UUUGGUUUGGGAA-GCUGGGAGGCAUC--AACUCCAAAACUAAACA--CGU-U-CCG-AGACCGAUAGU-G
CAC-C-AGUACGGUGACGGAAAG-CUGAAA-AGUGC-CCCGG-AAGGGG-G-GUU-AAAAGCGCUUGAAACC
A-GAUGGUUAUAG-ACG-UG-CAUG---GCUCAUAA-GGGAAG-----------------------------
------------------------------------------------------------------------
------------------------------------------------------------------------
------------------------------------------------------------------------
------------------------------------------------------------------------
------------------------------------------------------------------------
------------------------------------------------------------AU-UCCACCCG-
---AA-----------GG-AUGCCGUAGCGAUGCGGCGGU--------------------------------
------------------------------------------------------------------------
------------------------------------------------------------------------
------------------------------------------------------------------------
------------------------------------------------------------------------
-ACGA-------------------------------------------------------------------
------------------------------------------------------------------------
------------------------------------------------------------------------
------------------------------------------------------------------------
------------------------------------------------G-----------------------
------------------------------------------------------------------------
------------------------------------------------------------------------
------------------------------------------------------------------------
------------------------------------------------------------------------
------------------------------------------------------------------------
-------------------GGUGGA-GG-----------------ACUAG--G---GUCAUGCGAUCAGUCU
CGAAGCACUGGCCGGGGAGUUCA-C-UGUCAUGGCGA-GCUUAAGG--G-U--------GUUAA--------
G-CC--CGGA--GGCGAAGGGAAAC-CGA-----------------CAG----------------------G
CCCA-CAA-CGG----G-------------------------------------------------------
---G--------------------------------------------------------------------
--------------CAA--CCUG------------------------------------CGA-GGGGCG-GA
GUCU-----------------------------------GAAA--------------------------GG-
G-C-UC----------------------GGAGUCAUGGCGGUGA-GAAUAGAAACC-AGACGAUCUAGUCCU
GGGCAAGGCGAAGCCGGG-UGAAAA-CCCGGUGGAAGCCUGA--AGGGGUUCUGACGUGCAAUUCGA-UCCU
UUGACCUGGGAUUAGGGGCGAAAGACCAAUCUAGUCUGG-UGAUAGCUAUUUCCCUCCGAAG-UAUAUCGCA
GUAUAGUCUCGG-GCGA-------------------------------------------------------
----------------------GG-----UG-GCCGAUGGGGUAGAGC-ACCGAUU-GGG-AGAC-AAGGAC
U-C-GA-AA-GGGU-U-CACCUUCCUAUCAAACUCCGAAUCUGUCG--------------------------
------------------------------------------------------------------------
------------------G-CG-C--UG-AAGA--CCCCGGG-GGACGGUA-CUAG-G-GCG-UAAGGUCC
UA-AGGCCGAGAG-GGAAGC---AACCCAGACCGA-G-GCUAAGGUCCCUAAAU--GC-UGA-CU-AA---G
-U-GUGAGG--UAGAAGGGCGUCCUCAGCCAUAGACAGUGGGGAGGUAGGCUUAGAAGCAGCCA-CCCUUCA
AAGAAAGCGUAACAGCUCACCCACC-----GAGGCUGGGGGCCCCGAAAAU-AAAC-GGGACUCA-AGUCAG
```

```
C----------UACCGAG-CCCUCGGGGCACACCCGA-----------------------------------
------------------------------------------------------------------------
-----AG-----------------------------------------------------------------
-------------------------------GGGUGU-GAUC-CGGUAGGAGGGC-GUUCCGC---------
-----------U-U-GGGGA-GAAGCUG--G-GCCGUGAGGU--C-CAGUGGACCG-G-GCG-GAAUUGUAG
AUCC-CGGUGG-AAGUA-ACAGCA--UAGUC-AGGUGA--G---------------AAUCCU-GAC-CGCCG
A-AAGGGC-CAGGGUUCC-CCGGCGAUGCGUCAUCAGCCGGGGGUAAGUCGA--UC-CUAAGGCCGACCUU-
AAC-UGGG---U-AG---------------------------------------------------------
----------------------------------------GCC-GA-A-GGGGA-AACCGGUU-AAGA-UUC
CGGUA----------------------------------CCACG----GGGGUACA-------U--------
------------------------------------------------------------------------
-------GCGG--CAAC--------GC-AG-----GC--------------------CCUG-C-GUCU-GAC
GCUUCAG-------------GGUAGG-CUGAACGGG-GC-------CGUC-------------G-CCCCGU-
CUAA-C-CGC-UU-----A-A-------------------GC-----CU----------------GGGG-AG
U-G-CC--------------------GUAAU-GGCGAGAACC----------AGGCGA----AGGCCG-GGA-
AUG--GC-----------------------------------------------------------------
-----CUCCCG-UU-UAGGGGGGUU-C-AGC-C-G-AUCCCCGGAGCCA-GUGAAA-A-GGG-CGCAGGGAG
--------------------------------------------------------GGAUCC-UC-CGCGAC
CGUACCUAGAACCGAC------------------------------ACAGGUGU-CCCUAGGC-GAGAAACC
UAAGGC-GUAU-CGGGCAUAAUUCGUG-GCGAGGGAACUCGAC-AAAUUAGCCCCGUAACUUUGG-GAUAAG
GGGUG-CCUA-U-GG--UCUU--------------------------GGUCC--------------------
------------------------------------------------------------------------
------------------------------------------------------------------------
------------------------------------------------------------------------
------------------------------------------------------------------------
------------------------------------------------------------------------
------------------------------------------------------------------------
------------------------------------------------------------------------
------------------------------------------------------------------------
------------------------------------------------------------------------
------------------------------------------------------------------------
------------------------------------------------------------------------
------------------------------------------------------------------------
------------------------------------------------------------------------
----------------------------------------------------------UUGACU--GGGACC
GUA--GGUCGCAGUGGCAAAGGGGUGCCGA-----CUGUUUAAUAAAAACACAGGGGGCUGCUAGCGUG---
-AA-A-------------ACGUGUGU-ACAGCCCUUGA-AUCCUGGCCAGUGGCG-G-GAUC-UUAAACCUG
GU-------------UUCA-------------ACUGGGCGAAGGAC--CGCUAAAC-GCCGGGAGUAACUCU
GACUCUCUUAAGGUAGCCAAAUGCCUUGUCGGG-UAAGUUCCGACGUGCAUGAAUGGAUCAACGAGC--ACC
CCACUGUCCCCGCC----CGG-AGCCCGGUGAAACC---ACAUGACUCAGACGAACAGUCUGGGAUUUCCCA
GCGGGAAAAGAAGA-CCCCGUGGAGU-UUUACU--GUAGCUU-GUUGCUGGGAUGCGGCUGUCGUUGCAUAG
CGU-AGGCGGGAGCCGUCG----------AGGCCAGUCCC----------------------------UUCG
--------------------------------------GGGGCUGGCGGAG-G--C-GC-CAAUGGAACA-C
CGCC-CUUCGGCAGCUGCAUCUCUUACCG-GG--G------CG-A-A-------------------------
------------------------------------------------------------------------
```

```
------------------------------------------------------------------------
------------------------------------------------------------------------
------------------------------------------------------------------------
-------------------------------------------------------U----GC----------
------------------------------------------------UCCG-----GGACA-ACGGCAGGU
GGGCAGUUUGGCUGGGGCGGCACCCCCUUGA-AAACGAAUC-AAGGGGGCCCA-AAGGUCGGCUCAGGUGGG
ACAGAAACCCAUCGUA-GAGGG-CAAAGCCAA-AAGCCGGCCUGACU-GGAC--------------------
--------------CCUCCAAGGCAAGGGGUCCAGGAGCGAAA-GCUGGGCUUAGCGAUCUACC--------
---GA-G-----CCCCCAUUG--------------------AU-GGGGGCCGAUAGUGACAG-AA--AAAUU
ACCCCGGGG-GUAACAGGCUCGUCGCGGGUGAGAGCUCCCAUCGACCCCGCGGUUUGGUACCUCGCCGUCGG
CUCUUCCCAUCCUGGCGGUGCA-GCAGCCGCCAAGGGUAGGGCUGUUCGCCCAUUAAAGGGGAACGUGAGCU
GGGUUUAGACCGUCGUGAGACAGGUCGGAUUUUAUCUGCU--------------------G-GGA-------
-------------------------------AUGC--UGGUCGCUUGAGGGGAAGG-UGUCCUUAGUACGAG
AGGAA-CGGGGCGCCGUAGCCUA-CGGU-A-UACCGGUUGUCU-GAUA-AGGCAC-U-GCCGGGCAGCUGCG
CUAUA-UUGGAUAAGGGCUGAAUGCA-UC--UAAGCCCGAAACCAUACCCGA-A-AAU--AAGCGACC-AU-
--CGACAA-U-GA-U---------------------------------------------------------
------------------------------------------------------------------------
------------------------------------------------------------------------
------------------------------------------------------------------------
------------------------------------------------------------------------
------------------------------------------------------------------------
-----------------------------------------GAGAUUUG------GAAA----CAGG-UCUC
----------------------------U-------------------------------------------
---------------------AGUU------------------------GUGGA----CU-AGG------AC
UGGAAGAGAAGAUUCCGU-UGA-UGGAGU-GGA-GGUGUACG-CUG-G--------AAG-----GC------
-UUCGG-------CCGACCAGUUUAGCCU--G-CCAC-CCCCAAUC---GUCCAAGG--UGUUGGG------
------------------------------------------------------------------------
------------------------------------------------------------------------
------------------------------------------------------------------------
-----------------------------------CU------------
>LSUASG_KP308663.1:317300-320372/1-3073 Uncultured archaeon clone RBG_16_sc[illegible]
-----------------------ACCUGAAC----AC-GCAACGCUAUUU-GGUGAAUG-G-CUUGGC-UCG
G-G---AGCUGAAGAAGGGCGUGG--CAAGCUGCGAUAAGCCCAGGGUAGGCGCAUGCAGCCUU------AG
AUC-CUG-GGAUCCCCGAAUGGGACU-U--------CCUAUCAUUGGUCCG-CAAG-G----------A---
--------------------CU---C----G---UGA----U-G-UUCC-----------------------
-------------------U---UCA----------------------------------------------
-----------------------UG-G-AA-CGGGCACCUUCCGAACGGAAGCAUCUUAGUAGGAG--GAGG
AAGAGAAACCAA--UUGGGAUUCCCUUA--GU-AG-CGGCGAGCGAAAAGGGA-A-GA-GUUCAAACCGAAC
CCUG-CAGGGA--------AA------------C---UUG-CAGGGGAUGUAGUG--UAG------------
UAGG--GCC-UGG-CC------------------------------UCCGC--CUC-----------AC--C
G-------UCGAA--GCCG------AAGUGG-CUUG------------------------------------
-------GAA-CA-GCCUG-------CCAUAGAAGAUGACAGCUCUGUA---------GGC---G----UAA
G------ACGGUAGG-AGAUG---------------GGCU-UGUGU--CCCUGAGUAUUAUGCCUUG---GU
UUUGGCGUAAGAA-GCUGGGAGGCAUC--AACUUCCAAAGCUAAAUA--CAU-C-CCG-AGACCGAUAGC-G
CAC-U-AGUACGGUGACGGAAAG-CUGAAA-AGUAC-CCCGA-GAGGGG-G-GUA-AAAAGUGCCUGAAGCC
```

```
A-AAUAGUUAUAG-ACG-UG-CGCG---GCUUGAAA-GGGUGG-----------------------------
------------------------------------------------------------------------
------------------------------------------------------------------------
------------------------------------------------------------------------
------------------------------------------------------------------------
------------------------------------------------------------------------
------------------------------------------------------------AU-CCCUUCCG-
---AA-----------GG-A-------------------A------UUC-----------------GUG---
------------------------------------------------------------------------
--------------------------------------------------------------G--C-AA---
------------------------------------------------------------------------
-------------------------------------------------C-GC------GA--------UAG
UACGC-------------------------------------------------------------------
------------------------------------------------------------------------
------------------------------------------------------------------------
------------------------------------------------------------------------
------------------------------------------------A-----------------------
------------------------------------------------------------------------
------------------------------------------------------------------------
------------------------------------------------------------------------
------------------------------------------------------------------------
------------------------------------------------------------------------
-------------------GGAAGGGGG-----------------ACCAG--A---GUCGCGCGAUCCGUUU
AGAAAUACGGGCCGGGGAGGUUA-C-GGUUGUGGCGA-GUUUAAGA--G-U--------UCAAA--------
-GCU--CGGA--GGCGAAGGGAAAC-CGA-----------------CA-----------------------G
CCCG-CAGCCAG------------------------------------------------------------
------------------------------------------------------------------------
------------CUCGU--CUGG------------------------------------UAA-GGGGCG-GG
GUCU-----------------------------------GAAUA-------------------------GG-
G-C-CU----------------------GGAGCCACAACAGUAUUUACUAGAAACC-GGACGAUCUAGCCCU
GGGCAAGGUGAAGCCGGG-UGAAAA-CUCGGUGGAGGCCUGA--AAGGGUUCUGAUGUGCAAUUCGU-UCCC
CUGACCUGGGGCUAGGGGCCAAAGACCAAUCUAGUCCGG-UGAUAGCUAGUUCCCUCCGAAG-UAGGUCGCA
GCCUAGUCUUCC-CAGA-------------------------------------------------------
----------------------GG-----UA-GCCAGCGGGGUAGAGC-ACUGAUU-AGG-GAGU-AAGGUU
C-C-GA-AAGGAAC---CGCUUCCUUCUCAAACUCCGAAUCUGCUG--------------------------
------------------------------------------------------------------------
------------------G-CG-C--CG-UAGA--AGGGAGG-AAGCGGGUA-ACGG-G-GGG-UAAGCCCA
CG-U-GCCGAGAA-GGGAAC---AACCUAGACUGA-G-GUUAAGGCCCCCAAAU--GC-UGG-CU-AA---G
-U-GUCAAUC-AAGAAGGGCGUCCUCUGCCUAAGACAGUGGGGAGGUGGGCUCAGAAGCAGCCA-UCCUUCA
AAGAGUGCGUAACAGCUCACCCACC-----GAGGUCGAGGGCCCCGAAAAU-GGAC-GGGACUA--AGUCAG
C----------UGCCGAG-ACCUCAGGGUACAGACUC---------------------A-------------
------------------------------------------------------------------------
------------------------------------------------------------------------
------------------------------CGUCUGU-AAUC-CGGUAGGAGGGC-GUCCCAG---------
-----------U-U-GGGUA-GAAGCAG--G-GCUGUGAGGU--C-CUGUGGACCG-G-CUG-GGAUUGAAU
```

```
AUCC-CGGUGG-UAGUA-ACAGCA--AAAUC-AGGUGA--G---------------AAUCCU-GAU-CGCCG
A-AAGGGC-CAGGGUUCC-CCGGCAAUGCGUCGUCAGCCGGGGGUAAGUCGA--UC-CUAAGGUGGAACUU-
AAC-UGUA-----UCC--------------------------------------------------------
----------------------------------------GCC-GA-A-AGGGA-AACUGGUU-AAUA-UUC
CAGUA----------------------------------CCUUA----GAGGUACAC---------------
------------------------------------------------------------------------
-------GCGG--CAAC--------GC-AA-----GC--------------------UCGA-C-AGCC-GAC
GCUUCGA-------------GGUAGG-UCGAACAAG-GU--------CAUC-------------G-CCUUGU-
CUAA-C-UGU-CU-----A-A-------------------GU-----CU----------------AGGG-AG
U-G-CC--------------------GUAAU-GGCGAGAACU---------AGACGA----AUA-UA-GGA-
AUG--GC-----------------------------------------------------------------
-----CGCCCG-UCUAUGGGCGGUU-U-GGC-U-G-AUCCUCGGAGCCA-AUGAAA-A-AGU-UGUCGAGAU
--------------------------------------------------------GGAUCC-UC-UAAGAU
CGUACCUAGAACCGAC------------------------------ACUGGUGA-CCCUAGGU-GAGAAGCC
UAAGGC-GUGA-CAGGUAGACUUGA-G-GCGAGGGAACUCGGC-AAUCUAGCCCCGUAACUUCGG-GAUAAG
GGGUG-CCUA-C-GA--UUUUGGCG-----------------------------------------------
------------------------------------------------------------------------
------------------------------------------------------------------------
------------------------------------------------------------------------
------------------------------------------------------------------------
------------------------------------------------------------------------
------------------------------------------------------------------------
------------------------------------------------------------------------
------------------------------------------------------------------------
------------------------------------------------------------------------
------------------------------------------------------------------------
------------------------------------------------------------------------
------------------------------------------------------------------------
------------------------------------------------------------------------
----------------------------------------------------------CAAGCU--GGAAUC
AUA--GGUCGCAGUGACAAAGGGGUCCUGA-----CUGUUUAAUAAAAACAUAGGUGACUGCAAGUCCG---
-AA-A-------------GGAUGUGU-ACAGUCGCUGA-AUCCUGGCCAGUGGCG-G-CAUC-UGAAAUCUG
GU-------------UUCA-------------ACCGGGCUAAGGAC--CGCUAAAC-GCCGGGAGUAACUCU
GACUCUCUUAAGGUAGCCAAAUGCCUUGUCGGG-UAAGUUCCGACGCGCAUGAAUGGAUCAACUAGG--GCC
CCGCUGUCCCCGCC----UCG-AACCUGGUGAACCC-ACAU---AACCUGGAGAACGGUCCAGGAAAUCCCA
GCGGGAUAAGAAGA-CCCCGUGGAGU-UUUACU--GCAGCUU-GUCGCUGGGAUAUGGCCGCGGAUGCAGAG
UGU-AGGCGGGAGCCGUUA----------AUCCCGAUCUC----------------------------UCCG
--------------------------------------GGGGUCGGGGGAG-G--C-AA-CAAUGGAACA-C
CGCC-CUCUUGUGGCUAUGUCUCUUACUC----UAGC----GU---A-------------------------
------------------------------------------------------------------------
------------------------------------------------------------------------
------------------------------------------------------------------------
------------------------------------------------------------------------
-----------------------------------------------------------UGCUUCU------
---------------------------------------------------------GGACA-ACGGCAGGU
```

```
AGGCAGUUUGGCUGGGGCGGCACCCCCUUGA-AAAGGUAUC-GAGGGGGCCCA-AAGGUUGGCUCAGGUGGG
ACAGAAACCCACUGUA-GAGGG-CAAGGCCAA-AAGCCAGCUUGACU-GGAU--------------------
--------------CCUCCACAGUAAGGGAUCCAGCGGCGAAA-GCCUGGCCUAGCGAUCCUUC--------
---GU-G-----UCCUCCUUG--------------------GU-GGGGGCCGGAGGUGACAG-AA--AAAUU
ACCCCGGGG-AUAACAGGCUCGUCGCGGGCGAGAGUUCCUAUCGACCCCGCGGUUUGGUACCUCGUCGUCGG
CUCUUCCCAUCCUGGCGGUGCA-GCAGCCGCCAAGGGUAGGGCUGCUCGCCCAUUAAAGGGGAACGUGAGCU
GGGUUUAGACCGUCGUGAGACAGGUCGGACUUUAUCUGCU--------------------G-GGA-------
-------------------------------UUGC--UGGUCGCUUGAGGGGAAGC-UGUCCCUAGUACGAG
AGGAA-CGGGGUGGCGCGGCCUC-UAGU-U-UACCGGUUGUUC-GAUA-GAGCAU-U-GCCGGGCAGCCACG
CCGUA-GGGGAUAAGAGCUGAAAGCA-UC--UAAGCUCGAAACCCCUCCCGA-A-AAU--AGGCGGCC-AU-
--UCUUUGACACC-UAAUC-----------------------------------------------------
------------------------------------------------------------------------
------------------------------------------------------------------------
------------------------------------------------------------------------
------------------------------------------------------------------------
------------------------------------------------------------------------
----------------------------------------UGAG-CUUG------GUAA----CGAGCCCGG
----------------------------A-------------------------------------------
---------------------GGUC------------------------AAGGA----CG-AGG------GC
UCCCGUAGAAGACGGGUU-UGA-UGGAGC-GGG-GGUGUACG-CAU-C--------AAG-----GC------
-UUCGG-------UCGAGAUGUUCAGCCC--G-CCGC-CACCAAAC---GCCCAAGAUGUCAGGUCG-----
------------------------------------------------------------------------
------------------------------------------------------------------------
------------------------------------------------------------------------
-------------------------------------------------
>LSUASG_QMYW01000148.1:199-3284/1-3085 Candidatus Lokiarchaeota archaeon is
---------------------------------------------UGCAU-AGUGGAUG-U-CUCGGC-UUA
G-U---AUUCGAAGAAGGGCGUGA--CAAGCUGCGAUAAGCUCCGGGUAGGUGCACGUAACCGU------UG
AUC-CGG-AGAUUCCCGAAUGUGACA-UC-------------------------------------------
------------------------------------A----U-G-GUUA-GGGUAA---------------U
GCC--------U-----------UUC----------------------------------------------
--CAUCCCC----------GUCAGG-G-GA-GGGGAACGCGCUGAAGCGAAGCAUCAUAAUAAGCG--CAGG
AAAAGAAAGCAA--UAGCGAUUCCGUGA--GU-AG-CGGCGAGCGAAAACGGA-A-GA-GGCCAAACCGAAU
UCUC-CGAUGA--------AA------------G---UCG-GCGGAGAUGUGGAG--UGU------------
G-GA--CUCGUGU-CGAGC----------------------------------CUC-----------CG--U
G-------AAGAA--GUUG------AACCUU-UCUG------------------------------------
-------GAA-UG-UUAGA-------CCAUAGAGGGUGACAGUCCCGUA---------AAC---G----UAA
A------UCACAAGG-CUUA----------------UGGC-GAGUA--UCCCGAGUAGCGCGGGAUG---GU
UGUUCCGUGUGAA-UGUGCCUGGUACC--GACAGGCAAGCCUAAAUA--UAA-C-CUA-AGACCGAUAGC-G
CAU-U-AGUACUGUGAAGGAAAG-AUGAAA-AGCAC-CCAGG-AAUGGU-U-GUG-AAAAGUGCCUGAAACU
A-UGCAG-UGAAG-AGU-GA-AGCA---GUCUGAAA-GAAAUA-----------------------------
------------------------------------------------------------------------
------------------------------------------------------------------------
------------------------------------------------------------------------
------------------------------------------------------------------------
```

```
------------------------------------------------------------------------
------------------------------------------------------------AU-GCUUAUCG-
---AAGUUCGUACACGUG-A-------------------G-------UG-----------------UG----
------------------------------------------------------------------------
-------------------------------------------------------------------GU---
------------------------------------------------------------------------
-------------------------------------------------A-AC------GA--------GAU
G-----------------------------------------------------------------------
------------------------------------------------------------------------
------------------------------------------------------------------------
------------------------------------------------------------------------
------------------------------------------------------------------------
------------------------------------------------------------------------
------------------------------------------------------------------------
------------------------------------------------------------------------
------------------------------------------------------------------------
------------------------------------------------------------------------
-----------------------AGCGG-----------------AUCAG--G---AUUGCUUCUUUAUUCU
CGAAACAUUAGCCAGGGAGCUUA-U-GUUAGUGGCGA-GAUUAAGG--G-G--------AUCAC--------
-CCC--CGGA--GUCGCAGGGAAAC-CGA-----------------UAC----------------------A
UUCG-CAACUUC----U-------------------------------------------------------
---UU-------------------------------------------------------------------
----------------U--GAAG------------------------------------UGA-GGAAUG-CG
GUGU-----------------------------------GAAA--------------------------GC-
G-C-CU----------------------AUAGUCACUGAUGUAA-UGCCCGAAGUC-GGACGAUCUAGGCCU
GGUCAGGGCGAAGUGCGG-GGAAAC-CCGCAUGGAGGUCCGA--UAUCGUUCUGACGUGCAAUUCGU-UGAG
UUGAACUGGGUCUAGGCGGAAAAGCCAAAUCUAGUCCGA-GGAUGGCUGGUUCCCCUCGAAG-CUGCUCAAA
GGCAGGCCUCAU-ACGA-------------------------------------------------------
----------------------GA-----GU-GAUUGUGGUGUAGAGC-ACUAAUUCGCU-UGGU-ACGGGG
A-G-UA-AA-AUCC-U-CGCCGGCUUCUUAAACUCCGAAGCUGCAA--------------------------
------------------------------------------------------------------------
------------------U-CC-U--UG-UAGA--CUAUGGG-AGACGGC-U-UACCCG-CGG-UAAGCGUG
GG-U-AGCGAGAG-AGAAAC---AACUCAGACCUC-G-GUUAAGGUCCCAAAGU--GU-AGA-CU-AA---G
-U-GUUAAA--CCGAAUGGCGUCAUUUUUCGAAGACAGCAAAAAGGUAGGCUCAGAAACAGCCA-CCCUACA
AUGAGUACGUAACAGUUCAUUUGCU-----AAGAAAAAUGGCCCAGAAAAU-GGAC-GGGGCUCA-AGUCUA
C----------CACCGAU-ACCGGGGGGCAUACU--C-----------------------------------
------------------------------------------------------------------------
-----AGUAAA-------------------------------------------------------------
------------------------------G-AGUAU-GAUC-CGGUAGAGGGGC-GUGGGGC----------
-----------A-U-GAGAU-AAAGUCC--G-GUUGUGAAAU--C-GGAUGAAUCG-U-GCU-CCAGUGCAG
AUCC-UGGCAG-UAGUA-ACAGCU--GAGAU-ACGUGA--A---------------AAUCGU-AUC-CGCCG
A-AAGGCC-AAGGGUUUC-UUGGCAAUACGUCGUUAGCCAAGAGUUAGUCGU--UC-CUAAGGUCAUUCUU-
AAC-CGUC---U-UGA--------------------------------------------------------
----------------------------------------CCG-AA-A--GGGA-AACAGAUU-UAUA-UUU
CUGUA----------------------------------CUUGG----AAGUUAGG----U-AU--------
```

```
------------------U-----------------------------------------------------
--------CGG--CGAC--------GAACGG--------------------------GGGA-U-UGGU-GAC
AGAAGUG-------------GAUAGU-UCGAGCGGG-AU-------UGUCGG-------------UCCCGU-
UUAA-A-UGU-AU-----A-C-------------------CU-----CU----------------GGGG-UG
U-G-CC--------------------UUUGC-GGCGAGAACC---------AGACAA----AAG-CA-UGA-
AUA--GC-----------------------------------------------------------------
-----CGCCUU-AUUGUGGGCGGUU-CGAGA-C-G-ACUCCAAUUCUCA-AUGAAA-A-GCCAAUCU-----
------------------------------------------------U-----CAACGAAC-UU-CUAAGA
CGUACCCAGAACCGAC------------------------------ACAGGUGUGCCCAAGGU-GAGAAGCC
UCAGGC-GUGU-GGAA-UUACGGGU-U-GUUAGGGAAAUCGGC-AAAAUGGAGCGGAACCUUCGG-AAGAAC
GCUUG-CUUG-G-AG--CCUUUGCG----AAAG---------------------------------------
------------------------------------------------------------------------
------------------------------------------------------------------------
------------------------------------------------------------------------
------------------------------------------------------------------------
------------------------------------------------------------------------
------------------------------------------------------------------------
------------------------------------------------------------------------
------------------------------------------------------------------------
------------------------------------------------------------------------
------------------------------------------------------------------------
------------------------------------------------------------------------
------------------------------------------------------------------------
------------------------------------------------------------------------
-------------------------------------------------------------CGG--GGGGCU
CUA--AGUUGCAAUAACUAGUAUCACCCGA-----CUGUUAAAUACAAACAUAGCUUUCUGCAAGACCG---
-AA-A-------------GGUUGUGU-ACAGAAAGUGA-AUCCUGCCCAGUGCAA-G-UCAGUCAAAACCGG
GU-------------C-CA-------------ACCGGAAUAAGCUC--UUGUAAAC-GGCGGGAGUAACUAU
AACUCUCUUAAGGUAGCCAAAUGCCUUGUCGUA-UAAGUAACGACGUGCAUGAACGGACCAACGCGG--UGG
UAACUGUCCCAACA----UCC-UACUCCGUGAACCU---AC---AGGACGGCGAACGGUCCGUCACGCACCG
UUGGGAAGAGAAGA-CCCCAUGGAGU-UUUACU--GCAGCUU-GUUCUUGACAUACUCUGAUACAUGCAGAG
UGU-AGCUGGGAGC-------------GUCGAUGCUAUGC----------------------------UUUC
--------------------------------------GG-GUAUAGCGGA-G--C-GA-CAAUGGAACA-C
CAGC-CUUGUAUUGGGGCAUGUCUAAC--------------UUGC-A-------------------------
------------------------------------------------------------------------
------------------------------------------------------------------------
------------------------------------------------------------------------
------------------------------------------------------------------------
-------------------------------------------------------A----GU----------
------------------------------------------------CGCUUGCGAGGACA-GGUGCAGGU
GGGCAGUUUCGGUGGGGCGCCAGAGCUUGGA-AAAGAUAUC-CAAGCUACCCU-AAGGUCAACUCAACCGGG
ACAGAAAUCCGGUGAA-GAGUG-GAAAGGCAA-AAGUUGGCCUGAUG-GGAC--------------------
--------------CCUUGACAGCAAGGGGUCCCGAGACGAAA-GUCUGGCUUAACGAUCUUUC--------
---AU-U-----AACCU-UUG--------------------GU-AGGUGGUGGAAGUGAUAG-AA--AAAUU
ACCCUGGGG-GUAACAGAGUUGUCGCAGGCAAGAGUUCAUAUCGACCCUGCGGCUUGCUUCCUCGAUGUCGG
```

```
CUCUUCCUAUCCUGGCCAUGCA-GCAGUGGCCAAGGGUGGGGCUGUACGCCCAUUAAAAGGGAACGUGAGCU
GGGUUUAGACCGUCGUGAGACAGGUCGGAUUCUAUCUAAC--------------------G-GUG-------
-------------------------------CUGU--UGGGAGUUUGAGAGCAGCG-UGCGUCCAGUACGAG
AGGAA-CAACGCGCGGUCGCCUC-UGGU-G--AGGGGUUGUUC-GACA-GAGCAU---GCCCCACAGCUACG
CGGCA-AAUCAUAAACGCUGAAAGCA-UC--UAAGCGUGAAGGAUGCCUCGA-A-ACG--AGACUCCC-AA-
--UC-------------------------------------------------------------AAGA---
---------AC----A-CAUGCAAUGCGUGUCAUUCGUUUUGC-----------------------------
---------------------------------------------CGAACGGAU----GAGAAGCGCU----
-----------UGCAA--------------------------------------------------------
------------------------------------------------------------------------
------------------------------------------------------------------------
---------------------------------------ACGGGGACUG------GCGA----CAGUCACCG
----------------------------U-------------------------------------------
---------------------GGUUC------------------------AAGA----CG-AGC------UC
ACCGGUAAAAGACCGGGU-AGG-UGAGUC-AGGCGGUGUAAA-CAC-G--------GAG------------C
AUUU---------GCGACGUGCGUAGCCG--G-CCGA-UCUCAAUA---GAGCGAGUG-UUUGGAC------
------------------------------------------------------------------------
------------------------------------------------------------------------
------------------------------------------------------------------------
------------------------------------U------------
>LSUASG_CP042905.1:656088-659193/1-3106 Anaerobic archaeon MK-D1 chromosome
---------------------C-----------------CUACGCUGUAU-AGUGGAUG-U-CUCGGC-UUG
G-A---AUCUGAUGAAGGGCGUGA--CAAGCUGCGAUAAGCUCCGGGUAGGUGCAAGUAACCGU------CG
AUC-CGG-AGAUCCCCGAAUGUGACA-UC-------------------------------------------
------------------------------------A----U-A-ACUC-AAUUAA---------------U
AUU--------G-----------AGU----------------------------------------------
--UGCUCCCAUAAUCUUUGAUGAGG-G-AG-GAGGAACGUGCUGAAGCGAAGCAUCAUAAUAAGCG--CUGG
AAAAGAAAGCAA--AUGCGAUUCCGUUA--GU-AA-CAGCGAGCGAAAGCGGA-A-AA-GGUCAAACCGAAU
CCCCACGCGAG--------AA------------G---UGU-GGGGAGAUGUGGAG--UGU------------
G-GA--CUCGUGU-CAAAC----------------------------------CUA-----------AA--C
G-------AUAAA--GAUG------AACUAC-CCUG------------------------------------
-------GAA-UG-UCUGA-------CCAUAGAGGGUGAUAGUCCCGUA---------AUC---G----UAA
A------UCGUAAGG-UUUA----------------UGGC-GAGUA--UCCCGAGUAACGCGGGCUG---GU
UGCCUCGUGUGAA-UACACCUGAUACC--AACAGGUAAACCUAAAUA--UAA-C-CCA-AGACCGAUAGC-G
UAU-U-AGUACCGUGAGGGAAAG-AUGAAA-AGCAC-CCAGG-AAUGGU-U-GUG-AAAAGUGCCUGAAACU
A-UACAG-UAAAA-AGU-GA-AGCG---GUCCGAAA-GAGUUU-----------------------------
------------------------------------------------------------------------
------------------------------------------------------------------------
------------------------------------------------------------------------
------------------------------------------------------------------------
------------------------------------------------------------------------
------------------------------------------------------------GA-AUCUCCGU-
---GAAGUCUAAGGUGUG-A-------------------A-------CC-----------------GG----
------------------------------------------------------------------------
-------------------------------------------------------------------AC---
```

------------------------------------------------------------------------
-------------------------------------------------G-AC------GA--------AUG
G-----------------------------------------------------------------------
------------------------------------------------------------------------
------------------------------------------------------------------------
------------------------------------------------------------------------
------------------------------------------------------------------------
------------------------------------------------------------------------
------------------------------------------------------------------------
------------------------------------------------------------------------
------------------------------------------------------------------------
------------------------------------------------------------------------
-----------------------AGACG-----------------AUCAG--G---AUUGCUUCUUUAUUCU
AGAAACAUUAGCCAGGGAGCUCA-U-GUUAGUGGCGA-GAUUAAGG--A-G--------AUAAC--------
-CUC--CGUA--GUCGCAGGGAAAC-CAA-----------------UAU----------------------G
CUUG-UAAGUUCGAUUC-------------------------------------------------------
---GU-------------------------------------------------------------------
----------------C--GAGC------------------------------------CCA-GAAGCA-UG
GUGU-----------------------------------GAAA--------------------------GC-
G-C-CU----------------------AAAGUCACUGAUGUGA-UGCCCGAAGUC-GGACGAUCUAGGCCU
GGUCAGGAUGAAGUGCUU-CGAAAG-GAGCAUGGAGGUCCGA--UAUCGUUCUGACGUGCAAUUCGU-UGAG
UUGAACUGGGUCUAGGCGGAAAAGCCAAAUCUAGUCCGA-GGAUGGCUGGUUCCCCCGAAG-CUGCUCAAA
GGCAGGCCUCAG-ACGA-------------------------------------------------------
----------------------GU-----GU-GAACACGGUGUAGAGC-ACUAAUUCGCU-UGGU-ACGGAU
C-A-GA-AA-UGAU-U-CGCCGGCCUCUUAAACUCCGAAUCUGUGU--------------------------
------------------------------------------------------------------------
------------------U-CC-A--UG-UAGA--CUCUGGG-AGACGGC-U-AACCCG-CGA-UAAGCGUG
GG-U-GGCGAGAG-AGAAAC---AACUCAGACUUC-G-GUUAAGGUCCCUAAGU--GU-UGU-CU-AA---G
-U-GUUAAA--CCGAAUGGCGUUAAAUUUCAAAGACAGUAAAAAGGUAGGCUCAGAAACAGCCA-CCCUAUA
AUGAGUACGUAACAGUUCAUUUACU-----AAGAAAUUUAGCCAGAAAAU-GGAC-GGGGCUUA-AGACAA
C----------CACCGAU-ACCGAAGGGUAUACU--C-----------------------------------
------------------------------------------------------------------------
-----GGCAGA-------------------------------------------------------------
------------------------------G-AGUAU-AAUC-CGGUAGGGGGC-GUGGGGC----------
-----------U-C-GAGAU-AAAGCAG--G-ACAGUAAUGC--C-CUGUGAAUCG-U-ACC-CCAGUGCAG
AUCC-UGGCAG-UAGUA-ACAGCU--GAGAU-ACGUGA--A---------------AAUCGU-AUC-CGCCG
A-AAGGCC-AAGGGUUUC-UUGGCAAUACGUCGUUAGCCAAGAGUUAGUCGG--UC-CUAAGUGCAUUCUU-
AAA-CGUC---G-UGC--------------------------------------------------------
----------------------------------------ACG-AA-A--GGGA-AACAGAUU-UAUA-UUU
CUGUA----------------------------------CUUGG----AGCCUAAGAAAAU-UA--------
------------------U-----------------------------------------------------
--------CGG--CAAC--------GAAAGA--------------------------UGAA-U-UCGA-GAC
GAAAGUG-------------GAUAGC-UCGAACAGG-AC--------UGUCGU-------------UCCUGU-
UUAA-A-UGU-AU-----A-C--------------------CU-----CC----------------GGGA-AG
U-G-CC--------------------UUUAC-GGCGAGAACC---------GGACUA----AAG-CA-UGA-

```
AUA--GC-----------------------------------------------------------------
-----CACCCU-AUUGUGGGAGGUU-CGAGG-C-A-ACUCCAAUUUCCA-UUGAAA-A-UCGCAUUC-----
------------------------------------------------A-----UAAUGAAU-CU-CCAAAC
CGUACCAAGAACCGAC------------------------------ACAGGUGUG-CCAAGGU-GAGAAGCC
UAAGGC-GUGU-GGAA-GUAUAGAU-U-GUUAGGGAAAUCGGC-AAAAUAGAGCGGUAACUUCGG-AAUAAC
GCUUG-CUUG-G-AA--UCUUUGCGGUUAAACG---------------------------------------
------------------------------------------------------------------------
------------------------------------------------------------------------
------------------------------------------------------------------------
------------------------------------------------------------------------
------------------------------------------------------------------------
------------------------------------------------------------------------
------------------------------------------------------------------------
------------------------------------------------------------------------
------------------------------------------------------------------------
------------------------------------------------------------------------
------------------------------------------------------------------------
------------------------------------------------------------------------
------------------------------------------------------------------------
-------------------------------------------------------------CGG--GGGAUU
CUA--AGUUGCAAUAACUAUGUGCACCCGA-----CUGUUAAAUACAAACAUAGCUUUCUGCAAGACCG---
-AA-A-------------GGUUGUGU-ACAGAAAGUGA-AUCCUGCCAGUGGUA-G-UCAGUCAAAACCGG
GU-------------U-CA-------------ACCGGACUAAGCUC--UACUAAAC-GGCGGGAGUAACUAU
AACUCUCUUAAGGUAGCCAAAUGCCUUGUCGUG-UAAGUUACGACGUGCAUGAACGGAUCAACGCGG--UGC
AUACUGUCCCAACA----UUC-U-CUCCGUGAACCU---AC---AGGACGGCGAACGGUCCGUCACGCACCG
UUGGGAAGAGAAGA-CCCUAUGGAGU-UUUACU--GCAGCCA-AUUCCUGAAGUAUUUUUUUAGAUGCAGAG
UGU-AGCUGGGAGC-------------GGAGAUGCCGUGU----------------------------UUUC
--------------------------------------GG-ACAUGGCGGA-G--C-AA-CAAUGGAACA-C
CAGC-CUUCUAAAAGGAUAUUUCUAAC--------------UCGU-C-------------------------
------------------------------------------------------------------------
------------------------------------------------------------------------
------------------------------------------------------------------------
------------------------------------------------------------------------
-------------------------------------------------------A----CU----------
------------------------------------------------AGGUGGCGAGGACA-UGUGUUGGU
GGGCAGUUUCGGUGGGGCGCCAGACCUUGGA-AAAGAUAUC-CAAGGUACCCU-AAGGUCAACUCAAUCGGA
ACAGAAACCCGAUGAA-GAGUG-GAAAGGCAA-AAGUUGGCCUGAUG-GGAC--------------------
--------------CCUCGACAGCAAGGGGUCCCGAGACGAAA-GUCUGGCUUAACGAUCCUUU--------
---ACAG-----AAUUU-UUG--------------------GU-AGAUGGUAAAGGUGAUAG-AA--AAAUU
ACCCUAGGG-GUAACAGAGUUGUCACGGGCAAGAGUCCAUAUCGACCCCGUGGCUUGCUUCCUCGAUGUCGG
CUCUUCCUAUCCUGGCCGUGCA-CCAGCGGCCAAGGGUGGGGCUGUACGCCCAUUAAAAGGGAACGUGAGCU
GGGUUUAGACCGUCGUGAGACAGGUCGGAUUCUAUCUAAC--------------------G-GUG-------
-------------------------------CUGU--UGGGAGUUUGAGAGCAACG-UAUGUCCAGUACGAG
AGGAA-CAACAUGCGGUCGCCUC-UGGU-G--AGCAGUUGUCU-GACA-AGGCAU---GCUGCACAGCUACG
CGGCA-AAUCAUAAGCGCUGAAAGCA-UC--UAAGCGCGAAGGAUGCCUCGA-A-AAG--AGACUCCC-AU-
```

```
--UC-------------------------------------------------------------AAGA---
---------ACAGCAA-UAUGUAGUGAGCACAA---GUCUUGC-----------------------------
---------------------------------------------CGGAUUUGU----GUAAGUGAAA----
-----------UAU-A--------------------------------------------------------
------------------------------------------------------------------------
------------------------------------------------------------------------
---------------------------------------CCGGGGACUG------GUGA----CAGUAACCG
----------------------------G-------------------------------------------
---------------------GGUUC------------------------AAGA----CG-AGG------AC
ACGCGUAAAAGACGCGAU-AGG-UGAGAU-AGGCGGUGUAAA-CAC-G--------GAG------------C
AUUA---------GCGACGUGCGUAGCCG--G-CCAU-UUUCAAUA---GUCCGAGUG--------------
------------------------------------------------------------------------
------------------------------------------------------------------------
------------------------------------------------------------------------
-------------------------------------------------
>LSUASG_2026747_HEIM/1-3090 PBWW01000025.1 Candidatus Heimdallarchaeota arc
---------------------------------------GAACAGUAAAU-UAUGGAUA-C-CUUGGC-UUC
U-A---UACCGAAGAAGGGCGUGG--CAAGCUGCGAUAAUCUAUGGGUAGCCGCAUGCAGGCUA------UG
AUC-CAU-AGGUACCCGAAUUGGAAU-CCAUU----CAUU------GUC---CAAAAA-GCAA---------
-------------------------U----G--AAAG------G-CCCC-----------------------
-----------------------UUA----------------------------------------------
-----------------------GG-G-CC-GGGGAACCCUCCGAACGAAAGCAUUCAAGUAGGAG--GUGG
AAAAGAAAUCAAUUUCGAGAUUUCCUUA--GU-AA-CGGCGAGUAAAAAGGAA-A-CA-GGUCAAACCGAAU
GAGU-AUAAGA--------AA------------U---UAU-AUUCGGAUGUGGUG--UUGA-----------
--GG--ACU-------------------------------AGUGUCAACGCG-UAA-----------GU--U
U-------UAGAA--ACUGA-----AUUUUG-UUUG------------------------------------
-------GAA-CA-CAAAA-------CCAUAGAGGGUGACAGUCCCGUA---------AGU---G-UAUAAA
A------CAUGCUAC-GAGCA--------------------UUAGU-AUCCAGACUAGGGCGGAUUG---GU
AAUUCCGUCCAAA-CCUCCAGAACC----AACUGGAAAGCCUAAAUA--UUA-A-UAGAAGACCGAUAGC-G
AAC-U-AGUAACGUGAGUGAAAG-GUGAAA-AGUAG-UGGUA-AAACAC-A-GUG-AAAAGAGCUUGAAAUA
A-UUUACUUAAAG-AGU-GU-AAUG---GCGUCGAA-GGAAAG-----------------------------
------------------------------------------------------------------------
------------------------------------------------------------------------
------------------------------------------------------------------------
------------------------------------------------------------------------
------------------------------------------------------------------------
------------------------------------------------------------AG-ACUCAUCG-
---AAUCU-------------------------------A------ACG-----------------GUU---
------------------------------------------------------------------------
--------------------------------------------------------------GN---GA---
------------------------------------------------------------------------
-------------------------------------------------A-AC------CU--------UCU
UGAGA-------------------------------------------------------------------
------------------------------------------------------------------------
------------------------------------------------------------------------
```

```
------------------------------------------------------------------------
------------------------------------------------G-----------------------
------------------------------------------------------------------------
------------------------------------------------------------------------
------------------------------------------------------------------------
------------------------------------------------------------------------
------------------------------------------------------------------------
-------------------AUGAGU-UU-----------------ACCAG--C---GUCAUUGCGUCAGUCU
UGAAACACUGGCCAGGGAGUUUA-U-UUAUGUGGUGA-GGCUAAGA--UCA---------------------
U-AU--CUGU--GCCGUAGGGAAAC-CAA-----------------CCA----------------------G
CCCG-CAAUAUU------------------------------------------------------------
------------------------------------------------------------------------
----------------------A------------------------------------UGA-GGGGCG-AC
GUCU-----------------------------------GAAA--------------------------GG-
G-C-GU----------------------GAAAUCACAUGAAUAA-GAUGCGAAACG-AGUUGAUCUAUGCGU
GGGCAGGUUGAAGCCUUG-GGAAAC-UAAGGUGGAGGACCGA--ACGGUUGCUGAUAUACAACUCGC-UCCG
CUGACCUACGUAUAGUAGUCAAAGGCUAAUCUAAACUCG-UGAUAGCACAUUAAAACCGAAGUGGUGCCGCA
GACCAGCGUUGG-ACGA-------------------------------------------------------
----------------------AA-----GA-AAUUUGAAGGUAGAGU-ACAGACA-GAG-UCGG-AAAGGG
C-C-UUCAC-GGGC-U-CGCCACUCCAUCUAACUCCGAAUGUCAAA--------------------------
------------------------------------------------------------------------
------------------U-UC-U--UG-UAGA--CUCCAGC-AGUCGGGUG-GGUC-G-CCG-UAAGGGUG
AU-C-ACCGAGAA-GCGAAU---AAGCUUGAUCUG-G-GUUAAGGGCCCCAAAU--GU-CAG-UU-AA---G
-U-GUUAAG--CUGAAGGGCGUCGUAUCGCGAAGACAAUGAGGUGGUAAGCUUAGAAGCAGCUA-CCCAUUA
AAAAGUUCGUACUAGAUUACUCAUC-----GAGCGGUACGGCCCUGAAAAU-GGAC-GGGGCUCA-AACUGA
C----------UCCCGAU-ACCCAGAGGCGUAUA--U-----------------------------------
------------------------------------------------------------------------
------------------------------------------------------------------------
-------------------------------UAAUAC-GAUC-CGGUAGGUUUUC-GAUGGCA---------
-----------U-A-GGGUA-GAAGCAG--G-UUGGAGAUGA--C-CUGUGGACCU-U-UGU-CAUGUGCAG
AUCC-UGGUAG-UAGUA-GGCAGA-AAACAU-GGGUGA--A---------------AACCCC-AUG-GGUCA
A-AAAGGC-AAGGGUUCC-GAGGCAAUG-UCAAUCAGCCUCGGGUAAGCCGG--UC-CUAAGCGUAACCUU-
AUU-AUGG---U-CAU--------------------------------------------------------
----------------------------------------CGC-AA-A-AGGGA-AGCAGGUU-AAUA-UUC
CUGCG----------------------------------CUGCG----CCUG--------A-AC--------
------------------------------------------------------------------------
-------UCGG--UAAC--------GAUAGA--------------------------AUAU-G-AUCG-GAC
GCUUGUA-------------GAUAGG-UUCUUGUGG-AC-------CGUC-------------G-UCCAGA-
CUAA-G-UGU-AU-----A-A-------------------GU-----CC----------------UAGG-AG
U-U-GC--------------------GUUGA-GCAGAGAAUU---------GGACAA----AAU-CA-CGA-
AUG--GC-----------------------------------------------------------------
-----CACCCA-AUAGUGGGGGGUU-GAAUU-G-A-UUUCUAAAAGCCC-AUGAAA-A-CGA----------
------------------------------------------------UCAUAUUUACUAAU-GG-CGCAUC
CGUACCUA-AACCGAC------------------------------ACGGGUGUCCUUAGGCU-UAGAAGGC
UAAGAC-CUCU-UGGA-GUAAUGUA-G-UUAAGGGAAUUCGGC-AAAUUAGGCGAGUAACUUCGG-GAUAAU
```

```
CGCUG-CCUU-U-AG--ACGU---------------------------------------------------
------------------------------------------------------------------------
------------------------------------------------------------------------
------------------------------------------------------------------------
------------------------------------------------------------------------
------------------------------------------------------------------------
------------------------------------------------------------------------
------------------------------------------------------------------------
------------------------------------------------------------------------
------------------------------------------------------------------------
------------------------------------------------------------------------
------------------------------------------------------------------------
------------------------------------------------------------------------
------------------------------------------------------------------------
-------------------------------------------------------------UUU--GCGUCC
GAA--GGUGUCACUAACAUAGGGAACCCAA-----CUGUUUAACAAAAACAUAGCAAACUGCUACGCGG---
-AA-A-------------CGCUUUGU-AUAGUUUGUGA-AUCCUGCUCCGUGCUC-G-UAAG-UUAAAGCCG
GG-------------UACA-------------ACCGGCCUAAGCUC--GAGUUAAC-AGCGGGAGUAACCAU
GACUCUCUUAAGGUAGGCAAAUGCCUUGUCGGA-UAAUUACCGACGUCCAUGAAGGGAACAAUGAGG--UUC
CCCCUGUCCCUAAC----UAG-AAUCCAGUGAACCU------ACAGUCCGGCGAACAGUCCGGAGAGACCUG
ACGGGAAGAGAAGA-CCCUG-AGAGC-UUUACU--ACAGCUU-CCAGUUGUGUAUGCGGUUAUCAUACAGAG
CGU-AGUUAGGAGACUUUG----------AAGAUUU-AUC----------------------------CCNG
--------------------------------------GG-UAAAUUGGAG-U--C-AA-CAAUGGAACA-C
UAAC-UAUGAUGACUGCGUAUGCUCACCU-----------------G-------------------------
------------------------------------------------------------------------
------------------------------------------------------------------------
------------------------------------------------------------------------
------------------------------------------------------------------------
-------------------------------------------------------A---UGC----------
-----------------------------------------------GUCGG-----GGACA-CCUGGAGGU
GGGUAGUUUGGCUGGGGCGGCACGCCUCCGA-UAAGAAAAC-AGAGGCGCCCA-AAGGUUGACUCAGAAGGG
UCAGAAAUCCUUCGUA-GAGUG-CAAGGGCAU-AAGUCAGCUUUACU-GUGA--------------------
--------------GCACGAAAAUCCUGCUCGCUGUUACGAAA-GUAGGGCCUAGCGAUCCAUC--------
---CU-G----AUCCUCCUUG--------------------GU-GGGGGCGGGUGGUGACAG-AA--AAGUU
ACCUCAGGG-GUAACAGAGUUGUCGCGGGCUAAAGUUCACAUUGACCCCGCGGCUUGCUUCCUCGAUGUCGG
CUCUUCCUCGCUUGGUGAUGCA-CAAGUCACCUAGAGUAGGGUUGUUCGCCCAUUAAUAGGGAACGUGAGCU
GGGUUCAGACCGUUGUGAGACAGGUUGGUUUCUAUCUUUC--------------------A-GGU-------
-------------------------------CUGU--UGCGUAUUUGCAGGUAAAU-UGCCACCAGUACGAG
AGGAA-CGUGGGGAUGGGGCCUC-UGGU-C-GAUCGGUUGUCU-GACA-AGGCAC-U-GCCGAGUAGCUACG
CCCUU-CCAGAUAAGAGCUGAACGCA-UC--UAAGCUCGAAGCUGGUCCUAA-A-ACG--AAAUACGC-AU-
--UCAAUACCACA-----------------------------------------------------------
---------------GAUCUAUAAUGUUCAAAAA--------------------A----------------A
CAUAUCAUACCAUGAUAUU-----------------AUUAUUACAUUUAUUCAU------------------
-------UAUAUUUCA--------------------------------------------------------
------------------------------------------------------------------------
```

------------------------------------------------------------------------
--------------------------------------ACAUGGGGCGG------GUGA----CUGCAACAU
----------------------------G-------------------------------------------
--------------------GGUAU-------------------------UAGA----CG-ACU------CC
ACCCAAAGAAGAUGGGAU-UGA-UAGACA-GGU-UGUGUAAG-CGU-U--------GAG-------------
CCUCGG--------CGAAAUGUUUAGCAU--A-CCUU-UACUAACG---GAGAAGGUG-UUUUACG------
------------------------------------------------------------------------
------------------------------------------------------------------------
------------------------------------------------------------------------
-------------------------------------------------
>LSUASG_HEIM_B18G1/1-3101
----------------------------------------AACGCCAUCU-GGGGGAUG-G-AUAGGC-UUG
A-G---CGCCGAUGAAGGCCGUGG--CUAGCUGCGAUAAGCGCGGGGUAGUUGCAGGCAAACGU------CG
AUC-CCG-CGAUUGCCGAAUAGGAAU-UCCA-----UGU-------AU----GGA----------C------
----------------------------------CAA----U-A-AGCC-AUACAA---------------C
UCC--------G------------------------------------------------------------
---------------------UAAG-G-AG-GGGGAACGCACCGGACGGAAACAUCCAAGUAGGUG--CAGG
AAAAGAAAUCAC--CUGAGAUGCCCUGACUGU-AA-CGGCGAGUGAAAGGGGC-A-UA-GACCAAACUGAAU
CCUGGCGGGC---------GA------------C---CGU-UAGGAGAUGUGGUGU-UGU------------
G-GU--CUGUCAC-UAAAU----------------------------------UCU-----------GC--G
U-------AUGAU--AGUC------GAAUAU-CCUG------------------------------------
-------GAA-AG-UUUAG-------CCUUAGAAGGUGAUAGCCCUGUA---------GGC---G----AAA
UCAUA-CCCAGAAGG----------------------AGA-CAGUA--ACCAGAAUAACGGGGGUUG---GA
AAUCUCUCGUGAA-CUGGGGCACAACG--AGCU-CCAAGUCUAAAUA--CGU-C-UCA-AGACCGAUAGC-G
CAA-U-AGUAACGUGAGUGAAAG-CUGAAA-AGUAA-CCCGG-AAGGGU-G-GUG-AAAAGUGCCUGAAACC
A-GAUGGUUAUAG-ACG-UU-UGUG---GUCCGAAA-GCGUAG-----------------------------
------------------------------------------------------------------------
------------------------------------------------------------------------
------------------------------------------------------------------------
------------------------------------------------------------------------
------------------------------------------------------------------------
------------------------------------------------------------AG-UCUGGACG-
---AA----------GAG-A-------------------A------GGC-----------------UG----
------------------------------------------------------------------------
-------------------------------------------------------------------GU---
------------------------------------------------------------------------
-------------------------------------------------G-AC------GG--------CUG
GAUCG-------------------------------------------------------------------
------------------------------------------------------------------------
------------------------------------------------------------------------
------------------------------------------------------------------------
------------------------------------------------A-----------------------
------------------------------------------------------------------------
------------------------------------------------------------------------
------------------------------------------------------------------------

```
------------------------------------------------------------------------
------------------------------------------------------------------------
-------------------GUUCAGAGG-----------------AGCGG--G---AUCACAAGAUCCGUCU
AGAAACACGGGCCAGGGAGUUUG-C-CCUCGUGGCGA-GCUUAAGG--G-U--------UUCAA-------A
GACC--CGAA--GGCGGAGCGAAG-CGA------------------CAA----------------------G
UCCG-CAGCUUG------------------------------------------------------------
------------------------------------------------------------------------
-----------------------------------------------------------CAA-GGGACG-CC
GUCU-----------------------------------GAAA--------------------------GG-
G-C-GG----------------------GAAGUCACGGGGGUAA-GACCCGAAACC-GGACGAUCUAUCCGU
GGACAGGCUGAAGCCCGG-GGAAAC-UCGGGUGGAGGGCCGA--ACAAGUGUUGACGUGCAAUUCACGUUUG
UUGAUCUGCGGAUAGGGGCAAAAGACCAAUCUAGUCCGG-UGAUAGCUGGUUCCCAACGAAG-UGGACAU-A
GGCCAGCCUGAU-UUGA-------------------------------------------------------
----------------------GG-----UG-GCGCAUUCGGUAGAGC-AAGGAUU-CGU-GGUC-AUGGGG
A-G-UA-AA-AUCC-U-CACCCAUGAGUCAAACUACGAACGGAUGC--------------------------
------------------------------------------------------------------------
------------------G-UA-C--CA-UAGA--CGAUCAG-AGUCGGG-C-GUGCGU-GCG-UAAGGCAC
GU-G-GCCGAAAG-GGGAAU---AACCCAGACCAU-G-GUUAAGGACCCCAAGU--CU-CGA-UU-AA---G
-U-GUUAAAAUCGGAAGGGCGUCCUUGGACUAAGACAGUUAGGCGGUAGGCUCAGAAACAGCCA-CCCGCUA
AGCAACACGUAACAGUGGACUAACC-----GAGUCCUUGGGCCCCGAGAAU-GGAC-GGGGCU-A-AAUCGA
G----------CUCCGAG-ACCAUGGGGCACACU--G-----------------------------------
------------------------------------------------------------------------
-----CAA----------------------------------------------------------------
--------------------------------GGUGU-GAUC-CGGUAGUUGGGC-GAUCGGG---------
-----------U-U-GGGCG-GAAGUAG--G-GCAGUGAUGU--C-CUGUGGACCG-A-CCC-GAUUUGCAG
AUCC-UGGCGA-UAGUA-ACAGCA--AAGAG-AGGUGA--G---------------AAUCCU-CUC-CGCCG
A-AAGGGC-UAGGGUUUC-ACGGCAACGU-UCGUCAGCCGUGAGUUAGUCGA--UC-CUAAGGCCAGUCUU-
ACC-UGAG---C-UGG--------------------------------------------------------
----------------------------------------UCG-AA-A--GGGA-AACAGGUU-UAAA-UUC
CUGUA----------------------------------CUGCG----UGGAUACGCC--A-GA--------
------------------A-----------------------------------------------------
-------GCGG--CAAC--------GCAAAG--------------------------CUCA-UCUUCU-GAC
GCGACUG-------------GAUAGG-CUCUGUUAA-GC--------CAUCG--------------CUUAAU-
UUAA-C-UGU-AU-----A-A-------------------CC-----CU----------------CCGA-AG
U-A-CU--------------------GUGAC-AGUGAGAAGG---------AGGCGA----AAG-CA-GGA-
AUA--GG-----------------------------------------------------------------
-----CUGCCA-AUAGUGGCGGCUA-G-AGC-U-G-CUUCCACUCGCCC-AUGAAA-A-GGAAGAUG-----
------------------------------------------------A-----GAAGGAUU-CCACGUAAU
CGUACCGAGAACUGAC------------------------------ACAGGUGU-CCCUAGGU-GAGAAGCC
UAAGGC-GUGU-UCGG-UCAAAGAU-G-GUGAGGGAAAUCGGC-AAAUUCGCCCUGUAACUUUGG-GAUAAA
GGGUG-CCUG-U-GG--GCUU---------------------------------------------------
------------------------------------------------------------------------
------------------------------------------------------------------------
------------------------------------------------------------------------
------------------------------------------------------------------------
```

```
------------------------------------------------------------------------
------------------------------------------------------------------------
------------------------------------------------------------------------
------------------------------------------------------------------------
------------------------------------------------------------------------
------------------------------------------------------------------------
------------------------------------------------------------------------
------------------------------------------------------------------------
------------------------------------------------------------------------
------------------------------------------------------------GAAA--GGGCUC
GCA--GGUCGCAGUGACAAGAGGGGGGCAA-----CUGUUCAAUUCAAACAUAGCAAACUGCAAGAGCG---
-AA-A-------------GCUUGUGA-ACAGUUUGUGA-AUCCUGCUCAAUGCUG-G-UAAGUCAAAGCC-G
GU-------------UACA-------------ACCGGUCUAAGCUC--CAGUCAAU-AGCGGGAGUAACUAU
GACUCUUAUAAGGUAGCCAAAUGCCUUGUCAGA-UAAGUACUGACGUGCAUGAACGGAUCAAUGACC--UCC
CUACUGUCCCCACC----AUC-UACCGAGUGAAAUC---AC---UAUCCAGCGAACAGUCUGGAACCAACCA
GCGGGAAGAGAAGA-CCCCGUGGAGU-UUUACU--GCAGCCU-GUGCCUGUUGUACGGGUCGCAAUACAUAG
AGU-AGUCGGGAGC-------------GUUGAACCAAGUU----------------------------CUCC
--------------------------------------GG-GACUUGGGAAUG--C-GA-CAAUGAAACA-C
CGAC-UAUUGUUGCUUGUACGACUAACU--------------------------------------------
------------------------------------------------------------------------
------------------------------------------------------------------------
------------------------------------------------------------------------
------------------------------------------------------------------------
------------------------------------------------------------UU----------
------------------------------------------------CUGCGGAAAGGACA-AGUAUUGGU
GGGCAGUUUAGCUGGGGCGGCU-GCGCUUGA-AAAAGUAUC-GAGCGUGUCCA-AAGGUCAGCUCAGGAGGG
ACAGAAAUCCUCCGUA-GAGCA-UAAGAGCAA-AAGCUGGCUUGACA-GGGU--------------------
--------------UCUUACCCAUAGGGAACCCUGGAACGAAA-GUUGGGUCUAGCGAUCUGUC--------
---GA-G-----ACUUC-UCG--------------------UU-GAGGUCCGGCAGCGGCAG-AA--AAAUU
ACCCCGGGG-GUAACAGAGUCGUCGCAGGCGAGAGCUCACAUCGACCCUGCGGCUUGCUACCUCGAUGUCGG
CUCUUCCUCUCCUGGCGCCGCACAAGAC-GCCUAGGGUAGGGUUGUUCGCCCAUUAAAAGGGAACGUGAGCU
GGGUUUAGACCGUCGUGAGACAGGUCGGAUUCUUUCUACU--------------------G-GUU-------
-------------------------------GUGC--UGCGAAUCUGAGGGUAGGG-CCGGGUCGGUACGAG
AGGAC-CCCCCCGUCGGCGCCUC-UAGU-C-UACCGGUUGUCU-GAUA-AGGCAU-U-GCCGGGCAGCCACG
CGCCA-AGCGAUAAUAGCUGAAAGCA-UC--UAAGCUAGAAACGCGUCCUGA-A-ACG--AGAUUCGC-AU-
--UC-----UGGGGAUUAUUUG------GAGUUUGAGACCUGCUGCUCUUU------AGCUGUUGAGCAGCG
CU-------------UUUAGAUGUCAAGUGUAUGUCUUUUUUGGUCUCCUCCUGC----------------A
C-----------------------------------------------------------------------
------------------------------------------------------------------------
------------------------------------------------------------------------
------------------------------------------------------------------------
------------------------------------------GGGGCUG------GUGA----CAGCAACGU
----------------------------G-------------------------------------------
--------------------UCUUCCU-----------------------CAGA----CG-ACG------GU
UCCGAUACAAGAUCGGUU-UGA-AAGGCC-UGC-GGUGUACG-CGU-C--------GAG------------U
```

```
CUUCGG-------GCGAGAUGUUCAGCCG--GCAGGU-ACUUACCA---CCGA-------------------
------------------------------------------------------------------------
------------------------------------------------------------------------
------------------------------------------------------------------------
-------------------------------------------------
>LSUASG_1538547_LOKI/1-3298 JYIM01000373.1, Lokiarch_contig000373, whole ge
------------------------CCAAUAC----GC--CUCCGCUAUUU-GGUGGAUA-U-CUUGGC-UUG
G-G---AACCGAUGAAGAGCGUGA--CAAGCUGCGAUAAGCUCGGGUUAGUCGCAUGAAGACGU------UG
ACC-CCG-AGAUCCUCGAAUAGGACA-UCCA-----CUU-------GC----GGA----------C------
----------------------------------CAA----U-A-AUCC-GCAAGA---------------U
UCC--------C-----------UUA----------------------------------------------
--U--------------------GG-G-AA-GGGGAACCCUGAGAAGUAAAACAUUUUAGUAUCAG--GAGG
AAAAGAAAACAA--UAGUGAUUCCGUGA--GU-AA-CGGCGAGCGAAAGCGGA-A-CA-GAUCAAACCGAAU
CCUU-AGGGA---------AA------------C---CUA-UAGGAAAUGUGGUG--UGU------------
G-GA--CCC-CGU-ACGAC----------------------------------UUC-----------CC--U
A-------GAGAA--AAUG------AAUACU-CCUG------------------------------------
-------GAA-AG-UUGAG-------CCAUAGAGCGUGAAAGCCGCGUA---------AUU---G----CAA
U------CUAGAAAG-UUGC----------------UGCG-GGUGU--UCCCGAGUAGUGCGGGAUG---GU
UUUCUCGUACGAA-AAUGCCAGAUACA--CGCUGGCAAAUCUAAAUA--UAU-C-CCA-AGACCGAUAGC-G
AAA-U-AGUAGCGUGAGCGAAAG-AUGAAA-AGUAA-CCCGG-AAGGGU-C-GUG-AAAAGUGCUUGAAACC
A-AAUAGCUUAGU-AAG-GA-UGCA---GCCCGAAA-GGAUAG-----------------------------
------------------------------------------------------------------------
------------------------------------------------------------------------
------------------------------------------------------------------------
------------------------------------------------------------------------
------------------------------------------------------------------------
------------------------------------------------------------AU-UCUGCGCG-
---AA-----------UG-A-------------------G------UUC-----------------UG----
------------------------------------------------------------------------
-------------------------------------------------------------------GU---
------------------------------------------------------------------------
-------------------------------------------------G-AC------AG--------AAC
AUCAG-------------------------------------------------------------------
------------------------------------------------------------------------
------------------------------------------------------------------------
------------------------------------------------------------------------
------------------------------------------------A-----------------------
------------------------------------------------------------------------
------------------------------------------------------------------------
------------------------------------------------------------------------
------------------------------------------------------------------------
------------------------------------------------------------------------
-------------------GCACAGAGG-----------------ACUAG--G---GUUGCAUCAUCAGUCU
GGAAUCAUUGGCCAGGGAGUUUA-U-AGUAGAAGCAA-GCUUAAGU--G-G--------AUCAC--------
C-CA--CGGA--GGCAUAGGGAAAC-CGA-----------------UAU----------------------U
```

```
CUUG-CAACCUA----A-------------------------------------------------------
---UUAUCACAUUAGGAAGGUCUAAUC-------GUAAAAGAGGGGUUCAGCCUCUCG-GAUCACGUUUAGC
AAAUCCUUCUAAGGAGU--UAGG------------------------------------CCA-GAAGAA-UG
GUCU-----------------------------------AAAA--------------------------GG-
G-C-CG----------------------AAAGUUUCUGCUAUAA-UACCCGAAGCC-GGUCGAUCUAUUCCU
GGUCAAGAUGAAGUCGUU-CGAAAG-GAUGAUGGAAGUCUGC--ACCAGUUCUGAUGUGCAAAUCGA-UUGG
AUGAACUGGGAAUAGGAGCAAAAGGCUAAUCUAGAUCGG-UGAUAGCUGGUUCUCCCCGAAG-UUGGUGUCA
GCCAAGUCUCAG-ACGA-------------------------------------------------------
----------------------AG-----UC-GCCAGUGACGUAGAGC-ACUAAUU-GUG-UGGU-ACGGGG
G-A-GG-AA-UCCC-U-CGCCUCGCUCUUAAACUCCGAAAUUGCUG--------------------------
------------------------------------------------------------------------
------------------G-CA-C--UG-AAGA--CUCUGGG-AAACGGG-U-UGGUGG-GCG-UAAAGUCC
AU-C-GCCGAGAA-GGGAAC---AACCUAGACUUG-G-GUUAAGGUCCCUAAAU--GC-AGG-UU-AA---G
-U-GUUAAA--CGAAAGGGCGUUAUAUUUCUAAAACAGCAAAAGGAUAGGUAUAGAAGUAGCCA-UUCCCUA
ACGAGUGUGUAACAACUCAUUUGCC-----GAGAAAUAUAGCCCCGAAAAU-GGAC-GGGGCUCA-AAUCUG
C----------UACCGAG-ACCCAAGAGCACGGG--C-----------------------------------
------------------------------------------------------------------------
-----GGAAGA-------------------------------------------------------------
------------------------------ACCCCGU-GAUC-UGGUAGGGGAGC-GUCCGGA---------
-----------U-G-GGGUU-GAAGCAG--G-UGCGUGAGUU--C-CUGUGGACCC-G-UUC-GGAGUGAAA
AUCC-UGGUAA-UAGUA-AAAGCA--UAGAU-GGGUUA--G---------------AAUCCC-AUC-CGCCG
A-AAGGGC-AAGGGUUUC-UUUGCAAUGCGUCGUCAGCGAAGAGUUAGUCGA--UC-CUAAGCGUACCCUU-
AAC-UGGU---A-UGC--------------------------------------------------------
----------------------------------------GCG-AA-A--GGGA-AACAGGUU-UAUA-UUC
CUGUA----------------------------------CCACG----GCACUACG----A-AU--------
------------------A-----------------------------------------------------
-------GCGG--UAAC--------GCAUGG--------------------------UUUA-A-UGUU-GAC
AGUUCCG-------------GAUAGG-UGGAAUAGG-GU--------CGUCGAAUAUCU-------CCCUAU-
UUAA-A-UGU-AU-----A-A-------------------UG-----CU----------------GGUG-AA
U-A-GA--------------------GUAAA-UCUUAGACCC---------AGCUUA----AAG-CA-UGA-
AUA--GC-----------------------------------------------------------------
-----CAUCCA-AUUGUGGAAGGUU-U-CGC-C-G-AGUCCAGGAUUUA-UUAAAA-A-GAC-AUAA-----
------------------------------------------------A-----CAAGAAAU-GC-CGUGAU
CGUACCCAGAACCGAC------------------------------ACAGGUGC-CCCUAGGU-GAGUAGCC
UAAGGC-GUAU-UCGU-GUAAAGGA-G-GCUAGGGAAAUCGAC-AAAUUAGAGCCGUAACCUUGG-GAUAAG
GCUUG-CUUG-C-GA--ACUUCUUC----ACGCA-GCUUCG------GCAGU--------------------
------------------------------------------------------------------------
------------------------------------------------------------------------
------------------------------------------------------------------------
------------------------------------------------------------------------
------------------------------------------------------------------------
------------------------------------------------------------------------
------------------------------------------------------------------------
------------------------------------------------------------------------
------------------------------------------------------------------------
```

```
------------------------------------------------------------------------
------------------------------------------------------------------------
------------------------------------------------------------------------
------------------------------------------------------------------------
----------------------------------------------------------GUGUGG--GGGUUU
UCA--AGUUGCAAUAACUAGGGGGGGACGA-----CUGUUUAACAAAAACAUAGGUUUCUGAUAGAGUG---
-AA-A-------------ACUUUUUU-ACAGAAGCUGA-AACCUGGCCAGUGGCA-G-UUUGAUCAAUCCUG
GU-------------UACA-------------ACCAGAGUAAGUCC--UGCUAAAC-GCCGGGAGUAACUAU
AACUCUCUUAAGGUAGCCAAAUGCCUUGUCUGG-UAAGUUCAGACGUGCAUGAAUGGUCCAACGUUC--UCC
CUAUUGUCCCAGCC----UUC-UGACGAUUGAAAUU---AC---ACUCCGGCGAACGGUCCGGAGAAGCCCA
GCGGGACGAGUAGA-CCCCGUGGAGU-UUUACU--GCAGCCU-GUUGUUGCGAUUUUGGACUAAAUGCAGAG
UAU-AGUCGGUAGC-------------GUAGAUGCCGUGA----------------------------UCUC
--------------------------------------GG-UCACGGUGGA-G--C-AC-AAAUGGAAUA-C
CGGC-CUUUCCUUCUGAAAUCGCUAACGG-C--------A-UC-A-A--------------------------
------------------------------------------------------------------------
------------------------------------------------------------------------
------------------------------------------------------------------------
------------------------------------------------------------------------
-------------------------------------------------------U----AG----------
------------------------------------------------UGUC-----GAACA-CCAGCAGGU
GGGCAGUUUGGCUGGGGCGGCUCACUCUAGA-AACAAUAUC-UAGAGUAACCA-AAGAUCAGCUCAAGCGGG
UCAGGAAUCCGCUGUA-GAGUU-UAAGAGCAA-AAGCUGGUCUGACU-GGAU--------------------
--------------UCUGAUCACCAAGGAAUCCUGGGCUAAC-GCCUGGUCUAGCGAUCCUUA---------
---AC-U-----AACCU-CGU--------------------GU-GGGUGGUUAAGGUGAUAG-AA--AAAUU
ACCCCGGGG-GUAACAGAGUUGUCACGAGUAAGAGUUCCUAUCGACCUCGUGGCUUGCUUCCUCGAUGUCGG
CUCUUCCUAUCCUGGGAGCGCACAUAACUUCCAAGGGUGGGGCUGUACGCCCAUUAAAAGGGAACGUGAGCU
GGGUUCAGACCGACGUGAGUCAGGUCGGUUUCUAUCUGCU--------------------G-GGC-------
-------------------------------UUGU--AGGUUGACUGAGGGUAAGG-GAAAAUCAGUACGAG
AGGAA-CAUUUUGCCGGCGCCUC-UAGU-G-UACCGGUUAUCU-GACA-AGGUAU-C-GCCGGGCAGCCACG
CGCCA-AUGGAUAAGUGCUGAAAGCA-UC--UAAGCACGAAGCCAGACCCGA-A-AAU--AGUCAGCC-AA-
--UCUUAUUCCGU-AAUGUUGGGCUUUAUACUUUAGUAUUCUUGU-UGAUGU----UGGAAGGU-AAGA---
---------CC----G-CCUGUGAUGGUCGGAAGCCUUCCAAUCGAGUCUAUGCG----------------A
GGGAUUGCAU-----GGAU-----------------GGGAU----CGUCAUAAU----GAGAAUGGUGAAUA
UGAAGUACCAGCGCGA--------------------------------------------------------
------------------------------------------------------------------------
------------------------------------------------------------------------
---------------------------------------GCGGGGUGCG------GUGA----CGCAAACCG
----------------------------U-------------------------------------------
--------------------GGGAAC------------------------GAGA----CG-AGU------GC
ACCCGUAAAAGACGGGGU-UGG-UGAGGU-AGG-GGUGUACG-UAG-G--------AAG------------C
UUUCGG-------GCGACCUAUUCAGCCC--G-CUAC-UCUCAACC---GCACAAGUG-UAUGGC-------
------------------------------------------------------------------------
------------------------------------------------------------------------
------------------------------------------------------------------------
-------------------------------------------------
```

```
>LSUASG_SDNK01000007.1:46432-48422_SDNK01000056.1:1-1270/1-3259 Candidatus
----------------------------------------------AUUU-GGUGGAUA-U-CUUGGC-UUG
G-G---UACCGAUGAAGAGCGUGA--CAAGCUGCGAUAAGCUCGGGUUAGUCGCAUGUAGACGU------UG
ACC-CCG-AGAUUCUCUAAUUGGACA-UCCA-----CUU-------GC----GGC------------C------
----------------------------------CAA----U-A-AUCC-GCAAGA---------------U
UCC--------C-----------GAA----------------------------------------------
--A--------------------GG-G-AA-GGGGAACCCUGGGAAGUAAAACAUUUUAGUACCAG--GAGG
AAAAGAAAACAA--CAGUGAUUCCGUUA--GU-AA-CGGCGAGCGAAAGCGGA-A-UA-GAUCAAACCGAAU
CCCU-AGGGA---------AA------------C---CUA-GUGGAAAUGUGGUG--UGA------------
G-GA--CCU-UGU-ACGGC----------------------------------CGA-----------CU--C
A-------AAGAA--GGUG------AAUACU-UCUG------------------------------------
-------AAA-AG-CCGAG-------CCAUAGAGCGUGAUAGCCGCGUA---------ACC---G----UAA
U------UUGGAAGG-CUGC----------------UGCA-AGAGU--UCCCGAGUAAUGCGGGAUA---GU
UUUCUCGUAUGAA-CAAACCAGAUACA--CGCUGGUAAAUCUAAAUA--UAC-C-CCA-AGACCGAUAGC-G
AAA-U-AGUAGCGUGAGCAAAAG-AUGAAA-AGAAA-CCCGG-AAGGGU-C-GUG-AAAAGCGCUUGAAACC
A-AAUAGCUUAUU-AAG-GA-UGCA---GCCUGAAA-GGACAG-----------------------------
------------------------------------------------------------------------
------------------------------------------------------------------------
------------------------------------------------------------------------
------------------------------------------------------------------------
------------------------------------------------------------------------
------------------------------------------------------------AU-UCUGGACG-
---AA-----------UG-A-------------------G------UUC-----------------UG----
------------------------------------------------------------------------
-------------------------------------------------------------------GU---
------------------------------------------------------------------------
-------------------------------------------------G-AC------AG--------AAU
AUCAG-------------------------------------------------------------------
------------------------------------------------------------------------
------------------------------------------------------------------------
------------------------------------------------------------------------
------------------------------------------------A-----------------------
------------------------------------------------------------------------
------------------------------------------------------------------------
------------------------------------------------------------------------
------------------------------------------------------------------------
------------------------------------------------------------------------
-------------------GUUCAGAGG-----------------ACUAG--G---GUUGCAUCAUCAGUCU
GGAAUCAUUGGCCAGGGAGUUUA-U-AGUAGUGGCGA-GCUUAAAU--G-U--------UUAUC-------A
A-CA--UGUA--GGCAAAGGGAAAC-CGA-----------------UAU----------------------U
CCUG-CAACCUA----A-------------------------------------------------------
---UUAGCACAUUAGGACGAAUUAUGU-------ACUAAGGGGUGGUUCUGCUGCUCCAAAAGGA-CAUAAU
GAAUCCUUCUAAGGGAU--UAGG------------------------------------CUA-GAGGAA-UG
GUCU-----------------------------------UAAA--------------------------GG-
G-C-CA----------------------ACAGUCCCUGCUAUAA-UACCCGAAGCC-GGUCGAUCUAUUCCU
```

```
GGUCAAGAUGAAGUCGUU-CGAAAG-GAUGAUGGAGGUCUGC--ACCAGUGCUGACGUGCAAUUCGC-UUGG
AUGAACUGGGAAUAGGAGCAAAAGGCUAAUCUAGAUCGG-UGAUAGCUGGUUCUCCCCGAAG-UUGGUGUCA
GCCAAUCCUCAG-ACGA-------------------------------------------------------
----------------------AG-----UA-GCCAGUGGCGUAGAGC-ACUAAUU-GUG-GAGU-AAGGGG
A-A-GA-GA-UUCC-U-CCCUUCGCUCUUAAACUCCGAAACUGCUG--------------------------
------------------------------------------------------------------------
------------------G-CA-C--UG-UAGA--CUCUGGG-AGACGGG-U-UGGUGG-GCG-UAAAGUCC
AU-C-GCCGAGAA-GGGAAC---AACCUAGACUUG-G-GUUAAGGUCCCUAAGU--GC-AGG-UU-AA---G
-U-GUUAAA--CUGAAGGGCGUUAUAUUUCUAAGACAGCAAAAGGAUAGGUAUAGAAGUAGCCA-UUCCCUA
ACGAGUGUGUAACAACUCAUUUGCC-----GAGAAAUAUAGCCCCGAAAAU-GGAC-GGGGCUUA-AAUCUG
C----------CACCGAG-ACCCAAGAGCACGGG--U-----------------------------------
------------------------------------------------------------------------
-----GGAAGA-------------------------------------------------------------
------------------------------AUCCCGU-GAUC-UGGUAGGGGAGC-GUCCAGG---------
-----------U-U-GGGCA-GAAGCAG--G-UGCGUGAGUU--C-CUGUGGACCG-A-CCU-GGAGUGAAG
AUCC-UGGUAA-UAGUA-AAAGCA--UAGAU-GGGUUA--G---------------AAUCCC-AUC-CGCCG
A-AAGGGC-AAGGGUUUC-UUCGCAAUGCGUCGUCAGCGAAGAGUUAGUCGA--UC-CUAAGUGGACCCUU-
AAC-UGGU---A-ACC--------------------------------------------------------
----------------------------------------ACG-AA-A--GGGA-AACAGGUU-UAUA-UUC
CUGUA----------------------------------CCACG----GCGCUACG----A-AU--------
------------------A-----------------------------------------------------
-------GCGG--UGAC--------GCAUGG--------------------------UGUU-A-UGCU-GAC
AGUUCAG-------------GGUAAG-UGGAAUAGG-GC-------CGUCGAAUAUCU-------CCCUAU-
UUAA-A-UGUAAU-----A-A-------------------AG-----CU----------------GGUA-AA
U-A-GU--------------------GUAAA-ACUUAGACCC---------AGCUUA----AAG-CA-UGA-
AUA--GC-----------------------------------------------------------------
-----UAUCCA-AUUGUGGAAGGUU-C-CGC-U-G-AUCUCAGGACUUA-UUAAAA-A-AGC-AAAC-----
------------------------------------------------A-----CAAGGAAC-GC-CGUGAU
CGUACCCAGAACCGAC------------------------------ACAGGUGC-CCCUAGGU-GAGUAGCC
UAAGGC-GUAU-UCGU-GUAAAGGA-G-GCUAGGGAAAUCGAC-AAAUUAGAGCCGUAACUUUGG-GAUAAG
GCUUG-CUUG-C-GA--ACUUCUUC----ACGCA-GCUUCG------GCAGU--------------------
------------------------------------------------------------------------
------------------------------------------------------------------------
------------------------------------------------------------------------
------------------------------------------------------------------------
------------------------------------------------------------------------
------------------------------------------------------------------------
------------------------------------------------------------------------
------------------------------------------------------------------------
------------------------------------------------------------------------
------------------------------------------------------------------------
------------------------------------------------------------------------
------------------------------------------------------------------------
------------------------------------------------------------------------
----------------------------------------------------------GUGUAG--GGGCUC
```

```
UCA--AGUCGCAGUAACUGAGGGGGGACGA-----CUGUUUAAUAAAAACAUAGGUCUCUGAUAGAGCG---
-AA-A-------------GCUUGUUU-ACAGAGGCUGA-AACCUGGCCAGUGGCA-G-UUUGAUCAAUCCGG
GU-------------UACA-------------ACCCGAGUAAGUCC--UGCUAAAC-GCCGGGAGUAACUAU
AACUCUCUUAAGGUAGCCAAAUGCCUUGUCUGG-UAAGUUCAGACGUGCAUGAAUGGUCCAACGUUC--UCC
CUAUUGUCCCAGCC----UUC-UGACGAUUGAAAUU---AC---ACUCCGGCGAACGGUCCGGAGAUACCUG
GUGGGACGAGAAGA-CCCCGUGGAGU-UUUACU--GCAGCUU-GUUGUUGUGGUUUUGGGCUAGAUAUAGAG
CGU-AGCCGGGAGC-------------GUAGAUCCAUUCA----------------------------UUUC
--------------------------------------GG-UGGAUGGGGA-G--C-AU-CGUUGGAACA-C
CGGU-UUUCUUUCCUGGGAUCACUAACGG-C--------A-UCAC-A-------------------------
------------------------------------------------------------------------
------------------------------------------------------------------------
------------------------------------------------------------------------
------------------------------------------------------------------------
------------------------------------------------------------AA----------
------------------------------------------------UGUC-----GGACA-UCAGCAGGU
GGGCAGUUUGGCUGGGGCGGCUCUCUCUAGA-AAAGGUAUC-UAGAGAAACCA-AAGGUCAGCUCAAGCGGG
UCAGGAAUCCGCUGUA-GAGUU-UAAGAGCAA-AAGCUGGCCUGACU-GGAU--------------------
--------------UCUGAUUACUAAGGAAUCCUGAGGCUAAC-GCCUGGUCUAGCGAUCCUUA--------
---AC-A-----UACCU-CGU--------------------GU-GGGUUGUUGAGGUGAUAG-AA--AAAUU
ACCCCGGGG-GUAACAGAGUUGUCACGAGUAAGAGUCCCUAUCGACCUCGUGGCUUGCUUCCUCGAUGUCGG
CUCUUCCUAUCCUGGCAGCGCACAUAACUGCCAAGGGUGGGGCUGUACGCCCAUUAAAAGGGAACGUGAGCU
GGGUUUAGACCGUCGUGAGACAGGUCGGAUUCUAUCUACC--------------------A-GGU-------
-------------------------------AUGU--GGGAUGACUGAAGGUAAGG-GAAAAUCAGUACGAG
AGGAA-CAUUUUGCCGGCGCCUC-UGGU-G-UACCGGUUGUCU-GACA-AGGCAA-U-GCCGGGCAGCUACG
CGCUA-ACGGAUAAGUGCUGAAAGCA-UC--UAAGCACGAAGCCGGUCCUGA-A-AAU--AGUCAUCC-AA-
--UCUUAUCCCAU-AACGUUAUAAAAG----------AUGAGAUUUAUUUU------AAUUGAU-AAAGUGA
---GUAAGAUU----U-UCUGUGUAAAUCGGAUACUAACUUC----UU------------------------
-------------------------------------AGUAACUAU-CAUAAU------UAAAAUGAAUGUU
AUGUUGAAUAACAAGA--------------------------------------------------------
------------------------------------------------------------------------
------------------------------------------------------------------------
---------------------------------------GUGGGGCACG------GUGA----CGUGAACCA
----------------------------U-------------------------------------------
--------------------GGGGA-A-----------------------UAGA----CG-AGU------GC
ACCCGUAAAAGACGGGGU-UGC-UGAGAU-GGG-GGUGUACG-UAG-G--------AAG------------C
UCUCGG-------GCGACCUAUUUAGCCC--G-CCAU-UCUCAACA---GCGCGAGUG-UAUGGCA------
------------------------------------------------------------------------
------------------------------------------------------------------------
------------------------------------------------------------------------
------------------------------------AU-----------
>LSUASG_NJBH01000007.1:89956-93415/1-3275 Candidatus Lokiarchaeota archaeon
------------------------CCAAUAC----GC--CUCCGCUAUUU-GGUGAAUA-U-CUUGGC-UUG
G-G---CACCGAUGAAGAGCGUGA--CAAGCUGCGAUAAGCUCGGGUUAGUCGCAUGAAGACGC------UG
ACC-CCG-AGAUUCUCGAAUAGGACA-UCCG-----CUU-------GC----GGC----------U------
----------------------------------UAA----U-A-AUCC-GCAAGA---------------U
```

```
UCC--------C-----------GUC----------------------------------------------
--A--------------------GG-G-AA-GGGGAACCCUGGGAAGUAAAACAUUUUAGUACCAG--GAGG
AAAAGAAAACAA--UAGUGAUUCCGUAA--GU-AA-CGGCGAGCGAAAGCGGA-A-CA-GAUCAAACCGAAU
CCCA-AGGGA---------AA------------C---CUA-UGGGAAAUGUGGUG--UAU------------
G-GG--CCU-UGU-ACGGC----------------------------------CUC-----------UC--U
A-------GAGAA--GAUG------AAUACC-UCUG------------------------------------
-------GAA-UG-CUGAG-------CCAUAGAGCGUGACAGCCGCGUA---------AUC---G----UAA
U------CUAGAUGG-CUGC----------------UGCA-AGAGU--CCCUGAGUAGUGCGGGAUG---GU
UUUCUCGUACGAA-UGUGCCAGAUACA--CGCUGGCAAAUCUAAAUA--UGU-C-CCA-AGACCGAUAGC-G
UAG-U-AGUAGUGUGAACAAAAG-AUGAAA-AGUAA-CCCGG-AAGGGC-U-GUG-AAAAGUGCUUGAAACC
A-AAUAGCUUAUU-AAG-GA-UGCG---GCCUGAAA-GGACAG-----------------------------
------------------------------------------------------------------------
------------------------------------------------------------------------
------------------------------------------------------------------------
------------------------------------------------------------------------
------------------------------------------------------------------------
------------------------------------------------------------AU-UCUGCAUG-
---AA-----------UG-A-------------------G------UUC-----------------UG----
------------------------------------------------------------------------
-------------------------------------------------------------------GU---
------------------------------------------------------------------------
-------------------------------------------------G-AC------AG--------AAU
AUCAG-------------------------------------------------------------------
------------------------------------------------------------------------
------------------------------------------------------------------------
------------------------------------------------------------------------
------------------------------------------------A-----------------------
------------------------------------------------------------------------
------------------------------------------------------------------------
------------------------------------------------------------------------
------------------------------------------------------------------------
------------------------------------------------------------------------
-------------------AUACAGAGG-----------------ACUAG--G---GUCGCAUCAUCAGUCU
GGAAUCAUUGGCCAGGGAGUUUA-U-AGUAGUGGCAA-GCUUAAGC--G-U--------CUUGC-------G
G-CG--CGUA--GGCAUAGGGAAAC-CGA-----------------UAU----------------------U
CUUG-CAACCUA----A-------------------------------------------------------
---UUAUCACAUUAGGAAGACCUACUC-------GUUAACGAGUGGUUAUGCCAUUCUAACAACG-UUUAGC
UAAUCCUUCUAAGGAAU--UAGG------------------------------------CCA-GGAGAA-UG
GUCU-----------------------------------CAAA--------------------------GG-
G-C-CA----------------------AAAGUCUCUGCUAUAA-UACCCGAAGCC-GGUCGAUCUAUUCCU
GGUCAAGAUGAAGUCGUU-CGAAAG-GGCGAUGGAGGUCUGC--ACCAGUGCUGACGUGCAAUUCGC-UUGG
AUGAACUGGGAAUAGGAGCAAAAGGCUAAUCUAGAUCGG-UGAUAGCUGGUUCUCCCCGAAG-UUGGUAUCA
GCCAAGUCUCAG-ACGA-------------------------------------------------------
----------------------GG-----CA-GCCAGUGGCGUAGAGC-ACUAAUU-GCG-UGGU-AUGGGG
A-A-GA-AA-UUCC-U-CUCCCCGCUCUUAAACUCCGAAACUGCUG--------------------------
```

```
------------------------------------------------------------------------
------------------G-CA-C--UG-CAGA--CUCUGGG-AAACGGG-U-UGGUGG-GCG-UAAAGUCC
AU-C-GCCGAGAA-GGGAAC---AACCUAGACUUG-G-GUUAAGGUCCCUAAGU--GC-AGG-UU-AA---G
-U-GUUAAA--CAGAAGGGCGUUAUAUUUCUAAAACAGCAAAAGGAUAGGUAUAGAAGUAGCCA-UUCCCUA
ACGAGUGUGUAACAACUCAUUUGCC-----GAGAAAUAUAGCCCCGAAAAU-GGAC-GGGGCUUA-AAUCUG
C----------CACCGAG-ACCCAAGGGCACGGG--C-----------------------------------
------------------------------------------------------------------------
-----GGAAGA-------------------------------------------------------------
------------------------------ACCCCGU-GAUC-CGGUAGGGGAGC-GUCCGGG---------
-----------U-A-GAGUA-GAAGCAG--G-UGCGUGAGUU--C-CUGUGGAUCU-G-CCC-GGAGUGAAC
AUCC-UGGUAA-UAGUA-AAAGCA--UAGAU-GGGUUA--G---------------AAUCCC-AUC-CGCCG
A-AAGGGC-AAGGGUUUC-UUUGCAAUGCGUCGUCAGCGAAGAGUUAGUCGA--UC-CUAAGCGUACCCUU-
AAC-UGGU---A-UGC--------------------------------------------------------
----------------------------------------GCG-AA-A--GGGA-AACAGGUU-UAUA-UUC
CUGUA----------------------------------CCACG----GCACUACG----A-AU--------
------------------A-----------------------------------------------------
-------GCGG--UGAC--------GCACGA--------------------------CUUU-A-UGCU-GAC
GGUUCAG-------------GGUAGG-UGGAAUAGG-GC-------CGUCGAAUAUCU-------CCCUAU-
UUAA-A-UGC-AU-----A-A-------------------AG-----CU----------------GGUG-AA
U-A-GU--------------------GUAAA-ACUUAGACCC---------AGCUUA----AAC-CA-UGA-
AUA--GC-----------------------------------------------------------------
-----CAUCCA-AUUGUGGAAGGUU-C-CGC-C-G-AGCCCAGGACCUA-UUAAAA-A-GGC-AAAA-----
------------------------------------------------G-----UAAGAAU-GC-CGUGAU
CGUACCCAGAACCGAC------------------------------ACAGGUGC-CCCUAGGU-GAGUAGCC
UAAGGC-GUAU-UCGU-GUAAAGGA-G-GCUAGGGAAAUCGAC-AAAUUAGAGCCGUAACUUUGG-GAGAAG
GCUUG-CUUG-C-GA--ACUUCUUC----ACGCA-GCUUCG------GCAGU--------------------
------------------------------------------------------------------------
------------------------------------------------------------------------
------------------------------------------------------------------------
------------------------------------------------------------------------
------------------------------------------------------------------------
------------------------------------------------------------------------
------------------------------------------------------------------------
------------------------------------------------------------------------
------------------------------------------------------------------------
------------------------------------------------------------------------
------------------------------------------------------------------------
------------------------------------------------------------------------
------------------------------------------------------------------------
----------------------------------------------------------GUGAAG--GGGCUC
ACA--AGUUGCAAUAACUAGGGGGGGACGA-----CUGUUUAAUAAAAACAUAGGUCACUGAUAGAGUG---
-AA-A-------------ACUUGUUU-ACAGUGGCUGA-AACCUGGCCAGUGGCA-G-UCUGAUAAAUCCGG
GU-------------UACA-------------ACCCGAGUAAGUCC--UGCUAAAC-GCCGGGAGUAACUAU
AACUCUCUUAAGGUAGCCAAAUGCCUUGUCUGG-UAAGUUCAGACGUGCAUGAAUGGUCCAACGUUC--UCC
CCAUUGUCCAGCC----UUC-UGACGAUUGAAAUU---AC---ACUCCGGCGAACGGUCCGGAGAAACCCA
```

```
GUGGGACGAGAAGA-CCCCGUGGAGU-UUUACU--GCAGCCU-GUUGUUGUGGUCCUGGAUUGGAUACAGAG
UGU-AGUCGGGAGC-------------GUAGAUGCCGCGA----------------------------UUUC
--------------------------------------GG-UCACGGCGGA-G--C-AU-CAUUGGAACA-C
CGGC-UUUCCCCUCUGGUAUCACUAACGA-U--------G-CAAA-U-------------------------
------------------------------------------------------------------------
------------------------------------------------------------------------
------------------------------------------------------------------------
------------------------------------------------------------------------
------------------------------------------------------------CG----------
------------------------------------------------CAUC-----GAACA-CCCGCAGGU
GGGCAGUUUGGCUGGGGCGGCUCACUCUAGA-AACAAUAUC-UAGAGUAACCA-AUGAUCAGCUCAGGCGGG
UCAGGAAUCCGCUGAA-GAGUU-UAAGAGCAA-AAGCUGGUCUGACU-GGAU--------------------
--------------UCUGAUCACCAAGGAAUCCUGAGGCUAAC-GCCUGGUCUAGCGAUCCUUA--------
---AC-U-----GAGCU-CGU--------------------GU-GGCUGGUUAAGGUGAUAG-AA--AAAUU
ACCCCGGGG-GUAACAGAGUUGUCACGAGUAAGAGUUCCUAUCGACCUCGUGGCUUGCUUCCUCGAUGUCGG
CUCUUCCUAUCCUGGGAGCGCACAUAACUCCCAAGGGUGGGGCUGUACGCCCAUUAAAAGGGAACGUGAGCU
GGGUUUAGACCGUCGUGAAACAGGUCGGAUUCUAUCUACU--------------------G-GGU-------
-------------------------------UUGU--AGGUUGACUGAGGGUAAGG-GAAAAUCAGUACGAG
AGGAA-CAUUUUGCCGGCGCCUC-UAGU-G-UACCGGUUAUCU-GACA-AGGUAU-U-GCCGGGCAGCCACG
CGCCA-ACGGAUAAGUGCUGAAAGCA-UC--UAAGCACGAAGUCGGACCCGA-A-AAU--AGUCAGCC-AA-
--UCUCGUAUAGCGAGUGUU----------AAUGAGUGAGUGGUGUAUUUA------GGUUGGUGUUGGUUA
---GUAAGACA----CACCUGUGAUCGUCAGAAGCUAAUC--------------------------------
-----------------AU-----------------AAGAAUACAU-CAUAAU------CUAAAUGGAUUAU
UUGCGAGUUAACGAGA--------------------------------------------------------
------------------------------------------------------------------------
------------------------------------------------------------------------
---------------------------------------UCAGGGCGCG------GUGA----CGCGAACUG
----------------------------A-------------------------------------------
--------------------GUAUG-C-----------------------GAGA----CG-AGU------GC
ACCCGUAUAAGACGGGGU-UAG-UGAGGU-GGG-GGUGUACG-UAG-G--------GAG------------C
UUUCGG-------GCGACCUAUUCAGCCC--G-CCAC-UCUCAAUA---GCACGAGUG-UGUGGUG------
------------------------------------------------------------------------
------------------------------------------------------------------------
------------------------------------------------------------------------
------------------------------------U------------
>LSUASG_SDNY01000005.1:53412-56906/1-3294 Candidatus Lokiarchaeota archaeon
----------------------------------------------AUUU-GGUGGAUA-U-CUUGGC-UUG
G-G---CACCGAUGAAGAGCGUGA--CAAGCUGCGAUAAGCUUGGGUUAGUCGCACGUAGACUU------AG
ACC-CCA-AGAUUCUCGAAUCGGACA-UCCG-----CUU-------GC----GGC----------C------
----------------------------------CAA----U-A-AUCC-GCAAGA---------------U
UCC--------CAUAACCGUAAGGUG----------------------------------------------
--A--------------------GG-G-AA-GGGGAACUCUGAGAAGUAAAACAUUUUAGUAUCAG--AAGG
AAAAGAAAACAA--UAGUGAUUCCGUGA--GU-AA-CGGCGAGCGAAAGCGGA-A-CA-GGUCAAACCGAAU
CCGU-UGGAGA-------AGA------------U---CCC-UCGGAGAUGUGGUG--UAU------------
G-GA--CCU-CAU-ACGGC----------------------------------CUC-----------CC--U
```

```
A-------GAGAA--CAUG------AAAACC-UCUU------------------------------------
-------GAA-AG-UGGUG-------CCAUAGAGCGUGAUAGCCGCGUA---------AUG---G----UAA
U------CUAGAAGG-CUGA----------------UUUG-AGGUA--UCCUGAGUAGUGCGGGAUG---GU
UUUCUCGUACAAA-UGUGCCAAACACA--UGUUGGCAAACCUAAAUA--UGU-C-CCA-AGACCGAUAGC-G
UAA-U-AGUAGCGUGAGCGAAAG-AUGAAA-AGUAA-CCCGG-AAGGGU-C-GUG-AAAAGUGCUUGAAACC
A-AAUAGCUUAAU-AGG-GA-UGCA---GCCUGAAA-GGAGAG-----------------------------
------------------------------------------------------------------------
------------------------------------------------------------------------
------------------------------------------------------------------------
------------------------------------------------------------------------
------------------------------------------------------------------------
------------------------------------------------------------AU-UCUGAGCG-
---AA-----------UG-U-------------------G------UUC-----------------CG----
------------------------------------------------------------------------
-------------------------------------------------------------------GU---
------------------------------------------------------------------------
-------------------------------------------------A-AC------GG--------AAU
AUCAG-------------------------------------------------------------------
------------------------------------------------------------------------
------------------------------------------------------------------------
------------------------------------------------------------------------
------------------------------------------------A-----------------------
------------------------------------------------------------------------
------------------------------------------------------------------------
------------------------------------------------------------------------
------------------------------------------------------------------------
------------------------------------------------------------------------
-------------------GCACAGAGG-----------------ACUAG--G---GUUGCAUCAUCAGUCU
GGAAUCAUUGGCCAGGGAGUUUA-U-AGUAGAGGCGA-GCUUAAGC--G-U--------CUUAC-------G
G-CG--CGAA--GGCAUAGGGAAAC-CAA-----------------UAU----------------------U
CUUG-CAACCUA----A-------------------------------------------------------
---UUAUCACAUUAGGAAGAUCUAUUC-------GUCCUAGAUUGGUUUUGCCAUUCAAAUACGU-C-UAGU
GAAUCCUUCUAAGGGAU--UAGG------------------------------------CCA-GAAGAA-UG
GUCU-----------------------------------CAAA--------------------------GG-
G-C-CU----------------------AAAGUCUCUGUUAUAA-AACCCGAAGCC-GGUCGAUCUAUUCCU
GGCCAAGAUGAAGCCGUU-CGAAAG-GACGGUGGAGGUCUGC--AGUAGUGCUGACGUGCAAUUCGC-UUAC
AUGAGCUGGGAAUAGGAGCAAAAGGCUAAUCUAGAUCGG-UGAUAGCUGGUUCUCCCCGAAG-UUGGUAUCA
GCCAAGUCUCAG-ACGA-------------------------------------------------------
----------------------GG-----CA-GCCAGUGGUGUAGAGC-ACUAAUU-GUG-GAGC-AUGGGG
A-A-GA-AA-UUCC-U-CACUUUACUCUUAAACUCCGAAACUGCUG--------------------------
------------------------------------------------------------------------
------------------G-CA-C--UU-UAGA--CUCUGGG-AAACGGG-U-UGGUGG-GCG-UAAAGUCC
AU-C-GCCGAGAA-GGGAAC---AACCUAGACUUG-G-GUUAAGGUCCCAAAGU--GC-AGG-UU-AA---G
-U-GUUAAA--AUAAAGGGCGUUAUAUUUCUAAGACAGUAAAAGGAUAGGUAUAGAAGUAGCCA-UUCCCUA
ACGAGUGUGUAAUAACUCAUUUGCC-----GAGAAAUAUAGCCCCGAAAAU-GGAC-GGGGCUUA-AAUCUG
```

```
C----------CACCGAG-ACCCAAGGGCACGGG--U-----------------------------------
------------------------------------------------------------------------
-----AGAAGA-------------------------------------------------------------
------------------------------AUCCCGU-GAUC-CGGUAGGGGAGC-GUCCGAG---------
-----------U-U-GCAUA-GAAGCAG--G-UGCGCGAGUU--C-CUGUGGAGCG-G-CUC-GGAGUGAAG
AUCC-UGGUGA-UAGUA-AAAGCA--AAGAU-GGGUUA--G---------------AAUCCC-AUC-CGCCG
A-AAGGGC-AAGGGUUUC-UUCGCAAUGCGUCGUCAGCGAAGAGUUAGUCGG--UC-CUAAGCGCAGCCUU-
AAC-UGGU---AUUGC--------------------------------------------------------
----------------------------------------GCG-AA-A--GGGA-AACAGGUU-UAUA-UUC
CUGUA----------------------------------CCUCG----GUGCUAC-----A-CU--------
------------------A-----------------------------------------------------
-------GCGG--UAAC--------GCACGG--------------------------CGUU-G-UGCU-GAC
GGUUCCG-------------GGUAAG-UGGAAUAGG-GU-------CGUCGAAUAUCU-------CCCUAU-
UUAA-A-UGU-AU-----A-A-------------------AG-----CU----------------GGUA-AA
U-A-GA--------------------GUAAA-UCUUAGACCC---------AGCCAA----AAU-CA-UGA-
AUA--GC-----------------------------------------------------------------
-----CCCCCA-AUUGUGGGAGGUU-C-CGC-U-G-AUCCCAGGACCCA-AUGAAA-A-GGC-AAAC-----
------------------------------------------------G-----CAAGGAAC-AC-CGAGUC
CGUACCCAGAACUGACACAAGAGCUCUAUAACUGAGAGCGCUGUCAAAAGGUGC-CCCUAGGU-GAGUAGCC
UAAGGC-GUAU-UCGU-GUAAAGGA-G-GCUAGGGAAAUCGAC-AAAUUAGAGCCGUAACCUAGG-GAUAAG
GCUUG-CUUG-A-GA--ACUUCGUU----GCGC-----CUC------ACAGU--------------------
------------------------------------------------------------------------
------------------------------------------------------------------------
------------------------------------------------------------------------
------------------------------------------------------------------------
------------------------------------------------------------------------
------------------------------------------------------------------------
------------------------------------------------------------------------
------------------------------------------------------------------------
------------------------------------------------------------------------
------------------------------------------------------------------------
------------------------------------------------------------------------
------------------------------------------------------------------------
------------------------------------------------------------------------
----------------------------------------------------------GCUUCG--GGGUUU
CCA--AGUCGCAGUGACUAGGGGGGGGCGA-----CUGUUUAAUAAAAACAUAGGUCUCUGAUAGAGCG---
-AA-A-------------GCUUGUUU-ACAGAGGCUGA-CACCUGGCCAGUGGCA-G-UCUGAUAAAUCCGG
GU-------------UACA-------------ACCCGAGUAAGUCC--UGCUAAAC-GCCGGGAGUAACUAU
AACUCUCUUAAGGUAGCCAAAUGCCUUGUCUGG-UAAGUUCAGACGUGCAUGAACGGUCCAACGUCC--UCC
CUAUUGUCCCAGCC----UUC-UGACGAUUGAAAUC---AC---ACUCCGGCGAACGGUCCGGAGAAGCCCA
ACGGGACGAGAAGA-CCCCGUGGAGU-UUUACU--GCAGCCU-GUUGUUGCGGUUUUGAACUGGAUACAGAG
UGU-AGUCGGGAGC-------------GUAGAUGCUAUCA----------------------------UUUC
--------------------------------------GG-UGGUAGUGGA-G--C-GA-CUUUGGAAUA-C
CGGC-UUUCUUUUUUGAAAUCGCUAACGG-C--------G-UGAA-A-------------------------
------------------------------------------------------------------------
```

```
------------------------------------------------------------------------
------------------------------------------------------------------------
------------------------------------------------------------------------
------------------------------------------------------------AG----------
------------------------------------------------CGUC-----GGACA-CCCGCAGGU
GGGCAGUUUGGCUGGGGCGGCUCACUCCAGA-AAAAAUAUC-UGGAGUAACCA-AUGGUCAGCUCAAGCGGG
UCAGGAAUCCGCUGAA-GAGUU-UAAGAGCAA-AAGCUGGCCUGACU-GGAU--------------------
--------------UCUGAUCACCAAGGAAUCCUGAGGCGAAA-GCCUGGUCUAGCGAUCCUCU--------
---AU-U-----GUGCU-CAU--------------------GU-GGCUUGUAGAGGUGAUAG-AA--AAAUU
ACCCCGGGG-GUAACAGAGUUGUCACGAGUAAGAGCUCAUAUCGACCUCGUGGCUUGCUUCCUCGAUGUCGG
CUCUUCCUAUCCUGGGAGCGCACAUAACUCCCAAGGGUGGGGCUGUACGCCCAUUAAAAGGGAACGUGAGCU
GGGUUUAGACCGUCGUGAGACAGGUCGGAUUCUAUCUGUU--------------------G-GGC-------
-------------------------------UUGU--UGGAAGACUGAGGGUAAGG-GAAAAUCAGUACGAG
AGGAA-CAUUUUGCCGGCGCCUC-UGGU-G-UACCGGUUGUCU-GACA-AGGCAU-U-GCCGGGCAGCUACG
CGCUA-ACGGAUAAGUGCUGAAAGCA-UC--UAAGCACGAAGCCGGACCCGA-A-AAU--AGUCUUCC-AC-
--UCCUUUCUCAA-AUUGUU--------UGAUUGAUUUACUGGGACAUUUU------GGUUGGC-UGGAUUG
---AUAAGGCA----U-CCUGUGUAAGCCGGAAGUCGAUCCG----UA------------------------
---------------------------------------AAAAAGC-CAUAAU------CGGAAUGGUCGGU
GAGUUGGUUGACGUGC--------------------------------------------------------
------------------------------------------------------------------------
------------------------------------------------------------------------
---------------------------------------GUGGGGGCUG------GUGA----CAGUAACCA
----------------------------C-------------------------------------------
--------------------AGGGA-A-----------------------GGGA----CG-AGU------GC
ACCCAUAAAAGAUGGGGU-UGG-UGAGAC-GGG-GGUGUACG-UGC-C--------GAG------------C
UUUCGG-------GCGAGGUAUUUAGCCC--G-CCGU-UCUCAAUA---GCACGAGUG-UUUGGUG------
------------------------------------------------------------------------
------------------------------------------------------------------------
------------------------------------------------------------------------
-------------------------------------------------
>LSUASG_NODE_1830_length_806_cov_332.076416_g1424_i0_REV/1-806
------------------------------------------------------------------------
------------------------------------------------------------------------
------------------------------------------------------------------------
------------------------------------------------------------------------
------------------------------------------------------------------------
------------------------------------------------------------------------
------------------------------------------------------------------------
------------------------------------------------------------------------
------------------------------------------------------------------------
------------------------------------------------------------------------
------------------------------------------------------------------------
------------------------------------------------------------------------
------------------------------------------------------------------------
------------------------------------------------------------------------
```

```
----------------------------------------------------------------------
----------------------------------------------------------------------
----------------------------------------------------------------------
----------------------------------------------------------------------
----------------------------------------------------------------------
----------------------------------------------------------------------
----------------------------------------------------------------------
----------------------------------------------------------------------
----------------------------------------------------------------------
----------------------------------------------------------------------
----------------------------------------------------------------------
----------------------------------------------------------------------
----------------------------------------------------------------------
----------------------------------------------------------------------
----------------------------------------------------------------------
----------------------------------------------------------------------
----------------------------------------------------------------------
----------------------------------------------------------------------
----------------------------------------------------------------------
----------------------------------------------------------------------
----------------------------------------------------------------------
----------------------------------------------------------------------
----------------------------------------------------------------------
----------------------------------------------------------------------
----------------------------------------------------------------------
----------------------------------------------------------------------
----------------------------------------------------------------------
----------------------------------------------------------------------
----------------------------------------------------------------------
----------------------------------------------------------------------
----------------------------------------------------------------------
----------------------------------------------------------------------
----------------------------------------------------------------------
----------------------------------------------------------------------
----------------------------------------------------------------------
----------------------------------------------------------------------
----------------------------------------------------------------------
----------------------------------------------------------------------
----------------------------------------------------------------------
----------------------------------------------------------------------
----------------------------------------------------------------------
----------------------------------------------------------------------
----------------------------------------------------------------------
----------------------------------------------------------------------
----------------------------------------------------------------------
```

```
------------------------------------------------------------------------
------------------------------------------------------------------------
------------------------------------------------------------------------
------------------------------------------------------------------------
------------------------------------------------------------------------
------------------------------------------------------------------------
------------------------------------------------------------------------
------------------------------------------------------------------------
------------------------------------------------------------------------
------------------------------------------------------------------------
------------------------------------------------------------------------
------------------------------------------------------------------------
------------------------------------------------------------------------
------------------------------------------------------------------------
------------------------------------------------------------------------
------------------------------------------------------------------------
------------------------------------------------------------------------
------------------------------------------------------------------------
------------------------------------------------------------------------
------------------------------------------------------------------------
------------------------------------------------------------------------
------------------------------------------------------------------------
------------------------------------------------------------------------
------------------------------------------------------------------------
------------------------------------------------------------------------
------------------------------------------------------------------------
------------------------------------------------------------------------
------------------------------------------------------------------------
------------------------------------------------------------------------
------------------------------------------------------------------------
------------------------------------------------------------------------
------------------------------------------------------------------------
------------------------------------------------------------------------
------------------------------------------------------------------------
------------------------------------------------------------------------
------------------------------------------------------------------------
------------------------------------------------------------------------
------------------------------------------------------------------------
------------------------------------------------------------------------
------------------------------------------------------------------------
------------------------------------------------------------------------
------------------------------------------------------------------------
------------------------------------------------------------------------
------------------------------------------------------------------------
------------------------------------------------------------------------
```

```
------------------------------------------------------------------------
------------------------------------------------------------------------
------------------------------------------------------------------------
------------------------------------------------------------------------
------------------------------------------------------------------------
---------------------------------------------------------------ACGUGAGCU
GGGUUUAGACCGUCGUGAGACAGGUCGGAUUCUAUCUACU--------------------G-GGU-------
-------------------------------AUGU--GGGUUGACUGAGGGUAAGG-GAAAAUCAGUACGAG
AGGAA-CAUUUUGCCGGCGCCUC-UAGU-G-UACCGGUUAUCU-GACA-AGGUAU-U-GCCGGGCAGCCACG
CGCCA-ACGGAUAAGUGCUGAAAGCA-UC--UAAGCACGAAGCCGGACCCGA-A-AAU--AGUCAGCC-AA-
--UCUCGAUCAAU-AGUGCUAUGCUCUCAUGA-GCGUAUUUCAGUUGGUUG------AAUUGAC-AAGGC--
---------AG----U-CCUGUGAAAGCCGUAAGUCAGUUUGGAGUGUAGAGGUAGAACUUGGAUAUUAGAG
GAGACUUUAA-----UGUUUGAGUUUUGUCAGAAGCAUAA--AGCC-AUA--AU----UGUAAUGUGUGUUG
AGUGUUAAUAGCGAGA--------------------------------------------------------
------------------------------------------------------------------------
------------------------------------------------------------------------
---------------------------------------GCAGGGCGCG------GUGA----CGCGAACUG
----------------------------U-------------------------------------------
--------------------GGAAU-C-----------------------GAGA----CG-AGU------GC
ACCCGUAUAAGACGGGGU-UAG-UGAGGU-AGG-GGUGUACG-UAG-G--------AAG------------C
UUUCGG-------GCGACCUAUUUAGCCC--G-CUAC-UCUCAAUA---GCACGAGUG-UAUGGCUGUCGAA
AGAGGAGCAUAAGGUUCAGGCUUAGUCUGAAUCUUUCCUCAACUAGACGCCAAUACGCCUCCGCUAUUUGGU
GGAUAUCUUGGCUUGGGAACCGAUGAAGAGCGUGACAAGCUGCGAUAAGCUCGGGUUAGUCGCAUGAAGACG
UUGACCCCGAGAUUCUCGAAUAGGACAUCCGCUUGCGAACCAAUAAUUCGCAAGAUUCCCCCAU--------
---GGGAAGGGGAACCCUGAGAAGUAAAACAUUUUAGUAUCAGGAGGAA
>LSUASG_NODE_1615_length_777_cov_447.117808_g1100_i2/1-777
------------------------------------------------------------------------
------------------------------------------------------------------------
------------------------------------------------------------------------
------------------------------------------------------------------------
------------------------------------------------------------------------
------------------------------------------------------------------------
------------------------------------------------------------------------
------------------------------------------------------------------------
------------------------------------------------------------------------
------------------------------------------------------------------------
------------------------------------------------------------------------
------------------------------------------------------------------------
------------------------------------------------------------------------
------------------------------------------------------------------------
------------------------------------------------------------------------
------------------------------------------------------------------------
------------------------------------------------------------------------
------------------------------------------------------------------------
------------------------------------------------------------------------
```

```
--------------------------------------------------------------------------------
--------------------------------------------------------------------------------
--------------------------------------------------------------------------------
--------------------------------------------------------------------------------
--------------------------------------------------------------------------------
--------------------------------------------------------------------------------
--------------------------------------------------------------------------------
--------------------------------------------------------------------------------
--------------------------------------------------------------------------------
--------------------------------------------------------------------------------
--------------------------------------------------------------------------------
--------------------------------------------------------------------------------
--------------------------------------------------------------------------------
--------------------------------------------------------------------------------
--------------------------------------------------------------------------------
--------------------------------------------------------------------------------
--------------------------------------------------------------------------------
--------------------------------------------------------------------------------
--------------------------------------------------------------------------------
--------------------------------------------------------------------------------
--------------------------------------------------------------------------------
--------------------------------------------------------------------------------
--------------------------------------------------------------------------------
--------------------------------------------------------------------------------
--------------------------------------------------------------------------------
--------------------------------------------------------------------------------
--------------------------------------------------------------------------------
--------------------------------------------------------------------------------
--------------------------------------------------------------------------------
--------------------------------------------------------------------------------
--------------------------------------------------------------------------------
--------------------------------------------------------------------------------
--------------------------------------------------------------------------------
--------------------------------------------------------------------------------
--------------------------------------------------------------------------------
--------------------------------------------------------------------------------
--------------------------------------------------------------------------------
--------------------------------------------------------------------------------
--------------------------------------------------------------------------------
--------------------------------------------------------------------------------
--------------------------------------------------------------------------------
--------------------------------------------------------------------------------
--------------------------------------------------------------------------------
--------------------------------------------------------------------------------
--------------------------------------------------------------------------------
```

```
--------------------------------------------------------------------------------
--------------------------------------------------------------------------------
--------------------------------------------------------------------------------
--------------------------------------------------------------------------------
--------------------------------------------------------------------------------
--------------------------------------------------------------------------------
--------------------------------------------------------------------------------
--------------------------------------------------------------------------------
--------------------------------------------------------------------------------
--------------------------------------------------------------------------------
--------------------------------------------------------------------------------
--------------------------------------------------------------------------------
--------------------------------------------------------------------------------
--------------------------------------------------------------------------------
--------------------------------------------------------------------------------
--------------------------------------------------------------------------------
--------------------------------------------------------------------------------
--------------------------------------------------------------------------------
--------------------------------------------------------------------------------
--------------------------------------------------------------------------------
--------------------------------------------------------------------------------
--------------------------------------------------------------------------------
--------------------------------------------------------------------------------
--------------------------------------------------------------------------------
--------------------------------------------------------------------------------
--------------------------------------------------------------------------------
--------------------------------------------------------------------------------
--------------------------------------------------------------------------------
--------------------------------------------------------------------------------
--------------------------------------------------------------------------------
--------------------------------------------------------------------------------
--------------------------------------------------------------------------------
--------------------------------------------------------------------------------
--------------------------------------------------------------------------------
--------------------------------------------------------------------------------
--------------------------------------------------------------------------------
--------------------------------------------------------------------------------
--------------------------------------------------------------------------------
--------------------------------------------------------------------------------
--------------------------------------------------------------------------------
--------------------------------------------------------------------------------
--------------------------------------------------------------------------------
--------------------------------------------------------------------------------
--------------------------------------------------------------------------------
--------------------------------------------------------------------------------
```

--------------------------------------------------------------AACGUGAGCU
GGGUUUAGACCGUCGUGAGACAGGUCGGAUUCUAUCUGUU--------------------G-GGC-------
-------------------------------UUGU--CGGGGGAUUGAGGGUAAGG-GAAAAUCAGUACGAG
AGGAA-CAUUUUGCCGGCGCCUC-UAGU-G-UACCGGUUGUCU-GACA-AGGCAU-C-GCCGGGCAGCCACG
CGCUA-ACGGAUAAGUGCUGAAAGCA-UC--UAAGCACGAAGCCGGACCCGA-A-AAU--AGUCAGCC-AA-
--UCUUAUUCCGU-AAUGUUGGGCUUUAUGGUUGGGUAUUCUCGUUGAUGUU----GGGAGGU--AAGAC--
---------UG----C-C-UGUGAUGGUCGGAAGCCUUCCAAUCGAAUCC----------------------
-----UUAUU-----GCAUG--CUGCAGUGUGGAGGAAAG--CAUC-AUA--AU----GGGAAUGUUCAGUG
UAAAGUACCGACGCGA--------------------------------------------------------
------------------------------------------------------------------------
------------------------------------------------------------------------
---------------------------------------GCGGGGCGCG------GUGA----CGCGAACCG
----------------------------U-------------------------------------------
--------------------GGGAA-C-----------------------GAGA----CG-AGU------GC
ACCCGUAAAAGACGGGGU-UGG-UGAGGU-AGG-GGUGUACG-UAG-G--------AAG------------C
UUUCGG-------GCGACCUAUUCAGCCC--G-CUAC-UCUCAACA---GCACGAGUG-UAUGGCUGUCGAA
AGAGGAGCAUAAGGUUCAGGCUUAGUCUGAAUCUUUCCUCAACUAGACGCCAAUACGCCUCCGCUAUUUGGU
GGAUAUCUUGGCUUGGGAACCGAUGAAGAGCGUGACAAGCUGCGAUAAGCUCGGGUUAGUCGCAUGAAGACG
CUGACCCCGAGAUUCUCGAAUAGGACAUCCGCUUGCGAACCAAUAAUUCGCAAGAUUCCCCCAU--------
---GGGAAGGGGAACCCUGAGAAGUAAAACAUUUUAGUAUCAGGAGG--
>LSUASG_NODE_2003_length_785_cov_204.440379_g1570_i0_REV/1-785
------------------------------------------------------------------------
------------------------------------------------------------------------
------------------------------------------------------------------------
------------------------------------------------------------------------
------------------------------------------------------------------------
------------------------------------------------------------------------
------------------------------------------------------------------------
------------------------------------------------------------------------
------------------------------------------------------------------------
------------------------------------------------------------------------
------------------------------------------------------------------------
------------------------------------------------------------------------
------------------------------------------------------------------------
------------------------------------------------------------------------
------------------------------------------------------------------------
------------------------------------------------------------------------
------------------------------------------------------------------------
------------------------------------------------------------------------
------------------------------------------------------------------------
------------------------------------------------------------------------
------------------------------------------------------------------------
------------------------------------------------------------------------
------------------------------------------------------------------------
------------------------------------------------------------------------

```
------------------------------------------------------------------------
------------------------------------------------------------------------
------------------------------------------------------------------------
------------------------------------------------------------------------
------------------------------------------------------------------------
------------------------------------------------------------------------
------------------------------------------------------------------------
------------------------------------------------------------------------
------------------------------------------------------------------------
------------------------------------------------------------------------
------------------------------------------------------------------------
------------------------------------------------------------------------
------------------------------------------------------------------------
------------------------------------------------------------------------
------------------------------------------------------------------------
------------------------------------------------------------------------
------------------------------------------------------------------------
------------------------------------------------------------------------
------------------------------------------------------------------------
------------------------------------------------------------------------
------------------------------------------------------------------------
------------------------------------------------------------------------
------------------------------------------------------------------------
------------------------------------------------------------------------
------------------------------------------------------------------------
------------------------------------------------------------------------
------------------------------------------------------------------------
------------------------------------------------------------------------
------------------------------------------------------------------------
------------------------------------------------------------------------
------------------------------------------------------------------------
------------------------------------------------------------------------
------------------------------------------------------------------------
------------------------------------------------------------------------
------------------------------------------------------------------------
------------------------------------------------------------------------
------------------------------------------------------------------------
------------------------------------------------------------------------
------------------------------------------------------------------------
------------------------------------------------------------------------
------------------------------------------------------------------------
------------------------------------------------------------------------
------------------------------------------------------------------------
------------------------------------------------------------------------
------------------------------------------------------------------------
```

```
------------------------------------------------------------------------
------------------------------------------------------------------------
------------------------------------------------------------------------
------------------------------------------------------------------------
------------------------------------------------------------------------
------------------------------------------------------------------------
------------------------------------------------------------------------
------------------------------------------------------------------------
------------------------------------------------------------------------
------------------------------------------------------------------------
------------------------------------------------------------------------
------------------------------------------------------------------------
------------------------------------------------------------------------
------------------------------------------------------------------------
------------------------------------------------------------------------
------------------------------------------------------------------------
------------------------------------------------------------------------
------------------------------------------------------------------------
------------------------------------------------------------------------
------------------------------------------------------------------------
------------------------------------------------------------------------
------------------------------------------------------------------------
------------------------------------------------------------------------
------------------------------------------------------------------------
------------------------------------------------------------------------
------------------------------------------------------------------------
------------------------------------------------------------------------
------------------------------------------------------------------------
------------------------------------------------------------------------
------------------------------------------------------------------------
------------------------------------------------------------------------
------------------------------------------------------------------------
------------------------------------------------------------------------
------------------------------------------------------------------------
------------------------------------------------------------------------
------------------------------------------------------------------------
------------------------------------------------------------------------
------------------------------------------------------------------------
------------------------------------------------------------------------
------------------------------------------------------------------------
---------------------------------------------------------------ACGUGAGCU
GGGUUUAGACCGUCGUGAGACAGGUCGGAUUCUAUCUACU--------------------G-GGU-------
-------------------------------AUGU--GGGUUGACUGAGGGUAAGG-GAAAAUCAGUACGAG
AGGAA-CAUUUUGCCGGCGCCUC-UAGU-G-UACCGGUUGUCU-GACA-AGGCAC-C-GCCGGGCAGCCACG
CGCUA-ACAGAUAAGUGCUGAAAGCA-UC--UAAGCACGAAGCUGGCCCCGA-A-AAG--AGUCUUCC-A--
```

```
--UUCCAUUUCUA-UAGUGGACAUUGAUUUGAGAGGUAUUUUGGUUGGUGUU----GGGUUGAU-AAGGU--
---------CC----C-C-UGUGAAGACCGGAAGUCAGCCCUGAAUCGUAUUCUG-------------CGGU
GAAACGUGGG-----AUUG----------------CGAGGUAAAGCAUCAUAAU----UGAAGUGUUGAUUA
AA-UUGGUGGUCACGU--------------------------------------------------------
------------------------------------------------------------------------
------------------------------------------------------------------------
---------------------------------------GCGGGGCGCG------GUGA----CGCGAACCG
----------------------------U-------------------------------------------
--------------------GAGAG-A-----------------------GGGA----CG-AGU------GC
ACCCGUAAAAGACGGGGU-UGG-UGAGAC-GGG-GGUGUACG-UGC-G--------GAG------------C
UUUCGG-------GCGAGGUUUUUAGCCC--G-CCGU-UCUCAAAC---GCACGAGUG-UGUGGUG-UUGAA
AGAGGA-CUCGUGGUUCAGGCUUUGUCUGAACCUUUCCUUAAGUAGAUGCCAUUACGCCUCCGCUAUUUGGU
GGAUAUCUUGGCUUGGGUACCGAUGAAGAGCGUGACAAGCUGCGAUAAGCUUGGGUUAGUCGCACGUAGACU
UAGACCCCAAGAUUCUCGAAUAGGACAUCCACUUGUGGCCUAAUAAUCCACAAGAUUCCCAUAUCCGAAAGG
UGAGGGAAGGGGAACCCUGAGAAGUAAAACAUUUUAGUAUCAG------
>LSUASG_SHMU01000207.1:1-2097/1-1912 Candidatus Lokiarchaeota archaeon isol
------------------------------------------------------------------------
------------------------------------------------------------------------
------------------------------------------------------------------------
------------------------------------------------------------------------
------------------------------------------------------------------------
------------------------------------------------------------------------
------------------------------------------------------------------------
------------------------------------------------------------------------
------------------------------------------------------------------------
------------------------------------------------------------------------
------------------------------------------------------------------------
------------------------------------------------------------------------
------------------------------------------------------------------------
------------------------------------------------------------------------
------------------------------------------------------------------------
------------------------------------------------------------------------
------------------------------------------------------------------------
------------------------------------------------------------------------
------------------------------------------------------------------------
------------------------------------------------------------------------
------------------------------------------------------------------------
------------------------------------------------------------------------
------------------------------------------------------------------------
------------------------------------------------------------------------
------------------------------------------------------------------------
------------------------------------------------------------------------
------------------------------------------------------------------------
------------------------------------------------------------------------
------------------------------------------------------------------------
```

```
------------------------------------------------------------------------
------------------------------------------------------------------------
------------------------------------------------------------------------
------------------------------------------------------------------------
------------------------------------------------------------------------
------------------------------------------------------------------------
------------------------------------------------------------------------
------------------------------------------------------------------------
------------------------------------------------------------------------
------------------------------------------------------------------------
------------------------------------------------------------------------
------------------------------------------------------------------------
------------------------------------------------------------------------
------------------------------------------------------------------------
------------------------------------------------------------------------
------------------------------------------------------------------------
------------------------------------------------------------------------
------------------------------------------------------------------------
------------------------------------------------------------------------
------------------------------------------------------------------------
------------------------------------------------------------------------
------------------------------------------------------------------------
------------------------------------------------------------------------
------------------------------------------------------------------------
------------------------------------------------------------------------
------------------------------------------------------------------------
------------------------------------------------------------------------
------------------------------------------------------------------------
--------------------------------------GAUC-UGGUAGGAAGGC-GUCCGGG---------
-----------C-G-GGGUA-GAAGCAG--G-UGCGUGAGUU--C-CUGUGGACCG-G-CCU-GGAGUGAAG
AUCC-UGGCGA-CAGUA-AAAGCA--AAGAU-GGGUUA--G---------------AAUCCC-AUC-CGCCG
A-AAGGGC-AAGGGUUUC-UUCGCAAUGCGUCGUCAGCGAAGAGUUAGUCGA--UC-CUAAGCGUAUCCUU-
AAC-UGGC---G-UAC--------------------------------------------------------
----------------------------------------GCG-AA-A--GGGA-AACAGGUU-UAUA-UUC
CUGUA----------------------------------CCACG----GCACUACG----A-AU--------
------------------------------------------------------------------------
-------GCGG--UGAC--------GCACGG--------------------------CCUG-A-UGCU-GAC
AGUUUUG-------------GGUAGG-UGGAAUAGG-GC--------UGUCGAAUAUCU-------CCCUAU-
UUAA-A-UGC-AU-----A-A-------------------CG-----CU----------------GGCG-AA
U-A-GC--------------------GUAAA-GCUUAGACCC----------AGCCGA----ACC-CA-UGA-
AUA--AC-----------------------------------------------------------------
-----CAUCCA-AAUGUGGACGGUU-C-CGC-C-G-AGCCCAGGACUUA-AUAAAA-A-GGC-ACAG-----
------------------------------------------------G-----CAAGAACU-GC-CGUGAU
CGUACCCAGAACCGAC------------------------------ACAGGUGC-CCCUAGGU-GAGUAGCC
UUAGGC-GUGU-UCGU-GUAAAGGA-G-GCUAGGGAAAUCGAC-AAAAUAGAGCCGUAACUUUGG-GAUAAG
```

```
GCUUG-CUUG-C-GA--ACUUCCGC----ACGCA-UU-----------CAGU--------------------
------------------------------------------------------------------------
------------------------------------------------------------------------
------------------------------------------------------------------------
------------------------------------------------------------------------
------------------------------------------------------------------------
------------------------------------------------------------------------
------------------------------------------------------------------------
------------------------------------------------------------------------
------------------------------------------------------------------------
------------------------------------------------------------------------
------------------------------------------------------------------------
------------------------------------------------------------------------
------------------------------------------------------------------------
----------------------------------------------------------GUGCGG--GGGCUC
UCA--AGUCGCAGUGACUGGGGGGGGACGA-----CUGUUUAAUAAAAACAUAGGUCUCUGAUAGAGUG---
-AA-A-------------ACUUUUUU-ACAGAGGCUGA-UACCUGGCCAGUGCCA-G-UCAGAUAAACACGG
GU-------------UACA-------------ACCCGAAUAAGUCC--UGGUAAAC-GCCGGGAGUAACUAU
AACUCUCUUAAGGUAGCCAUAUGCCUUGUCUGG-UAAGUUCAGACGUGCAUGAACGGUCCAACGUUC--UCC
CCAUUGUCCCAGCC----UUC-UGACGAUUGAAAUC---AC---ACUCCGGCGAACGGUCCGGAGAUACCUA
GUGGGACGAGAAGA-CCCCGUGGAGU-UUUACU--GCAACUU-GUUGCUGUAGAUUGGAAUGGGAUACAGAG
UGU-AGUCGGGAGC-------------GGAGAUGCCUUAA----------------------------UUUC
--------------------------------------GG-UUAAGGCGGA-G--C-GA-UCUUGGAAUA-C
CGGC-UUUCUCAUUCUGGUCUGCUAACGG-C--------G-UCAU-G-------------------------
------------------------------------------------------------------------
------------------------------------------------------------------------
------------------------------------------------------------------------
------------------------------------------------------------------------
-------------------------------------------------------A----GG----------
------------------------------------------------CGUC-----GGACA-CCAGCAGGC
GGGCAGUUUGGCUGGGGCGGCUCACUCCAGA-AACAAUAUC-UGGAGUAACCA-AAGGUCAGCUCAGGCGGG
CCAGGAAUCCGCUGUA-GAGUU-UAAGAGCAA-AAGCUGGCCUGACU-GGAU--------------------
--------------UCUGAGCACCAAGGAAUCCUGAGGCUAAC-GCCGGGUCUAGCGAUCCUUA--------
---AC-A-----AUGUU-CAU--------------------GU-GGCUUGUUGAGGUGAUAG-AA--AAAUU
ACCCCGGGG-GUAACAGAGUUGUCACGAGUGAGAGUUCCUAUCGACCUCGUGGCUUGCUUCCUCGAUGUCGG
CUCUUCCUAUCCUGGGAGCGCACAGAACUUCCAAGGUGGGGCUGUACGCCCAUUAAAAGGGAACGUGAGCU
GGGUUUAGACCGUCGUGAGACAGGUCGGAUUCUAUCUACU--------------------A-GGU-------
-------------------------------ACGU--GGGGAGACUGAGGGUAACG-GAAAAUCAGUACGAG
AGGAA-CAUUUUGCGGGCGCCUC-UGGU-G-UACCGGUUGUCU-GACA-AGGCAC-U-GCCGGGCAGCUACG
CGCCA-ACGGAUAAGUGCUGAAAGCA-UC--UAAGCACGAAGCCGGUCCCGA-A-AAG--AGUCUCCC-AC-
--UCUCGUUUUAUAGAUGUA--------AAGCUUAUUGUUGGGUAUGUAUUCUGACGAGUUGAUAUGAAACA
CCUGUGAGUUU----C-UCUAUUGACUUGAGGUGCCGAGUAUUGCGUGUUAC--------------------
-----------------GU-----------------GAUGAGGCGUCCGUCAUAGGAUACAUGCCGGAGGCG
AUGAGCGUUGCAGAGA--------------------------------------------------------
------------------------------------------------------------------------
```

```
------------------------------------------------------------------------
---------------------------------------GCGGGGUGCG------GUGA----CGCAUACCG
----------------------------U-------------------------------------------
--------------------GGGAAGC-----------------------GAGA----CG-AGC------GC
ACCCGUAAAAGACGGGGU-UAG-UGAGGC-AGG-GGUGUACG-UGG-A--------AAG------------C
UUUCGG-------GCGAUCCAUUCAGCCC--GUCUGC-UCUCAAUA---GCGCGAGUG-UGUGCUA------
------------------------------------------------------------------------
------------------------------------------------------------------------
------------------------------------------------------------------------
-------------------------------------------------
>LSUASG_F3H4_B5_WNEK01000002.1:3579-251/1-3300 23S ribosomal RNA  no intron
----------------------CGCCAAUAC----GC--CUACGCUAUUU-GGUGGAUA-U-CUUGGC-UUG
G-G---CACCGAUGAAGAGCGUGA--CAAGCUGCGAUAAGCUCGGGUUAGUCGCAUGUAGACGU------UU
ACC-CCG-AGAUUCUCGAAUCGGACA-UCCG-----CUU-------GC----GGC----------C------
----------------------------------CAA----U-A-AUCC-GCAAGA---------------U
UCC--------C-----------GUC----------------------------------------------
--A--------------------GG-G-AA-AGGGAACCCUGGGAAGUAAAACAUUUUAGUACCAG--GAGG
AAAAGAAAACAA--UCGUGAUUCCGUUA--GU-AA-CGGCGAGCGAAAGCGGA-A-CA-GAUCAAACCGAAU
CCCU-UGGGGA-------UAA------------C---CCA-AGGGAGAUGUGGAG--UGU------------
G-GA--CCU-CGU-ACGGC----------------------------------CUC-----------CC--C
A-------AAAAA--GGUG------AAUACC-CCUG------------------------------------
-------AAA-AG-UCGAG-------CCAUAGAGCGUGAUAGCCGCUUA---------ACC---G----UAA
U------UUGGAAGG-CUGC----------------UGCG-AGGUA--UCCCGAGUAGUGCGGGAUA---GU
UUUCUCGUACGAA-UGUGCCAGAUACA--CGCUGGCAAAUCUAAAUA--UGU-C-CCA-AGACCGAUAGU-G
AAA-U-AGUAGCGUGAGCAAAAG-AUGAAA-AGAAA-CCCGG-AAGGGU-C-GUG-AAAAGCGCUUGAAACC
A-AAUAGUUUAGU-AAG-GA-UGCA---GCCCGAAA-GGAUAG-----------------------------
------------------------------------------------------------------------
------------------------------------------------------------------------
------------------------------------------------------------------------
------------------------------------------------------------------------
------------------------------------------------------------------------
------------------------------------------------------------AU-UCUGGACG-
---AA-----------UG-A-------------------G------UUC-----------------UG----
------------------------------------------------------------------------
-------------------------------------------------------------------GU---
------------------------------------------------------------------------
-------------------------------------------------G-AC------AG--------AAU
AUCAG-------------------------------------------------------------------
------------------------------------------------------------------------
------------------------------------------------------------------------
------------------------------------------------------------------------
------------------------------------------------A-----------------------
------------------------------------------------------------------------
------------------------------------------------------------------------
------------------------------------------------------------------------
```

```
------------------------------------------------------------------------
------------------------------------------------------------------------
-------------------GUUCAGAGG-----------------ACUAG--G---GUUGCAUCAUCAGUCU
GGAAUCAUUGGCCAGGGAGUUUA-U-AGUAGUGGCGA-GCUUAAGU--G-U--------CUCGC-------G
G-CA--CGUA--GGCAUAGGGAAAC-CAA-----------------UAU----------------------U
CUUG-CAAUCUA----A-------------------------------------------------------
---UUAUCACAUUAGGACGAUUUGCGU-------ACCACGGAGUAGUUUUGCUAUUCC-AAGGGC-CGUAAU
GAAUCCUUCUAAGGAGU--UAGA------------------------------------CUA-GAAGAA-UG
GUCU-----------------------------------CAAA--------------------------GG-
G-C-CU----------------------AAAGUCACUGCUGUAA-UACCCGAAGCC-GGUCGAUCUAUUCCU
GGUCAAGAUGAAGUCGCU-CGAAAG-GGUGAUGGAAGUCUGC--ACCAGUGAUGAUGUGCAAUUCUC-UUGG
AUGAACUGGGAAUAGGAGCAAAAGGCUAAUCUAGAUCGG-UGAUAGCUGGUUCUCCCCGAAG-UUGGUGUCA
GCCAAUCCUCAG-GCGA-------------------------------------------------------
----------------------GG-----UC-GCUAGUGGUGUAGAGC-ACUAAUU-GCA-AAGU-AAGGGG
A-A-GA-AA-UUCC-U-CGCUUCGCUCUUAAACUCCGAAUCUGCUA--------------------------
------------------------------------------------------------------------
------------------G-CA-C--UG-UAGA--CCCUGGG-AGACGGG-U-UAGUGG-GCG-UAAAGUCC
AU-U-GCCGAGAA-GGGAAC---AACCUAGACUUG-G-GUUAAGGUCCCUAAGU--GC-AGG-UU-AA---G
-U-GUUAAA--AGAAAGGGCGUUAUAUUUCUAAAACAGCAAAAGGAUAGGUAUAGAAGUAGCCA-CUCCCUA
ACGAGUGUGUAACAACUCAUUUGCC-----GAGAAAUAUAGCCCCGAAAAU-GGAC-GGGGCUUA-AAUCUG
C----------CACCGAG-ACCCAAGAGCACGGG--U-----------------------------------
------------------------------------------------------------------------
-----GGAAGA-------------------------------------------------------------
------------------------------AUCCCGU-GAUC-UGGUAGGGGAGC-GUCCAGG---------
-----------U-U-GGGCA-GAAGCAG--G-UGCGUGAGUU--C-CUGUGGACCG-U-CCU-GGACCGCAG
AUCC-UGGUAA-UAGUA-AAAGCA--UAGAU-GGGUUA--G---------------AAUCCC-AUC-CGCCG
A-AAGGGC-AAGGGUUUC-UUCGCAAUGCGUCGUCAGCGAAGAGUUAGUCGA--UC-CUAAGCGCACCCUU-
AAC-UGGU---A-UGC--------------------------------------------------------
----------------------------------------GCG-AA-A--GGGA-AACAGGUU-UAUA-UUC
CUGUA----------------------------------CCACG----GUGCUACA----A-AU--------
------------------A-----------------------------------------------------
-------GCGG--UAAC--------GCACGG--------------------------CUGU-G-UGCU-AAC
AAUUCCG-------------GGUAAG-UGAAAUAGG-GU--------CGUCGAAUAUCU-------CCCUAU-
UUAA-A-UGUAAU-----A-A-------------------AG-----CU----------------GGUA-AA
U-A-GU--------------------GUAAA-ACUUAGACCC---------AGCCCA----AAG-CA-UGA-
AUA--GC-----------------------------------------------------------------
-----CAUCCA-AUUGUGGAAGGUU-U-CGC-U-G-AUCCCAGGAUUUA-UUAAAA-A-GGC-AACA-----
------------------------------------------------G-----CAAGAAAC-AC-CGUGUU
CGUACCCAGAACCGAC------------------------------ACAGGUGC-CCCUAGGU-GAGUAGCC
UAAGGC-GUAU-UCGU-GGAAAGGA-G-GCUAGGGAAAUCGAC-AAAUUAGAGCCGUAACUUUGG-GAUAAG
GCUUG-CUUG-A-GA--ACUUCUUC----ACGCA-GCUUCG------GCAGU--------------------
------------------------------------------------------------------------
------------------------------------------------------------------------
------------------------------------------------------------------------
------------------------------------------------------------------------
```

```
------------------------------------------------------------------------
------------------------------------------------------------------------
------------------------------------------------------------------------
------------------------------------------------------------------------
------------------------------------------------------------------------
------------------------------------------------------------------------
------------------------------------------------------------------------
------------------------------------------------------------------------
------------------------------------------------------------------------
----------------------------------------------------------GUGCGG--GGGCUC
UCA--AGUCGCAGUAACUAGGGGGGGACGA-----CUGUUUAAUAAAAACAUAGGUCACUGAUAGAGCG---
-AA-A-------------GCUUGUUU-ACAGUGGCUGA-AACCUGGCCAGUGGCA-G-UUUGAUAAAUCCGG
GU-------------UACA-------------ACCCGAGUAAGUCC--UGCUAAAC-GCCGGGAGUAACUAU
AACUCUCUUAAGGUAGCCAAAUGCCUUGUCUGG-UAAGUUCAGACGUGCAUGAAUGGUCCAACGUUC--UCC
CUAUUGUCCCAGCC----UUC-UGACGAUUGAAAUU---AC---ACUCCGGCGAACGGUCCGGAGAUACCUG
GUGGGACGAGAAGA-CCCCGUGGAGU-UUUACU--GCAGCCU-GUUGUUGUGGUCUUGAGCUGGAUACAGAG
UGU-AGUCGGGAGC-------------GUAGAUGCCGUAA----------------------------UUUC
--------------------------------------GG-UUACGGCGGA-G--C-AA-CUUUGGAAUA-C
CGGC-UUUCCUUCUUGAAAUCACUAACGG-C--------G-CU-U-A-------------------------
------------------------------------------------------------------------
------------------------------------------------------------------------
------------------------------------------------------------------------
------------------------------------------------------------------------
------------------------------------------------------------AG----------
------------------------------------------------CGUC-----GGACA-UCCGCAGGU
GGGCAGUUUGGCUGGGGCGGCUCUCUCUAGA-AAAGAUAUC-UAGAGAAACCA-AAGGUCAGCUCAGGCGGG
UCAGGAAUCCGCUGAA-GAGUU-UAAGAGCAA-AAGCUGGCCUGACU-GGAU--------------------
--------------UCUGAUUACCAAGGAAUCCUGAGGCUAAC-GCCUGGUCUAGCGAUCCUUA--------
---AC-U-----GUGCU-CGU--------------------GU-GGCUGGUUGAGGUGAUAG-AA--AAAUU
ACCCCGGGG-GUAACAGAGUUGUCACGAGUAAGAGUUCCUAUCGACCUCGUGGCUUGCUUCCUCGAUGUCGG
CUCUUCCUAUCCUGGGAGCGCACAUAACUCCCAAGGGUGGGGCUGUACGCCCAUUAAAAGGGAACGUGAGCU
GGGUUUAGACCGUCGUGAGACAGGUCGGAUUCUAUCUACC--------------------A-GGU-------
-------------------------------AUGU--GGGAUGACUGAGGGAAAGG-GAAAAUCAGUACGAG
AGGAA-CAUUUUGCCGGCGCCUC-UGGU-G-UACCGGUUGUCU-GACA-AGGCAU-C-GCCGGGCAGCUACG
CGCUA-ACUGAUAAGUGCUGAAAGCA-UC--UAAGCACGAAGCAGAUCCCGA-A-AAG--AGUCAUCC-AA-
--UCUUACUUCAU-AUUGUUCUUGAGGCGAGUCGAUUAUUUCGUUUAGUUU------GGUUGUU-AAGGU--
---------UC----C-CCUGUGAGGGCCGGAUGACGACCUG----CUUGCGCGU----------------G
UAAAAGCGCG-----UAAG-----------------AAGAAAAGGC-UAUAAA------CGGAAUGUUUGGC
UUGCGGAGAGACGAGA--------------------------------------------------------
------------------------------------------------------------------------
------------------------------------------------------------------------
---------------------------------------GCGGGGCGCG------GUGA----CGCGAACCG
----------------------------U-------------------------------------------
--------------------GGGAG-C-----------------------GAGA----CG-AGC------GC
ACCCGUAAAAGACGGGUU-UGG-UGAGAU-GGG-GGUGUACG-UGC-C--------AAG------------C
```

```
UUUCGG-------GCGAGGUAUUCAGCCC--G-CCAU-UCUCAACA---GCGCAAGUG-UGUGGCA------
------------------------------------------------------------------------
------------------------------------------------------------------------
------------------------------------------------------------------------
-------------------------------------------------
>LSUe_9606_HUMAN/1-5227
---------------------------------------CGACUCUUAGC-GGUGGAUC-A-CUCGGC-UCG
U-G---CGUCGAUGAAGAACGCAG--CUAGCUGCGAGAAUUAAUGUGAAU--U---GCA--GGA-CA--CAU
UGAUCAUCGACACUUCGAACGCA-CU-U--------GC--------GG----CCCC-G-GGU---------U
C-C-----------UCC---C----G----G---GGC----U-A-CGCC-UGU-----CUGAGCGUCG-CUU
-----------------------------------------------------------CGCGAC-CUCA-G
AUC--------------------AG-A-CG-UGGCGACCCGCUGAAUUUAAGCAUAUUAGUCAGCG--GAGG
AGAAGAAACUAA--CCAGGAUUCCCUCA--GU-AA-CGGCGAGUGAACAGGGA-A-GA-GCCCAGCGCCGAA
UCCC-CGCCCCGCG-GC-GGG------------G---CGC--GGGACAUGUGGCG--UA-------------
--CG-GAAG-ACC-C--GCUC-------------------------CCCGG--CGC-----------CG---
CUCGUG--GGGGG--CCC-------AAGUCC-UUCU------------------------------------
-------GAU-CG-AGGCCC---AGCCCGUGGACGGUGUGAGGCCGGUA---------GCG---G----CCC
CCG---GC--GCGCC-GGGC----------------CCGG-GUCUU--CCCGGAGUCGGGUUGCUUG---GG
AAUGCAGCCCAAA-G-CGGGUGGUAAA-CUCCAUCUAAGGCUAAAUACCGG--C-ACG-AGACCGAUAGU-C
AAC-A-AGUACCGUAAGGGAAAG-UUGAAA-AGAAC-UUUGA-AGAGAG-A-GUUCAAGAGGGCGUGAAACC
G-UUAAGAGG--U-AAA-CG-GGUG---GGGU-CCGCGCAGU--------------------CC--------
-------------------------------------------GCCCGGAG--GAUUCAACCC--GGCG---
------GCG--GGUCCGGCCGUGU-CGGCGG--CCCGGCG-GAUCUUUCCCGCCCCCCGUUCCUCCCGACCC
CUCCACCCGCCC--------------UCCCUUCC-CC-CGCCGCCC---CUCCUCC----UCCUCCCC----
-----------------------------------------GGAG-GGGGCGGGCUCCGGCGGGUGC---GG
GGGUGGGCGGGCGGGGCCGGGGGUGGGGUCGGCGGGGGACCG-UCCCCCG--ACCG-GCGACCGGCCGCCGC
C-----GGGCGCA-UUUCC---ACCG-CG--GC----------GGU--------GCGCCGCGACCGGCUCC-
---GG-----------GA-C-------------------G------GCU-----------------GGGAAG
GCC-----------------CGGC-------GGGGAAGGUGGCUCGGGG-------------------GGCC
CCGUCCGUCCGUCCGUCCGUCCUCCUCCUCCCCCGUCUCCGCCCCCCGGCCCCGCGUCCUCCC--U-CGGGA
GGGCGCGCGGGUCGGGGCGGCGGCGGCGGCGGCGGUGGCGGCGGCGGCGGC---------GGCGGCGGGACC
GAAACCCCCCCCGAGUGUUACAGCCCCCCCGGCAGCAGC------ACUCG-CCGA--AUCC--------CGG
GGCCGAGGGAGC--GAGACCCGUC----------------GCCGCGCUCUCCCCCCU--------CCCGGCG
CC------------------CACCCCCGCGGGGA---------------------------------AUCCC
CCG-----------------------------------CGAGGGGGGUC----------------------U
CC-----CCCGCGGGGG---CG-----------CGC------------------CGGC--------------
GUCUCC-------------------------------------UCGUGGGGGGC-----CGGGCCA------
-CCCCUC------------CCACGGCGC-------GACCGCUCUCCCACCCCUCCUCCCCGCG----CCCCC
GCCCCGGCGACGGGGG--GGGUGCCGCGCGCGGGUCGGGGGGC---G-GGGCG----GACUGUC--------
-----CCCAGUG--CGC-CCC---GG-----GCG-GGUC--------GCGCCGUC--GGGCCCGGGGG---A
GGUUCUCUCGGGGCCACGCGCGCG--------------------------------------------UCCC
CC-----------GAAGAGGGGGACGGCGGAG-CGAGCGCA-------------------------------
----------------CGGGGUCGGCGG------------CGACGUCGGCU-A---CCCACCCGACCCGUCU
UGAAACACGGACCAAGGAGUCUA-A-CACGUGCGCGA-GUCGGGGG--C-UCG-------CACG------AA
A-GC--CGCCGUGGCGCAAUGAAGG-UGA-----------------AGGCCGGCGCGCU--CGCCGGCCGAG
```

```
GUGG-GAUCCCG----AGGCCUC-----UC--------------------------CAGUCCGC--------
CGA---------------------------------------------------------------------
-------------------GGG--------------------------------------CGCACCACC-GG
CCCGUC----UCGCCCG-C--------------------CGCG----------------CCGGGGAGG-UG-
GAG-CA----------------------CGAGCGCACGUGUUAG-GACCCGAAAGA-UGGUGAACUAUGCCU
GGGCAGGGCGAAGCCAGA-GGAAAC-UCUGGUGGAGGUCCGU--AGCGGUCCUGACGUGCAAAUCGG-UCGU
CCGACCUGGGUAUAGGGGCGAAAGACUAAUCGAACCAUC-UAGUAGCUGGUUCCCUCCGAAG-UUUCCCUCA
GGAUAGCUGGCG-CUCU-----------------------------------------------------CG
CAGACCCGACGCACCCCCGCCACG-----CAGUUUUAUCCGGUAAAGCGAAUGAUU-AGA-GGUCUUGG-GG
C-C-GAAAC-GAUC-U-CAACCUAUUCUCAAACUUUAAAUGGGUAAGAA-----------------------
------------------GCCCG-GC-----UCG-CU--GGCG--U--GGA---------------------
GC---CGGG-C---GUGGA-AU-G--C-----G--AGUGCCU-AGUGGGC-C-ACUU-U-UGG-UAAGCAGA
AC-U-GGCGCUGC-GGGAUG---AACCGAACGCCG-G-GUUAAGGCGCCCGAUG--CCGACG-CU-CA---U
-C-AGACCCC-AGAAAAGGUGUUGGUUGAUAUAGACAGCAGGACGGUGGCCAUGGAAGUCGGAA-UCCGCUA
AGGAGUGUGUAACAACUCACCUGCC-----GAAUCAACUAGCCCUGAAAAU-GGAU-GGCGCUGG-AGCGUC
G----------GGCCCAU-ACCCGGCCGUCGCCG--GCAGUCGAGAGNGGACGGG-----AGCGGCGGGGGC
GGCGCGCGCNCGCGCGCGUGUG---------GUGUGCGUCGGAGGGCGGCGGCGGCGGCGGCGGCGGGGGUG
---UGGGGUCCUUCCCC-CGCCCCCC--CCC---CCACGCCUCCUCC--CCUCCUCCCGCC--CAC--GCCC
CGCUCCCCGCCCCCGGAGCCCCGCGGACGCUACGCCG-CGAC-GAGUAGGAGGGCCGCUGCGG---------
-----------U-GAGCCUU-GAAGCCUA-G-GGCGCGGGCC--C-GGGUGGAGCC-G-CCG-CAGGUGCAG
AUCU-UGGUGG-UAGUA-GCAAAU---AUUC-AAACGA--G---------------AACUUU-GAA-GGCCG
AAGUGGAG-AAGGGUUCC-AUGUGAACA-GCAGUUGAACAUGGGUCAGUCGG--UC-CUGAGAGAUGGGCG-
A-G-CGCC-----GUUCCGA--------AGGGACGGGCG----AUGG---CCUC-----------------C
GU---------------UGCCCUC------GGCC----GA-UC-GA-A-AGGGA-GUCGGGU-CAGA-UCC
CCGAA----------------------------------UCCGGAGUGGCG------------GAGAUGGGC
G----------------CC-GCGA-----------------------------------GGCGUCC------
--AGUG-CGG---UAAC--------GC-GA-----CC--------------------GAUC-C-CGGA-GAA
GCCGGCG-------------GGAGCC-CCGGGGAGA-GU------UCUCU-------------U-UUCUUU-
GUGAAG-GGC-AG-----G-G-------------------CG-----CC----------------CUGG-AA
U-G-GG--------------------UUCGC-CCCGAGAGAG---------GGGCCC----GUG-CC-UUG-
GAA--AG-----------------------------------------------------------------
----CGUCGCG-GUUCCGGCGGCGU-C-CGGUGAG-CUCUCGCUGGC-CCUUGAAA-A-UCC-GGGGGAGAG
-G-------------------------------------------GUGU-----AAAUCUCG-CG-CCGGGC
CGUACCCAUAUCCGCA------------------------------GCAGGUC--UCCAAGGU-GAACAGCC
UCUGGCAU-GU-UGGA-ACAAUGUA-G-GUAAGGGAAGUCGGC-AAGCCGGAUCCGUAACUUCGG-GAUAAG
GAUUG-GCUC-U-AA--GGGCUG-G----GUCGG-UC------GGGCUG--GGG-CGCGAA-GC--GGG-GC
UGGG------CGCGCGCCGCG---GC----------------------------UGGACGAGGC-GCCG--C
CGCC-CCCCCC-ACGCCCGG-GGC-ACC--CCCCUCG-----CGGCCCUCCCCC-----GCCCCACCCCGCG
-CGCGCCGCUCGC-------UCCCUCCCCGCCCCGCGCCCUCUCUCUCUCUCUC--UCCCCCGCUCC-----
------------------------CCGUC------------------------C-UCCCCCCUCC-----CC
GGGGG-------------AGCG---CCGCGU-------------------GGGG--GCGGC-GGCGGGG-GG
AGAAGGG-------UCGGGGC-------G--GCAGGG-----GCCGGCGGCG---G-CCCGCCGCGG--GGC
CCCGG---CGGCGGGGGCACGGUCCCCCGCGAGG-------GGG-GCCCGG-GCACCCGGGGGG--CCGGCG
-G-CGGCGGCGACUCUGGACGCGAG-CCGG-GCCCUUC----------------------------------
---CCG----UG-GAUCGC-----CC-CAGCUGCG-GCGGGCGUCGCG-GCCGCCCCCGGGGAGC--CCGGC
```

```
GGGCGCCGGCGCGCCCCCCCCCCCACCCCA---------CGUCUCGUCGCGCGCGUCCGCUGGGGGCGGG
GAGCGGU-----------------CGGG-CGGCGG--CGGUCGGCGGGCGGCGGGGC--GGGG---------
---------------CGGUUCGUCCCCCGCCCUACCCCCC-GGCCCCGUCCGCCCCCCGUUCCCCCUCC
UCCUCGGCGCG--CGGCGGCGGCGGCGGCAGGCGGCGGAGGGGCCGCGGGCCGGUCC-CCCCCGCCGGUCC
GCCCCCGGGGCCGCGGU--UCCGCGCGGCGCCUC-GCCU----CGGCCG-GCG--CCUAGCAGC--CGACUU
AGA--ACUGGUGCGGACCAGGGGAAUCCGA-----CUGUUUAAUUAAAACAAAGCAUCGCGAAGGCCCG---
-CG-G------------CGGGUGUUG-ACGCGAUGUGA-UUUCUGCCCAGUGCUCUG-AAUG-UCAAAGUGA
A--------------GAAA--------------UUCAAUGAAGCGC--GGGUAAAC-GGCGGGAGUAACUAU
GACUCUCUUAAGGUAGCCAAAUGCCUCGUCAUC-UAAUUAGUGACGCGCAUGAAUGGAUGAACGAGA--UUC
CCACUGUCCCUACC----UAC-UAUCCAGCGAAACC---AC---AGCCAAGGGAACGGGCUUGGCGGAAUCA
GCGGGGAAAGAAGA-CCCUGUUGAGC-UUGACU--CUAGUCU-GGCACGGUGAAGAGACAUGAGAGGUGUAG
AAU-AAGUGGGAGGCCCCC---------GGCGC--------------------CCCCCGGUGUCCCCGCGA
GGGG-C-CCGGGGCGG----------------GGUCCGCCGGCCCUGCGGG-C--C-GC-CGGUGAAAUA-C
CACUACUCUGAUCGUUUUUUCACUGACCC-GG-------U-GA-G-GCGGG----GGG-----------GC-
------GA--GC-CC--------CGAG-GGGCU-CU-CG-------------CUUCU---------------
-----------GGCG--------------------------------------CCAA--GC---------GC
------------------C--------------------------------------------CG-------
---GCCG------------CGC---------GCC---GGC-----CGG------------------------
----------GCG------CGA------------------------------CCCG----CU----------
------------------------------------------------CCGG-----GGACA-GUGCCAGGU
GGGGAGUUUGACUGGGGCGGUACACCUGUCA-AACGGUAAC-GCAGGUGUCCU-AAGGCGAGCUCAGGGAGG
ACAGAAACCUCCCGUG-GAGCA-GAAGGGCAA-AAGCUCGCUUGAUC-UUGA--------------------
--------------UUUUCA--GUACGAAUACAGACCGUGAAA-GCGGGGCCUCACGAUCCUUC--------
---UG-A-----CCUUUUGGG--------------------UU-UUAAGCAGGAGGUGUCAG-AA--AAGUU
ACCACAGGG-AUAACUGGCUUGUGGCGGCCAAGCGUUCAUAGCGACGUCGCUUUUUGAUCCUUCGAUGUCGG
CUCUUCCUAUCAUUGUGAAGCA-GAAUUCACCAAGCGUUGGAUUGUUCACCCACUAAUAGGGAACGUGAGCU
GGGUUUAGACCGUCGUGAGACAGGUUAGUUUUACCCUACU--------------------G-AUG-------
-------------------------------AU--G-UGUUGUUGCCAUGGU-AAU-CCUGCUCAGUACGAG
AGGAA-CCGCAGGUUCAGACAUU-UGGU-GUAUGUGCUUGGCU-GAGG-AGCCAA-U-GGGGCGAAGCUACC
AUCUG-UGGGAUUAUGACUGAACGCC-UC--UAAGUCAGAAUCCCGCCCAG-----GC--GGAACGAUACGG
CAGCGCCG-CGGAGCCU-------------------------------------------------------
------------------------------------------------------------------------
------------------------------------------------------------------------
---------------------CGGUUGGCCUCG----------GAUAGCCGGUCCCCCGCCUGUCCCCGCCG
GCGGGCCGCCCCCCCCCUCCACGCGCCC---CGCGCG---C--GCGGGAGGGCGC-------GUG-------
---CCCCG--CC-------GC--GCGCCGGGACCGGGUCCG------------------GUGCGGA---G-
--UGCCCUUCGU--CCU--GG-----GAAACGGG-----GCGCGGCCG-------GAGAGG--CGGCCGCCC
CC--------------------------U----C-GCCCGUCACGC-----ACCGNACGUU----------C
G-----------------UGGGGAA---------------------C-CUGGCGCU-----AAA------CC
AUUCGUAGACGACCUGCU-UCU-GGGUCGGGGU-UUCGUACG-UAGCA--------GAGCAGC-UC------
----CCU--CGCUGCGAUCUAUUGAAAGUCAG-CCCU-CGACACAAG--GGUUUGU-C--------------
------------------------------------------------------------------------
------------------------------------------------------------------------
------------------------------------------------------------------------
-------------------------------------------------
```

```
>LSUe_45157_CYAME/1-3640
---------------------------AAUC----UA--CAGCUCCUGGC-GGUGGAUA-U-CUAGGC-UCU
C-G---CAGCGAUGAAGAACGCAG--CAAAGCGCGAUAGUUGGUGCGAUU--U---GCA--GGU-UU--CGU
GAAUCAUCGAGUCUUUGAACGCAAGU-G--------GC--------GC----UCGC-G-GG---------UU
C-G------------CU---C----G----C---GAG----C-A-UGCC-UGU-----UUGAGUGUCA-GCA
ACCUUGCUCCU----------------CCGCGGCGGGCGCGGGU------------G--CGUGAC-CUCA-A
AUC--------------------AG-G-CA-AGAUGACCCGCUGAAUUUAAGCAUAUCACUAAGCG--GAGG
AGAAGAAAAUAA--CCAUGAUUCCCCUA--GU-AG-CGGCGAGCGAAGCGGGA-G-GA-GCUCACGCUGGGA
ACCG-GGCGCCU------CGC------------G---CGU--CCGGGUUGUAGCC--UU-------------
--GC-GAGA-CGG-------C-------------------------GCCAG--UGC-----------CG---
CG------CCGGA--CCC-------AAGUCG-CUUG------------------------------------
-------GAA-GA-GCGCAU------CCGGAGAGGGUGAAAGUCCCGUC---------GGU---G-G-UCCG
GAGG-GCG--GUGCG-CAGCG---------------GGCC-GCGCU--CUACGAGUCGGGUUGUUUG---GG
AAUGCAGCCCCAA-G-UGGGUGGUAAA-CUUCACCUAAGGCUAAAUACCGG--C-GAG-AGACCGAUAGC-G
AAC-A-AGUACCGUGAGGGAAAG-AUGAAA-AGCAC-UUUGA-AAAGAG-G-GUGAAA-AGCGCUUGAAACC
G-CCAGGAGG--G-AAG-CG-AAGG---GAGU-CGA-UG-----------------------CC--------
-------------------------------------------GCGCCGGGCCG------CGC--GCCG---
------------------CAGC---UGC----AAGA-CGCCAU-----CCGGC-------------------
----------------------------------------------------------------GCC-----
------------------------------------------------------------------------
----------------------------------GCCAGGCAACUGG---------------CAGCAGCGGG
C-----GCG-------CGC---CCGG--CCUGC----------GG----------------G-UCCGCGUC-
---GG--------------------------------------------------------------CUCGC
GGC-----------------GCAGGAGCG--------GCUGCUGGCAAGGUAGCCGGCC----C--------
-------------------------------------------------------------GG--U-CG---
------------------------------------------------------------------------
----------------------------CCGGGUCUGGC------GCUAU--AGGCCAGCGGAGCCGGCUGU
GCGGCGGCC---------------------------------------------------------------
------------------------------------------------------------------------
------------------------------------------------------------------------
------------------------------------------------------------------------
-------------------------------------------------GGGG-------------------
----------------------CGACGC--------------------------------------------
-------------------------------------------------CUCG-----------GC------
---AA---------CGG-------G----------------------GCGC---------------------
------------------------------------------------------------------------
------------------------------------------------------------------------
-----------------GACGCGGGCGC-----------------CAUGG--C---UCUCUCCGACCCGUCU
UGAAACACGGACCAAGGAGUCUA-C-CGCGCAAGCGA-GUAUUCGG--G-C---------GGCAA--------
C-CC--CGAG--UGCGCAACGAAAG-UGG-----------------AGGGCGCGCCC---AGCGCGCCCGAG
CUGG-GAUCGGC----GGCGGAG---------------------------------------CG--------
CCG---------------------------------------------------------------------
-------------------CCG--------------------------------------UGCACCAGC-GG
CCGGCC----AU-GAAC----------------------UUGCA----------------GUGAACGG-CCC
GCG-CG----------------------GGAGCCUGCGCGGUGG-GACCCGAAAGA-UGGUGAACUAUGCCU
```

```
GAGUAGGGCGAAGCCAGA-GGAAACUUCUGGUGGAGGCUCGU--AGCGAUUCUGACGUGCAAAUCGA-UCGU
CGAACUUGGGUAUAGGGGCGAAAGACUCAUCGAACCAUC-UAGUAGCUGGUUCCCUCCGAAG-UUUCCCUCA
GGAUAGCCGGAG-CAGG-------------------------------------------------------
-------------------CGGUC-----CAGUUUUACCGGGUAAAGCGAAUGACU-CGA-GGCCUCGG-GG
G-A-GCGCA-UCCC-U-CGACCUAGUCUCAAACUCUAAAUCGGUAAGGA-----------------------
------------------GCGCG-CG-----CUG-CU--UUGG--U--GAG---------------------
CG---CGCG-C---GGACA-AU-G--GA----C--UGCUCCG-AGUGGGC-C-AUUU-U-UGG-UAAGCAGA
AC-U-GGCGAUGC-GGGAUG---CUCCGAAAGCGG-C-GUUAAGGUGCCGAGUG--GC-ACG-CU-CA---U
-GCAGACCCC-ACAAAAGGUGUUGAUGCAUUUAGACAGCAGGACGGUGGCCAUGGAAGUCGGCA-CCCGCUA
AGGAGUGUGUAACAACUCACCUGCC-----GAAUGCCUCAGCCCUGAAAAU-GGAU-GGCGCUCC-AGCGUG
C----------AACCGAC-ACGCCGCCACCGGCG--CGAG--------------------------------
------------------------------------------------------------------------
------------------------------------------------------------------------
---------------------------CGGU-UGCGC-CGGU-GUGUAGGAGGGC-GUGGGGC---------
-----------C-C-GUGCC-GAAGCGCC-G-GGCGCGAGCC--U-GCGUGGAACG-G-GCC-CCAGUGCGG
AUCU-UGGUGG-GAGUA-GCAAAU---AUUC-AAGUGC--G---------------AUCCUU-GAA-GGCCG
AAGGGGAG-AAGGGUUCC-AUGUGAACC-GCGAUGGGACAUGGGUGAGUCGG--UC-CUAAGGGGCAGGCC-
A-A-CGCC-----GAUGCGA--------GUG-----GCG-----CG--------------------------
-----------------UGAGCGC-------CGC----AC-CC-GA-A-AGGGA-AUCGGGUU-AACA-UUC
CCGAA----------------------------------CCGGGC---GACG--------G-AG--------
G----------------UA-----------------------------------------------------
----CG-CGG---CGAC--------GC-CC-----AG--------------------GAGC-C-UGGG-GAC
GCCGGCG-------------CGAGUC-CCGGCGAGC-GU------UGUCU-------------G-UGCUUG-
UUGACU-GCC--U-----G-A-------------------CG-----CC----------------CUGG-AA
G-C-AG--------------------CUGAC-CUGGAGAUAG---------GGCCGA----CUG-GC-AGG-
CAG--AG-----------------------------------------------------------------
-----CCCUGC-AAGUGCGCGGGGU-C-CGGUGCG-CUCGCGACGGC-CCGUGAAA-A-CCC-AGGGGACGG
-A---------------------------------------------UU-----UCCUCUCG-UG-CCCGGC
CGUACCCAUAACCGCA------------------------------UCCGGUC--UCCAAGGU-GAGCAGCC
UCUGGC-C-CA-UAGA-ACAAUGUA-G-AUAAGGGAAGUCGGC-AAAAUGGAUCCGUAACCUCGG-GAAAAG
GAUUG-GCUC-U-AA--GGGUUG-G----GCCGG-GG------GCGCCGGCGC---------GC--GGC---
-CGC------C------UGCG---GC------------------------------G--------CCUG---
----------------------------------------------------------------------CG
-CCU----------------AGC-------------------------------------------------
---------------------------------------------------------GA-----------GC
GAUC--------------GC-------------------------------GGG--GC--------------
---------------------------------------------------------------GCGG-----
------------------------------------------------------------------------
-G-CG---------------CU--G-CG---GGUGUGC----------------------------------
---GCG----------CGG-----CG--GG------------------------------------------
----------------------------------------------------------------AGCG----
--------------------------------------------------------G--GCGC---------
---------------G------------------------------AGCC---C------------------
------------------------------------------------------------------------
----GC-----------------------GUG-----------CCGCCG-GCG--GCGAACAAC--CGACUU
```

```
AGA--ACUGGUACGGACAAGGGGAAUCCGA-----CUGUUUAAUUAAAACAUAGCAUUGCGAUGGUCGG---
-AA-A-------------CGAUGCUG-ACGCAAUGUGA-UUUCUGCCCAGUGCUCUG-AAUG-UCAAAGUGA
A--------------GAAA--------------UUCAACCAAGCGC--GGGUAAAC-GGCGGGAGUAACUAU
GACUCUCUUAAGGUAGCCAAAUGCCUCGUCAUC-UAAUUAGUGACGCGCAUGAAUGGAUCAACGAGA--UUC
CCACUGUCCCUAUC----UAC-UAUCUAGCGAAACC---AC---AGCCAAGGGAACGGGCUUGGCAGAAUCA
GCGGGGAAAGAAGA-CCCUGUUGAGC-UUGACU--CUAGUCC-GACUCUGUGAAACGAGUUAGGAGGUGUAG
CAU-AAGUGGGAGCCG----------------------------------------------------CUCG
-------------------------------------------------CG-G--C-GA-UCCUGAAAUA-C
CACUACUCCUAGUCUUGUUUUACUUACUC-CA-------U-GA-G-GCAGA------G-----------AC-
------GG--GC-GA--------AAG---CCCU-G--C---------------UUCA---------------
-----------AAAG--------------------------------------CCAC--GC---------CG
------------------------------------------------------------------------
---GCG---------------------------C---CGC--------------------------------
----------CGG------CCA------------------------------UCUG----CG----------
------------------------------------------------UGGA-----GGACA-AGGUCAGGU
GGGGAGUUUGGCUGGGGCGGCACAUCUGCUA-AACGAUAAC-GCAGAUGUCCU-AAGGUGAGCUCAAUGAGA
ACGGAAAUCUCAUGUG-GAACA-AAAGGGUAA-AAGCUCACUUGAUU-UGGA--------------------
--------------UUUUCA--GUACGAAUACAAACUGUGAAA-GCAUGGCCUAUCGAUCCUUU--------
---GG-A-----CGUUCGAAG--------------------UU-UGAAGCCAGAGGUGUCAGAAA--AAGUU
ACCACAGGGAAUAACUGGCUUGUGGCAGCCAAGCGUUCAUAGCGACGUUGCUUUUUGAUCCUUCGAUGUCGG
CUCUUCCUAUCAUUGUGAAGCA-GAAUUCACCAAGUGUUGGAUUGUUCACCCACUAAUAGGGAACGUGAGCU
GGGUUUAGACCGUCGUGAGACAGGUUAGUUUUACCCUACU--------------------G-AUG-------
-------------------------------CG--U-CGUUGUCGCGAUAGU-AAU-CCAGCUUAGUACGAG
AGGAA-CCGCUGGUUCAGAUACU-UGGC-UC-AGCACUUGGCU-GAAC-AGCCAG-U-GGUGCGACGCUACC
AUCUG-UCGGAUUAUGGCUGAACGCC-UC--UAAGCCAGAAUCCGUGCUAG-----AA--GCGACGAU-ACC
UUUCUCGC-CCGGACCG-------------------------------------------------------
------------------------------------------------------------------------
------------------------------------------------------------------------
---------------------CUUCGAGCCGGU-----------CUA-------------------------
-------------------------------GGGCGG------------CGCGC------------------
----------GC-------CG--CC-----------GCCCCA------------------GGGCCAU---A-
------GAGCAC-------GG-----CGUCGGGU---G-CUGCGCGUC-------GAAGCG--CGCGCGGCG
CC--------------------------C----C-GCCG-CAG--------AGCGCC-AUU----------G
C-----------------ACGGCAC---------------------G-GGCGGGAC-----AAA------UC
GCCUGUAGACGACUUGAA-UG--GGAACGCAGG-GAUGUACG-CAUCU--------GAGUAGC-CU------
----UGC--UGCUACGAAAUGCUGAGUCUUAG-CUGC-UGUUCCGG---GAUUUGUGA--------------
------------------------------------------------------------------------
------------------------------------------------------------------------
------------------------------------------------------------------------
-------------------------------------------------
>LSUe_10116_RAT/1-4942
---------------------------------------CGACUCUUAGC-GGUGGAUC-A-CUCGGC-UCG
U-G---CGUCGAUGAAGAACGCAG--CUAGCUGCGAGAAUUAAUGUGAAU--U---GCA--GGA-CA--CAU
UGAUCAUCGACACUUCGAACGCA-CU-U--------GC--------GG----CCCC-G-GGU---------U
C-C-----------UCC---C----G----G---GGC----U-A-CGCC-UGU-----CUGAGCGUCG-CU-
```

```
-----------------------------------------------------------CGCGAC-CUCA-G
AUC--------------------AG-A-CG-UGGCGACCCGCUGAAUUUAAGCAUAUUAGUCAGCG--GAGG
AAAAGAAACUAA--CCAGGAUUCCCUCA--GU-AA-CGGCGAGUGAACAGGGA-A-GA-GCCCAGCGCCGAA
UCCC-CGCCGCGCG-CC-GCG------------G---CGC--GGGAAAUGUGGCG--UA-------------
--CG-GAAG-ACC-C--ACUC-------------------------CCCGG--CGC-----------CG---
CUCGUG--GGGGG--CCC-------AAGUCC-UUCU------------------------------------
-------GAU-CG-AGGCCC---AGCCCGUGGACGGUGUGAGGCCGGUA---------GCG---G-----CC
CCG---GC--GCGCC-GGGC----------------CCGG-GUCUU--CCCGGAGUCGGGUUGCUUG---GG
AAUGCAGCCCAAA-G-CGGGUGGUAAA-CUCCAUCUAAGGCUAAAUACCGG--C-ACG-AGACCGAUAGC-C
AAC-A-AGUACCGUAAGGGAAAG-UUGAAA-AGAAC-UUUGA-AGAGAG-A-GUUCAAGAGGGCGUGAAACC
G-UUAAGAGG--U-AAA-CG-GGUG---GGGU-CCGCGCAGU--------------------CC--------
-------------------------------------------GCCCGGAG--GAUUCAACCC--GGCG---
------GCGC-GCGCCGGCCGG-C-CGGUGGU-CCCGGCG-GAUCUUUCCCGCUCCCCGUUCCUCCCGACCC
CUCCACCCGCGCGUCUCUCUCCCCCCUCCCCGCGUCC-CGCCGUCGCCGUCCCCGCUCCUCCCUCCGG----
-----------------------GGGGGUGUCGGCGGGCGCUCCG-GCGGCGGG---CGCGGGGUGUGGUGG
GGGCGCGCGGGCGGGGCCGGGGGUGGGGUCGGCGGGGGACCG--CCCCCG--GUCG-GCGACCGGCCGCCGC
C-----GGGCGCA-CUUCC---ACCG-UG--GC----------GGU--------GCGCCGCGACCGGCUCC-
---GG-----------GA-C-------------------G------GCU-----------------GGGAAG
GCC-----------------CGGC-------GGGGAAGGUGGCUCGGGG-------------------GGGG
CGG--CGU-----------------------------------------------------CA--C-CCG--
-----------------------------------------------------------UGGGCGCCGGACC
--ACCCCGCCCCGAGUGUUACAG-CCCCCCGGCAGCAGC------GCUCG-CCGA--AUCC--------CGG
GGCCGAGGGAGCCGGAUACCCGUC----------------GCCGCGCUCUCCCCCG---------GCCUCUC
C---------------------CCUCCC---------------------------------------GCCCC
UCC-----------------------------------CCGUGGG---------------------------
-----------UGGCGGA--AA-----------GGG------------------GGGCG-------------
GUCGC--------------------------------------------GGGGGC----CGGGCCG------
-CCCCUC------------CCACGGCGC-------GACCGCUCUCCCACCCCCCGUCGCCUC------CGUCG
-UCCCUCU--CGGGGGUCCGG--------G-GGCCCGGGGGGC---G-GGGCG----GACUGUC--------
-----CCCAGUG--CGC-CCC---GG-----GCGUCGUC--------GCGCCGUC--GGG-CCCGGGG---G
G--------GCCGUC-GUCACGCG--------------------------------------------CUCU
CCCUCCCCUUCUCGGGGUGGGGGGGAGCGAAGCCGAGCGCA-------------------------------
----------------CGGGGUCGGCGG------------CGAUGUCGGC--U---ACCACCCGACCCGUCU
UGAAACACGGACCAAGGAGUCUA-A-CGCGUGCGCGA-GUCAGGGG--C-UCG-------UCCG------AA
A-GC--CGCCGUGGCGCAAUGAAGG-UGA-----------------AGGGCCCCGUUCCCGGGGGCCCCGAG
GUGG-GAUCCCG----AGGCCUC-----UC--------------------------CAGUCCGC--------
CGA---------------------------------------------------------------------
-------------------GGG--------------------------------------CGCACCACC-GG
CCCGUC----UCGCCCG-C--------------------CGCG----------------CCGGGGAGG-UG-
GAG-CA----------------------CGAGCGUACGCGUUAG-GACCCGAAAGA-UGGUGAACUAUGCUU
GGGCAGGGCGAAG-CAGA-GGAAAC-UCUGGUGGAGGUCCGU--AGCGGUCCUGACGUGCAAAUCGG-UCGU
CCGACUUGGGUAUAGGGGCGAAAGACUAAUCGAACCAUC-UAGUAGCUGGUUCCCUCCGAAG-UUUCCUCA
GGAUAGCUGGCG-CUCU------------------------------------------------------C
GCAACGCGUUCGCUCGACAACCCG-----CAGUUUUAUCCGGUAAAGCGAAUGAUU-AGA-GGUCUUGG-GG
C-C-GAAAC-GAUC-U-CAACCUAUUCUCAAACUUUAAAUGGGUAAGAA-----------------------
```

```
------------------GCCCG-GC-----UCG-CU--GGCG--U--GGA---------------------
GC---CGGG-C---GUGG--AU-G--C-----G--AGUGCCU-AGUGGGC-C-ACUU-U-UGG-UAAGCAGA
AC-U-GGCGCUGC-GGGAUG---AACCGAACGCCG-G-GUUAAGGCGCCCGAUG--CCGACG-CU-CA---U
-C-AGACCCC-AGAAAAGGUGUUGGUUGAUAUAGACAGCAGGACGGUGGCCAUGGAAGUCGGAA-UCCGCUA
AGGAGUGUGUAACAACUCACCUGCC-----GAAUCAACUAGCCCUGAAAAU-GGAU-GGCGCUGG-AGCGUC
G----------GGCCCAU-ACCCGGCCGUCGCCG--GCAGUCGGAACGGGACGGG-----AGCGGCCGCGGG
CGCGCGACCCC------------------------------GGGGGCCG----------GGCGG---CGU--
----CGGCUUCGGCCGG-CCGCCGCC--CGU---CCACCCCCGGGGC--UCCCCCCGCGGCGUCGG--GCCC
CGC---------------------GGAGCCUACGCCG-CGAC-GAGUAGGAGGGCGCUGCGG----------
-----------U-GAGCCUU-GAAGCCUA-G-GGCGCGGGCC--C-GGGUGGAGCC-G-CCG-CAGGUGCAG
AUCU-UGGUGG-UAGUA-GCAAAU---AUUC-AAACGA--G---------------AACUUU-GAA-GGCCG
AAGUGGAG-AAGGGUUCC-AUGUGAACA-GCAGUUGAACAUGGGUCAGUCGG--UC-CUGAGAGAUGGGCG-
A-G-UGCC-----GUUCCGA--------AGGGACGGGCG----AUGG---CCUC-----------------C
GU---------------UGCCCUC------AGCC----GA-UC-GA-A-AGGGA-GUCGGGUU-CAGA-UCC
CCGAA----------------------------------UCCGGAGUGGCG------------GAGAUGGGC
G----------------CC-GCGA-----------------------------------GGCGUCC------
--AGUGCCGG---UAAC--------GC-GA-----CC--------------------GAUC-C-CGGA-GAA
GCCGGCG-------------GGAG-C-CCGGGGAGA-GU------UCUCU-------------U-UUCUUU-
GUGAAG-GGC-AG-----G-G-------------------CG-----CC----------------UUGG-AA
U-G-GG--------------------UUCGC-CCCGAGAGAG---------GGGCCC----GUG-CC-UUG-
GAA--AG-----------------------------------------------------------------
----CGUCGCA-GUUCCGGCGGCGU-C-CGGUGAG-CUCUCGCUGGC-CCUUGAAA-A-UCC-GGGGGAGAG
-G-------------------------------------------GUGU-----AAAUCUCG-CG-CCGGGC
CGUACCCAUAUCCGCA------------------------------GCAGGUC--UCCAAGGU-GAAC-GCC
UCUGGCAU-GU-UGGA-ACAAUGUA-G-GUAAGGGAAGUCGGC-AAGCCGGAUCCGUAACUUCGG-GAUAAG
GAUUG-GCUC-U-AA--GGGCUG-G----GUCGG-UC------GGGCUG--GGG-CGCGAAGGC--GGG-GC
UGGG------CGCGCGCCGCG---GC----------------------------UGGACGAGGC-GCCG--C
CGCCCCCCUCCCACGUCCGG-GGAGAC---CCCCUCCU--UUCCGCCCGGGCCC-----GCC---CUCCCCU
-CUCCCCGCGGGG-------CCCCG--CCGUCCCCGCGUCGU-CGCCGUGGUC--CCCUCCUCCCU-----
------------------------CCCUU------------------------CUUCCCCGUCC------GC
GGGGGG---------GACGGGG----CGGGU-------------------GCGG--GGGGG-CGCGCGC-GC
GCGCGG------CCCAGGGGC-------G--GC-GGGUCCAACCCCGCGCGG---G--CCGGAGCGG--GG-
---GGAACCCGCGGGCCCCGGU----GGG--GG-------GGG-GCCCGG-ACACCCGGGGGGGACCGGCG
-G-CGGCGGCGACUCUGGACGCGAG-CCGG-GCCCUUC----------------------------------
---CCG----UG-GAUCGC-----CC-CAGCUGCG-GCGGGCGUCGCG-GCCGCUCCCGGGGAGC--CCGGC
GGG--UCGCC-----------------------------CGGCGGGGUUU-UCCUCC-------GGCC-UCG
UCCU-------------------------CCCCC-------------------UUCC--CCCU---------
---------------CC-------GCGGGGU--------------CGGGG---G------------------
UUCCCGGGGUU--CGGGGUUCUCCUCC------------------GCGCGGCGGUUC--CCCCGCCGGGUGC
GCCCCCCGGG-CGCGGU--UUCCCGCGGCCGCCC-GCCU----CGGCCG-GCG--CCUAGCAGC--CGACUU
AGA--ACUGGUGCGGACUAGGGGAAUCCGA-----CUGUUUAAUUAAAACAAAGCAUCGCGAAGGCCCG---
-CG-G------------CGGGUGUUG-ACGCGAUGUGA-UUUCUGCCAGUGCUCAG-AAUG-UCAAAGUGA
A--------------GAAA--------------UUCAAUGAAGCGC--GGGUAAAC-GGCGGGAGUAACUAU
GACUCUCUUAAGGUAGCCAAAUGCCUCGUCAUC-UAAUUAGUGACGCGCAUGAAUGGAUGAACGAGA--UUC
CCACUGUCCCUACC----UAC-UAUCCAGCGAAACC---AC---AGCCAAGGAACGGCUUGGCGGAAUCA
```

```
GCGGGGAAAGAAGA-CCCUGUUGAGC-UUGACU--CUAGUCU-GGCACGGUGAAGAGACAUGAGAGGUGUAG
AAU-AAGUGGGAGGCCCCC---------GGCGCCC------------CCCCGUUCCCC----------GCGA
-------------------GGGGUCGGGGCGGGGUCCGCCGGCCUCGCGGG-C--C-GC-CGGUGAAAUA-C
CACUACUCUCAUCGUUUUUUCACUGACCC-GG-------U-GA-G-GCGGG----GGG-----------GC-
------GA--GC-CC--------CGAG-GGGCU-CU-CG-------------CUUCU---------------
-----------GGCG--------------------------------------CCGAACGC---------GU
------------------C--------------------------------------------CG-------
---CGCG------------------------CGC---GGG-----CGG------------------------
----------GCG------CGA------------------------------CCCG----CU----------
------------------------------------------------CCGG-----GGACA-GUGCCAGGU
GGGGAGUUUGACUGGGGCGGUACACCUGUCA-AACGGUAAC-GCAGGUGUCCU-AAGGCGAGCUCAGGGAGG
ACAGAAACCUCCCGUG-GAGCA-GAAGGGCAA-AAGCUCGCUUGAUC-UUGA--------------------
--------------UUUUCA--GUACGAAUACAGACCGUGAAA-GCGGGGCCUCACGAUCCUUC--------
---UG-A-----CCUUUUGGG--------------------UU-UUAAGCAGGAGGUGUCAG-AA--AAGUU
ACCACAGGG-AUAACUGGCUUGUGGCGGCCAAGCGUUCAUAGCGACGUCGCUUUUUGAUCCUUCGAUGUCGG
CUCUUCCUAUCAUUGUGAAGCA-GAAUUCACCAAGCGUUGGAUUGUUCACCCACUAAUAGGGAACGUGAGCU
GGGUUUAGACCGUCGUGAGACAGGUUAGUUUACCCUACU---------------------G-AUG-------
-------------------------------AU--G-UGUUGUUGCCAUGGU-AAU-CCUGCUCAGUACGAG
AGGAA-CCGCAGGUUCAGACAUU-UGGU-GUAUGUGCUUGGCU-GAGG-AGCCAA-U-GGGGCGAAGCUACC
AUCUG-UGGGAUUAUGACUGAACGCC-UC--UAAGUCAGAAUCC-GCCCAA-----GC--GGAACGAUACGG
CAGCGCCGAAGGAGCCU-------------------------------------------------------
------------------------------------------------------------------------
------------------------------------------------------------------------
---------------------CGGUUGGCCCG-----------GAUAGCCGGCUCCCCGUCCGUCCC-----
-----------------------CGUCC---GGCGGGUCCCCGCCUCGUCGCCCCCCCGGGUGCGGGGCGGG
U-CCCCCG--CC-------GG--GCGUCGGGACCGGGUCCG-------------------GUGCGGA---G-
--AGCCAUUCGU--CCC--GG-----GAAACGGG-----GUGCGGCCG-------GAAAGGGGGCCGCCCUC
----------------------------U----C-GCCCGUCACGCU---UAACGCACGUU----------C
G-----------------UGUGGAA---------------------C-UUGGCGCU-----AAA------CC
AUUCGUAGACGACCUGCU-UCU-GGGUCGGGGU-UUCGUACG-UAGCA--------GAGCAGC-UC------
----CCU--CGCUGCGAUCUAUUGAAAGUCAG-CCCU-CGACACAAG--GGUUUG--U--------------
------------------------------------------------------------------------
------------------------------------------------------------------------
------------------------------------------------------------------------
-------------------------------------------------
>LSUe_10090_MOUSE/1-4888
----------------------------------------GACUCUUAGC-GGUGGAUC-A-CUCGGC-UCG
U-G---CGUCGAUGAAGAACGCAG--CUAGCUGCGAGAAUUAAUGUGAAU--U---GCA--GGA-CA--CAU
UGAUCAUCGACACUUCGAACGCA-CU-U--------GC--------GG----CCCC-G-GGU---------U
C-C-----------UCC---C----G----G---GGC----U-A-CGCC-UGU-----CUGAGCGUCG-CUU
--------------------------------------------------------G--CGCGAC-CUCA-G
AUC--------------------AG-A-CG-UGGCGACCCGCUGAAUUUAANCAUAUUAGUCAGCG--GAGG
AAAAGAAACUAA--CCAGGAUUCCCUCA--GU-AA-CGGCGAGUGAACAGGGA-A-GA-GCCCAGCGCCGAA
UCCC-CGCCGCGCG-UC-GCG------------G---CGU--GGGAAAUGUGGCG--UA-------------
--CG-GAAG-ACC-C--ACUC-------------------------CCCGG--CGC-----------CG---
```

```
CUCGUG--GGGGG--CCC-------AAGUCC-UUCU------------------------------------
-------GAU-CG-AGGCCC---AGCCCGUGGACGGUGUGAGGCCGGUA---------GCG---G----CCC
CCG---GC--GCGCC-GGGC----------------UCGG-GUCUU--CCCGGAGUCGGGUUGCUUG---GG
AAUGCAGCCAAA-G-CGGGUGGUAAA-CUCCAUCUAAGGCUAAAUACCGG--C-ACG-AGACCGAUAGU-C
AAC-A-AGUACCGUAAGGGAAAG-UUGAAA-AGAAC-UUUGA-AGAGAG-A-GUUCAAGAGGGCGUGAAACC
G-UUAAGAGG--U-AAA-CG-GGUG---GGGU-CCGCGCAGU-------------------CC---------
-------------------------------------------GCCCGGAG--GAUUCAACCC--GGCG---
------GCGCGCGUCCGGCCGUGC-CGGUGGU-CCCGGCG-GAUCUUCCCGCUCCCCGUUCCUCCCGACCC
CUCCACCCGCGCGUCG---------UUCCCCUCUUCC-UCCCCGCG-----UCCGG----CGCUCCGG----
-----------------------C-----------------GGCG-GGCGCGG----GGGUGGUGUGGUGG
UGGCGCGCGGGCGGGGCCGGGGGUGGGGUCGGCGGGGGACCG-CCCCCCG--GCCG-GCGACCGGCCGCCGC
C-----GGGCGCA-CUUCC---ACCG-UG--GC----------GGU--------GCGCCGCGACCGGCUCC-
---GG-----------GA-C-------------------G------GCC-----------------GGGAAG
GCC-----------------CGGU-------GGGGAAGGUGGCUCGGGG-------------------GGGG
CGGCGCGUCU---------------------------------------------------CA--G-G----
------------------------------------------------------------GCGCGCCGAACC
--ACCUCACCCCGAGUGUUACAG-CCCUCCGGCCGCAGC------UUUCG-CCGA--AUCC--------CGG
GGCCGAGGAAGCCAGAUACCCGUC----------------GCCGCGCUCUCCCUCU-------------CCC
C---------------------CCGUCC---------------------------------------GCCUC
CC-----------------------------------------GGG--------------------------
-----------CGGGCG----------------UGG------------------GGGU--------------
-------------------------------------------------GGGGGC----CGGGCCG------
-CCCCUC------------CCACGGCGC-------GACCGCUCUCCACCCC------CCUC------CGUCG
-CCUCUCU--CGGGGC-CCGG--------------UGGGGGGC---G-GGGCG----GACUGUC--------
-----CCCAGUG--CGC-CCC---GG-----GCGUCGUC--------GCGCCGUC--GGGUCCGGGG---G
G--------ACCGUCGGUCACGCG--------------------------------------------UCUC
CC-----------------------GACGAAGCCGAGCGCA-------------------------------
----------------CGGGGUCGGCGG------------CGAUGUCGGC--U---ACCACCCGACCCGUCU
UGAAACACGGACCAAGGAGUCUA-A-CGCGUGCGCGA-GUCAGGGG--C-UCG-------UCCG------AA
A-GC--CGCCGUGGCGCAAUGAAGG-UGA-----------------AGGGCCCGCCC---GGGGGCCCGAG
GUGG-GAUCCG----AGGCCUC-----UC---------------------------CAGUCCGC--------
CGA---------------------------------------------------------------------
-------------------GGG--------------------------------------CGCACCACC-GG
CCCGUC----UCGCCCG-C--------------------CGCG----------------CCGGGGAGG-UG-
GAG-CA----------------------CGAGCGUACGCGUUAG-GACCCGAAAGA-UGGUGAACUAUGCCU
GGGCAGGGCGAAGCCAGA-GGAAAC-UCUGGUGGAGGUCCGU--AGCGGUCCUGACGUGCAAAUCGG-UCGU
CCGACCUGGGUAUAGGGGCGAAAGACUAAUCGAACCAUC-UAGUAGCUGGUUCCCUCCGAAG-UUUCCUCA
GGAUAGCUGGCG-CUCU------------------------------------------------------C
GCUCUUUUUUUUUNCCGACGUACG-----CAGUUUUAUCCGGUAAAGCGAAUGAUU-AGA-GGUCUUGG-GG
C-C-GAAAC-GAUC-U-CAACCUAUUCUCAAACUUUAAAUGGGUAAGAA-----------------------
------------------GCCCG-GC-----UCG-CU--GGCG--U--GGA---------------------
GC---CGGG-C---GUGGA-AU-G--C-----G--AGUGCCU-AGUGGGC-C-ACUU-U-UGG-UAAGCAGA
AC-U-GGCGCUGC-GGGAUG---AACCGAACGCCG-G-GUUAAGGCGCCCGAUG--CCGACG-CU-CA---U
-C-AGACCCC-AGAAAAGGUGUUGGUUGAUAUAGACAGCAGGACGGUGGCCAUGGAAGUCGGAA-UCCGCUA
AGGAGUGUGUAACAACUCACCUGCC-----GAAUCAACUAGCCCUGAAAAU-GGAU-GGCGCUGG-AGCGUC
```

```
G----------GGCCCAU-ACCCGGCCGUCGCCG--GCAGUCGGAACGGAACGGGACGGGAGCGGCCGCGGG
UGCUCGUCUCU-----------------------------UGGGGUCUG----------GGGGGCGUGGC--
----GGGGGCCCGUCCC-CCGCCUCC--CCU---CCGCGCGCCGGGUUUCGCCCCCGCGGCGUCGG--GCCC
CGC---------------------GGACGCUACGCCG-CGAC-G-GUAGGAGGGCCGCUGCGG---------
-----------U-GAGCCUU-GAAGCCUA-G-GGCGCGGGCC--C-GGGUGGAGCC-G-CCG-CAGGUGCAG
AUCU-UGGUGG-UAGUA-GCAAAU---AUUC-AAACGA--G---------------AACUUU-GAA-GGCCG
AAGUGGAG-AAGGGUUCC-AUGUGAACA-GCAGUUGAACAUGGGUCAGUCGG--UC-CUGAGAGAUGGGCG-
A-G-UGCC-----GUUCCGA--------AGGGACGGGCG----AUGG---CCUC-----------------C
GU---------------UGCCCUC------GGCC----GA-UC-GA-A-AGGGA-GUCGGGUU-CAGA-UCC
CCGAA----------------------------------UCCGGAGUGGCG------------GAGAUGGGC
G----------------CC-GCGA-----------------------------------GGCGUCC------
--AGUG-CGG---UAAC--------GC-GA-----CC--------------------GAUC-C-CGGA-GAA
GCCGGCG-------------GGAGCC-CCGGGGAGA-GU------UCUCU-------------U-UUCUUU-
GUGAAG-GGC-AG-----G-G-------------------CG-----CC----------------CUGG-AA
U-G-GG--------------------UUCGC-CCCGAGAGAG---------GGGCCC----GUG-CC-UUG-
GAA--AG-----------------------------------------------------------------
----CGUCGCG-GUUCCGGCGGCGU-C-CGGUGAG-CUCUCGCUGGC-CCUUGAAA-A-UCC-GGGGGAGAG
-G-------------------------------------------GUGU-----AAAUCUCG-CG-CCGGGC
CGUACCCAUAUCCGCA------------------------------GCAGGUC--UCCAAGGU-GAACAGCC
UCUGGCAU-GU-UGGA-ACAAUGUA-G-GUAAGGGAAGUCGGC-AAGCCGGAUCCGUAACUUCGG-GAUAAG
GAUUG-GCUC-U-AA--GGGCUG-G----GUCGG-UC------GGGCUG--GGG-CGCGAA-GC--GGG-GC
UGGG------CGCGCGCCGCG---GC----------------------------UGGACGAGGC-GCCG--C
CGCC-CUCUCCCACGUCCGG-GGAGACCC-CCCGUCCU--UUCCGCCCGGGCCC-----GCCC--UCCCCUC
-UUCCCCGCGGGG-------CCCCG--UCGUCCCCCGCGUC----GUCGC--------CACCUCUCU-----
------------------------UCCCC--------------------------CCCUCCUUCUUCCCGUC
GGGG--------------GGC------GGGU--------------------CGG--GGGUC-GGCGCGC--G
GCGCGGG-----CUCCGGGGC-------G--GC-GGGUCCAACCCCGCGGGG---GUUCCGGAGCGG--GA-
---GGAACCAGCGGGCCCCCGGU----GGGGCGG-------GGG-GCCCGG-ACACUCGGGGGGG-CCGGCG
-G-CGGCGGCGACUCUGGACGCGAG-CCGG-GCCCUUC----------------------------------
---CCG----UG-GAUCGC-----CU-CAGCUGCG-GCGGGCGUCGCG-GCCGCUCCCGGGGAGC--CCGGC
GGGUGCCGGC--G--------------------------CGGGUCCCCUC-CCCGCG------GGGCC-UCG
CUCCA------------------------CCCCC------CCA----------UCGC--CUCU---------
---------------CC------CGAGGUGCGU----------GGCGGGG---GCGG-GCGGGCGUC-----
----CCGCGCGUGUGGGGGGAACCUCC------------------GCGUCGGUGUUC--CCCCGCCGGGUCC
GCCCCCCGGGCCGCGGUUUUCCGCGCGGCGCCUC-GCCU----CGGCCG-GCG--CCUAGCAGC--CGACUU
AGA--ACUGGUGCGGACCAGGGGAAUCCGA-----CUGUUUAAUUAAAACAAAGCAUCGCGAAGGCCCG---
-CG-G------------CGGGUGUUG-ACGCGAUGUGA-UUUCUGCCCAGUGCUCUG-AAUG-UCAAAGUGA
A--------------GAAA--------------UUCAAUGAAGCGC--GGGUAAAC-GGCGGGAGUAACUAU
GACUCUCUUAAGGUAGCCAAAUGCCUCGUCAUC-UAAUUAGUGACGCGCAUGAAUGGAUGAACGAGA--UUC
CCACUGUCCCUACC----UAC-UAUCCAGCGAAACC---AC---AGCCAAGGGAACGGGCUUGGCGGAAUCA
GCGGGGAAAGAAGA-CCCUGUUGAGC-UUGACU--CUAGUCU-GGCACGGUGAAGAGACAUGAGAGGUGUAG
AAU-AAGUGGGAGGCCCCC---------GGCGCCCG-----------GCCCCGUCCUC----------GCGU
-----------------------CGGGGUCGGGGCACGCCGGCCUCGCGGG-C--C-GC-CGGUGAAAUA-C
CACUACUCUCAUCGUUUUUUCACUGACCC-GG-------U-GA-G-GCGGG----GGG-----------GC-
------GA--GC-CC--------CGAG-GGGCU-CU-CG-------------CUUCU---------------
```

-----------GGCG--------------------------------------CCAA--GC---------GU
------------------C--------------------------------------------CGU------
---CCCG------------CGC--------GUGC---GGG-----CGG------------------------
----------GCG------CGA------------------------------CCCG----CU----------
------------------------------------------------CCGG-----GGACA-GUGCCAGGU
GGGGAGUUUGACUGGGGCGGUACACCUGUCA-AACGGUAAC-GCAGGUGUCCU-AAGGCGAGCUCAGGGAGG
ACAGAAACCUCCCGUG-GAGCA-GAAGGGCAA-AAGCUCGCUUGAUC-UUGA--------------------
--------------UUUUCA--GUACGAAUACAGACCGUGAAA-GCGGGGCCUCACGAUCCUUC--------
---UG-A-----CCUUUUGGG--------------------UU-UUAAGCAGGAGGUGUCAG-AA--AAGUU
ACCACAGGG-AUAACUGGCUUGUGGCGGCCAAGCGUUCAUAGCGACGUCGCUUUUUGAUCCUUCGAUGUCGG
CUCUUCCUAUCAUUGUGAAGCA-GAAUUCACCAAGCGUUGGAUUGUUCACCCACUAAUAGGGAACGUGAGCU
GGGUUUAGACCGUCGUGAGACAGGUUAGUUUUACCCUACU--------------------G-AUG-------
-------------------------------AU--G-UGUUGUUGCCAUGGU-AAU-CCUGCUCAGUACGAG
AGGAA-CCGCAGGUUCAGACAUU-UGGU-GUAUGUGCUUGGCU-GAGG-AGCCAA-U-GGGGCGAAGCUACC
AUCUG-UGGGAUUAUGACUGAACGCC-UC--UAAGUCAGAAUCCCGCCCAG-----GC--GGAACGAUACGG
CAGCGCCGAAGGAGCCU-------------------------------------------------------
------------------------------------------------------------------------
------------------------------------------------------------------------
---------------------CGGUUGGCCCG-----------GAUAGCCGGGUCCCCGUCCGUCCC-----
-----------------------CGCUC---GGCGGGGUCC--CCGCGUCGUCCCCCGCGGCGGCGCGGGGUC
UCCCCCCG--CC-------GG--GCGUCGGGACCGGGGUCCG------------------GUGCGGA---G-
--AGCCGUUCGU--CUU--GG-----GAAACGGG-----GUGCGGCCG-------GAAAGGGGGCCGCCCUC
----------------------------U----C-GCCCGUCACGUU---GAACGCACGUU----------C
G-----------------UGUGGAA---------------------C-CUGGCGCU-----AAA------CC
AUUCGUAGACGACCUGCU-UCU-GGGUCGGGGU-UUCGUACG-UAGCA--------GAGCAGC-UC------
----CCU--CGCUGCGAUCUAUUGAAAGUCAG-CCCU-CGACACAAG--GGUUUGUCU--------------
------------------------------------------------------------------------
------------------------------------------------------------------------
------------------------------------------------------------------------
-------------------------------------------------
>LSUe_13616_MONDO/1-5115
----------------------------------------AACUCUUAGCGGGUGGAUCUA-CUCGGC-UCG
U-G---CGUCGAUGAAGAACGCAG--CUAGCUGCGAGAAUUAAUGUGAAU--U---GCA--GGA-CAC-CAU
UGAUCAUCGACACUUCGAACGCA-CU-U--------GC--------GG----CCCC-G-GGU---------U
C-C-----------UCC---C----G----G---GGC----U-A-CGCC-UGU-----CUGAGCGUCG-CUU
------------------------------------------------------------GCGAC-CUCA-G
AUC--------------------AG-A-CG-UGGCGACCCGCUGAAUUUAAGCAUAUUAGUCAGCGAGAGAG
AAAAGAAACUAA--CCAGGAUUCCCUCA--GU-AA-CGGCGAGUGAACAGGGA-A-GA-GCCCAGCGCCGAA
UCCC-CGCCCCGCG-GU-GGG------------G---CGC--GGGCCAUGUGGCG--UA-------------
--CG-GAAG-ACC-C--ACUC-------------------------CCCGG--UGC-----------CG---
CUCUCGGUGGGGG--CCC-------AAGUCC-UUCU------------------------------------
-------GAU-CG-AGGCAC---AGCCCGUGGACGGUGUGAGGCCGGUA---------GCG---G----CCC
CCG---GC--GCGCCGGGGA----------------CCGG-GUCUU--CUCGGAGUCGGGUUGCUUG---GG
AAUGCAGCCCAAA-G-CGGGUGGUAAA-CUCCAUCUAAGGCUAAAUACCGG--C-ACG-AGACCGAUAGUCC
AAC-A-AGUACCGUAAGGGAAAG-UUGAAA-AGAACUUUUGA-AGAGAG-A-GUUCAAGAGGGCGUGAAACC

```
G-UUGAGAGG--U-AAA-CGAGGUG---GGGU-CCGCGCAGU--------------------CC--------
-------------------------------------------GCCCGGAG--GAUUCAACCC--GGCG---
------GUCCCUGGCCGGCCGUGC-CGGUGGUUUCCGGCGAGAUCUUUCCCGCC--CCU------CGUUCUU
C----CCGACCC---------------CUCCA---CCACGUCGC-------CCCG----------CGA----
------------------------------------------GGG-GCGGCGGG---CGA------------
GGGC---CGGG-GAG--U-GAGGUG---CCCGCGGGGGACCGCCCCCCCG--GACG-GAGCCCGGCCGCCGC
C-----GGGCGCA-UUUCC---ACCG-AG--GC----------GGU--------GCGCCGCGACCGUCUCC-
---GG-----------GA-C-------------------G------GCU-----------------GGGAAG
GCC-----------------GGGCUUUUCCCUGGGCAGGUGGCUCGGGGGUGGCGGGGG----CCAGAGGGG
GCCCGGGUCGCCGUGUGGGGUGGCGGGG------------------------------UCGGU--C-GGU--
----CCGCCGGCCGGCCUCGUCUUCCCCUCGGCGCCCGCCUCUC-------GUGGGUCCUCCGCCGUCG---
----CAGCCCCCGAGUGUUAUAG-CCCCCCGGCAGUAGC------UCUCG-CCAGC-AUCC--------CGG
GGUCGAGGAAA---GAUGACCGCC----------------GCCGCGGCUUUCCCCCG--------CCGGAGC
CC------------------CUCCGCCGCCCGUC--------------------------------GCCGCC
CCU-----------------------------------CGCGGGUGG-------------------------
--------GGGCGGGUG---CG----------CCGGU---------------GCGGGC--------------
-------------------------------------------------CCGGGUU-CCCGGGGGACGGGCC
GCCCCUC------------CCCCGGCGC-------GACUGCUCU----CCCC--------------------
-------------------------------------GAAACC---G-GGGCG----GACUGUC--------
-----CUCAGUG--CGC-CCC---GA-----CCG-CGUC--------GCGCCGCC--GGG---CUCGG---G
G--------ACCGCU--GCUCUCG--------------------------------------------UCGC
-------------------------------ACGAGGCGCC-------------------------------
----------------CGGGGUCAGCGG------------CGAUGUCGGC--C---CCCACCCGACCCGUCU
UGAAACACGGACCAAGGAGUCUA-A-CACGCGCGCGA-GUCAGGGG--C-UCCG------CACG-----AAA
G-CC--ACCCGUGGCGCAAUGAAGGAUGA-----------------AGGCCCGGCGCC---CGCCGGCCGAG
GUGG-GAUCCCG----AGGCCUC-------------------GGUCCU-UCU-UCCCGUCCUGC--------
CGA---------------------------------------------------------------------
-------------------GGG--------------------------------------CGCACCACC-GG
CCCGUC----UCGCCCG-C--------------------CCCG----------------CCGGGGAGG-UG-
GAG-CA----------------------UGAGCGUGUGUGUUAG-GACCCGAAAGA-UGGUGAACUAUGCCU
GGGCAGGGCGAAGCCAGA-GGAAAC-UCUGGUGGAGGUCCGU--AGCGGUCCUGACGUGCAAAUCGG-UCGU
CCGACCUGGGUAUAGGGGCGAAAGACUAAUCGAACCAUC-UAGUAGCUGGUUCCCUCCGAAG-UUUCCCUCA
GGAUAGCUGGCG-CUCUCGCAGUGGUCGUCCGGUCGGAGCGGGGGAGCGCCCCCCUCCCUCGGGGGGCGGGG
GUUCCCCUGGCACCCGCCGCCACG-----CAGUUUUAUCCGGUCAAGCGAAUGAUU-AGA-GGUCUUGG-GG
C-C-GAAAC-GAUC-U-CAACCUAUUCUCAAACUUUAAAUGGGUAAGAA-----------------------
------------------GCCCG-GC-----UCG-CU--GGCU--U--GGA---------------------
GC---CGGG-C---GUGGA-AU-G--C-----G--AGUGCCU-AGUGGGC-C-ACUU-U-UGG-UAAGCAGA
AC-U-GGCGCUGC-GGGAUG---AACCGAACGCCG-G-GUUAAGGCGCCCGAUG--CCGACG-CU-CA---U
-C-AGACCCC-AGAAAAGGUGUUGGUUGAUAUAGACAGCAGGACGGUGGCCAUGGAAGUCGGAA-UCCGCUA
AGGAGUGUGUAACAACUCACCUGCC-----GAAUCAACUAGCCCUGAAAAU-GGAU-GGCGCUGG-AGCGUC
G----------GGCCCAU-ACCCGGCCGUCGCUG--GCAAUGGGCAAAAGCAG-------------------
-------------------------------------------------------------------GAA--
----CAGAAACGUGUUGUUUCCCGAGAACGGAUUCCGCGCGCGGGAGGAAG-----GCGCGAACUC--ACUC
CUC---------------------GGGGGCUAGGCCG-CGAC-GAGUAGGAGGGCCGCCGCGG---------
-----------U-GGGCCUG-GAAGCCUA-G-GGCGUGGGCC--C-GGGUGGAGCC-G-CCG-CGGGUGCAG
```

```
AUCU-UGGUGG-UAGUA-GCAAAU---AUUC-AAACGA--G---------------AACUUU-GAA-GGCCG
AAGUGGAG-AAGGGUUCC-AUGUGAACA-GCAGUUGAACAUGGGUCAGUCGG--UC-CUGAGAGAUAGGCG-
A-G-CGCC-----GUUCCGA--------AGGGACGGGCG----AUGG---CCUC-----------------C
GU---------------UGCCCUC------GGCC----GA-UC-GA-A-AGGGA-GUCGGGUU-CAGA-UCC
CCGAA----------------------------------UCCGGAGUGGCG------------GAGACGGGC
G----------------CC-GUGA-----------------------------------GGCGUCC------
--AGUG-CGG---UAAC--------GC-GA-----CC--------------------GAUC-C-CGGA-GAA
GCCGGCG-------------GGAGCC-CCGGGGAGA-GU------UCUCU-------------U-UUCUUU-
GUGAAG-GGC-AG-----G-G------------------CCG-----CC----------------CUGG-AA
U-G-GG--------------------UUCGC-CCCGAGAGAG---------GGGCCC----GCG-CC-UUG-
GAA--AG-----------------------------------------------------------------
----CGUCGCG-GUUCCGGCGGCGU-C-CGGUGAG-CUCUCGCUGGC-CCUUGAAA-A-UCC-GGGGGAGAG
-G-------------------------------------------GUGU-----AAAUCUCG-CG-CCGGGC
CGUACCCAUAUCCGCA------------------------------GCAGGUC--UCCAAGGU-GAACAGCC
UCUGGCAU-GU-UGGA-ACAAUGUA-G-GUAAGGGAAGUCGGC-AAGCCGGAUCCGUAACUUCGG-GAUAAG
GAUUG-GCUC-U-AA--GGGCUG-G----GUCGG-UC------GGGCUG--GGG-CGCGAA-GC--GGG-GC
UGGG------CGCGCGCCGCG---GC----------------------------UGGACGAGGC-GCCG--C
CGCU-CCCCCU----CCCGGCG----CUCUCCCGCCGG--ACGCC---UCCCCGC---GGCCCCCCCGCCCG
-CUCCCCGCCGGGUUCCCUUCCCCG--CGGCUCCCCUGGCCUC-CCCCCCUCCCACGCCUCCCCGCCGGCCA
AACCGCGGCGGUGGCGGGCGGGGGGGGACGGGGGGGGCGCGCGGGGGGUCGGGGGCGCGGGUCCC----GCC
CGCGUCUCCCCUCCCUCCCGCCCCCCCUCGCUCCUCUCCCACCCUCCCCACCGCG-GCGGUGGGGGGGCAGG
GGGGGGGAUCGGCGCCAGGGGGUUCCAAGUCGC-GGGGUCGGCCCGGCGGGGGUUG-GUCGGUGCGGUCGG-
---GG-----UCCGGGGUUCGG-----GCCCGGC-------GGGUGCCCGG-GAAC-CGGGGG---UCGGCG
-G-CGGCGGCGACUCUGGACGCGAG-CCGG-GCCCUUC----------------------------------
---CCG----UG-GAUCGC-----CC-CAGCUGCG-GCGGGCGUCGCG-GCCGGGCCCGGGGAGC--CCGGG
GGC--GGGCC-UGCUCGGCCCGCCGGCCU----------CGGGGGGGUCU-GCCCUG------GGGCGCGCG
CGGC-------------------------UCCCCG-----CCCG--CG-----GCCC--CCCU---------
---------------CC-------GGGGGGUGU------C---GGCGGCG---G-GGUGGGUCCGCGCG---
----CCCCG-----GGUGCC--CCUCC---GGCCGCG--GGA--UCUGUGGCGGCCCUCCCCCCCCGGGUGC
-CCCCCCGGGCCUCGGU---CCCCGCGGCGUCCC-GCCU----CGGCCG-GCG--CCUAGCAGC--UGACUU
AGA--ACUGGUGCGGACCAGGGGAAUCCGA-----CUGUUUAAUUAAAACAAAGCAUCGCGAGGGCCCG---
-CG-G------------CGGGUGUUG-ACGCGAUGUGA-UUUCUGCCCAGUGCUCUG-AAUG-UCAAAGUGA
A--------------GAAA--------------UUCAAUGAAGCGC--GGGUAAAC-GGCGGGAGUAACUAU
GACUCUCUUAAGGUAGCCAAAUGCCUCGUCAUCUUAAUUAGUGACGCGCAUGAAUGGAUGAACGAAGAUUCC
CCACUGUCCCUACC----UACUUAUCUAGCGAAACC---AC---AGCCAAGGGAACGGGCUUGGCGGAAUCA
GCGGGGAAAGAAGACCCCUGUUGAGC-UUGACU--CUAGUCU-GGCACGGUGAAGAGACAUGAGAGGUGUAG
AAUAAAGUGGGAGGCCCCCC-------AGGUGCCCCCCGGGGCCCCGCCCCCGGGCUCCCCCGCGAGGAGGA
UCCCCCGGGUCGGCGGGCCCCCCCCCGGGGUCGGCGAGCCGGCCUUCCGGG-C--C-GC-CGGUGAAAUACC
CACUACUCUGAUCGUUUUUUCACUGACCC-GG------U-GA-G-GCGGG---GGGA------------GC-
------GA--GC-CC--------CGAG-GGGCU-CU-CG-------------CUUCU---------------
-----------GGCG--------------------------------------CCAA--GC---------GC
------------------C--------------------------------------------CG-------
---GCCG------------CAC--------GGCG---CGC-----CGG------------------------
----------GCG------CGA------------------------------CCCG----CU----------
------------------------------------------------CCGG-----GGACA-GUGUCAGGU
```

```
GGGGAGUUUGACUGGGGCGGUACACCUGUCACAACCGUAACGGCAGGUGUCCU-AAGGCGAGCUCAGGGAGG
ACAGAAACCUCCCGUG-GAGCA-GAAGGGCAA-AAGCUCGCUUGAUC-UUGA--------------------
--------------UUUUCA--GUAUGAAUACAGACCGUGAAA-GCGGGGCCUCACGAUCCUUC--------
---UG-A-----CCUUUGGGG--------------------UU-UUAAGCAGGAGGUGUCAG-AA--AAGUU
ACCACAGGG-AUAACUGGCUUGUGGCGGCCAAGCGUUCAUAGCGACGUCGCUUUUGAUCCUUCGAUGUCGG
CUCUUCCUAUCAUUGUGAAGCA-GAAUUCACCAAGCGUUGGAUUGUUCACCCACUAAUAGGGAACGUGAGCU
GGGUUUAGACCGUCGUGAGACAGGUUAGUUUACCCUACU---------------------G-AUG-------
-------------------------------AG--G-GGUUGUCGCCAUAGU-AAU-CCUGCUCAGUACGAG
AGGAA-CCGCAGGUUCAGACAUUAUGGU-GUAUGUGCUUGGCU-GAGG-AGCCAA-U-GGGGCGAAGCUACC
AUCUG-UGGGAUUAUGACUGAACGCC-UCCGUAAGUCAGAAUCCCCCCUAA-----GC--GUGACGAUACCG
CAGCGCCG-AGGAGCCU-------------------------------------------------------
------------------------------------------------------------------------
------------------------------------------------------------------------
---------------------CGGUGGGCCACG----------GAUAGCCGGG---CUGCCCGUGCUCCCU-
-----------------------CCCUC------------------------------UGUUUCG-------
---GCGGG--GG-------GG--CCUCGGGGGCCGGGGCCCG------------------GUGCGGA---G-
--GGCCGUUCGC--GCC--GG-----GGGCUGGA-----GAGCGGCCGC------GAAG-GGGGGCCGCCUC
----------------------------U----C-ACCCGGAGCGC-----AGCGCAUGUU----------C
G-----------------UGGGGCA---------------------C-CUGGUGCU-----AAA------UC
AUUCGUAGACGACCUGCU-UCU-GGGUCGGGGU-UUCGUGCG-UAGCA---------GAGCAGC-UC------
----CCU--CGCUGCGAUCUAUUGAAAGUCAG-CCCU-UGACACAAG--CUUUUGU-C--------------
------------------------------------------------------------------------
------------------------------------------------------------------------
------------------------------------------------------------------------
-------------------------------------------------
>LSUe_28377_ANOCA/1-4085
------------------------------------------CUCUUAGC-GGUGGAUC-A-CUCGGC-UCG
U-G---CGUCGAUGAAGAACGCAG--CUAGCUGCGAGAAUUAAUGUGAAU--U---GCA--GGA-CA--CAU
UGAUCAUCGACACUUCGAACGCA-CU-U--------GC--------GG----CCCC-G-GGU---------U
C-C-----------UCC---C----G----G---GGC----U-A-CGCC-UGU-----CUGAGCGUCG-CU-
-----------------------------------------------------------CGCGAC-CUCA-G
AUC--------------------AG-A-CG-UGGCGACCCGCUGAAUUUAAGCAUUAGUCAGCG--GAGG
AAAAGAAACUAA--CCAGGAUUCCCUCA--GU-AA-CGGCGAGUGAACAGGGA-A-GA-GCCCAGCGCCGAA
UCCC-CGUCCCGCG-GC-GGG------------G---CGC--GGGAAAUGUGGCG--UA-------------
--CG-GAAG-ACC-C--CCUC-------------------------CCCGG--CGC-----------CG---
CUCCCUCGGGGGG--CCC-------AAGUCC-UUCU------------------------------------
-------GAU-CG-AGGCAC---AGCCCGCGGACGGUGUGAGGCCGGUA---------GCG---G----CCC
CCG---GC--GCGUC-GGGA----------------CCGG-GUCUU--CUCGGAGUCGGGUUGCUUG---GG
AAUGCAGCCCAAA-G-CGGGUGGUAAA-CUCCAUCUAAGGCUAAAUACCGG--C-ACG-AGACCGAUAGU-C
GAC-A-AGUACCGUAAGGGAAAG-UUGAAA-AGAAC-UUUGA-AGAGAG-A-GUUCAAGAGGGCGUGAAACC
G-UUAAGAGG--U-AAA-CG-GGUG---GGGC-CCGCGCAGU---------------------CC--------
-------------------------------------------GCCCGGAG--GAUUCAACCC--GGCG---
------GGCCGUCGUCGGCCGCGC-CGG-GCG---CG-CG-GAU----CCC---------------------
-----CCCCCCG---------------CCCC--------G---------GACCCA---------CCUC----
------------------------------------------GGG-GAAG-GGGUUCC--------------
```

```
-------CGGG------------------A----GGGGACCG-CCGCCCG--GGCG-GAG-CCGGCCCCCGU
C-----GGGCGCA-UUUCC---GCCG-GG--GC----------GGU--------GCGCCGCGACCGGCUCC-
---GG-----------GU-C-------------------G------GCC-----------------GGGAAG
GCC-----------------CC-GGUUC----GGGCAGGUGGCCCGCCG-----------------------
----------------------------------------------------------CCUCG--C-CC---
------------------------------------------------------------------------
------GGCGGCGGGUGUUACAG-CCCCCGGGCCUCAGC------CAUCG-CCGC--AUCC--------CGG
GGCCGAGGGA----GAAGACCGCC----------------GCCGCGCCCUCCCCCCG--------CCGGCCC
C---------------------CCGUCC---------------------------------------CUCCC
CCU-----------------------------------CGCGGGG---------------------------
-----------CGGGCGGCGCGGG-----GGCCUCC------------------GCGC--------------
-------------------------------------------------GGGGGA----CGGGCCC------
-CCCCGC------------UCCCGGCGC-------GACUGU-C-----------------------------
---------------------------------------AACC---G-GGGCG----GACUGCC--------
-----CUCAGUG--CGC-CCC---GA-----CCG-CGUC--------GCGCCGCC--GGG---CGGGG---A
G--------------------GCG--------------------------------------------CCAC
-------------------------------GCCGGGCGCC-------------------------------
----------------CGGGGUCCGCGG------------CGACGUCGGC--C---CCCACCCGACCCGUCU
UGAAACACGGACCAAGGAGUCUA-A-CACGCGCGCGA-GUCGGAGG--C-UGA-------GCGC------GA
A-AG--CCCCGCGGCGCAAUGAAGG-UGA-----------------GGGCCGGCGCG----CGCCGGCCGAG
GUGG-GAUCCCG----UGUCCGCCG---CC------------------------------GAGC--------
GGG---------------------------------------------------------------------
-------------------GGG--------------------------------------CGCACCACC-GG
CCCGUC----UCGCCCG-C--------------------CCCG----------------UCGGGGAGG-UG-
GAG-CG----------------------UGAGCGCGCGUGCUAG-GACCCGAAAGA-UGGUGAACUAUGCCC
GGGCAGGGCGAAGCCAGA-GGAAAC-UCUGGUGGAGGUCCGU--AGCGGUCCUGACGUGCAAAUCGG-UCGU
CCGACCUGGGUAUAGGGGCGAAAGACUAAUCGAACCGUC-UAGUAGCUGGUUCCCUCCGAAG-UUUCCCUCA
GGAUAGCUGGCG-CUCG-------------------------------------------------------
-------------UCCGUCCGUCG-----CAGUUUUAUCCGGUAAAGCGAAUGAUU-AGA-GGUCUUGG-GG
C-C-GAAAC-GAUC-U-CAACCUAUUCUCAAACUUUAAAUGGGUAAGAA-----------------------
------------------GCCCG-GC-----UCG-CU--GGCC--U--GGA---------------------
GC---CGGG-C---GUGGA-AU-G--CG----A--GCCGCCC-AGUGGGC-C-ACUU-U-UGG-UAAGCAGA
AC-U-GGCGCUGC-GGGAUG---AACCGAACGCCG-G-GUUAAGGCGCCCGAUG--CCGACG-CU-CA---U
-C-AGACCCC-AGAAAAGGUGUUGGUCGAUCUAGACAGCAGGACGGUGGCCAUGGAAGUCGGAA-UCCGCCA
AGGAGUGUGUAACAACUCACCUGCC-----GAAUCGACUAGCCCUGAAAAU-GGAU-GGCGCUGG-AGCGUC
G----------GGCCCAU-ACCCGGCCGUCGCGG--GCGAUGCGG---------------------------
--------------------------------------------------------------------GC--
----CGC-----------------------------------------------------------------
------------------------GGGGGCUAGGCCG-CGAC-GAGUAGGAGGGCCGCCGCGG---------
-----------U-GGGCCUU-GAAGCCUA-G-GGCGCGGGCC--C-GGGUGGAGCC-G-CCG-CGGGUGCAG
AUCU-UGGUGG-UAGUA-GCAAAU---AUUC-AAACGA--G---------------AACUUU-GAA-GGCCG
AAGUGGAG-AAGGGUUCC-AUGUGAACA-GCAGUUGAACAUGGGUCAGUCGG--UC-CUAAGAGAUAGGCG-
A-G-CGCC-----GUUCCGA--------AGGGACGGGCG----AUGG---CCUC-----------------C
GU---------------CGCCCUC------GGCC----GA-UC-GA-A-AGGGA-GUCGGGUU-CAGA-UCC
CCGAA----------------------------------UCCGGAGCGGCG------------GAGACGGGC
```

```
G----------------CC-GCGA-----------------------------------GGCGUCC------
--AGUG-CGG---UAAC--------GC-GA-----CC--------------------GAUC-C-CGGA-GAA
GCCGGCG-------------GGAGCC-CCGGGGAGA-GU------UCUCU-------------U-UUCUUC-
GUGAAG-GGC-AG-----G-G-------------------CG-----CC----------------CUGG-AA
C-G-GG--------------------UUCGC-CCCGAGAGAG---------GGGCCC----GCG-CC-UUG-
GAA--AG-----------------------------------------------------------------
----CGUCGCG-GUUCCGGCGGCGU-C-CGGUGAG-CUCUCGCCGGC-CCUUGAAA-A-UCC-GGGGGAGAG
-G-------------------------------------------GUGU-----AAAUCUCG-CG-CCGGGC
CGUACCCAUAUCCGCA------------------------------GCAGGUC--UCCAAGGU-GAACAGCC
UCUGGCAU-GU-UAGA-ACAACGUA-G-GUAAGGGAAGUCGGC-AAGCCGGAUCCGUAACUUCGG-GAUAAG
GAUUG-GCUC-U-GA--GGGCUG-G----GCCGG-UC------GGGCUG--GGG-CGCGAA-GC--GGG-GC
UGGG------CGCGCGCCGCG---GC----------------------------CGGAGGAGGC-GCC----
----------------------------------------------------------------------CG
-CUCCC--------------CCCCG-----------------------------------------------
-------------------------------------------------------UGCGC----------CC
CGCG--------------UGC------------------------------CGG--GG--------------
-----------------------------------------------------------------GGC----
------------------------------------------------------------------------
-G-CGGCGGCGACUCUGGACGCGCG-CCGG-GCCCUUC----------------------------------
---CCG----UG-GAUCGC-----CC-CAGCUGCG-GCGGGCGCGG--------------------------
-----CCG--------------------------------------------------------GGCC----
---------------------------------C-------------------UCCC--CCCU---------
---------------CG---------------------------GCGGGG---G------------------
------ACG-----GGU------------------------------------------------CGGG---
----CC--------------------GGCGUCCC-GCCU----CGGCCG-GCG--CCUAGCAGC--CGACUU
AGA--ACUGGCGCGGACCAGGGGAAUCCGA-----CUGUUUAAUUAAAACAAAGCAUCGCGAAGGCCCG---
-CG-G-----------CGGGUGUUG-ACGCGAUGUGA-UUUCUGCCCAGUGCUCUG-AAUG-UCAAAGUGA
A--------------GAAA--------------UUCAAUGAAGCGC--GGGUAAAC-GGCGGGAGUAACUAU
GACUCUCUUAAGGUAGCCAAAUGCCUCGUCAUC-UAAUUAGUGACGCGCAUGAAUGGAUGAACGAGA--UUC
CCACUGUCCCUACC----UAC-UAUCUAGCGAAACC---AC---AGCCAAGGGAACGGGCUUGGCGGAAUCA
GCGGGGAAAGAAGA-CCCUGUUGAGC-UUGACU--CUAGUCU-GGCCCUGUGAAGAGACAUGAGAGGCGUAG
AAU-AAGUGGGAGGCCCG--------------------------------------------------CCCG
U----------------------------------------------CGGG-C--C-GC-CGGUG-AAUA-C
CACUACUCUGAUCGUUUUUUCACUUACCC-GG-------U-GA-G-GCGGG----GGG-----------GC-
------GA--GC-CC--------CGAG-GGGCU-CU-CG-------------CUUCU---------------
-----------GGCU--------------------------------------GCAA--GC---------GC
------------------C--------------------------------------------CG-------
---GC--------------------------GCG---GGC-----CGG------------------------
----------GCG------CGA------------------------------CCCG----CU----------
------------------------------------------------CCGG-----GGACA-GCGUCAGGU
GGGGAGUUUGACUGGGGCGGUACACCUGUCA-AAGCGUAAC-GCAGGUGUCCU-AAGGCGAGCUCAGGGAGG
ACAGAAACCUCCCGCG-GAGCA-GAAGGGCAA-AAGCUCGCUUGAUC-UUGA--------------------
--------------UUUUCA--GUACGAAUACAGACCGUGAAA-GCGGGG-CUCACGAUCCUUC--------
---UG-A-----CUUUUUGGG--------------------UU-UUAAGCAGGAGGUGUCAG-AA--AAGUU
ACCACAGGG-AUAACUGGCUUGUGGCGGCCAAGCGUUCAUAGCGACGUCGCUUUUUGAUCCUUCGAUGUCGG
```

```
CUCUUCCUAUCAUUGUGAAGCA-GAAUUCACCAAGCGUUGGAUUGUUCACCCACUAAUAGGGAACGUGAGCU
GGGUUUAGACCGUCGUGAGACAGGUUAGUUUUACCCUACU--------------------G-AUG-------
-------------------------------AG--G-CGUCGUUGCGCUAGU-AAU-CCUGCUCAGUACGAG
AGGAA-CCGCAGGUUCAGACAUU-UGGU-GUAUGUGCUUGGCU-GAGG-AGCCAA-U-GGGGCGAAGCUACC
AUCUG-UGGGAUUAUGACUGAACGCC-UC--UAAGUCAGAAUCCCCCCUAA-----AG--GGAACGAUACCG
CAGCGCCG-AGGAGCCU-------------------------------------------------------
------------------------------------------------------------------------
------------------------------------------------------------------------
---------------------CGGUGGCCCGG------------AUAGCCG---------------------
-------------------------------GGCCGG------------CCCUCC-----------------
---CGGGG--GC-------CG--GC-------------CCCG------------------GUGCGGA---G-
--AGCCGUCCGU--CUC--GG-----GAG-CGGA-----GCGCGGCCG-------GAAGGGGG--CCGCCUC
----------------------------U----C-GCCCGUAGCGC-----AACGCACGUU----------C
G-----------------CGGGGAA---------------------C-CCGGCGCU-----AAA------UC
AUUCGUAGACGACCUGAU-UCU-GGGUCGGGGU-UUCGUACG-UAGCA--------GAGCAGC-UC------
----CCU--CGCUGCGAUCUAUUGAAAGUCAG-CCCU-CGACACAAG--CUUUUGU-C--------------
------------------------------------------------------------------------
------------------------------------------------------------------------
------------------------------------------------------------------------
-------------------------------------------------
>LSUe_9031_CHICK/1-4610
---------------------------------------CAACUCUUAGC-GGUGGAUC-A-CUCGGC-UCG
U-G---CGUCGAUGAAGAACGCAG--CUAGCUGCGAGAAUUAAUGUGAAU--U---GCA--GGA-CA--CAU
UGAUCAUCGACACUUCGAACGCA-CU-U--------GC--------GG----CCCC-G-GGU---------U
C-C-----------UCC---C----G----G---GGC----U-A-CGCC-UGC-----CUGAGCGUCG-CUU
UCCCGU--------------------------------UUCCGU------------U--CGCGAC-CUCA-G
GUC--------------------AG-A-CG-UGGCGACCCCUUUAAUUUAAGCAUAUUAGUCAGCG--GAGG
AAAAGAAACUAC--CUAGGAUUCCCUCA--GU-AA-CGGCGAGUGAAGAGGGA-A-GA-GCCCAGCGCCGAA
UCCC-CGCCCCGCG-GU-GGG------------G---CGC--GGGAGGUGUGGCG--UA-------------
--CG-GAAG-CCC-C--CAUC-------------------------CCCGG--CGC-----------CG---
CUCUCG--GGGGG--CCC-------AAGUCC-UUCU------------------------------------
-------GAU-CG-AGGCCC---AGCCCGCGGACGGUGUGAGGCCGGUA---------GCG---G-C--CCC
CCG---GC--GCGCC-GGGC----------------CCGG-GGCUU--CUCGGAGUCGGGUUGCUUG---GG
AAUGCAGCCCAAA-G-CGGGUGGUAAA-CUCCAUCUAAGGCUAAAUACCGG--C-ACG-AGACCGAUAGC-C
AAC-A-AGUACCGUAAGGGAAAG-UUGAAA-AGAAC-UUUGA-AGAGAG-A-GUUCAAGAGGGCGUGAAACC
G-UUAAGAGG--U-AAA-CG-GGUG---GGGU-CCGCGCAGU--------------------CG--------
-------------------------------------------GCCCGGAG--GAUUCAACCC--GGCG---
------GGCCAAGGUCGGCCGG-CGCGG-GCG--CCGUCG-GAUC---CCCGCC------------------
-----UCCGCCU---------------CCCC----UC-CGUCCCUC-----CCC--------UUCGCC----
-----------------------------------------GGGGCGGGGCGGGC-CC-A-----------G
GGG-GGGCGGGCGGG-------------CC----GGGGACCG-CCGCCCG--GCCG-GCGUCCGGCCCCCGU
C-----GGGCGCA-UUUCC---UCCG-CG--GC----------GGU--------GCGCCGCGACCGGCUCC-
---GG-----------GA-C-------------------G------GCU-----------------GGGAAG
GGC-----------------UGCCGG-----CGGGCAGGUGGCCCG----------------------GCGC
CGC---------------------------------------------------------GCG--A------
```

```
-------------------------------------------------------------GCGGCCG----
----------CCGGGUGUUAUAGCCGCCGGGCCCGGAUC------G-UCG-CCGA--AUCC--------CGG
GGCCGAGGGA----GAGGACCGCCGCCGCGCCCUCCCCCGGAGGGGGCGGCCCCCCGGAGGGCCCCCCGCGG
CCGGACCGGCGUCGGGCCGGCCGCGCCGCGCGCGUCCGCGCCGCCGCCGUACGCCGCCGCUCGCUCUCUC
UCCGUUCCCCGCCCCGGGUCCGUCCCGGGGCGCGGGGGCGGGGGGGGUCGGGUGUCCGGCGCGCGGCUGGGC
GCGGCGCCGCGCGUGUGGCGCGCAUCCAGCCCGGCGCGGGCGAGGCCGCGGGGGGCGCCGGGGGGGAACC
UUCCCCCUUCUGUUCGGGCCGCCUCCGUUCCCGCGGGGGCGGCCCGUUCGGGGGA----CGGGCCC------
-GCCGGC------------CCCCGGCGC-------CGCUGUCC-----------------------------
---------------------------------------GACC---G-GGGCG----GACUGCG--------
-----CUCAGUG--CGC-CCC---GA-----CCG-CGCG--------GCGCCGCC--GGG---CCGGGCUCG
G--------GCC---------ACG--------------------------------------------CC--
---------------------------------AGGGCGCC-------------------------------
----------------CGGGGUCCGCGG------------CGACGUCGGC--U---ACCACCCGACCCGUCU
UGAAACACGGACCAAGGAGUCUA-G-CACGCGCGCGA-GUCGGCGG--C-UCG-------CGCG------AA
A-GC--CCGC--GGCGCAAUGAAGG-UGA-----------------GGGCCGGCGCG----CGCCGGCUGAG
GUGG-GAUCCCG----GGGCGGCAG---GC--------------------------CGGAAGGC--------
CCC---------------------------------------------------------------------
-------------------GGG--------------------------------------CGCACCACC-GG
CCCGUC----UCGCCCG-C--------------------CUCG----------------CCGGGGAGG-UG-
GAG-CA----------------------UGAGCGCGCGUGCUAG-GACCCGAAAGA-UGGUGAACUAUGCCU
GGGCAGGGCGAAGCCAGA-GGAAAC-UCUGGUGGAGGUCCGU--AGCGGUCCUGACGUGCAAAUCGG-UCGU
CCGACCCGGGUAUAGGGGCGAAAGACUAAUCGAACCAUC-UAGUAGCUGGUUCCCUCCGAAG-UUUCCCUCA
GGAUAGCUGGCG-CUCG-------------------------------------------------------
-------------GGGCGGCGGUG-----CAGUUUACCCGGUAAAGCGAAUGAUU-AGA-GGUCUUGG-GG
C-C-GAAAC-GAUC-U-CAACCUAUUCUCAAACUUUCAAUGGGUAAGAC-----------------------
------------------GCCCG-GC-----UCG-CU--GGCG--U--GGA---------------------
GC---CGGGCC---GUGGA-AU-G--C-----G--AGCGCUC-AGUGGGC-C-ACUU-U-UGG-UAAGCAGA
AC-U-GGCGCUGC-GGGAUG---AACCGAACGCCG-G-GUUAAGGCGCCCGAUG--CCGACG-CU-CA---U
-C-AGAGCCC-AGAAAAGGUGUUGGUUGAUCUAGACAGCAGGACGGUGGCCAUGGAAGUCGGAA-CCCGCUA
AGGAGUGUGUAACAACUCACCUGCC-----GAAUCAACUAGCCCUGAAAAU-GGAU-GGCGCUGG-AGCGUC
A----------GGCCCAU-ACCCGGCCGUCGCCG--GCGGUGCG----------------------------
-------------------------------------------------------------------AGC--
----CGC-----------------------------------------------------------------
------------------------GGGGGCUACGCCG-CGAC-GAGUAGGAGGGCCGCUGCGG---------
-----------U-GCGCCUG-GAAGCCUG-G-GGCGCGGGCC--C-GGGUGGAGCC-G-CCG-CAGGUGCAG
AUCU-UGGUGG-UAGUA-GCAACU---AUUC-AAACGA--G---------------AGCUUU-GAA-GGCCG
AAGUGGAG-CAGGGUUCC-AUGUGAACA-GCAGUUGAACAUGGGUCAGUUGG--UC-CUAAGCGAUAGGCG-
A-G-CGCC-----GUUCCGA--------AGGGACGGGCG----AUAG---CCUC-----------------C
GU---------------UGCCCUC------AGCC----GA-UC-GA-A-AGGGA-GUCGGGUU-CAGA-UCC
CCGAA----------------------------------UCCGGGGCGGCG------------GAGACGGGC
G----------------CC-GCGA-----------------------------------GGCGCCC------
--AGUG-CGG---UAAC--------GC-AA-----GC--------------------GAUC-C-CGGA-GAA
GCCGGCG-------------GGAGCC-CCGGGGAGA-GU------UCUCU-------------U-UUCUUU-
GUGAAG-GGC-CG-----G-G-------------------CG-----CC----------------CUGG-AA
C-G-GG--------------------UUCGC-CCCGAGAGAG---------GGGCCC----GCG-CC-UUG-
```

```
GAA--AG-----------------------------------------------------------------
----CGUCGCG-GUUCCGGCGGCGU-C-CGGUGAG-CUCUCGCUGGC-CCGUGAAA-A-UCC-GGGGGAGAG
-G-------------------------------------------GUGU-----AAAUCUCG-CG-CCGGGC
CGUACCCAUAUCCGCA------------------------------GCAGGUC--UCCAAGGU-GAACAGCC
UCUGGCAU-GU-UGGA-CCAAUGUA-G-GUAAGGGAAGUCGGC-AAGCCGGAUCCGUAACUUCGG-GAUAAG
GAUUG-GCUC-U-AA--GGGCUG-G----GUCGG-UC------GGGCUG--GGG-CGCGAA-GC--GGG-GC
UGGG------CGCGCGCCGCG---GC----------------------------UGGACGAGGC-GCCG--C
CC---------------------------------------------------------------GCCCCCG
-CCCCCCCUUU---------CCCCG-----------------------------------------------
-------------------------------------------------------CUCCCGCUCG-----CC
GGGG--------------CGC------------------------------CGG--GGG-------------
----------------GGGGG-------G-----------------------------UCA--GCGG-----
--------------------------------GC-------GGC-------------------------GCG
-G-CGGCGGCGACUCUGGACGCGCG-CCGG-GCCCUUC----------------------------------
---CCG----UG-GAUCGC-----CC-CAGCUGCG-GCGGGCGCCGCUCGCCCCCCUCCUUGCCCCUCCGCC
CCC--CCGCU--CC-------------------------CGGCGCCCC-U-CCCGUC-------GGCCGUCG
UCCGGG------------------CCGC-CCCCCG-----UCC---CG----AGCGC--CCUC---------
---------------CCC-----GCGAGGGCGCGAGGGGCGGCGGCGGCG---GCCGCGGGCGCGGCGG---
----CGGCG-G---GGGGGGGCCCGCC---GGCGGCGCCGG----GCGGGGCGGUCC----CGGGCGGG---
----GGGGGUCUCCGG-------GCCGGCGCCCC-GCCU----CGGCCG-GCG--CCUAGCAGC--CGGCUU
AGA--ACUGGUGCGGACCAGGGGAAUCCGA-----CUGUUUAAUUAAAACAAAGCAUCGCGAAGGCCCG---
-CG-G------------CGGGUGUUG-ACGCGAUGUGA-UUUCUGCCCAGUGCUCUG-AAUG-UCAAAGUGA
A--------------GAAA--------------UUCAAUGAAGCGC--GGGUAAAC-GGCGGGAGUAACUAU
GACUCUCUUAAGGUAGCCAAAUGCCUCGUCAUC-UAAUUAGUGACGCGCAUGAAUGGAUGAACGAGA--UUC
CCACUGUCCCUACC----UAC-UCUCCAGCGAAACC---AC---AGCCAAGGGAACGGGCUUGGCGGAAUCA
GCGGGGAAAGAAGA-CCCUGUUGAGC-UUGACU--CUAGUCU-GGCGCUGUGAAGAGACAUGAGAGGUGUAG
AAU-AAGUGGGAGGCC---C-------CGCGGUCGCG-------------------------------CGAC
CC-----------------------------------GCGCCGCGGCCCGG-C--C-GC-CGGUGAAAUA-C
CACUACUCUGAUCGUUUUUUCACUUACCC-GG-------U-GA-G-GCGGG----GGG-----------GC-
------GA--GC-CC--------CGAG-GGGCU-CU-CG-------------CUUCU---------------
-----------GGCG--------------------------------------CCAA--GC---------GC
------------------C--------------------------------------------CG-------
---GC--------------------------GCG---CGC-----CGG------------------------
----------GCG------CGA------------------------------CCCG----CU----------
------------------------------------------------CCGG-----GGACA-GCGUCAGGU
GGGGAGUUUGACUGGGGCGGUACACCUGUCA-AAGCGUAAC-GCAGGUGUCCU-AAGGCGAGCUCAGGGAGG
CCAGAAACCUCCCGUG-GAGCA-GAAGGGCAA-AAGCUCGCUUGAUC-UUGA--------------------
--------------UUUUCA--GUACGAAUACAGACCGUGAAA-GCGGGGCCUCACGAUCCUUC--------
---UG-A-----CUUUUGGG---------------------UU-UUAAGCAGGAGGUGUCAG-AA--AAGUU
ACCACAGGG-AUAACUGGCUUGUGGCGGCCAAGCGUUCAUAGCGACGUCGCUUUUUGAUCCUUCGAUGUCGG
CUCUUCCUAUCAUUGUGAAGCA-GAAUUCACCAAGCGUUGGAUUGUUCACCCACUAAUAGGGAACGUGAGCU
GGGUUUAGACCGUCGUGAGACAGGUUAGUUUUACCCUACU--------------------G-AUG-------
-------------------------------AU--G-UGUUGUUGCGCUAGU-AAU-CCUGCUCAGUACGAG
AGGAA-CCGCAGGUUCAGACAUU-UGGU-GUAUGUGCUUGGCU-GAGG-AGCCAC-U-GGAGCGAGGCUACC
AUCUG-UGGGAUUAUGACUGAACGCC-UC--UAAGUCAGAAUCCCCCCUAA-----AC--GUAGCGAUACCG
```

```
CAGCGCCG-AGGCGCCU-------------------------------------------------------
------------------------------------------------------------------------
------------------------------------------------------------------------
---------------------CGGUGGGCUCGC----------GAUAG------------------------
--------------------------CC---GGCCGC------------CGCCCCC----------------
---CUCGG--GC-------GG--GC------------GGUCG------------------GUGCGGA---G-
--CGCCGCUCGUG-GUC--GG-----GAC-CGGA-----GCGCGGACA-------GAUGUG-GCGCCGCCUC
----------------------------U----C-CCCCGCCGCGU-----ACCGCAUGUU----------C
G-----------------UGGGGAA---------------------C-CCGGUGCU-----AAA------UC
AUUCGUAGACGACCUGAU-UCU-GGGUCGGGGU-UUCGUACG-UAGCA--------GAGCAGC-UC------
----CCU--CGCUGCGAUCUAUUGAGAGUCAG-CCCU-CGACACAA---GCUUUUGUC--------------
------------------------------------------------------------------------
------------------------------------------------------------------------
------------------------------------------------------------------------
-------------------------------------------------
>LSUe_7897_LATCH/1-3903
------------------------------U----GA--CAACACUUAGC-AGUGGAUC-A-CUCAGC-UCG
U-G---CGUCGAUGAAGAACUCAG--CUAGCUGCGAGAAUUAGUGCGAAU--U---GCA--GGA-CA--CGU
UGAUCAUCAACACUUCAAACGCACCU-U--------CC--------GG----CCCC-G-GGU---------U
A-C-----------UCC---C----G----G---GGC----U-A-UGCC-UGU-----CUGAGGAUCA-UUU
-----------------------------------------------------------UAAUCAACUCA-G
AUC--------------------AG-A-CG-UGACAACCCACUGAAUUUAAGCAUAUUACUAAGCG--GAGG
AAAAGAAACUAA--CCAGGAUUCCCUCA--GU-AA-CGGCGAGUGAAGAGGGA-A-GA-GCCCAGCGCCGAA
UCCC-CGCUCGUCU-GG-CGG-------------G---CGC--GGGAAAUGUGGCG--UA-------------
--CA-GAAG-ACC-C-ACCUC-------------------------CCCGG--CGC-----------CG---
CUC-----GGGGG--CCC-------AAGUCC-UUCU------------------------------------
-------GAU-UG-AGGCUU---AGCCCACGGACGGUGCGAGGCCGGUA---------GCG---G----CCC
CGG---GC--GCGCC-GGGA----------------CCGG-GUCUU--CUCGGAGUCGGGUUGUUUG---GG
AAUGCAGCCCAAA-G-CGGGUGGUAAA-CUCCAUCUAAGGCUAAAUACCGG--C-ACG-AGACCGAUAGU-C
GAC-A-AGUACCGU-AGGGAAAG-UUGAAA-AGAAC-UUUGA-AGAGAG-A-GUUCAAGAGGGCGUGAAACC
G-UUGAGAGG--U-AAA-CG-GGUG---GGGU-CCGCGCAGU--------------------CC--------
-------------------------------------------GCCCGGAG--GAUUCAACUC--GGCG---
------UGCC-GUGCCGGCCGC-CCCGGUGUG-----GCG-GAU----CCC---------------------
----------------------------------------------------CUC-----------GC----
------------------------------------------------------------------------
----------------------------------GGGAA-CG--CCCCCGUCGGCGUG---CGGGCCCCCGC
C-----GGGUGCA-UUUCC---UCCG-CG--GU----------GGU--------GCGCCGCGACCGGCUCC-
---GG-----------GU-C-------------------G------CCU-----------------GGGAUG
GCU-----------------GCGGG-------CGG-AGGUAGCU-GGC------------------------
-----------------------------------------------------------CUCU--C-CG---
------------------------------------------------------------------------
-------GAGAUCAGUGUCU--GCCCCCC-UGCCGCA--------CCUCG-GCGU--CUCC--------CGG
GGCCGUGGAA----GAUGACCGCC----------------GCCGCGCCCUCCUC------------------
-------------------------------------------------------------------UCCCC
CCU---------------------------------------------------------------------
```

```
---------------------------------CGG------------------GGGC--------------
-------------------------------------------------UGAGA-----CAGGGGC------
-CCCCGC------------UGCCGGCGC-------GACUGU-C-----------------------------
---------------------------------------GAAU---G-GGGCG----GACUGUC--------
-----CUCAGUG--CGC-GCC---GG-----ACG-CGUC--------UCGCCGCC--CGG---CCGGG---A
G--------GCU-----------------------------------------------------------CUCU
-------------------------------AUCGGGCGCC-------------------------------
----------------AGGGGUCUGCGG------------CGAUGUCGGU--U---UCCACCCGACCCGUCU
UGAAACACGGACCAAGGAGUCUA-A-CACGUGCGCGA-GUCGAAGG--G-CAG-------AGCG------AA
A-CC--CCAU--GGCGCAAUGAAAG-UGA-----------------GGGCCGGCGCG----GGCCGGCUGAG
GUGG-GAUCCCC----ACGCCCC-------------------------------------GUGC--------
GGG---------------------------------------------------------------------
-------------------GGG--------------------------------------CGCACCACC-AG
CCCGUC----UCACCCG-C--------------------AGCG----------------UCGGGGAGG-UG-
GAG-CA----------------------UGAGCGUGCGUGAUAG-GACCCGAAAGA-UGGUGAACUAUGCCU
GGGCAGGGCGAAGCCAGA-GGAAAC-UCUGGUGGAGGUCCGU--AGCGGUCCUGACGUGCAAAUCGG-UCGU
CCGACCUGGGUAUAGGGGCGAAAGACUAAUCGAACCAUC-UAGUAGCUGGUUCCCUCCGAAG-UUUCCCUCA
GGAUAGCUGGUG-CUCG-------------------------------------------------------
----------------AGCGAACG-----CAGUUUUAUCUGGUAAAGCGAAUGACU-AGA-GGUCUUGG-GG
C-C-GAAAC-GAUC-U-CAACCUAUUCUCAAACUUUAAAUGGGUAAGAA-----------------------
------------------GCCCG-GC-----UCG-CU--GGCU--U--GGA---------------------
GC---CGGG-C---GUGGA-AU-G--C-----G--AGUGCCU-AGUGGGC-C-ACUU-U-UGG-UAAGCAGA
AC-U-GGCGCUGC-GGGAUG---AACCGAACGCCG-G-GUUGUGGCGCCCGAUG--CCGACG-CU-CA---U
-C-AGACCCC-AGAAAAGGUGUUGGUUGAUAUAGACAGCAGGACGGUGGCCAUGGAAGUCGGAA-UCCGCUA
AGGAGUGUGUAACAACUCACCUGCC-----GAAUCAACUAGCCCUGAAAAU-GGAU-GGCGCUGG-AGCGUC
G----------GGCCCAU-ACCCGGCCGUCGCGG--GCGACCGG----------------------------
-----------------------------------------------------------------GGC--
----CAC-----------------------------------------------------------------
------------------------GAUGGCUAUCCCG-CGAC-GAGUAGGAGGGCCGCCGCGG---------
-----------U-GAGCACU-GAAGCCUA-G-GGCGCGGGCC--C-GGGUGGAGCU-G-CCG-CGGGUGCAG
AUCU-UGGUGG-UAGUA-GCAAAU---AUUC-AAACGA--G---------------AACUUU-GAA-GGCCG
AAGUGGAG-AAGGGUUCC-AUGUGAACA-GCAGUUGAACAUGGGUCAGUCGG--UC-CUAAGAGAUGGGCG-
A-A-CGCC-----GUUCCGA--------AGGGACGGGCG----AUGG---CCUC-----------------C
GU---------------CGCCCUC------AGCC----GA-UC-GA-A-AGGGA-GUUGGGUU-CAGA-UCC
CCGAA-----------------------------------UCCGGAAUGGCG------------GAGAGGGGC
G----------------CC-GCUGUU--------------------------------GGCGUCC------
--AGUG-CGC---CAAC--------GC-AA-----AC-------------------GAUC-C-CGGA-GAA
GCCGGCG-------------GGAGCC-CCGGGGAGA-GU------UCUCU-------------U-UUCUUU-
GUGAAG-GGC-AG-----G-G-------------------CG-----CC----------------CUGG-AA
U-G-GG---------------------UUUGC-ACCGAGAGAG---------GGGCCC----GCG-CC-CUG-
GAA--AG-----------------------------------------------------------------
----CGUCGCG-GUUCCGGCGGCGU-C-CGGUGAG-CUCUCGCUGGC-CCUUGAAA-A-UCC-GGGGGAGAA
-G-------------------------------------------GUGU-----AAAUCUCG-CG-CCGGGC
CGUACCCAUAUCCGCA------------------------------GCAGGUC--UCCAAGGU-GAACAGCC
UCUGGCAU-GU-UAGA-ACAAUGUA-G-GUAAGGGAAGUCGGC-AAGCCAGAUCCGUAACUUCGG-GAUAAG
```

```
GAUUG-GCUC-U-AA--GGGCUG-G----GUCGG-UC------GGGCCG--GGG-UGCGAA-GC--GGU-GC
UGGG------UACGCGCCGCG---GC----------------------------UGGACGAGGC-GCCG---
------------------------------------------------------------------------
---------------------CCUC-----------------------------------------------
------------------------------------------------------------------------
---------------------------------------------------CGG------------------
------------------------------------------------------------------------
------------------------------------------------------------------------
-G-CGGCGGCGACUCUGGACGCGAG-CCGG-ACCCUUC----------------------------------
---CUG----UG-GAUCGC-----CU-CAGCUGCG-GCGCGCGC----------------------------
---------------------------------------------------------------GGGC-----
-----------------------------------------------------------CCCC---------
---------------CG-------------------------------GG---G------------------
------------------------------------------------------------------------
----CU--------------------GGCGUCUC-GCCU----CGGCCG-GCG--CCUAGCAGC--UGACUU
AGA--ACUGGUGCGGACCAGGGGAAUCCGA-----CUGUUUAAUUAAAACAAAGCAUCGCGAAGGCCCG---
-CG-G------------UGGGUGUUG-ACGCGAUGUGA-UUUCUGCCCAGUGCUCUG-AAUG-UCAAAGUGA
A--------------GAAA--------------UUCAAUGAAGCGC--GGGUAAAC-GGCGGGAGUAACUAU
GACUCUCUUAAGGUAGCCAAAUGCCUCGUCAUC-UAAUUAGUGACGCGCAUGAAUGGAUGAACGAGA--UUC
CCACUGUCCCUACC----UAC-UAUCUAGCGAAACC---AC---AGCCAAGGGAACGGGCUUGGCAGAAUCA
GCGGGGAAAGAAGA-CCCUGUUGAGC-UUGACU--CUAGUCU-GGCACUGUGAAGAGACAUGAGAGGUGUAG
GAU-AAGUGGAGGC------------------------------------------------------CUCG
--------------------------------------------------G-C--C-GC-CGGUGAAAUA-C
CACUACUCUUAUCGUUUUUUCACUUACCC-GG-------U-GA-G-GCGGG----GGG-----------GC-
------GA--GU-CC--------CGAG-GGGCU-CU-CG-------------AUUCU---------------
-----------GGUG--------------------------------------ACAA--GC---------GC
------------------C--------------------------------------------CG-------
---GUU---------------------------C---GGU-----CGG------------------------
----------GUG------CGA------------------------------CCCG----CU----------
------------------------------------------------CCGG-----GGACA-GUGGCAGGU
GGGGAGUUUGACUGGGGCGGUACACCUGUCA-AACCGUAAC-GCAGGUGUCCU-AAGGCGAGCUCAGGGAGG
ACAGAAACCUCCCGUG-GAGCA-GAAGGGCAA-AAGCUCGCUUGAUC-UUGA--------------------
--------------UUUUCA--GUAUGAGUACAGACCGUGAAA-GCGGGGCCUCACGAUCCUUC--------
---UG-A-----CUUUUUGGG--------------------UU-UUAAGCAGGAGGUGUCAG-AA--AAGUU
ACCACAGGG-AUAACUGGCUUGUGGCGGCCAAGCGUUCAUAGCGACGUCGCUUUUUGAUCCUUCGAUGUCGG
CUCUUCCUAUCAUUGUGAAGCA-GAAUUCACCAAGCGUUGGAUUGUUCACCCACUAAUAGGGAACGUGAGCU
GGGUUUAGACCGUCGUGAGACAGGUUAGUUUACCCUACU---------------------G-AUG-------
-------------------------------AU--G-UGUUGUUGCAAUAGU-AAU-CCUGCUCAGUACGAG
AGGAA-CCGCAGGUUCAGACAUU-UGGU-GUAUGUGCUUGGCU-GAGG-AGCCAA-U-GGUGCGAAGCUACC
AUCUG-UAGGAUUAUGACUGAACGCC-UC--UAAGUCAGAAUCCUCCCUAA-----AC--GUAACAAU-UAA
UAGC-CUG------UUU-------------------------------------------------------
------------------------------------------------------------------------
------------------------------------------------------------------------
---------------------AGCUCCACCCUUU---------GAUUGU-----------------------
----------------------AGCACU---AGUAGG------ACAUUUGAACAA-----------------
```

```
---UUUCAUCAUA------CC--UU------------ACCUG------------------GU----------
-----CUCGCUU--UUU--UG-----AAACCUCA-----GAAAUCCUA-------AAAUGGGGGUUUGUCCC
----------------------------A----G-GGGUGCAACA------GGUAC---------------U
G---------------------------------------------C-AUUGCACU-----A--------CC
A--UGUGGGAGAUCAGGGCUCA-AUCCCUGUUG-UGGAUCUC-CCAC---------AGGCAAC---------
----UAA--C--ACCAAGA--------------CACA-UUACAAAGG--CAAUGCU-C--------------
------------------------------------------------------------------------
------------------------------------------------------------------------
------------------------------------------------------------------------
-------------------------------------------------
>LSUe_7955_DANRE/1-4272
---------------------------------------CAACUCUUAGC-GGUGGAUC-A-CUCGGC-UCG
U-G---CGUCGAUGAAGAACGCAG--CUAGCUGCGAGAACUAAUGUGAAU--U---GCA--GGA-CA--CAU
CGAUCAUCGACACUUCGAACGCACUU-U--------GC--------GG----CCCC-G-GGU---------U
C-C-----------UCC---C----G----G---GGC----C-A-CGCC-UGU-----CUGAGCGUCG-CUU
------------------------------------------UC------------U--UACGAC-CUCA-G
AUC--------------------AG-A-CG-AGACAACCCGCUGAAUUUAAGCAUAUUACUAAGCG--GAGG
AAAAGAAACUAA--CCAGGAUUCCCUCA--GU-AG-CGGCGAGCGAAGAGGGA-A-GA-GCCCAGCGCCGAA
UCCC-CGCCCCGUG-CCAGGG------------G---CGA--GGGAAAUGUGGCG--UA-------------
--CG-GAAG-AUC-GCUCUCU-------------------------CUCGG--CGC-----------GG---
GCC-----GGGGG--UCU-------AAGUCC-UUCU------------------------------------
-------GAU-GG-AGGCUU---AGCCCGUGGACGGUGUGAGGCCGGUA---------ACG---G----CCC
CCG---CC--CCGCC-GGGG----------------UACG-GUCUU--CCCGGAGUCGGGUUGUUUG---GG
AAUGCAGCCCAAA-G-CGGGUGGUAAA-CUCCAUCUAAGGCUAAAUACCGG--C-ACG-AGACCGAUAGU-C
GAC-A-AGUACCGUGAGGGAAAG-UUGAAA-AGAAC-UUUGA-AGAGAG-A-GUUCAACAGGGCGUGAAACC
G-UUGAGAGG--U-AAA-CG-GGUG---GGGU-CCGCACGGU--------------------CC--------
-------------------------------------------GCCCGGAG--GAUUCAACCC--GGCG---
------GGGCUGGUCGGCCCGU-C-CGGUGCG---C-UCG-GAU----CCC---------------------
-----CCC-------------------------------------------CCUC-----------UC----
------------------------------------------GGG-GGAG----------------------
----------------------------------GGGGACCG-GCGCCCG--GACG-GAGCUCGGCCGCCGC
C-----GGGCGCA-CUUCC---UCCGUCG--GC-----------GGU--------GCGCCGCGACCGGCUCC-
---GG-----------UU-C-------------------G------GCU-----------------UGGAAG
G-----------------------GUUCGGGGGCGAAGGUGGCUCGCGGC----------------------
--------------------------------------------------------------U--C-CG---
------------------------------------------------------------------------
-------GCCUCGAGCUUUACAGCGCCCCCCCGCCCCGA------CUUCG-CCGC--UUGCUACC----CGG
GGCCGCGGGC----AGUGUCC-------------------UCCGCGCCUUCUCU------------------
----------------------CCGGCC---------------------------------------CCUCU
CUC--------------------------------------GGGGG--------------------------
------------GGGC-----------------UCG------------------------------------
-------------------------------------------------GAGGGA----CGGGGCC------
-CCUCGC------------UCCCGGCGC-------GUGCGU-C-----------------------------
---------------------------------------GACC---G-GAACG----GACUGUC--------
-----CUCAGUC--CGUUUCC---GA-----CCG-CGCC--------GUGCCGCCCAGGG---CGGGGACAG
```

```
G--------CCC---------ACG--------------------------------------------UCCA
-------------------------------AAAGGGCGCC-------------------------------
----------------CGAGGUCCGCGG------------UGAUGCCGGC--C---ACCACCCGACCCGUCU
UGAAACACGGACCAAGGAGUCUA-A-CGCGCGCGCGA-GUCAGAGG--G-UGC-------CCUC------GA
G-CC--CCCAC-GGCGCAAUGAAGG-UGA-----------------AGGCCGGCGCG----CGCCGGCCCAG
GUGG-GAUCCCC----CCGCCCC--------------------------------------GGC--------
GGG---------------------------------------------------------------------
-------------------GGG--------------------------------------CGCACCACC-GG
CCCGUC----UCGUCCG-C--------------------UCUG----------------UCGGGGAGG-UG-
GAG-CU----------------------AGAGCGCGUGCGAUGG-UACCCGAAAGA-UGGUGAACUAUGCCU
GGGCAGGGCGAAGCCAGA-GGAAAC-UCUGGUGGAGGCCCGC--AGCGGUCCUGACGUGCAAAUCGG-UCGU
CCGACCUGGGUAUAGGGGCGAAAGACUAAUCGAACCAUC-UAGUAGCUGGUUCCCUCCGAAG-UUUCCCUCA
GGAUAGCUGGCG-CUCG-------------------------------------------------------
-------------------CCACG-----CAGUUUUAUCCGGUAAAGCGAAUGACU-AGA-GGCCUUGG-GG
C-C-GAAAC-GACC-U-CAACCUAUUCUCAAACUUUAAAUGGGUAAGAA-----------------------
------------------GCCCG-GC-----UCG-CU--GGCU--U--GGA---------------------
GC---CGGG-C---GUGGA-AU-G--CG----A--GACGCCC-AGUGGGC-C-UUUU-U-UGG-UAAGCAGA
AC-U-GGCGCUGC-GGGAUG---AACCGAACGCCG-G-GUUAAGGCGCCCGAUG--CCGACG-CU-CA---U
-C-AGACCCC-AGAAAAGGUGUUGGUUGAUAUAGACAGCAGGACGGUGGCCAUGGAAGUCGGAA-UCCGCUA
AGGAGUGUGUAACAACUCACCUGCC-----GAAUCAACUAGCCCUGAAAAU-GGAU-GGCGCUGG-AGCGUC
G----------GGCCCAU-ACCCGGCCGUCGACG--GCAGAAGGA---------------------------
-----------------------------------------------GC------------------CCC--
----CCCCUCC-------GGGGGGGG----------------------------------------------
------------------------AAAGGGUAAGCCU-CGAC-GAGUAGGAGGGCCGCCGCGG---------
-----------U-G-GCGCG-GAAGCCUA-G-GGCGUGGGCC--C-GGGUGGAGCC-G-CCG-CGGGUGCAG
AUCU-UGGUGG-UAGUA-GCAAAU---AUUC-AAACGA--G---------------AGCUUU-GAA-GGCCG
AAGUGGAG-AAGGGUUCC-AUGUGAACA-GCAGUUGAACAUGGGUCAGUCGG--UC-CUAAGGGAUGGGCG-
A-A-CGCC-----GUUCGGA--------AGGGAGGGGCG----AUGG---CCUC-----------------C
GU---------------CGCCCCC------GGCC----GA-UC-GA-A-AGGGA-GUCGGGUU-CAGA-UCC
CCGAA----------------------------------CCCGGAGUGGCG------------GAGACGGGC
G----------------CC-GCGA-----------------------------------GGCGCCC------
--AGUG-CGG---UGAC--------GC-AA-----AC--------------------GAAC-C-CGGA-GAA
GCCGGCG-------------GGAGCC-CCGGGAAGA-GU------UCUCU-------------U-UUCUUU-
GUGAAG-GGC-AG-----G-G-------------------CG-----CC----------------CUGG-AA
U-G-GG--------------------UUCGC-CCCGAGAG-----------GGGCCC----GCG-CC-CUG-
GAA--AG-----------------------------------------------------------------
----CGCCGCG-GUUCCGGCGGCGU-C-CGGUGAG-CUCUCGUCGGC-CCUUGAAA-A-UCC-GGGGGAGAA
-G-------------------------------------------GUGU-----AAGUCUCG-CG-CCGGGC
CGUACCCAUAUCCGCA------------------------------GCAGGUC--UCCAAGGU-GAACAGCC
UCUGGCGU-GU-UGGA-ACAAGGCA-G-AUAAGGGAAGUCGGC-AAGUCAGAUCCGUAACUUCGG-GAUAAG
GAUUG-GCUC-U-AA--GGGCUG-G----GCCGG-UC------GGGCUG--AGG-UGCGAA-GC--GGG-GC
UGGG------CCCGCGCCGCG---GC----------------------------UGGGGGAG-CGGCCG--C
CCCG-CGGCGC----CCCCU----------CUCGUUC-----CCG---UCUCCGG---AGCACG-UCGUGCG
-GCCCCUACC----------CCCCG--UCGCCCGCCUCUUG----CUCGC--------GUCCGUCC------
-------------------------GGUCG-----------------------CCCCCGCCUUCG----GGU
```

```
GGGGGA------------GGC------------------------------CGG--GCGGCUCGUGGGC---
UCGGGG----------UGGGU-------GUUGC-GGG------UAGGGGGGG--UG-UCCGCUGCGG--CG-
---CG-----CCGAGGG-UCGG-----GGCCUGC-------GGG-GGGGGAUGC----GCUGC-----GGUG
UG-CGGCGGCGACUCUGGACGUGCG-CCGG-GCCCUUC----------------------------------
---UCG----CG-GAUCUC-----CC-CAGCUACG-GCCCGCGUCG--------------GGACC--U----
-----CCGUC-----------------------------UGGC---------------------GUCU----
-----------------------------------------------------UCUC--CUCU---------
---------------CG---------GGGG----------------GAGG---G------------------
-----CACG-----GGCG-------------------------------GGCGG-------CCUCC------
----CC--------------------GGCGCGGC-GCCU----CGGCCG-GCG--CCUAGCAGC--UGGCUU
AGA--ACUGGUGCGGACCAGGGGAAUCCGA-----CUGUUUAAUUAAAACAAAGCAUCGCGAAGGCCCG---
-UG-G-----------CGGGUGUUG-ACGCGAUGUGA-UUUCUGCCCAGUGCUCUG-AAUG-UCAAAGUGA
A--------------GAAA--------------UUCAAUGAAGCGC--GGGUAAAC-GGCGGGAGUAACUAU
GACUCUCUUAAGGUAGCCAAAUGCCUCGUCAUC-UAAUUAGUGACGCGCAUGAAUGGAUGAACGAGA--UUC
CCACUGUCCCUACC----UGC-UAUCUAGCGAAACC---AC---AGCCAAGGGAACGGGCUUGGCAGAAUCA
GCGGGGAAAGAAGA-CCCUGUUGAGC-UUGACU--CUAGUCU-GGCACUGUGAAGAGACAUGAGGGGUGUAG
AAU-AAGUGGGAGGC-CCUG---------GGGCC----------------------------------CUCG
--------------------------------------GGCCCAGCGGGCG-C--C-GC-CGGUGAAAUA-C
CACUACUCUUAUCGUUUCCUCACUUACCC-GG-------U-UG-G-GCAGG----GAG-----------GC-
------GA--GC-CC-------CCGGC-GGGCU-CC-UG-------------CUUCU---------------
-----------GGCG--------------------------------------UCAA--GC---------GC
------------------C-------------------------------------------UCGG------
---GGCC------------CAU--------CGCG---GUCC----CGG------------------------
----------GCG------CGA------------------------------CCUG----CC----------
------------------------------------------------CCGG-----GGACA-GUGGCAGGU
GGGGAGUUUGACUGGGGCGGUACACCUGUCA-AACGGUAAC-GCAGGUGUCCU-AAGGCGAGCUCAGGGAGG
ACAGAAACCUCCCGCG-GAGCA-GAAGGGCAA-AAGCUCGCUUGAUC-UUGA--------------------
--------------UUUUCA--GUAUGAGUACGGACCGUGAAA-GCGGGGCCUCACGAUCCUUC--------
---UG-G-----CUUUUGGG---------------------UU-UUAAGCAGGAGGUGUCAG-AA--AAGUU
ACCACAGGG-AUAACUGGCUUGUGGCGGCCAAGCGUUCAUAGCGACGUCGCUUUUUGAUCCUUCGAUGUCGG
CUCUUCCUAUCAUUGUGAAGCA-GAAUUCACCAAGCGUUGGAUUGUUCACCCACUAAUAGGGAACGUGAGCU
GGGUUUAGACCGUCGUGAGACAGGUUAGUUUUACCCUACU--------------------G-AUGU------
-------------------------------CC--G-UGUUGUUGCAAUAGU-AAU-CCUGCUCAGUACGAG
AGGAA-CCGCAGGUUCAGACAUU-UGGU-GCGUGUGCUUGGCU-GAGG-AGCCAC-U-GGUGCGAAGCUACC
AUCUG-UGGGAUUAUGACUGAACGCC-UC--UAAGUCAGAAUCCCGCCUAA-----AC--GUAACGAUACCG
CAGCGCCG-CGGGGCUU-------------------------------------------------------
------------------------------------------------------------------------
------------------------------------------------------------------------
---------------------CGAUUGGCCUGG----------GAUAG------------------------
--------------------------CC---GGCCGG------------GGUUC------------------
----------GC-------CC--UC------------GGUCG------------------GUGCGGA---G-
--AGCCAGCCGU--GAC--GG-----GAC-CGGG-----GCGCGGCC--------GAACGA-GCGCCGCCCC
----------------------------U----C-UUCCGGUACGC-----ACCGCAUGUU----------U
G-----------------UGGGGAA---------------------C-CCGGUGCU-----AAA------UG
ACUCGUAGACGACCUGAU-UCU-GGGUCAGGGU-UUCGUGCG-UGGCA--------GAGCAGC-UC------
```

```
----CAU--CGCUGCGAUCCAUUGAAAGUCAG-CCCU-CGAUCCAAG--UUUUGUCGG--------------
------------------------------------------------------------------------
------------------------------------------------------------------------
------------------------------------------------------------------------
-------------------------------------------------
>LSUe_9598_PANTR/1-5004
----------------------------------------UGACUUUUAGC-GGUGGAUC-A-CUUGGC-UCC
U-G---CGUUGAUGAAGAAUGCAG--CUAGCUGUGAGAAUUAAUGUGAAU--U---GCA--GGA-CA--CAU
UGAUCAUCGACACUUCGAACGCA-CU-U--------GC--------GG-----CCC-G-GGU---------U
C-C-----------UCC---C----A----G---GGC----U-A-UGCC-UGU-----CUGAGCUUUC-CUU
--------------------------------------------------------G--CGUGAC-AUCA-G
AUC--------------------AG-A-CG-UGGUGAUCCAUUGAAUUUAAGCGUAUUAGUGAGCG--GAGG
AAAAGAGACCAA--CCAGGAUUCCCUUA--GU-AA-CAGUGAGUGAAGAGGGA-A-GA-GCCCAGUGCUGAA
UCCC-UGCCUUGUG-GU-GGG------------G---UGU--GGGAAAUGUUGCA--UA-------------
--UG-GAAG-GCC-C--UGUC-------------------------CCCGG--CAC-----------CG---
CUGGUG--GGUGG--CCC-------AAGUCC----U------------------------------------
-------GAU-CU-AGGCCA---AGACUGUUGGUGGUGUGAGGCUGGUA---------GCC---A----CCC
CUG---GC--GCGCU-GGGC----------------CCGG-GUCUU--CCUGGAGUUGGGUUGCUUG---GG
AAUGCAGACCAAA-G-GGGGUGGUAAA----CAUCUAAGACUAAAUACUGA--C-ACG-AGAGCGAUAGU-C
AAC-A-AGUACCGUAAGGGAAAG-UUGAAA-AGAAC-UUUGA-AGAGAG-A-GUUCAAGAGGGCGUGAAACC
G-UUAAGAGG--U-AAA-CG-GGUG---GGGU-CCGCGCAGU--------------------CC--------
-------------------------------------------GCCCGGAG--GAUUCAACCC--GGCG---
------GCG--GGUCCGGCCGUGU-CGGCGG--CCCGGCG-GAUCUUUCCCGCCCCCCGUUCCUCCCGACCC
CUCCACCCGCCC--------------UCCCUUCCCCC-CGCCGCCC---CUCCUCC----UCCUCCCC----
-----------------------------------------GGAG-GGGGCGGGCUCCGGCGGGUGC---GG
GGGUGGGCGGGCGGGGCCGGGGGUGGGGUCGGCGGGGGACCG-UCCCCCG--ACCG-GCGACCGGCCGCCGC
C-----GGGCGCA-UUUCC---ACCG-CG--GC----------GGU--------GCGCCGCGACCGGCUCC-
---GG-----------GA-C-------------------G------GCU-----------------GGGAAG
GCC-----------------CGGC-------GGGGAAGGUGGCUCGGGG-------------------GGCC
CCGUCCGUCC---------UCCUCCUCCCCCCCCGUCUCCGCCCCCCGGCCCCGCGUCCUCCC--C-CGGGA
GGGCGCGCGGGUCGGGGCGGUGGCGGCGGCGGC---GGCGGUGGCGGCGGU---------GGCGGCGGGACC
GAAACCCCCCCCGAGUGUUACAG-CCCCCCGGCAGCAGC------ACUCG-CCGA--AUCC--------CGG
GGCCGAGGGAGC--GAGACCCGUC----------------GCCGCGCUCUCCCCCCU--------CCCGGCG
CC------------------CACCCCCGCGGGGA--------------------------------UAUCCU
CCG-----------------------------------CGAGGGGGGUC----------------------U
CC-----CCCGCGGGGG---CG-----------CGC------------------CGGC--------------
GUCUCC-------------------------------------UCGUGGGGGGGC----CGGGCCA------
-CCCCUC------------CCACGGCGC-------GACCGCUCUCCCACCCCUCCUCCCCGCAACCCCCCUC
-UCCCGGCGACGGGGA--GGG--CCGCGCGCGGGUCGGGGGGC---G-GGGCG----GACUGUC--------
-----CCCAGUG--CGC-CCC---GG-----GCG-GGUC--------GCGCCGUC--GGGCCCGGGGG---A
GGUUCUCUCGGGGCCACGCGCGCG--------------------------------------------UCCC
CC-----------GAAGAGGGGGACGGCGGAGCCGAGCGCA-------------------------------
----------------CGGGGUCGGCGG------------CGAUGUCGGC--C---ACCACCCGACCCGUCU
UGAAACACGGACCAAGGAGUCUA-A-CACGUGCGCGA-GUCGGGGG--C-UCG-------CACG------AA
A-GC--CGCCGUGGCGCAAUGAAGG-UGA-----------------AGGCCGGCGCGCU--CGCCGGCCGAG
```

```
GUGG-GAUCCGA----GGCCUC------UC--------------------------CAGUCCGC--------
CGA---------------------------------------------------------------------
-------------------GGG--------------------------------------CGCACCACC-GG
CCCGUC----UCGCCCG-C--------------------CGCG----------------CCGGGGAGG-UG-
GAG-CA----------------------CGAGCGCACGUGUUAG-GACCCGAAAGA-UGGUGAACUAUGCCU
GGGCAGGGC-AAGCCAGA-GGAAAC-UCUGGUGGAGGUCCGU--AGCGGUCCUGACGUGCAAAUCGG-UCGU
CCGACCUGGGUAUAGGGGCGAAAGACUAAUCGAACCAUC-UAGUAGCUGGUUCCCUCCGAAG-UUUCCCUCA
GGAUAGCUGGCG-CUCU----------------------------------------------------CGC
AGACCCGACGCACACCCCCCCACG-----CAGUUUUAUCCGGUAAAGCGAAUGAUU-AGA-GGUCUUGG-GG
C-C-GAAAC-GAUC-U-CAACCUAUUCUCAAACUUUAAAUGGGUAAGAA-----------------------
------------------GCCCG-GC-----UCG-CU--GGCG--U--GGA---------------------
GC---CGGG-----GUGGA-AU-G--C-----G--AGUGCCU-AGUGGGC-C-ACUU-U-UGG-UAAGCAGA
AC-U-GGCGCUGC-GGGAUG---AACCGAACGCCG-G-GUUAAGGCGCCCGAUG--CCGACG-CU-CA---U
-C-AGACCCC-AGAAAAGGUGUUGGUUGAUAUAGACAGCAGGACGGUGGCCAUGGAAGUCGGAA-UCCGCUA
AGGAGUGUGUAACAACUCACCUGCC-----GAAUCAACUAGCCCUGAAAAU-GGAU-GGCGCUGG-AGCGUC
G----------GGCCCAU-ACCCGGCCGUCGCCG--GCAGUCGAGAGUGGACGGG-----AGCGGCGGGGGC
GGCGCGGGCGUGUGCGCGCGCGCGUGUGUGCGUGUGUGUCGGAGGGCGGCGGCGGU---GGCGGCGGGGG--
---UGGGGUCCUCCCCCUCCCCCACG--CCG---CCUCCCCUCCUCCCACCCACCACCGCCGCCGCCACCCC
CGCUCCCCGCCCCCGGAGCCCCGCGGACGCUACGCCG-CGAC-GAGUAGGAGGGCCGCUGCGG---------
-----------U-GAGCCUU-GAAGCCUA-G-GGCGCGGGCC--C-GGGUGGAGCC-G-CCG-CAGGUGCAG
AUCU-UGGUGG-UAGUA-GCAAAU---AUUC-AAACGA--G---------------AACUUU-GAA-GGCCG
AAGUGGAG-AAGGGUUCC-AUGUGAACA-GCAGUUGAACAUGGGUCAGUCGG--UC-CUGAGAGAUGGGCG-
A-G-CGCC-----GUUCCGA--------AGGGACGGGCG----AUGG---CCUC-----------------C
GU---------------UGCCCUC-------GGCC----GA-UC-GA-A-AGGGA-GUCGGGUU-CAGA-UCC
CCGAA----------------------------------UCCGGAGUGGCG------------GAGAUGGGC
G----------------CC-GCGA-----------------------------------GGCGUCC------
--AGUG-CGG---UAAC--------GC-GA-----CC--------------------GAUC-C-CGGA-GAA
GCCGGCG-------------GGAGCC-CCGGGGAGA-GU-------UCUCU-------------U-UUCUUU-
GUGAAG-GGC-AG-----G-G-------------------CG-----CC----------------CUGG-AA
U-G-GG--------------------UUCGC-CCCGAGAGAG---------GGGCCC----GUG-CC-UUG-
GAA--AG-----------------------------------------------------------------
----CGUCGCG-GUUCCGGCGGCGU-C-CGGUGAG-CUCUCGCUGGC-CCUUGAAA-A-UCC-GGGGGAGAG
-G-------------------------------------------GUGU-----AAAUCUCG-CG-CCGGGC
CGUACCCAUAUCCGCA------------------------------GCAGGUC--UCCAAGGU-GAACAGCC
UCUGGCAU-GU-UGGA-ACAAUGUA-G-GUAAGGGAAGUCGGC-AAGCCGGAUCCGUAACUUCGG-GAUAAG
UAUUG-GCUG-U-AA--GGGCUG-G----GUCGG-UU------GGGCUG--GGU--------GC--AUG---
-CCA------C------CACU---GG----------------------------AAGA---GGC-CCUG--C
AGCG-CCCCC--AUGCCCCA-GGG-AUG--CCUGUCGUGGCCCUCCUCUGCCCCAACCCGCGUG-CUGCUCG
-CUCCCUCCCC---------UGCCC--CCUCU-----------------------------CUCCCUG----
-------------------------------------------------------UCCUCCCCCUC--U-CU
GGAGGU------------GGC------GUG---------------------UGG--GGG-------------
---CAGA---------GGGGU-------G--GC-GGUGGCCAUACGGCCCCG-----------GUGG--CA-
---GG-----GCA------UGAUCUCCCGUGAGG-------GGG-GGCU-------------------GGUG
-G-UGGUGGUGACUCUGGACCCGAG-CCGG-GCCCUUCCCAUGGAUUGCCC-----------CAGUU-----
--GUGG----CC-GCCCAU-----GGGGAGCC-CA-GUGGGUGCUG---GCGUGUCCUGUCUUGC-UCCGUG
```

```
-----CUGUCAUGCGUUUCAGCGCGGGCAGGGAGUGAUUGGGUGUCG-----GUGGGU------GGUGGGCG
GGGUGGUUCAUUCCCCCACCUCCCCCGCGGCCCCAUCUGUCUCG--UG----UCCCC--CCCU---------
---------------CC---------------------------GUGGAG---G------------------
----UGGCG-----GGCC---------------------------ACGGGCUGGUCC-CCACCACCGGG---
----GCGG-----CGGUU-------CCGUGCAGC-GCCU----UGGCCG-GUG--CUUAGCAGU--CGACUU
AUAGAACUGGUGUGCACCAGGGGAAUCCGA-----CUGUUUAAUUAAAACAAAGCAUCGCGAAGGCCCU---
-CG-G------------GGGGUGUU----GCAAAGUGA-UUUCUGCCCAGUGCUCUG-AAUG-UCAAAAUGA
A--------------GAAA--------------UUCAAUGAAGUAU--GGGUAAAU-GGCAGAAGUAACUAU
GGCUCUCUUAAGGUAGCCAAAUGCCUCGUCAUC-UAACUAAUGACGUGCAUGAAUGGAUGAAUGAGA--UUC
CCACUGUCCCUACCUACUUAC-UAUCCAGUGAAACC---AC---AGCCAAGGGAAUGGGGUUGGUGGAAUCA
CUGGGGAAAGAAGA-ACCUGUUGAGC-UUGACU--GUAGUCU-GGCAUGGUGAAGAGACAUGAGAGGUGUAG
AAU-AAGUGGGAGGUCCCC---------GGUGCCC-------------CCUAUCCCU-----------GUGA
----------------------GGGGGUGGGGUGG-----GGUCCUGUGGG-C--U-GC-CCAUGGAAUA-C
UACUACUAUGAUCAUUUUUUCACUGACCU-GA-------U-GA-G-GCAGG-----GG-----------GU-
------GA--GC-CC--------CAAG-AGGCU-CU-CC-------------CUUCU---------------
-----------GGCG--------------------------------------CUAA--GU---------GC
------------------C--------------------------------------------CA-------
---GAUG------------CAU---------GCU---GGC-----CGG------------------------
----------GUGU-----UGA------------------------------CCCA----UU----------
------------------------------------------------CCGA-----GGACA-GUGCCAGGU
GGGGAGUUUGACUGGGGCAGUACACCUGUCG-AACAGUAAU-ACAGGUGUCCU-AAGGCGAGCUCAGGGAGG
ACAGAAACCUCUUGUG-GAGCA-GAAGGACAA-AAGCUCGCUUGAUC-GUGA--------------------
--------------UUUUCA--GUGCGAACACAGACUG-AAAG-CCGGGCUUUGACGAUCAUUC--------
---UG-A-----CGU--------------------------UU-UUAAGCAGGAGCUGUCAG-AG--AAGUU
ACCACGGGG-AUAACUGGCUCGUUGUGGCCAAGCGUUCAUAGGGAUGUUGCUUUUUGAUCCUUUGAUGUUGG
UUCUUCCUAUCAUUGUGAAGUA-GAAUUCACCAAGCAUUGGAUUGUUCACCCACUAAUGGGGCAUGUGAGCU
GGAUUUAGACCUUCGUGAGACAGGUUAGUUUACCCUCCG---------------------U-GAU-------
-------------------------------GU--G-UGUUGUUGCCAUGGU-AAU-CCUGCUCAGCAUGAG
AGGAA-CCACAGGUUCAGAC-----------AUGUGCUUGGUU-GAGG-AGCCAA-U-GGGGUGAAGCUACA
GUCUG-UUGGAUUGUGACUGAACGCC-UC--UAAGUCAGAAUCCCGCCCAA-----GA--GGAAGGAUACAG
CAGCGCCG-UGGAGCCU-------------------------------------------------------
------------------------------------------------------------------------
------------------------------------------------------------------------
---------------------UGGUUGGCCUUGGCUGGCCUUGGCUAGCCGGUUCCCCGCCCCCGCUGGU--
-----------------------GGGCU---GCCUCG---C--CCUGGGUGGGGU-------GUG-------
---CCCCA--CC-------AU--GUGCCAGGACU-GCGUCCG------------------GUGCAGA---G-
--UGCCCUUCUU--CCU--GG-----GAA---GG-----GUGUGGCA--------GAAAA--GGCGGCCUCC
----------------------------U----U-GCCUGUCACGC-----AACACACGUU----------C
G-----------------UGGGGAA---------------------C-CUGGUGCU-----AAA------CC
AUUGGUAGACGACCUGCU-UCC-GGGUUGGAGU-UUCAUAAG-UAGCA--------GGGCAGC-UC------
----CCU--UGCUGUGAUCUAUUGAAAGUCAG-CCCU-UGACACAAG--GGUUUGU-C--------------
------------------------------------------------------------------------
------------------------------------------------------------------------
------------------------------------------------------------------------
-------------------------------------------------
```

```
>LSUe_8355_XENLA/1-4247
------------------------------U----CG--CGACUCUUAGC-GGUGGAUC-A-CUCGGC-UCG
U-G---CGUCGAUGAAGAACGCAG--CUAGCUGCGAGAAUUAGUGUGAAU--U---GCA--GGA-CA--CAU
UGAUCAUCGACACUUCGAACGCACCU-U--------GC--------GG----CCCC-G-GGU---------U
C-C-----------UCC---C----G----G---GGC----C-A-CGCC-UGU-----CUGAGGGUCG-CUC
--------------------------------------------------------C--UCAGAC-CUCA-G
AUC--------------------AG-A-CG-CGGCGACCCGCUGAAUUUAAGCAUAUUACUAAGCG--GAGG
AAAAGAAACUAA--CCAGGAUUCCCCCA--GU-AA-CGGCGAGUGAAGAGGGA-A-GA-GCCCAGCGCCGAA
UCCC-GCGCCCGC-----CGG------------G---CGC--GGGACGUGUGGCG--UA-------------
--CG-GGAG-ACC-GGACCCC-------------------------CCCGG--CGC-----------GG---
CUC-----GGGGG--CCC-------AAGUCC-UUCU------------------------------------
-------GAU-CG-AGGCCC---AGCCCGCGGACGGUGUUAGGCCGGUG---------GGC---G-G--CCC
CCG---GC--GCGGC-GGGA----------------CCCG-GUCUC--CUCGGAGUCGGGUUGUUUG---GG
AAUGCAGCCCAAA-G-CGGGUGGUAAA-CUCCAUCUAAGGCUAAAUACCGG--C-ACG-AGACCGAUAGC-G
GAC-A-AGUACCGUAAGG-AAAG-UUGAAA-AGAAC-UUUGA-AGAGAG-A-GUUCAAGAGGGCGUGAAACC
G-UUAAGAGG--U-AAA-CG-GGUG---GGGC-CGU-GCGGU--------------------CC--------
-------------------------------------------GCCCGGAG--GAUUCAACCC--GGCG---
------GGUCAGCGCCGCCGGGAC-CGG-GCCACUCGGCG-GAC----CCC---------------------
-----CCCGCCG---------------GCCC----------CCU-------CCCC--------UCGCC----
------------------------------------------GGG-AGGGCGG--UC--GCG----------
-------CGGG-----------------------GGGGA-CG-CGGCCCG--GGCG--------CCCGGCGC
C-----GCG-----UUUCC---UCCG-CG--GC----------GGU--------GCGC--GCGCCGGCUCC-
---GG-----------GC-C-------------------G------CGU-----------------GGGAAG
GCC-----------------CAGGGGUUCA-GGGGAAGGUGGCCGGC-------------------------
-----------------------------------------------------------CGCC--C-CC---
------------------------------------------------------------------------
---------GCGCGCGGCUACAGCCCCCCCCC-AGCAGC------AGCACUCGCCGUCGCC--------CGG
GGCCGAGGGA----GACGCCGGCC----------------UCCGCGGUCCUCCUCCC--------CGGAGCG
CG------------------UCCCGCCG---------------------------------------CUCCC
CCC-----------------------------------CG-GGGGGG-------------------------
--------CGGCGCGCG---GGG----------CGG------------------GGAA--------------
-------------------------------------------------GGGGGA----AGGGGCC------
-CCCCGC------------UCCCGGCGC-------GGCUGU-C-----------------------------
---------------------------------------AACC---G-GGGCG----GACUGCC--------
-----CCCAGUG--CGC-CCC---GU-----CCG-CGCC--------GCGCCGCC--GAG---GCGGG---A
G--------GGC----------CG--------------------------------------------CCG-
---------------------------------GGAGCCGC-------------------------------
----------------CAGGGUCCGCGG------------CGAUGUCGGU--G---UCCACCCGACCCGUCU
UGAAACACGGACCAAGGAGUCUA-A-CGCGCGCGCGA-GUCGGAGG--G-ACU------CUGCG------CG
A-AA--CCCUGUGGCGCAAUGAAGG-UGA-----------------GGGCCGGGGCG----CCCCGGCUGAG
GUGG-GAUCCCG----CCGCCCCUC---CCUCCGCCCCCCCGGGGGCG-GGG---GGGGGGGCGC--------
CGG---------------------------------------------------------------------
-------------------CGG--------------------------------------GCCACCACC-GG
CCCGUC----UCGCCCG-C--------------------CCCG-----------------UCGGGGGU-GG-
GCG-UG----------------------AGCGCGCGCG-AUUAG-GACCCGAAAGA-UGGUGAACUAUGCCU
```

```
GGGCA-GGCGAAGCCAGA-GGAAAC-UCUGGUGGAGGUCCGU--AGCGGUCCUGACGUGCAAAUCGG-UCGU
CCGACCU-GGUAUAGGGGCGAAAGACGAAUCGAACCAUC-UAGUAGCUGGUUCCCUCCGAAG-UUUCCCUCA
GGAUAGCUGGCG-CUGU-------------------------------------------------------
------------------CCGUCG-----CAGUUUUAUCCGGUAAAGCGAAUGAUU-AGA-GGUCUUGG-GG
C-CGAA-AUCGAUC-U-CAACCUAUUCUCAAACUUUAAAUGGGUAAGAA-----------------------
------------------GCCCG-GC-----UCG-CU--GGCU--U--GGA---------------------
GC---CGGGGC---GUGGA-AU-G--CG----G--CACGCCAUAGUGGGC-C-ACUU-U-UGG-UAAGCAGA
AC-U-GGCGCUGC-GGGAUG---AACCGAACGCCG-G-GUUAAGGCGCCCGAUG--CCGACG-CU-CA---U
-C-AGACCCC-AGAAAAGGUGUUGGUUGAUAUAGACAGCAGGACGGUGGCCAUGGAAGUCGGAA-UCCGCUA
AGGAGUGUGUAACAACUCACCUGCC-----GAAUCAACUAGCCCUGAAAAU-GGAU-GGCGCUGG-AGCGUC
G----------GGCCCAU-ACCCGGCCGUCGCCG--GCGCUGGGU---------------------------
-----------------------------------------------CA------------------GUC--
----CGC-----------------------------------------------------------------
------------------------GGGGGCUAGGCCG-CGACUGAGUAGGAGGGCCCCGGCGG---------
-----------U-GGGCGCG-GAAGCGC--G-CGCGAGGGCC--C-GGGUGGAGCC-G-CCG-CGGGUGCAG
AUCU-UGGUGG-UAGUA-GCAAAU---AUUC-AAACGA--G---------------AACUUU-GAA-GGCCG
AAGUGGAG-AAGGGUUCC-AUGUGAACA-GCAGUUGAACAUGGGUCAGUCGG--UC-CUAAGAGAUGGGCG-
A-G-CGCC-----GUUCGGA--------AGGGACGGCG----AUGG---CCUC------------------C
GU---------------CGCCCUC------GGCC----GA-UC-GA-A-AGGGA-GUCGGGUU-CAGA-UCC
CCGAA----------------------------------CCCGGAGUGGCG------------GAGACGGGC
G----------------CCCGCGGCCCCCCCCGACGCCUCGCGGCGGCGGGGGGGCGGGGGCGUCC------
--AGUG-CGG---CGAC--------GC-GA-----CC--------------------GAUC-C-CGGA-GAA
GCCGGGG-------------GAGCCC-GGGAGAAGA-GU------UCUCU-------------U-UUCUUU-
GUGAAG-GGC-AG-----G-G-------------------CC-----GC----------------CCUG-AA
U-G-GG--------------------UUCGC-CCCGAGAGAG---------GGGCCC----GCG-CCGUUG-
GAA--AG-----------------------------------------------------------------
---CGUCGCGG-UUCCGGCGGCGUC-C-GGUGAGC-UUCUCGCUGGC-CCUUGAAA-A-UCC-GGGGGAGAG
-G-------------------------------------------GUGU-----AAAUCUCUGCG-CCGGGC
CGUACCCAUAUCCGCA------------------------------GCAGGUC--UCCAAGGU-GAACAGCC
UCUGGCAU-GU-UAGA-ACAAUGUA-G-GUAAGGGAAGUCGGC-AAGUCAGAUCCGUAACUUCGG-GAUAAG
GAUUG-GCUC-U-AA--GGGCUG-G----GUCGG-UC------GGGCUG--GGG-CGCGAA-GC--GGG-GC
UGGG------C-CGCGCCGCG---GC----------------------------UGGACGAAGGCGCCC--C
CGUG-GCGCU--------------------CUU----------------UCCCU-----------CCUCCCG
-CCCCCCUCU----------CUCUC--CGG------------------------------------------
-------------------------------------------------------CCCCCCCUC------GC
GGGG--------------GGC-------CG---------------------CGG--GGG-------------
---------------CGGGGG-------G--GC------------------------------GCGG--GG-
---GG-------------CCG-----------GG-------AGC-GCC----------------------CG
-G-CGGCGGCGACUCUGGACGCGCG-CCGG-GCCCUUC----------------------------------
---CUG----UG-GAUCGC-----CC-CAGCUGCG-GCGCGCCUCU-CCC-----------------CCGCG
-----CCGUC-----------------------------CCCCUCC------------------UGCG----
--------------------------------CC------------------UCCCC--CCGU---------
---------------CA--------------------------GGGGACG---G------------------
-----GGCGC----GGCG----------------------------CGGGGCGG----------CCGGG---
----GC---------------------GGGCCCG-GCCU----CGGCCG-GCG--CCUAGCAGC--UGACUU
```

AGA--ACUGGUGCGGAC--AGGGAAUCCGA-----CUGUUUAAUUAAAACAAAGCAUCGCGAAGGCCCG---
-AG-G------------CGGGUGUUG-ACGCGAUGUGA-UUUCUGCCCAGUGCUCUG-AAUG-UCAAAGUGA
A--------------GAAA--------------UUCAAUGAAGCGC--GGGUAAAC-GGCGGGAGUAACUAU
GACUCUCUUAAGGUAGCCAAAUGCCUCGUCAUC-UAAUUAGUGACGCGCAUGAAUGGAUGAACGAGA--UUC
CCACUGUCCCUACC----UAC-UAUCUAGCGAAACC---AC---AGCCAAGGGAACGGGCUUGGCGGAAUCA
GCGGGGAAAGAAGA-CCCUGUUGAGC-UUGACU--CUAGUCU-GCAACUGUGAAGAGACAUGAGAGGUGUAG
GAU-AAGUGGGAGGCCCCC------------------------------GCGCUCGUC----------GCAA
-A-------------------------------------------GGGGCG-C--C-GC-CGGUGAAAUA-C
CACUACUCUUAUCGUUUUUUCACUUACCC-GG-------U-GA-G-GCGGG------G-----------GC-
------GA--GC-CC--------CGAG-GGGCU-CU-CG-------------CUUCU---------------
-----------GGAC--------------------------------------CCAA--GC---------GC
---------------------------------------------------------------CG-------
---GCCC------------------------CCG---CGC-----CGG------------------------
----------GCG------CGA------------------------------CCCG----CU----------
------------------------------------------------CCGA-----GGACA-GUGGCAGGU
GGGGAGUUUGACUGGGGCGGUACACCUGUCA-AACCGUAAC-GCAGGUGUCCU-AAGGCGAGCUCAGGGAGC
ACAGAAACCUCCCGUG-GAGCA-GAAGGGCAA-AAGCUCGCUUGAUC-UUGA--------------------
--------------UUUUCA--GUAUGAAUACAGACCGUGAAACGCGGGGCCUCACGAUCCUUC--------
---UG-A-----CUUUUUGGG--------------------UU-UUAAGCAGGAGGUGUCAG-AA--AAGUU
ACCACAGGG-AUAACUGGCUUGUGGCGGCCAAGCGUUCAUAGCGACGUCGCUUUUUGAUCCUUCGAUGUCGG
CUCUUCCUAUCAUUGUGAAGCA-GAAUUCACCAAGCGUUGGAUUGUUCACCCACUAAUAGGGAACGUGAGCU
GGGUUUAGACCGUCGUGAGACAGGUUAGUUUUACCCUACU--------------------G-AUG-------
-------------------------------AU--G-UGUUGUCGCAAUAGU-AAU-CCUGCUCAGUACGAG
AGGAA-CCGCAGGUUCAGACAUU-UGGU-GUAUGUGCUUGGCU-GAGG-AGCCAA-U-GGGGCGAAGCUACC
AUCUG-UGGGAUUAUGACUGAACGCC-UC--UAAGUCAGAAUCCCCCCUAA-----AC--GUGACGAUACCG
CAGCGCCG-CGGAGCCU-------------------------------------------------------
------------------------------------------------------------------------
------------------------------------------------------------------------
---------------------CGGUCGGCCUCG----------GAU---------------------U----
-------------------------------AGCCGG------------CGCCCC-----------------
---CCCCG--GG-------GG--GC-------------GCCG------------------GCGGGCA---G-
--AGCCGCUCGC--CUC--GG-----GAC-CGGA-----GCGCGGAC--------GAAAGGGGG-CCGCCUC
----------------------------U----C-UCCCGGAGCGC-----ACCGCACGUU----------C
G-----------------UGGGGAA---------------------C-CUGGUGCU-----AAA------UC
AUUCGUAGACGACCUGAU-UCU-GGGUCAGGGU-UUCGUGCG-UAGCA--------GAGCAGC-UA------
----CCU--CGCUGCGAUCUAUUGAAAGUCAU-CCCU-UGGCCAAGC--UUUUGU--C--------------
------------------------------------------------------------------------
------------------------------------------------------------------------
------------------------------------------------------------------------
-------------------------------------------------
>LSUe_6238_CAEBR/1-3661
---------------------------------------CUAGCUUCAGC-GAUGGAUC-G-GUUGCA-UCG
A-G---UAUCGAUGAAGAACGCAG--CUUGCUGCGUUACUUACCACGAAU--U---GCA--GAC-G---CUU
AGAGUGGUGAAAUUUCGAACGC-AN--A--------GC--------AC----CGAC-U-GG---------GU
C-U------------CC---A----G----U---CGG----U-A-CGUC-UGG-----UUCAGGGUUG-UU-

```
-----------------------------------------------------------AUCAAC-CUGA-A
CUC--------------------AG-U-CG-UGAUUACCCGCUGAACUUAAGCAUAUCAUUUAGCG--GAGG
AAAAGAAACUAA--AAAGGAUUCCCUUA--GU-AA-CGGCGAGUGAAACGGGA-U-GA-GCCCAGCGCCGAA
UCGA-UCAGCCUA-----UGG------------U---UGC-UUCGAAAUGUGGCG--UA-------------
--UA-GGUG-UAG-U----CU-------------------------UCCAG--CAG-----------UG---
UC------GUGAG--UCCG------AAGUCC-UUAC------------------------------------
-------GAU-UG-AGGCCAU--AGACCAGAGAGGGUGCGAGUCCCGUC---------CUG---G----AUU
GCG---AU--GCUGUUGGUU----------------CGAU-UGCUC--CUUGGAGUCGGGUUGCUUG---AA
AGUGCAGCCUAAA-G-UGGGUGAUAAA-CUUCAUCUAAGGCUAAAUAUUGA--C-UCG-AUUGCGAUAGC-G
AAC-A-AGUACCGUGAGGGAAAG-UUGCAA-AGGAC-UUUGA-AGAGAG-A-GUUCAAGAGAACGUGAAAUC
G-CUGGAGUG--G-AAC-CG-GAGA---CAGU-UGA-UG-----------------------UU--------
-------------------------------------------GCUUGGAG--AC--AAGUUU--GGUG---
--------------------GC---UGG--------------------UCGC--------------------
-------------------------------------------------------------UUAGUUG----
------------------------------------------------------------------------
----------------------------------UGA----------UCG------------------UCGC
C-----GGAUG---UCGUU---UCCU-AU--GC----------------------------U-ACGCCGAC-
---GG--------------C-------------------GUUGGUCGCU-----------------CGUUCU
GGC-----------------CCGA----------UAGUGUUGCCAGC-------------------------
-------------------------------------------------------------UC--G-CAA--
------------------------------------------------------------------------
------------------------GAGUCGGUGUCUUG-----------------UUAUCGGUCGUUGGUUC
GUGGUGGCUAGCGUU---------------------------------------------------------
------------------------------------------------------------------------
------------------------------------------------------------------------
------------------------------------------------------------------------
------------------------------------------------------------------------
---------------------UAGUUAC--------------------------------------------
--------------------------------------------------GC--------------------
-------UAGUG--UGU------------------------------GUGAC--------------------
------------------------------------------------------------------------
------------------------------------------------------------------------
--------------------GUCGGUGU------------GAAAGUCGAC--G---ACUUUCCGACCCGUCU
UGAAACACGGAUUGCGGAGUGCU-U-GUCUACUGCGA-GUCAAAGG--G-UG--------UUAA-------A
A-CC--UUGC--GGCGAAAUGAAAG-UGA-----------------AGGUCAGUCUU---GAAUUGGCCGAC
GUGG-GAUCGGU----GCUCCUCG--------------------------------------GA--------
GUG---------------------------------------------------------------------
-------------------CUG--------------------------------------CGCACCACG-GC
CCUGUG----CG-UGUC----------------------ACUUG----------------UGACUGUG-CA-
GAG-GU----------------------UGAGCAGUUGGCAAAC-GACCCGAAAGA-UGGUGAACUAUGCCU
GAGCAGGAUGAAGCCAGA-GGAAAC-UCUGGUGGAAGUCCGU--AUCGGUUCUGACGUGCAAAUCGA-UCGA
UAGACUUGGGUAUAGGGGCGAAAGACUAAUCGAACCAUC-UAGUAGCUGGUUCCUUCCGAAG-UUUCCCUCA
GGAUAGCUGGAU-CUCC-------------------------------------------------------
----------------------GG-----CAGUUAUAUUCGGUAAAGCUAAUGAUU-AGA-GGCCUUGG-GG
A-C-GUAAU-GUCC-U-CAACCUAUUCUCAAACUUUCAAUGGAUAUGAA-----------------------
```

```
------------------GUUGC-AG-----UUU-CU--UUAG--U--GAA---------------------
CU---GUCA-A---CGUGA-AU-G--C-----G--AGGUCCA-AGUGGGC-C-AUUU-U-UGG-UAAGCAGA
AC-U-GGCGCUGU-GGGAUG---AACCAAACGUGG-A-GUUAAGGUGCCUAACU--UC-UCG-CU-CA---U
-G-AGACCCC-AUAAAAGGUGUUGGUUGAUAUUGACAGCAGGACGGUGGCCAUGGAAGUCGGUA-UCCGCUA
AGGAGUGUGUAACAACUCACCUGCC-----GAAUCAACUAGCCCUGAAAAU-GGAU-GGCGCUUA-AGCGAG
A----------GACCUAU-ACUCCGCCGUUGCGG--CAUGU-------------------------------
--------------------------------------------------------------------GC--
----GUUGU---------------------------------------------------------------
-----------------------CUAGCCAGGCCG-UAAC-GAGUAGGAAGGUCGUGGCGG-----------
-----------U-U-GCUUU-GAAGGCUAUG-AGCGUGGGCU--C-GGCUGGAGCU-U-CCG-UCAGUGCAG
AUCG-UAAUGG-UAGUA-GCAAAU---AUUC-AAGUUC--G---------------AUCCUU-GAA-GACUG
AAGUGGAG-AAGGGUUCC-ACGUGAACA-GUAGUUGGAUGUGGGUCAGUCGA--UC-CUAAGGUACUGGCG-
A-A-CGCC--UU-GUAUCAU--------CGGU----GGCGA--AAAG---CUUG----------------CU
UU---------------UGGUCCC------CGUU----UG-UC-GA-A-AGGGA-AUAGGGUU-AAUA-UUC
CCUAA----------------------------------CUGAGAUGCAAAG--------A-UUGUG-----
U----------------UC-UUCG-----------------------------------GAGCACA------
--AGCG-CGG---UAAC--------GC-AU-----UC--------------------GAAC-U-UGGU-UA-
GUCGCUC-------------AAAGAC-CGAGCUAGA-GU------UUUCU-------------U-CUCUAG-
UUAAGG-AAC--G-----G-A-------------------CU-----CC----------------CUGG-AA
U-U-GG--------------------UUCAG-CCAGAGAUGG---------GGACGU----UGU-UU-CCG-
AAA--AG-----------------------------------------------------------------
----CACCGCG-GUUUCUGUGGUGU-C-UCGUGCU-CUUUGAACGGC-CCUU-AAA-A-CAC-CAAGGGAGG
-C---------------------------------------------UA-----UUAAUUUG-CA-CUCAAU
CGUACCGAUAUCCGCA------------------------------UUAGGUC--UCCAAGGU-GAACAGCC
UCUAGU-C-GA-UAGA-AUAAUGUA-G-GUAAGGGAAGUCGGC-AAACUAGAUCCGUAACUUCGG-GAAAAG
GAUUG-GCUC-C-AG--UGGUUG-G----AACGA-UU------GGCCAGUUGGU---UGAU-GCUUGUC---
-CGG-------------CGCU---AU---------------------------------------UCUG--U
C---------------------------------------------------------------------UG
-CUUG------A--------UGCUC-----------------------------------------------
----------------------------------------------------------------------UC
GGGCUG----------AUG---------------------------------GC--GG--------------
---------------------------------------------------------------ACUA-----
------------------------------------------------------------------------
-G-UG---------------AU--A-GU---GCUU-GC----------------------------------
---UUG----UG-GA-CGC-----UUUCUGGU----GUG---------------------------------
----------------------------------------------------------------UGCU----
------------------------------------------------------AGG--ACCU---------
---------------C------------------------------GGUU---C------------------
------------------------------------------------------------------------
----UA-----------------------GU----AUCC----UGAUCGCUCAU-CUAAACAAC--CGUACU
GGA--ACCGGUACGGACUCAGGGAAUCCGA-----CUGUCUAAUUAAAACAGAGGUGACAGAUGGUCCU---
-UG-C-------------GGACGCUG-ACUGUCACUGA-UUUCUGCCCAGUGCUCUG-AAUG-UUAAAUCGU
A--------------GUAA--------------UUCGAGUAAGCGC--GGGUAAAC-GGCGGGAGUAACUAU
GACUCUCUUAAGGUAGCCAAAUGCCUCGUCAUU-UAAUUGUUGACGCGCAUGAAUGGAUUAACGAGA--UUC
CUACUGUCCCUAAC----UAC-UAUCUAGCGAAACC---AC---AGCCAAGGGAACGGGCUUGGCAAAAAUA
```

```
GCGGGGAAAGAAGA-CCCUGUUGAGC-UUGACU--CUAGUUU-GACAUUGUGAAGAGUCAUGAGAGGUGUAG
CAU-AGGUGGGAGAC-----------------------------------------------------UUCG
--------------------------------------------------G-U--C-GC-CGGUGAAAUA-C
CACCACUUUCAUCGACUCUUUACUUAUUC-GG-------U-UA-A-AAGAG------A-----------AU-
------UG--GC-UU--------CAC---GGCC-U--U--------------UUUUC--------------G
----------AAGCA--------------------------------------UUAA--GC---------GG
-----------------AG--------------------------------------------CC-------
---AUUU--------------------------U---AUG-----GCA------------------------
----------CCG------UGA------------------------------CUCU----CC----------
------------------------------------------------UCGA-----AGACA-GUGUCAAGC
GGGGAGUUUGACUGGGGCGGUACAUCUAUCA-AAUCGUAAC-GUAGGUGUCCU-AAGGCGAGCUCAGAGAGG
ACGGAAACCUCUCGUA-GAGCA-AAAGGGCAA-AAGCUUGCUUGAUC-UUGA--------------------
--------------CUUUCA--GUACGAGUACAGACCGCGAAA-GCGUGGCCUAUCGAUCCUUU--------
---UA-A-----UCCUGAUUG--------------------UU-UCAGGUAAGAGGUGUCAG-AA--AAGUU
ACCACAGGG-AUAACUGGCUUGUGGCAGCCAAGCGUCCAUAGCGACGUUGCUUUUGAUCCUUCGAUGUCGG
CUCUUCCUAUCAUUGCGAAGCA-GAAUUCGCCAAGCGUUGGAUUGUUCACCCACUAAUAGGGAACGUGAGCU
GGGUUUAGACCGUCGUGAGACAGGUUAGUUUUACCCUACU--------------------G-UUG-------
-------------------------------AC--U-UGUUAUUGCGAAAGU-AAU-CCUGCUUAGUACGAG
AGGAA-CAGCGGGUUCAAACAUU-UGGU-UCAUAGACUUGAUC-GACA-GAUCAA-U-GGUCUGAAGCUACC
AUUUG-AGAGAUUAUAACUGAACGCC-UC--UAAGUUAGAAUCUCGCCUUG-----UC--AAGGCGAAAAUU
UCUUGCUU-CCCGGUGU-------------------------------------------------------
------------------------------------------------------------------------
------------------------------------------------------------------------
---------------------CGGGAGGCAUCU-----------------------CU--------------
-------------------------------AUCUCG------------UGGCAA-----------------
-------C--AC-------GA--GA-------------GCUU------------------AUGC--------
-----CCUAUGU--A---UGG-----CCU-UGGC-----UUCGUAGU--------GAAC---UCUGCGAAGC
----------------------------U----U-GCCAACGCCA-----GAUCACU-CUG----------G
U-----------------UCAAUGU---------------------C-GGGGCGCU-----AAA------UC
ACUUGCAUACGACUUGGUCUCU-UGGUCAAGGU-GUUGUAUU-CAGUA--------GAGCAGU-CC------
----UCUUAUACUGCGAUCUGUUGAGACU-AU-CCUU-UGAUUGAG---UUAGUU--G--------------
------------------------------------------------------------------------
------------------------------------------------------------------------
------------------------------------------------------------------------
-------------------------------------------------
>LSUe_6239_CAEEL/1-3661
---------------------------------------CUAGCUUCAGC-GAUGGAUC-G-GUUGCA-UCG
A-G---UAUCGAUGAAGAACGCAG--CUUGCUGCGUUACUUACCACGAAU--U---GCA--GAC-G---CUU
AGAGUGGUGAAAUUUCGAACGC-AU--A--------GC--------AC----CAAC-U-GG---------GC
C-U------------CC---A----G----U---UGG----U-A-CGUC-UGG-----UUCAGGGUUG-UU-
-----------------------------------------------------------CUCAAC-CUGA-A
CUC--------------------AG-U-CG-UGAUUACCCGCUGAACUUAAGCAUAUCAUUUAGCG--GAGG
AAAAGAAACUAA--AAAGGAUUCCCUUA--GU-AA-CGGCGAGUGAAACGGGA-A-GA-GCCCAGCGCCGAA
UCGA-UCAGUCUU-----UGG------------C---UGC-UUCGAAAUGUGGCG--UA-------------
--UA-GGUG-UAA-G----UU-------------------------UCCAG--CAG-----------UG---
```

```
UC------GUAUG--UCCG------AAGUCC-UUAC------------------------------------
-------GAU-UG-AGGCCAU--AAACCAGAGAGGGUGCGAGCCCCGUU---------CUG---G----AUA
GCG---GC--ACUGUUGGUU----------------CGCU-UGCUC--CUUGGAGUCGGGUUGCUUG---AA
AGUGCAGCCUAAA-G-UGGGUGAUAAA-CUUCAUCUAAGGCUAAAUAUCGA--C-UCG-AUUGCGAUAGC-G
AAC-A-AGUACCGUGAGGGAAAG-UUGCAA-AGGAC-UUUGA-AGAGAG-A-GUUCAAGAGAACGUGAAAUC
G-CUGGAGUG--G-AAC-CG-GAGA---CAGU-UGA-UG-----------------------UU--------
-------------------------------------------GCUUGGAG--AC--AAGCUU--GGUG---
--------------------AC---UGG--------------------UCGC--------------------
-------------------------------------------------------------UUAGUUG----
------------------------------------------------------------------------
----------------------------------UGA----------UCG------------------UUGC
C-----GGGUG---UCGUU---UCCU-AU--GC----------------------------U-ACGCCGAC-
---GG--------------C-------------------GUUGGCUGCU-----------------CGUUCU
AGC-----------------CCGA----------CAGUGUUGCCCAUC------------------------
-------------------------------------------------------------UC--G-CAA--
------------------------------------------------------------------------
------------------------GAGAAGGUGUCUUG-----------------CUGGCGGUAGUGGGUUC
GUGGCGGCUAGCGUU---------------------------------------------------------
------------------------------------------------------------------------
------------------------------------------------------------------------
------------------------------------------------------------------------
------------------------------------------------------------------------
---------------------UAGUUAC--------------------------------------------
--------------------------------------------------GC--------------------
-------UAGUG--UGU------------------------------GUGAC--------------------
------------------------------------------------------------------------
------------------------------------------------------------------------
--------------------GUCGGUGU------------GAAAGUCGAC--G---ACUUUCCGACCCGUCU
UGAAACACGGAUUGCGGAGUGCU-U-GUCUACUGCGA-GUCAAAGG--G-UG--------UUAA-------A
A-CC--UUGC--GGCGAAAUGAAAG-UAA-----------------AGGUCAGUCUC---GAAUUGGCCGAC
GUGG-GAUCUGU----GUUCUUCG---------------------------------------GA--------
GUG---------------------------------------------------------------------
-------------------CAG--------------------------------------CGCACCACG-GC
CCUGUG----CG-UGUC----------------------ACUUG----------------UGACUGUG-CA-
GAG-GU----------------------UGAGCAGUUGGCAAAC-GACCCGAAAGA-UGGUGAACUAUGCCU
GAGCAGGAUGAAGCCAGA-GGAAAC-UCUGGUGGAAGUCCGU--AUCGGUUCUGACGUGCAAAUCGA-UCGA
UAGACUUGGGUAUAGGGGCGAAAGACUAAUCGAACCAUC-UAGUAGCUGGUUCCUUCCGAAG-UUUCCCUCA
GGAUAGCUGGAU-CUCA-------------------------------------------------------
----------------------GG-----CAGUUAUAUUCGGUAAAGCUAAUGAUU-AGA-GGCCUUGG-GG
A-C-GUAAU-GUCC-U-CAACCUAUUCUCAAACUUUCAAUGGAUAUGAA-----------------------
------------------GUUGC-AG-----UUU-CU--UUAG--U--GAA---------------------
CU---GUCA-A---CGUGA-AU-G--C-----G--AGGUCCA-AGUGGGC-C-AUUU-U-UGG-UAAGCAGA
AC-U-GGCGCUGU-GGGAUG---AACCAAACGUGG-A-GUUAAGGUGCCUAACU--UC-UCG-CU-CA---U
-G-AGACCCC-AUAAAAGGUGUUGGUUGAUAUUGACAGCAGGACGGUGGCCAUGGAAGUCGGUA-UCCGCUA
AGGAGUGUGUAACAACUCACCUGCC-----GAAUCAACUAGCCCUGAAAAU-GGAU-GGCGCUUA-AGCGAG
```

```
A----------GACCUAU-ACUCCGCCGUUGCGA--CAUGU-------------------------------
--------------------------------------------------------------------GC--
----GUUGU---------------------------------------------------------------
-----------------------CUAGCGCCAGGUCG-UAAC-GAGUAGGAAGGUCGUGGCGG---------
-----------U-U-GCGUU-GAAGGCUAUG-AGCGUAGGCU--C-GGCUGGAGCU-U-CCG-UCAGUGCAG
AUCG-UAAUGG-UAGUA-GCAAAU---AUUC-AAGUUC--G---------------AUCCUU-GAA-GACUG
AAGUGGAG-AAGGGUUCC-ACGUGAACA-GUAGUUGGAUGUGGGUCAGUCGA--UC-CUAAGGUACUGGCG-
A-A-CGCC--UU-GUAUCAU--------CGGU----GGCGA--AAAG---CUUG----------------CU
UU---------------UAGUCCC------CGCU----UG-UC-GA-A-AGGGA-AUAGGGUU-AAUA-UUC
CCUAA----------------------------------CUGAGAUGCAAAG--------A-UUGUG-----
U----------------UC-UUCG-----------------------------------GAGCACA------
--AGCG-CGG---UAAC--------GC-AU-----UC--------------------GAAC-U-UGGU-UA-
GUCGCUC-------------AAAGAC-CGAGCUAGA-GU------UUUCU-------------U-CUCUAG-
UUAAGG-AAC--G-----G-A-------------------CU-----CC----------------CUGG-AA
U-U-GG--------------------UUCAG-CCAGAGAUGG----------GGACGU----UGU-UU-CCG-
AAA--AG-----------------------------------------------------------------
----CACCGCG-GUUUCUGUGGUGU-C-UCGUGCU-CUUUGAACGGC-CCUU-AAA-A-CAC-CAAGGGAGG
-C---------------------------------------------UA-----UUAAUUUG-CA-CUCAAU
CGUACCGAUAUCCGCA------------------------------UUAGGUC--UCCAAGGU-GAACAGCC
UCUAGU-C-GA-UAGA-AUAAUGUA-G-GUAAGGGAAGUCGGC-AAACUAGAUCCGUAACUUCGG-GAAAAG
GAUUG-GCUC-C-AG--UGGUUG-G----AACGG-UU------GGCCAGUUGGU---UGAU-GCUUGUC---
-CGG-------------CGCA---GU---------------------------------------UCUG--U
C---------------------------------------------------------------------UG
-CUUG------A--------UACUU-----------------------------------------------
----------------------------------------------------------------------UC
GGGUUG----------AUG---------------------------------GC--GG--------------
---------------------------------------------------------------ACUA-----
------------------------------------------------------------------------
-G-UG---------------AU--U-GU---GCUU-GC----------------------------------
---UUG----CG-GA-CGC-----UUUCUGGU----GUG---------------------------------
----------------------------------------------------------------UGCU----
------------------------------------------------------UGG--ACCU---------
---------------C------------------------------GGUU---C------------------
------------------------------------------------------------------------
----UA-----------------------GU----AUCC----UGAUCGCUCAU-CUAAACAAC--CGUACU
GGA--ACCGGUACGGACUCAGGGAAUCCGA-----CUGUCUAAUUAAAACAGAGGUGACAGAUGGUCCU---
-UG-C-------------GGACGUUG-ACUGUCACUGA-UUUCUGCCCAGUGCUCUG-AAUG-UUAAAUCGU
A--------------GUAA--------------UUCGAGUAAGCGC--GGGUAAAC-GGCGGGAGUAACUAU
GACUCUCUUAAGGUAGCCAAAUGCCUCGUCAUU-UAAUUGUUGACGCGCAUGAAUGGAUUAACGAGA--UUC
CUACUGUCCCUAAC----UAC-UUUCUAGCGAAACC---AC---AGCCAAGGGAACGGGCUUGGCAAAAAUA
GCGGGGAAAGAAGA-CCCUGUUGAGC-UUGACU--CUAGUUU-GACAUUGUGAAGAGUCAUGAGAGGUGUAG
CAU-AGGUGGGAGUC-----------------------------------------------------UUCG
--------------------------------------------------G-A--C-GA-CAGUGAAAUA-C
CACCACUUUCAUCGACUCUUUACUUAUUC-GG-------U-UA-A-AAGAG------A-----------AU-
------UG--GC-UU--------CAC---GGCC-U--U--------------UUUUC--------------G
```

```
----------AAGCA--------------------------------------UUAA--GC---------GG
-----------------AG--------------------------------------------CC-------
---AUUU--------------------------U---AUG-----GCA------------------------
----------CCG------UGA------------------------------CUCU----CC----------
------------------------------------------------UCGA-----AGACA-GUGUCAAGC
GGGGAGUUUGACUGGGGCGGUACAUCUAUCA-AAUCGUAAC-GUAGGUGUCCU-AAGGCGAGCUCAGAGAGG
ACGGAAACCUCUCGUA-GAGCA-AAAGGGCAA-AAGCUUGCUUGAUC-UUGA--------------------
--------------CUUUCA--GUACGAGUACAGACCGCGAAA-GCGUGGCCUAUCGAUCCUUU--------
---UA-A-----UCCUGAUUG--------------------UU-UCAGGUAAGAGGUGUCAG-AA--AAGUU
ACCACAGGG-AUAACUGGCUUGUGGCAGCCAAGCGUCCAUAGCGACGUUGCUUUUUGAUCCUUCGAUGUCGG
CUCUUCCUAUCAUUGCGAAGCA-GAAUUCGCCAAGCGUUGGAUUGUUCACCCACUAAUAGGGAACGUGAGCU
GGGUUUAGACCGUCGUGAGACAGGUUAGUUUUACCCUACU--------------------G-UUG-------
-------------------------------AC--U-UGUUAUUGCGAAAGU-AAU-CCUGCUUAGUACGAG
AGGAA-CAGCGGGUUCAAACAUU-UGGU-UCAUAGACUUGAUC-GACA-GAUCAA-U-GGUCUGAAGCUACC
AUUUG-AGAGAUUAUAACUGAACGCC-UC--UAAGUUAGAAUCUCGCCUUG-----UC--AAGGCGAAAAUU
UCUUGCUU-CCCGGUGU-------------------------------------------------------
------------------------------------------------------------------------
------------------------------------------------------------------------
---------------------CGGGAGGCAUCU-----------------------CU--------------
-------------------------------AUCUCG------------UGGCAA-----------------
-------C--AC-------GA--GA-------------GCUU------------------AUGC--------
-----CCUAUGU--A---UGG-----CCU-UGGC-----GUCGUAGU--------GAAU---UCUGCGACGC
----------------------------U----U-GCCAACGCCA-----GAUCACU-CUG----------G
U-----------------UCAAUGU---------------------C-GGGGCGCU-----AAA------UC
ACUUGCAUACGACUUGGUCUCU-UGGUCAAGGU-GUUGUAUU-CAGUA--------GAGCAGU-CC------
----UUUUAUACUGCGAUCUGUUGAGACU-AU-CCUU-UGAUUGAG---UUUUUU--G--------------
------------------------------------------------------------------------
------------------------------------------------------------------------
------------------------------------------------------------------------
-------------------------------------------------
>LSUe_554453_PLAMI/1-3641 GU001169.1:379-530,927-4415 Planomonas micra 28S
------------------------------------------CUUUUAAC-AAUGGAUA-U-CUUGGU-UCU
C-G---CAACGAUGAAGAACGCAG--CGAAAUGCGAUACGUAAUGUGAAU--U---GCA--ACG--CC-AGU
GAAUCAUCAAAUCUUUGAACGCAUAU-U--------GC--------GC-----UUG-G-GGG---------C
A-U-----------UCC---C----C----C---AAG----C-A-CGUC-UGU------UUGAGUGUC-----
--------------------------------------------------------U--AUGGAC-CUCA-A
AUC--------------------AG-A-CG-GGGCUACCCGCUGAAUUUAAGCAUAUUAAUAAGCG--GAGG
AAAAGAAACUAA--CCAGGAUUCCCUA--GU-AA-CGGCGAGUGAAGCGGGA-A-GA-GCUCAAAUUUAAA
AUCU-UGGCGCUUC-------------------G---CGU-CAUGAAUUGUAGUU--UU-------------
--GA-GAUG-CUG-U-----G-------------------------UCUGU--GUA-----------CA---
A-------CCGGG--UCC-------AAGUUC-UUUG------------------------------------
-------GAA-CA-AGACA-------UCAGAGAGGGUGAGAAUCCCGUC---------UUU---GGC-CUGG
CGC---UG--GCGCU-GUACGAU-------------ACA---GUUU--CGAAGAGUCGCGUUGCUUG---GG
AAUGCAGCGCUAA-A-UGGGUGGUAAA-CUUCAUCUAAAGCUAAAUAUUGG--C-GAG-AGACCGAUAGC-G
AAC-A-AGUACCGUGAGGGAAAG-AUGCAA-AGAAC-UUUGA-AAAGAG-A-GUCAAAUAGUACUUGAAAUU
```

GUUAGAAGGA--A-AAG-CG-AUUG---AAAU-GAG-UG------------------------UAUCGCGCG
UACAAA--------CUUAUCGGUGUGGACCUGAAUCCGCGGCGUUCCGGAA--UC------GC---AAG---
------UUC----------------CGU--------------------GCGU--------------------
-------------------------------------------------------------CGUGU------
------------CAGGAGGGUC-UG-----------------------------------------------
----------------------------------GCC-----------------------------------
------GUGCA------CU---UUUG--U--G-----------CGUGCG-----------GG-UCAAGAUG-
---GA----------------------------------------UUGC-----------------GUACCC
AGG---------------AGAUGGU------------------GGCCGUGAAGGUAGGC--CCGGCGC----
--------------------------------------------------------------U--U-GC---
------------------------------------------------------------------------
------------------------------GUUGGGCUGUUAUAGCACGG---UGGCUGGACUGGGCGCGUG
AUUGAGGAA---------------------------------------------------------------
------------------------------------------------------------------------
------------------------------------------------------------------------
------------------------------------------------------------------------
--------------------------------------------------U---------------------
-----------------------GUAGG--------------------------------------------
------------------------------------------------------------------------
-----------------------------------------------AUGCGCGG----------------C
A----------------CUAGCCUUGCCGGCGUGGCCCUUUGUUUCGUAUCGACUGUGGACUGGAAACCAGU
GU----------------------------GCGGACGGUGCGGGGCUUACAGGUUGCGUGCGGGCAGAAGCG
CGUCU-----------GGAUCUUGACGA-----------------AAUGG--U---UUCAAUUGACCCGUCU
UGAAACACGGACCAAGGAGUCUA-A-CACGUAUGCGA-GUGUUUGG--G-U--------GGAAA--------
A-CCC-GGGA--CGCGUAAAGAAAU-UAAAG-GCCCCGUUGGGGGCAGGA---------------------G
GUAG-GAAUCC---------------------------------------------------GU--------
AA----------------------------------------------------------------------
-------------------GGA--------------------------------------GGCACUACC-GA
CCAACC----AU-GAUAU---------------------U--C----------------UUUGAAAGGUUU-
GAG-UA----------------------AGAGCAUAUUUGUUGG-GACCCGAAAGA-UGGUGAACUAUGCCU
GAAUAGGGUGAAGUCAGA-GGAAAC-UCUGAUGGAGGCUCGU--AGCGAUUCUGACGUGCAAAUCGA-UCGU
CAAAUUUGGGUAUAGGGGCGAAAGACUAAUCGAACCAUC-UAGUAGCUGGUUCCCUCCGAAG-UUUCCCUCA
GGAUAGCUGGAA-CUCG-------------------------------------------------------
-----------------------UAAGAAAAGUUUUGUCAGGUAAAGCGAAUGAUU-AGA-GGCCUUGG-GG
A-U-GAAAC-AUCC-U-CAACCUAUUCUCAAACUUUAAAUAGGCAAGAU-----------------------
------------------GCCGC-GG-----UUG-CU--UUAG--U--GAA---------------------
CC---GUAG-C---GAAGA-AU-U--U-----G--AGUUCUU-AGUGGGC-C-AUUU-U-UGG-UAAGCAGA
AC-U-GGCGAUGC-GGGAUG---AACCGAACGUUG-C-GUUAAGGUGCCCAACU--GC-GCG-CU-CA---U
-GUAGAUACC-ACAAAAGGUGUUGAUUCAUUCAGACAGCAGGACGGUGGCCAUGGAAGUCGGAA-UCCGCUA
AGGAGUGUGUAACAACCCACCUGCC-----GAAUGAAUUAGCCCUGAAAAU-GGAU-GGCGCUAA-AGCGCG
C----------GACCGAU-ACGCGACCAUUGGUG--CAAA--------------------------------
------------------------------------------------------------------------
------------------------------------------------------------------------
---------------------------UGCGAGGCAC-CAAU-GAGUAGGAGGGC-GUGGGGU---------
-----------U-U-GUGAC-GAAGCUUA-G-GCCGUAAGGC--C-GAGUCGAACG-G-ACC-CUAGUGCAG

```
AUCU-UGGUGA-UAGUA-GCAAAU---AUUC-AAGUGA--G---------------AACCUU-GAA-GACUG
AAGUGGAG-AAAGGUUCC-AUGUGAACA-GCAAUUGGACAUGGGUUAGUCGA--UC-CUAAGAGAUAGGGA-
A-A-UUCC-----GUAUGAA--------GGG-----CGC-----CAU---UAUU-----------------C
GG---------------CG----C------CGCC----UA-UC-GA-A-AGGGA-AUCGGGUU-AAUA-UUC
CCGAA----------------------------------CCAGAA---UGUG--------G-AU--------
A----------------UG-----------------------------------------------------
---GUA-UGG---UAAC--------AU-AA-----AU--------------------GAAC-U-CGGA-GAC
GCCUGCG------------CGGAGCC-CCGGGAAGA-GU------UCUCU-------------U-UUCUUU-
UUAACA-AC---U----UU-U-------------------UA-----CC----------------CCGG-AA
U-C-AG--------------------AUUGG-CUGGAGAAAG---------GGUUAC----AUG-GU-UGG-
GAA--AG-----------------------------------------------------------------
----CACUUCA-AUUCUCGAAGUGU-C-CGGUGCG-CUCUCGUCGGU-CCUUGAAA-A-UCC-GAGGGAAAG
-G---------------------------------------------AU-----AAUUUUCA-CG-UCUGGU
CGUACUCAUAACCGCA------------------------------UCAGGUC--UCCAAGGU-GAACAGCC
UCUAGU-U-GA-UAGA-ACAAUGUA-G-AUAAGGGAAGUCGGC-AAAAUAGAUCCGUAACUUCGG-GAAAAG
GAUUG-GCUC-U-UA--GGGUUG-G----GGUAC-GG------GGUC----CCU---------G--GCG---
-GAACGGCGUGG-ACUGACCG---AGGACUACUG-------------------GCUG---------CUC---
----------------------------------------------------------------------AC
GCGG-----------------CU-------------------------------------------------
------------------------------------------------------------------------
--GG--------------C---------------------------------GG--AC--------------
---------------------------------------------------------------UUUG-----
--------------------------------------------------G---------------------
-A------------------CG--G-AC---UGCGUUC----------------------------------
---UGC----------CCG-----CCUUGGAA----------------------------------------
-----------------------------------------------------------------GCG----
-------------------------------------------------------------CU---------
---------------U-------------------------------GCG----------------------
------------------------------------------------------------------------
----CA-----------------------UU------------CCUGUG-CCG--AUGAACAAC--CGACUA
AGA--ACUGGUACGGACAAGGGGAAUCCGA-----CUGUUUAAUUAAAACAAAGCAUUGCGAAGGCCGG---
-AA-A-------------UGGUGUUG-ACGCAAUGUGA-UUUCUGCCCAGUGCUCUG-AAUG-UCAAAGUGA
A--------------GAAA--------------UUCAACCAAGCGC--GGGUAAAC-GGCGGGAGUAACUAU
GACUCUCUUAAGGUAGCCAAAUGCCUCGUCAUC-UAAUUAGUGACGCGCAUGAAUGGAUUAACGAGA--UUC
CCACUGUCCCUAUC----UAC-UAUCUAGCGAAACC---AC---AGCCAAGGGAACGGGCUUGGAAGAAUCA
GCGGGGAAAGAAGA-CCCUGUUGAGC-UUGACU--CUAGUCC-GACUUUGUGAAAUGACAUAAGAGGUGUAG
AAU-AAGUGGAGC-------------------------------------------------------UUCG
----------------------------------------------------G--C-GC-CGGUGAAAUA-C
CACUACUUUUAUUGUCGUUUUACUUAUCC-AA-------U-UA-G-UUAGA------GA----------UG-
------GACCUC-----------------GCGGUC--UU-------------UUUCU---------------
-----------GGAC--------------------------------------UUAA--G-----------C
------------------------------------------------------------------------
---GUC---------------------------UUU-ACG--------------------------------
----------GCG------CGA------------------------------UCUA----CG----------
------------------------------------------------UUGG-----AGACA-CGGUCAGGU
```

```
GGGGAGUUUGGCUGGGGCGGCACAUCUGUUA-AAAGAUAAC-GCAGGUGUCCU-AAGAUGAGCUCAAUGAGA
ACAGAAAUCUCAUGUG-GAACA-AAAGGGUAA-AAGCUCAUUUGAUU-UUGA--------------------
--------------UUUUCA--GUAUGAAUACAAACUGUGAAA-GCAUGGCCUAUCGAUCCUUU--------
---AG-U-----UUUUAGAAG--------------------UU-UUAAGCUAGAGGUGUCAG-AA--AAGUU
ACCACAGGG-AUAACUGGCUUGUGGCAGCCAAGCGUUCAUAGCGACGUUGCUUUUGAUCCUUCGAUGUCGG
CUCUUCCUAUCAUUGCGAAGCA-GAAUUCGCCAAGUGUUGGAUUGUUCACCCACUAAUAGGGAACGUGAGCU
GGGUUUAGACCGUCGUGAGACAGGUUAGUUUUACCCUACU--------------------G-AUG-------
-------------------------------AA--G-UGUUGUUGCAAUAGU-AAU-CCAACUUAGUACGAG
AGGAA-CCGUUGGUUCAGUUAUU-UGGUCCU-UGCGGUUGGCU-GAAA-AGCCAG-U-GCCGCGAAGCUACC
AACUG-UAGGAUUAUGGCUGAACGCC-UC--UAAGUCAGAAUCCAUGCUAG-----AA-CGUGACGAU-GUU
UACCCCUC-CUGUAUUG-------------------------------------------------------
------------------------------------------------------------------------
------------------------------------------------------------------------
---------------------AGAC-----------------------------------------------
--------------------------------GAAUA-----------------------------------
----------CAA-----AUA--GA-----------------------------------GGCGGC--UCGU
CCGUCUUCG---AUA-----UCAUAGCACU--GGGA--ACGCGCGUGUG--CGGGGAAAA-GCCGUGCGCGU
GUG-------------------------U----C-CUUG-----------GAUGUUA-AUU----------U
C-----------------AAAUAUC---------------------G-GGAGGGAU-----AAA------UC
CUUUGUAGACGACUUAAA-UCA-AGAAC--------------------------------------------
------------------------------------------------------------------------
------------------------------------------------------------------------
------------------------------------------------------------------------
------------------------------------------------------------------------
--------------------------------------------------
>LSUe_2044563_ANTWI/1-3706 MG202010.1:3400-3576,3799-7327 Ancoracysta twist
AAC------------------------UAUA----UA--AAACUCUAAAC-AACGGAUA-U-CUUGGC-UCU
U-G---CAACGAUGAAGAACGCAG--CGAAAUGCGAUACUUAAUGUGAAU--U---GCA--GAA-CCU-UGU
GAAUCAUCAAAACUUUGAACGCAUAU-U--------GC--------AC-----AUC-C-AUU---------U
A-A-----------UCG---U----G----G---AUG----U-A-CAUC-UGU-----UUGAGGGUCA----
-GUAU------------------------------------AUCUUCUAC------A--CUUGAC-CUCA-A
AUC--------------------AG-G-UG-GGGCUACCCGCUGAAUUUAAGCAUAUAAUUAAGCG--GAGG
AAAAGAAACUAA--CAAGGAUUCCCCUA--GU-AA-CGGCGAGUGAAGCGGGA-A-GA-GCUCAAAAUGUAA
AUCU-CUGAGGUUC-------------------C---CUC-AGCGAAUUGUAGUC--UA-------------
--GA-GAAG-CAU-G-----U-------------------------UCUGU--GAC-----------GG---
C-------UUAGG--UAU-------AAAUCU-UUUG------------------------------------
-------GAA-CA-AGGUA-------UCAUAGAGGGUGAGAAUCCCGUC---------CAU---GAU-CUAA
GCU---UA--GUCAC-UUUUGAU-------------AUG---CUUU--CAAAGAGUCGGGUUGCUUG---GG
AGUGCAGCCUUAA-A-UGGGUGGUAAA-UUUCAUCUAAAGCUAAAUAUGGG--C-AAG-AGACCGAUAGC-G
AAC-A-AGUACCGUGAGGGAAAG-AUGCAA-AGAAC-UUUGA-AAAGAG-A-GUUAAAAAGUGCUUGAAAUU
GUUUAGAGGA--A-AAA-CG-AACG---GAUU-CAG-UG------------------------UUGCUUGUU
GUUCAAUCAGUUUCCUCAGCCGAAAUGAGCGU---------------GUAU--GU------GG---AGG---
------UCA----------------AAC--------------------CUGC--------------------
-------------------------------------------------------------UGCGCGUU---
----------UACUGGAGGUUCGAG-----------------------------------------------
```

```
----------------------------------GGG-----------------------------------
------AUGCA------CU---UGGG--C--AG----------UAGGCA-----------GG-UCAAGAUC-
---AG----------------------------------------UUUG-----------------CAUUCU
GAG---------------AGAAAAC------------------UUGGGGGAAGGUAGGUUGCCUGUUU----
--------------------------------------------------------------C--G-GC---
------------------------------------------------------------------------
------------------------------AGGGGAUUGUUAUAGCCCUC---GAGAGUUGUCAGAGUGUGG
ACUGAGG-U---------------------------------------------------------------
------------------------------------------------------------------------
------------------------------------------------------------------------
------------------------------------------------------------------------
--------------------------------------------------C---------------------
-----------------------ACGGU--------------------------------------------
------------------------------------------------------------------------
-----------------------------------------------GCGCGCCA----------------C
A----------------UGG-------CGAGGCAGCUGGUUCCUGCGUGGGUGUCACGACCUGGA-------
CU----------------------------AUAAAUAGUGCUGGAUGUGGCAUCUGCGUACAGAGCUGGUCA
CCUUCUGCGUUCC-CAUGAUCUUGACAA-----------------AAUGG--A---UCUGUCCGACCCGUCU
UGAAACACGGACCAAGGAGUCUA-A-CAUAUAUGCGA-GUAUUUGG--G-U--------GUCAA--------
A-CCUAUUGA--UGCGUAAUGAAAG-UAAA------------------AA---------------------G
GUGG-GAUCCC---------------------------------------------------GC--------
AA----------------------------------------------------------------------
-------------------GGG--------------------------------------UGCACCAUC-GA
CCGAUC----UG-GAUCU---------------------U-------------------UGUGAAAGAUUU-
GAG-UA----------------------AGAGCAUAUCUGUUGG-GACCCGAAAGA-UGGUGAACUAUGCCU
GAGUAGGGCGAAGCCAGA-GGAAAC-UCUGGUGGAGGCUCGU--AGCGGUACUGACGUGCAAAUCGU-UCGU
CAAACUUGGGUAUAGGGGCGAAAGACUAAUCGAACCAUC-UAGUAGCUGGUUCCCUCCGAAG-UUUCCCUCA
GGAUAGCUGGAA-CUCA-------------------------------------------------------
--------------------------GAUGAGUUUUAUCAGGUAAAGCGAAUGAUU-AGA-GGCCUUAG-GG
A-U-GAAAC-AUCC-U-UAACCUAUUCUCAAACUUUAAAUAGGUAAGAA-----------------------
------------------GCUUA-GG-----CUG-CU--UAAU--U--GAG---------------------
CU---UAGG-C---GUGGA-AU-C--A-----G--AGUUCUU-AGUGGGC-C-AUUU-U-UGG-UAAGCAGA
AC-U-GGCGAUGC-GGGAUG---AACCGAACGUCG-A-GUUAAGGUGCCUAACU--AC-GCG-CU-CA---U
--GAGAUCCC-ACAAAAGGUGUUGAUUCAUCGAGACAGCAGGACGGUGGCCAUGGAAGUCGGAA-UCCGCUA
AGGAGUGUGUAACAACUCACCUGCC-----GAAUGAAUCAGCCCUGAAAAA-GGAU-GGCGCUCA-AGCGUG
U----------GACCUAU-ACUCGACCAUUGGAG--UAAA--------------------------------
------------------------------------------------------------------------
------------------------------------------------------------------------
---------------------------UGCGAAGCUC-CAAU-GAGUAGGAGGGC-GUGAAGG---------
-----------U-U-GUGAA-GCAGCUCGAU-CUCGUGAGAG--G-GAGUGAAACA-G-CCU-UUAGUGCAG
AUCU-UGGUGG-UAGUA-GCAAAU---AUUC-AAAUGA--G---------------AACUUU-GAA-GACUG
AAGUGGAG-AAAGGUUCC-AUGUGAACA-GCAAUUGGACAUGGGUUAGUCGA--UC-CUAAGAGAUAGGGA-
A-A-CUCC-----GUUAUAA--------AGU-----GCG-----CGA---U-UU-----------------U
CG---------------CG----C------CGCC----UA-UC-GA-A-AGGGA-AUCGGGUU-AAUA-UUC
CCGAA----------------------------------CCUGAA---UACG--------G-AU--------
```

```
A----------------UU-----------------------------------------------------
---GUA-UGG---UAAC--------AU-AA-----AU--------------------GAAC-U-CGGA-GAC
GUCAGCA-------------GGAGCC-CUGGGAAGA-GU------UCUCU-------------U-UUCUUU-
UUAACU-GC---U----UA-A-------------------UA-----CC----------------CUGA-AA
U-C-GC--------------------AUUAA-GCGGAGAUAG---------GGUUAC----AUG-GC-AGG-
GAA--AG-----------------------------------------------------------------
----CACCUUA-AGUCUUGAGGUGU-C-CGGUGCG-CUCUUGAUGGC-CCUUGAAA-A-UCC-GAGGGAAAG
-U---------------------------------------------AU-----UAUUCUCG-UA-UCAGGU
CGUACUCAUAACCGCA------------------------------UCAGGUC--UCCAAGGU-GAACAGCC
UCUAGU-C-GA-UAGA-ACAAUGUA-G-AUAAGGGAAGUCGGC-AAAAUAGAUCCGUAACUUCGG-GAAAAG
GAUUG-GCUC-U-GA--GGGUUG-G----GUCUA-GG------GGUC----CUU---------G--UCU---
-AGAAGCUUGGG-AUAGUCGA---UGAGCUAUU---------------------CGG---------CCU---
----------------------------------------------------------------------UC
GCGG-----------------GCU------------------------------------------------
-----------------------------------------------------------------------G
--AA--------------C---------------------------------GG--AG--------------
---------------------------------------------------------------CGUU-----
--------------------------------------------------G---------------------
-A------------------CC--G-UC---GCCUCGU----------------------------------
---GGA----------CGG-----GCUUGGAA----------------------------------------
----------------------------------------------------------------CGAG----
-------------------------------------------------------------CU---------
---------------U-------------------------------GCU----------------------
------------------------------------------------------------------------
----CU-----------------------UU------------CCCUAG-GCA--AUGAACAAC--CAACUC
AGA--ACUGGUACGGACAAGGGGAAUCCGA-----CUGUUUAAUUAAAACAAAGCAUUGCGAUGGCCAC---
-AA-C-------------UGGUGUUA-ACGCAAUGUGA-UUUCUGCCCAGUGCUCUG-AAUG-UCAAAGUGA
A--------------GAAA--------------UUCAACCAAGCGC--GGGUAAAC-GGCGGGAGUAACUAU
GACUCUCUUAAGGUAGCCAAAUGCCUCGUCAUC-UAAUUAGUGACGCGCAUGAAUGGAUUAACGAGA--UUC
CCACUGUCCCUAUC----UAC-UAUCUAGCGAAACC---AC---AGCCAAGGGAACGGGCUUGGAAAAAUCA
GCGGGGAAAGAAGA-CCCUGUUGAGC-UUGACU--CUAGUCU-GACUUUGUGAAAUGACUUAGAAGGUGUAG
AAU-AAGUGGGAGC------------------------------------------------------UUCG
----------------------------------------------------G--C-GU-CGGUGAAAUA-C
CACUACUUUUAUUGUCGUUUUACUUAUAU-AC-------U-UA-A-AUAGA------AG----------CG-
------GCCUC--------------------GGCC--UU-------------UUUCU---------------
-----------AGAA--------------------------------------UUAA--C-----------G
------------------------------------------------------------------------
---UCC---------------------------UUU-ACG--------------------------------
----------GGC------GGA------------------------------UCUA----GG----------
------------------------------------------------GUAU-----AGACA-UUGUCAGGU
GGGGAGUUUGGCUGGGGCGGCACAUCUGUUA-AAUGAUAAC-GCAGGUGUCCU-AAGAUGAGCUCAAUGAGA
ACAGAAAUCUCAUGUG-GAACA-AAAGGGUAA-AAGCUCAUUUGAUU-UUGA--------------------
--------------UUUUCA--GUAUGAAUACAAACUGUGAAA-GCAUGGCCUAUCGAUCCUUU--------
---AG-U-----CUUUAGAAA--------------------UU-UUAAGCUAGAGGUGUCAG-AA--AAGUU
ACCACAGGG-AUAACUGGCUUGUGGCAGCCAAGCGUUCAUAGCGACGUUGCUUUUUGAUCCUUCGAUGUCGG
```

```
CUCUUCCUAUCAUUGUGAAGCA-GAAUUCACCAAGCGUUGGAUUGUUCACCCACUAACAGGGAACGUGAGCU
GGGUUUAGACCGUCGUGAGACAGGUUAGUUUUACCCUACU--------------------G-AUG-------
-------------------------------AA----CGUUGUCGCAAUAGU-AAU-ACAACUUAGUACGAG
AGGAA-CCGUUGUUUCAGAUAAU-UGGUAUU-UGCUCUUAGCU-GAAA-AGCUAA-U-GGGGCGAAGCUACC
AUCUG-UAGGAUUAUGACUGAACGCC-UC--UAAGUCAGAAUCCAUGCUAG-----AA-AGUGACGAU-AUC
UACCUCUC-CCAAUUUU-------------------------------------------------------
------------------------------------------------------------------------
------------------------------------------------------------------------
---------------------ACAU-----------------------------------------------
--------------------------------GAAUA-----------------------------------
----------ACA-----AUA--G------------------------------------------------
----------GCU--UCGGCC-----CA---AAA-----UCAUACUACG------GGAA-CUA-GCUAGUAG
CAGAACAAGC----UGCUGGUGGA----C----U-UCCG-----------GAACACA-AUU----------U
C-----------------AAAUAUU---------------------G-GGAGAGAU-----AAA------UC
CUUUGUAGACGACUUAAA-UAU-GGAACCAGGU-AUUGUAAG-UAGCA--------GAGUAGC-CU------
----UGU--UGCUACGAUCUACUGAGAUUAAG-CCUG-UGUUUCGA---AGAUUUGUU--------------
------------------------------------------------------------------------
------------------------------------------------------------------------
------------------------------------------------------------------------
---------------------------------CUUU------------
>LSUe_1333499_NAOCE/1-3550 MPCS01000111.1:589422-589583;589966-593354 Nanno
----------------------------------------AACUUUCAGC-AACGGAUG-U-CUUGGC-UCC
C-A---CAACGAUGAAGAACGCAG--CGAAAUGCGAUACGUAAUGCGAAU--U---GCA--GAA-UUC-CGC
GAGUCAUCAAACCUUUGAACGCACCU-U--------GC--------GC----UUUC-G-GGA---------U
A-U-----------GCC---C----G----U---UAG----C-A-UGUU-UGU-----UGGAGUGUCU----
-GUUA------------------------------------ACC------------U--UCGGAC-CUCC-A
AUC--------------------AA-G-CA-AGGCUACCCGCUGAAUUUAAGCAUAUAACUAAGCG--GAGG
AAAAGAAACUAA--CCAGGAUUCCCCUA--GU-AA-CGGCGAGUGAAGCGGGA-A-GA-GCUCAACUUGAAA
AUCU-GGGUCGUUUUAC-G-------------------GC-UCCGAAUUGUAGUC--UA-------------
--AA-GAAG-UGC-A-----G-------------------------CCAUC--GGC-----------CG---
A-------CGCGA--GAC-------AAGUCC-CUUG------------------------------------
-------GAA-AA-GGGCG-------CCCCAGAGGGUGACAGUCCCGUA---------CAU---G-C-UUGC
GUG---GG--UCCGU-CUACGGC-------------GUAU---UUU--CGACGAGUCGAGUUGCUUG---GG
AUUGCAGCUCAAA-G-UGGGUGGUAAA-UUCCAUCUAAGGCUAAAUAUCGG--U-GGG-AGACCGAUAGC-G
AAC-A-AGUACCGUGAGGGAAAG-AUGCAA-AGAAC-UUUGA-AAAGAG-A-GUUAAAGAGUGCCUGAAAUU
G-CUGAAAGG--G-AAG-GA-AAGG---AAAC-CAG-UG---------------------------------
-------------------------------------------UAAUGCUG--UG------GC---A-----
-------UU----------------UCC--------------------CCGC--------------------
-------------------------------------------------------------UGGACGU----
-----------------------UG-----------------------------------------------
----------------------------------GAC-----------------------------------
------UGCGG------UU---GUGC--C--AU----------GGCGUU-----------GG-UCAAGAUG-
---GG----------------------------------------UUCG-----------------ACUCGC
AGG---------------AUUAGGG------------------UGAGAGGGAGGUGGUU----CGCAC----
--------------------------------------------------------------U--U-CG---
```

```
------------------------------------------------------------------------
------------------------------GUGUGAAUGUUAUAUCCUUG---CACCUUUGCUGUGGGUUGG
ACCGAGGUC---------------------------------------------------------------
------------------------------------------------------------------------
------------------------------------------------------------------------
------------------------------------------------------------------------
--------------------------------------------------A---------------------
-----------------------ACCUU--------------------------------------------
------------------------------------------------------------------------
-----------------------------------------------GCGCCUCU----------------G
G----------------CUGAGCU--------------------------------------------CUGC
UC----------------------------AUGCAAGGUAC-------------------------------
----------------GGAUCUUGACGA-----------------AAUGG--U---UUUCUUUCACCCGUCU
UGAAACACGGACCAAGGAGUCUA-A-CAUAUAUGCGA-GUACAGGG--G-U--------GGCAA--------
A-CC--CCCG--UGCGCAAUGAAAG-UAAAGGGCUGCUUCUGCGGUCUGA---------------------G
GUGG-GAUCUG---------------------------------------------------UC--------
AA----------------------------------------------------------------------
-------------------AGG--------------------------------------UGCACCAUC-GA
CCGACC----AU-GAUUC---------------------UUCG----------------GAUGAAAGGUUU-
GAG-UG----------------------UGAGCAUUUAUGUUGG-GACCCGAAAGA-UGGUGAACUAUGCCU
GAGUAGGGCGAAGCCAGG-GGAAAC-UCUGGUGGAGGCUCGU--AGCGAUUGUGACGUGCAAAUCCA-UCGU
GGAAUUUGGGUAUAGGGGCGAAAGACUAAUCGAACCAUC-UAGUAGCUGGUUCCCUCCGAAA-UUUCCCUCA
GGAUAGCUGCAA-CCUA-------------------------------------------------------
-----------------------U---GCCAGUUUUAUCAGGUAAAGCGAAUGAUU-AGA-GGCCUCGG-GG
G-C-GAAAU-GCCC-U-CGACCUAUUCUCAAACUUUAAAUGGGUAAGAG-----------------------
------------------CCUGG-GG-----UUG-CU--UAAU--U--GAA---------------------
CC---CUAC-G---GUCGA-AU-G--A-----A--GGUUGCU-AGUGGGC-C-AACU-G-GGG-UAAGCUCG
GU-U-GGCGAUGC-GGGAUG---AUCCGAACGCUG-A-GUUAAAGUGCCAAACU--AC-CCG-CU-CA---C
-CUAGAUCCC-ACAAAAGGUGUUGGUUCAUUUAGACAGCAGGACGGUGGCCAUGGAAGUCGGAA-ACCGCUA
AGGAGUGUGUAACAACUCACCUGCC-----GAAUGAACUAGCCCUGAAAAU-GGAU-GGCGCUGA-AGCGAG
U----------GACUUAU-ACUCAGCCAUCAUGG--CAAG--------------------------------
------------------------------------------------------------------------
------------------------------------------------------------------------
---------------------------UGCGAGGCCG-UGAU-GAGUAGGAGGGC-GUGGAGU---------
-----------U-C-GUAGC-GCAUCCUG-U-GGCGUAAGCC--A-AGGUGAAACG-G-ACU-CUAGUGCAG
AUCU-UGGUGG-UAGUA-GCAAAU---AUUC-AACUGA--G---------------AACUUU-GAA-GGCCG
ACGUGGAG-AAAGGUUCC-AUGUGAACA-GCAAUUGGACAUGGGUUAGUCGA--UC-CUAAGGGAUAGGAG-
A-A-AUCC-----GUU--AG--------AAG-----GCG-----CGU---UCUU-----------------C
CG---------------AACG--C------CGUC----UA-CC-GA-A-AGGGA-AGAGGGUU-AAUA-UUC
CCUCA----------------------------------CCAGGA---CGUG--------G-AU--------
GU---------------UU-----------------------------------------------------
---GAG-UGG---CAAC--------AC-AA-----AC--------------------AAAA-C-CGGG-GAC
GUCGGCA-------------GGAGUU-CCGGGAAGA-GU------UCUCU-------------U-UUCUUC-
UUAACA-ACG-CG----AU-A-------------------GA-----CC----------------CUGG-AA
U-C-GG--------------------AUUGC-CCGGCGAUAG---------GGUGUA----UUG-GU-UGG-
```

```
GAA--AG-----------------------------------------------------------------
----CGGCGUA-AUUUUUGCGCCGU-C-CGGUGCG-CUCUUGACGGC-CCUUGAAA-A-CCC-GGUGGCGUG
------------------------------------------------------UAUUCGCA-CC-CCUGGU
CGUACUCAUAACCGCA------------------------------UCAGGUC--UCCAAGGU-GUGCAGCC
UCUGGU-C-GA-UAGA-AGAAUGUA-G-AUAAGGGAAGUCGGC-AAAAUAGAUCCGUAACCUCGG-GAAAAG
GAUUG-GCUC-U-GA--GGGUUG-G----GCCUA-GG------GGCU----UCU---------U--GGC---
-CAACGGAUGUG-GCUCGCUA---CA-----------------------------UGU---------GCC---
----------------------------------------------------------------------UG
ACGG-----------------UCU------------------------------------------------
-----------------------------------------------------------------------G
GCAA--------------C---------------------------------GG--AC--------------
---------------------------------------------------------------UAUU-----
---GG---------------------------GC-------GCGUGGUUG----------------------
-C------------------AG--G-CU---CUCUUGC----------------------------------
---GGC----------CUU-----GAGGAUUG----------------------------------------
----------------------------------------------------------------GUUU----
-------------------------------------------------------------CG---------
---------------G-------------------------------CCA----------------------
------------------------------------------------------------------------
----UC-----------------------UU------------CCCUAGUGCG--UGGAACAAC--CAGCUC
AGA--ACUGGAGCGGACAAGGGGAAUCCGA-----CUGUUUAAUUAAAACAAAGCAUUGCGAUGGUCGG---
-AG-A-------------CGAUGUUG-ACGCAAUGUGA-UUUCUGCCCAGUGCUCUG-AAUG-UCAAAGUGA
A--------------GGAA--------------UUCAACCAAGCGC--GGGUAAAC-GGCGGGAGUAACUAU
GACUCUCUUAAGGUAGCCAAAUGCCUCGUCAUC-UAAUUAGUGACGCGCAUGAAUGGAUUAAUGAGA--UUC
CCACUGUCCCUAUC----UGC-CAUCUAGCGAACCC---AC---AGCCAAGGGAACGGGCUUGGAAUAAUCA
GCGGGGAAAGAAGA-CCCUGUUGAGC-UUGACU--CUAGUCC-GACGUUGUGAAAUGACUUAUGAGGUGUAG
CAU-AAGUGGGAGC------------------------------------------------------UCCG
----------------------------------------------------G--C-GA-CGAUGAAAUA-C
CACUACUCGUAAUGUUGUUUUACUUAUUC-CG-------U-GG-C-GUAGA------GU----------CC-
------AGUCG--CU-------UCAC---GGCGCC--UG-------------GUUCU---------------
-----------AGAU--------------------------------------UUAA--AC---------GG
------------------------------------------------------------------------
---CCC---------------------------UCGAGGC--------------------------------
----------CCG------UGA------------------------------UCUA----CG----------
------------------------------------------------CGGG-----AGACA-ACGUCAGGU
GGGGAGUUUGGCUGGGGCGGCACAUCUGUUA-AACCAUAAC-ACAGGUGUCCU-AAGGUUAGCUCAAUGAGA
ACAGAAAUCUCAUGUA-GAACA-AAAGGGUAA-AAGCUAACUUGAUU-UUGA--------------------
--------------UUUUCA--GUACGAAUACAAACUGUGAAA-GCAUGGCCUAUCGAUCCUUU--------
----A-G-----UUUUUAGAA--------------------UU-UUAAGCUAGAGGUGUCAG-AA--AAGUU
ACCACAGGG-AUAACUGGCUUGUGGCAGCCAAGCGUUCAUAGCGACGUUGCUUUUUGAUUCUUCGAUGUCGG
CUCUUCCUAUCAUUGUGUCGCA-GAAGUCACCAAGUGUAGGAUUGUUCACCCACUAACAGGGAACGUGAGCU
GGGUUUAGACCGUCGUGAGACAGGUUAGUUUUACCCUACU--------------------G-AUG-------
-------------------------------AG--U-UGUUGUCGCGACAGU-AAU-ACAACUUAGUACGAG
AGGAA-CCGUUGUUUCAGAUAAU-UGGUAAA-UGCGGUUGGCU-GAAC-AGCCAG-U-GCCGCGAAGCUACC
AUCUG-UGGGAUUAUGACUGAACGCC-UC--UAAGUCAGAAUCCUUGCUGA-----AGAUGCGACGAU-AUU
```

```
CACCUCUU-CUGAUGCU-------------------------------------------------------
------------------------------------------------------------------------
------------------------------------------------------------------------
---------------------ACGC-----------------------------------------------
--------------------------------GUAUC-----------------------------------
----------GCG-----AUA--G------------------------------------------------
---------------------------------------GCUUCGGC--------CCAGC--AUCAUACG--
----------------------------U----C-----------------AUCAUC-AUU----------U
C-----------------AAUCAUU---------------------A-GAAGAGAU-----AAA------UC
CUCUGUAGACGACUUUGA-UAU-GGAACGGGGU-AUUGUAAG-UGUGA--------GAGUAGG-CU------
----UUG--UCCUACGAUCCACUGAGAUUCAG-CCUCUUGUUCCGU---CGAUUU-----------------
------------------------------------------------------------------------
------------------------------------------------------------------------
------------------------------------------------------------------------
-------------------------------------------------
>LSUe_1315715_PIJUD/1-3514 JX988758.1:2007-2187,2512-5844 Picomonas judraske
ACC------------------------CAAC----CA--AAACUCCUAAC-GAUGGAUA-U-CUUGGU-UCU
C-G---CAACGAUGAAGAACGCAG--CGAAAUGCGAUACGCAGUGCGAAU--U---GCA--GAA-UUC-AGU
GAAUCAUCAGAACUUUGAACGCAUAU-U--------GC--------GC-----UCU-G-AGG---------U
A-U-----------UCC---U----C----A---GAG----C-A-UGUU-UGU-----CUCAGUGGCA----
-UCUC------------------------------------CCCCUCUCCCCCUUAU--CUUGCC-CUGA-G
AUC--------------------AA-G-CA-AGGCUACCCGCUGAAUUUAAGCAUAUAACUAAGCG--GAGG
AAAAGAAACUAA--CAAGGAUUCCCCUA--GU-AA-GGGCGACUGAAGCGGGA-A-GA-GCUCAAGCCUAGA
AUCU-GCAUGUUUC-------------------G---CAU-GCCGAAUUGUAGUC--UA-------------
--UC-GAGU-UGUCG-----U-------------------------UCUCC--GGC-----------GG---
C-------GCAGG--UAU-------AAGUCU-UUUG------------------------------------
-------GAA-CA-AGGCA-------UCAUAGAGGGUGAGAAUCCCGUU---------UGU---GAC-UUGC
GUG---CG--UCCGG-UCUUGUA-------------UCGA-CAUCU--UGGCGAGUCAGGUUGUUCG---AG
AUUGCAGCCUAAA-A-UGGGUGGUAAA-UUUCAUCUAAAGCUAAAUAUAGG--C-GAG-AGACCGAUAGC-G
AAC-A-AGUACCGUGAGGGAAAG-AUGCAA-AGAAC-UUUGA-AAAGAG-A-GUUAAAAAGUGCUUGAAAUU
G-UUAGGAGG--G-AAG-CG-GAUC---GAAC-CAG-UG------------------------UUGC-----
------------------GCAGUAAGGACCGG---------------AGGC--CU------GG---CGC---
------UUC----------------CAU--------------------ACGU--------------------
-------------------------------------------------------------CUUGC------
------------------------------------------------------------------------
------------------------------------------------------------------------
-------------------------------------------UGCGCA-----------UG-CCAGCAUC-
---AG----------------------------------------U--------------------CGUUCG
GCA---------------GUAUAA------------------------------------------------
------------------------------------------------------------------------
------------------------------------------------------------------------
------------------------------AGCGGAUUCUGACCAGUUUC---CUGUGCUGCUGCUGAUAAG
ACUGAGGAG---------------------------------------------------------------
------------------------------------------------------------------------
------------------------------------------------------------------------
```

```
------------------------------------------------------------------------
--------------------------------------------------U---------------------
-----------------------UCGAC--------------------------------------------
------------------------------------------------------------------------
-----------------------------------------------GGGCGCUU----------------U
A----------------U----------GCGUUACUGUGUCUUUCAGUGCUCACCAUUUCUGGGA-------
CU----------------------------ACAUGUAGUGCCGGUUUG----UCUGGGCGCUCUGGCUGCAG
UUUUCUGCCCGUC-GGCGAUGUUGGCAA-----------------AAUGC--U---UCGCUCCGGCCCGUCU
UGAAACACGGACCAAGGAGUCUA-A-CAUGUGUGCGA-GUAUUGUG--G-U--------GGCAA--------
A-CC--AUGG--UGCGCAAUGAAAG-UAAA------------------AG---------------------G
GU----------------------------------------------------------------------
------------------------------------------------------------------------
-------------------GGG--------------------------------------UGCACCGCC-GA
CCGACC----AU-GAUCU---------------------U--C----------------UGUGAAAGGUUU-
GAG-UA----------------------AGAGCAUGCCUGUUGG-GACCCGAAAGA-UGGUGAACUAUGCCU
GAGUAGGGUGAAGCCAGA-GGAAAC-UCUGGUGGAGGCUCGU--AGCGAUACUGACGUGCAAAUCGU-UCGU
CAAACUUGGGUAUAGGGGCGAAAGACUAAUCGAACUAUC-UAGUAGCUGGUUCCCUCCGAAG-UUUCCCUCA
GGAUAGCUGGAG-CUCA-------------------------------------------------------
---------------------------AUGAGUUUUAUCAGGUAAAGCGAAUGAUU-AGA-GGCAUUGG-GG
U-U-GUAAC-AACC-U-CGACCUAUUCUCAAACUUUAAAUGGGUAAGAU-----------------------
------------------GCCUU-CG-----CUU-CC--UAAC--C--GAG---------------------
CG---UGGG-C---GUGCA-AU-C--G-----G--AGCUCUU-AGUGGGC-C-AUUU-U-UGG-UAAGCAGA
AC-U-GGCGAUGC-GGGAUG---AACCGAAAGCCG-A-GUUAAAGUGCCGGACU--GC-ACG-CU-CA---U
UCCAGAUACC-ACAAAAGGUGUUGAUUCAUACUGACAGCAGGACGGUGGUCAUGGAAGUCGAAA-UCCGCUA
AGGAGUGUGUAACAACUCACCUGCC-----GAAUGAAUUAGCCCUGAAAAU-GGAU-GGCGCUCA-AGCGUG
C----------GACUCAU-ACUCGGCCAUCAGGG--CAAA--------------------------------
------------------------------------------------------------------------
------------------------------------------------------------------------
---------------------------UGCGAUGCCC-UGAU-GAGUAGGAGGGC-GUGGGGC---------
-----------U-C-GUGAA-GCAGCCCG-C-GGCGUGAGCC--G-GGGUGAAACG-U-GCU-CUAGUGCAG
AUCU-UGGUGG-UAGUA-GCAAAU---AUUC-AAAUGA--G---------------AACUUU-GAA-GACUG
AAGUGGAG-AAAGGUUCC-AUGUGAACA-GCACUUGGACAUGGGUUAGUCGG--UC-CUAAGCGAUAGGGA-
A-A-CUCC-----GUUUUAA--------AGA-----CGC-----CUU---U-UU-----------------U
GG---------------CG----U------CAUA----GC-GC-GA-A-AGGGA-AUCGGGUU-AAUA-UUC
CCGAA----------------------------------CGGGGA---UGUG--------G-GU--------
A----------------AU-----------------------------------------------------
---GUG-UGG---UAAC--------AC-AA-----CA--------------------GAAC-G-CGGA-GAC
GUCGGCG-------------GGAGCC-CUGGGAAGA-GU------UCUCU-------------U-UUCUUU-
UUAACU-GC---C----UC-U-------------------UA-----CC----------------CUGG-AA
U-C-AG--------------------AUUAC-CUGGAGAUAG---------GGUUAC----ACG-GC-AGG-
GAA--AG-----------------------------------------------------------------
----CACCUUA-CGUCUUGAGGUGU-C-CGGUGCG-CUCUCGACGGC-CCUUGAAA-A-UCC-GCGGGACAG
-G---------------------------------------------AU-----UAUCAUCA-CG-CCCCGC
CGUACUCAUAACCGCA------------------------------UCAGGUC--UCCAAGGU-GAACAGCC
UCUAGU-C-GA-UAGA-ACAAUGUA-G-GUAAGGAAGUCGGC-AAAAUAGAUCCGUAACUUCGG-GAAAAG
```

```
GAUUG-GCUC-U-AA--GGGUUG-G----GUCUA-GG------GGUC----UGC---------G--GCA---
-AGAAGCCGGAG-GCUGUGUG---CGGACUAGCG-------------------GCGG---------CCU---
----------------------------------------------------------------------UC
ACGG-----------------GCU------------------------------------------------
-----------------------------------------------------------------------G
CUGU--------------C---------------------------------GG--AC--------------
---------------------------------------------------------------CGCG-----
--------------------------------------------------U---------------------
-A------------------CG--G-CC---GAAACGC----------------------------------
---GGA----------CGG-----CCGCAGAA----------------------------------------
----------------------------------------------------------------CGCU----
-------------------------------------------------------------UC---------
---------------A-------------------------------CGG----------------------
------------------------------------------------------------------------
----CU-----------------------UU------------CCCUAG-GCA--AUGAACAAC--CGACUU
AGA--ACUGGUACGGACAAGGGGAAUCCGA-----CUGUUUAAUUAAAACAAAGCAUUGCGAUGGCCGC---
-AA-C-------------CGGUGUUG-ACGCAAUGUGA-UUUCUGCCCAGUGCUCUG-AAUG-UCAAAGUGA
A--------------GAAA--------------UUCAACCAAGCGC--GGGUAAAC-GGCGGGAGUAACUAU
GACUCUCUUAAGGUAGCCAAAUGCCUCGUCAUC-UAAUUAGUGACGCGCAUGAAUGGAUUAACGAGA--UUC
CCACUGUCCCUAUC----UAC-UAUCUAGCGAAACC---AC---AGCCAAGGGAACGGGCUUGGAAUAAUCA
GCGGGGAAAGAAGA-CCCUGUUGAGC-UUGACU--CUAGUCU-GACUUUGUGAAAUGAGCUUCGGGGUGUAG
CAU-AAGUGGGAGC------------------------------------------------------UCCG
----------------------------------------------------G--C-GC-CAAUGAAAUA-C
CACUACUCGUUGUCUCGUUUUACUUAUUC-CA-------U-GA-U-GUAGA------AG----------CG-
------GUCUC-------------------UGACC--UC-------------CUUCU---------------
-----------AGCA--------------------------------------UUAA---------------
------------------------------------------------------------------------
-------------------------------------GCA--------------------------------
----------CUG------CGA------------------------------UCUA----AG----------
------------------------------------------------UGGA-----AGACA-UUGUCAGGU
GGGGAGUUUGGCUGGGGCGGCACAUCUGUUA-AACAAUAAC-GCAGGUGUCCU-AAGAUGAGCUCAAUGAGA
ACAGAAAUCUCAUGUG-GAACA-AAAGGGUAA-AAGCUCAUUUGAUU-UUGA--------------------
--------------UUUUCA--GUACGAAUACAAACUGCGAAA-GCAUGGCCUAUCGAUCCUUU--------
---AG-C-----CUUUAGAAA--------------------UU-UUAAGCUAGAGGUGUCAG-AA--AAGUU
ACCACAGGG-AUAACUGGCUUGUGGCAGCCAAGCGUUCAUAGCGACGUUGCUUUUUGAUCCUUCGAUGUCGG
CUCUUCCUAUCAUUGUGACGCA-GAAGUCACCAAGUGUCGGAUUGUUCACCCGCCAAUAGGGAACGUGAGCU
GGGUUUAGACCGUCGUGAGACAGGUUAGUUUUAUCCUACU--------------------G-AUG-------
-------------------------------AA--G-UGUUAUCGCAAUAGU-AAU-UCAACUUAGUACGAG
AGGAA-CCGUUGAUUCACAUACU-UGGUAUU-UUCACUUAGCU-GAAC-AGCUAA-U-GGUGAGAAGCUAUC
AUGUG-UAGGAUUACGGCUGAACGCC-UC--UAAGCCGGAAUCCAUGCUAG-----AU-CGCGAUGAU-UAU
UACCGCUC-UCUCUAUC-------------------------------------------------------
------------------------------------------------------------------------
------------------------------------------------------------------------
---------------------ACAU-----------------------------------------------
--------------------------------GCGCA-----------------------------------
```

```
----------ACA-----AUA--G------------------------------------------------
---------------------------------------GCUUCGGC--------CCAAC--GCCAUACU--
---------------------------------------------------GAUUUC-AAU----------C
G-----------------AAAUAUU---------------------G-AGAGCGAU-----AAA------UC
CUCUGCAGACGACUU-AG-CAG-GGAACAGAGU-ACUGUAAG-GAGUA--------GAGUAGC-CU------
----UGU--UGCUA----------------------------------------------------------
------------------------------------------------------------------------
------------------------------------------------------------------------
------------------------------------------------------------------------
-------------------------------------------------
>LSUe_35144_PRYPA/1-3366 AF289038.1:392-553,816-4019 Prymnesium patelliferu
------------------------------A----CA--CAACUCUUGUC-GAUGGAUA-U-CUUGGC-UCU
C-G---CAACGAUGAAGAACGCAG--CGAAAUGCGAUACGUAAUGCGAAU--U---GCA--GAA-UUC-AGU
GAAUCAUCGAACUUUUGAACGCAACU-G--------GC--------GC----UUCC-A-GGU---------U
C-C-----------GCC---U----G----G---GAG----C-A-UGUU-UCU-----UCGAGUGGCG-C--
--------------------------------------------------------CUAUCCGCC-AUCG-A
AUG--------------------AA-G-CA-AGACUACCCGCUGAACUUAAGCAUAUCACUAAGCG--GAGG
AGAAGAAACUAA--CCAGGAUGCCCCUA--GU-AA-UGGCGAAUGAAGCGGGC-A-UA-GCUCACGCCGCCA
AUCU-GGAGGUC------UC-----UACGACGAC---CCC--CCGAAUUGUAGUC--UA-------------
--GA-AAGG-CGC-------C-------------------------AUCGG--CCG-----------CG---
UC------CGCCG--CUC-------AAAUCC-CCUG------------------------------------
-------GAA-CG-GGGUA-------UCGGAGAAGGUGAGAAUCCUGUG---------CCG--------UGG
CGG---UU--CGCGG-UCGCACG-------------AUGC-GCUCU--UGCAGAGUCGGGUUGCUUG---GG
AUUGCAGCUUGAA-U-UGGGUGGUAAA-UUCCAUCUAAAGCUAAAUACUGG--C-GAG-AGACCGAUAGC-G
AAC-A-AGUACCGUGAGGGAAAG-AUGCAA-AGAAC-UUUGA-AAAGAG-A-GUUAAAGAGUGCUUGAAAUC
G-GCAAGAGG--A-AAC-CC-GUUU---GCGC-CCG-GUG--------------------------------
------------------------------------------------------------------------
------------------------------------------------------------------------
------------------------------------------------------------------------
------------------------------------------------------------------------
----------------------------------UCG--------UCUCU-----------------UCGGC
U-----CAA-------------CCAU--G--CCUCUUGGUGUGGU---------------GCACUUCGGAG-
---AG----------------------------------AUGGGUCAGCUCGCGCUGCGUGGGCGUCACACA
CAU-----------------CCAGGU----------------------------------------------
------------------------------------------------------------------------
------------------------------------------------------------------------
------------------------------------------------------------------------
------------------------------------------------------------------------
------------------------------------------------------------------------
------------------------------------------------------------------------
------------------------------------------------------------------------
-------------------------------------------------AGGGCU-----------------
------C------------CGGUGAGC---------CUGGUUG-----------------------------
----------------------------------------GAU---G-GCGCG------CG-----------
-----CUCGG--------------------------------------------------------------
```

```
------------------------------------------------------------------------
------------------------------------------------------------------------
-------------------CGUCGGCUG----------------ACGGAA--CGGGCGCAAUCGACCCGUCU
AGAAACACGGACCAAGGAGUCUG-A-CACGUGUGCGA-GUACCCGG--G-UG-------GCAAG-------C
C-C----GCG--UGCGCAAUGAAAG-UGA-----------------AG-----------------------G
GCG---------------------------------------------------------------------
------------------------------------------------------------------------
------------------------------------------------------------UUCGGCGCC-AA
CCGACC----UU-GAUC----------------------UUCU-----------------GUGAAGGGUUU-
GAG-U-----------------------GGAGCACACUUGUCGG-GACCCGAAAGA-UGGUGAACUAUGCCU
GAGGAGGGUGAAGCCAGA-GGAAAC-UCUGGUGGAGGCUCGU--AGCGAUACUGACGUGCAAAUCGU-UCGU
CGAACUUGGGUAUAGGGGCGAAAGACUAAUCGAACCAUC-UAGUAGCUGGUUCCCUCCGAAG-UUUCCCUCA
GGAUAGCUUGAA-CG---------------------------------------------------------
--------------------CCGA-----CGGUUUUGUCAGGUAAAGCGAAUGAUU-AGA-GGCCUUGG-GG
C-U-GAAAC-AGCC-U-UAACCUAUUCUCAAACUUUAAAUGGGCAAGAG-----------------------
------------------GCGUG-GG-----AUG-CU--UGAG--U--GAU---------------------
CC---CACG-C---GGCGA-AC-G--GC----G--CGUUCAA-AGUGGGC-C-AUUU-U-UGG-UAAGCAGA
AC-U-GGCGAUGC-GGGAUG---AACCGAACGCCG-C-GCUAAGGUACCCAAGU--GU-CCG-CU-CA---U
-C-AGAGACC-ACAAAAGGUGUUGGUACCUCUAGACAGCAGGACGGUGGCCAUGGAAGUCGGAA-UCCGCUA
AGGAGUGUGUAACAACUCACCUGCC-----GAA-GUACUAGCCCUGAAAU-GGAU-GGCGCUCA-AGCGGA
C----------CACCGAU-GCGCGGCCGCCGGCG--CGAG--------------------------------
------------------------------------------------------------------------
------------------------------------------------------------------------
---------------------------CGCGAGGCGC-CGGC-GAGUAGGGGGGC-GCGGGGC---------
-----------U-C-GCGAC-GCAGCCUU-G-GGCGUGAGCC--U-GGGUGAAGCG-G-GCU-CUGGUGCAG
AUCU-UGGUGG-UAGUA-GCAAAU---AUUC-AAAUGA--G---------------AACUUU-GAA-GACUG
AAGUGGAG-AAAGGUUCC-GUGUGAACA-GCAGUUGGACACGGGUGAGCCGA--CC-CUAAGAGACAGGAG-
A-G-CUCC-----GCCGAAC--------G-------UGG-----CCG---CUUCU-----------------
----------------GUCGGCCC-------GUC----CG-UC-GA-A-AGGGA-AUCGGGUU-AAUA-UUC
CCGAG----------------------------------CCAGUG---CGUG--------GAUG--------
G----------------UG-----------------------------------------------------
----GG-CGG---CGAC--------GC-GA-----AC--------------------GAAC-C-CGGA-GAC
GCCGGCG-------------GGAGCC-CUGGGAAGA-GU-------UCUCU-------------U-UUCUUC-
UUAACU-GCC--U-------C------------------UUG-----CC----------------CUGG-AA
U-C-GG--------------------UUCGG-CCGGAGAUAG----------GGCUAU----ACG-GC-AGG-
CAA--AG-----------------------------------------------------------------
----CACCUUA-AGUCUUGAGGUGU-C-CGGCGCG-CUCCCGACGGC-CCGUGAAA-A-UCC-GGGGGAGCG
-G---------------------------------------------UU-----CGUUCUCA-CG-CCUGGU
CGUACUCAUAACCGCA------------------------------UCAGGUC--UCCAAGGU-GAACAGCC
UCUAGU-C-GA-UAGA-GCAAUGUA-G-AUAAGGGAAGUCGGC-AAAAUGGUUCCGCAACUUCGG-GAGAAG
GAAUG-GCUC-U-GA--AGGCUG-G----GCGUA-GG------GGUCCU--AGG-UCCGAA-GU--CCA---
-AGG------C------UGCG---CG----------------------------------------CGG--A
C---------------------------------------------------------------------UA
------------------------------------------------------------------------
------------------------------------------------------------------------
```

```
--CU--------------G---------------------------------GCAACG--------------
---------------------------------------------------------------GCGGA----
------------------------------------------------------------------------
-C-UG---------------CG--C-GCG--GCCG-AC----------------------------------
---CGA----CG-GA-UGGC----CU-AGGAGC---------------------------------------
----------------------------------------------------------------GCUC----
-----------------------------------------------------------UC-----------
------------------------------------------------GG---G------------------
------------------------------------------------------------------------
----CU-----------------------UU------------CCCUGC-GCG--AUGAACAGC--CACUUC
AGA--ACUGGCGCGGACAAGGGGAAUCCGA-----CUGUUUAAUUAAAACAAAGCAUUGCGAGGGCCGG---
-GG-A-------------CGGUGCUG-ACGCAAUGUGA-UUUCUGCCCAGUGCUCUG-AAUG-UCAAAGUGA
A--------------GAAA--------------UUCAACCAAGCGC--GGGUAAAC-GGCGGGAGUAACUAU
GACUCUCUUAAGGUAGCCAAAUGCCUCGUCAUC-CAAUUAGUGACGCGCAUGAAUGGAUUCGUGAGA--UUC
CCACUGUCCCUAUC----UAC-UGUCUAGCGAAACC---AC---AGCCAAGGGAACGGGCUUGGAAGAAUCA
GCGGGGAAAGAAGA-CCCUGUUGAGC-UUGACU--CUAGUCU-GUUGC-GCAGGACGACUUGAGAGGUGUAG
AAU-AAGUGGGAGGA-----------------------------------------------------GCAA
--------------------------------------------------U-C--C-GC-CGGUGAAAUA-C
CACUACUUUCAACGUCGUCCUACUUACCC-GG-------U-GA-A-GCGGA---------------------
------GU--GC-AC--------CUC---GGUG-C--U--------------CUUCU---------------
-----------GGAC--------------------------------------UUAA--CC---------C-
------------------------------------------------------------------------
---------------------------------------C-----UC-------------------------
----------GGG------GGA------------------------------UCCG----CG----------
------------------------------------------------UCGG-----GGACA-CACACAGGU
GGGGAGUUUGGCUGGGGCGGCACGUCUGUUA-AAAAAUAAC-GCAGGCGCCCG-AUUUUGAGCUCAAUGAGA
ACAGAAAUCUCAUGUG-GACCG-AAAGGGGAA-AAGCUCGGUUGAUU-UUGA--------------------
--------------UUUUCA--GUACGAAUACAAACCGCGAAA-GCGUGGCCUAUCGAUCCUUU--------
---AG-U-----CUUUGGAA---------------------UU-UCAAGCUAGAGGUGUCAG-AA--AAGUU
ACCACAGGG-AUAACUGGCUUGUGGCUGCCAAGCGUUCAUAGCGACGUAGCUUUUUGAUCCUUCGAUGUCGG
CUCUUCCUAUCACUGCCUCGCA-GCGGGGGCCAAAUGCAGGAUUGUUCACCCACCAAUAGGGAACGUGAGCU
GGGUUUAGACCGUCGUGAGACAGGUUAGUUUUACCCUACU--------------------G-CUG-------
--------------------------------C----UCGUUUUUCAACGGU-AAU-UUGGUUCAGUACGAG
AGGAA-CCACCAAUUCGCACCAU-CGGU-UCACCCGC-CGCCC-GAUA-GGGCGA-U-GCGGGGUCGCUGCC
AUGCG-CCGGAUUACAACUGAACGCC-UC--UAAGUUGGAAUCCAUGCCGG-----AA--GAGAAGC----G
GUCCACCG-CGCGG----------------------------------------------------------
------------------------------------------------------------------------
------------------------------------------------------------------------
------------------------------------------------------------------------
--------------------------------GGG----------A--------------------------
----------AG-------CC--ACG---------AGCUGCC------------------GU---AU---C-
---------------------------------------GC-ACGAGCC------GUGA-----GCG-ACCA
----------------------------C-------AA-------------UUUCCU-GAC----------C
C-----------------CGCGCGA---------------------U--------------GAA------UC
CCUUGCAGACGACUUGAA-UAA-GGAACGGGGU-AUUGUACG-CGGCA--------GAGUGGC-CU------
```

```
----UGU--UGCCA----------------------------------------------------------
------------------------------------------------------------------------
------------------------------------------------------------------------
------------------------------------------------------------------------
-------------------------------------------------
>LSUe_55529_GUITH/1-3866
-----------------------------UU----CC--UAGCCCCCAAU-GGUGGAUA-U-CUUGGC-UCC
U-G---UAUCGAUGACGAACGCAG--CAGCGCGCGAUAGGUGUUGUGAUC--C---GCA--AAA-CUAUAUU
UGAUCAUCGAGUCUUCGAACGCAAAC-G--------GC--------AC----GCUC-C-UCAC-------CA
U-C---------CG-GU---G----G-AGUA---GCG----U-G-CGCC-CGU-----CUGAGCAUCG-UUU
--------------------------------------------------------A--CACGGU-CUCA-G
GUC--------------------GG-G-CG-GGAAUACCCGUCGAACUUAAGCAUAUCAAUAGACG--GAGG
AAAAGAAACUAA--CCAGGAUUCCCUUA--GU-AG-CGGCGAGCGAAUCGGGA-U-CA-GCUCAAGAUUUUA
AUCG-GCAGUGGUUCCU-AUG-------------AUC-CACCGCUGAGAUGUAGUC--CU-------------
--AA-GUGG-CGC-A-----C-------------------------UCGGG--AGA-----------AG---
U-------GGUAU--AUU-------AAGUCU-UUUG------------------------------------
-------GAA-CA-AAGCG-------CCAUAGAGGGUGAUAGCCCCGUC---------UGU---G---UAUA
CCA---AG--UCGUC-UGAGCCUCUCCCUGCCG---AUGC-GCCAC--UCGUGAGUCGGGUUGCUUG---GG
AGUGCAGCCUAAA-U-AGGGUGGUAUA-CUUCAUCCAAAGCUAAAUAUGGA--C-AGG-AGACCGAUAGC-G
AAC-A-AGUACCGCGAGGGAAAG-AUGAAA-AGAAC-UUUGG-AAAGAG-A-GUGAAAUAGUGCUUGAAACC
A-UUGGGGGG--A-AAG-CG-AAUG---GAGC-UUC-CG-----------------------UA--------
-------------------------------------------AUAUCGAA--GG---AUCUC--G------
-------------------------AGC--------------------CAGC--------------------
---------------------------------------------------------------GCUUU----
-----------------------GG-----------------------------------------------
----------------------------------UUG--------------------------------GAU
C-----UGGUGUAAUGACU---UCGA--U--G----------------------------AUGCGUCCGUG-
---AGUUUUUU-----CCCU-------------------UCUCGUGCGU-----------------GCGUUG
AAC-----------------ACAACGAAA--------GAA-UUCACUGGCUU--------------------
--------------------------------------------------------------U--C-CG---
------------------------------------------------------------------------
-------UGUGAGGCAGAAGCUUUGAGCUGGAGUCUUUC------CUUUUUGUUGUUGUUGGAAUCGACACA
GAAGGGCUCUUC------------------------------------------------------------
------------------------------------------------------------------------
------------------------------------------------------------------------
-----------------------------------GU-----------------GA----------------
-------------------------------------------------GAGU-------------------
--------------------UUCGAUUU-------UUGUUU-------------------------------
------------------------------------------C---U-CCACCUCUCCCUUUUUUUU-----
-----CCAAAGGUUGGU-GGA---GGU--------CCAA--------AGAUU--------------------
------------------------------------------------------------------------
------------------------------------------------------------------------
-----------------------GGCAA-----------UGGACGUAGGG--G---CUCCUUCGACCCGUCU
UGAAACACGGACCAAGGAGUCCG-G-UAUGAGCGCGA-GUGCUCGG--G-AA-------AGUUA-------C
A-CC--CAUG--CGCGUAGCGAAAG-CAA-----------------UGACUGGAAUCUGUUCAUCGAGAACG
```

```
CUGAUAGACAUU----UCCCUCGUUACGAGAGAUCUCUA---GAGUCUUUCAAUCUUCUCCUUG--------
AAU---------------------------------------------------------------------
-------------------AGGUAGGUUCUUAAGGUGAGAAAUUUGGCUAAGCUCCAUAUUGCAUCAUC-GU
CCAACC----UA-UCUG-CC-------------------CUUAG--------------GGUCGAUUGGCUU-
GUG-AU----------------------UGAGCGCUCAUACUGG-GACCCGAAAGA-UGGUGAACUAUGCCU
GAGCAGGAUGAAGCCAGA-GGAAAC-UCUGGUGGAGGUCCGA--AGGCGUUCUGACGUGCAAAUCGA-UGCU
CAGACUUGGGUAUAGGGGCGAAAGACCAAUCGAACCGUC-UAGUAGCUGGUUCCCUCCGAAG-UUUCCCUCA
GGAUAGCUGGAG-CUGU-------------------------------------------------------
----------------------AG-----CAGUCUUAUCCGGUAAAGCGAAUGAUU-AGA-GGAAUCGG-AG
A-GUCC-AAACUCU-U-UGACCUAUUCUCAAACUUUAAACAGGUAAGGA-----------------------
------------------AGUUG-UG-----UUG-CU--UAGA--U--GAA---------------------
CA---CAGC-U---GGACG-AU-G--GU----U--AGCUCUU-AGUGGGC-C-AUUU-U-UGG-UAAGCAGA
AC-U-GGCGAUGC-GGGAUG---AACCGAGAGUGG-U-GUUAAGGCGCCCAACU--AC-GCG-CU-CA---U
-U-AGAUCCC-AGGAAAGGUGUAGGUGCGUAUGGACAGCAGGACGGUGGUCAUGGAGGUCGACA-UCCGCUA
AGGAGUGUGUAACAACUCACCUGCC-----GAACGCAUCAGCCCUGAAAAU-GGAU-GGCGCUCA-AGCGCG
U----------GGCCGAU-ACACCACCGCCGGCG--UACA--------------------------------
------------------------------------------------------------------------
------------------------------------------------------------------------
---------------------------GAAUUUGCUC-CGGU-GUGUAGGAGGGC-GUCGUGC---------
----------CU-UUGCGUC-GAAGCCUU-U-GGCGUGAGCC--U-UGGUGGAGCA-G-GCU-CGAGUGCAG
AUCU-UGGUGG-UAGUA-GCAAAA---AUUC-AAGCGA--G---------------AAUCUU-GAG-GACUG
GAGUGGAG-AAGGUUUCC-AUGCAAACU-GUGUUUGAGCAUGGGUGAGUCGA--UC-CUAAGAGAAUGGGU-
A-A-UUCC-----GUUAGAG--------UU------GUG-----CUU-------------------------
-----------------CGGGCAU------GUUC----CA-UC-GA-A-AGGGA-AACGGGUU-AAUA-UUC
CCGUA----------------------------------CCAUGA---UGCG--------G-AG--------
U----------------UU-----------------------------------------------------
---AAG-CGG---CAAC--------GC-AA-----AC--------------------GAUC-U-UGGG-GAC
AUCGGUG-------------UGGACA-CUGGAGAGA-GU------ACUCU-------------U-UUCUAA-
UUAACA-GCA-CC-----A-U------------------CAA-----CC----------------AUGG-AA
U-C-AG--------------------GUUGU-CUGGAGAAGU---------GGUGGU----GUG-UC-UGG-
AAG--AG-----------------------------------------------------------------
-----CGGUGC-CUUCGCGCAUCGU-C-UGGUUAG-UUCAUAACGAU-CCUUGAAA-A-CCC-GAGGGAGAA
-A-------------------------------------------AUCU-----GACUCCCG-CG-CAUGGU
CGUACCCAGAACCGCA------------------------------UCAGGUC--UCCAAGGU-GAGUAGCC
UCGAGU-C-GG-UAGA-CGAAGGUA-A-GUAAGGGAAGUCGGC-AAAAUGGAUCCGUAACUUCGG-GACAAG
GAUUG-GCUC-U-AA--GGGUUG-A----GCAUC-GUC-----GCAC----GUU--------UC--CUG---
-CGG------U---UCUUGAGC--GC----------------------------GUGUU------CGAG---
----------------------------------------------------------------------AG
-AC------------------UC-------------------------------------------------
---------------------------------------------------------GAUGCUC--------
--UC--------------G---------------------------------AG--UC--------------
---------------------------------------------------------------UCUC-----
------------------------------------------------------------------------
-CUCG---------------GC--U-CAU--ACCAUGC---------UUCU---------------------
---UUG----------CAG-----GA--GGCU----AUAGA-G-----------------------------
```

```
-----AGAUA----------------------------UAGG----------------------AGUU----
------------------------------------------------------CUC--GGAU---------
---------------CUUC--------------------CUCUUCCACUG---A------------------
-----GA-------ACCA------------------------------------------------------
----UC-------------------UGAACUA-U-GGUG----CACGAUUGUGAAUAAAACAAC--CAGCUU
AGA--ACUGGUACGGACAAGGGGAAUCCGA-----CUGUUUAAUUAAAACAUAGCAUUGCGAUGGCCGG---
-AA-A------------UCGGUGUUG-ACGCAAUGUGA-AUUCUGCCCAGUGCUCUG-AAUG-UUAAUGUGA
A--------------GAAA--------------UUCAAGGAAGCGC--GGGUAAAC-GGCGGGAGUAACUAU
GACUCUCUUAAGGUAGCCAAAUGCCUCGUCAUC-UAAUUAGUGACGCGCAUGAAUGGAUGAACGAGA--UUC
CCACUGUCCCUACU----UAC-CAUCUAGCGAAACC---AC---AGCCAAGGGAACGGGCUUGGAAGAAUCA
GCGGGGAAAGAAGA-CCCUGUUGAGCUUUGACU--CUAGUCU-GGUUUUGUGGAAUGAGUCGAGAGGUGUAG
CAU-AAGUGGGAGC------------------------------------------------------UUCG
----------------------------------------------------G--C-AA-CAGUGAAAUA-C
CACUACUCUUGGUCUUUUUCCACUCACCC-CG-------U-UA-G-UCCGG------A-----------GG-
------CG--GC-CUCGGUUGUUUACCUAGGCU-C--UC-------------CUUUAUUGUUU---------
----------GUGUA-------------------------------------AUCUG--GG---------AG
------------------AACA----------------------------------------UCU-------
---CCCU------------------------CUC---GGGAAG--UUU------------------------
----------CUC------CUA------------------------------CCGG----UA----------
------------------------------------------------CGGA-----GGACA-UAACCAGGU
GGGGAGUUUGGCUGGGGCGGCACAUCUGCUA-AAAGAUAAC-GCAGGUGUCCU-AAGAUGAGCUCAGUGAGA
ACAGCAAUCUCAUGUG-GAGUA-GAAGGGCAA-AAGCUCAUUUGAUU-UGGA--------------------
--------------UUUUCA--GUACGAAUACAAACUGUGAAA-GCAUGGCCUAUCGAUCCUUU--------
---AC-G-----ACUUCGAAG--------------------UA-UGAAGCUAGAGGUGUCAG-AA--AAGUU
ACCACAGGG-AUAACUGGCUUGUGGCAGCCGAGCGUUCAUAGCGACGUUGCUUUUUGAUCCUUCGAUGUCGG
CUCUUCCUAUCAUUGUGAAGCA-GAAUUCACAAAGCGUUGGAUUGUUCACCCACUGACAGGGAACGUGAGCU
GGGUUUAGACCGUCGUGAGACAGGUUAGUUUUACCCUACU--------------------G-AUG-------
-------------------------------AC--A-GGUCGUCGUGAUAGU-AAU-UCAGCUCAGUACGAG
AGGAA-CCGUUGAUUCAGAUAAU-UGGU-CUUGGCACUUGGCU-GAGA-AGCUAU-C-GGUGCGGAGCUACC
AUCUG-UCGGAUUAUGGCUGAACGCC-UC--UAAGCCAGAAGCCUUGCUAG-----AA--ACGACGAUAGAC
AAUCUCUC-CUGGUUUC-------------------------------------------------------
------------------------------------------------------------------------
------------------------------------------------------------------------
---------------------GAUGUGAUAUCC-----------ACAG--GGC-------------------
--------------------AUAGUGUA---AGAGCA---------AUGGGGGUAAA---------------
------CC--CUU------UU--GC-------------UCUU------------------GUGCCGA---AU
--GUACCCGUGUCAAUCAGAGUAA---CGGUGA------GCGUGGGAGUGA----GUGA-UGACUCGUUGUG
----------------------------UA---U-AUCU---------------------------------
---------------------------------------------------UACUU-----CUU--------
--UUCGUUAAGAAUGGAG-----AGGACAC----AUCGU--G-UUUUC--------AGUAUAU-CU------
----C----UACGAGGAUAAACGGGAGGGUAC-AAAG-GGGUUGCUU--UUUCCCUUU--------------
------------------------------------------------------------------------
------------------------------------------------------------------------
------------------------------------------------------------------------
-------------------------------------------------
```

```
>LSUe_36329_PLAF7/1-3943
-----------------------------UA----GC--ACAGUCUUAAC-GAUGGAUG-U-CUUGGU-UCC
U-A---CAGCGAUGAAGGCCGCAG--CAAAAUGCGAUACGCAAUGAAAAU--U---GCA--GUG-AC--UGU
GAAUCAUCAGAAUGCUGAAUGUAAAC-U--------AC--------AC----CAAU-U-C-----------C
U-U-------------C---G----G----G---GAA----U-A-GUGG-UAC-----UCCUACAGAA-AAU
--------------------------------------------------------G--UUUGAU-UUGA-U
GUU--------------------AG-G-AG-GGCAAAUCCGCCGAAUUUAAGCAUAUAAUUAAGCG--GAAG
AAAAGAAAAUAA--CUAUGAUUCCUUUA--GU-AA-CGGCGAGUGAACAAGGA-C-CA-GCUCAAACGGUUA
AUCC-UUCGUUUGGUGCAAAC------------CUUUCGA--CGGAAUUGUCGU------------------
---U-GAAU-UA--------U-------------------------UCCAU--GAA-----------UG---
CCUUUAA-UUUGA--GCA-------AAGUGA-GCAG------------------------------------
-------GAA-UG-CUUCC-------CCAUAGAGGGUGAAAGGCCCAUA---------GUU---C----UUU
UUAA-UUU--AUUCA-UUGGA----------------AUU-UUUCA--UGCCGAGUCGUGUUCUUUG---AG
AUUGGAGCACUAAUA-CGUGUGAUACA-UUUCACAUAAAGCUAAAUAUGUG--U-AGG-AGACCGAUAGC-A
AAC-A-AGUACCGUGAGGGAAAG-AUGAAAUAGUAC-UCAGG-AAUGAGCA-AUUAAAUAGUACCUGAAAUC
G-UUAAGAUG--G-AAC-GG-AUUA---AGAG-AGA-AAA-----------C---------AAG--------
-------------------------------------------UAAAGAGG--GG----------AAUU---
------UU----------------------------------------UAAU--------------------
------------------------------------------------------------UUUUUUUG----
------------------------------------------------------------------------
----------------------------------UUA-----------------------------UAAUUC
U---------------CUU---CUUU--A--UU----------A----------------AAAGAAACAUC-
---AG---------------------------------------UGAUU-----------------AAUUUA
AUU-----------------UCAAUAAAGCAAUCCCCUGAAAUUCAAAAUUUCUUAAUU----UUCUUUUCA
CUUUCUCCCC--------------------------------------------------GCA--C-UAAUG
U-----------------------------------------------------------------------
-------------------GGGGAAAACUGGCUUUAUU----------UCUUCAAUUAUUUUUUUUGCU---
------------------------------------------------------------------------
------------------------------------------------------------------------
------------------------------------------------------------------------
------------------------------------------------------------------------
-------------------------------------------------GAGGAA-----------------
------U------------UUUAAAAUU-------AUU----------------------------------
---------------------------------------------------UG----AAUUUU---------
--------------UCU-UCUU---------------AU--------AAUUUAAGU----------------
------------------------------------------------------------------------
------------------------------------------------------------------------
-------------------UGUUUCUAU---------------AUAGUAC-UU---UCUAACCCACUCGUCU
UGAAACACGGACCAAGGAGUCUA-G-CAAAUGUGCAA-GUGUAUAU--G-AUUCUUUAAACAUUUCUCUUUU
U-AA--UAUA--CGCAUAAUUAAUG-UAA-----------------UAUGUUUCUUU------AUUGUAGAU
UUGU-GUGGUGU----UUAAUUU---------------------------------------UU--------
AUU---------------------------------------------------------------------
-------------------AAAUC------------------------------------CCCACUUUG-CA
UACAUC----CG-GUAA-G--------------------CAAU----------------UAUGCUUUA-UU-
GAG-UA----------------------CGAGCAUAUUUGGUAG-GACCCGAGAGG-CUUUGAACUAAGCGU
```

```
GAUGAGAUUGAAGUCAGA-CGAAAG-UCUGAUGGAGGAUCGA--GUUGAUACUGACGUGCAAAUCGU-UCAU
UUCAAUCACGUUUAGGGGCGAAAGACUAAUCGAAAAGCC-UAUUAGCUGGUUAUUUCGAAA-GAUCUCUCA
GGAUCGCUGGAG-UUGA-------------------------------------------------------
-----------------AUUAAUA-----UAAUUUUACCAGGUAGAGAUAAUGAUU-AGA-GGAAUCAG-GG
A-A-AUCGU-UUUCCU-UGACCUAUUCUCAAACUCCCAAUAGGUAAAAA-----------------------
------------------GGGCG-AA-----AUAACU--CAAC--UU-GUAUU-------------------
UC--ACUCA-U---AUUAA-AU-A--AAA---U--AACUCCA-AGUGGGC-C-AUUU-U-UGG-UAAGCAGA
AC-U-GGCGAUGA-GGGAUG---CUCCUAACGCUU-G-GAUAAGGUGCCUAAAU--AU-UCG-CU-CA---U
-G-AGAUCCC-AUAAAAGGUGUUGGUUCAUAAUGACAGUAGGACGGUGGUCAUGGAAGUCGAAA-UCCGCUA
AGGAGUGUGUAACAACUCACCUACC-----GAAUGAACUAGCCCUGAAAAU-GGAU-GGCGCUAA-AGCGGA
U----------UACCGAU-ACCAAGCCAUAAAAA--GAA---------------------------------
-----------------------------------------------CA------------------GAA--
----AAAAUUUA------UUUUUU------------------------------------------------
------------------------UUUUAGAAUCUUU-UUAU-GAGUAGAAAAUC-GUGGGGU---------
-----------U-U-GUGUU-GAAGCGAA-A-UACGUGAGUU--U-UCGUGGAACA-UCUCC-CUAGUGCAG
AUCU-UGGUGG-AAGUA-GCAACU---AUUCAAAAUGA--G---------------AACUUU-GAA-GACUG
AAGUGGAG-AAGGGUUUC-UUGUCAACU-GUGAUUGAACAAGAGUUAGCCGC--UC-CUAAGGGAUAGCUG-
A-A-AAGU-----GUUUAAA--------AG------GGG-----GUU-------------------------
-----------------CCUUCCC--------CG----UC-UC-AA-A-AGGGA-AACAGGUU-GAUA-UUC
CUGUG----------------------------------CCAAUA---GUAU--------U-AUGAGUUU--
C----------------UU-----------------------------------------------------
---AGA-UGG---UAACAUAU--AUAU-AA-----AU--------------------GAAC-U-CCUU-UAC
AUAGGCU-------------UUACAC-UCGGGGUGC-GU------UUUCU-------------U-UGCACU-
UUACCU-UUA-UA-----A-C------------------AAA-----CC----------------UUGG-AA
U-C-AA--------------------UUUAC-UUGGAGAAGA---------GGUUCG----UUG-AA-CUC-
AAU--UCAAAAAAUUUCUUGAACGGGGUUUUCGGAUUCAGUUCAUUUUAUUUUUUGUUUUGUAGCAAUAG--
---UAAUUCGU-UUUUAUGAAUUAU-C-CGAAGUG-GUAAGGACUAU-CCUUGAAA-A-AAG-GAGGGAACG
-GCUUUGUAAACUUGGUUUCUUCUGAUUCCAUUUUGCUUUGCUAUACUC-----UUAAUACU-UG-UAUGAG
CGUACCAACAACCGCA------------------------------UCAGGUC--UCCAAGGU-UAACAGCC
UCUGGU-UAAA-UAGA-AAAAAGUA-G-GUAAGGGAAGUCGGCAAAAAUAGAUCCGUAACUUCGG-GAAAAG
GAUUG-GCUC-U-GA--GGACAUUA----GAAAA-GA------GAAGA---AAA--------AA--AGA---
-GGG------U------CGAA---AA----------------------------UAAA---AUU-GCAG--U
C---------------------------------------------------------------------UU
-AUU----------------UGCUU-----------------------------------------------
---------------------------------------------------------UUCUUUC--------
--GAUU-----------UGCU------------------------------UGU--AAU-------------
---------------------------------------------------------------CUGCU-U--
------------------------------------------------------------------------
-U-UU-------CU------UU--U-UCUU-CUUU-CU----------------------------------
---UUU----U-----UUCUCCCCUCUUUUCGCCUUCA--C-------------------------------
----------------------------------------------------------------UUUA----
----------------------------------------------------UUGUA--AUUU---------
---------------U-------------------------------AUU---A------------------
------------------------------------------------------------------------
----CU-----------------------UU------------AAUUUG-AUAC-CUAUAAUGU--UAACUC
```

```
AGA--ACUGAAACGGACAAGGGGAAUCCGA-----CUGUUUAAUUAAAACAUAGCAUUGUGAAAAGCCA---
-CA-A------------CUGGUAUUA-ACACAAUGUGA-UUUCUGCCCAGUGCUUUG-AAUG-UCAACUUGA
U--------------GAAA--------------UUCAAUCAAGCGC--AGGUAAAC-GGCGGGAGUAACUAU
GACUCUCUUAAGGUAGCCAAAUGCCUCGUCAUC-UAAUUAGUGACGCGCAUGAAUGGAUUAACGAGA--UUC
CCACUGUCCCUACU----UGC-UAUCUAGCGAAACC---AC---AGCCAAGGGAACGGGCUUGGCAAAAUCA
GCGGGGAAAGAAGA-CCCUGUUGAGC-UUUACU--CUAGUCU-GGCUUUGUGAAACGACUUAAGAGGUGUAG
CAU-AAGUGGGAGUAG-------------AAAC----------------------------------UGAAA
UAU------------------------------------------GUUUUU-A--C-GA-CAGUGAAAUA-C
CACUACUUUUAAAGUUGUUUUACUUAUCC-AU-------U-UA-U-AGGGA----AAU-----------AUA
U----UAU--GC-UUU-AUCCUUCGGGAAGGCA-UUCUG-------------AUAUU--AUUU---------
----------GAAUU--------------------------------------UUUA--AG---------AA
------------------------------------------------------------------------
---ACU---------------------------U---GUU-----UC-------------------------
----------UUAU---UUUUC------------------------------UCCCAUUUCU----------
------------------------------------------------AUGG-----AGACA-UAGCCAGGU
GGGGAGUUUGACUGGGGCGGUACAUCUGUUA-AAUAUUAAC-GCAGAUGUCCA-AAGACAAGCUCAAAGAGA
ACAGAAAUCUCUUGUA-GACUA-AAAGGGGAA-AAGCUUGUUUGAUU-UCUA--------------------
--------------UUUUCA--GAACAAGUAGAAAACGUGAAA-GCGUGGCCUAUCGAUCCUUU--------
---AU-A-----UUUGCAAAAUGACGUAAAUUACUUACUACUG-UGCAUAUAGAGGUGUCUG-AA--AAGUU
ACCACAGGG-AUAACUGGCUUGUGGCUGCCAAGCGCUCCUAGCGACGUAGCUUUUUGAUCCUUCGAUGUCGG
CUCUUCCUAUCAUUGGGACGCA-GAAGUCUCAAAGUGUCGGAUUGUUCACCCGCUAAUAGGGAACGUGAGCU
GGGUUUAGACCGUCGUGAGACAGGUUAGUUUUACCCUACU--------------------G-AUGAAUUU--
----------UAUUAUAUUUUUUAUAUAUACAU--A-UAGUAUUGUGACAGU-AAU-CCAACUUGGUACGAG
AGGAU-UAGUUGGUUCAGACAAU-UGGU-AC-AGCAAUUGGUU-GACA-AACCAG-U-GUUGCGAACGUA-A
GUCUG-UUGGAUAAUGGCUGAACGCC-UCU-UAAGCCAGAAACCAUGCUGAUU---AA--ACAAUACU-AUU
UCGUUCUU-UUUUGUUU-------------------------------------------------------
------------------------------------------------------------------------
------------------------------------------------------------------------
---------------------CCUUUUUUACUGC------UAAUGUAAAAGA--------------------
--------------------GAAAUUAU---AUAUGU------------ACCUUU-----------------
---UUUUG--GU-------AC--AU------------AAAUA------------------AU----------
----CCUAUCAC---CUUUUU-----CG--UGA------GCUGUGAGCU------GUGA-----GCUAGGCA
A---------------------------U----A-GUUC-------------------UUU-----------
-A----------------ACAACA----------------------A-AAAGAAUU-----GCA------CA
AACUGUAGACGACUUU---UUU-GCCUCAGGGU-GCUGUAAA-CAUGA--------AAGUAAA-CU------
----UUG--UUUUACGAUCUGUUGAGGCUUAU-CCCU-AGUGGUAA---AGUGUU--U--------------
------------------------------------------------------------------------
------------------------------------------------------------------------
------------------------------------------------------------------------
-------------------------------------------------
>LSUe_505711_ANDGO/1-3715 VRVR01000052.1:1-3557,3948-4105 Andalucia godoyi
---------------------------------------CAACUCUUAGC-AAUGGAUC-U-CUUGGU-UCU
C-G---CAUCGAUGAAGAACGCAG--CGAAAUGCGAUACGUGAUGUGAAU--U---GCA--GAC-UA--CGU
GAAUCAUCGAAUCUUUGAACGCAUAU-G--------GC--------AC----UCCU-U-GGC--------UU
G--------------CC---A----A----G---GAG----U-A-UGUC-UGU-----UUGAGGGUCA-GUU
```

```
-UUUC------------------------------------------------------AAUGAC-CUCA-A
AUC--------------------AG-G-CA-AGAGUACCCGCUGAACUUAAGCAUAUCAGUAAGCG--GAGG
AAAAGAAACUAA--CUAGGAUUCCCUCA--GU-AA-CGGCGAGUGAAGCGGGA-A-GA-GCCCAGCAUGAAA
AUCG-GACAUCUUU--------G----------G---UGU--UCGAGUUGUAGGG--UG-------------
--AA-GGAG-CGAUU-----C-------------------------CCUCA--GGC-----------CG---
G-------CCGCG--UAU-------AAGUCC-CAUG------------------------------------
-------GAA-AG-GGGCA--------UCAGAGAGGGUGACAAUCCCGUU---------CAU---GAC-GCGC
GCUU--GG--UCUGG-UUGUGGA-------------GCG---UUUC--CAGAGAGUCAGGUUCCUUG---GG
AGUGGAGCCCAAA-A-UGGGUGGUAGA-UCUCAUCUAAGGCUAAAUAUAGG--C-GAG-AGACCGAUAGC-G
AAC-A-AGUACCGUGAGGGAAAG-AUGAAA-AGCAC-UUUGA-AAAGAG-A-GUUAAA-AGGGCGUGAAAUU
G-CUAAGAGG--U-GAA-GG-AAUG---GCUG-CCU-UGGGU--------------------ACGCCGUACG
GUUUUAGUCGUGG------------------------------CUGUGGAG-------G-GGC--AUCG---
-------------------------UAC------GU-CAGAGA-----GCGCA-------------------
----------------------------------------------------------AGCCUCGCGU----
------------------------------------------------------------------------
-------------------------------------------------------------------GCGGG
U-----CGGCGA---------AGCAG--C-UGC-GGUGAUCACCGUGCG-----GUGU--GGGUCAGCAUC-
---GG----------------------------------UUUUUCCGGU--------GGCAGACGCACAGCA
GGA---------------AGAUGGCU-----------------CGUGGAAUUCUUCGGAAGUCUGCUU----
-----------------------------------------------------------GCUU--U-AUAG-
------------------------------------------------------------------------
---------------------AC-------------------------------UGCUGAUAACGCGGUCGG
CCGGGAAGACC-------------------------------------------------------------
------------------------------------------------------------------------
------------------------------------------------------------------------
------------------------------------------------------------------------
-------------------------------------------------GAGGCC-----------------
------UACAAAGGCUUGGCUGCGGAUC--------------------------------------------
------------------------------------------------------------------------
-----------------------------------------------GUGGCAGG----------------G
C----------------CUCUUAGGGCUGCAUUAGCAACUGUUAAGCCGGUGGCGUCGACCAGAA-------
CG----------------------------GAAAACAGCGUUGAUG----CAGCUCUGGGGAGAGCUGGUUA
CGUGAAGCGUGCCAAGGAUGCUGGCGC------------------AUGGG--U---AGCGAUUCACCCGUCU
UGAAACACGGACCAAGGAGUCUU-A-UACGUGUGCGA-GUUGUAUG--G-U--------GGAAA--------
A-CCAAUGCA--AGCGUAAUGAAAG-UAA----------------------------------------GAG
GUGG-GAUUCGU----CGCA----------------------------------------------------
AGG---------------------------------------------------------------------
-------------------CGA--------------------------------------UGCACCAUC-GA
CCGGUC----UA-GAGCC---------------------CUCUG---------------UGGUAAAGGUACC
GAG-UU----------------------GGAGCACACUUAUAGG-GACCCGAAAGA-UGGUGAACUAUGCCU
GAGCAGGGUGAAGCCAGA-GGAAAC-UCUGGUGGAGGCUCGU--CGCGGUGCUAACGUGCAAAUUGC-UCGU
CGGACUUGGGUAUAGGGGCGAAAGACUAAUCGAACCAUC-UAGUAGCUGGUUCCCUCCGAAG-UUUCCCUCA
GGAUAGCUGGCA-CUCG-------------------------------------------------------
--------------------UUACG----GAGUUUUAUCAGGUAAAGCGAAUGAUU-AGA-GGUACCGG-GA
A-U-GUAAC-AUUC-U-UGACCUAUUCUCAAACUUUCAAUGGGUAAGAA-----------------------
```

```
------------------GGGGU-GG-----UUG-CU--UAAU--U--GAA---------------------
CC---ACCA-A--CCUCGA-AU-C--CA----G--AGUGCUA-AGUGGGC-C-GUUU-U-UGG-UAAGCAGA
AC-U-GGCGACGC-GGGAUG---AACCGAUAGCUG-U-GUUAAAGUGCCCGAGU--GC-ACG-CU-CA---U
--GAGAUCCC-ACAAAGGGUGUUGGUUCAUCUAGACAGCAGGACGGUGGCCAUGGAAGUCGGAA-ACCGCUA
AGGAGUGUGUAACAACUCACCUGCC-----GAAUGAACCAGUCCCGAAAAU-UGAU-GGCGCUCA-AGCGUG
C----------CACUUAU-ACACAGCCACGGAAG--CAAA--------------------------------
------------------------------------------------------------------------
------------------------------------------------------------------------
---------------------------UGCGAAGCUU-CCGU-GUGUAGGAGGGC-GUGGAGCAC-------
---------------GUGGC-GAAGGCGA-A-UGCGUAAGCA--U-GGCUGGAACGUG-GCU-CUAGUGCAG
AUCU-UGGUGG-UAGUA-GCAAAU---AUUC-AAGUGA--G---------------AACCUU-GAA-GGCUG
AAGUGGAG-AAAGGUUCC-AUGGGAACC-GUAGUGGGCCAUGGGUAAGUCGG--UC-CUAAGGAACAGGCC-
A-A-CGCU-----GCGCGGC--------AAGC----GGA-----CAU---UUCGU----------------C
UG---------------CAAGUUC------------------C-GA-A-AGGGA-AGCAGGUU-AAAA-UUC
CUGCACUAUGCAUGGGGGUGCGUGCGCGUACGUGCUUUGCCGAAA---GGUG--------GGGU--------
GGCGGCGCGCAUUGCGGUG-----------------------------------------------------
---GGG-UGG---UAAC--------AC-GA-----AC--------------------GAAC-C-AGAA-GAC
GUCAGCG-------------GCAGUC-CCGGGAAGA-GU------UGUCU-------------U-UUCUUC-
UUAACA-GUC-UU----GA-AGCAUGCUAGACAUGCCAUCGA-----CC----------------CUGG-AA
U-C-AG--------------------AUUGG-CUGGAGAAGG---------GGUUGG----CGG-AC-UGG-
AAA--AG-----------------------------------------------------------------
----CAGCGCG-C-CUUUGCGCUGU-C-UGGUGCA-CUGUCGAUGAU-CCAUGAAA-A-UUC-UGGGGAGCG
-A--------------------------------------------GAU-----GCUCUACA-UG-CAUAGC
CGUACUCAUCACCGCA------------------------------UCAGGUC--UCCAAGGU-GAAUAGCC
UCUAGU-C-AAUUAGA-ACAAUGUA-G-GUAAGGGAAGUCGGC-AAAAUAGAUCCGUAACUUCGG-GAGAAG
GAUUG-GCUC-U-AA--AGGUUG-A----GCCGU-GG------GAUCCAG-GGU------------------
-----------------------------------------------------GCAG---------------
----------------------------------------------------------------------UG
GCUC----------------UGCU------------------------------------------------
---------------------------------------------------------GCC----------AG
ACGG--------------U---------------------------------GGCUAC--------------
------------------GCA-------A----------------------------------GUGGA----
------------------------------------------------------------------------
--------------------CGCUG-UUA--GGCGGGC-----------------------------------
---GGG----UGUCA-CGG-----CCCUGGUCA---------------------------------------
----------------------------------------------------------------GGCG----
------------------------------------------------------------------------
---------------C----------------------------AAGCCG---G------------------
------------------------------------------------------------------------
-------------------------------------------CCCACG-GCA--ACGAACAAC--CAAUUU
AGA--ACUGGUACGGACAAGGGGAAUCCGA-----CUGUUUAAUAAAAACAAAGCAUUGUGAUGGCCUU---
-UG-C-------------AGGUGUUG-ACACAAUGUGA-UUUCUGCCAGUGCUCUG-AAUG-UCAAAGUGA
A--------------GAAA--------------UUCAACCAAGCGC--GGGUAAAC-GGCGGGAGUAACUAU
GACUCUCUUAAGGUAGCCAAAUGCCUCGUCAUC-UAAUUAGUGACGCGCAUGAAUGGAUUAACGAGA--UUC
CCACUGUCCCUAUC----UAC-UAUCUAGCGAACCC---AC---AGCCAAGGGAACGGGCUUGGGACUAACA
```

```
GCGGGGAAAGAAGA-CCCUGUUGAGC-UUGACU--CUAGUCC-GACUUUGUGAAAUGACUUAGAGGGUGUAG
AAU-AGGUGGGAGCGC--------------------------------------------------------
-------------------------------------------------AA-G--C-GA-UCGUGAAAUA-C
CACUACCUUUAAUGUUAUUUUACUUAUUG-GG-------U-AA-G-GUGGA------GU----------CG-
-----------------------------GUGGCC--AGAAAUGGUCGCUG-CUUCU---------------
-----------GGAA--------------------------------------UUAA--GC---------CC
---------------------------------------------------------------CC-------
---UCU-------------------------------UGU-----GG-------------------------
----------GGG------UGA------------------------------UCCA----UG----------
------------------------------------------------CCCA-----AGACA-UUGUCAGGU
GGGGAGUUUGGCUGGGGCGGCACAUCUGUUA-AACGGUAAC-GCAGAUGUCCU-AAGAUGAGCUCAAUGAGA
ACAGAAAUCUCAUGUA-GAACA-AAAGGGUAA-AAGCUCAUUUGAUU-UUGA--------------------
--------------UUUUCA--GUACGAAUCCAAACUGUGAAA-GCAUGGCCUAUCGAUCCUUU--------
---AG-U-----CAUUCGAGG--------------------UC-UGAAGCUAGAGGUGUCAG-AA--AAGUU
ACCACAGGG-AUAACUGGCUUGUGGCAGCCAAGCGCUCAUAGCGACGUUGCUUUUUGAUCCUUCGAUGUCGG
CUCUUCCUAUCAUUGUGAAGCA-GAAUUCACAAAGUGUCGGAUUGUUCACCCGCUAACAGGGAACGUGAGCU
GGGUUUAGACCGUCGUGAGACAGGUUAGUUUUACCCUACU--------------------G-UUA-------
-------------------------------GG--A-GGUUAUCGCCACAGU-AAU-GCAACCCAGUACGAG
AGGAA-CCGUUGUGUCGGAUCAC-UGGU-AUUUGCCCUUCGCU-GAAC-AGCGAA-U-GGGGCGAAGCUACC
AUCCG-UUGGAUUAUGACUGAACGCC-UC--UAAGUCAGAAUCCAUGCUGA-----GA--GCGAUGAU-AUC
UACCUGCU-GCGGAUUA-------------------------------------------------------
------------------------------------------------------------------------
------------------------------------------------------------------------
---------------------GCCG------------------------CA-----CA--------------
-------------------------------CGGAUA-----G-----------------------------
------------------------------------------------------------GCACUAC---U-
--------------------------UGUUGGUGCC----CUGAGUCGUAG----GUAA----GUCGAG---
------------------------------------------------------------------------
------------------AAGCGGC---------------------G-CGGCGGAA-----GGG------CC
GUGAGCGCAUCGGCAGAG-UAC-AAGAUGGAGU-AGC---AG-CAGGG--------GACAUAC-CU------
----U-------------------------------------------------------------------
------------------------------------------------------------------------
------------------------------------------------------------------------
------------------------------------------------------------------------
-------------------------------------------------
>LSUe_857967_ICHMG/1-3507 AEDN01002274.1:1-4270 Ichthyophthirius multifilii
------------------------------A----GA--AAAUUUUCAAC-GGUGGAUA-U-CUAGGU-UCC
C-G---UGACGAUGAAGAACGCAG--CGAAAUGCGAUACGCAAUGCGAAU--U---GCA--GAAUC---CGC
GAGUCAUCAGAUCUUUGAACGCAAGU-G--------GU--------GG----AG---G------------CU
A-A-----------------A----A----C---CUU----C-A-UGUU-UGU-----AUCAGUGUGGAAA-
-----------------------------------------------------------UUACAC-CUGA-U
A-U--------------------AA-G-CA-AGAUUACCCGCUGAACUUAAGCAUAUCAGUAAGCG--GAGG
AAAAGAAAUUAA--CGAAGAUAGCCUUA--GU-AA-CUGCGAGUGAACAGGCU-G-AA-GCUCAAAGUGUAA
AUCU-GCGUGGGUC----AAC------------C---ACA--AAGAAUUGUAAUC--UA-------------
--UA-GGGU-UAA-U-----C-------------------------CGAAG--GUA-----------AG---
```

```
CU------ACUCG--CAU-------AAGUUU-UUUG------------------------------------
-------GAA-CA-AGACG-------UCAAAGAGGGUGACAACCCCGUA---------GUC---G----GCG
A-GU--AU--GCUGC-UGUAG---------------GAGA-GGCUU--CGAAGAGUCGGGUUGUUUG---GG
ACUGCAGCCCUAA-G-UGGGAGAUAAA-CUUCUUCUAAAGCUAAAUAUUCA--C-GGG-AGACCGAUAGC-G
AAC-A-AGUACUGCGAAGGAAAG-AUGAAA-AGAAC-UUUGA-AAAGAG-G-GUUAA--AAGACUUGAAACC
G-UUGAGAAG--G-AAG-CG-GUAG---AAGA-GCAAUAAACUGGCUUCGGUGUAAACCGUCUG--------
-------------------------------------------AUGAUACU---------CAC--UGCU---
------G------------------UAU--------------------AGAU--------------------
------------------------------------------------------------ACG--AAA----
-----------------------GG-----------------------------------------------
----------------------------------UCG----------GUG-------------------CGG
U----GUGAAG------------AGG--UCAUC----------A---------------AGCGGUGCUACUG
G--AG----------------------------------------------------------------UUA
GGAGACAAAAUGAUGUUUGGUUGAAGGAGUCACC---UGAGAUU-CGUA-----------------------
-----------------------------------------------------------------G-C----
------------------------------------------------------------------------
--------------------AUCAAAAGGAAAACUUCAA------UCGGGCAU-------------------
------------------------------------------------------------------------
------------------------------------------------------------------------
------------------------------------------------------------------------
------------------------------------------------------------------------
-------------------------------------------------GAGGGG----C------------
------C------------UAAGGGCGA--------------------------------------------
------------------------------------------------------------------------
---------------------------------------U------UUUGUC--------------------
------------------------------------------------------------------------
------------------------------------------------------------------------
-------------------------------------------AAAAUGG--C---UUCUACUGACCCGUCU
UGAAACACGGACCAAGGAGUCUA-U-CAAUUAAGCGA-GUGACAGG--G-UG--------GAAA-------A
A-CC--CGUC--CGCAAAACGAAG-UGA------------------GUA-------------------CAAG
GUGC-CAA-CC---------------------------------------------------GC--------
AA----------------------------------------------------------------------
-------------------GGU---------------------------------------GCAGCAUC-AC
CCGGCC----UC-AAGU----------------------CUCCG----------------ACAAGAGGAUC-
GAG-GA----------------------AGAGCUUAAUUGAUAG-GACCCGAAAGA-UGGUGAACUACGCUU
GAAUAGGGUGAAGCCAGG-GGAAAC-UCUGGUGGAAGCUCGU--AGCGAUACUGACGUGCAAAUCGU-UCGU
CAAAUUUGAGUGUAGGGGCGAAAGACUAAUCGAACCAUC-UAGUAGCUGGUUCCCUCCGAAG-UUUCUCUCA
GGAUAGCAAGGG-CAAU-------------------------------------------------------
--------------------AACG-----CAGUUUUAUUAGGUAAAGCGAAUGAUU-AGA-GGAUUUGG-GG
U-U-CCAAG-AAUC-U-CGACCUAUUCUCAAACUUUAAAUUGGUAAGAG-CC--------------------
------------------GUAGA-GU-----UUU-CU--UAAU--U--GAA---------------------
CU---CUCG-G---GUAGA-AU-G--CA----G--UGCUCUU-AGUGGGC-C-AUUU-U-UGG-UAAGCAGA
AC-U-GGCGAUGA-GGGAUG---AACCUAACGCUU-A-GAUAAGGCGCCCAAAU--AC-ACG-CU-CA---U
-C-AGAUACC-AAAAAAGGUGUUGGUUCAUUUAGACAGCAGGACGGUGGCUAUGGAAGUUAGAA-CCCGCUA
AGGAGUGUGUAACAACUCACCUGCC-----GAAUGAACUAGCCCUGAAAAU-GGAU-GGCGCUGA-AGCGUG
```

```
U----------UGCCGAU-ACUAAGCCAUUAGAG--CAAA--------------------------------
------------------------------------------------------------------------
------------------------------------------------------------------------
---------------------------UGCGAGGCUU-UGAU-GAGUAGGAGGC-GUGAACA----------
-----------U-C-GCCUA-GAAGUCUU-G-GGUGUGAACC--U-GGAUGGAGCG-G-UGU-UUAGUGCAG
AUCU-UGGUGG-UAGUA-GCAAAU---AUUC-AAAUGA--G---------------AACUUU-GAA-GACCG
AAGUGGAG-AAGGGUUCC-AUGAGAACA-GCAUUUGUUCAUGGGUUACUCGA--UC-CUAAGACAUAGGUU-
A-A-CUCC-----UUGUAAU--------ACAA----GCG-----GAC----CUC------------------
-----------------GGUCUGC--------GU----UG-UC-AA-A-AGGGA-AUGAGGUU-AAUA-UUC
CUCAA----------------------------------GCUGGA---UGUG--------GAUG--------
U----------------UG-----------------------------------------------------
----AG-UGG---UAAC--------AC-AA-----AG--------------------AAAC-C-CGGA-GAC
GUCAGCA-------------GGUGUC-ACUGGAAGA-GU------UAUCU-------------U-UUCUUU-
UUAACA-UAC--U-----A-U-------------------GG-----CC----------------UUGA-AG
--U-GG--------------------AUUAU-CCAGAGA-AG--------AGGCUGU----ACG-UA-UGG-
CAG--AG-----------------------------------------------------------------
----CAGCUCA-CCCUAAGAGCUGU-C-AGUUGCG-CAUCUGAUGAC-CCUUGAAA-A-UCU-GGGGGAGAC
UA----------------------------------------------------UCAUAUCA-CA-CCAGGU
CGUACCCAUAACCGCA------------------------------UCAGGUC--UCCAAGGU-UAGCAGCC
UCUGGU-C-AA-UAGA-ACAAUGUA-G-AUAAGGGAAGUCGGC-AUAUUGGAUCCGUAACUUCGG-GAAAAG
GAUUG-GCUC-U-GA--GGAUUG-G----GUGAAUGG------GGCUU---UAC------------------
--AU------UG----AACUCUAAGC----------------------------UUG--------UGCA---
------------------------------------------------------------------------
-UGG-----------------GCG------------------------------------------------
---------------------------------------------------------UG-----------GA
AACA--------------U---------------------------------GC--CU--------------
---------------------------------------------------------------GCGC-----
------------------------------------------------------------------------
--------------------AUAAG-CUGCCGAGG-------------------------------------
-------------AAUUUG-----UA-GAGU-----------------------------------------
----------------------------------------------------------------GGCU----
------------------------------------------------------------------------
-----------------------------------------------UCG---G------------------
------------------------------------------------------------------------
----CC-----------------------UU------------CCCUUC-ACA--AUGAACAAU--CAACUC
AGA--ACUGAAGCGGACAAAGGGAAUCCGA-----CUGUUUAAUAAAAACAAAGCAUUGUGACGGCCUC---
-AA-C-------------AGGUGUUG-ACACAAUGUGA-UUUCUGCCCAGUGCUCUG-AAUG-UCAAAGUGA
C--------------GCAA--------------UUCAACCAAGCGC--GGGUAAAC-GGCGGGAGUAACUAU
GACUCUCUUAAGGUAGCCAAAUGCCUCGUCAUC-UAAUUAGUGACGCGCAUGAAUGGAUUAAUGAGA--UUC
CCACUGUCCCUAUC----UAC-UAUCUAGCGAACCC---AC---AGCUAAGGGAACGGGCUUAGAAUAAUCA
GCGGGGAAAGAAGA-CCCUGUUGAGC-UUGACU--CUAGUCU-AACUUUGUGAAAUGGCACGGGAGGUAUAG
CCU-AGGUGGGAGA------------------------------------------------------GUAA
----------------------------------------------------U--C-AA-UCCUGUAAAA-C
CACCACUUCCGGGGUCAUUUGCUUAUUU-CG--------U-GA-A-GAAAU------G-----------GC-
------UG--GC-GU--------AAG---CCAG-U--U-----------------CU---------------
```

```
----------AAAAG--------------------------------------UUAA--GG---------GC
------------------------------------------------------------------------
---GAU---------------------------U---UUC--------------------------------
----------GUC------UGA------------------------------UUUU----UG----------
------------------------------------------------CGAA-----AGACA-UGGUUAGGG
GGGGAGUUUGUCUGGGGCGGAAUGCCUGUUA-AACCAUAAC-GCAGGCGUCCU-AAGGUGAGCUCAAUGAGA
ACGGAAAUCUCAUGUA-GAAUA-AAAGGGUAA-AAGCUCACUUGAUU-UUGA--------------------
--------------UUUUCA--GUAUGAAUACAAACCGCGAAA-GCGUGGCCUAUCGAUCCUUU--------
----A-A-----CUUUACAAG--------------------UU-UUAAGCUAGAGGUGUCAG-AA--AAGUU
ACCACAGGG-AUAACUGGCUUGUGGCAGCCAAGAGUUCAUAUCGACGUUGCUUUUGAUCCUUCGAUGUCGG
CUCUUCCUAUCAUUGUGAAGCA-GAAUUCACAACGUGUCGGAUUGUUCACCCGCUAAUAGGGAACGUGAGCU
GGGUUUAGACCGUCGUGAGACAGGUUAGUUUUACCCUACU--------------------G-AUG-------
-------------------------------AA--A-UGAUAUUGCGACAGU-AAU-UUAACUUAGUACGAG
AGGAA-CCGUUAAAUCAGAUAAU-UGGU-AAAUACGGUUGUCU-GAAA-AGACAA-U-GCCGUGAAGCUACC
AUCUG-UAAGAUUAUGACUGAAGGCC-UC--UAAGUCAGAAUCUAUGCUGGA----AA--GCAAUGUC-CUG
--GUGA------------------------------------------------------------------
------------------------------------------------------------------------
------------------------------------------------------------------------
------------------------------------------------------------------------
-------------------------------UGAUAA-----------ACGAAAAA----------------
--A-------AUAAGCGCAAG--CU--------------UAA------------------GUUCGAAUUGG-
-------------------UG-----CGA-GUCG-----GCUGAA-GAC------GAAA-----GAGUCCAG
GCU-------------------------CAU--CGCAUG-----------AAUCGUA-AUU----------U
C-----------------GAAUUAU---------------------C-AUCAACAU-----AAA------UC
UUUUGUAGACGACUUAGU--UCAGGAGA-GGGU-AUUGUAAG-UAUGA--------GAGUAGA-AU------
----U----UUCUACGAUCUGCUGAGAUUCAU-CCCU-UUUCCUAA---GAUUUAUCU--------------
------------------------------------------------------------------------
------------------------------------------------------------------------
------------------------------------------------------------------------
-------------------------------------------------
>LSUe_7227_DROME/1-4077
----------------------------------------AACUCUAAGC-GGUGGAUC-A-CUCGGC-UCA
U-G---GGUCGAUGAAGAACGCAG--CAAACUGUGCGUCAUCGUGUGAAC--U---GCA--GGA-C---ACA
UGAACAUCGACAUUUUGAACGCAUAU-C--------GC--------AG----UCCA-U-GC----------U
G-U------------GC---U----U----G---GAC----U-A-CAUA-UGG-----UUGAGGGUUG-UAU
-----------------------------------------UAU------------A--UACAAC-CUCA-A
CUC--------------------AU-A-UG-GGACUACCCCCUGAAUUUAAGCAUAUUAAUUAGGG--GAGG
AAAAGAAACUAA--CAAGGAUUUUCUUA--GU-AG-CGGCGAGCGAAAAGAAA-A-CA-GUUCAGCACUAAG
UCAC-UUUGUCUAUA---UGG------------C---AAA-UGUGAGAUGCAGUG--UA-------------
--UG-GAGC-GUC-A--AUAU-------------------------UCUAG--UAU-----------GA---
GAAAUU--AACGA--UUU-------AAGUCCUUCUU------------------------------------
-------AAA-UG-AGGCCAUU-UACCCAUAGAGGGUGCCAGGCCCGUA---------UAA--------CGU
UAA---UG--AUUAC-UAGA----------------UGAU-GUUUC--CAAAGAGUCGUGUUGCUUG---AU
AGUGCAGCACUAA-G-UGGGUGGUAAA-CUCCAUCUAAAACUAAAUAUAAC--C-AUG-AGACCGAUAGU-A
AAC-A-AGUACCGUGAGGGAAAG-UUGAAA-AGAAC-UCUGA-AUAGAG-A-GUUAAACAGUACGUGAAACU
```

```
G-CUUAGAGG--UUAAG-CCCGAUG---AACC-UGA-AUAUC--------------------CG--------
-------------------------------------------UUAUGGAA--AAUUCAUCAU--UAAA---
------AU-----------------UGU------------A-------AUAU--------------------
--------------------------------------------------------------UUAAAU----
-----------------------AA-----------------------------------------------
----------------------------------UAU---------UAUG-----------------AGAAU
A-----GUGUGCA-UUUUU---UCCA--U--AU-----------------------AAGGACAUUGUAAUC-
---UA-------------UU-------------------AGCAUAUACC-----------------AAAUUU
AUC-----------------AUAAAAUAUAACUUAUAGUUAUUCCAAU------------------------
--------------------------------------------------------------U--A-A----
------------------------------------------------------------------------
----AUUGCUUGCAUUUUAACACAGAAUAAAUGUUAUUA------AUUUGAUAAAGUGCUGAUAGAUU----
------------------------------------------------------------------------
------------------------------------------------------------------------
------------------------------------------------------------------------
------------------------------------------------------------------------
-------------------------------------------------UAUAUG-----------------
------A------------UUACAGUGC----GUUAAUUUUC-----------------------------G
------------------------------------GAAUUAU---A-UAAUGG---CAUAAUUAUCA----
UUGAUUUUUGUGUUUAU-UAU---AU---------------------GCACUUGU-----------------
------------------------------------------------------------------------
------------------------------------------------------------------------
-----------------AUGAUUAACAAUGCGA------AAGAUUCAGGA--U---ACUUCGGGACCCGUCU
UGAAACACGGACCAAGGAGUCUA-A-CAUAUGUGCAA-GUUAUUGG--G-A--------UAUAA--------
A-CC--UAAU--AGCGUAAUUACU-UGA------------------CUAAUAAU-----------------G
GGAU-UAGUUUU----UUAGCUAUUU------------------------------------AU--------
AGC---------------------------------------------------------------------
-------------------UAAUU---------------------------------AACACAAUCCCG-GG
GCGUUCUAUAUA-GUUA-UGUAUAAUGUAUA--------UUUA-UAUUAUUUAUGCCUCUAACUGGAA-CG-
UAC-CU----------------------UGAGCAUAUAUGCUGU-GACCCGAAAGA-UGGUGAACUAUACUU
GAUCAGGUUGAAGUCAGG-GGAAAC-CCUGAUGGAAGACCGA--AACAGUUCUGACGUGCAAAUCGA-UUGU
CAGAAUUGAGUAUAGGGGCGAAAGACCAAUCGAACCAUC-UAGUAGCUGGUUCCUUCCGAAG-UUUCCCUCA
GGAUAGCUGGUG-CAUU-------------------------------------------------------
--------UUAAUAUUAUAUAAAA-----UAAUCUUAUCUGGUAAAGCGAAUGAUU-AGA-GGCCUUAG-GG
U-C-GAAAC-GAUC-U-UAACCUAUUCUCAAACUUUAAAUGGGUAAGAA-----------------------
------------------CCUUA-AC-----UUU-CUUGAUAU--G--AAG---------------------
AU---CAAG-G---UUAUG-AU-A--UA----A--AUGUCCC-AGUGGGC-C-ACUU-U-UGG-UAAGCAGA
AC-U-GGCGCUGU-GGGAUG---AACCAAACGUAA-U-GUUACGUGCCCAAAUU--AA-CAA-CU-CA---U
-GCAGAUACC-AUGAAAGGCGUUGGUUGCUUAAAACAGCAGGACGGUGAUCAUGGAAGUCGAAA-UCCGCUA
AGGAGUGUGUAACAACUCACCUGCC-----GAAGCAACUAGCCCUUAAAAU-GGAU-GGCGCUUA-AGUUGU
A----------UACCUAU-ACAUUACCGCUAAAG--UAGAU-------------------------------
-----------------------------------------------GA------------------UUU-A
----UAUUACUU------GUGAUAUA----------------------------------------------
------------------------AAUUUUGAAACUU-UAGU-GAGUAGGAAGGU-ACAAUGG----------
-----------U-AUGCGUA-GAAGUGUU-U-GGCGUAAGCC--U-GCAUGGAGCU-G-CCA-UUGGUACAG
```

```
AUCU-UGGUGGAUAGUA-GCAAAU---AAUC-GAAUGA--G---------------AGCCUUGGAG-GACUG
AAGUGGAG-AAGGGUUUC-GUGUGAACA-GUGGUUGAUCACGAGUUAGUCGG--UC-CUAAGUUCAAGGCG-
A-A-AGC------GAAAAUU--------UUCA----AGU----AAAA---CA--------------------
--------------------UAAU-------UUU----GA-AC-GA-A-AGGGA-AUACGGUU-CCAA-UUC
CGUAA----------------------------------CCUGUUGAGUAUC--------C-GUU-------
U----------------GUUAUUA--------------------------------AAUAUGGGCCUCGUGC
UCAUCC-UGG---CAAC--------AG-GA-----AC--------------------GACC-A-UAAA-GAA
GCCGUCG-------------AGAGAU-AUCGGAAGA-GU------UUUCU-------------U-UUCUGU-
UUUAUA-GCC-GU-----A-C-------------------UA-----CC----------------AUGG-AA
G-U-CU--------------------UUCGC-AGAGAGAUAU---------GGUAGA----UGG-GC-UAG-
AAG--AG-----------------------------------------------------------------
-----CAUGAC-AUAUACUGUUGUG-U-CGAUAUU-UUCUCCUCGGA-CCUUGAAA-A-UUU-AUGGUGGGG
-A------------------------------------------CACGC-----AAACUUCU-CA-ACAGGC
CGUACCAAUAUCCGCA------------------------------GCUGGUC--UCCAAGGU-GAAGAGUC
UCUAGU-C-GA-UAGA-AUAAUGUA-G-GUAAGGGAAGUCGGC-AAAUUAGAUCCGUAACUUCGG-GAUAAG
GAUUG-GCUC-U-GA--AGAUUG-A----GAUAG-UC------GGCCUU--GAU-UGGGAA-AC--AAUAAC
AUGG------U-----UUAUGU--GC----------------------------UCGU-------UCUG--G
G-UA--------------------------------------------------------------AAUAGA
-GUU-----------------UCU------------------------------------------------
---------------------------------------------------------AGCAUUUA-------
UGUU--------------AGU------------------------------UAC--UU--------------
-----------------GUUC-------C----------------------------------CCGG-----
------------------------------------------------------------------------
-A-UAG-------UU----UAGUUA-CGU--AGCC-AA----------------------------------
---UUG----UG-GAACUU-----UC-UUGCUAAA-AUUUU-------------------------------
-----UAA--------------------------------GAA--------------------UACUA----
------------------------------------------------------UUU--GGGU---------
---------------U----------------------------AAACCA---A------------------
----UUA-------GU--------------------------------------------------------
----UC-----------------------UUAUU-AAUUAUAACGAUUA-UCA--AUUAACAAU--CAAUUC
AGA--ACUGGCACGGACUUGGGGAAUCCGA-----CUGUCUAAUUAAAACAAAGCAUUGUGAUGGCCCU---
-AG-C-------------GGGUGUUG-ACACAAUGUGA-UUUCUGCCCAGUGCUCUG-AAUG-UCAAAGUGA
A--------------GAAA--------------UUCAAGUAAGCGC--GGGUCAAC-GGCGGGAGUAACUAU
GACUCUCUUAAGGUAGCCAAAUGCCUCGUCAUC-UAAUUAGUGACGCGCAUGAAUGGAUUAACGAGA--UUC
CUACUGUCCCUAUC----UAC-UAUCUAGCGAAACC---AC---AGCCAAGGGAACGGGCUUGGAAUAAUUA
GCGGGGAAAGAAGA-CCCUUUUGAGC-UUGACU--CUAAUCU-GGCAGUGUAAGGAGACAUAAGAGGUGUAG
AAU-AAGUGGGAGAUAUUAG-------AC---------------------------------------CUCG
-------------------------------------------GUUUGGUA-U--C-GU-CAAUGAAAUA-C
CACUACUCUUAUUGUUUCCUUACUUACUU-GA-------U-UA-A-AUGGAACGUGUAUCAUUUCCUAGCCA
UUAUACGG--AU-AU-----AUUUAUU-AUAUC-UUAUG-------------GUAUUGGGUUUUGAUGCAAG
-CUUC---UUGAUCA----------AAGU-----------------------AUCAC--GA---------GU
------------------UUGUUAUAUAAUCGC-----AAAC--AAAUUC-------------UUUAAUAAA
ACGAUGC------------AUU---------UAU---GUA-----UUUU-----------------------
------UGAUUUGAAAAUUUGGUA-UA-----------------------ACUCCAA---UUAC--------
--------------------------------UCAGGUAUGAUCCAAUUCAA-----GGACA-UUGCCAGGU
```

```
AGGGAGUUUGACUGGGGCGGUACAUCUCUCA-AAUAAUAC-GGAGGUGUCCC-AAGGCCAGCUCAGUGCGG
ACAGAAACCACACAUA-GAGCA-AAAGGGCAA-AUGCUGACUUGAUC-UCGG--------------------
--------------UGUUCA--GUACACACAGGGACAGCAAAA-GCUCGGCCUAUCGAUCCUUU--------
---UG-G-----UUUAAAGAG--------------------UU-UUUAACAAGAGGUGUCAG-AA--AAGUU
ACCAUAGGG-AUAACUGGCUUGUGGCGGCCAAGCGUUCAUAGCGACGUCGCUUUUUGAUCCUUCGAUGUCGG
CUCUUCCUAUCAUUGUGAAGCA-AAAUUCACCAAGCGUUGGAUUGUUCACCCA-UGCAAGGGAACGUGAGCU
GGGUUUAGACCGUCGUGAGACAGGUUAGUUUUACCCUACU--------------------A-AUGACA----
-------------------------------AA--A-CGUUGUUGCGACAGC-AUU-CCUGCGUAGUACGAG
AGGAACCCGCAGGUACGGACCAA-UGGC-AC-AAUACUUGUUC-GAGC-GAACAG-U-GGUAUGACGCUA-C
GUCCG-UUGGAUUAUGCCUGAACGCC-UC--UAAGGUCGUAUCCGUGCUGGA----CU--GCAAUGAUAAAU
AAGGGGCA-AUUUGCAU-------------------------------------------------------
------------------------------------------------------------------------
------------------------------------------------------------------------
---------------------UGUAUGGCUUCU---------------------------------------
--------------------AAACCAUUUAAAGUUUA------------UAAUUU-----------------
---ACUUU--AU-------AA--ACG---------ACAAUGG------------------AUGUGAU---G-
-----CCAAUGUA-AUUUGU-AACAUA------------GUAAAUUGGGAGG--AUCUUCGA--UCACCUGA
----------------------------UGCCGC-GCUAGUUACAUAUAAAAGCAUU-AUUUA--------A
U-----------------ACAAUGA---------------------C-AAAGCCUA-----GAA------UC
AAUUGUAAACGACUUUUG-UAA-CAGGCAAGGU-GUUGUAAG-UGGUU---------GAGCAGC-UG------
----CCA--UACUGCGAUCCACUGAAGCUUAU-CCUU-UGCUUGAU---GAUUCG--A--------------
------------------------------------------------------------------------
------------------------------------------------------------------------
------------------------------------------------------------------------
-------------------------------------------------
>LSUe_1928728_PAMIC/1-3671 BJOX01000850.1:27283-27444,27918-31426 Paulinell
---------------------------------------CAACUCCUAGC-GACGGAUA-U-CUUGGU-UCU
C-G---CAACGAUGAAGAACGCAG--CGAAAUGCGAUACGUAAUGCGAAU--U---GCA--GAA-UUC-AGU
GAAUCAUCAAAUCUUCGAACGCAGAC-U--------GC--------GC----UUUC-G-GGA---------U
A-A-----------CCC---U----G----G---AAG----C-A-CGCC-UCU-----UUCAGUGUCG----
---------------------------------------------ACUAC------A--CUAGAC-CUGA-G
AUG--------------------AA-G-UG-AGACUACCCGCUGAAUUUAAGCAUAUUAUUAAGCG--GAGG
AAAAGAAACUAA--CCAGGAUUCCCUCA--GU-AA-CGGCGAGUGAAGCGGGA-A-GA-GCUCAAAGUGAAA
AUCG-GUCACCCAG----GCG------------G---GUG-UUCGAGUUGUAAUC--UA-------------
--GA-GACG-UAUGG-----U-------------------------CUGCG--CGC-----------GA---
C-------CUGGG--UAU-------AAGUCC-UUUG------------------------------------
-------GAA-CA-GGGCG--------UCAUAGAGGGUGAGAACCCCGUG---------CAU---GAC-CCGG
AGU---UC--GCCGC-GUACGAU-------------AUG---CGUU--CGAUGAGUCGUGUUGCCUG---GG
AUCGCAACACUAA-A-UGGGUGGUAAA-UCUCAUCUAAAGCUAAAUACUGG--C-GAG-AGACCGAUAGC-G
AAC-A-AGUAGCGUGAGCGAAAG-AUGAAA-AGAAC-UUUGA-AAAGAG-A-GUUAAACAGUACUUGAAAUC
G-CUAGGAGG--G-AAG-UU-AUGG---AAA-----------------------------------------
----------------------------CUAG-----------UUUCAGCA--UC------AC---UUGGGC
AUUGCCGGA----------------GAU--------------------GCGC--------------------
-------------------------------------------------------------CGCGGGAAGGU
GGUAGCUUGCUACUUUAUAGCC--------------------------------------------------
```

------------------------------------------------------------------------
-------------------------------------------CGUGGU-----------UG-CGACGGUG-
---GG----------------------------------------CCGG-----------------AGUGAG
GAA---------------GGAGAA------------------------GGCAUCUAAGGCACCAGUGC----
--------------------------------------------------------------A---------
------------------------------------------------------------------------
--------------------------------------------------------------UGCUCGCCGG
GCGG--------------------------------------------------------------------
------------------------------------------------------------------------
------------------------------------------------------------------------
------------------------------------------------------------------------
------------------------------------------------------------------------
-----------------------UCAGG--------------------------------------------
------------------------------------------------------------------------
-----------------------------------------------GACAGCAA----------------A
G----------------CUGUGUCUGGUUGGUCGGCGUAUUUGCGCGGGGUUCGCUAUU-------------
------------------------------------------------------------------------
-----CGUGCCUU-UGCGAUGCUGAGUA-----------------AAUGG--U---UUUCAUUGACCCGUCU
UGAAACACGGACCAAGGAGUCUA-A-CAUAUAUGCGA-GUAUUUGGGUG-U--------GUCAA--------
A-CC--CUGG--UGCGCAAUGAAAG-UGAAGGGCCGC----AAGGCCUGA---------------------G
GUAG-GAACCG---------------------------------------------------UCUUUAACGG
GG----------------------------------------------------------------------
-------------------CGG--------------------------------------GGCACUAUC-GA
CCGAUC----GC-CGC--------------------------U----------------UGCGGUAGAUUU-
GAG-UA----------------------AGAGCAUAUCUGUUGG-GACCCGAAAGA-UGGUGAUCUAUGCCU
GAACAGGGUGAAGCCAGG-GGAAAC-UCUGGUGGAAGCUCGU--AGCGGUACUGACGUGCAAAUCGU-UCGU
CUGAUUUGGGUAUAGGGGCGAAAGACUAAUCGAACCAUC-UAGUAGCUGGUUCCCUCCGAAG-UUUCCCUCA
GGAUAGCUGGAG-CUCA-------------------------------------------------------
-----------------------GUUAAUCAGUUUUAUCAGGUAAAGCUAAUGAUU-AGA-GGUACCGG-GG
G-C-GAAAU-GCCC-U-UGACCUAUUCUCAAACUUUAAAUAGGUAAGAA-----------------------
------------------CCCCG-UGG----UUA-UU--UGAU--U--GAG---------------------
CC---CGGG-G---AUUGA-AU-G--A-----ACGAGUUCUU-AGUGGGC-C-AUUU-U-UGG-UAAGCAGA
AC-U-GGCGAUGC-GGGAUC---AUCCGGACGCAU-GAGUUAAGGUGUCCAAGU--GU-UGG-UU-CA---U
-CCAGAUUAC-GCAAAAGGUGUUGGUUGAUAGAGACAGUUGGACGGUGGUCAUGGAAGUCGAAA-UCCGCUA
AGGAGUGUGUAACAACUCACCAACC-----GAAUCAAUUAGCCCUGAAAAU-GGAU-GACGCUAA-AACCAA
C----------CACCUAA-ACUCAGCCACCAGCG--CAAG--------------------------------
------------------------------------------------------------------------
------------------------------------------------------------------------
---------------------------UGCGAGGCGU-UGGU-GAGUAGGAGGGC-GUGGGGGU--------
----------CU-C-GUGAC-GAAGGCUA-G-GGCGUGAGCC--C-GGCUGAAACG-G-CCC-CUAGUGCAG
AUCU-UGGUGG-UAGUA-GCAAAU---AUUC-AAAUGA--G---------------AACUUU-GAA-GACUG
AAGUGGAG-AAAGGUUCC-AUGUGAACA-GCAAUUGGACAUGGGUUAGUCGA--UC-CUAAGAGAUAGGGA-
A-A-CUCC-----GUUCAA---------AGU-----GCG-----CA----CUGC-----------------C
UG---------------CG----C------CGCC----UA-UC-GA-A-AGGGA-ACCGGGU-AAUA-UUC
CCGGA----------------------------------CCAGGA---CGUG--------G-AU--------

```
A----------------UU-----------------------------------------------------
---GUG-UGG---CAAC--------AC-AA-----GU--------------------GAAC-U-CGGA-GAA
GCCAGCG-------------GGAGUU-CCGGGAAGA-GU------UCUCU-------------U-UUCUUU-
UUAACC-GCC--C----AG-C-------------------GA-----CC----------------CUGG-AA
G-C-UG--------------------UUUAG-CAGGAGAUAG----------GGUGGC----AUG-GC-GGG-
GAA--AG-----------------------------------------------------------------
----CGCCGCA-UCUC-UGCGGCGU-C-CGGUGCA-CUCCCGACGGU-CCGUGAAA-A-UCC-GAGGGAAAG
-A---------------------------------------------CU-----UAUUCGCA-CC-CCUGGU
CGUACUCAUAACCGCA------------------------------UCAGGUC--UCCAAGGU-GAAAAGCC
UCUAGU-C-GA-UAGA-AUAAUGUA-G-GUAAGGGAAGUCGGC-AAAUUAGAUCCGUAACUUCGG-GAUAAG
GAUUG-GCUC-U-AA--GGGUUG-A----GUGGU-UG------GAGC----UGG---------C--GCU---
-CCGAGCCCGGU-GACUGCGG---GGGACUACUGCGGACCCCUCACGGGGACUGCGG---------CGG---
----------------------------------------------------------------------AC
CCCU-----------------GCG------------------------------------------------
-----------------------------------------------------------------------G
GCGC--------------C---------------------------------GG--AG--------------
---------------------------------------------------------------GACG-----
---AG---------------------------UU---------ACGGU--C---------------------
-G------------------GG--C-UU---GCCGCGG----------------------------------
---GAG----------UGC-----GCUUGGGU----------------------------------------
----------------------------------------------------------------GGCC----
-------------------------------------------------------------UCACGGCCGCC
CGGGCGUGCCGCGGU--------------------------------GCG----------------------
------------------------------------------------------------------------
----GC-----------------------UU------------UCCUCC-ACA--AAGAACAAC--CAACUU
AGA--ACUGGAGCGGACAAAGGGAAUCCGA-----CUGUUUAAUUAAAACAAAGCAUUGCGAUGGUUCG---
-AA-A-------------GAAUGUUG-ACGCAAUGUGA-UUUCUGCCCAGUGCUCUG-AAUG-UCAAAGUGG
A--------------GAAA--------------UUCAAUGAAGCGC--GGGUAAAC-GGCGGGAGUAACUAU
GACUCUCUUAAGGUAGCCAAAUGCCUCGUCAUC-UAAUUAGUGACGCGCAUGAAUGGAUUAACGAGA--UUC
CCACUGUCCCUAUC----UAC-UAUCUAGCGAAACC---AC---AGCCAAGGGAACGGGCUUGGAAAAAUCA
GCGGGGAAAGAAGA-CCCUGUUGAGC-UUGACU--CUAGUUU-GACUUUGUGAACUGACUUGAGAGGUGUAG
AAU-AAGUGGGAGA------------------------------------------------------UGCA
----------------------------------------------------AGUC-GA-CGAUGAAAUA-C
CACUACUCUUGACGUCAGUUUACUUAUUU-CG-------U-UA-C-CUAGA------AC----------CG-
------UGGCCCCGU-------UUAC---GCGGGU--AGCCAU-----UUGUCUUCU---------------
-----------GGAC--------------------------------------UUAA--GCGCCUGGCACUU
------------------------------------------------------------------------
---GUG---------------------------CCU-CGG--------------------------------
----------GCG------CGA------------------------------UCUA----CG----------
------------------------------------------------CGAG-----AGACA-UUGUCAGAU
GGGGAGUUUGGCUGGGGCGGUACAUCUGUUA-AACGAUAAC-ACAGGUGUCCU-AAGAGAAGCUCAAUGAGA
ACAGAAAUCUCAUGUA-GAACA-AAAGGGUAA-AAGCUUCUUUGAUU-UUGA--------------------
--------------UUUUCA--GUACGAAUACAAACUGUGAAA-GCAUGGCCUAUCGAUCCUUU--------
---AG-C-----CUUGGAGG---------------------UUCUCAAGCUAGAGGUGUCAG-AA--AAGUU
ACCACAGGG-AUAACUGGCUUGUGGCAGCCAAGCGUUCAUAGCGACGUUGCUUUUUGAUUCUUCGAUGUCGG
```

```
CUCUUCCUAUCAUGCGCCCGCA-GCAGAGCGCAAGUGUCGGAUUGUUCAUCCACCAACAGGGAACGUGAGCU
GGGUUUAGACCGUCGUGAGACAGGUUAGUUUUACCCUACU--------------------G-AUG-------
-------------------------------AAG-U-CGUCACCGCGACAGU-AAU-UCAAUUUAGUACGAG
AGGAA-CCGUUGAUUCGCAUACU-UGGCAAA-UUCGGUUGAUU-GAAA-CGUCAA-U-GCUGAGAAGCUAUC
AUGCG-CAGGAGUAAGGCUGAACGCC-UC--UAAGCCCGAACCCAGGCUGG-----CA-AGCGGUGAU-AUG
UUGCUCCC-UCUGUGUU-------------------------------------------------------
------------------------------------------------------------------------
------------------------------------------------------------------------
---------------------ACAC-----------------------------------------------
--------------------------------GGAUA-----------------------------------
----------AGA-----AUA--GG--------------C--------------------------------
-----ACU--UGUGCCCA---AGA--UCAUAACAUGGCCGGACGAAGCCAC----GAAA--GGCGGACCAGG
CC------------------------------------G-----------AAUGACA-AGU----------U
C-----------------GAAUAUG---------------------GCAGGGAGAC-----AAA------UC
CUUUGCAGACGACUUAAA-U---GGAACGGAGG-GUUGUAAG-CAUGA--------GAGUGUG-CC------
----UUG--UGCCACGAUCUGCUGAGACUUAG-CUCC-UG-UUCUU---CGAUUU-----------------
------------------------------------------------------------------------
------------------------------------------------------------------------
------------------------------------------------------------------------
-------------------------------------------------
>LSUe_420245_LEIBR/1-4081
-----------------------------------------ACGUGUCGC-GAUGGAUG-A-CUUGGCUUCC
U-A---UUUCGUUGAAGAACGCAG--UAAAGUGCGAUAAGUGGUAUCAAU--U---GCA--GAA-UC--AUU
CAAUUACCGAAUCUUUGAACGCAAAC-G--------GC--------GC----AUGG-G-AGAAGCUC---UA
U-U-------GUGUCAU--CC----C----C---GUG----C-A-UGCC-AUAU--U-CUCAGUGUCG-AGU
-------------------------------------------U------------U--ACAGAC-CUGA-G
U-G--------------------UG-G-CA-GGACUACCCGCCGAACUUAAGCAUAUUACUCAGCG--GAGG
AAAAGAAAACAA--CCGUGAUUCUCUCA--GUGAG-CGGCGAGCGAAGAGGGA-ACGA-ACUCGUUGCCGAA
UCGG-GUCUUAC-------AA------------G---GCC--UUGAGUUGUGGCA--CAA------------
--AU-AUCA-UUU-C--GUGG-------------------------UUGGU--GCA-----------GG---
--------AGUGG--CGA-------AAUUCA-AGCG-CAAAGCAAACCCGUUGCUGAA------UACAACCC
UUCAUGUGAG-UA-UUGAG-------CCAAAGAAGGUGUUAGCCCAUUG---------AGC---C----AUA
AACCU-GA--GCGCCUCAUG----------------AACU-GUAUU--AAGCGAGUAGCACUGUUUG---GG
AAUGCAGUGUCAAGU-UGGCAGGUAUU-UGUCUGCCAAAGUUAAAUACAGAGUA-GGA-AGACCGAUAGC-G
AAC-A-AGUAGCGUGAGCGAAAGUUUGAAA-AGCAC-UUUGG-AAAGAG-A-GUGAAAUAGAACCUGAAGUC
G-UGACAAG--A-CAA-UG-GAAG---UACCUCCAUUUCAA-------------------------------
------------------------------------------------------------------------
------------------------------------------------------------------------
------------------------------------------------------------------------
------------------------------------------------------------------------
------------------------------------------------------------------------
----------------------------------------------------------------------C-
---GG-------------------------------------------------------------------
--------------------CCAGA------------------ACCAGCCCUACUGCAU----UAUCCUGUC
GGGGGUCAAAA---------------------------------------------------U--C-CG---
```

```
------------------------------------------------------------------------
------------GCGGGCAACGAAGUGCAAGAAUCUUGG------GUCUUGGUG------------------
------------------------------------------------------------------------
------------------------------------------------------------------------
------------------------------------------------------------------------
----------------------------------------------------------A-------------
-----------------------------------------------AAUGAG-------------------
----------------------CGGAGUA---UUUGAGCU--------------------------------
------------------------------------------------GUGCA----CGGCGC-----A--A
--------GCCGCACGC-ACU---GG----------UUUC------UUCUCCGUG-----------------
---------------------AUG--------------------------------------------CU--
------------------------------------------------------------------------
----------------CCAUGUUGGCGA-------------UGGAAAUGG--G---GUGCCCCCACCCGUCU
UGAAACACGGACCAAGGAGUCAA-A-CAGACGCGCAA-GGGAGGAG--A-UAUG-----GUUGC-----CAG
U-AC--UUCU--CUCGUACUGAAAG-GGA-----------------GAUGCAA-------AAUGCAUCUGUG
GUGU-GCACACG----U----------------------------------------------CA--------
AAU---------------------------------------------------------------------
-------------------CCG--------------------------------------UUU-GAUUU-GC
CCAACA----CCGACCG----------------------GCCC-----------------UCCGUGGG-UU-
GCG-UU----------------------GGAGUGUGCCUGUUUG-GACCCGAAAGG-UGGUGAACUAUGCCU
GAACAAGGUGAAGCCCGG-GGAAAC-CCAGGUGGAGGCCUGU--AGCGAUGCUGACGUGCAAAUCGC-UCGU
AUGAUUUGGGUAUCGGAGCGAAAGACUCAUCGAACCACC-UAGUAGCUGGUUCACAUCGAAG-UUUCCCUCA
GGAUAGCUGGUG-CUAG-------------------------------------------------------
---------------------UAG-----AAGUAUUGUGCGGUAAAGCAAAUGAUU-AGA-GGCAUUGG-UG
U-U-CUUAG-AAUA-U-CGACCUAUUCUCAAACUUUAAAUGUGCAAACA-----------------------
------------------AACCGUGC-----CUUAGC--CAAC--U--GCAAG-------------------
GC---CGGA-AAGAAAAGA-AU-U--AG----A--GGCACCA-AGUGGGC-C-UUCU-U-CGG-UAAGCGGA
GC-A-GGCGAUGC-GGAAUG---AACCGCUUAGGG-A-UAUGUGGCGUCAAAACUUAU-GGG-CU-CAUUCU
-C-GAUACGU-GAAAAGGGUGUUGGUGGAUAUGGACAGUUGGACGGUGGCCAUGGAAGUCGGCA-UCCGCUA
AGGAGUGUGUAACAACUCACCAACC-----GAAUCCACCGGCCCCGAAAAU-GGAU-GACGCUUA-AGCCCA
A--UUAGUGAUUGCCCAUUAUUCCUUUGGUAAGG--GCGGA-------------------------------
-----------------------------------------------AA-----------------CAAC--
----UCUUG-----------GAGA------------------------------------------------
-------------------------AAGUGUAGCCCU-UCCG-AUGUAGAUGUGGCCUGGAGG---------
-----------U-U-AGGAC-GAAGCUUA-U-GGCGUGAGCC--UAAGAUGGACCG-G-CCU-CUAGUGCAG
AUCU-UGGUUG-GCGUA-GCAAAG---AUCU-AACGGA--GAUAUACUCAACAUGCAACGUU-GGA-UACUG
GAGCGGGG-AAGGAUUUC-GUGCCAACG-GCACUCGUACACGAGUUGUUCGG--AAACUGAGCACAACGUU-
A-U-AUCGUUUU-GUUAGGAAAGUGAAGGUG----UGUCGGCGGCAG---CUGCUCUC-----------GGG
UA---------------GCUGUUCG--ACUGGCCA---UA-ACUGA-A-AAGGG-GCAACA---GAGA---A
CC----------------------------------------------------------------------
-------------------------------UGGGAUUAUAUUCCAAAAAGAAAUUGCAUGUGGGU------
--AGUGGUAA---UGCG--------AA-AC-----AC--------------------UUGC-C-AGGU-AAC
AAAUCA--------------AUCC-U-CCCCCGGUG-AG-C----UUUCU-------------U-UUCACC-
AUAAUC-CAC-AU-----C-U-------------------CC-----GG----------------CUUU-GC
U-G-GGCUUGGGCCUUUUUACUUCUCGCGU-UGUUCGGAGC----------GGGGGC----UCA-AG-AUU-
```

```
GAA--AA----------------------------------------------------------------A
--UGCAGCUCU-CCCUACGUACUGU-C-A-UUGUU-GUGAGUUCUGC-GCAUUAAA-G-CAA-AAACCUGGG
-G----------------------------------------------------UGUCCCAA-CU-GCAGAC
CGUACUCAUCACCGCA------------------------------UCAGGUC--CCCAAGCA-UAGAAUGC
UCUGGU-A-CA-UAGA-GAAAGGUA-CACUCAGGGAAGUCGGC-AAAAUAGAUGCGUAAGUUCGC-AAGAAG
CAUUG-GCUC-U-GA-GGGCAAA-G----UCAGA-GA------AAACCG--GAA-GGCCAGUG---UCC-UC
UCGC--------AUGCAGCC----GU----------------------------UCU--------UCAG---
--------------------------------------------------------------------AUG-
-UU-----------------GUUU------------------------------------------------
-------------------------------------------------------UGGCGCUCG------CC
GCCUU-------------A---------------------------------GC--AC--------------
---------------------------------------------------------------GCUGA-UA-
--------------------------------GC-------GACUGUAC---------------------UG
-U-CG--------UCUG--UGAAGG-CUA--CCUU-CCCGAGGC---GUC----GUAACC------------
---CGG----CG-U--CUCA----CUCUGGCAC---GGG---------------------------------
-----CC---------------------------------------------------------GGUC----
----------------------------------------------------UUGGC--GGGU---------
---------------U---------------------------CACGCUC---G------------------
------------------------------------------------------------------------
----UCUU---------------------GUG-----------CCGCGU-CCC--ACAAAUUGC--CAACUC
AGA--ACUGCUUACGGC--GGGGAAUCCAA-----CUGUAUAAUUAAAACAUAGGUUUGUGAUGCAUCC---
-AA-G------------UGGUGUAUA-UCGCAAACUGA-UUUCUGCCCAGUGCUCUG-AAUG-UCAACGUGA
C--------------GAGA--------------UUCACCGACGCGC--GGGUAAAC-GGCGGGAGUAACUAU
GACUCUCUUAAGGUAGCCAAAUGCCUCGUCUUC-CAAUUAGAGACGCGCAUGAAUGGAUUAAUGAGA--UUC
CCUCUGUCCCGAGU----UAC-UAUCUAGCGAAACG---AC---AGUCAAGGGAACGGACUUGAAGGGCUAA
GCGGGGAAAGAAGA-CCCUGUUGAGU-UUGACU--CCAGUCU-GGCUCUGUGCGGCGACAUCUGAGGUGUAG
UAU-AGGUGGAAGC------------------------------------------------------GCAA
----------------------------------------------------G--C-GC-AAAUGAAAUA-C
CACCACUCGGAACGUUGCUUCACUUAUCG-AA-------U-GA-A-GAGAC-CAAUGGGUUAA------GCG
------UA--GU-CU--------UCG---GGCU-A--UG-------------CACCG---UCU-------AG
-GUUU-----GGGAGUAAUUUAUGGCGGUA-----------------ACCCCGUCCC-CAC---------GG
------------------GAAAGG-GU---GG---------------------UGUGCUUUUUCUGCGU--G
C--GUCUUCAC--------------GGGCGUCGC---GCGGGCCCCGCUCUUUUUUAUUGCGUUUCCUUCGG
GGGGCGUGACUCG------UGGGUA---------------------------CCCAA---CCGUGCUGUAA-
ACUGAACCAACGGGAUUGC-----CUCGU-GCAGUUCCGGAGUCUUGUUUCG-----AGACAUCUGCCAGAU
GGGGAGUUUGGCUGGGGCGGCAUAUCUGUUA-CACGACAAC-GCAGGUGUCCU-AAGGCGAGCUCAGUGGGA
ACAGAAAUCUCACGUA-GAACA-CAAGCGUAA-AAGCUUGCUUGAUUAACGA--------------------
--------------UUUCCA--GUACGAAUCGAGACUGCGAAA-GCAAGGCCAG-CGAUCCUUUGCGAAGAA
UAGGA-A-----AUAUGAACA--------------------AA-UCAUACCAGAGGUGUCAG-AA--UAAUU
ACCACAGGG-AUAACUGGCUUGUGGCGGCCAAGCGUUCAUAGCGACGUCGCUUUUUGAUCCUUCGAUGUCGG
CUCUUCCUAACCUAGCGCCGCA-GAAGACGCUAAGGGUUGGAUUGUUCACCCACUGACAGGGAACGUGAGCU
GGGUUUAGACCGUCGUGAGACAGGUUGGUGAGAUUGUGAA--------------------A-GGGAU-----
-----------------------------CUCG--C-AGGUAUCGUGAGGGA-AGUAUUGGGGUAGUACGAG
AGGAA-CUCCCAUGCCGUGCCUC-UAGU-UUCUGGGGUUUGUC-GAAC-GGCAAG-U-GCCCCGAAGCCAUC
GCACG-GUGGUUCUCGGCUGAACGCC-UC--UAAGCCAGAAGCCAAUCCCAAG---AC--CAGAUGCC-CAC
```

```
CUCCCAUG-CCGUGCCU-------------------------------------------------------
------------------------------------------------------------------------
------------------------------------------------------------------------
---------------------CUAGUUUC-------------------------------------------
-------------------------------UGGGGU------------UUGUCGAA---------------
------CG--GC-------AAGUGC-------------CCCG------------------AAGCCA------
----UCGCACGG--------------UGGUUCUCGGCUGAACGCCUCUAA-GCCAGAAGCCAAUCCCAAGAC
CA---------------------------------GAUGCCCACUCAUCGAAUCGCCACCUACAAGACUGGA
GCUUGCUCCCUCGAAGGCGCCAAGUAUAUUCAUGAUCACAAGACAGU-CCCUCUCC-----AAA------CG
AGAGAACAUGCAUGGGCU-GGC-AUGAGCGGCA-UGCUUCAC-UUCGGUGGGGCUCGAGGGGC-AU------
----UUA--CGUCCCGAGGCGCUGAACCUUGA-GGCC-UGAAAUUUC--AUGCUCA-G--------------
------------------------------------------------------------------------
------------------------------------------------------------------------
------------------------------------------------------------------------
-------------------------------------------------
>LSUe_5911_TETTH/1-3491
------------------------------A----GA--AAAUUUUCAAC-GGUGGAUA-U-CUAGGU-UCC
C-G---UGACGAUGAAGAACGCAG--CGAAAUGCGAUACGCAAUGCGAAU--U---GCA--GAA-C---CGC
GAGUCAUCAGAUCUUUGAACGCAAGU-G--------GU--------GG----AGGU-G------------UA
A-A-----------------A----A----C---CUU----C-A-UGUU-UGU-----UUCAGUGUGG-AA-
-----------------------------------------------------------CUACAC-CUGA-A
A-C--------------------AA-G-CA-AGAUUACCCGCUGAACUUAAGCAUAUCAGUAAGCG--GAGG
AAAAGAAACUAA--CUAGGAUAGCCCCA--GU-AA-UGGCGAAUGAACAGGCU-A-AA-GCUCAAAGUGAAA
AUCU-GAAGUGGUC----AAC------------A---CAA--CAGAAUUGUAAUC--UA-------------
--AA-GGGU-CAA-C-----C-------------------------UGAAA--CUA-----------AG---
CU------CCUCU--CAU-------AAGUUC-CUUG------------------------------------
-------GAA-CA-GGACG-------UCAAAGAGGGUGACAACCCCGUA---------GUC---G----GAG
AGGA--AG--GCUGG-UGUAA---------------GGGA-GAUUU--CAAAGAGUCGGGUUGUUUG---GG
AUUGCAGCCCUAA-G-UGGGAGAUAAA-CUUCUUCUAAAGCUAAAUAUACA--C-GGG-AGACCGAUAGC-G
AAC-A-AGUACUGCGAAGGAAAG-AUGAAA-AGAAC-UUUGA-AAAGAG-G-GCUAA--AAGACUUGAAACC
G-UUGAGAAG--G-AAG-CU-GUAG---AAGA-GCAAUAAACUGGACGGCGCAUAAG-GGGGAA--------
-------------------------------------------GUACUAAU---------CAC--UGCA---
------G------------------AGU--------------------CGAU--------------------
------------------------------------------------------------ACGUAAAA----
-----------------------GG-----------------------------------------------
----------------------------------UCG----------AUG-------------------AGU
A----AGGAAA------------UGG--U--AC----------A---------------GAACUUGCUACGC
C--GG----------------------------------------------------------------UUC
AGA-----------------UUGAAGGAGUCACC---UGAGAUCGGGCA-----------------------
-----------------------------------------------------------------G-CAA--
------------------------------------------------------------------------
---------------UGCAGAUCAAAAGGAAAACUUCAA------ACUGGACU-------------------
------------------------------------------------------------------------
------------------------------------------------------------------------
------------------------------------------------------------------------
```

```
------------------------------------------------------------------------
-------------------------------------------------GAGGGG----C------------
------C------------UAAGGGCGA--------------------------------------------
------------------------------------------------------------------------
---------------------------------------U------UUUGUC--------------------
------------------------------------------------------------------------
------------------------------------------------------------------------
-------------------------------------------AAAAUGG--C---UUCUACUGACCCGUCU
UGAAACACGGACCAAGGAGUCUA-U-CAAUUAAGCGA-GUGAUAGG--G-UG--------GAAA-------A
A-CC--CGUC--CGCGAAACGAAAG-UGA-----------------GUA-------------------CAAG
GUGC-CAAGCC---------------------------------------------------GC--------
AA----------------------------------------------------------------------
-------------------GGU--------------------------------------AGCAGCAUC-AC
CCGACC----UU-GAGU----------------------CUCCG----------------ACGAAGGGUUC-
GAG-GA----------------------AGAGCUUAAUUGUUAG-GACCCGAAAGA-UGGUGAACUACGCUU
GAAUAGGGUGAAGCCAGG-GGAAAC-UCUGGUGGAAGCUCGU--AGCGAUACUGACGUGCAAAUCGU-UCGU
CAAAUUUGAGUGUAGGGGCGAAAGACUAAUCGAACCAUC-UAGUAGCUGGUUCCCUCCGAAG-UUUCUCUCA
GGAUAGCAAGAG-CAAG-------------------------------------------------------
--------------------UACG-----CAGUUUUAUUAGGUAAAGCGAAUGAUU-AGA-GGACUCGG-GG
U-U-CCAAG-AAUC-U-CGACCUAUUCUCAAACUUUAAAUUGGUAAGAG-CC--------------------
------------------GCGGA-GU-----UUA-CU--UAAA--U--GAA---------------------
CU---CUCG-G---GAAGA-AU-G--CA----G--UGCUCUU-AGUGGGC-C-AUUU-U-UGG-UAAGCAGA
AC-U-GGCGAUGA-GGGAUG---AACCUAACGUUG-A-GAUAAGGCGCCCAAAU--GC-ACG-CU-CA---U
-C-AGAUACC-ACAAAAGGUGUUGGUUCAUACGGACAGCAGGACGGUGGCUAUGGAAGUUAGAA-UCCGCUA
AGGAGUGUGUAACAACUCACCUGCC-----GAAUGAACUAGCCCUGAAAAU-GGAU-GGCGCUGA-AGCGUG
U----------UGCCGAU-ACUCAACCAUCAGAG--CAAA--------------------------------
------------------------------------------------------------------------
------------------------------------------------------------------------
---------------------------UGCGAGGCUU-UGAU-GAGUAGGAGGGC-GUGAUCG---------
-----------U-U-GCCUA-GAAGUAUU-G-GGCGUGAGCC--U-AUAUGGAGCA-G-CGA-UUAGUGCAG
AUCU-UGGUGG-UAGUA-GCAAAU---AUUC-AAAUGA--G---------------AACUUU-GAA-GACCG
AAGUGGAG-AAGGGUUCC-AUGAGAACA-GCAAUUGUUCAUGGGUUACUCGA--UC-CUAAGACAUAGGUU-
A-A-CUCC-----UUGCAAU--------ACAA----GAA-----GAC----AUU------------------
-----------------CGUUUUC--------GU----UG-UC-AA-A-AGGGA-AUGAGGUU-AAUA-UUC
CUCAA----------------------------------GCUGGA---CGUG--------G-UA--------
U----------------AG-----------------------------------------------------
----AG-CGG---UAAC--------GC-AA-----AG--------------------AAAC-C-CGGA-GAC
GUCAGCA-------------GGUGUC-ACUGGAAGA-GU------UAUCU-------------U-UUCUUU-
UUAACA-UAC--U-----A-U-------------------GG-----CC----------------AUGA-AA
U-U-GG--------------------AUUAU-CCAGAGAUAU---------CGGCUGU----AUG-UA-UGG-
CAG--AG-----------------------------------------------------------------
----CAGCUCA-CCCUAAGAGCUGU-C-AGUUGCG-CACCUGAUGAC-CCUUGAAA-A-UCU-GGGGGAGAC
-A----------------------------------------------------UAAUUUCA-CG-CCAGUU
CGUACCCAUAACCGCA------------------------------UCAGGUC--UCCAAGGU-UAGCAGCC
UCUGGU-C-CA-UAGA-ACAAUGUA-G-AUAAGGGAAGUCGGC-AAAUUGGAUCCGUAACUUCGG-GAUAAG
```

GAUUG-GCUC-U-GA--GGAUCG-G----GUAUA-AA------GGCUU---UGU------------------
--AA------UG----AUAUCCAAGC----------------------------UUG--------UUUG---
------------------------------------------------------------------------
-UUA-----------------GUG------------------------------------------------
---------------------------------------------------------UG-----------GC
AACA--------------U---------------------------------GC--UG--------------
---------------------------------------------------------------AUAG-----
------------------------------------------------------------------------
--------------------ACUUG-CGAACGAUG-------------------------------------
-------------AAUUUG-----CA-AGGUA----------------------------------------
----------------------------------------------------------------GGUU----
------------------------------------------------------------------------
-----------------------------------------------UCG---G------------------
------------------------------------------------------------------------
----CC-----------------------GU------------CUUUAU-ACA--AUUAACGAU--CAACUC
AGA--ACUGAAGCGGACAAAGGUAAUCCGA-----CUGUUUAAUAAAAACAAAGCAUUGUGACGGCCUC---
-AA-C-------------AGGUGAUG-ACACAAUGUGA-UUUCUGCCCAGUGCUCUG-AAUG-UCAAAGUGA
C--------------GCAA--------------UUCAACCAAGCGC--GGGUAAAC-GGCGGGAGUAACUAU
GACUCUCUUAAGGUAGCCAAAUGCCUCGUCAUC-UAAUUAGUGACGCGCAUGAAUGGAUUAAUGAGA--UUA
CCACUGUCCCUAUC----UAC-UAUCUAGCGAACCC---AC---AGCUAAGGGAACGGGCUUAGAAUAAUCA
GCGGGGAAAGAAGA-CCCUGUUGAGC-UUGACU--CUAGUCU-AACUUUGUGAAAUGGCACGUGGGGUAUAG
CCU-AGGUGGGAGA------------------------------------------------------GCAA
----------------------------------------------------U--C-GA-UCCUGUAAAA-C
CACUACCCACGUAGUCAUUUUGCUUAUUU-CG-------U-GA-A-GAAAA------G-----------AC-
------UG--GU-GC--------AAA---CCAG-U--U-----------------CU---------------
-----------AAAG--------------------------------------UUAA--GG---------UC
------------------------------------------------------------------------
---AUU---------------------------U---AUU--------------------------------
----------GAC------UGA------------------------------UUUU----UG----------
------------------------------------------------CGAA-----AGACA-UGGUUAGGG
GGGGAGUUUGUCUGGGGCGGAAUGCCUGUUA-AACCAUAAC-GCAGGCGUCCU-AAGUGUAGCUCAGUGAGA
ACGGAAAUCUCACGUA-GAACA-AAAGGGUAA-AAGCUACAUUGAUU-UUGA--------------------
--------------UUUUCA--GUAGGAAUACAAACCGCGAAA-GCGUGGCCUAUCGAUCCUUU--------
----A-A-----CUUUACAAG--------------------UU-UUAAGCUAGAGGUGUCAG-AA--AAGUU
ACCACAGGG-AUAACUGGCUUGUGGCAGCCAAGAGUUCAUAUCGACGUUGCUUUUUGAUCCUUCGAUGUCGG
CUCUUCCUAUCAUUGUGAAGCA-GAAUUCACAACGUGUCGGAUUGUUCACCCGCUAAUAGGGAACGUGAGCU
GGGUUUAGACCGUCGUGAGACAGGUUAGUUUUACCCUACU--------------------G-AUG-------
-------------------------------AA--A-CGAUGUUGCGACAGU-AAU-UUAAGUUAGUACGAG
AGGAA-CACUUAAAUCAGAUAAU-UGGU-AAAUACGGUUGUCU-GAAA-AGACAA-U-GCCGUGAAGCUACC
AUCUG-UUGGAUUAUGACUGAAGGCC-UC--UAAGUCAGAAUCCAUGCUGGA----AA--GCAAUGUC-UAA
GUGUGA------------------------------------------------------------------
------------------------------------------------------------------------
------------------------------------------------------------------------
------------------------------------------------------------------------
-------------------------------UGAUAA-----------ACGAAAAA----------------

```
--A-------AUA-----AAA--AU--------------UAA------------------GUUCGAA---A-
-------------------GG-----UAG-AGCG-----GGG-AAGAGC------GAAAA--A-GCUUGACC
U---------------------------UAA--C-UGCU-----------AAUCGUA-AUU----------C
C-----------------AAAUUAU---------------------C-AUCUACGU-----AAA------UC
UUUUGUAGACGACUUAAC--AU-GGAAC-GGGU-AUUGUAAG-CAUGA--------GAGUAGA-AU------
----U----UCCUACGAUCUGCUGAGAUUCAG-CCCG-UCUCCUUA---GAUUUAUCU--------------
------------------------------------------------------------------------
------------------------------------------------------------------------
------------------------------------------------------------------------
-------------------------------------------------
>LSUe_33169_EREGO/1-3547
------------------------------------A--AAACUUUCAAC-AACGGAUC-U-CUUGGU-UCU
C-G---CAUCGAUGAAGAACGCAG--CGAAUUGCGAUAAGUAUUGUGAAU--U---GCA--GAU-UUU-CGU
GAAUCAUCGAAUCUUUGAACGCACAU-U--------GC--------GU----CCUC-U-GG---------UA
U-U------------CC---A----G----G---GGG----C-A-UGCC-UGU-----UUGAGCGUCA-U--
--------------------------------------------------------G--UUUGAC-CUCA-A
AUC--------------------AG-G-UA-GGAUUACCCGCUGAACUUAAGCAUAUCAAUAAGCG--GAGG
AAAAGAAACCAA--CCGGGAUUGCCUUA--GU-AA-CGGCGAGUGAAGCGGCA-A-AA-GCUCAAAUUUGAA
AUCU-GGCGCCU------UCG------------G---CGU--CCGAGUUGUAAUU--UG-------------
--AA-GAAA-GUA-C-----C-------------------------UUGGU--UGC-----------UA---
GU------CCCUG--UCU-------AUGUUC-CUUG------------------------------------
-------GAA-CA-GGACG-------UCAUAGAGGGUGAGAAUCCCGUC---------UGG---C----GGG
GGU---GC--UAGUG-CCAUCUA-------------AGGU-UCUUU--CGACGAGUCGAGUUGUUUG---GG
AAUGCAGCUCUAA-G-UGGGUGGUAAA-UUCCAUCUAAAGCUAAAUAUUGG--C-GAG-AGACCGAUAGC-G
AAC-A-AGUACAGUGAUGGAAAG-AUGAAA-AGAAC-UUUGA-AAAGAG-A-GUGAAAAAGUACGUGAAAUU
G-UUGAAAGG--G-AAG-GG-CAUU---UGAU-CAG-ACAUG--------------------GU--------
-------------------------------------------GUUUGCG--G------CCC---------
------------------------------------------------UCGC--------------------
------------------------------------------------------------UCCUUGUG----
-----------------------GG-----------------------------------------------
----------------------------------UGG-----------------------------------
------GGUCA------CC---GCAGUUC--AC----------UG---------------GG-CCAGCAUC-
---AG----------------------------------UUUUGGUGGC-----------------AGCAGA
AAC-----------------CCUUAGGAA--------UG-------UGACUUUCUC----------------
--------------------------------------------------------------U--U-CG---
------------------------------------------------------------------------
------------------------GAGGAAGUGUUAUAG------CC---------UAAGGUUAUACUGCCA
ACCGGGACU---------------------------------------------------------------
------------------------------------------------------------------------
------------------------------------------------------------------------
------------------------------------------------------------------------
-------------------------------------------------GAGGAC-----------------
--------------------UGCGGCUU--------------------------------------------
------------------------------------------------------------------------
-----------------------------------------------CGGCCAAG-----------------
```

------------------------------------------------------------------------
------------------------------------------------------------------------
-----------------GAUGCUGGCAU-----------------AAUGG--U---UAAAUGCCGCCCGUCU
UGAAACACGGACCAAGGAGUCUA-A-CGUCUAUGCGA-GUGUUUGG--G-U--------GUAAA--------
A-CC--CGUA--CGCGUAAUGAAAG-UGA-----------------ACGU--------------------AG
GUGA-GGGCCUC--------------------------------------------------UU--------
UAG---------------------------------------------------------------------
-------------------AGG--------------------------------------UGCAUCAUC-GA
CCGAUC----CU-GAUG----------------------UCUU-----------------CGGAUGGAUUU-
GAG-UA----------------------AGAGCAUAGCUGUUGG-GACCCGAAAGA-UGGUGAACUAUGCCU
GAAUAGGGUGAAGCCAGA-GGAAAC-UCUGGUGGAGGCUCGU--AGCGGUUCUGACGUGCAAAUCGA-UCGU
CGAAUUUGGGUAUAGGGGCGAAAGACUAAUCGAACCAUC-UAGUAGCUGGUUCCUGCCGAAG-UUUCCCUCA
GGAUAGCAGAAG-CUCG-------------------------------------------------------
---------------------UAU-----CAGUUUUAUGAGGUAAAGCGAAUGAUU-AGA-GGUACCGG-GG
U-U-GAAAU-GACC-U-UGACCUAUUCUCAAACUUUAAAUAUGUAAGAA-----------------------
------------------GUCCU-UG-----UUG-CU--UAAU--U--GAA---------------------
CG---UGGA-C--AUAUGA-AU-G--AA----G--AGCUUUU-AGUGGGC-C-AUUU-U-UGG-UAAGCAGA
AC-U-GGCGAUGC-GGGAUG---AACCGAACGUGG-A-GUUAAGGUGCCAGAAU--AC-ACG-CU-CA---U
-C-AGACACC-ACAAAAGGUGUUAGUUCAUCUAGACAGCCGGACGGUGGCCAUGGAAGUCGGAA-UCCGCUA
AGGAGUGUGUAACAACUCACCGGCC-----GAAUGAACUAGCCCUGAAAAU-GGAU-GGCGCUCA-AGCGUG
U----------UACCUAU-ACUCCACCGUCAGGG--CAAA---------------------------------
------------------------------------------------------------------------
------------------------------------------------------------------------
---------------------------UAUGACGCCC-UGAC-GAGUAGGCAGGC-GUGGAGG---------
-----------U-CAGUGAC-GAAGCCUA-G-GCUGUAAAGC--U-GGGUCGAACG-G-CCU-CUAGUGCAG
AUCU-UGGUGG-UAGUA-GCAAAU---AUUC-AAAUGA--G---------------AACUUU-GAA-GACUG
AAGUGGGG-AAAGGUUCC-ACGUCAACA-GCAGUUGGACGUGGGUUAGUCGA--UC-CUAAGAGAUGGGGA-
A-G-CUCC-----GUUCAA---------AG------GCC-----UGA---UUU-------------------
-----------------UCCAGGC-------CAC----CA-UC-GA-A-AGGGA-AUCCGGUU-AAUA-UUC
CGGAA----------------------------------CCUGGA---UAUG--------G-AU--------
U----------------CU-----------------------------------------------------
---UCA-CGG---CAAC--------GU-AA-----CU--------------------GAGU-G-UGGA-GAC
GUCGGCG-------------UGAGCC-CUGGGAGGA-GU------UAUCU-------------U-UUCUUC-
UUAACA-GCU--U-----A-U-------------------CA-----CC----------------CCGG-AA
U-U-GG--------------------UUUAU-CCGGAGAUGG----------GGUCUU----AUG-GC-UGG-
AAG--AG-----------------------------------------------------------------
-----CCCAGC-AUCUAUGCUGGGU-C-CGGUGCG-CUUACGACGGC-CCUUGAAA-A-UCC-ACAGGAAGG
-A---------------------------------------------AU-----AGUUUUCA-UG-CCAGGU
CGUACUGAUAACCGCA------------------------------GCAGGUC--UCCAAGGU-GAACAGCC
UCUAGU-U-GA-UAGA-AUAAUGUA-G-AUAAGGGAAGUCGGC-AAAAUAGAUCCGUAACUUCGG-GAUAAG
GAUUG-GCUC-U-AA--GGAUCG-G----GUAGU-GA------GGGCCUU-GGU--CAGAC-GC--GGC---
-AAG-------------UGUG----C----------------------------UUG---------UGG--U
C---------------------------------------------------------------------UG
-UCC-----------------UCG------------------------------------------------
---------------------------------------------------------GGGGCUU--------

```
GCUC--------------CU--------------------------------GG--GG--------------
---------------------------------------------------------------ACGG-----
------------------------------------------------------------------------
-A-CU---------------GCUUG-CGU--GCUC-UG----------------------------------
---UCG----UA-GA-CGG-----CCUUGGUA----------------------------------------
----------------------------------------------------------------GACC----
------------------------------------------------------------AUC---------
---------------U-------------------------------CUG---G------------------
------------------------------------------------------------------------
----UC-----------------------GUCG----------CUUGCU-ACA--AUUAACGAU--CAACUU
AGA--ACUGGUACGGACAAGGGGAAUCUGA-----CUGUCUAAUUAAAACAUAGCAUUGCGAUGGUCAG---
-AA-A------------GUGAUGUUG-ACGCAAUGUGA-UUUCUGCCCAGUGCUCUG-AAUG-UCAAAGUGA
A--------------GAAA--------------UUCAACCAAGCGC--GGGUAAAC-GGCGGGAGUAACUAU
GACUCUCUUAAGGUAGCCAAAUGCCUCGUCAUC-UAAUUAGUGACGCGCAUGAAUGGAUUAACGAGA--UUC
CCACUGUCCCUAUC----UAC-UAUCUAGCGAAACC---AC---AGCCAAGGGAACGGGCUUGGCAGAAUCA
GCGGGGAAAGAAGA-CCCUGUUGAGC-UUGACU--CUAGUUU-GACAUUGUGAAGAGACAUAGAGGGUGUAG
AAU-AAGUGGGAGC------------------------------------------------------UUCG
----------------------------------------------------G--C-GC-CAGUGAAAUA-C
CACUACCUUUAUAGUUUCUUUACUUAUUC-AA-------U-UA-A-GCGGA------G-----------CU-
------GG--AA-UU--------CAU---UUUC-C--AC-------------CUUCU---------------
-----------AGCA-------------------------------------UUUAA--AG---------UC
---------------------------------------------------------------CU-------
---AUAC--------------------------------G--------------------------------
----------GGC------UGA------------------------------UCCG----GG----------
------------------------------------------------UUGA-----AGACA-UUGUCAGGU
GGGGAGUUUGGCUGGGGCGGCACAUCUGUUA-AACGAUAAC-GCAGAUGUCCU-AAGGGGGACUCAUGGAGA
ACAGAAAUCUCCAGUA-GAACA-AAAGGGUAA-AAGUCCCCUUGAUU-UUGA--------------------
--------------UUUUCA--GUGUGAAUACAAACCAUGAAA-GUGUGGCCUAUCGAUCCUUU--------
---AG-U-----UCCUCGGAG--------------------UU-UGAGGCUAGAGGUGCCAG-AA--AAGUU
ACCACAGGG-AUAACUGGCUUGUGGCAGUCAAGCGUUCAUAGCGACAUUGCUUUUUGAUUCUUCGAUGUCGG
CUCUUCCUAUCAUACCGAAGCA-GAAUUCGGUAAGCGUUGGAUUGUUCACCCACUAAUAGGGAACGUGAGCU
GGGUUUAGACCGUCGUGAGACAGGUUAGUUUACCCUACU---------------------G-AUG-------
-------------------------------AA----UGUUAUCGCAAUAGU-AAU-UGAACUUAGUACGAG
AGGAA-CAGUUCAUUCGGAUAAU-UGGU-UUUUGCGGCUGUCC-GACC-GGGCAU-U-GCCGCGAAGCUACC
AUCCG-CUGGAUUAUGGCUGAACGCC-UC--UAAGUCAGAAUCCAUGCUAGA----AC--GCGAUGAU-UCU
UUUUCUCG-CACAUUAU-------------------------------------------------------
------------------------------------------------------------------------
------------------------------------------------------------------------
---------------------AGAUGGAUACGA-----------AUA-------------------------
-------------------------------AGGUGC-----------------------------------
----UUUU--AG-------CA--UC--------------GCU------------------GAACCAU---A-
------GCAGGC------CGG-----CAACUGGU-----GUU-CAGACG------GAAA-GGUC-UGGGCGC
----------------------------G----U-GCCGGCGG--------AUUGC--AAU----------G
U-----------------CAUACUG---------------------C-GCGAGAGU-----AAA------UC
AUUUGUACACGACUUAGA-UGU-ACAACAGGGU-AUUGUAAG-CAGUA--------GAGUAGC-CU------
```

```
----UGU--UGUUACGAUCUGCUGAGAUUAAG-CCUU-CGUUGUCU---GAUUUG--U--------------
------------------------------------------------------------------------
------------------------------------------------------------------------
------------------------------------------------------------------------
-------------------------------------------------
>LSUe_559292_YEAST/1-3554
---------------------------------------AAACUUUCAAC-AACGGAUC-U-CUUGGU-UCU
C-G---CAUCGAUGAAGAACGCAG--CGAAAUGCGAUACGUAAUGUGAAU--U---GCA--GAA-UUC-CGU
GAAUCAUCGAAUCUUUGAACGCACAU-U--------GC--------GC----CCCU-U-GG---------UA
U-U------------CC---A----G----G---GGG----C-A-UGCC-UGU-----UUGAGCGUCA-UUU
--------------------------------------------------------G--UUUGAC-CUCA-A
AUC--------------------AG-G-UA-GGAGUACCCGCUGAACUUAAGCAUAUCAAUAAGCG--GAGG
AAAAGAAACCAA--CCGGGAUUGCCUUA--GU-AA-CGGCGAGUGAAGCGGCA-A-AA-GCUCAAAUUUGAA
AUCU-GGUACCU------UCG------------G---UGC--CCGAGUUGUAAUU--UG-------------
--GA-GAGG-GCA-A-----C-------------------------UUUGG--GGC-----------CG---
UU------CCUUG--UCU-------AUGUUC-CUUG------------------------------------
-------GAA-CA-GGACG-------UCAUAGAGGGUGAGAAUCCCGUG---------UGG---C----GAG
GAG---UG--CGGUU-CUUUGUA-------------AAGU-GCCUU--CGAAGAGUCGAGUUGUUUG---GG
AAUGCAGCUCUAA-G-UGGGUGGUAAA-UUCCAUCUAAAGCUAAAUAUUGG--C-GAG-AGACCGAUAGC-G
AAC-A-AGUACAGUGAUGGAAAG-AUGAAA-AGAAC-UUUGA-AAAGAG-A-GUGAAAAAGUACGUGAAAUU
G-UUGAAAGG--G-AAG-GG-CAUU---UGAU-CAG-ACAUG--------------------GU--------
-------------------------------------------GUUUUGUG--C------CCU---------
------------------------------------------------CUGC--------------------
------------------------------------------------------------UCCUUGUG----
-----------------------GG-----------------------------------------------
----------------------------------UAG-----------------------------------
------GGGAA----UCUC---GCAUUUC--AC----------UG---------------GG-CCAGCAUC-
---AG----------------------------------UUUUGGUGGC-----------------AGGAUA
AAU-----------------CCAUAGGAA--------UG-------UAGCUUGC------------------
--------------------------------------------------------------C--U-CG---
------------------------------------------------------------------------
--------------------------GUAAGUAUUAUAG------CC---------UGUGGGAAUACUGCCA
GCUGGGACU---------------------------------------------------------------
------------------------------------------------------------------------
------------------------------------------------------------------------
------------------------------------------------------------------------
-------------------------------------------------GAGGAC-----------------
--------------------UGCGACGU--------------------------------------------
------------------------------------------------------------------------
-----------------------------------------------AAGUCAAG-----------------
------------------------------------------------------------------------
------------------------------------------------------------------------
-----------------GAUGCUGGCAU-----------------AAUGG--U---UAUAUGCCGCCCGUCU
UGAAACACGGACCAAGGAGUCUA-A-CGUCUAUGCGA-GUGUUUGG--G-U--------GUAAA--------
A-CC--CAUA--CGCGUAAUGAAAG-UGA-----------------ACGU--------------------AG
```

```
GUUG-GGGCCUC--------------------------------------------------GC--------
AAG---------------------------------------------------------------------
-------------------AGG--------------------------------------UGCACAAUC-GA
CCGAUC----CU-GAUG----------------------UCUU-----------------CGGAUGGAUUU-
GAG-UA----------------------AGAGCAUAGCUGUUGG-GACCCGAAAGA-UGGUGAACUAUGCCU
GAAUAGGGUGAAGCCAGA-GGAAAC-UCUGGUGGAGGCUCGU--AGCGGUUCUGACGUGCAAAUCGA-UCGU
CGAAUUUGGGUAUAGGGGCGAAAGACUAAUCGAACCAUC-UAGUAGCUGGUUCCUGCCGAAG-UUUCCCUCA
GGAUAGCAGAAG-CUCG-------------------------------------------------------
---------------------UAU-----CAGUUUUAUGAGGUAAAGCGAAUGAUU-AGA-GGUUCCGG-GG
U-C-GAAAU-GACC-U-UGACCUAUUCUCAAACUUUAAAUAUGUAAGAA-----------------------
------------------GUCCU-UG-----UUA-CU--UAAU--U--GAA---------------------
CG---UGGA-C--AUUUGA-AU-G--AA----G--AGCUUUU-AGUGGGC-C-AUUU-U-UGG-UAAGCAGA
AC-U-GGCGAUGC-GGGAUG---AACCGAACGUAG-A-GUUAAGGUGCCGGAAU--AC-ACG-CU-CA---U
-C-AGACACC-ACAAAAGGUGUUAGUUCAUCUAGACAGCCGGACGGUGGCCAUGGAAGUCGGAA-UCCGCUA
AGGAGUGUGUAACAACUCACCGGCC-----GAAUGAACUAGCCCUGAAAAU-GGAU-GGCGCUCA-AGCGUG
U----------UACCUAU-ACUCUACCGUCAGGG--UUGA--------------------------------
------------------------------------------------------------------------
------------------------------------------------------------------------
---------------------------UAUGAUGCCC-UGAC-GAGUAGGCAGGC-GUGGAGG---------
-----------U-CAGUGAC-GAAGCCUA-G-ACCGUAAGGU--C-GGGUCGAACG-G-CCU-CUAGUGCAG
AUCU-UGGUGG-UAGUA-GCAAAU---AUUC-AAAUGA--G---------------AACUUU-GAA-GACUG
AAGUGGGG-AAAGGUUCC-ACGUCAACA-GCAGUUGGACGUGGGUUAGUCGA--UC-CUAAGAGAUGGGGA-
A-G-CUCC-----GUUUCAA--------AG------GCC-----UGA---UUUU-----------------A
-----------------UGCAGGC-------CAC----CA-UC-GA-A-AGGGA-AUCCGGUU-AAGA-UUC
CGGAA----------------------------------CCUGGA---UAUG--------G-AU--------
U----------------CU-----------------------------------------------------
---UCA-CGG---UAAC--------GU-AA-----CU--------------------GAAU-G-UGGA-GAC
GUCGGCG-------------CGAGCC-CUGGGAGGA-GU------UAUCU-------------U-UUCUUC-
UUAACA-GCU--U-----A-U-------------------CA-----CC----------------CCGG-AA
U-U-GG--------------------UUUAU-CCGGAGAUGG---------GGUCUU----AUG-GC-UGG-
AAG--AG-----------------------------------------------------------------
-----GCCAGC-ACCUUUGCUGGCU-C-CGGUGCG-CUUGUGACGGC-CCGUGAAA-A-UCC-ACAGGAAGG
-A---------------------------------------------AU-----AGUUUUCA-UG-CCAGGU
CGUACUGAUAACCGCA------------------------------GCAGGUC--UCCAAGGU-GAACAGCC
UCUAGU-U-GA-UAGA-AUAAUGUA-G-AUAAGGGAAGUCGGC-AAAAUAGAUCCGUAACUUCGG-GAUAAG
GAUUG-GCUC-U-AA--GGGUCG-G----GUAGU-GA------GGGCCUU-GGU--CAGAC-GC--AGC---
-GGG-------------CGUG----C----------------------------UUG---------UGG--A
C---------------------------------------------------------------------UG
-CUU-----------------GGU------------------------------------------------
---------------------------------------------------------GGGGCUU--------
GCUC--------------UG--------------------------------CU--AG--------------
---------------------------------------------------------------GCGG-----
------------------------------------------------------------------------
-A-CU---------------ACUUG-CGU--GCCU-UG----------------------------------
---UUG----UA-GA-CGG-----CCUUGGUA----------------------------------------
```

```
----------------------------------------------------------------GGUC----
------------------------------------------------------------UCU---------
---------------U------------------------------GUAG---A------------------
------------------------------------------------------------------------
----CC-----------------------GUCG----------CUUGCU-ACA--AUUAACGAU--CAACUU
AGA--ACUGGUACGGACAAGGGGAAUCUGA-----CUGUCUAAUUAAAACAUAGCAUUGCGAUGGUCAG---
-AA-A------------GUGAUGUUG-ACGCAAUGUGA-UUUCUGCCCAGUGCUCUG-AAUG-UCAAAGUGA
A--------------GAAA--------------UUCAACCAAGCGC--GGGUAAAC-GGCGGGAGUAACUAU
GACUCUCUUAAGGUAGCCAAAUGCCUCGUCAUC-UAAUUAGUGACGCGCAUGAAUGGAUUAACGAGA--UUC
CCACUGUCCCUAUC----UAC-UAUCUAGCGAAACC---AC---AGCCAAGGGAACGGGCUUGGCAGAAUCA
GCGGGGAAAGAAGA-CCCUGUUGAGC-UUGACU--CUAGUUU-GACAUUGUGAAGAGACAUAGAGGGUGUAG
AAU-AAGUGGGAGC------------------------------------------------------UUCG
----------------------------------------------------G--C-GC-CAGUGAAAUA-C
CACUACCUUUAUAGUUUCUUUACUUAUUC-AA-------U-GA-A-GCGGA------G-----------CU-
------GG--AA-UU--------CAU---UUUC-C--AC-------------GUUCU---------------
-----------AGCA-------------------------------------UUCAA--GG---------UC
---------------------------------------------------------------CC-------
---AUUC-------------------------------GG--------------------------------
----------GGC------UGA------------------------------UCCG----GG----------
------------------------------------------------UUGA-----AGACA-UUGUCAGGU
GGGGAGUUUGGCUGGGGCGGCACAUCUGUUA-AACGAUAAC-GCAGAUGUCCU-AAGGGGGGCUCAUGGAGA
ACAGAAAUCUCCAGUA-GAACA-AAAGGGUAA-AAGCCCCCUUGAUU-UUGA--------------------
--------------UUUUCA--GUGUGAAUACAAACCAUGAAA-GUGUGGCCUAUCGAUCCUUU--------
---AG-U-----CCCUCGGAA--------------------UU-UGAGGCUAGAGGUGCCAG-AA--AAGUU
ACCACAGGG-AUAACUGGCUUGUGGCAGUCAAGCGUUCAUAGCGACAUUGCUUUUUGAUUCUUCGAUGUCGG
CUCUUCCUAUCAUACCGAAGCA-GAAUUCGGUAAGCGUUGGAUUGUUCACCCACUAAUAGGGAACGUGAGCU
GGGUUUAGACCGUCGUGAGACAGGUUAGUUUACCCUACU---------------------G-AUG-------
-------------------------------AA----UGUUACCGCAAUAGU-AAU-UGAACUUAGUACGAG
AGGAA-CAGUUCAUUCGGAUAAU-UGGU-UUUUGCGGCUGUCU-GAUC-AGGCAU-U-GCCGCGAAGCUACC
AUCCG-CUGGAUUAUGGCUGAACGCC-UC--UAAGUCAGAAUCCAUGCUAGA----AC--GCGGUGAUUUCU
UUGCUCCA-CACAAUAU-------------------------------------------------------
------------------------------------------------------------------------
------------------------------------------------------------------------
---------------------AGAUGGAUACGA-----------AUA-------------------------
-------------------------------AGGCGU------------C----------------------
---CUUGU--GG-------CG--UC--------------GCU------------------GAACCAU---A-
------GCAGGC------UAG-----CAACGGUG-----CAC-UUGGCG------GAAA-GGCCUUGGGUGC
----------------------------U----U-GCUGGCGA--------AUUGC--AAU----------G
U-----------------CAUUUG----------------------C-GUGGGGAU-----AAA------UC
AUUUGUAUACGACUUAGA-UGU-ACAACGGGGU-AUUGUAAG-CAGUA--------GAGUAGC-CU------
----UGU--UGUUACGAUCUGCUGAGAUUAAG-CCUU-UGUUGUCU---GAUUUG--U--------------
------------------------------------------------------------------------
------------------------------------------------------------------------
------------------------------------------------------------------------
-------------------------------------------------
```

```
>LSUe_284812_SCHPO/1-3654
---------------------------------------AAACUUUCAGC-AACGGAUC-U-CUUGGG-UCU
C-G---CAUCGAUGAAGAACGCAG--CGAAAUGCGAUACGUAAUGUGAAU--U---GCA--GAA-UUC-CGU
GAAUCAUCGAAUCUUGAACGCACAU-U---------GC--------GC----CUUU-G-GG---------UU
C-U------------AC---C----A----A---AGG----C-A-UGCC-UGU-----UUGAGUGUCA-UUA
-----------------------------------------------------------UUUGAC-CUCA-A
AUC--------------------AG-G-UA-GGACUACGCGCUGAACUUAAGCAUAUCAAUAAGCG--CAGG
AAAAGAAAAUAA--CCAUGAUUCCCUCA--GU-AA-CGGCGAGUGAAGCGGGA-A-AA-GCUCAAAUUUGAA
AUCU-GGCAACAUUUCUUUUG------------U---UGU--CCGAGUUGUAAUU--UC-------------
--AA-GAAG-CUG-C-----U-------------------------UUGAG--UGU-----------AG---
ACG-----AUCGG--UCU-------AAGUUC-CUUG------------------------------------
-------GAA-CA-GGACG-------UCAGAGAGGGUGAGAACCCCGUC---------UUU---G----GUC
GAUU--GG--AUAUG-CCAUAUA-------------AAGC-GCUUU--CGAAGAGUCGAGUUGUUUG---GG
AAUGCAGCUCUAA-A-UGGGUGGUAAA-UUUCAUCUAAAGCUAAAUAUUGG--C-GAG-AGACCGAUAGC-G
AAC-A-AGUAGAGUGAUCGAAAG-AUGAAA-AGAAC-UUUGA-AAAGAG-A-GUUAAAUAGUACGUGAAAUU
G-CUGAAAGG--G-AAG-CA-UUGG---AAAU-CAG-UC-----------------------UU--------
-------------------------------------------ACCUGGGU--GA----------GAUC---
------AG-----------------UAG--------------------UCUC--------------------
---------------------------------------------------------------UUCGC----
-----------------------G------------------------------------------------
----------------------------------AGA-----------------------------------
------CUAUGCA--CUCUGAACCUG-UG--GU----------A----------------GG-UCAGCAUC-
---AG--------------U-------------------UUUCGGGGC------------------GGAAAA
AGA-----------------AUAAGGGAA--------GG-------UGGCUUUCCGGG--------------
--------------------------------------------------------------U--U-CUG--
------------------------------------------------------------------------
---------------------CCUGGGGAGUGUUUAUAG------CCC-------UUGUUGUAAUACGUCCA
CUGGGGACU---------------------------------------------------------------
------------------------------------------------------------------------
------------------------------------------------------------------------
------------------------------------------------------------------------
-------------------------------------------------GAGGAC-----------------
--------------------UGCGGCUU--------------------------------------------
------------------------------------------------------------------------
----------------C------------------------------GUGCCAAG-----------------
------------------------------------------------------------------------
------------------------------------------------------------------------
-----------------GAUGCUGACAU-----------------AAUGG--U---UUUCAAUGGCCCGUCU
UGAAACACGGACCAAGGAGUCUA-G-CAUCUAUGCGA-GUGUUUGG--G-UG-------AUGAA-------A
A-CC--CAUC--CGCGAAAUGAAAG-UGA-----------------AUGC--------------------AG
GUGG-GAACGCC----C---------------------------------------------UU--------
GUG---------------------------------------------------------------------
-------------------GCG--------------------------------------UGCACCAUC-GA
CCGACC----CG-GAAG-UU-------------------UGUC--------------A-AUGGAAGGGUUU-
GAG-UA----------------------AGAGCAUAGCUGUUGG-GACCCGAAAGA-UGGUGAACUAUGCCU
```

```
GAAUAGGGUGAAGCCAGA-GGAAAC-UCUGGUGGAGGCUCGU--AGAGAUUCUGACGUGCAAAUCGA-UCUU
CAAAUUUGGGUAUAGGGGCGAAAGACUAAUCGAACCAUC-UAGUAGCUGGUUCCUGCCGAAG-UUUCCCUCA
GGAUAGCAGAAA-CUCA-------------------------------------------------------
---------------------GAU-----CAGUUUUAUGAGGUAAAGCGAAUGAUU-AGA-GGUCUUGG-GG
A-A-GGAAU-UUCC-U-CAACCUAUUCUCAAACUUUAAAUAUGUAAGAC-----------------------
------------------GCCCU-UG-----UCG-CU--UAAU--U--GGA---------------------
CG---UGGG-C--CAUCGA-AU-G--A-----G--AGUUUCU-AGUGGGC-C-AUUU-U-UGG-UAAGCAGA
AC-U-GGCGAUGC-GGGAUG---AACCGAACGUGA-G-GUUAAGGUGCCGGAAU--GU-ACG-CU-CA---U
-C-AGACACC-AGAAAAGGUGUUAGUUCAUCUAGACAGCAGGACGGUGGCCAUGGAAGUCGGAA-UCCGCUA
AGGAGUGUGUAACAACUCACCUGCC-----GAAUGAACUAGCCCUGAAAAU-GGAU-GGCGCUUA-AGCGUA
C----------UACCCAU-ACCUCACCGUCUGGG--UUAGC-------------------------------
-------------------------------------------------------U----------------
------------------------------------------------------------------------
--------------------------UUGAGAAGCUC-AGAC-GAGUAGGCAGGC-GUGGAGG---------
-----------U-UUGUGAC-GAAGCCUU-G-GGCGUGAGCC--U-GGGUCGAACA-G-CCU-CUAGUGCAG
AUCU-UGGUGG-AAGUA-GCAAAU---AUUC-AAAUGA--G---------------AACUUU-GAA-GACUG
AAGUGGGG-AAAGGUUCC-AUGUGAACA-GCAGUUGGACAUGGGUUAGUCGA--UC-CUAAGAGAUAGGGA-
A-G-CUCC-----GUAUGAA--------AGUU----GCA-----CGA---UUU-------------------
-----------------UUCGUGC------CUCC----UA-UC-GA-A-AGGGA-AUCCGGUU-AAUA-UUC
CGGAA----------------------------------CCAGAA---GGUG--------G-AA--------
U----------------CA-----------------------------------------------------
---ACA-CGG---CAAC--------GU-AA-----AU--------------------GAAG-U-UGGA-GAC
GUCGGCG-------------GGAGCC-CUGGGAAGA-GU------UCUCU-------------U-UUCUUU-
UUAACA-AAC--C-----A-UU--------------GAACUA-----CC----------------CUGA-AA
U-C-GG--------------------UUUAU-CCGGAGCUAG---------GGUAUG----GUG-UU-UGG-
AAG--AG-----------------------------------------------------------------
----UUCAGCG-CCUCAUGCUGAAU-C-CGGUGCG-CUCUCGACGGC-CCUUGAAA-A-UCC-AACGGAAGA
-AUGGACCUU-------------------------------CGGGUCCU-----UGUUUUCA-CA-UCUGGU
CGUACUCAUAACCGCA------------------------------GCAGGUC--UCCAAGGU-GAACAGCC
UCUAGU-U-GA-UAGA-ACAAUGUA-G-AUAAGGGAAGUCGGC-AAAAUGGAUCCGUAACUUCGG-GAUAAG
GAUUG-GCUC-U-AA--GGGUUG-G----GUACG-UU------GGGCCUU-GGAACCUGA--AC--GGU---
-UGC------U--GGACUGAG---CG----------------------------UGG--------ACCG--A
UG--------------------------------------------------------------------UC
-UUU-----------------UCUC-----------------------------------------------
---------------------------------------------------------GCCUUUCG-------
GGGU--------------GA--------------------------------GA--AG--------------
-----------------GGAU-------G----------------------------------UUGGA----
------------------------------------------------------------------------
-C-CUG--------------CUUGGACCUU-GGCG-GC----------------------------------
----CG-----G-GA-AGU-----CCUUGGUC----G-----------------------------------
----------------------------------------------------------------GGCU----
-----------------------------------------------------UUUC--UCCU---------
---------------UCU---------------------------CGGGG---A------------------
----UUAU------G---------------------------------------------------------
----CU-----------------------CU--U-AC------UGGCGU-ACG--UUUAACAAC--CAACUU
```

```
AGA--ACUGGUACGGACAAGGGGAAUCUGA-----CUGUCUAAUUAAAACAUAGCAUUGCGAUGGCCAG---
-AA-A------------GUGGUGUUG-ACGCAAUGUGA-UUUCUGCCCAGUGCUCUG-AAUG-UCAAAGUGA
A--------------GAAA--------------UUCAACCAAGCGC--GGGUAAAC-GGCGGGAGUAACUAU
GACUCUCUUAAGGUAGCCAAAUGCCUCGUCAUC-UAACUAGUGACGCGCAUGAAUGGAUUAACGAGA--UUC
CCACUGUCCCUAUC----UAC-UAUCUAGCGAAACC---AC---AGCCUGGGGAACGGGCCAGGCAAAAUCA
GCGGGGAAAGAAGA-CCCUGUUGAGC-UUGACU--CUAGUUU-GACAUUGUGAAGAGACAUAGAGGGUGUAG
GAU-AAGUGGGAGUAUGU--------------------------------------------------UUCG
-----------------------------------------------GCAU-A--C-GC-CGGUGAAAUA-C
CACUACCUUUAUCGUUUCUUUACUUAAUC-AA-------U-GA-A-GCGGA------A-----------UU-
------GG--GA-UU--------UAU---UUCC-C--AU-------------AUUCU---------------
-----------AGCG--------------------------------------UUAA--AG---------UU
---------------------------------------------------------------UC-------
-------------------------------UUC---GCG--------------------------------
----------AAC------UGA------------------------------UCCG----CG----------
------------------------------------------------UUGA-----UGACA-UUGUCAGGU
GGGGAGUUUGGCUGGGGCGGCACAUCUGUUA-AAAGAUAAC-GCAGGUGUCCU-AAGGGGGACUCAUCGAGA
ACAGAAAUCUCGAGUA-GAAUA-AAAGGGUAA-AAGUCCCCUUGAUU-UUGA--------------------
--------------UUUUCA--GUGUGAAUACAAACCAUGAAA-GUGUGGCCUAUCGAUCCUUU--------
---GU-U-----CCCUCGAAA--------------------UU-UGAGGACAGAGGUGCCAG-AA--AAGUU
ACCACAGGG-AUAACUGGCUUGUGGCAGCCAAGCGUUCAUAGCGACGUUGCUUUUUGAUUCUUCGAUGUCGG
CUCUUCCUAUCAUACCGAAGCA-GAAUUCGGUAAGCGUUGGAUUGUUCACCCACUAAUAGGGAACGUGAGCU
GGGUUUAGACCGUCGUGAGACAGGUUAGUUUUACCCUACU--------------------G-AUG-------
-------------------------------AA--G-UGUCGUCGCAAUGGU-AAU-UCAACUUAGUACGAG
AGGAA-CCGUUGAUUCAGAUCAU-UGGU-AUUUGCGGCUGCCU-GACA-AGGCAA-U-GCCGCGGAGCUAUC
AUCUG-CCGGAUAACGGCUGAACGCC-UC--UAAGCCAGAAUCCGUGCCAGA----AA--GCGACGAU-UUU
UUGGUCCG-CAUGAUUU-------------------------------------------------------
------------------------------------------------------------------------
------------------------------------------------------------------------
---------------------AUAUGUAUAAAA-----------AUAG------------------------
-------------------------------AGGUAG-------------GACUU-----------------
---GUUCC--UA-------CU--CU--------------CCU------------------GUAUCGU---A-
------GAAGAU------GGG-----CGA-UGGU---U-GAU-GAAACG------GAAG-UGUUUUAUUGAC
----------------------------U----U-GUCCAUGA--------AAUUCC-AUU----------G
A-----------------AAUCUUG---------------------U-GCGGAAUC-----GAA------UC
CAUUGCAUACGACUUUAA-UGU-GGAACGGGGU-AUUGUAAG-CAGUA--------GAGUAGC-CU------
----UGU--UGUUACGAUCUGCUGAGAUUAAG-CCUU-UGUUUCCC---AAGAUU--U--------------
------------------------------------------------------------------------
------------------------------------------------------------------------
------------------------------------------------------------------------
-------------------------------------------------
>LSUe_3055_CHLRE/1-3506 ABCN02001174.1:150850-151006,151252-154600
---------------------------------------CAACUCUCAAC-AACGGAUA-U-CUUGGC-UCU
C-G---GAUCGAUGAAGAACGCAG--CGAAAUGCGAUACGUAGUGUGAAU--U---GCA--GAA-AUA-CGU
GAAUCAUCGAAUCUUGAACGCAUAU-U---------GC--------GC-----UCG-A-GGC---------U
U-C-----------GGC---C----A----A---GAG----C-A-UGUC-UGC-----CUCAGCGUCG----
```

-G------------------------------------------------------G--CUCGAC-CUGA-G
CUC--------------------AG-G-CA-AGAUUACCCGCUGAACUUAAGCAUAUCAAUAAGCG--GAGG
AAAAGAAACUAA--CAAGGAUUCCCCUA--GU-AA-CGGCGAGCGAACCGGGA-A-UA-GCCCAACUUGAAA
AUC--------UCC-------------------C---UUU-GGAGAAUUGUAGUC--UA-------------
--GA-GAAG-CGC-U-----U-------------------------UCUAG--GGC-----------UG---
G-------CGGAA--CUC-------AAGUCG-GAUC------------------------------------
-------GAA-UG-CCGCG-------UCAGAGAGGGUGAUAACCCCGUC---------GGU-----U-CCUG
CUU---AG--UCCUU-CCACGAA-------------GUG---CUUU--CCACGAGUCGGGUUGUUUG---GG
AAUGCAGCCCUAA-U-UUGGAGGUAAA-UCCCUUCUAAGGCUAAAUACUGC--C-GAG-AGACCGAUAGC-G
AAC-A-AGUACCGUGAGGGAAAG-AUGAAA-AGAAC-UUUGA-AAAGAG-A-GUUAAA-AGUGCUUGAAAUU
G-UUGAGAGG--G-AAG-CG-AUUG---GCGC-UCG-UA------------------------G--------
-----------------------------------------------GUGC--GC------CC---AGG---
------CUU----------------AAG--------------------CGGU--------------------
-------------------------------------------------------------CCUAA------
-----------------------CG-----------------------------------------------
----------------------------------GCC-----------------------------------
------CGUUG------AA---UGUG--C--UG----------GGUGCU-----------GG-UCAGAAUG-
---GG----------------------------------------UUGA-----------------GUUGGC
GGG---------------ACAAAAG------------------CUGGGUCCACCCAGGUAACCCG-------
------------------------------------------------------------------------
------------------------------------------------------------------------
------------------------------------------------------GCCGAUGCCGCCGACUCG
ACCAAGGCG---------------------------------------------------------------
------------------------------------------------------------------------
------------------------------------------------------------------------
------------------------------------------------------------------------
--------------------------------------------------U---------------------
-----------------------AAAGA--------------------------------------------
------------------------------------------------------------------------
-----------------------------------------------GUACCUUG----------------U
C----------------CUU----------------------------------------CGGGA-------
------------------------------------------------------------------------
---UCUGUGCUCU-AAAGAUUCUGGCAG-----------------AAGAG--C---GUCAAUCGACCCGUCU
UGAAACACGGACCAAGGAGUCUA-A-CAUGUAUGCGA-GUUGGCGG--G-U--------GGAAA--------
A-CC--CGUA--AGCGCAAGUAACC-UGAC---------UGGUGGGAUGG---------------------G
GUAA-AACCC--------------------------------------------------------------
------------------------------------------------------------------------
------------------------------------------------------------UGCACCAUC-GA
CCGACC----AU-GUUGU---------------------UU-C----------------UACGAAAGGUUU-
GAG-UG----------------------CGAGCAUACCUGUUGG-GACCCGAAAGA-UGGUGAACUAUGCCU
GAGCAGGGUGAAGCCAGA-GGAAAC-UCUGGUGGAGGCUCGU--AGAUGUGCUGACGUGCAAAUCGC-UUUU
CAGACUUGGGUAUAGGGGCGAAAGACUAAUCGAACCAUC-UAGUAGCUGGUUCCCUCCGAAG-UUUCCCCCA
GGAUAGCUGGAG-CUUG-------------------------------------------------------
---------------------------AUCAGUUUUAUCGGGUAAAGCGAAUGAUU-AGA-GGUUCGGG-GG
A-U-GAAAC-AUCC-U-UCACCUAUUCUCAAACUUUAAAUAGGUAAGAC-----------------------

```
------------------GUGUC-GG-----UUG-CU--UAAU--U--GAA---------------------
CC---GGCA-C---AUUCA-AU-G--U-----G--AGCUCCA-AGUGGGC-C-AUUU-U-UGG-UAAGCAGA
AC-U-GGCGAUGC-GGGAUG---AACCGAUAGUCG-A-GUUAAGGUGCCAAACU--AC-GCG-CU-AA---C
-CUAGAUCCC-ACAAAGGGUGUUGAUUGAUAUAAACAGCAGGACGGUGGUCAUGGAAGUCGAAA-UCCGCUA
AGGAGUGUGUAACAACUCACCUGCC-----GAAUCAAUUAGCCCCGAAAAU-GGAU-GGCGCUUA-AGCGCG
U----------GACCUAU-ACUCGGCCAUGGAAG--CAAG--------------------------------
------------------------------------------------------------------------
------------------------------------------------------------------------
---------------------------UGCGACGCUU-CCAU-GAGUAGGAGGGC-GUGGGUG---------
-----------U-C-GAGAC-UAAGCCUC-U-GGCGUGAGCC--U-GGGUGAAUCG-G-CAU-CUAGUGCAG
AUCU-UGGUGG-UAGUA-GCAAAU---AUUC-AAAUGA--G---------------AACUUU-GAA-GACUG
AAGUGGAG-AAAGGUUCC-AUGUGAACA-GCAAUUGGACAUGGGUUAGUCGA--UC-CUAAGAGAUGGGGU-
A-A-UCCU-----GUGUGAA--------GAG-----CGC-----GAU---U---------------------
CG---------------CG----C------UGCC----CA-UC-GA-A-AGGGA-AAAGGGUU-AAGA-UUC
CCUUA----------------------------------CUUGGA---CAAG--------G-CG--------
G----------------CU-----------------------------------------------------
---G-G-CGG---UAAC--------GC-AA-----GC--------------------GAGC-C-CGGA-GAC
AUCGGCA-------------UCGGCC-CUGGGAAGA-GU------UCUCU-------------U-UUCUUU-
UUAACA-ACG--C----GA-A-------------------GG-----CC----------------CUGG-AA
U-C-GA--------------------AUCAU-UCGGAGAUAG---------GGCUCA----GAC-GU-UGG-
UAA--AG-----------------------------------------------------------------
----CACCGCA-CUUCUCGCGGUGU-C-CGGCGCG-CCGUUGACGGU-CCUUGAAA-A-UCC-GGGGGAGCA
-U----------------------------------------------U-----CCCGAUCU-UG-CCAAGU
CGUACUCAUAACCGCA------------------------------UCAGGUC--UCCAAGGU-GAACAGCC
UCUAGU-C-GA-UAGA-ACAAUGUA-G-AUAAGGGAAGUCGGC-AAAAUGGAUCCGUAACUUCGG-GAAAAG
GAUUG-GCUC-U-GA--GGGCUG-G----GCCUA-GG------GGUC----UGC---------A--GCU---
-GCGAAGCUCGG-GACUGCGG---UGGUCUACC---------------------CAG---------CUG---
----------------------------------------------------------------------GA
AACG-----------------GCU------------------------------------------------
-----------------------------------------------------------------------G
--GG--------------C---------------------------------GG--AC--------------
---------------------------------------------------------------UGCU-----
--------------------------------------------------G---------------------
-C------------------GU--G-UC----CUGGGU----------------------------------
---GGA----------CGG-----CUGUAGAA----------------------------------------
-------------------------------------------------------------------G----
-------------------------------------------------------------CU---------
---------------U-------------------------------CGG----------------------
------------------------------------------------------------------------
----CG-----------------------UU------------CCCUAG-GCG--ACGAACAGC--CAACUC
AGA--ACUGGUACGGACAAGGGGAAUCCGA-----CUGUUUAAUUAAAACAAAGCAUUGUGAUGGUCCU---
-AA-A-------------GGAUGUUG-ACACAAUGUGA-UUUCUGCCCAGUGCUCUG-AAUG-UCAAAGUGA
A--------------GAAA--------------UUCAACCAAGCGC--GGGUAAAC-GGCGGGAGUAACUAU
GACUCUCUUAAGGUAGCCAAAUGCCUCGUCAUC-UAAUUAGUGACGCGCAUGAAUGGAUUAACGAGA--UUC
CCACUGUCCCUAUC----UAC-UAUCUAGCGAAACC---AC---AGCCAAGGGAACGGGCUUGGAAUAAACA
```

```
GCGGGGAAAGAAGA-CCCUGUUGAGC-UUGACU--CUAGUCC-GACUUUGUGAAAUAACUUAAGAGGUGUAG
AAU-AAGUGGGAGC------------------------------------------------------UUCG
----------------------------------------------------G--C-GA-CGGUGAAAUA-C
CACUACUUUUAACGUUGUUUACUUAUUC-CA--------U-UA-C-UUGGA------GG----------CG-
------GGACUC-----------------UGUCCC--UG-------------CUUCU---------------
-----------AGCU--------------------------------------CUAA--GA---------CG
------------------------------------------------------------------------
---GCU---------------------------UUU-GCA--------------------------------
----------CGU------CGA------------------------------UCCA----GG----------
------------------------------------------------UGGA-----AGACA-UUGUCAGGU
GGGGAGUUUGGCUGGGGCGGCACAUCUGUUA-AAAGAUAAC-GCAGGUGUCCU-AAGAUGAGCUCAACGAGA
ACAGAAAUCUCGUGUA-GAACA-AAAGGGUAA-AAGCUCAUUUGAUU-UUGA--------------------
--------------UUUUCA--GUACGAAUACAAACUGUGAAA-GCAUGGCCUAUCGAUCCUUU--------
---AG-C-----CUUUCGGGA--------------------UU-UGAAGCUAGAGGUGUCAG-AA--AAGUU
ACCACAGGG-AUAACUGGCUUGUGGCAGCCAAGCGUUCAUAGCGACGUUGCUUUUUGAUCCUUCGAUGUCGG
CUCUUCCUAUCAUUGUGAAGCA-GCAUUCACCAAGCGUUGGAUUGUUCACCCACUAAUAGGGAACGUGAGCU
GGGUUUAGACCGUCGUGAGACAGGUUAGUUUUACCCUACU--------------------G-UUG-------
-------------------------------GA--C-CGAUUCCGCCAUAGU-AAU-UCGGCUCAGUACGAG
AGGAA-CCGCCGAGUCAGAUAAU-UGGUA-A-UGCCCUUGUCU-GAAA-AGACAA-U-GGGGCGAAGCUAAC
AUCUG-UAGUCUAAUGACUGAACGCC-UC--UAAGUCAGAAGACGUGCUAG-----G--UGCGGAGUC-ACU
UACCC--A-AUGAUGUC-------------------------------------------------------
------------------------------------------------------------------------
------------------------------------------------------------------------
---------------------ACCC-----------------------------------------------
--------------------------------GACUA-----------------------------------
----------AGG-----AUA--CA--------------UCCGCCUGUGCGGAUGCUGGAGCAUACC---C-
-------GUUGGU--UCCCCUGU---UA---GGU-----CCACAUGGCC------GAAG----CAGGCGCCA
AGCAUGAC--------------------------------------------------AAU----------U
C-----------------CACUCGU---------------------C-AUUGGGU------AAA------UC
CUCUGUAGACGACUUUGU-U---GCAACUGGGU-AUUGUAAG-UGGUA--------GAGUGGC-CU------
----UGC--UGCUACGAUCCACUGAGAUUCAU-CCCG-UGUUGCUA---AGAUUUGU---------------
------------------------------------------------------------------------
------------------------------------------------------------------------
------------------------------------------------------------------------
-------------------------------------------------
>LSUe_284591_YARLI/1-3338
------------------------------------A--UAACUAUCAAC-AACGGAUC-U-CUUGGC-UCU
C-A---CAUCGAUGAAGAACGCAG--CGAACCGCGAUAUUUUUUGUGACU--U---GCA--GAU------GU
GAAUCAUC-AAUCUUUGAACGCACAU-U--------GC--------GC----GGUA-U-GG----------CA
U-U------------CC---G----U----A---CCG----C-A-CGGA-UGG-----AGGAGCGUGU-U--
------------------------------------------------------------UUCAC-CUCC-U
U-C--------------------AU-C-CG-AGAUUACCCGCUGAACUUAAGCAUAUCAAUAAGCG--GAGG
AAAAGAAACCAA--CAGGGAUUGCCUCA--GU-AA-CGGCGAGUGAAGCGGCA-A-AA-GCUCAAAUUUGAA
ACCC-UCG------------------------------------GGAUUGUAAU--UG--------------
--AA-GAUU-UGG-------C-------------------------AUUGG--AGA-----------A----
```

```
A-------GCUAA--CCC-------AAGUUG-CUUG------------------------------------
-------GAA-UA-GUACG-------UCAUAGAGGGUGACAACCCCGUC---------UGG---C----UAA
CCG---UU--CUCCA-UGUA----------------UUGC-CUUAU--CAAAGAGUCGAGUUGUUUG---GG
AAUGCAGCUCAAA-G-UGGGUGGUAAA-CUCCAUCUAAAGCUAAAUACUGG--U-GAG-AGACCGAUAGC-G
AAC-A-AGUACUGUGAAGGAAAG-GUGAAA-AGAAC-UUUGA-AAAGAG-A-GUGAAAUAGUAUGUGAAAUU
G-UUGAUAGG--G-AAG-GA-AAUG---AGUG-GAGAGUGGC------------------------------
-----------------------------------------------CGAG--GUUUCA-------------
--------------GCCGCCCC--------------------------------------------------
----------------------------------------------------------------UCGU----
------------------------------------------------------------------------
------------------------------------------------------------------------
------GGGCGG--UGUAC---UGCCGAC-------------------------GCCGAGUCAUCGAUAGC-
---GA-------------------------------------------G-----------------ACGAGG
GUU-----------------ACAAAU-----------GGGAGCGCC--------------------------
-------------------------------------------------------------UU--U-CG---
------------------------------------------------------------------------
-----------------------------------------------------GGCGUUCUCCCCUAACCCU
CCACACUGCCA-------------------------------------------------------------
------------------------------------------------------------------------
------------------------------------------------------------------------
------------------------------------------------------------------------
------------------------------------------------------------------------
------------------------------------------------------------------------
------------------------------------------------------------------------
------------------------------------------------------------------------
------------------------------------------------------------------------
------------------------------------------------------------------------
------------------------CCGA------------CGACAUAAUC--C---ACCAUUUCACCCGUCU
UGAAACACGGACCAAGGAGUCUA-A-UGGAUAUGUGA-GUGUUAGG--G-U--------GGCAA--------
A-CC--CCAG--CGCGCAAUGAAAG-UGA-----------------AUG----------------------G
AUUC--------------------------------------------------------------------
------------------------------------------------------------------------
---------------------G--------------------------------------UUCAGAAUC-GA
CCGAAC----UU-GAU-----------------------UAUUA-----------------UGACAGUUUU-
GAG-UA----------------------A-A-CACAUCCAUUGG-GACCCGAAAGA-UGGUGAACUAUGCCU
GGAUAGGGUGAAGUCAGA-GGAAAC-UCUGAUGGAGGCUCGU--AGCGGUUCUGACGUGCAAAUCGA-UCGU
AGGAUCUGGGUAUAGGGGCGAAAGACUAAUCGAACCAUC-UAGUAGCUGGUUCCUGCCGAAG-UUUCCCUCA
GGAUAGCAGAAG-CUCA-------------------------------------------------------
---------------------UAU-----CAGUUUUAUGAGGUAAAGCGAAUGAUU-AGA-AGUAUUGG-GG
G-C-GAAAU-GCCC-U-CGGCUUAUUCUCAAACUUUAAAUAUGUAAGAA-----------------------
------------------GCCUU-GG-----UUA-CU--UAAU--C--GAA---------------------
CC---GUGG-C---UACGA-AU-G--AA----G--AGCUUCU-AGUGGGC-C-AUUU-U-UGG-UAAGCAGA
AC-U-GGCGAUGC-GGGAUG---AACCGAACGUGG-A-GUUAAGGUGCCGGAAU--AC-ACG-CU-CA---U
-C-AGACACC-ACAAAAGGUGUUAGUUUAUCUAGACAGCCGGACGGUGGCCAUGGAAGUCGGAA-UCCGCUA
AGGAGUGUGUCACAACUCACCGGCC-----GAAUGAACUAGCCCUGAAAAU-GGAU-GGCGCUUA-AGCGUG
```

```
U----------UACCUAU-ACUCUACCGAGAGGA--G-----------------------------------
------------------------------------------------------------------------
------------------------------------------------------------------------
-------------------------------GUUUCC-UCUC-GAGUAGGCAGGC-GUGGGGG---------
-----------U-U-GUUGA-GAAGCGUU-G-GCCGAGAAGC--U-GCGUCGAACG-G-CCC-CUAGUGCAG
AUCU-UGGUGG-UAGUA-GCAAAU---AUUC-AAAUGA--G---------------AACUUU-GAA-GACUG
AAGUGGGG-AAAGGUUCC-GUGUGAACA-GCAGUUGGACACGGGUAAGUCGA--UC-CUAAGGGGUGGCAU-
A-A-CUGU-----CGCGUAC--------G-------GCC-----CGA-------------------------
-----------------UAAGGGC-------CUU----CU-CC-AA-A-AGGGA-AGCCGGUU-GAAA-UUC
CGGCA----------------------------------CUUGGA---UGUG--------G-AU--------
U----------------CU-----------------------------------------------------
---CCA-CGG---CAAC--------GU-AA-----CU--------------------GAAU-G-UGGG-GAC
GGUGGCA-------------CAAGUC-UUGGAAGGA-GU------UAUCU-------------U-UUCUUU-
UUAACG-GAG-UC-----A-A-------------------CA-----CC----------------CUGG-AA
U-U-AG--------------------UUUGU-CUAGAGAUAG---------GGUAUC-----GU-UC-CGG-
AAG--AG-----------------------------------------------------------------
-----GGGGGC-AGCUUUGUCCCCU-C-CGAUGCA-CUUGUGACGCC-CCUUGAAA-A-CCC-GCAGGAAGG
-A---------------------------------------------AU-----AGUUUUCA-CG-CCAAGU
CGUACUGAUAACCGCA------------------------------GCAGGUC--UCCAAGGU-GAACAGCC
UCUAGU-U-GA-UAGA-AUAAUGUA-G-AUAAGGGAAGUCGGC-AAAAUAGAUCCGUAACUUCGG-GAUAAG
GAUUG-GCUC-U-GG--GGGUUG-G-----UGGA-UG------GAAG----CGU------------------
-GGG------AG-ACCCCAAG---GG---------------------------------------ACUG---
------------------------------------------------------------------------
-GCA-----------------GCU------------------------------------------------
---------------------------------------------------------GG-----------GC
AACU--------------G---------------------------------GC--AG--------------
---------------------------------------------------------------CCGG-----
------------------------------------------------------------------------
-------------------------------ACCC-GC----------------------------------
------------------------------------------------------------------------
----------------------------------------------------------------GGCA----
------------------------------------------------------------------------
-------------------------------------------GACACUG---C------------------
--------------GU--------------------------------------------------------
----CG-----------------------CU------------CCGUCC-ACA---UCAUCAAC--CGCCCC
AGA--ACUGGUACGGACAAGGGGAAUCUGA-----CUGUCUAAUUAAAACAUAGCUUUGCGAUGGUUGU---
-AA-A------------ACAAUGUUG-ACGCAAAGUGA-UUUCUGCCCAGUGCUCUG-AAUG-UCAAAGUGA
A--------------GAAA--------------UUCAACCAAGCGCGCGGGUAAAC-GGCGGGAGUAACUAU
-GCUCUCUUAAGGUAGCCAAAUGCCUCGUCAUC-UAAUUAGUGACGCGCAUGAAUGGAUUAACGAGA--UUC
CCACUGUCCCUAUC----UAC-UAUGUAGCGAAACC---AC---AGCCAAGGGAACGGGCUUGGCAGAAUCA
GCGGGGAAAGAAGA-CCCUGUUGAGC-UUGACU--CUAGUUU-GACAUUGUGAAGAGACAUAGGGGGUGUAG
AAU-AAGUGGGAGC------------------------------------------------------UUCG
----------------------------------------------------G--C-GC-CGGUGAAAUA-C
CACUACCCUUAUCGUUUCUUUACUUAUUU-AG-------U-AA---GUGGA------A-----------GU-
------GG--UU-UA--------ACA---ACCA-U--U---------------UUCU---------------
```

```
-----------AGCA--------------------------------------UU-----------------
---------------------------------------------------------------CC-------
---UU-------------------------------------------------------------------
----------------------------------------------------UCCA----GG----------
------------------------------------------------CUGA-----AGACA-UUGUCAGGU
GGGGAGUUUGGCUGGGGCGGCACAUCUGUUA-AAAGAUAAC-GCAGAUGUCCU-AAGGGGGACUCAAUGAGA
ACAGAAAUCUCAUGUA-GAACA-AAAGGGUAA-AAGUCCCUUGAUU-UUGA---------------------
--------------UUUUCA--GUGUGAAUACAAACCAUGAAA-GUGUGGCCUAUCGAUCCUUU--------
---AG-U-----UGUUCGGAG--------------------UU-UGAACCUAGAGGUGCCAG-AA--AAGUU
ACCACAGGG-AUAACUGGCUUGUGGCAGUCAAGCGUUCAUAGCGACAUUGCUUUUUGAUCCUUCGAUGUCGG
CUCUUCCUAUCAUACCGAAGCA-GAAUUCGGUAAGCGUUGGAUUGUUCACCCACUAAUAGGGAACGUGAGCU
GGGUUUAGACCGUCGUGAGACAGGUUAGUUUUACCCUACU--------------------G-AUG-------
-------------------------------GA-----CUUGUUGCAAUAGU-AAU-UGAACUUAGUACGAG
AGGAA-CAGUUCAUUCGGAUAAU-UGGU-GUUUGCUGCUGUCU-GACC-AGGCAA-U-GCAGCGAAGC-ACC
ACCCG-CUGGGUUAUGGCCGAACGCC-UC--CAAGUCAGAACCCAUGCCAGA----AG--GGAAGAAU-CAG
GGGGAAGG----------------------------------------------------------------
------------------------------------------------------------------------
------------------------------------------------------------------------
-----------------------AGGGAUAUGA---------------------------------------
-------------------------------AGA--------------------------------------
----------AG-------UA--CC--------------GCA------------------GUACCGG---A-
-------------------GG-----GGG-AGGG-----GGG-GUGGAUAAG---GAAA---CCGCCCGCCC
----------------------------C----C-CCCCG------------------ACU----------G
GA----------------AAGACCC---------------------A-CCCUUGUG-----AAA------UC
CAUUGUAGACGACUUUAG-UAU-GCGACGAGGU-AUUGUAAG-UAGUA--------GAGUAGC-CU------
----UGU--UGUUACGAUCUAUUGAGAUUAAG-CCUU-UGUUGUUUA--GAUUCG--A--------------
------------------------------------------------------------------------
------------------------------------------------------------------------
------------------------------------------------------------------------
-------------------------------------------------
>LSUe_104782_ADIVA/1-3945
---------------------------------------UAAUUCUGUAC-GGUGGAUC-A-CUCGGC-UCG
U-U---AUUCGAUGAAGAGCGCAG--CAAAGUGCGAGAAAGCAAUGCGAA--U---UGC--AGG-ACA-CAU
UGAGCAUGAAUAACUCGAACGCAUCU-A--------GC--------AG----CCAA-G-AC---------AU
U-A------------GU---C----G----A---GGC----U-A-AAUU-UGC-----UUCAGUCUUG-UUU
--------------------------------------------------------U--UUCAAG-CUGA-A
GUC--------------------AA-A-UU-AGAAUACUUGCUGAACUUAAGCAUAUUAAUAAGCA--AAGG
AAAAGAAAUCAA--CUGAGAUUGCCUUA--GU-AA-UGGCGAAUGAACCGGCA-A-AA-GCUCAUUGCUGAA
GCGU-GUUAGAUUUAC--UUC------------U---GAC--AUGCAUUGUAGCA--UU-------------
--UG-GGGU-AAC-U-----A-------------------------CUGUA--UGG-----------UG---
CCU-----GUUCG--UGU-------GAGUUC-CUUU------------------------------------
-------GAU-CA-GGGCUU---UAACCAUAGAAGGUGCUAGUCCUAUA---------AUC---A----CGA
AUAGUUCU--GUGCC-GUACA---------------UUGU-CUACC--UUUCGAGUAGCGUAGUUUG---AG
AUUGCUACGCCAA-CGUGGGACGUAAA-CUUGUUCCAAAGCUAAAUAUUAU--A-ACG-AGACCGAUAGC-G
AAC-A-AGUACCGUGAGGGAAAG-UUGAAA-AGCAC-UCUGA-AGAGAG-A-GUUAAAUAGUACGUAAAACC
```

```
G-UACAGAAG--A-AAA-CG-AAUG---GCCG-CAU-UCAUCAG----------A-------UU--------
-------------------------------------------ACUUUGGA----UUCAAUUC--GAUU---
------UCUAUAUGAUGGUAAU-A-CAU-GUU---------GAUUUGGAAC---------------------
------------------------------------------UU-------CAAC-------AGUCCA----
------------------------------------------------------------------------
----------------------------------UUG--------AUAUG--UUCU------UGCCUAGAAU
C-----GGAUGCA---UUU---UCAA--A--GU----------A----------UUAACUGAACCUGGAUC-
---UA-----------UAAC-------------------AUUACAGCUA-----------------GUCGAA
UCA-----------------GUCUAUUUUGGUAGCGACUUUUAGUCGUG-----------------------
------------------------------------------------------------UUA--A-A----
------------------------------------------------------------------------
-------AAUAUGACUUGCAUAAAUAACAAAUUCGACU------------UGGCGUUAUAGGUAAA------
------------------------------------------------------------------------
------------------------------------------------------------------------
------------------------------------CUGACGUGUUCUUUGAUU------------------
-----------UCAGUA---------------CUUUC-----------------GAGU--------------
------------------------------------------------ACGG--------------------
--------------------CACGCCAU-------GAGCAUUGUUU--------------------------
---------------------------------------GUGU---U-ACAUG----AGUAACUAAUAUCUA
AACAAUAUUAGUUUUAU-UCAUU-GU------CA-AUGCCUA-----AUGUUUUG--UGG---CUGUC---A
A-----------------------------------------------------------------------
----------------------------------------U-------------------------------
----------------GAAAUCGGCAAGUCAACAGGAGUUUUGAAUGCCG--A---GUUAUUCGACCCGUCU
UGAAACACGGUCUAAGGAGUUCA-C-UGCAUGUGCAA-GAGUUCGG--C-UGU------AUGA-------AA
A-GC--CUUG--CUCGUAAUGAAAG-UAA-----------------AGAACAUUCUUGC---AAUGUUUGAA
GUGU-GAUUUCU----GUCAUCUG----GUGCAA----------------------------AU--------
CAU---------------------------------------------------------------------
-------------------GACAAUG--------------------------------AACGCAACACU-GA
CAGGUU----CUAUUGC----------------------CGCUG----------------GCAAAGAG-CU-
UAG-UA----------------------AGAGUAUAUGCAAUGA-GACCCGAAAGA-UGGUGAACUUAACGU
GUGUAGAAUGAAGCCAGG-GGAAAC-CCUGGUGGAAGUUCGU--AGCGGUUCUGACGUGCAAAUCGA-UCGU
CAAACAUGCGUUAAGGGGCGAAAGACUAAUCGAACCAUC-UAGUAGCUGGUUCCGUCCGAAG-UUUCCCUCA
GGAUAGCUGGCA-UUCA-------------------------------------------------------
-----------------AUUUUCA-----CAGUUAUAUCCGGUAAAGCGAAUGUUU-AGA-GGCAUUGG-GC
G-U-AAAAU-ACGC-U-CAACCUAUUGACAAACUUUAAAUGGGUAUGAA-----------------------
------------------GUUCA-UU-----UUU-CUA-CUUCAUU--GAG---------------------
AA---UGAA-C---AGCGA-AU-G--U-----G--AGUGCCA-AGUGGGC-C-AUUU-U-UGG-UAAGCAGA
AC-U-GGCGCUGU-UGGAUG---CACAAAACGUCA-A-GAUAAAGAGCCAGAUG--GUCUCG-CU-CA---U
-C-AGAUACC-AUAAAAGGUGUUGGUCGAUAUAGACAGCAGGACGGUGGCCAUGGAAGUCGGAA-UCCGUUA
AGGAGUGUGUAACAACUCACCUGCC-----GAAUCGACUAGUCCUAAAAAU-GGAU-GGCGCUCG-AGCGAG
C----------CUCUUAU-ACUUGACUGUUACGA--------------------------------------
--------------------------------------------------------------------UACA
ACAAGCCAUUAA------------------------------------------------------------
-----------------------CGGACAUCAUGUCG-UAAC-AAGUAGGACGGUUGAUAUGG---------
-----------U-U-GCGUC-GAAGGCUA-A-CACGUGAGUG--U-GGCUUGAGCA-G-UCA-UAUGUACAG
```

```
AUCU-UAGUGG-UAGUA-GCAACU---AUUC-AAAUGA--G---------------AACUUU-GAA-GACUG
ACGUGGAG-AAAGGUUCC-AUGUGAACA-GCUAUUGGACAUGGGUCAGUCAG--UC-CUAAGAUGCGCUCG-
A-A-AGAG---G-GUUCCGAAC------GAG-----GAAUAUCUAUAUUCAUAUC---------------AU
AU--------------UUAUAUUC-----CAUCA----AA-UC-GA-A-AGGGA-AACAAGUU-AAUA-UUC
UUGUA----------------------------------CUAAGU---GUCG--------G-AGAUUGUU--
U----------------GAUUUCGU----------------------------------GUCAAACAU----
---AUG-UGG---UAAC--------AC-AA-----AC--------------------AAGC-U-CAAA-UAC
AUCACAA-------------GUGGUC-CUGGAAAGA-GU------UUUCU-------------C-UUCUUU-
GUUACU-UCA-UC----AG-A------------------CGA-----CC----------------CUGG-AA
U-C-AG--------------------UUCAA-CUGGAGAUAG---------GGUUCA----AAU-GA-AGG-
UAA--AG-----------------------------------------------------------------
--AGUAGCCGU-UCUUGCUACUUCC-U-CGGUGCA-UCAUUUGUGAU-CCUUGAAA-A-UUU-GAGCGAGAU
-A----------------------------------------------UU-----CAAUUCG-CA-CUUAUC
UGUACUCUAAACCGCA------------------------------UCAGGUC--UCCAAGGU-GAAAAGCC
UCUAGU-CAAU-UAGA-ACAAGGUA-GCACAAGGGAAGUCGGC-AAAAUCGAUCGGCAACUUCGG-GAUAAC
GAUUG-GCUC-U-AA--GUGCUA-G----GGUGU-UC------GGGCUU--GAU-GAUAAA-GC--AGU---
-AGA------C-----UUGCU---AU----------------------------UGGA-------GCUG--A
G---------------------------------------------------------------------CU
UUCG-----------------GAUG-----------------------------------------------
---------------------------------------------------------CA-----------GC
AAUGUG------------UC-------------------------------UGAUCGU--------------
---------------------------------------------------------------UCGGA----
------------------------------------------------------------------------
-C-UA---------------GU--G-GCUU-GUCU-UG----------------------------------
---CUG----UG-GAAUCG-----UU-UUGCUACG-GUA---------------------------------
----------------------------------------------------------------UGCG----
------------------------------------------------------GUU--CGCU---------
---------------U-------------------------------GCG---G------------------
--------------G---------------------------------------------------------
----UC-----------------------GUAUU-ACUU----CGGACA-UCG--UUUAAUAGC--AAACUU
AGA--ACUCGCUGCGUCUAGGGGAAUCCGA-----CUGUCUAAUUAAAACAGAGCUUUGCGAUUGGCCG---
-GA-AA---------ACUUGGUGUUG-ACGCAAAGUGA-UUUCUGCCCAGUGCUCUG-AAUG-UUAACGUGA
A--------------GAAA--------------UUCAAAUAAGCGC--GGGUAAAC-GGCGGGAGUAACUAU
GACUCUCUUCUGGUAGCCAAAUGCCUCGUCAUC-UAAUUAGUGACGCGCAUGAAUGGAUUAACGAGA--UUC
CCACUGUCCCUGUC----UAC-UAUCUAGCGAAACC---AC---AGCCAAGGGAACGGGCUUGGACUUCAAU
GCGGGGAAAGAAGA-CCCUGUUGAGC-UUGACU--CUAAUCU-GUCUUCGUAAAGAGACUUGAAAAGUGUAG
CGU-AGAUGGGAGUG-----------------------------------------------------GCAA
--------------------------------------------------C-A--C-GU-CAAUGAAACA-C
CAUUAUUUCAUUGUUUCUUUACUUACUU-GG--------U-UA-U-ACAGA-------G-----------AU-
------UG--GA-UU-------UUAC---AUCC-A--GU-------------GUUUG---------------
-----------AGUG--------------------------------------ACAA--AC---------GU
---------------------------------------------------------------CG-------
---AUA-------------------------CUU---GUA-----UCA------------------------
----------ACG------UGA------------------------------UCUG----CU----------
------------------------------------------------UCAA-----GGAAA-GAGACAGGU
```

GGGGAGUUUGGCUGGGGCGGUACAUCUAUCA-AAUGUUAAC-GUAGGUGUCCA-AAAGUUGACUCAGUGCGG
ACAGAAACCACACAUU-GAGUA-AAAGGGCAU-AAGUCAGCUUGAUU-UAGA--------------------
--------------UUUUCA--GUACGAAUACAGACGGAGAAA-UCCGGGCCUAUCGAUCCUUU--------
---UG-A-----UUUAACGAG--------------------UU-UUUUGCAAGAGGUGACAG-AA--AAGUU
ACCACAGGG-AUAACUGGCUUGUGGCGGCCAAGCGUUCAUAGCGACGUCGCUUUUUGAUCCUUCGAUGUCGG
CUCUUCCUAUCAUUGCGAAGCA-GACUUCGCAAAGUGUUGGAUUGUUCACCCACCAAUAGGGAACGUGAGCU
GGGUUUAGACCGUCGUGAGACAGGUUAGUUUACCCUACA---------------------U-UGGA------
-------------------------------UU--U-UGUCAUUGAUAUAGG-AAU-CGUACUUAGUACGAG
AGGAA-CCGUACGUUCGGACAUU-UGGU-UUACGUAUUUGUUC-GAAA-GGACAA-U-GAUGCGAAGCUACC
AUCCG-ACGGCUUACAACUGAACGCC-UC--UAAGUUGGAAUCCGAUCUAG-----AC--UCAAUGAUUUUU
UGGCACAU-UGGUUGUC-------------------------------------------------------
------------------------------------------------------------------------
------------------------------------------------------------------------
---------------------GUCAGCACAUUU---------------------------------------
-----------------------AUAUA---UGUUCA---A-------ACUGUUG-----------------
---AUCAA--GU-------GA--AC-------------AUAU------------------GAAUGAA---G-
-----CUACCGU-----AU--UG--UUGCCUCG-UCA--UAAUCGAAAU------GAGA---UUGUGAUU--
----------------------------G----C-AACGAUGC--------AUUUUUAGUC----------G
AA----------------ACAACAC---------------------U-AAUGGCUA-----GAA------UC
GAUUGUAAACGACUUUCU-UCA-GUAACAGAGU-AACGUACA-CGUGA--------GAGCAAC-CG------
----CCA--CAUUGCGAUCCGCUGAGUUUCUG-CUCG-AUGUUAUG---CUGAUU--C--------------
------------------------------------------------------------------------
------------------------------------------------------------------------
------------------------------------------------------------------------
-------------------------------------------------
>LSUe_352472_DICDI/1-3902
-------------------------------------------CAUAAAC-GGUGAAUA-C-CUCGAC-UCC
U-A---AAUCGAUGAAGACCGUAG--CAAACUGCGAUAAUUCACUUGAAU--U---GCA--GCC-UA--CUG
GGAUAGUUGAAAUGUUGAACGCACAU-G--------AU--------GA----CAUC-G-GUCC-------UU
U-C---------GGAUU---A----G----G---UGU----U-A-UACU-UGG-----GUGAGAGUGG-UCC
U------------------------------------------G------------A--UUCCGC-CUCA-C
C-U--------------------UU-G-UA-AGAUUACCCGCUGAACUUAAGCAUAUCAGUAAGCG--GAGG
AAAAGAAACUAA--CUAGGAUUCCGUCA--GU-AA-CGGCGAGUGAAGACGGA-A-UA-GCCCAAGGUUCAA
ACCU-GGAUCUCUU----CGA------------G---GUU--AGGUGAUGUGACC--UA-------------
--UG-GACU-GAU-G---GAG-------------------------CCCGC--UGU-----------UG---
--------UGACU--GCU-------AAUUCCGUUUG------------------------------------
-------GAA-UU-UCGAG-------UCGUAGAAGGUGAUAACCCUGUU---------CGC---A----GUA
UCA---CA--ACAGU-UGGACUUUG-----------CCAU-UAGCU--CCACGAGUAGGAAUGUCUG---AA
AUUGCAUUCUGAA---UGGGUGAUAAG-AUUCAUCCAAGGCUAAAUAUAUG-UU-AGG-AGAUCGAUAGC-A
UAC-A-AGUACCGUGAGGGAAAG-GUGAAA-AGAAC-UUUGA-AAAAAG---GUUUAAAAGUAUUUGACACC
G-UUUAUGUG--G-AAG-CG-UUUACUUGGAC-CCCGAU--------------UAAU-GACGUC--------
-------------------------------------------GGUUUAGCUCUA---AUUCUUAGGUG---
------GCC--------AAAGUAG-AGU--------GUUAC-------GUGC--------------------
-----------------------------------------------------------UGAUCAAAA----
-----------------------GG-----------------------------------------------

```
----------------------------------UAACGG-----ACAUU--UGAU-----UCAUUGGUUAU
CGACGAGGAAGG--UACUC---UAAA-----UC----------G--------GCCAGUUACUAACGGGUGA-
---GA-------------------------------------------U-----------------CUGAUG
UUU-----------------AUAAAA-----------UGGGGGAUGAGGCUUAUCGGCU----UGC------
-------------------------------------------------------------UG--G-UGG--
------------------------------------------------------------------------
--------------------------------CUCGCUC------UCAAUAAUGGAUAUUGGGUUUCAUCAA
------------------------------------------------------------------------
------------------------------------------------------------------------
------------------------------------------------------------------------
------------------------------------------------------------------------
-------------------------------------------------GAGUGCAAAAUGGUGGC--AAUU
CACUAUU------------AGUGGUUAUUAAUUUUGUUUGCGU----------------------------G
GCU------------------------------------UGGCCUUGUCUACA----GGUUAUCUUCG----
--GA---UGGCU--UGU-AGCUUUGUUGAACGCGUGGGCUUAAUGUUGUGAUUCU---AG------------
------------------------------------------------------------------------
------------------------------------------------------------------------
----------------UAGCGUUACCAUAUCG-------UUAGAGUGGGU--U---CAUAAAUGUCCCGUCU
UGAAACACGGAUCAAGGAGGCCG-U-UUUGUGUGCGA-GUGUAAGA--G-UA-------AUUAA-------A
A-CU--CUGA--CGCGUAUUGAAAG-AAA-----------------GAAU---------------------A
CUCCAAAAGAUC----GUAACU----------------------------------------AC--------
GGU---------------------------------------------------------------------
-------------------UACCU---------------------------------UCUGUAAGGAGU-GC
CCGAAU----CA-UGAG-AAC------------------UCUG--------------UUUCGAAAGGAUUU-
GCG-GU----------------------UGAGCACCUAGAAUGG-GACCCGAAAGG-UUGUGAACUAUGCCU
GAGGAAGGCGAAGUCAGG-GGAAAC-UCUGAUGGAGGCUUGU--CGCAAUGCUGACGUGCAAAUCGC-UUGU
CUAACUUGGGUAUAGGGGCGAAAGACUAAUCGAACAACC-UAGUAGCUGGUUCCUUCCGAAG-UUUCCCUCA
GGAUAGCUGGAG-CAGU--------------------------------------------------------
--------------------AUUC-----UAGUUCCAUCUUGUAAAGACAAUGAUU-AGC-AGUUUCGG-GG
G-C-GUAAU-GCUC-U-CAGCUGAUUCUCAAACUCUGAACGGGUGGGUA-UC-A------------------
------------------UUUUA-AU-----UCA-CU--UAAU--U--GGAUUUUAAAAUUAAAUUGCACAU
GU---GCAA-U---GAAAA-AU-A--G-----G--AGCUCUU-AGUGGGC-C-AUUU-U-UGG-UAAGCAGA
AC-U-GGCGAUGU-GGGUUG---AACCAAAUAUUG-G-GAUAAGACGUCUAAC---AU-UCA-CU-AA---U
---AGAUACC-ACAAAAGGUGUUAGUUCAUUAAGACAGCAGGACGGUGGCCAUGGAAGUCGGUA-UCCGCUA
AGGAGUGUGUAACAACUCACCUGCC-----AAAUGGACUAGCCCUGAAAAU-GGAU-GACGCUAGCAGUGGA
U----------GGUCGAU-GCCCAAUCGUUAAAA--GAAGUGAUA---------------------------
-------------------------------------------------------------------AUA--
----C-------------------------------------------------------------------
----------------------------------UUU-UAAC-GUGUAGGAAGGC-GUGAAGG---------
-----------U-A-ACGUA-GAAGCUU--G-AAUGUGAAUU--C-GAGUGGAGUU-G-UCU-UUAGUGCAG
AUCU-UGAUGG-UAGUA-GCAAAU---AUUC-AAAAGA--AUU-------------UACUUU-GAA-GGCCG
AAGUGGGG-AAGGGUUCC-AUAACAAUG-GAAUUCACUUAUGGGUGAGUCGA--UC-CUAAGGUUUGGGUU-
A-A-CUCU-----CUCUAAU--------AAG-----GUUACU-AGGU---CAUUGGAU-CGAAAGUGAAGGU
GGCU------UUAACACUAGUGAC-----UUUAUA---GG-CC-GA-A-AGGGA-AGCGGGUU-AAAA-UUC
CUGCA----------------------------------CCAUCG---AAUG--------G-GA--------
```

U----------------AU-----------------------------------------------------
----UA-GGG---UAAC---------C-GA-----UC--------------------GUAA-U-CCGG-GAC
AUCAAUU-------------GGCGGU-CGAGGAAGA-GU------UAUCU-------------U-UUCUUG-
UUAACA-UUG-UC--UUGG-G-------------------GU-----CC----------------UCCG-AA
U-C-AG--------------------GUCAA-CUGGAGACGA---------GGAUUCAUCGCAC-AA-UGG-
AAG--AG-----------------------------------------------------------------
----CACAGUC-CUUUGGAUUGGGU-C-UCGCAUC-CGCUAAAUGGU-CCUUGAAA-A-CCG-GAUUAUGGU
-A--------------------------------------------UUU-----AAUCCUAU-UU-GGUGUU
CGUACCAAUAACCACA------------------------------UCAGGUC--UCCAAGGU-GAAUAGCC
UCUGGU-CAAA-UGUA-UUAAUGUA-G-AUAAGGGAAGUCGGC-AAAACCGAUCUGUAACUUCGG-GAUAAG
GAUUG-GCUC-U-AA--AGGCUG------GUGGA-GU------GGACA---UAU---------U--G-----
-GAG------U-----UUGCU---AU----------------------------UUG---------UUUUUA
C---------------------------------------------------------------------UU
-UUA-----------------GGAU-----------------------------------------------
---------------------------------------------------------GG-----------GC
AACU--------------GU--------------------------------UU--UG--------------
---------------------------------------------------------------AAGG-----
------------------------------------------------------------------------
-U-UU---------------AA--G-AUG--GGUG-GU----------------------------------
----AAUU-CU-----UUC-----CA-AUGUG----------------------------------------
----------------------------------------------------------------AGGG----
-------------------------------------------------------------CU---------
---------------U-------------------------------GCU---C------------------
--------------GU--------------------------------------------------------
----UC-------------------------------------UGCUUU-ACG--AUUAACAGC--UAAUUU
AGA--ACUGUGACGAUCACCGGGAAUCCAA-----CUGUUUAAUUAAAACAAAGCAUUGCGAUAAGCUU---
-AA-A-------------AGCUUUUG-ACGCAAUGUGA-UUUCUGCCCAGUGCUCUG-AAUG-UCAAAGUGA
A--------------GAGA--------------UUCAACCUAGCAC--GGGUAAAC-GGCGGGAGUAACUAU
GACUCUCUUAAGGUAGCCAAAUGCCUCGUCAUC-UAAUUAGUGACGCGCAUGAAUGGAUCAAUGAGA--UUC
CCACUGUCCCUAAC----UAC-UAUACAGCGAAACC---AC---UGCAAGGGGAACGGGCCUUGCAAAAACA
GCGGGGAAAGAAGA-CCCUGUUGAGC-UUGACU--CUAGUCU-GAUAUUGCAUAGUGACCUAAAAGGUGUAG
AAU-AGGUGGGAGGG-----------------------------------------------------GCAA
--------------------------------------------------C-C--C-GA-CGGUGAAAUA-C
CACCCCUUUUGGCGUUACUUUGCUAACUU-GG-------A-AU-A-ACAGU------A-----------CC-
------UC--AU-AA------UUCAUU-UUAUG-A--UG-------------GUUUU-GGU----------G
-AAU------AAGCG----------G--------------------------AUCAACCAC---------GG
------------------GUGAAA--------------------------------------UCUGUGC--A
A--AUUG------------GGC---AACUGAUUU---GUA-----UAGCA----------------------
---AAGUAGUCCC----UCUGGU-----------------------------CCCGUA--UUAU--------
-----GUCGA--------------------------------------CCAA-----GAACA-GUUUCAGGU
GGGGAGUUUGGCUGGGGCGGCACAUUUGUUA-AAAGAUAAC-GCAAGUGUCCA-AAGGCAGGCUCAGUGAGA
ACAGAAAUCUCACGUA-GAGUA-AAAGGGCAA-AAGCCUGCUUGAUU-CUGA--------------------
--------------UUUUCA--GUACUAAUCGGAACUGGGAAA-CCAGGGCCUAUCGAUCCUUU--------
---AU-G-----UGCUUAAAU--------------------CU-UAAGCCUAGAGGUGUCAG-AA--AAGUU
ACCACAGGG-AUAACUGGCUUGUGGCAGCCAAGCGCUCAUAGCGACGCUGCUUUUUGAUCCUUCGAUGUCGG

```
CUCUUCUUAUCAUUGUGAAGCA-GAAUUCACAAAGUGUUGGAUUGUUCACCCACUAACAAGGAACGUGAGCU
GGGUUUAGACCGUCGUGAGACAGGUUAGUUUUACCCUACU--------------------G-UUGUC-----
-------------------------------AA--U-UGUUUGCGUAAUAGU-AGC-AUGAUUUAGUACGAG
AGGAA-CUGUCAUGCCGGAUCAC-UGGU-CUGUAGGUUUAUUU-GACA-AAAUAG-U-GACCUGCCGCUACC
AUCCG-UUGGAUAAUGGCUGAACGCC-UC--UAAGUCAGAAUCCAUUCUAGAA---AC--GCAAACCAAAUG
CUUUAGAG-UGUGAAUG-------------------------------------------------------
------------------------------------------------------------------------
------------------------------------------------------------------------
---------------------UUGUAGGUA------------------------------------------
-----------------------ACAUU---AGGUUG---------UUGGUGGGGA----------------
-----CCA--CUU-----UCA--AC--------------UUU------------------------------
----------------------------------------AAACCAUAU------GAUUA-AUCGCUGUUAC
A-------------------------------CUGCAGUUCC--------UUCCGGUUAU-----------G
UG---------------GUGGGUGGCUAAAUUCUAAUUUAUAUCCUC-GUUCCGCU-----CAAC--UCUUC
GAUUGUAGACGACUAUCA--AA-UGAACUAGGU-GCUGUAAG-CUUCC--------GAGUAGC-GU------
----UCA---GUUACGAGGGGUUGAGGCUUUU-CCAU-UA---------GUUCUU--U--------------
------------------------------------------------------------------------
------------------------------------------------------------------------
------------------------------------------------------------------------
-------------------------------------------------
>LSUe_753081_BIGNC/1-3492 ADNK01002490.1:55802-55961,56311-59642 Bigelowiel
---------------------------------------UGACCCUUAGC-GAAGGAUG-U-CUUGGU-UCC
G-G---CAACGAUGAAGAACGCAG--CGAAAUGCGAUACGUAAUGCGAAU--U---GCA--GAA-UUC-AGU
GAAUCAUCAAAUUUUUGAACGCAAAU-U--------GC--------GC----UCCG-U-AGG----------U
GUC-----------CCC---U----U----G---GAG----C-A-UGCC-UCC------UUCAGUGUCG----
---------------------------------------------UGUG-----------UUGAC-CUGA-A
GUG--------------------AG-G-UG-AGAGGACCGGCCGAAUUUAAGCAUAUUAAUAAGCC--GAGG
AAAAGAAACUAA--CAAGGAUUCCCCUA--GU-AA-CGGCGAGUGAAGCGGGA-A-GA-GCUCAAGUUGAUA
AGCG-UUGUCACUU----GUG------------G-----C-AUCGCGUUGUAAAC--UG-------------
--AU-GAGC-GUGGG-----U-------------------------CAGUG--GUG-----------GA---
C-------CUGGG--GGC-------AGGUUG-UUUG------------------------------------
-------GAA-CA-ACGCG--------UUAUAAAGGGUGAGAACCCCGUA---------CAG---ACC-CCGG
AGA---UC--CCCAC-GUACGAC-------------UAC---GUUU--CGAAGAGUCGUGUUGUUUG---GG
AUUGCAGCACUAA-U-UGGGAGGUAAG-CUUCUUCCAAAGCUAAAUAUCAG--C-CGG-AGACCGAUAGC-G
AAC-A-AGUACCGUGAGGGAAAG-AUGAAA-AGAAC-UUUGG-AAAGAG-A-GUUAAAAAGUACCUGAAAUU
G-CUAAGGGG--G-AAG-UG-AUGG---AAAG-UAA-CA------------------------UUGU-----
------------------------AGGUUCAG-----------UUCUAGGU--UC------GU---CUG---
------CUA----------------UAU--------------------CCGU--------------------
-------------------------------------------------------------CGCG-------
-------------UGUCUAACC--------------------------------------------------
------------------------------------------------------------------------
------------------------GG--C--CG----------CAUGGU-----------GG-CAAAGGCG-
---GU----------------------------------------CUG------------------------
------------------------------------------------------------------------
------------------------------------------------------------------------
```

```
------------------------------------------------------------------------
---------------------------------------------------------------AUUGGCUGG
AUUU--------------------------------------------------------------------
------------------------------------------------------------------------
------------------------------------------------------------------------
------------------------------------------------------------------------
------------------------------------------------------------------------
-----------------------GCUGU--------------------------------------------
------------------------------------------------------------------------
-----------------------------------------------GUCAGCUU----------------G
G----------------CUUUUCUUCGUGGGAUAAUGCCUAGUCUAUUGACGACGGCGACUGCCA-------
--------------------------------------UGUUGG----------------------------
---GGAGUUUUUU-UUAAAAGCUGGCGA-----------------AAUGG--U---UUUCAUCGACCCGUCU
UGAAACACGGACCAAGGAGUCUA-A-CAUGUGUGCGA-GUGUUUGG--G-U--------GGUAA--------
A-CC--CUGA--CGCGGAAUGAAAG-UGAAAGGGUG------------------------------------
------------------------------------------------------------------------
------------------------------------------------------------------------
--------------------GG--------------------------------------UUCACCACC-GA
CCGGCC----GU-GGC--------------------------U----------------UGCCAUUGGUCU-
GAG-UA----------------------AGAGCAGAUAUGUUGG-GACCCGAAAGA-UGGUGAACUAUGCCU
GAACAGGGUGAAGCCAGG-GGAAAC-UCUGGUGGAGGCUCGU--AGCGAUACUGACGUGCAAAUCGU-UCGU
CUGAUUUGGGUAUAGGGGCGAAAGACUAAUCGAACCAUC-UAGUAGCUGGUUUCCUCCGAAG-UUUCCCUCA
GGAUAGCUGGAG-CUU--------------------------------------------------------
-----------------------GUAGUACAGUUUUAUCAGGUAAAGCUAAUGAUU-AGA-GGAUGUUG-AG
U-U-GAAAC-AACU-U-GCACCUAUUCUCAAACUUUAAAUAGGUAAGAA-----------------------
------------------GCCGU-UUGAUUUUUU-CU--UGAU--U--GAU---------------------
CA---UCGG-C---CUCGA-AU-G--G-----A--AGCUCUU-AGUGGGC-C-AUUU-U-UGG-UAAGCAGA
AC-U-GGCGAUGC-GGGAUG---AACCGGACGUUU-G-GUUAAGGUGCCCAACU--GG-UCA-CU-CA---U
-CUAGAUACC-AUAAAAGGUGUUGGUUGAUCUGGACAGCAGGACGGUGGUCAUGGAAGUCGAAA-UCCGCUA
AGGAGUGUGUAACAACUCACCUGCC-----GAAUCAACUAGCCCUGAAAAU-GGAU-GGCGCUGA-AGUGAU
C----------GACCUAU-ACCGGACCAUCGGAG--CGAG--------------------------------
------------------------------------------------------------------------
------------------------------------------------------------------------
---------------------------UGCGAAGCUUCCGAU-GAGUAGGAGGAC-GUGGGGG---------
-----------C-G-GUGAC-GCAGCUCG-U-GGCGUGAGCC--A-GAGUGAAACC-A-CCC-CUAGUGCAG
AUCU-UGGUGG-UAGUA-GCAAAU---AUUC-AAAUGA--G---------------AACUUU-GAA-GACUG
AAGUGGAG-AAAGGUUCC-AUGUGAACA-GCAAUUGGACAUGGGUUAGUCGA--UC-CUAAGAGAUAGGGA-
A-A-UUCC-----GUAUGAA--------AGU-----GUG-----CAU---UUAU-----------------U
UG---------------CG----C------CGCC----UA-UC-GA-A-AGGGA-AACAGGUU-AAUA-UUC
CUGUA----------------------------------CCUGGG---CGUG--------G-AU--------
A-----------------A-----------------------------------------------------
---GGG-UGG---CAAC--------AC-AA-----GU--------------------GAAC-U-UGCC-GAC
GGCAGCG-------------GGGAGU-CCGGGAAGA-GU------UAUCU-------------U-UUCUUU-
UUAACA-GCC--U-----G-U-------------------AG-----CC----------------CUGG-AA
U-C-GG--------------------AUUAC-CCGGAGAUAG---------GGUGAU----GUG-GC-UGG-
```

```
UAA--AG-----------------------------------------------------------------
----CACCUCA-CGUCUUGAGGUGU-C-CGGUGCC-UCUCCGACGCC-CCUUGAAA-A-GGC-AAGGGAAGG
-A----------------------------------------------U-----UAUUUGCA-CC-CCAGGU
CGUACUCAUAACCGCA------------------------------UCAGGUC--UCCAAGGU-GAACAGCC
UCUAGU-C-GA-UAGA-ACAAUGUA-G-GUAAGGGAAGUCGGC-AAAAUAGAUCCGUAACUUCGG-GAAAAG
GAUUG-GCUC-U-UA--GGGCUG-G----GGAGG-UU------GGG--------------------------
-------CCUUA-GAUUAGAA---GGAGCCACCUCGAGCC--------------------------------
------------------------------------------------------------------------
----------------------GG------------------------------------------------
-----------------------------------------------------------------------A
UCGG--------------C---------------------------------GG--AA--------------
---------------------------------------------------------------CGCA-----
---AG---------------------------UUUAGCUGAGAACGCU--U---------------------
-G------------------UU--G-GU---GCGCUGU----------------------------------
---GGA----------CGG-----UCUUUGGU----------------------------------------
----------------------------------------------------------------AGUC----
-------------------------------------------------------------UU---------
------------------------------------------------CG----------------------
------------------------------------------------------------------------
----GG-----------------------CG------------UCCUCC-UCC--UAGUACAGC--UAACUA
AGA--ACUGGAGCGGACAAGGGGAAUCCGA-----CUGUUUAAUUAAAACAAAGCAUUGUGAUGGCCCU---
-CA-C-------------GGGUGUUG-ACACAAUGUGA-UUUCUGCCCAGUGCUCUG-AAUG-UCAACGUGA
A--------------GAGA--------------UUCAAGCAAGCGC--GGGUAAAC-GGCGGGAGUAACUAU
GACUCUCUUAAGGUAGCCAAAUGCCUCGUCAUC-UAAUUAGUGACGCGCAUGAAUGGAUUAACGAGA--UUC
CCACUGUCCCUAUC----UAC-UAUCUAGCGAAACC---AC---AGCCAAGGGAACGGCUUGGAAUAAUCA
GCGGGGAAAGAAGA-CCCUGUUGAGC-UUGACU--CUAGUCC-GACUUUGUGAACUCACCUAAGAGGUGUAG
AAU-AAGUGGGAGC------------------------------------------------------UUCG
----------------------------------------------------G--C-GC-CGGUGAAAUA-C
CACUACUCUUGGAGUGAGUUUACUCAUUA-CA-------U-UA-U-GCAAG------GC----------CG-
------CG-----GU-------UCGC---GCCGCU--GG--------------UUCU---------------
-----------AGCG--------------------------------------UUA-----------CGUCG
------------------------------------------------------------------------
---GUU---------------------------UCG-UGC--------------------------------
----------CGG------CAA------------------------------CUUG----CG----------
------------------------------------------------UGUA-----AGACA-AAGUCAGGU
GGGGAGUUUGGCUGGGGCGGCACAUCUGUUA-AACGAUAAC-ACAGGUGUCCU-AAGGUAAGCUCAAUGAGA
ACAGAAAUCUCAUGUA-GAACA-AAAGGGUAA-AAGCUUACUUGAUU-CUGA-------------------
--------------UUUUCA--GUAUGAAUACGAACCGUGAAA-GCGUGGCCUAUCGAUCCUUU--------
---AG-A-----UUUUAGAAG--------------------CU-UUAAGCUAGAGGUGUCAG-AA--AAGUU
ACCACAGGG-AUAACUGGCUUGUGGCAGCCAAGCGUUCAUAGCGACGUUGCUUUUGAUUCUUCGAUGUCGG
CUCUUCCUAUCAUUGGCCCGCA-GCAGAGCCAAAGUGCAGGAUUGUUCAUCCACCAACAGGGAACGUGAGCU
GGGUUUAGACCGUCGUGAGACAGGUUAGUUUUACCCUACU--------------------G-AUG-------
-------------------------------AA--U-GGUCGUGGCGACAGU-AAU-CCAAUUUAGUACGAG
AGGAA-CCGUUGGUUCACGCAUU-UGGC-AA-UGCGGUUGAUU-GAAA-CGUCAA-U-GCCGCGAAGCUACC
ACGUG-CAGGAUUAAGGCUGAACGCC-UC--UAAGCCCGAAUCCAGGCUGG-----AC-AGCCACGAU-GUG
```

```
CGCACUCC-GAUGUA---------------------------------------------------------
------------------------------------------------------------------------
------------------------------------------------------------------------
-----------------------GA-----------------------------------------------
--------------------------------GACUA-----------------------------------
----------AGU-----AUA--G------------------------------------------------
-----------CC--UUGGC------CA---AGA----GUUACCAGCC-------GACAAAA-CGGCGAGCG
UUG------------------------------------------------AUCACA-AGUGUAC------U
C-----------------AAACAUC---------------------G-GAAGUGAC-----AAA------UC
CCUUGUAGACGACUUGAC-A---UGAAAGAGGU-AUUGUAAG-CUGGA--------GAGUGC--CA------
----UUU--UGGCACGAUCCGCUGAGAUUAAG-CCUU-CGUUUCUU---CGAUUUGUC--------------
------------------------------------------------------------------------
------------------------------------------------------------------------
------------------------------------------------------------------------
-------------------------------------------------
>LSUe_296543_THAPS/1-3538
---------------------------CUAU----UA--CAACUUUCAGC-AAUGGAUG-U-CUUGGA-UCC
C-A---CAACGAUGAAGAACGCAG--CGAAAUGCGAUACGUAAUGCGAAU--U---GCA--GGA-CCU-CGU
GAAUCAUCAAACUUUUGAACGCACAU-U--------GC--------GC----UUUU-G-GGU---------A
A-U-----------GCC---C----A----A---GAG----C-A-UGCU-UAU-----CUGAGUGUCC-CGA
AUUUC------------------------------------AUC------------U--AUGGAU-CUCA-G
AUU--------------------AA-G-UA-AGACUACCCGCUGAAUUUAAGCAUAUUAAUAAGCG--GAGG
AAAAGAAACUAA--CAAGGAUUCCCCUA--GU-AA-CGGCGAGUGAAGCGGGA-A-GA-GCUCACCAUGUGA
AUCU-AUGUAACCCUUC-GGG------------G---UGC-AUCGAAUUGUGGUC--UG-------------
--GA-GAAG-UAC-U-----G-------------------------UCGGC--CGU-----------GG---
U-------UCCGG--GCC-------AAGUCU-CUUG------------------------------------
-------GAA-AA-GAGCA-------GCUGAGAGGGUGAAACUCCCGUU---------CUU---G-C-CUGG
AAA---CA--UUGUG-CCUUGGC-------------AUAU-GCUUU--CUAUGAGUCGAGUUGCUUG---GG
AUUGCAGCUCAAA-U-UUGGUGGUAAA-UUCCAUCUAAAGCUAAAUAUUGG--U-GGG-AUACCGAUAGU-G
CAC-A-AGUACCGUGAGGGAAAG-AUGCAA-AGAAC-UUUGA-AAAGAG-A-GUUAAAGAGUACCUGAAAUU
G-UUGAAAGG--G-AAG-CG-AAGG---AAAC-CAG-UG------------------------C--------
-------------------------------------------CAACGUUG--UG------UC---AUA---
------CUU----------------UCU--------------------CUGC--------------------
-------------------------------------------------------------CGCUUGC----
-----------------------GG-----------------------------------------------
----------------------------------UAG-----------------------------------
------GGACG------CU---GUGG--C--GC----------UACAUG-----------GG-UCAGCAUC-
---GG----------------------------------------CUUU-----------------GUGCUU
GGG-----------------UCAAA------------------CGUUCAGUUGGUAGGC----GGCUC----
--------------------------------------------------------------C--U-CG---
------------------------------------------------------------------------
------------------------------GAGUUAGUG------CCUUC---UGUUCAUGUCCUGGGUCGG
GCUGAGGUC---------------------------------------------------------------
------------------------------------------------------------------------
------------------------------------------------------------------------
```

```
------------------------------------------------------------------------
--------------------------------------------------AG--------------------
---------------------UCACUAU--------------------------------------------
------------------------------------------------------------------------
-----------------------------------------------GUGCUCGU-----------------
------------------------------------------------------------------------
------------------------------------------------------------------------
-----------------GAUGCUGACGA-----------------AAUGG--U---UUUCUUUACCCCGUCU
UGAAACACGGACCAAGGAGUCUA-A-CAUAUGUGCGA-GUACAAAG--G-U--------GGCAA--------
A-CC--UGCG--UGCGCAAUGAAAG-UAA-----------------CAGU---------------------G
GUUG-GAUCCC---------------------------------------------------UC--------
CG----------------------------------------------------------------------
-------------------GGG---------------------------------------UGCACAAUC-CG
CCGGCC----AA-UAUCC---------------------UUCG----------------GGAGAAAGGUCU-
GAG-UG----------------------UGAGCAUACAUGUUGG-GACCCGAAAGA-UGGUGAACUAUGCCU
GAGUAGGGCGAAGCCAGG-GGAAAC-UCUGGUGGAGGCUCGU--AGUGAUUCUGACGUGCAAAUCGA-UCAU
CAAAUUUGGGUAUAGGGGCGAAAGACUAAUCGAACCAUC-UAGUAGCUGGUUCCCUCCGAAA-UUUCCCUCA
GGAUAGCUGCAA-UUAA-------------------------------------------------------
-----------------------U-----CAAUUUUAUCAGGUAAAGCGAAUGAUU-AGA-GGUUCCGG-GG
G-C-GUAAU-GCCC-U-CGGCCUAUUCUCAAACUUUAAAUGGGUAAGAA-----------------------
------------------CCCAG-GG-----UUG-CU--UAAU--U--GAA---------------------
CC---CUGG-C---GUCGA-AU-G--A-----A--GGUUGCU-AGUGGGC-C-ACAC-C-GGG-UAAGCUGG
UU-U-GGCGAUGC-GGGAUG---AUCCGAACGUUU-A-GUUAAAGUGCCAAAGU--AU-UCG-CU-CA---C
-CUAGAUCCC-ACAAAAGGUGUUGGUCAUUUAGACAGCAGGACGGUGGCCAUGGAAGUCGGAA-UCCGCUA
AGGAGUGUGUAACAACUCACCUGCC-----GAAUGAACUAGCCCUGAAAAU-GGAU-GGCGCUGA-AGCGAA
U----------CACUUAU-ACUAAACCACCAGGC--CAAG--------------------------------
------------------------------------------------------------------------
------------------------------------------------------------------------
---------------------------UGCCAAGGCC-UGGU-GAGUAGGAGGGC-GUGAAGG---------
-----------U-U-GUUGU-GCAGCUUA-U-ACCGUGAGGU--U-AAGUGAAACG-G-CCU-UUAGUGCAG
AUCU-UGGUGG-UAGUA-GCAAAU---AUUC-AAAUGA--G---------------AACUUU-GAA-GACUG
AAGUGGGG-AAAGGUUCC-ACGUGAACA-GCAAUUGGACGUGGGUUAGUCGA--UC-CUAAGAGAUAGGAG-
A-A-AUCC-----GUUUCAA--------AGU-----GAG-----UGU---UCUU-----------------C
GG---------------UUCGCUC------CGCC----UA-UC-GA-A-AGGGA-AUCUGGUU-AAUA-UUC
CAGAA----------------------------------CCAGGA---CGUG--------G-AU--------
A----------------UU-----------------------------------------------------
---GAG-UGG---UAAC---------AC-AA-----GC--------------------AAAG-C-CGGG-UAC
GAUGGCG-------------GGAGCC-CUGAGGAGA-GU------UCUCU-------------C-UUCUUC-
UUAACA-AGC-AC----AC-U--------------------AA-----CC----------------CUGG-AA
U-C-GG--------------------AUUGA-CCGGAGAUAG----------GGUUGG----AGU-CU-UGG-
GAA--AG-----------------------------------------------------------------
----GUGGAUA-GUUUUUGUCUGCU-U-CGGUGCG-CUCUCGUUGUC-CCUUGAAA-C-CCC-GGUGGCGUG
-A---------------------------------------------CU-----UAUUCGCA-CC-CCUGGU
CGUACUGAUAACCGCA------------------------------UCAGGUC--UCCAAGGU-UAGCAGCC
UCUAGU-U-GA-UAGA-AUAAUGUA-G-GUAAGGGAAGUCGGC-AAAAUAGAUCCGUAACUUCGG-GAAAAG
```

```
GAUUG-GCUC-U-GA--GGGUUG-G----GCCUG-GG------GGUC----CCU---------C--GUU---
-UGA------UG-UUGCGAGG---AC----------------------------UAG---------GAG---
----------------------------------------------------------------------AG
ACUG-----------------GCU------------------------------------------------
-----------------------------------------------------------------------C
ACGC--------------U---------------------------------GG--AC--------------
---------------------------------------------------------------UUUC-----
------------------------------------------------------------------------
-C------------------AG--G-UC---UUCCUUC----------------------------------
---GGA----------CGG-----CGGGGGA-----------------------------------------
----------------------------------------------------------------UGUG----
-------------------------------------------------------------CU---------
---------------U-------------------------------GCA----------------------
------------------------------------------------------------------------
----CU-----------------------UU------------CCCCAGUGCG--UAGAACAAC--CAGCUC
AGA--ACUGGAGCGGACAAGGGGAAUCCGA-----CUGUUUAAUUAAAACAAAGCAUUGCGACGGCCGG---
-AG-A-------------CGGUGUUG-ACGCAAUGUGA-UUUCUGCCCAGUGCUCUG-AAUG-UCAAAGUGA
A--------------GGAA--------------UUCAACCAAGCGC--GGGUAAAC-GGCGGGAGUAACUAU
GACUCUCUUAAGGUAGCCAAAUGCCUCGUCAUC-UAAUUAGUGACGCGCAUGAAUGGAUUAAUGAGA--UUC
CCACUGUCCCUAUC----UAC-UAUCUAGCGAACCC---AC---AGCCAAGGGAACGGGCUUGGAAUAAUCA
GCGGGGAAAGAAGA-CCCUGUUGAGC-UUGACU--CUAGUCC-GACGUUGUGAAAUGACUUAUGAGGUGUAG
AAU-AAGUGGGAGC------------------------------------------------------UUCG
----------------------------------------------------G--C-GC-ACAUGAAAUA-C
CACUACUCGUAAUGUUGUUUACUUAUCA-UG--------U-GA-C-GUAGC------AC----------CC-
------AGCCGU-CU-------UCAC--GGGCGCC--UG-------------GUUCU---------------
-----------AGAU--------------------------------------UUAA--GU---------CA
------------------------------------------------------------------------
---ACU---------------------------U---GUU--------------------------------
----------UGG------CGA------------------------------GCUA----CG----------
------------------------------------------------CAUG-----AGACA-ACGUCAGGU
GGGGAGUUUGGCUGGGGCGGCACAUCUGUUA-AACAAUAAC-ACAGGUGUCCU-AAGGUCAGCUCAAUGAGA
ACAGAAAUCUCAUGUA-GAACA-AAAGGGUAA-AAGCUGACUUGAUU-UUGA--------------------
--------------UUUUCA--GUACGAAUACAAACCGGGAAA-CCGUGGCCUAUCGAUCCUUU--------
---CA-G-----UUUUUAGAA--------------------UU-UGAAGUGAGAGGUGUCAG-AA--AAGUU
ACCACAGGG-AUAACUGGCUUGUGGCAGCCAAGCGUUCAUAGCGACGUUGCUUUUUGAUUCUUCGAUGUCGG
CUCUUCCUAUCAUUGUGUCGCA-GAAGUCACCAAGUGUAGGAUUGUUCACCCACUAACAGGGAACGUGAGCU
GGGUUUAGACCGUCGUGAGACAGGUUAGUUUACCCUACU---------------------G-AUG-------
-------------------------------AA--U-UGUUGUCCCGACAGU-AAU-CCAACUUAGUACGAG
AGGAA-CCGUUGGUUCAGAUAUU-UGGU-AA-UGCGGUUGGCU-GAAC-AGCCAA-U-GCUGCAAAGCUACC
AUCUG-UAGGAUUAUGACUGAACGCC-UC--UAAGUCAGAAUCCAUGCUGG-----CA--GGGAUGAU-AAC
CCCCUUGA-CCACUGUU-------------------------------------------------------
------------------------------------------------------------------------
------------------------------------------------------------------------
---------------------ACCU-----------------------------------------------
--------------------------------GAGUA-----------------------------------
```

```
----------GCA-----AUA--GU--------------CAC------------------GUAUCAA---C-
-------ACCACC--CCCUGG-----UA---GCG--CUGUACACUAAAGU-----ACAA-----GCUUGC--
-------------------------------------CG-----------AAACACA-AUU----------U
C-----------------CAAUGAU---------------------G-GUCAAGUC-----AAA------UC
CUUUGUAGACGACUUAAU-UAA-GGAACGUGGU-ACUGUAAG-CAUGA--------GAGUAGC-CU------
----CUG--UGCUACGAUCUGCUGAGGUUCUG-CCUC-UGUUCCUG---UGAUUUGUG--------------
------------------------------------------------------------------------
------------------------------------------------------------------------
------------------------------------------------------------------------
-------------------------------------------------
>LSUe_238746_MICPO/1-3597 GU001165.1:375-526,975-4419 Micronuclearia podove
------------------------------------------CUUCUAAC-AACGGAUG-U-CUUGGC-UCU
C-G---CAUCGAUGAAGAACGCAG--CGAAAUGCGAUACGUAAUGUGAAU--U---GCA--GAA-UU--CGU
GAAUCAUCGAGUUUUUGAACGCAUAU-U--------GC--------AC----UUGC-C-AGU--------UU
A--------------CU---G----G----U---GAG----U-A-UGUC-UUC-----UUGAGCGUCU----
-----------------------------------------------------------CCGGAC-CUCA-A
GUG--------------------AG-A-CA-AGGCUACCCGCUGAAUUUAAGCAUAUAAAUAAGCG--GAGG
AAAAGAAACUAA--CAAGGAUUGCCUUA--GU-AA-CGGCGAGUGAAGCGGCA-A-AA-GCUCAAGGUUAAA
AUCU-GGACUUUC--------------------A---GUC-UCCGAAUUGUAGUC--UA-------------
--AA-GAGG-CAA-------G-------------------------UUUGA--CGA-----------CG---
CG------UCUGG--UAU-------AAGUCC-CUUG------------------------------------
-------GAA-UA-GGGUA-------CCAUAGAGGGUGAGAGUCCCGUA---------CAU---GAC-UGGA
AGU---UG--CGUUG-UUAUAUA-------------AACU-GCUUU--CGAAGAGUCGGGUUGUUUG---AG
AUCGCAGCCCAAA-G-UGGGAGGUAAA-UUCCUUCUAAAGCUAAAUAACGG--C-GAG-AGACCGAUAGC-G
AAC-A-AGUAGAGUGAUCGAAAG-AUGAAA-AGAAC-UUUGA-AAAGAG-A-GUUAAA-AGGACCUGAAAUU
G-UUAGAAGG--G-AAG-CG-AUUG---CAAG-CAGUGGAUU--------------------GUUUUUUGGA
AAUUAAGUCGGGGCAUUGGUAGCGGGUGUUUG-----------AUUUAAGA-------U-CGC--AAGA---
------CUU-----AGGUUAGA---UAU--------------------------------------------
-------------------------------------------------------------CUUACGUUAUU
A----------------------UG-----------------------------------------------
----------------------------------UGA-----------------------------------
------CGGUGA--UUUUU---CCAG-----------------AGAACA-----------GG-UCAGCAUG-
---GC--------------U-------------------UCGAUCCUUU---------GCAAGAGCGUUUCG
GGA---------------AGGUAGGU----------------CUCAGAGCAAUCUGGAC-------------
------------------------------------------------------------UGU--U-AUAG-
------------------------------------------------------------------------
---------------------AC------------------------------CGUUACGGGCAGCAGGAGG
AUUGGAGUG---------------------------------------------------------------
------------------------------------------------------------------------
------------------------------------------------------------------------
------------------------------------------------------------------------
-------------------------------------------------AGGAAA-----------------
------UAAA------UUAUUUCGGGAU--------------------------------------------
------------------------------------------------------------------------
-----------------------------------------------AUGCCGCA----A-----------G
```

```
G----------------CGUUCUUU-----------------------------------------------
--------------------------------------UAUUGA----------------------------
---ACGAUAUCCC-CAAGAUGCUGGCGA-----------------AAUGC--U---UGUAAUUGACCCGUCU
UGAAACACGGACCAAGGAGUCUA-A-CAUGUGUGCGA-GUGUUCGG--G-UG-------AUUGA-------A
A-CC--CGUA--CGCGAAGUGAAAG-CGA----------------------------------------GAG
GUGG-GAGUCCA----AGU-------------------------------------------UU------AA
ACA---------------------------------------------------------------------
-------------------GGA--------------------------------------UGCACCAUC-GG
CCGACC----AU-GACAA---------------------CAUUG---------------UUCGAACGGUUC-
GAG-CA----------------------GGAGCAUACACGUUGG-GACCCGAAAGA-CGGUGAACUAUGCCC
GAAUAGGAUGAAGCCAGA-GGAAAC-UCUGGUGGAGGUCCGU--AGAGAUUCUGACGCGCAAAUCGA-UCUU
CAAAUUUGGGUAUAGGGGCGAAAGACUAAUCGAACCGUC-UAGUAGCUGGUUCCCUCCGAAG-UUUCCCUCA
GGAUAGCUGGAA-CUUG-------------------------------------------------------
-------------------UAGACA----CAGUUUUAUCCGGUAAAGCGAAUGAUU-AGC-GGUUUGGG-GG
A-U-GAAAC-AUCC-U-CCGCCGAUUCUCAAACUUUAAAUGGGUAAGCU-----------------------
------------------GCUGU-GG-----UUG-CU--UAAG--U--GAA---------------------
CC---AUGG-C---GUAGA-AU-G--GA----A--AGUUCCA-AGUGGGC-C-AUUU-U-UGG-UAAGCAGA
AC-U-GGCGAUGC-GGGAUG---AACCGAACGUUG-U-GUUAAGGCGCCUAAAU--GU-GCG-CU-CA---U
--GAGACCCC-ACAAAAGGUGUUGGUUCAUUUAGACAGCAGGACGGUGGCCAUGGAAGUCGGAA-UCCGCUA
AGGAGUGUGUAACAACUCACCUGCC-----GAAUGAACUAGCCCUGAAAAU-GGAU-GGCGCUGC-AGCGUG
C----------UGCCUAU-ACACAACCAUGGUUG--CAAA--------------------------------
------------------------------------------------------------------------
------------------------------------------------------------------------
----------------------------UGCGAGGCAG-CCAU-GAGUAGGAGGC-GUGGAGAUCCUGAUAU
GCAAAUAUCAUU-AUGCGAC-GAAGGCUU-G-UGCGUAAGCG--C-GGCUGGAGCG-G-UCU-CUAGUGCAG
AUCU-UGGUGG-UAGUA-GCAAAU---AUUC-AAGUGA--G---------------AUCCUU-GAA-GACUG
AAGUGGAG-AAAGGUUUC-AUGUGAACA-GUGAUUGAACAUGAGUUAGUCGA--UC-CUAAGAUAUAGGGU-
A-A-UACC-----GUUUAAA--------AGU-----GGG-----CUA---UAAUU----------------G
UG---------------UAGCCCC------GCCU----UA-UC-GA-A-AGGGA-AGCAGGUU-AAUA-UUC
CUGCA----------------------------------CCGAAC---GGCG--------G-AU--------
A----------------CU-----------------------------------------------------
---GAG-CGG---CAAC--------GC-AA-----AU--------------------GAAC-U-CGCA-GAC
AUUGGCG-------------AAAGUC-CUGGCAAGA-GU------UCUCU-------------U-UUCUUG-
UUAACA-GUC--C----GA-G------------------GCG-----CC----------------AUGG-AA
U-C-GG--------------------AUUAU-CCGGAGAUAU---------GGCUAU----GUG-GC-UGG-
UAA--AG-----------------------------------------------------------------
----CACCCUG-C-UUAAGGGGUGU-C-CGGAGCG-CUUUCGACAAU-CCUUGAAA-A-UGC-GAGGGAAUG
-G---------------------------------------------AU-----UAUUUUCU-CG-UUCGGU
CGUACUCAUAACCGCA------------------------------UCAGGUC--UCCAAGGU-GAGUAGCC
UCUAGU-C-GA-UAGA-AGAAUGUA-G-AUAAGGGAAGUCGGC-AAAAUAGAUCUGUAACUUCGG-GAAAAA
GAUUG-GCUC-U-AA--GGGUUG-G----GUGCU-GG------CGGACC--GAUACUCGAG-GC--GAC---
-GGA--------------------------------------------------------------CUG--A
C-----------------------------------------------------------------------
--------------------UGUU------------------------------------------------
------------------------------------------------------------------------
```

----------------------------------------------------GG--AC--------------
--------------------U-------A----------------------------------CCGG-----
--------------------------------------------------U---------------------
-A-CG---------------CA-AGUGCU--GGCGGAC----------------------------------
---UGG----CG-GA-CGGA----UCGACGAA----------------------------------------
----------------------------------------------------------------GACG----
------------------------------------------------------GUG--UCGUGUG------
---------------U-G-------------------------AGGAGCA---A------------------
----U-------------------------------------------------------------------
----CU-----------------------UUACUUCA------CUGCUGCACA--UGAAACAAC--CAACUU
AGA--ACUGGUACGGACAAGGGGAAUCCGA-----CUGUCUAAUUAAAACAAAGCGUUGCGAUGGGGUC---
-AG-A------------GACCUGUUG-ACGCAACGUGA-UUUCUGCCCAGUGCUCUG-AAUG-UCAAAGUGA
A--------------GAAA--------------UUCAACCAAGCGC--GGGUAAAC-GGCGGGAGUAACUAU
GACUCUCUUAAGGUAGCCAAAUGCCUCGUCAUC-UAAUUAGUGACGCGCAUGAAUGGAUUAACGAGA--UUC
CCACUGUCCCUAAC----UAC-UAUCUAGCGAAACC---AC---AGCUAGGGGAACGGGCCUAGCAGAAUCA
GCGGGGAAAGAAGA-CCCUGUUGAGC-UUGACU--CUAGUCC-GACUUUGUGGAAUGACUUGAGAGGUGUAG
AAU-AAGUGGGAGC------------------------------------------------------UUCG
----------------------------------------------------G--C-AA-UCUUGAAAUA-C
CACUACUUUCAAUGUUAUUCUACUUAUUU-AG-------U-GA-G-GUGGA------AG----------CG-
------GGAGUAAUC-------CCAG--------------------------UUUCU---------------
-----------AGCA--------------------------------------UUAC--GC-----------
------------------------------------------------------------------------
---GUA---------------------------UUU-ACU--------------------------------
----------ACG------UCA------------------------------UCCA----AG----------
------------------------------------------------CUAA-----AGACA-UAGUCAGGU
GGGGAGUUUGGCUGGGGCGGCACAUCUGUUA-AACGAUAAC-GCAGGUGUCCU-AAGGUGAGCUCAAUGAGA
ACAGAAAUCUCAUGUA-GAACA-AAAGGGUAA-AAGCUCACUUGAUU-UUGA--------------------
--------------UCUUCA--GUACUGAUACAAACUGUGAAA-GCAUGGCCUAUCGAUCCUUU--------
---GG-U-----CCUCCGGAA--------------------UU-UGGAGCUAGAGGUGUCAG-AA--AAGUU
ACCACAGGG-AUAACUGGCUUGUGGCAGCCAAGCGUUCAUAGCGACGUUGCUUUUUGAUCCUUCGAUGUCGG
CUCUUCCUAUCAUUACGAAGCA-GAAUUCGUAAAGUGUUGGAUUGUUCACCCACUAACAGGGAACGUGAGCU
GGGUUUAGACCGUCGUGAGGCAGGUUAGUUUUACCCUACU--------------------G-AUG-------
-------------------------------AA--G-UGAUGUCCCAAUAGU-AAU-UCAACUUAGUACGAG
AGGAA-CAGUUGAUCCGCGCAAU-UGGU-AA-UGCGGUUGUUU-GAAA-AGACAU-U-GCCGCAAAGCUAUC
ACGCG-UAGGAAAAUGACUGAACGCC-UC--UAAGUCAGAAUCCAUGCUAG-----AA-CGGGACGUU-GUC
UCCCCCGG-GCAUGUUU-------------------------------------------------------
------------------------------------------------------------------------
------------------------------------------------------------------------
---------------------AGUU-----------------------------------------------
------------------------------------------------------------------------
----------------------------------------------------------GUGUAUCAA---UA
GGGCU--UCGGCCCAGAAACCA--UACCAUUAAU-CAA-GUUGUUGGAC------GGAA---GCGUCUGAUG
G---------------------------C----U-AUUAGUGA--------ACCUCA-AUU----------C
C-----------------AAAUAUU---------------------G-CCUGGGAU-----AAA------UC
CUUCGCAGACGACCUAGU-U---GGAAC--------------------------------------------

```
------------------------------------------------------------------------
------------------------------------------------------------------------
------------------------------------------------------------------------
------------------------------------------------------------------------
-------------------------------------------------
>LSUe_353151_CRYHT/1-3519
---------------------A----ACACC----UA--ACACUUUAAGU-AAUGGAUG-U-CUUGGU-UCU
C-A---UAACGAUGAAGGACGCAG--CAAAGUGCGAUAAGCAAUGUGAAC--U---GCA--UAA-UUC-UAU
GAACAAUCAGAUCUCUCAACGCAAAU-G--------GC--------AG----UAAU-U-------------U
U-A-----------------U----A----U---UAC----U-A-UAUU-UAU-----AUCAGUGUCU-AUU
GACUACUUCA-------------------UAUAAUAUAAUGUUU------------A--AAAGAC-CUGA-U
AUU--------------------AA-G-UA-AGAUUACCCACUGAACUUAAGCAUAUUAUUAAGUG--GAGG
AAAAGAAACUAA--CUAGGAUUUCCUUA--GU-AA-CUGCGAGUGAACAGGAA-A-GA-GCUCAACAUGAAA
AUCA-AUAAUAAUUAUAUUUU------------A---UUA--UUGACUUGUAGUA--UC-------------
--UA-GAAG-UAA-------U-------------------------GCCAG--UGA-----------AU---
UA------AUAAA--AUU-------AAGUGU-UUUG------------------------------------
-------GAA-UA-AACCA-------CCAUAGAGGGUGAAAGUCCCGUC---------UAU---A-U-UUUA
UUUU-UAA--UUCAC-AUACG---------------GUUU-ACUAU--CAAAGAGUCAUAUUCCUUG---GG
AAUGGAGUAUAAA-A-UGGGUGGUAAU-UUUCAUCCAAAGCUAAAUAUUGA--U-GAG-AAACCGAUAGC-A
AAC-A-AGUACCGUGAGGGAAAG-AUGAAA-AGAAC-UUUGA-AAAGAG-A-GUUAAA-AGUACCUGAAAUU
A-CUUAAAGG--G-AAG-CA-AAUA---AAAC-CUA-AC---------------------------------
-------------------------------------------AUUUAAAA--AG----------G------
------------------------------------------------------------------------
-----------------------------------------------------------------UA-----
------------------------------------------------------------------------
---------------------------------------------------------------------AAU
U---------------CUU---UUAG--A--AU----------UGUUAUA----------GUUUUAAUAUU-
-UAAA--------------------------------------------------------------UAUAA
GAA-----------------AUGAAUGAGGGAA---------------------------------------
-------------------------------------------------------------UG--U-AUA--
------------------------------------------------------------------------
-----------AACAUUCAAAGAAUAAUAAUUUUUAAUA------UAUUAUUAUAUUAUUGUU---------
------------------------------------------------------------------------
------------------------------------------------------------------------
------------------------------------------------------------------------
------------------------------------------------------------------------
-------------------------------------------------A----------------------
------------------------------------------------------------------------
------------------------------------------------------------------------
-------------------------------------------------UUCUAA-----------------
------------------------------------------------------------------------
------------------------------------------------------------------------
-----------------GAUAUUGGUAU---------------AAAAUAG--G---UUUAUCUGACCCGUCU
UGAAACACGGACCAAGGAGUCUA-A-CAUGUGUGCAA-GUAUAUAA--G-UA-------AUUAA-------A
A-CU--UAUA--UGCAUAAUGAAAG-UGA-----------------CAAU----------------------
```

```
-UGG-GAUUUUU----UUUA------------------------------------------AA--------
AGA---------------------------------------------------------------------
-------------------AAA--------------------------------------UGCACCAAU-GA
CCGACA----UU-AAUU-UCA------------------UUUU--------------UUGAAUUAAAGUUU-
GAG-UA----------------------UGAGCAUAAAUGUUAG-GACCCGAAAGA-UGGUGAACUAUGCCU
GAGUAGGGUGAAGCCAGG-GGAAAC-UCUGGUGGAAGCUCGC--AGCGAUACUGACGUGCAAAUCGU-UCGU
CAAACUUGGGUAUAGGGGCGAAAGACUAAUCGAACCAUC-UAGUAGCUGGUUCCCUCCGAAG-UUUCCCUCA
GGAUAGCUGGAG-UUUA-------------------------------------------------------
---------------UUUUUAAUU-----UAGUUUUGUCAGGUAGAGAUAAUGAUU-AGA-GGCAUCUG-GG
A-C-AUGUU-GUCC-U-AGACCUAUUCUCAAACUUCGAAUAGGCAAGUA-----------------------
------------------CUAAU-AA-----UUA-CU--UAAU--G--AAU---------------------
UA---UUAG-A----AUUA-AU-A--U-----U--AACUCUU-AGUGGGC-C-AUUU-U-UGG-UAAGCAGA
AC-U-GGCGAUGA-GGGAUG---CUCCUAACGUUU-A-GUUAAGGUGCCUAAAU--GU-AUG-CU-CA---U
-C-AGAUACC-AUAAAAGGUGUUGGUUCAUUUAAACAGCAGGACGGUGGUCAUGGAAGUCGAAA-UCCGCUA
AGGAGUGUGUAACAACUCACCUGCC-----GAAUGAACCAGCCCUGAAAAU-GGAU-GGCGCUGA-AGCAUA
U----------UACCGAU-ACUAAACCAUUAUUU--UUAA--------------------------------
------------------------------------------------------------------------
------------------------------------------------------------------------
---------------------------UUAUAUAAAA-UAAU-GUGUAGGAGGGC-GUAAAAG---------
-----------U-U-GUUUU-GAAGUAUA-A-GAUGUAAAUC--U-AUAUGGAACG-G-CUU-UUAGUGCGG
AUCU-UGGUGG-UAGUA-GCAAUU---AUUC-AAAUGA--G---------------AACUUU-GAA-GACUG
ACGUGGAG-CAGGGUUCC-AUGAGAACA-GUAUUUGGACAUGGGUUAGUCGG--UC-CUAAGUGAGAGGUU-
A-A-CACC-----AAUAAAA--------UA------AAU----UAUU---UUU-------------------
-----------------UAUAUUU------AUAC----UC-AC-AA-A-AGGGA-AUCAGGUU-GAUA-UUC
CUGAA----------------------------------CCAGAA---UAUG--------G-AU--A-----
U----------------UU-----------------------------------------------------
---AUA-UGG---UAAC--------AU-AA-----AA--------------------GAAC-U-CUAA-AAC
ACAAGUA-------------UAAAUC-CCGAAUAGC-GU------UAUCU-------------UUUGCUAU-
UUAACA-AUC--U-----A-U-------------------UA-----CC----------------AUGG-AA
U-U-AG--------------------UUCAG-CUAGAGAAAU---------GGUUAU----AUG-AU-UGG-
AAU--AG-----------------------------------------------------------------
-----AAUUCU-CUUUGUGGAAUUU-U-CGGUGCA-UUUAUAAUUGU-CCUUGAAA-A-UUA-GAGGGAUAG
-A---------------------------------------------UU-----UAUUUUCA-UA-UCUGGC
CGUACCGAUAACCGCA------------------------------UCAGGUC--UCCAAGGU-UAACAGCC
UCUGGU-C-AA-UAGA-UUAAUGUA-G-AUAAGGGAAGUCGGC-AAAAUAGAUCCGUAACUUCGGGGAAAAG
GAUUG-GCUC-U-AA--GGGUUG-A----GUAUA-UA------GACC----CAU---------C--AAU---
-UAU-------------UAUA---UU----------------------------AAG---------CUG---
----------------------------------------------------------------------UU
-UUUUUGACUU---------UAUA------------------------------------------------
---------------------------------------------------------AUCUUU-------AU
UGGA------------------------------------------------UU--AU--------------
--------------UUUGAAU------------------------------------------UUAA-----
------------------------------------------------------------------------
-A------------------AC--G-GCAA-AAUA-------------------------------------
----UA----------UAA-----UU----------------------------------------------
```

------------------------------------------------------------------------
------------------------------------------------------------------------
----------------------------------------------AAUG---G------------------
--------------AU--------------------------------------------------------
----UC-----------------------AU------------CUAUAU-ACA--UUGAACAAC--UAACUU
AGA--ACUGGAGCGGACAAGGGGAAUCCGA-----CUGUUUAAUUAAAACAUAGCAUUGCGAUGGCCAG---
-AA-A------------AUGGUGUUG-ACGCAAUGUGA-UUUCUGCCCAGUGCUCUG-AAUG-UCAAAGUGA
A--------------GAAA--------------UUCAAACAAGCGC--GGGUAAAC-GGCGGGAGUAACUAU
GACUCUCUUAAGGUAGCCAAAUGCCUCGUCAUC-UAAUUAGUGACGCGCAUGAAUGGAUUAACGAGA--UUC
CCACUGUCCCUAUC----UAC-UAUCUAGCGAACCC---AC---AGCCAAGGGAACGGGCUUGGAAUAAUCA
GCGGGGAAAGAAGA-CCCUGUUGAGC-UUGACU--CUAGUUU-GACUUUGUGAAAUGGCUUAGAAGGUGUAG
CAU-AAGUGGGAGC------------------------------------------------------UACG
----------------------------------------------------G--C-GA-UCUUGAAAUA-C
CACUACUUUUAAUGUCAUUUUACUUAUUC-CG-------U-UA-U-AUUGA---------------------
------GC--CU-CA--------UUG-----CU---------------------------------------
------------------------------------------------------------------------
------------------------------------------------------------------------
------------------------------------------------------------------------
--------------------GA------------------------------UCUU----CG----------
------------------------------------------------CGGA-----AGACA-UAGUCAGAU
GGGGAGUUUGGCUGGGGCGGCACAUCUGUUA-AAAGGUAAC-GCAGGUGUCCU-AAGGUAAGCUCAAUGAGA
ACAGAAAUCUCAUGUA-GACCA-AAAGGGGAA-AAGCUUACUUGAUU-UUGAUUUUCAGUACGAAUACAAAC
UGUGAAAGCAUGGCUUUUCA--GUACGAAUACAAACUGUGAAA-GCAUGGCCUAUCGAUCCUUU--------
---AG-A-----UCUGUGAAA--------------------AU-UACAGCUAGAGGUGUCAG-AA--AAGUU
ACCACAGGG-AUAACUGGCUUGUGGCAGCCAAGCGUUCAUAGCGACGUUGCUUUUUGAUCCUUCGAUGUCGG
CUCUUCCUAUCAUUGGGAUGCA-GAAGUCCCAAAGUGUCGGAUUGUUCACCCGCUAAUAGGGAACGUGAGCU
GGGUUUAGACCGUCGUGAGACAGGUUAGUUUUACCCUACU--------------------G-AUGA------
-------------------------------AU--A-UGAUAUUGCGACAGU-AAU-CUAACUUAGUACGAG
AGGAA-CAGUUAGGUCACAUAAU-UGGU-UAAUACAGCUAGUU-GAAA-AGCUAG-U-GCUGUGAAGCUACC
AUGUG-UUGGAUUAUGACUGAACGCC-UC--UAAGUCAGAAUCCAUACUGAAU---AU--GCAAUGUU-AUU
CACCAAUU-UUAAUGUU-------------------------------------------------------
------------------------------------------------------------------------
------------------------------------------------------------------------
---------------------ACAUGUAAACAA-----------AU--------------------------
-------------------------------AGGUUA-----------------------------------
--UUUUUU--AA-------CC--CU------------ACUUC------------------AUACUA------
---------------------------------------AGGUA-----------UCAA----CGCUUAUUA
CG--------------------------U----U-AUACUAAA--------AAAAUA-AUU----------U
C-----------------AAAUAUU---------------------A-AAAUUGAU-----ACA------UC
CUCUGUAUACGACUUUGA---U-GCAUCCAGGU-AUUGUAAG-UACGA--------GAGUAAG-CU------
----UUG--UCUUACGAUCUACUGAGAUUCAU-CCUG-UGUUCUAA---AGUGUGUAU--------------
------------------------------------------------------------------------
------------------------------------------------------------------------
------------------------------------------------------------------------
-------------------------------------------------

```
>LSUe_280463_EMIHU/1-3419 AHAL01003460.1:c35260-38519,38810-38968 Emiliania
---------------------------------------CAACUCCUGCC-GAUGGAUA-U-CUUGGC-UCU
C-G---CAACGAUGAAGAACGCAG--CGAAAUGCGAUACGUAGUGCGAAU--U---GCA--GAA-UUC-AGU
GAAUCAUCGAACCUUUGAACGCAAAU-G--------GC--------GC----AUCC-G-GGC---------U
C-C-----------GCU---C----G----G---ACG----C-A-UGUC-UCU-----UCGAGGGGCG-C--
--------------------------------------------------------CUACCCGCC-AUCG-A
AUG--------------------AG-G-CA-AGACUACCCGCUGAACUUAAGCAUAUCACUAAGCG--GAGG
AGAAGAAACUAA--CCAGGAUGCCCCUA--GU-AA-UGGCGAAUGAAGCGGGC-U-GA-GCUCACGCUGCCA
AUCU-GGAGGCC------UCGCGUAUGCAGCG-C---CCC--CCGAAUUGUGGUC--UA-------------
--GA-GAGG-CGU-------U-------------------------AUCGG--CCG-----------CG---
CC------CGCCG--CUC-------AAGUCC-CUUG------------------------------------
-------GAA-CA-GGGUG-------UCGGAGAGGGUGAGAAUCCCGUA---------CCG--------UGG
CGG---GU--UGCGG-CCUUACG-------------AUGC-GUCCU--CGUCGAGUCGGGUUGCUUG---GG
AUUGCAGCUUGAA-U-UGGGUGGUAAA-UUCCAUCUAAAGCUAAAUACUGG--C-GAG-AGACCGAUAGC-G
AAC-A-AGUACCGUGAGGGAAAG-AUGCAA-AGAAC-UUUGA-AAAGAG-A-GUCAAAGAGUGCUUGAAAUC
G-GUAGGAGG--A-AAC-CC-GGAU---GGGC-CCG-GCU--------------------------------
------------------------------------------------------------------------
------------------------------------------------------------------------
------------------------------------------------------------------------
------------------------------------------------------------------------
----------------------------------UCG--------GCGCG-----------------CCGAC
U-----CAG-------------CCUG--G--GGUUCUCCCCAGGU---------------GCACUUCUGCG-
---CG----------------------------------CCGGGUCAGC-CGCGCCUCGGCGCCGCGCCUCA
ACC-----------------CGGGAG----------------------------------------------
------------------------------------------------------------------------
------------------------------------------------------------------------
------------------------------------------------------------------------
------------------------------------------------------------------------
------------------------------------------------------------------------
------------------------------------------------------------------------
------------------------------------------------------------------------
-------------------------------------------------CGGGGC-----------------
------U------------UUACGGCCA-------GCUCCCGU-----------------------------
----------------------------------------CCU---G-GCGCG------GC-----------
-----CCGGG--------------------------------------------------------------
------------------------------------------------------------------------
------------------------------------------------------------------------
-------------------GCCCGGCUG----------------ACGGAA--CGGGCCCCUCCGACCCGUCU
AGAAACACGGACCAAGGAGUCUG-A-CACGUGUGCGA-GUACCCGG--G-UG-------GCAAG-------C
C-C----GCG--UGCGGAAUGAAAG-UGA-----------------AG-----------------------G
GCA---------------------------------------------------------------------
------------------------------------------------------------------------
------------------------------------------------------------UGCCCUGCC-AA
CCGACC----UU-GACC----------------------UUCU-----------------GUGAGGGGUUU-
GAG-U-----------------------GGAGCACACCUGUCGG-GACCCGAAAGA-UGGUGAACUAUGCCU
```

```
GGGUAGGGUGAAGCCAGA-GGAAAC-UCUGGUGGAGGCUCGU--AGCGAUACUGACGUGCAAAUCGU-UCGU
CGAAUCUGGGUAUAGGGGCGAAAGACUAAUCGAACCAUC-UAGUAGCUGGUUCCCUCCGAAG-UUUCCCUCA
GGAUAGCUUGAA-CG---------------------------------------------------------
---------------------CAA-----CGGUUUUGUCAGGUAAAGCGAAUGAUU-AGA-GGCCUGUG-GG
C-U-GAAAC-AGCU-U-GCACCUAUUCUCAAACUUUAAAUGGGCAAGAG-----------------------
------------------GCCCG-GG-----AUG-CU--UGAG--U--GAU---------------------
CC---CGGG-C---GGCGA-AC-G--GC----G--CGUUCAA-AGUGGGC-C-AUUU-U-UGG-UAAGCAGA
AC-U-GGCGAUGC-GGGAUG---AACCGAACGCCG-C-GCUAAGGUACCCAAGU--GU-CCG-CU-CA---U
-C-AGAUACC-ACAAAAGGCGUUGGUACCUCCAGACAGCAGGACGGUGGCCAUGGAAGUCGGAA-CCCGCUA
AGGAGUGUGUAACAACUCACCUGCC-----GAAGGUACUAGCCCUGAAAAU-GGAU-GGCGCUCA-AGCGGA
C----------CACCGAU-GCGCGGCCGCCGGCG--CGAG--------------------------------
------------------------------------------------------------------------
------------------------------------------------------------------------
---------------------------CGCGAGGCGC-CGGC-GAGUAGGAGGGC-GCGGGGC---------
-----------U-C-GCGGC-GCAGCCUU-G-GGCGCGAGCC--U-GGGUGAAGCG-G-GCC-CCGGUGCAG
AUCU-UGGUGG-UAGUA-GCAAAU---AUUC-AAAUGA--G---------------AACUUU-GAA-GACUG
AAGUGGAG-AAAGGUUCC-GUGUGAACA-GCAGUUGGACACGGGUGAGCCGA--CC-CUAAGAGACAGGAA-
A-A-CUCC-----GCCGAAC--------G-------CGG-----CUG---GUUCU-----------------
----------------GUCAGCCC-------GUC----CG-UC-GA-A-AGGGA-AUCGGGUU-AAUA-UUC
CCGAG----------------------------------CCAGCG---CGUG--------GAUG--------
G----------------UG-----------------------------------------------------
----GG-CGG---CGAC--------GC-GA-----AC--------------------GAAC-C-CGGA-GAC
GCCGGCG-------------GGAGCC-CUGGGAAGA-GU------UCUCU-------------U-UUCUUC-
UUAACU-GCC--U-------C------------------CUG-----CC----------------CUGG-AA
U-C-GG--------------------UUUAG-CCGGAGCGAG---------GGCUAC----ACG-GC-AGG-
CAA--AG-----------------------------------------------------------------
----CACCCUA-AGUCUUGGGGUGU-C-CGGCGCG-CUCCCGACGGC-CCGUGAAA-A-UCC-GGGGGAGCG
-G---------------------------------------------UU-----CGUUCUCA-CG-CCUGGU
CGUACUCAUAACCGCA------------------------------UCAGGUC--UCCAAGGU-GAACAGCC
UCUAGU-C-GA-UAGA-GCAAUGUA-G-GUAAGGGAAGUCGGC-AAAAUGGUUCCGCAACCUCGG-GAGAAG
GAAUG-GCUC-U-GA--AGGCUG-G----GCGCA-GG------GGUCCU--AGG-CUCGAG-GC--CGG---
CUGG------C------UGCG---CG----------------------------------------GGG--A
C---------------------------------------------------------------------UA
-C----------------------------------------------------------------------
-----------------------------------------------------------------------G
GCCU--------------C---------------------------------ACGGCU--------------
---------------------------------------------------------------GCGGA----
------------------------------------------------------------------------
-U-CC---------------CG--C-GCG--GCCG-UC----------------------------------
---CGG----CG-GA-CGGC----CU-AGGAGC---------------------------------------
----------------------------------------------------------------GCUC----
-----------------------------------------------------------C------------
-------------------------------------------------G---G------------------
------------------------------------------------------------------------
----CU-----------------------CU------------CCCUGC-GCG--ACGAACAGC--CGCUUC
```

```
AGA--ACUGGCGCGGACAAGGGGAAUCCGA-----CUGUUUAAUUAAAACAAAGCAUUGCGAUGGCCCG---
-CG-A-------------GGGUGCUG-ACGCAAUGUGA-UUUCUGCCCAGUGCUCUG-AAUG-UCAAAGUGA
A--------------GAAA--------------UUCAACCAAGCGC--GGGUAAAC-GGCGGGAGUAACUAU
GACUCUCUUAAGGUAGCCAAAUGCCUCGUCAUC-CAAUUAGUGACGCGCAUGAAUGGAUUCGUGAGA--UUC
CCACUGUCCCUAUC----UAC-UGUCUAGCGAAACC---AC---AGCCAAGGGAACGGGCUUGGAAGAAUCA
GCGGGGAAAGAAGA-CCCUGUUGAGC-UUGACU--CUAGUCU-GUUACUGUGGAACGACUUGAUGGGUGUAG
AAU-AUGCGGAGGG------------------------------------------------------GCAA
--------------------------------------------------C-C--C-GA-CAGUGAAAUA-C
CGCAACCGUCAACGUUGUUUUACUUACUC-GG-------U-GG-C-GCGGA----------------------
------GG--GA-GG--------CUC---GCCU-C--A--------------CUUCU---------------
-----------GGAU--------------------------------------CCAA--GC---------CG
---------------------------------------------------------------C--------
-----U---------------------------U---CGU-----GC-------------------------
----------GCG------CGA------------------------------UCCG----CG----------
------------------------------------------------UCGG-----GGACA-CUAACAGGU
GGGGAGUUUGGCUGGGGCGGCACGUCUGUUA-AAAAAUAAC-GCAGGCGCCCG-AUUUUGAGCUCAAUGAGA
ACAGAAAUCUCAUGUG-GACCG-AAAGGGGAA-AAGCUCGGUUGAUU-UUGA--------------------
--------------UUUUCA--GUACGAAUACAAACCGCGAAA-GCGUGGCCUAUCGAUCCUCU--------
---CG-U-----CUUUGGAA---------------------UU-UCAAGCUAGAGGUGUCAG-AA--AAGUU
ACCACAGGG-AUAACUGGCUUGUGGCUGCCAAGCGUUCAUAGCGACGUAGCUUUUUGAUCCUUCGAUGUCGG
CUCUUCCUAUCACUGCCUCGCA-GCGGGGGCCAAAUGCAGGAUUGUUCACCCACCAACAGGGAACGUGAGCU
GGGUUUAGACCGUCGUGAGACAGGUUAGUUUUACCCUACU--------------------G-CUG-------
--------------------------------U----CUGUUUUUCUACGGU-AAU-UUGGUUCAGUACGAG
AGGAA-CCACCAAUUCGCAUCAU-CGGU-AUAGCCGC-CGCCG-GAAA-CGGCGA-U-GCGGCGAAGCUGCC
AUGCG-CCGGAUUAUAACUGAACGCC-UC--UAAGUUAGAAUCCAUGCCGG-----AA--GGAAAGC----C
ACCUACCG-GCCCG----------------------------------------------------------
------------------------------------------------------------------------
------------------------------------------------------------------------
------------------------------------------------------------------------
--------------------------------GGG----------A--------------------------
----------CG-------CC--ACG---------AGCAACG------------------GUAGCAC---U-
---------------------------------------GCCUGAG---------CCGCG-AGCGAC-----
-------------------------------------CA-CAA--------UUCACG-GAC----------C
C-----------------GGGCCGA---------------------C--------------GAA------UC
CCUUGCAGACGACUUGAA-UAA-GGAACGGGGU-AUUGUACG-CGGCA--------GAGUGGC-CU------
----UGU--UGCCACGAUCCGCUGAGAUUCAG-CCUU-GGUUCUGAC--GAUUUGU----------------
------------------------------------------------------------------------
------------------------------------------------------------------------
------------------------------------------------------------------------
------------------------------------------------------------
>LSUe_3702_ARATH/1-3538
-----------------------------AA----AA--CGACUCUCGGC-AACGGAUA-U-CUCGGC-UCU
C-G---CAUCGAUGAAGAACGUAG--CGAAAUGCGAUACUUGGUGUGAAU--U---GCA--GAA-UCC-CGU
GAACCAUCGAGUCUUUGAACGCAAGU-U--------GC--------GC----CCCA-A-GCC--------UU
C-U-----------GGC---C----G----A---GGG----C-A-CGUC-UGC-----CUGGGUGUCA-C--
```

```
--------------------------------------------------------A--AGCGAC-CCCA-G
GUC--------------------AG-G-CG-GGAUUACCCGCUGAGUUUAAGCAUAUCAAUAAGCG--GAGG
AAAAGAAACUAA--CAAGGAUUCCCUUA--GU-AA-CGGCGAGCGAACCGGGA-A-GA-GCCCAGCUUGAAA
AUCG-GACGUCU------UCG------------G---CGU--UCGAAUUGUAGUC--UG-------------
--GA-GAAG-CGU-------C-------------------------CUCAG--CGA-----------CG---
GA------CCGGG--CCU-------AAGUUC-CCUG------------------------------------
-------GAA-AG-GGGCG-------CCAGAGAGGGUGAGAGCCC-GUC---------GUG---C----CCG
GAC---CC--UGUCG-CACCACG-------------AGGC-GCUGU--CUACGAGUCGGGUUGUUUG---GG
AAUGCAGCCCCAA-U-CGGGCGGUAAA-UUCCGUCCAAGGCUAAAUACGGG--C-GAG-AGACCGAUAGC-G
AAC-A-AGUACCGCGAGGUAAAG-AUGAAA-AGGAC-UUUGA-AAAGAG-A-GUCAAAGAGUGCUUGAAAUU
G-UCGGGAGG--G-AAG-CG-GAUG---GGGG-CCG-GCGAU--------------------GC--------
-------------------------------------------GUCCUGG----------UCG--GAUG---
-------------------------CGG-----------AA-------CGGA--------------------
---------------------------------------------------------------GCAAU----
------------------------------------------------------------------------
----------------------------------CCG--------GUCCG-----------------CCGAU
C-----GAU-------------UCGG--G--GC----------GU---------------GGACCGACGCG-
---GA----------------------------------UUACGGUGGC-----------------GGCCUA
AGC-----------------CCGGGC----------------------------------------------
-----------------------------------------------------------UUUU--G-AUAC-
------------------------------------------------------------------------
------------------------------------------------------GCUUGUGGAGACGUCGCU
GCCGUGAUC---------------------------------------------------------------
------------------------------------------------------------------------
------------------------------------------------------------------------
------------------------------------------------------------------------
-------------------------------------------------GUGGUC-----------------
--------------------UGCAGCAC-------GCGCCU-------------------------------
----------------------------------------AAC---G-GCGUG------C------------
-----CUCGG----CAU------------------CAGC--------GUGCUCCG-----------------
------------------------------------------------------------------------
------------------------------------------------------------------------
-----------------GGCGUCGGCCU----------------GUGGGC--U---CCCAUUCGACCCGUCU
UGAAACACGGACCAAGGAGUCUG-A-CAUGUGUGCGA-GUCAACGG--G-UG-------AGUAA-------A
C-CC--GUAA--GGCGCAAGGAAGC-UGA-----------------UU-----------------------G
GCGG-GAUCCU---------------------------------------------------CG--------
C-----------------------------------------------------------------------
-------------------GGG--------------------------------------UGCACCGCC-GA
CCGACC----UU-GAUC----------------------UUCU-----------------GAGAAGGGUUC-
GAG-UG----------------------UGAGCAUGCCUGUCGG-GACCCGAAAGA-UGGUGAACUAUGCCU
GAGCGGGGUAAAGCCAGA-GGAAAC-UCUGGUGGAAGCCCGC--AGCGAUACUGACGUGCAAAUCGU-UCGU
CUGACUUGGGUAUAGGGGCGAAAGACUAAUCGAACCAUC-UAGUAGCUGGUUCCCUCCGAAG-UUUCCUCA
GGAUAGCUGGAG-CUCG-------------------------------------------------------
-------------------GACGC-----GAGUUCUAUCGGGUAAAGCCAAUGAUU-AGA-GGCAUUGG-GG
G-C-GCAAC-G-CC-U-CGACCUAUUCUCAAACUUUAAAUAGGUAGGAC-----------------------
```

```
------------------GUGUC-GG-----CUG-CU--UUGU--U--GAG---------------------
CC---GUCA-C---ACGGA-AU-C--GA----G--AGCUCCA-AGUGGGC-C-AUUU-U-UGG-UAAGCAGA
AC-U-GGCGAUGC-GGGAUG---AACCGGAAGCCG-G-GUUACGGUGCCCAACU--GC-GCG-CU-AA---C
-CUAGAACCC-ACAAAGGGUGUUGGUCGAUUAAGACAGCAGGACGGUGGUCAUGGAAGUCGAAA-UCCGCUA
AGGAGUGUGUAACAACUCACCUGCC-----GAAUCAACUAGCCCCGAAAAU-GGAU-GGCGCUUA-AGCGC-
-----------GACCUAU-ACCCGGCCGUCGGGG--CAAG--------------------------------
------------------------------------------------------------------------
------------------------------------------------------------------------
---------------------------AGCCAGGCCU-CGAU-GAGUAGGAGGGC-GCGGCG----------
-----------U-C-GCUGC-AAAACCUA-G-GGCGCGAG------GCGCGGAGCG-G-CCG-UCGGUGCAG
AUCU-UGGUGG-UAGUA-GCAAAU---AUUC-AAAUGA--G---------------AACUUU-GAA-GGCCG
AAGAGGGG-AAAGGUUCC-AUGUGAACG-GCACUUGCACAUGGGUUAGUCGA--UC-CUAAGAGUCGGGGG-
A-A-ACCC-----GUCUGAU--------A-------GCG-----CU--------------------------
-----------------UAAGCGA---------A----CU-UC-GA-A-AGGGG-AUCCGGUU-AAAA-UUC
CGGAA----------------------------------CCGGGA---CGUG--------G-CG--------
G----------------UU-----------------------------------------------------
----GA-CGG---CAAC--------GU-UA-----GG--------------------GAGU-C-CGGA-GAC
GUCGGCG-------------GGGGCC-UCGGGAAGA-GU------UAUCU-------------U-UUCUGU-
UUACA-GCC--U-----G-C------------------CCA-----CC----------------CUGG-AA
A-C-GG--------------------CUCAG-CCGGAGGUAG---------GGUCCA----GCG-GC-UGG-
AAG--AG-----------------------------------------------------------------
----CACCGCA-CGUCGCGUGGUGU-C-CGGUGCG-CCCCCGGGCGC-CCUUGAAA-A-UCC-GGAGGACCG
-A---------------------------------------------GU-----GCCGCUCA-CG-CCCGGU
CGUACUCAUAACCGCA------------------------------UCAGGUC--UCCAAGGU-GAACAGCC
UCUGGU-C-GA-UGGA-ACAAUGUA-G-GCAAGGGAAGUCGGC-AAAAUGGAUCCGUAACUUCGG-GAAAAG
GAUUG-GCUC-U-GA--GGGCUG-G----GCUCG-GG------GGUCCC--AGU-UCCGAA-CC--CGU---
-CGG------C------UGUC---AG----------------------------------------CGG--A
C---------------------------------------------------------------------UG
-CUC-------G--------AGCU------------------------------------------------
---------------------------------------------------------GC-----------UU
CCGC--------------G---------------------------------GCGAGA--------------
---------------------------------------------------------------GCGGG----
------------------------------------------------------------------------
-U-CG---------------CC--G-GCU--GCCG-GC----------------------------------
---CGG----GG-GA-CGA-----CU-GGGAAC---------------------------------------
----------------------------------------------------------------GGCU----
-----------------------------------------------------------CUCU---------
---------------C------------------------------GGGA---G------------------
------------------------------------------------------------------------
----CU-----------------------UU------------CCCCGG-GCG--UCGAACAGU--CAGCUC
AGA--ACUGGUACGGACAAGGGGAAUCCGA-----CUGUUUAAUUAAAACAAAGCAUUGCGAUGGUCCC---
-UG-C-------------GGAUGCUA-ACGCAAUGUGA-UUUCUGCCCAGUGCUCUG-AAUG-UCAAAGUGA
A--------------GAAA--------------UUCAACCAAGCGC--GGGUAAAC-GGCGGGAGUAACUAU
GACUCUCUUAAGGUAGCCAAAUGCCUCGUCAUC-UAAUUAGUGACGCGCAUGAAUGGAUUAACGAGA--UUC
CCACUGUCCCUGUC----UAC-UAUCCAGCGAAACC---AC---AGCCAAGGAACGGGCUUGGCAGAAUCA
```

GCGGGGAAAGAAGA-CCCUGUUGAGC-UUGACU--CUAGUCC-GACUUUGUGAAAUGACUUGAGAGGUGUAG
GAU-AAGUGGGAGC------------------------------------------------------UUCG
----------------------------------------------------G--C-GC-AAGUGAAAUA-C
CACUACUUUUAACGUUAUUUACUUACUC-CG--------U-GA-A-UCGGA------G-----------GCC
------GG--GG-UA--------CAA---CCCC-U--G--------------UUUUU---------------
-----------GGUC--------------------------------------CCAA--GG---------CU
---------------------------------------------------------------CG-------
---CUU-------------------------------CGG-----CG-------------------------
----------GGU------CGA------------------------------UCCG----GG----------
------------------------------------------------CGGA-----GGACA-UUGUCAGGU
GGGGAGUUUGGCUGGGGCGGCACAUCUGUUA-AAAGAUAAC-GCAGGUGUCCU-AAGAUGAGCUCAACGAGA
ACAGAAAUCUCGUGUG-GAACA-AAAGGGUAA-AAGCUCGUUUGAUU-CUGA--------------------
--------------UUUUCA--GUACGAAUACGAACCGUGAAA-GCGUGGCCUAUCGAUCCUUU--------
---AG-A------CUUCGGAA--------------------UU-UGAAGCUAGAGGUGUCAG-AA--AAGUU
ACCACAGGG-AUAACUGGCUUGUGGCAGCCAAGCGUUCAUAGCGACGUUGCUUUUUGAUCCUUCGAUGUCGG
CUCUUCCUAUCAUUGUGAAGCA-GAAUUCACCAAGUGUUGGAUUGUUCACCCACCAAUAGGGAACGUGAGCU
GGGUUUAGACCGUCGUGAGACAGGUUAGUUUUACCCUACU--------------------G-AUG-------
--------------------------------C----CCGCGUCGCGAUAGU-AAU-UCAACCUAGUACGAG
AGGAA-CCGUUGAUUCGCACAAU-UGGU-CAUCGCGCUUGGUU-GAAA-AGCCAG-U-GGCGCGAAGCUACC
GUGCG-CUGGAUUAUGACUGAACGCC-UC--UAAGUCAGAAUCCGGGCUAG-----AA--GCGACGCA-UGC
GCCCGCCG-CCCGAUU--------------------------------------------------------
------------------------------------------------------------------------
------------------------------------------------------------------------
-----------------------GCCGACCCUC----------AGUA-------------------------
--------------------------------GGAGC-----UU-A--------------------------
----------GG-------CU--CCA---------AAGGCAC------------------GUGUCGU---U-
---GGCUAAGUC------CGU-----U------------CGGCG-----------GAAC--GG-UCGU-UCG
GA--------------------------C----C-GCCU-UGA--------AUUAUA-AUU----------A
C-----------------CACCGAG---------------------C-GGCGGGUA-----GAA------UC
CUUUGCAGACGACUUAAA-UAC-GCGACGGGGU-AUUGUAAG-UGGCA--------GAGUGGC-CU------
----UGC--UGCCACGAUCCACUGAGAUUCAG-CCCU-UUGUCGCUA--AGAUUCG-A--------------
------------------------------------------------------------------------
------------------------------------------------------------------------
------------------------------------------------------------------------
-------------------------------------------------
>LSUe_39947_ORYSJ/1-3567
-----------------------------CA----CA--CGACUCUCGGC-AACGGAUA-U-CUCGGC-UCU
C-G---CAUCGAUGAAGAACGUAG--CGAAAUGCGAUACCUGGUGUGAAU--U---GCA--GAA-UCC-CGU
GAACCAUCGAGUCUUUGAACGCAAGU-U--------GC--------GC----CCGA-G-GCC--------AU
C-C-----------GGC---C----G----A---GGG----C-A-CGCC-UGC-----CUGGGCGUCA-C--
------------------------------------------------------------GCGAC-CCCA-G
GUC--------------------AG-G-CG-GGACUACCCGCUGAGUUUAAGCAUAUAAAUAAGCG--GAGG
AGAAGAAACUUA--CGAGGAUUCCCCUA--GU-AA-CGGCGAGCGAACCGGGA-G-AU-GCCCAGCUUGAGA
AUCG-GGCGGCC------GCG------------C---CGU--CCGAAUUGUAGUC--UG-------------
--GA-GAGG-CGU-------C-------------------------CUCAG--CGA-----------CG---

```
GA------CCGGG--CCC-------AAGUCC-CCUG------------------------------------
-------GAA-AG-GGGCG-------CCUGGGAGGGUGAGAGCCCCGUC---------CGG---C----CCG
GAC---CC--UGUCG-CCCCACG-------------AGGC-GCCGU--CAACGAGUCGGGUUGUUUG---GG
AAUGCAGCCCAAA-U-CGGGCGGUAAA-CUCCGUCCAAGGCUAAAUACAGG--C-GAG-AGACCGAUAGC-G
AAC-A-AGUACCGCGAGGGAAAG-AUGAAA-AGGAC-UUUGA-AAAGAG-A-GUCAAAGAGUGCUUGAAAUU
G-CCGGGAGG--G-AAG-CG-GAUG---GGGG-CCG-GCGAU--------------------GC--------
-------------------------------------------GCCCCGG----------CCG--UAUG---
-------------------------CGG-----------AA-------CGG---------------------
---------------------------------------------------------------CUUCG----
-----------------------G------------------------------------------------
----------------------------------CUG--------GUCCG-----------------CCGAU
C-----GGC-------------UCGG--G--GC----------GU---------------GGACUGUUGUC-
---GG----------------------------------CCGCGCCGGC-----------------GGCCAA
AGC-----------------CCGGGGGC--------------------------------------------
-------------------------------------------------------------UC--C-GC---
------------------------------------------------------------------------
------------------------------------------------------GCCCCCGGC---------
------------------------------------------------------------------------
------------------------------------------------------------------------
------------------------------------------------------------------------
------------------------------------------------------------------------
-------------------------------------------------AGCCG------------------
------U------------CGUCGGCGC-------AGCCGGUC-----------------------------
----------------------------------------ACC-----GCGCG------CC-----------
-----UCUGGCG--CGC-CCCUCGGG-------G-CGCU--------GCGCCGCA----A------------
------------------------------------------------------------------------
------------------------------------------------------------------------
----------------------CGGCCU----------------GCGGGC--U---CCCAUCCGACCCGUCU
UGAAACACGGACCAAGGAGUCUG-A-CAUGCGUGCGA-GUCGACGG--G-UU-------CUGAA-------A
C-CU--GGGA--UGCGCAAGGAAGC-UGA-----------------CG-----------------------A
GCGG-GAGGCCC--------------------------------------------------UC--------
ACG---------------------------------------------------------------------
-------------------GGC--------------------------------------CGCACCGCU-GG
CCGACC----CU-GAUC----------------------UUCU-----------------GUGAAGGGUUC-
GAG-UU----------------------GGAGCACGCCUGUCGG-GACCCGAAAGA-UGGUGAACUAUGCCU
GAGCGGGGCGAAGCCAGA-GGAAAC-UCUGGUGGAGGCUCGA--AGCGAUACUGACGUGCAAAUCGU-UCGU
CUGACUUGGGUAUAGGGGCGAAAGACUAAUCGAACCAUC-UAGUAGCUGGUUCCCUCCGAAG-UUUCCCUCA
GGAUAGCUGGAG-CCCA-------------------------------------------------------
--------------------UUAC-----GAGUUCUAUCGGGUAAAGCCAAUGAUU-AGA-GGCAUCGG-GG
G-C-GCAAC-GCCC-U-CGACCUAUUCUCAAACUUUAAAUAGGUAGGAC-----------------------
------------------GGCGC-GG-----CUG-CU--CCGG--U--GAG---------------------
CC---GCGC-C---ACGGA-AU-C--GG----G--AGCUCCA-AGUGGGC-C-AUUU-U-UGG-UAAGCAGA
AC-U-GGCGAUGC-GGGAUG---AACCGGAAGCCU-G-GUUACGGUGCCGAACU--GC-GCG-CU-AA---C
-CUAGAACCC-ACAAAGGGUGUUGGUCGAUUAAGACAGCAGGACGGUGGUCAUGGAAGUCGAAA-UCCGCUA
AGGAGUGUGUAACAACUCACCUGCC-----GAAUCAACUAGCCCCGAAAAU-GGAU-GGCGCUGA-AGCGCG
```

```
C----------GACCCAC-ACCAGGCCAUCUGGG--CGAG--------------------------------
------------------------------------------------------------------------
------------------------------------------------------------------------
---------------------------CGCCAUGCCC-CGAU-GAGUAGGAGGGC-GCGGCGG---------
-----------C-C-GCCGC-AAAACCCG-G-GGCGCGAGCC--C-GGGCGGAGCG-G-CCG-UCGGUGCAG
AUCU-UGGUGG-UAGUA-GCAAAU---AUUC-AAAUGA--G---------------AACUUU-GAA-GGCCG
AAGAGGAG-AAAGGUUCC-AUGUGAACG-GCACUUGCACAUGGGUAAGCCGA--UC-CUAAGGGACGGGGU-
A-A-CCCC-----GGCAGAG--------A-------GCG-----CGA-------------------------
----------------CCACGCGC-------GUG----CC-CC-GA-A-AGGGA-AUCGGGUU-AAGAUUUC
CCGAG----------------------------------CCGGGA---CGUG--------G-CG--------
G----------------UU-----------------------------------------------------
----GA-CGG---CGAC--------GU-UA-----GG--------------------AAGU-C-CGGA-GAC
GCCGGCG-------------GGGGCC-UCGGGAAGA-GU------UAUCU-------------U-UUCUGC-
UUAACG-GCC--C-----G-C------------------CAA-----CC----------------CUGG-AA
A-C-GG--------------------UUCAG-CCGGAGGUAG---------GGUCCA----GCG-GC-CGG-
AAG--AG-----------------------------------------------------------------
----CACCGCA-CGUCGCGCGGUGU-C-CGGUGCG-CCCCCGGCGGC-CCUUGAAA-A-UCC-GGAGGACCG
-A---------------------------------------------GU-----ACCGUCCA-CG-CCCGGU
CGUACUCAUAACCGCA------------------------------UCAGGUC--UCCAAGGU-GAACAGCC
UCUGGC-C-AA-UGGA-ACAAUGUA-G-GCAAGGGAAGUCGGC-AAAACGGAUCCGUAACUUCGG-GAAAAG
GAUUG-GCUC-U-GA--GGGCUG-G----GCUCG-GG------GGUCCC--GGC-CCCGAA-CC--CGU---
-CGG------C------UGCC---GG----------------------------------------CGG--A
C---------------------------------------------------------------------UG
-CUC-------G--------AGCU------------------------------------------------
---------------------------------------------------------GC-----------UC
GCGC--------------G---------------------------------GCGAGA--------------
---------------------------------------------------------------GCGGG----
------------------------------------------------------------------------
-C-CG---------------CC--G-CGU--GCCG-GC----------------------------------
---CGG----GG-GA-CGGA----CC-GGGAAC---------------------------------------
----------------------------------------------------------------GGCC----
-----------------------------------------------------------CCCU---------
---------------C------------------------------GGGG---G------------------
------------------------------------------------------------------------
----CC-----------------------UU------------CCCCGG-GCG--UCGAACAGC--CGACUC
AGA--ACUGGUACGGACAAGGGGAAUCCGA-----CUGUUUAAUUAAAACAAAGCAUUGCGAUGGUCCU---
-CG-CGGAUGCUGACGCAAGUGAUUC-ACGCAAUGUGA-UUUCUGCCCAGUGCUCUG-AAUG-UCAAAGUGA
A--------------GAAA--------------UUCAACCAAGCGC--GGGUAAAC-GGCGGGAGUAACUAU
GACUCUCUUAAGGUAGCCAAAUGCCUCGUCAUC-UAAUUAGUGACGCGCAUGAAUGGAUUAACGAGA--UUC
CCACUGUCCCUGUC----UAC-UAUCCAGCGAAACC---AC---AGCCAAGGGAACGGGCUUGGCGGAAUCA
GCGGGGAAAGAAGA-CCCUGUUGAGC-UUGACU--CUAGUCC-GACUUUGUGAAAUGACUUGAGAGGUGUAG
GAU-AAGUGGGAGCC-----------------------------------------------------CUCG
--------------------------------------------------G-G--C-GC-AAGUGAAAUA-C
CACUACUUUUAACGUUAUUUUACUUAUUC-CG-------U-GA-G-UCGGA------A-----------GC-
------GG--GG-CC--------UGG---CCCC-U--C--------------CUUUU---------------
```

```
-----------GGCU--------------------------------------CUAA--GG---------CC
---------------------------------------------------------------CG-------
---AGU---------------------------C---CCU-----CG-------------------------
----------GGC------CGA------------------------------UCCG----GG----------
------------------------------------------------CGGA-----AGACA-UUGUCAGGU
GGGGAGUUUGGCUGGGGCGGCACAUCUGUUA-AAAGAUAAC-GCAGGUGUCCU-AAGAUGAGCUCAACGAGA
ACAGAAAUCUCGUGUG-GAACA-AAAGGGUAA-AAGCUCGUUUGAUU-CUGA--------------------
--------------UUUCCA--GUACGAAUACGAACCGUGAAA-GCGUGGCCUAUCGAUCCUUU--------
---AG-A-----CCUUCGGAG--------------------UU-UGAAGCUAGAGGUGUCAG-AA--AAGUU
ACCACAGGG-AUAACUGGCUUGUGGCAGCCAAGCGUUCAUAGCGACGUUGCUUUUUGAUCCUUCGAUGUCGG
CUCUUCCUAUCAUUGUGAAGCA-GAAUUCACCAAGUGUUGGAUUGUUCACCCACCAAUAGGGAACGUGAGCU
GGGUUUAGACCGUCGUGAGACAGGUUAGUUUUACCCUACU--------------------G-AUG-------
--------------------------------A----CCGUGCCGCGAUAGU-AAU-UCAACCUAGUACGAG
AGGAA-CCGUUGAUUCACACAAU-UGGU-CAUCGCGCUUGGUU-GAAA-AGCCAG-U-GGCGCGAAGCUACC
GUGUG-CCGGAUUAUGACUGAACGCC-UC--UAAGUCAGAAUCCAAGCUAGC----AA--GCGGCGCC-UGC
GCCCGCCG-CCCGC----------------------------------------------------------
------------------------------------------------------------------------
------------------------------------------------------------------------
-----------------------CCCGACCCAC----------GUUAGGGG-----CU--------------
-------------------------------AGGGGC-----GCAA--------------------------
----------GC-------CC--CCA---------AGGGCCC------------------GUGCCAC---C-
---GGCCAAGCC------GGC-----C------------CGGCC-----------GACG--CG-CCGCGGCC
GG--------------------------C----C-GCCU-CGA--------AGCUCC-CUU----------C
C-----------------CAACGGG---------------------C-GGCGGGCU-----GAA------UC
CUUUGCAGACGACUUAAA-UAC-GCGACGGGGC-AUUGUAAG-UGGCA--------GAGUGGC-CU------
----UGC--UGCCACGAUCCACUGAGAUCCAG-CCCC-GCGUCGCAC-GGAUUCGUCC--------------
------------------------------------------------------------------------
------------------------------------------------------------------------
------------------------------------------------------------------------
-------------------------------------------------
>LSUe_7160_AEDAL/1-4267
-----------------------------------------ACCCUAGGC-AGGGGAUC-A-CUCGGC-UCG
U-G---GAUCGAUGAAGACCGCAG--CUAAAUGCGCGUCAGAAUGUGAAC--U---GCA--GGA-C---ACA
UGAACACCGACACGUUGAACGCAUAU-U--------GC--------AC----AUCG-U-ACC---------U
C-C-----------AGU---A----C----G---AUG----U-A-CACA-UUU-----UUGAGUGCCU-AUA
UUUU-------------------------------------AUC------------A--GUAGGC-CUCA-A
AUA--------------------AU-G-UG-UGACUACCCCCUAAAUUUAAGCAUAUUAAUAAGGG--GAGG
AAAAGAAACUAA--CAAGGAUUCCCUGA--GU-AG-CUGCGAGCGAAACGGGA-A-GA-GCUCAGCACGCAG
GGGC-GAUGCACUG----GUG------------U---AUC-UGCCCGGUUCCGUG--UA-------------
--CU-GGAG-AGU-U--CGUU-------------------------AUCUG--CGU-----------AA---
CCGGUC--GCUGG--UUC-------AAGUUCAACUA------------------------------------
-------GAA-UG-UGGCUUUUACUCCCAGAGAGGGUGAUAGGCCCGUA---------GAG---C----GGC
GCA---CC--GAUGG-UAGA----------------AGAU-CUUUC--CAUGGAGUCGUGUUGCUUG---AU
AGUGCAGCACAAA-G-UGGGAGGUAAA-CUCCUUCUAAAGCUAAAUAUCAC--C-ACG-AGACCGAUAGC-G
AAC-A-AGUACCGUGAGG-AAAG-UUGAAA-AGCAC-UCUGA-AUAGAG-A-GUCAAAAAGUACGUGAAACU
```

```
G-CCUAGGGC--UCAAG-CCCGUUG---AACU-CGA-UUAUCC-------------------GA--------
-------------------------------------------GCG-GAGA--CAUUCA--CC--UGCG---
------GU-----------------UGG--------------------CCG---------------------
----------------------------------------------------------------GCGA----
------------------------------------------------------------------------
----------------------------------CGG---------CACG-----------------GCCGC
A-----GGGCA---CUUGU---CUCU--C--GC----------ACU----AGCCAAGAGGACACUGCGAUC-
---CA-----------------------------------UUACGAAAC-----------------AGCUUU
CGC-----------------GCAAG-------CGCAAGGUCGCCCGACAACUGCCCCC--------------
------------------------------------------------------------UGG--U-GCU--
----------------------------------------------GGUUGCUUGCCCCACAGUAGCGACGC
UCAGUUCUGAAGGCCUGUGCCGCGAGGUGGGGCUUACUG------CACGUG-GUGUUCUAGCAGUCGGGCGC
GUGAUGGAU------UGAUGGAU-------------------------------------------------
------------------------------------------------------------------------
------------------------------------------------------------------------
------------------------------------------------------------------------
-------------------------------------------------UCCCCC-----GGA---------
------C------------ACCGGGUGG--------------------------------------------
------------------------------------------UC----UUCCCG----------------UA
A-------------GGG-------------------GCC--------AC-CGGAC-----------------
----------------------UG--------------------------------------------UCG-
------------------------------------------------------------------------
--------------------AUCGGCAG--UGA------AAGAAUCGAGG--U---ACUUCGGGACCCGUCU
UGAAACACGGACCAAGAAGUCUA-U-CUUGCGCGCGA-GCCAAUGG--U-CGUC-----CUCCG-----GAG
C-AC--CAAA--GGCGCAGAAAACA-UAA-----------------CUU-----------GAGUUGUGC-GG
GAUU-ACGGGCG----CGGCUCU---------------------------------------CU--------
GCU---------------------------------------------------------------------
-------------------CGU-----------------------------------CCCUCCAUCCCC-GG
GUGUUGUAAUCG-GCAU-GCGGCUACCGGGACCAAGGCUUCAC---GGCCCCCGGCACCGUGCCACAA-CA-
UAC-CG----------------------UGAGUGUGCAGGAUGU-GACCCGAAAGA-UGGUGAACUAUGCCU
GAUCAGGUUGAAGUCAGG-GGAAAC-CCUGAUGGAGGACCGA--AGCAGUUCUGACGUGCAAAUCGA-UUGU
CAGAAUUGGGCAUAGGGGCGAAAGACCAAUCGAACCAUC-UAGUAGCUGGUUCCCUCCGAAG-UUUCCCUCA
GGAUAGCUGGAG-CACG--------------------------------------------------------
---------CAACGUUUCGAAUCC-----UAUUCUUAUCUGGUAAAGCGAAUGAUU-AGA-GGCCUUAG-GU
U-C-GAAAU-GAUC-U-UAACCUAUUCUCAAACUAUAAAUGGGUACGUACCAUAGCAUUCUUGCAUGAUGCU
GUUGCAAACGUUGGUAAACGCCG-GG-----CCGUCC--CCUG--UCAGGGCG------------------U
GCC-ACGGCGU---AGAAG-AU-A--UCU---G--UGUGCUU-AGUGGGC-CAAGUU-U-UGG-UAAGCAGA
AC-U-GGUGCUGU-GGGAUG---AACCAAACGUAA-U-GUUACGGCGCCUAAAUAAACGACG-CA-UC---A
-U-AGAUACC-AUGAAAGGUGUUGAUUGCUACAGACAGCAGGACGGUGGACAUGGAAGUUGUCA-UCCGCUA
AGGAGUGUGUAACAACUCACCUGCC-----GAAGCAAUUAGCCCUUAAAAU-GGAU-GGCGCUUA-AGUCGU
U----------UGCCUAU-ACAUUACCGUGGCGC--ACAAG-------------------------------
----------------------------------------------UGG------------------GGC-A
----CGCCUCGA------GCGCGU------------------------------------------------
------------------------CUGCCCUUGAGGCG-CCAC-GAGUAGGAGGGU-CUGGUG---------
-----------U-GUGCGUU-GAAGUGCC-U-GGCGUAAGCC--G-ACAUGGAGCC-G-CCA-CUAGCACAG
```

```
AUCU-UGGUGG-UAGUA-GCAAAU---AUUC-GAAUGA--G---------------AUCUUG-GAU-GACUG
AAGUGGAG-GAGGGUUUC-GUGUCAACA-GCAGUUGAACACGAGUUAGCCAA--UC-CUAAGCUCUAUGGG-
A-A-ACCU-----GAUAUAUAUUAA---GCAU----UUAA----CCA--AAUACAACAACGUAUACCAGCGU
GUUGAUGCAACCGAUACGAUAUAU-------AUUGAAUGA-GC-GA-A-AGGGA-AUCCGGUU-ACAA-UUC
CGGAG----------------------------------CCUGUUGGGUAUA--------C-GUU-------
U----------------GCAUUGGCGUCCC---AUACC----GGC-----------AACGGUAGCGCCUUUG
UAAUCA-UGG---CAAC--------AU-GAA--UCCU--------------------UUUC-U-UCGA-GAA
GCCAACA-------------AGAGAU-AUCGGAAGA-GU------UCUCU-------------U-UUCUGU-
UUUACA-GUC-AC----AC-U-------------------AG-----CC----------------AUGG-AA
G-U-CU--------------------UUCAU-AGAGAGAUAU---------GGUUGGA---CAG-GC-UGG-
UAG--AG-----------------------------------------------------------------
----CAUGGUA-UUAAAUUGCUGUG-U-CGAUAUU-CUCUUCUUGGA-CCUUGAAA-A-UCG-AAGACUGGG
-G-------------------------------------------CACG-----CAAACUCC-CA-ACAGCU
UGUACCGA-AUCCGCA------------------------------GCAGGUC--UCCAAGGU-UUAGAGUC
UCUAGU-C-GA-UAGA-UUAAUGUA-G-GUAAGGGAAGUCGGC-AAAUUAGAUCCGUAACUUCGG-GAUAAG
GAUUG-GCUC-U-GA--AGACUG-G----GAUGA-CUC-----GGGCUAU-ACC--C-----UC--CCG---
-GGG------U-----UCGCC---GC----------------------------------------GGG---
----------------------------------------------------------------------UG
-CCG-----------------GCC------------------------------------------------
---------------------------------------------------------CGCCU----------
--CG-----------------------------------------------CGC--GG--------------
---------------------------------------------------------------GUGC-----
------------------------------------------------------------------------
-C------------------UC--G-GU---CCCC-GG----------------------------------
----GGUC-C------GGG-----GUUUGCCUCAACGCGGU-------------------------------
-----CCG--------------------------------------------------------GGCCG---
-----------------------------UUUCG-------------------UGUG--GUGU---------
---------------UCG-------------------------CGCACCG---C------------------
----CGGCG-----GUCG------------------------------------------------------
----CCG----------------------GGGCC-GCUG----CAGUCA-UCA--AUAAACAGU--CAAUUC
AGA--ACUGGCACGG-CUGAGGGAAUCCGA-----CUGUCUAAUUAAAACAAAGCAUUGUGAUGGCCUC---
-AA-A-------------AGGUGUUG-ACACAAUGUGA-UUUCUGCCCAGUGCUCUG-AAUG-UCAACGUGA
A--------------GAAA--------------UUCAAGCAAGCGC--GGGUAAAC-GGCGGGAGUAACUAU
GACUCUCUUAAGGUAGCCAAAUGCCUCGUCAUC-UAAUUAGUGACGCGCAUGAAUGGAUUAACGAGA--UUC
CCUCUGUCCCUAUC----UAC-UAUCUAGCGAAACC---AC---AGCCAAGGGAACGGGCUUGGAAACACUA
GCGGGGAAAGAAGA-CCCUGUUGAGC-UUGACU--CUAGUCC-GGCAUUGUAAGGCGAUAUAAGAGGUGCAG
AAU-AGGUGGGAGCCGGGG------------------------------------------UAAAACAUUAC
CU--------------------------------------------CCCCG-G--C-AC-CAAUGAGAUA-C
CACCACUCUUACUGUUGCCUUACUUACAU-GA-------U-CA-G-GUGGAACAAGUG-----------CCA
------AG--GU-UA-------UUGC---AACC-UCCUG-------------GCAUAAGGUUUCUUGUUCAG
CGUUCAGUCAUGUCGUCAUU-CCUGGCCACAAGGCAGAAGACCGAGCCUGCGGUCAU-CGC---------GU
CCGUUGCGCGGCUCCCGGGUGUGC-GUUCCGG-------------------------------CCGGGUGCG
UUCGCGGCUGCCCCAGUAGGAUCCCGUCCGGUUU---GGGU----UCGCGU---------------------
-----GCGG-GCCA-----CGGCCGCAGUUUUGCUCAACUUUCACACCGUACGCCGA---UGACAG------
--UAAGACA------------------------UCUGGAUACUCCAAGUCAU-----GGACA-GUGCCAGGU
```

```
GCGGAGUUUGACUGGGGCGGUACAUCUCCAA-AACAAUAAC-GGAGGUGUCCA-AAGGUCAGCUCAGUGUGG
ACAGAAACCACACGCU-GAGCA-UAAGGACAA-AAGCUGGCUUGAUC-UCGA--------------------
--------------UGUUCA--GUACAUAUUGAGACAGCGAAA-GCUAGGCCUCACGAUCCUUU--------
---UG-G-----UUUAAAGAG--------------------UU-UUUAGCAAGAGGUGUCAG-AA--AAGUU
ACCACAGGG-AUAACUGGCUUGUGGCCGCCAAGCGUUCAUAGCGACGUGGCUUUUGAUCCUUCGAUGUCGG
CUCUUCCUAUCAUUGUGAAGCA-AAAUUCACAAAGCGUAGGAUUGUUCACCCUUUCA-AGGGAACGUGAGCU
GGGUUUAGACCGUCGUGAGACAGGUUAGUUUUACCCUACU--------------------G-GUGUG-----
--------------------CGAACGAACACGA--A-CGCUAUCUUAACGGA-AAU-GCUGUGCAGUACGAG
AGGAA-CCACAGCUUCGAACCAA-UGGC-UCAAUACCUAGUUC-GACC-GGACUU-U-GGUAUAACGCUA-C
GUUCG-UUGGAUUAUGCCUGAACGCC-UC--UAAGGUCGUAACCAAACCGAGC---UG--AUAGUGUC-UCC
UUAUAGGU------CGU-------------------------------------------------------
------------------------------------------------------------------------
------------------------------------------------------------------------
---------------------CGGCGAUCAUGU----------GGCAAAAAAC---CU--------------
--------------------ACAAGACC---UGCUAG------------CGUGAU-----------------
---CUCAC--GU-------UG--GCA-----UCUUAUUGCAU------------------GCACGCA---GA
CAAAGCCCCGCG-CGCGCAC-----CAUACAGC-----AAG--UAUCUA----AGAUA-----CUGGAACG
----------------------------C----U-GGUGGCGCUGCU---CAGCAUC-AAU----------A
U-----------------AUGAUUU---------------------C-GAUACCUG-----AGA------CC
CCCUACAAACGAUAGGUU-UAC-AGGCUG-GGA-GUUGCGAG-UUGCA--------GAGAGGU-GU------
----A----CAUUUCGAUCCUCAGUCGUU----UCUG-UUGUCGAC---GGGUCU--A--------------
------------------------------------------------------------------------
------------------------------------------------------------------------
------------------------------------------------------------------------
-------------------------------------------------
>LSUe_185431_TRYB2/1-4086
----------------------------------------AACGUGUCGC-GAUGGAUG-A-CUUGGCUUCC
U-A---UUUCGUUGAAGAACGCAG--CAAAGUGCGAUAAGUGGUAUCAAU--U---GCA--GAA-UC--AUU
UCAUUGCCCAAUCUUUGAACGCAAAC-G--------GC--------GC----AUGG-G-AGAAGCUC---UC
U-C-------GAGCCAU--CC----C----C---GUG----C-A-UGCC-ACAU--UUCUCAGUGUCG-AAU
-----------------------------------------------------------UCAGAC-CUGA-G
U-G--------------------UG-G-CA-GGACCACCCGCUAAACUUAAGCAUAUUACUCAGCG--GAGG
AAAAGAAAACAA--CCGUGAUUCUCUCA--GUCAG-CGGCGAGCGAAGAGGGA-UUCA-ACUCGUUGCCGAA
UCGG-GUCUUAC-------AU------------G---ACC--UCGAACUGUGGCA--CCAU-----UUUUGU
GGCC-GUUC-UUA-U--AGGG-------------------------CUGGU--GCA-----------GG---
--------CGUGG--CGG-------AAUUCA-AUUG-CAAAGCAAACCCGUUGCUGAACACAUUUACAACCC
UUCAUGUGAG-UA-UUGAG-------CCAAAGAAGGUGUCAGCCCAUUACAUUCCUUAGCC---A----CGG
AGCCU-AC--GCGCCUCCUCAU--------------GUUC-GGUCC--AAGAGAGUAGCACUGUUUG---GG
AAUGCAGUGUCAAGC-CGGCAGGUAUU-UGUCUGCCAAGGUUAAAUAUAGAGUA-GGA-AGACCGAUAGC-G
AAC-A-AGUAGCGUGAGCGAAAGUUUGAAA-AGUAC-UUUGG-AAAGAG-A-GUGACAUAGAACCUGAAAUC
G-UGACAACG--A-CAA-UG-GAAG---UACCUUCAUUUCUUU-----------------------------
------------------------------------------------------------------------
------------------------------------------------------------------------
------------------------------------------------------------------------
------------------------------------------------------------------------
```

```
------------------------------------------------------------------------
----------------------------------------------------------------UUUUUAC-
---GG-------------------------------------------------------------------
--------------------CCAGA------------------CCCAACCUCGCCGCGC----AAUUUCCUU
CCGGGGC-------------------------------------------------------U--C-AA---
------------------------------------------------------------------------
----------CAUCCGGGUUUCACUGUGCGGGUUUCUC-------UUUUG--AGGUCAUGGGGUGCGUGAGU
GGGCGGAAU---------------------------------------------------------------
------------------------------------------------------------------------
---------------------------------------AAAGGG---------------------------
------------------------------------------------------------------------
-------------------------------------------------U----------------------
----------------------CGGUGU-------GUU----------------------------------
------------------------------------------------GUGUG----UGGGUU---------
-----UUUCUCU--CAC------------------UCUCU------UCCUCCUUC-----------------
---------CCUGUC---CGUGCG--------------------------------------------UCU-
------------------------------------------------------------------------
----------------CGACGCUGGCGA-------------UGGAAAUGG--G---GUGCCCCCACCCGUCU
UGAAACACGGACCAAGGAGUCAA-A-CAGACGCGCAA-GGGAGGAG--U-AUGU------AUGU-----GAG
A-AC--UCUUUUCUCGUACUGAAAG-GGA-----------------GGUGUGAU------UUUACACCCGUG
GCGU-GCGCAAU----U---------------------------------------------UC--------
UCU---------------------------------------------------------------------
-------------------UUUU-------------------------------------UGUGUUUUU-GC
CCAACG----CCGGCCG----------------------GCCC-----------------CCCGUGGG-CU-
GCG-CG----------------------CGAGUGUGCCUGUUUG-GACCCGAAAGG-UGGUGAACUAUGCCU
GAACAAGGUGAAGCCCGG-GGAAAC-CCCGGUGGAGGCCUGU--AGCGAUGCUGACGUGCAAAUCGC-UCGU
ACGAUUUGGGUAUCGGAGCGAAAGACUCAUCGAACCACC-UAGUAGCUGGUUCACGUCGAAG-UUUCCCUCA
GGAUAGCUGGUG-CUUG-------------------------------------------------------
----------------UCUUUAAG-----AAGUAUUGUGCGGUAAAGCAAAUGAUU-AGA-GGCAUUGG-UG
U-U-CUUGU-AAUA-U-CGACCUAUUCUCAAACUUUAAAUGUGCAAACA-----------------------
------------------ACCCA-GC-----CUUAGU--UAUU--UUAACAAG-------------------
GCUUUUGGG-UACAUAAGA-AU-U--G-----A--GGCACCA-AGUGGGC-C-UUCU-U-CGG-UAAGCGGA
GC-A-GGCGAUGC-GGAAUG---AACCGCUUAGGG-A-UAUGUGGCGUCAAAACUUGU-GGG-CU-CAUUCU
-C-GAUACGU-GAAAAGGGUGUUGGUGGAUAGGGACAGUUGGACGGUGGCCAUGGAAGUCGGCA-UCCGCUA
AGGAGUGUGUAACAACUCACCAACC-----GAAUCCACCGGCCCCGAAAAU-GGAU-GACGCUUA-AGCCCC
AUGUGCGUGAUUGCCCAUUAUUCCUUUGGUGAGG--GCGGC-------------------------------
-----------------------------------------------UA-----------------UAUU--
----UCUUU---------GUGAGA------------------------------------------------
-------------------------AAGUGUAGCCCU-CCCG-AUGUAGACGUGGCGUGGGGG---------
-----------U-G-AGGAC-GAAGCCGA-U-GGCGCGAGCC--UCGGAUGGACCC-G-CCU-CUAGUGCAG
AUCU-UGGUUG-GCGUA-GCAAAG---AUCU-AACGGAAUAUUUUUUCUUAAAUGCAACGUU-GGA-UACUG
GAGCGGGG-AAGGAUUUC-GUGCCAACG-GCACUCGUACACGAGUUGUUCGG--UCACUGAGCACAACGUU-
A-C-ACCGUUUU-GUUAGGAAAGUGAAGGUG-----GUCGGC-GCACUU-CUCCCUUUUU---------UGG
GGU-------------UUGUGUGUUUUACCGGCU----UA-UCUGA-A-AAGGG-GCAACA---GAGA---A
CC----------------------------------------------------------------------
```

```
----------------------------------------------------------------AC------
--UGUGGAAA---UGCG--------AA-AC-----AC--------------------UUGC-C-AGGU-GAC
CCAUCC--------------AUCC-U-CCCACGGUG-AG-C----UUUCU-------------U-UUCACC-
AUAAUC-CAC-AU-----C-U-------------------CC-----GG----------------CCUU-GC
U-G-GGGCUUGGGCCUCUCGACUUCUCGCGU-UACUCGGAGC---------GGGGGC----UCA-AG-AUU-
GAA--AA----------------------------------------------------------------A
--UGCAGCUCA-CCCUACGUACUGU-C-CUUUGUU-GUGAGUUCGGC-GCAUUAAA-G-CAA-AACCUGGG
-G----------------------------------------------------UGUCCCAA-CU-GCAGAC
CGUACUCAUCACCGCA------------------------------UCAGGUC--CCCAAGCA-UCGAAUGC
UCUGGU-A-CA-UAGA-GAAAGGUA-CACUCAGGGAAGUCGGC-AAAAUAGAUGCGUAAGUUCGC-AAGAAG
CAUUG-GCUC-U-GA-GGGCAAA-G----UGGGG-GU------CCACAG--GAG-GGCCAGUG---UCC-UU
UCGU--------GCGCGGCC----GU----------------------------UU---------UCUG--C
GUUU-----------------------------------------------------------------UG-
-UU-----------------GUUU------------------------------------------------
-------------------------------------------------------UGGCGUUCG-------C
GCCUU-------------A---------------------------------GC--AC--------------
----------------AUAC-------------------------------------------GCGG-----
---------------------------------C-------GACCGUUU---------------------UG
-CUCGG-------AACG--UGAAGG-CUA--CCCU-CCCGAGCC---GUAUCUACUAACCUCC--UUCUGGU
GUGUUGUUUUUA-CGUCUCA----CUCCCGC-----GCA---------------------------------
----------------------------------------------------------------GGUG----
-----------------------------UU---------------------UUCGCUUCCUU---------
---------------UUU-----------------------------UGGU-UG------------------
------------------------------------------------------------------------
----CG-------------------------------------CA-----CCU--ACAAAUUGC--CAACUC
AGA--ACUGCUUACGGC--GGGGAAUCCAA-----CUGUAUAAUUAAAACAUAGGUUUGUGAUGCAUCC---
-AA-G------------UGGUGUAUG-UCGCAAACUGA-UUUCUGCCCAGUGCUCUG-AAUG-UCAACGUGA
C--------------GAGA--------------UUCACCGACGCGC--GGGUAAAC-GGCGGGAGUAACUAU
GACUCUCUUAAGGUAGCCAAAUGCCUCGUCUUC-CAAUUAGAGACGCGCAUGAAUGGAUUAAUGAGA--UUC
CCUCUGUCCCGAGU----UAC-UAUCUAGCGAAACG---AC---AGUCAAGGGAACGGACUUGAAGGGCUAA
GCGGGGAAAGAAGA-CCCUGUUGAGU-UUGACU--CCAGUCU-GGCUCUGUGCGGCGACUUUGGAGGUGUAG
UAU-AGGUGGAAGC------------------------------------------------------GCAA
----------------------------------------------------G--C-UC-AAAUGAAAUA-C
CACCACUCGGAACGUUGCUUCACUUAUCG-AA-------U-GA-A-GAGAC-CAAUGGUUGUUUU----GCG
------CA--GU-CU--------UCG---GGCU-G--UG-------------CGCCG---UCU-------AG
-GCU------GGGCAUAAUU--GUGGUGUUU----------------CUUUCACCGC-CAC---------GG
------------------GAGGGGUGU--UGGUUUUUUAUCCUUCUCUCCGUUUCUCCUUU-UCUCUCUUUC
U--CCCUUU---------------------UUUU---GGGCGUUUGAGGGUU--------------------
----GGUGACCCG------UGGGCG---------------------------CCCAG---CCGUGCGGAACU
UCUGCGCCAACGGGGUGGGAACUUUUCGUUUCUUCUCCGGAGUCUUGUUUCG-----AGACAUCUGCCAGAU
GGGGAGUUUGGCUGGGGCGGCAUAUCUGUUA-CACGACAAC-GCAGGUGUCCU-AAGGCGAGCUCAGUGGGA
ACAGAAAUCUCACGUG-GAACA-CAAGCGUAA-AUGCUUGCUUGAUUAACGA--------------------
--------------UUUCCA--GUACGAAUCGAGACUGCGAAA-GCAAGGCCAG-CGAUCCUUUGCGAAGAA
CAGGA-A-----AUAUGAACA--------------------AA-UCAUACCAGAGGUGUCAG-AA--UAAUU
ACCACAGGG-AUAACUGGCUUGUGGCGGCCAAGCGUUCAUAGCGACGUCGCUUUUUGAUCCUUCGAUGUCGG
```

```
CUCUUCCUAACCUAGCGCCGCA-GAAGACGCUAAGGGUUGGAUUGUUCACCCACUGACAGGGAACGUGAGCU
GGGUUUAGACCGUCGUGAGACAGGUUGGUUUACCCUACUUGGCUGGAGAUUGCGAGUAAA-AGAUUAUCCG
CGCAAAGCGGAGUGAGAUUGUGAAGGGAUCUCG--C-AGGCAUCGUGAGGGA-AGU-AUGGGGUAGUACGAG
AGGAA-CUCCCAUGCCGUGCCUC-UGGU-UUCUGGAGUUUGUC-GAAG-GGCAAG-U-GCUCCGACGCUAUC
GCACG-GUGGUUCUCGGCUGAACGCC-UC--UAAGCCAGAAACCAGUCCCAAG---AC--CGGGUGCC-CGU
UCUCGAAU-CGCCACUU-------------------------------------------------------
------------------------------------------------------------------------
------------------------------------------------------------------------
---------------------CUCUUAUC-------------------------------------------
-------------------------------UGGUGG------------GCUCU------------------
----------GC-------CC--GCCCU---------UGUCG------------------GCGCC-------
---------------------------------------CAGU---------------------ACCUUCAU
UU--------------------------U----C-AC-----------------------------------
---------------AAGAUCUUUUG--------------------U-CCCUCUCC-----AAA------CG
AGAGUACAUGCAUGGGCU-GGC-AUGAGCGGCA-CGCUCCAC-GACCGUGGGGCUCGAGGGGCACU------
----UUA--CGUCCCGAGGCGCUGAACCUUGA-UGCU-UGGAAUUUC--AUGCUCAGG--------------
------------------------------------------------------------------------
------------------------------------------------------------------------
------------------------------------------------------------------------
-------------------------------------------------
>LSUe_28002_HEXIN/1-3047 KC249997.1:1-144,195-3097 Hexamita inflata METAMONA
----------------------------AUC--------UAACGGAUAAU-AGUGGAUC-A-CUAGGC-UUG
G----ACAUCGACGAAGAGCGUAGAACAGAGCGAAAUUUGUGGCGACGAGCCUCAGCCC------UAU-GCG
AGU-UGU-CAAUUCUCGAAUACCUCA--------GGCC--------GC------------------------
------------------------------G---AGGCCG-C-G-CACC-----------------------
-----------G-----------UACA-------------------------------GAGCGAC-CUGU--
------------GC---------GGUA-CG-AGACUACGGGCCGAACUUAAGCAUAUCAGUAAGCC--CAGG
AAAAGAAAUCAAA-GCGAGAUUCCCCCA--GU-AG-CGGCGAGCGAACAGGGA---CGUGCCAGGCCUGAAU
UCCA-----------------------------GUUGUGGGCGUGCAUUGCAGCG-----------------
-----GGGG---------------------------------------------------------------
-----------------------CAAAGUGCCGUUG--------AACCGGCC--------------------
-------GGACCA-AACAGUG-----CAACUCUGGUUGGCCCCUUGCACU-------AGC-----------
-----------------------------------------------UGGCGUAGUAGUACUGCUUG---AG
AGUGUAGUAUGAA-UACGGGUGGUAUU-CCUCAUCCAAGGCCAAAUACAGGU---CCA-AGUCCGAUAGC-G
UAU-CGAGUAACGUGAGUGAAAG-GUGAAA-AGUAG-ACUCA-AAGGGGUU-GCU-UAAAAGACGUGAAACU
AUUAUUUUGAAGG-GUAAGC-GGGGGCAGCGC----------------------------------------
------------------------------------------------------------------------
------------------------------------------------------------------------
------------------------------------------------------------------------
------------------------------------------------------------------------
------------------------------------------------------------------------
---------------------AUCUG-----CC---------------------------GAACCUCUCCU-
--AGA----------GGG------------------------------------------------------
------------------------------------------------------------------------
-----------------------------------------------------------------C------
```

```
------------------------------------------------------------------------
------------------------------------------------------------------------
------------------------------------------------------------------------
------------------------------------------------------------------------
------------------------------------------------------------------------
------------------------------------------------------------------------
------------------------------------------------------------------------
------------------------------------------------------------------------
------------------------------------------------------------------------
--------------CGC-------------------------------------------------------
------------------------------------------------------------------------
------------------------------------------------------------------------
--------------------------------------------------------CGCUUUCCGCCCGUCU
CGAAACACGGAUUAAGGAGUUAC-G-UUAGACUGCGA-GCCAAUGC--G---------UGCAAA--------
------CGCAAGGGCGCAGCGAAAG-CGGAAAUACU------------------------------------
-----CAGCCAUGU----------------------------------------------------------
------------------------------------------------------------------------
-------------------UAGGGUGCACC----------------------------------------GG
UUCACC---------------------------------------------------------------GG-
GGC-----------------UCCUGUACAGAGUUGUCUGAUGUA-AACCCGAAAGG-UAGUGAUCUUUACAC
GGCGAGGAUGAAGCCAGU-CGAAAG-GCUGGUGGAGGUCCGC--AGCAAUACUGACGUGCAAAUCGU-UUGU
CAGAGCUGUGUAAAGGGGCGAAAGACUCAUCGAACUUCC-AAGUAGCUGGUUACGUUCGAAA-UUUCUCUCA
GGAUAGCAGCAG-----------------------------------------------U------------
----------------------GA---UUUAGCUCCGACAGGUAGAGAUAAUGAUU-GGU-GGAAAGUGGCC
UCC-------GGGUAU-GCCCCAACUCUCAAACUUCGAAUAGUCGGGAU-----------------------
-----------------CGCGGG--------UCUUCCG-CAAG-----GAG---------------------
CCG-------CCGCGGUGGAGU---AUG-------CACUGCU-AGUGGGCCC-CUCU---GGG-UAAGCCGG
AU-G-GGCAAGGA-GGAUUC---AGCCUUAAGUGG-A-GUUACGGUGCGCAACA-----GGGGCG-CA----
-UGAGA-----AAAAACGACGUGUGCUGAUACAGACAGCAGGAAGGUGGCCAUAAAAGUCGGAA-GCCUCUA
AGGAGUGUGUUACAACUCACCUGCC-----GAAUCAGUACGCUCGGAAAAU-GGAU-CGCGCUGC-AGCCUC
U----------GACCUAC-ACUCCACUGCACGCC------------------------G-------------
------------------------------------------------------------------------
------------------------------------------------------------------------
--------------------------------------UGCC-AGGUAGAACGUG-GUGGUGC---------
-------------GGGCGUC-GAAGCUUU---CUCGUGAGAGG---GGGCGGAGCG-C-GCA-CUAGUCCAG
AUCU-UAAUUG-GAGUA-GCCAAU---AAUCUGCAUGC--A----------------------GAC-CACUG
AAGAGGAG-AAGGAUUCC-GCAGCCACU-GAAACAGUCUGCGGGUGACUCGG--AC-CUAAGCGCUCGGCC-
A---CGGCC----AUUUAAG----------------------------------------------------
-------C--------------------------------GUC-GA-A-AGGGA-AGCAGGCU-AAGA-UCC
CUGCA-----------GGC--------------------CCGCC---------------CACAU--------
------------------------------------------------------------------------
-------UCGG--CAACG---------------------------------------------------GAC
GGCGCAGUUCACGGUCUCUCGGUCGGCGCGGUCGGA---------CCAUC-------------GCACCGGC-
UUAAUGCUCC--------A-G-------------------GG-----CGU---------------GGGG-AA
G---AG--------------------CUAAG-CUCUAGAGCCACGUUUCGCAGCUAGA---AUA-CGUCCGC
```

```
GUAC-AC--------------------------------------------------------------GGA
--------------------------CGCGU-GCGGGGCCGAGAGACGCCUCAAAAUGGACU----------
-GC---------------------------------------------------GCGACUCG-GGCGGGCAC
CGUACUGAUAACCGCA------------------------------UCAGGUC--UCCUAGGC-AAGAAACC
UCUAGU-G----AGAAGAUAACACG-A-CCAAGGGAAGUCGGC-AAAUCGGCUCUGUAACCUUGG-GAUAAA
GAGUG-GCUC-U-GA-AUGCUCGUG--CCGGCGGCGCU------------------------GC------AU
UGAG--------------------AC---------------------------CUCGGGGAAGC-GUCG---
------------------------------------------------------------------------
------------------------------------------------------------------------
------------------------------------------------------------------------
------------------------------------------------------------------------
---------------------------AA----------------------------------GUGGG----
--------------------------------GC--------------------------------------
-------------------------------AACCUGC----------------------------------
---CAG---AUGUGAGCUG-----------------------------------------------------
----------------------------------------------------------------GGUG----
-----------------------------------------------------------AUCU---------
---------------UUGC---------------------------------AG------------------
------------------------------------------------------------------------
-------------------------------------------CCUCUA-ACG--CACCAUGGG--CAAUUC
AGA--ACUGCUACUGACUAGAAGAACCCGA-----CUGUAUAAUUAAAACAUAGCAUCUCGAUAGCCGC---
----AG-----------CCGGUUUUG-ACGAGAUGUGA-AUCCUGCCAGUGCUAUG-ACUG-UCAACGUGA
CGAG----------------------------AGUCAUUCAAGCUC--UGGUAAAC-GGCGGGAGUAACUAU
GACUCUUUUAAGGUAGCCAAAUGCCUCGUCCAG-UAAUUUUGGACGUGCAUGAAUGGAUUAAUGAGG--UUC
UUACUGUCCCUGGU----CGU-AUCUUCGCGAAACC---AC-AGACUCGGGAACGGGCGAGUGUAACCCUCA
GCGGGGCAAGAAGA-CCCUGUUGAGC-UUGACU--CCAGCUUGGGAAUUGUAGAACGGUGAUGGUGGUAGAG
CGU-ACAGGGGAGG------------------------------------------------------GCGA
----------------------------------------------------C--C-GCAGCAUGUAACA-C
CCUGACACUACUCAUUGUUCUGCUAACCC-GG-------U-AA---CCGGC---------------------
-------------CC---------------------------------------------------------
----------------------------------------------------------GGC-----------
------------------------------------------------------------------------
------------------------------------------------------------------------
-----------------UUUGG------------------------------CCUG----GGGCUUG-----
------------------------------------------------UCGG-----GGACA-UUUCUAGGU
GGGGAGUUUGGCUGGGGCGGCGCGCCUGCUA-AAGCGCAAC-GCAGGCGUCCU-AAAACAAGCUCAGACAGG
ACGGAAACCUGUCGCA-GAGCA-AAAGAGCAA-AAGCUUGUCUGAUU-GCGU--------------------
--------------GCGUCA--GUGGCGUAUGCAACCGUGAAA-GCGUGGUCUAUCGAUCCUUUAC------
---GA-------UUCUUGGAA-----------------------UGAAUCUGGAGGUGACAG-AA--AAGUU
ACCACAGGG-AUAACUGGCUUGUGGCCGCCAAGCGACCAAAGCGACGUGGCUUUUUGAUUCUUCGAUGUCGG
CUCUUCCUAUCAUUCUGAAGCA-GCAUUCAGAAAGUGUUGGAUUGUUCACCCA-UGCAAGGGAACGUGAGCU
GGGUUUAGACCGUCGCGAGACAGGUUAGUUUUAUCCUACU--------------------G-AGGACC----
-------------------------------GCGCA--------CCGCUAGU-AAU-CUAGAUUAGUACGAG
AGGAA-CAUUUUUAUCAAAGCAC-UGGU-U-UACAGCCUUGUUUGACA-AAGCAG-U-UGGCUGAAGCCACC
CUUUG-GCAUGUUAUGGCUGACAGCC-UC--UAAGUGAGAAUCUCCCCUACAGGUGUG-UAUGGUUUCACC-
```

```
------------------------------------------------------------------------
------------------------------------------------------------------------
------------------------------------------------------------------------
------------------------------------------------------------------------
-------------------------------UGCUUA-----------------------------------
------------------------------------------------------------------------
------------------------------------------------------------------------
----------------------------UGA---------------------------AUUAGAAG------
----------------------------------------------------------CG-AAACCGCCGCU
CUCCAUAGACCAUGGAGCUCGAGUGCACUGUGA-GCUAUAAG-------------UAAA-------------
------------UUCAAGUAGUUGAGGUGCCA-CUCUGUGUUAUUAUUCAUUUUGCUG--------------
------------------------------------------------------------------------
------------------------------------------------------------------------
------------------------------------------------------------------------
-------------------------------------------------
```
