## Supplemental Data S4 for "Supersized ribosomal RNA expansion segments in Asgard archaea"

### Report

| | transcripts |
|---|---|
| # contigs (>= 0 bp) | 3258 |
| # contigs (>= 1000 bp) | 82 |
| # contigs (>= 5000 bp) | 1 |
| # contigs (>= 10000 bp) | 0 |
| # contigs (>= 25000 bp) | 0 |
| # contigs (>= 50000 bp) | 0 |
| Total length (>= 0 bp) | 1047911 |
| Total length (>= 1000 bp) | 129286 |
| Total length (>= 5000 bp) | 5447 |
| Total length (>= 10000 bp) | 0 |
| Total length (>= 25000 bp) | 0 |
| Total length (>= 50000 bp) | 0 |
| # contigs | 395 |
| Largest contig | 5447 |
| Total length | 333673 |
| GC (%) | 48.26 |
| N50 | 799 |
| N75 | 622 |
| L50 | 125 |
| L75 | 244 |
| # N's per 100 kbp | 212.78 |
| # predicted rRNA genes | 0 + 15 part |

All statistics are based on contigs of size >= 500 bp, unless otherwise noted (e.g., "# contigs (>= 0 bp)" and "Total length (>= 0 bp)" include all contigs).

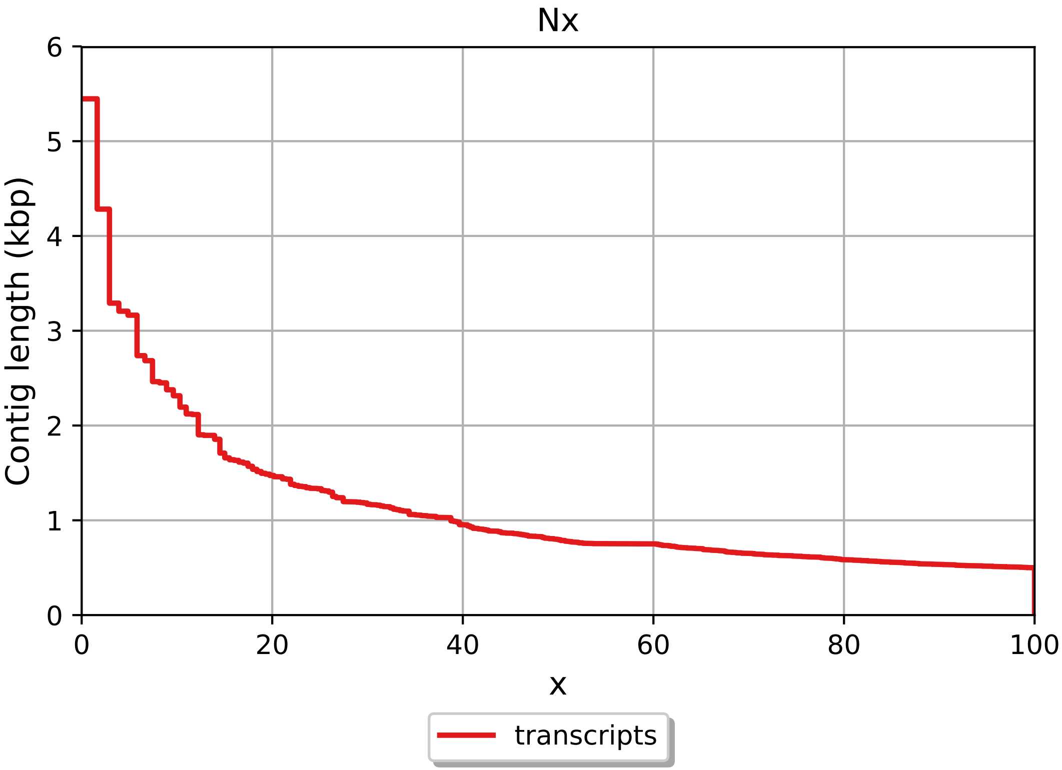
Nx
Contig length (kbp)
0
1
2
3
4
5
6
0
20
40
60
80
100
x
transcripts

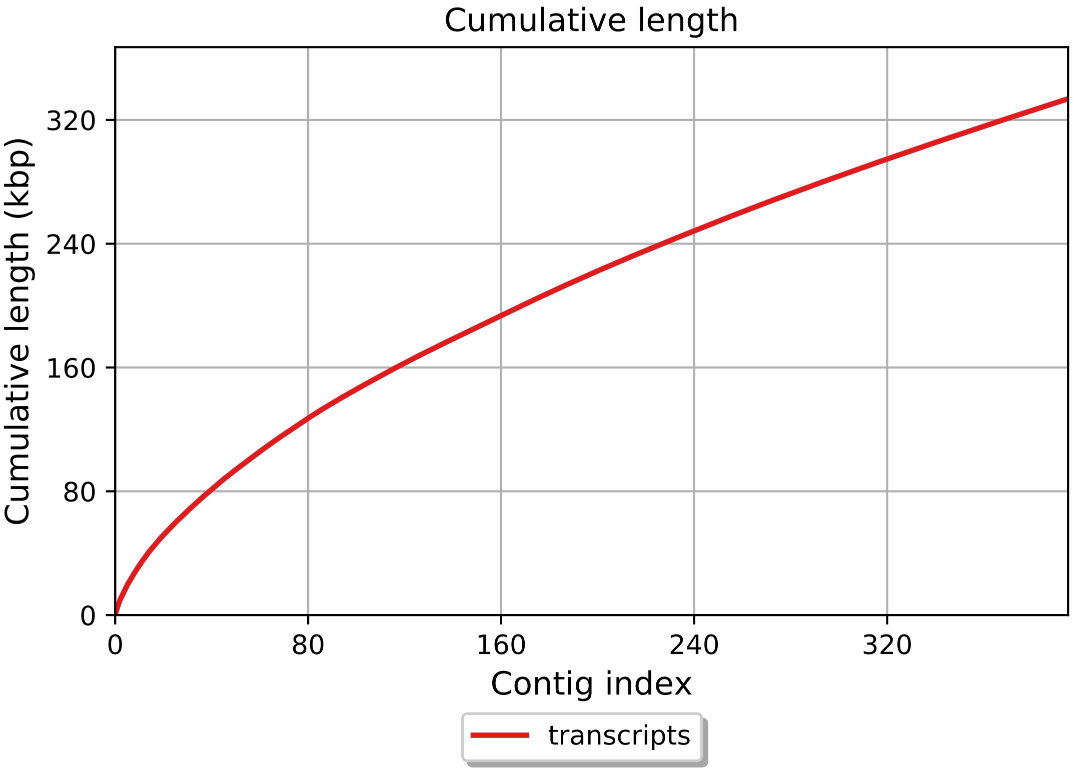
Cumulative length
Cumulative length (kbp)
0
80
160
240
320
0
80
160
240
320
Contig index
transcripts

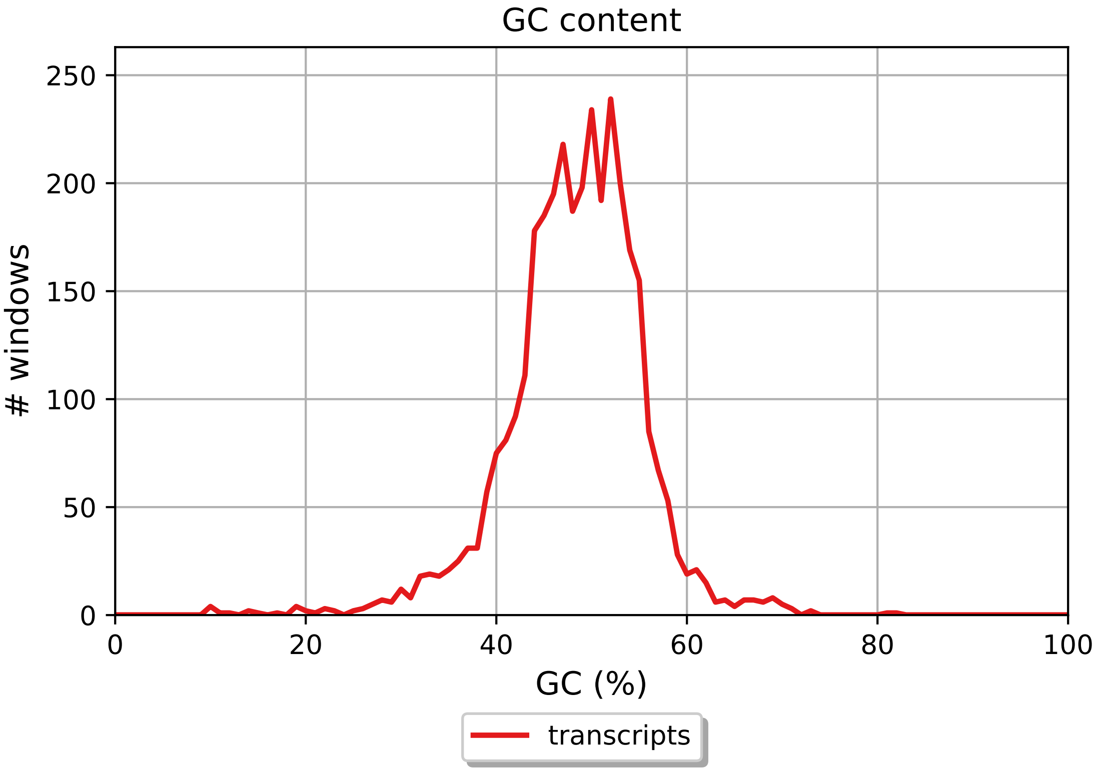
GC content
### windows
250
200
150
100
50
0
0
20
40
60
80
100
GC (%)
transcripts

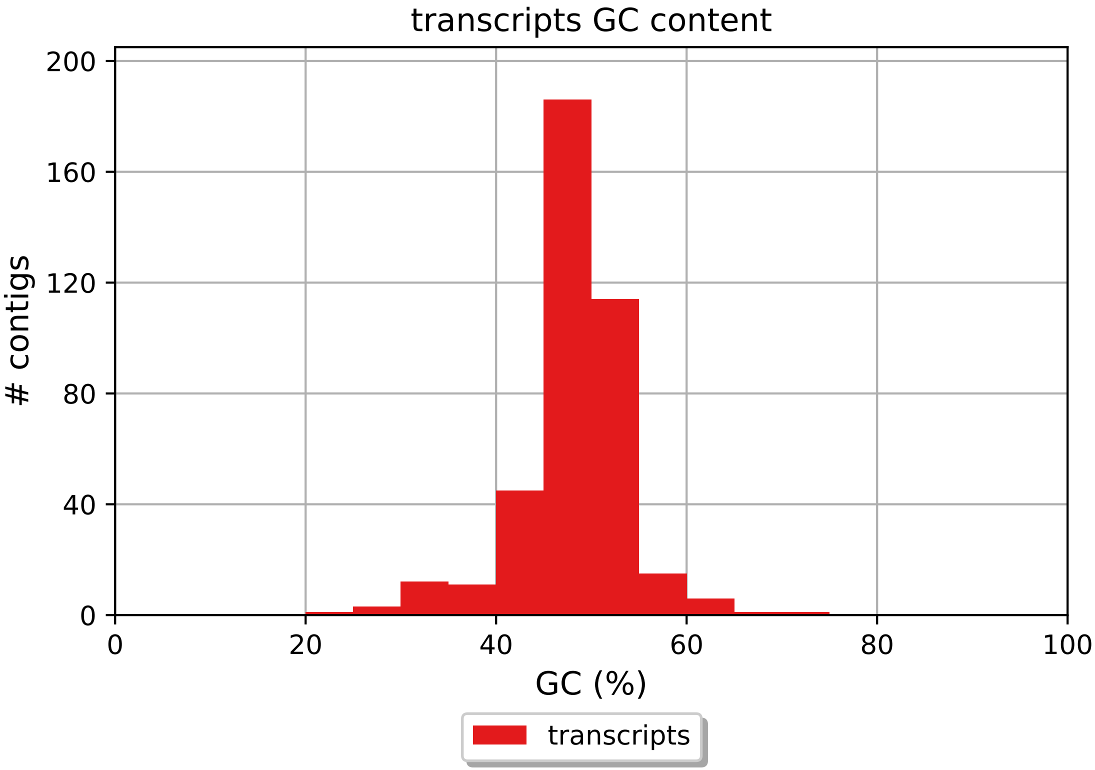
transcripts GC content
### contigs
GC (%)
0
40
80
120
160
200
0
20
40
60
80
100
transcripts

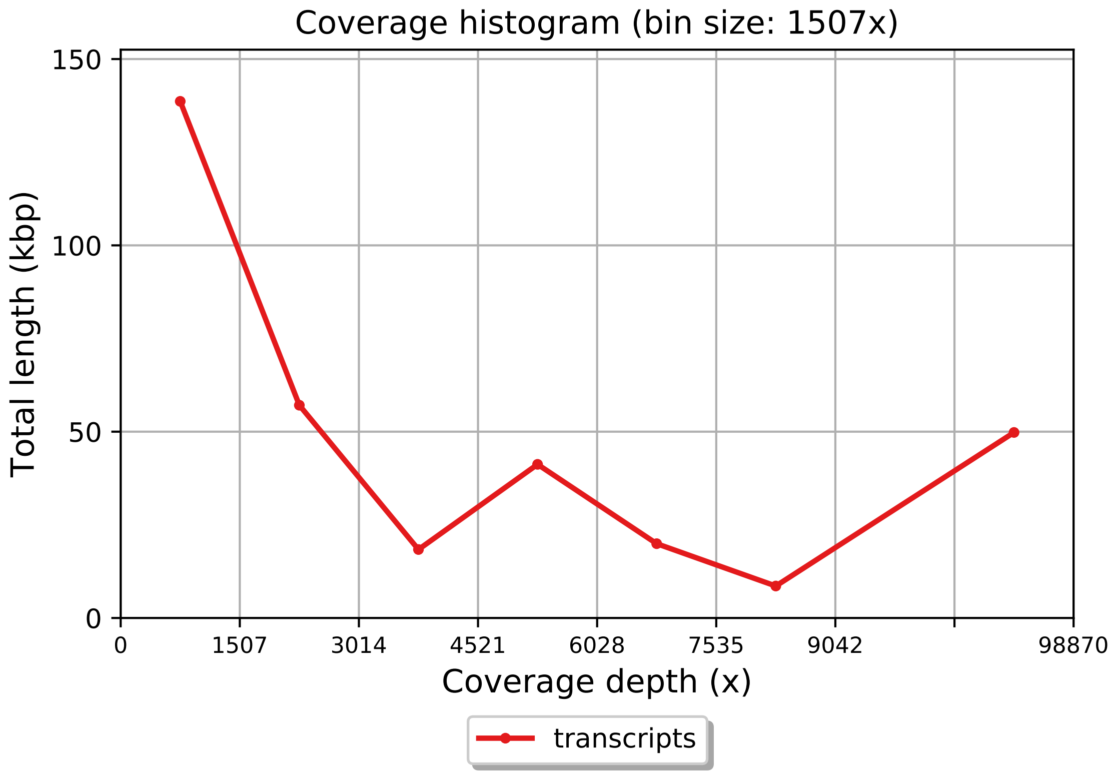
Coverage histogram (bin size: 1507x)
150
100
50
0
Total length (kbp)
0
1507
3014
4521
6028
7535
9042
98870
Coverage depth (x)
transcripts

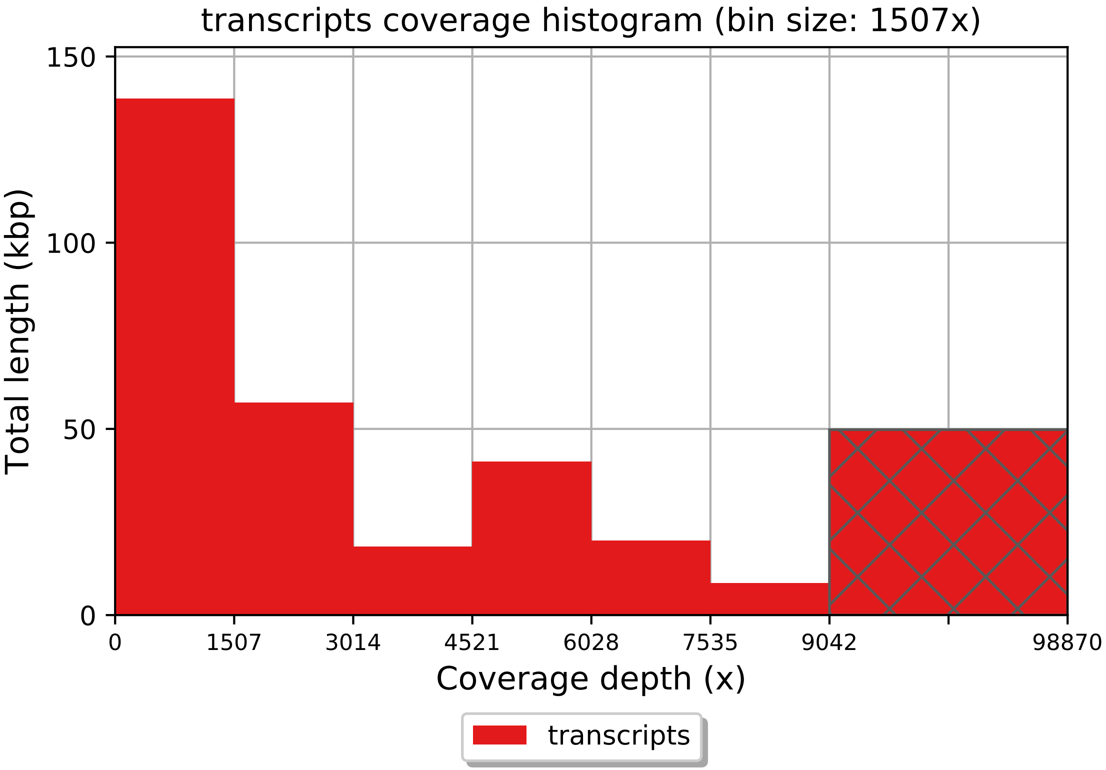
transcripts coverage histogram (bin size: 1507x)
Total length (kbp)
150
100
50
0
0
1507
3014
4521
6028
7535
9042
98870
Coverage depth (x)
transcripts
