## Supplemental Data S5 for "Supersized ribosomal RNA expansion segments in Asgard archaea"

### Report

| | contigs |
|---|---|
| # contigs (>= 0 bp) | 4102 |
| # contigs (>= 1000 bp) | 4 |
| # contigs (>= 5000 bp) | 1 |
| # contigs (>= 10000 bp) | 0 |
| # contigs (>= 25000 bp) | 0 |
| # contigs (>= 50000 bp) | 0 |
| Total length (>= 0 bp) | 589007 |
| Total length (>= 1000 bp) | 8931 |
| Total length (>= 5000 bp) | 5441 |
| Total length (>= 10000 bp) | 0 |
| Total length (>= 25000 bp) | 0 |
| Total length (>= 50000 bp) | 0 |
| # contigs | 83 |
| Largest contig | 5441 |
| Total length | 58393 |
| GC (%) | 46.72 |
| N50 | 628 |
| N75 | 556 |
| L50 | 31 |
| L75 | 56 |
| # N's per 100 kbp | 0.00 |
| # predicted rRNA genes | 0 + 1 part |

All statistics are based on contigs of size >= 500 bp, unless otherwise noted (e.g., "# contigs (>= 0 bp)" and "Total length (>= 0 bp)" include all contigs).

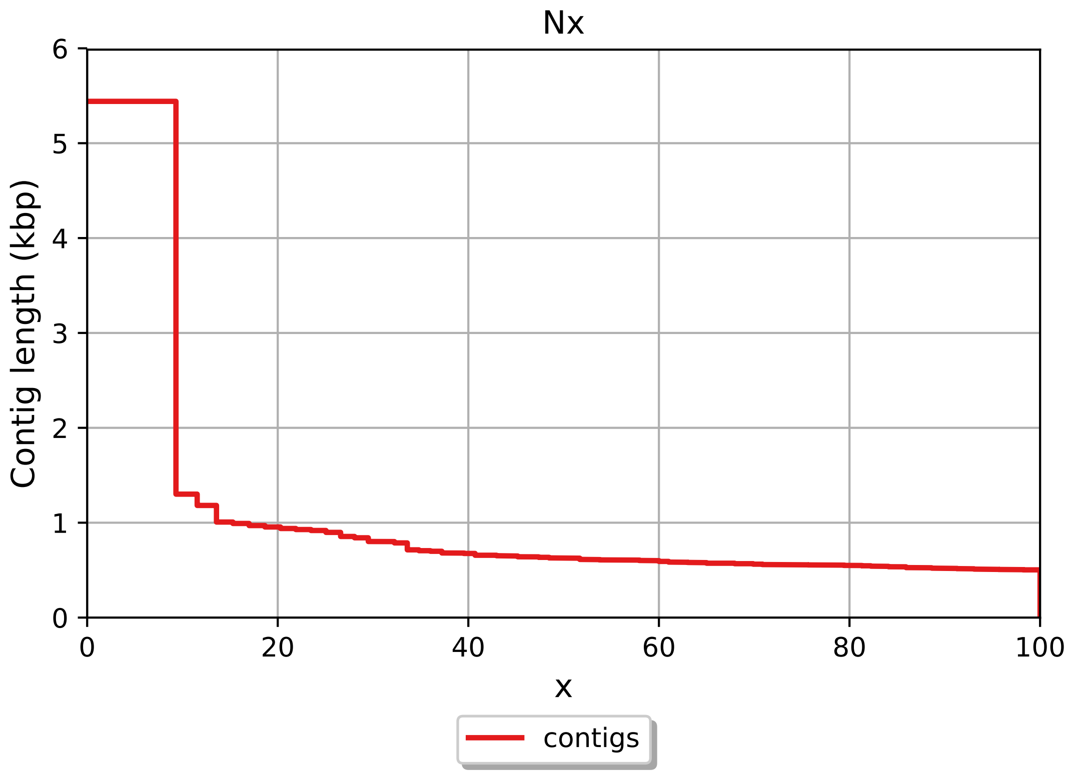

Nx
6
5
4
3
2
1
0
Contig length (kbp)
0
20
40
60
80
100
x
contigs

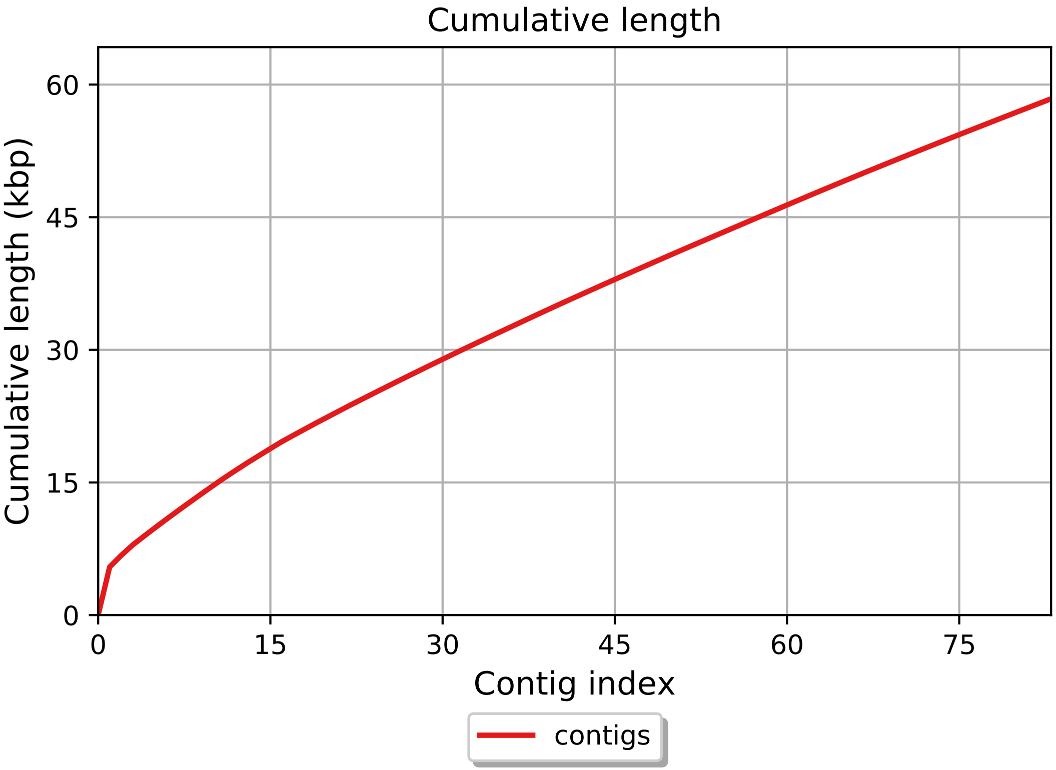

Cumulative length
Cumulative length (kbp)
60
45
30
15
0
0
15
30
45
60
75
Contig index
contigs

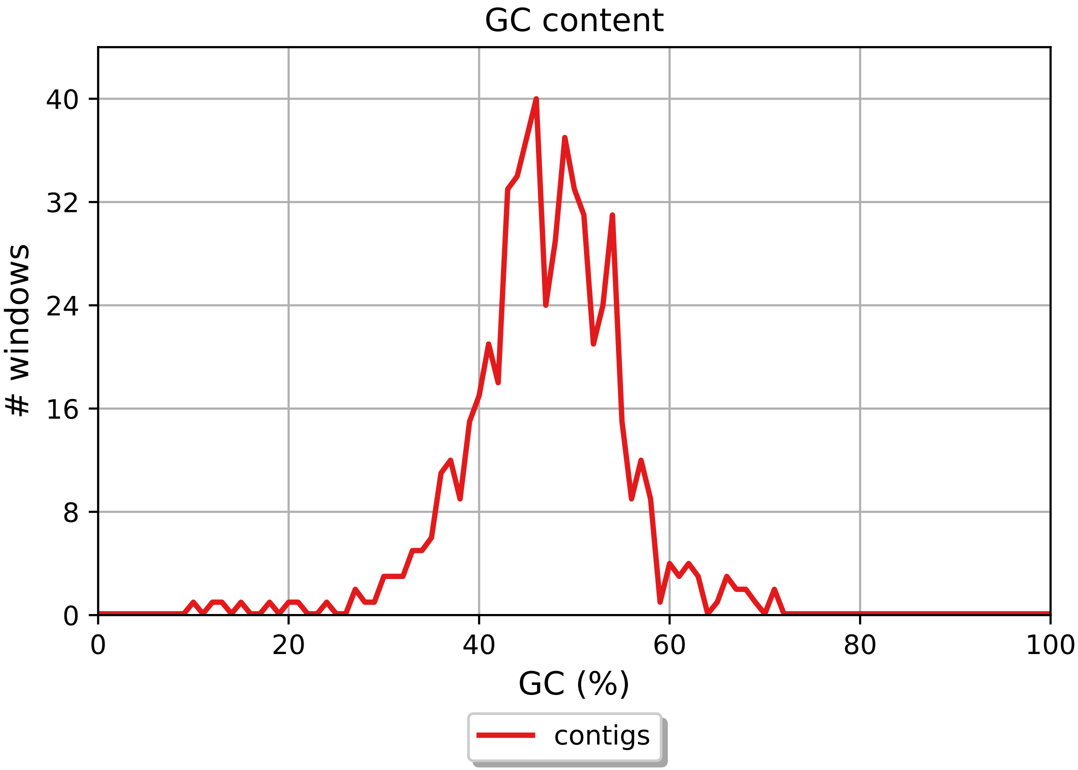
GC content
### windows
0
8
16
24
32
40
0
20
40
60
80
100
GC (%)
contigs

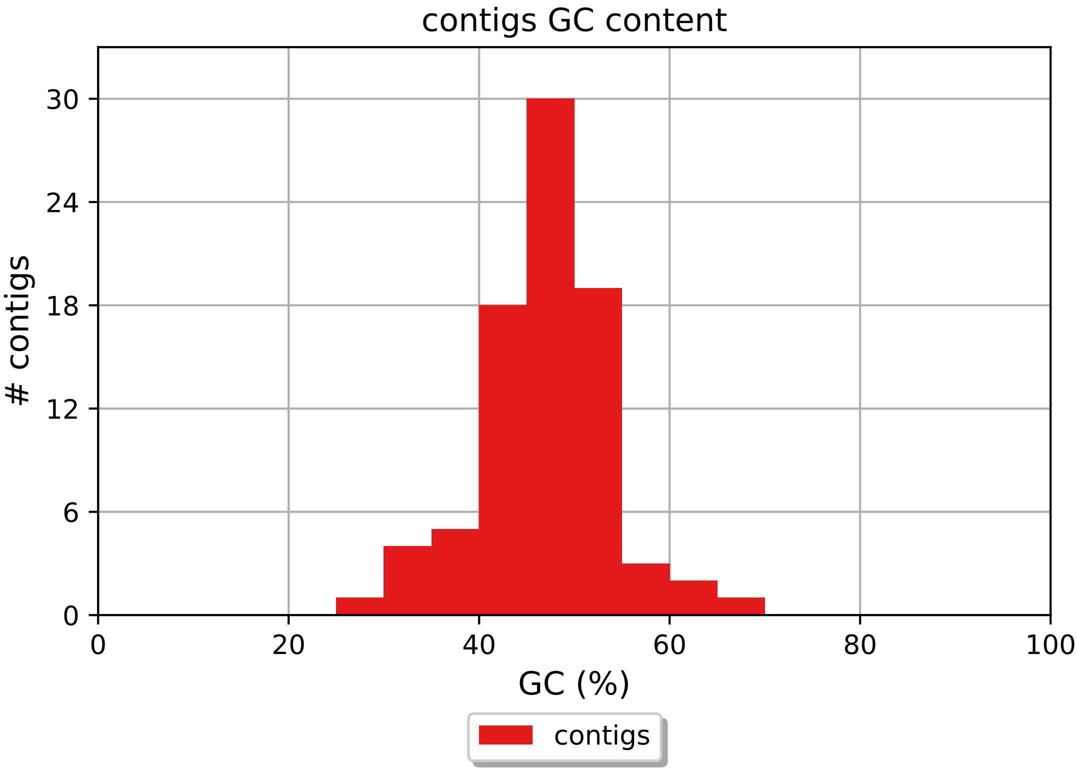

contigs GC content
### contigs
30
24
18
12
6
0
0
20
40
60
80
100
GC (%)
contigs

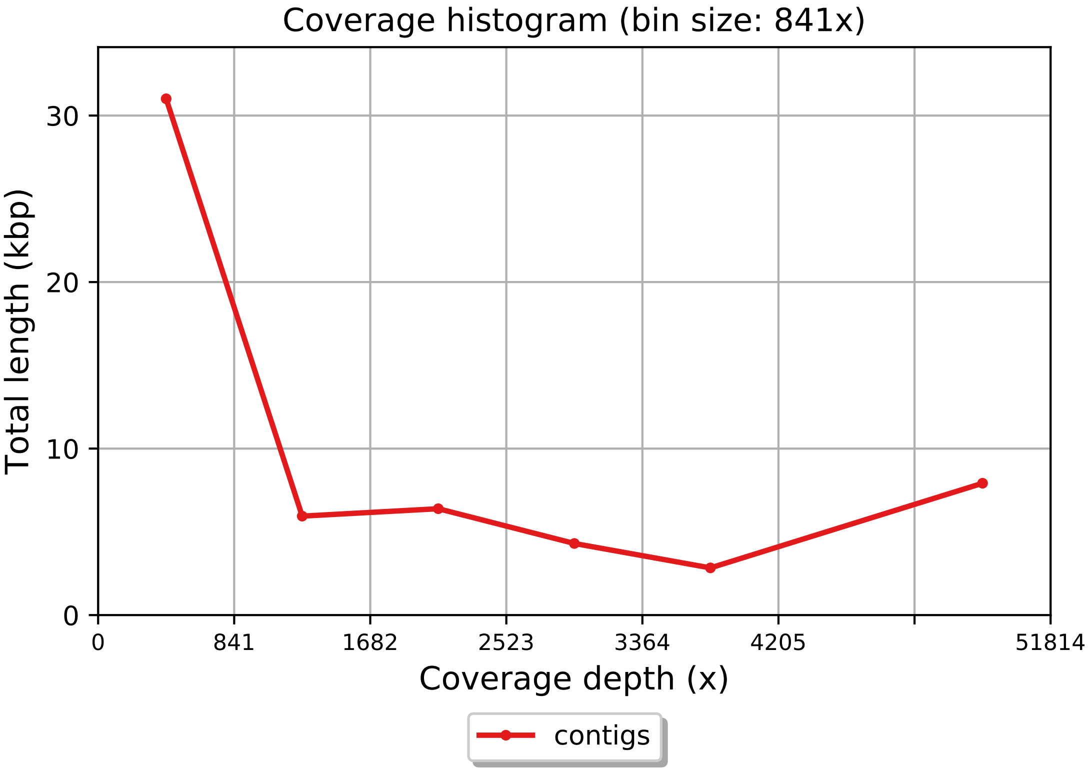

Coverage histogram (bin size: 841x)
Total length (kbp)
30
20
10
0
0
841
1682
2523
3364
4205
51814
Coverage depth (x)
contigs

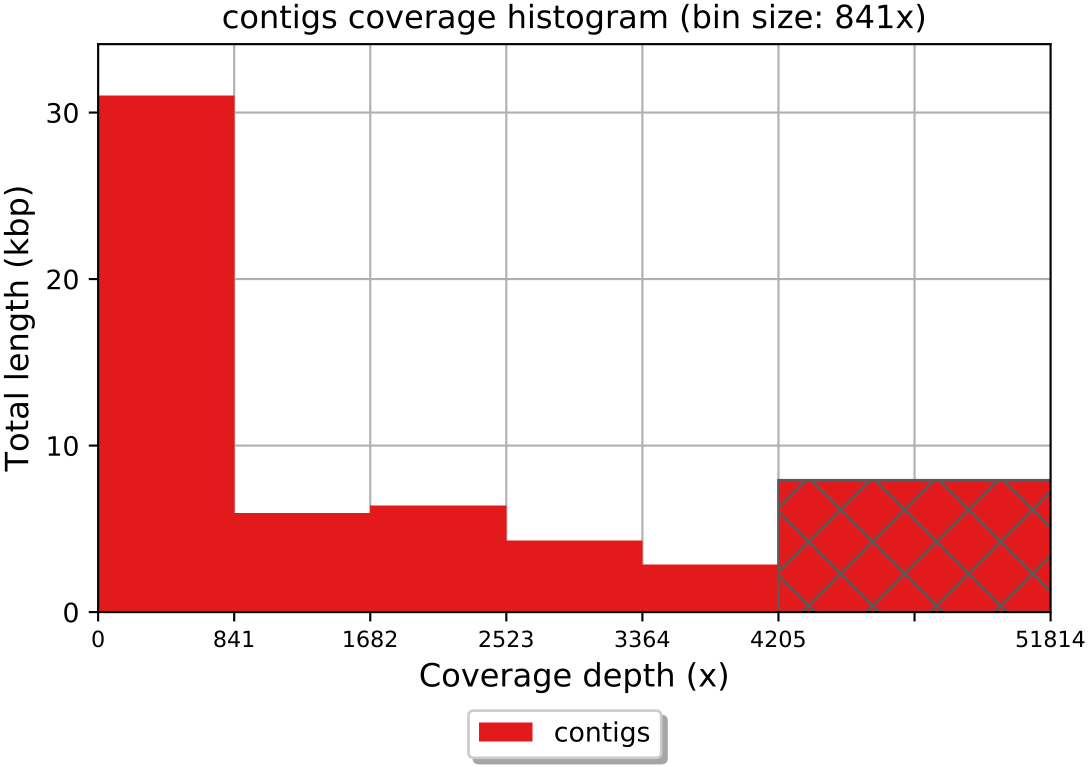

contigs coverage histogram (bin size: 841x)
Total length (kbp)
0
10
20
30
0
841
1682
2523
3364
4205
51814
Coverage depth (x)
contigs
